# Single-cell Landscape Analysis Unravels Molecular Programming of the Human B Cell Compartment in Chronic GVHD

**DOI:** 10.1101/2022.10.13.512162

**Authors:** Jonathan C. Poe, Jiyuan Fang, Dadong Zhang, Marissa R. Lee, Rachel A. DiCioccio, Hsuan Su, Xiaodi Qin, Jennifer Zhang, Jonathan Visentin, Sonali J. Bracken, Vincent T. Ho, Kathy S. Wang, Jeremy J. Rose, Steven Z. Pavletic, Frances T. Hakim, Wei Jia, Amy N. Suthers, Itaevia Curry-Chisolm, Mitchell E. Horwitz, David A. Rizzieri, William McManigle, Nelson J. Chao, Adela R. Cardones, Jichun Xie, Kouros Owzar, Stefanie Sarantopoulos

## Abstract

Alloreactivity can drive autoimmune syndromes. After allogeneic hematopoietic stem cell transplantation (allo-HCT) chronic graft-versus-host disease (cGVHD), a B cell-mediated autoimmune-like syndrome, commonly occurs. Because donor-derived B cells continually develop under selective pressure from host alloantigens, aberrant B Cell Receptor (BCR)-activation and IgG production can emerge and contribute to cGVHD pathobiology. To better understand molecular programing of B cells under selective pressure of alloantigens, we performed scRNA-Seq analysis on high numbers of purified B cells from allo-HCT patients. An unsupervised analysis revealed 10 clusters, distinguishable by signature genes for maturation, activation and memory. We found striking transcriptional differences in the ‘memory’ B cell compartment after allo-HCT compared to healthy or infected individuals. To identify intrinsic properties when B-cell tolerance is lost after allo-HCT, we then assessed clusters for differentially expressed genes (DEGs) between patients with vs. without autoimmune-like manifestations (Active cGVHD vs. No cGVHD, respectively). DEGs were found in Active cGVHD in both naïve and BCR-activated clusters, suggesting functional diversity. Some DEGs were also differentially expressed across most clusters, suggesting common molecular programs that may promote B cell ‘plasticity’. Our study of human allo-HCT and cGVHD provides new understanding of B-cell ‘memory’ in the face of chronic alloantigen stimulation.

**One Sentence Summary:** Our scRNA-Seq study of purified B cells after allo-HCT clarifies molecular differences in human B cell subsets when immune tolerance is lost or maintained.

**Graphical Abstract:** 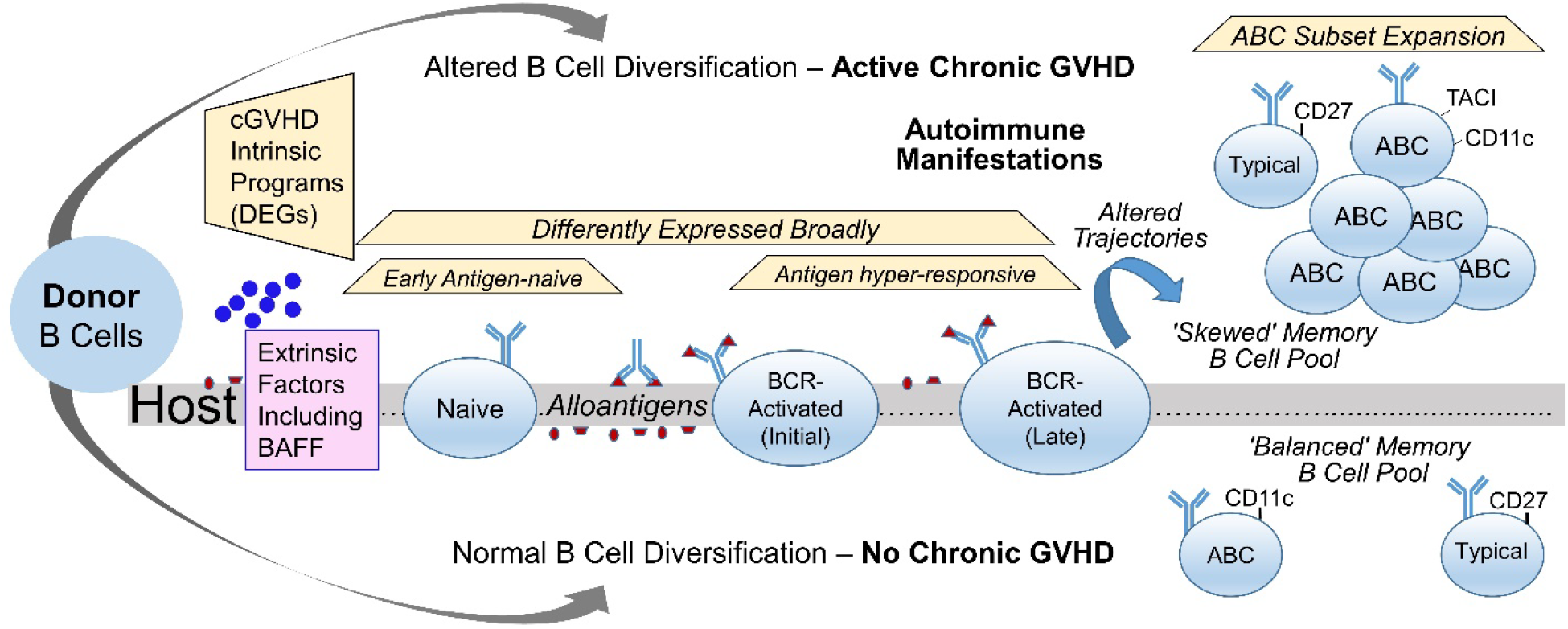

**The circulating human B cell compartment under the selective pressure of allogeneic hematopoietic stem cell transplantation (Allo-HCT) has an intrinsically altered memory B cell pool.** In allo-HCT, genetically disparate donor stem and progenitor cells engraft, regenerating a new peripheral B cell compartment in the host. During ongoing B lymphopoiesis and diversification, selective pressure from alloantigens and other extrinsic factors, including B Cell Activating Factor (BAFF), result in either B cell maturation and immune tolerance (No Chronic GVHD) or altered B cell homeostasis and autoimmune manifestations (Active Chronic GVHD). B cells within defined subsets in patients with Active GVHD have distinct intrinsic programs delineated by differentially expressed genes (DEGs). Some DEGs occur across nearly all B cell subsets (‘Differentially Expressed Broadly’), while other DEGs are more restricted within only one or a few B cell subsets (‘Naïve’, ‘BCR-activated’, or ‘Memory’). We affirm altered trajectories for diversification (blue arrow) and enrichment of an atypical memory B cell (ABC) pool, with intrinsic differences when chronic GVHD occurs.

## Introduction

B cell tolerance checkpoints normally function to silence the high proportion of potentially self-reactive peripheral B cells that recirculate and can mediate autoimmunity (*1*). Chronic graft-versus-host disease (cGVHD) is a T-cell incited autoimmune-like syndrome acquired by some, but not all patients, months after allogeneic hematopoietic stem cell transplantation (allo-HCT). B cells have a substantiated role in cGVHD genesis and perpetuation (*2, 3*). Since blocking T cells potentially attenuates anti-tumor effects after allo-HCT, aberrant peripheral B cell activation, survival, and maturation signaling pathways have become promising therapeutic targets for cGVHD patients (*4, 5*). B cell recovery and homeostasis after allo-HCT are altered in cGVHD, whereby extrinsic factors including BAFF (B cell-activating factor) contribute to a break in B cell tolerance (*6–11*).

B cells continually regenerate. After allo-HCT, B cells from donor stem cells and progenitor cells replace the recipient (host) immune system. Prior to allo-HCT, high dose chemotherapy and/or radiation conditioning creates “space” in the B cell compartment whereby excess BAFF promotes antigen-activated B cells (*10*). Alloantigen and BAFF operate together to promote B cell tolerance loss (*7*). After allo-HCT, donor T cells recognize host polymorphic or alloantigens in a coordinated T-B response (*12*), and follicular helper T cells are required for B cell anti-host reactivity and cGVHD development (*13*). The peripheral B cell compartment in patients with active cGVHD manifestations becomes enriched for B-cell receptor (BCR)-stimulated and IgG-secreting populations (*2, 3, 6, 8, 14-17*). Yet, whether constant exposure to alloantigens and high BAFF alters intrinsic B cell pathways in human allo-HCT remains unknown.

The advent of single-cell RNA-sequencing (scRNA-Seq) enables identification of intrinsic B cell pathways that potentially underpin pathological functions in autoimmune syndromes. We utilized scRNA-Seq to delineate transcriptional programs within 10 peripheral B cell clusters resolved in allo-HCT patients, and identified differentially expressed genes (DEGs) within these clusters that were associated with cGVHD. Some DEGs were common across these B cell populations, while others were restricted to certain subsets. Homing and cell cycle genes including *GPR183* and *CKS2*, respectively, were overexpressed in circulating transitional, activated and memory subsets, suggesting potential roles in B cell dysfunction from the time B cells first emerge in the periphery, and then after alloantigen BCR stimulation. In cGVHD, we found an expanded population of potentially-pathogenic ‘atypical/age-related’ B cells (ABC) (*18, 19*), and known drivers of ABCs including *TBX21* (*TBET)* and *ZEB2* (*18, 20, 21*) that were amongst the increased DEGs. By comparing our allo-HCT scRNA-Seq data with a publically-available dataset on blood B cells from normal healthy donors (HDs) and non-HCT patient groups (HIV, malaria), we found distinct signature gene profiles unique to the allo-HCT setting within these ABC and other memory subsets. Thus, our study provides new insight into altered transcriptional programs found in B cell subsets when B-cell homeostasis occurs after allo-HCT, and identifies potential molecular targets for future study in alleviating cGVHD and other B cell-mediated autoimmune diseases.

## Results

### The circulating B cell compartment after allo-HCT comprises transitional, naïve, antigen-stimulated, and memory single-cell signature gene profiles

Because allo-HCT affords an opportunity to better understand the intrinsic programming of B cell subsets when B cell tolerance is lost or maintained, we analyzed molecular features of a high number of highly purified viable human B cells (**Figure 1A**) from 8 allo-HCT patients > 9 months after engraftment (**Supplemental Table 1**). To confirm cell types, inferences were made by examining the relative expression of genes representing B cells (*PAX5*, *CD22*), and rare residual T cells (*CD3E*) and monocytes (*LYZ*) (**Supplemental Figure 1**). Unsupervised clustering analysis of the 8-patient scRNA-Seq dataset identified 10 clusters of B cells (**Figure 1B**). All 10 B cell clusters were represented in both the four Active and four No cGVHD patients (**Figure 1C**), and were numbered 1-10 from based on largest to smallest total number of B cells in both patient groups combined (**Figure 1D**).

To validate known and potentially novel B cell subsets in allo-HCT patients, we identified hallmark ‘signature’ (marker) genes to distinguish each cluster, and these signature genes were remarkably homogeneous within clusters among all 8 allo-HCT patients. We then plotted log2 normalized expression values for 16 major B cell development and function genes from all 8 patients (**Figure 1, E-H**), each validated using ‘findMarkers’ in the R package Seurat (*22*). Levels of the transitional B cell molecules *VPREB3*, *CD24* and *IRF8* were highest in Clusters 1 and 2 and lowest in clusters 9 and 10 (**Figure 1E**). Comparatively, Cluster 3 had lower expression of *CD24* and *IRF8*, and higher *IRF4*, suggesting a more mature B cell stage. Elevated *LTB* further supported that Clusters 1 and 2 represent recent bone marrow (BM) emigrants because lymphotoxin beta is required for follicle development in the spleen (*23*) and expressed highest in late transitional B cells (immgen.org). Clusters 1, 2 and 3 expressed the highest levels of IgM heavy chain (*IGHM*). Clusters 1 and 3 primarily expressed kappa Ig light chain and Cluster 2 primarily expressed lambda Ig light chain (*IGKC* and *IGLC3*, respectively, **Figure 1F**). Clusters 9 and 10 expressed similarly high levels of microRNA-155 (*MIR155HG*, **Figure 1G**), essential for B cell antibody isotype switching (*24*), and similarly high expression of *MYC* and *NFKB1*, suggesting these B cells may be poised to enter the cell cycle. Cluster 10 had uniquely high levels of the chemokine *CCL22*, indicating B cells poised to interact with T-follicular helper cells within the secondary lymphoid organ (SLO) microenvironment. By contrast, Cluster 5 expressed high levels of *CD27*, along with IgG1 and IgG3 heavy chain genes (*IGHG1* and *IGHG3*, respectively), indicating an enrichment for isotype-switched memory B cells (**Figure 1H**). Cluster 5 also expressed the highest *ITGAX* (*CD11C*), a distinguishing marker of ABCs (*25, 26*). Since cluster proximity can reflect signature program relatedness, we also depicted 3D UMAP plots to further highlight spatial relationships among neighboring B cell clusters (**Figure 1I**). Together, these data allowed us to define molecular pathways in B cell subsets that regenerate and circulate in an alloantigen-rich environment.

**Figure 1.**
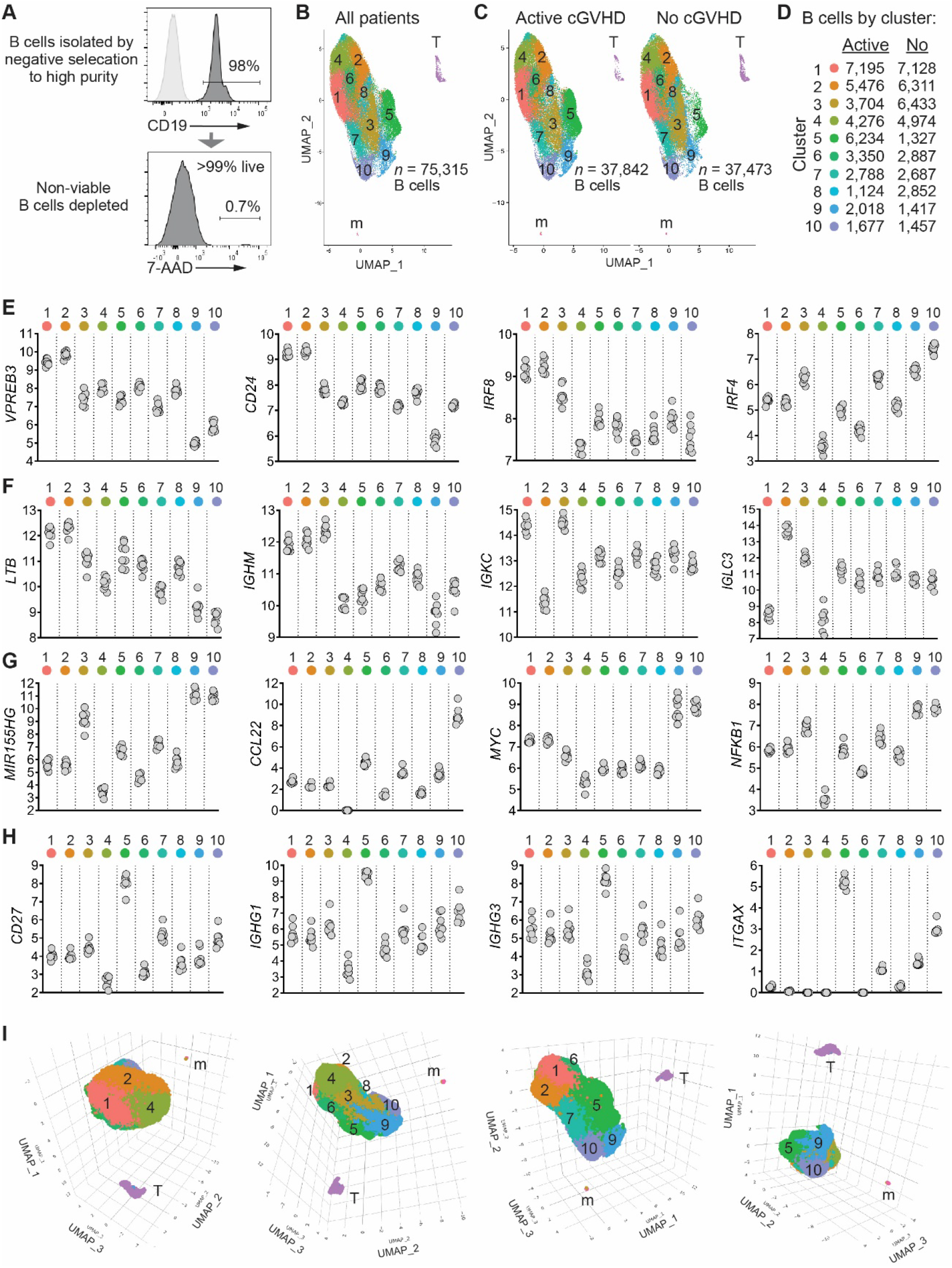
Unsupervised clustering analysis of scRNA-Seq data reveals multiple B cell subsets characterized by signature genes in allo-HCT patients. (**A**) Representative flow cytometry histograms showing B cell purity and viability from an allo-HCT patient. 10,000 high-quality B cells per patient sample isolated in the same manner were targeted for 10X Genomics single-cell library construction (*n*=4 No cGVHD, *n*=4 Active cGVHD). (**B**) 2D Uniform Manifold Approximation and Projection (UMAP) expression profiles of all untreated cells from the 8 allo-HCT patients. Numbers indicate each of the 10 major clusters identified as B cells, which were distinct spatially from residual cells identified as T cells (T) and monocytes (m). As indicated, 75,315 high-quality B cells were analyzed. (**C**) UMAP as in (**B**), with B cells partitioned by patient group. Total high-quality B cells per group are indicated. (**D**) Number of B cells mapping to each of the 10 B cell clusters by patient group. (**E-H**) Log2 normalized expression of genes indicative of B cell maturity (**E, F**), along with activation, antibody production potential, and memory markers (**G, H**), by B cell cluster. Each symbol (gray circle) represents regularized log-transformed gene counts (*27*) [y-axes] from one of the 8 allo-HCT patients. (**I**) 3D UMAP projections of B cells from all patients viewed at different angles of rotation.

### Allo-HCT B cell clusters can be further delineated via established signature pathways

We first reordered the 10 clusters based on the unsupervised clustering in **Figure 1** and published literature of known B cell subsets that enter the periphery and those that eventually encounter antigen/co-stimulatory signals. As shown in **Figure 1A**, we henceforth ordered from least mature/activated first and ending with memory signature gene profiles as follows: Cluster 1, 2, 4, 6, 8, 7, 3, 9, 10 and 5. In **Figure 2A,C-H** colored squares in the heat maps indicate signature genes of interest (rows) reaching significance (*P_adj_* <0.05) in the cluster indicated (columns), as weighted against all other clusters. Red hues indicate increased signature gene expression, and blue hues indicate decreased signature gene expression (log FC). Notably, Clusters 1, 2, 4, 6 and 8 retained one or more hallmark genes of B cell immaturity, while Clusters 7, 3, 9, 10 and 5 expressed markers of antigen activation, cytoskeletal activity, or the capacity to produce antibodies (**Figure 2A,B**). We further validated this ordering of the 10 B cell clusters by interrogating the Kyoto Encyclopedia of Genes and Genomes [KEGG] (*28*) and Gene Ontology [GO] (*29*) databases using signature genes of interest for major molecular pathways (**Figure 2C-H** heat maps). We found that Clusters 9 and 10 showed high expression of genes involved in B cell–T cell interactions, transcription, proliferation, survival, and metabolic processes (**Figure 2A,C-H**). Clusters 7 and 10 shared multiple ribosomal signature genes (**Figure 2G**), suggesting increased protein synthesis potential. Overall, Clusters 6 and 8 had relatively few signature gene increases, suggesting relative quiescence, although Cluster 6 had strong expression of actin cytoskeleton organizer *CDC42SE1* and the proto-oncogene *MENT* (previously *C1orf56,* **Figure 2F**). The less mature/less activated subsets (Clusters 1, 2, 4 and 6) tended to have more decreased signature genes (‘Down’, **Figure 2I**), while the activated/memory subsets (Clusters 3, 9, 10 and 5) tended to have more increased signature genes (‘Up’, **Figure 2I**). Thus, our data elucidate a hierarchy and relatedness among peripheral B cell subsets in human allo-HCT.

**Figure 2.**
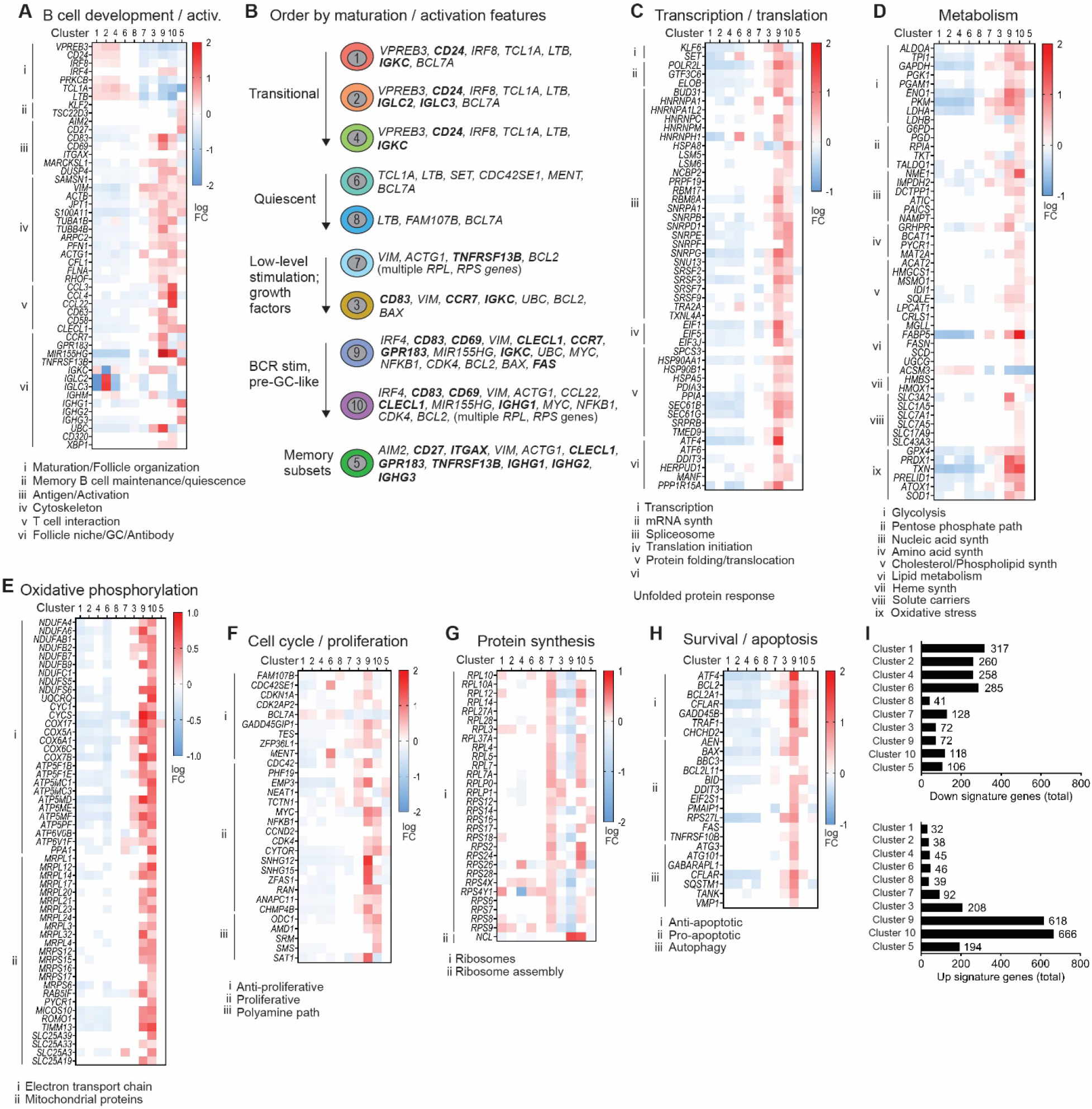
Signature genes empirically assigned to biological pathways corroborate the inter-relatedness and features of B cell development and activation among the 10 B cell clusters. (**A**) Signature (marker) genes related to B cell development and activation assessed in the 8 allo-HCT patients in the scRNA-Seq data set by manually interrogating the KEGG and GO databases. Each colored square represents a significant (*P_adj_* <0.05) log fold change (log FC) value for the gene indicated (rows), in the B cell cluster indicated (columns). Red shading indicates increased signature gene expression, while blue shading indicates decreased signature gene expression, as weighted against all other clusters. Genes were subdivided according to specific pathways as indicated by the roman numerals at left and corresponding key below each heatmap. (**B**) Cartoon depiction of B cell clusters and their associated signature genes that may help categorize these subsets (as labeled at left). Signature genes encoding surface proteins are highlighted in bold font. (**C-H**) Signature genes were assessed in the major biological pathways indicated above each heat map, and subdivided by the more specific pathways indicated by the roman numerals at left and corresponding keys below. (**I**) Total numbers of signature genes reaching significance within each cluster, being either decreased (‘Down signature genes’, top graph) or increased (‘Up signature genes’, bottom graph).

### The circulating ‘memory’ B cell compartment in allo-HCT patients is distinct from HDs and non-HCT patients with chronic infections

To pursue the hypothesis that chronic exposure to alloantigens results in intrinsic alterations of known B cell subsets, we compared gene profiles in allo-HCT patients to HDs and individuals with non-HCT chronic diseases. Since typical, functional memory B cell recovery is delayed after transplant (*30*), we assessed ‘memory’ B cell signatures. We leveraged a similar, publicly-available scRNA-Seq dataset generated from blood B cells of HDs (*n*=3) and non-HCT patients chronically exposed to HIV (*n*=3) or malaria (*n*=3), published by Holla et al. (*25*). We first identified total *CD27^+^ITGAX^−^*, *CD27^−^ITGAX^+^*, and *CD27^+^ITGAX^+^* B cells from all patient and HD groups. We then assessed the frequency of B cells within these Cluster 5 subsets expressing other genes associated with the function or additional phenotype(s) of memory B cells (*31–34*). For most of these ‘memory’ signature genes, the proportion of positive B cells in the 3 memory subsets was similar among all 5 groups (**Supplemental Figure 2**). However, certain genes were distinct in allo-HCT patients, irrespective of cGVHD status (**Figure 3**). B cells expressing *ADA*, associated with a form of SCID with disrupted B cell tolerance (*35*), the T cell-attracting chemokine *CCL22* (*36*), and three G-protein coupled receptors (GPCRs) important for B cell exit from the BM and/or homing to SLO niches, *CCR7*, *S1PR1*, and *S1PR2* (*37–41*), were all increased in proportion across *CD27*/*ITGAX* subsets in allo-HCT patients. Allo-HCT patients also had broadly increased proportions of B cells in these subsets expressing survival regulators with roles in memory B cell maintenance including *BCL2* (*42*) and *TRADD* (*43*). By contrast, allo-HCT patients had a paucity in B cells after expressing *FOS*, important during the GC response and terminal differentiation to plasma cells (*44*), and *ITGB2*, which pairs with various integrin alpha chains to mediate migration (*25*). Together these data reveal that the memory B cell pool in allo-HCT possesses intrinsic molecular hallmarks of altered tolerance, migration, SLO homing, and survival.

**Figure 3.**
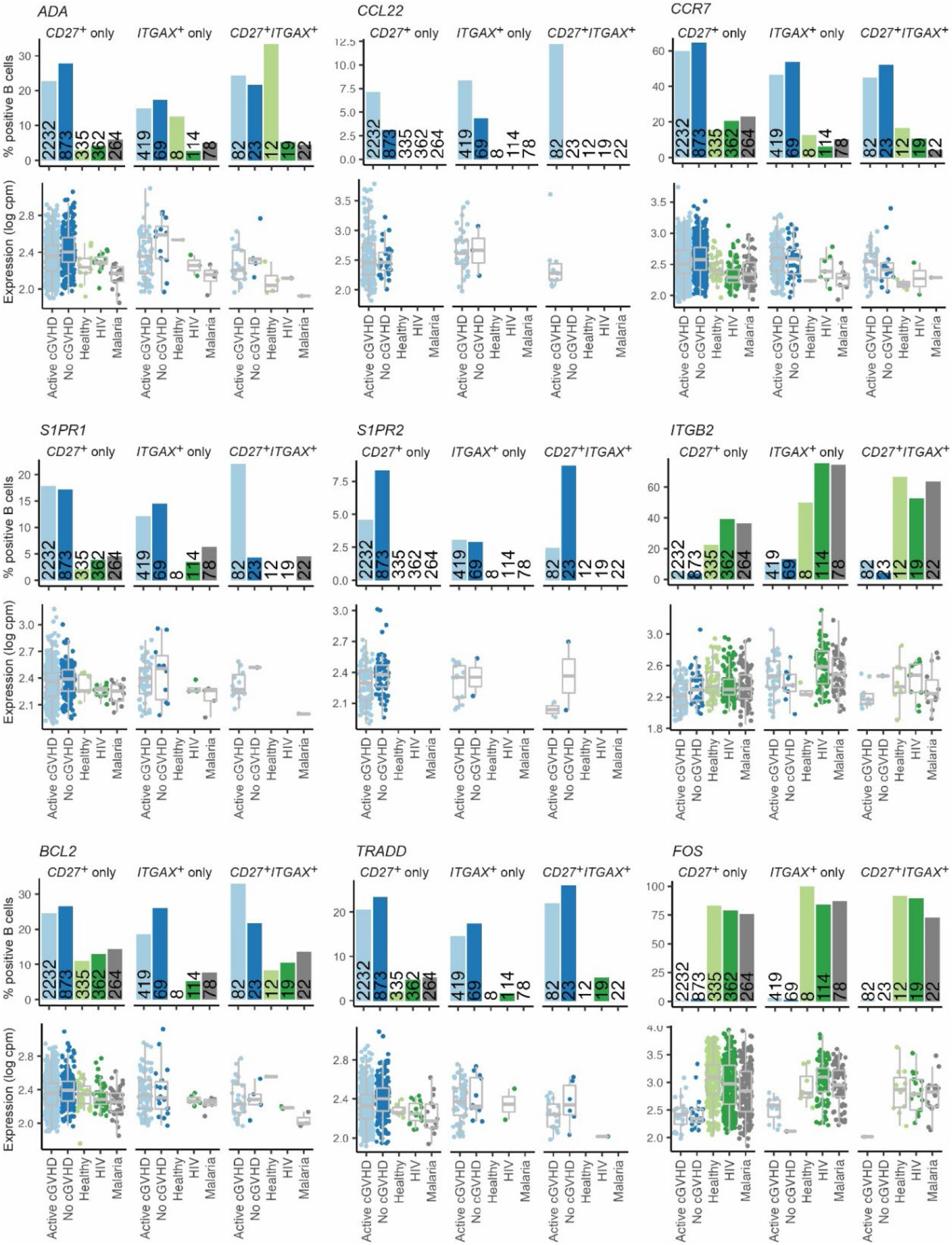
‘Memory’ *CD27* and *ITGAX* B cell subsets are transcriptionally unique in allo-HCT patients compared to HDs and non-HCT patients. Shown are results for 9 ‘memory’ B cell genes of interest displaying differences in the proportion of positive B cells between allo-HCT patients compared to HD (Healthy) and non-HCT patient groups (HIV, Malaria). Bar graphs represent the % positive B cells for the population indicated. Numbers embedded within the bars indicate the total number of B cells identified within the subset indicated, for the patient or HD group. In the gene expression graphs below each bar graph, each point represents an individual B cell, with the data normalized to show expression values for the gene of interest per million total mapped reads in the same B cell, expressed on a log scale (log CPM).

### In Active cGVHD the BCR-experienced ‘memory’ Cluster 5 includes a distinct CD27^+^ population and an expanded extrafollicular-like ABC population

Since antigen-experienced ‘memory’ B cells likely include pathogenic B cells that emerge in cGVHD, as occurs in other diseases such as lupus (*1, 45*), we further interrogated ‘memory’ Cluster 5. In addition to a CD27-expressing population, we found that Cluster 5 harbored B cells expressing *ITGAX* that encodes CD11c, which broadly marks ABC populations that also lack CD21 (CD11c^+^CD21^−^). ABCs have a pathogenic role in both autoimmunity and chronic inflammation, revealing a functional complexity of these ‘memory’ B cells (*25, 46, 47*). ABCs are often termed either ‘atypical’ or ‘age-related’ B cells, the latter term originally given because they accumulate in aged mice (*48, 49*). ABCs are considered ‘atypical’ because they undergo activation and differentiation outside of the germinal center (GC), rapidly homing to the outer follicle as ‘extrafollicular’ B cells (*50, 51, 52*). To elucidate these ‘memory’ subpopulations in cGVHD, we examined the pattern of *CD27* and *ITGAX* expression in Cluster 5. As shown in **Figure 4A,B**, cells expressing either *CD27* (left UMAPs) or *ITGAX* (right UMAPs) were largely spatially segregated, suggesting largely independent subsets. To validate this finding, we performed flow cytometry analysis on blood B cells from an independent cohort of allo-HCT patients. As shown in **Figure 4B**, allo-HCT B cell subsets primarily expressed either CD27 or CD11c alone, with a smaller subset expressing both markers (double-positive, ‘DP’). Importantly, CD11c^+^CD27^−^ ABCs were significantly expanded in Active cGVHD patients compared to No cGVHD patients and HDs (**Figure 4B,C**). Interestingly, the CD11c^+^CD27^+^ ‘DP’ subset was proportionally lower in the allo-HCT environment compared to HDs (**Figure 4B,C**). Overall these data are consistent with previous work showing that some CD11c^+^CD27^−^ ABC subsets are expanded in cGVHD and other diseases, sometimes exhibiting an ‘exhausted’ phenotype (*53–56*). We also examined a subset of CD11c^+^CD21^−^ ABCs that lacks both CD27 and IgD expression (CD11c^+^CD21^−^CD27^−^IgD^−^), called ‘DN2’, that is prevalent in lupus and can produce disease-associated antibodies (*52, 57, 58*). The emergence of pathogenic ABCs (including DN2) can be driven by one or more costimulatory signals that cooperate with BCR stimulation (*52, 59*). We found DN2 ABCs were markedly expanded in Active cGVHD patients relative to No cGVHD patients (**Figure 4D,E**). These data affirm our scRNA-seq data and advance previous work (*52, 55–58*) suggesting that potentially pathologic ABC subsets arise under constant alloantigen exposure.

To assess B cell surface phenotypes more broadly in allo-HCT and potentially elucidate subsets responsive to excess BAFF (*10*), we examined cell surface CD24 and TACI (TNFRSF13B) expression in the context of CD27, IgD, CD11c, CD21 (**Figure 4F**). Using high-dimensional flow cytometry [PhenoGraph, (*60, 61*)] we identified discreet B cell populations in an unbiased manner from concatenated allo-HCT samples. PhenoGraph analysis resolved 15 clusters, designated by lower case letters ‘a-o’ (**Figure 4G**). CD11c^+^ ABCs were primarily resolved by Cluster ‘a’ (**Supplemental Figure 3**), significantly expanded in Active cGVHD (**Figure 4H,I**). By contrast, four clusters consisting of IgD^+^CD27^-^ naïve-like B cells (Clusters ‘b, c, d and e’, **Supplemental Figure 3**) were decreased in Active cGVHD patients (**Figure 4H**). ABC Cluster ‘a’ was also TACI^+^, with a mix of IgD^+/–^ B cells (**Supplemental Figure 3**). Six clusters expressed high levels of surface TACI (Clusters ‘a, h, j, k, l and n’, **Figure 4J** **and Supplemental Figure 3**). Cluster ‘n’ expressed substantially higher surface TACI in some Active cGVHD patients (**Figure 4J**, arrow), and mapped adjacent to ABC Cluster ‘a’ in the PhenoGraph UMAP (**Figure 4G**). Clusters ‘j’ and ‘k’ likewise had elevated TACI in some Active cGVHD patients (**Figure 4J**, arrows). Accordingly, regions of B cells in the scRNA-Seq data having increased *TNFRSF13B (TACI)* transcripts were readily evident in Active cGVHD (**Figure. 4K**, UMAP boxed region II). These results are consistent with the previous observations that BAFF is a driver of B cell hyper-responsiveness and alloantibody production in cGVHD (*6, 7, 9, 10*).

To examine how allo-HCT compares to the non-HCT setting, we also performed PhenoGraph analysis on HD blood B cells using the same flow cytometry panel. The HD analysis also predicted 15 clusters, but HDs had a greater number of B cell clusters marking with CD27 (**Supplemental Figure 4**, Clusters ‘m,n,o’) compared to allo-HCT patients (**Supplemental Figure 3**, Clusters ‘l,o’ only). Furthermore, in allo-HCT patients Cluster ‘o’ represented mostly CD27^bright^ B cells that lacked CD21, CD24, IgD and CD11c expression, suggestive of isotype-switched plasmablasts (“PB”, **Supplemental Figure 3**). These were among the rarest in both allo-HCT groups (**Figure 4H**), and this cluster was notably absent in HDs (**Supplemental Figure 4**). These data reveal that CD27^+^ B cell subsets after allo-HCT are abnormal or lacking, extending previous observations (*53*). The relative reduction in CD27^+^ B cells in allo-HCT with reciprocally expanded ABCs in Active cGVHD is consistent with PhenoGraph findings recently reported in pediatric patients with CVID and related syndromes (*62*). Together, data affirm heterogeneity of the post-transplant ‘memory’ B cell pool and begin to validate our scRNA-Seq data by showing TACI expression increases in ABCs or their precursors in cGVHD.

**Figure 4.**
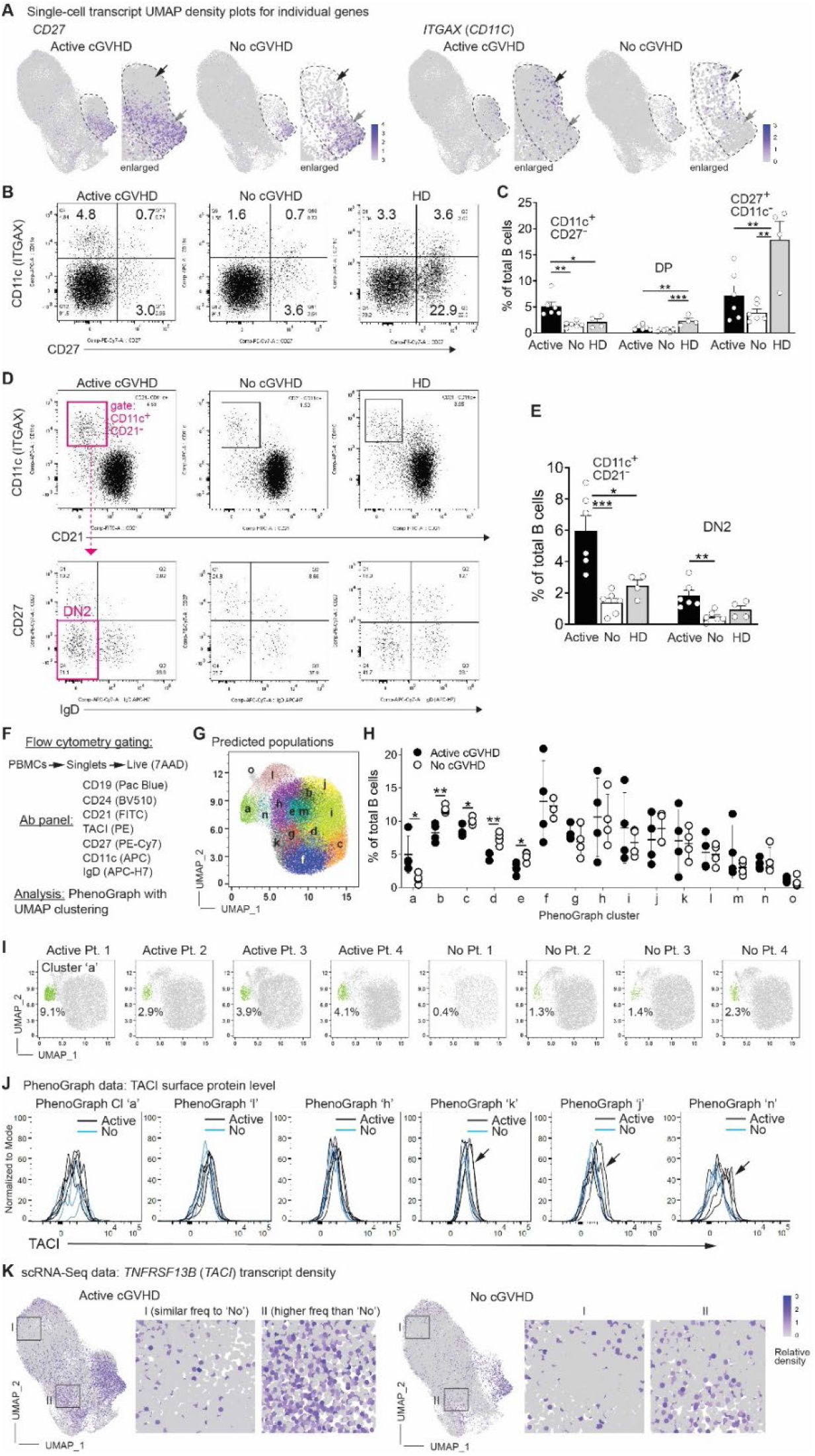
Cluster 5 contains both ‘typical’ CD27^+^ and ABC-like memory populations that are expanded in Active cGVHD. (**A**) B cells expressing *CD27* or *ITGAX* transcripts are enriched in Cluster 5 in two separate regions. Shown are UMAP density plots from the scRNA-Seq dataset displaying relative transcript density for the gene indicated, at the single-cell level. The dashed line approximates the perimeter of Cluster 5 (also enlarged for better clarity). Gray arrows indicate a region enriched for *CD27* expressing B cells, and black arrows indicate a region enriched for *ITGAX* expressing B cells. (**B,C**) Flow cytometric analysis to identify ABCs, CD27^+^ memory B cells, and a small population of CD11c^+^CD27^+^ memory B cells in PBMC samples from a cohort of allo-HCT patients with either Active cGVHD (*n*=6) or No cGVHD (*n*=6), or from HDs (*n*=4). PBMCs were pre-gated on live (7-AAD^−^) CD19^+^ B cells. Dot plots in (**B**) show representative individuals analyzed from each group. In (**C**), results from all groups for the CD11c^+^CD27^−^ population, CD11c^+^CD27^+^ (DP) population, and CD11c^−^CD27^+^ population (as gated in [**B**]) are represented. Statistical comparison was performed using a one-way ANOVA with Tukey’s multiple comparisons test (GraphPad Prism 9 software; *, p<0.05; **, p<0.01; ***, p<0.001). (**D,E**) DN2 B cells are expanded in allo-HCT patients with Active cGVHD. (**D**) Representative FACS plots from PBMCs of allo-HCT patients with Active or No cGVHD, and from a HD, generated after first gating on viable (7-AAD^-^), CD19^+^ B cells, with CD11c^+^CD21^−^ B cells (upper panel gate) and then DN2 B cells (CD11c^+^CD21^−^CD27^−^IgD^−^) identified in all groups as depicted in magenta for the Active cGVHD sample. (**E**) Statistical comparison between groups (Active cGVHD, n=6; No cGVHD n=6, HDs, n=4) was performed using a one-way ANOVA with Tukey’s multiple comparisons test (GraphPad Prism 9 software; *, p<0.05; **, p<0.01; ***, p<0.001). (**F-J**) Flow cytometry plus PhenoGraph analysis performed on blood B cells from Active cGVHD (*n*=4) or No cGVHD (*n*=4) patients. Shown in (**F**) is the gating strategy to distinguish B cells (top line), the B cell antibody panel used (middle), and the analysis platform (bottom). Shown in (**G**) is the concatenated PhenoGraph UMAP plot from all 8 patients, with 15 clusters identified (letters). The graph in (**H**) indicates B cell frequency within each of the 15 clusters by patient group. Each symbol represents results from one of the 8 patients assessed. Statistical comparisons were performed using a two-tailed, unpaired t-test (GraphPad Prism 9 software; *, *p*<0.05; **, *p*<0.01). The PhenoGraph UMAP plots in (**I**) show the position (green) and frequency (%) of B cells for Cluster ‘a’, which represents a population of ABCs (**Supplemental Figure 3**). In (**J**), histogram overlays for TACI surface protein expression on B cells in the six clusters from the PhenoGraph assessment described as being designated TACI^Br^ (**Supplemental Figure 3**), from the 4 allo-HCT patients in each group. Arrows for Clusters ‘k’, ‘j’ and ‘n’ highlight elevated TACI expression levels observed in some Active cGVHD patients. (**K**) B cells positive for *TNFRSF13B* (*TACI*) transcripts appear elevated in Active cGVHD patients. Normalized expression of *TNFRSF13B* in UMAP space are shown, separated by patient group. Boxed areas in representative regions (I, II) are enlarged to show detail.

### Genes critical for B cell fate and function are differentially expressed in Active cGVHD

We next asked whether intrinsic, aberrant B cell programs were found in Active cGVHD patients by examining DEGs between allo-HCT patient groups in the scRNA-Seq data (**Supplemental Table 3**). Bar graphs in **Figure 5** represent total DEGs by cluster, either increased (Up, **Figure 5A**) or decreased (Down, **Figure 5B**) in Active cGVHD compared to No cGVHD. Not surprisingly, ‘BCR-activated’ Clusters 9 and 10, and ‘memory’ Cluster 5, collectively had the most DEGs (Up or Down) compared to the more naïve, resting subsets (**Figure 5A,B**).

Given a potentially prominent role for BAFF in the genesis and progression of cGVHD (*6, 7, 9, 10, 63*), it was notable that the BAFF receptor *TNFRSF13B* (*TACI*) was increased in Clusters 8 and 10 (**Figure 5C,E**). Other members of the TNF superfamily (TNFSF) or TNF-receptor superfamily (TNFRSF) involved in key B cell pathways were likewise Up DEGs in Active cGVHD B cells. These included *TNFRSF14* (*LIGHTR*, *CD270*), *TNFSF10* (*TRAIL*, *CD253*), *TNFRSF12A* (*FN14*, *TWEAKR*, *CD266*), *TNFRSF1B* (*TNFR2*, *CD120b*), and *TNFRSF10B* (*TRAILR2*, *CD262*), **Figure 5C,E**. Interestingly, *GPR183* (*EBI2*) was overexpressed in Cluster 8 along with *TNFRSF13B* (**Figure 5C,E**; **Supplemental Figure 5A**), possibly reflecting a positive influence by BAFF on EBI2 expression, as reported (*64*). Co-expression of *GPR183* and *TNFRSF13B* indeed occurred in some B cells, which were expanded 4-fold in Active cGVHD patients (**Supplemental Figure 5B,C**). Indeed EBI2, like TACI, was expressed at a high level on the surface of CD11c^+^ ABCs by flow cytometry (**Supplemental Figure 5D**), while follicular-like naïve B cell subsets were generally low/negative for EBI2 and TACI (**Supplemental Figure 5E**). These data help validate the recent finding that *TNFRSF13B* and *GPR183* are major DEGs defining a B cell branch point that immediately precedes ABC expansion in malaria patients (*25*).

From the total list of DEGs (**Supplemental Table 3**), those depicted in **Figure 5C-E** represent our focus on genes with known functional roles in B cells (‘Select’ DEGs). A relatively large number of select DEGs occurred in at least one B cell cluster. Regulators of upstream BCR signaling, *BTK* and *BLNK*, were Up DEGs in Active cGVHD, including ‘memory’ Cluster 5 (**Figure 5C,E**). This is consistent with work showing BAFF-dependent increases in BLNK protein in B cells from Active cGVHD patients and mice (*7, 14*). The transcription factor *ZBTB20* was increased in pre-GC-like Cluster 10, and this zinc finger protein is essential for the conversion of BCR-activated B cells to antibody-secreting cells (ASCs) and maintenance of long-lived ASCs (*65*). *CAV1* (*CAVEOLIN-1*), which controls BCR compartmentalization on the plasma membrane (*66*), was decreased in Clusters 1 and 10 (**Figure 5D,E**). Accordingly, *Cav1* deficiency in mice leads to altered nanoscale organization of IgM-BCRs and a skewed Ig repertoire with features of poly-reactivity and auto-reactivity (*66*). Molecules functionally important in ABC expansion or function were also significantly affected. *ZC3H12A*, which encodes the RNA-binding protein REGNASE-1, was decreased in Cluster 1 (**Figure 5D,E**), and Regnase-1 deficiency at earlier stages of B cell development in mice produces severe autoimmune immunopathology and early mortality, accompanied by ABC expansion (*67*). *TBX21* (encoding *TBET*), *ZEB2* and *EGR3* are important and potential master regulators of ABC development and/or function in mice and humans (*18, 20, 21, 59*), and each were Up DEGs in either ‘memory’ Cluster 5 (*TBX21*, *ZEB2*) or activated Cluster 9 (*EGR3*), **Figure 5C,E**. Cluster 5 was enriched for ABCs (**Figure 4A**), suggesting that dysregulated *TBET*, *ZEB2* and *EGR3* are potential drivers of the observed ABC expansion in Active cGVHD patients (**Figure 4B-I**). Thus, key DEGs in Active cGVHD suggest potential functional diversity before and after BCR engagement.

Migration and maturation programs were altered, with Active cGVHD B cells comprising Up DEGs for *GPR183/EBI2* and other GPCRs known to regulate B cell homing and retention within essential SLO niches, including *P2RY8*, *S1PR2*, and *CXCR3* (*39, 68–71*). *GPR183/EBI2* was notably increased across five B cell clusters (**Figure 5C,E**), including the transitional-like subsets (Clusters 1 and 2) and the memory/ABC-like subset (Cluster 5). This was a potentially important finding because EBI2 has an intricate role in regulating B cell movement within the follicle before and following the GC reaction (*68, 71*). *P2RY8* and *S1PR2* were each Down DEGs in both the activated Cluster 9 and/or the ‘memory’ Cluster 5 (**Figure 5D,E**). This is an intriguing finding since these molecules are also pivotal in B cell homing and confinement to follicular niches or the GC reaction (*39, 70, 72*), and decreased *P2RY8* expression is linked to pathogenic antibody production and expansion of plasma cells and ABCs in humans and mice (*73*). These DEG data thus suggest altered follicle and GC movement of B cells in human cGVHD, consistent with findings in cGVHD mouse studies (*3, 15, 74*). Finally, genes associated with cell cycle were among the observed DEGs in Active cGVHD. These included *CDKN1B* and *CCND3* (Up, **Figure 5C,E**), and *CDKN1A* (Down, **Figure 5D,E**). This was consistent with our previous observations that Active cGVHD B cells demonstrate enhanced proliferation to some important stimuli, including low-dose surrogate antigen and NOTCH ligand (*8*).

*CXCR4*, a regulator of B cell migration from the BM to the periphery (*75*), was increased in transitional B cell Clusters 1, 2 and 4 in Active cGVHD (**Figure 5C,E**). This further suggested aberrant migration and homing, and compelled analysis of cGVHD skin B cells. We performed scRNA-Seq on a punch biopsy of lesional skin (dermis) from a patient with sclerodermatous cGVHD and compared it with a biopsy of normal dermal skin from a HD undergoing abdominoplasty. Dermal B cells were detected in both samples, and were distinguishable by a single UMAP cell cluster plus signature gene profile (**Supplemental Table 4**), which indicated the presence of Igκ (*IGKC*), mostly isotype-switched (*IGHG1*, *IGHG3*, *IGHG4*, *IGHA1*) B cells. Other notable signature genes included *GPR183* (*EBI2*) and receptors for BAFF (*TNFRSF13B* [*TACI*] and *TNFRSF13C* [*BAFFR*]). Remarkably, multiple Up DEGs in blood B cell clusters in Active cGVHD patients (**Figure 5C,E**) were also Up DEGs in Active cGVHD sclerodermatous skin B cells (**Supplemental Table 5**). *POU2F2* (*OCT2*), important for generation of ASCs (*76*), *SWAP70*, a molecule phosphorylated by SYK that influences B cell migration and ABC expansion (*77, 78*), and *AIM2*, which has a positive role in the expansion of autoreactive B cells in a mouse lupus model (*79*), were increased in both blood and skin B cells in Active cGVHD. Not surprisingly, most DEGs observed in sclerodermatous skin that were also DEGs in blood B cells of Active cGVHD patients occurred in the most activated and differentiated blood B cell clusters, primarily Clusters 3, 9, 10 and 5 (**Figure 5C,E**, **Supplemental Table 3**, **Supplemental Table 5**). Thus, lesional skin B cells in Active cGVHD potentially share characteristics of hyper-activity with circulating B cell subsets. Together these data both validate our previous work, and now elucidate distinct maturation, activation, survival and homing pathway alterations of B cell subsets when B cell tolerance is not achieved in the form of cGVHD.

**Figure 5.**
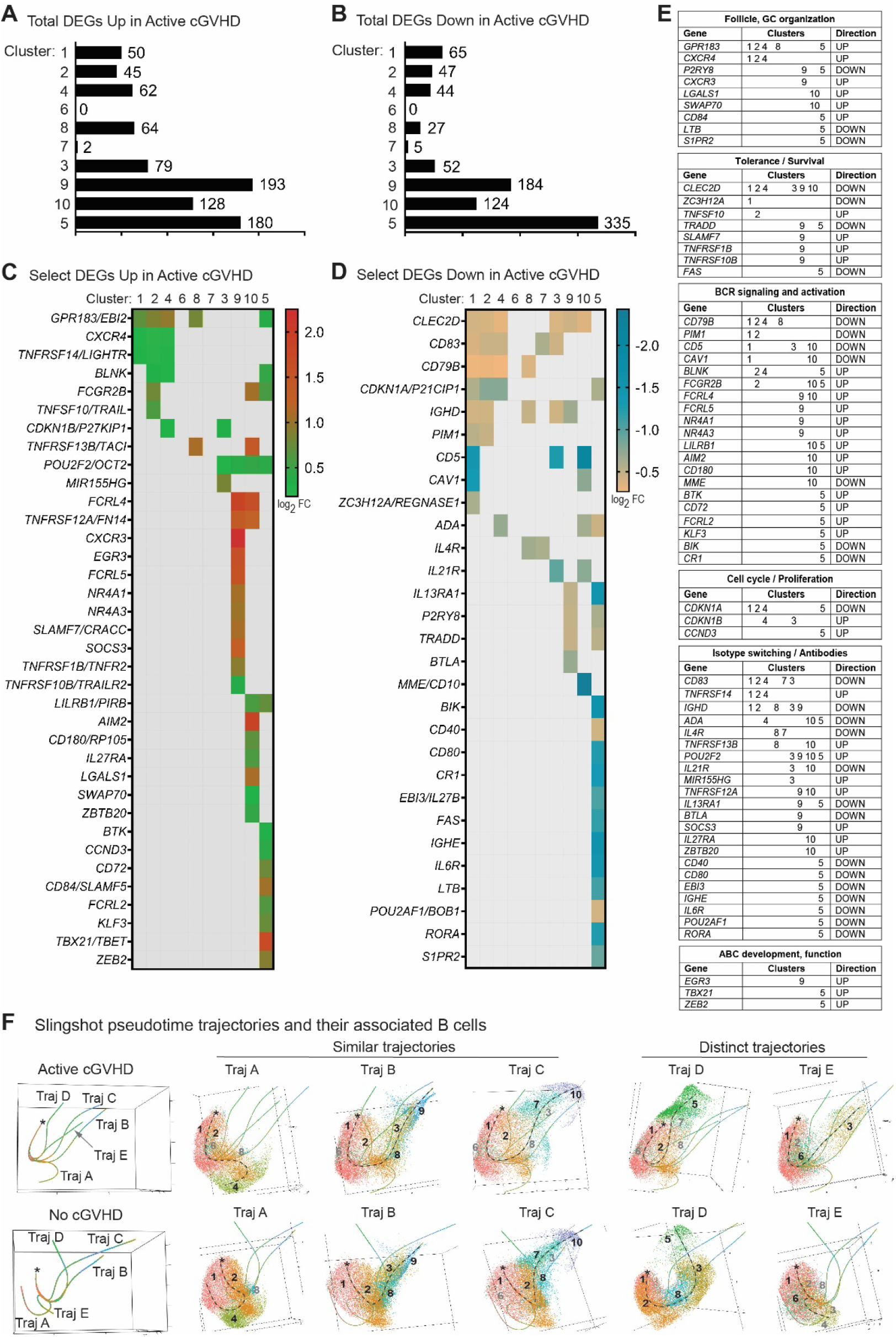
Molecules critical to B cell function are altered within clusters in Active cGVHD. (**A-D**) DEG analysis within the 10 B cell clusters in the scRNA-Seq dataset based on disease status (as described in Figure 1: Active cGVHD, *n*=4; No cGVHD, *n*=4). Bar graphs indicate total DEG number by cluster Up (**A**) or Down (**B**) in Active cGVHD. The heat maps in (**C**) and (**D**) depict DEGs selected from the entire dataset (**Supplemental Table 3**), critical for various aspects of B cell function (‘Select’ DEGs). Colored squares represent significant (*P_adj_* <0.05) log2 fold change (log2 FC) values for DEGs shown (rows) within the cluster(s) indicated (columns), either Up (**C**) or Down (**D**) in Active cGVHD B cells. (**E**) All genes from (**C**) and (**D**) were grouped into the categories shown based on their known roles in B cell function. Clusters with DEGs are summarized in the middle column, and direction of change (Up or Down) in Active cGVHD B cells relative to No cGVHD B cells is indicated in the right column. (**F**) Panels at left show Slingshot pseudotime trajectory predictions (Traj A-E) for untreated B cells from Active cGVHD patients (top) and No cGVHD patients (bottom). The origin of the pseudotime analysis (asterisks) was set as Cluster 1 based on our knowledge of its transitional, Igκ signature gene profile (**Figures 1E,F and 2A,B**), suggesting it is the earliest peripheral B cell population emerging from the bone marrow. Panels at right represent each trajectory in isolation (dashed line for reference), along with its associated B cells. B cell clusters are colored and numbered per original unbiased clustering (**Figure 1B,C,I**). Black numbers indicate major clusters that lie along each trajectory, while clusters present but having only a small number of B cells represented are indicated in gray.

### Trajectories for B cell subset diversification in Active cGVHD patients reflect potentially reversible maturation defects

Our past findings suggest that a maturation block is linked to aberrant activation of circulating Active cGVHD B cells via a NOTCH2-BCR signaling axis (*8*). Thus, we performed pseudotime trajectory analysis of our scRNA-Seq dataset using Slingshot (*80*), which may provide insight into the diversification of B cell subsets as they diversify or reach various states of activation. As shown in **Figure 5F** (far left panels), Slingshot predicted 5 pseudotime trajectories (designated Traj A-E) for each allo-HCT patient group. Asterisks indicate the origin (starting point) for the trajectories based on ‘biological knowledge’, assigned to Cluster 1 because of its transitional-like, *IGKC* (Igκ^+^) phenotype. Traj A, B and C were remarkably similar between allo-HCT patient groups (**Figure 5F**, ‘Similar trajectories’). Traj A consisted primarily of B cell Clusters 1, 2, and 4 only, possibly indicating termination at Cluster 4 that had generally low transcripts, consistent with a state of B cell anergy, (**Figure 1E**). Traj B proceeded through Clusters 1, 2, 8, 3, and 9. Traj C proceeded through Clusters 1, 2, 8, 7, and 10, although in Active cGVHD there were notably fewer B cells in Cluster 8. Traj D and E were markedly different between patient groups (**Figure 5F**, ‘Distinct trajectories’). In No cGVHD, Traj D had clear progression through Clusters 1, 2, 8, 3 and 5. By contrast in Active cGVHD, Traj D completely lacked Cluster 3 and had minimal Cluster 8 B cells, proceeding primarily through Clusters 1 and 2, then directly to Cluster 5. Traj E had major contributions from Clusters 1 and 6 in both groups, while Cluster 3 was uniquely prominent in Active cGVHD. These observations suggest that the generation of Cluster 5 ‘memory’ B cells in Active cGVHD patients can potentially occur along a diversification pathway that bypasses tolerance checkpoints, represented by Cluster 8 and Cluster 3 in No cGVHD when tolerance is achieved. Together these DEGs and altered trajectories described above expand on our previous work demonstrating altered maturation in circulating B cells in Active cGVHD (*8*).

Previously, we utilized all-*trans* retinoic acid (ATRA) as a tool *in vitro* to ‘mature’ Active cGVHD B cells by restoring a normal IRF4/IRF8 ratio, attenuating hyper-responsiveness to BCR and NOTCH co-stimulation (*8*). Thus, we assessed the effects of ATRA on B cell clustering, DEGs, and trajectory inferences. A total of 10,000 ATRA-treated B cells per sample were targeted for scRNA-Seq analysis, from the same 8 allo-HCT donor patients described in **Figure 1** and **Supplemental Table 1**. ATRA treatment efficacy was validated by the robust upregulation of known ATRA-responsive genes across multiple B cell clusters including *PLAAT4* (i.e., *RIG1*, *Retinoid-Inducible Gene 1*) and *ASB2* (*81*) [**Supplemental Table 6**]. ATRA treatment resulted in four predicted pseudotime trajectories (designated A-D) in both allo-HCT patient groups (**Figure 6A,B**), compared to the five trajectories for untreated B cells (**Figure 5F**), suggesting a potential narrowing of transcriptional diversity. Supporting this, Cluster 8 was notably absent from the post-ATRA UMAP clusters, while the nine remaining clusters were present (**Figure 6C,D**). Remarkably, the trajectory leading to Cluster 5 in ATRA-treated Active cGVHD B cells (Traj D, **Figure 6A**) included Clusters 1, 2, 7, 3, 10 and 5, which was a markedly different profile from the trajectory leading to Cluster 5 in untreated Active cGVHD B cells (Traj D, **Figure 5F** top panel). This suggests that ATRA, to some extent, normalized the trajectory leading to ABCs and other ‘memory’ B cells when cGVHD was clinically apparent. In all scenarios with ATRA treatment, Cluster 3 preceded Clusters 9, 10 and 5. Our data implicate Cluster 3 as a pivotal precursor memory checkpoint population, and reveal important relatedness and potential plasticity amongst post-BCR-activated B cell subsets.

Interestingly, we found that ATRA affected the distribution of some B cell populations differently between patient groups. The ratio of B cells mapping to Clusters 1 and 2 was much lower for No cGVHD B cells (0.38 and 0.43, respectively) compared to Active cGVHD B cells (0.84 and 0.79, respectively) following ATRA treatment (**Figure 6C,D**). The ratios of B cells mapping to Clusters 7 and 3 were reciprocally changed between groups by ATRA, decreasing in No cGVHD (0.76 and 0.93, respectively) and increasing in Active cGVHD (1.23 and 1.51, respectively). These differences may be explained, at least in part, by reduced viability following ATRA treatment in No cGVHD compared to Active cGVHD B cells (ATRA/Untr total B cell ratio of 0.70 for No cGVHD vs. ATRA/Untr total B cell ratio of 0.93 for Active cGVHD, **Figure 6C,D**). These data affirm important maturation potential of the post-transplant B cell compartment.

Finally, we assessed DEGs following ATRA treatment and found altered expression of a multitude of genes compared to untreated B cells from all 8 allo-HCT patients (**Supplemental Table 6**), although these DEG changes were statistically indistinguishable between groups based on cGVHD disease status. Total DEGs decreased or increased by ATRA are represented numerically in **Figure 6E** and **6G**, respectively. Cluster 7 was most affected by ATRA, with 1,673 Down DEGs compared to untreated B cells (**Figure 6E**, **Supplemental Table 6**). Interestingly, some DEGs in untreated B cells from Active cGVHD patients (Up or Down compared to No cGVHD, **Figure 5C-E**) were significantly changed in the opposite direction by ATRA (**Figure 6F,H**), hinting at potential corrective effects by agents that promote B cell maturation. Both *GPR183* (*EBI2*) and *TNFRSF13B* (*TACI*) were decreased upon ATRA treatment across multiple clusters, representing both ‘transitional’ Clusters 1 and 2, and ‘memory’ Cluster 5 (**Figure 6F**). Thus, ATRA both reshaped the distribution of B cells among clusters and impacted the transcription of some DEGs observed in Active cGVHD.

**Figure 6.**
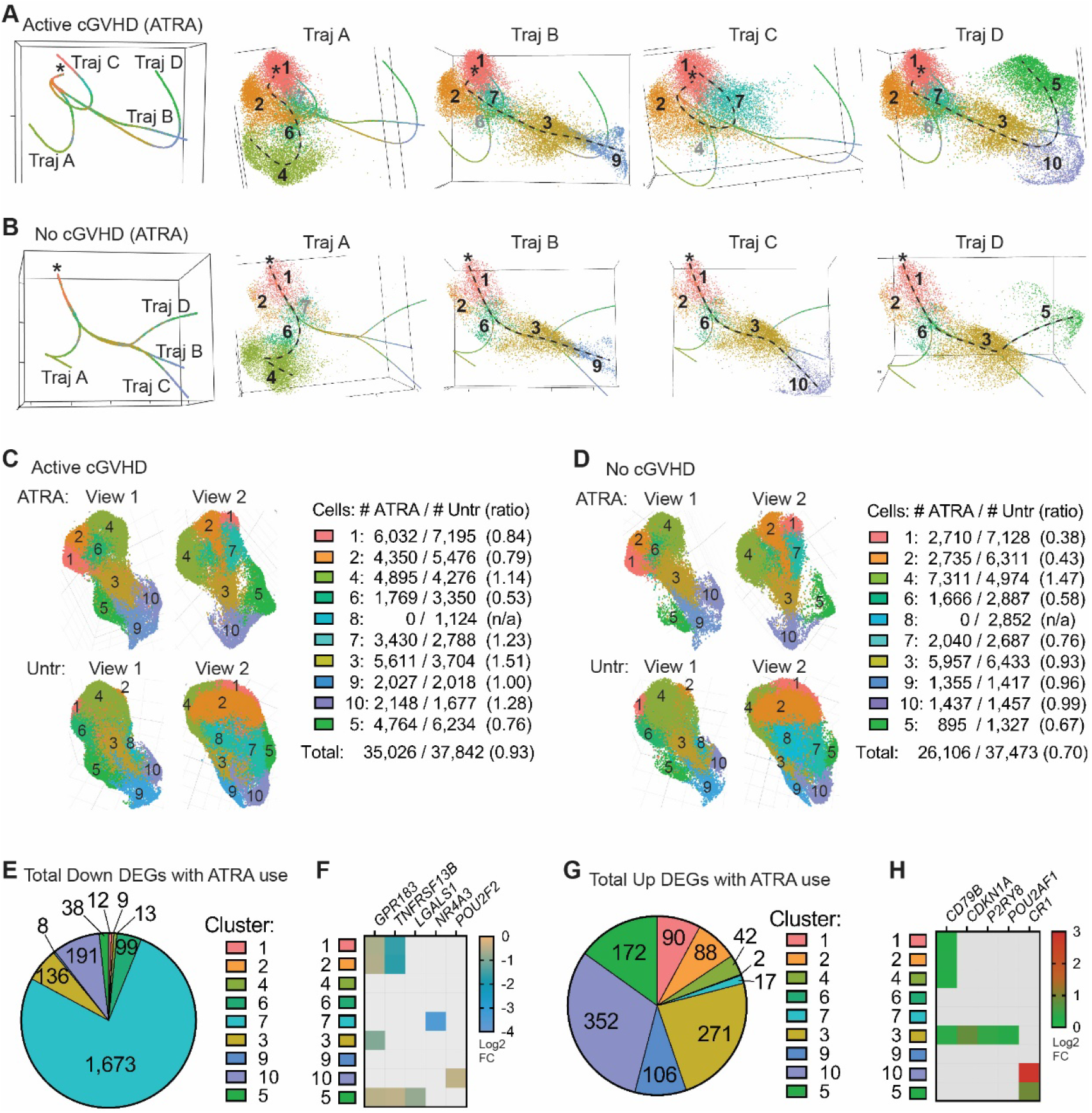
An inducer of cell differentiation, all-*trans* retinoic acid (ATRA), differentially influences B cell pseudotime trajectories and cluster distribution in Active cGVHD. **(A,B)** Panels at left show trajectory estimates (Traj A-D) for B cells from Active cGVHD patients (A) and No cGVHD patients (B) after ATRA treatment. The origin (asterisks) was set as Cluster 1, as in Figure 5F. Panels at right for each patient group represent each trajectory in isolation (dashed line for reference), along with all of its associated B cells (colored and numbered by the cluster to which they belong). Black numbers indicate major clusters that lie along each trajectory, while clusters present but having a small number of B cells represented are indicated in gray. (C,D) ATRA effects on B cell distribution among clusters relative to untreated B cells. UMAP cluster projection and B cell distribution per cluster among ATRA-treated and untreated (Untr) Active cGVHD samples (C), and among ATRA-treated and untreated (Untr) No cGVHD samples (D). Total B cell numbers within each cluster for each treatment group and patient group are indicated. For ATRA-treated B cells, the 9 clusters shown were identified as corresponding to the same 9 clusters in the untreated groups based on signature gene profiles. Ratios represent the number of B cells with ATRA treatment divided by untreated cells (#ATRA / #Untr). (E-H) DEGs induced by ATRA in B cells from all 8 allo-HCT patients in the single-cell RNA-Seq dataset, compared to untreated B cells from all 8 patients. Pie charts indicate the total number of DEGs significantly decreased (E) or increased (G) after ATRA treatment in the cluster indicated. Heat maps (F and H) represent DEGs from Figure 5 that were significantly altered in the opposite direction following ATRA treatment. Colored squares represent significant (*P_adj_* <0.05) log2 FC value for the gene and cluster indicated, for ATRA-treated B cells (all 8 samples) compared to untreated B cells (all 8 samples).

### Broadly dysregulated genes with constitutive activation of associated pathways suggest a common program may underpin B cell diversification in Active cGVHD patients

Since B cell hyper-responsiveness to surrogate antigen is found in a high proportion of total B cells taken from Active cGVHD (*8, 14*), we hypothesized that some DEGs occur broadly in this disease, which might affect B cells at multiple stages of peripheral diversification. As shown in **Figure 7A**, we identified Up DEGs (left) and Down DEGs (right) in the scRNA-Seq dataset that occurred in at least 4 B cell clusters in untreated Active cGVHD B cells. DEGs were ranked based on occurrence in the most clusters, and then by their first occurrence in the least mature/activated clusters. Twenty-nine DEGs were increased and 34 DEGs were decreased in 4 or more clusters in Active cGVHD. Several of these DEGs were already depicted in **Figure 5C-E** because of their known role in B cell function (bold font in **Figure 7A**). To corroborate broad dysregulation we also subjected the scRNA-Seq dataset to a ‘bulk-like’ DEG analysis (all B cells, **Figure 7B**). Indeed, numerous DEGs from **Figure 7A** were also significant the bulk-like analysis (**Figure 7B**, asterisks). We grouped the DEGs from **Figure 7B** empirically into GO-annotated cellular pathways to further highlight potential function (**Supplemental Table 7**). *ARRDC3* was among the most broadly decreased DEGs in Active cGVHD B cells (**Figure 7A,B**). Although the function of this α-arrestin in B cells is unknown, this finding further highlights the potential for dysregulation of GPCRs in Active cGVHD. Also notable, *NFKBIA* (encoding IκBα, the inhibitor of NFκB signaling), was decreased in eight clusters.

*CKS2* stood out among the most broadly increased DEGs in Active cGVHD B cells, reaching significance in eight clusters (**Figure 7A**) and in a bulk-like analysis (**Figure 7B**). We further validated this *CKS2* finding by qPCR analysis on purified B cells from an independent cohort of allo-HCT patients (**Figure 7C**, **Supplemental Table 2**). CKS2 serves as a co-activator of cyclin-dependent kinases (CDKs) (*82*). Alternatively, CKS2 can prevent or delay cell cycle progression by protecting the CDK inhibitor P27^KIP1^ (CDKN1B) from proteasomal degradation via competition for P27^KIP1^ binding with proteins that recruit ubiquitin ligases (*83*). This protection of P27^KIP1^ from degradation by CKS2 averts premature cell cycle entry, preventing apoptosis (*84*). UMAP plot comparisons clearly depict the widespread increase in *CKS2*-positive B cells in Active cGVHD (**Figure 7D**). Likewise, when displayed as normalized expression values (**Figure 7E**), *CKS2* was uniformly increased in Active cGVHD samples in the eight clusters reaching significance (Clusters 1,2,4,8,3,9,10&5).

Given that Active cGVHD B cells have enhanced survival in culture (*6*), the broad increase in *CKS2* hinted that P27^KIP1^ may be more protected from degradation. Employing unbiased protein phosphoarrays, we tested purified B cell lysates from an independent cohort of allo-HCT patients and found phospho-T198 was detected at a 6-to 10-fold greater level in Active cGVHD B cells compared to No cGVHD B cells (**Figure 7F,G** and **Supplemental Figure 6**). Active-site phosphorylation on the kinases AMPK and RSK, known to phosphorylate P27^KIP1^ T198 (*85, 86*), were similar between allo-HCT patient B cells, suggesting increased kinase activity was not responsible for enhanced phospho-T198 in Active cGVHD. Western blot analysis of total protein supported the concept of P27^KIP1^ being protected in Active cGVHD B cells, which had significantly higher levels compared to No cGVHD B cells (**Figure 7H**). These data enable formulation of a model for potential P27^KIP1^ regulation in Active cGVHD B cells (**Figure 7I**), whereby broadly increased CKS2 expression, combined with increased *CDKN1B* (*P27KIP1*) transcription in some subsets (**Figure 5C,E**), may lead to P27^KIP1^ accumulation. Together our dataset allows us to begin to determine functional contributors of enhanced B cell survival in Active cGVHD, and supports prior work showing that cGVHD B cells are primed for proliferation and protected from apoptosis (*6, 8, 14*).

**Figure 7.**
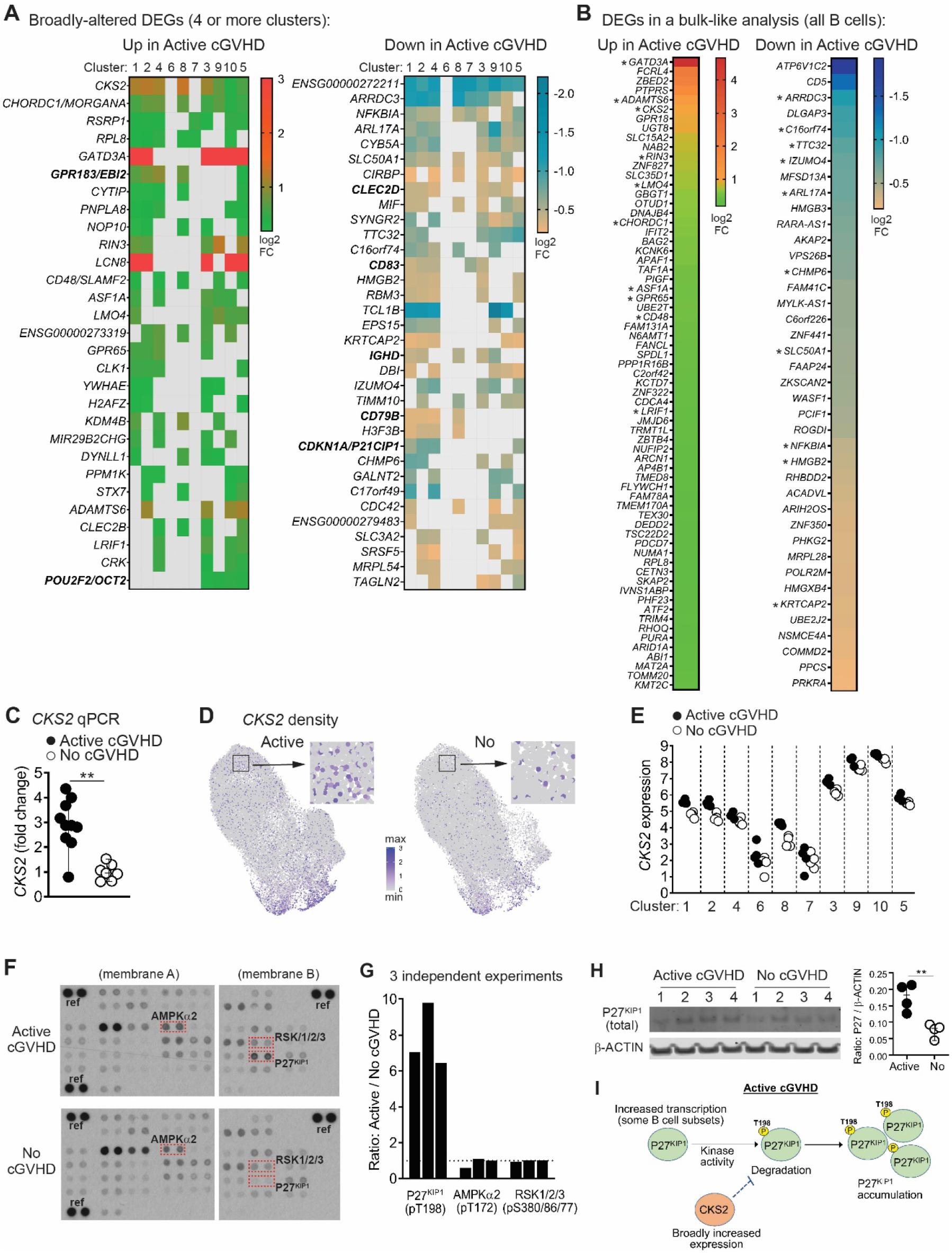
Assessment of DEGs present across the majority of clusters provides additional insight into altered B cell functions in Active cGVHD. (A) DEGs in the scRNA-Seq dataset reaching significance (*P_adj_* <0.05) in four or more B cell clusters as described in Figure 5, either Up (left heat map) or Down (right heat map) in Active cGVHD. Colored squares indicate significance, with log2 FC values as indicated. No DEGs mapped to Cluster 6. DEGs also depicted in Figure 5C-E are shown in bold font. (**B**) DEG analysis performed on total untreated B cells. Heat maps show log2 FC values for genes statistically different (*P_adj_* <0.05) between allo-HCT patient groups (Up or Down in Active cGVHD). (**C**) qPCR analysis of *CKS2* was performed on untreated B cells from a different allo-HCT patient cohort having Active cGVHD (*n*=10) or No cGVHD (*n*=7). Results indicate fold change in *CKS2* expression based on the mean value in the No cGVHD group normalized to 1. *ACTB* (β-ACTIN) was the housekeeping gene. Statistical comparison was performed using a two-tailed Mann-Whitney test (GraphPad Prism 9 software; **, *p*<0.01). (**D**) Normalized expression density UMAP plots for *CKS2* between Active cGVHD (Active) and No cGVHD (No). Representative regions (boxes) were chosen randomly and enlarged (arrows) to visualize single B cells. (**E**) Normalized *CKS2* expression values across all 10 B cell clusters for all 8 allo-HCT patients, separated by disease group. (**F**) Representative phosphoprotein capture arrays for detection of various intracellular signaling molecules phosphorylated on key sites, performed on whole cell lysates of purified, untreated B cells isolated from Active cGVHD (*n*=3) and No cGVHD (*n*=3) patient blood samples (see also **Supplemental Figure 6**). Dashed boxes and protein IDs indicate the location (duplicate spots) of P27^KIP1^ (phospho-T198), AMPKα2 (phospho-T172), and RSK1/2/3 (phospho-S380/S386/S377, respectively). Reference control spots are indicated (‘ref’). (**G**) Combined density results from the 3 independent phosphoprotein array assays shown in (**F**) and **Supplemental Figure 6**. Each bar indicates the results from one experiment and represents the ratio of the average dual spot intensity for Active cGVHD B cells over No cGVHD B cells (dashed line represents a ratio of 1 as a guide). (**H**) Western blot analysis of total P27^KIP1^ protein relative to β-ACTIN in whole cell lysates of B cells isolated from Active cGVHD (*n*=4) and No cGVHD (*n*=4) patient blood samples. Statistical comparisons were performed using a two-tailed, unpaired t-test (GraphPad Prism 9 software; **, *p*<0.01). (**I**) Depiction of a potential model showing how CKS2 expression in Active cGVHD B cells may maintain P27^KIP1^ and lead to accumulation.

## Discussion

Our scRNA-seq analysis reveals and details the breadth of B cell subset and gene expression abnormalities in human cGVHD. While immediate BCR hyper-responsiveness and aberrant upstream signaling in B cells from patients who manifest cGVHD has been shown (*6, 8, 14*), comprehensive analyses of the peripheral B cell compartment have been lacking. The clarity provided herein identifying signature genes that define individual B cell subpopulations, and DEGs associated with Active cGVHD, may support the development of improved targeted therapies for patients. Our work will enable new opportunities for exploitation of the identified molecular pathways to further understand tolerance loss in B cells, in allo-HCT and beyond.

We found that some key DEGs will almost certainly alter the normal trajectory of B cells as they diversify that are likely reflected in additional DEGs that lead to tolerance loss and host reactivity. DEGs in Active cGVHD could be subdivided into two major categories (**Figure 8**, right) that comprise intrinsic differences from a B-cell tolerant state (No cGVHD). The first category of DEGs emerge in B cells in early maturation states that are not constitutively activated (‘DEGs in Naïve Subsets’, Clusters 1, 2, 4, 6, and 8), with some of these DEGs maintained throughout B cell diversification (‘Broadly-Altered DEGs’). The second category of DEGs emerge only after some degree of BCR stimulation (‘DEGs in BCR-Activated Subsets’, Clusters 3, 7, 9, 10 and 5). Examples of unique transcriptional changes included *CKS2* and *GPR183* (*EBI2*), each overexpressed in Active cGVHD from the earliest transitional/naïve B cell subsets to the most differentiated subset, Cluster 5, enriched for ABCs and CD27^+^ memory B cells. Such DEGs occurring from (at least) the time B cells first enter the circulation may cause early epigenetic changes that alter subsequent tolerance checkpoints and potentially affect clonal diversity.

**Figure 8.**
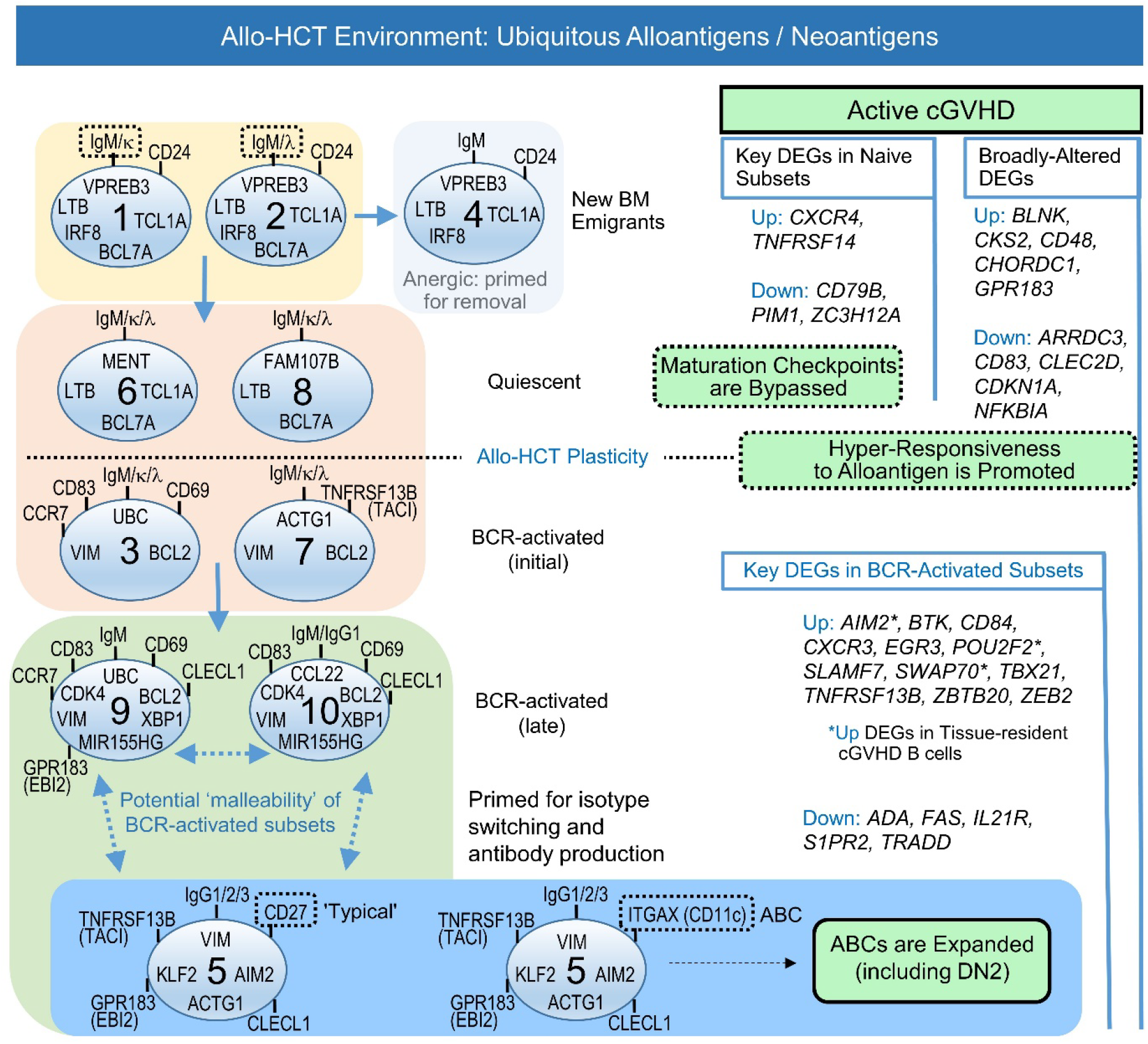
Working model for extrinsic activation signals and intrinsic gene changes influencing B cell maturation, survival, and proliferative capacity that lead to the expansion of pathologic ‘memory’ B cells in Active cGVHD. Cluster-defining ‘signature’ genes (left side) and trajectory analyses (Figure 5F) led to assignment of the 10 clusters along a logical pathway of B cell diversification in the periphery of allo-HCT patients, from newly-emigrated populations, through proposed checkpoint steps where survival decisions occur, and finally to BCR-activated and ‘memory’ states. Together, data suggest that extrinsic activation signals, including ubiquitous alloantigens and cytokines like BAFF, enable gene changes generally in the allo-HCT environment relative to the non-HCT setting that leads to ‘plasticity’, and potential therapeutic ‘malleability’, of the B cell compartment. DEGs that influence diversification checkpoints and BCR hyper-responsiveness can occur early, broadly or exclusively after BCR stimulation, leading to pathogenesis characterized by ABC expansion in Active cGVHD (right side).

Extrinsic factors in microenvironmental niches may be the initial drivers of such DEGs that lead to early loss in peripheral B cell tolerance, and that can be elucidated through studies of circulating B cells.

The observed expansion of potentially pathogenic ABCs in Active cGVHD patients (**Figure 4**, and as modeled in **Figure 8**: “ABCs Expanded (including DN2)”) is likely a consequence of gene alterations before and after BCR-engagement occurring in a cumulative way. ABCs have emerged as important in both normal immunity and autoimmunity (*20, 21, 25, 26*). These subsets likely directly contribute to pathogenesis by producing alloantibodies after migrating to tissue sites where cGVHD manifests (*10*). The expansion of ABCs (including DN2) is driven by one or more costimulatory signals that cooperate with BCR stimulation (*87–90*). Indeed, B cells isolated from Active cGVHD patients are hyper-responsive to minimal BCR ligation in synergy with co-stimulatory molecules including NOTCH2 and BAFF (*8*). These processes are likely driven in part by the increased BAFF-to-B cell ratio that occurs in cGVHD patients (*9, 10, 63, 91*), and by niche production of BAFF in the SLO, confirmed recently in a mouse cGVHD model (*7*). BAFF influence may additionally propagate by increases in BAFF receptors such as TACI (encoded by the *TNFRSF13B* gene), as published (*10*) and herein described for specific B cell subsets. It is also noteworthy that multiple other TNF superfamily and TNF-receptor superfamily members were increased DEGs in cGVHD B cell clusters, including the metabolically active Clusters 9 and 10. Therefore, ongoing studies of these molecules as potential drivers of ABC expansion in cGVHD is warranted.

‘Plasticity’ or ‘malleability’ has been demonstrated in newly formed B cells and during the generation of memory (*92, 93*). Our scRNA-Seq dataset and supporting work provides new insight into these processes (**Figure 8**, ‘Allo-HCT Plasticity’). Signature gene profiles defined 10 distinct B cell subsets (**Figure 8**, left) that continually develop and circulate under constant exposure to alloantigens. Furthermore, Allo-HCT patient ‘memory’ B cell subsets had some remarkably distinct gene features, regardless of cGVHD status, compared to the same subsets from healthy individuals or patients chronically exposed to microbial antigens (**Figure 3**). This direct comparison of our allo-HCT patient scRNA-Seq dataset with a publicly-available dataset on blood B cells isolated from HDs and individuals with chronic infections (HIV, malaria), provided new insight into the unique aspects of the B cell pool that persist following immune reconstitution with allo-HCT therapy. In allo-HCT, striking differences in the proportion of B cells expressing some molecules that orchestrate exit from the BM and homing to/establishment in SLO niches, along with molecules affecting survival, suggest an impact on the unique clinical manifestations of cGVHD. These observations may additionally reflect the apparent maturation block associated with B cell hyper-responsiveness in cGVHD patients (*8*), and may partially explain the paucity of CD27^+^ follicular memory B cells in allo-HCT patients compared to HDs (**Figure 4B,C**). These data are also consistent with published data showing that the peripheral B cell pool in allo-HCT patients is potentially ‘malleable’ (**Figure 8**, ‘Potential malleability of BCR-activated subsets’), and as such is possibly amenable to novel treatment strategies to prevent or treat cGVHD, while maintaining immunity against pathogens and possibly malignant cells (*8*). This unique plasticity of the B cell compartment in an allo-HCT environment can now be modeled based on our scRNA-seq data.

Identification in this study of certain molecules as dysregulated in subsets of Active cGVHD B cells is potentially novel, particularly those that regulate B cell movement and positioning within SLO. The identification of *GPR183* (*EBI2*), *P2RY8*, and *S1PR2* as DEGs in one or more clusters of B cells is particularly remarkable, as each molecule is important for follicular movement and GC positioning of B cells (*39, 68–71*). The increase in *GPR183* across five B cell clusters, including transitional-like subsets, suggests that strictly regulated tolerance mechanisms that are initiated as B cells first recirculate and enter the SLO microenvironment may be disrupted in cGVHD patients. This is consistent with the notion that altered intrinsic functions of lymphocytes and stromal cells within SLOs disrupt normal architecture during follicle organization and GC development, potentially leading to premature displacement of B cells to extrafollicular spaces and altered tolerance (*74*). Remarkably, *GPR183*, *P2RY8*, and *S1PR2* were each DEGs in Cluster 5, which was enriched for ABCs and DN2. *P2RY8* expression was additionally decreased in metabolically active Cluster 9, intriguing given recent evidence that decreased B cell *P2RY8* propagates pathogenic antibody-producing plasma cells and ABCs in humans with lupus or antiphospholipid syndrome (*73*). The ligands for each of these molecules are known, and in some cases inhibitors and agonists have been described, permitting future studies in mouse models of cGVHD and eventually clinical trials. This concept is supported by an observation in a mouse lung transplant model of bronchiolitis obliterans (BO), a condition that frequently causes lung damage in cGVHD patients, where inhibition of GPR183 with a highly selective inhibitor reduced pulmonary lymphoid lesions and lung damage (*94*). Elucidating B cell transcriptional changes that help incite cGVHD development, versus transcriptional changes that occur after cGVHD genesis and nevertheless contribute to ongoing pathogenesis in an unrelenting cycle, will remain an area of future investigation.

While the full implications of the findings herein remain to be fully realized, our data provide a foundation for future assessment of novel, potentially targetable pathways that alter B cell homeostasis during the diversification of B cells in the periphery when alloantigens or neo-autoantigens are prevalent. Mouse studies can now be launched that evaluate the hierarchy of these molecules in initiating or propagating loss of B cell tolerance. The ultimate benefit of this research will be the ability to design tailored therapies that eliminate or prevent the emergence of pathogenic B cells without disrupting normal humoral immune responses required for long-term health.

## Methods

### Patient blood and skin samples

De-identified whole blood samples or apheresis samples were obtained from allo-HCT patients under IRB protocols from Duke University, the National Cancer Institute (NIH), or the Dana-Farber Cancer Institute. Informed consent was obtained from all patients, with the scope of the research and the minimal risks fully explained. Beyond some basic inclusion and exclusion criteria given to clinicians for obtaining PBMC samples for study (including cGVHD status), samples were obtained and chosen for use in a blinded fashion. For all supporting experiments, allo-HCT patient samples were likewise obtained and chosen for use in a blinded fashion. A skin punch biopsy was collected from an adult allo-HCT patient with Active cGVHD in accordance with an IRB protocol approved by Duke University. Likewise, a skin punch biopsy from surgically discarded abdominoplasty tissue from a HD was obtained in accordance with an IRB protocol approved by Duke University.

### Experimental procedures

See Supplementary Methods for information on B cell isolation and culture, scRNA-Seq sample preparation, additional information on scRNA-Seq data analysis, dermal skin B cell isolation and scRNA-Seq analysis, qPCR, flow cytometry and phenograph analyses, phosphoarray sample preparation and analysis, and Western blot analysis.

### Statistical analysis of single-cell RNA-Seq data

Clustering analysis was conducted using R statistical environment and extension package Seurat v3.1.4 (*22*). DEG analysis with respect to disease status (Active cGVHD vs. No cGVHD) within each cluster was performed based on the read counts across all cells within each sample using the Bioconductor package DESeq2 (version 1.26.0). Global interaction between disease status and the B cell clusters also was analyzed. Multiple testing was accounted for within the framework of control of the false discovery rate (FDR) using the Q-value approach (version 2.14.1) (*95*).

### Data and Code availability

All blood B cell scRNA-Seq library data from the 8 allo-HCT patients (16 samples) are available through the Gene Expression Omnibus (GEO) database (https://www.ncbi.nlm.nih.gov/geo/) under accession code GSE161343. The R scripts to reproduce the analyses of these scRNA-Seq data are available at this site: https://gitlab.oit.duke.edu/dcibioinformatics/pubs/sarantopoulos-10x-cgvhd/.

## Supplemental Materials

Supplemental Figures 1-6

Supplemental Tables 1, 2, 4, 5 and 6 (combined PDF)

Supplemental Tables 3 and 6 (individual spreadsheets)

Supplemental Methods

## Acknowledgments

The authors wish to thank Laney Weber, Certified Editor in the Life Sciences, for the critical review of this manuscript and assistance with formatting. The authors also wish to thank Lauren S. Riley and Serach Patterson for assistance with proofreading.

## Author contributions

Conceptualization: JCP, SS

Data curation: JF, MRL, KO

Formal analysis: JF, DZ, MRL, XQ, JX, KO

Funding acquisition: SS, SJB, JV

Investigation: JCP, RAD, HS, JZ, ANS, ICC

Methodology: JF, DZ, MRL, RAD, XQ, JV, JX, KO

Resources: VTH, KSW, JJR, SZP, FTH, MEH, DAR, ARC

Software: JF, DZ, MRL, XQ, JX, KO

Supervision: SS, KO Validation: JF, MRL, XQ, KO

Visualization: JCP, JF, JV, KO, SS

Writing – original draft: JCP, SS, KO

Writing – review & editing: JF, DZ, MRL, RAD, XQ, JZ, JV, SJB, WJ, WM, NJC, ARC, JX

