## Supplemental Figures 1-6 for "Single-cell Landscape Analysis Unravels Molecular Programming of the Human B Cell Compartment in Chronic GVHD"

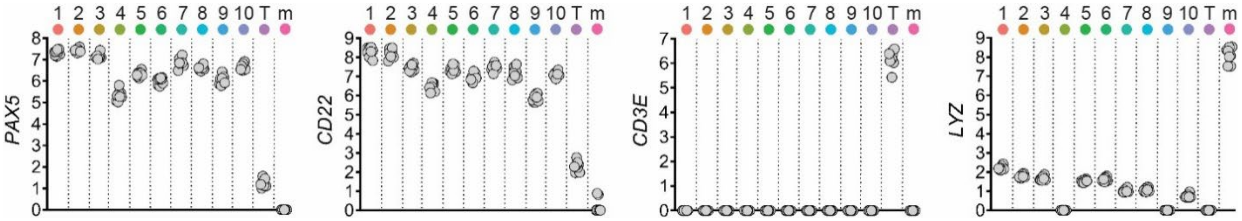


**Supplemental Figure 1. Expression of genes with known specificity for B cells (*PAX5*, *CD22*), T cells (*CD3E*), or monocytes (*LYZ*) confirms cluster lineage identity.** Log2 normalized expression values for lineage-specific genes of interest. Numbers and colored dots at top represent the corresponding B cell cluster, with letters and colored dots representing clusters for residual T cells (T) and monocytes (m). Each symbol (gray circle) represents log2 normalized expression results from one of the 8 total allo-HCT patients assessed.

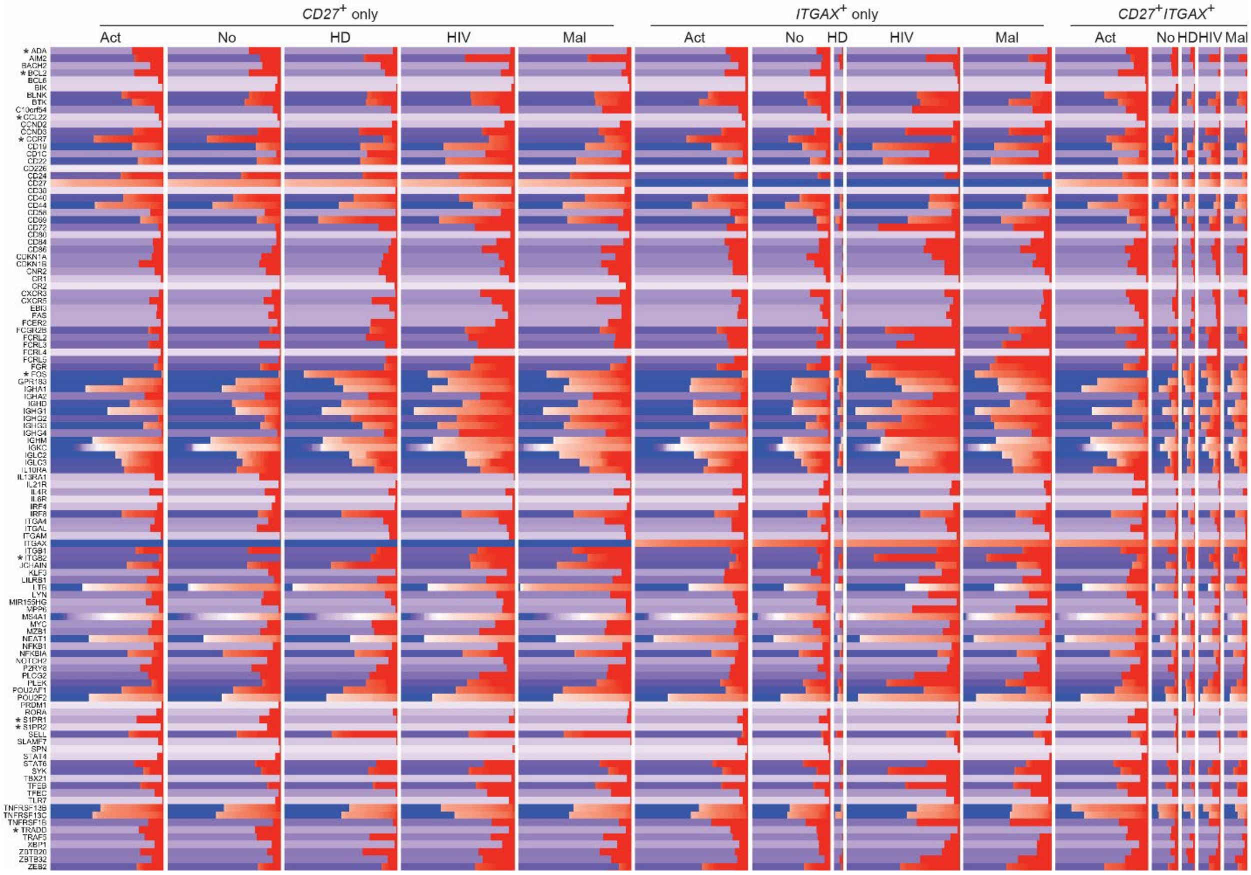


**Supplemental Figure 2. The proportion of B cells expressing signature genes that define memory B cells subsets in allo-HCT patients compared to healthy donors (HD) or patients with infectious disease (HIV, Malaria [Mal]).** Genes on the Y-axis were chosen based on a thorough search of the literature for those known to be associated with memory B cell subsets. Memory B cell subsets were defined by *CD27* expression only (*CD27*$^{+}$*ITGAX*$^{-}$, left 5 columns), by *ITGAX* (*CD11C*) expression only (*CD27*$^{-}$*ITGAX*$^{+}$, middle 5 columns), or co-expression of both (*CD27*$^{+}$*ITGAX*$^{+}$, right 5 columns). Individual columns represent the patient or healthy donor group indicated. Colors represent log counts for each gene per million total reads in the same B cell, reported as log counts per million (log CPM). Red hues indicate that a B cell was positive for the gene of interest, and so the relative amount of red to blue is representative of the % positive B cells for that gene within the corresponding subset and group. Each column represents a maximum of 100 B cells, with fewer B cells shown in groups where < 100 B cells mapped to the subset of interest (indicated by a narrower column width). Where > 100 B cells were identified, 'downsampling' was performed to reduce the maximum number B cells shown in the column to 100 for visual purposes, as described in the Methods section. Asterisks indicate those genes also depicted in **Figure 3** of the main paper.

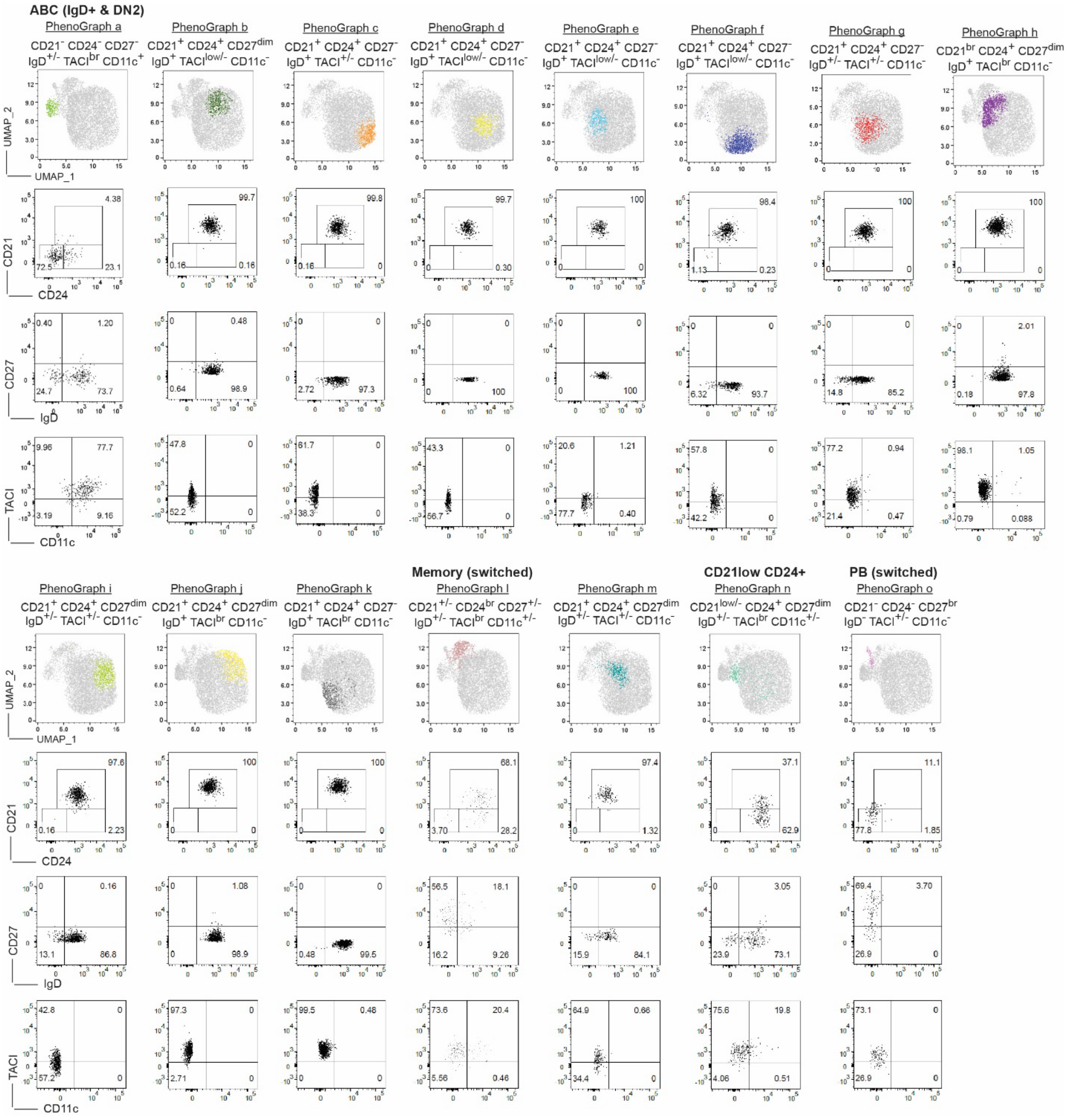


**Supplemental Figure 3. Surface protein analysis by flow cytometry and PhenoGraph provides novel resolution of B cell subpopulations in allo-HCT patients.** PhenoGraph results as described in **Figure 4F-J** for the 15 identified B cell clusters from a representative allo-HCT patient in the 8-allo-HCT patient cohort. PhenoGraph UMAP projections are shown in the top panels, with the cluster of focus shown in color. The corresponding flow cytometric results for that cluster are shown in the dot plots below each UMAP. Numbers in the dot plots represent the frequency of B cells within the corresponding gate. ABC, memory, and other populations of interest including plasmablasts (PB) are labeled above their corresponding PhenoGraphs.

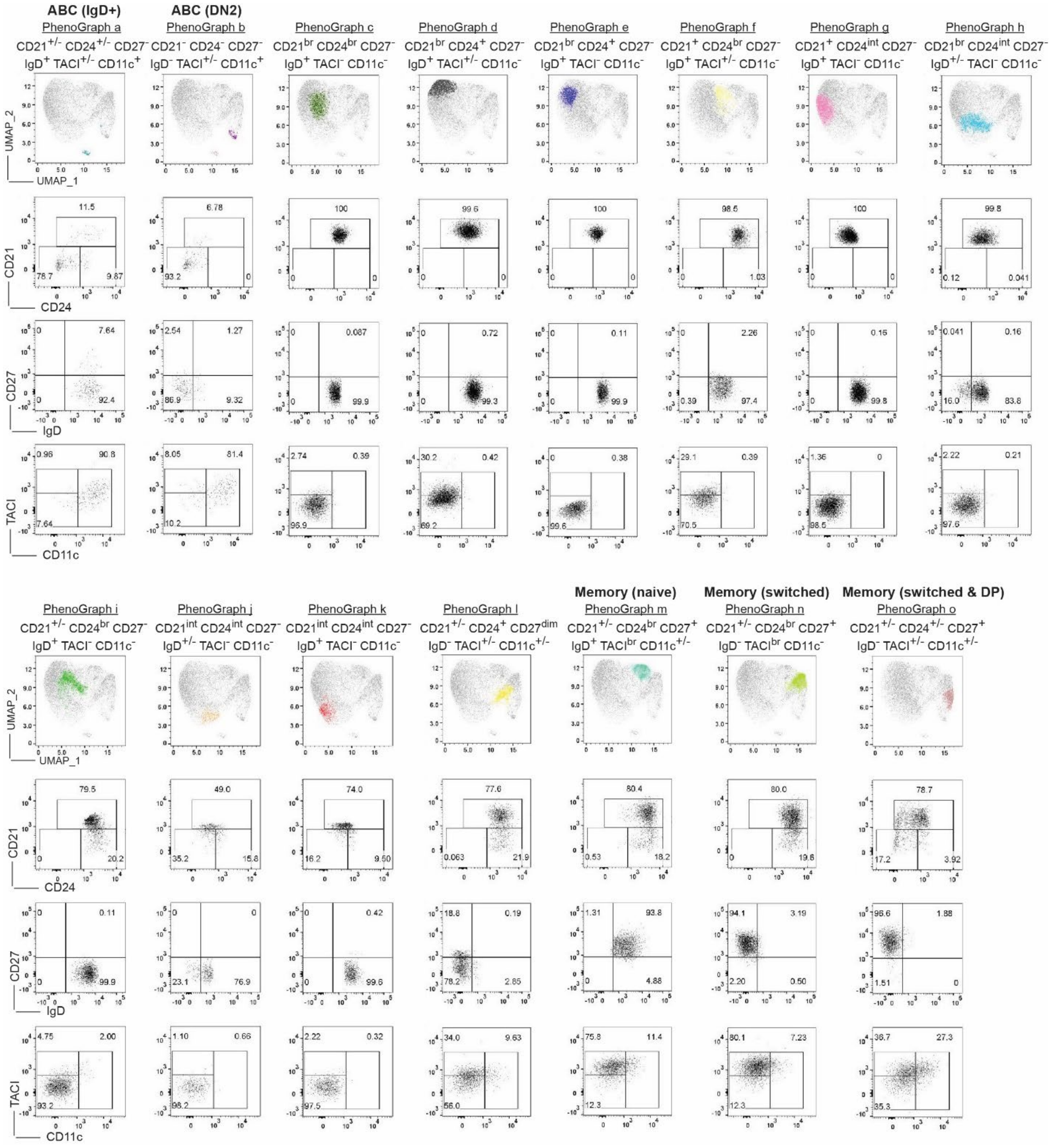


**Supplemental Figure 4. Surface protein analysis by flow cytometry and PhenoGraph on HD peripheral B cells reveals subset distinctions with allo-HCT patients.** PhenoGraph results as described in **Figure 4F-J** and **Supplemental Figure 3** showing representative results from 1 of 4 healthy donor (HD) PBMC samples assessed. ABC and memory populations of interest are labeled above corresponding PhenoGraphs.

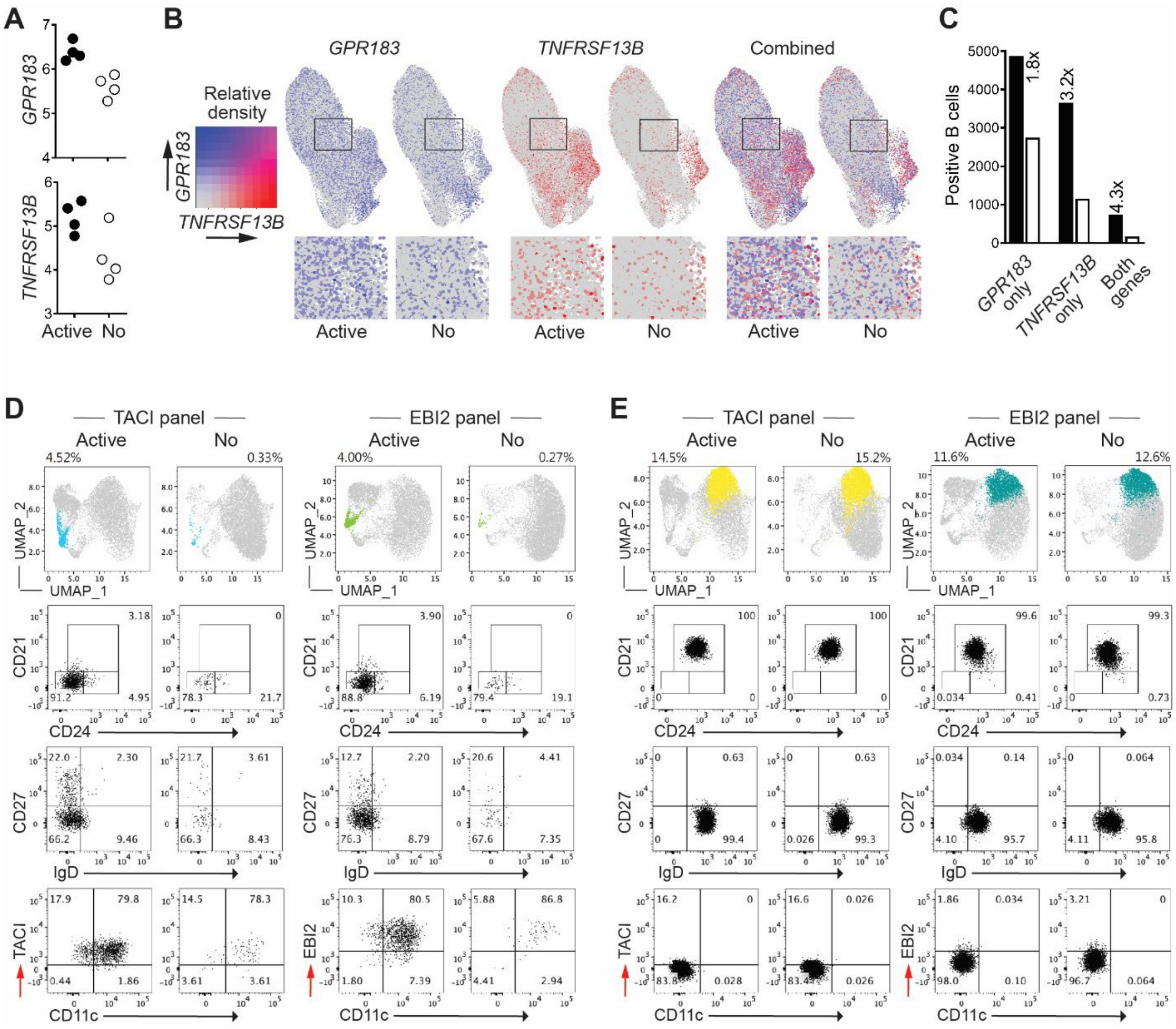


**Supplemental Figure 5. B cells expressing *GPR183, TNFRSF13B* or both genes are increased in Active cGVHD.** (**A**) Log2 normalized expression values for *GPR183* (*EBI2*) and *TNFRSF13B* (*TACI*) within Cluster 8 are shown, correlating with the observed significant difference between allo-HCT patient groups within this cluster (**Figure 5A**). Each symbol represents one patient within the group indicated (Active, Active cGVHD; No, No cGVHD). (**B,C**) Co-expression analysis of *GPR183* and *TNFRSF13B* using Seurat provides insight into numbers of B cells separately expressing and co-expressing these genes. In (**B**), normalized expression density UMAP plots are shown for *GPR183* only (left panels), *TNFRSF13B* only (middle panels), and both genes in combination (right panels). The relative expression level of each gene in the density plots is based on the minimum to maximum color threshold range shown at the left. B cells co-expressing the genes at a high threshold level in the combination plots are depicted by magenta color. The boxed areas in the density plots approximate the B cells in Cluster 8 and are enlarged at the bottom to more easily visualize single B cells individually expressing or co-expressing the two genes. In (**C**), all B cells in the scRNA-Seq dataset individually expressing or co-expressing *TNFRSF13B* and *GPR183* were quantitated using this available function in Seurat and separated by patient group. Bars represent total B cells expressing one or both genes as indicated by the X-axis labels, with the ratio of Active cGVHD B cells (filled bars) to No cGVHD B cells (open bars) indicated by the number above each condition, highlighting the increase in Active cGVHD. (**D,E**) PhenoGraph analyses of representative allo-HCT patients as described in **Figure 4** and **Supplemental Figure 3** for identical flow cytometry panels with the exception of the inclusion of antibodies to either TACI (left plots) or EBI2 (right plots), as indicated by the red arrows. Cluster prediction was similar for the 2 panels, with 15 clusters predicted in both cases (UMAP plots at top, and not shown), enabling the comparison of similar clusters of B cells between panels. In (**D**), a similarly positioned and shaped cluster representing ABCs is present for both the TACI panel (blue) and EBI2 panel (green) that is otherwise phenotypically similar based on the other markers used, and is increased in frequency in Active cGVHD B cells (percentages indicated above each UMAP plot, and as with ABCs described in **Figure 4** and **Supplemental Figure 3**). For comparison, (**E**) shows a different cluster in the PhenoGraph projections (yellow, TACI panel; aqua, EBI2 panel) that has a naïve follicular B cell phenotype, lacks expression of TACI and EBI2, and is similar in frequency between patient groups.

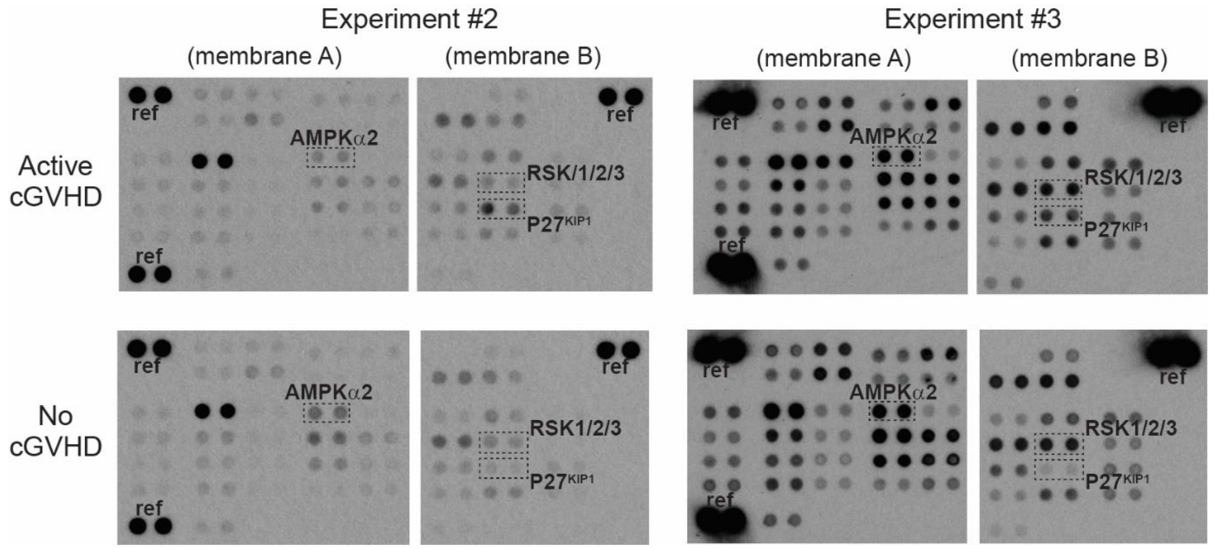


**Supplemental Figure 6. The 2 additional experiments represented in the bar graph in Fig. 7G (along with the experiment shown in Fig. 7F) demonstrating P27$^{Kip1}$ phosphorylation at a major regulatory site is enhanced in Active cGVHD patient B cells.** Phosphoprotein capture arrays (Proteome Profiler™ Human Phospho-Kinase Array) for detection of various intracellular signaling molecules phosphorylated on key sites involved in their regulatory activity, performed on whole cell lysates of purified, unstimulated B cells isolated from Active cGVHD and No cGVHD patient PBMC samples. Dashed boxes and protein IDs indicate the location and assay results for duplicate spots of capture antibodies against P27$^{KIP1}$ (phospho-T198), AMPKα2 (phospho-T172), and RSK1/2/3 (phospho-S380/S386/S377, respectively). Reference control spots on the arrays are indicated (ref).
