## Supplemental Tables 1,2,4,5 and 7 for "Single-cell Landscape Analysis Unravels Molecular Programming of the Human B Cell Compartment in Chronic GVHD"

**Supplemental Table 1. Characteristics of patients whose samples were used to generate the single-cell RNA-Seq dataset.**

| Characteristic | No cGVHD (*n*=4) | Active cGVHD (*n*=4) | *p* |
|---|---|---|---|
| Median age, years (range) | 37 (30-48) | 47 (22-61) | 0.54 |
| Sex, no. (%) of males | 2 (50) | 4 (100) | 0.43 |
| Median time after transplant, mos (range) | 11.5 (10.0-12.3) | 12.6 (11.1-13.9) | 0.16 |
| *Conditioning regimen (%)* | | | 1 |
| Non-myeloablative | 4 (100) | 4 (100) | |
| *Source of graft (%)* | | | 1 |
| Peripheral blood | 4 (100) | 4 (100) | |
| *HLA matching (%)* | | | 1 |
| Matched (MRD, MUD) | 4 (100) | 4 (100) | |
| Mismatched | 0 (0) | 0 (0) | |
| *Immunosuppressive treatment (%)* | | | |
| Steroidal (Prednisone) | 0 (0) | 2 (50) | 0.43 |
| Other IST | 1 (25) | 4 (100) | 0.14 |
| *Initial disease (%)* | | | |
| HD | 3 (75) | 1 (25) | |
| Thalassemia | 1 (25) | 0 (0) | |
| CML | 0 (0) | 1 (25) | |
| NHL | 0 (0) | 1 (25) | |
| Diamond Blackfan Anemia | 0 (0) | 1 (25) | |

All patients provided consent, and all studies were approved under IRB protocols of Duke University, The National Institutes of Health, and The Dana-Farber Cancer Institute. Statistical comparisons between groups were performed using two-tailed, unpaired Student's t-test (age, time after transplant) or Fisher's exact test (other comparisons). cGVHD, chronic graft versus host disease; HLA, human leukocyte antigen; MRD, matched related donor; MUD, matched unrelated donor; IST, immunosuppressive therapy; HD, Hodgkin's disease; CML, chronic myeloid leukemia; NHL, non-Hodgkin lymphoma.

**Supplemental Table 2. Characteristics of patients who provided samples for supporting experiments.**

| Characteristic | No cGVHD (*n*=19) | Active cGVHD (*n*=14) | *p* |
|---|---|---|---|
| Median age, year (range) | 53 (24-73) | 52 (22-69) | 0.88 |
| Sex, no. (%) of males | 10 (58) | 9 (64) | 0.72 |
| Median time after transplant, mos (range) | 19 (11-73) | 48 (13-112) | 0.013 |
| *Conditioning regimen (%)* | | | 0.73 |
| Myeloablative | 11 (58) | 7 (50) | |
| Non-myeloablative/reduced | 8 (42) | 7 (50) | |
| *Source of graft (%)* | | | 1 |
| Peripheral blood | 17 (89) | 12 (86) | |
| Bone marrow | 2 (11) | 2 (14) | |
| *HLA matching (%)* | | | 0.42 |
| Matched (MRD, MUD) | 19 (100) | 13 (93) | |
| Mismatched | 0 (0) | 1 (7) | |
| *Immunosuppressive treatment (%)* | | | |
| Steroidal (Prednisone) | 2 (11) | 8 (57) | 0.007 |
| Other IST | 1 (5) | 7 (50) | 0.005 |
| *Initial disease* | | | |
| ALL | 2 | 1 | |
| AML/AML from MDS | 6 | 3 | |
| Burkitt's lymphoma | 0 | 1 | |
| CLL | 1 | 0 | |
| CML | 1 | 1 | |
| DLBCL | 1 | 0 | |
| MDS/MF | 4 | 3 | |
| MM | 1 | 0 | |
| T-Cell lymphoma | 1 | 1 | |
| Other | 2 | 4 | |

All patients provided consent, and all studies were approved under IRB protocols of Duke University, The National Institutes of Health, and The Dana-Farber Cancer Institute. Statistical comparisons between groups using two-tailed, unpaired Student's t-test (age, time after transplant) or Fisher's exact test (other comparisons). cGVHD, chronic graft versus host disease; HLA, human leukocyte antigen; MRD, matched related donor; MUD, matched unrelated donor; IST, immunosuppressive therapy; ALL, acute lymphoblastic leukemia; AML, acute myeloid leukemia; MDS, myelodysplastic syndrome; CLL, chronic lymphocytic leukemia; CML, chronic myeloid leukemia; DLBCL, diffuse large B-cell lymphoma; MF, myelofibrosis; MM, multiple myeloma.

**Supplemental Table 4. Signature genes reaching significance in the skin cell scRNA-Seq dataset for the cluster identified as B cells.** LogFC values represent the expression level of the gene of interest in the B cell cluster relative to clusters for all other skin cell lineages. Some genes with particular relevance to the B cell lineage are highlighted in yellow.

| gene | avg_logFC | pct.1 | pct.2 | p_val | p_val_adj | Additional description |
|---|---|---|---|---|---|---|
| *IGKC* | 5.840324113 | 0.5 | 0.041 | 5.91E-69 | 1.35E-64 | Ig kappa light chain |
| *IGHG1* | 4.48271604 | 0.167 | 0.002 | 1.01E-127 | 2.31E-123 | IgG1 heavy chain |
| *IGHG3* | 3.890657725 | 0.241 | 0.003 | 6.89E-187 | 1.57E-182 | IgG3 heavy chain |
| *GZMB* | 2.908180123 | 0.556 | 0.049 | 6.89E-70 | 1.57E-65 | |
| *PLAC8* | 2.838899102 | 0.556 | 0.023 | 5.90E-146 | 1.35E-141 | |
| *IGHG4* | 2.568618359 | 0.167 | 0.004 | 2.98E-82 | 6.81E-78 | IgG4 heavy chain |
| *TCL1A* | 2.384288612 | 0.222 | 0.002 | 6.96E-224 | 1.59E-219 | |
| *LTB* | 2.31135052 | 0.537 | 0.097 | 3.62E-32 | 8.26E-28 | |
| *IRF8* | 2.065864406 | 0.37 | 0.071 | 2.23E-19 | 5.08E-15 | |
| *RGS2* | 1.968658624 | 0.426 | 0.175 | 8.35E-09 | 0.000190494 | |
| *ALOX5AP* | 1.923027434 | 0.407 | 0.098 | 1.78E-15 | 4.07E-11 | |
| *IGHM* | 1.877102155 | 0.204 | 0.005 | 4.60E-91 | 1.05E-86 | IgM heavy chain |
| *SPIB* | 1.844424329 | 0.296 | 0.012 | 1.58E-77 | 3.61E-73 | |
| *IRF7* | 1.835136805 | 0.352 | 0.12 | 6.22E-09 | 0.000141869 | |
| *PLD4* | 1.828996003 | 0.204 | 0.018 | 2.76E-24 | 6.30E-20 | |
| *GPR183* | 1.728443737 | 0.407 | 0.121 | 1.66E-11 | 3.79E-07 | a.k.a. EBI2 |
| *MZB1* | 1.690247628 | 0.241 | 0.012 | 4.90E-54 | 1.12E-49 | Marginal Zone B and B1 Cell-Specific Protein |
| *ITM2C* | 1.655431722 | 0.481 | 0.14 | 1.21E-14 | 2.77E-10 | |
| *LILRA4* | 1.632328535 | 0.167 | 0.007 | 1.81E-46 | 4.13E-42 | |
| *AREG* | 1.587745467 | 0.315 | 0.089 | 1.18E-09 | 2.70E-05 | |
| *UCP2* | 1.483850276 | 0.407 | 0.125 | 9.11E-11 | 2.08E-06 | |
| *CYBA* | 1.446793637 | 0.611 | 0.334 | 2.11E-10 | 4.82E-06 | |
| *NCF1* | 1.393078587 | 0.37 | 0.059 | 1.48E-22 | 3.38E-18 | |
| *FCHSD2* | 1.382376321 | 0.278 | 0.09 | 4.37E-07 | 0.009977268 | |
| *BCL11A* | 1.355831393 | 0.241 | 0.024 | 6.30E-26 | 1.44E-21 | |
| *MPEG1* | 1.270548263 | 0.167 | 0.041 | 2.11E-06 | 0.048204372 | |
| *GAPT* | 1.242264259 | 0.204 | 0.009 | 1.19E-50 | 2.71E-46 | |
| *SELL* | 1.223975793 | 0.204 | 0.044 | 6.14E-09 | 0.000140149 | |
| *TYROBP* | 1.199836161 | 0.296 | 0.111 | 2.13E-06 | 0.048492729 | |
| *MS4A1* | 1.158058622 | 0.185 | 0.004 | 2.23E-87 | 5.08E-83 | a.k.a. CD20 |
| *DERL3* | 1.094782318 | 0.222 | 0.005 | 1.49E-105 | 3.40E-101 | |
| *MARCHF1* | 1.094421244 | 0.241 | 0.046 | 4.01E-12 | 9.16E-08 | |
| *CD79A* | 1.057818646 | 0.259 | 0.002 | 1.26E-295 | 2.88E-291 | Immunoglobulin-Associated Alpha |
| *PRKCB* | 1.041053875 | 0.222 | 0.063 | 6.85E-07 | 0.015628941 | |
| *SIDT1* | 1.034585484 | 0.111 | 0.018 | 1.36E-07 | 0.003106374 | |
| *IRF4* | 1.032109579 | 0.185 | 0.028 | 3.30E-12 | 7.53E-08 | |
| *BANK1* | 1.023041358 | 0.204 | 0.016 | 1.68E-28 | 3.83E-24 | B-Cell Scaffold Protein with Ankyrin Repeats |
| *ADAM19* | 1.017732552 | 0.204 | 0.033 | 1.79E-12 | 4.08E-08 | |
| *NEK8* | 0.977455409 | 0.111 | 0.019 | 4.18E-07 | 0.009546737 | |
| *NUP210* | 0.89893479 | 0.148 | 0.026 | 1.45E-08 | 0.000331422 | |
| *CYBB* | 0.873497434 | 0.185 | 0.044 | 4.22E-07 | 0.009631 | |
| *FAM129C* | 0.859258535 | 0.167 | 0.003 | 1.15E-101 | 2.63E-97 | |
| *ARHGAP24* | 0.805663503 | 0.185 | 0.045 | 3.13E-07 | 0.007149293 | |
| *LY9* | 0.799524896 | 0.185 | 0.025 | 4.51E-14 | 1.03E-09 | |
| *TRAF4* | 0.792841267 | 0.204 | 0.04 | 8.04E-10 | 1.83E-05 | |
| *EVI2B* | 0.787922656 | 0.278 | 0.092 | 1.02E-06 | 0.023281725 | |
| *CD74* | 0.777773836 | 0.833 | 0.555 | 2.82E-09 | 6.42E-05 | |
| *RALGPS2* | 0.769373394 | 0.259 | 0.03 | 8.14E-23 | 1.86E-18 | |
| *HLA-B* | 0.684407312 | 0.907 | 0.753 | 5.73E-08 | 0.001307338 | |
| *BTK* | 0.639524637 | 0.185 | 0.034 | 6.32E-10 | 1.44E-05 | Bruton Tyrosine Kinase |
| *TLR10* | 0.61921292 | 0.167 | 0.004 | 9.48E-69 | 2.16E-64 | |

| | | | | | | |
|---|---|---|---|---|---|---|
| *BLK* | 0.616529536 | 0.148 | 0.002 | 1.12E-109 | 2.55E-105 | B Lymphocyte Kinase |
| *LY86* | 0.58109155 | 0.167 | 0.025 | 1.60E-11 | 3.66E-07 | |
| *BLNK* | 0.574324462 | 0.167 | 0.019 | 7.72E-16 | 1.76E-11 | B Cell Linker |
| *DOK3* | 0.539394416 | 0.111 | 0.014 | 2.42E-09 | 5.53E-05 | |
| *ATP2A3* | 0.528046235 | 0.222 | 0.049 | 4.35E-09 | 9.94E-05 | |
| *ITGB7* | 0.462564875 | 0.185 | 0.046 | 1.20E-06 | 0.02727237 | |
| *GPR18* | 0.420638934 | 0.13 | 0.011 | 3.96E-16 | 9.04E-12 | |
| *LINC01781* | 0.411192206 | 0.111 | 0 | 2.61E-305 | 5.95E-301 | |
| *IGHA1* | 0.384504615 | 0.111 | 0.001 | 3.75E-159 | 8.57E-155 | IgA heavy chain |
| *MIR29B2CHG* | 0.37431651 | 0.185 | 0.041 | 4.87E-08 | 0.001110869 | |
| *HSH2D* | 0.366454394 | 0.241 | 0.031 | 2.65E-19 | 6.05E-15 | |
| *CD22* | 0.345489373 | 0.148 | 0.003 | 5.25E-75 | 1.20E-70 | |
| *TNFRSF13C* | 0.344446007 | 0.167 | 0.002 | 3.69E-113 | 8.43E-109 | a.k.a. BAFFR |
| *TNFRSF13B* | 0.335683275 | 0.222 | 0.006 | 8.11E-82 | 1.85E-77 | a.k.a. TACI |
| *FAM30A* | 0.313279998 | 0.148 | 0.001 | 9.17E-191 | 2.09E-186 | |
| *LINC00926* | 0.308727601 | 0.167 | 0.011 | 2.74E-26 | 6.26E-22 | |
| *ADAM28* | 0.306994007 | 0.185 | 0.019 | 3.27E-19 | 7.45E-15 | |
| *POU2AF1* | 0.268814264 | 0.204 | 0.001 | 1.23E-254 | 2.81E-250 | a.k.a. BOB.1 |
| *TMEM156* | 0.266320283 | 0.204 | 0.016 | 1.56E-27 | 3.55E-23 | |

**Supplemental Table 5. All DEGs reaching significance in the skin B cell scRNA-Seq dataset.** LogFC values represent the change in Active cGVHD skin B cells compared to healthy donor skin B cells. Skin DEGs that were also DEGs in the blood B cell scRNA-Seq dataset are highlighted yellow, with the clusters reaching significance as indicated: 'Finding in Active cGVHD Blood B cells'.

| Gene | avg_logFC | pct.1 | pct.2 | p_val | p_val_adj | Finding in Active cGVHD Blood B cells | Additional description |
|---|---|---|---|---|---|---|---|
| *PLEKHO1* | 1.731832406 | 0.833 | 0 | 1.25E-10 | 2.85E-06 | | |
| *MS4A1* | 1.970212715 | 0.75 | 0.024 | 2.29E-08 | 0.000522363 | | a.k.a. CD20 |
| *AES* | 1.886489815 | 0.75 | 0.071 | 8.76E-07 | 0.019996573 | | |
| *SH3BGRL3* | 1.804945432 | 0.833 | 0.095 | 6.16E-08 | 0.001404755 | | |
| *POU2F2* | 1.748266571 | 0.75 | 0.024 | 9.35E-09 | 0.000213346 | **Up in Clusters 3, 9, 10 and 5** | a.k.a. OCT2 |
| *COTL1* | 1.693307692 | 0.75 | 0.048 | 5.78E-08 | 0.001317887 | | |
| *CD69* | 1.606726935 | 0.75 | 0.071 | 2.07E-06 | 0.047341368 | | |
| *GMFG* | 1.594499772 | 0.75 | 0.024 | 9.35E-09 | 0.000213346 | | |
| *COMMD6* | 1.587647041 | 0.917 | 0.119 | 2.23E-08 | 0.00050919 | **Up in Cluster 8** | |
| *GGA2* | 1.573872227 | 0.5 | 0 | 1.71E-06 | 0.038925645 | | |
| *HLA-DRB5* | 1.547592835 | 0.917 | 0.048 | 7.27E-09 | 0.000165891 | | |
| *NOP14* | 1.536044844 | 0.5 | 0 | 1.71E-06 | 0.038925645 | | |
| *PRR13* | 1.502403175 | 0.667 | 0 | 1.71E-08 | 0.000389903 | | |
| *CD79B* | 1.500873877 | 0.75 | 0.119 | 1.53E-06 | 0.035026517 | | Ig-Beta |
| *HNRNPDL* | 1.479502198 | 0.833 | 0.095 | 6.16E-08 | 0.001404755 | | |
| *UIMC1* | 1.441756297 | 0.583 | 0 | 1.77E-07 | 0.004041391 | | |
| *STX12* | 1.432046578 | 0.667 | 0 | 1.71E-08 | 0.000389903 | | |
| *RPL36A* | 1.425296202 | 0.833 | 0.095 | 3.90E-08 | 0.000890461 | | |
| *BASP1* | 1.401746836 | 0.75 | 0.048 | 1.11E-07 | 0.002538172 | | |
| *ATAD1* | 1.375997106 | 0.5 | 0 | 1.71E-06 | 0.038925645 | | |
| *CRIP1* | 1.323870643 | 0.75 | 0.024 | 1.60E-08 | 0.000366112 | | |
| *ATP2B1* | 1.286889212 | 0.583 | 0.024 | 1.25E-06 | 0.028457219 | | |
| *HIST1H4C* | 1.283036004 | 0.75 | 0.071 | 2.66E-07 | 0.006063075 | **Up in Clusters 9 and 10** | |
| *GPR18* | 1.277021074 | 0.583 | 0 | 1.77E-07 | 0.004041391 | **Up in Clusters 1 and 5** | |
| *RPL23A* | 1.271659998 | 1 | 0.405 | 2.08E-06 | 0.047378048 | **Up in Cluster 8** | |
| *BTF3* | 1.220760448 | 0.917 | 0.167 | 7.05E-07 | 0.016078078 | | |
| *PNISR* | 1.215349733 | 0.833 | 0.095 | 1.30E-07 | 0.00296091 | | |
| *CD40* | 1.215322917 | 0.833 | 0.095 | 1.30E-07 | 0.00296091 | | |
| *SIPA1* | 1.185170632 | 0.667 | 0 | 1.71E-08 | 0.000389903 | | |
| *CAPZA1* | 1.182468362 | 0.667 | 0.048 | 5.15E-07 | 0.011758898 | | |
| *EIF4A1* | 1.178192543 | 0.75 | 0.071 | 3.09E-07 | 0.007057947 | | |
| *CD24* | 1.156117001 | 0.667 | 0.024 | 2.26E-07 | 0.005161888 | | |
| *BANK1* | 1.146682611 | 0.75 | 0.048 | 2.48E-07 | 0.005649835 | | B-Cell Scaffold Protein with Ankyrin Repeats |
| *UQCRB* | 1.138665328 | 0.917 | 0.143 | 6.92E-07 | 0.015788036 | | |
| *PDCD4* | 1.121603609 | 0.833 | 0.071 | 9.26E-08 | 0.002113134 | | |
| *EZR* | 1.112715616 | 0.833 | 0.119 | 1.97E-06 | 0.044950962 | | |
| *NDUFC2* | 1.105321475 | 0.75 | 0.071 | 2.28E-07 | 0.005204314 | | |
| *C4orf48* | 1.091497768 | 0.583 | 0.024 | 1.25E-06 | 0.028457219 | | |
| *IRF1* | 1.081771718 | 0.833 | 0.119 | 1.33E-06 | 0.030299525 | | |
| *TRAF3IP3* | 1.061680721 | 0.667 | 0.024 | 1.13E-07 | 0.002576638 | | |
| *U2SURP* | 1.056527054 | 0.833 | 0.095 | 1.12E-07 | 0.002554318 | | |
| *RSRC1* | 1.045776177 | 0.583 | 0.024 | 1.25E-06 | 0.028457219 | | |
| *GRK2* | 1.032408611 | 0.583 | 0.024 | 2.06E-06 | 0.047109747 | | |
| *ZFAND6* | 1.027058439 | 0.583 | 0.024 | 2.06E-06 | 0.047109747 | | |
| *MARCKSL1* | 1.006574335 | 0.667 | 0.024 | 3.77E-07 | 0.008601702 | | |
| *TNFRSF13C* | 1.003644959 | 0.667 | 0.024 | 1.60E-07 | 0.003654234 | | a.k.a BAFFR |
| *TOMM22* | 0.995554389 | 0.75 | 0.024 | 4.61E-08 | 0.001051747 | | |
| *RPL17* | 0.980823901 | 0.75 | 0.119 | 1.34E-06 | 0.030592196 | | |
| *MYCBP2* | 0.977666984 | 0.75 | 0.095 | 1.70E-06 | 0.038863573 | | |
| *CMTM6* | 0.969105333 | 0.75 | 0.071 | 1.56E-06 | 0.035632785 | | |
| *ADAM28* | 0.969034967 | 0.667 | 0.048 | 6.04E-07 | 0.0137911 | | |
| *FCRL2* | 0.960641873 | 0.583 | 0 | 1.77E-07 | 0.004041391 | **Up in Cluster 5** | |
| *LINC00926* | 0.95768142 | 0.667 | 0.024 | 1.90E-07 | 0.00434529 | | |
| *TELO2* | 0.934612879 | 0.5 | 0 | 1.71E-06 | 0.038925645 | | |
| *ARF6* | 0.901562563 | 0.667 | 0.048 | 5.15E-07 | 0.011758898 | | |
| *PCSK7* | 0.900022264 | 0.583 | 0 | 1.77E-07 | 0.004041391 | | |
| *BDP1* | 0.899996906 | 0.75 | 0.071 | 6.53E-07 | 0.014909069 | **Up in Cluster 9** | |
| *ACTR3* | 0.891973675 | 0.833 | 0.095 | 6.29E-07 | 0.014348891 | | |
| *TRAF5* | 0.882415435 | 0.583 | 0 | 1.77E-07 | 0.004041391 | | |
| *SP100* | 0.881179509 | 0.75 | 0.048 | 2.11E-07 | 0.004822465 | | |
| *FAM107B* | 0.880310967 | 0.667 | 0.048 | 7.08E-07 | 0.016159769 | | |
| *SP140* | 0.873552264 | 0.583 | 0 | 1.77E-07 | 0.004041391 | | |
| *GRK6* | 0.865769615 | 0.667 | 0.071 | 2.08E-06 | 0.047445504 | | |
| *DRAM2* | 0.851895555 | 0.667 | 0.048 | 5.15E-07 | 0.011758898 | | |
| *TMBIM4* | 0.849998071 | 0.75 | 0.071 | 3.60E-07 | 0.008209564 | | |

| | | | | | | | |
|---|---|---|---|---|---|---|---|
| *CAPG* | 0.848095752 | 0.75 | 0.071 | 5.63E-07 | 0.012858288 | | |
| *LIMD2* | 0.843313942 | 0.75 | 0.071 | 5.63E-07 | 0.012858288 | | |
| *COX8A* | 0.841398255 | 0.833 | 0.095 | 4.75E-07 | 0.010838304 | | |
| *ZNF655* | 0.84068727 | 0.667 | 0.024 | 4.46E-07 | 0.010177604 | | |
| *LAT2* | 0.832692064 | 0.583 | 0.024 | 1.25E-06 | 0.028457219 | | |
| *PHF20* | 0.795379869 | 0.75 | 0.048 | 1.80E-07 | 0.004112769 | | |
| *SNX5* | 0.792810403 | 0.5 | 0 | 1.71E-06 | 0.038925645 | | |
| *PSMA3-AS1* | 0.786434561 | 0.583 | 0 | 1.77E-07 | 0.004041391 | | |
| *RAP1B* | 0.764140085 | 0.75 | 0.071 | 1.01E-06 | 0.023130948 | | |
| *IFT57* | 0.761529181 | 0.5 | 0 | 1.71E-06 | 0.038925645 | **Up in Cluster 3** | |
| *SNX2* | 0.736198813 | 0.75 | 0.071 | 7.57E-07 | 0.017273273 | | |
| *TCP1* | 0.72581675 | 0.667 | 0.048 | 1.13E-06 | 0.025856449 | | |
| *DGKD* | 0.664724383 | 0.5 | 0 | 1.71E-06 | 0.038925645 | | |
| *TSC22D4* | 0.664253013 | 0.5 | 0 | 1.71E-06 | 0.038925645 | | |
| *IRF2* | 0.655922604 | 0.5 | 0 | 1.71E-06 | 0.038925645 | | |
| *TMC6* | 0.644492191 | 0.583 | 0 | 1.77E-07 | 0.004041391 | | |
| *PXK* | 0.64226044 | 0.5 | 0 | 1.71E-06 | 0.038925645 | | |
| *SSBP1* | 0.636605768 | 0.833 | 0.048 | 2.28E-08 | 0.000519848 | | |
| *CCDC88B* | 0.629988365 | 0.583 | 0.024 | 1.25E-06 | 0.028457219 | | |
| *HIGD2A* | 0.629139979 | 0.833 | 0.143 | 1.74E-06 | 0.039774971 | | |
| *ARHGAP25* | 0.624924796 | 0.5 | 0 | 1.71E-06 | 0.038925645 | | |
| *CMTM7* | 0.605603609 | 0.5 | 0 | 1.71E-06 | 0.038925645 | | |
| *BRD4* | 0.602992705 | 0.5 | 0 | 1.71E-06 | 0.038925645 | | |
| *ANKRD10* | 0.602691643 | 0.667 | 0 | 1.71E-08 | 0.000389903 | | |
| *SWAP70* | 0.590166501 | 0.75 | 0.024 | 4.61E-08 | 0.001051747 | **Up in Cluster 10** | Switching B-Cell Complex Subunit 70 |
| *PLEKHB2* | 0.578750717 | 0.5 | 0 | 1.71E-06 | 0.038925645 | | |
| *FCRLA* | 0.576012988 | 0.5 | 0 | 1.71E-06 | 0.038925645 | | |
| *TERF2* | 0.57600478 | 0.583 | 0 | 1.77E-07 | 0.004041391 | | |
| *ARFGAP2* | 0.572533511 | 0.583 | 0 | 1.77E-07 | 0.004041391 | | |
| *LINC00513* | 0.571454147 | 0.583 | 0 | 1.77E-07 | 0.004041391 | | |
| *IL16* | 0.563591483 | 0.5 | 0 | 1.71E-06 | 0.038925645 | **Up in Cluster 4** | Interleukin 16 |
| *REL* | 0.548459613 | 0.75 | 0.048 | 7.33E-07 | 0.016715073 | | |
| *AIM2* | 0.543726595 | 0.583 | 0.024 | 1.25E-06 | 0.028457219 | **Up in Cluster 10** | Absent In Melanoma 2 |
| *PHTF2* | 0.517004237 | 0.5 | 0 | 1.71E-06 | 0.038925645 | | |
| *SFSWAP* | 0.514465126 | 0.583 | 0 | 1.77E-07 | 0.004041391 | | |
| *MED28* | 0.508644339 | 0.583 | 0 | 1.77E-07 | 0.004041391 | | |
| *PTPN11* | 0.494422297 | 0.5 | 0 | 1.71E-06 | 0.038925645 | | |
| *RAB5IF* | 0.47342014 | 0.583 | 0.024 | 1.25E-06 | 0.028457219 | | |
| *PLEKHG1* | 0.467787339 | 0.5 | 0 | 1.71E-06 | 0.038925645 | | |
| *GABARAPL2* | 0.44397426 | 0.75 | 0.071 | 5.63E-07 | 0.012858288 | | |
| *PTBP3* | 0.427655806 | 0.5 | 0 | 1.71E-06 | 0.038925645 | | |
| *DAZAP2* | 0.413067189 | 0.75 | 0.071 | 6.53E-07 | 0.014909069 | | |
| *SELL* | 0.35536355 | 0.75 | 0.048 | 7.33E-07 | 0.016715073 | | |
| *GGA1* | 0.35047393 | 0.667 | 0.048 | 1.80E-06 | 0.041033934 | | |
| *LY86* | 0.322450642 | 0.667 | 0.024 | 4.46E-07 | 0.010177604 | | |

**Supplemental Table 7. DEGs identified in the bulk-like analysis shown in Fig. 7B, assigned to their respective biological pathways after interrogation of the GO database.** *Direction of impact on cell activation, differentiation or survival based on GO annotation (? = potential direction based on available literature). DEGs from **Fig. 7B** without an assigned classification to at least one GO pathway were omitted from the table.

| | **Up in Active cGVHD** | | **Down in Active cGVHD** | |
|---|---|---|---|---|
| **Pathway** | **Gene ID** | **Impact*** | **Gene ID** | **Impact*** |
| Cell Cycle | *CKS2*<br>*SPDL1*<br>*CDCA4*<br>*NUMA1*<br>*CETN3* | Positive<br>Positive<br>Positive<br>Positive<br>Positive? | (no genes) | |
| Cytoskeletal Reorganization | *UGT8*<br>*SKAP2*<br>*RHOQ*<br>*ABI1* | Positive<br>Positive<br>Positive<br>Positive | *WASF1* | Positive |
| G Protein-Coupled Receptor / GTPase | *GPR18*<br>*RIN3*<br>*GPR65* | Positive<br>Positive?<br>Negative? | *ARRDC3* | Negative |
| Immune Response | *FCRL4*<br>*CD48* | Positive<br>Positive | *CD5* | Negative |
| Kinase/Phosphatase Activity | *PTPRS*<br>*PPP1R16B* | Negative<br>Positive? | *PRKRA*<br>*PHKG2* | Negative<br>Positive? |
| mRNA Splicing; Translation | *PDCD7*<br>*RPL8*<br>*IVNS1ABP* | Negative?<br>Positive?<br>Positive | *MRPL28* | Positive? |
| Protein/Amino Acid Transport | *SLC15A2*<br>*SLC35D1*<br>*TRMT1L*<br>*ARCN1*<br>*AP4B1*<br>*TOMM20* | Positive<br>Positive?<br>Positive?<br>Positive?<br>Positive?<br>Positive | *CHMP6*<br>*VPS26B*<br>*ARL17A* | Positive?<br>Positive?<br>Positive? |
| Protein Stability; Proteolysis | *ADAMTS6*<br>*OTUD1*<br>*DNAJB4*<br>*CHORDC1*<br>*BAG2*<br>*UBE2T*<br>*FANCL*<br>*PHF23*<br>*TRIM4* | Negative?<br>Positive?<br>Negative?<br>Negative?<br>Positive?<br>Positive<br>Positive<br>Positive?<br>Negative? | *UBE2J2*<br>*RHBDD2* | Positive?<br>Positive? |
| Survival | *IFIT2*<br>*APAF1*<br>*DEDD2* | Negative<br>Negative<br>Negative | (no genes) | |
| Transcriptional Regulation | *ZBED2*<br>*NAB2*<br>*ZNF827*<br>*LMO4*<br>*TAF1A*<br>*ASF1A*<br>*N6AMT1*<br>*ZNF322*<br>*LRIF1*<br>*JMJD6*<br>*ZBTB4*<br>*TSC22D2*<br>*ATF2*<br>*PURA*<br>*ARID1A*<br>*KMT2C* | Positive<br>Negative<br>Positive<br>Positive<br>Positive<br>Positive<br>Positive<br>Positive<br>Positive<br>Positive<br>Negative<br>Negative?<br>Positive<br>Positive?<br>Positive<br>Positive | *POLR2M*<br>*ZNF350*<br>*HMGB2*<br>*NFKBIA*<br>*ZKSCAN2*<br>*ZNF441*<br>*HMGB3* | Negative<br>Negative<br>Positive?<br>Negative<br>Negative?<br>Negative?<br>Positive? |
