## Supplemental Table 3 for "Single-cell Landscape Analysis Unravels Molecular Programming of the Human B Cell Compartment in Chronic GVHD"

**Supplemental Table 3. All DEGs by cluster in the blood B cell scRNA-Seq dataset for untreated B cells.** Log2FC values represent the change in expression for the gene of interest in untreated B cells from patients with Active cGVHD compared to No cGVHD, along with associated statistical data.

| Cluster 1 | log2FC | lfcSE | stat | pvalue | padj |
|---|---|---|---|---|---|
| *ARRDC3* | -1.16089 | 0.124256 | -9.342741 | < 2.22e-16 | < 2.22e-16 |
| *CKS2* | 1.150264 | 0.145341 | 7.914258 | 2.49E-15 | 1.17E-11 |
| *SMIM10* | -4.719045 | 0.714476 | -6.604905 | 3.98E-11 | NA |
| *NFKBIA* | -0.633411 | 0.10013 | -6.325907 | 2.52E-10 | 7.88E-07 |
| *GATD3A* | 4.647591 | 0.850573 | 5.464067 | 4.65E-08 | 0.00010924 |
| *CXCR4* | 0.256004 | 0.048661 | 5.260934 | 1.43E-07 | 0.000269166 |
| *RSRP1* | 0.390152 | 0.074935 | 5.206521 | 1.92E-07 | 0.000281759 |
| *C7orf50* | -0.470123 | 0.090579 | -5.190221 | 2.10E-07 | 0.000281759 |
| *SLC50A1* | -0.579278 | 0.113403 | -5.108134 | 3.25E-07 | 0.000381886 |
| *C16orf74* | -0.70939 | 0.141126 | -5.026657 | 4.99E-07 | 0.000520732 |
| *ACADVL* | -0.343337 | 0.068627 | -5.002962 | 5.65E-07 | 0.000530124 |
| *SYNGR2* | -0.729816 | 0.146819 | -4.970843 | 6.67E-07 | 0.000539799 |
| *ZBED2* | 2.186817 | 0.440846 | 4.9605 | 7.03E-07 | 0.000539799 |
| *CYTIP* | 0.273594 | 0.055287 | 4.948643 | 7.47E-07 | 0.000539799 |
| *GPR183* | 0.68349 | 0.138905 | 4.920551 | 8.63E-07 | 0.000550377 |
| *C17orf49* | -0.94241 | 0.191667 | -4.916913 | 8.79E-07 | 0.000550377 |
| *CLK1* | 0.541952 | 0.111031 | 4.881087 | 1.06E-06 | 0.000612724 |
| *LMO4* | 0.689222 | 0.14149 | 4.871187 | 1.11E-06 | 0.000612724 |
| *ENSG00000272211* | -1.16566 | 0.240687 | -4.843062 | 1.28E-06 | 0.000632555 |
| *RIN3* | 0.762537 | 0.157466 | 4.842555 | 1.28E-06 | 0.000632555 |
| *CD5* | -1.485943 | 0.30748 | -4.832647 | 1.35E-06 | 0.000632555 |
| *ENSG00000274265* | -0.550603 | 0.116453 | -4.728101 | 2.27E-06 | 0.001013358 |
| *ENSG00000275964* | -1.161019 | 0.246299 | -4.713867 | 2.43E-06 | 0.00103742 |
| *CHMP6* | -0.829528 | 0.177124 | -4.683326 | 2.82E-06 | 0.001152347 |
| *CD79B* | -0.290299 | 0.062331 | -4.657387 | 3.20E-06 | 0.001252968 |
| *HMGB2* | -0.382977 | 0.0825 | -4.642141 | 3.45E-06 | 0.001255193 |
| *H3F3B* | -0.299 | 0.064433 | -4.64051 | 3.48E-06 | 0.001255193 |
| *GLRX* | 0.413958 | 0.090174 | 4.590641 | 4.42E-06 | 0.001536785 |
| *CHORDC1* | 0.587313 | 0.128611 | 4.566594 | 4.96E-06 | 0.001633973 |
| *MIF* | -0.415094 | 0.090973 | -4.562852 | 5.05E-06 | 0.001633973 |
| *HINT2* | -0.434598 | 0.095437 | -4.553756 | 5.27E-06 | 0.001649399 |
| *CD48* | 0.394216 | 0.087627 | 4.49882 | 6.83E-06 | 0.002011967 |
| *IGHD* | -0.420601 | 0.093507 | -4.498094 | 6.86E-06 | 0.002011967 |
| *CAV1* | -1.331937 | 0.300417 | -4.433623 | 9.27E-06 | 0.002636676 |
| *HSPD1* | 0.432091 | 0.098449 | 4.388966 | 1.14E-05 | 0.003087995 |
| *VMAC* | -0.831149 | 0.189472 | -4.386667 | 1.15E-05 | 0.003087995 |
| *RBM3* | -0.42132 | 0.096375 | -4.371671 | 1.23E-05 | 0.003216054 |
| *KRTCAP2* | -0.258614 | 0.059542 | -4.343415 | 1.40E-05 | 0.0035602 |
| *ZC3H12A* | -0.618369 | 0.142572 | -4.337239 | 1.44E-05 | 0.003565332 |
| *GTF3C6* | -0.396704 | 0.09224 | -4.300789 | 1.70E-05 | 0.004097685 |
| *GPR18* | 0.895258 | 0.210281 | 4.257429 | 2.07E-05 | 0.004854436 |

| | | | | | |
|---|---|---|---|---|---|
| *KRT10* | -0.337307 | 0.079644 | -4.235192 | 2.28E-05 | 0.00488225 |
| *RPL8* | 0.357522 | 0.084461 | 4.232968 | 2.31E-05 | 0.00488225 |
| *ARF5* | -0.425702 | 0.100766 | -4.224671 | 2.39E-05 | 0.00488225 |
| *SUSD3* | 0.576231 | 0.136416 | 4.224075 | 2.40E-05 | 0.00488225 |
| *UGT8* | 0.87686 | 0.207602 | 4.223757 | 2.40E-05 | 0.00488225 |
| *NUP210* | 0.444684 | 0.105367 | 4.22032 | 2.44E-05 | 0.00488225 |
| *DBI* | -0.322718 | 0.076475 | -4.219935 | 2.44E-05 | 0.00488225 |
| *LCN8* | 3.644744 | 0.870617 | 4.18639 | 2.83E-05 | 0.005544514 |
| *CYB5A* | -0.767566 | 0.184549 | -4.159138 | 3.19E-05 | 0.006121721 |
| *NINJ1* | -1.082314 | 0.26119 | -4.143781 | 3.42E-05 | 0.006415748 |
| *CLEC2D* | -0.42138 | 0.101973 | -4.132261 | 3.59E-05 | 0.006613734 |
| *SLC5A3* | -0.534456 | 0.131281 | -4.071085 | 4.68E-05 | 0.008450031 |
| *GPR65* | 0.579118 | 0.142868 | 4.053528 | 5.05E-05 | 0.008938394 |
| *INKA1* | 0.604139 | 0.149323 | 4.045864 | 5.21E-05 | 0.00906491 |
| *ZBTB25* | 0.524798 | 0.130158 | 4.032025 | 5.53E-05 | 0.009440941 |
| *ENSG00000279483* | -0.350686 | 0.087464 | -4.009477 | 6.09E-05 | 0.010203814 |
| *TNFRSF14* | 0.290898 | 0.072707 | 4.000988 | 6.31E-05 | 0.010391356 |
| *NSUN7* | -1.03129 | 0.260893 | -3.952917 | 7.72E-05 | 0.012499074 |
| *DYNLL1* | 0.411037 | 0.104805 | 3.921903 | 8.79E-05 | 0.013981949 |
| *ASF1A* | 0.571849 | 0.146569 | 3.901559 | 9.56E-05 | 0.014795111 |
| *PIM1* | -0.54679 | 0.140195 | -3.9002 | 9.61E-05 | 0.014795111 |
| *TCL1B* | -1.290914 | 0.331564 | -3.89341 | 9.88E-05 | 0.014970205 |
| *GTF2IRD2* | -0.504434 | 0.130221 | -3.873685 | 0.000107202 | 0.015978195 |
| *TMEM170A* | 0.448666 | 0.116308 | 3.857573 | 0.000114518 | 0.01647536 |
| *NUFIP2* | 0.422939 | 0.109695 | 3.855601 | 0.000115446 | 0.01647536 |
| *ZNF350* | -0.45989 | 0.119302 | -3.854849 | 0.000115801 | 0.01647536 |
| *ZFAND2A* | -0.53185 | 0.138426 | -3.842116 | 0.000121978 | 0.017095139 |
| *GRAMD1C* | -0.771036 | 0.201458 | -3.827275 | 0.00012957 | 0.01789209 |
| *NELFE* | -0.374344 | 0.098337 | -3.806738 | 0.000140812 | 0.019032576 |
| *ZNF564* | -0.797105 | 0.209496 | -3.804863 | 0.000141883 | 0.019032576 |
| *EPS15* | -0.678831 | 0.178621 | -3.800395 | 0.000144466 | 0.019106088 |
| *H2AFZ* | 0.266131 | 0.070243 | 3.788698 | 0.000151439 | 0.019750181 |
| *NOP10* | 0.199173 | 0.052859 | 3.76803 | 0.000164541 | 0.021164922 |
| *MIR29B2CHG* | 0.540707 | 0.144595 | 3.739462 | 0.000184415 | 0.023400737 |
| *APOBEC3C* | -0.29942 | 0.080151 | -3.735707 | 0.000187188 | 0.023435964 |
| *KDM4B* | 0.682874 | 0.184351 | 3.704215 | 0.000212046 | 0.026160816 |
| *YWHAE* | 0.343801 | 0.092887 | 3.701269 | 0.000214524 | 0.026160816 |
| *DEDD2* | 0.448125 | 0.122142 | 3.668885 | 0.000243611 | 0.029327001 |
| *FRG1* | -0.27148 | 0.07433 | -3.652339 | 0.000259862 | 0.030839417 |
| *EVI2A* | 0.596672 | 0.163494 | 3.649508 | 0.000262743 | 0.030839417 |
| *ENSG00000260879* | -1.428487 | 0.39218 | -3.642427 | 0.00027008 | NA |
| *TRMT112* | 0.262871 | 0.07225 | 3.638378 | 0.00027436 | 0.031805473 |
| *DNAJB4* | 0.912539 | 0.251043 | 3.634993 | 0.000277988 | 0.031832997 |
| *ATF7IP2* | -0.543517 | 0.149881 | -3.626315 | 0.000287494 | 0.032524966 |
| *TNFAIP8L2* | 0.517053 | 0.142818 | 3.620355 | 0.000294199 | 0.032887273 |
| *ENSG00000254397* | 1.73244 | 0.479891 | 3.610068 | 0.000306116 | 0.033816855 |
| *RPS9* | 0.291026 | 0.080799 | 3.601869 | 0.000315938 | 0.034496028 |

| | | | | | |
|---|---|---|---|---|---|
| *CMTM7* | -0.444415 | 0.123653 | -3.594058 | 0.000325568 | 0.035138871 |
| *SKAP2* | 0.312678 | 0.08711 | 3.589457 | 0.000331367 | 0.035358398 |
| *ODF3B* | -0.704025 | 0.196646 | -3.580167 | 0.000343375 | 0.035826235 |
| *RPS5* | 0.228828 | 0.06393 | 3.579355 | 0.000344443 | 0.035826235 |
| *TXNL4A* | -0.438732 | 0.122644 | -3.577273 | 0.000347198 | 0.035826235 |
| *ORMDL2* | -0.381928 | 0.106964 | -3.57061 | 0.000356151 | 0.036309812 |
| *MRPL28* | -0.333947 | 0.093634 | -3.566514 | 0.000361762 | 0.036309812 |
| *DDB2* | -0.486155 | 0.136359 | -3.565268 | 0.000363485 | 0.036309812 |
| *CD83* | -0.390139 | 0.109527 | -3.562039 | 0.000367985 | 0.036372415 |
| *METTL26* | -0.287434 | 0.081341 | -3.533709 | 0.000409772 | 0.040080791 |
| *THEMIS2* | 1.146557 | 0.324875 | 3.529229 | 0.000416772 | 0.040345243 |
| *GAPT* | 0.38751 | 0.110015 | 3.522343 | 0.00042775 | 0.040468656 |
| *MICOS13* | -0.256183 | 0.072784 | -3.519775 | 0.000431913 | 0.040468656 |
| *BCL7C* | -0.536615 | 0.152504 | -3.518699 | 0.000433669 | 0.040468656 |
| *PNPLA8* | 0.352473 | 0.1002 | 3.517711 | 0.000435286 | 0.040468656 |
| *INTS13* | 0.65668 | 0.188391 | 3.485727 | 0.000490802 | 0.044849206 |
| *SGK1* | -0.956973 | 0.274652 | -3.484314 | 0.0004934 | 0.044849206 |
| *ZNF688* | -0.464641 | 0.133421 | -3.482512 | 0.000496732 | 0.044849206 |
| *ARL17A* | -0.574469 | 0.165372 | -3.473793 | 0.000513156 | 0.045890795 |
| *YARS* | -0.974461 | 0.280885 | -3.469255 | 0.000521903 | 0.046232726 |
| *GBGT1* | 0.701681 | 0.202602 | 3.463339 | 0.000533517 | 0.046411383 |
| *THAP9.AS1* | -0.613626 | 0.177185 | -3.463193 | 0.000533805 | 0.046411383 |
| *LINC00638* | 1.573401 | 0.45509 | 3.457337 | 0.000545542 | NA |
| *LPAR5* | 0.748113 | 0.216636 | 3.453321 | 0.000553731 | 0.047009118 |
| *TMEM141* | -0.376051 | 0.108913 | -3.452763 | 0.000554876 | 0.047009118 |
| *PRDX3* | 0.379255 | 0.109854 | 3.452363 | 0.000555699 | 0.047009118 |
| *TTC32* | -0.632841 | 0.183627 | -3.446337 | 0.000568241 | 0.047269271 |
| *CDKN1A* | -0.581922 | 0.168963 | -3.44409 | 0.000572986 | 0.047269271 |
| *ZNF91* | 0.394216 | 0.114476 | 3.44367 | 0.000573876 | 0.047269271 |
| *CDC42* | -0.410693 | 0.119491 | -3.437033 | 0.000588125 | 0.048021683 |
| *WNT10A* | -0.970705 | 0.283539 | -3.423537 | 0.000618117 | 0.050035525 |
| *ENSG00000254614* | -0.753886 | 0.220615 | -3.417195 | 0.000632699 | 0.050736973 |
| *KRCC1* | 0.347691 | 0.101863 | 3.413326 | 0.000641751 | 0.050736973 |
| *MT1X* | -1.355045 | 0.397048 | -3.412799 | 0.000642993 | 0.050736973 |
| *ENSG00000267002* | 0.419032 | 0.122934 | 3.408591 | 0.000652993 | 0.051096713 |
| *TAGLN2* | -0.334047 | 0.098089 | -3.405563 | 0.000660278 | 0.051239728 |
| *STARD10* | -0.533181 | 0.156871 | -3.398851 | 0.000676697 | 0.051894883 |
| *ATF2* | 0.52874 | 0.155621 | 3.39761 | 0.000679773 | 0.051894883 |
| *SOAT1* | 0.662492 | 0.195159 | 3.394624 | 0.000687229 | 0.052040985 |
| *NDUFB7* | -0.273641 | 0.080731 | -3.389543 | 0.000700093 | 0.052590966 |
| *ZKSCAN1* | -0.376423 | 0.111438 | -3.377865 | 0.000730508 | 0.053362415 |
| *ZNF860* | 0.645387 | 0.191179 | 3.375817 | 0.000735968 | 0.053362415 |
| *TSC22D1* | -0.646647 | 0.191589 | -3.375176 | 0.000737684 | 0.053362415 |
| *CRK* | 0.445801 | 0.132207 | 3.371982 | 0.000746292 | 0.053362415 |
| *ENSG00000224505* | -0.389297 | 0.115471 | -3.371386 | 0.00074791 | 0.053362415 |
| *FKBP5* | 1.225567 | 0.363773 | 3.369044 | 0.000754293 | 0.053362415 |
| *CDCA4* | 0.589569 | 0.175057 | 3.367867 | 0.000757522 | 0.053362415 |

| | | | | | |
|---|---|---|---|---|---|
| *AKAP13* | 0.225168 | 0.066866 | 3.367451 | 0.000758666 | 0.053362415 |
| *UROD* | -0.377402 | 0.112108 | -3.366419 | 0.000761508 | 0.053362415 |
| *RENBP* | 0.758214 | 0.225797 | 3.357946 | 0.000785239 | 0.054617722 |
| *FANCL* | 0.743954 | 0.221754 | 3.354855 | 0.000794066 | 0.054825597 |
| *EGLN2* | 0.407101 | 0.12175 | 3.343738 | 0.000826579 | 0.056653844 |
| *ROCK1* | 0.27931 | 0.083615 | 3.340449 | 0.00083643 | 0.056913626 |
| *ENSG00000237596* | -0.929469 | 0.279221 | -3.328788 | 0.000872246 | 0.058546903 |
| *IL23A* | -1.068503 | 0.321089 | -3.327751 | 0.0008755 | 0.058546903 |
| *ENSG00000273319* | 0.580923 | 0.174659 | 3.326042 | 0.000880886 | 0.058546903 |
| *MICAL1* | -0.646335 | 0.194408 | -3.324625 | 0.000885374 | 0.058546903 |
| *NFKBIE* | -0.436676 | 0.131566 | -3.319074 | 0.000903165 | 0.059236291 |
| *BLNK* | 0.257367 | 0.077646 | 3.31463 | 0.000917646 | 0.059236291 |
| *MYBL2* | -1.852426 | 0.558952 | -3.314105 | 0.00091937 | NA |
| *RPL30* | 0.153139 | 0.046209 | 3.314045 | 0.000919569 | 0.059236291 |
| *HLA.A* | -0.463547 | 0.139892 | -3.3136 | 0.000921033 | 0.059236291 |
| *HHIP.AS1* | -2.030877 | 0.614239 | -3.306327 | 0.000945276 | NA |
| *MFSD13A* | -0.961639 | 0.291074 | -3.303764 | 0.000953961 | 0.060936709 |
| *SPINT2* | -0.280067 | 0.084863 | -3.300236 | 0.000966036 | 0.061291046 |
| *LIMD2* | -0.236235 | 0.071842 | -3.288253 | 0.001008114 | 0.063531451 |
| *FGL2* | -1.625232 | 0.494597 | -3.285975 | 0.001016301 | NA |
| *EYA3* | 0.756162 | 0.230172 | 3.285209 | 0.001019066 | 0.063793522 |
| *PILRB* | 0.721077 | 0.219838 | 3.280029 | 0.001037964 | 0.064546227 |
| *GPR137* | -0.59461 | 0.181538 | -3.275396 | 0.00105514 | 0.064871871 |
| *PYROXD1* | -0.56969 | 0.174027 | -3.273581 | 0.001061941 | 0.064871871 |
| *RAF1* | 0.37146 | 0.11349 | 3.273051 | 0.001063934 | 0.064871871 |
| *TCP1* | 0.297048 | 0.090806 | 3.271223 | 0.001070835 | 0.064871871 |
| *HMGB3* | -1.141141 | 0.348875 | -3.270915 | 0.001072001 | NA |
| *COMMD3* | -0.344209 | 0.105288 | -3.269206 | 0.001078499 | 0.064917329 |
| *ENSG00000254837* | 1.194193 | 0.365596 | 3.266425 | 0.001089147 | NA |
| *CRIPT* | 0.417244 | 0.127995 | 3.259845 | 0.001114731 | 0.066670854 |
| *ZC3H12D* | -0.647001 | 0.198631 | -3.257304 | 0.00112476 | 0.066844912 |
| *CASP4* | 0.243324 | 0.074754 | 3.255009 | 0.001133887 | 0.066963505 |
| *NAB2* | 1.055125 | 0.325418 | 3.242363 | 0.00118543 | 0.069500561 |
| *ZFYVE26* | 0.924577 | 0.285381 | 3.239798 | 0.001196144 | 0.069500561 |
| *CHP1* | -0.339326 | 0.104814 | -3.237403 | 0.00120623 | 0.069500561 |
| *TSN* | -0.340065 | 0.105104 | -3.235495 | 0.001214322 | 0.069500561 |
| *ENSG00000278834* | -1.04798 | 0.32401 | -3.234401 | 0.001218984 | NA |
| *RHOBTB2* | 0.965666 | 0.298631 | 3.23364 | 0.001222235 | 0.069500561 |
| *SERPINI1* | 1.024205 | 0.316875 | 3.232202 | 0.001228401 | 0.069500561 |
| *ADAMTS6* | 0.972409 | 0.300856 | 3.232143 | 0.001228657 | 0.069500561 |
| *RHOH* | 0.246444 | 0.076412 | 3.225203 | 0.001258833 | 0.070525326 |
| *CCT8* | 0.456192 | 0.141476 | 3.22453 | 0.001261795 | 0.070525326 |
| *MTPN* | 0.256935 | 0.079766 | 3.221124 | 0.001276889 | 0.070946669 |
| *ENSG00000166927* | -1.03188 | 0.322399 | -3.200628 | 0.001371285 | 0.075743323 |
| *ENSG00000268858* | -0.878811 | 0.275171 | -3.193692 | 0.00140466 | 0.076852121 |
| *NRL* | -1.502583 | 0.470491 | -3.193651 | 0.001404856 | NA |
| *PTGES2* | -0.47533 | 0.148863 | -3.193062 | 0.001407728 | 0.076852121 |

| | | | | | |
|---|---|---|---|---|---|
| *ENSG00000239636* | -0.804073 | 0.252183 | -3.188455 | 0.001430353 | 0.077600443 |
| *DNAJB1* | 0.524837 | 0.164685 | 3.18692 | 0.001437963 | 0.077600443 |
| *DAP* | -0.650767 | 0.204534 | -3.181714 | 0.001464065 | 0.07855755 |
| *C6orf203* | 0.546628 | 0.172073 | 3.176724 | 0.001489487 | 0.079016803 |
| *CUTC* | -0.397087 | 0.125 | -3.1767 | 0.001489612 | 0.079016803 |
| *SCRN1* | 0.926439 | 0.291783 | 3.175096 | 0.001497869 | 0.079016803 |
| *EIF4A2* | 0.178043 | 0.056194 | 3.168354 | 0.00153305 | 0.080420886 |
| *RFTN1* | -0.468597 | 0.148154 | -3.16291 | 0.001562008 | 0.081484733 |
| *IZUMO4* | -0.566745 | 0.179396 | -3.159177 | 0.001582154 | 0.082079726 |
| *CFAP410* | -0.374775 | 0.11879 | -3.15495 | 0.001605256 | 0.082820627 |
| *PTPMT1* | -0.340336 | 0.107957 | -3.152527 | 0.001618637 | 0.083054652 |
| *POGZ* | 0.363958 | 0.115919 | 3.13976 | 0.001690861 | 0.085895694 |
| *ROGDI* | -0.612072 | 0.195131 | -3.136717 | 0.001708507 | 0.085895694 |
| *SYNJ2BP* | 0.339509 | 0.108268 | 3.135821 | 0.001713736 | 0.085895694 |
| *ENSG00000237491* | -0.670294 | 0.213783 | -3.135387 | 0.001716273 | 0.085895694 |
| *MAML3* | -1.034544 | 0.33002 | -3.134795 | 0.001719743 | 0.085895694 |
| *RBBP8* | -0.770184 | 0.246901 | -3.119408 | 0.001812148 | 0.089963196 |
| *ATP6V1C2* | -2.14975 | 0.689292 | -3.118777 | 0.001816033 | NA |
| *SMCR8* | 0.610329 | 0.195739 | 3.118079 | 0.001820342 | 0.089963196 |
| *JADE1* | 0.35726 | 0.114949 | 3.107975 | 0.001883742 | 0.091571792 |
| *JMJD6* | 0.654045 | 0.210452 | 3.107816 | 0.001884753 | 0.091571792 |
| *TNFSF10* | 0.582584 | 0.187465 | 3.107699 | 0.001885501 | 0.091571792 |
| *PPCS* | -0.241844 | 0.077846 | -3.106698 | 0.001891899 | 0.091571792 |
| *CTNNBIP1* | -1.011588 | 0.325967 | -3.103348 | 0.001913445 | 0.092049805 |
| *BOLA2B* | -0.611408 | 0.197093 | -3.102123 | 0.00192138 | 0.092049805 |
| *FGFBP3* | -1.060911 | 0.342838 | -3.094493 | 0.001971496 | NA |
| *PXMP2* | -0.680085 | 0.219864 | -3.093201 | 0.001980102 | 0.094381518 |
| *TCOF1* | -0.406225 | 0.13149 | -3.089393 | 0.002005657 | 0.094649807 |
| *BBX* | 0.242398 | 0.078462 | 3.089359 | 0.00200589 | 0.094649807 |
| *MCRIP1* | -0.223996 | 0.072687 | -3.081656 | 0.002058528 | 0.096647869 |
| *ZNF253* | 0.521255 | 0.169349 | 3.077994 | 0.002083992 | 0.097356644 |
| *CYB561D2* | -0.37341 | 0.121481 | -3.073813 | 0.002113416 | 0.098242473 |
| *ZNF442* | -1.370983 | 0.446494 | -3.070549 | 0.002136657 | NA |
| *TMEM179B* | -0.317544 | 0.103449 | -3.069581 | 0.002143595 | 0.099078874 |
| *SRSF5* | -0.173124 | 0.056423 | -3.06834 | 0.002152512 | 0.099078874 |
| *UTRN* | 0.396235 | 0.129255 | 3.065539 | 0.002172783 | 0.099524074 |
| *PCIF1* | -0.494578 | 0.161724 | -3.058156 | 0.002227038 | 0.101514031 |
| *FGFR1OP2* | 0.337413 | 0.110454 | 3.054776 | 0.002252284 | 0.102065614 |
| *CD180* | 0.60242 | 0.19728 | 3.053634 | 0.002260878 | 0.102065614 |
| *METTL7A* | 0.358239 | 0.117513 | 3.048501 | 0.002299864 | 0.10273556 |
| *FMNL2* | -1.869366 | 0.613231 | -3.048387 | 0.002300733 | NA |
| *ENSG00000282988* | 0.803505 | 0.263897 | 3.044768 | 0.002328598 | 0.10273556 |
| *PFN1* | -0.189443 | 0.062225 | -3.044463 | 0.002330959 | 0.10273556 |
| *YIPF6* | -0.43366 | 0.142462 | -3.044046 | 0.002334191 | 0.10273556 |
| *IGHG1* | 1.111677 | 0.365289 | 3.043277 | 0.002340167 | 0.10273556 |
| *POLR2E* | -0.248482 | 0.081654 | -3.043123 | 0.002341364 | 0.10273556 |
| *SLC3A2* | -0.408761 | 0.134551 | -3.037966 | 0.002381811 | 0.104024218 |

| | | | | | |
|---|---|---|---|---|---|
| *RMND5A* | 0.536121 | 0.177 | 3.028926 | 0.002454244 | 0.106659345 |
| *SETD1B* | 0.690387 | 0.228029 | 3.027622 | 0.002464865 | 0.106659345 |
| *ZBTB20* | 0.31137 | 0.10309 | 3.020378 | 0.002524593 | 0.108287638 |
| *ADO* | -0.734679 | 0.24325 | -3.020262 | 0.002525558 | 0.108287638 |
| *ICA1L* | -0.474282 | 0.15751 | -3.011118 | 0.002602879 | 0.110822514 |
| *RAB11A* | -0.199029 | 0.066112 | -3.010488 | 0.002608283 | 0.110822514 |
| *POLH* | 0.524935 | 0.175031 | 2.999092 | 0.002707857 | 0.113973575 |
| *CHTOP* | 0.247792 | 0.082648 | 2.998179 | 0.002715984 | 0.113973575 |
| *AKAP2* | -0.704735 | 0.23508 | -2.997856 | 0.002718858 | 0.113973575 |
| *DUSP12* | -0.3299 | 0.110253 | -2.992208 | 0.002769675 | 0.115587751 |
| *LRIF1* | 0.346047 | 0.115728 | 2.990174 | 0.002788188 | 0.11584551 |
| *ZC3H6* | 0.425917 | 0.142588 | 2.98705 | 0.002816835 | 0.116520191 |
| *PTMS* | -1.661519 | 0.556945 | -2.983272 | 0.002851847 | NA |
| *ADA* | -0.572731 | 0.192011 | -2.982805 | 0.002856203 | 0.117630466 |
| *MRPL32* | 0.39143 | 0.131713 | 2.97184 | 0.002960206 | 0.120475892 |
| *TBC1D10A* | 0.473991 | 0.159511 | 2.971531 | 0.002963187 | 0.120475892 |
| *COL19A1* | -0.483238 | 0.162783 | -2.968605 | 0.002991553 | 0.120475892 |
| *MAP7D1* | -0.42946 | 0.144672 | -2.968521 | 0.002992367 | 0.120475892 |
| *ASPSCR1* | -0.462098 | 0.15567 | -2.968452 | 0.002993043 | 0.120475892 |
| *FAM41C* | -0.810986 | 0.273307 | -2.967308 | 0.0030042 | 0.120475892 |
| *TMEM208* | -0.324331 | 0.109361 | -2.96569 | 0.003020051 | 0.120475892 |
| *CLECL1* | 1.516478 | 0.512001 | 2.961869 | 0.003057781 | 0.120475892 |
| *SYNC* | -0.502881 | 0.169819 | -2.961275 | 0.003063685 | 0.120475892 |
| *FXYD2* | -1.035424 | 0.349836 | -2.959736 | 0.003079027 | 0.120475892 |
| *LRRC61* | -0.667859 | 0.2257 | -2.959056 | 0.003085833 | 0.120475892 |
| *BRK1* | -0.195204 | 0.065973 | -2.958862 | 0.003087776 | 0.120475892 |
| *MTHFR* | 0.581089 | 0.196481 | 2.957478 | 0.003101665 | 0.120475892 |
| *PDE6D* | -0.418857 | 0.141642 | -2.957155 | 0.003104916 | 0.120475892 |
| *RPS6KB2* | -0.271591 | 0.092002 | -2.952023 | 0.003156998 | 0.121768294 |
| *DUSP22* | -0.355981 | 0.120618 | -2.951323 | 0.00316416 | 0.121768294 |
| *TSEN15* | -0.507926 | 0.172267 | -2.948473 | 0.003193478 | 0.121987831 |
| *NDUFS8* | -0.231341 | 0.078467 | -2.948244 | 0.003195847 | 0.121987831 |
| *TUBB4B* | 0.373356 | 0.126958 | 2.940785 | 0.003273816 | 0.124050513 |
| *FCRL3* | -1.024275 | 0.348328 | -2.940549 | 0.003276307 | 0.124050513 |
| *ETS2* | -1.290828 | 0.438984 | -2.940493 | 0.003276906 | NA |
| *IL21R* | -0.623476 | 0.212223 | -2.937831 | 0.00330517 | 0.124640764 |
| *C1orf43* | 0.242806 | 0.082825 | 2.931542 | 0.003372838 | 0.12543106 |
| *LRIG1* | -0.754339 | 0.257342 | -2.931266 | 0.003375835 | 0.12543106 |
| *FNTA* | 0.250265 | 0.085381 | 2.931163 | 0.003376958 | 0.12543106 |
| *ALDH9A1* | 0.388372 | 0.132508 | 2.930924 | 0.003379559 | 0.12543106 |
| *DNAAF4* | -1.619068 | 0.553301 | -2.926198 | 0.003431325 | 0.126386548 |
| *FXYD7* | 0.703499 | 0.240465 | 2.925578 | 0.003438169 | 0.126386548 |
| *PRICKLE1* | 0.595171 | 0.203535 | 2.924164 | 0.003453827 | 0.126386548 |
| *ENSG00000225963* | 0.92584 | 0.316896 | 2.921584 | 0.003482563 | 0.126386548 |
| *STAM2* | 0.679355 | 0.232566 | 2.921126 | 0.003487689 | 0.126386548 |
| *RPL24* | 0.173892 | 0.059538 | 2.920676 | 0.003492725 | 0.126386548 |
| *GDF11* | 0.450888 | 0.15441 | 2.920071 | 0.003499521 | 0.126386548 |

| | | | | | |
|---|---|---|---|---|---|
| *VPREB1* | -1.814691 | 0.622183 | -2.916651 | 0.003538109 | 0.127290575 |
| *FBXO4* | -0.485592 | 0.166634 | -2.914124 | 0.003566886 | 0.127836097 |
| *IER3* | -1.127319 | 0.387392 | -2.910022 | 0.003614029 | NA |
| *FAM177B* | 0.623079 | 0.21412 | 2.90995 | 0.003614866 | 0.12804914 |
| *XIAP* | 0.427322 | 0.146857 | 2.909788 | 0.003616737 | 0.12804914 |
| *DFFB* | 0.895908 | 0.307937 | 2.90939 | 0.003621348 | NA |
| *ORMDL3* | -0.449711 | 0.154599 | -2.908879 | 0.003627274 | 0.12804914 |
| *TMEM243* | 0.236022 | 0.081139 | 2.90887 | 0.003627377 | 0.12804914 |
| *SIGLEC5* | 1.555229 | 0.53558 | 2.903822 | 0.003686377 | NA |
| *CHMP5* | 0.267365 | 0.092107 | 2.902757 | 0.003698938 | 0.130086256 |
| *DCAF13* | 0.368919 | 0.12725 | 2.899167 | 0.003741551 | 0.131093911 |
| *TAPBP* | -0.299784 | 0.103476 | -2.897133 | 0.003765895 | 0.131456344 |
| *RNF43* | -0.801607 | 0.276826 | -2.895709 | 0.00378303 | 0.131565361 |
| *LY9* | 0.458417 | 0.158422 | 2.893642 | 0.003808016 | 0.13194565 |
| *H1FX* | -0.888601 | 0.307293 | -2.891704 | 0.003831588 | 0.132274298 |
| *POLE3* | -0.306222 | 0.105942 | -2.890461 | 0.003846771 | 0.13231201 |
| *RPS28* | 0.183031 | 0.06341 | 2.886489 | 0.003895659 | 0.133504528 |
| *PIGX* | -0.552344 | 0.191528 | -2.883888 | 0.00392798 | 0.134122646 |
| *TPM4* | 0.314647 | 0.109189 | 2.881684 | 0.003955564 | 0.134575166 |
| *POR* | -0.464993 | 0.161732 | -2.875081 | 0.004039241 | 0.136568436 |
| *GDI1* | 0.311317 | 0.108293 | 2.874768 | 0.00404324 | 0.136568436 |
| *PTGIR* | -1.387579 | 0.48322 | -2.871528 | 0.004084922 | 0.137481772 |
| *LINC01991* | -1.107371 | 0.385675 | -2.871256 | 0.004088437 | NA |
| *EIF5* | 0.184059 | 0.064222 | 2.865981 | 0.004157195 | 0.139269814 |
| *LYRM2* | -0.277301 | 0.096783 | -2.865181 | 0.004167712 | 0.139269814 |
| *RPL19* | 0.118167 | 0.041297 | 2.861415 | 0.00421754 | 0.140435103 |
| *AIM2* | 1.751814 | 0.613558 | 2.855174 | 0.004301326 | NA |
| *IFIT2* | 0.533366 | 0.186818 | 2.854997 | 0.00430373 | 0.142798664 |
| *TDG* | 0.323483 | 0.113422 | 2.852035 | 0.004344033 | 0.143529531 |
| *RBM43* | -0.632853 | 0.221965 | -2.851136 | 0.004356328 | 0.143529531 |
| *ENSG00000272277* | -1.05399 | 0.369818 | -2.850019 | 0.004371657 | NA |
| *PIGF* | 0.53759 | 0.188804 | 2.847344 | 0.004408572 | 0.144572504 |
| *LINC00662* | 0.298325 | 0.1048 | 2.846608 | 0.004418776 | 0.144572504 |
| *TTC21A* | 0.337012 | 0.118777 | 2.837341 | 0.004549103 | 0.148003859 |
| *HLA.C* | -0.485777 | 0.171234 | -2.836915 | 0.004555177 | 0.148003859 |
| *ENSG00000280433* | -0.894275 | 0.315624 | -2.833357 | 0.004606196 | 0.149145451 |
| *ENSG00000273145* | -1.583334 | 0.559347 | -2.830684 | 0.004644856 | NA |
| *SMIM2.AS1* | -2.109612 | 0.745598 | -2.829422 | 0.004663222 | NA |
| *PDCL3* | -0.50087 | 0.177037 | -2.829185 | 0.004666677 | 0.150584541 |
| *GALNT2* | -0.415248 | 0.147154 | -2.82185 | 0.004774757 | 0.153278685 |
| *LAMC1* | 1.018833 | 0.361121 | 2.821309 | 0.004782817 | 0.153278685 |
| *CREBRF* | 0.287077 | 0.101849 | 2.818651 | 0.004822591 | 0.154027667 |
| *BIN1* | 0.317453 | 0.112798 | 2.814354 | 0.004887532 | 0.155344062 |
| *ST13* | 0.134099 | 0.047659 | 2.813739 | 0.004896895 | 0.155344062 |
| *IL16* | 0.243047 | 0.08642 | 2.812392 | 0.004917455 | 0.155471063 |
| *TRMT13* | 0.341244 | 0.121396 | 2.810989 | 0.004938948 | 0.155626591 |
| *SLC16A7* | 0.420343 | 0.149638 | 2.809067 | 0.004968526 | 0.156034965 |

| | | | | | |
|---|---|---|---|---|---|
| *SAFB* | 0.312617 | 0.111345 | 2.807648 | 0.004990476 | 0.156201886 |
| *TAGAP* | -0.396166 | 0.141493 | -2.799898 | 0.005111882 | 0.159470348 |
| *YWHAG* | 0.39984 | 0.143094 | 2.794255 | 0.005201947 | 0.160496171 |
| *PPM1K* | 0.197311 | 0.07062 | 2.794004 | 0.00520599 | 0.160496171 |
| *EFCAB11* | -0.694267 | 0.248488 | -2.793971 | 0.005206511 | 0.160496171 |
| *CFAP73* | -1.323146 | 0.473663 | -2.793435 | 0.005215155 | 0.160496171 |
| *SCN3A* | 1.542525 | 0.552389 | 2.792461 | 0.005230875 | NA |
| *PPP1R15A* | -0.581857 | 0.208493 | -2.790768 | 0.005258307 | 0.160496171 |
| *NDUFS7* | -0.208876 | 0.074848 | -2.790673 | 0.005259853 | 0.160496171 |
| *LINC02422* | 0.65955 | 0.236448 | 2.789406 | 0.005280481 | 0.160496171 |
| *OCLN* | -0.817582 | 0.293171 | -2.788753 | 0.005291143 | 0.160496171 |
| *SNRPA1* | -0.301484 | 0.108125 | -2.788297 | 0.005298596 | 0.160496171 |
| *CHL1* | 1.339659 | 0.480531 | 2.787873 | 0.005305528 | NA |
| *SNAP23* | -0.230134 | 0.082667 | -2.78386 | 0.005371628 | 0.162185163 |
| *NDUFB11* | 0.280297 | 0.100732 | 2.782586 | 0.005392753 | 0.162301123 |
| *SLAMF1* | 0.468927 | 0.168596 | 2.781372 | 0.005412967 | 0.162389003 |
| *COL9A3* | -1.00434 | 0.361582 | -2.777629 | 0.005475705 | 0.163246405 |
| *TMEM99* | 0.609994 | 0.219613 | 2.777593 | 0.005476317 | 0.163246405 |
| *TMEM205* | -0.348325 | 0.125555 | -2.774279 | 0.005532424 | 0.164397032 |
| *DHPS* | 0.232991 | 0.084027 | 2.772821 | 0.005557261 | 0.164489914 |
| *ENSG00000259943* | -0.619443 | 0.223461 | -2.772042 | 0.005570585 | 0.164489914 |
| *KLHDC7B* | -0.921422 | 0.333223 | -2.765184 | 0.005689065 | 0.166407883 |
| *SREBF2.AS1* | -0.692699 | 0.250569 | -2.764501 | 0.005700996 | 0.166407883 |
| *CEP104* | 0.389294 | 0.140822 | 2.764452 | 0.005701853 | 0.166407883 |
| *SMIM4* | -0.41057 | 0.148559 | -2.763693 | 0.005715121 | 0.166407883 |
| *PODXL2* | -1.312078 | 0.474882 | -2.762957 | 0.005728028 | NA |
| *FAAP24* | -0.749711 | 0.271374 | -2.762652 | 0.005733388 | 0.166407883 |
| *TAX1BP3* | -0.360475 | 0.130504 | -2.762169 | 0.005741869 | 0.166407883 |
| *FCGR2B* | 0.804798 | 0.291929 | 2.756825 | 0.005836552 | 0.16821394 |
| *RETREG1* | -0.659536 | 0.239382 | -2.755157 | 0.005866391 | 0.16821394 |
| *POLR2B* | -0.274625 | 0.099727 | -2.753771 | 0.005891296 | 0.16821394 |
| *PPP1R35* | -0.549201 | 0.199467 | -2.753344 | 0.005898985 | 0.16821394 |
| *DMAC2L* | -0.260385 | 0.094571 | -2.753333 | 0.005899194 | 0.16821394 |
| *ENSG00000272606* | -0.505394 | 0.183603 | -2.752641 | 0.005911672 | 0.16821394 |
| *MRPL50* | 0.294232 | 0.106988 | 2.750146 | 0.005956875 | 0.168988095 |
| *MRPL23* | -0.281482 | 0.102476 | -2.746807 | 0.006017852 | 0.170203696 |
| *ZBTB16* | 1.855098 | 0.676434 | 2.742466 | 0.006097971 | 0.171553146 |
| *ADAL* | -0.516844 | 0.188475 | -2.742244 | 0.006102103 | 0.171553146 |
| *PPP1R16B* | 0.371481 | 0.135543 | 2.740693 | 0.00613098 | 0.171850446 |
| *ENSG00000262580* | 1.188904 | 0.433851 | 2.740349 | 0.006137398 | NA |
| *MPND* | -1.084342 | 0.395719 | -2.740179 | 0.006140568 | NA |
| *C1GALT1* | 0.368354 | 0.134645 | 2.735747 | 0.006223882 | 0.173935261 |
| *SELENOW* | -0.372056 | 0.136133 | -2.733036 | 0.006275346 | 0.174117038 |
| *SRSF10* | 0.190367 | 0.069664 | 2.732647 | 0.006282761 | 0.174117038 |
| *DUSP14* | 0.64163 | 0.234835 | 2.732255 | 0.00629025 | 0.174117038 |
| *HMGXB4* | -0.40231 | 0.147301 | -2.731219 | 0.006310061 | 0.174117038 |
| *VOPP1* | -0.328164 | 0.120183 | -2.730538 | 0.0063231 | 0.174117038 |

| | | | | | |
|---|---|---|---|---|---|
| *CIRBP* | -0.175971 | 0.064559 | -2.725738 | 0.006415795 | 0.176152962 |
| *RWDD2A* | -0.873893 | 0.321161 | -2.721039 | 0.0065077 | NA |
| *ZNF721* | -0.322834 | 0.118717 | -2.719352 | 0.006541003 | 0.178631109 |
| *HERC2* | -0.319741 | 0.117587 | -2.719195 | 0.0065441 | 0.178631109 |
| *C9orf72* | -0.449174 | 0.165301 | -2.717309 | 0.006581522 | 0.179131849 |
| *TNFRSF17* | 1.265323 | 0.465922 | 2.715739 | 0.006612811 | 0.179291827 |
| *TMEM140* | 0.413952 | 0.15247 | 2.714981 | 0.00662796 | 0.179291827 |
| *ZNF557* | 0.474963 | 0.174995 | 2.714146 | 0.006644681 | 0.179291827 |
| *AGAP9* | 1.524093 | 0.561719 | 2.713267 | 0.006662339 | NA |
| *ATP6V0E2* | -0.673254 | 0.248239 | -2.712121 | 0.006685427 | 0.179874381 |
| *ZNF628* | -1.364092 | 0.503248 | -2.710576 | 0.00671664 | NA |
| *RPL9* | 0.15455 | 0.057057 | 2.708717 | 0.006754389 | 0.181210599 |
| *OPTN* | -0.736532 | 0.272036 | -2.707474 | 0.006779731 | 0.181372292 |
| *XXYLT1.AS2* | 1.133009 | 0.418782 | 2.705485 | 0.006820481 | 0.18194407 |
| *RAB11FIP2* | -0.457953 | 0.169364 | -2.703959 | 0.006851875 | 0.182263754 |
| *TCTN1* | -0.798005 | 0.296151 | -2.694588 | 0.007047575 | 0.186919045 |
| *ENSG00000262292* | 0.711962 | 0.264535 | 2.691373 | 0.007115869 | 0.186919045 |
| *RPS23* | 0.139149 | 0.051706 | 2.691125 | 0.007121148 | 0.186919045 |
| *PFKP* | -0.763444 | 0.283779 | -2.690277 | 0.007139276 | 0.186919045 |
| *NT5E* | 0.668207 | 0.248467 | 2.689315 | 0.00715989 | 0.186919045 |
| *AGMAT* | 0.729028 | 0.271131 | 2.688842 | 0.007170028 | 0.186919045 |
| *PARP9* | 0.561489 | 0.208945 | 2.68725 | 0.007204308 | 0.186919045 |
| *GLS* | 0.407385 | 0.151604 | 2.687164 | 0.007206162 | 0.186919045 |
| *PHKG1* | -0.524366 | 0.195146 | -2.687038 | 0.007208882 | 0.186919045 |
| *RPL35A* | 0.15235 | 0.056743 | 2.68492 | 0.007254726 | 0.186919045 |
| *PFDN2* | 0.222657 | 0.082933 | 2.684795 | 0.007257424 | 0.186919045 |
| *EMP3* | -0.344669 | 0.128397 | -2.684412 | 0.007265756 | 0.186919045 |
| *MED23* | -0.482284 | 0.17982 | -2.682037 | 0.007317539 | 0.187736858 |
| *AAK1* | -0.913817 | 0.341002 | -2.679797 | 0.007366692 | 0.187826387 |
| *TMEM60* | 0.330271 | 0.123274 | 2.679162 | 0.007380657 | 0.187826387 |
| *ZNF830* | 0.281757 | 0.105196 | 2.678408 | 0.007397305 | 0.187826387 |
| *LILRB1* | 0.414567 | 0.154827 | 2.67761 | 0.007414945 | 0.187826387 |
| *XRRA1* | -0.65964 | 0.246379 | -2.677335 | 0.007421043 | 0.187826387 |
| *CENPS* | -0.494336 | 0.184813 | -2.67479 | 0.007477606 | 0.188455976 |
| *SYNE2* | -0.37654 | 0.140794 | -2.674411 | 0.007486057 | 0.188455976 |
| *CLCF1* | 0.531298 | 0.19874 | 2.673329 | 0.007510254 | 0.188559576 |
| *IFT57* | 0.448623 | 0.168013 | 2.670165 | 0.007581398 | 0.189576885 |
| *TNFAIP8* | 0.300739 | 0.112654 | 2.669578 | 0.00759466 | 0.189576885 |
| *ZNF816* | 0.618176 | 0.231627 | 2.668841 | 0.00761134 | 0.189576885 |
| *LONP2* | -0.260358 | 0.097594 | -2.66777 | 0.00763566 | 0.189679492 |
| *ZC3H7A* | -0.467025 | 0.175312 | -2.663961 | 0.007722649 | 0.191334235 |
| *PDPK1* | 0.372638 | 0.139957 | 2.662519 | 0.007755812 | 0.191380149 |
| *TMEM14B* | -0.30359 | 0.114071 | -2.661402 | 0.00778159 | 0.191380149 |
| *BAG2* | 0.672967 | 0.252885 | 2.661154 | 0.007787341 | 0.191380149 |
| *COG3* | -0.481723 | 0.18108 | -2.66027 | 0.007807795 | 0.191380149 |
| *PTP4A1* | -0.382614 | 0.143869 | -2.659468 | 0.007826409 | 0.191380149 |
| *GSTO1* | 0.295132 | 0.111172 | 2.654724 | 0.007937331 | 0.193588409 |

| | | | | | |
|---|---|---|---|---|---|
| *IL18BP* | -1.472422 | 0.555331 | -2.651432 | 0.008015117 | NA |
| *NAA50* | -0.258821 | 0.097628 | -2.651091 | 0.008023231 | 0.195141217 |
| *ZNRD2* | -0.281016 | 0.106033 | -2.650278 | 0.008042561 | 0.195141217 |
| *CCDC191* | -0.817398 | 0.308659 | -2.648222 | 0.008091639 | 0.195826 |
| *SUGCT* | -2.789715 | 1.055027 | -2.644213 | 0.008188119 | NA |
| *RDH5* | 0.497628 | 0.188221 | 2.643855 | 0.008196778 | 0.197860524 |
| *ZNF322* | 0.474622 | 0.179611 | 2.642502 | 0.008229588 | 0.198107503 |
| *BCL6* | -0.491069 | 0.185892 | -2.641696 | 0.008249205 | 0.198107503 |
| *PSMB10* | -0.343732 | 0.130202 | -2.640003 | 0.008290534 | 0.198586326 |
| *TPI1* | -0.171074 | 0.064855 | -2.637797 | 0.00834466 | 0.198586326 |
| *ZNF568* | 0.468811 | 0.177746 | 2.637524 | 0.008351367 | 0.198586326 |
| *PSMA4* | -0.190331 | 0.072165 | -2.637428 | 0.008353738 | 0.198586326 |
| *ENSG00000216895* | -0.528582 | 0.200521 | -2.636038 | 0.008388044 | 0.198898306 |
| *ENSG00000271746* | -1.420261 | 0.539478 | -2.63266 | 0.008471916 | NA |
| *ZNF595* | 0.829007 | 0.314926 | 2.632385 | 0.008478775 | 0.200543308 |
| *GID8* | -0.272751 | 0.103674 | -2.630866 | 0.00851677 | 0.20093586 |
| *CLTA* | 0.188596 | 0.071713 | 2.629872 | 0.008541706 | 0.201019106 |
| *HAUS3* | 0.517933 | 0.197188 | 2.62659 | 0.00862453 | 0.201088846 |
| *TMX4* | -0.324487 | 0.123569 | -2.625954 | 0.00864065 | 0.201088846 |
| *TSPAN33* | -0.421031 | 0.16035 | -2.6257 | 0.008647111 | 0.201088846 |
| *OTUD1* | 0.59483 | 0.226624 | 2.624737 | 0.008671582 | 0.201088846 |
| *RHBDD2* | -0.354183 | 0.135 | -2.623582 | 0.008701038 | 0.201088846 |
| *SNHG9* | -0.333345 | 0.127091 | -2.622893 | 0.008718673 | 0.201088846 |
| *NUAK2* | -0.290662 | 0.110839 | -2.62237 | 0.008732061 | 0.201088846 |
| *LINC01215* | -0.326311 | 0.124455 | -2.621925 | 0.008743464 | 0.201088846 |
| *GPX1* | -0.24121 | 0.092023 | -2.621196 | 0.008762199 | 0.201088846 |
| *MAGEH1* | -0.497388 | 0.189823 | -2.620271 | 0.008785995 | 0.201088846 |
| *SMAP2* | -0.298928 | 0.114122 | -2.619371 | 0.008809209 | 0.201088846 |
| *ELOB* | -0.167336 | 0.063906 | -2.618498 | 0.00883178 | 0.201088846 |
| *ZNF213.AS1* | -0.62556 | 0.238914 | -2.618351 | 0.00883558 | 0.201088846 |
| *LETMD1* | 0.270743 | 0.103416 | 2.618008 | 0.008844483 | 0.201088846 |
| *VSIR* | -0.774977 | 0.296121 | -2.617095 | 0.008868159 | 0.201140117 |
| *DDX6* | -0.348896 | 0.133408 | -2.615256 | 0.00891605 | 0.201628635 |
| *ZNF267* | -0.436942 | 0.167138 | -2.614258 | 0.008942155 | 0.201628635 |
| *SNHG32* | 0.193928 | 0.07421 | 2.613222 | 0.008969295 | 0.201628635 |
| *HSPH1* | 0.395268 | 0.151308 | 2.612338 | 0.008992535 | 0.201628635 |
| *TPMT* | -0.580867 | 0.222426 | -2.611503 | 0.009014509 | 0.201628635 |
| *TTC9C* | 0.301035 | 0.115325 | 2.610325 | 0.009045619 | 0.201628635 |
| *LDHA* | -0.32796 | 0.12566 | -2.609904 | 0.009056773 | 0.201628635 |
| *CYC1* | -0.239816 | 0.091893 | -2.609726 | 0.009061479 | 0.201628635 |
| *SORD* | -0.546811 | 0.209621 | -2.608569 | 0.009092158 | 0.201833014 |
| *MVP* | -0.420726 | 0.16137 | -2.607211 | 0.00912832 | 0.202099302 |
| *LIPT1* | 0.496987 | 0.190679 | 2.606411 | 0.009149669 | 0.202099302 |
| *HAUS8* | 0.490332 | 0.188177 | 2.605698 | 0.009168722 | 0.202099302 |
| *SF1* | -0.201569 | 0.077442 | -2.60285 | 0.009245241 | 0.203050879 |
| *ELF1* | 0.244474 | 0.094003 | 2.600702 | 0.009303324 | 0.203050879 |
| *BET1* | 0.345708 | 0.132979 | 2.599709 | 0.009330286 | 0.203050879 |

| | | | | | |
|---|---|---|---|---|---|
| *RPL7L1* | 0.194288 | 0.074746 | 2.599316 | 0.009340981 | 0.203050879 |
| *C2orf42* | 0.531729 | 0.204565 | 2.599311 | 0.009341113 | 0.203050879 |
| *NME2* | -0.885503 | 0.340671 | -2.599292 | 0.009341638 | 0.203050879 |
| *RFX1* | 0.330303 | 0.127124 | 2.598269 | 0.009369507 | 0.203186316 |
| *ENSG00000257354* | -0.714341 | 0.275084 | -2.596809 | 0.009409434 | 0.203581997 |
| *FRY.AS1* | -1.437212 | 0.553805 | -2.595157 | 0.009454761 | NA |
| *MFNG* | 0.350123 | 0.134983 | 2.593837 | 0.009491134 | 0.204877577 |
| *RPL38* | 0.150979 | 0.058283 | 2.590456 | 0.009584899 | 0.206427069 |
| *EIPR1* | -0.391308 | 0.151129 | -2.58924 | 0.009618804 | 0.206683218 |
| *PTPRS* | 1.54415 | 0.597224 | 2.585544 | 0.009722537 | 0.208017312 |
| *ENSG00000267458* | -1.419959 | 0.549212 | -2.58545 | 0.009725197 | 0.208017312 |
| *GLUL* | -0.563213 | 0.21791 | -2.584611 | 0.00974889 | 0.208050169 |
| *CCT5* | 0.335435 | 0.129866 | 2.582934 | 0.009796415 | 0.208079446 |
| *CTH* | 1.411147 | 0.546348 | 2.582871 | 0.009798204 | NA |
| *RCC1* | 0.409393 | 0.158526 | 2.5825 | 0.009808743 | 0.208079446 |
| *ASB16.AS1* | 0.396384 | 0.153572 | 2.581099 | 0.009848626 | 0.208079446 |
| *PHF21A* | -0.316506 | 0.122646 | -2.580647 | 0.009861543 | 0.208079446 |
| *HSPBAP1* | -0.783509 | 0.303669 | -2.580147 | 0.009875837 | 0.208079446 |
| *TIMM10* | -0.365184 | 0.141569 | -2.579545 | 0.00989306 | 0.208079446 |
| *RNF34* | 0.327712 | 0.127083 | 2.578734 | 0.009916308 | 0.208079446 |
| *NDUFB10* | -0.196159 | 0.076091 | -2.57796 | 0.00993856 | 0.208079446 |
| *SLC37A3* | 0.653456 | 0.253589 | 2.576835 | 0.009970958 | 0.208079446 |
| *MAVS* | 0.340464 | 0.132127 | 2.576804 | 0.009971858 | 0.208079446 |
| *AP1S1* | -0.355763 | 0.138254 | -2.57326 | 0.010074557 | 0.209756306 |
| *POLR2J* | -0.179786 | 0.069895 | -2.572241 | 0.010104249 | 0.209909066 |
| *PRMT7* | 0.5307 | 0.20649 | 2.570096 | 0.010167026 | 0.210746954 |
| *BLOC1S1* | -0.17318 | 0.067506 | -2.565394 | 0.010305858 | 0.213154189 |
| *PRDM8* | -0.770938 | 0.300827 | -2.562726 | 0.010385394 | 0.213924413 |
| *ERP29* | 0.152301 | 0.059442 | 2.562178 | 0.010401814 | 0.213924413 |
| *CUX1* | -0.349117 | 0.136333 | -2.560769 | 0.010444071 | 0.213924413 |
| *MRPS15* | -0.242209 | 0.094593 | -2.560547 | 0.010450762 | 0.213924413 |
| *TAF10* | -0.550206 | 0.214896 | -2.560339 | 0.010457008 | 0.213924413 |
| *SAFB2* | 0.244349 | 0.095469 | 2.559464 | 0.010483382 | 0.213997733 |
| *TLR7* | 0.78002 | 0.304912 | 2.558184 | 0.010522047 | 0.214056319 |
| *MED6* | -0.314018 | 0.122766 | -2.55786 | 0.010531844 | 0.214056319 |
| *ZNF714* | 0.599237 | 0.234595 | 2.554341 | 0.010638897 | 0.215765096 |
| *GLMN* | -0.393196 | 0.15413 | -2.551077 | 0.010739049 | 0.217215091 |
| *MPHOSPH8* | 0.156331 | 0.061294 | 2.550506 | 0.010756658 | 0.217215091 |
| *TRIM69* | 0.402412 | 0.158003 | 2.546864 | 0.010869577 | 0.219024307 |
| *PHF23* | 0.331322 | 0.130311 | 2.542546 | 0.011004814 | 0.2212158 |
| *MAP3K8* | -0.544835 | 0.214346 | -2.54185 | 0.011026753 | 0.2212158 |
| *TOMM20* | 0.175173 | 0.068945 | 2.540749 | 0.011061522 | 0.2212158 |
| *WASHC1* | -0.461397 | 0.181641 | -2.540154 | 0.011080366 | 0.2212158 |
| *MT.CO1* | 0.224598 | 0.088436 | 2.539657 | 0.011096128 | 0.2212158 |
| *HLA.G* | -1.937256 | 0.763262 | -2.538126 | 0.011144792 | NA |
| *GIMAP2* | 0.489658 | 0.192924 | 2.538095 | 0.011145774 | 0.221734785 |
| *KCTD7* | 0.496738 | 0.195813 | 2.536796 | 0.011187224 | 0.222088856 |

| | | | | | |
|---|---|---|---|---|---|
| *PRKRA* | -0.255192 | 0.100626 | -2.536053 | 0.011210987 | 0.222091066 |
| *KCTD3* | -1.225784 | 0.483854 | -2.533377 | 0.011296931 | NA |
| *BZW1* | -0.226015 | 0.089232 | -2.532887 | 0.011312743 | 0.223220306 |
| *UQCRH* | 0.158358 | 0.062527 | 2.532627 | 0.011321147 | 0.223220306 |
| *SPATA2L* | 0.559054 | 0.22079 | 2.532065 | 0.011339306 | 0.223220306 |
| *MED14OS* | -0.787357 | 0.311081 | -2.531035 | 0.011372639 | 0.22340812 |
| *KPNB1* | 0.215964 | 0.085393 | 2.529051 | 0.011437143 | 0.223543046 |
| *CCDC138* | -0.693338 | 0.274161 | -2.528941 | 0.01144073 | 0.223543046 |
| *FBXO28* | 0.596044 | 0.235718 | 2.528628 | 0.011450927 | 0.223543046 |
| *FAM184B* | -1.590337 | 0.629145 | -2.527777 | 0.011478734 | 0.223620985 |
| *TNFRSF13B* | 0.931101 | 0.368493 | 2.526783 | 0.011511276 | 0.223790638 |
| *HHEX* | 0.242992 | 0.096251 | 2.524558 | 0.011584386 | 0.224267304 |
| *RIOX1* | -0.72167 | 0.28592 | -2.524025 | 0.011601969 | 0.224267304 |
| *CYSTM1* | -0.321824 | 0.127514 | -2.523836 | 0.011608223 | 0.224267304 |
| *PGGHG* | -0.460717 | 0.182597 | -2.523136 | 0.011631329 | 0.224267304 |
| *KMT2E.AS1* | -0.513062 | 0.203521 | -2.520932 | 0.011704435 | 0.224737411 |
| *ORAI3* | -0.235793 | 0.093536 | -2.52087 | 0.011706493 | 0.224737411 |
| *CBLN3* | -0.568582 | 0.225607 | -2.520239 | 0.011727511 | 0.224737411 |
| *PPP1CB* | -0.309529 | 0.122886 | -2.518828 | 0.011774607 | 0.225180374 |
| *IGLC7* | -2.060806 | 0.818588 | -2.517514 | 0.011818625 | 0.225515616 |
| *HSD17B11* | 0.139016 | 0.055243 | 2.516469 | 0.011853739 | 0.225515616 |
| *SMPD4* | -0.416541 | 0.165546 | -2.516158 | 0.011864187 | 0.225515616 |
| *ENSG00000187186* | -0.979523 | 0.389313 | -2.516029 | 0.011868538 | NA |
| *NPRL2* | 0.391221 | 0.155665 | 2.513229 | 0.011963158 | 0.226707989 |
| *MYC* | 0.270865 | 0.107791 | 2.512874 | 0.011975204 | 0.226707989 |
| *ENSG00000257839* | -1.291571 | 0.514145 | -2.512078 | 0.012002254 | NA |
| *RALGPS2* | 0.315108 | 0.12555 | 2.509831 | 0.012078886 | 0.228062454 |
| *CD40* | -0.153249 | 0.061071 | -2.509351 | 0.012095325 | 0.228062454 |
| *MYCBP2* | 0.168394 | 0.067193 | 2.506136 | 0.012205867 | 0.228841308 |
| *HSBP1* | 0.204431 | 0.081586 | 2.505697 | 0.012221016 | 0.228841308 |
| *PPP4C* | -0.183206 | 0.073133 | -2.505094 | 0.012241883 | 0.228841308 |
| *RAB34* | -0.643444 | 0.256882 | -2.504823 | 0.012251252 | 0.228841308 |
| *DLGAP3* | -0.866635 | 0.346073 | -2.504201 | 0.012272832 | NA |
| *SIDT1.AS1* | -0.592577 | 0.236644 | -2.504086 | 0.012276804 | 0.228841308 |
| *MARCHF1* | 0.404454 | 0.161529 | 2.503912 | 0.012282856 | 0.228841308 |
| *FIS1* | -0.267853 | 0.107041 | -2.502326 | 0.012338013 | 0.229413747 |
| *DPM1* | 0.281586 | 0.112595 | 2.500874 | 0.01238871 | 0.22951289 |
| *UBE2L3* | -0.179493 | 0.071775 | -2.500774 | 0.01239223 | 0.22951289 |
| *LINC00680* | -0.461139 | 0.184489 | -2.499547 | 0.012435206 | 0.229855474 |
| *H6PD* | -0.464914 | 0.186083 | -2.498424 | 0.012474671 | 0.230131953 |
| *MRPL41* | -0.257272 | 0.103073 | -2.496009 | 0.012559959 | 0.231251002 |
| *ENSG00000260349* | -0.578667 | 0.231967 | -2.494607 | 0.012609671 | 0.231711961 |
| *PKNOX1* | -0.433922 | 0.174084 | -2.492603 | 0.012681072 | 0.232568881 |
| *HRK* | -1.252097 | 0.502537 | -2.491554 | 0.012718556 | 0.232801648 |
| *NCBP2AS2* | -0.277258 | 0.111317 | -2.490718 | 0.012748513 | 0.232895993 |
| *RPL13A* | 0.119521 | 0.048043 | 2.487772 | 0.012854623 | 0.234239177 |
| *RASAL1* | -1.320389 | 0.530772 | -2.487674 | 0.012858141 | NA |

| | | | | | |
|---|---|---|---|---|---|
| *SHARPIN* | -0.263292 | 0.105855 | -2.487293 | 0.012871929 | 0.234239177 |
| *ZNF835* | 0.769677 | 0.309675 | 2.485436 | 0.012939282 | NA |
| *CARMIL1* | -0.940716 | 0.378945 | -2.482462 | 0.013047814 | 0.236980598 |
| *OSTF1* | 0.34996 | 0.141028 | 2.481484 | 0.013083645 | 0.237172634 |
| *LIMD1* | -0.816246 | 0.329592 | -2.476536 | 0.013266408 | 0.238923469 |
| *SLC9A9* | -0.502482 | 0.202928 | -2.476162 | 0.013280321 | 0.238923469 |
| *KIAA0040* | 0.46311 | 0.187187 | 2.474044 | 0.013359319 | 0.238923469 |
| *BHLHE41* | 1.756752 | 0.710126 | 2.473859 | 0.013366261 | NA |
| *MGAT1* | 0.25301 | 0.102287 | 2.473532 | 0.013378494 | 0.238923469 |
| *SHF* | -0.751675 | 0.304032 | -2.472354 | 0.013422649 | 0.238923469 |
| *ELMSAN1* | 0.318229 | 0.128723 | 2.472205 | 0.013428254 | 0.238923469 |
| *LINC02201* | -1.237143 | 0.500463 | -2.471998 | 0.013436017 | 0.238923469 |
| *MAN1B1.DT* | -0.621304 | 0.2514 | -2.471375 | 0.013459468 | 0.238923469 |
| *LPIN2* | 0.488001 | 0.197466 | 2.47132 | 0.013461538 | 0.238923469 |
| *FAM117A* | 0.302043 | 0.122233 | 2.471037 | 0.013472196 | 0.238923469 |
| *EEA1* | 0.404775 | 0.16383 | 2.470696 | 0.01348503 | 0.238923469 |
| *DDX5* | -0.289184 | 0.117046 | -2.470682 | 0.013485563 | 0.238923469 |
| *SLC2A14* | -3.695871 | 1.496675 | -2.469388 | 0.013534429 | NA |
| *NEK1* | 0.432185 | 0.175079 | 2.468513 | 0.013567576 | 0.239923794 |
| *ACAP1* | 0.284352 | 0.115228 | 2.467734 | 0.013597114 | 0.239994172 |
| *JRK* | 0.470737 | 0.190826 | 2.466834 | 0.013631357 | 0.240147163 |
| *ZNF605* | -0.402652 | 0.163312 | -2.465544 | 0.013680538 | 0.240562274 |
| *CCDC126* | 0.436936 | 0.177301 | 2.464369 | 0.013725485 | 0.240901501 |
| *DENND2D* | 0.390674 | 0.158639 | 2.462659 | 0.013791116 | 0.241194735 |
| *CD58* | -0.569048 | 0.231119 | -2.462141 | 0.013811051 | 0.241194735 |
| *JOSD2* | -0.417872 | 0.169734 | -2.461928 | 0.013819251 | 0.241194735 |
| *FAM204A* | -0.225313 | 0.091657 | -2.458233 | 0.013962265 | 0.24270801 |
| *GCNT1* | 0.795572 | 0.323643 | 2.458178 | 0.013964376 | 0.24270801 |
| *ENSG00000268400* | -1.209155 | 0.492 | -2.457634 | 0.01398557 | NA |
| *PLEKHO2* | 0.883051 | 0.359375 | 2.457187 | 0.014002992 | 0.24270801 |
| *ABTB1* | -0.269904 | 0.10985 | -2.457024 | 0.014009344 | 0.24270801 |
| *SLC38A11* | 1.622383 | 0.66075 | 2.455368 | 0.014074058 | NA |
| *CELF2* | 0.286704 | 0.116788 | 2.454916 | 0.014091759 | 0.243686212 |
| *PDE4B* | 0.226047 | 0.092192 | 2.451905 | 0.014210199 | 0.2448464 |
| *LAMTOR1* | -0.191297 | 0.078028 | -2.451626 | 0.014221242 | 0.2448464 |
| *PSPC1* | -0.308529 | 0.125867 | -2.451225 | 0.014237075 | 0.2448464 |
| *RBMX* | 0.129714 | 0.052976 | 2.448528 | 0.014344123 | 0.246236405 |
| *ZNF580* | -0.264121 | 0.107899 | -2.447864 | 0.014370596 | 0.246240689 |
| *MX1* | 0.702208 | 0.287038 | 2.446394 | 0.014429304 | 0.246516039 |
| *MINDY2* | 0.33607 | 0.137387 | 2.446148 | 0.014439172 | 0.246516039 |
| *ANKRD28* | -0.593943 | 0.242932 | -2.444893 | 0.014489526 | 0.24692676 |
| *UFD1* | -0.324147 | 0.132716 | -2.442405 | 0.014589774 | 0.248184738 |
| *MPP7* | -0.768844 | 0.314984 | -2.440898 | 0.014650791 | 0.248772015 |
| *CNOT6L* | -0.609132 | 0.249759 | -2.43888 | 0.014732849 | 0.24971382 |
| *JUND* | -0.320256 | 0.131418 | -2.436929 | 0.014812601 | 0.250222592 |
| *FAM131A* | 0.553611 | 0.227184 | 2.436842 | 0.014816162 | 0.250222592 |
| *NDUFB8* | -0.167939 | 0.069094 | -2.430584 | 0.01507453 | 0.254128977 |

| | | | | | |
|---|---|---|---|---|---|
| *OR2A1.AS1* | -0.915744 | 0.377083 | -2.428491 | 0.015161789 | NA |
| *MRPL54* | -0.272029 | 0.112049 | -2.427778 | 0.015191646 | 0.255644366 |
| *SNAP29* | 0.296079 | 0.121994 | 2.426996 | 0.015224406 | 0.255737343 |
| *PCF11* | 0.199098 | 0.082129 | 2.424206 | 0.015341894 | 0.257250694 |
| *COX16* | -0.348454 | 0.14378 | -2.423525 | 0.015370716 | 0.257274546 |
| *SLC35D1* | 0.579094 | 0.23918 | 2.421162 | 0.015470999 | 0.25849232 |
| *FBXO2* | -0.868107 | 0.358694 | -2.42019 | 0.015512397 | 0.258723631 |
| *ENSG00000276509* | 0.549238 | 0.227041 | 2.419116 | 0.015558267 | 0.258754598 |
| *ENSG00000271204* | 0.446237 | 0.184483 | 2.418857 | 0.015569366 | 0.258754598 |
| *PFDN1* | -0.268706 | 0.111151 | -2.417483 | 0.015628257 | 0.259005292 |
| *OSTM1* | 0.342269 | 0.141596 | 2.417219 | 0.015639617 | 0.259005292 |
| *USP53* | 0.951023 | 0.393614 | 2.416128 | 0.015686564 | 0.25917512 |
| *CAPN12* | 0.520111 | 0.215339 | 2.415315 | 0.01572159 | 0.25917512 |
| *IGHG3* | 1.031544 | 0.427422 | 2.413408 | 0.015804118 | 0.25917512 |
| *PPM1D* | 0.515373 | 0.21356 | 2.413248 | 0.015811045 | 0.25917512 |
| *CYP4V2* | 0.647396 | 0.268272 | 2.413205 | 0.015812925 | 0.25917512 |
| *NECAP2* | -0.224156 | 0.092912 | -2.412575 | 0.015840279 | 0.25917512 |
| *SLC25A26* | -0.281432 | 0.116682 | -2.41195 | 0.015867457 | 0.25917512 |
| *ENSG00000228463* | 0.475214 | 0.197031 | 2.411876 | 0.015870681 | 0.25917512 |
| *PCNX4* | 0.366652 | 0.15213 | 2.41013 | 0.015946849 | 0.259966853 |
| *ZSCAN18* | 0.234655 | 0.0974 | 2.409186 | 0.015988136 | 0.259997185 |
| *C9orf16* | -0.177 | 0.07348 | -2.408822 | 0.016004087 | 0.259997185 |
| *TBC1D8* | -1.141503 | 0.474211 | -2.407162 | 0.016077021 | NA |
| *ISCA2* | -0.231081 | 0.095998 | -2.407139 | 0.016078055 | 0.260010225 |
| *UBE2E1* | 0.330835 | 0.137468 | 2.406638 | 0.0161001 | 0.260010225 |
| *ENSG00000247373* | 0.828262 | 0.344283 | 2.405762 | 0.016138775 | 0.260010225 |
| *SCP2* | -0.237532 | 0.098768 | -2.404946 | 0.016174852 | 0.260010225 |
| *RAB5C* | -0.186187 | 0.077423 | -2.404805 | 0.016181119 | 0.260010225 |
| *ANP32B* | 0.159874 | 0.066488 | 2.404559 | 0.01619201 | 0.260010225 |
| *SENP3* | -0.527752 | 0.219494 | -2.404407 | 0.01619872 | 0.260010225 |
| *APOBEC3G* | -0.221621 | 0.092247 | -2.402466 | 0.016284945 | 0.26051831 |
| *NOTCH2* | 0.316716 | 0.131834 | 2.402393 | 0.016288191 | 0.26051831 |
| *SPOP* | 0.230537 | 0.095984 | 2.401823 | 0.016313607 | 0.26051831 |
| *FIBP* | -0.206006 | 0.085922 | -2.397583 | 0.016503629 | 0.262789448 |
| *IRF3* | 0.190959 | 0.079652 | 2.397402 | 0.016511797 | 0.262789448 |
| *DSCAML1* | -1.257975 | 0.5253 | -2.394773 | 0.016630648 | NA |
| *SMIM14* | -0.261254 | 0.109127 | -2.394033 | 0.016664259 | 0.264611317 |
| *ZBTB8OS* | -0.258481 | 0.107987 | -2.393629 | 0.01668263 | 0.264611317 |
| *POLR2M* | -0.27314 | 0.114172 | -2.392363 | 0.016740277 | 0.265077915 |
| *WASH9P* | -0.436919 | 0.182695 | -2.391521 | 0.016778721 | 0.265239374 |
| *PSMD8* | -0.1616 | 0.067636 | -2.389283 | 0.016881292 | 0.26566487 |
| *TRA2B* | -0.169214 | 0.070852 | -2.388264 | 0.016928187 | 0.26566487 |
| *SLA* | 0.395691 | 0.165689 | 2.388146 | 0.016933632 | 0.26566487 |
| *DRG2* | -0.374524 | 0.156849 | -2.387801 | 0.016949526 | 0.26566487 |
| *MME* | -1.99894 | 0.837224 | -2.387582 | 0.016959634 | NA |
| *STIP1* | 0.337602 | 0.14141 | 2.387394 | 0.016968297 | 0.26566487 |
| *SLC2A1* | -0.317901 | 0.133172 | -2.387137 | 0.016980174 | 0.26566487 |

| | | | | | |
|---|---|---|---|---|---|
| *COMMD2* | -0.249164 | 0.1044 | -2.386628 | 0.017003683 | 0.26566487 |
| *ERV3.1* | -0.308685 | 0.129504 | -2.383605 | 0.017143971 | 0.266991678 |
| *ENSG00000125726* | -1.513501 | 0.634972 | -2.383573 | 0.017145472 | 0.266991678 |
| *TMEM165* | -0.33557 | 0.140836 | -2.382706 | 0.017185928 | 0.267178577 |
| *RPRD1A* | -0.253143 | 0.106299 | -2.381423 | 0.017245898 | 0.267667743 |
| *UNC119* | -0.320824 | 0.134824 | -2.379565 | 0.017333103 | 0.267670508 |
| *DEAF1* | -0.559263 | 0.235061 | -2.379227 | 0.017349005 | 0.267670508 |
| *CPT1A* | -0.508516 | 0.213751 | -2.379011 | 0.017359154 | 0.267670508 |
| *FH* | 0.399364 | 0.167871 | 2.378991 | 0.0173601 | 0.267670508 |
| *RNF144B* | 0.740134 | 0.311525 | 2.375841 | 0.017508987 | 0.269523588 |
| *VDAC2* | 0.191583 | 0.080751 | 2.372531 | 0.01766669 | 0.271506087 |
| *LINC02035* | 0.853765 | 0.359949 | 2.371907 | 0.017696552 | NA |
| *GPS2* | -0.252839 | 0.106614 | -2.37153 | 0.017714626 | 0.271579817 |
| *SENP2* | -0.476655 | 0.201042 | -2.370915 | 0.01774409 | 0.271579817 |
| *ZNF776* | 0.483814 | 0.204095 | 2.370535 | 0.017762351 | 0.271579817 |
| *PET100* | -0.237262 | 0.10011 | -2.370019 | 0.017787177 | 0.271579817 |
| *ADGRB2* | -1.314415 | 0.555073 | -2.368005 | 0.017884304 | NA |
| *PLXNB2* | -1.140005 | 0.481665 | -2.366798 | 0.017942718 | NA |
| *CEP170* | -0.321295 | 0.135904 | -2.364123 | 0.018072807 | 0.275444519 |
| *ABHD3* | 0.379083 | 0.160436 | 2.362833 | 0.018135847 | 0.275444519 |
| *RPLP2* | 0.183504 | 0.077674 | 2.362488 | 0.018152716 | 0.275444519 |
| *WDR33* | 0.255383 | 0.108104 | 2.362388 | 0.018157631 | 0.275444519 |
| *ZKSCAN2* | -0.714877 | 0.302723 | -2.361487 | 0.018201827 | NA |
| *S100PBP* | -0.346852 | 0.146939 | -2.360516 | 0.018249543 | 0.276392277 |
| *ENSG00000226571* | 1.153364 | 0.488752 | 2.359815 | 0.018284059 | NA |
| *BCAP31* | -0.217763 | 0.092295 | -2.359418 | 0.018303611 | 0.276764745 |
| *COA6.AS1* | -0.507295 | 0.215083 | -2.358601 | 0.018343975 | 0.27692914 |
| *CALM3* | -0.275938 | 0.117195 | -2.354518 | 0.018546757 | 0.279541012 |
| *DDX55* | 0.398966 | 0.169857 | 2.348835 | 0.01883226 | 0.283389304 |
| *HEBP2* | -0.517344 | 0.220341 | -2.347926 | 0.018878282 | 0.283627308 |
| *MPLKIP* | 0.214016 | 0.091265 | 2.344994 | 0.0190274 | 0.285088417 |
| *SOX4* | -0.63265 | 0.269834 | -2.344591 | 0.019047978 | 0.285088417 |
| *ATP5ME* | -0.156031 | 0.06656 | -2.344226 | 0.019066616 | 0.285088417 |
| *RPL32* | 0.126919 | 0.054157 | 2.343551 | 0.019101136 | 0.285150498 |
| *MFAP1* | 0.382812 | 0.163438 | 2.342249 | 0.019167902 | 0.285693013 |
| *APOL6* | 0.745414 | 0.318619 | 2.339516 | 0.019308758 | 0.286759656 |
| *BUD13* | 0.394329 | 0.168588 | 2.339007 | 0.019335048 | 0.286759656 |
| *PHYH* | -0.725462 | 0.310159 | -2.339002 | 0.019335316 | 0.286759656 |
| *ARCN1* | 0.421139 | 0.18009 | 2.338494 | 0.019361621 | 0.286759656 |
| *ZFYVE27* | 0.325612 | 0.139387 | 2.336023 | 0.019490049 | 0.288207178 |
| *HSD17B4* | 0.418195 | 0.179128 | 2.334611 | 0.019563744 | 0.288394435 |
| *ZNF680* | 0.304522 | 0.130445 | 2.33448 | 0.019570626 | 0.288394435 |
| *CCL5* | -1.362285 | 0.583795 | -2.333497 | 0.019622064 | 0.288394435 |
| *ZDHHC23* | -0.450347 | 0.192998 | -2.33343 | 0.019625564 | 0.288394435 |
| *ATP6V1C1* | 0.347022 | 0.14888 | 2.330885 | 0.019759418 | 0.289266442 |
| *ENSG00000224746* | -1.046286 | 0.449189 | -2.329279 | 0.019844304 | NA |
| *HSH2D* | -0.294635 | 0.126541 | -2.328374 | 0.019892251 | 0.289266442 |

| | | | | | |
|---|---|---|---|---|---|
| *NPM1* | 0.127144 | 0.054626 | 2.327564 | 0.019935279 | 0.289266442 |
| *ENSG00000267390* | 0.472164 | 0.202874 | 2.327372 | 0.01994548 | 0.289266442 |
| *IDNK* | -0.486209 | 0.208932 | -2.327119 | 0.019958928 | 0.289266442 |
| *YY1* | 0.177093 | 0.076113 | 2.326715 | 0.019980414 | 0.289266442 |
| *RALA* | -0.290484 | 0.12486 | -2.326468 | 0.019993593 | 0.289266442 |
| *ACYP2* | -0.300867 | 0.129329 | -2.326366 | 0.019999008 | 0.289266442 |
| *DUS2* | 0.340592 | 0.146414 | 2.326236 | 0.020005974 | 0.289266442 |
| *C12orf49* | -0.531167 | 0.228368 | -2.325921 | 0.020022764 | 0.289266442 |
| *XYLT2* | -0.707896 | 0.304353 | -2.325902 | 0.020023769 | 0.289266442 |
| *KIAA0930* | 0.523746 | 0.225397 | 2.323659 | 0.020143784 | 0.290553203 |
| *PPP4R1* | -0.524484 | 0.226113 | -2.319569 | 0.020364217 | 0.293187165 |
| *CFL2* | -0.380287 | 0.16398 | -2.319114 | 0.020388841 | 0.293187165 |
| *ARL11* | 0.561761 | 0.242338 | 2.318087 | 0.020444599 | 0.29353943 |
| *PEX13* | 0.590358 | 0.254844 | 2.316544 | 0.020528594 | 0.294295422 |
| *FXYD5* | 0.161398 | 0.069749 | 2.314007 | 0.020667361 | 0.295833113 |
| *ENSG00000274400* | 1.581173 | 0.683324 | 2.313942 | 0.0206709 | NA |
| *RER1* | -0.224169 | 0.096954 | -2.312121 | 0.020770994 | 0.296711637 |
| *SMARCD1* | -0.273623 | 0.118362 | -2.311741 | 0.020791934 | 0.296711637 |
| *GSTP1* | -0.19264 | 0.083357 | -2.311011 | 0.020832223 | 0.296835464 |
| *GUK1* | -0.200605 | 0.086842 | -2.310007 | 0.020887758 | 0.297175825 |
| *DUS4L* | -0.495216 | 0.214529 | -2.308381 | 0.020977949 | 0.297900264 |
| *CCNQ* | -0.316439 | 0.137187 | -2.306625 | 0.021075746 | 0.297900264 |
| *FHL3* | -0.814912 | 0.353332 | -2.30636 | 0.021090507 | 0.297900264 |
| *CLNS1A* | 0.329814 | 0.143022 | 2.306033 | 0.021108768 | 0.297900264 |
| *NUP160* | 0.256503 | 0.111236 | 2.305925 | 0.021114847 | 0.297900264 |
| *LENG1* | 0.277682 | 0.120441 | 2.305556 | 0.021135451 | 0.297900264 |
| *A4GALT* | 1.441525 | 0.625285 | 2.305387 | 0.021144911 | NA |
| *RTCA* | 0.374968 | 0.162696 | 2.304719 | 0.021182295 | 0.297900264 |
| *HGSNAT* | -0.411296 | 0.178484 | -2.304387 | 0.021200956 | 0.297900264 |
| *POU2AF1* | -0.19466 | 0.084489 | -2.303972 | 0.021224204 | 0.297900264 |
| *ENSG00000275441* | -0.663706 | 0.288347 | -2.301759 | 0.021348747 | NA |
| *ZNF526* | 0.388417 | 0.16878 | 2.301315 | 0.021373832 | 0.299552659 |
| *PCMTD2* | -0.265623 | 0.115619 | -2.297403 | 0.021595809 | 0.302212589 |
| *ENSG00000170846* | -0.832788 | 0.362727 | -2.29591 | 0.021681039 | NA |
| *CEP152* | 0.420129 | 0.183025 | 2.295471 | 0.02170615 | 0.302890942 |
| *LAMTOR4* | -0.156069 | 0.067992 | -2.295425 | 0.021708797 | 0.302890942 |
| *FAM111A.DT* | 0.323071 | 0.140842 | 2.293852 | 0.02179899 | 0.303698096 |
| *ZNF579* | -0.932055 | 0.406461 | -2.2931 | 0.021842273 | NA |
| *ING3* | -0.330455 | 0.144174 | -2.292064 | 0.021901975 | 0.304191004 |
| *FOXN2* | 0.395889 | 0.172728 | 2.291985 | 0.021906482 | 0.304191004 |
| *LDHB* | 0.173779 | 0.075852 | 2.291035 | 0.021961417 | 0.304191004 |
| *PTK2B* | 0.23838 | 0.104051 | 2.290991 | 0.021963951 | 0.304191004 |
| *POLA1* | -0.790528 | 0.34525 | -2.289727 | 0.022037176 | NA |
| *PSPH* | -0.749264 | 0.327712 | -2.286353 | 0.022233632 | NA |
| *PFKFB3* | -0.314353 | 0.137499 | -2.286218 | 0.022241515 | 0.306542512 |
| *PCYT2* | -0.470029 | 0.205632 | -2.285773 | 0.022267538 | 0.306542512 |
| *MSL3* | 0.25067 | 0.109699 | 2.285062 | 0.022309167 | 0.306542512 |

| | | | | | |
|---|---|---|---|---|---|
| *ENSG00000272941* | 0.956558 | 0.418614 | 2.285058 | 0.022309431 | NA |
| *OPN3* | -0.489965 | 0.214443 | -2.28482 | 0.022323372 | 0.306542512 |
| *HCK* | -0.463672 | 0.202992 | -2.284191 | 0.02236033 | 0.306542512 |
| *MAP4* | -0.234469 | 0.102653 | -2.284089 | 0.022366326 | 0.306542512 |
| *LMF1* | -0.607699 | 0.266065 | -2.284028 | 0.022369889 | 0.306542512 |
| *STK16* | -0.242341 | 0.10615 | -2.283009 | 0.02242985 | 0.306542512 |
| *ABCG1* | -0.628384 | 0.275523 | -2.280694 | 0.022566551 | 0.306542512 |
| *SLC15A2* | 0.868573 | 0.380848 | 2.28063 | 0.022570371 | 0.306542512 |
| *WDR11* | -0.330921 | 0.145144 | -2.279952 | 0.022610557 | 0.306542512 |
| *LDLRAP1* | 0.481656 | 0.211263 | 2.279888 | 0.022614316 | 0.306542512 |
| *CYBC1* | -0.169237 | 0.074238 | -2.279638 | 0.022629158 | 0.306542512 |
| *RPSA* | 0.191754 | 0.084117 | 2.279626 | 0.022629903 | 0.306542512 |
| *ENSG00000251661* | -1.525007 | 0.668974 | -2.279621 | 0.022630155 | NA |
| *TMEM242* | -0.290094 | 0.12728 | -2.279175 | 0.02265664 | 0.306542512 |
| *SNHG25* | -0.332722 | 0.146011 | -2.278744 | 0.02268227 | 0.306542512 |
| *TBCB* | -0.20982 | 0.092089 | -2.278433 | 0.022700804 | 0.306542512 |
| *BARD1* | 0.370916 | 0.162819 | 2.278088 | 0.022721362 | 0.306542512 |
| *LZTS3* | -1.388416 | 0.609656 | -2.277377 | 0.022763702 | NA |
| *PDCD4* | 0.43204 | 0.189776 | 2.276583 | 0.022811143 | 0.307093248 |
| *RPL3* | 0.116615 | 0.05123 | 2.276308 | 0.022827592 | 0.307093248 |
| *MIR210HG* | -1.389388 | 0.610397 | -2.276204 | 0.022833796 | NA |
| *STK38L* | -0.496584 | 0.218409 | -2.273646 | 0.02298728 | 0.308799089 |
| *AAAS* | -0.417062 | 0.183618 | -2.271355 | 0.023125473 | 0.310054802 |
| *PHF7* | -0.471557 | 0.207643 | -2.271003 | 0.023146796 | 0.310054802 |
| *WDFY2* | 0.288527 | 0.127196 | 2.268369 | 0.023306753 | 0.311665409 |
| *HNRNPD* | -0.185858 | 0.08195 | -2.267931 | 0.023333417 | 0.311665409 |
| *ZNF827* | 1.028256 | 0.453654 | 2.26661 | 0.023414041 | NA |
| *PPWD1* | 0.304475 | 0.134392 | 2.265576 | 0.023477372 | 0.31314279 |
| *KIAA1324L* | 1.020768 | 0.450816 | 2.264268 | 0.02355761 | NA |
| *SUMO1* | -0.163519 | 0.07224 | -2.263552 | 0.023601656 | 0.314353976 |
| *ICMT* | -0.677418 | 0.299376 | -2.262767 | 0.023650032 | 0.314552125 |
| *NMI* | 0.249371 | 0.110241 | 2.262051 | 0.023694234 | 0.314613259 |
| *STX7* | 0.219384 | 0.09702 | 2.261225 | 0.023745335 | 0.314613259 |
| *DNAAF2* | 0.299748 | 0.132569 | 2.261066 | 0.023755144 | 0.314613259 |
| *ACSM1* | -0.805007 | 0.35607 | -2.260811 | 0.023770933 | NA |
| *TLE1* | -0.693686 | 0.306972 | -2.259769 | 0.023835613 | 0.315234377 |
| *VPS9D1* | -1.096947 | 0.486001 | -2.257086 | 0.024002674 | NA |
| *SORL1* | -0.295079 | 0.130786 | -2.256201 | 0.024058036 | 0.317728492 |
| *USPL1* | 0.503223 | 0.223132 | 2.255271 | 0.024116298 | 0.31805061 |
| *TOLLIP.AS1* | -0.898371 | 0.39844 | -2.254719 | 0.024150981 | NA |
| *ARRDC4* | -1.91203 | 0.848174 | -2.254289 | 0.024177995 | NA |
| *NAT14* | -0.629125 | 0.279285 | -2.252626 | 0.024282751 | 0.319796678 |
| *ENSG00000253948* | -0.563772 | 0.250357 | -2.251878 | 0.024329957 | 0.319969603 |
| *HFE* | -1.034696 | 0.459758 | -2.250524 | 0.024415711 | NA |
| *SLC30A1* | -1.149455 | 0.510771 | -2.250431 | 0.02442158 | NA |
| *DHRS4L2* | 0.452477 | 0.201067 | 2.25038 | 0.024424852 | 0.320768331 |
| *KMT2C* | 0.243814 | 0.108383 | 2.249567 | 0.02447643 | 0.320996754 |

| | | | | | |
|---|---|---|---|---|---|
| *GM2A* | 0.5507 | 0.244887 | 2.248797 | 0.02452539 | 0.321190254 |
| *RHOC* | -0.581917 | 0.258912 | -2.247547 | 0.024605072 | 0.321355972 |
| *STX5* | -0.240923 | 0.107195 | -2.247525 | 0.02460649 | 0.321355972 |
| *ADRB2* | 0.882184 | 0.392541 | 2.247367 | 0.024616572 | NA |
| *ZNF90* | -0.331622 | 0.147592 | -2.246886 | 0.024647303 | 0.321441907 |
| *OAS2* | 0.496481 | 0.221024 | 2.246278 | 0.024686198 | 0.32150263 |
| *UBA52* | 0.124058 | 0.05534 | 2.24175 | 0.024977528 | 0.324846243 |
| *GATM* | 0.953598 | 0.425551 | 2.240854 | 0.025035539 | 0.325048523 |
| *TOPORS* | 0.237629 | 0.106063 | 2.240441 | 0.025062314 | 0.325048523 |
| *ENSG00000228106* | -0.348156 | 0.155527 | -2.238559 | 0.025184599 | 0.326183978 |
| *MED11* | 0.243303 | 0.108719 | 2.237911 | 0.025226847 | 0.326281121 |
| *MXD4* | 0.419217 | 0.187411 | 2.236889 | 0.025293564 | 0.326609567 |
| *CNN3* | -0.633211 | 0.283131 | -2.236458 | 0.025321807 | 0.326609567 |
| *MRPS36* | 0.18978 | 0.084896 | 2.235447 | 0.025388029 | 0.326810273 |
| *IFT88* | -0.379353 | 0.169721 | -2.235158 | 0.025406975 | 0.326810273 |
| *PPP1CA* | -0.136524 | 0.061106 | -2.234228 | 0.025468081 | 0.327148129 |
| *IGHV3.15* | -1.135136 | 0.508247 | -2.233432 | 0.025520472 | NA |
| *STAC3* | -0.993426 | 0.444856 | -2.233138 | 0.025539858 | NA |
| *C1orf21* | -1.810576 | 0.810999 | -2.232526 | 0.025580243 | NA |
| *SPAG7* | -0.170155 | 0.076279 | -2.230677 | 0.025702501 | 0.329708306 |
| *KLHDC10* | -0.483652 | 0.216928 | -2.229556 | 0.025776932 | 0.33021199 |
| *APH1A* | -0.158532 | 0.071126 | -2.228907 | 0.025820125 | 0.3302735 |
| *FBXO22* | 0.275965 | 0.123866 | 2.22793 | 0.02588518 | 0.3302735 |
| *SLC25A5* | -0.152667 | 0.068529 | -2.227766 | 0.025896151 | 0.3302735 |
| *HDDC2* | -0.212119 | 0.095247 | -2.227031 | 0.025945221 | 0.3302735 |
| *ZNF718* | 0.804765 | 0.361425 | 2.226646 | 0.025970953 | 0.3302735 |
| *PELI2* | -0.608531 | 0.273344 | -2.22625 | 0.025997471 | 0.3302735 |
| *IFNLR1* | -0.490539 | 0.220388 | -2.225795 | 0.026027944 | 0.3302735 |
| *ENSG00000270820* | 0.764293 | 0.343462 | 2.225265 | 0.026063458 | 0.330277831 |
| *PLAAT3* | -1.543901 | 0.693936 | -2.224846 | 0.026091608 | NA |
| *TRIP11* | 0.274642 | 0.123558 | 2.222767 | 0.026231492 | 0.331846604 |
| *PDE4D* | -0.79384 | 0.35722 | -2.222273 | 0.026264858 | 0.331846604 |
| *C3orf33* | -0.813403 | 0.36608 | -2.221928 | 0.0262882 | NA |
| *JUP* | -0.694638 | 0.312639 | -2.221852 | 0.026293277 | 0.331846604 |
| *PIGZ* | -1.225145 | 0.551469 | -2.221605 | 0.026310031 | NA |
| *RPS16* | 0.151937 | 0.068453 | 2.219569 | 0.026448067 | 0.333304107 |
| *TXNL1* | 0.174406 | 0.078593 | 2.219103 | 0.026479751 | 0.333304107 |
| *CMTM3* | -0.82268 | 0.370826 | -2.218507 | 0.026520262 | 0.33336715 |
| *PTPN20* | -1.69566 | 0.764934 | -2.216739 | 0.026640919 | NA |
| *MAPKAPK2* | -0.355317 | 0.160304 | -2.216518 | 0.026656025 | 0.334625124 |
| *CYP3A5* | 0.468584 | 0.211519 | 2.215329 | 0.026737479 | 0.334625124 |
| *RAB31* | 0.640004 | 0.288923 | 2.215135 | 0.02675082 | 0.334625124 |
| *RPS8* | 0.170743 | 0.077086 | 2.214959 | 0.026762883 | 0.334625124 |
| *WIPF2* | 0.280429 | 0.12667 | 2.213844 | 0.026839494 | 0.3348799 |
| *IMPACT* | -0.459489 | 0.20758 | -2.213554 | 0.026859485 | 0.3348799 |
| *CUL2* | 0.373228 | 0.168644 | 2.213107 | 0.02689025 | 0.3348799 |
| *GNAI3* | -0.210352 | 0.095082 | -2.212322 | 0.026944447 | 0.335110404 |

| | | | | | |
|---|---|---|---|---|---|
| *UCP2* | -0.280931 | 0.127087 | -2.210532 | 0.027068292 | 0.336205379 |
| *CD69* | -0.305024 | 0.138049 | -2.209531 | 0.027137711 | 0.336622333 |
| *BASP1* | -0.325277 | 0.14748 | -2.205569 | 0.027414195 | 0.339603283 |
| *ATP6V0A1* | 0.430274 | 0.195139 | 2.204956 | 0.027457194 | 0.33968781 |
| *FGR* | -0.524923 | 0.238211 | -2.20361 | 0.027551795 | 0.340409672 |
| *LINC00926* | -0.215194 | 0.097722 | -2.202105 | 0.027657864 | 0.340605158 |
| *NDUFA6* | -0.175399 | 0.079657 | -2.201933 | 0.027670058 | 0.340605158 |
| *SERPINF1* | -0.873992 | 0.397 | -2.20149 | 0.027701351 | 0.340605158 |
| *YTHDC1* | 0.177826 | 0.080781 | 2.201329 | 0.027712709 | 0.340605158 |
| *FDXR* | -0.730491 | 0.332017 | -2.200162 | 0.027795368 | NA |
| *ALAS1* | 0.48397 | 0.220036 | 2.199501 | 0.027842283 | 0.341364533 |
| *POLR3K* | 0.253452 | 0.115249 | 2.199171 | 0.02786575 | 0.341364533 |
| *EPB41L2* | -0.569273 | 0.258887 | -2.198921 | 0.027883557 | 0.341364533 |
| *TRG.AS1* | 1.510739 | 0.687268 | 2.198181 | 0.02793621 | NA |
| *CCDC130* | 0.268664 | 0.122283 | 2.197058 | 0.028016291 | 0.342542929 |
| *ROR1* | -0.878881 | 0.400053 | -2.196912 | 0.028026751 | NA |
| *RAB13* | -0.823281 | 0.375187 | -2.194321 | 0.028212334 | 0.343634365 |
| *METTL15* | -0.312593 | 0.142473 | -2.194059 | 0.028231195 | 0.343634365 |
| *NFKB2* | -0.336243 | 0.153253 | -2.194042 | 0.028232418 | 0.343634365 |
| *NBPF9* | 0.513445 | 0.234063 | 2.193618 | 0.028262895 | 0.343634365 |
| *CARM1* | 0.489756 | 0.2233 | 2.193262 | 0.028288537 | 0.343634365 |
| *SWAP70* | 0.209191 | 0.095406 | 2.192631 | 0.02833397 | 0.343741576 |
| *NLRC5* | -0.44029 | 0.200875 | -2.191867 | 0.028389107 | 0.343966085 |
| *HIBCH* | 0.301183 | 0.137508 | 2.190294 | 0.028502952 | 0.344900408 |
| *THRAP3* | -0.230928 | 0.105489 | -2.189117 | 0.028588341 | 0.345264161 |
| *SINHCAF* | 0.24091 | 0.11009 | 2.188297 | 0.028647997 | 0.345264161 |
| *LMBRD1* | -0.188708 | 0.08624 | -2.188185 | 0.028656114 | 0.345264161 |
| *DAZAP1* | -0.303799 | 0.13888 | -2.187489 | 0.028706873 | 0.345264161 |
| *RPL34* | 0.119191 | 0.054491 | 2.187352 | 0.028716859 | 0.345264161 |
| *ENSG00000258302* | 1.439461 | 0.658133 | 2.187188 | 0.028728827 | NA |
| *NDUFAF4* | 0.294009 | 0.134481 | 2.186246 | 0.028797642 | 0.345353679 |
| *SHLD1* | -0.450191 | 0.20592 | -2.186243 | 0.028797863 | 0.345353679 |
| *APP* | 0.32251 | 0.147582 | 2.185293 | 0.028867346 | 0.345636404 |
| *RPS15A* | 0.152591 | 0.069839 | 2.184915 | 0.028895056 | 0.345636404 |
| *CRYM* | -1.951094 | 0.893163 | -2.184476 | 0.028927321 | NA |
| *EXOSC3* | 0.284212 | 0.130111 | 2.184374 | 0.028934751 | 0.345670884 |
| *METRNL* | 1.174133 | 0.537878 | 2.1829 | 0.02904317 | NA |
| *CTSH* | -0.299355 | 0.137228 | -2.181435 | 0.029151264 | 0.347814955 |
| *ACAA2* | -0.307573 | 0.141038 | -2.180779 | 0.029199748 | 0.347932515 |
| *PSMG4* | -0.315321 | 0.144623 | -2.1803 | 0.029235224 | 0.347932515 |
| *SAP18* | -0.124049 | 0.056964 | -2.17767 | 0.029430644 | 0.349802547 |
| *MGME1* | 0.268137 | 0.123158 | 2.177184 | 0.02946686 | 0.349802547 |
| *ZNF345* | 0.573071 | 0.263378 | 2.175851 | 0.029566382 | 0.349940697 |
| *CERS4* | -0.287415 | 0.13212 | -2.175407 | 0.029599604 | 0.349940697 |
| *ASCL2* | -1.215169 | 0.558738 | -2.174847 | 0.029641566 | NA |
| *RPS12* | 0.136187 | 0.062631 | 2.174449 | 0.029671439 | 0.349940697 |
| *FNBP1* | -0.289253 | 0.133032 | -2.174322 | 0.029680949 | 0.349940697 |

| | | | | | |
|---|---|---|---|---|---|
| *DCXR* | -0.27333 | 0.12571 | -2.174289 | 0.029683438 | 0.349940697 |
| *CABLES1* | -0.62706 | 0.288431 | -2.17404 | 0.029702102 | 0.349940697 |
| *ZNF358* | -0.444606 | 0.204617 | -2.172871 | 0.029790005 | 0.350536526 |
| *MED17* | 0.254988 | 0.117502 | 2.170082 | 0.030000611 | 0.352013862 |
| *MRPL43* | -0.192535 | 0.088762 | -2.169123 | 0.030073324 | 0.352013862 |
| *SNX5* | 0.225561 | 0.103995 | 2.16896 | 0.030085715 | 0.352013862 |
| *ST3GAL1* | 0.42875 | 0.197694 | 2.168756 | 0.030101211 | 0.352013862 |
| *ELOVL5* | 0.186207 | 0.085872 | 2.168437 | 0.030125482 | 0.352013862 |
| *IKZF3* | 0.165099 | 0.076144 | 2.168239 | 0.030140484 | 0.352013862 |
| *ENSG00000271895* | -0.896038 | 0.414089 | -2.163876 | 0.030473838 | NA |
| *KIF9.AS1* | 1.160133 | 0.536293 | 2.163245 | 0.030522378 | NA |
| *UQCRFS1* | -0.192724 | 0.089138 | -2.162092 | 0.030611063 | 0.35706569 |
| *ODC1* | -0.736137 | 0.340751 | -2.160335 | 0.030746733 | 0.358203253 |
| *PREX1* | 0.4147 | 0.192473 | 2.154593 | 0.031193688 | 0.362960012 |
| *CTSB* | 0.261021 | 0.121259 | 2.152599 | 0.031350225 | 0.364329971 |
| *MBP* | -0.228751 | 0.1064 | -2.149927 | 0.031561023 | 0.365704186 |
| *PPID* | 0.361163 | 0.167991 | 2.149892 | 0.031563795 | 0.365704186 |
| *ENSG00000253982* | -0.336155 | 0.156379 | -2.14962 | 0.031585314 | 0.365704186 |
| *CBY1* | 0.578267 | 0.269151 | 2.148487 | 0.031675116 | 0.36629229 |
| *LRRC47* | -0.555334 | 0.258637 | -2.147153 | 0.031781104 | 0.367065884 |
| *PRR5L* | -1.608997 | 0.749366 | -2.147143 | 0.031781881 | NA |
| *GNG7* | -0.166578 | 0.077633 | -2.145704 | 0.031896632 | 0.367947636 |
| *ERICH1* | 0.183139 | 0.085385 | 2.144861 | 0.031963923 | 0.368271461 |
| *KLB* | -1.568704 | 0.731463 | -2.144613 | 0.031983811 | NA |
| *POLR3D* | 0.376195 | 0.175485 | 2.143747 | 0.032053163 | 0.368847055 |
| *L1CAM* | -0.832631 | 0.388655 | -2.14234 | 0.032166153 | NA |
| *COG8* | 0.334498 | 0.156174 | 2.141824 | 0.032207637 | 0.36949918 |
| *MATR3* | -0.710117 | 0.331551 | -2.141806 | 0.032209061 | 0.36949918 |
| *THRB* | 1.551006 | 0.724241 | 2.141559 | 0.032228961 | NA |
| *ABRACL* | 0.336099 | 0.156981 | 2.141015 | 0.032272789 | 0.36949918 |
| *MIEN1* | -0.187822 | 0.087756 | -2.14027 | 0.032332928 | 0.36949918 |
| *PCYT1A* | 0.407309 | 0.190336 | 2.139942 | 0.03235947 | 0.36949918 |
| *BCL2L13* | -0.396896 | 0.185512 | -2.13946 | 0.032398424 | 0.36949918 |
| *SCAMP4* | -0.400914 | 0.1874 | -2.139346 | 0.03240765 | 0.36949918 |
| *RPL35* | 0.137769 | 0.064422 | 2.138549 | 0.032472213 | 0.36949918 |
| *RELB* | -0.337908 | 0.158017 | -2.138423 | 0.032482428 | 0.36949918 |
| *DDX23* | 0.339865 | 0.158996 | 2.137574 | 0.032551369 | 0.36949918 |
| *CYTH3* | -0.568964 | 0.26629 | -2.136633 | 0.032627884 | 0.36949918 |
| *ST6GALNAC6* | -0.240997 | 0.112817 | -2.136186 | 0.032664235 | 0.36949918 |
| *ZNF500* | 0.439966 | 0.205963 | 2.13614 | 0.032667985 | 0.36949918 |
| *ENSG00000228434* | 0.34828 | 0.163055 | 2.135969 | 0.032681918 | 0.36949918 |
| *SERPINB1* | -0.2132 | 0.099825 | -2.135746 | 0.032700087 | 0.36949918 |
| *TNKS1BP1* | -1.716027 | 0.803574 | -2.135493 | 0.032720768 | NA |
| *JTB* | -0.105467 | 0.049395 | -2.135188 | 0.032745667 | 0.369569482 |
| *CCDC121* | 1.336731 | 0.626266 | 2.134447 | 0.03280618 | NA |
| *MYO18A* | 1.010107 | 0.473288 | 2.134231 | 0.032823873 | NA |
| *HCST* | 0.852019 | 0.399592 | 2.132224 | 0.032988468 | 0.371721707 |

| | | | | | |
|---|---|---|---|---|---|
| *PEX16* | -0.187587 | 0.087991 | -2.131894 | 0.033015538 | 0.371721707 |
| *ENSG00000235501* | -0.9729 | 0.456605 | -2.130725 | 0.033111803 | NA |
| *NR6A1* | -0.789484 | 0.37053 | -2.130689 | 0.033114804 | NA |
| *NR2F6* | 0.974482 | 0.457364 | 2.130648 | 0.033118171 | NA |
| *ENSG00000267519* | -0.502407 | 0.23599 | -2.128934 | 0.033259735 | 0.374022651 |
| *CGAS* | 0.414205 | 0.194658 | 2.127864 | 0.033348372 | 0.37403956 |
| *DNAJA1* | 0.30746 | 0.144539 | 2.127173 | 0.033405718 | 0.37403956 |
| *SLC7A6* | 0.448629 | 0.210978 | 2.126426 | 0.033467791 | 0.37403956 |
| *GLTP* | -0.229345 | 0.107894 | -2.125657 | 0.033531808 | 0.37403956 |
| *ENSG00000272221* | 0.719362 | 0.338429 | 2.125593 | 0.033537169 | 0.37403956 |
| *PARP15* | 0.53892 | 0.253559 | 2.125417 | 0.033551796 | 0.37403956 |
| *LRRC59* | 0.351973 | 0.165622 | 2.125161 | 0.033573196 | 0.37403956 |
| *ARPC1A* | 0.229952 | 0.108208 | 2.12508 | 0.033579909 | 0.37403956 |
| *SMG9* | -0.297196 | 0.140003 | -2.122777 | 0.03377251 | 0.375657565 |
| *ENSG00000283013* | -0.265646 | 0.125185 | -2.122035 | 0.033834824 | 0.375657565 |
| *GATD1* | 0.33595 | 0.158324 | 2.121911 | 0.033845186 | 0.375657565 |
| *FAM149B1* | -0.692567 | 0.32651 | -2.121119 | 0.033911813 | NA |
| *ZBED5.AS1* | -0.339829 | 0.160216 | -2.121069 | 0.03391601 | 0.375724302 |
| *HELZ2* | 0.487976 | 0.230081 | 2.120888 | 0.033931226 | 0.375724302 |
| *FAM114A2* | 0.420066 | 0.198174 | 2.119684 | 0.034032664 | 0.3760526 |
| *CNR1* | 0.677918 | 0.319835 | 2.119586 | 0.03404097 | 0.3760526 |
| *IFRD2* | -0.231184 | 0.109103 | -2.118938 | 0.034095677 | 0.376214346 |
| *ABI1* | 0.22921 | 0.108286 | 2.116713 | 0.03428423 | 0.377095867 |
| *SIPA1L3* | -0.300803 | 0.142152 | -2.116064 | 0.034339356 | 0.377095867 |
| *SCNM1* | -0.170765 | 0.080714 | -2.115663 | 0.034373438 | 0.377095867 |
| *ENSG00000277654* | -0.473586 | 0.223871 | -2.115442 | 0.03439226 | 0.377095867 |
| *ZNF234* | 0.35334 | 0.167135 | 2.114093 | 0.034507365 | 0.377095867 |
| *HMGA1* | -0.274342 | 0.129781 | -2.113882 | 0.034525349 | 0.377095867 |
| *PTPRA* | -0.558446 | 0.264214 | -2.113613 | 0.03454834 | 0.377095867 |
| *CLMN* | 0.477013 | 0.225713 | 2.113362 | 0.034569793 | 0.377095867 |
| *PDE3B* | -0.414958 | 0.196354 | -2.113317 | 0.034573684 | 0.377095867 |
| *AGTRAP* | -0.380857 | 0.180221 | -2.113276 | 0.034577161 | 0.377095867 |
| *MPP6* | 0.36802 | 0.174232 | 2.112247 | 0.034665232 | 0.377357199 |
| *DPP3* | -0.471278 | 0.223137 | -2.112058 | 0.034681498 | 0.377357199 |
| *MDFIC* | 0.791488 | 0.374761 | 2.111981 | 0.034688095 | NA |
| *DYNC1LI2* | 0.299869 | 0.142031 | 2.111282 | 0.034748095 | 0.377644224 |
| *APOBEC3B* | -2.108066 | 0.99951 | -2.109099 | 0.034935996 | NA |
| *IL13RA1* | -0.614724 | 0.291512 | -2.108745 | 0.03496659 | 0.37944706 |
| *SNX8* | -0.401353 | 0.190363 | -2.108355 | 0.035000299 | 0.37944706 |
| *PXK* | 0.220265 | 0.104509 | 2.107616 | 0.03506423 | 0.37944706 |
| *APLP2* | -0.419192 | 0.198906 | -2.107484 | 0.035075617 | 0.37944706 |
| *ECI1* | -0.51488 | 0.244404 | -2.106678 | 0.035145497 | 0.379574887 |
| *INAFM1* | -0.525864 | 0.249649 | -2.106415 | 0.03516828 | 0.379574887 |
| *AFF2* | -1.131643 | 0.537358 | -2.105941 | 0.035209519 | NA |
| *ENSG00000273748* | -0.836944 | 0.397433 | -2.105874 | 0.035215341 | 0.379646441 |
| *PHF19* | -0.655375 | 0.311409 | -2.104547 | 0.035330746 | 0.380453793 |
| *ZFP82* | 0.365894 | 0.173942 | 2.103547 | 0.035417996 | 0.380956457 |

| | | | | | |
|---|---|---|---|---|---|
| *ENSG00000263080* | -1.273838 | 0.606006 | -2.102021 | 0.035551448 | NA |
| *IGLC2* | -0.665612 | 0.31672 | -2.10158 | 0.035590085 | 0.382016851 |
| *MIS18BP1* | 0.172707 | 0.082201 | 2.10105 | 0.035636545 | 0.382016851 |
| *ZNF622* | 0.251015 | 0.119473 | 2.101027 | 0.035638633 | 0.382016851 |
| *RPL5* | 0.135118 | 0.064373 | 2.098996 | 0.035817253 | 0.38349374 |
| *RP2* | 0.537969 | 0.256435 | 2.097877 | 0.035916027 | 0.384113322 |
| *INAVA* | -2.001256 | 0.95432 | -2.097048 | 0.035989299 | NA |
| *RNF26* | -0.339252 | 0.16179 | -2.096868 | 0.036005241 | 0.384629364 |
| *UBE2T* | 0.487243 | 0.232478 | 2.095868 | 0.03609394 | 0.385138743 |
| *TCL1A* | -0.353778 | 0.168852 | -2.095191 | 0.036154026 | 0.385237027 |
| *TP73.AS1* | -0.451535 | 0.215558 | -2.094726 | 0.036195315 | 0.385237027 |
| *SMIM27* | -0.257464 | 0.122931 | -2.094379 | 0.036226229 | 0.385237027 |
| *WDR66* | -0.819242 | 0.391374 | -2.093246 | 0.036327192 | 0.38566652 |
| *NUMA1* | 0.358348 | 0.171266 | 2.092352 | 0.036407015 | 0.38566652 |
| *RAC1* | -0.353913 | 0.169148 | -2.092324 | 0.036409499 | 0.38566652 |
| *ARPC5* | -0.151219 | 0.072295 | -2.091708 | 0.036464617 | 0.38566652 |
| *FADS3* | -0.264256 | 0.126362 | -2.091252 | 0.036505495 | 0.38566652 |
| *KHDRBS1* | -0.348965 | 0.166969 | -2.090003 | 0.036617572 | 0.38566652 |
| *CFL1* | -0.180319 | 0.086296 | -2.08955 | 0.036658274 | 0.38566652 |
| *SMARCE1* | -0.203028 | 0.097168 | -2.089442 | 0.036667957 | 0.38566652 |
| *NUPL2* | 0.254747 | 0.121923 | 2.089419 | 0.036669989 | 0.38566652 |
| *GCC2* | 0.228526 | 0.109377 | 2.089338 | 0.036677338 | 0.38566652 |
| *INTS3* | 0.346863 | 0.166103 | 2.088244 | 0.03677581 | 0.386269416 |
| *ZDHHC3* | -0.261446 | 0.125306 | -2.086462 | 0.036936796 | 0.387151784 |
| *PCNA* | 0.486717 | 0.233281 | 2.086401 | 0.036942279 | 0.387151784 |
| *RNF181* | -0.17129 | 0.082179 | -2.084344 | 0.037128865 | 0.388673404 |
| *ZBTB44* | 0.190833 | 0.091599 | 2.083351 | 0.03721921 | 0.38918528 |
| *LAPTM5* | -0.177142 | 0.085071 | -2.082277 | 0.037317198 | 0.389775845 |
| *ADPRHL2* | 0.279609 | 0.134351 | 2.08119 | 0.037416487 | 0.390152468 |
| *SPPL2A* | -0.20385 | 0.097959 | -2.080973 | 0.037436355 | 0.390152468 |
| *ACAP3* | 0.541 | 0.26008 | 2.080127 | 0.037513911 | 0.390279819 |
| *RBM25* | 0.164788 | 0.079235 | 2.079745 | 0.037548927 | 0.390279819 |
| *AUP1* | -0.163006 | 0.078419 | -2.078659 | 0.037648721 | 0.390279819 |
| *TRIB3* | 0.443367 | 0.213299 | 2.078619 | 0.037652387 | 0.390279819 |
| *PDE7B* | -0.472164 | 0.227158 | -2.078575 | 0.037656392 | 0.390279819 |
| *SLC5A6* | -0.409269 | 0.19697 | -2.077827 | 0.037725287 | 0.390562787 |
| *LACTB2* | 0.403507 | 0.194282 | 2.076917 | 0.037809183 | 0.391000247 |
| *ARID5B* | -0.183721 | 0.088501 | -2.075935 | 0.037899976 | 0.391280365 |
| *ABHD15* | -0.307562 | 0.148171 | -2.075723 | 0.037919609 | 0.391280365 |
| *GOSR2* | -0.322543 | 0.155482 | -2.074475 | 0.038035202 | 0.392042309 |
| *ZNHIT3* | 0.170507 | 0.082225 | 2.07367 | 0.038109936 | 0.392381904 |
| *ATRAID* | -0.17861 | 0.086208 | -2.071854 | 0.038279081 | 0.393487111 |
| *PFDN5* | 0.103412 | 0.049924 | 2.071402 | 0.038321252 | 0.393487111 |
| *GPSM3* | -0.098943 | 0.047772 | -2.071169 | 0.038342993 | 0.393487111 |
| *HINFP* | 0.342721 | 0.16551 | 2.070692 | 0.038387621 | 0.39351502 |
| *RAP2A* | 0.762256 | 0.368301 | 2.069657 | 0.038484445 | NA |
| *ENSG00000270189* | 0.55138 | 0.266682 | 2.067555 | 0.038681901 | 0.395219312 |

| | | | | | |
|---|---|---|---|---|---|
| *HSPA5* | 0.334863 | 0.161983 | 2.067276 | 0.038708155 | 0.395219312 |
| *SRPK1* | -0.273897 | 0.132493 | -2.067256 | 0.038710068 | 0.395219312 |
| *ERGIC2* | 0.221042 | 0.106932 | 2.067127 | 0.038722233 | 0.395219312 |
| *HNRNPA0* | -0.112035 | 0.054214 | -2.066553 | 0.038776274 | 0.395341161 |
| *POLDIP2* | -0.242816 | 0.117594 | -2.064865 | 0.03893576 | 0.396536646 |
| *RAB11FIP5* | -1.070835 | 0.518828 | -2.06395 | 0.039022479 | NA |
| *ABHD12* | -0.439992 | 0.213318 | -2.062612 | 0.03914948 | 0.39815827 |
| *SLC25A1* | -0.316675 | 0.153555 | -2.062294 | 0.039179791 | 0.39815827 |
| *GKAP1* | 0.288437 | 0.139913 | 2.061551 | 0.039250492 | 0.39827542 |
| *AHCYL1* | 0.270482 | 0.131224 | 2.061225 | 0.039281537 | 0.39827542 |
| *BRWD1* | -0.219949 | 0.106728 | -2.060837 | 0.039318564 | 0.39827542 |
| *CRNDE* | -1.620745 | 0.787312 | -2.058579 | 0.039534576 | NA |
| *ENSG00000268516* | 0.355781 | 0.172881 | 2.057952 | 0.039594724 | 0.400640579 |
| *ENSG00000262222* | -0.600937 | 0.292107 | -2.057246 | 0.039662613 | 0.400895523 |
| *DLG2* | 1.15147 | 0.559793 | 2.056955 | 0.039690545 | NA |
| *DTX3L* | 0.516388 | 0.251104 | 2.056468 | 0.039737448 | 0.401100313 |
| *UBQLN1* | 0.265586 | 0.129167 | 2.056147 | 0.039768306 | 0.401100313 |
| *CHCHD3* | -0.220864 | 0.107462 | -2.055275 | 0.039852417 | 0.401435006 |
| *RBM12B* | 0.376819 | 0.183386 | 2.054791 | 0.039899162 | 0.401435006 |
| *DHX57* | -0.377611 | 0.183907 | -2.05327 | 0.040046439 | 0.401435006 |
| *MRPL14* | -0.211942 | 0.103237 | -2.052972 | 0.040075316 | 0.401435006 |
| *TERF2IP* | 0.221451 | 0.107874 | 2.052861 | 0.040086033 | 0.401435006 |
| *BAG1* | -0.190218 | 0.092665 | -2.052758 | 0.040096102 | 0.401435006 |
| *CDK5RAP2* | -0.321694 | 0.156717 | -2.05271 | 0.040100749 | 0.401435006 |
| *DOCK8* | 0.284733 | 0.138764 | 2.051927 | 0.040176786 | 0.4015242 |
| *UBE2J2* | -0.210887 | 0.102784 | -2.051737 | 0.040195181 | 0.4015242 |
| *LPCAT4* | 0.345506 | 0.168736 | 2.047608 | 0.040598428 | 0.405121402 |
| *NUDT12* | 0.8107 | 0.396047 | 2.04698 | 0.040660086 | NA |
| *CHCHD5* | -0.294622 | 0.14393 | -2.046973 | 0.040660719 | 0.405312264 |
| *TMEM231* | -1.153115 | 0.563584 | -2.046039 | 0.040752513 | NA |
| *PRR13* | -0.145965 | 0.071372 | -2.045114 | 0.040843586 | 0.4053796 |
| *CYB5R3* | 0.24117 | 0.117943 | 2.044796 | 0.040874939 | 0.4053796 |
| *KLC4* | 0.428557 | 0.209721 | 2.043461 | 0.041006792 | 0.4053796 |
| *HIST1H2BN* | 0.537565 | 0.263091 | 2.043267 | 0.04102599 | 0.4053796 |
| *CHMP2B* | 0.224817 | 0.11004 | 2.043058 | 0.04104666 | 0.4053796 |
| *RSBN1L* | 0.180861 | 0.088525 | 2.043049 | 0.041047562 | 0.4053796 |
| *MZF1* | 0.409398 | 0.200526 | 2.041622 | 0.041189088 | 0.4053796 |
| *PF4* | 3.158337 | 1.546978 | 2.041617 | 0.0411895 | NA |
| *RACK1* | 0.116613 | 0.057123 | 2.041433 | 0.041207838 | 0.4053796 |
| *CEP57* | 0.185106 | 0.090686 | 2.041179 | 0.041233018 | 0.4053796 |
| *ZNF865* | -0.548254 | 0.268644 | -2.040817 | 0.041269055 | 0.4053796 |
| *AZIN1* | -0.260124 | 0.127463 | -2.040782 | 0.041272465 | 0.4053796 |
| *APH1B* | -0.40471 | 0.198318 | -2.040716 | 0.041279068 | 0.4053796 |
| *RAB9A* | 0.342886 | 0.168023 | 2.040704 | 0.041280241 | 0.4053796 |
| *ZBTB1* | 0.296125 | 0.145139 | 2.040286 | 0.041321809 | 0.4053796 |
| *DAD1* | -0.169094 | 0.082881 | -2.04019 | 0.041331437 | 0.4053796 |
| *ENSG00000224521* | -1.733902 | 0.849903 | -2.040118 | 0.041338565 | NA |

| | | | | | |
|---|---|---|---|---|---|
| *TMED4* | -0.208415 | 0.102168 | -2.039921 | 0.041358217 | 0.4053796 |
| *ZMIZ2* | -0.396226 | 0.194288 | -2.03937 | 0.041413147 | 0.405494737 |
| *EFHB* | 1.581973 | 0.775981 | 2.038674 | 0.041482624 | NA |
| *FUT7* | 1.506134 | 0.739024 | 2.038004 | 0.041549517 | NA |
| *NUP188* | 0.430951 | 0.2115 | 2.037595 | 0.041590454 | 0.406298289 |
| *RAB30* | 0.176617 | 0.086684 | 2.037496 | 0.041600387 | 0.406298289 |
| *DRAM1* | -0.518963 | 0.254737 | -2.03725 | 0.041625022 | 0.406298289 |
| *USP44* | 0.681509 | 0.334635 | 2.036576 | 0.041692524 | NA |
| *HNRNPA1L2* | -0.321812 | 0.158031 | -2.036385 | 0.04171167 | 0.406721273 |
| *CENPF* | -0.748229 | 0.367497 | -2.036015 | 0.041748876 | NA |
| *TUNAR* | -1.372752 | 0.674335 | -2.035713 | 0.041779171 | NA |
| *RUBCNL* | -0.21954 | 0.107853 | -2.035547 | 0.04179588 | 0.407119618 |
| *SERGEF* | 0.37133 | 0.182601 | 2.033562 | 0.041995823 | 0.408643297 |
| *MED21* | 0.356508 | 0.17551 | 2.031275 | 0.042227128 | 0.409558319 |
| *CD2* | -2.600764 | 1.28053 | -2.031006 | 0.042254393 | NA |
| *HNRNPL* | -0.266722 | 0.131377 | -2.030197 | 0.042336471 | 0.409558319 |
| *MYL12A* | 0.1294 | 0.06374 | 2.030111 | 0.042345287 | 0.409558319 |
| *EEFSEC* | -0.323095 | 0.159164 | -2.029953 | 0.042361326 | 0.409558319 |
| *KLC1* | 0.301492 | 0.148545 | 2.029633 | 0.04239388 | 0.409558319 |
| *RPF2* | 0.307267 | 0.151415 | 2.029312 | 0.042426495 | 0.409558319 |
| *ENSG00000246790* | -0.623532 | 0.307271 | -2.029255 | 0.042432326 | 0.409558319 |
| *MRPL21* | 0.23687 | 0.116737 | 2.029095 | 0.042448581 | 0.409558319 |
| *HSPE1* | 0.160059 | 0.078903 | 2.028542 | 0.042504932 | 0.409558319 |
| *MAU2* | -0.362268 | 0.178618 | -2.028176 | 0.04254227 | 0.409558319 |
| *CNBD2* | -0.451097 | 0.222445 | -2.027908 | 0.04256964 | 0.409558319 |
| *ENSG00000259038* | -0.675706 | 0.333315 | -2.027229 | 0.042639016 | NA |
| *NACA* | 0.12199 | 0.060214 | 2.025926 | 0.042772392 | 0.410323033 |
| *DHX15* | 0.257623 | 0.127165 | 2.025885 | 0.042776532 | 0.410323033 |
| *ZBED1* | -0.368215 | 0.181758 | -2.025849 | 0.042780218 | 0.410323033 |
| *VPS37C* | -0.66734 | 0.32955 | -2.025005 | 0.042866847 | NA |
| *MEX3C* | 0.294154 | 0.145345 | 2.023838 | 0.042986827 | 0.410428198 |
| *ENSG00000283103* | 0.311725 | 0.154046 | 2.023591 | 0.043012232 | 0.410428198 |
| *ENSG00000273270* | -0.471284 | 0.232908 | -2.023481 | 0.043023581 | 0.410428198 |
| *INPP5B* | 0.346979 | 0.171485 | 2.02338 | 0.043034027 | 0.410428198 |
| *KCNK6* | 0.458079 | 0.2264 | 2.023315 | 0.043040724 | 0.410428198 |
| *COPG2* | -0.585705 | 0.289496 | -2.023191 | 0.043053437 | 0.410428198 |
| *ITGB2.AS1* | -1.8836 | 0.931132 | -2.022914 | 0.043082027 | NA |
| *C1orf35* | 0.202304 | 0.100044 | 2.022151 | 0.043160803 | 0.410545591 |
| *ANKLE2* | -0.330305 | 0.163349 | -2.022079 | 0.043168238 | 0.410545591 |
| *PMF1* | -0.222601 | 0.1101 | -2.021801 | 0.043196916 | 0.410545591 |
| *MIB1* | 0.542843 | 0.268606 | 2.020965 | 0.043283426 | 0.410659524 |
| *HIP1R* | -0.269899 | 0.133558 | -2.02084 | 0.043296372 | 0.410659524 |
| *ZNF74* | 0.855972 | 0.423813 | 2.019695 | 0.043415008 | NA |
| *ENSG00000261770* | 0.890543 | 0.4411 | 2.018914 | 0.043496146 | NA |
| *SPN* | -0.868668 | 0.430413 | -2.018221 | 0.043568221 | NA |
| *TMUB1* | -0.223652 | 0.110824 | -2.018096 | 0.043581316 | 0.412928502 |
| *ENSG00000271133* | -1.70875 | 0.846858 | -2.017754 | 0.043616939 | NA |

| | | | | | |
|---|---|---|---|---|---|
| *ATP5IF1* | -0.124159 | 0.06155 | -2.017208 | 0.043673761 | 0.412928502 |
| *CBX6* | -0.173391 | 0.085967 | -2.016961 | 0.043699625 | 0.412928502 |
| *ZBTB43* | -0.347655 | 0.172376 | -2.016847 | 0.043711494 | 0.412928502 |
| *HSPA8* | 0.257992 | 0.127962 | 2.016158 | 0.043783442 | 0.41319248 |
| *ARL2BP* | -0.257589 | 0.127824 | -2.015185 | 0.043885239 | 0.413737344 |
| *IGHA2* | -0.968913 | 0.480942 | -2.014615 | 0.043945053 | 0.413752165 |
| *ZNF174* | 0.491474 | 0.243989 | 2.01433 | 0.043974937 | 0.413752165 |
| *ENSG00000234936* | -0.847418 | 0.421195 | -2.011939 | 0.044226389 | NA |
| *SIGIRR* | -0.217889 | 0.108312 | -2.01169 | 0.044252668 | 0.415948503 |
| *CEP295* | 0.32575 | 0.162008 | 2.010709 | 0.044356173 | 0.416103904 |
| *FGD2* | 0.201837 | 0.100382 | 2.010694 | 0.044357828 | 0.416103904 |
| *CEP95* | 0.304247 | 0.151403 | 2.009516 | 0.044482441 | 0.416603229 |
| *MANF* | 0.240253 | 0.119567 | 2.009352 | 0.044499791 | 0.416603229 |
| *ZNF138* | 0.370673 | 0.184644 | 2.0075 | 0.04469647 | 0.417650493 |
| *THOC6* | -0.277715 | 0.138342 | -2.007461 | 0.044700612 | 0.417650493 |
| *INPP5K* | -0.388239 | 0.19352 | -2.006198 | 0.044835093 | 0.418021336 |
| *VDAC1* | 0.313071 | 0.156088 | 2.005726 | 0.0448855 | 0.418021336 |
| *FNIP1* | -0.265683 | 0.132469 | -2.005629 | 0.044895855 | 0.418021336 |
| *ZNF785* | 0.340177 | 0.169651 | 2.005164 | 0.044945528 | 0.418021336 |
| *TXLNG* | 0.349464 | 0.174296 | 2.005001 | 0.044962891 | 0.418021336 |
| *COLCA2* | -0.888989 | 0.443481 | -2.004571 | 0.045008941 | NA |
| *ANKRD12* | -0.178774 | 0.089232 | -2.00347 | 0.045126864 | 0.418624462 |
| *TEX264* | -0.214608 | 0.107157 | -2.002751 | 0.045204007 | 0.418624462 |
| *ACSF3* | 0.276478 | 0.138055 | 2.002664 | 0.045213386 | 0.418624462 |
| *ALG13* | -0.231193 | 0.115445 | -2.002628 | 0.045217218 | 0.418624462 |
| *CLSPN* | -1.136596 | 0.567576 | -2.002544 | 0.045226309 | NA |
| *BCR* | 0.451057 | 0.225281 | 2.002198 | 0.045263466 | 0.418624462 |
| *DTD2* | -0.473358 | 0.236454 | -2.001902 | 0.045295256 | 0.418624462 |
| *AEBP2* | 0.349603 | 0.174708 | 2.001074 | 0.04538438 | 0.419035718 |
| *TAF6L* | -0.682853 | 0.341438 | -1.999931 | 0.045507764 | 0.419762188 |
| *GINM1* | 0.305392 | 0.152792 | 1.998745 | 0.045635906 | 0.420230332 |
| *MGAT5B* | -0.594044 | 0.297225 | -1.998633 | 0.045648023 | 0.420230332 |
| *ZPBP2* | -1.135336 | 0.569019 | -1.995253 | 0.046015245 | NA |
| *NINL* | -1.296684 | 0.65003 | -1.994805 | 0.046064104 | NA |
| *DPH5* | 0.293308 | 0.14711 | 1.993798 | 0.046174082 | 0.42436526 |
| *MIS12* | 0.322968 | 0.161996 | 1.993675 | 0.046187571 | 0.42436526 |
| *SOCS4* | 0.423835 | 0.212727 | 1.992389 | 0.04632841 | 0.425243177 |
| *CDKN1B* | 0.180564 | 0.090681 | 1.991198 | 0.046459149 | 0.425903445 |
| *CTDP1* | -0.648776 | 0.325897 | -1.990737 | 0.046509789 | 0.425903445 |
| *NOP9* | 0.638462 | 0.320787 | 1.990301 | 0.046557817 | NA |
| *SURF4* | -0.252835 | 0.127044 | -1.990133 | 0.046576309 | 0.425903445 |
| *RPL21* | 0.121054 | 0.060838 | 1.989762 | 0.046617126 | 0.425903445 |
| *ARHGAP30* | 0.204894 | 0.102979 | 1.989672 | 0.046627129 | 0.425903445 |
| *LDB1* | 0.767572 | 0.38578 | 1.98966 | 0.046628353 | NA |
| *TTC7A* | -0.374097 | 0.188098 | -1.988845 | 0.046718282 | 0.426321352 |
| *P2RX1* | -0.350855 | 0.176514 | -1.98769 | 0.046846 | 0.42707179 |
| *GATC* | 0.338515 | 0.170347 | 1.98721 | 0.046899162 | 0.42714174 |

| | | | | | |
|---|---|---|---|---|---|
| *BTBD3* | -0.709347 | 0.35709 | -1.986467 | 0.046981447 | NA |
| *ELMO3* | -0.759993 | 0.382653 | -1.986117 | 0.047020336 | NA |
| *KMT2A* | 0.198344 | 0.099866 | 1.986102 | 0.047021946 | 0.427845034 |
| *TAF9* | -0.213637 | 0.107623 | -1.985039 | 0.047140136 | 0.428505207 |
| *HMGCL* | 0.319202 | 0.160865 | 1.984287 | 0.047223874 | 0.428851236 |
| *TCF25* | 0.152731 | 0.076999 | 1.983543 | 0.047306848 | 0.429078598 |
| *PRADC1* | -0.476493 | 0.240288 | -1.983007 | 0.047366667 | 0.429078598 |
| *IGIP* | 0.918936 | 0.463447 | 1.982829 | 0.047386521 | NA |
| *INSR* | -0.590533 | 0.29788 | -1.982451 | 0.047428746 | 0.429078598 |
| *TRAF3IP1* | -0.683633 | 0.34485 | -1.982408 | 0.04743358 | NA |
| *SYK* | 0.173307 | 0.087427 | 1.982304 | 0.047445246 | 0.429078598 |
| *NIPAL2* | 0.784683 | 0.395861 | 1.982218 | 0.047454874 | NA |
| *BSDC1* | 0.199557 | 0.100684 | 1.982017 | 0.047477387 | 0.429078598 |
| *IER2* | -0.177019 | 0.089357 | -1.981042 | 0.047586522 | 0.429647581 |
| *EXOSC7* | -0.27955 | 0.141141 | -1.980638 | 0.047631856 | 0.429647581 |
| *NREP* | -0.40684 | 0.205528 | -1.979489 | 0.047761032 | 0.430399316 |
| *NDC80* | 1.199463 | 0.60616 | 1.978789 | 0.047839804 | NA |
| *PSME3* | -0.359801 | 0.181907 | -1.977939 | 0.047935604 | 0.430913836 |
| *SLTM* | 0.149808 | 0.07574 | 1.977925 | 0.04793712 | 0.430913836 |
| *SNHG30* | -0.262755 | 0.132855 | -1.97776 | 0.0479558 | 0.430913836 |
| *P4HB* | -0.193851 | 0.098045 | -1.977163 | 0.048023217 | 0.431107081 |
| *CDK6* | -0.533176 | 0.269922 | -1.975296 | 0.048234592 | 0.432455343 |
| *VAMP2* | -0.153303 | 0.077621 | -1.975023 | 0.048265516 | 0.432455343 |
| *KLHL5* | -0.372646 | 0.188746 | -1.974321 | 0.048345284 | 0.432757122 |
| *TOMM34* | 0.329322 | 0.166912 | 1.97303 | 0.048492097 | 0.433590982 |
| *ZNF28* | 0.655319 | 0.332195 | 1.972691 | 0.04853079 | 0.433590982 |
| *SAR1B* | 0.307308 | 0.155892 | 1.971285 | 0.048691247 | 0.433851034 |
| *TMEM187* | 0.510546 | 0.259 | 1.97122 | 0.048698742 | 0.433851034 |
| *RND1* | -0.577673 | 0.293069 | -1.971118 | 0.048710405 | 0.433851034 |
| *BET1L* | 0.298595 | 0.151508 | 1.970818 | 0.048744711 | 0.433851034 |
| *NIPSNAP2* | -0.188966 | 0.095934 | -1.969752 | 0.048866781 | 0.434525635 |
| *CLK3* | 0.182957 | 0.092935 | 1.968656 | 0.04899264 | 0.435232632 |
| *SCAF11* | 0.121093 | 0.061589 | 1.966159 | 0.049280288 | 0.437374202 |
| *PAQR4* | -1.475493 | 0.750484 | -1.966054 | 0.04929243 | NA |
| *NAGK* | 0.245674 | 0.125043 | 1.964717 | 0.049446995 | 0.438037442 |
| *IARS2* | 0.303225 | 0.154336 | 1.964706 | 0.049448316 | 0.438037442 |
| *MREG* | -0.733682 | 0.373648 | -1.963565 | 0.049580566 | NA |
| *MYL6* | -0.135468 | 0.069009 | -1.963049 | 0.049640463 | 0.438342841 |
| *SMIM12* | -0.188649 | 0.096102 | -1.963015 | 0.049644406 | 0.438342841 |
| *ARRDC2* | -0.258979 | 0.131937 | -1.962889 | 0.049659089 | 0.438342841 |
| *RAN* | 0.130253 | 0.066361 | 1.962799 | 0.049669519 | 0.438342841 |
| *ZFP62* | -0.296203 | 0.150986 | -1.961785 | 0.049787473 | 0.438391688 |
| *ADAMTS7* | -0.740964 | 0.377717 | -1.961692 | 0.049798326 | 0.438391688 |
| *TSGA10* | 0.485954 | 0.247759 | 1.961398 | 0.049832555 | 0.438391688 |
| *CLOCK* | -0.288394 | 0.147054 | -1.961148 | 0.049861802 | 0.438391688 |

| Cluster 2 | log2FC | lfcSE | stat | pvalue | padj |
|---|---|---|---|---|---|
| *ARRDC3* | -1.177752 | 0.110552 | -10.65342 | < 2.22e-16 | < 2.22e-16 |
| *NFKBIA* | -0.519261 | 0.069716 | -7.448257 | 9.46E-14 | 3.14E-10 |
| *H3F3B* | -0.3961 | 0.054845 | -7.222115 | 5.12E-13 | 1.13E-09 |
| *CD83* | -0.465405 | 0.065285 | -7.128808 | 1.01E-12 | 1.68E-09 |
| *CKS2* | 1.220771 | 0.183998 | 6.634679 | 3.25E-11 | 4.32E-08 |
| *GPR183* | 0.855737 | 0.146652 | 5.835136 | 5.37E-09 | 5.95E-06 |
| *CHORDC1* | 0.751185 | 0.135081 | 5.560986 | 2.68E-08 | 2.54E-05 |
| *CHMP6* | -0.798552 | 0.148419 | -5.380405 | 7.43E-08 | 6.17E-05 |
| *RSRP1* | 0.398926 | 0.074696 | 5.340668 | 9.26E-08 | 6.83E-05 |
| *SMIM10* | -3.791334 | 0.710869 | -5.333376 | 9.64E-08 | NA |
| *GATD3A* | 4.608235 | 0.895634 | 5.145222 | 2.67E-07 | 0.00017737 |
| *CDKN1A* | -0.777446 | 0.152389 | -5.101708 | 3.37E-07 | 0.000203124 |
| *CD79B* | -0.30552 | 0.061409 | -4.9752 | 6.52E-07 | 0.000336284 |
| *CLK1* | 0.536565 | 0.107891 | 4.973194 | 6.59E-07 | 0.000336284 |
| *NOP10* | 0.268179 | 0.055204 | 4.857916 | 1.19E-06 | 0.000562466 |
| *HMGB2* | -0.404119 | 0.086356 | -4.679673 | 2.87E-06 | 0.001271543 |
| *ARL17A* | -0.719897 | 0.154759 | -4.651723 | 3.29E-06 | 0.001365657 |
| *TAGAP* | -0.52709 | 0.113688 | -4.636295 | 3.55E-06 | 0.001385039 |
| *BLNK* | 0.327873 | 0.072122 | 4.546109 | 5.46E-06 | 0.001997552 |
| *ERP29* | 0.28176 | 0.062109 | 4.536572 | 5.72E-06 | 0.001997552 |
| *ATF7IP2* | -0.688388 | 0.152688 | -4.508469 | 6.53E-06 | 0.002167211 |
| *CXCR4* | 0.180207 | 0.040347 | 4.466365 | 7.96E-06 | 0.002514843 |
| *KRTCAP2* | -0.298987 | 0.067355 | -4.438965 | 9.04E-06 | 0.002633078 |
| *IGHD* | -0.443708 | 0.100003 | -4.436971 | 9.12E-06 | 0.002633078 |
| *MIF* | -0.405706 | 0.092884 | -4.367875 | 1.25E-05 | 0.003470044 |
| *SLC3A2* | -0.485364 | 0.112259 | -4.323628 | 1.53E-05 | 0.003894959 |
| *TTC32* | -0.70869 | 0.16416 | -4.31706 | 1.58E-05 | 0.003894959 |
| *TIMM10* | -0.522847 | 0.121124 | -4.316633 | 1.58E-05 | 0.003894959 |
| *SNHG9* | -0.479621 | 0.112649 | -4.257672 | 2.07E-05 | 0.004897097 |
| *EMP3* | -0.395834 | 0.09465 | -4.182092 | 2.89E-05 | 0.006265733 |
| *HSBP1* | 0.355446 | 0.085127 | 4.175497 | 2.97E-05 | 0.006265733 |
| *RPL8* | 0.378487 | 0.090702 | 4.172846 | 3.01E-05 | 0.006265733 |
| *IGLC2* | -0.417639 | 0.100107 | -4.171912 | 3.02E-05 | 0.006265733 |
| *LCN8* | 3.779297 | 0.91018 | 4.15225 | 3.29E-05 | 0.006622356 |
| *C16orf74* | -0.626001 | 0.151076 | -4.143606 | 3.42E-05 | 0.00662472 |
| *C7orf50* | -0.46358 | 0.112011 | -4.138687 | 3.49E-05 | 0.00662472 |
| *ENSG00000273319* | 0.595867 | 0.144949 | 4.110884 | 3.94E-05 | 0.007118565 |
| *PIM1* | -0.484403 | 0.117878 | -4.109343 | 3.97E-05 | 0.007118565 |
| *JMJD6* | 0.803095 | 0.196072 | 4.095909 | 4.21E-05 | 0.007345732 |
| *SRSF5* | -0.310027 | 0.075984 | -4.080149 | 4.50E-05 | 0.007660381 |
| *ENSG00000272211* | -1.000086 | 0.247674 | -4.03792 | 5.39E-05 | 0.008949206 |
| *DNAJA1* | 0.449953 | 0.11191 | 4.020684 | 5.80E-05 | 0.009364755 |
| *TCL1B* | -1.397902 | 0.348103 | -4.015767 | 5.93E-05 | 0.009364755 |
| *RIN3* | 0.699153 | 0.174454 | 4.00766 | 6.13E-05 | 0.009466618 |
| *CLEC2D* | -0.4074 | 0.102081 | -3.99093 | 6.58E-05 | 0.009929051 |
| *GPR18* | 0.915276 | 0.230187 | 3.976223 | 7.00E-05 | NA |

| | | | | | |
|---|---|---|---|---|---|
| *SLC50A1* | -0.471271 | 0.118949 | -3.961948 | 7.43E-05 | 0.010966079 |
| *CCT8* | 0.604433 | 0.152816 | 3.955306 | 7.64E-05 | 0.011030136 |
| *ENSG00000235078* | 0.897166 | 0.227923 | 3.936271 | 8.28E-05 | NA |
| *RBM3* | -0.381252 | 0.097665 | -3.903668 | 9.47E-05 | 0.013381326 |
| *H2AFZ* | 0.270115 | 0.069346 | 3.895181 | 9.81E-05 | 0.01356992 |
| *ENSG00000224505* | -0.518755 | 0.134323 | -3.862001 | 0.00011246 | 0.015156338 |
| *CIRBP* | -0.20865 | 0.054078 | -3.858332 | 0.00011416 | 0.015156338 |
| *ADAMTS6* | 1.038987 | 0.270359 | 3.842992 | 0.00012154 | 0.015819682 |
| *GPR65* | 0.582748 | 0.1529 | 3.811294 | 0.00013824 | 0.017402028 |
| *TCTN1* | -1.118132 | 0.293763 | -3.806232 | 0.0001411 | 0.017402028 |
| *PNPLA8* | 0.372045 | 0.097767 | 3.805418 | 0.00014157 | 0.017402028 |
| *YWHAE* | 0.332198 | 0.087523 | 3.795569 | 0.00014731 | 0.01753509 |
| *C6orf226* | -0.657625 | 0.173309 | -3.794517 | 0.00014793 | 0.01753509 |
| *SRSF11* | 0.243417 | 0.06425 | 3.788566 | 0.00015152 | 0.017645355 |
| *FCGR2B* | 0.781868 | 0.207361 | 3.770556 | 0.00016288 | 0.018635642 |
| *GAPT* | 0.454182 | 0.120589 | 3.766371 | 0.00016564 | 0.018635642 |
| *MED21* | 0.590251 | 0.157641 | 3.744268 | 0.00018092 | 0.019823043 |
| *GLRX* | 0.411067 | 0.109836 | 3.742547 | 0.00018216 | 0.019823043 |
| *ATP6V0E2* | -0.748252 | 0.202734 | -3.690805 | 0.00022355 | 0.023933828 |
| *TDG* | 0.414617 | 0.113062 | 3.667155 | 0.00024526 | 0.025842274 |
| *CYB5A* | -0.615565 | 0.168883 | -3.644924 | 0.00026747 | 0.027691383 |
| *SCRN1* | 1.100879 | 0.30231 | 3.641552 | 0.000271 | NA |
| *MRPL32* | 0.368212 | 0.101178 | 3.639261 | 0.00027342 | 0.027691383 |
| *ZBED2* | 1.836315 | 0.505329 | 3.633903 | 0.00027917 | NA |
| *EPS15* | -0.598349 | 0.164815 | -3.630421 | 0.00028296 | 0.027691383 |
| *ZFAND2A* | -0.550626 | 0.151758 | -3.628322 | 0.00028527 | 0.027691383 |
| *PPM1K* | 0.261652 | 0.072149 | 3.626555 | 0.00028723 | 0.027691383 |
| *PMF1* | -0.388777 | 0.107219 | -3.626002 | 0.00028784 | 0.027691383 |
| *SMIM4* | -0.430031 | 0.119555 | -3.596922 | 0.00032201 | 0.030535312 |
| *PTGIR* | -1.757506 | 0.490371 | -3.584032 | 0.00033833 | NA |
| *TNFSF10* | 0.593193 | 0.16605 | 3.572367 | 0.00035377 | 0.033074967 |
| *JADE1* | 0.430241 | 0.1214 | 3.543992 | 0.00039412 | 0.035693292 |
| *TMEM9B* | -0.392535 | 0.110957 | -3.537736 | 0.00040357 | 0.035693292 |
| *MRPL54* | -0.395262 | 0.111741 | -3.537295 | 0.00040425 | 0.035693292 |
| *UBE2L3* | -0.254563 | 0.072021 | -3.534546 | 0.00040848 | 0.035693292 |
| *SLC16A7* | 0.448153 | 0.126796 | 3.534427 | 0.00040866 | 0.035693292 |
| *BOLA2B* | -0.656056 | 0.186723 | -3.513531 | 0.00044219 | 0.038057031 |
| *PSMA4* | -0.262787 | 0.074857 | -3.510544 | 0.00044719 | 0.038057031 |
| *STX7* | 0.275863 | 0.07883 | 3.499448 | 0.00046622 | 0.039174463 |
| *EXOSC3* | 0.511944 | 0.146527 | 3.493869 | 0.00047608 | 0.039502285 |
| *CREBRF* | 0.307288 | 0.088038 | 3.490393 | 0.00048231 | 0.03952566 |
| *PIGF* | 0.670396 | 0.19241 | 3.484197 | 0.00049362 | 0.03995878 |
| *HSPE1* | 0.251856 | 0.072368 | 3.480204 | 0.00050103 | 0.040070533 |
| *ZCCHC9* | 0.472834 | 0.136043 | 3.475622 | 0.00050967 | 0.040276106 |
| *TNFRSF14* | 0.216856 | 0.062655 | 3.461119 | 0.00053793 | 0.041575926 |
| *SNAPC3* | 0.435739 | 0.125908 | 3.460764 | 0.00053865 | 0.041575926 |
| *AP4B1* | 0.930623 | 0.269796 | 3.449359 | 0.00056192 | NA |

| | | | | | |
|---|---|---|---|---|---|
| *PDE4DIP* | -0.730859 | 0.212573 | -3.438153 | 0.0005857 | 0.044440773 |
| *IL21R* | -0.841727 | 0.244958 | -3.436215 | 0.0005899 | NA |
| *COMMD3* | -0.391135 | 0.113848 | -3.435582 | 0.00059128 | 0.044440773 |
| *IFIT2* | 0.696895 | 0.203159 | 3.430294 | 0.00060293 | 0.044440773 |
| *CYTIP* | 0.236633 | 0.06899 | 3.429976 | 0.00060363 | 0.044440773 |
| *ELMSAN1* | 0.389727 | 0.113707 | 3.427469 | 0.00060924 | 0.044440773 |
| *ROR1* | -1.299246 | 0.379567 | -3.422969 | 0.00061941 | NA |
| *IZUMO4* | -0.707578 | 0.207504 | -3.409944 | 0.00064976 | 0.046881794 |
| *RPL30* | 0.155733 | 0.046092 | 3.378734 | 0.0007282 | 0.051161528 |
| *RHOH* | 0.241595 | 0.07151 | 3.378473 | 0.0007289 | 0.051161528 |
| *DMAC2L* | -0.333608 | 0.098861 | -3.374534 | 0.00073941 | 0.051161528 |
| *ZNF441* | -0.663633 | 0.196749 | -3.372991 | 0.00074356 | 0.051161528 |
| *UCP2* | -0.344525 | 0.102237 | -3.369861 | 0.00075206 | 0.051161528 |
| *CD48* | 0.333025 | 0.09886 | 3.368668 | 0.00075532 | 0.051161528 |
| *WDR13* | -0.358117 | 0.106677 | -3.357012 | 0.0007879 | 0.052828943 |
| *SGK1* | -1.622911 | 0.484192 | -3.351795 | 0.0008029 | NA |
| *EIF4A2* | 0.220356 | 0.066073 | 3.335052 | 0.00085284 | 0.056611173 |
| *ENSG00000237596* | -1.005042 | 0.30171 | -3.331156 | 0.00086486 | NA |
| *IL23A* | -1.016229 | 0.305228 | -3.329407 | 0.00087031 | 0.056663465 |
| *IL2RG* | -0.410417 | 0.123275 | -3.329284 | 0.0008707 | 0.056663465 |
| *CRIPT* | 0.406671 | 0.122514 | 3.319372 | 0.0009022 | 0.058143904 |
| *POU2AF1* | -0.343366 | 0.103625 | -3.313531 | 0.00092126 | 0.058292334 |
| *UBE2J2* | -0.331044 | 0.099914 | -3.313285 | 0.00092207 | 0.058292334 |
| *ENSG00000237491* | -0.854833 | 0.258775 | -3.303381 | 0.00095527 | NA |
| *PRICKLE1* | 0.844305 | 0.255656 | 3.302497 | 0.00095828 | 0.059695566 |
| *LIMD2* | -0.235713 | 0.071399 | -3.301337 | 0.00096225 | 0.059695566 |
| *NFX1* | 0.386971 | 0.117462 | 3.294432 | 0.00098621 | 0.060406824 |
| *LRIF1* | 0.481814 | 0.146412 | 3.290818 | 0.00099897 | 0.060406824 |
| *TMEM243* | 0.245528 | 0.074623 | 3.290241 | 0.00100102 | 0.060406824 |
| *RPS9* | 0.296649 | 0.090295 | 3.285353 | 0.00101855 | 0.060910845 |
| *ZNF564* | -0.9478 | 0.290249 | -3.265477 | 0.0010928 | NA |
| *MSMP* | -0.692931 | 0.212248 | -3.264732 | 0.00109568 | 0.0643658 |
| *CYTH1* | -0.319651 | 0.097911 | -3.264723 | 0.00109571 | 0.0643658 |
| *MME* | -2.329112 | 0.717147 | -3.247748 | 0.00116322 | NA |
| *TPI1* | -0.177085 | 0.054558 | -3.245794 | 0.00117123 | 0.068198679 |
| *SAYSD1* | 0.441649 | 0.136225 | 3.242062 | 0.00118668 | 0.068497346 |
| *ANAPC15* | -0.519052 | 0.160439 | -3.235196 | 0.00121559 | 0.069561277 |
| *TAGLN2* | -0.282729 | 0.087631 | -3.226362 | 0.00125375 | 0.071131305 |
| *PTPMT1* | -0.407196 | 0.126389 | -3.221766 | 0.00127403 | 0.071669699 |
| *SCN3A* | 1.319171 | 0.410129 | 3.216475 | 0.00129776 | NA |
| *ENSG00000259001* | 1.439227 | 0.448179 | 3.211276 | 0.00132147 | NA |
| *DBI* | -0.256013 | 0.079723 | -3.211269 | 0.0013215 | 0.07371532 |
| *NRROS* | -0.546232 | 0.170326 | -3.206977 | 0.00134138 | 0.074200514 |
| *DTX3L* | 0.548116 | 0.171045 | 3.204511 | 0.00135292 | 0.074220598 |
| *KRCC1* | 0.274702 | 0.085806 | 3.201446 | 0.0013674 | 0.074399878 |
| *KIAA0040* | 0.594552 | 0.186143 | 3.194059 | 0.00140287 | 0.075709556 |
| *MRNIP* | 0.362635 | 0.113625 | 3.19151 | 0.00141531 | 0.075764782 |

| | | | | | |
|---|---|---|---|---|---|
| *ZC3H12A* | -0.503614 | 0.158215 | -3.183092 | 0.00145711 | 0.077378534 |
| *DYNLL1* | 0.403626 | 0.126941 | 3.179623 | 0.00147467 | 0.07768936 |
| *DOCK9.DT* | -1.802569 | 0.56736 | -3.177116 | 0.00148748 | NA |
| *FADS1* | -0.571351 | 0.180151 | -3.171514 | 0.00151647 | 0.079262185 |
| *ING3* | -0.339441 | 0.10733 | -3.162604 | 0.00156365 | 0.081089727 |
| *ABTB1* | -0.280605 | 0.088842 | -3.158466 | 0.00158602 | 0.081612388 |
| *ZBTB16* | 1.920903 | 0.608894 | 3.154742 | 0.0016064 | NA |
| *TMEM60* | 0.398212 | 0.126299 | 3.152917 | 0.00161648 | 0.082360607 |
| *IGHG1* | 1.33429 | 0.423433 | 3.15112 | 0.00162646 | 0.082360607 |
| *THAP5* | 0.456929 | 0.145122 | 3.148577 | 0.00164067 | 0.082360607 |
| *TNFAIP8L2* | 0.587545 | 0.186707 | 3.146887 | 0.00165019 | 0.082360607 |
| *MARCHF7* | -0.312542 | 0.099685 | -3.135289 | 0.00171685 | 0.085048152 |
| *SIGLEC5* | 1.883248 | 0.600694 | 3.135122 | 0.00171783 | NA |
| *HHEX* | 0.388525 | 0.124006 | 3.133102 | 0.0017297 | 0.0850498 |
| *TLE1* | -0.793759 | 0.2537 | -3.128734 | 0.00175561 | 0.085643246 |
| *UBE2I* | -0.165457 | 0.052917 | -3.126739 | 0.00176757 | 0.085643246 |
| *SYNGR2* | -0.553756 | 0.177597 | -3.118047 | 0.00182054 | 0.087215452 |
| *LMO4* | 0.438017 | 0.140574 | 3.115929 | 0.00183366 | 0.087215452 |
| *NAGK* | 0.403179 | 0.129431 | 3.115002 | 0.00183943 | 0.087215452 |
| *SAP18* | -0.209103 | 0.067292 | -3.107399 | 0.00188741 | 0.088087234 |
| *CDC42* | -0.282256 | 0.090835 | -3.107333 | 0.00188784 | 0.088087234 |
| *UROD* | -0.426598 | 0.137385 | -3.105126 | 0.00190198 | 0.088087234 |
| *UQCRH* | 0.196568 | 0.063333 | 3.103742 | 0.0019109 | 0.088087234 |
| *KLHDC7B* | -0.903374 | 0.291816 | -3.095695 | 0.00196352 | NA |
| *RPL35A* | 0.189463 | 0.061345 | 3.088475 | 0.00201187 | 0.092101847 |
| *SYNCRIP* | 0.391192 | 0.126746 | 3.086426 | 0.00202579 | 0.09210388 |
| *EVI2A* | 0.503732 | 0.16355 | 3.079991 | 0.00207007 | 0.093476862 |
| *MEGF6* | -1.059687 | 0.344472 | -3.076263 | 0.00209613 | NA |
| *CLEC2B* | 0.417045 | 0.135657 | 3.07426 | 0.00211025 | 0.094336515 |
| *LYRM7* | 0.311791 | 0.101485 | 3.072298 | 0.00212417 | 0.094336515 |
| *MAVS* | 0.403705 | 0.131447 | 3.071237 | 0.00213174 | 0.094336515 |
| *THEMIS2* | 0.971331 | 0.316823 | 3.065851 | 0.00217052 | 0.09541645 |
| *PHKG1* | -0.512084 | 0.167215 | -3.062425 | 0.00219552 | 0.09588051 |
| *NREP* | -0.876796 | 0.286688 | -3.058364 | 0.00222549 | NA |
| *IGHG3* | 1.527124 | 0.499509 | 3.057251 | 0.00223377 | 0.096582783 |
| *MARCHF1* | 0.401624 | 0.131408 | 3.056323 | 0.0022407 | 0.096582783 |
| *AUP1* | -0.243508 | 0.079827 | -3.050461 | 0.0022849 | 0.097295085 |
| *NDUFS8* | -0.248862 | 0.081588 | -3.050246 | 0.00228654 | 0.097295085 |
| *ENSG00000239636* | -0.751415 | 0.246867 | -3.043809 | 0.00233604 | 0.097914563 |
| *HLA.A* | -0.463217 | 0.152185 | -3.043781 | 0.00233625 | 0.097914563 |
| *HIPK2* | 0.943667 | 0.310042 | 3.043672 | 0.0023371 | NA |
| *TTC7A* | -0.5576 | 0.183264 | -3.042612 | 0.00234535 | 0.097914563 |
| *GINS2* | -1.102848 | 0.363504 | -3.033936 | 0.00241386 | NA |
| *ZNF557* | 0.565987 | 0.186713 | 3.031324 | 0.00243484 | 0.10101523 |
| *ENSG00000279483* | -0.285339 | 0.094472 | -3.020356 | 0.00252478 | 0.104096145 |
| *MT.CO1* | 0.222812 | 0.073816 | 3.018474 | 0.00254052 | 0.104098419 |
| *ENSG00000263394* | 0.631845 | 0.209598 | 3.014551 | 0.0025736 | 0.104807086 |

| | | | | | |
|---|---|---|---|---|---|
| *COLCA2* | -1.381221 | 0.458436 | -3.0129 | 0.00258764 | NA |
| *ARL6IP6* | 0.369167 | 0.122548 | 3.012426 | 0.00259169 | 0.104900226 |
| *BLOC1S1* | -0.211451 | 0.070291 | -3.008202 | 0.00262798 | 0.105724474 |
| *CAMTA1* | -0.306899 | 0.102261 | -3.001143 | 0.00268968 | 0.107554905 |
| *ENSG00000275964* | -1.020822 | 0.34018 | -3.000831 | 0.00269244 | NA |
| *BLVRB* | 0.661982 | 0.22106 | 2.994584 | 0.0027482 | 0.108673679 |
| *ACAP1* | 0.369048 | 0.123333 | 2.992299 | 0.00276885 | 0.108673679 |
| *C17orf49* | -0.819006 | 0.273953 | -2.98959 | 0.00279353 | 0.108673679 |
| *ARHGAP30* | 0.274711 | 0.091903 | 2.989135 | 0.00279768 | 0.108673679 |
| *TMEM138* | -0.352863 | 0.118056 | -2.988935 | 0.00279952 | 0.108673679 |
| *PARVB* | 0.646323 | 0.21641 | 2.986565 | 0.0028213 | 0.108882652 |
| *EBLN2* | 0.594972 | 0.199451 | 2.983051 | 0.0028539 | 0.109066659 |
| *ARID5B* | -0.325892 | 0.109268 | -2.982512 | 0.00285893 | 0.109066659 |
| *FXYD7* | 0.816187 | 0.274116 | 2.977521 | 0.0029059 | 0.110224898 |
| *ZNHIT3* | 0.261632 | 0.088185 | 2.966872 | 0.00300846 | 0.111288028 |
| *HNRNPA0* | -0.141821 | 0.047827 | -2.965296 | 0.00302392 | 0.111288028 |
| *PILRB* | 0.694089 | 0.234122 | 2.964648 | 0.00303029 | 0.111288028 |
| *IFIT5* | 0.529993 | 0.178795 | 2.964253 | 0.00303418 | 0.111288028 |
| *RUBCNL* | -0.262173 | 0.088473 | -2.963328 | 0.00304332 | 0.111288028 |
| *TMC6* | -0.339092 | 0.11443 | -2.963306 | 0.00304354 | 0.111288028 |
| *METTL26* | -0.272233 | 0.091892 | -2.962524 | 0.00305128 | 0.111288028 |
| *RPS5* | 0.229804 | 0.077673 | 2.958613 | 0.00309027 | 0.112093934 |
| *ENSG00000271204* | 0.524355 | 0.17746 | 2.954779 | 0.00312893 | 0.112879456 |
| *SNRPA1* | -0.287459 | 0.097347 | -2.952928 | 0.00314776 | 0.112944981 |
| *PPP1R16A* | -0.673571 | 0.228767 | -2.94435 | 0.00323634 | NA |
| *MAPK14* | 0.601882 | 0.204535 | 2.942679 | 0.00325386 | 0.116095692 |
| *HRK* | -1.430781 | 0.486479 | -2.941094 | 0.00327055 | 0.116095692 |
| *FIS1* | -0.29289 | 0.099731 | -2.936808 | 0.00331609 | 0.116279999 |
| *SP140* | 0.287727 | 0.097986 | 2.936405 | 0.0033204 | 0.116279999 |
| *ST6GALNAC6* | -0.355432 | 0.121074 | -2.935669 | 0.00332829 | 0.116279999 |
| *CERS4* | -0.324136 | 0.110501 | -2.93333 | 0.00335347 | 0.116546196 |
| *DDB2* | -0.517366 | 0.176624 | -2.929196 | 0.0033984 | 0.117492558 |
| *PTPRA* | -0.788848 | 0.26933 | -2.928922 | 0.0034014 | NA |
| *GLRX2* | 0.512648 | 0.175477 | 2.921449 | 0.00348408 | 0.119830523 |
| *NBPF9* | 0.876661 | 0.300284 | 2.919444 | 0.00350656 | NA |
| *RPL19* | 0.142523 | 0.048872 | 2.916232 | 0.00354287 | 0.121160197 |
| *PHF23* | 0.395438 | 0.135666 | 2.914793 | 0.00355924 | 0.121160197 |
| *RTL10* | 0.732556 | 0.251731 | 2.91007 | 0.00361348 | NA |
| *LIMD1* | -0.907512 | 0.312354 | -2.9054 | 0.00366784 | 0.12386072 |
| *SRSF6* | -0.342588 | 0.117942 | -2.904714 | 0.00367589 | 0.12386072 |
| *DNAAF2* | 0.459293 | 0.158231 | 2.902675 | 0.0036999 | 0.12404008 |
| *SAP25* | 0.730464 | 0.252072 | 2.897844 | 0.00375738 | NA |
| *RACK1* | 0.159888 | 0.055175 | 2.897813 | 0.00375774 | 0.125346221 |
| *ENSG00000266844* | -1.715952 | 0.592891 | -2.89421 | 0.00380114 | NA |
| *ZNF426* | 0.407027 | 0.140649 | 2.893927 | 0.00380457 | 0.125763709 |
| *CFL1* | -0.215241 | 0.074384 | -2.893631 | 0.00380815 | 0.125763709 |
| *SLC2A14* | -3.605707 | 1.248446 | -2.888156 | 0.00387507 | NA |

| | | | | | |
|---|---|---|---|---|---|
| *ACTN1* | 2.844679 | 0.985118 | 2.887652 | 0.00388129 | NA |
| *GPS2* | -0.367503 | 0.127399 | -2.884668 | 0.00391826 | 0.128672843 |
| *ENSG00000256092* | -0.550564 | 0.190948 | -2.883325 | 0.00393501 | 0.128672843 |
| *C6orf203* | 0.499207 | 0.17331 | 2.880423 | 0.00397143 | 0.129227039 |
| *UXT* | 0.209451 | 0.072955 | 2.87096 | 0.00409228 | 0.132509952 |
| *JUND* | -0.31461 | 0.109662 | -2.868914 | 0.00411884 | 0.132722622 |
| *ENSG00000254614* | -0.76304 | 0.266145 | -2.867012 | 0.00414367 | NA |
| *ADA* | -0.602061 | 0.210216 | -2.864013 | 0.00418311 | 0.133465057 |
| *SLC5A3* | -0.418507 | 0.146194 | -2.862684 | 0.0042007 | 0.133465057 |
| *MAML3* | -1.107093 | 0.386748 | -2.86257 | 0.0042022 | 0.133465057 |
| *PEX10* | -0.586348 | 0.204967 | -2.860699 | 0.00422708 | 0.133615873 |
| *THEM6* | 0.789358 | 0.276362 | 2.856247 | 0.00428682 | NA |
| *EPB41L2* | -0.590689 | 0.206834 | -2.855857 | 0.00429208 | 0.135027707 |
| *ENSG00000089127* | 0.482521 | 0.169054 | 2.854244 | 0.00431394 | 0.135075254 |
| *RPL38* | 0.169596 | 0.059546 | 2.848156 | 0.00439734 | 0.13704021 |
| *SCAND1* | -0.223233 | 0.07842 | -2.846619 | 0.00441863 | 0.13706001 |
| *MED23* | -0.578377 | 0.203351 | -2.844227 | 0.00445193 | 0.137450794 |
| *SLC38A2* | 0.364225 | 0.128349 | 2.837764 | 0.00454307 | 0.139506764 |
| *EIF3E* | 0.208432 | 0.073481 | 2.836534 | 0.00456062 | 0.139506764 |
| *CAB39L* | -1.175181 | 0.414606 | -2.834452 | 0.00459043 | NA |
| *ASPSCR1* | -0.48013 | 0.169424 | -2.833898 | 0.0045984 | 0.139506764 |
| *MUC20.OT1* | -0.372565 | 0.131481 | -2.833607 | 0.00460259 | 0.139506764 |
| *DRG2* | -0.420148 | 0.148409 | -2.831014 | 0.00464006 | 0.139856574 |
| *PDE4B* | 0.25732 | 0.090929 | 2.829899 | 0.00465627 | 0.139856574 |
| *ATP6V1C1* | 0.453115 | 0.16038 | 2.825252 | 0.00472435 | 0.14126237 |
| *CHM* | 0.631983 | 0.223919 | 2.822373 | 0.00476698 | 0.141897675 |
| *MYCBP2* | 0.241982 | 0.085831 | 2.819274 | 0.00481324 | 0.142635222 |
| *NAE1* | -0.365645 | 0.129831 | -2.816309 | 0.00485789 | 0.143318471 |
| *PTPN20* | -2.618712 | 0.930067 | -2.815619 | 0.00486834 | NA |
| *VAV3.AS1* | -1.212294 | 0.430636 | -2.815124 | 0.00487584 | NA |
| *FAM111A.DT* | 0.526128 | 0.187411 | 2.807351 | 0.00499508 | 0.146388181 |
| *GM2A* | 0.828694 | 0.295388 | 2.805438 | 0.00502483 | 0.146388181 |
| *SNHG30* | -0.374087 | 0.133353 | -2.805228 | 0.0050281 | 0.146388181 |
| *HLA.C* | -0.473834 | 0.169468 | -2.796013 | 0.00517373 | 0.14940857 |
| *SPATC1L* | -0.576267 | 0.206118 | -2.795818 | 0.00517686 | 0.14940857 |
| *MGST2* | -0.93523 | 0.334741 | -2.79389 | 0.00520782 | NA |
| *SNHG32* | 0.217864 | 0.078053 | 2.791226 | 0.00525088 | 0.150384931 |
| *ST13* | 0.137263 | 0.049182 | 2.790911 | 0.005256 | 0.150384931 |
| *FAM184B* | -1.703321 | 0.610989 | -2.78781 | 0.00530657 | 0.151180355 |
| *TOMM20* | 0.231839 | 0.083218 | 2.785939 | 0.00533728 | 0.151405511 |
| *PDE6A* | -1.360102 | 0.488534 | -2.784048 | 0.0053685 | NA |
| *NKIRAS2* | -0.335691 | 0.120585 | -2.783859 | 0.00537164 | 0.151731799 |
| *CLECL1* | 1.392572 | 0.500736 | 2.781049 | 0.00541835 | NA |
| *HSPD1* | 0.293641 | 0.105789 | 2.775714 | 0.00550807 | 0.154926122 |
| *CREG1* | 0.891341 | 0.321585 | 2.77171 | 0.00557627 | NA |
| *EIF5* | 0.220205 | 0.079486 | 2.770369 | 0.00559928 | 0.156827039 |
| *KCP* | -2.303337 | 0.83183 | -2.768998 | 0.00562289 | NA |

| | | | | | |
|---|---|---|---|---|---|
| *ASF1A* | 0.446854 | 0.161519 | 2.766574 | 0.00566487 | 0.157997421 |
| *MRPL35* | 0.465375 | 0.168536 | 2.761274 | 0.00575764 | 0.159383877 |
| *TIAL1* | 0.27525 | 0.099729 | 2.759986 | 0.00578038 | 0.159383877 |
| *ICA1L* | -0.456645 | 0.165473 | -2.759634 | 0.00578661 | 0.159383877 |
| *PRMT7* | 0.552167 | 0.200281 | 2.756961 | 0.00583413 | 0.160028659 |
| *ENSG00000225342* | -0.695043 | 0.252229 | -2.755598 | 0.0058585 | NA |
| *IL27RA* | 0.419602 | 0.152427 | 2.752797 | 0.00590885 | 0.160755563 |
| *PRPF38A* | 0.256866 | 0.093316 | 2.752661 | 0.00591131 | 0.160755563 |
| *CDCA4* | 0.495646 | 0.180253 | 2.749724 | 0.00596455 | 0.160755563 |
| *CFAP410* | -0.346298 | 0.125973 | -2.74899 | 0.00597792 | 0.160755563 |
| *MTHFR* | 0.465091 | 0.169199 | 2.748782 | 0.00598172 | 0.160755563 |
| *INSR* | -1.042519 | 0.379873 | -2.744386 | 0.00606242 | NA |
| *GPS1* | -0.391351 | 0.143236 | -2.732221 | 0.00629089 | 0.16838263 |
| *LINC00926* | -0.217827 | 0.079778 | -2.730403 | 0.00632569 | 0.168634229 |
| *ENSG00000224746* | -1.240809 | 0.455704 | -2.722839 | 0.00647235 | NA |
| *ISG20L2* | 0.346455 | 0.127337 | 2.72077 | 0.006513 | 0.172933276 |
| *GATM* | 0.932994 | 0.343118 | 2.719162 | 0.00654475 | NA |
| *KLK1* | 1.288407 | 0.474082 | 2.717687 | 0.00657399 | NA |
| *MRPL17* | 0.464257 | 0.170914 | 2.716324 | 0.00660112 | 0.174574687 |
| *COL9A3* | -1.250024 | 0.460569 | -2.714083 | 0.00664595 | 0.175062854 |
| *TNFAIP1* | 0.677419 | 0.249686 | 2.713089 | 0.00666591 | NA |
| *APOOL* | 0.630127 | 0.232283 | 2.712748 | 0.00667277 | NA |
| *SUSD3* | 0.359441 | 0.132563 | 2.711466 | 0.00669865 | 0.175142349 |
| *ORMDL3* | -0.407972 | 0.150504 | -2.710702 | 0.0067141 | 0.175142349 |
| *RPS28* | 0.176311 | 0.065059 | 2.71001 | 0.00672813 | 0.175142349 |
| *CHL1* | 1.830075 | 0.675749 | 2.708216 | 0.0067646 | NA |
| *SLC23A3* | 1.006803 | 0.372102 | 2.705719 | 0.00681566 | NA |
| *ENSG00000267002* | 0.337488 | 0.124895 | 2.702178 | 0.00688869 | 0.177541415 |
| *POLR2M* | -0.320334 | 0.118553 | -2.702019 | 0.00689198 | 0.177541415 |
| *LTBP3* | 0.335759 | 0.124281 | 2.701607 | 0.00690053 | 0.177541415 |
| *IGLV6.57* | -0.994622 | 0.368324 | -2.700401 | 0.0069256 | NA |
| *FKBP5* | 1.031473 | 0.382043 | 2.69989 | 0.00693625 | NA |
| *FNIP1* | -0.40962 | 0.151736 | -2.699558 | 0.00694317 | 0.177948853 |
| *ENSG00000274265* | -0.276371 | 0.102437 | -2.69797 | 0.00697637 | 0.178111979 |
| *IFI44* | 1.015015 | 0.37812 | 2.68437 | 0.00726666 | NA |
| *LHFPL4* | 1.162362 | 0.433869 | 2.679063 | 0.00738286 | NA |
| *RAB9A* | 0.457246 | 0.170753 | 2.677822 | 0.00741027 | 0.18844412 |
| *MAP3K9* | 0.762881 | 0.284943 | 2.677316 | 0.00742146 | NA |
| *CCDC66* | 0.386815 | 0.144519 | 2.676577 | 0.00743784 | 0.18844412 |
| *NAB2* | 0.847115 | 0.317023 | 2.672096 | 0.00753792 | NA |
| *GRAP2* | 1.087227 | 0.407108 | 2.670612 | 0.00757132 | NA |
| *TNFAIP8* | 0.320975 | 0.120221 | 2.669878 | 0.00758788 | 0.191514586 |
| *LILRB1* | 0.467936 | 0.175652 | 2.663989 | 0.00772201 | 0.194161632 |
| *SERP1* | 0.14527 | 0.054579 | 2.661655 | 0.00777574 | 0.194775026 |
| *CAV1* | -1.120652 | 0.421164 | -2.660842 | 0.00779455 | NA |
| *LDHA* | -0.343209 | 0.129074 | -2.659009 | 0.00783708 | 0.19557356 |
| *CD180* | 0.42468 | 0.159837 | 2.656963 | 0.00788481 | 0.196027543 |

| | | | | | |
|---|---|---|---|---|---|
| *NPM1* | 0.183985 | 0.069302 | 2.654816 | 0.00793516 | 0.196416569 |
| *RNASEK* | -0.416085 | 0.15679 | -2.653777 | 0.00795964 | 0.196416569 |
| *RING1* | -0.328167 | 0.123832 | -2.650103 | 0.00804673 | 0.1978303 |
| *POLE3* | -0.256062 | 0.096679 | -2.64857 | 0.0080833 | 0.197996135 |
| *LETMD1* | 0.284733 | 0.107587 | 2.646539 | 0.00813202 | 0.198457271 |
| *DHPS* | 0.223618 | 0.084591 | 2.643519 | 0.00820492 | 0.199022222 |
| *TRIM69* | 0.435499 | 0.164787 | 2.642804 | 0.00822225 | 0.199022222 |
| *PFKP* | -0.942396 | 0.356716 | -2.641864 | 0.00824512 | 0.199022222 |
| *IFIT3* | 0.711901 | 0.269514 | 2.64142 | 0.00825592 | NA |
| *LY9* | 0.449578 | 0.170317 | 2.639656 | 0.00829903 | 0.19918574 |
| *ENSG00000253948* | -0.793616 | 0.300801 | -2.638342 | 0.00833126 | NA |
| *COL19A1* | -0.483302 | 0.183213 | -2.637916 | 0.00834173 | 0.19918574 |
| *RSBN1L* | 0.242608 | 0.09197 | 2.637908 | 0.00834192 | 0.19918574 |
| *NEIL1* | -0.362759 | 0.137666 | -2.63506 | 0.00841224 | 0.200144919 |
| *NSMCE1* | -0.267368 | 0.101546 | -2.632964 | 0.00846434 | 0.200312071 |
| *ARF5* | -0.300007 | 0.113974 | -2.632244 | 0.0084823 | 0.200312071 |
| *C15orf61* | 0.429294 | 0.163158 | 2.631144 | 0.00850979 | 0.200312071 |
| *WIPF2* | 0.371925 | 0.141549 | 2.627526 | 0.00860082 | 0.201574678 |
| *RNF144B* | 0.767639 | 0.29219 | 2.627193 | 0.00860925 | NA |
| *RHOC* | -0.67186 | 0.25579 | -2.626604 | 0.00862417 | 0.201574678 |
| *RP9* | 0.51712 | 0.197189 | 2.622459 | 0.00872977 | 0.20332695 |
| *PODXL2* | -1.428765 | 0.54505 | -2.621347 | 0.00875831 | NA |
| *OTUD1* | 0.666187 | 0.254198 | 2.620737 | 0.00877399 | NA |
| *HMG20B* | -0.252825 | 0.096491 | -2.620206 | 0.00878768 | 0.203960128 |
| *PPP4R2* | 0.31663 | 0.121068 | 2.615314 | 0.00891456 | 0.206184049 |
| *NDUFB5* | 0.222033 | 0.08504 | 2.610924 | 0.0090298 | 0.20812444 |
| *XKR6* | 1.134278 | 0.434976 | 2.607677 | 0.00911589 | NA |
| *ENSG00000272994* | -0.402656 | 0.154419 | -2.607552 | 0.00911923 | 0.208973808 |
| *HCK* | -0.630644 | 0.241889 | -2.607162 | 0.00912962 | 0.208973808 |
| *PSMG3* | -0.401087 | 0.154017 | -2.604174 | 0.00920961 | 0.209697096 |
| *PHF20* | 0.214318 | 0.082315 | 2.603624 | 0.0092244 | 0.209697096 |
| *BSG* | -0.209967 | 0.08075 | -2.60022 | 0.00931639 | 0.210434413 |
| *ZBTB25* | 0.453066 | 0.174251 | 2.600079 | 0.00932024 | 0.210434413 |
| *POU2F2* | 0.275913 | 0.106266 | 2.596436 | 0.00941965 | 0.211958171 |
| *CCDC120* | 0.982708 | 0.378604 | 2.595608 | 0.00944237 | NA |
| *MORC3* | 0.341039 | 0.131579 | 2.5919 | 0.00954476 | 0.213837884 |
| *DDX6* | -0.283449 | 0.109394 | -2.591077 | 0.00956762 | 0.213837884 |
| *NAA35* | 0.580176 | 0.223926 | 2.590935 | 0.00957157 | NA |
| *XPO4* | -0.420877 | 0.162759 | -2.585887 | 0.00971287 | 0.21613234 |
| *ASNSD1* | 0.309805 | 0.119843 | 2.585089 | 0.0097354 | 0.21613234 |
| *USP49* | -0.866485 | 0.336194 | -2.577338 | 0.00995646 | NA |
| *NFE2L2* | 0.279894 | 0.108661 | 2.575853 | 0.00999933 | 0.221251798 |
| *NACA* | 0.142626 | 0.055409 | 2.574069 | 0.01005104 | 0.221657066 |
| *LINC01250* | -2.722735 | 1.057815 | -2.573925 | 0.01005522 | NA |
| *MRPL28* | -0.241128 | 0.093748 | -2.572088 | 0.01010873 | 0.222191214 |
| *MINPP1* | 0.721592 | 0.280701 | 2.570672 | 0.01015013 | NA |
| *PCMTD2* | -0.366568 | 0.142913 | -2.564976 | 0.01031831 | 0.225225992 |

| | | | | | |
|---|---|---|---|---|---|
| *LINC01215* | -0.311626 | 0.121517 | -2.564463 | 0.01033357 | 0.225225992 |
| *MTPN* | 0.221086 | 0.086229 | 2.563959 | 0.01034859 | 0.225225992 |
| *MGAT3* | 1.228248 | 0.479077 | 2.563777 | 0.01035402 | NA |
| *LILRA4* | 1.941654 | 0.757507 | 2.563216 | 0.01037076 | NA |
| *SLC25A45* | 0.567909 | 0.221708 | 2.561521 | 0.01042149 | 0.225868897 |
| *SKAP2* | 0.245428 | 0.095844 | 2.560699 | 0.01044618 | 0.225868897 |
| *VMAC* | -0.558851 | 0.218247 | -2.560639 | 0.01044798 | NA |
| *GALNT2* | -0.537012 | 0.209981 | -2.557435 | 0.01054472 | 0.226632175 |
| *CCDC191* | -0.757304 | 0.296246 | -2.556336 | 0.01057809 | 0.226632175 |
| *CGGBP1* | -0.244446 | 0.095631 | -2.556145 | 0.01058391 | 0.226632175 |
| *ENSG00000261087* | 0.868756 | 0.340427 | 2.55196 | 0.01071187 | NA |
| *FGFR1OP2* | 0.283897 | 0.111335 | 2.549929 | 0.01077448 | 0.22997095 |
| *ALOX5* | -0.274426 | 0.107723 | -2.547506 | 0.0108496 | 0.230832149 |
| *CHPF2* | -0.543453 | 0.213641 | -2.543766 | 0.01096644 | 0.232572694 |
| *GNG11* | 1.179621 | 0.463731 | 2.543763 | 0.01096654 | NA |
| *SDHAF4* | 0.957667 | 0.376538 | 2.543345 | 0.01097968 | NA |
| *REST* | 0.282099 | 0.110976 | 2.541996 | 0.01102216 | 0.232610752 |
| *AGTRAP* | -0.459464 | 0.180786 | -2.541483 | 0.01103832 | 0.232610752 |
| *RNF43* | -0.739766 | 0.291237 | -2.540087 | 0.01108249 | NA |
| *ZEB1* | -0.310664 | 0.122449 | -2.537096 | 0.01117765 | 0.234131411 |
| *GTF2E2* | 0.344685 | 0.135864 | 2.53699 | 0.01118103 | 0.234131411 |
| *ENSG00000278834* | -1.030856 | 0.407344 | -2.530677 | 0.01138426 | NA |
| *RPL9* | 0.155734 | 0.061601 | 2.528113 | 0.01146774 | 0.238632913 |
| *TUT7* | 0.325736 | 0.128846 | 2.528108 | 0.0114679 | 0.238632913 |
| *CHTF18* | -0.70264 | 0.278506 | -2.522887 | 0.0116396 | NA |
| *RHBDD2* | -0.394263 | 0.15628 | -2.522792 | 0.01164273 | 0.240353431 |
| *IL13RA1* | -0.85679 | 0.339659 | -2.522503 | 0.0116523 | 0.240353431 |
| *SORL1* | -0.493487 | 0.19565 | -2.522294 | 0.01165921 | 0.240353431 |
| *JUNB* | -0.29076 | 0.115477 | -2.517901 | 0.01180566 | 0.242462605 |
| *GSDMD* | -0.346503 | 0.137678 | -2.516761 | 0.01184392 | 0.242462605 |
| *ENSG00000166685* | -0.461943 | 0.183606 | -2.515953 | 0.0118711 | 0.242462605 |
| *SNAP23* | -0.21855 | 0.086907 | -2.514759 | 0.01191138 | 0.242536696 |
| *MRTO4* | -0.387307 | 0.154148 | -2.512571 | 0.0119855 | 0.242536696 |
| *SPESP1* | 1.527076 | 0.607803 | 2.512455 | 0.01198945 | NA |
| *MREG* | -0.863258 | 0.343685 | -2.51177 | 0.01201273 | NA |
| *ENSG00000258757* | 0.584023 | 0.232524 | 2.511662 | 0.01201643 | 0.242536696 |
| *MED11* | 0.242545 | 0.096573 | 2.511531 | 0.01202088 | 0.242536696 |
| *FHL3* | -0.786236 | 0.313554 | -2.507498 | 0.01215892 | NA |
| *TAF9* | -0.285624 | 0.113952 | -2.506515 | 0.01219277 | 0.245259486 |
| *MINDY2* | 0.381266 | 0.152302 | 2.503346 | 0.01230251 | 0.246080967 |
| *CD69* | -0.390544 | 0.156018 | -2.503195 | 0.01230776 | 0.246080967 |
| *UGT8* | 0.738404 | 0.29499 | 2.503148 | 0.0123094 | NA |
| *PPARD* | -0.493075 | 0.197322 | -2.498832 | 0.01246033 | 0.248383381 |
| *BCAS4* | 0.36509 | 0.146208 | 2.49706 | 0.01252277 | 0.248636132 |
| *MRPS14* | 0.251776 | 0.100886 | 2.495656 | 0.01257244 | 0.248636132 |
| *RPSA* | 0.23382 | 0.093705 | 2.495291 | 0.01258538 | 0.248636132 |
| *OPTN* | -0.753233 | 0.302442 | -2.490501 | 0.01275632 | 0.250413166 |

| | | | | | |
|---|---|---|---|---|---|
| *RPS15A* | 0.172929 | 0.069436 | 2.490485 | 0.0127569 | 0.250413166 |
| *CD40* | -0.206197 | 0.082823 | -2.489605 | 0.0127885 | 0.250413166 |
| *SLC2A1* | -0.313428 | 0.126122 | -2.485116 | 0.01295093 | 0.252847765 |
| *EMC9* | -0.486767 | 0.195902 | -2.484743 | 0.0129645 | NA |
| *IFIT1* | 2.018428 | 0.81241 | 2.484492 | 0.01297363 | NA |
| *RAB30* | 0.212917 | 0.085742 | 2.483242 | 0.01301927 | 0.253409359 |
| *ENSG00000237773* | -0.913858 | 0.368054 | -2.482946 | 0.01303007 | NA |
| *FASTK* | -0.245613 | 0.098948 | -2.482237 | 0.01305604 | 0.253409359 |
| *FLYWCH1* | 0.54623 | 0.220291 | 2.479587 | 0.01315346 | NA |
| *DOCK8* | 0.340323 | 0.137279 | 2.479058 | 0.013173 | 0.254933974 |
| *USF1* | -0.667133 | 0.269478 | -2.475652 | 0.01329932 | NA |
| *CFAP73* | -1.047877 | 0.42334 | -2.475259 | 0.01331396 | NA |
| *GCNT2* | 0.684373 | 0.276925 | 2.471335 | 0.01346097 | 0.259539909 |
| *CETN3* | 0.439238 | 0.177838 | 2.46988 | 0.01351585 | 0.259539909 |
| *JAM3* | 0.718412 | 0.290901 | 2.469608 | 0.01352612 | NA |
| *RANGRF* | -0.280088 | 0.113417 | -2.46955 | 0.01352829 | 0.259539909 |
| *GGCX* | -0.499102 | 0.202299 | -2.467147 | 0.01361943 | 0.260535326 |
| *HSPA8* | 0.245918 | 0.099747 | 2.465403 | 0.01368593 | 0.261055248 |
| *ADHFE1* | 1.471291 | 0.597106 | 2.464038 | 0.01373816 | NA |
| *RND1* | -0.884731 | 0.359115 | -2.463639 | 0.01375344 | NA |
| *ATP6V1D* | 0.457626 | 0.185789 | 2.463147 | 0.01377234 | 0.261333908 |
| *POLR3K* | 0.2964 | 0.120372 | 2.462377 | 0.01380194 | 0.261333908 |
| *RPL24* | 0.160687 | 0.065276 | 2.461667 | 0.01382931 | 0.261333908 |
| *GCNT1* | 0.837723 | 0.34057 | 2.459765 | 0.0139028 | NA |
| *C2orf69* | 0.385022 | 0.156529 | 2.459749 | 0.01390342 | 0.261333908 |
| *ARPC3* | -0.156184 | 0.063503 | -2.459456 | 0.01391476 | 0.261333908 |
| *ACADVL* | -0.212839 | 0.086559 | -2.458889 | 0.01393676 | 0.261333908 |
| *RAB33A* | 1.575588 | 0.641604 | 2.455702 | 0.01406098 | NA |
| *TRMT13* | 0.290659 | 0.118365 | 2.455618 | 0.01406424 | 0.262981451 |
| *RNF181* | -0.19372 | 0.078929 | -2.454359 | 0.01411362 | 0.263163409 |
| *ABHD12* | -0.601078 | 0.245126 | -2.452114 | 0.01420197 | NA |
| *ZNF860* | 0.492524 | 0.200894 | 2.451654 | 0.01422012 | NA |
| *PURA* | 0.356434 | 0.145464 | 2.450323 | 0.01427282 | 0.264829124 |
| *TTC19* | -0.26316 | 0.107409 | -2.450073 | 0.01428274 | 0.264829124 |
| *ATP5F1B* | -0.230273 | 0.094166 | -2.445401 | 0.01446911 | 0.266398949 |
| *LINC00662* | 0.296198 | 0.121131 | 2.445274 | 0.01447422 | 0.266398949 |
| *SLC25A37* | 0.330138 | 0.135029 | 2.444936 | 0.0144878 | 0.266398949 |
| *ENSG00000271646* | -0.577288 | 0.236195 | -2.444113 | 0.01452089 | NA |
| *RPRML* | -1.961666 | 0.806212 | -2.43319 | 0.01496646 | NA |
| *VPS26B* | -0.807113 | 0.332356 | -2.428458 | 0.01516319 | NA |
| *DPY19L3* | -0.754895 | 0.311465 | -2.423687 | 0.01536385 | NA |
| *ENSG00000223478* | -0.58899 | 0.243223 | -2.421605 | 0.01545212 | NA |
| *ZNF260* | 0.473219 | 0.195753 | 2.417423 | 0.01563084 | NA |
| *SYNJ2BP* | 0.277652 | 0.114862 | 2.417265 | 0.01563763 | 0.2863145 |
| *SIDT1.AS1* | -0.621963 | 0.257349 | -2.416811 | 0.01565715 | 0.2863145 |
| *ENSG00000235501* | -1.158859 | 0.479514 | -2.416736 | 0.01566038 | NA |
| *LAPTM5* | -0.20122 | 0.083305 | -2.415463 | 0.0157152 | 0.286586553 |

| | | | | | |
|---|---|---|---|---|---|
| *BAX* | -0.312829 | 0.129653 | -2.41281 | 0.01583007 | 0.286993728 |
| *PIGS* | 0.441336 | 0.18297 | 2.412073 | 0.01586213 | 0.286993728 |
| *HIRA* | -0.552858 | 0.229216 | -2.411955 | 0.01586723 | 0.286993728 |
| *CDK7* | 0.554761 | 0.230136 | 2.410577 | 0.01592732 | NA |
| *ALAS1* | 0.518025 | 0.214938 | 2.410115 | 0.0159475 | NA |
| *ENSG00000204802* | -1.014382 | 0.420999 | -2.409464 | 0.01597597 | NA |
| *PURB* | 0.450359 | 0.187004 | 2.408288 | 0.01602752 | 0.287192474 |
| *ERH* | -0.218321 | 0.090664 | -2.408018 | 0.0160394 | 0.287192474 |
| *LPAR5* | 0.469961 | 0.195167 | 2.40799 | 0.01604061 | 0.287192474 |
| *CLMN* | 0.645887 | 0.268253 | 2.407747 | 0.01605128 | 0.287192474 |
| *CD5* | -0.85502 | 0.355695 | -2.403802 | 0.01622557 | NA |
| *PYGO2* | 0.371257 | 0.154463 | 2.403534 | 0.01623744 | 0.28951641 |
| *KMT2A* | 0.211491 | 0.088017 | 2.402838 | 0.0162684 | 0.28951641 |
| *RHOBTB3* | 1.759852 | 0.732812 | 2.401503 | 0.01632786 | NA |
| *HIST1H2BN* | 0.871351 | 0.362849 | 2.401413 | 0.0163319 | NA |
| *ARL11* | 0.626704 | 0.261087 | 2.400364 | 0.01637878 | 0.290701388 |
| *PIAS2* | -0.45379 | 0.189358 | -2.396465 | 0.01655406 | 0.292553132 |
| *YDJC* | -0.219516 | 0.091614 | -2.396085 | 0.01657125 | 0.292553132 |
| *IRF5* | 0.613587 | 0.256123 | 2.395673 | 0.01658988 | NA |
| *FANCL* | 0.589617 | 0.246167 | 2.395195 | 0.01661153 | NA |
| *MORF4L2* | 0.254457 | 0.106241 | 2.395107 | 0.01661551 | 0.292556413 |
| *NINJ1* | -0.882172 | 0.368325 | -2.39509 | 0.01661627 | NA |
| *DNAJB4* | 0.613456 | 0.256273 | 2.39376 | 0.01667664 | NA |
| *AKAP12* | -1.938961 | 0.810069 | -2.393575 | 0.01668505 | NA |
| *CCDC51* | 0.603294 | 0.252257 | 2.391581 | 0.01677598 | NA |
| *FBXL6* | -1.259591 | 0.526687 | -2.391537 | 0.01677797 | NA |
| *APBB2* | -0.739353 | 0.309188 | -2.391271 | 0.01679015 | 0.294849165 |
| *TRMT112* | 0.206043 | 0.086248 | 2.388959 | 0.01689621 | 0.295350411 |
| *ITGB7* | 0.524978 | 0.219775 | 2.388709 | 0.01690768 | 0.295350411 |
| *ZNF827* | 0.734557 | 0.307704 | 2.387221 | 0.01697629 | NA |
| *TMX2* | 0.268946 | 0.112667 | 2.387096 | 0.01698206 | 0.29587112 |
| *ENSG00000258017* | -1.234596 | 0.517453 | -2.385908 | 0.01703701 | NA |
| *SLC15A2* | 0.833893 | 0.349513 | 2.385873 | 0.01703866 | NA |
| *GPR137* | -0.355168 | 0.148973 | -2.384118 | 0.01712012 | 0.297433914 |
| *TIPARP* | -0.414776 | 0.174064 | -2.382894 | 0.01717712 | 0.297433914 |
| *TLR7* | 0.77367 | 0.324741 | 2.382419 | 0.0171993 | NA |
| *COA5* | 0.269944 | 0.113314 | 2.382272 | 0.01720618 | 0.297433914 |
| *CYB5RL* | 0.956372 | 0.401465 | 2.382203 | 0.01720942 | NA |
| *INKA1* | 0.419128 | 0.176217 | 2.378472 | 0.01738456 | 0.299100679 |
| *MCM7* | -0.353132 | 0.148481 | -2.378299 | 0.01739272 | 0.299100679 |
| *DDX5* | -0.290524 | 0.122282 | -2.37585 | 0.01750856 | 0.300314765 |
| *ARL4D* | 1.062334 | 0.447179 | 2.375636 | 0.01751874 | NA |
| *VCPIP1* | 0.390673 | 0.164581 | 2.373742 | 0.01760887 | 0.300501029 |
| *RPL5* | 0.167745 | 0.070681 | 2.373289 | 0.01763047 | 0.300501029 |
| *RPL37* | 0.129717 | 0.054669 | 2.372771 | 0.01765523 | 0.300501029 |
| *ECHDC2* | 1.158543 | 0.488589 | 2.371201 | 0.01773038 | NA |
| *INTS13* | 0.505084 | 0.213072 | 2.370486 | 0.01776472 | NA |

| | | | | | |
|---|---|---|---|---|---|
| *SWAP70* | 0.245541 | 0.103603 | 2.370011 | 0.01778756 | 0.300710814 |
| *EMC8* | 0.349106 | 0.147329 | 2.369565 | 0.01780901 | 0.300710814 |
| *ATP5IF1* | -0.166884 | 0.07047 | -2.368145 | 0.01787753 | 0.300710814 |
| *TNFRSF13B* | 1.080443 | 0.456265 | 2.368018 | 0.01788368 | 0.300710814 |
| *PPP1CA* | -0.16852 | 0.071171 | -2.367803 | 0.01789406 | 0.300710814 |
| *RPS6KB2.AS1* | -0.910507 | 0.385323 | -2.362971 | 0.01812907 | NA |
| *MAD2L2* | -0.289822 | 0.122717 | -2.361706 | 0.01819105 | 0.303966517 |
| *RENBP* | 0.596236 | 0.252622 | 2.360189 | 0.01826561 | 0.303966517 |
| *RNF19B* | 0.949342 | 0.40234 | 2.359554 | 0.01829693 | NA |
| *LMBRD1* | -0.258254 | 0.109461 | -2.359312 | 0.01830888 | 0.303966517 |
| *BCAP31* | -0.255473 | 0.108287 | -2.359223 | 0.01831324 | 0.303966517 |
| *PEX13* | 0.46454 | 0.19691 | 2.359152 | 0.01831675 | 0.303966517 |
| *FAXDC2* | 1.519125 | 0.644405 | 2.357407 | 0.01840309 | NA |
| *ENSG00000272630* | 1.104355 | 0.468514 | 2.357147 | 0.01841598 | NA |
| *POLD3* | -0.528557 | 0.224274 | -2.35675 | 0.01843568 | NA |
| *SLC35D1* | 0.741048 | 0.314585 | 2.355635 | 0.01849109 | NA |
| *PPOX* | -0.412682 | 0.175212 | -2.355335 | 0.01850599 | 0.306341118 |
| *SKI* | 0.576502 | 0.244936 | 2.353684 | 0.01858841 | NA |
| *ZNF667.AS1* | 0.491333 | 0.208761 | 2.353566 | 0.0185943 | 0.307037184 |
| *SAMHD1* | -0.95498 | 0.406065 | -2.351792 | 0.01868321 | NA |
| *BATF* | 1.120456 | 0.476749 | 2.350202 | 0.01876324 | NA |
| *ARHGEF3* | -0.447753 | 0.190548 | -2.34981 | 0.01878302 | 0.309383806 |
| *CCDC157* | -0.892233 | 0.379859 | -2.348852 | 0.01883141 | NA |
| *SNHG25* | -0.336021 | 0.143075 | -2.348564 | 0.01884595 | 0.30965201 |
| *NPHP3* | 0.442287 | 0.188426 | 2.347279 | 0.01891109 | 0.30995506 |
| *NDUFB3* | 0.194162 | 0.082779 | 2.345554 | 0.01899882 | 0.310625941 |
| *CCDC144A* | 1.423199 | 0.607817 | 2.341493 | 0.01920676 | NA |
| *CRIP3* | 0.698797 | 0.298453 | 2.341401 | 0.01921151 | NA |
| *HELZ* | 0.358712 | 0.153213 | 2.341263 | 0.01921864 | 0.313447925 |
| *KLC4* | 0.482429 | 0.206175 | 2.339902 | 0.01928879 | NA |
| *BIN1* | 0.260565 | 0.111393 | 2.339141 | 0.01932814 | 0.313701161 |
| *MHENCR* | -0.208641 | 0.089196 | -2.339131 | 0.01932868 | 0.313701161 |
| *IGHV3.30* | 1.295754 | 0.554235 | 2.337916 | 0.01939159 | NA |
| *HNRNPDL* | -0.177524 | 0.076004 | -2.335708 | 0.01950646 | 0.314370415 |
| *TRADD* | -0.30522 | 0.130741 | -2.334538 | 0.01956758 | 0.314370415 |
| *PFKFB3* | -0.387987 | 0.166214 | -2.334257 | 0.01958229 | 0.314370415 |
| *TMX4* | -0.277785 | 0.11905 | -2.333349 | 0.01962981 | 0.314370415 |
| *RPL3* | 0.133846 | 0.057402 | 2.331723 | 0.01971529 | 0.314370415 |
| *LPIN1* | 0.56204 | 0.241186 | 2.330321 | 0.01978919 | 0.314370415 |
| *ENSG00000260409* | -2.001854 | 0.859048 | -2.330318 | 0.01978935 | NA |
| *LDHB* | 0.186209 | 0.079962 | 2.328703 | 0.01987479 | 0.314370415 |
| *LSM6* | 0.277774 | 0.119286 | 2.328639 | 0.01987819 | 0.314370415 |
| *ZC3H12D* | -0.625287 | 0.268559 | -2.328301 | 0.0198961 | 0.314370415 |
| *MRPL50* | 0.274045 | 0.117712 | 2.328101 | 0.01990674 | 0.314370415 |
| *MYC* | 0.288218 | 0.123892 | 2.326363 | 0.01999922 | 0.314370415 |
| *FANCF* | 0.353307 | 0.151912 | 2.325733 | 0.02003281 | 0.314370415 |
| *TOMM7* | 0.20183 | 0.086801 | 2.3252 | 0.0200613 | 0.314370415 |

| | | | | | |
|---|---|---|---|---|---|
| *RAF1* | 0.297629 | 0.12801 | 2.325035 | 0.02007008 | 0.314370415 |
| *SFR1* | -0.329746 | 0.141836 | -2.324844 | 0.0200803 | 0.314370415 |
| *SGPP1* | 1.040489 | 0.447668 | 2.324242 | 0.02011252 | NA |
| *LACTB2* | 0.409416 | 0.176294 | 2.322348 | 0.02021419 | 0.315721794 |
| *OAS2* | 0.398018 | 0.171525 | 2.320476 | 0.02031516 | 0.316554081 |
| *EIF4EBP1* | -0.298688 | 0.128795 | -2.31909 | 0.02039016 | 0.316978607 |
| *DOK3* | 0.829356 | 0.357718 | 2.318461 | 0.02042426 | NA |
| *WRB* | 0.375994 | 0.162209 | 2.317956 | 0.02045173 | 0.317192954 |
| *TSC22D1* | -0.599387 | 0.25865 | -2.317367 | 0.02048377 | NA |
| *RPL26* | 0.146212 | 0.063172 | 2.314514 | 0.02063957 | 0.318640168 |
| *DEDD2* | 0.284471 | 0.122909 | 2.314487 | 0.02064105 | 0.318640168 |
| *KBTBD8* | 0.870275 | 0.376142 | 2.313691 | 0.02068467 | NA |
| *WASH9P* | -0.419451 | 0.181349 | -2.312948 | 0.02072548 | 0.319201251 |
| *ENSG00000272106* | -0.459681 | 0.198849 | -2.311708 | 0.02079378 | 0.31951177 |
| *MPEG1* | -0.490849 | 0.212573 | -2.309084 | 0.02093894 | 0.320999336 |
| *GATD3B* | -1.275884 | 0.552572 | -2.308992 | 0.02094403 | NA |
| *SPN* | -1.036413 | 0.449064 | -2.307939 | 0.02100253 | NA |
| *CLCF1* | 0.40249 | 0.174416 | 2.307638 | 0.02101927 | 0.321488235 |
| *NR2F6* | 1.099224 | 0.476779 | 2.30552 | 0.02113745 | NA |
| *POLDIP2* | -0.26453 | 0.114791 | -2.304451 | 0.02119734 | 0.323466525 |
| *IFT22* | -0.361579 | 0.157068 | -2.302046 | 0.02133258 | 0.324409994 |
| *TOR3A* | -0.28128 | 0.12221 | -2.301615 | 0.02135691 | 0.324409994 |
| *TPM4* | 0.237969 | 0.103544 | 2.29824 | 0.02154812 | 0.326567185 |
| *FAM13A.AS1* | -0.791225 | 0.344429 | -2.29721 | 0.02160682 | NA |
| *H6PD* | -0.430523 | 0.187481 | -2.296354 | 0.02165562 | 0.326762007 |
| *CFLAR* | 0.284277 | 0.123799 | 2.296288 | 0.02165943 | 0.326762007 |
| *LY6E.DT* | 1.308133 | 0.569716 | 2.296114 | 0.02166938 | NA |
| *IGHV3.15* | -1.437448 | 0.626262 | -2.29528 | 0.02171707 | NA |
| *AASDH* | 0.406424 | 0.17707 | 2.295268 | 0.02171778 | 0.326899294 |
| *C22orf34* | 0.808 | 0.352154 | 2.294452 | 0.02176457 | NA |
| *SEC24A* | 0.845711 | 0.368713 | 2.293683 | 0.02180869 | NA |
| *KRT10* | -0.263607 | 0.114999 | -2.292262 | 0.02189055 | 0.328754427 |
| *MAT2A* | 0.246138 | 0.10745 | 2.29072 | 0.02197959 | 0.329346518 |
| *ENSG00000254397* | 1.082559 | 0.47268 | 2.290258 | 0.02200634 | NA |
| *HMGB3* | -1.106553 | 0.483661 | -2.287871 | 0.02214503 | NA |
| *INPP5K* | -0.472332 | 0.206667 | -2.285476 | 0.02228495 | 0.332620607 |
| *PRDM8* | -0.719139 | 0.314712 | -2.285068 | 0.02230884 | NA |
| *MED10* | -0.157061 | 0.068743 | -2.284752 | 0.02232737 | 0.332620607 |
| *LTBP4* | 0.518644 | 0.227038 | 2.284393 | 0.02234842 | 0.332620607 |
| *ASAP1* | 0.567859 | 0.248948 | 2.28104 | 0.02254606 | NA |
| *FXYD5* | 0.202793 | 0.089043 | 2.277468 | 0.02275828 | 0.337963014 |
| *STARD9* | 0.64671 | 0.283996 | 2.277182 | 0.02277534 | NA |
| *MAP3K8* | -0.484887 | 0.213069 | -2.275728 | 0.02286229 | 0.338043062 |
| *P2RY10* | 0.200851 | 0.08826 | 2.275674 | 0.02286552 | 0.338043062 |
| *SRA1* | -0.307967 | 0.135384 | -2.274759 | 0.02292036 | 0.338100837 |
| *RPL13A* | 0.118537 | 0.052143 | 2.273299 | 0.02300816 | 0.338643379 |
| *SERPINB6* | 0.426027 | 0.187837 | 2.268069 | 0.02332502 | 0.342547585 |

| | | | | | |
|---|---|---|---|---|---|
| *TRIM32* | 0.901458 | 0.397723 | 2.266547 | 0.02341794 | NA |
| *ENSA* | -0.163036 | 0.071956 | -2.265786 | 0.0234645 | 0.343238718 |
| *TTC21A* | 0.312417 | 0.137895 | 2.265606 | 0.0234755 | 0.343238718 |
| *EXD3* | -0.425364 | 0.187854 | -2.264325 | 0.02355412 | 0.34363135 |
| *UTP14C* | 1.01663 | 0.449 | 2.264211 | 0.02356111 | NA |
| *ENSG00000274922* | -1.438869 | 0.635843 | -2.26293 | 0.02363998 | NA |
| *CERS6* | 0.956849 | 0.422891 | 2.262638 | 0.02365801 | NA |
| *FNTA* | 0.267567 | 0.118271 | 2.262323 | 0.02367745 | 0.344431669 |
| *NSMCE4A* | -0.277282 | 0.122596 | -2.261752 | 0.02371276 | 0.344431669 |
| *TCL1A* | -0.37924 | 0.167845 | -2.25947 | 0.02385414 | 0.344648239 |
| *NSMAF* | 0.601224 | 0.266148 | 2.258985 | 0.02388431 | 0.344648239 |
| *ZC3H7A* | -0.353375 | 0.156445 | -2.258785 | 0.02389677 | 0.344648239 |
| *COX17* | 0.234878 | 0.104013 | 2.258165 | 0.02393535 | 0.344648239 |
| *ENSG00000272758* | 0.590359 | 0.261488 | 2.257694 | 0.02396472 | NA |
| *ZNF747* | -0.521229 | 0.230993 | -2.256464 | 0.02404156 | 0.345428318 |
| *ENSG00000275799* | 0.417137 | 0.185033 | 2.254389 | 0.02417171 | 0.345485311 |
| *TMEM9* | -0.414781 | 0.18409 | -2.253143 | 0.02425015 | 0.345485311 |
| *BNIP3* | 0.369202 | 0.16392 | 2.252329 | 0.02430148 | 0.345485311 |
| *PDHA1* | -0.370658 | 0.164622 | -2.251572 | 0.02434932 | 0.345485311 |
| *TPCN1* | -0.337874 | 0.150061 | -2.251572 | 0.0243493 | 0.345485311 |
| *ALDH9A1* | 0.386823 | 0.171812 | 2.251438 | 0.02435781 | 0.345485311 |
| *KLF8* | -0.327316 | 0.14561 | -2.247896 | 0.02458281 | 0.346749515 |
| *NOTCH2* | 0.319627 | 0.142204 | 2.247667 | 0.0245974 | 0.346749515 |
| *CDKN1B* | 0.2471 | 0.109941 | 2.24757 | 0.02460365 | 0.346749515 |
| *GRAMD1C* | -0.600071 | 0.267232 | -2.245505 | 0.02473575 | 0.347872692 |
| *FXYD2* | -0.886314 | 0.394896 | -2.244423 | 0.02480518 | 0.348111562 |
| *BANK1* | 0.225442 | 0.100497 | 2.243268 | 0.02487955 | 0.348313878 |
| *AFG3L2* | -0.391107 | 0.174534 | -2.240872 | 0.02503437 | 0.348313878 |
| *PTRH1* | -0.472092 | 0.210699 | -2.2406 | 0.02505199 | 0.348313878 |
| *DDX21* | 0.199816 | 0.089188 | 2.240399 | 0.02506502 | 0.348313878 |
| *IER2* | -0.210297 | 0.093877 | -2.240138 | 0.02508196 | 0.348313878 |
| *RPS8* | 0.194387 | 0.086836 | 2.238565 | 0.02518421 | 0.349003781 |
| *RAB11FIP1* | 0.317289 | 0.141832 | 2.237076 | 0.02528138 | 0.349539074 |
| *FRG1* | -0.220349 | 0.09853 | -2.236361 | 0.02532816 | 0.349539074 |
| *RFX7* | 0.565215 | 0.25284 | 2.23546 | 0.02538714 | NA |
| *HSPBAP1* | -0.888631 | 0.397576 | -2.235122 | 0.02540933 | 0.349548696 |
| *TDP1* | 0.496407 | 0.222105 | 2.235011 | 0.02541663 | NA |
| *EIF4B* | 0.148473 | 0.066439 | 2.234744 | 0.02543417 | 0.349548696 |
| *FAM177B* | 0.569952 | 0.255266 | 2.232779 | 0.02556352 | 0.350600448 |
| *THRB* | 1.757718 | 0.787868 | 2.23098 | 0.02568243 | NA |
| *CBL* | 0.417025 | 0.186934 | 2.230868 | 0.02568988 | 0.350995918 |
| *PRNP* | -0.375732 | 0.168433 | -2.230744 | 0.0256981 | 0.350995918 |
| *MANF* | 0.296335 | 0.132913 | 2.229546 | 0.0257776 | 0.351358738 |
| *OAZ2* | -0.178658 | 0.080232 | -2.226782 | 0.02596188 | 0.352901902 |
| *MIR762HG* | -0.727742 | 0.327033 | -2.225283 | 0.02606222 | NA |
| *ZNF844* | -0.477927 | 0.214796 | -2.225025 | 0.02607953 | 0.352901902 |
| *PCIF1* | -0.325859 | 0.146461 | -2.224892 | 0.02608851 | 0.352901902 |

| | | | | | |
|---|---|---|---|---|---|
| *POLR3C* | -0.455669 | 0.204826 | -2.224669 | 0.02610347 | 0.352901902 |
| *FALEC* | -0.478433 | 0.215199 | -2.223209 | 0.02620173 | 0.35329158 |
| *ENSG00000272990* | -0.890217 | 0.400567 | -2.222391 | 0.02625689 | NA |
| *C9orf72* | -0.365745 | 0.164596 | -2.222081 | 0.02627781 | 0.35329158 |
| *VTA1* | 0.401166 | 0.180609 | 2.221187 | 0.02633829 | 0.35329158 |
| *KMT2E.AS1* | -0.420725 | 0.189423 | -2.221085 | 0.02634518 | 0.35329158 |
| *TMSB10* | -0.173529 | 0.078164 | -2.220053 | 0.02641518 | 0.35351607 |
| *NFU1* | -0.272249 | 0.12268 | -2.219186 | 0.0264741 | 0.353591689 |
| *DUSP12* | -0.31519 | 0.142129 | -2.217628 | 0.02658021 | 0.354177454 |
| *APAF1* | 0.558833 | 0.252099 | 2.216722 | 0.02664207 | NA |
| *WASF1* | -0.580722 | 0.261996 | -2.216527 | 0.02665545 | NA |
| *SNX8* | -0.408588 | 0.18434 | -2.216492 | 0.02665785 | 0.354177454 |
| *HES6* | -0.680461 | 0.307029 | -2.216273 | 0.02667279 | NA |
| *TRMT1L* | 0.414968 | 0.187243 | 2.216197 | 0.02667803 | 0.354177454 |
| *LONP2* | -0.253145 | 0.114265 | -2.215418 | 0.02673138 | 0.354177454 |
| *MOAP1* | -0.30311 | 0.136868 | -2.214623 | 0.02678596 | 0.354193649 |
| *ZNF257* | 0.615014 | 0.277874 | 2.213283 | 0.02687811 | NA |
| *CRY1* | -0.584073 | 0.263931 | -2.212977 | 0.02689924 | NA |
| *CRK* | 0.270578 | 0.122276 | 2.212836 | 0.02690894 | 0.354307089 |
| *TCP1* | 0.239004 | 0.108018 | 2.212621 | 0.0269238 | 0.354307089 |
| *HMGN2* | -0.146321 | 0.066148 | -2.212027 | 0.02696482 | 0.354307089 |
| *DYRK2* | 0.395976 | 0.179204 | 2.209634 | 0.02713056 | 0.354307089 |
| *OVCH1.AS1* | -1.670878 | 0.756456 | -2.208823 | 0.02718693 | NA |
| *EPB42* | -0.435922 | 0.197428 | -2.208004 | 0.02724403 | 0.354307089 |
| *HGSNAT* | -0.44682 | 0.202367 | -2.207969 | 0.02724642 | 0.354307089 |
| *TMEM208* | -0.244037 | 0.110608 | -2.206319 | 0.02736169 | 0.354307089 |
| *P4HB* | -0.206133 | 0.093436 | -2.206128 | 0.02737506 | 0.354307089 |
| *AKAP9* | 0.14973 | 0.067879 | 2.205849 | 0.02739461 | 0.354307089 |
| *ICAM3* | 0.22577 | 0.102417 | 2.204421 | 0.02749475 | 0.354307089 |
| *LEAP2* | 0.363248 | 0.164799 | 2.20419 | 0.02751097 | 0.354307089 |
| *LASP1* | 0.246169 | 0.111714 | 2.203571 | 0.02755455 | 0.354307089 |
| *MYO1C* | -0.430923 | 0.195579 | -2.20332 | 0.02757224 | 0.354307089 |
| *TSC22D2* | 0.348475 | 0.15816 | 2.203304 | 0.02757333 | 0.354307089 |
| *CD72* | 0.30898 | 0.140346 | 2.201569 | 0.02769575 | 0.354307089 |
| *EYA3* | 0.783415 | 0.355865 | 2.201435 | 0.02770527 | 0.354307089 |
| *ATP1A1* | -0.288533 | 0.131084 | -2.201141 | 0.02772606 | 0.354307089 |
| *TSEN15* | -0.488987 | 0.22226 | -2.200066 | 0.02780222 | 0.354307089 |
| *C12orf49* | -0.577584 | 0.26255 | -2.199901 | 0.02781392 | 0.354307089 |
| *SUMO1* | -0.179856 | 0.081793 | -2.198911 | 0.02788428 | 0.354307089 |
| *GTF2IRD2* | -0.438613 | 0.199508 | -2.198473 | 0.02791543 | 0.354307089 |
| *NCBP2* | -0.213097 | 0.096996 | -2.196968 | 0.02802273 | 0.354665457 |
| *PARL* | -0.275675 | 0.125559 | -2.195587 | 0.02812153 | 0.354665457 |
| *KLHL2* | -0.574913 | 0.261888 | -2.195265 | 0.02814462 | 0.354665457 |
| *CD82* | -0.282619 | 0.128751 | -2.195087 | 0.02815738 | 0.354665457 |
| *MAP11* | -0.547699 | 0.249589 | -2.194402 | 0.02820654 | NA |
| *SLC16A1.AS1* | 0.329744 | 0.150288 | 2.194085 | 0.02822928 | 0.354897646 |
| *DHFR2* | 0.547235 | 0.249593 | 2.192512 | 0.02834253 | NA |

| | | | | | |
|---|---|---|---|---|---|
| *RBM8A* | -0.135569 | 0.061844 | -2.192109 | 0.0283716 | 0.356012668 |
| *INO80* | -0.322349 | 0.147206 | -2.189788 | 0.02853964 | 0.356529296 |
| *EPHB6* | -0.417276 | 0.190587 | -2.189419 | 0.0285664 | 0.356529296 |
| *ATG16L2* | -0.322414 | 0.147267 | -2.189316 | 0.02857391 | 0.356529296 |
| *MYL6* | -0.138589 | 0.063404 | -2.185796 | 0.02883056 | 0.358427579 |
| *DENR* | -0.185675 | 0.084948 | -2.185748 | 0.02883404 | 0.358427579 |
| *BBX* | 0.182217 | 0.083399 | 2.184892 | 0.0288968 | 0.358536352 |
| *ZNF506* | 0.343448 | 0.157301 | 2.183377 | 0.02900803 | 0.359244953 |
| *PPP1R15A* | -0.524557 | 0.240361 | -2.182376 | 0.02908178 | 0.359487572 |
| *WAC* | 0.178485 | 0.081841 | 2.180872 | 0.02919286 | 0.359590228 |
| *TMED2* | -0.207501 | 0.095149 | -2.180797 | 0.02919842 | 0.359590228 |
| *ENSG00000275418* | -1.465782 | 0.672512 | -2.179563 | 0.0292899 | NA |
| *MMP28* | 1.609095 | 0.738813 | 2.177946 | 0.02941006 | NA |
| *RPL36* | 0.107235 | 0.049272 | 2.176397 | 0.02952556 | 0.362140364 |
| *RIMS3* | -1.560775 | 0.717173 | -2.176287 | 0.02953379 | NA |
| *ENSG00000273748* | -0.903295 | 0.415119 | -2.175992 | 0.02955584 | 0.362140364 |
| *POLM* | -0.266141 | 0.122318 | -2.175814 | 0.02956916 | 0.362140364 |
| *PF4* | 2.393039 | 1.100456 | 2.174588 | 0.02966104 | NA |
| *TTC9C* | 0.250948 | 0.115519 | 2.172347 | 0.02982952 | 0.36465622 |
| *C9orf66* | -1.382959 | 0.63668 | -2.172143 | 0.02984491 | NA |
| *PPP1CB* | -0.222237 | 0.10241 | -2.170058 | 0.03000242 | 0.365672932 |
| *ODF3B* | -0.550128 | 0.25354 | -2.169789 | 0.03002286 | 0.365672932 |
| *RPH3AL* | -1.042184 | 0.480456 | -2.169155 | 0.03007096 | NA |
| *ENSG00000274015* | -1.705455 | 0.786423 | -2.168621 | 0.03011146 | NA |
| *RBM26.AS1* | -0.693014 | 0.319775 | -2.167191 | 0.03022032 | NA |
| *GOLGA6L9* | -1.112481 | 0.513553 | -2.166244 | 0.03029254 | NA |
| *PPWD1* | 0.292306 | 0.135048 | 2.16446 | 0.03042907 | 0.368088243 |
| *YWHAG* | 0.318446 | 0.147129 | 2.164395 | 0.03043401 | 0.368088243 |
| *DAZAP1* | -0.363682 | 0.168032 | -2.164363 | 0.03043651 | 0.368088243 |
| *MIIP* | -0.325787 | 0.150529 | -2.164279 | 0.03044297 | 0.368088243 |
| *GSTP1* | -0.19029 | 0.087982 | -2.162824 | 0.03055474 | 0.368767939 |
| *DUSP2* | -1.898357 | 0.878374 | -2.161217 | 0.03067855 | NA |
| *CNOT6L* | -0.391928 | 0.181368 | -2.160957 | 0.03069869 | 0.369832861 |
| *MPP5* | 0.60044 | 0.278043 | 2.159522 | 0.03080969 | NA |
| *ST20.AS1* | -0.884263 | 0.409559 | -2.159059 | 0.03084556 | NA |
| *THRAP3* | -0.206011 | 0.095431 | -2.158742 | 0.03087016 | 0.370817737 |
| *APOBEC3H* | -1.825928 | 0.845867 | -2.158648 | 0.0308775 | NA |
| *C15orf62* | -0.677962 | 0.314095 | -2.15846 | 0.03089207 | NA |
| *TRIM23* | 0.519078 | 0.240486 | 2.158459 | 0.03089217 | 0.370817737 |
| *LRP5L* | -0.594752 | 0.275611 | -2.157936 | 0.03093279 | NA |
| *ENSG00000261353* | -1.040004 | 0.482125 | -2.157124 | 0.03099598 | NA |
| *TMEM14B* | -0.23109 | 0.107154 | -2.156625 | 0.03103489 | 0.37185853 |
| *CAPN12* | 0.370543 | 0.171891 | 2.155684 | 0.03110831 | 0.372066617 |
| *FAM117B* | -0.542172 | 0.251686 | -2.154164 | 0.03122728 | 0.372216546 |
| *TRIM4* | 0.264445 | 0.122792 | 2.153602 | 0.03127136 | 0.372216546 |
| *PET100* | -0.182837 | 0.084928 | -2.152851 | 0.03133035 | 0.372216546 |
| *ELF1* | 0.213387 | 0.099127 | 2.152663 | 0.03134514 | 0.372216546 |

| | | | | | |
|---|---|---|---|---|---|
| *RPS23* | 0.131221 | 0.061006 | 2.150935 | 0.03148131 | 0.373165934 |
| *CR1* | 0.962701 | 0.44762 | 2.150709 | 0.03149917 | NA |
| *SPTLC1* | -0.64257 | 0.298854 | -2.150109 | 0.03154656 | NA |
| *CALHM6* | 0.263532 | 0.122609 | 2.14937 | 0.03160509 | 0.373965361 |
| *TRIM21* | 0.585018 | 0.272488 | 2.146952 | 0.0317971 | NA |
| *GFM2* | 0.457375 | 0.213192 | 2.145363 | 0.03192387 | NA |
| *UNC50* | 0.313216 | 0.146167 | 2.14286 | 0.03212432 | 0.377867817 |
| *ENSG00000261840* | -0.627235 | 0.292831 | -2.141969 | 0.03219595 | NA |
| *CUL4B* | 0.451479 | 0.210845 | 2.141288 | 0.03225086 | 0.377867817 |
| *DCXR* | -0.312946 | 0.146155 | -2.141194 | 0.03225836 | 0.377867817 |
| *PEX16* | -0.227036 | 0.106038 | -2.141082 | 0.03226745 | 0.377867817 |
| *BCL7A* | -0.476373 | 0.222531 | -2.140705 | 0.03229786 | 0.377867817 |
| *LINC00324* | 0.448688 | 0.209729 | 2.13937 | 0.03240571 | 0.377867817 |
| *ZNF667* | 0.527666 | 0.246701 | 2.138887 | 0.03244485 | NA |
| *MFAP1* | 0.410441 | 0.191917 | 2.138633 | 0.03246537 | 0.377867817 |
| *TBCC* | -0.214757 | 0.100425 | -2.138475 | 0.03247818 | 0.377867817 |
| *SHMT2* | 0.205667 | 0.096179 | 2.138383 | 0.03248565 | 0.377867817 |
| *COASY* | -0.300235 | 0.140418 | -2.138155 | 0.03250415 | 0.377867817 |
| *ASCC1* | -0.429156 | 0.200875 | -2.136439 | 0.03264361 | 0.378825674 |
| *PSMC5* | -0.174202 | 0.081671 | -2.132966 | 0.03292751 | 0.381453425 |
| *HADHA* | 0.229774 | 0.107872 | 2.130056 | 0.03316697 | 0.38335407 |
| *C1GALT1* | 0.307636 | 0.144459 | 2.129571 | 0.03320708 | 0.38335407 |
| *ABHD17A* | 0.380972 | 0.179091 | 2.127249 | 0.03339937 | 0.384904499 |
| *RANBP1* | -0.230047 | 0.108179 | -2.126541 | 0.03345823 | 0.384914624 |
| *IMPACT* | -0.489426 | 0.230288 | -2.125273 | 0.03356386 | NA |
| *ENSG00000282988* | 0.703786 | 0.331256 | 2.124597 | 0.03362028 | 0.385636007 |
| *TMED8* | 0.322393 | 0.151758 | 2.124395 | 0.03363713 | 0.385636007 |
| *DNAJB1* | 0.403781 | 0.190308 | 2.121722 | 0.03386105 | 0.387185545 |
| *EEA1* | 0.385714 | 0.181821 | 2.121391 | 0.03388894 | 0.387185545 |
| *ENSG00000125726* | -1.396224 | 0.658449 | -2.120473 | 0.0339662 | NA |
| *CD24* | -0.171514 | 0.080971 | -2.118223 | 0.03415621 | 0.388701076 |
| *PHF21A* | -0.302211 | 0.142688 | -2.117991 | 0.03417579 | 0.388701076 |
| *FADS3* | -0.223492 | 0.105554 | -2.11733 | 0.03423188 | 0.388701076 |
| *NDUFA13* | -0.186902 | 0.088284 | -2.117047 | 0.03425582 | 0.388701076 |
| *ZNF48* | 0.554701 | 0.262053 | 2.116753 | 0.03428081 | NA |
| *ENSG00000250790* | 1.248255 | 0.589934 | 2.115925 | 0.03435122 | NA |
| *NR6A1* | -1.209176 | 0.571484 | -2.115852 | 0.03435739 | NA |
| *PTPRS* | 1.333231 | 0.630141 | 2.115766 | 0.03436469 | NA |
| *MDK* | -0.675252 | 0.319182 | -2.115568 | 0.03438154 | NA |
| *PDCD7* | 0.439589 | 0.207838 | 2.115055 | 0.03442527 | 0.389957217 |
| *TNFRSF13C* | -0.124297 | 0.058791 | -2.114223 | 0.03449624 | 0.390095491 |
| *SOX4* | -0.510952 | 0.241857 | -2.112618 | 0.03463351 | 0.390494268 |
| *RSPRY1* | 0.377127 | 0.178527 | 2.112435 | 0.03464916 | 0.390494268 |
| *ANKDD1A* | -1.444866 | 0.684181 | -2.111816 | 0.0347022 | NA |
| *PPCS* | -0.195168 | 0.092426 | -2.111622 | 0.03471891 | 0.390617131 |
| *RAB1B* | -0.295767 | 0.140165 | -2.110124 | 0.03484767 | 0.391402417 |
| *RMDN3* | 0.508325 | 0.240915 | 2.109978 | 0.03486023 | NA |

| | | | | | |
|---|---|---|---|---|---|
| *ERV3.1* | -0.246754 | 0.117055 | -2.108018 | 0.03502942 | 0.392239895 |
| *SSU72* | -0.207258 | 0.098325 | -2.107891 | 0.03504041 | 0.392239895 |
| *MAP4K3.DT* | -0.523107 | 0.248215 | -2.107473 | 0.03507664 | NA |
| *LINC02201* | -1.567934 | 0.744408 | -2.106283 | 0.03517978 | NA |
| *PUS3* | 0.413404 | 0.19646 | 2.10427 | 0.03535489 | 0.395093883 |
| *RPL18* | 0.137837 | 0.065567 | 2.10223 | 0.0355331 | 0.396418041 |
| *ZNF646* | 0.687893 | 0.327434 | 2.100856 | 0.03565362 | NA |
| *GCFC2* | -0.449286 | 0.213863 | -2.100813 | 0.03565742 | NA |
| *CEP97* | 0.524226 | 0.249695 | 2.099467 | 0.03577579 | NA |
| *FAM41C* | -0.57364 | 0.27333 | -2.098711 | 0.0358424 | NA |
| *PIN4* | -0.187102 | 0.089187 | -2.097865 | 0.03591704 | 0.400029037 |
| *ENSG00000274605* | 0.742668 | 0.35419 | 2.096809 | 0.03601053 | NA |
| *FAM200A* | 0.59494 | 0.283872 | 2.095804 | 0.03609956 | NA |
| *SEC11C* | -0.258311 | 0.123267 | -2.095547 | 0.03612238 | 0.401642148 |
| *CLPX* | -0.313276 | 0.149636 | -2.093591 | 0.03629641 | 0.402902325 |
| *ENSG00000261505* | -0.636747 | 0.304543 | -2.090826 | 0.03654365 | NA |
| *ATP2A3* | -0.287922 | 0.137768 | -2.089898 | 0.03662697 | 0.405892841 |
| *IFI44L* | 1.039675 | 0.497806 | 2.088515 | 0.0367514 | 0.406490785 |
| *ACP1* | -0.151095 | 0.072377 | -2.087609 | 0.03683315 | 0.406490785 |
| *ELF2* | -0.191329 | 0.091665 | -2.08726 | 0.03686464 | 0.406490785 |
| *RXRA* | -0.876269 | 0.41993 | -2.086701 | 0.03691518 | NA |
| *ENSG00000272335* | 0.971244 | 0.465553 | 2.086216 | 0.03695908 | NA |
| *TUBD1* | 0.34803 | 0.166864 | 2.085715 | 0.03700442 | 0.406929587 |
| *ERICH1* | 0.190517 | 0.09136 | 2.085338 | 0.03703865 | 0.406929587 |
| *MRPL14* | -0.245325 | 0.117674 | -2.08479 | 0.03708834 | 0.406929587 |
| *FKBP3* | -0.185465 | 0.089083 | -2.081951 | 0.03734697 | 0.409091029 |
| *AKAP13* | 0.181504 | 0.087252 | 2.080234 | 0.03750403 | 0.409831096 |
| *IKZF2* | -0.825451 | 0.396863 | -2.079941 | 0.03753092 | NA |
| *ZNF138* | 0.434833 | 0.209068 | 2.079864 | 0.03753801 | 0.409831096 |
| *EVA1B* | -0.783345 | 0.376716 | -2.079407 | 0.03757992 | NA |
| *TM4SF19.AS1* | -1.282044 | 0.61686 | -2.078337 | 0.03767834 | NA |
| *ATF4* | -0.169699 | 0.081666 | -2.077967 | 0.0377124 | 0.410213859 |
| *ZC3H6* | 0.329175 | 0.158424 | 2.077807 | 0.03772713 | 0.410213859 |
| *NAA10* | -0.215631 | 0.103795 | -2.077467 | 0.03775846 | 0.410213859 |
| *FUT2* | 1.548763 | 0.745556 | 2.077325 | 0.03777154 | NA |
| *PLA2G6* | -0.48034 | 0.231276 | -2.076908 | 0.03781004 | NA |
| *ENSG00000270562* | 0.578285 | 0.27859 | 2.075761 | 0.03791609 | NA |
| *FAU* | 0.113768 | 0.054827 | 2.075061 | 0.03798089 | 0.411956075 |
| *TMEM99* | 0.48254 | 0.232634 | 2.07425 | 0.0380561 | NA |
| *POLB* | 0.51607 | 0.248896 | 2.07344 | 0.03813132 | NA |
| *ENSG00000224790* | -1.273637 | 0.614278 | -2.073389 | 0.03813611 | NA |
| *ENSG00000237357* | 1.113593 | 0.537102 | 2.073336 | 0.03814104 | NA |
| *SLC15A3* | -0.651355 | 0.314178 | -2.0732 | 0.03815365 | NA |
| *WDR46* | -0.279631 | 0.134883 | -2.073139 | 0.03815932 | 0.412997172 |
| *ENSG00000257275* | -0.335579 | 0.161905 | -2.072688 | 0.03820131 | 0.412997172 |
| *MAP1LC3A* | -0.693302 | 0.33456 | -2.072276 | 0.03823973 | NA |
| *TCOF1* | -0.2803 | 0.135312 | -2.071507 | 0.03831143 | 0.413514221 |

| | | | | | |
|---|---|---|---|---|---|
| *IGLV3.21* | -1.187244 | 0.573458 | -2.070325 | 0.03842197 | NA |
| *CHRAC1* | -0.236516 | 0.114261 | -2.069966 | 0.03845553 | 0.414395841 |
| *ENSG00000227468* | -1.508145 | 0.729175 | -2.06829 | 0.03861273 | NA |
| *CACYBP* | 0.161629 | 0.078162 | 2.067859 | 0.03865326 | 0.415851437 |
| *TOMM40L* | -0.884936 | 0.427985 | -2.06768 | 0.03867015 | NA |
| *SYNC* | -0.260785 | 0.126228 | -2.065988 | 0.03882965 | 0.415984225 |
| *ISOC2* | -0.398447 | 0.192873 | -2.065851 | 0.03884252 | 0.415984225 |
| *RARA.AS1* | -0.624547 | 0.302336 | -2.065734 | 0.0388536 | 0.415984225 |
| *NACC1* | 0.550731 | 0.266721 | 2.064824 | 0.03893966 | NA |
| *XRRA1* | -0.631194 | 0.305709 | -2.064691 | 0.03895225 | 0.416368826 |
| *CHMP5* | 0.209471 | 0.101521 | 2.063332 | 0.03908109 | 0.417074363 |
| *DCUN1D5* | 0.43368 | 0.21033 | 2.061906 | 0.03921668 | 0.417849671 |
| *FUT4* | -1.384376 | 0.671783 | -2.060749 | 0.03932696 | NA |
| *NEMP1* | 0.676984 | 0.328566 | 2.060424 | 0.03935802 | NA |
| *THOC3* | -0.422708 | 0.20523 | -2.059684 | 0.03942881 | 0.419436546 |
| *CCZ1B* | -0.63728 | 0.30955 | -2.058727 | 0.03952036 | 0.41973784 |
| *SPEF2* | -1.410971 | 0.685649 | -2.057862 | 0.03960341 | NA |
| *FAM86C1* | -0.843412 | 0.410289 | -2.055654 | 0.03981585 | NA |
| *CHCHD5* | -0.254126 | 0.123623 | -2.055646 | 0.03981666 | 0.42220925 |
| *LINC01970* | -0.890605 | 0.433269 | -2.055548 | 0.03982612 | NA |
| *IL10RB* | 0.377243 | 0.183593 | 2.054779 | 0.03990032 | 0.422209674 |
| *TMEM170A* | 0.240952 | 0.11731 | 2.053979 | 0.03997777 | 0.422209674 |
| *ENSG00000262420* | -1.235971 | 0.601753 | -2.053951 | 0.03998048 | NA |
| *PSMB10* | -0.21918 | 0.106726 | -2.053671 | 0.04000752 | 0.422209674 |
| *DOK1* | 0.328708 | 0.160277 | 2.050876 | 0.04027902 | 0.423919206 |
| *GMFG* | -0.1261 | 0.061491 | -2.050689 | 0.04029723 | 0.423919206 |
| *NBDY* | -0.179734 | 0.087683 | -2.049808 | 0.04038313 | 0.424006027 |
| *TTLL12* | -0.643269 | 0.313854 | -2.049582 | 0.04040525 | NA |
| *TELO2* | -0.424981 | 0.207379 | -2.049295 | 0.04043324 | 0.424006027 |
| *NSUN7* | -0.678539 | 0.331114 | -2.04926 | 0.04043674 | NA |
| *RAB3A* | 1.062736 | 0.518812 | 2.048403 | 0.04052056 | NA |
| *STARD10* | -0.378432 | 0.184751 | -2.048338 | 0.04052692 | 0.424318103 |
| *CD99L2* | -0.631275 | 0.308269 | -2.047806 | 0.040579 | NA |
| *GTPBP6* | -0.574254 | 0.280515 | -2.047143 | 0.04064403 | 0.424590754 |
| *TAF1C* | -0.388739 | 0.189993 | -2.046069 | 0.04074957 | 0.424590754 |
| *TAPBP* | -0.233335 | 0.114063 | -2.045672 | 0.04078863 | 0.424590754 |
| *TMEM179B* | -0.223179 | 0.109109 | -2.045467 | 0.04080881 | 0.424590754 |
| *SNAI3* | 0.716255 | 0.350209 | 2.045221 | 0.04083307 | NA |
| *UBAP2* | -0.421248 | 0.206011 | -2.04479 | 0.04087556 | 0.424619668 |
| *ENSG00000260572* | -1.579772 | 0.772673 | -2.044554 | 0.04089881 | NA |
| *COMMD1* | -0.3176 | 0.155493 | -2.04253 | 0.04109903 | 0.425382421 |
| *RYBP* | 0.405155 | 0.198387 | 2.04225 | 0.04112672 | 0.425382421 |
| *EFHB* | 1.573414 | 0.770622 | 2.041744 | 0.04117695 | NA |
| *MSRB1* | -0.307995 | 0.15091 | -2.040919 | 0.04125888 | 0.425382421 |
| *MAPRE2* | 0.26686 | 0.13077 | 2.040684 | 0.04128227 | 0.425382421 |
| *KIAA1191* | 0.327508 | 0.160554 | 2.03986 | 0.04136431 | 0.425382421 |
| *HSD17B7* | 0.309794 | 0.151902 | 2.039437 | 0.04140644 | 0.425382421 |

| | | | | | |
|---|---|---|---|---|---|
| *ZNF397* | 0.191922 | 0.09415 | 2.038465 | 0.04150343 | 0.425382421 |
| *CD2BP2* | -0.220013 | 0.107949 | -2.038128 | 0.04153714 | 0.425382421 |
| *NFKB2* | -0.31114 | 0.152715 | -2.03739 | 0.04161102 | 0.425382421 |
| *FAM161B* | -0.953978 | 0.46825 | -2.037327 | 0.04161725 | NA |
| *ING1* | 0.370143 | 0.181686 | 2.037268 | 0.04162314 | 0.425382421 |
| *NOC2L* | 0.21945 | 0.107766 | 2.036348 | 0.04171539 | 0.425382421 |
| *FAM107B* | -0.277558 | 0.136304 | -2.036323 | 0.04171798 | 0.425382421 |
| *LINC02413* | 1.321044 | 0.648742 | 2.036316 | 0.04171861 | NA |
| *LINC02550* | 1.360633 | 0.668589 | 2.035082 | 0.04184266 | NA |
| *CD8B* | -2.908313 | 1.429836 | -2.034018 | 0.04194976 | NA |
| *PSPC1* | -0.302681 | 0.148815 | -2.033944 | 0.04195728 | 0.426975521 |
| *MYO5C* | -1.382105 | 0.679624 | -2.033632 | 0.04198868 | NA |
| *QDPR* | 0.689701 | 0.339281 | 2.032831 | 0.04206957 | NA |
| *BOLA2.SMG1P6* | -0.532937 | 0.262205 | -2.03252 | 0.04210104 | NA |
| *CIAO2B* | 0.171633 | 0.084453 | 2.032279 | 0.04212539 | 0.426975521 |
| *ARID5A* | -0.376573 | 0.185306 | -2.032168 | 0.04213664 | 0.426975521 |
| *PNRC2* | 0.151481 | 0.074556 | 2.031758 | 0.04217817 | 0.426975521 |
| *PIGT* | -0.279894 | 0.137771 | -2.031584 | 0.04219583 | 0.426975521 |
| *C15orf40* | -0.230515 | 0.113552 | -2.030033 | 0.04235317 | 0.427556799 |
| *CPM* | -0.721817 | 0.355594 | -2.029889 | 0.04236784 | NA |
| *GSTO1* | 0.237936 | 0.117225 | 2.029749 | 0.0423821 | 0.427556799 |
| *IGLV4.69* | -1.244647 | 0.613425 | -2.029012 | 0.04245712 | NA |
| *SERPINI1* | 0.582509 | 0.287294 | 2.027575 | 0.04260367 | NA |
| *NDUFV2.AS1* | -0.386991 | 0.190865 | -2.027565 | 0.04260462 | 0.429149379 |
| *PRRT3* | 0.588855 | 0.290536 | 2.026791 | 0.04268381 | NA |
| *GLUL* | -0.489441 | 0.241526 | -2.026451 | 0.04271856 | 0.429645136 |
| *AMIGO2* | 0.950752 | 0.469377 | 2.025563 | 0.0428096 | NA |
| *HSD17B11* | 0.135569 | 0.066932 | 2.025493 | 0.04281677 | 0.429819387 |
| *DNAJC7* | -0.196225 | 0.0969 | -2.025019 | 0.04286539 | 0.429819387 |
| *WNT10A* | -0.650252 | 0.321152 | -2.02475 | 0.04289309 | NA |
| *NDUFA4* | -0.109192 | 0.053956 | -2.023731 | 0.04299784 | 0.430029262 |
| *XG* | 1.851903 | 0.91519 | 2.023518 | 0.04301981 | NA |
| *ENSG00000247765* | -1.459896 | 0.721593 | -2.023158 | 0.04305685 | NA |
| *GEMIN6* | 0.257083 | 0.127077 | 2.02305 | 0.043068 | 0.430029262 |
| *CLASRP* | -0.251755 | 0.124451 | -2.022927 | 0.04308067 | 0.430029262 |
| *CNR1* | 0.716045 | 0.354001 | 2.022719 | 0.04310213 | NA |
| *CLOCK* | -0.342908 | 0.169688 | -2.020817 | 0.04329867 | 0.431556447 |
| *C12orf76* | 0.247853 | 0.122762 | 2.018969 | 0.04349048 | 0.432247825 |
| *TP53INP1* | 0.39232 | 0.194324 | 2.018894 | 0.04349828 | 0.432247825 |
| *APH1A* | -0.178357 | 0.088399 | -2.017645 | 0.0436282 | 0.432279608 |
| *ATF7* | -0.396479 | 0.196509 | -2.017612 | 0.04363172 | 0.432279608 |
| *EHD3* | -0.614361 | 0.304513 | -2.017522 | 0.04364108 | NA |
| *TOMM5* | -0.240312 | 0.119152 | -2.016857 | 0.04371049 | 0.432414614 |
| *CXorf65* | -0.690161 | 0.342415 | -2.015571 | 0.04384489 | NA |
| *BCL6* | -0.359361 | 0.178358 | -2.014824 | 0.04392308 | 0.432984833 |
| *ERGIC2* | 0.208406 | 0.103442 | 2.0147 | 0.04393612 | 0.432984833 |
| *N6AMT1* | 0.557884 | 0.276923 | 2.014582 | 0.04394851 | NA |

| | | | | | |
|---|---|---|---|---|---|
| *HIBCH* | 0.243815 | 0.121034 | 2.014436 | 0.04396381 | 0.432984833 |
| *CYHR1* | 0.238549 | 0.118514 | 2.012838 | 0.04413171 | 0.433994483 |
| *ANKMY1* | -0.456868 | 0.227045 | -2.012236 | 0.04419506 | NA |
| *ZNF786* | 0.549369 | 0.273143 | 2.011287 | 0.04429511 | NA |
| *ENSG00000197302* | 0.245058 | 0.121897 | 2.010373 | 0.04439175 | 0.434957504 |
| *POLR2I* | -0.225578 | 0.112223 | -2.010085 | 0.04442217 | 0.434957504 |
| *PRDX6* | -0.144163 | 0.071741 | -2.009504 | 0.04448374 | 0.434957504 |
| *KDM4B* | 0.336336 | 0.167379 | 2.009428 | 0.04449174 | 0.434957504 |
| *ARMC10* | -0.705312 | 0.351496 | -2.0066 | 0.04479223 | NA |
| *TSEN54* | -0.530212 | 0.26425 | -2.00648 | 0.04480511 | NA |
| *LTA4H* | 0.190418 | 0.094932 | 2.005841 | 0.04487319 | 0.438041487 |
| *HCG11* | -0.43486 | 0.216839 | -2.005445 | 0.04491546 | NA |
| *TAF6L* | -0.659766 | 0.329128 | -2.00459 | 0.04500692 | NA |
| *PRR4* | 0.364545 | 0.181925 | 2.003819 | 0.04508944 | 0.439506197 |
| *MAGED1* | -0.582391 | 0.290831 | -2.002509 | 0.04523004 | NA |
| *G6PC3* | -0.490079 | 0.244808 | -2.001894 | 0.04529609 | NA |
| *ATP11A* | 0.542068 | 0.270896 | 2.00102 | 0.04539023 | NA |
| *NUP210* | 0.204825 | 0.102401 | 2.000218 | 0.0454767 | 0.442331376 |
| *MRPS36* | 0.180506 | 0.090258 | 1.999886 | 0.04551255 | 0.442331376 |
| *SMIM20* | 0.333107 | 0.166624 | 1.999157 | 0.04559133 | 0.442449213 |
| *POLE2* | 0.87002 | 0.435503 | 1.997736 | 0.04574526 | NA |
| *ERF* | 0.475606 | 0.23808 | 1.997673 | 0.04575215 | NA |
| *ZNF628* | -0.931539 | 0.466626 | -1.996326 | 0.04589845 | NA |
| *PCID2* | 0.25292 | 0.126739 | 1.9956 | 0.04597748 | 0.443733612 |
| *RALY* | -0.154601 | 0.077473 | -1.995535 | 0.04598453 | 0.443733612 |
| *NELFE* | -0.173316 | 0.086864 | -1.995246 | 0.04601601 | 0.443733612 |
| *LAMC1* | 0.806688 | 0.404358 | 1.994983 | 0.04604475 | NA |
| *CTNNBIP1* | -0.688917 | 0.345339 | -1.994899 | 0.04605384 | NA |
| *IL6* | -0.780578 | 0.391386 | -1.994396 | 0.04610885 | NA |
| *ENSG00000267040* | 0.238568 | 0.119622 | 1.994348 | 0.04611402 | 0.443733612 |
| *SLA* | 0.380222 | 0.190655 | 1.994287 | 0.04612073 | 0.443733612 |
| *ZPR1* | 0.2415 | 0.121132 | 1.993694 | 0.04618546 | 0.443733612 |
| *FLCN* | 0.328253 | 0.16465 | 1.993638 | 0.04619161 | 0.443733612 |
| *DPH3* | -0.305803 | 0.153443 | -1.992942 | 0.0462678 | 0.443823168 |
| *CEMIP2* | 0.505914 | 0.254027 | 1.991575 | 0.0464177 | 0.44455636 |
| *CCL5* | -1.14281 | 0.573874 | -1.991396 | 0.04643732 | NA |
| *GPR160* | 0.57876 | 0.290684 | 1.991025 | 0.04647817 | 0.44455636 |
| *GTF3C6* | -0.205495 | 0.103252 | -1.990222 | 0.04656646 | 0.444759915 |
| *SENP6* | 0.208346 | 0.104726 | 1.98945 | 0.04665154 | 0.444932378 |
| *CCNG2* | 0.376171 | 0.189158 | 1.988656 | 0.0467392 | 0.445128842 |
| *TFDP2* | -0.277722 | 0.139812 | -1.986391 | 0.04698997 | 0.44687599 |
| *HIST1H3H* | 1.286066 | 0.647511 | 1.986167 | 0.04701474 | NA |
| *GTF2H2C* | -0.380469 | 0.191925 | -1.98238 | 0.04743675 | 0.449014872 |
| *RBAK.RBAKDN* | 0.345426 | 0.174249 | 1.982373 | 0.0474375 | 0.449014872 |
| *USP53* | 1.123284 | 0.566678 | 1.982225 | 0.04745401 | 0.449014872 |
| *CPQ* | -0.641696 | 0.323737 | -1.982155 | 0.04746195 | NA |
| *MKNK1* | 0.283301 | 0.142976 | 1.981467 | 0.0475389 | 0.449014872 |

| | | | | | |
|---|---|---|---|---|---|
| *PSPH* | -0.829097 | 0.41843 | -1.981446 | 0.04754123 | NA |
| *HMGXB4* | -0.241921 | 0.1221 | -1.981341 | 0.0475531 | 0.449014872 |
| *UBA6* | 0.27138 | 0.137014 | 1.980675 | 0.04762771 | 0.44908061 |
| *DAPK2* | -0.77251 | 0.390102 | -1.980275 | 0.04767258 | NA |
| *DLGAP3* | -0.921743 | 0.465591 | -1.979728 | 0.04773415 | NA |
| *TTC9* | 0.412173 | 0.20825 | 1.979219 | 0.04779137 | NA |
| *FXYD1* | 0.530461 | 0.2681 | 1.978591 | 0.0478621 | 0.450315583 |
| *PPM1N* | -0.438653 | 0.221732 | -1.978304 | 0.04789437 | 0.450315583 |
| *PTPRC* | 0.134667 | 0.068104 | 1.977365 | 0.04800039 | 0.450674131 |
| *NPEPL1* | -0.523817 | 0.264926 | -1.977217 | 0.04801708 | NA |
| *LINC00663* | 0.696683 | 0.352402 | 1.976953 | 0.04804695 | NA |
| *C1GALT1C1* | 0.423372 | 0.214188 | 1.976639 | 0.04808241 | NA |
| *IRAK3* | 1.184596 | 0.599576 | 1.975722 | 0.04818622 | NA |
| *MRPL39* | 0.318243 | 0.161108 | 1.975344 | 0.04822916 | 0.452182477 |
| *CCDC28A* | 0.27493 | 0.139236 | 1.974566 | 0.04831738 | 0.452370592 |
| *MEX3A* | 1.040362 | 0.527247 | 1.973197 | 0.0484731 | NA |
| *MDFIC* | 0.749214 | 0.379794 | 1.972684 | 0.04853158 | NA |
| *ZNF346* | 0.544996 | 0.276313 | 1.972387 | 0.04856542 | NA |
| *FNDC3B* | -1.180799 | 0.598745 | -1.972122 | 0.04859564 | NA |
| *GDF11* | 0.295078 | 0.149662 | 1.971633 | 0.04865148 | 0.453961136 |
| *NMT2* | -0.269505 | 0.136719 | -1.971224 | 0.0486983 | 0.453961136 |
| *MKLN1* | 0.292251 | 0.148306 | 1.9706 | 0.0487697 | 0.453961136 |
| *NDUFB11* | 0.170917 | 0.086796 | 1.969172 | 0.04893332 | 0.453961136 |
| *CARD8* | -0.231938 | 0.117803 | -1.968862 | 0.04896889 | 0.453961136 |
| *SIKE1* | 0.268053 | 0.136175 | 1.968452 | 0.04901604 | 0.453961136 |
| *SPAG1* | -0.585204 | 0.297343 | -1.968112 | 0.04905513 | NA |
| *SPCS1* | -0.16638 | 0.084549 | -1.967861 | 0.04908408 | 0.453961136 |
| *NARS* | 0.213928 | 0.108729 | 1.967526 | 0.04912256 | 0.453961136 |
| *SOCS4* | 0.466069 | 0.236885 | 1.967491 | 0.04912658 | NA |
| *RPA2* | 0.192852 | 0.098031 | 1.967247 | 0.04915474 | 0.453961136 |
| *STMN1* | -0.218426 | 0.111039 | -1.967105 | 0.04917114 | 0.453961136 |
| *LRRC8B* | 0.618861 | 0.314659 | 1.966768 | 0.04920992 | NA |
| *SENP3* | -0.518271 | 0.263675 | -1.965569 | 0.04934846 | NA |
| *ENSG00000105705* | -0.392993 | 0.199969 | -1.965269 | 0.0493831 | 0.455284759 |
| *YJEFN3* | 0.833037 | 0.423913 | 1.965111 | 0.04940144 | NA |
| *MFSD6* | 0.759613 | 0.386652 | 1.964589 | 0.04946188 | NA |
| *PSMB3* | -0.175919 | 0.089623 | -1.96287 | 0.04966132 | 0.456832518 |
| *ENSG00000231760* | -1.121619 | 0.571722 | -1.961826 | 0.04978277 | NA |
| *NPC2* | 0.29125 | 0.148519 | 1.96103 | 0.04987553 | 0.456832518 |
| *RPLP2* | 0.160744 | 0.08197 | 1.961008 | 0.04987805 | 0.456832518 |
| *CREBL2* | 0.359342 | 0.18328 | 1.960616 | 0.04992384 | 0.456832518 |
| *CLNS1A* | 0.306302 | 0.156234 | 1.960534 | 0.04993337 | 0.456832518 |

| Cluster 3 | log2FC | lfcSE | stat | pvalue | padj |
|---|---|---|---|---|---|
| *ARRDC3* | -0.906905 | 0.105263 | -8.615607 | < 2.22e-16 | 6.71E-14 |
| *CKS2* | 1.060892 | 0.156998 | 6.757345 | 1.41E-11 | 6.77E-08 |
| *ZBED2* | 2.066521 | 0.311917 | 6.625237 | 3.47E-11 | 1.11E-07 |
| *SMIM10* | -4.230547 | 0.66977 | -6.316416 | 2.68E-10 | NA |
| *LRIF1* | 0.666533 | 0.107266 | 6.213866 | 5.17E-10 | 1.25E-06 |
| *CD83* | -0.444947 | 0.073151 | -6.08258 | 1.18E-09 | 2.28E-06 |
| *RIN3* | 0.695222 | 0.122011 | 5.698023 | 1.21E-08 | 1.95E-05 |
| *GATD3A* | 5.237077 | 0.938273 | 5.581615 | 2.38E-08 | 3.28E-05 |
| *ASF1A* | 0.675453 | 0.132032 | 5.115829 | 3.12E-07 | 0.000348589 |
| *NFKBIA* | -0.405485 | 0.079381 | -5.108082 | 3.25E-07 | 0.000348589 |
| *SCRN1* | 1.075203 | 0.214137 | 5.021097 | 5.14E-07 | 0.000495276 |
| *AIM2* | 1.736807 | 0.347411 | 4.999281 | 5.75E-07 | NA |
| *GPR65* | 0.543458 | 0.109702 | 4.953934 | 7.27E-07 | 0.000637361 |
| *DYNLL1* | 0.411612 | 0.083675 | 4.919141 | 8.69E-07 | 0.000698298 |
| *ARL17A* | -0.614261 | 0.126289 | -4.863922 | 1.15E-06 | 0.000734787 |
| *CIRBP* | -0.333865 | 0.068691 | -4.860373 | 1.17E-06 | 0.000734787 |
| *ENSG00000273319* | 0.612141 | 0.126004 | 4.858106 | 1.19E-06 | 0.000734787 |
| *CHORDC1* | 0.556131 | 0.114826 | 4.843251 | 1.28E-06 | 0.000734787 |
| *ATF7IP2* | -0.651997 | 0.134699 | -4.840399 | 1.30E-06 | 0.000734787 |
| *LMO4* | 0.60854 | 0.127267 | 4.781616 | 1.74E-06 | 0.000931285 |
| *ZNF441* | -0.722512 | 0.151552 | -4.767413 | 1.87E-06 | 0.000946783 |
| *TTC32* | -0.676371 | 0.145857 | -4.637225 | 3.53E-06 | 0.001640089 |
| *MS4A1* | -0.402119 | 0.086864 | -4.629271 | 3.67E-06 | 0.001640089 |
| *KRCC1* | 0.380676 | 0.082357 | 4.62226 | 3.80E-06 | 0.001640089 |
| *RBM3* | -0.430066 | 0.093325 | -4.608245 | 4.06E-06 | 0.001640089 |
| *GALNT2* | -0.6758 | 0.146687 | -4.607101 | 4.08E-06 | 0.001640089 |
| *RHOH* | 0.373675 | 0.081743 | 4.571317 | 4.85E-06 | 0.001868877 |
| *DNAAF4* | -1.166947 | 0.258729 | -4.510303 | 6.47E-06 | 0.002367313 |
| *SLC3A2* | -0.499769 | 0.110931 | -4.505219 | 6.63E-06 | 0.002367313 |
| *ENSG00000226571* | 1.235661 | 0.278802 | 4.432042 | 9.33E-06 | 0.003213735 |
| *SIGLEC5* | 1.782323 | 0.402959 | 4.423083 | 9.73E-06 | 0.003234457 |
| *SNAP23* | -0.293072 | 0.067151 | -4.36434 | 1.28E-05 | 0.004084577 |
| *FAM131A* | 0.782884 | 0.180143 | 4.345911 | 1.39E-05 | 0.004084577 |
| *CFLAR* | 0.389848 | 0.08978 | 4.342266 | 1.41E-05 | 0.004084577 |
| *CLECL1* | 1.747045 | 0.402629 | 4.339094 | 1.43E-05 | 0.004084577 |
| *ARID5A* | -0.621658 | 0.143319 | -4.337577 | 1.44E-05 | 0.004084577 |
| *ENSG00000254397* | 1.389237 | 0.321424 | 4.322136 | 1.55E-05 | 0.004223669 |
| *GLUL* | -0.641415 | 0.148558 | -4.317606 | 1.58E-05 | 0.004223669 |
| *TNFAIP8L2* | 0.679909 | 0.157704 | 4.311298 | 1.62E-05 | 0.004228543 |
| *PCIF1* | -0.455296 | 0.106268 | -4.284405 | 1.83E-05 | 0.004648238 |
| *HHEX* | 0.360067 | 0.084353 | 4.268581 | 1.97E-05 | 0.004727466 |
| *CDKN1B* | 0.332463 | 0.077921 | 4.266657 | 1.98E-05 | 0.004727466 |
| *HMGB2* | -0.378623 | 0.088801 | -4.263706 | 2.01E-05 | 0.004727466 |
| *NOP10* | 0.294742 | 0.069322 | 4.251756 | 2.12E-05 | 0.004795266 |
| *DUSP14* | 0.66526 | 0.156537 | 4.249868 | 2.14E-05 | 0.004795266 |
| *CLEC2D* | -0.386421 | 0.092384 | -4.182786 | 2.88E-05 | 0.006308894 |

| | | | | | |
|---|---|---|---|---|---|
| *PIGF* | 0.688922 | 0.165275 | 4.168329 | 3.07E-05 | 0.006444678 |
| *IZUMO4* | -0.629073 | 0.150936 | -4.16782 | 3.08E-05 | 0.006444678 |
| *SNAI3* | 0.997271 | 0.240037 | 4.154662 | 3.26E-05 | 0.00668171 |
| *RNF34* | 0.435951 | 0.105574 | 4.129333 | 3.64E-05 | 0.007306662 |
| *JADE1* | 0.445268 | 0.108046 | 4.121105 | 3.77E-05 | 0.007418079 |
| *MIR29B2CHG* | 0.465665 | 0.11402 | 4.084077 | 4.43E-05 | 0.008531866 |
| *KIAA0040* | 0.686825 | 0.168627 | 4.073045 | 4.64E-05 | 0.008770964 |
| *MDFIC* | 0.984743 | 0.243154 | 4.049878 | 5.12E-05 | 0.009499929 |
| *MRPS36* | 0.255238 | 0.063632 | 4.011149 | 6.04E-05 | 0.010990331 |
| *ARL6IP6* | 0.42088 | 0.10511 | 4.004196 | 6.22E-05 | 0.011108984 |
| *RFTN1* | -0.479819 | 0.120415 | -3.984725 | 6.76E-05 | 0.011841146 |
| *ADAMTS6* | 0.986226 | 0.247795 | 3.980015 | 6.89E-05 | 0.01186254 |
| *SORD* | -0.844154 | 0.213829 | -3.947806 | 7.89E-05 | 0.013204865 |
| *POU2F2* | 0.285831 | 0.072434 | 3.946058 | 7.94E-05 | 0.013204865 |
| *NT5E* | 1.02071 | 0.259458 | 3.934011 | 8.35E-05 | 0.013649571 |
| *TXN* | 0.486873 | 0.123957 | 3.927768 | 8.57E-05 | 0.013660976 |
| *ZBTB25* | 0.461119 | 0.117563 | 3.922304 | 8.77E-05 | 0.013660976 |
| *NAB2* | 0.929734 | 0.237064 | 3.921879 | 8.79E-05 | 0.013660976 |
| *MTMR6* | -0.600899 | 0.154273 | -3.895035 | 9.82E-05 | 0.014663406 |
| *ERV3.1* | -0.38838 | 0.099768 | -3.892825 | 9.91E-05 | 0.014663406 |
| *MIF* | -0.347729 | 0.089397 | -3.889719 | 0.00010036 | 0.014663406 |
| *CCDC117* | 0.557054 | 0.143215 | 3.889641 | 0.000100393 | 0.014663406 |
| *YWHAE* | 0.355407 | 0.091824 | 3.870519 | 0.000108604 | 0.015625973 |
| *HLA.G* | -1.71986 | 0.445191 | -3.863198 | 0.000111912 | NA |
| *NUFIP2* | 0.462702 | 0.120234 | 3.848356 | 0.000118913 | 0.016857684 |
| *CYTOR* | 1.331962 | 0.347611 | 3.831758 | 0.000127231 | 0.017775459 |
| *IGHD* | -0.442154 | 0.115836 | -3.817062 | 0.00013505 | 0.018350263 |
| *KCNK6* | 0.59021 | 0.154712 | 3.814893 | 0.000136242 | 0.018350263 |
| *SYNGR2* | -0.564226 | 0.148074 | -3.810442 | 0.000138718 | 0.018350263 |
| *APBB2* | -1.03418 | 0.27157 | -3.808158 | 0.000140006 | 0.018350263 |
| *H2AFZ* | 0.309938 | 0.08142 | 3.806648 | 0.000140863 | 0.018350263 |
| *TMEM170A* | 0.354453 | 0.093204 | 3.802985 | 0.000142963 | 0.01837554 |
| *SMIM20* | 0.454421 | 0.119613 | 3.799095 | 0.000145226 | 0.018407057 |
| *OCLN* | -1.057143 | 0.278486 | -3.796037 | 0.000147027 | 0.018407057 |
| *SLC50A1* | -0.350839 | 0.092632 | -3.787454 | 0.000152199 | 0.018587949 |
| *CRIPT* | 0.395735 | 0.104515 | 3.786377 | 0.00015286 | 0.018587949 |
| *TM2D3* | 0.396752 | 0.104847 | 3.784114 | 0.000154257 | 0.018587949 |
| *ATP6V1C2* | -2.678555 | 0.708674 | -3.779672 | 0.000157035 | NA |
| *LCN8* | 3.906921 | 1.033802 | 3.779178 | 0.000157347 | 0.01872625 |
| *SKAP2* | 0.273097 | 0.072546 | 3.764445 | 0.000166919 | 0.01960483 |
| *IL21R* | -0.963932 | 0.256253 | -3.761649 | 0.000168797 | 0.01960483 |
| *SLC2A14* | -3.80731 | 1.012741 | -3.759412 | 0.000170313 | NA |
| *GLS* | 0.421884 | 0.11264 | 3.745437 | 0.00018008 | 0.020666332 |
| *ENSG00000272221* | 0.858746 | 0.230673 | 3.722779 | 0.000197042 | 0.022346868 |
| *CLEC2B* | 0.463362 | 0.124583 | 3.719303 | 0.000199773 | 0.022393175 |
| *RAB31* | 0.970033 | 0.261755 | 3.705882 | 0.000210657 | 0.023341733 |
| *EGLN2* | 0.312647 | 0.084478 | 3.700947 | 0.000214796 | 0.023529961 |

| | | | | | |
|---|---|---|---|---|---|
| *HLX* | 1.210844 | 0.328047 | 3.691072 | 0.000223311 | 0.024100005 |
| *HSBP1* | 0.321188 | 0.087063 | 3.689155 | 0.000225 | 0.024100005 |
| *MIR155HG* | 0.830094 | 0.225835 | 3.675673 | 0.000237223 | 0.025130023 |
| *CRK* | 0.377479 | 0.102893 | 3.668673 | 0.000243813 | 0.025547364 |
| *ACADVL* | -0.270675 | 0.07417 | -3.649383 | 0.000262871 | 0.027248098 |
| *SCN3A* | 1.352025 | 0.370853 | 3.645716 | 0.000266649 | NA |
| *ABTB1* | -0.29489 | 0.080896 | -3.645284 | 0.000267097 | 0.02726419 |
| *CYB5A* | -0.49586 | 0.136085 | -3.643762 | 0.000268682 | 0.02726419 |
| *PCNX4* | 0.413661 | 0.113789 | 3.635331 | 0.000277624 | 0.027878105 |
| *MPLKIP* | 0.231161 | 0.063715 | 3.628079 | 0.000285538 | 0.028377211 |
| *GLRX2* | 0.564294 | 0.155782 | 3.622327 | 0.000291965 | 0.028719828 |
| *TAGLN2* | -0.252827 | 0.069977 | -3.612983 | 0.000302695 | 0.029474552 |
| *TNFAIP1* | 0.595717 | 0.165602 | 3.597274 | 0.00032157 | 0.030733531 |
| *TRIR* | 0.147309 | 0.040954 | 3.596925 | 0.000322001 | 0.030733531 |
| *TAF1D* | -0.182812 | 0.050932 | -3.589331 | 0.000331528 | 0.031332605 |
| *RSRP1* | 0.283335 | 0.079185 | 3.578134 | 0.000346056 | 0.032388164 |
| *RAF1* | 0.322477 | 0.090209 | 3.574763 | 0.000350545 | 0.03249284 |
| *YJEFN3* | 0.927618 | 0.261254 | 3.550633 | 0.000384306 | 0.035282935 |
| *MT1X* | -0.879434 | 0.247879 | -3.54784 | 0.000388404 | 0.035322756 |
| *LHFPL4* | 1.11981 | 0.31593 | 3.544483 | 0.000393384 | NA |
| *SFR1* | -0.410136 | 0.115795 | -3.541921 | 0.000397224 | 0.035787254 |
| *XIAP* | 0.388941 | 0.109911 | 3.5387 | 0.000402103 | 0.035891399 |
| *TIMM10* | -0.497742 | 0.140926 | -3.531939 | 0.000412524 | 0.035939054 |
| *TTC17* | -0.319771 | 0.090552 | -3.531368 | 0.000413417 | 0.035939054 |
| *CNFN* | -0.429206 | 0.121585 | -3.530097 | 0.000415407 | 0.035939054 |
| *MME* | -2.306357 | 0.653879 | -3.527192 | 0.000419992 | NA |
| *CHMP6* | -0.496426 | 0.140764 | -3.526658 | 0.000420841 | 0.035939054 |
| *SESN3* | 0.309394 | 0.087782 | 3.524573 | 0.000424165 | 0.035939054 |
| *ENSG00000272211* | -1.021569 | 0.289885 | -3.524049 | 0.000425005 | 0.035939054 |
| *PTPRS* | 2.08247 | 0.591799 | 3.518879 | 0.000433374 | NA |
| *SNRPN* | 0.260136 | 0.074103 | 3.510469 | 0.000447317 | 0.03749685 |
| *MYBL2* | -1.938921 | 0.552882 | -3.506933 | 0.000453303 | NA |
| *TMEM243* | 0.338596 | 0.096971 | 3.491723 | 0.000479916 | 0.039882676 |
| *MAT2A* | 0.283832 | 0.081397 | 3.487015 | 0.000488444 | 0.040226173 |
| *ENSG00000282988* | 0.854927 | 0.245347 | 3.484568 | 0.000492932 | 0.040226173 |
| *IFIT5* | 0.473834 | 0.13609 | 3.481769 | 0.000498114 | 0.040226173 |
| *CHRAC1* | -0.321039 | 0.092243 | -3.48036 | 0.000500741 | 0.040226173 |
| *COL9A3* | -1.723667 | 0.495742 | -3.476946 | 0.00050716 | 0.040405121 |
| *GPR146* | -0.931862 | 0.268656 | -3.468609 | 0.000523159 | 0.04130651 |
| *IFT57* | 0.52268 | 0.150775 | 3.466621 | 0.000527044 | 0.04130651 |
| *FNBP1* | -0.345967 | 0.100181 | -3.453404 | 0.000553559 | 0.043034768 |
| *GTF3C5* | -0.395877 | 0.114785 | -3.448845 | 0.000562991 | 0.043417837 |
| *MRPL28* | -0.287606 | 0.083553 | -3.442193 | 0.000577018 | 0.044146471 |
| *INKA1* | 0.485292 | 0.141294 | 3.434636 | 0.00059335 | 0.045038523 |
| *CHP1* | -0.296851 | 0.086627 | -3.426761 | 0.000610828 | 0.046002948 |
| *GPAA1* | -0.415578 | 0.121522 | -3.419764 | 0.000626756 | 0.046253119 |
| *ZCCHC9* | 0.404566 | 0.118308 | 3.419601 | 0.00062713 | 0.046253119 |

| | | | | | |
|---|---|---|---|---|---|
| *CD5* | -1.268106 | 0.370901 | -3.418989 | 0.000628543 | 0.046253119 |
| *AZIN1.AS1* | -1.860333 | 0.547489 | -3.397937 | 0.00067896 | NA |
| *A4GALT* | 1.296368 | 0.38187 | 3.394786 | 0.000686822 | 0.049948755 |
| *CDCA4* | 0.4629 | 0.13641 | 3.393437 | 0.000690214 | 0.049948755 |
| *TAPBP* | -0.249669 | 0.073609 | -3.391817 | 0.000694308 | 0.049948755 |
| *NR6A1* | -1.036896 | 0.306379 | -3.384359 | 0.000713448 | 0.050945455 |
| *LINC02422* | 0.665729 | 0.197012 | 3.37913 | 0.000727156 | 0.051542512 |
| *LINC00525* | -2.81289 | 0.834013 | -3.372718 | 0.000744302 | NA |
| *BHLHE40* | -0.488228 | 0.14477 | -3.372431 | 0.000745078 | 0.052086733 |
| *ENSG00000173727* | 0.296929 | 0.088051 | 3.372223 | 0.00074564 | 0.052086733 |
| *RCC1* | 0.475178 | 0.141189 | 3.365539 | 0.000763942 | 0.052981283 |
| *PFKL* | -0.330825 | 0.098442 | -3.36061 | 0.000777705 | 0.053550518 |
| *DSCAML1* | -1.500081 | 0.446759 | -3.357695 | 0.000785954 | NA |
| *HSPE1* | 0.232143 | 0.069277 | 3.350959 | 0.000805321 | 0.054759772 |
| *MIR181A1HG* | -0.573768 | 0.171248 | -3.350511 | 0.000806627 | 0.054759772 |
| *TEX30* | 0.492295 | 0.147091 | 3.346862 | 0.000817319 | 0.055097555 |
| *ENSG00000257275* | -0.606902 | 0.181441 | -3.344906 | 0.000823104 | 0.055102233 |
| *ENSG00000263394* | 0.715972 | 0.214804 | 3.333141 | 0.000858714 | 0.057089679 |
| *PNRC2* | 0.258484 | 0.07767 | 3.328001 | 0.000874716 | 0.05775521 |
| *MANF* | 0.374779 | 0.11285 | 3.321036 | 0.00089684 | 0.058813208 |
| *ODF3B* | -0.731436 | 0.220995 | -3.30974 | 0.000933826 | 0.060824852 |
| *PIM1* | -0.361973 | 0.10956 | -3.303873 | 0.00095359 | 0.061695357 |
| *ENSG00000185527* | -0.997815 | 0.30221 | -3.301733 | 0.000960896 | NA |
| *LARP4B* | 0.452635 | 0.137148 | 3.300344 | 0.000965663 | 0.062059912 |
| *DPP7* | -0.353173 | 0.107489 | -3.285661 | 0.001017433 | 0.064697222 |
| *HIPK2* | 0.589507 | 0.179459 | 3.284918 | 0.001020122 | 0.064697222 |
| *EHD1* | -0.370326 | 0.112806 | -3.282847 | 0.001027645 | 0.064748329 |
| *NUAK2* | -0.335724 | 0.102562 | -3.273388 | 0.001062666 | 0.06618916 |
| *LAPTM5* | -0.335023 | 0.102361 | -3.272968 | 0.001064245 | 0.06618916 |
| *ENSG00000259976* | 0.656715 | 0.200787 | 3.2707 | 0.001072815 | 0.066294474 |
| *TMEM138* | -0.295569 | 0.090478 | -3.266734 | 0.001087959 | 0.066301985 |
| *IL4R* | -0.475145 | 0.145487 | -3.265901 | 0.001091163 | 0.066301985 |
| *ROCK1* | 0.234812 | 0.071912 | 3.265277 | 0.00109357 | 0.066301985 |
| *ABHD17A* | 0.437069 | 0.134055 | 3.260384 | 0.001112615 | 0.06703507 |
| *ENSG00000254614* | -0.853697 | 0.262221 | -3.255638 | 0.001131379 | 0.06732656 |
| *CD48* | 0.354668 | 0.10894 | 3.255627 | 0.001131421 | 0.06732656 |
| *MCCC2* | -0.485503 | 0.149513 | -3.247231 | 0.001165336 | 0.068919278 |
| *MYO1C* | -0.404901 | 0.124832 | -3.243563 | 0.001180449 | 0.069280545 |
| *MAP3K9* | 0.776167 | 0.239471 | 3.241173 | 0.001190388 | 0.069280545 |
| *FAM184B* | -1.726672 | 0.532834 | -3.240547 | 0.001193005 | 0.069280545 |
| *LSM6* | 0.344573 | 0.106648 | 3.230945 | 0.001233817 | 0.071221538 |
| *IFIT2* | 0.615786 | 0.190877 | 3.226088 | 0.001254949 | 0.072010193 |
| *ZNF350* | -0.306739 | 0.095222 | -3.221302 | 0.001276098 | 0.072431836 |
| *ADA* | -0.551165 | 0.171115 | -3.221026 | 0.001277325 | 0.072431836 |
| *ZNF267* | -0.269864 | 0.084063 | -3.210256 | 0.001326168 | 0.074761735 |
| *KIAA0930* | 0.857553 | 0.26737 | 3.207363 | 0.001339579 | 0.075078709 |
| *UBAP1L* | 1.048304 | 0.327239 | 3.203487 | 0.00135774 | NA |

| | | | | | |
|---|---|---|---|---|---|
| *PRICKLE1* | 0.606399 | 0.189304 | 3.203311 | 0.001358571 | 0.07570304 |
| *HCK* | -0.748083 | 0.233762 | -3.200183 | 0.001373403 | 0.076089706 |
| *TBC1D9* | 0.538531 | 0.168391 | 3.198094 | 0.001383389 | 0.076204995 |
| *RUBCNL* | -0.305857 | 0.095794 | -3.192875 | 0.001408639 | 0.077155005 |
| *SSPN* | 2.283536 | 0.71591 | 3.189698 | 0.001424214 | NA |
| *ROR1* | -1.163179 | 0.365244 | -3.184662 | 0.001449232 | 0.078929939 |
| *CDKN1A* | -0.362959 | 0.114067 | -3.181968 | 0.001462782 | 0.079099654 |
| *INPP5K* | -0.435088 | 0.136786 | -3.180786 | 0.001468759 | 0.079099654 |
| *GLRX* | 0.287717 | 0.090509 | 3.178872 | 0.001478491 | 0.079181424 |
| *ENSG00000225302* | 2.371706 | 0.746567 | 3.176815 | 0.001489021 | NA |
| *C7orf50* | -0.5609 | 0.176716 | -3.174025 | 0.001503406 | 0.080012062 |
| *UGT8* | 0.813983 | 0.256563 | 3.172639 | 0.001510601 | 0.080012062 |
| *ZNF726* | -1.094163 | 0.345157 | -3.170043 | 0.001524166 | NA |
| *GPS2* | -0.339433 | 0.107123 | -3.168633 | 0.001531575 | 0.08067971 |
| *IL23A* | -0.669339 | 0.211622 | -3.1629 | 0.001562061 | 0.081838415 |
| *MED21* | 0.416483 | 0.131915 | 3.157196 | 0.001592945 | 0.083005336 |
| *FAM41C* | -0.705204 | 0.223537 | -3.154757 | 0.00160632 | 0.083252289 |
| *ITGB7* | 0.371445 | 0.118449 | 3.135913 | 0.001713201 | 0.086683873 |
| *APOL6* | 0.86288 | 0.275165 | 3.135862 | 0.001713497 | 0.086683873 |
| *ZNF580* | -0.317757 | 0.101373 | -3.134528 | 0.001721306 | 0.086683873 |
| *GPATCH11* | 0.466294 | 0.148782 | 3.134069 | 0.001724001 | 0.086683873 |
| *DDX5* | -0.371748 | 0.118643 | -3.133326 | 0.001728372 | 0.086683873 |
| *SPINT2* | -0.227372 | 0.072611 | -3.131369 | 0.001739931 | 0.086683873 |
| *TSC22D2* | 0.385875 | 0.123275 | 3.130192 | 0.001746921 | 0.086683873 |
| *ENSG00000166927* | -0.762691 | 0.243658 | -3.130165 | 0.001747084 | 0.086683873 |
| *SUSD3* | 0.612465 | 0.195732 | 3.129094 | 0.00175346 | 0.086683873 |
| *LYRM7* | 0.292592 | 0.093577 | 3.126742 | 0.001767551 | 0.086934662 |
| *HMGXB4* | -0.300102 | 0.096115 | -3.122313 | 0.001794359 | 0.087805176 |
| *PRCP* | -0.396809 | 0.12773 | -3.106619 | 0.001892401 | 0.092135083 |
| *TMEM187* | 0.755501 | 0.243515 | 3.102482 | 0.001919055 | 0.092587706 |
| *MAGEE1* | 1.023203 | 0.329909 | 3.101471 | 0.001925618 | NA |
| *ZNF486* | 0.937018 | 0.302243 | 3.100209 | 0.001933841 | 0.092587706 |
| *ENSG00000258757* | 0.735415 | 0.237259 | 3.099625 | 0.00193766 | 0.092587706 |
| *FAM200A* | 0.57884 | 0.186794 | 3.09882 | 0.001942931 | 0.092587706 |
| *TMEM64* | 0.666633 | 0.215223 | 3.097402 | 0.001952251 | 0.092587706 |
| *NSUN7* | -1.029441 | 0.332472 | -3.09633 | 0.001959325 | 0.092587706 |
| *IFI16* | 0.194621 | 0.062941 | 3.092105 | 0.001987426 | 0.093457485 |
| *YTHDC1* | 0.21842 | 0.07073 | 3.088103 | 0.002014387 | 0.093629524 |
| *CLK1* | 0.259635 | 0.084148 | 3.08545 | 0.002032441 | 0.093629524 |
| *PPP1R9A* | 1.528429 | 0.495517 | 3.084515 | 0.002038841 | NA |
| *CHTOP* | 0.260678 | 0.08453 | 3.083866 | 0.002043296 | 0.093629524 |
| *IGHG1* | 1.273043 | 0.412811 | 3.083838 | 0.002043487 | 0.093629524 |
| *DDAH2* | 0.366539 | 0.11886 | 3.083773 | 0.002043936 | 0.093629524 |
| *EPS15* | -0.428854 | 0.139104 | -3.082984 | 0.00204936 | 0.093629524 |
| *USP28* | -0.359864 | 0.116801 | -3.081007 | 0.002063021 | 0.093809079 |
| *TNFRSF13B* | 1.121829 | 0.364598 | 3.076896 | 0.002091686 | 0.094069044 |
| *DDX19B* | -0.330851 | 0.107535 | -3.076679 | 0.002093209 | 0.094069044 |

| | | | | | |
|---|---|---|---|---|---|
| *SF3B4* | -0.398188 | 0.129498 | -3.074844 | 0.002106127 | 0.094069044 |
| *RRM2B* | -0.383902 | 0.124862 | -3.074611 | 0.002107771 | 0.094069044 |
| *ALOX5* | -0.300147 | 0.097738 | -3.070925 | 0.002133968 | 0.094799334 |
| *NSMAF* | 0.464512 | 0.151868 | 3.058661 | 0.002223288 | 0.098314225 |
| *ENSG00000272172* | -1.522937 | 0.498043 | -3.057843 | 0.002229361 | NA |
| *INAVA* | -2.995991 | 0.980522 | -3.055507 | 0.002246801 | NA |
| *PDE4B* | 0.211589 | 0.069285 | 3.053884 | 0.002258996 | 0.09847402 |
| *CABLES1* | -0.693697 | 0.227169 | -3.053662 | 0.002260663 | 0.09847402 |
| *KPNA2* | -0.44671 | 0.146368 | -3.051958 | 0.002273537 | 0.09847402 |
| *ELP3* | -0.563254 | 0.184604 | -3.051148 | 0.002279684 | 0.09847402 |
| *MFSD13A* | -0.70715 | 0.231806 | -3.050618 | 0.002283711 | 0.09847402 |
| *TNFAIP3* | -0.63185 | 0.207162 | -3.050029 | 0.002288193 | 0.09847402 |
| *RAB9A* | 0.407206 | 0.133711 | 3.045423 | 0.002323533 | 0.099204193 |
| *AASDH* | 0.391809 | 0.128667 | 3.045137 | 0.002325741 | 0.099204193 |
| *TAF11* | 0.312455 | 0.102692 | 3.042659 | 0.002344981 | 0.099584215 |
| *PFKFB3* | -0.490162 | 0.161224 | -3.040247 | 0.002363844 | 0.099868223 |
| *STX7* | 0.237964 | 0.078299 | 3.03916 | 0.002372388 | 0.099868223 |
| *TMC6* | -0.288386 | 0.094932 | -3.037808 | 0.002383058 | 0.099881219 |
| *SPPL2A* | -0.3066 | 0.10104 | -3.034443 | 0.0024098 | 0.099921447 |
| *ZNF860* | 0.519527 | 0.17121 | 3.034437 | 0.002409853 | 0.099921447 |
| *PNPLA8* | 0.268386 | 0.088466 | 3.033779 | 0.002415114 | 0.099921447 |
| *SARS2* | -0.697553 | 0.230052 | -3.032155 | 0.002428146 | 0.100031305 |
| *TAGAP* | -0.492943 | 0.16273 | -3.029217 | 0.002451886 | 0.100087824 |
| *ENSG00000237491* | -0.671006 | 0.221545 | -3.028755 | 0.002455634 | 0.100087824 |
| *NOP16* | 0.466188 | 0.153969 | 3.027796 | 0.00246344 | 0.100087824 |
| *ADRB2* | 0.823436 | 0.27207 | 3.026559 | 0.002473549 | 0.100087824 |
| *GPR18* | 0.629552 | 0.208075 | 3.025597 | 0.00248143 | 0.100087824 |
| *SERPINB6* | 0.501514 | 0.166032 | 3.020582 | 0.002522896 | 0.101336335 |
| *HLA.C* | -0.485151 | 0.160839 | -3.016376 | 0.002558159 | 0.101919252 |
| *ACAP1* | 0.293331 | 0.097248 | 3.016329 | 0.002558554 | 0.101919252 |
| *PTPN20* | -1.630377 | 0.541231 | -3.01235 | 0.002592333 | NA |
| *MREG* | -0.624325 | 0.2077 | -3.005905 | 0.002647915 | 0.105044848 |
| *MINDY2* | 0.378593 | 0.126093 | 3.002486 | 0.002677843 | 0.105796765 |
| *PPM1K* | 0.22176 | 0.073958 | 2.998476 | 0.002713335 | 0.106761407 |
| *RBM15* | 0.402534 | 0.134337 | 2.996445 | 0.002731474 | 0.107038268 |
| *REEP3* | 0.452765 | 0.151178 | 2.994911 | 0.002745249 | 0.107142494 |
| *THG1L* | 0.570651 | 0.190701 | 2.992388 | 0.002768045 | 0.107596593 |
| *CD27* | 1.127606 | 0.377183 | 2.989546 | 0.002793923 | 0.108166345 |
| *CCDC78* | -1.867102 | 0.625101 | -2.986881 | 0.002818397 | NA |
| *DUSP12* | -0.310487 | 0.104004 | -2.985341 | 0.002832624 | 0.109091027 |
| *CARD8* | -0.276231 | 0.092591 | -2.983338 | 0.002851232 | 0.109091027 |
| *P2RY8* | -0.336082 | 0.112655 | -2.983281 | 0.002851757 | 0.109091027 |
| *SPOP* | 0.259784 | 0.087273 | 2.976699 | 0.002913699 | 0.111019975 |
| *NMT2* | -0.382223 | 0.128459 | -2.975449 | 0.002925599 | 0.111034548 |
| *LILRB1* | 0.393952 | 0.132513 | 2.972941 | 0.002949607 | 0.111304639 |
| *SLC25A26* | -0.264806 | 0.089138 | -2.970753 | 0.002970705 | 0.111304639 |
| *STN1* | 0.44127 | 0.148579 | 2.969943 | 0.002978551 | 0.111304639 |

| | | | | | |
|---|---|---|---|---|---|
| *TRAF3* | -0.261284 | 0.087977 | -2.969907 | 0.0029789 | 0.111304639 |
| *LY6E* | 0.349634 | 0.117799 | 2.968055 | 0.002996906 | 0.111545056 |
| *POLE3* | -0.271571 | 0.091552 | -2.966306 | 0.003014005 | 0.111750015 |
| *MRPL54* | -0.268261 | 0.09048 | -2.96487 | 0.003028105 | 0.11184267 |
| *ENSG00000259001* | 1.094851 | 0.369705 | 2.961419 | 0.003062252 | NA |
| *HVCN1* | -0.396961 | 0.134054 | -2.961197 | 0.00306446 | 0.112451453 |
| *EIF2AK4* | -0.44711 | 0.15105 | -2.96001 | 0.003076291 | 0.112451453 |
| *SERTAD2* | 0.351269 | 0.118685 | 2.95968 | 0.003079583 | 0.112451453 |
| *MED11* | 0.223049 | 0.075395 | 2.958423 | 0.003092175 | 0.112485168 |
| *HLA.A* | -0.469464 | 0.158929 | -2.953931 | 0.003137545 | 0.11370652 |
| *ARCN1* | 0.413952 | 0.140445 | 2.947435 | 0.003204226 | 0.115688147 |
| *TMEM120A* | -0.470106 | 0.159661 | -2.944412 | 0.003235694 | 0.116388398 |
| *TMEM70* | 0.349462 | 0.118898 | 2.939169 | 0.003290933 | 0.117824309 |
| *ARHGEF3* | -0.342433 | 0.116541 | -2.938311 | 0.003300058 | 0.117824309 |
| *ENSG00000278384* | 1.814486 | 0.617706 | 2.937459 | 0.003309138 | NA |
| *ENSG00000267002* | 0.31354 | 0.106783 | 2.936251 | 0.003322056 | 0.117866197 |
| *LTBP3* | 0.310247 | 0.105697 | 2.935257 | 0.003332711 | 0.117866197 |
| *DHRS4L2* | 0.450895 | 0.153639 | 2.934774 | 0.003337912 | 0.117866197 |
| *TOE1* | -0.430269 | 0.147113 | -2.924748 | 0.003447357 | 0.120829305 |
| *SLA* | 0.375959 | 0.128603 | 2.923413 | 0.003462167 | 0.120829305 |
| *SNX8* | -0.357695 | 0.122398 | -2.922396 | 0.003473492 | 0.120829305 |
| *ARSD* | 0.711152 | 0.243472 | 2.920876 | 0.003490489 | 0.120829305 |
| *C21orf62.AS1* | 1.057755 | 0.362201 | 2.920354 | 0.003496341 | NA |
| *SMIM4* | -0.406814 | 0.139398 | -2.918367 | 0.003518701 | 0.120829305 |
| *TRIM56* | 0.225729 | 0.077362 | 2.917835 | 0.003524706 | 0.120829305 |
| *AK6* | 0.355372 | 0.12185 | 2.91648 | 0.003540057 | 0.120829305 |
| *FAM120A* | 0.336977 | 0.115592 | 2.915224 | 0.003554339 | 0.120829305 |
| *FAM102A* | 0.592718 | 0.203328 | 2.915086 | 0.003555901 | 0.120829305 |
| *FNIP1* | -0.382095 | 0.1311 | -2.91453 | 0.00356225 | 0.120829305 |
| *HIST1H3E* | 0.754127 | 0.258977 | 2.911946 | 0.003591851 | 0.120829305 |
| *GATM* | 0.938007 | 0.322181 | 2.911428 | 0.003597809 | 0.120829305 |
| *FGR* | -0.735434 | 0.25274 | -2.909841 | 0.003616123 | 0.120829305 |
| *NBPF9* | 0.608436 | 0.20912 | 2.909505 | 0.003620012 | 0.120829305 |
| *RABEP2* | 0.250293 | 0.08603 | 2.909379 | 0.00362147 | 0.120829305 |
| *PARP9* | 0.387314 | 0.133166 | 2.908507 | 0.003631593 | 0.120829305 |
| *RBAK.RBAKDN* | 0.35122 | 0.120768 | 2.908222 | 0.003634906 | 0.120829305 |
| *ENSG00000265218* | -2.08549 | 0.717473 | -2.906715 | 0.00365246 | NA |
| *RAB3GAP1* | -0.312244 | 0.107533 | -2.903702 | 0.003687794 | 0.121428324 |
| *TACC1* | 0.361836 | 0.124622 | 2.903472 | 0.003690498 | 0.121428324 |
| *TYSND1* | -0.514334 | 0.177169 | -2.903069 | 0.003695253 | 0.121428324 |
| *TPMT* | -0.477976 | 0.164781 | -2.900684 | 0.003723488 | 0.121428324 |
| *GM2A* | 0.543719 | 0.187464 | 2.900385 | 0.003727047 | 0.121428324 |
| *BAG2* | 0.527044 | 0.181723 | 2.900262 | 0.003728505 | 0.121428324 |
| *RMND5A* | 0.453612 | 0.156587 | 2.896867 | 0.003769096 | 0.122202518 |
| *USP53* | 0.768912 | 0.265494 | 2.896157 | 0.00377763 | 0.122202518 |
| *ZNF667.AS1* | 0.422151 | 0.145846 | 2.894491 | 0.003797735 | 0.122442026 |
| *FADS3* | -0.26965 | 0.093264 | -2.891248 | 0.003837147 | 0.123300334 |

| | | | | | |
|---|---|---|---|---|---|
| *CALM3* | -0.284303 | 0.098426 | -2.888485 | 0.003871022 | 0.1239756 |
| *FCRL4* | 1.750294 | 0.606847 | 2.88424 | 0.00392359 | NA |
| *CHCHD5* | -0.307109 | 0.106546 | -2.882412 | 0.003946439 | 0.125581419 |
| *SUN1* | -0.310729 | 0.107804 | -2.882349 | 0.003947217 | 0.125581419 |
| *SLC25A43* | 0.424961 | 0.147492 | 2.881253 | 0.003960978 | 0.125604682 |
| *KLB* | -2.062507 | 0.715961 | -2.880753 | 0.003967258 | NA |
| *CCDC51* | 0.50831 | 0.176623 | 2.877944 | 0.00400276 | 0.12596178 |
| *DEDD2* | 0.290403 | 0.100917 | 2.877642 | 0.004006593 | 0.12596178 |
| *METTL7A* | 0.298127 | 0.103615 | 2.877261 | 0.004011438 | 0.12596178 |
| *CRTC2* | -0.32478 | 0.11297 | -2.874927 | 0.004041209 | 0.126484603 |
| *FAM122A* | 0.301002 | 0.104818 | 2.871672 | 0.00408306 | 0.12738091 |
| *AMD1* | 0.345662 | 0.120452 | 2.869705 | 0.00410855 | 0.127762667 |
| *PILRB* | 0.481528 | 0.167941 | 2.867238 | 0.004140713 | 0.128348791 |
| *THRB* | 1.669661 | 0.58237 | 2.867013 | 0.00414366 | NA |
| *MAPRE2* | 0.397558 | 0.138745 | 2.865385 | 0.00416502 | 0.128688426 |
| *CREB3L2* | -0.395342 | 0.138107 | -2.862579 | 0.004202091 | 0.129419042 |
| *RBIS* | 0.239542 | 0.083716 | 2.86137 | 0.00421814 | 0.129499579 |
| *PEX16* | -0.231391 | 0.080922 | -2.859419 | 0.004244181 | 0.129785168 |
| *ENSG00000280433* | -0.838318 | 0.293181 | -2.859384 | 0.004244646 | NA |
| *CCDC126* | 0.484819 | 0.169597 | 2.858658 | 0.004254369 | 0.129785168 |
| *C9orf72* | -0.513412 | 0.18012 | -2.850396 | 0.004366479 | 0.13240725 |
| *ZBTB3* | -0.732261 | 0.256988 | -2.849401 | 0.004380162 | 0.13240725 |
| *WDR55* | -0.324152 | 0.113765 | -2.849302 | 0.004381526 | 0.13240725 |
| *PHF23* | 0.296513 | 0.104144 | 2.847154 | 0.004411208 | 0.13288765 |
| *ZNF330* | 0.363477 | 0.127755 | 2.845117 | 0.004439505 | 0.133323439 |
| *CNTRL* | 0.210615 | 0.074067 | 2.84359 | 0.004460842 | 0.133341068 |
| *SPATA24* | -0.667122 | 0.234646 | -2.843096 | 0.004467756 | 0.133341068 |
| *ALOX12.AS1* | 0.358796 | 0.126298 | 2.84086 | 0.004499204 | 0.133865192 |
| *ZNF835* | 0.889371 | 0.313136 | 2.840208 | 0.004508408 | NA |
| *ATP6V0A1* | 0.432192 | 0.152457 | 2.83484 | 0.004584871 | 0.135994324 |
| *BCL2L11* | -0.615994 | 0.21762 | -2.830595 | 0.00464615 | 0.137185549 |
| *CD79B* | -0.233717 | 0.082613 | -2.829057 | 0.004668536 | 0.137185549 |
| *POLR2L* | 0.289306 | 0.102304 | 2.827896 | 0.004685509 | 0.137185549 |
| *MTHFR* | 0.403748 | 0.142791 | 2.827539 | 0.00469073 | 0.137185549 |
| *HRK* | -1.662404 | 0.588011 | -2.827167 | 0.004696186 | 0.137185549 |
| *ZNF827* | 0.796675 | 0.281891 | 2.826177 | 0.004710719 | 0.137194357 |
| *TMEM205* | -0.316397 | 0.112002 | -2.824914 | 0.004729325 | 0.137321376 |
| *CAV1* | -0.639229 | 0.226838 | -2.818 | 0.004832386 | 0.139332861 |
| *AK1* | -0.580421 | 0.206047 | -2.816937 | 0.004848398 | 0.139332861 |
| *TMEM19* | 0.238673 | 0.084792 | 2.814809 | 0.004880617 | 0.139332861 |
| *DBNL* | -0.252878 | 0.089842 | -2.81471 | 0.004882125 | 0.139332861 |
| *ENSG00000275464* | -0.898276 | 0.31916 | -2.814502 | 0.00488529 | 0.139332861 |
| *LIMD2* | -0.229136 | 0.081413 | -2.8145 | 0.004885322 | 0.139332861 |
| *MACF1* | 0.297323 | 0.105831 | 2.809415 | 0.004963163 | 0.141135363 |
| *ENSG00000279483* | -0.30216 | 0.107605 | -2.808062 | 0.004984062 | 0.141312825 |
| *UPB1* | -1.49338 | 0.532397 | -2.805013 | 0.005031453 | NA |
| *ZNF391* | 1.351697 | 0.48246 | 2.801676 | 0.005083794 | NA |

| | | | | | |
|---|---|---|---|---|---|
| *ENSG00000261786* | 2.142904 | 0.766699 | 2.794975 | 0.005190376 | NA |
| *GRN* | -0.285108 | 0.10209 | -2.79271 | 0.005226847 | 0.147761898 |
| *TTC39B* | -1.214583 | 0.435049 | -2.791831 | 0.005241073 | NA |
| *NDUFV3* | 0.29424 | 0.105436 | 2.790686 | 0.005259652 | 0.148254514 |
| *FRY.AS1* | -1.11033 | 0.398511 | -2.786196 | 0.005333066 | NA |
| *AAK1* | -0.665325 | 0.238825 | -2.785828 | 0.005339126 | 0.14968864 |
| *ENSG00000270562* | 0.63918 | 0.229452 | 2.785678 | 0.005341586 | 0.14968864 |
| *TAF1A* | 0.480443 | 0.17253 | 2.784689 | 0.005357902 | 0.149710648 |
| *ENSG00000277369* | -2.602331 | 0.934549 | -2.784584 | 0.005359636 | NA |
| *RHBDD2* | -0.354405 | 0.127361 | -2.782689 | 0.005391038 | 0.150201165 |
| *CARMIL1* | -1.004674 | 0.361598 | -2.778427 | 0.005462276 | 0.150764885 |
| *RPL8* | 0.30357 | 0.109287 | 2.777734 | 0.005473946 | 0.150764885 |
| *CMTM7* | -0.395453 | 0.142388 | -2.77728 | 0.005481602 | 0.150764885 |
| *CKS1B* | 0.233624 | 0.084133 | 2.776843 | 0.005488961 | 0.150764885 |
| *FOXO3* | 0.473644 | 0.170647 | 2.775585 | 0.00551025 | 0.150764885 |
| *ERGIC2* | 0.232212 | 0.083678 | 2.775074 | 0.005518907 | 0.150764885 |
| *ENSG00000268400* | -1.32119 | 0.476102 | -2.775013 | 0.005519944 | NA |
| *MED13* | 0.342148 | 0.123298 | 2.774966 | 0.005520747 | 0.150764885 |
| *MYL12A* | 0.199378 | 0.071985 | 2.769706 | 0.005610697 | 0.151881812 |
| *BNIP3* | 0.298852 | 0.107904 | 2.769613 | 0.005612295 | 0.151881812 |
| *PMVK* | 0.297232 | 0.107322 | 2.769544 | 0.005613481 | 0.151881812 |
| *VPREB1* | -1.621823 | 0.585841 | -2.768369 | 0.005633758 | 0.151881812 |
| *ZNF470* | 0.772951 | 0.279252 | 2.767936 | 0.005641259 | 0.151881812 |
| *UBC* | -0.366361 | 0.132418 | -2.766711 | 0.005662497 | 0.151881812 |
| *GNG11* | 1.085592 | 0.392394 | 2.766589 | 0.005664604 | NA |
| *DDB2* | -0.283271 | 0.102416 | -2.765895 | 0.005676684 | 0.151881812 |
| *EIF5* | 0.192797 | 0.069721 | 2.765263 | 0.00568769 | 0.151881812 |
| *PROC* | 1.354931 | 0.489999 | 2.765169 | 0.00568933 | NA |
| *ZC3H7A* | -0.413084 | 0.149472 | -2.763629 | 0.005716257 | 0.152222973 |
| *SGMS1.AS1* | -0.686578 | 0.248719 | -2.760457 | 0.005772061 | 0.153285594 |
| *C3orf14* | 1.848453 | 0.66969 | 2.760162 | 0.005777278 | NA |
| *ZNF628* | -0.862717 | 0.31274 | -2.758579 | 0.005805333 | 0.153441072 |
| *ARL4C* | -0.632539 | 0.229319 | -2.75833 | 0.00580975 | 0.153441072 |
| *GOPC* | 0.338593 | 0.122893 | 2.755183 | 0.005865935 | 0.154501665 |
| *CNOT1* | -0.261178 | 0.094876 | -2.75283 | 0.005908254 | 0.155122606 |
| *NELFE* | -0.212901 | 0.07736 | -2.752086 | 0.005921693 | 0.155122606 |
| *ENSG00000277476* | 0.964681 | 0.350569 | 2.751755 | 0.005927678 | NA |
| *NIPAL3* | 0.400066 | 0.145436 | 2.750807 | 0.005944872 | 0.155304347 |
| *BLOC1S3* | 0.407396 | 0.148148 | 2.749927 | 0.005960852 | 0.155304347 |
| *FGFR1OP2* | 0.268088 | 0.097523 | 2.748982 | 0.00597806 | 0.155332886 |
| *PM20D2* | 0.520528 | 0.189872 | 2.741464 | 0.006116615 | 0.158505822 |
| *MED23* | -0.382389 | 0.139704 | -2.73714 | 0.006197593 | 0.159948626 |
| *ELOA* | 0.297589 | 0.108739 | 2.736722 | 0.006205476 | 0.159948626 |
| *APOBEC3H* | -0.973374 | 0.355874 | -2.735169 | 0.006234825 | 0.160276579 |
| *CTDSP2* | 0.281041 | 0.10285 | 2.732549 | 0.006284639 | 0.160817091 |
| *SNHG9* | -0.365774 | 0.133886 | -2.731975 | 0.006295587 | 0.160817091 |
| *UPP1* | 0.655629 | 0.240031 | 2.731436 | 0.006305898 | 0.160817091 |

| | | | | | |
|---|---|---|---|---|---|
| *ENSG00000007237* | -0.791293 | 0.289908 | -2.729459 | 0.006343836 | 0.16132299 |
| *ENSG00000228463* | 0.413991 | 0.151719 | 2.728661 | 0.006359205 | 0.16132299 |
| *ENSG00000245904* | 0.289023 | 0.105958 | 2.727705 | 0.006377659 | 0.161366498 |
| *ELMSAN1* | 0.322584 | 0.118312 | 2.726555 | 0.006399931 | 0.161506109 |
| *CHID1* | -0.374345 | 0.137476 | -2.722976 | 0.00646967 | 0.162839735 |
| *ERRFI1* | 1.42442 | 0.523333 | 2.721822 | 0.006492319 | NA |
| *ENSG00000273748* | -0.934016 | 0.343283 | -2.720833 | 0.00651176 | 0.163325071 |
| *ENSG00000244701* | -0.563501 | 0.207149 | -2.720271 | 0.006522837 | 0.163325071 |
| *GBP2* | 0.351092 | 0.12911 | 2.719322 | 0.006541584 | 0.163370117 |
| *LINC00638* | 1.186595 | 0.436454 | 2.718717 | 0.006553575 | NA |
| *SEL1L3* | 0.354316 | 0.130521 | 2.714628 | 0.00663503 | 0.165275683 |
| *GCNT1* | 0.701622 | 0.258602 | 2.713139 | 0.006664906 | 0.165591997 |
| *SF1* | -0.137746 | 0.050819 | -2.710519 | 0.006717791 | 0.166372407 |
| *FCGR2B* | 0.713676 | 0.263528 | 2.708157 | 0.006765809 | 0.166372407 |
| *TMED8* | 0.355321 | 0.131225 | 2.707728 | 0.006774553 | 0.166372407 |
| *GSTO1* | 0.261215 | 0.096472 | 2.707666 | 0.006775813 | 0.166372407 |
| *TNFRSF12A* | 1.066839 | 0.394055 | 2.707333 | 0.00678261 | 0.166372407 |
| *BBX* | 0.201828 | 0.074594 | 2.705692 | 0.006816214 | 0.166772342 |
| *ZNF419* | 0.427381 | 0.158098 | 2.703273 | 0.006866026 | 0.167565802 |
| *DNAJA1* | 0.268408 | 0.099328 | 2.702249 | 0.006887206 | 0.167658254 |
| *TBPL1* | 0.232037 | 0.086031 | 2.697136 | 0.006993877 | 0.169354717 |
| *PTGIR* | -1.054023 | 0.390854 | -2.696721 | 0.007002587 | 0.169354717 |
| *PHKG1* | -0.432647 | 0.160475 | -2.696044 | 0.007016835 | 0.169354717 |
| *TOB2* | 0.267048 | 0.09907 | 2.695554 | 0.007027167 | 0.169354717 |
| *FBXW9* | -0.89046 | 0.330713 | -2.692547 | 0.007090844 | NA |
| *GPM6B* | 0.304518 | 0.113204 | 2.689998 | 0.007145242 | 0.171770916 |
| *EIF3J.DT* | 0.257369 | 0.095724 | 2.688668 | 0.007173775 | 0.171940186 |
| *LINC01355* | 0.5143 | 0.191331 | 2.688009 | 0.007187956 | 0.171940186 |
| *FAM78A* | 0.390355 | 0.145323 | 2.686127 | 0.007228566 | 0.172218935 |
| *PRDX3* | 0.241097 | 0.089799 | 2.684861 | 0.007256004 | 0.172218935 |
| *BYSL* | 0.684816 | 0.255122 | 2.684265 | 0.007268941 | 0.172218935 |
| *SDE2* | -0.507355 | 0.189065 | -2.683502 | 0.007285561 | 0.172218935 |
| *ENSG00000272831* | -0.429619 | 0.160106 | -2.683347 | 0.007288934 | 0.172218935 |
| *DDX49* | -0.382366 | 0.142589 | -2.681598 | 0.007327135 | 0.17269825 |
| *GTF2IRD2* | -0.384834 | 0.143592 | -2.680051 | 0.0073611 | 0.173075624 |
| *CDC26* | 0.238739 | 0.089379 | 2.671099 | 0.007560341 | 0.177327716 |
| *ZRANB3* | -0.634184 | 0.237575 | -2.669402 | 0.007598646 | 0.177703698 |
| *FXYD2* | -0.772116 | 0.289317 | -2.668758 | 0.007613239 | 0.177703698 |
| *UNC45A* | -0.282765 | 0.106014 | -2.667233 | 0.007647863 | 0.178080686 |
| *PPBP* | 2.854967 | 1.071457 | 2.664564 | NA | NA |
| *DHX30* | -0.26267 | 0.098718 | -2.66082 | 0.007795052 | 0.181070595 |
| *DDX6* | -0.230462 | 0.086654 | -2.659576 | 0.007823908 | 0.18130403 |
| *TERF2IP* | 0.22002 | 0.08286 | 2.655332 | 0.007923045 | 0.182819627 |
| *YTHDF3* | 0.412165 | 0.155232 | 2.655153 | 0.007927241 | 0.182819627 |
| *PIGT* | -0.329406 | 0.124161 | -2.65306 | 0.00797657 | 0.183518229 |
| *RTP4* | 0.609309 | 0.229788 | 2.65161 | 0.008010899 | 0.183869208 |
| *NCBP2* | -0.210297 | 0.079348 | -2.650317 | 0.008041635 | 0.184136245 |

| | | | | | |
|---|---|---|---|---|---|
| *UPF2* | -0.138622 | 0.052323 | -2.649365 | 0.008064314 | 0.184217987 |
| *HDDC2* | -0.253771 | 0.095854 | -2.647463 | 0.008109829 | 0.184423313 |
| *GOLGA7* | -0.247379 | 0.093443 | -2.64739 | 0.008111565 | 0.184423313 |
| *DPH5* | 0.300297 | 0.113551 | 2.64461 | 0.008178521 | 0.185508094 |
| *ZNF473* | 0.680745 | 0.257575 | 2.642903 | 0.008219867 | 0.186008262 |
| *TRMT112* | 0.218873 | 0.0829 | 2.640207 | 0.008285546 | 0.186570187 |
| *GID8* | -0.227424 | 0.086161 | -2.639523 | 0.008302291 | 0.186570187 |
| *TSHZ1* | 0.676657 | 0.256429 | 2.638772 | 0.008320696 | 0.186570187 |
| *NOL12* | -0.393043 | 0.149021 | -2.637496 | 0.008352062 | 0.186570187 |
| *IGHA1* | -0.452862 | 0.171704 | -2.637449 | 0.008353209 | 0.186570187 |
| *C16orf74* | -0.406055 | 0.153976 | -2.63714 | 0.008360822 | 0.186570187 |
| *RNF19A* | 0.348502 | 0.13225 | 2.635182 | 0.008409212 | 0.186855397 |
| *UCP2* | -0.259322 | 0.098412 | -2.635055 | 0.00841237 | 0.186855397 |
| *ZNF784* | -0.6876 | 0.26122 | -2.632263 | 0.008481809 | 0.187559523 |
| *BARD1* | 0.367529 | 0.139647 | 2.631833 | 0.008492556 | 0.187559523 |
| *CCDC191* | -0.713952 | 0.271316 | -2.631438 | 0.008502439 | 0.187559523 |
| *ENSG00000272379* | 1.410762 | 0.536303 | 2.630531 | 0.008525169 | NA |
| *CLHC1* | 0.74967 | 0.285154 | 2.629002 | 0.008563594 | 0.188400341 |
| *ACO2* | -0.219637 | 0.083564 | -2.628356 | 0.008579871 | 0.188400341 |
| *C6orf47* | 0.247582 | 0.09423 | 2.62741 | 0.008603762 | 0.188400341 |
| *MOB4* | 0.258686 | 0.098479 | 2.626818 | 0.008618729 | 0.188400341 |
| *FNBP1L* | 0.586361 | 0.223386 | 2.624877 | 0.008668016 | 0.188791429 |
| *LRP5L* | -0.661872 | 0.252183 | -2.624567 | 0.008675931 | 0.188791429 |
| *TNFRSF17* | 0.958871 | 0.365505 | 2.623412 | 0.00870539 | 0.188791429 |
| *FAM111A.DT* | 0.361052 | 0.137646 | 2.623038 | 0.008714957 | 0.188791429 |
| *DNASE1* | 0.360106 | 0.137345 | 2.621913 | 0.008743764 | 0.188990778 |
| *PDE4DIP* | -0.425812 | 0.16272 | -2.616836 | 0.008874888 | 0.1913958 |
| *LDOC1* | 0.832412 | 0.318239 | 2.615685 | 0.008904855 | 0.191613406 |
| *PDE6D* | -0.307802 | 0.117794 | -2.613049 | 0.008973851 | 0.192667987 |
| *MALSU1* | 0.29483 | 0.112974 | 2.609728 | 0.009061428 | 0.194115934 |
| *NRDC* | -0.256242 | 0.098284 | -2.607169 | 0.009129434 | 0.195139123 |
| *MGAT5B* | -0.736478 | 0.282587 | -2.606202 | 0.009155235 | 0.195257659 |
| *TBC1D8* | -0.964129 | 0.370294 | -2.603685 | 0.009222741 | NA |
| *DPY30* | 0.196672 | 0.075575 | 2.602332 | 0.009259225 | 0.197039568 |
| *CYBC1* | -0.220977 | 0.085104 | -2.596541 | 0.009416757 | 0.199733478 |
| *GNPDA1* | 1.002097 | 0.385992 | 2.596159 | 0.009427254 | 0.199733478 |
| *GATC* | 0.341663 | 0.131705 | 2.594163 | 0.009482148 | 0.200455945 |
| *C2orf42* | 0.50367 | 0.194282 | 2.592467 | 0.009529043 | 0.200984148 |
| *ENSG00000254837* | 0.85048 | 0.328127 | 2.591925 | 0.009544063 | NA |
| *ENSG00000266918* | -1.067828 | 0.412064 | -2.591414 | 0.009558235 | NA |
| *IL2RA* | -0.828927 | 0.319916 | -2.591079 | 0.009567557 | 0.200984148 |
| *STMN1* | -0.317673 | 0.122642 | -2.590241 | 0.009590875 | 0.200984148 |
| *AGTRAP* | -0.379794 | 0.14664 | -2.589969 | 0.009598461 | 0.200984148 |
| *NRP2* | -0.604964 | 0.233679 | -2.588872 | 0.009629083 | 0.200984148 |
| *C9orf64* | 0.665563 | 0.257097 | 2.58876 | 0.009632228 | 0.200984148 |
| *ZNF442* | -1.361937 | 0.526219 | -2.588158 | 0.009649079 | NA |
| *SORL1* | -0.308906 | 0.11939 | -2.587367 | 0.009671254 | 0.201362614 |

| | | | | | |
|---|---|---|---|---|---|
| *PSMA8* | 1.558479 | 0.602757 | 2.585586 | 0.009721367 | NA |
| *PIGX* | -0.441953 | 0.170935 | -2.585498 | 0.009723848 | 0.202021325 |
| *RDX* | 0.324749 | 0.125723 | 2.583049 | 0.009793134 | 0.202878717 |
| *XRRA1* | -0.882864 | 0.341857 | -2.582554 | 0.009807208 | 0.202878717 |
| *DBI* | -0.205654 | 0.079657 | -2.58176 | 0.009829805 | 0.202910753 |
| *C9orf16* | -0.198219 | 0.076817 | -2.580393 | 0.009868781 | 0.203280024 |
| *ZNF747* | -0.390453 | 0.151578 | -2.575917 | 0.009997454 | 0.205491385 |
| *PURB* | 0.37356 | 0.14517 | 2.573258 | 0.0100746 | 0.206354794 |
| *LRRC61* | -0.610744 | 0.23738 | -2.572849 | 0.010086513 | 0.206354794 |
| *LPAR5* | 0.412034 | 0.160201 | 2.571978 | 0.010111923 | 0.206354794 |
| *TRIM4* | 0.261697 | 0.101767 | 2.571528 | 0.010125085 | 0.206354794 |
| *ENSG00000244459* | 0.530167 | 0.206399 | 2.568645 | 0.010209695 | 0.207399409 |
| *HSPA1A* | -1.423805 | 0.554373 | -2.568317 | 0.010219369 | 0.207399409 |
| *FKBP5* | 0.620247 | 0.241574 | 2.56753 | 0.010242586 | 0.207433885 |
| *MIS12* | 0.286223 | 0.111559 | 2.565674 | 0.010297558 | 0.208109977 |
| *JADE3* | 0.535892 | 0.208999 | 2.564091 | 0.010344653 | 0.208370527 |
| *TMEM60* | 0.311244 | 0.1214 | 2.563788 | 0.010353681 | 0.208370527 |
| *S100A13* | -0.699847 | 0.273071 | -2.562881 | 0.010380777 | 0.208480606 |
| *TPM4* | 0.241441 | 0.094294 | 2.560506 | 0.010451995 | 0.209474485 |
| *UQCRC2* | -0.2253 | 0.088035 | -2.559205 | 0.010491191 | 0.209771825 |
| *PLXNA1* | -0.734738 | 0.287223 | -2.558071 | 0.010525467 | NA |
| *APEH* | -0.384043 | 0.15013 | -2.55807 | 0.010525485 | 0.209771825 |
| *MSL3* | 0.219271 | 0.085725 | 2.557851 | 0.010532112 | 0.209771825 |
| *GCFC2* | -0.420248 | 0.164351 | -2.557011 | 0.01055759 | 0.209845716 |
| *TPI1* | -0.150772 | 0.058987 | -2.55603 | 0.010587394 | 0.210005101 |
| *LUCAT1* | 1.151097 | 0.450596 | 2.554612 | 0.010630613 | NA |
| *RAB23* | 0.915486 | 0.35877 | 2.551738 | 0.010718724 | NA |
| *MLST8* | -0.318165 | 0.12469 | -2.551641 | 0.010721683 | 0.212232086 |
| *POLR3K* | 0.235166 | 0.092203 | 2.550525 | 0.010756089 | 0.212418654 |
| *MARCHF1* | 0.45882 | 0.179936 | 2.549907 | 0.010775179 | 0.212418654 |
| *CASP4* | 0.171507 | 0.067302 | 2.548309 | 0.010824646 | 0.212958337 |
| *S1PR4* | 0.332309 | 0.130475 | 2.546905 | 0.010868312 | 0.213381934 |
| *ENSG00000272426* | 0.899759 | 0.353559 | 2.54486 | 0.010932168 | NA |
| *BET1* | 0.255193 | 0.100279 | 2.544843 | 0.01093269 | 0.214209609 |
| *KLK4* | 1.932528 | 0.759725 | 2.543721 | 0.010967851 | NA |
| *MOB3B* | 0.659278 | 0.259186 | 2.543648 | 0.010970172 | 0.214439137 |
| *CERS4* | -0.314954 | 0.123849 | -2.543052 | 0.010988894 | 0.214439137 |
| *SCAMP4* | -0.462194 | 0.181921 | -2.540626 | 0.011065413 | 0.215496133 |
| *UACA* | 1.588208 | 0.625325 | 2.539814 | 0.011091157 | NA |
| *NDUFAF4* | 0.273037 | 0.107581 | 2.537963 | 0.011149982 | 0.216705291 |
| *ENSG00000271204* | 0.434991 | 0.171604 | 2.534845 | 0.01124972 | 0.217784411 |
| *PHF20* | 0.178383 | 0.070375 | 2.534742 | 0.011253005 | 0.217784411 |
| *CSDE1* | -0.151986 | 0.059993 | -2.53338 | 0.011296842 | 0.217784411 |
| *PARP15* | 0.327502 | 0.129309 | 2.532706 | 0.011318592 | 0.217784411 |
| *ARHGAP17* | 0.292886 | 0.115699 | 2.531461 | 0.011358861 | 0.217784411 |
| *NFKB2* | -0.395099 | 0.156118 | -2.530763 | 0.011381457 | 0.217784411 |
| *ILK* | 0.200973 | 0.079414 | 2.530684 | 0.011384035 | 0.217784411 |

| | | | | | |
|---|---|---|---|---|---|
| *CCDC71* | -0.366438 | 0.144802 | -2.530616 | 0.011386239 | 0.217784411 |
| *ZKSCAN1* | -0.232638 | 0.091967 | -2.529587 | 0.011419687 | 0.217991657 |
| *DHTKD1* | -0.346789 | 0.137171 | -2.528143 | 0.011466753 | 0.218311765 |
| *TMF1* | 0.202155 | 0.079986 | 2.527396 | 0.011491182 | 0.218311765 |
| *ENSG00000223881* | -1.074169 | 0.425048 | -2.527172 | 0.011498503 | NA |
| *ZNF557* | 0.461746 | 0.182726 | 2.526993 | 0.011504396 | 0.218311765 |
| *WBP1L* | -0.298134 | 0.118082 | -2.524798 | 0.011576495 | 0.219041528 |
| *TRIM69* | 0.340218 | 0.13477 | 2.524439 | 0.011588297 | 0.219041528 |
| *DCUN1D5* | 0.275544 | 0.109208 | 2.523098 | 0.011632592 | 0.21944851 |
| *CREBRF* | 0.205851 | 0.081615 | 2.522232 | 0.011661284 | 0.219529162 |
| *CD44* | -0.23864 | 0.094638 | -2.521595 | 0.011682413 | 0.219529162 |
| *SPINK2* | -1.885229 | 0.747874 | -2.520784 | 0.011709382 | NA |
| *ENSG00000267042* | 1.739171 | 0.690157 | 2.519964 | 0.011736688 | NA |
| *IGLV4.69* | -2.097872 | 0.832677 | -2.519431 | 0.011754468 | NA |
| *TRADD* | -0.243154 | 0.0966 | -2.517132 | 0.011831445 | 0.220628521 |
| *RALGPS2* | 0.300132 | 0.119245 | 2.516939 | 0.011837917 | 0.220628521 |
| *TCTN1* | -0.763839 | 0.303542 | -2.516419 | 0.011855413 | 0.220628521 |
| *GBGT1* | 0.581525 | 0.231151 | 2.515784 | 0.01187678 | 0.220628521 |
| *VPS26C* | 0.277213 | 0.110193 | 2.515706 | 0.011879429 | 0.220628521 |
| *DIP2B* | 0.368645 | 0.146579 | 2.514995 | 0.011903409 | 0.220628521 |
| *CXorf21* | 0.374027 | 0.148752 | 2.514433 | 0.011922403 | 0.220628521 |
| *RBBP8* | -0.472804 | 0.188039 | -2.514385 | 0.01192401 | 0.220628521 |
| *SLC38A2* | 0.24901 | 0.099074 | 2.513375 | 0.011958212 | 0.220837472 |
| *AP4M1* | -0.317901 | 0.12652 | -2.512659 | 0.011982516 | 0.220863207 |
| *LZTS3* | -1.217939 | 0.485052 | -2.510943 | 0.012040926 | NA |
| *SAMD9L* | 0.323284 | 0.128811 | 2.509761 | 0.012081284 | 0.221508603 |
| *GPATCH2* | -0.402455 | 0.160364 | -2.509624 | 0.012085985 | 0.221508603 |
| *PAFAH2* | 0.686438 | 0.273524 | 2.50961 | 0.012086465 | 0.221508603 |
| *ZNRD1ASP* | -0.441665 | 0.176213 | -2.506427 | 0.012195814 | 0.222587568 |
| *CPLANE1* | -0.443497 | 0.176957 | -2.506233 | 0.012202504 | 0.222587568 |
| *NDUFAF1* | 0.357688 | 0.142739 | 2.505883 | 0.012214608 | 0.222587568 |
| *FYCO1* | 0.937562 | 0.374401 | 2.504165 | 0.012274086 | NA |
| *FMNL2* | -1.466461 | 0.585966 | -2.50264 | 0.012327089 | NA |
| *ENSG00000275964* | -0.652133 | 0.260591 | -2.502512 | 0.012331528 | 0.223958297 |
| *ENSG00000268516* | 0.387623 | 0.154902 | 2.502376 | 0.012336292 | 0.223958297 |
| *C6orf226* | -0.406671 | 0.162563 | -2.501625 | 0.012362462 | 0.224011529 |
| *CD37* | -0.201285 | 0.080515 | -2.499962 | 0.012420665 | 0.224529212 |
| *PDCL3* | -0.332311 | 0.132959 | -2.49935 | 0.012442126 | 0.224529212 |
| *JMJD6* | 0.31584 | 0.126396 | 2.498816 | 0.012460905 | 0.224529212 |
| *HSH2D* | -0.328509 | 0.131521 | -2.497762 | 0.012497997 | 0.224777413 |
| *PLAAT3* | -0.903806 | 0.362394 | -2.493986 | 0.012631769 | 0.226404768 |
| *NUCB2* | -0.540416 | 0.216729 | -2.493508 | 0.012648786 | 0.226404768 |
| *HNRNPDL* | -0.184246 | 0.073903 | -2.493088 | 0.012663762 | 0.226404768 |
| *TSPAN33* | -0.336321 | 0.13493 | -2.492555 | 0.012682758 | 0.226404768 |
| *RHOT2* | -0.2119 | 0.085035 | -2.491908 | 0.012705911 | 0.226404768 |
| *CLNS1A* | 0.259339 | 0.104151 | 2.490029 | 0.012773278 | 0.226877522 |
| *SCP2* | -0.290299 | 0.116593 | -2.489855 | 0.012779512 | 0.226877522 |

| | | | | | |
|---|---|---|---|---|---|
| *COX17* | 0.314517 | 0.126393 | 2.488412 | 0.012831512 | 0.227369764 |
| *TPRA1* | 0.406747 | 0.163498 | 2.487778 | 0.012854411 | 0.227369764 |
| *C17orf49* | -0.462936 | 0.186175 | -2.486559 | 0.012898512 | 0.227731976 |
| *LINC00662* | 0.294875 | 0.118802 | 2.482062 | 0.013062439 | 0.229859586 |
| *ENSG00000255240* | -0.549757 | 0.22154 | -2.481523 | 0.01308221 | 0.229859586 |
| *SNAPC1* | 0.318273 | 0.128269 | 2.481296 | 0.013090551 | 0.229859586 |
| *CNOT6L* | -0.450012 | 0.181575 | -2.478388 | 0.013197764 | 0.230861454 |
| *GADD45B* | -0.344762 | 0.139107 | -2.478386 | 0.013197826 | 0.230861454 |
| *EIF4E* | 0.197487 | 0.079711 | 2.477529 | 0.013229569 | 0.230861454 |
| *GINS4* | -0.552733 | 0.223132 | -2.477156 | 0.013243401 | 0.230861454 |
| *IKZF2* | -0.611851 | 0.247179 | -2.475335 | 0.013311124 | 0.231615204 |
| *OPTN* | -0.522105 | 0.210977 | -2.474703 | 0.013334693 | 0.231615204 |
| *RHEB* | 0.226147 | 0.091431 | 2.473409 | 0.013383079 | 0.231847429 |
| *UNC50* | 0.291963 | 0.118082 | 2.47254 | 0.013415667 | 0.231847429 |
| *ENSG00000258056* | 0.579013 | 0.234225 | 2.472034 | 0.01343467 | 0.231847429 |
| *NAA10* | -0.158491 | 0.06412 | -2.471779 | 0.013444265 | 0.231847429 |
| *C19orf12* | 0.336793 | 0.136318 | 2.470638 | 0.013487213 | 0.232172742 |
| *BHLHE41* | 1.149076 | 0.465155 | 2.470308 | 0.013499662 | NA |
| *NFE2L3* | 1.184714 | 0.479868 | 2.468833 | 0.01355546 | NA |
| *ZNF506* | 0.331767 | 0.134403 | 2.468445 | 0.013570156 | 0.233184137 |
| *IRF2BPL* | 0.36505 | 0.147964 | 2.467156 | 0.01361911 | 0.233608934 |
| *ETS2* | -1.043487 | 0.423533 | -2.463769 | 0.013748457 | NA |
| *C12orf65* | 0.24897 | 0.101067 | 2.46341 | 0.01376223 | 0.235285613 |
| *TOB1* | 0.347784 | 0.141185 | 2.463321 | 0.013765673 | 0.235285613 |
| *DTX3L* | 0.341194 | 0.138673 | 2.460415 | 0.013877626 | 0.236779312 |
| *DHX9* | -0.223124 | 0.090772 | -2.458082 | 0.013968143 | 0.23786215 |
| *BMI1* | 0.30248 | 0.123084 | 2.457509 | 0.01399044 | 0.23786215 |
| *IGHV3.30* | 1.111375 | 0.453197 | 2.452299 | 0.014194672 | NA |
| *CYTIP* | 0.183134 | 0.074685 | 2.452074 | 0.014203564 | 0.240221556 |
| *S100A11* | 0.336669 | 0.137311 | 2.451878 | 0.014211275 | 0.240221556 |
| *LCLAT1* | 0.642027 | 0.261878 | 2.451624 | 0.014221329 | 0.240221556 |
| *PPP2R2A* | 0.225114 | 0.091844 | 2.451045 | 0.014244211 | 0.240221556 |
| *CA2* | 0.720318 | 0.29389 | 2.450978 | 0.014246865 | NA |
| *TCL1A* | -0.430437 | 0.175631 | -2.450803 | 0.01425381 | 0.240221556 |
| *TMEM101* | 0.283717 | 0.115796 | 2.450138 | 0.014280139 | 0.240232301 |
| *CDC42EP3* | 0.684629 | 0.279536 | 2.449164 | 0.014318812 | 0.240232301 |
| *CEP135* | -0.21056 | 0.085981 | -2.448903 | 0.014329209 | 0.240232301 |
| *EEFSEC* | -0.339293 | 0.138601 | -2.447982 | 0.0143659 | 0.240429296 |
| *RHBDF2* | -0.231711 | 0.094709 | -2.44656 | 0.014422664 | 0.240661341 |
| *NDUFB11* | 0.207247 | 0.084729 | 2.445992 | 0.014445401 | 0.240661341 |
| *MRPL32* | 0.224443 | 0.091768 | 2.445761 | 0.014454659 | 0.240661341 |
| *PRDX2* | -0.407855 | 0.166988 | -2.442414 | 0.014589402 | 0.241349731 |
| *NRROS* | -0.270755 | 0.110887 | -2.441712 | 0.014617823 | 0.241349731 |
| *LDHA* | -0.298754 | 0.122397 | -2.440864 | 0.014652169 | 0.241349731 |
| *ENSG00000166685* | -0.377964 | 0.154861 | -2.440669 | 0.014660091 | 0.241349731 |
| *EXOSC3* | 0.242486 | 0.099353 | 2.440661 | 0.014660421 | 0.241349731 |
| *NDUFA8* | 0.211023 | 0.086482 | 2.440069 | 0.01468446 | 0.241349731 |

| | | | | | |
|---|---|---|---|---|---|
| *TAOK3* | 0.154402 | 0.063287 | 2.439727 | 0.014698356 | 0.241349731 |
| *ENSG00000278869* | -1.372303 | 0.562529 | -2.439522 | 0.014706694 | NA |
| *RAB14* | 0.201863 | 0.082758 | 2.439191 | 0.014720187 | 0.241349731 |
| *TMEM99* | 0.505968 | 0.207435 | 2.439163 | 0.014721332 | 0.241349731 |
| *TMEM44* | 1.176217 | 0.482455 | 2.437981 | 0.014769565 | NA |
| *TPP1* | 0.220439 | 0.090531 | 2.434945 | 0.014894063 | 0.24291553 |
| *ACTN1* | 2.025524 | 0.832182 | 2.433992 | 0.014933329 | NA |
| *ATP6V0E2* | -0.546584 | 0.224584 | -2.433756 | 0.014943063 | 0.24291553 |
| *CLPTM1* | -0.225675 | 0.092732 | -2.433631 | 0.014948227 | 0.24291553 |
| *CTSH* | -0.183722 | 0.075496 | -2.433539 | 0.014952041 | 0.24291553 |
| *RAB10* | 0.251984 | 0.103577 | 2.432818 | 0.014981819 | 0.24291553 |
| *HLA.DMB* | -0.257051 | 0.105663 | -2.43275 | 0.014984655 | 0.24291553 |
| *HMGN2* | -0.173934 | 0.071503 | -2.432543 | 0.01499323 | 0.24291553 |
| *POR* | -0.32243 | 0.13264 | -2.430865 | 0.015062806 | 0.243633305 |
| *ENSG00000228172* | -1.550922 | 0.63839 | -2.429427 | 0.015122704 | NA |
| *GPSM2* | -0.444644 | 0.183043 | -2.429179 | 0.015133065 | 0.244255482 |
| *RRP1B* | 0.375036 | 0.154458 | 2.428079 | 0.01517904 | 0.244255482 |
| *NRDE2* | -0.27923 | 0.115026 | -2.427534 | 0.015201861 | 0.244255482 |
| *RNF38* | 0.365068 | 0.150388 | 2.427516 | 0.015202623 | 0.244255482 |
| *SERTAD1* | 0.408418 | 0.168326 | 2.426347 | 0.015251677 | 0.244568455 |
| *PPIL4* | 0.257809 | 0.106308 | 2.425116 | 0.015303487 | 0.244568455 |
| *LSR* | 0.68907 | 0.28417 | 2.424855 | 0.015314523 | 0.244568455 |
| *SLTM* | 0.181095 | 0.07469 | 2.42464 | 0.015323584 | 0.244568455 |
| *ZNF547* | 0.822169 | 0.339424 | 2.422247 | 0.015424852 | NA |
| *ASB16.AS1* | 0.399857 | 0.165083 | 2.422155 | 0.015428773 | 0.245659567 |
| *ENSG00000235078* | 0.473279 | 0.195461 | 2.421352 | 0.015462902 | 0.245659567 |
| *USF1* | -0.573339 | 0.236797 | -2.421223 | 0.015468398 | 0.245659567 |
| *TNRC6C* | 0.344169 | 0.142303 | 2.418568 | 0.015581723 | 0.247052322 |
| *LANCL2* | -0.713504 | 0.295266 | -2.41648 | 0.015671393 | NA |
| *CDC42SE2* | 0.206113 | 0.085305 | 2.416191 | 0.01568382 | 0.247573742 |
| *PSMA5* | 0.177786 | 0.073596 | 2.415692 | 0.015705323 | 0.247573742 |
| *PFDN2* | 0.166644 | 0.068991 | 2.415455 | 0.015715551 | 0.247573742 |
| *XKR8* | 0.43758 | 0.181187 | 2.415074 | 0.015732015 | 0.247573742 |
| *SRSF6* | -0.285443 | 0.118205 | -2.414819 | 0.015743019 | 0.247573742 |
| *GPR183* | 0.393364 | 0.163176 | 2.410671 | 0.01592319 | 0.249867998 |
| *C8orf37* | 0.674546 | 0.279863 | 2.410273 | 0.015940592 | NA |
| *TMC8* | -0.345587 | 0.143396 | -2.410021 | 0.015951589 | 0.249867998 |
| *KIF1BP* | 0.477009 | 0.197956 | 2.409677 | 0.015966669 | 0.249867998 |
| *WDR92* | 0.416019 | 0.172823 | 2.407196 | 0.016075547 | 0.251164146 |
| *ATP13A3* | 0.399144 | 0.165858 | 2.406533 | 0.016104758 | 0.25121338 |
| *LIMD1* | -0.74012 | 0.307796 | -2.404576 | 0.016191239 | 0.251656337 |
| *ZNF608* | -0.591373 | 0.245988 | -2.40407 | 0.016213661 | 0.251656337 |
| *APAF1* | 0.454102 | 0.188889 | 2.404065 | 0.016213899 | 0.251656337 |
| *YARS* | -0.387903 | 0.161426 | -2.402979 | 0.016262105 | 0.251656337 |
| *PFKP* | -0.559377 | 0.232788 | -2.402944 | 0.016263682 | 0.251656337 |
| *ENSG00000228242* | 1.294571 | 0.538786 | 2.402755 | 0.016272081 | NA |
| *TNFRSF14* | 0.173708 | 0.072338 | 2.401326 | 0.016335784 | 0.252080189 |

| | | | | | |
|---|---|---|---|---|---|
| *MBP* | -0.230101 | 0.09585 | -2.400647 | 0.016366127 | 0.252080189 |
| *ZC3H6* | 0.487046 | 0.202887 | 2.400571 | 0.016369523 | 0.252080189 |
| *MAP3K4* | 0.333038 | 0.138771 | 2.399902 | 0.016399481 | 0.252138751 |
| *HIP1R* | -0.334217 | 0.139311 | -2.39907 | 0.016436772 | 0.252309685 |
| *ENSG00000276449* | 0.928079 | 0.387069 | 2.397711 | 0.016497887 | NA |
| *MICAL1* | -0.346141 | 0.144457 | -2.396148 | 0.016568421 | 0.253650009 |
| *NEIL1* | -0.629627 | 0.262899 | -2.394942 | 0.016622975 | 0.253650009 |
| *GATD1* | 0.276208 | 0.115338 | 2.394774 | 0.016630615 | 0.253650009 |
| *BAG1* | -0.194403 | 0.081196 | -2.394241 | 0.016654806 | 0.253650009 |
| *JARID2* | -0.291537 | 0.121782 | -2.393921 | 0.016669355 | 0.253650009 |
| *POLR2J* | -0.178853 | 0.074751 | -2.392657 | 0.016726877 | 0.253650009 |
| *BIN1* | 0.359564 | 0.150354 | 2.391451 | 0.016781943 | 0.253650009 |
| *C4orf36* | 0.593021 | 0.248005 | 2.39117 | 0.016794784 | 0.253650009 |
| *ZFP37* | 1.171485 | 0.490119 | 2.390204 | 0.016839027 | NA |
| *ZNF138* | 0.335426 | 0.140363 | 2.389704 | 0.016861978 | 0.253650009 |
| *PCYT1A* | 0.364373 | 0.152478 | 2.389675 | 0.016863285 | 0.253650009 |
| *HSPBAP1* | -1.060638 | 0.443857 | -2.389597 | 0.016866881 | 0.253650009 |
| *DIPK1A* | -0.447733 | 0.18746 | -2.388418 | 0.01692109 | 0.253650009 |
| *ASCC2* | -0.25646 | 0.107406 | -2.387765 | 0.016951169 | 0.253650009 |
| *LLGL1* | -0.676866 | 0.283549 | -2.387123 | 0.016980787 | 0.253650009 |
| *ZNF594* | 0.609196 | 0.255225 | 2.3869 | 0.0169911 | 0.253650009 |
| *ENSG00000188206* | 0.302909 | 0.126907 | 2.386862 | 0.016992878 | 0.253650009 |
| *MED25* | -0.385915 | 0.161701 | -2.386599 | 0.017005033 | 0.253650009 |
| *BCS1L* | 0.394452 | 0.165295 | 2.386355 | 0.017016328 | 0.253650009 |
| *ZBTB43* | -0.317587 | 0.133094 | -2.386189 | 0.01702402 | 0.253650009 |
| *TUNAR* | -1.520323 | 0.637413 | -2.385145 | 0.017072391 | NA |
| *RASSF1* | 0.220894 | 0.09262 | 2.38496 | 0.017080979 | 0.253716467 |
| *SDAD1* | -0.261782 | 0.109764 | -2.384957 | 0.017081119 | 0.253716467 |
| *ADD2* | 0.841739 | 0.353134 | 2.383626 | 0.017143034 | NA |
| *RPRD1A* | -0.243313 | 0.102119 | -2.38265 | 0.017188513 | 0.254550782 |
| *GPN3* | 0.36739 | 0.154262 | 2.381598 | 0.017237691 | 0.254550782 |
| *ENSG00000272758* | 0.613999 | 0.257818 | 2.381524 | 0.017241187 | 0.254550782 |
| *ENSG00000261669* | 0.768193 | 0.32262 | 2.381107 | 0.017260701 | 0.254550782 |
| *PSMA4* | -0.130101 | 0.054643 | -2.380923 | 0.017269317 | 0.254550782 |
| *ZBTB20.AS2* | -1.076662 | 0.452221 | -2.380829 | 0.017273713 | NA |
| *RNF43* | -0.662986 | 0.278576 | -2.379915 | 0.017316623 | 0.254587651 |
| *LTBP4* | 0.380352 | 0.159829 | 2.379745 | 0.017324637 | 0.254587651 |
| *TMA16* | 0.283821 | 0.119366 | 2.377739 | 0.017419133 | 0.254922136 |
| *RBM28* | 0.272153 | 0.114482 | 2.377252 | 0.017442177 | 0.254922136 |
| *RPL24* | 0.203397 | 0.085567 | 2.377042 | 0.01745209 | 0.254922136 |
| *ATG101* | 0.26315 | 0.110706 | 2.377019 | 0.017453175 | 0.254922136 |
| *UBE2J2* | -0.181608 | 0.076421 | -2.376398 | 0.017482584 | 0.254965375 |
| *SNRPG* | 0.160576 | 0.06763 | 2.374335 | 0.017580576 | 0.255669147 |
| *ZBTB20* | 0.248874 | 0.104821 | 2.374266 | 0.017583884 | 0.255669147 |
| *TSFM* | 0.339043 | 0.142973 | 2.371383 | 0.017721649 | 0.25728418 |
| *PLTP* | -1.234125 | 0.52096 | -2.368943 | 0.017839006 | NA |
| *TOX4* | -0.209834 | 0.088633 | -2.367449 | 0.017911167 | 0.259644587 |

| | | | | | |
|---|---|---|---|---|---|
| *ZDHHC16* | 0.297357 | 0.125658 | 2.366398 | 0.01796212 | 0.259733531 |
| *BDP1* | 0.145741 | 0.061592 | 2.366211 | 0.017971189 | 0.259733531 |
| *RGMB* | 0.810702 | 0.342719 | 2.365503 | 0.018005583 | NA |
| *TCL1B* | -1.242108 | 0.525821 | -2.362224 | 0.018165646 | 0.261976473 |
| *HNRNPH3* | 0.180874 | 0.076593 | 2.361504 | 0.018200989 | 0.261976473 |
| *ECE1* | -0.389026 | 0.164754 | -2.361249 | 0.018213511 | 0.261976473 |
| *PSMC3* | -0.207839 | 0.088045 | -2.360592 | 0.018245773 | 0.261976473 |
| *AGMAT* | 0.576338 | 0.244185 | 2.360257 | 0.01826226 | 0.261976473 |
| *OTUD6B.AS1* | 0.215057 | 0.091145 | 2.359495 | 0.018299804 | 0.262124025 |
| *BCLAF1* | -0.162901 | 0.069057 | -2.358946 | 0.018326929 | 0.262124025 |
| *YY1* | 0.152605 | 0.064759 | 2.356512 | 0.018447479 | 0.262741848 |
| *OTUD1* | 0.504246 | 0.213987 | 2.356436 | 0.018451268 | 0.262741848 |
| *UBALD2* | -0.249223 | 0.105763 | -2.356423 | 0.018451891 | 0.262741848 |
| *BEND4* | 0.516975 | 0.219631 | 2.353837 | 0.018580775 | 0.264186824 |
| *NPHP4* | -1.115287 | 0.474093 | -2.352464 | 0.018649495 | NA |
| *PGPEP1* | 0.430555 | 0.183035 | 2.352301 | 0.018657646 | 0.264889119 |
| *PRDX6* | -0.174361 | 0.074186 | -2.350337 | 0.018756438 | 0.265900088 |
| *DTNBP1* | -0.200815 | 0.085589 | -2.346281 | 0.018961814 | 0.268378487 |
| *ENSG00000239636* | -0.59951 | 0.255569 | -2.345787 | 0.018986943 | 0.268378487 |
| *ENSG00000277007* | 0.900116 | 0.383833 | 2.34507 | 0.019023521 | NA |
| *DDX23* | 0.30078 | 0.128267 | 2.344958 | 0.019029209 | 0.268582097 |
| *ENSG00000276075* | -1.077995 | 0.460021 | -2.343359 | 0.019110978 | NA |
| *WAC* | 0.155672 | 0.066475 | 2.341806 | 0.019190696 | 0.270076051 |
| *BOLA2B* | -0.360762 | 0.154053 | -2.341798 | 0.019191089 | 0.270076051 |
| *LINC01250* | -2.012553 | 0.859833 | -2.340633 | 0.01925107 | NA |
| *UBE2L3* | -0.146382 | 0.062579 | -2.339145 | 0.019327932 | 0.270989764 |
| *RAE1* | -0.336519 | 0.143893 | -2.338676 | 0.01935221 | 0.270989764 |
| *DPYD* | -0.410374 | 0.175505 | -2.338239 | 0.019374846 | 0.270989764 |
| *OGT* | 0.272605 | 0.116593 | 2.338098 | 0.019382155 | 0.270989764 |
| *GNG3* | -1.343126 | 0.574478 | -2.337994 | 0.019387557 | NA |
| *RBM18* | 0.289409 | 0.123794 | 2.33782 | 0.01939657 | 0.270989764 |
| *ZSCAN29* | 0.457447 | 0.195803 | 2.336269 | 0.019477218 | 0.27165467 |
| *CDC37* | -0.166878 | 0.071443 | -2.335822 | 0.019500522 | 0.27165467 |
| *ENSG00000267169* | -0.639668 | 0.273919 | -2.335243 | 0.019530744 | NA |
| *ZBTB16* | 1.469758 | 0.629481 | 2.33487 | 0.019550198 | NA |
| *LPP.AS2* | -0.849675 | 0.36395 | -2.334594 | 0.019564615 | 0.272154239 |
| *ARID1A* | 0.238832 | 0.102327 | 2.334014 | 0.019594986 | 0.272183953 |
| *AKAP12* | -1.735568 | 0.743802 | -2.333373 | 0.019628559 | NA |
| *HSPA8* | 0.300879 | 0.128959 | 2.333129 | 0.019641355 | 0.272435492 |
| *KPNA6* | 0.219499 | 0.094122 | 2.332081 | 0.019696417 | 0.272758104 |
| *ING1* | 0.299385 | 0.128437 | 2.330985 | 0.019754125 | 0.272758104 |
| *ENSG00000272106* | -0.391122 | 0.167819 | -2.330615 | 0.019773689 | 0.272758104 |
| *SDF2L1* | 0.24965 | 0.107143 | 2.330073 | 0.019802318 | 0.272758104 |
| *LPIN1* | 0.50362 | 0.216146 | 2.330001 | 0.019806086 | 0.272758104 |
| *DDI2* | 0.334699 | 0.143827 | 2.327091 | 0.019960419 | 0.274491361 |
| *PSMB10* | -0.211002 | 0.090702 | -2.326322 | 0.020001393 | 0.274663006 |
| *JUP* | -1.006515 | 0.433331 | -2.32274 | 0.020193105 | 0.276478586 |

| | | | | | |
|---|---|---|---|---|---|
| *ARHGAP30* | 0.182219 | 0.078475 | 2.322007 | 0.020232537 | 0.276478586 |
| *ZNF2* | 0.592888 | 0.255348 | 2.321885 | 0.020239133 | 0.276478586 |
| *ENSG00000224086* | -0.387217 | 0.166787 | -2.321634 | 0.020252658 | 0.276478586 |
| *SLC25A38* | 0.231989 | 0.099959 | 2.320838 | 0.020295595 | 0.276478586 |
| *GRAMD1C* | -0.33736 | 0.145373 | -2.320651 | 0.020305689 | 0.276478586 |
| *DLGAP3* | -0.789879 | 0.340678 | -2.318547 | 0.020419634 | NA |
| *SP2* | -0.302041 | 0.130292 | -2.318183 | 0.020439358 | 0.277557786 |
| *ENSG00000229539* | 0.361154 | 0.155819 | 2.317786 | 0.020460963 | 0.277557786 |
| *ZNF284* | 0.501504 | 0.21639 | 2.317595 | 0.020471326 | 0.277557786 |
| *UBE2E3* | -0.214999 | 0.09281 | -2.316562 | 0.020527594 | 0.277929781 |
| *OAS2* | 0.392717 | 0.169574 | 2.315905 | 0.020563471 | 0.278025045 |
| *NAA60* | -0.259469 | 0.112146 | -2.313664 | 0.020686177 | 0.278806703 |
| *SFXN3* | 0.549334 | 0.237445 | 2.31352 | 0.020694089 | 0.278806703 |
| *CCDC88B* | 0.320902 | 0.138722 | 2.313265 | 0.02070805 | 0.278806703 |
| *ZNF395* | 0.264439 | 0.114474 | 2.310035 | 0.020886235 | 0.280792593 |
| *ARPC3* | -0.132395 | 0.057325 | -2.309537 | 0.020913805 | 0.280792593 |
| *TYW3* | 0.301001 | 0.130384 | 2.308579 | 0.020966935 | 0.281114406 |
| *ZNF420* | 0.348108 | 0.150861 | 2.307471 | 0.021028591 | 0.281147259 |
| *PSAP* | -0.153908 | 0.066713 | -2.307023 | 0.021053565 | 0.281147259 |
| *APIP* | 0.237781 | 0.103071 | 2.306963 | 0.02105688 | 0.281147259 |
| *ATP6V0A2* | -0.35007 | 0.151809 | -2.305993 | 0.021110997 | 0.281479954 |
| *CTBP1.AS* | -1.276876 | 0.554036 | -2.30468 | 0.021184476 | NA |
| *CIPC* | 0.385058 | 0.167084 | 2.304574 | 0.021190455 | 0.282052038 |
| *ZC3H12D* | -0.351046 | 0.15238 | -2.303756 | 0.021236317 | 0.282052038 |
| *UBLCP1* | -0.316544 | 0.137409 | -2.303661 | 0.021241678 | 0.282052038 |
| *CNR1* | 0.711396 | 0.309004 | 2.302224 | 0.021322528 | 0.282531849 |
| *ZNF184* | 0.440309 | 0.191288 | 2.301813 | 0.021345735 | 0.282531849 |
| *GINM1* | 0.303166 | 0.131728 | 2.301458 | 0.021365738 | 0.282531849 |
| *PJA1* | 0.365318 | 0.158791 | 2.300625 | 0.021412836 | 0.28276676 |
| *CEMIP2* | 0.543599 | 0.23637 | 2.299783 | 0.021460501 | 0.283008528 |
| *AP1S3* | -0.509132 | 0.221461 | -2.298968 | 0.021506738 | 0.28323081 |
| *NPHP3* | 0.343873 | 0.1497 | 2.297072 | 0.02161469 | 0.283966938 |
| *PPP4R2* | 0.237276 | 0.103301 | 2.296951 | 0.021621549 | 0.283966938 |
| *POLR2B* | -0.23662 | 0.103068 | -2.295762 | 0.021689499 | 0.284471791 |
| *ENSG00000277654* | -0.464981 | 0.202597 | -2.2951 | 0.021727413 | 0.284581871 |
| *DNAJA2* | -0.204152 | 0.088976 | -2.294452 | 0.021764561 | 0.284681644 |
| *AGO2* | -0.242292 | 0.105656 | -2.293211 | 0.021835865 | 0.284872375 |
| *HINT1* | 0.233137 | 0.101668 | 2.293119 | 0.021841178 | 0.284872375 |
| *MSL2* | 0.282448 | 0.12328 | 2.291119 | 0.021956546 | 0.284872375 |
| *FRG1* | -0.210053 | 0.091707 | -2.290486 | 0.021993131 | 0.284872375 |
| *HLA.E* | -0.260839 | 0.11388 | -2.290466 | 0.021994343 | 0.284872375 |
| *SETDB1* | -0.360352 | 0.157335 | -2.290355 | 0.022000719 | 0.284872375 |
| *ELP6* | -0.320438 | 0.139967 | -2.289387 | 0.022056886 | 0.284872375 |
| *FAM214B* | 0.5619 | 0.24553 | 2.288522 | 0.022107138 | 0.284872375 |
| *CDC42* | -0.159678 | 0.069774 | -2.288494 | 0.022108772 | 0.284872375 |
| *TAOK1* | 0.274947 | 0.120144 | 2.28848 | 0.022109562 | 0.284872375 |
| *NUDCD2* | 0.232226 | 0.101487 | 2.288239 | 0.022123615 | 0.284872375 |

| | | | | | |
|---|---|---|---|---|---|
| *CNIH4* | 0.218412 | 0.095457 | 2.288065 | 0.022133756 | 0.284872375 |
| *HIST1H2BN* | 0.652152 | 0.285194 | 2.286699 | 0.02221338 | 0.285515979 |
| *PRPF18* | -0.28806 | 0.126005 | -2.286094 | 0.022248737 | 0.285589644 |
| *RNASEL* | 0.534892 | 0.23403 | 2.28557 | 0.022279407 | 0.285603038 |
| *ENSG00000267213* | -1.820087 | 0.796399 | -2.285396 | 0.022289628 | NA |
| *NOTCH2* | 0.281947 | 0.123438 | 2.284117 | 0.022364656 | 0.286315116 |
| *TMUB1* | -0.227589 | 0.099677 | -2.283261 | 0.022415009 | 0.286579165 |
| *PCBD1* | 0.413756 | 0.181327 | 2.281825 | 0.022499647 | 0.28728026 |
| *ABI1* | 0.192018 | 0.084217 | 2.280047 | 0.022604891 | 0.288242262 |
| *TSPO* | 0.19151 | 0.084036 | 2.278891 | 0.022673533 | 0.288546732 |
| *CISD1* | 0.255499 | 0.112128 | 2.278637 | 0.022688633 | 0.288546732 |
| *FAM89A* | 1.187535 | 0.521219 | 2.27838 | 0.022703957 | NA |
| *PAK1IP1* | 0.341597 | 0.150008 | 2.277195 | 0.022774584 | 0.289258221 |
| *DGAT2* | 0.681154 | 0.299129 | 2.277123 | 0.022778854 | NA |
| *ARHGAP20* | 1.291142 | 0.567362 | 2.275694 | 0.022864352 | NA |
| *NCBP2AS2* | -0.241269 | 0.106165 | -2.272594 | 0.023050653 | 0.291859637 |
| *MRNIP* | 0.23175 | 0.101994 | 2.272191 | 0.023074987 | 0.291859637 |
| *THEMIS2* | 0.708246 | 0.311718 | 2.272069 | 0.023082319 | 0.291859637 |
| *KLF6* | 0.304123 | 0.133871 | 2.271768 | 0.023100509 | 0.291859637 |
| *HIPK1* | 0.355692 | 0.156641 | 2.270745 | 0.023162403 | 0.292258596 |
| *BATF2* | 1.14306 | 0.503455 | 2.27043 | 0.023181527 | NA |
| *ZNF613* | 0.725898 | 0.31975 | 2.270203 | 0.023195269 | NA |
| *TTC21A* | 0.312017 | 0.137544 | 2.268493 | 0.023299188 | 0.29360023 |
| *ATP6V1C1* | 0.268665 | 0.118523 | 2.26677 | 0.023404261 | 0.294539269 |
| *STT3B* | -0.329412 | 0.145408 | -2.265426 | 0.02348653 | 0.294654164 |
| *NUP107* | -0.229407 | 0.101297 | -2.26469 | 0.023531687 | 0.294654164 |
| *JRKL* | 0.402949 | 0.177928 | 2.264678 | 0.02353247 | 0.294654164 |
| *KLHL21* | 0.50577 | 0.223375 | 2.264216 | 0.023560819 | 0.294654164 |
| *NAALADL1* | 0.442018 | 0.195227 | 2.264128 | 0.02356622 | 0.294654164 |
| *HIST1H2BH* | 0.830854 | 0.367453 | 2.261117 | 0.023752043 | NA |
| *SLC15A2* | 0.790525 | 0.349701 | 2.260577 | 0.023785441 | NA |
| *CBFA2T3* | 0.571699 | 0.25323 | 2.257631 | 0.023968687 | 0.297451548 |
| *MED30* | 0.235782 | 0.104444 | 2.257493 | 0.02397727 | 0.297451548 |
| *REL* | -0.271584 | 0.120304 | -2.257483 | 0.023977917 | 0.297451548 |
| *VPS11* | -0.364878 | 0.161681 | -2.256779 | 0.024021884 | 0.297451548 |
| *SUB1* | 0.13462 | 0.059662 | 2.256385 | 0.024046516 | 0.297451548 |
| *MIATNB* | 0.490883 | 0.217558 | 2.25633 | 0.024049979 | 0.297451548 |
| *TSNAX* | -0.261451 | 0.115891 | -2.256009 | 0.024070052 | 0.297451548 |
| *CYC1* | -0.170618 | 0.075671 | -2.254741 | 0.024149567 | 0.297451548 |
| *IGKV3.20* | -0.82903 | 0.367708 | -2.254585 | 0.024159371 | NA |
| *UGDH* | 0.280876 | 0.124592 | 2.254367 | 0.02417307 | 0.297451548 |
| *LINC02362* | -1.4649 | 0.649837 | -2.254258 | 0.024179917 | NA |
| *TNFRSF10D* | 0.591574 | 0.262528 | 2.253374 | 0.024235552 | 0.297451548 |
| *COMMD6* | 0.126199 | 0.056005 | 2.253337 | 0.024237938 | 0.297451548 |
| *FLYWCH1* | 0.353324 | 0.156803 | 2.253293 | 0.02424066 | 0.297451548 |
| *GIT2* | 0.194001 | 0.0861 | 2.253199 | 0.024246621 | 0.297451548 |
| *PRMT1* | -0.189263 | 0.084026 | -2.252442 | 0.024294377 | 0.297451548 |

| | | | | | |
|---|---|---|---|---|---|
| *TSC22D1* | -0.424737 | 0.188574 | -2.252369 | 0.02429899 | 0.297451548 |
| *ELOC* | 0.192633 | 0.085529 | 2.252252 | 0.024306336 | 0.297451548 |
| *MAP3K8* | -0.452597 | 0.201016 | -2.251546 | 0.024351004 | 0.297451548 |
| *PYROXD1* | -0.363854 | 0.161652 | -2.25084 | 0.024395684 | 0.297451548 |
| *ATP1A1* | -0.197422 | 0.087711 | -2.250834 | 0.024396055 | 0.297451548 |
| *ENSG00000267390* | 0.462101 | 0.205318 | 2.25066 | 0.024407072 | 0.297451548 |
| *HIRA* | -0.451131 | 0.200504 | -2.249979 | 0.024450296 | 0.297602093 |
| *ENSG00000232611* | 0.643721 | 0.286158 | 2.249528 | 0.024478895 | NA |
| *PCGF1* | -0.428319 | 0.190459 | -2.24888 | 0.024520163 | 0.298076125 |
| *TAT.AS1* | 0.951655 | 0.423227 | 2.248568 | 0.024540007 | NA |
| *ENSG00000258308* | -1.509776 | 0.671605 | -2.24801 | 0.024575521 | NA |
| *EYA3* | 0.559103 | 0.24876 | 2.247559 | 0.024604345 | 0.298165384 |
| *THAP2* | 0.331562 | 0.147537 | 2.247324 | 0.024619328 | 0.298165384 |
| *IKZF5* | 0.333449 | 0.148412 | 2.246781 | 0.024653995 | 0.298165384 |
| *TRA2B* | -0.155683 | 0.069307 | -2.246262 | 0.024687246 | 0.298165384 |
| *CD82* | -0.216672 | 0.096469 | -2.24602 | 0.024702756 | 0.298165384 |
| *RNH1* | -0.230852 | 0.10279 | -2.245858 | 0.024713085 | 0.298165384 |
| *SSBP4* | -0.289651 | 0.129012 | -2.245149 | 0.024758573 | 0.2983408 |
| *NEDD8* | 0.119982 | 0.05347 | 2.243918 | 0.024837679 | 0.298571996 |
| *ACTL6A* | 0.28044 | 0.124991 | 2.24367 | 0.024853623 | 0.298571996 |
| *RYK* | 0.376165 | 0.167676 | 2.243406 | 0.024870676 | 0.298571996 |
| *THNSL1* | 0.816568 | 0.364098 | 2.242717 | 0.024915038 | NA |
| *CD38* | -0.512464 | 0.228621 | -2.241549 | 0.024990555 | 0.299027299 |
| *IL2RG* | -0.249508 | 0.111313 | -2.2415 | 0.024993683 | 0.299027299 |
| *LACTB2* | 0.33353 | 0.148806 | 2.241377 | 0.02500166 | 0.299027299 |
| *ZNF709* | -1.397995 | 0.623948 | -2.240564 | 0.02505432 | NA |
| *BAZ2B* | -0.341969 | 0.152678 | -2.239808 | 0.025103394 | 0.299872019 |
| *RASGEF1A* | -0.834402 | 0.372668 | -2.238992 | 0.025156447 | NA |
| *HHIP.AS1* | -1.451788 | 0.648716 | -2.237941 | 0.025224935 | NA |
| *PRRT3* | 0.550644 | 0.246075 | 2.23771 | 0.025239977 | 0.301130422 |
| *HES6* | -0.537488 | 0.240289 | -2.236841 | 0.025296739 | 0.301434562 |
| *ZBTB4* | 0.304131 | 0.136013 | 2.236045 | 0.025348799 | 0.301682 |
| *SRSF11* | 0.15124 | 0.067701 | 2.233939 | 0.025487108 | 0.30233239 |
| *ST3GAL1* | 0.3436 | 0.15383 | 2.23364 | 0.025506766 | 0.30233239 |
| *SPDL1* | 0.518211 | 0.232029 | 2.233386 | 0.025523495 | 0.30233239 |
| *PRR4* | 0.376771 | 0.168706 | 2.233304 | 0.025528897 | 0.30233239 |
| *CENPM* | -0.379531 | 0.170032 | -2.232113 | 0.02560749 | 0.302626271 |
| *POLH* | 0.40591 | 0.181861 | 2.231977 | 0.025616498 | 0.302626271 |
| *DCXR.DT* | -1.107495 | 0.496595 | -2.230175 | 0.025735846 | NA |
| *RCSD1* | -0.287432 | 0.128919 | -2.229559 | 0.025776758 | 0.30360294 |
| *MTHFD2L* | -0.341373 | 0.153131 | -2.229294 | 0.025794335 | 0.30360294 |
| *PDCD4* | 0.426457 | 0.191347 | 2.228714 | 0.025832921 | 0.30360294 |
| *RAD23A* | 0.134139 | 0.060196 | 2.228386 | 0.025854817 | 0.30360294 |
| *UBB* | -0.208779 | 0.093692 | -2.228358 | 0.02585664 | 0.30360294 |
| *ENSG00000271971* | 1.008736 | 0.452708 | 2.228226 | 0.025865456 | NA |
| *TBC1D12* | 1.86663 | 0.837923 | 2.227688 | 0.025901352 | NA |
| *HMOX1* | 0.773206 | 0.347168 | 2.227183 | 0.025935055 | 0.303929723 |

| | | | | | |
|---|---|---|---|---|---|
| *FZR1* | -0.307976 | 0.138315 | -2.226627 | 0.025972226 | 0.303929723 |
| *PSMC1* | -0.162944 | 0.073183 | -2.226525 | 0.025979055 | 0.303929723 |
| *ENSG00000264443* | 0.55707 | 0.250298 | 2.225628 | 0.026039126 | 0.304052359 |
| *SLC16A1.AS1* | 0.323992 | 0.145591 | 2.225362 | 0.026056928 | 0.304052359 |
| *ENSG00000279927* | -1.17333 | 0.527316 | -2.2251 | 0.026074504 | NA |
| *QTRT1* | -0.330967 | 0.148752 | -2.224956 | 0.02608416 | 0.304052359 |
| *ZNF146* | 0.257077 | 0.115677 | 2.222358 | 0.026259113 | 0.305722036 |
| *FGL2* | -1.169023 | 0.526216 | -2.221564 | 0.026312777 | NA |
| *NUP43* | 0.386396 | 0.173931 | 2.221554 | 0.026313463 | 0.305985267 |
| *VSIR* | -0.65088 | 0.293198 | -2.219932 | 0.02642335 | 0.306892888 |
| *ZNF322* | 0.314464 | 0.141729 | 2.218777 | 0.026501868 | 0.306992141 |
| *CDK5R1* | 1.98572 | 0.895079 | 2.218486 | 0.026521736 | NA |
| *CEP85L* | 0.354008 | 0.159591 | 2.218223 | 0.026539625 | 0.306992141 |
| *UBA6* | 0.27765 | 0.125176 | 2.218071 | 0.026550012 | 0.306992141 |
| *EPB41L2* | -0.423495 | 0.190945 | -2.217894 | 0.026562064 | 0.306992141 |
| *FCGRT* | -0.291407 | 0.131419 | -2.217387 | 0.02659667 | 0.306992141 |
| *DR1* | 0.200082 | 0.090249 | 2.217002 | 0.02662297 | 0.306992141 |
| *RIPK2* | 0.267373 | 0.120661 | 2.215909 | 0.026697751 | 0.307279515 |
| *ENSG00000261087* | 0.643231 | 0.290305 | 2.215706 | 0.026711642 | 0.307279515 |
| *ENSG00000274213* | -0.783953 | 0.354057 | -2.214202 | 0.026814854 | NA |
| *ZHX2* | -0.279018 | 0.126014 | -2.214187 | 0.026815924 | 0.308010351 |
| *DYNC1LI2* | 0.244152 | 0.110284 | 2.21385 | 0.026839076 | 0.308010351 |
| *PELI3* | -0.708255 | 0.320038 | -2.213031 | 0.026895536 | NA |
| *CHD1* | 0.194021 | 0.087686 | 2.212681 | 0.026919675 | 0.308432203 |
| *MRPL18* | 0.201664 | 0.091161 | 2.212181 | 0.026954134 | 0.308432203 |
| *EP300* | 0.207195 | 0.093674 | 2.211869 | 0.026975696 | 0.308432203 |
| *TMEM140* | 0.408961 | 0.184955 | 2.211138 | 0.027026285 | 0.308432203 |
| *PHF13* | 0.495761 | 0.224225 | 2.211 | 0.02703581 | 0.308432203 |
| *GTF2IRD2B* | -0.433868 | 0.196447 | -2.208572 | 0.027204417 | 0.309988868 |
| *CFAP73* | -1.35794 | 0.61562 | -2.205808 | 0.027397455 | NA |
| *RPL36* | 0.12709 | 0.057622 | 2.205589 | 0.027412797 | 0.311693955 |
| *CLMN* | 0.486069 | 0.220389 | 2.205504 | 0.027418721 | 0.311693955 |
| *WDR11* | -0.220339 | 0.099999 | -2.203412 | 0.027565724 | 0.312995972 |
| *AVP* | -2.254481 | 1.023476 | -2.202769 | 0.027611059 | NA |
| *KARS* | -0.192369 | 0.087337 | -2.202609 | 0.027622348 | 0.313089243 |
| *ENSG00000125726* | -0.921826 | 0.41856 | -2.202374 | 0.027638895 | 0.313089243 |
| *FBXL8* | -0.822997 | 0.374051 | -2.200226 | 0.027790871 | NA |
| *ZXDA* | -0.710118 | 0.322774 | -2.200046 | 0.027803609 | NA |
| *TWISTNB* | -0.328444 | 0.149301 | -2.199883 | 0.027815221 | 0.313917916 |
| *HMGB3* | -0.51531 | 0.234346 | -2.198929 | 0.027882968 | 0.313917916 |
| *SPHK1* | 0.716971 | 0.326061 | 2.198887 | 0.027885941 | 0.313917916 |
| *RBX1* | 0.168303 | 0.076554 | 2.198503 | 0.027913296 | 0.313917916 |
| *HDHD3* | 0.239255 | 0.108846 | 2.198098 | 0.027942145 | 0.313917916 |
| *MAVS* | 0.230229 | 0.10474 | 2.198098 | 0.02794213 | 0.313917916 |
| *MYO1F* | 0.374427 | 0.170359 | 2.197878 | 0.027957799 | 0.313917916 |
| *DLGAP1.AS2* | 0.671586 | 0.30559 | 2.197671 | 0.027972561 | 0.313917916 |
| *C1GALT1* | 0.259914 | 0.118321 | 2.196696 | 0.028042178 | 0.31411531 |

| | | | | | |
|---|---|---|---|---|---|
| *CD55* | -0.265692 | 0.120961 | -2.196512 | 0.02805532 | 0.31411531 |
| *MINK1* | -0.430959 | 0.196483 | -2.193367 | 0.028280921 | 0.316273868 |
| *CBLN3* | -0.451568 | 0.20615 | -2.190483 | 0.028489202 | 0.318233959 |
| *MEGF11* | -0.888699 | 0.405771 | -2.190146 | 0.028513638 | NA |
| *MED6* | -0.208103 | 0.095027 | -2.18993 | 0.028529331 | 0.318313373 |
| *C8orf82* | -0.709761 | 0.324131 | -2.189734 | 0.02854355 | NA |
| *PPP2R5A* | 0.396011 | 0.180896 | 2.189162 | 0.028585067 | 0.318566531 |
| *SUGCT* | -2.539144 | 1.160063 | -2.188799 | 0.028611477 | NA |
| *TNFRSF21* | -2.074996 | 0.948378 | -2.187941 | 0.028673934 | NA |
| *WASF1* | -0.432681 | 0.197836 | -2.187068 | 0.028737524 | 0.319895764 |
| *DIRAS1* | -0.545263 | 0.249381 | -2.186463 | 0.028781761 | 0.320018658 |
| *MGST3* | 0.351708 | 0.160893 | 2.185976 | 0.028817394 | 0.320045709 |
| *DUSP22* | -0.187738 | 0.085928 | -2.184838 | 0.028900762 | 0.320236933 |
| *SRPRA* | -0.274886 | 0.125815 | -2.184834 | 0.028901051 | 0.320236933 |
| *CYTH3* | -0.634104 | 0.290363 | -2.183833 | 0.028974494 | 0.320682115 |
| *TTC9* | 0.34512 | 0.158134 | 2.182453 | 0.029076103 | 0.321102803 |
| *P4HB* | -0.16099 | 0.073767 | -2.182412 | 0.029079123 | 0.321102803 |
| *PCED1B* | 0.531251 | 0.243586 | 2.180963 | 0.029186142 | 0.321222433 |
| *JAK3* | -0.310337 | 0.1423 | -2.180868 | 0.0291932 | 0.321222433 |
| *ASPSCR1* | -0.314458 | 0.144194 | -2.1808 | 0.029198206 | 0.321222433 |
| *NDUFC1* | 0.190264 | 0.087258 | 2.180462 | 0.029223244 | 0.321222433 |
| *ZNF362* | -0.584645 | 0.268197 | -2.179907 | 0.029264386 | 0.321308292 |
| *IFI27L2* | 0.235215 | 0.107977 | 2.178392 | 0.029376845 | 0.322081135 |
| *HINFP* | 0.265841 | 0.122054 | 2.17806 | 0.029401597 | 0.322081135 |
| *HSPD1* | 0.219025 | 0.10068 | 2.175448 | 0.029596526 | 0.323536399 |
| *PHF12* | 0.28942 | 0.133055 | 2.175191 | 0.029615818 | 0.323536399 |
| *ZNF614* | 0.494253 | 0.22725 | 2.174933 | 0.029635129 | 0.323536399 |
| *RHOBTB1* | -1.194767 | 0.549635 | -2.173745 | 0.029724322 | NA |
| *NDUFB3* | 0.168528 | 0.077612 | 2.17142 | 0.029899459 | 0.326052921 |
| *MCM6* | 0.604955 | 0.278714 | 2.170522 | 0.029967317 | 0.326179362 |
| *RUVBL2* | -0.281551 | 0.129748 | -2.169989 | 0.030007679 | 0.326179362 |
| *CTDP1* | -0.645131 | 0.297306 | -2.169925 | 0.030012562 | 0.326179362 |
| *ENSG00000261386* | 0.491195 | 0.226496 | 2.168673 | 0.030107496 | 0.326842631 |
| *MDM2* | -0.253873 | 0.117131 | -2.167434 | 0.030201805 | 0.327497638 |
| *KLHDC2* | -0.202737 | 0.093572 | -2.166656 | 0.030261119 | 0.327772116 |
| *APH1A* | -0.123338 | 0.056942 | -2.166041 | 0.030308019 | 0.327911672 |
| *SNHG25* | -0.327898 | 0.151433 | -2.165305 | 0.030364299 | 0.328022832 |
| *CASP10* | 0.495096 | 0.228686 | 2.164966 | 0.030390327 | 0.328022832 |
| *FBXO44* | -0.301516 | 0.139297 | -2.16455 | 0.030422141 | 0.328022832 |
| *ZNF43* | -0.290612 | 0.134324 | -2.163521 | 0.03050115 | 0.328022832 |
| *UQCRH* | 0.142912 | 0.06606 | 2.163371 | 0.030512623 | 0.328022832 |
| *LENG8* | -0.232076 | 0.107282 | -2.163243 | 0.030522457 | 0.328022832 |
| *JHY* | -0.639168 | 0.295508 | -2.162944 | 0.030545483 | NA |
| *RMDN2* | -0.496016 | 0.229383 | -2.162396 | 0.030587646 | 0.328165501 |
| *WDR81* | -0.416668 | 0.192707 | -2.162186 | 0.030603816 | 0.328165501 |
| *TSTD1* | -0.233083 | 0.107843 | -2.161306 | 0.030671698 | 0.328500971 |
| *ENSG00000268352* | 1.133751 | 0.524806 | 2.160325 | 0.030747489 | NA |

| | | | | | |
|---|---|---|---|---|---|
| *WNT10A* | -0.564208 | 0.261191 | -2.160139 | 0.030761908 | 0.328500971 |
| *SEMA4F* | 0.688794 | 0.318923 | 2.159748 | 0.030792158 | 0.328500971 |
| *MIR22HG* | 0.440212 | 0.203829 | 2.159712 | 0.030794943 | 0.328500971 |
| *PDCD6IP* | -0.17373 | 0.080459 | -2.159241 | 0.03083147 | 0.328500971 |
| *ENSG00000262873* | -1.359977 | 0.629854 | -2.159193 | 0.030835195 | NA |
| *PLAGL1* | -0.351309 | 0.162708 | -2.159137 | 0.030839562 | 0.328500971 |
| *SLC38A11* | 1.453132 | 0.673026 | 2.159102 | 0.030842251 | NA |
| *NDUFB5* | 0.151574 | 0.070255 | 2.157489 | 0.03096756 | 0.328995326 |
| *PTPRA* | -0.498999 | 0.231299 | -2.157376 | 0.030976378 | 0.328995326 |
| *COG3* | -0.415359 | 0.192543 | -2.157222 | 0.030988356 | 0.328995326 |
| *BMPR2* | 0.311729 | 0.144741 | 2.153704 | 0.031263397 | 0.331550222 |
| *PNPLA4* | 0.467974 | 0.217361 | 2.152976 | 0.031320598 | 0.331791826 |
| *RBL1* | 0.421783 | 0.196139 | 2.150429 | 0.031521267 | 0.33250536 |
| *N6AMT1* | 0.510091 | 0.2373 | 2.149556 | 0.031590336 | 0.33250536 |
| *NUDCD3* | -0.2412 | 0.112228 | -2.149203 | 0.031618341 | 0.33250536 |
| *ENSG00000131408* | 0.148525 | 0.069108 | 2.149173 | 0.031620649 | 0.33250536 |
| *DUSP11* | 0.166521 | 0.077488 | 2.149009 | 0.031633695 | 0.33250536 |
| *BABAM2* | -0.322384 | 0.150029 | -2.148817 | 0.031648902 | 0.33250536 |
| *TMEM123* | -0.229511 | 0.106841 | -2.14816 | 0.03170103 | 0.33250536 |
| *GRAMD1A* | -0.472477 | 0.219955 | -2.148063 | 0.031708723 | 0.33250536 |
| *VPS16* | -0.322287 | 0.150055 | -2.1478 | 0.03172965 | 0.33250536 |
| *SCYL2* | -0.287665 | 0.133949 | -2.147576 | 0.031747423 | 0.33250536 |
| *GCSAM* | -0.569786 | 0.265347 | -2.147326 | 0.031767369 | 0.33250536 |
| *ENSG00000260572* | -1.731671 | 0.80655 | -2.147011 | 0.03179242 | NA |
| *ZFAND2A* | -0.390162 | 0.181784 | -2.146296 | 0.03184935 | 0.333001879 |
| *C3orf52* | -0.719095 | 0.33516 | -2.145525 | 0.031910913 | NA |
| *ACAT1* | -0.36852 | 0.171796 | -2.145104 | 0.031944554 | 0.333631524 |
| *SWAP70* | 0.181967 | 0.084846 | 2.144676 | 0.031978789 | 0.333631524 |
| *CCL4* | -0.87465 | 0.408022 | -2.143635 | 0.032062137 | 0.334139457 |
| *CHAF1A* | 0.583891 | 0.272439 | 2.143197 | 0.032097269 | 0.334144359 |
| *KIAA1324L* | 0.863807 | 0.403119 | 2.142809 | 0.032128464 | NA |
| *PEA15* | -0.33333 | 0.155612 | -2.142062 | 0.032188525 | 0.334478349 |
| *IL13RA1* | -0.51055 | 0.238359 | -2.141935 | 0.032198746 | 0.334478349 |
| *ENSG00000274184* | 0.486035 | 0.226965 | 2.141451 | 0.032237693 | 0.334511165 |
| *BRF2* | -0.431648 | 0.201607 | -2.141034 | 0.032271305 | 0.334511165 |
| *RIF1* | 0.230304 | 0.107631 | 2.139765 | 0.032373768 | 0.335133245 |
| *APMAP* | -0.327311 | 0.15299 | -2.13943 | 0.032400849 | 0.335133245 |
| *CROCC* | -0.620521 | 0.290353 | -2.137126 | 0.032587737 | NA |
| *TOMM20* | 0.17154 | 0.080294 | 2.136408 | 0.032646168 | 0.337308745 |
| *TLR4* | -1.950021 | 0.912977 | -2.135893 | 0.032688167 | NA |
| *NFE2L2* | 0.202254 | 0.094742 | 2.134799 | 0.032777432 | 0.338093876 |
| *AKIRIN2* | 0.224006 | 0.10494 | 2.134617 | 0.0327923 | 0.338093876 |
| *IGHG3* | 1.169622 | 0.548233 | 2.133439 | 0.032888704 | 0.338725544 |
| *ENSG00000268205* | 0.2713 | 0.127204 | 2.132797 | 0.032941375 | 0.338905931 |
| *IRGQ* | 0.496552 | 0.233017 | 2.13097 | 0.033091582 | 0.340088324 |
| *RIMKLB* | -0.321313 | 0.15092 | -2.12903 | 0.033251785 | 0.340730747 |
| *NOL10* | -0.257202 | 0.12083 | -2.128635 | 0.033284506 | 0.340730747 |

| | | | | | |
|---|---|---|---|---|---|
| *CLN5* | -0.275445 | 0.129401 | -2.128615 | 0.033286093 | 0.340730747 |
| *STRBP* | -0.151441 | 0.071162 | -2.128112 | 0.03332782 | 0.340730747 |
| *AGO1* | -0.224371 | 0.105435 | -2.128053 | 0.033332659 | 0.340730747 |
| *ADGRG5* | 1.028748 | 0.483504 | 2.127691 | 0.033362678 | NA |
| *ZNHIT3* | 0.180843 | 0.084996 | 2.127649 | 0.033366164 | 0.340730747 |
| *ITPR3* | 0.469352 | 0.220695 | 2.126699 | 0.033445117 | 0.341066389 |
| *NR1D2* | 0.316516 | 0.14885 | 2.126402 | 0.033469793 | 0.341066389 |
| *MT.ATP8* | -0.431748 | 0.203106 | -2.125728 | 0.033525947 | 0.341277851 |
| *CCDC82* | 0.232357 | 0.109361 | 2.124671 | 0.033614068 | 0.34135726 |
| *PCYOX1* | 0.328305 | 0.154531 | 2.124524 | 0.033626334 | 0.34135726 |
| *ISG15* | 0.229931 | 0.108236 | 2.124354 | 0.033640547 | 0.34135726 |
| *ITGA6.AS1* | -0.495875 | 0.233503 | -2.123634 | 0.033700764 | 0.34135726 |
| *EML3* | -0.272171 | 0.128189 | -2.123198 | 0.03373728 | 0.34135726 |
| *HMOX2* | 0.255038 | 0.120126 | 2.123091 | 0.03374621 | 0.34135726 |
| *ENSG00000237499* | -0.523609 | 0.246705 | -2.122407 | 0.033803606 | 0.341579415 |
| *IFT22* | -0.244022 | 0.115025 | -2.121464 | 0.033882738 | 0.34165741 |
| *C12orf49* | -0.490729 | 0.231354 | -2.121118 | 0.03391191 | 0.34165741 |
| *DCXR* | -0.247531 | 0.116712 | -2.120883 | 0.033931666 | 0.34165741 |
| *SRD5A1* | -0.32284 | 0.15227 | -2.120187 | 0.033990239 | 0.34165741 |
| *AGPAT4* | -0.753415 | 0.355384 | -2.120002 | 0.034005842 | NA |
| *ADAL* | -0.285006 | 0.134447 | -2.119837 | 0.034019815 | 0.34165741 |
| *ZEB1.AS1* | -0.463689 | 0.21878 | -2.119433 | 0.034053864 | 0.34165741 |
| *COMMD2* | -0.171732 | 0.08103 | -2.119368 | 0.034059416 | 0.34165741 |
| *ANKS6* | -1.068145 | 0.504234 | -2.118349 | 0.034145514 | NA |
| *PPIL3* | 0.36322 | 0.171525 | 2.117598 | 0.034209148 | 0.342802688 |
| *LYAR* | 0.418968 | 0.198021 | 2.115777 | 0.034363801 | 0.343994852 |
| *ZFAS1* | 0.246542 | 0.11655 | 2.115331 | 0.034401704 | 0.344017042 |
| *HTR3A* | -0.790885 | 0.374104 | -2.114077 | 0.034508678 | NA |
| *SH3RF1* | -1.817151 | 0.860022 | -2.112912 | 0.034608307 | NA |
| *ENSG00000251661* | -1.520032 | 0.720442 | -2.10986 | 0.034870382 | NA |
| *UTP6* | 0.232646 | 0.11027 | 2.109787 | 0.034876721 | 0.348037981 |
| *SNX2* | -0.20963 | 0.09937 | -2.109595 | 0.034893258 | 0.348037981 |
| *OTUD6B* | -0.399278 | 0.189287 | -2.109376 | 0.034912109 | 0.348037981 |
| *FGFBP2* | -2.663971 | 1.263113 | -2.109052 | 0.034940057 | NA |
| *MED18* | 0.435071 | 0.206392 | 2.107983 | 0.03503243 | 0.348876681 |
| *ZNF33A* | 0.182803 | 0.086747 | 2.107303 | 0.035091355 | 0.349102852 |
| *GLMN* | -0.323939 | 0.153794 | -2.106321 | 0.035176519 | 0.349589326 |
| *SPRYD7* | 0.368192 | 0.175008 | 2.103862 | 0.035390477 | 0.351353448 |
| *CD52* | 0.245508 | 0.116819 | 2.101615 | 0.035586984 | 0.352940869 |
| *TBCA* | 0.153523 | 0.073072 | 2.100996 | 0.035641284 | 0.353116116 |
| *AEBP1* | -0.441841 | 0.210384 | -2.100163 | 0.035714506 | 0.353354196 |
| *SVBP* | -0.154407 | 0.073531 | -2.099889 | 0.035738625 | 0.353354196 |
| *CD3D* | -2.240022 | 1.067439 | -2.098502 | 0.035860853 | NA |
| *GTF2E2* | 0.226854 | 0.108118 | 2.098203 | 0.035887191 | 0.354459552 |
| *RHEBL1* | 0.399921 | 0.190718 | 2.096922 | 0.036000449 | 0.355023388 |
| *CETN3* | 0.264037 | 0.125928 | 2.096725 | 0.036017933 | 0.355023388 |
| *FOXN2* | 0.368216 | 0.175699 | 2.095717 | 0.036107271 | 0.355540438 |

| | | | | | |
|---|---|---|---|---|---|
| *DNAAF2* | 0.221248 | 0.105684 | 2.093499 | 0.036304609 | 0.357118805 |
| *ZBTB11.AS1* | -0.541334 | 0.258638 | -2.093017 | 0.036347644 | 0.357177668 |
| *GNMT* | -1.446775 | 0.691298 | -2.092838 | 0.036363628 | NA |
| *ABCG1* | -0.606045 | 0.289621 | -2.092546 | 0.036389738 | 0.357227161 |
| *FXYD7* | 0.551685 | 0.263748 | 2.091715 | 0.036464013 | 0.357248238 |
| *LINC02193* | -0.657285 | 0.314268 | -2.091475 | 0.036485482 | 0.357248238 |
| *VPS52* | -0.440399 | 0.210588 | -2.091279 | 0.036503062 | 0.357248238 |
| *LAMP1* | -0.366507 | 0.175348 | -2.090168 | 0.03660274 | 0.357860457 |
| *IRAK3* | 0.712035 | 0.340891 | 2.088747 | 0.036730473 | NA |
| *CD40* | -0.1748 | 0.083698 | -2.088466 | 0.036755841 | 0.358212414 |
| *GATD3B* | -1.252563 | 0.599814 | -2.088254 | 0.036774911 | NA |
| *XRN2* | -0.174252 | 0.083469 | -2.087616 | 0.036832523 | 0.358212414 |
| *PLEKHG1* | -0.259257 | 0.124216 | -2.08715 | 0.036874572 | 0.358212414 |
| *IKBKE* | -0.516027 | 0.247294 | -2.0867 | 0.036915298 | 0.358212414 |
| *SIPA1L3* | -0.240402 | 0.115209 | -2.086669 | 0.036918073 | 0.358212414 |
| *SLC35D1* | 0.494818 | 0.237195 | 2.086121 | 0.036967638 | 0.358212414 |
| *DNAJC18* | -0.395655 | 0.18967 | -2.086022 | 0.036976639 | 0.358212414 |
| *DEGS2* | 0.94486 | 0.452961 | 2.085963 | 0.036981996 | NA |
| *AVPI1* | 0.753446 | 0.361215 | 2.085866 | 0.036990784 | NA |
| *GART* | -0.30725 | 0.14733 | -2.085453 | 0.037028244 | 0.358212414 |
| *PIK3CD.AS2* | 0.947144 | 0.454213 | 2.085243 | 0.03704727 | NA |
| *FAM177B* | 0.453161 | 0.217371 | 2.08473 | 0.037093774 | 0.358212414 |
| *ARMH3* | -0.371897 | 0.178398 | -2.084653 | 0.037100808 | 0.358212414 |
| *METTL8* | 0.461096 | 0.221219 | 2.084344 | 0.037128857 | 0.358212414 |
| *BORCS6* | 0.242596 | 0.116396 | 2.084222 | 0.037139943 | 0.358212414 |
| *RTL8C* | -0.312748 | 0.150065 | -2.084081 | 0.037152811 | 0.358212414 |
| *SEC61A2* | -0.402739 | 0.193252 | -2.084013 | 0.037158964 | 0.358212414 |
| *SMAD5* | 0.25151 | 0.12083 | 2.081522 | 0.03738613 | 0.359651966 |
| *EXOG* | -0.243059 | 0.116775 | -2.081437 | 0.037393882 | 0.359651966 |
| *LYSMD3* | 0.333254 | 0.160145 | 2.080952 | 0.037438271 | 0.359651966 |
| *LDLRAP1* | 0.360362 | 0.173189 | 2.080742 | 0.037457528 | 0.359651966 |
| *DOK7* | 1.245203 | 0.598643 | 2.080041 | 0.037521788 | NA |
| *FAIM* | 0.346979 | 0.166828 | 2.079861 | 0.037538319 | 0.360010554 |
| *ORMDL3* | -0.3354 | 0.161359 | -2.078599 | 0.037654234 | 0.360010554 |
| *MAP3K5* | 0.489627 | 0.235604 | 2.078181 | 0.037692693 | 0.360010554 |
| *ZFAND6* | 0.253796 | 0.122136 | 2.077986 | 0.03771064 | 0.360010554 |
| *IRF2BP2* | -0.269366 | 0.129639 | -2.077817 | 0.037726179 | 0.360010554 |
| *COL19A1* | -0.341599 | 0.16442 | -2.077597 | 0.03774651 | 0.360010554 |
| *TBC1D20* | 0.240144 | 0.115609 | 2.077212 | 0.037781985 | 0.360010554 |
| *DYNLT3* | 0.218709 | 0.105315 | 2.076712 | 0.037828174 | 0.360010554 |
| *PEX6* | 0.593562 | 0.285877 | 2.076283 | 0.03786779 | 0.360010554 |
| *ZNF100* | -0.424713 | 0.204555 | -2.076277 | 0.03786833 | 0.360010554 |
| *WIPF2* | 0.227145 | 0.109456 | 2.07521 | 0.037967092 | 0.360292921 |
| *NUP50* | 0.22539 | 0.108614 | 2.075148 | 0.037972781 | 0.360292921 |
| *TNRC6B* | 0.126742 | 0.061135 | 2.073168 | 0.038156681 | 0.361302189 |
| *RGS14* | 0.317448 | 0.153147 | 2.072823 | 0.038188714 | 0.361302189 |
| *EMC1* | 0.374244 | 0.180564 | 2.07264 | 0.038205824 | 0.361302189 |

| | | | | | |
|---|---|---|---|---|---|
| *LPCAT4* | 0.26755 | 0.129102 | 2.07239 | 0.03822907 | 0.361302189 |
| *CGAS* | 0.255697 | 0.123428 | 2.071625 | 0.038300421 | 0.361621999 |
| *ZKSCAN3* | 0.668229 | 0.322599 | 2.071392 | 0.03832219 | NA |
| *PHOSPHO2* | 0.40115 | 0.193859 | 2.069292 | 0.038518716 | 0.363327221 |
| *ENSG00000258017* | 0.776977 | 0.375602 | 2.068618 | 0.03858198 | NA |
| *STRIP1* | -0.316818 | 0.153232 | -2.067578 | 0.038679683 | 0.364360472 |
| *PIP4K2A* | 0.372118 | 0.180022 | 2.067066 | 0.038727923 | 0.364360472 |
| *LINC00663* | 0.66765 | 0.323035 | 2.066801 | 0.038752876 | NA |
| *BMT2* | -0.404984 | 0.195966 | -2.066604 | 0.038771443 | 0.364360472 |
| *UBE2E1* | 0.211873 | 0.102527 | 2.06652 | 0.038779444 | 0.364360472 |
| *ENSG00000080031* | -1.663115 | 0.805292 | -2.065232 | 0.038901086 | NA |
| *ATG16L2* | -0.219452 | 0.10626 | -2.065228 | 0.038901413 | 0.365150558 |
| *TMED4* | -0.203024 | 0.098333 | -2.064656 | 0.038955546 | 0.365302981 |
| *ZNF79* | 0.76227 | 0.36948 | 2.063088 | 0.039104261 | NA |
| *NFIX* | 0.439114 | 0.212898 | 2.062556 | 0.039154826 | 0.366814894 |
| *HIST1H2BJ* | 0.498825 | 0.241957 | 2.061628 | 0.039243172 | 0.36728561 |
| *HIST1H3H* | 1.261566 | 0.6121 | 2.061046 | 0.039298658 | NA |
| *PPL* | 1.098883 | 0.533565 | 2.059513 | 0.039445164 | NA |
| *DGCR6* | -0.997567 | 0.484372 | -2.059506 | 0.039445775 | NA |
| *SYNGAP1.AS1* | 0.675878 | 0.328217 | 2.059242 | 0.039471051 | NA |
| *ZNF627* | -0.497227 | 0.241558 | -2.058417 | 0.039550093 | 0.369434699 |
| *BCAT1* | -1.482462 | 0.720279 | -2.058178 | 0.03957308 | NA |
| *MRPS18B* | 0.145754 | 0.070831 | 2.057783 | 0.039610934 | 0.369434699 |
| *ABHD6* | 0.46755 | 0.227253 | 2.057397 | 0.039648092 | 0.369434699 |
| *CARD11* | 0.365937 | 0.177865 | 2.057381 | 0.039649632 | 0.369434699 |
| *PITPNA* | 0.332484 | 0.161618 | 2.057227 | 0.03966441 | 0.369434699 |
| *CAPN12* | 0.400382 | 0.194839 | 2.054935 | 0.039885236 | 0.371132893 |
| *ITGA10* | 0.549348 | 0.267499 | 2.053647 | 0.040009909 | 0.371605004 |
| *GGCT* | 0.24197 | 0.117839 | 2.053399 | 0.040033856 | 0.371605004 |
| *ARID5B* | -0.195328 | 0.095133 | -2.053216 | 0.040051618 | 0.371605004 |
| *CALHM2* | 0.618803 | 0.301454 | 2.052729 | 0.040098858 | 0.37168557 |
| *PNO1* | -0.33366 | 0.162622 | -2.05175 | 0.040193924 | 0.37211293 |
| *LSM10* | 0.14234 | 0.069385 | 2.05146 | 0.040222165 | 0.37211293 |
| *R3HDM2* | -0.327076 | 0.159471 | -2.050999 | 0.040267061 | 0.372171115 |
| *DGLUCY* | 0.247904 | 0.120963 | 2.049412 | 0.040421816 | 0.373243592 |
| *AIDA* | 0.216109 | 0.105555 | 2.047354 | 0.040623352 | 0.374422661 |
| *GMNN* | 0.481236 | 0.235087 | 2.047059 | 0.040652316 | 0.374422661 |
| *TRIB3* | 0.409721 | 0.200165 | 2.046919 | 0.04066603 | 0.374422661 |
| *KLF12* | 0.294009 | 0.143787 | 2.044753 | 0.040879179 | 0.376026033 |
| *ZKSCAN2* | -0.60906 | 0.29805 | -2.043483 | 0.041004684 | NA |
| *ZNRD2* | -0.206959 | 0.101301 | -2.04301 | 0.041051476 | 0.377250934 |
| *IGHV3.15* | -1.016542 | 0.498042 | -2.041075 | 0.041243408 | NA |
| *MXD1* | 0.425268 | 0.208379 | 2.040839 | 0.041266851 | 0.378516777 |
| *ARL2* | 0.328917 | 0.161168 | 2.04083 | 0.041267752 | 0.378516777 |
| *AMIGO2* | 0.920519 | 0.451139 | 2.040434 | 0.041307098 | NA |
| *ATP5PD* | -0.127802 | 0.062642 | -2.040192 | 0.041331187 | 0.378738257 |
| *CFP* | -1.647387 | 0.807652 | -2.039722 | 0.041377992 | NA |

| | | | | | |
|---|---|---|---|---|---|
| *MTPN* | 0.156917 | 0.076933 | 2.039661 | 0.041384063 | 0.378862645 |
| *STYX* | 0.296963 | 0.145687 | 2.038365 | 0.041513474 | 0.379441949 |
| *STX5* | -0.182042 | 0.089313 | -2.038239 | 0.041526064 | 0.379441949 |
| *MRPS21* | -0.156072 | 0.076612 | -2.037184 | 0.041631624 | 0.380046262 |
| *ACTR1B* | -0.193178 | 0.094873 | -2.036171 | 0.041733165 | 0.380330394 |
| *GSE1* | 0.447144 | 0.219673 | 2.035502 | 0.041800339 | 0.380330394 |
| *VPS25* | -0.240741 | 0.118278 | -2.035375 | 0.041813116 | 0.380330394 |
| *PLA2G6* | -0.463567 | 0.227767 | -2.035266 | 0.041824122 | 0.380330394 |
| *PLOD3* | 0.334642 | 0.16445 | 2.034909 | 0.041860015 | 0.380330394 |
| *TMEM167B* | -0.189311 | 0.093065 | -2.034171 | 0.041934349 | 0.380647006 |
| *TNFSF10* | 0.3564 | 0.175266 | 2.033483 | 0.042003761 | 0.380818631 |
| *ARF5* | -0.242767 | 0.119403 | -2.033167 | 0.042035656 | 0.380818631 |
| *CLTC* | 0.233343 | 0.114788 | 2.03281 | 0.042071768 | 0.380818631 |
| *POGLUT3* | 0.900104 | 0.442831 | 2.032614 | 0.042091527 | NA |
| *AKIRIN1* | 0.164488 | 0.080935 | 2.032359 | 0.042117346 | 0.380873559 |
| *MYL6B* | -0.211233 | 0.104042 | -2.030265 | 0.042329592 | 0.381482826 |
| *SRSF2* | -0.121404 | 0.059802 | -2.030084 | 0.042348022 | 0.381482826 |
| *MAX* | 0.176452 | 0.086925 | 2.029932 | 0.042363455 | 0.381482826 |
| *SF3B2* | -0.13961 | 0.068779 | -2.029824 | 0.042374406 | 0.381482826 |
| *GABPB2* | 0.344738 | 0.169843 | 2.029739 | 0.042383084 | 0.381482826 |
| *LINC00954* | -0.326522 | 0.1609 | -2.029355 | 0.042422157 | 0.381482826 |
| *SEPHS1* | 0.416583 | 0.205362 | 2.028534 | 0.042505733 | 0.381878162 |
| *SLFN12* | 0.724823 | 0.357548 | 2.027205 | 0.042641422 | NA |
| *POLM* | -0.221996 | 0.109524 | -2.02691 | 0.042671652 | 0.382657319 |
| *ENSG00000245869* | 0.466044 | 0.229949 | 2.026726 | 0.042690466 | 0.382657319 |
| *BAD* | -0.192507 | 0.095003 | -2.026336 | 0.042730315 | 0.382657319 |
| *MAP3K2* | 0.206896 | 0.102114 | 2.026132 | 0.042751238 | 0.382657319 |
| *AKAP10* | 0.196495 | 0.097061 | 2.024448 | 0.042924079 | 0.383669117 |
| *VCPIP1* | 0.260831 | 0.128853 | 2.024255 | 0.042943877 | 0.383669117 |
| *ENSG00000257194* | -1.412965 | 0.698482 | -2.02291 | 0.043082435 | NA |
| *NKG7* | -1.799232 | 0.889561 | -2.022607 | 0.043113729 | NA |
| *ERLIN1* | 0.506621 | 0.25053 | 2.022192 | 0.043156495 | 0.384793427 |
| *RIPOR1* | -0.190794 | 0.094387 | -2.021403 | 0.043238086 | 0.384793427 |
| *SQOR* | -0.257544 | 0.127419 | -2.021239 | 0.043254988 | 0.384793427 |
| *FAM71D* | -0.598012 | 0.295874 | -2.021173 | 0.04326183 | 0.384793427 |
| *KCTD7* | 0.317491 | 0.157088 | 2.021101 | 0.043269302 | 0.384793427 |
| *MYLK.AS1* | -0.541303 | 0.267974 | -2.019986 | 0.043384825 | 0.385465171 |
| *ESYT2* | 0.434154 | 0.214972 | 2.019587 | 0.043426248 | 0.385477927 |
| *ACSM3* | -0.410769 | 0.203452 | -2.019001 | 0.043487097 | 0.385662937 |
| *MIR4435.2HG* | 1.141831 | 0.565865 | 2.017851 | 0.043606798 | NA |
| *ENSG00000273002* | 0.554073 | 0.274661 | 2.017296 | 0.043664591 | 0.385952631 |
| *HLA.DMA* | -0.262497 | 0.130157 | -2.016771 | 0.043719396 | 0.385952631 |
| *UBL4A* | 0.219545 | 0.10886 | 2.016768 | 0.043719686 | 0.385952631 |
| *FAM8A1* | 0.571606 | 0.283427 | 2.016765 | 0.043720075 | NA |
| *IDH1* | 0.257305 | 0.127583 | 2.016764 | 0.043720115 | 0.385952631 |
| *SEMA4A* | -1.040722 | 0.516097 | -2.016523 | 0.04374532 | NA |
| *TIPARP* | -0.233262 | 0.115711 | -2.015911 | 0.043809267 | 0.385952631 |

| | | | | | |
|---|---|---|---|---|---|
| *PCF11* | 0.162865 | 0.080813 | 2.015338 | 0.043869231 | 0.385952631 |
| *PWWP2B* | 0.59519 | 0.295333 | 2.015314 | 0.043871735 | 0.385952631 |
| *VPS37B* | 0.196684 | 0.097596 | 2.015282 | 0.043875119 | 0.385952631 |
| *ANXA6* | -0.241566 | 0.11987 | -2.015234 | 0.043880092 | 0.385952631 |
| *CISD2* | 0.213511 | 0.106075 | 2.012819 | 0.044133717 | 0.387829566 |
| *TCOF1* | -0.240661 | 0.11963 | -2.011711 | 0.044250368 | 0.388500498 |
| *EIF2B5* | -0.226047 | 0.11243 | -2.010552 | 0.044372759 | 0.389220558 |
| *MBOAT2* | -1.046021 | 0.520485 | -2.009706 | 0.044462273 | NA |
| *FUOM* | -0.390634 | 0.194461 | -2.008805 | 0.044557868 | 0.39048895 |
| *SAMD10* | 0.579118 | 0.288325 | 2.008563 | 0.044583475 | NA |
| *DRG2* | -0.230472 | 0.114765 | -2.008201 | 0.044621947 | 0.390570353 |
| *PPP2R5B* | 0.255491 | 0.127239 | 2.007954 | 0.044648188 | 0.390570353 |
| *PLXNC1* | 0.813193 | 0.405154 | 2.007121 | 0.044736832 | NA |
| *DAZAP1* | -0.275083 | 0.137084 | -2.006673 | 0.044784472 | 0.391057623 |
| *ENSG00000186019* | 0.950143 | 0.473598 | 2.006221 | 0.044832631 | NA |
| *ZNF417* | 0.545855 | 0.272149 | 2.005721 | 0.044886006 | 0.391057623 |
| *MORC3* | 0.207808 | 0.103612 | 2.005637 | 0.044895037 | 0.391057623 |
| *SRSF5* | -0.182219 | 0.090872 | -2.005221 | 0.044939414 | 0.391057623 |
| *TCEAL4* | 0.243137 | 0.121256 | 2.005151 | 0.044946876 | 0.391057623 |
| *SYNE2* | -0.288137 | 0.143706 | -2.005046 | 0.044958121 | 0.391057623 |
| *EVI2A* | 0.314154 | 0.156703 | 2.004768 | 0.044987853 | 0.391057623 |
| *RPSA* | 0.221315 | 0.110418 | 2.004343 | 0.045033306 | 0.391100066 |
| *ENSG00000236723* | 1.027524 | 0.512734 | 2.004011 | 0.045068887 | NA |
| *KMT2A* | 0.197977 | 0.098835 | 2.003109 | 0.04516557 | 0.391782285 |
| *NINJ1* | -0.435747 | 0.217563 | -2.002852 | 0.045193143 | 0.391782285 |
| *GLB1L* | 0.582102 | 0.290667 | 2.002641 | 0.045215866 | NA |
| *LAGE3* | 0.177089 | 0.088453 | 2.00206 | 0.045278325 | 0.392168059 |
| *AP1B1* | -0.344892 | 0.172356 | -2.001046 | 0.045387436 | 0.392760215 |
| *ERGIC1* | -0.206242 | 0.103088 | -2.000637 | 0.045431563 | 0.392789476 |
| *ECI2* | 0.220699 | 0.110356 | 1.999881 | 0.04551311 | 0.393141917 |
| *ENSG00000204802* | -0.710427 | 0.355302 | -1.9995 | 0.045554241 | NA |
| *LINC00667* | 0.222098 | 0.111156 | 1.998078 | 0.045708153 | 0.394473224 |
| *ENSG00000271869* | 0.358652 | 0.179622 | 1.996708 | 0.045856938 | 0.39540329 |
| *MAMLD1* | -1.021688 | 0.512056 | -1.995268 | 0.046013712 | NA |
| *ENSG00000256690* | 0.353092 | 0.17697 | 1.99521 | 0.046019959 | 0.395906872 |
| *ERO1A* | 0.310329 | 0.155565 | 1.994857 | 0.04605852 | 0.395906872 |
| *CXCR5* | 0.196094 | 0.098302 | 1.994816 | 0.046062988 | 0.395906872 |
| *ZEB1* | -0.270537 | 0.13563 | -1.994663 | 0.046079617 | 0.395906872 |
| *LRMP* | 0.271657 | 0.136286 | 1.993292 | 0.046229507 | 0.396640625 |
| *WASIR2* | 0.660196 | 0.331288 | 1.992814 | 0.046281782 | 0.396640625 |
| *CENPS.CORT* | -0.998298 | 0.500956 | -1.992785 | 0.046285048 | NA |
| *DPP3* | -0.353491 | 0.177388 | -1.992753 | 0.046288455 | 0.396640625 |
| *NMNAT3* | -1.011631 | 0.507856 | -1.991966 | 0.046374821 | NA |
| *GTF3C6* | -0.134324 | 0.067538 | -1.988874 | 0.046715099 | 0.399940993 |
| *AGAP3* | 1.647152 | 0.828234 | 1.988752 | 0.046728623 | NA |
| *POLR2H* | -0.232122 | 0.116739 | -1.988388 | 0.046768755 | 0.400045075 |
| *IP6K2* | -0.202639 | 0.102009 | -1.986479 | 0.046980149 | 0.400082168 |

| | | | | | |
|---|---|---|---|---|---|
| *MGME1* | 0.254364 | 0.128072 | 1.986106 | 0.047021568 | 0.400082168 |
| *ETFRF1* | 0.236662 | 0.119213 | 1.985204 | 0.047121758 | 0.400082168 |
| *DHRS3* | 2.263718 | 1.14047 | 1.984899 | 0.047155739 | 0.400082168 |
| *PAFAH1B1* | 0.14299 | 0.07205 | 1.984584 | 0.04719075 | 0.400082168 |
| *REEP4* | 0.309959 | 0.156187 | 1.984538 | 0.047195865 | 0.400082168 |
| *GDI2* | -0.168399 | 0.084872 | -1.984151 | 0.047239025 | 0.400082168 |
| *RAB11B* | -0.158072 | 0.079674 | -1.983982 | 0.047257833 | 0.400082168 |
| *FAAP24* | -0.493202 | 0.248631 | -1.98367 | 0.04729261 | 0.400082168 |
| *IRF5* | 0.316422 | 0.159554 | 1.98316 | 0.04734954 | 0.400082168 |
| *UBE4B* | 0.241019 | 0.121543 | 1.982997 | 0.047367796 | 0.400082168 |
| *XRCC4* | -0.353502 | 0.178274 | -1.982915 | 0.047376963 | 0.400082168 |
| *PLD4* | -0.587347 | 0.296209 | -1.982881 | 0.047380699 | 0.400082168 |
| *CAMTA1* | -0.163788 | 0.082613 | -1.982591 | 0.047413125 | 0.400082168 |
| *C1orf35* | 0.193305 | 0.097503 | 1.982563 | 0.047416246 | 0.400082168 |
| *ZBTB1* | 0.210769 | 0.106321 | 1.982376 | 0.047437128 | 0.400082168 |
| *ENSG00000257354* | -0.503089 | 0.253865 | -1.981721 | 0.047510479 | 0.400350543 |
| *IL27RA* | 0.213954 | 0.108007 | 1.980931 | 0.04759901 | 0.40074625 |
| *PITPNM2* | -0.709471 | 0.358294 | -1.980136 | 0.047688288 | NA |
| *ITPR1.DT* | 1.349532 | 0.682147 | 1.97836 | 0.047888136 | NA |
| *XXYLT1.AS2* | 0.800203 | 0.404714 | 1.977208 | 0.048018106 | 0.403921939 |
| *PPTC7* | 0.2526 | 0.127788 | 1.976714 | 0.048073963 | 0.403953132 |
| *USP40* | 0.576646 | 0.291794 | 1.976207 | 0.048131339 | 0.403953132 |
| *MAD2L2* | -0.181604 | 0.091902 | -1.976064 | 0.048147526 | 0.403953132 |
| *NDUFS6* | 0.12979 | 0.065717 | 1.974962 | 0.048272432 | 0.404626884 |
| *LINC02285* | -1.006182 | 0.509547 | -1.97466 | 0.048306707 | NA |
| *EEA1* | 0.260421 | 0.131898 | 1.974411 | 0.048335067 | 0.404626884 |
| *HIST4H4* | 0.548887 | 0.278005 | 1.974378 | 0.048338828 | NA |
| *C22orf46* | 0.393635 | 0.199418 | 1.973917 | 0.048391143 | 0.404626884 |
| *TMEM179B* | -0.174632 | 0.088485 | -1.973572 | 0.048430462 | 0.404626884 |
| *ATG5* | 0.181015 | 0.091736 | 1.973217 | 0.04847086 | 0.404626884 |
| *UBE2W* | 0.19221 | 0.097413 | 1.973139 | 0.048479673 | 0.404626884 |
| *CETP* | -0.812432 | 0.411995 | -1.971945 | 0.048615901 | NA |
| *CCDC152* | 0.381457 | 0.193442 | 1.97194 | 0.048616476 | 0.405417674 |
| *RPS6KB2.AS1* | -0.64505 | 0.327321 | -1.970699 | 0.048758353 | NA |
| *ENSG00000256591* | -0.716611 | 0.363722 | -1.970219 | 0.048813312 | NA |
| *PATJ* | 0.373495 | 0.189573 | 1.970191 | 0.048816479 | 0.406391041 |
| *KANK2* | -0.987851 | 0.5014 | -1.970184 | 0.048817282 | NA |
| *FYN* | 0.415787 | 0.21104 | 1.970182 | 0.048817513 | 0.406391041 |
| *TOLLIP.AS1* | -1.071128 | 0.543839 | -1.969569 | 0.04888778 | NA |
| *C1orf216* | 0.509095 | 0.258611 | 1.968571 | 0.049002377 | 0.407578013 |
| *ENSG00000267939* | -1.76805 | 0.898636 | -1.967482 | 0.049127677 | NA |
| *HIST2H2BE* | 0.413122 | 0.209987 | 1.967367 | 0.049140874 | 0.408138074 |
| *RPAP2* | 0.210956 | 0.107238 | 1.967171 | 0.049163504 | 0.408138074 |
| *ENSG00000272870* | 0.628355 | 0.319451 | 1.966987 | 0.049184663 | NA |
| *SEC13* | -0.255711 | 0.130008 | -1.966883 | 0.049196726 | 0.408138074 |
| *FAHD2B* | -0.853381 | 0.433929 | -1.966638 | 0.049225008 | NA |
| *HCCS* | 0.248304 | 0.126303 | 1.965935 | 0.049306076 | 0.40826996 |

| | | | | | |
|---|---|---|---|---|---|
| *E2F6* | 0.32729 | 0.166504 | 1.965659 | 0.049338002 | 0.40826996 |
| *INO80* | -0.271897 | 0.138327 | -1.96561 | 0.049343716 | 0.40826996 |
| *DUSP18* | 0.323468 | 0.164592 | 1.965278 | 0.04938203 | 0.40826996 |
| *C7orf61* | 0.847245 | 0.431158 | 1.965044 | 0.049409114 | NA |
| *ORAI2* | 0.136245 | 0.069343 | 1.964798 | 0.049437568 | 0.408378883 |
| *LPCAT2* | -1.153615 | 0.587204 | -1.96459 | 0.049461711 | NA |
| *RMND1* | -0.332451 | 0.169296 | -1.96372 | 0.04956259 | 0.409061104 |
| *PIGC* | -0.21394 | 0.109 | -1.962748 | 0.049675521 | 0.409563033 |
| *TRIM52* | 0.274312 | 0.139779 | 1.962465 | 0.049708376 | 0.409563033 |
| *CLIC1* | 0.185901 | 0.0948 | 1.960976 | 0.049881821 | 0.410385651 |
| *ADH5* | -0.19598 | 0.099945 | -1.960877 | 0.049893359 | 0.410385651 |

| Cluster 4 | log2FC | lfcSE | stat | pvalue | padj |
|---|---|---|---|---|---|
| *ARRDC3* | -1.093058 | 0.159098 | -6.870345 | 6.40E-12 | 3.13E-08 |
| *GPR183* | 0.96683 | 0.165013 | 5.859129 | 4.65E-09 | 8.55E-06 |
| *C7orf50* | -0.510511 | 0.087428 | -5.839218 | 5.24E-09 | 8.55E-06 |
| *SMIM10* | -3.760857 | 0.646237 | -5.819627 | 5.90E-09 | NA |
| *MT1X* | -1.838662 | 0.323822 | -5.678001 | 1.36E-08 | NA |
| *SYNGR2* | -0.76872 | 0.138034 | -5.569053 | 2.56E-08 | 3.13E-05 |
| *CLK1* | 0.558842 | 0.104823 | 5.331317 | 9.75E-08 | 9.54E-05 |
| *ENSG00000273319* | 0.725078 | 0.139862 | 5.184235 | 2.17E-07 | 0.000176885 |
| *LRIF1* | 0.552042 | 0.10815 | 5.104394 | 3.32E-07 | 0.000231967 |
| *ZBTB1* | 0.638495 | 0.127926 | 4.991123 | 6.00E-07 | 0.000332472 |
| *CYTIP* | 0.267219 | 0.053577 | 4.987538 | 6.12E-07 | 0.000332472 |
| *CD83* | -0.470277 | 0.095109 | -4.94463 | 7.63E-07 | 0.00037328 |
| *NFKBIA* | -0.499838 | 0.101575 | -4.920864 | 8.62E-07 | 0.000383268 |
| *CKS2* | 1.120976 | 0.229744 | 4.879249 | 1.06E-06 | 0.00041957 |
| *ZC3H6* | 0.698181 | 0.143357 | 4.87022 | 1.11E-06 | 0.00041957 |
| *CDKN1A* | -0.787616 | 0.165683 | -4.753741 | 2.00E-06 | 0.000697906 |
| *GATD3A* | 4.371743 | 0.928638 | 4.707696 | 2.51E-06 | NA |
| *CHMP6* | -0.771983 | 0.165038 | -4.677617 | 2.90E-06 | NA |
| *RPS5* | 0.328483 | 0.071666 | 4.583509 | 4.57E-06 | 0.001455455 |
| *CRIPT* | 0.523886 | 0.114507 | 4.575127 | 4.76E-06 | 0.001455455 |
| *CHORDC1* | 0.772773 | 0.171623 | 4.502747 | 6.71E-06 | 0.001930744 |
| *DBI* | -0.329261 | 0.073967 | -4.45147 | 8.53E-06 | 0.002293783 |
| *RPL8* | 0.467311 | 0.1052 | 4.442137 | 8.91E-06 | 0.002293783 |
| *PDE4B* | 0.388308 | 0.088625 | 4.381483 | 1.18E-05 | 0.002668076 |
| *PNPLA8* | 0.483716 | 0.110466 | 4.378877 | 1.19E-05 | 0.002668076 |
| *IZUMO4* | -0.787927 | 0.179988 | -4.377656 | 1.20E-05 | 0.002668076 |
| *IL16* | 0.325289 | 0.075833 | 4.289552 | 1.79E-05 | 0.003719356 |
| *H3F3B* | -0.381837 | 0.089241 | -4.278705 | 1.88E-05 | 0.003719356 |
| *GPR65* | 0.735149 | 0.171913 | 4.276289 | 1.90E-05 | 0.003719356 |
| *TTC32* | -0.737522 | 0.17261 | -4.272771 | 1.93E-05 | NA |
| *HHEX* | 0.299358 | 0.070253 | 4.261117 | 2.03E-05 | 0.003827968 |
| *CXCR4* | 0.233225 | 0.054846 | 4.252374 | 2.12E-05 | 0.003833151 |
| *MRPL54* | -0.374501 | 0.088664 | -4.223842 | 2.40E-05 | 0.004197022 |
| *TCTN1* | -0.818617 | 0.194231 | -4.214647 | 2.50E-05 | 0.004220955 |
| *TAGLN2* | -0.352291 | 0.084397 | -4.174201 | 2.99E-05 | 0.004877244 |
| *CD5* | -1.539502 | 0.370699 | -4.152975 | 3.28E-05 | NA |
| *ENSG00000272211* | -1.215828 | 0.293575 | -4.141449 | 3.45E-05 | 0.005447301 |
| *CRK* | 0.524754 | 0.12822 | 4.092613 | 4.27E-05 | 0.006377082 |
| *ZC3H12D* | -0.885557 | 0.216481 | -4.09069 | 4.30E-05 | 0.006377082 |
| *RPS9* | 0.367874 | 0.090905 | 4.046809 | 5.19E-05 | 0.007471984 |
| *TMEM243* | 0.368468 | 0.091278 | 4.036785 | 5.42E-05 | 0.007575573 |
| *HHIP.AS1* | -2.757367 | 0.684093 | -4.030691 | 5.56E-05 | NA |
| *SGK1* | -1.312415 | 0.326232 | -4.022949 | 5.75E-05 | NA |
| *NPM1* | 0.251972 | 0.062941 | 4.00333 | 6.25E-05 | 0.008488956 |
| *RHOH* | 0.313857 | 0.078709 | 3.987579 | 6.68E-05 | 0.008827378 |
| *ASF1A* | 0.516179 | 0.130493 | 3.955596 | 7.63E-05 | 0.009667011 |

| | | | | | |
|---|---|---|---|---|---|
| *CIRBP* | -0.221754 | 0.056092 | -3.95339 | 7.71E-05 | 0.009667011 |
| *CD79B* | -0.267773 | 0.068073 | -3.933594 | 8.37E-05 | 0.009909599 |
| *RPS23* | 0.201496 | 0.051273 | 3.929904 | 8.50E-05 | 0.009909599 |
| *LETMD1* | 0.396342 | 0.100899 | 3.928091 | 8.56E-05 | 0.009909599 |
| *RPL13A* | 0.189182 | 0.048211 | 3.924013 | 8.71E-05 | 0.009909599 |
| *SLC3A2* | -0.466697 | 0.11978 | -3.896275 | 9.77E-05 | 0.010862825 |
| *LINC01215* | -0.476098 | 0.124651 | -3.819456 | 0.00013375 | 0.014542706 |
| *LRRC61* | -0.99072 | 0.259955 | -3.811123 | 0.00013834 | NA |
| *RPL7A* | 0.183581 | 0.04829 | 3.801651 | 0.00014374 | 0.015044042 |
| *SNAI3* | 1.347743 | 0.3548 | 3.798596 | 0.00014552 | NA |
| *RPL30* | 0.208312 | 0.054928 | 3.79249 | 0.00014914 | 0.015044042 |
| *KDM4B* | 0.555616 | 0.146581 | 3.790499 | 0.00015035 | 0.015044042 |
| *CCDC191* | -1.283915 | 0.338765 | -3.789986 | 0.00015066 | 0.015044042 |
| *RPS8* | 0.262593 | 0.069939 | 3.754612 | 0.00017361 | 0.016989515 |
| *RSRP1* | 0.391818 | 0.105313 | 3.7205 | 0.00019883 | 0.019075832 |
| *ZBTB25* | 0.485386 | 0.1309 | 3.708074 | 0.00020884 | 0.019651197 |
| *ENSG00000261386* | 0.711354 | 0.19227 | 3.699765 | 0.0002158 | 0.019797206 |
| *CHCHD5* | -0.40761 | 0.110331 | -3.694435 | 0.00022038 | 0.019797206 |
| *TCL1B* | -1.245556 | 0.337736 | -3.687953 | 0.00022607 | 0.019797206 |
| *SLC50A1* | -0.497085 | 0.134807 | -3.687377 | 0.00022658 | 0.019797206 |
| *TNFRSF14* | 0.265524 | 0.072173 | 3.679004 | 0.00023415 | 0.020099615 |
| *KRTCAP2* | -0.320078 | 0.088092 | -3.633442 | 0.00027967 | 0.023153585 |
| *MIR29B2CHG* | 0.497632 | 0.136968 | 3.633198 | 0.00027993 | 0.023153585 |
| *FGFR1OP2* | 0.417302 | 0.114974 | 3.629547 | 0.00028392 | 0.023153585 |
| *RPL36* | 0.151161 | 0.041698 | 3.62516 | 0.00028878 | 0.023164186 |
| *ARPC3* | -0.207215 | 0.057507 | -3.603321 | 0.00031418 | 0.024794632 |
| *ZBED2* | 2.042977 | 0.571681 | 3.573633 | 0.00035206 | NA |
| *C16orf74* | -0.494382 | 0.138619 | -3.566491 | 0.00036179 | 0.028099316 |
| *ENSG00000261505* | -0.912922 | 0.256074 | -3.565066 | 0.00036377 | NA |
| *PPM1K* | 0.261894 | 0.074005 | 3.53886 | 0.00040186 | 0.030043256 |
| *PNRC2* | 0.280407 | 0.079258 | 3.5379 | 0.00040332 | 0.030043256 |
| *LPAR5* | 0.667879 | 0.188845 | 3.536646 | 0.00040524 | 0.030043256 |
| *C17orf49* | -0.973275 | 0.276885 | -3.515092 | 0.0004396 | 0.032104083 |
| *RPSA* | 0.302235 | 0.086137 | 3.508774 | 0.00045018 | 0.03239285 |
| *CD48* | 0.334635 | 0.095525 | 3.503137 | 0.00045981 | 0.032606736 |
| *NUP210* | 0.460743 | 0.132151 | 3.486482 | 0.00048942 | 0.033158573 |
| *PPP1CA* | -0.169175 | 0.048537 | -3.485508 | 0.0004912 | 0.033158573 |
| *RPL38* | 0.221852 | 0.063673 | 3.484257 | 0.00049351 | 0.033158573 |
| *CLEC2B* | 0.392925 | 0.112826 | 3.482572 | 0.00049662 | 0.033158573 |
| *RPL9* | 0.224606 | 0.064543 | 3.479965 | 0.00050148 | 0.033158573 |
| *PRICKLE1* | 0.71638 | 0.205928 | 3.478789 | 0.00050368 | NA |
| *FGR* | -0.939853 | 0.270302 | -3.477044 | 0.00050697 | NA |
| *TPI1* | -0.232038 | 0.06684 | -3.471533 | 0.0005175 | 0.033529992 |
| *RBM3* | -0.34656 | 0.099878 | -3.469823 | 0.0005208 | 0.033529992 |
| *EPS15* | -0.591244 | 0.171341 | -3.45069 | 0.00055916 | 0.035107659 |
| *RPL32* | 0.163843 | 0.047562 | 3.44486 | 0.00057136 | 0.035107659 |
| *ADA* | -0.701694 | 0.203741 | -3.444048 | 0.00057307 | 0.035107659 |

| | | | | | |
|---|---|---|---|---|---|
| *NDUFB5* | 0.297633 | 0.08643 | 3.443608 | 0.00057401 | 0.035107659 |
| *HMGB2* | -0.377199 | 0.109819 | -3.434741 | 0.00059312 | 0.035828879 |
| *PSMD8* | -0.245944 | 0.071851 | -3.422963 | 0.00061943 | 0.036727913 |
| *DYNLT1* | 0.503514 | 0.147166 | 3.421391 | 0.00062302 | 0.036727913 |
| *SCRN1* | 1.094996 | 0.320881 | 3.412465 | 0.00064378 | NA |
| *BLNK* | 0.269009 | 0.078995 | 3.405379 | 0.00066072 | 0.038262808 |
| *APH1A* | -0.27726 | 0.081472 | -3.40311 | 0.00066624 | 0.038262808 |
| *MCRIP1* | -0.267807 | 0.078754 | -3.400547 | 0.00067251 | 0.038262808 |
| *SCN3A* | 1.614904 | 0.475378 | 3.397093 | 0.00068106 | NA |
| *RPL3* | 0.191296 | 0.056317 | 3.39677 | 0.00068186 | 0.038348887 |
| *PFDN5* | 0.136899 | 0.040385 | 3.389816 | 0.0006994 | 0.038887962 |
| *GALNT2* | -0.573459 | 0.169338 | -3.386483 | 0.00070795 | 0.038921093 |
| *EIF4A2* | 0.271207 | 0.080172 | 3.382832 | 0.00071743 | 0.039003995 |
| *CLEC2D* | -0.324356 | 0.096008 | -3.378428 | 0.00072902 | 0.03919858 |
| *ENSG00000266844* | -1.940438 | 0.576394 | -3.366511 | 0.00076126 | NA |
| *NEIL1* | -0.495229 | 0.147137 | -3.365767 | 0.00076331 | 0.040582092 |
| *RPL24* | 0.224521 | 0.066764 | 3.362882 | 0.00077133 | 0.040582092 |
| *CELF1* | 0.264683 | 0.078986 | 3.350993 | 0.00080522 | 0.041914451 |
| *RNF43* | -1.01973 | 0.30488 | -3.344693 | 0.00082374 | NA |
| *CYB5A* | -0.795223 | 0.237863 | -3.343194 | 0.0008282 | 0.042656694 |
| *FAU* | 0.163075 | 0.04884 | 3.33896 | 0.00084093 | 0.042860988 |
| *ODF3B* | -0.700625 | 0.210202 | -3.333111 | 0.00085881 | NA |
| *TAGAP* | -0.414947 | 0.124867 | -3.323112 | 0.00089019 | 0.044904126 |
| *HHLA3* | 0.773717 | 0.232877 | 3.322436 | 0.00089235 | NA |
| *GPR18* | 0.607875 | 0.183046 | 3.320887 | 0.00089732 | NA |
| *CDKN1B* | 0.288132 | 0.087012 | 3.311391 | 0.00092833 | 0.046212037 |
| *NUFIP2* | 0.362653 | 0.109583 | 3.309386 | 0.00093501 | 0.046212037 |
| *WRB* | 0.554147 | 0.167931 | 3.299844 | 0.00096739 | NA |
| *ARL17A* | -0.642947 | 0.195193 | -3.293904 | 0.00098806 | 0.048345877 |
| *FXYD2* | -1.132432 | 0.344316 | -3.288932 | 0.00100568 | NA |
| *RMND5A* | 0.539333 | 0.164063 | 3.287352 | 0.00101134 | 0.048403941 |
| *RNF34* | 0.500672 | 0.152344 | 3.286454 | 0.00101457 | 0.048403941 |
| *TRAF4* | -0.361903 | 0.11016 | -3.285248 | 0.00101893 | 0.048403941 |
| *SRSF5* | -0.214314 | 0.06541 | -3.276474 | 0.00105112 | 0.049453151 |
| *PTPMT1* | -0.326816 | 0.099849 | -3.273094 | 0.00106377 | 0.049499404 |
| *TRMT13* | 0.40174 | 0.122825 | 3.270827 | 0.00107234 | 0.049499404 |
| *DNAJB4* | 0.795668 | 0.243279 | 3.270596 | 0.00107321 | NA |
| *ZNF250* | 0.744533 | 0.227668 | 3.27026 | 0.00107449 | NA |
| *LINC02201* | -1.671062 | 0.511111 | -3.269472 | 0.00107749 | NA |
| *ORMDL3* | -0.657683 | 0.201815 | -3.258837 | 0.0011187 | 0.051156917 |
| *SNX15* | -1.439564 | 0.443814 | -3.243623 | 0.0011802 | NA |
| *RHOC* | -0.671051 | 0.207118 | -3.239944 | 0.00119553 | NA |
| *CDC42* | -0.274011 | 0.084799 | -3.231305 | 0.00123226 | 0.055828425 |
| *ENSG00000256092* | -0.622792 | 0.192845 | -3.2295 | 0.00124007 | NA |
| *TNFSF10* | 0.750956 | 0.233104 | 3.22155 | 0.00127499 | 0.057234314 |
| *LCN8* | 3.047177 | 0.951215 | 3.203457 | 0.00135788 | 0.060094407 |
| *H2AFZ* | 0.285445 | 0.089137 | 3.202317 | 0.00136327 | 0.060094407 |

| | | | | | |
|---|---|---|---|---|---|
| *TSPAN33* | -0.533479 | 0.166663 | -3.200951 | 0.00136975 | NA |
| *ACADVL* | -0.283078 | 0.088489 | -3.199017 | 0.00137897 | 0.060240479 |
| *RPS28* | 0.194913 | 0.060978 | 3.196469 | 0.00139121 | 0.060240479 |
| *PLAAT3* | -0.879527 | 0.275932 | -3.187479 | 0.00143519 | NA |
| *RPS6* | 0.198205 | 0.062226 | 3.185257 | 0.00144626 | 0.062074775 |
| *MAML3* | -1.127562 | 0.354737 | -3.178587 | 0.00147995 | NA |
| *IGLV6.57* | -1.292565 | 0.406772 | -3.177617 | 0.00148491 | NA |
| *ARF5* | -0.34252 | 0.107889 | -3.174743 | 0.00149969 | 0.063463521 |
| *HSH2D* | -0.316101 | 0.099625 | -3.172895 | 0.00150927 | 0.063463521 |
| *HSPD1* | 0.325575 | 0.102706 | 3.169981 | 0.00152449 | 0.063463521 |
| *IGHD* | -0.311222 | 0.098272 | -3.166954 | 0.00154045 | 0.063463521 |
| *ELF1* | 0.284993 | 0.090006 | 3.166385 | 0.00154346 | 0.063463521 |
| *H1FX* | -0.841793 | 0.266156 | -3.162773 | 0.00156274 | NA |
| *TIMM10* | -0.466356 | 0.147899 | -3.1532 | 0.00161491 | 0.065847903 |
| *HLA.A* | -0.48811 | 0.155191 | -3.145216 | 0.00165964 | 0.066734957 |
| *DEDD2* | 0.373975 | 0.118986 | 3.143012 | 0.00167219 | 0.066734957 |
| *RPL11* | 0.17566 | 0.055922 | 3.141162 | 0.00168279 | 0.066734957 |
| *RPL19* | 0.174982 | 0.055732 | 3.139698 | 0.00169122 | 0.066734957 |
| *TOE1* | -0.525842 | 0.16765 | -3.13654 | 0.00170954 | NA |
| *HSD17B11* | 0.225734 | 0.071969 | 3.13654 | 0.00170954 | 0.066918293 |
| *ENSG00000280433* | -1.035115 | 0.330889 | -3.128285 | 0.00175829 | NA |
| *S1PR4* | 0.334145 | 0.106935 | 3.12474 | 0.00177962 | 0.069108695 |
| *CD82* | -0.453641 | 0.145515 | -3.11748 | 0.00182404 | 0.069941038 |
| *CFAP410* | -0.353813 | 0.113536 | -3.116309 | 0.0018313 | 0.069941038 |
| *PRPF38A* | 0.267702 | 0.08596 | 3.114281 | 0.00184394 | 0.069941038 |
| *ENSG00000275964* | -0.777304 | 0.249614 | -3.114021 | 0.00184556 | NA |
| *XXYLT1.AS2* | 1.363731 | 0.438451 | 3.110336 | 0.00186875 | NA |
| *PHF23* | 0.402011 | 0.129262 | 3.110054 | 0.00187053 | 0.070403929 |
| *STX7* | 0.242498 | 0.078181 | 3.101743 | 0.00192385 | 0.071857913 |
| *LYRM7* | 0.38125 | 0.123316 | 3.091657 | 0.00199042 | 0.073781386 |
| *PILRB* | 0.788754 | 0.255835 | 3.083062 | 0.00204882 | 0.075315587 |
| *RPL23A* | 0.201632 | 0.065442 | 3.081068 | 0.0020626 | 0.075315587 |
| *ENSG00000254397* | 1.823559 | 0.592292 | 3.078818 | 0.00207824 | NA |
| *SPINT2* | -0.295527 | 0.096286 | -3.069263 | 0.00214588 | 0.077776047 |
| *NFKB2* | -0.583608 | 0.190647 | -3.061202 | 0.0022045 | 0.079313463 |
| *TAPBP* | -0.323273 | 0.10571 | -3.058101 | 0.00222745 | 0.079553971 |
| *ENSG00000166927* | -1.052166 | 0.344448 | -3.05464 | 0.00225331 | 0.079686918 |
| *IL23A* | -0.942947 | 0.308746 | -3.054122 | 0.0022572 | NA |
| *HSBP1* | 0.262303 | 0.085909 | 3.053254 | 0.00226374 | 0.079686918 |
| *PCMTD2* | -0.384406 | 0.126111 | -3.048158 | 0.00230249 | 0.080019533 |
| *LDHA* | -0.326486 | 0.107125 | -3.047713 | 0.0023059 | 0.080019533 |
| *ROR1* | -1.186801 | 0.391034 | -3.035032 | 0.0024051 | NA |
| *RPL35A* | 0.205765 | 0.067807 | 3.034584 | 0.00240868 | 0.082407157 |
| *RPL10* | 0.231465 | 0.076286 | 3.034188 | 0.00241185 | 0.082407157 |
| *ENSG00000282988* | 0.739522 | 0.243864 | 3.032518 | 0.00242523 | 0.082407157 |
| *NIPAL3* | 0.476726 | 0.15732 | 3.0303 | 0.00244311 | 0.082442266 |
| *RPS18* | 0.2255 | 0.074541 | 3.025194 | 0.00248474 | 0.083272686 |

| | | | | | |
|---|---|---|---|---|---|
| *SRA1* | -0.430777 | 0.14273 | -3.018135 | 0.00254335 | 0.084657294 |
| *IL21R* | -1.125275 | 0.372887 | -3.017737 | 0.0025467 | NA |
| *ENSG00000273748* | -1.525363 | 0.50591 | -3.015087 | 0.00256905 | 0.08493499 |
| *TCOF1* | -0.419291 | 0.139407 | -3.007673 | 0.00263256 | 0.086450426 |
| *DDB2* | -0.496002 | 0.165111 | -3.004043 | 0.00266417 | 0.086905343 |
| *ELMSAN1* | 0.380013 | 0.126616 | 3.001288 | 0.0026884 | 0.087074039 |
| *TNRC6C* | 0.5103 | 0.170071 | 3.000504 | 0.00269533 | NA |
| *TDG* | 0.305235 | 0.101855 | 2.996769 | 0.00272857 | 0.087074039 |
| *PCIF1* | -0.393408 | 0.131431 | -2.993279 | 0.00275997 | 0.087074039 |
| *GCC2* | 0.263848 | 0.088149 | 2.993201 | 0.00276068 | 0.087074039 |
| *RPLP2* | 0.24274 | 0.081112 | 2.992641 | 0.00276575 | 0.087074039 |
| *MRPL28* | -0.286764 | 0.09586 | -2.991499 | 0.00277612 | 0.087074039 |
| *RHBDD2* | -0.500153 | 0.167411 | -2.987582 | 0.00281194 | 0.087635666 |
| *AASDH* | 0.382776 | 0.128361 | 2.982025 | 0.00286349 | 0.088677528 |
| *RPS13* | 0.149323 | 0.050206 | 2.974185 | 0.00293768 | 0.090402951 |
| *CRLF3* | 0.270125 | 0.090932 | 2.970624 | 0.00297196 | 0.090886182 |
| *TMEM170A* | 0.384487 | 0.129542 | 2.968052 | 0.00299694 | 0.091080836 |
| *CMTM7* | -0.37344 | 0.126134 | -2.960668 | 0.00306973 | 0.092554195 |
| *RIN3* | 0.716283 | 0.242034 | 2.959429 | 0.0030821 | NA |
| *SRP72* | -0.35572 | 0.120203 | -2.959314 | 0.00308325 | 0.092554195 |
| *CCL4* | -1.705615 | 0.57698 | -2.956108 | 0.00311548 | NA |
| *RPL34* | 0.15883 | 0.05384 | 2.950051 | 0.00317722 | 0.094793416 |
| *RPL21* | 0.191804 | 0.065263 | 2.938915 | 0.00329364 | 0.097671311 |
| *MME* | -2.060243 | 0.701642 | -2.936317 | 0.00332135 | NA |
| *COLCA2* | -1.373355 | 0.467992 | -2.934569 | 0.00334012 | NA |
| *PURA* | 0.493217 | 0.168154 | 2.93313 | 0.00335563 | 0.097868847 |
| *PRDX3* | 0.428249 | 0.146052 | 2.932166 | 0.00336606 | 0.097868847 |
| *MAPKAPK2* | -0.453663 | 0.154867 | -2.929369 | 0.0033965 | 0.097868847 |
| *PSPC1* | -0.406329 | 0.138719 | -2.929142 | 0.003399 | 0.097868847 |
| *POLR2J* | -0.270736 | 0.092432 | -2.929022 | 0.00340031 | 0.097868847 |
| *ZNF16* | -0.651843 | 0.222714 | -2.926816 | 0.00342451 | NA |
| *LIMD2* | -0.255781 | 0.087633 | -2.918778 | 0.00351406 | 0.099441379 |
| *PCGF5* | -0.287392 | 0.098463 | -2.918774 | 0.00351411 | 0.099441379 |
| *RPL37* | 0.142625 | 0.048867 | 2.918614 | 0.00351591 | 0.099441379 |
| *ENSG00000130313* | -0.230551 | 0.079121 | -2.913899 | 0.00356945 | 0.100278419 |
| *FAM184B* | -1.755632 | 0.60281 | -2.912412 | 0.0035865 | 0.100278419 |
| *ZC3H7A* | -0.374955 | 0.129031 | -2.905923 | 0.00366172 | 0.101235053 |
| *MIF* | -0.356909 | 0.122823 | -2.905891 | 0.00366209 | 0.101235053 |
| *ADAMTS6* | 0.901601 | 0.310914 | 2.899836 | 0.00373358 | NA |
| *TNFAIP8L2* | 0.551683 | 0.190483 | 2.896234 | 0.00377671 | NA |
| *RPL35* | 0.169675 | 0.058619 | 2.894539 | 0.00379715 | 0.104379059 |
| *SVBP* | -0.243834 | 0.084301 | -2.892421 | 0.00382285 | 0.104498456 |
| *TM2D3* | 0.501328 | 0.173548 | 2.888691 | 0.00386849 | 0.105158555 |
| *VOPP1* | -0.382596 | 0.13265 | -2.884248 | 0.00392349 | 0.106064304 |
| *SLC25A25* | 0.636256 | 0.220833 | 2.881164 | 0.0039621 | NA |
| *LILRB1* | 0.496608 | 0.17276 | 2.874558 | 0.00404594 | 0.10877356 |
| *SEC61B* | -0.207005 | 0.072317 | -2.862458 | 0.00420369 | 0.112228979 |

| | | | | | |
|---|---|---|---|---|---|
| *MRPL41* | -0.291403 | 0.101846 | -2.861205 | 0.00422034 | 0.112228979 |
| *ATF7IP2* | -0.437288 | 0.152942 | -2.85917 | 0.00424751 | 0.112340831 |
| *CAV1* | -1.194325 | 0.41795 | -2.857577 | 0.00426889 | NA |
| *ENSG00000272754* | -0.743623 | 0.260326 | -2.856502 | 0.00428337 | NA |
| *SESN3* | 0.306248 | 0.10733 | 2.85334 | 0.00432623 | 0.11380773 |
| *GOPC* | 0.394866 | 0.138553 | 2.849933 | 0.00437284 | 0.114418726 |
| *SETD5* | 0.254646 | 0.089505 | 2.845045 | 0.00444051 | 0.115312441 |
| *FAM98A* | 0.497794 | 0.175039 | 2.843904 | 0.00445645 | NA |
| *RPS15A* | 0.205888 | 0.072402 | 2.843665 | 0.00445979 | 0.115312441 |
| *ACTR1B* | -0.283076 | 0.099591 | -2.842388 | 0.0044777 | 0.115312441 |
| *LNPEP* | 0.366092 | 0.128977 | 2.838425 | 0.00453367 | 0.116142742 |
| *NOP10* | 0.19167 | 0.067576 | 2.836369 | 0.00456297 | 0.116213014 |
| *AEN* | -0.57529 | 0.20291 | -2.8352 | 0.00457969 | NA |
| *PTGES2* | -0.437927 | 0.15455 | -2.833558 | 0.0046033 | 0.116213014 |
| *TIAL1* | 0.237631 | 0.083933 | 2.831198 | 0.0046374 | 0.116213014 |
| *INKA1* | 0.475206 | 0.167912 | 2.830087 | 0.00465353 | 0.116213014 |
| *SUMO3* | -0.422274 | 0.149237 | -2.829557 | 0.00466125 | 0.116213014 |
| *ZNF253* | 0.551765 | 0.195023 | 2.829235 | 0.00466594 | NA |
| *CYBC1* | -0.189521 | 0.067008 | -2.828346 | 0.00467892 | 0.116213014 |
| *CLECL1* | 1.554259 | 0.549767 | 2.827121 | 0.00469685 | NA |
| *RPL39* | 0.18079 | 0.063997 | 2.82499 | 0.00472821 | 0.116844021 |
| *RPS12* | 0.189623 | 0.067167 | 2.823153 | 0.00475539 | 0.116925297 |
| *UQCRH* | 0.152193 | 0.054 | 2.818374 | 0.00482676 | 0.118086557 |
| *ZNF557* | 0.433904 | 0.15411 | 2.815549 | 0.00486939 | 0.11853699 |
| *EIF3E* | 0.256169 | 0.091168 | 2.809868 | 0.00495618 | 0.1196862 |
| *YWHAE* | 0.34407 | 0.122477 | 2.809262 | 0.00496552 | 0.1196862 |
| *ENSG00000245904* | 0.246239 | 0.087772 | 2.805448 | 0.00502468 | 0.120518344 |
| *CPM* | -1.058451 | 0.377783 | -2.801742 | 0.00508275 | NA |
| *AKAP13* | 0.242941 | 0.086908 | 2.795369 | 0.00518404 | 0.123692773 |
| *CYTH1* | -0.245946 | 0.08803 | -2.793905 | 0.00520759 | 0.123692773 |
| *SLC15A2* | 0.786214 | 0.281523 | 2.792714 | 0.00522678 | NA |
| *MLST8* | -0.394124 | 0.141246 | -2.790328 | 0.00526546 | 0.124463333 |
| *PTP4A1* | -0.34734 | 0.124874 | -2.781521 | 0.00541048 | 0.127276274 |
| *LINC02256* | -0.537581 | 0.193402 | -2.779603 | 0.00544255 | NA |
| *ADRB2* | 1.090669 | 0.393052 | 2.774874 | 0.0055223 | NA |
| *RPL5* | 0.178347 | 0.064296 | 2.773852 | 0.00553969 | 0.129474663 |
| *RPL26* | 0.160762 | 0.05801 | 2.77131 | 0.00558313 | 0.129474663 |
| *APOBEC3C* | -0.276212 | 0.099669 | -2.771299 | 0.00558331 | 0.129474663 |
| *HCK* | -0.816004 | 0.294459 | -2.771199 | 0.00558503 | NA |
| *C6orf226* | -0.52381 | 0.189086 | -2.770215 | 0.00560193 | NA |
| *TTC9C* | 0.320794 | 0.115897 | 2.767925 | 0.00564144 | 0.130183803 |
| *RPS27L* | -0.326341 | 0.117964 | -2.766445 | 0.00566711 | 0.130183803 |
| *RYK* | 0.538182 | 0.194859 | 2.761908 | 0.00574647 | NA |
| *POU2F2* | 0.221034 | 0.080064 | 2.760725 | 0.00576732 | 0.1309594 |
| *SLC25A5* | -0.178823 | 0.064788 | -2.760103 | 0.00577831 | 0.1309594 |
| *VPREB1* | -1.983685 | 0.71898 | -2.759027 | 0.00579738 | 0.1309594 |
| *GAPT* | 0.317165 | 0.115066 | 2.756373 | 0.00584463 | 0.1309594 |

| | | | | | |
|---|---|---|---|---|---|
| *ABTB1* | -0.273906 | 0.099476 | -2.753481 | 0.00589653 | 0.1309594 |
| *HRK* | -1.300865 | 0.472458 | -2.753397 | 0.00589803 | 0.1309594 |
| *MFAP1* | 0.427017 | 0.155097 | 2.75322 | 0.00590122 | 0.1309594 |
| *PSMB10* | -0.318096 | 0.115733 | -2.748524 | 0.00598643 | 0.1309594 |
| *EGLN2* | 0.274869 | 0.100019 | 2.748168 | 0.00599294 | 0.1309594 |
| *BET1* | 0.325685 | 0.118557 | 2.747086 | 0.00601274 | 0.1309594 |
| *TRAM1* | -0.185794 | 0.067656 | -2.746158 | 0.00602978 | 0.1309594 |
| *SYNE2* | -0.422918 | 0.154009 | -2.746061 | 0.00603155 | 0.1309594 |
| *SCAND1* | -0.222559 | 0.081074 | -2.745124 | 0.00604881 | 0.1309594 |
| *NT5E* | 0.899723 | 0.327816 | 2.744596 | 0.00605854 | NA |
| *ATP6V0A2* | -0.770433 | 0.280977 | -2.741978 | 0.00610705 | NA |
| *CRIP1* | -0.563519 | 0.205696 | -2.739571 | 0.00615194 | 0.131908708 |
| *ITGB7* | 0.445843 | 0.162795 | 2.738677 | 0.00616869 | 0.131908708 |
| *RANGRF* | -0.29113 | 0.106313 | -2.738419 | 0.00617353 | 0.131908708 |
| *BIN1* | 0.397057 | 0.145105 | 2.73635 | 0.0062125 | 0.132164144 |
| *GTF3C6* | -0.260674 | 0.095321 | -2.734688 | 0.00624394 | 0.132258025 |
| *PDPK1* | 0.383146 | 0.140236 | 2.732153 | 0.0062922 | 0.132705683 |
| *RPL13* | 0.193309 | 0.070909 | 2.726172 | 0.00640736 | 0.134554481 |
| *ENSA* | -0.167654 | 0.061575 | -2.722777 | 0.00647357 | 0.135364026 |
| *RPS15* | 0.142783 | 0.052471 | 2.721161 | 0.00650531 | 0.135448812 |
| *ENSG00000263004* | -1.062851 | 0.391107 | -2.717544 | 0.00657684 | NA |
| *UBE2L3* | -0.212085 | 0.078046 | -2.717448 | 0.00657874 | 0.136038619 |
| *YTHDF3* | 0.53961 | 0.198618 | 2.716819 | 0.00659126 | NA |
| *EEF1B2* | 0.286283 | 0.105401 | 2.716133 | 0.00660495 | 0.136038619 |
| *UBA52* | 0.125479 | 0.046208 | 2.715527 | 0.00661704 | 0.136038619 |
| *VPS25* | -0.384442 | 0.141728 | -2.712529 | 0.00667719 | 0.136700872 |
| *FBXO4* | -0.489094 | 0.18045 | -2.710417 | 0.00671986 | NA |
| *INTS5* | 0.593392 | 0.219082 | 2.708535 | 0.0067581 | NA |
| *ATP6V1C2* | -1.757281 | 0.649031 | -2.707545 | 0.00677829 | NA |
| *PFKFB3* | -0.42051 | 0.155317 | -2.707427 | 0.00678071 | 0.138241642 |
| *SLCO5A1* | -1.131897 | 0.418556 | -2.704288 | 0.00684509 | NA |
| *RACK1* | 0.179328 | 0.066359 | 2.702394 | 0.00688421 | 0.139474999 |
| *DENND2D* | 0.494061 | 0.182869 | 2.701719 | 0.00689821 | 0.139474999 |
| *ENSG00000260879* | -1.134608 | 0.420333 | -2.699308 | 0.00694839 | NA |
| *UCP2* | -0.298811 | 0.110771 | -2.697552 | 0.00698514 | 0.140651475 |
| *MYL12A* | 0.174549 | 0.064795 | 2.693868 | 0.00706282 | 0.141292516 |
| *UBE2E3* | -0.310966 | 0.115459 | -2.693306 | 0.00707473 | 0.141292516 |
| *LACC1* | -0.831987 | 0.30913 | -2.691377 | 0.00711577 | NA |
| *FCRL2* | 0.367777 | 0.136673 | 2.690914 | 0.00712566 | 0.141731174 |
| *ENSG00000278158* | 0.280179 | 0.10437 | 2.684487 | 0.00726411 | 0.143899991 |
| *ALOX5* | -0.306124 | 0.114164 | -2.681448 | 0.00733044 | 0.144628401 |
| *RPL18* | 0.187302 | 0.069909 | 2.679234 | 0.00737909 | 0.144952071 |
| *NDUFS8* | -0.279515 | 0.104374 | -2.67801 | 0.00740609 | 0.144952071 |
| *PIP4K2A* | 0.335144 | 0.125522 | 2.67 | 0.00758514 | 0.147864808 |
| *SPHK2* | -0.31189 | 0.117137 | -2.662617 | 0.00775357 | 0.149994386 |
| *ID3* | -0.317455 | 0.119248 | -2.662152 | 0.00776429 | 0.149994386 |
| *HSPE1* | 0.163839 | 0.061566 | 2.661197 | 0.00778634 | 0.149994386 |

| | | | | | |
|---|---|---|---|---|---|
| *FNTA* | 0.311237 | 0.117106 | 2.657749 | 0.00786644 | 0.150943179 |
| *NFKBIB* | -0.398174 | 0.149909 | -2.656117 | 0.00790463 | NA |
| *ENSG00000275799* | 0.600337 | 0.226095 | 2.655239 | 0.00792523 | 0.151477045 |
| *CCNG1* | 0.238444 | 0.089906 | 2.652145 | 0.00799822 | 0.152277394 |
| *ENSG00000239636* | -0.611627 | 0.230822 | -2.649785 | 0.0080543 | NA |
| *THAP2* | 0.418559 | 0.15798 | 2.649438 | 0.00806258 | NA |
| *CLCF1* | 0.548929 | 0.207306 | 2.64792 | 0.00809887 | 0.153244288 |
| *PLTP* | -1.516412 | 0.572743 | -2.647632 | 0.00810578 | NA |
| *PF4* | 2.595135 | 0.980214 | 2.647517 | 0.00810852 | NA |
| *ENSG00000271204* | 0.544914 | 0.205831 | 2.647387 | 0.00811164 | 0.153244288 |
| *C11orf24* | -0.416076 | 0.157279 | -2.645465 | 0.00815786 | NA |
| *PPP4C* | -0.181597 | 0.068761 | -2.640997 | 0.00826623 | 0.155564172 |
| *ATP6V1C1* | 0.376196 | 0.142712 | 2.636051 | 0.00838772 | NA |
| *PFKP* | -0.85598 | 0.325288 | -2.631451 | 0.00850211 | NA |
| *ZNRD2* | -0.342515 | 0.130443 | -2.625785 | 0.00864495 | 0.162064417 |
| *RPS27* | 0.157738 | 0.06012 | 2.623714 | 0.00869768 | 0.162064417 |
| *NDUFB8* | -0.151631 | 0.057804 | -2.623193 | 0.008711 | 0.162064417 |
| *PIM1* | -0.377791 | 0.144357 | -2.617052 | 0.00886929 | 0.164384309 |
| *TOMM20* | 0.240327 | 0.091908 | 2.614873 | 0.00892606 | 0.164812027 |
| *DNAAF4* | -1.198195 | 0.458246 | -2.614741 | 0.00892951 | NA |
| *RBBP8* | -0.645056 | 0.246755 | -2.614156 | 0.00894481 | NA |
| *SPACA9* | -0.57916 | 0.221558 | -2.61403 | 0.00894811 | NA |
| *UBE2J2* | -0.265526 | 0.101645 | -2.612293 | 0.0089937 | 0.165436778 |
| *NUAK2* | -0.243456 | 0.093275 | -2.610094 | 0.00905172 | 0.165880454 |
| *CHRAC1* | -0.394821 | 0.151412 | -2.607587 | 0.00911829 | 0.16647674 |
| *ENSG00000231856* | -0.935858 | 0.359122 | -2.605961 | 0.0091617 | NA |
| *DNAAF5* | -0.91531 | 0.351264 | -2.605764 | 0.00916696 | NA |
| *MYC* | 0.379556 | 0.145798 | 2.603309 | 0.00923288 | 0.167929264 |
| *HSPH1* | 0.43613 | 0.167721 | 2.600334 | 0.0093133 | 0.167929264 |
| *BBC3* | -1.118981 | 0.430397 | -2.59988 | 0.00932563 | NA |
| *CAPN12* | 0.440776 | 0.16954 | 2.599839 | 0.00932674 | 0.167929264 |
| *SLC16A1.AS1* | 0.445365 | 0.171325 | 2.599531 | 0.00933512 | 0.167929264 |
| *FUOM* | -0.543878 | 0.209222 | -2.599526 | 0.00933527 | NA |
| *ENSG00000185527* | -0.963706 | 0.371097 | -2.596908 | 0.0094067 | NA |
| *XIAP* | 0.370507 | 0.142697 | 2.596455 | 0.00941912 | 0.16881961 |
| *FAM217B* | 0.393687 | 0.151827 | 2.593005 | 0.00951414 | 0.169900311 |
| *ENSG00000179094* | -0.784403 | 0.302812 | -2.590394 | 0.0095866 | NA |
| *ENSG00000262089* | 0.643232 | 0.24832 | 2.590333 | 0.0095883 | NA |
| *PTPN20* | -1.27365 | 0.491973 | -2.58886 | 0.00962942 | NA |
| *HSPA4* | -0.359719 | 0.139095 | -2.586129 | 0.00970607 | 0.172697399 |
| *LINC02422* | 0.787421 | 0.304825 | 2.583188 | 0.0097892 | NA |
| *OTUB1* | -0.222 | 0.085975 | -2.58216 | 0.0098184 | 0.173495571 |
| *RPS16* | 0.172607 | 0.066849 | 2.582039 | 0.00982184 | 0.173495571 |
| *LHFPL2* | -0.464941 | 0.180134 | -2.581083 | 0.00984909 | NA |
| *NEMP1* | 0.741633 | 0.287391 | 2.580575 | 0.00986361 | NA |
| *LINC02397* | 0.281732 | 0.109292 | 2.577777 | 0.00994383 | 0.174082295 |
| *FCGR2B* | 0.672499 | 0.260887 | 2.57774 | 0.00994488 | 0.174082295 |

| | | | | | |
|---|---|---|---|---|---|
| *PDE6D* | -0.381198 | 0.147914 | -2.577153 | 0.00996179 | 0.174082295 |
| *UNC50* | 0.318851 | 0.123849 | 2.574509 | 0.01003826 | 0.174794303 |
| *RPS3A* | 0.167783 | 0.065258 | 2.571072 | 0.01013842 | 0.175812233 |
| *FZD3* | 1.107262 | 0.430665 | 2.571052 | 0.010139 | NA |
| *ERGIC2* | 0.282611 | 0.110004 | 2.569093 | 0.01019651 | 0.175812233 |
| *DYNLL1* | 0.315994 | 0.123055 | 2.567914 | 0.01023126 | 0.175812233 |
| *BBX* | 0.231082 | 0.089999 | 2.567603 | 0.01024044 | 0.175812233 |
| *ZNF350* | -0.260766 | 0.101683 | -2.564492 | 0.01033268 | 0.176634813 |
| *MAT2A* | 0.243615 | 0.095049 | 2.563038 | 0.01037605 | 0.176634813 |
| *GLRX* | 0.306484 | 0.119612 | 2.562324 | 0.01039742 | 0.176634813 |
| *DLGAP3* | -1.01477 | 0.39612 | -2.561772 | 0.01041397 | NA |
| *RPL29* | 0.169651 | 0.066259 | 2.560415 | 0.01045474 | 0.176634813 |
| *MAD2L2* | -0.353588 | 0.138264 | -2.557342 | 0.01054753 | 0.176634813 |
| *CCNG2* | 0.571025 | 0.223312 | 2.557076 | 0.01055563 | 0.176634813 |
| *NUP50* | 0.352357 | 0.137812 | 2.556802 | 0.01056394 | 0.176634813 |
| *ATRAID* | -0.20057 | 0.078459 | -2.556367 | 0.01057715 | 0.176634813 |
| *ST13* | 0.185407 | 0.072644 | 2.552262 | 0.01070262 | 0.178122098 |
| *FHL3* | -0.914414 | 0.358412 | -2.551292 | 0.01073243 | NA |
| *IGHV1.2* | 1.61346 | 0.632743 | 2.549946 | 0.01077395 | NA |
| *KRCC1* | 0.250183 | 0.098177 | 2.548279 | 0.01082559 | 0.179558034 |
| *RPN1* | -0.453929 | 0.178354 | -2.545098 | 0.01092469 | 0.180268041 |
| *IFIT2* | 0.479433 | 0.188384 | 2.544975 | 0.01092855 | NA |
| *STK17A* | 0.268257 | 0.105439 | 2.544192 | 0.01095308 | 0.180268041 |
| *HINT2* | -0.219144 | 0.086163 | -2.543369 | 0.01097892 | 0.180268041 |
| *BATF* | 1.104642 | 0.434625 | 2.541597 | 0.01103473 | NA |
| *CD40* | -0.233378 | 0.091867 | -2.540385 | 0.01107304 | 0.181205321 |
| *FAM43A* | 0.710271 | 0.279846 | 2.538077 | 0.01114636 | 0.1817971 |
| *DTX3L* | 0.466191 | 0.183842 | 2.53583 | 0.01121811 | NA |
| *TMEM205* | -0.331599 | 0.130965 | -2.531972 | 0.01134232 | 0.184378613 |
| *FBXO8* | 0.5057 | 0.199768 | 2.531436 | 0.01135964 | NA |
| *SOX4* | -0.873724 | 0.345358 | -2.529911 | 0.01140914 | NA |
| *NGLY1* | 0.199051 | 0.078686 | 2.529688 | 0.01141639 | 0.184660001 |
| *PPP1R15A* | -0.556058 | 0.219905 | -2.528629 | 0.0114509 | 0.184660001 |
| *ECI2* | 0.323018 | 0.127785 | 2.527824 | 0.01147719 | 0.184660001 |
| *SLK* | -0.489084 | 0.193505 | -2.527501 | 0.01148775 | NA |
| *SKAP2* | 0.297638 | 0.117792 | 2.526804 | 0.01151059 | 0.184660001 |
| *MED21* | 0.397974 | 0.157584 | 2.525474 | 0.01155423 | NA |
| *IKBIP* | 0.293966 | 0.116638 | 2.520326 | 0.01172461 | 0.187478867 |
| *APBB2* | -0.846972 | 0.336169 | -2.519485 | 0.01175266 | NA |
| *GRAMD1C* | -0.571206 | 0.22692 | -2.517208 | 0.01182891 | 0.188530403 |
| *YARS* | -0.566445 | 0.225029 | -2.517204 | 0.01182902 | NA |
| *CYSTM1* | -0.321657 | 0.127958 | -2.51378 | 0.01194451 | 0.189523685 |
| *CFL1* | -0.169002 | 0.067254 | -2.512884 | 0.01197487 | 0.189523685 |
| *RRP8* | 0.358214 | 0.142565 | 2.512634 | 0.01198337 | NA |
| *RPL27A* | 0.139133 | 0.055389 | 2.511926 | 0.01200743 | 0.189523685 |
| *FNBP1* | -0.331856 | 0.132229 | -2.509698 | 0.01208345 | 0.190110347 |
| *LINC00638* | 0.976702 | 0.3894 | 2.508226 | 0.01213392 | NA |

| | | | | | |
|---|---|---|---|---|---|
| *MIS18BP1* | 0.182819 | 0.072906 | 2.507597 | 0.01215552 | 0.190631231 |
| *ENSG00000270562* | 0.618622 | 0.247183 | 2.502686 | 0.01232548 | NA |
| *ZBTB20* | 0.258248 | 0.10326 | 2.500939 | 0.01238645 | 0.193632305 |
| *KLHL2* | -0.727038 | 0.29076 | -2.500474 | 0.01240273 | NA |
| *ENSG00000268516* | 0.456051 | 0.182396 | 2.500337 | 0.01240753 | NA |
| *EXOG* | -0.432704 | 0.17346 | -2.494542 | 0.01261198 | 0.196530062 |
| *MICOS13* | -0.213222 | 0.085535 | -2.492805 | 0.01267385 | 0.196867091 |
| *CFAP73* | -1.104437 | 0.44357 | -2.489883 | 0.01277851 | NA |
| *HIP1R* | -0.289572 | 0.116436 | -2.486968 | 0.01288371 | 0.19895543 |
| *ZNF138* | 0.414299 | 0.166704 | 2.485235 | 0.0129466 | 0.19895543 |
| *CCT8* | 0.440732 | 0.177377 | 2.484713 | 0.01296558 | 0.19895543 |
| *FLII* | -0.35755 | 0.143961 | -2.483667 | 0.01300372 | 0.19895543 |
| *MT.CO1* | 0.211661 | 0.085229 | 2.483452 | 0.0130116 | 0.19895543 |
| *SCAMP4* | -0.606467 | 0.244284 | -2.482632 | 0.01304158 | NA |
| *SSR2* | 0.148378 | 0.059794 | 2.481469 | 0.0130842 | 0.199442258 |
| *LSM6* | 0.19613 | 0.079093 | 2.479752 | 0.01314738 | 0.199782989 |
| *ENSG00000260349* | -0.795281 | 0.320771 | -2.479276 | 0.01316496 | NA |
| *HSP90AB1* | 0.223881 | 0.090354 | 2.477813 | 0.01321906 | 0.200250286 |
| *PIP4P2* | 0.50201 | 0.202765 | 2.47582 | 0.01329305 | NA |
| *PDCL3* | -0.345681 | 0.139626 | -2.475757 | 0.01329539 | 0.200785026 |
| *ENSG00000235560* | 0.510569 | 0.206589 | 2.47143 | 0.01345739 | NA |
| *ZBTB16* | 2.010131 | 0.813463 | 2.47108 | 0.01347058 | NA |
| *CAST* | -0.309502 | 0.125292 | -2.47025 | 0.01350187 | 0.202514433 |
| *PHPT1* | -0.466565 | 0.18889 | -2.470041 | 0.01350974 | 0.202514433 |
| *KMT2A* | 0.232561 | 0.094177 | 2.469398 | 0.01353407 | 0.202514433 |
| *UNC119* | -0.335012 | 0.135785 | -2.46722 | 0.01361668 | 0.202532344 |
| *PTTG1IP* | 0.396481 | 0.160714 | 2.46699 | 0.01362541 | 0.202532344 |
| *RNF146* | 0.283151 | 0.114817 | 2.466097 | 0.01365945 | 0.202532344 |
| *RND1* | -0.839869 | 0.34071 | -2.465051 | 0.01369936 | NA |
| *MTHFR* | 0.332436 | 0.134886 | 2.464565 | 0.01371795 | 0.202785345 |
| *ENSG00000274922* | -1.550244 | 0.629042 | -2.464453 | 0.01372226 | NA |
| *SAP18* | -0.166721 | 0.067806 | -2.458789 | 0.01394066 | 0.205456695 |
| *THNSL1* | 0.846126 | 0.344461 | 2.456378 | 0.01403455 | NA |
| *EBLN2* | 0.453626 | 0.184717 | 2.455783 | 0.01405779 | NA |
| *NDUFB7* | -0.160348 | 0.065295 | -2.455735 | 0.0140597 | 0.206508251 |
| *RPL37A* | 0.129089 | 0.052602 | 2.454074 | 0.01412481 | 0.206508251 |
| *CHMP2B* | 0.240972 | 0.098207 | 2.453722 | 0.01413862 | 0.206508251 |
| *KCTD5* | 0.379322 | 0.154662 | 2.452585 | 0.01418338 | NA |
| *TBPL1* | 0.314131 | 0.128207 | 2.450194 | 0.01427794 | 0.207922462 |
| *ENSG00000227495* | -1.145911 | 0.467994 | -2.448557 | 0.01434297 | NA |
| *ZNF593* | -0.328721 | 0.134281 | -2.448 | 0.01436518 | 0.208572193 |
| *PTPN7* | -0.511668 | 0.209259 | -2.445136 | 0.01447974 | 0.209012677 |
| *GOLGA3* | -0.45082 | 0.184377 | -2.445106 | 0.01448095 | 0.209012677 |
| *CREBL2* | 0.38466 | 0.157571 | 2.441188 | 0.01463903 | NA |
| *NHSL2* | -0.519081 | 0.212773 | -2.439602 | 0.01470345 | 0.2115999 |
| *ORMDL2* | -0.290892 | 0.119514 | -2.433956 | 0.01493482 | 0.214299339 |
| *YJEFN3* | 0.976485 | 0.401583 | 2.431587 | 0.01503284 | NA |

| | | | | | |
|---|---|---|---|---|---|
| *CARD8* | -0.311804 | 0.128235 | -2.431506 | 0.01503621 | 0.215123365 |
| *ENSG00000183308* | -1.429081 | 0.588032 | -2.430277 | 0.01508727 | NA |
| *SRD5A3* | 0.905191 | 0.372491 | 2.430103 | 0.01509455 | NA |
| *RDH5* | 0.463849 | 0.190885 | 2.429986 | 0.01509941 | 0.215218649 |
| *FRY.AS1* | -1.566976 | 0.645049 | -2.429236 | 0.01513069 | NA |
| *GPSM3* | -0.119232 | 0.049091 | -2.4288 | 0.0151489 | 0.215218649 |
| *DBF4B* | 0.763726 | 0.31456 | 2.427915 | 0.01518589 | NA |
| *FXYD7* | 0.683703 | 0.281642 | 2.427559 | 0.01520083 | 0.215218649 |
| *PRKRA* | -0.230909 | 0.095141 | -2.427015 | 0.01522364 | 0.215218649 |
| *UFD1* | -0.35994 | 0.148363 | -2.426083 | 0.0152628 | 0.215218649 |
| *TM9SF4* | -0.432016 | 0.178132 | -2.425255 | 0.01529763 | NA |
| *PRDX2* | -0.329769 | 0.136163 | -2.421876 | 0.0154406 | 0.217100147 |
| *MAP1LC3B2* | -0.639375 | 0.264072 | -2.421215 | 0.01546874 | NA |
| *PTPRS* | 1.571937 | 0.649256 | 2.421136 | 0.0154721 | NA |
| *ZNF827* | 0.833749 | 0.344484 | 2.420286 | 0.0155083 | NA |
| *PTGIR* | -1.230154 | 0.508733 | -2.418073 | 0.01560295 | NA |
| *ENSG00000233178* | -0.615145 | 0.254418 | -2.417858 | 0.01561218 | NA |
| *LANCL2* | -0.858447 | 0.35536 | -2.415713 | 0.01570443 | NA |
| *TNFAIP8* | 0.216674 | 0.089723 | 2.414913 | 0.01573898 | 0.220082337 |
| *KIAA0040* | 0.538618 | 0.223046 | 2.414828 | 0.01574266 | 0.220082337 |
| *GNG11* | 0.903484 | 0.374196 | 2.41447 | 0.01575812 | NA |
| *CNOT1* | -0.259623 | 0.107588 | -2.413135 | 0.01581596 | 0.220477139 |
| *STARD10* | -0.385568 | 0.15995 | -2.410552 | 0.0159284 | NA |
| *CHURC1* | 0.175808 | 0.072977 | 2.409091 | 0.01599231 | 0.22194853 |
| *WDR18* | -0.361166 | 0.150015 | -2.407536 | 0.0160606 | 0.22194853 |
| *TOR3A* | -0.308968 | 0.128372 | -2.406822 | 0.016092 | 0.22194853 |
| *MICAL1* | -0.325634 | 0.135318 | -2.406426 | 0.01610947 | NA |
| *SAYSD1* | 0.333674 | 0.1387 | 2.405729 | 0.01614022 | 0.22194853 |
| *WDR83* | -0.281258 | 0.11692 | -2.405546 | 0.01614831 | 0.22194853 |
| *RAB33B* | 0.620829 | 0.258085 | 2.405519 | 0.0161495 | NA |
| *FAM161A* | 0.498276 | 0.207158 | 2.405291 | 0.01615959 | NA |
| *KRT10* | -0.340039 | 0.141472 | -2.403581 | 0.01623538 | 0.222520185 |
| *TAF8* | -0.3502 | 0.145941 | -2.399593 | 0.0164133 | 0.224330318 |
| *CD53* | 0.193268 | 0.080611 | 2.397526 | 0.01650621 | 0.224408682 |
| *ARID1A* | 0.331538 | 0.138289 | 2.397425 | 0.01651076 | 0.224408682 |
| *ZNF345* | 0.43757 | 0.182628 | 2.395962 | 0.01657682 | NA |
| *DUSP12* | -0.286062 | 0.119436 | -2.395109 | 0.0166154 | 0.225205463 |
| *TSPAN17* | -0.454327 | 0.189719 | -2.394735 | 0.01663237 | NA |
| *CCDC122* | -0.555979 | 0.232249 | -2.393891 | 0.0166707 | NA |
| *CXorf40B* | 0.495943 | 0.207249 | 2.39298 | 0.01671213 | NA |
| *PHKG2* | -0.328147 | 0.137191 | -2.391904 | 0.01676122 | 0.226554276 |
| *NUCB2* | -0.544639 | 0.227708 | -2.391831 | 0.01676458 | NA |
| *CBLN3* | -0.713134 | 0.298305 | -2.390622 | 0.01681986 | NA |
| *COMMD3* | -0.25651 | 0.107391 | -2.388559 | 0.0169146 | 0.227582354 |
| *SRGAP2B* | 0.652928 | 0.273396 | 2.388218 | 0.0169303 | 0.227582354 |
| *TXLNB* | 1.174999 | 0.492096 | 2.387742 | 0.01695224 | NA |
| *DNMT1* | -0.351861 | 0.147542 | -2.38481 | 0.01708794 | 0.229072012 |

| | | | | | |
|---|---|---|---|---|---|
| *TXNDC11* | 0.396606 | 0.166352 | 2.384134 | 0.01711938 | NA |
| *SNRPB* | -0.184089 | 0.077262 | -2.382673 | 0.01718745 | 0.22946159 |
| *PIK3CD.AS1* | -0.87956 | 0.369179 | -2.382477 | 0.01719661 | NA |
| *HLA.C* | -0.463901 | 0.194788 | -2.381563 | 0.01723937 | 0.22946159 |
| *EVI2A* | 0.473635 | 0.198908 | 2.381171 | 0.01725769 | 0.22946159 |
| *OSGEPL1* | 0.469026 | 0.196995 | 2.380898 | 0.01727048 | NA |
| *ENSG00000268400* | -1.498302 | 0.629528 | -2.380039 | 0.01731082 | NA |
| *XPO5* | -0.719274 | 0.302277 | -2.379515 | 0.01733542 | NA |
| *TCTN2* | -0.883452 | 0.371296 | -2.379374 | 0.01734205 | NA |
| *RBAK.RBAKDN* | 0.68464 | 0.287818 | 2.378729 | 0.01737245 | NA |
| *SLBP* | 0.485784 | 0.204244 | 2.378447 | 0.01738574 | 0.230537765 |
| *RPS4X* | 0.262034 | 0.11032 | 2.375218 | 0.01753858 | 0.231548941 |
| *MOB4* | 0.21705 | 0.091396 | 2.374838 | 0.01755664 | 0.231548941 |
| *SEZ6* | -0.62298 | 0.262478 | -2.373452 | 0.0176227 | NA |
| *ENSG00000198106* | -0.33641 | 0.141802 | -2.372397 | 0.01767311 | 0.232458438 |
| *MPND* | -1.03488 | 0.436724 | -2.369642 | 0.01780529 | NA |
| *SUSD3* | 0.334266 | 0.141122 | 2.368633 | 0.01785398 | 0.234207785 |
| *TPRA1* | 0.642237 | 0.271297 | 2.367286 | 0.01791908 | NA |
| *MYBL2* | -1.412934 | 0.596871 | -2.367236 | 0.01792152 | NA |
| *SLTM* | 0.173806 | 0.073447 | 2.366406 | 0.01796171 | 0.234991059 |
| *ZNF232* | 0.476996 | 0.201852 | 2.3631 | 0.01812278 | NA |
| *SRSF11* | 0.203342 | 0.086127 | 2.360957 | 0.01822785 | 0.237837043 |
| *MBLAC1* | -1.210656 | 0.512967 | -2.360104 | 0.01826983 | NA |
| *DUS3L* | -0.448706 | 0.190377 | -2.356929 | 0.01842677 | NA |
| *METTL8* | 0.726364 | 0.308189 | 2.356878 | 0.01842929 | 0.239825825 |
| *ACAP1* | 0.300548 | 0.127708 | 2.353392 | 0.01860303 | 0.241296592 |
| *IGHA1* | -0.552741 | 0.234946 | -2.352635 | 0.01864094 | 0.241296592 |
| *DOCK8* | 0.292542 | 0.124726 | 2.345467 | 0.01900326 | 0.245337584 |
| *DRG2* | -0.291938 | 0.124598 | -2.34304 | 0.01912735 | 0.246289816 |
| *ICA1L* | -0.450706 | 0.192484 | -2.341527 | 0.01920503 | 0.246640996 |
| *NAA60* | -0.383607 | 0.1639 | -2.340495 | 0.01925818 | 0.246676132 |
| *MUC20.OT1* | -0.293479 | 0.125565 | -2.337273 | 0.01942498 | 0.248162998 |
| *CLPP* | -0.211328 | 0.09048 | -2.335634 | 0.01951033 | 0.248604245 |
| *PTBP1* | -0.279724 | 0.11984 | -2.334137 | 0.01958856 | 0.248952839 |
| *LACTB2.AS1* | -0.800263 | 0.343216 | -2.331661 | 0.01971855 | NA |
| *UBE2T* | 0.519094 | 0.222756 | 2.330326 | 0.01978893 | NA |
| *FOXK1* | -0.352347 | 0.151267 | -2.329303 | 0.01984304 | 0.251185624 |
| *ZNF564* | -0.489022 | 0.209974 | -2.328965 | 0.01986094 | NA |
| *KRIT1* | 0.338284 | 0.145303 | 2.328129 | 0.01990524 | 0.251185624 |
| *NDUFA1* | -0.144179 | 0.061936 | -2.327884 | 0.01991826 | 0.251185624 |
| *RPS27A* | 0.155948 | 0.067051 | 2.325811 | 0.02002863 | 0.251928234 |
| *FBXL22* | 0.805855 | 0.346575 | 2.325194 | 0.02006156 | NA |
| *RPS14* | 0.212395 | 0.091366 | 2.324657 | 0.02009029 | 0.252055916 |
| *CYP3A5* | 0.521592 | 0.22458 | 2.322517 | 0.02020511 | NA |
| *LRRC8A* | 0.71105 | 0.306187 | 2.322274 | 0.02021818 | NA |
| *LTA4H* | 0.310699 | 0.133827 | 2.321646 | 0.02025202 | 0.252897875 |
| *GYPC* | 0.181567 | 0.078215 | 2.321378 | 0.02026644 | 0.252897875 |

| | | | | | |
|---|---|---|---|---|---|
| *DHX40* | -0.536479 | 0.231115 | -2.321267 | 0.02027243 | NA |
| *PSMB3* | -0.186209 | 0.080268 | -2.319845 | 0.02034927 | 0.252897875 |
| *NKAPL* | -1.088834 | 0.469404 | -2.319609 | 0.02036206 | NA |
| *DDX21* | 0.188854 | 0.081417 | 2.31957 | 0.02036415 | 0.252897875 |
| *ENSG00000270659* | -0.674386 | 0.290772 | -2.319294 | 0.02037908 | NA |
| *FRG1* | -0.183293 | 0.079105 | -2.31709 | 0.02049883 | 0.25392604 |
| *CABLES1* | -0.897406 | 0.387923 | -2.313363 | 0.0207027 | NA |
| *NPIPB11* | -1.145564 | 0.495286 | -2.312936 | 0.02072614 | NA |
| *ARIH2OS* | -0.425634 | 0.184083 | -2.312181 | 0.02076771 | NA |
| *MYL6* | -0.148029 | 0.064032 | -2.311803 | 0.02078853 | 0.256838118 |
| *RPL22* | 0.124476 | 0.053896 | 2.309568 | 0.02091208 | 0.256838118 |
| *RPS3* | 0.167087 | 0.072354 | 2.309297 | 0.02092713 | 0.256838118 |
| *SEC13* | -0.333994 | 0.144649 | -2.308995 | 0.02094388 | 0.256838118 |
| *TMEM140* | 0.444785 | 0.192677 | 2.308452 | 0.020974 | NA |
| *ENSG00000226571* | 1.196915 | 0.518634 | 2.307822 | 0.02100904 | NA |
| *MARCHF1* | 0.290273 | 0.125793 | 2.307549 | 0.02102421 | 0.257178696 |
| *MSL3* | 0.286326 | 0.124159 | 2.306124 | 0.02110372 | 0.257499905 |
| *RPL7* | 0.179694 | 0.077952 | 2.305194 | 0.02115573 | 0.257499905 |
| *ST3GAL6* | -1.521413 | 0.660082 | -2.304883 | 0.02117312 | NA |
| *BHLHE40* | -0.465646 | 0.202114 | -2.303879 | 0.02122942 | 0.257755692 |
| *RARA* | -0.355178 | 0.154287 | -2.302056 | 0.02133204 | 0.258360597 |
| *KCTD17* | -0.433665 | 0.188599 | -2.299406 | 0.02148189 | NA |
| *POLR2B* | -0.246923 | 0.107426 | -2.298536 | 0.02153129 | 0.259454462 |
| *PFKL* | -0.235724 | 0.102556 | -2.298492 | 0.02153381 | 0.259454462 |
| *COL9A3* | -0.960622 | 0.418088 | -2.297655 | 0.02158144 | 0.259454462 |
| *RTL10* | 0.678327 | 0.295234 | 2.297589 | 0.02158518 | NA |
| *B3GNT7* | -0.816925 | 0.356391 | -2.292219 | 0.02189304 | NA |
| *HAGHL* | 0.355292 | 0.155036 | 2.291674 | 0.02192446 | 0.262932262 |
| *RTL8C* | -0.508968 | 0.222147 | -2.291135 | 0.02195561 | NA |
| *MMP7* | -1.844977 | 0.80539 | -2.290787 | 0.02197573 | NA |
| *TMEM60* | 0.343031 | 0.149786 | 2.290143 | 0.02201304 | 0.263132239 |
| *ENSG00000235078* | 0.45469 | 0.19858 | 2.28971 | 0.02203815 | NA |
| *MRPS15* | -0.206158 | 0.090044 | -2.289528 | 0.02204869 | 0.263132239 |
| *CLIC4* | -0.36264 | 0.158495 | -2.288025 | 0.02213606 | 0.26353222 |
| *TNFAIP1* | 0.472304 | 0.206446 | 2.287785 | 0.02215004 | NA |
| *KLHL8* | 0.410014 | 0.179294 | 2.286826 | 0.02220601 | NA |
| *ENSG00000267390* | 0.434521 | 0.190177 | 2.284826 | 0.02232305 | NA |
| *PDE4D* | -1.042291 | 0.456879 | -2.28133 | 0.02252893 | NA |
| *STK38L* | -0.476664 | 0.209065 | -2.279976 | 0.0226091 | NA |
| *DHX9* | -0.242077 | 0.106184 | -2.279783 | 0.02262056 | 0.26864665 |
| *HLA.F* | -0.357597 | 0.156989 | -2.277842 | 0.02273601 | 0.269267721 |
| *DHX15* | 0.313154 | 0.137535 | 2.276906 | 0.02279186 | 0.269267721 |
| *FDX2* | -0.344718 | 0.151438 | -2.276293 | 0.02282846 | NA |
| *IFT57* | 0.463853 | 0.20379 | 2.276135 | 0.02283795 | 0.269267721 |
| *PIGF* | 0.390657 | 0.171642 | 2.275992 | 0.02284651 | NA |
| *HSD17B7* | 0.334744 | 0.147133 | 2.275117 | 0.02289891 | 0.269337356 |
| *ZNF585A* | 0.482038 | 0.211967 | 2.274123 | 0.02295858 | NA |

| | | | | | |
|---|---|---|---|---|---|
| *ENOSF1* | 1.10238 | 0.48515 | 2.272247 | 0.02307157 | NA |
| *LINC01250* | -2.270184 | 0.999293 | -2.27179 | 0.02309922 | NA |
| *ZNF704* | -1.167461 | 0.514078 | -2.27098 | 0.02314822 | NA |
| *NUP43* | 0.330958 | 0.14577 | 2.270415 | 0.02318244 | NA |
| *HDAC9* | -0.516153 | 0.227345 | -2.270345 | 0.02318666 | NA |
| *APPL2* | -0.54794 | 0.241625 | -2.267733 | 0.02334546 | NA |
| *G6PC3* | -0.533845 | 0.23543 | -2.26753 | 0.02335787 | NA |
| *ADAM28* | 0.319272 | 0.140846 | 2.266819 | 0.02340128 | 0.274501693 |
| *SORL1* | -0.345991 | 0.152687 | -2.26602 | 0.02345018 | 0.274501693 |
| *DOK1* | 0.429448 | 0.189612 | 2.264882 | 0.02351994 | 0.274661282 |
| *THAP9.AS1* | -0.443426 | 0.195998 | -2.262396 | 0.02367296 | 0.275359084 |
| *LMO4* | 0.362146 | 0.160094 | 2.262083 | 0.02369225 | 0.275359084 |
| *ZNF707* | -0.530556 | 0.23477 | -2.259895 | 0.02382778 | NA |
| *GPM6B* | 0.301194 | 0.133327 | 2.259062 | 0.02387955 | 0.276878244 |
| *UGT8* | 0.605173 | 0.268747 | 2.251827 | 0.02433319 | NA |
| *CIZ1* | -0.25951 | 0.115287 | -2.250997 | 0.0243857 | 0.282078587 |
| *CLNS1A* | 0.386815 | 0.171962 | 2.249419 | 0.02448586 | 0.282569158 |
| *ZNF100* | -0.558243 | 0.248298 | -2.24828 | 0.02455837 | NA |
| *CSE1L* | 0.389061 | 0.173074 | 2.247952 | 0.02457925 | NA |
| *IFIT5* | 0.384217 | 0.170986 | 2.247069 | 0.02463562 | NA |
| *APOBEC3H* | -1.064776 | 0.473937 | -2.246659 | 0.02466186 | NA |
| *SPATC1L* | -0.553716 | 0.246502 | -2.24629 | 0.02468547 | 0.283400481 |
| *FNIP1* | -0.336788 | 0.149948 | -2.246027 | 0.02470229 | 0.283400481 |
| *RFC4* | 0.468067 | 0.208403 | 2.245969 | 0.024706 | NA |
| *SNHG32* | 0.197083 | 0.087765 | 2.245569 | 0.02473166 | 0.283400481 |
| *ABHD5* | -0.833997 | 0.371713 | -2.243658 | 0.02485444 | NA |
| *FLYWCH2* | -0.270263 | 0.120571 | -2.241528 | 0.0249919 | 0.285237149 |
| *EXOSC8* | 0.228365 | 0.101891 | 2.241271 | 0.02500853 | 0.285237149 |
| *COCH* | -0.797252 | 0.355866 | -2.240317 | 0.02507032 | NA |
| *FADS2* | -1.343051 | 0.599647 | -2.239737 | 0.02510797 | NA |
| *CYB561D2* | -0.272959 | 0.121945 | -2.238384 | 0.02519603 | 0.28670734 |
| *ZDHHC19* | 1.12934 | 0.505013 | 2.236259 | 0.02533483 | NA |
| *PTPRJ* | -0.889542 | 0.397792 | -2.236198 | 0.02533883 | NA |
| *UBE2I* | -0.126909 | 0.056771 | -2.23545 | 0.02538783 | 0.28821965 |
| *DDX23* | 0.363832 | 0.162756 | 2.235449 | 0.0253879 | NA |
| *UBC* | -0.197142 | 0.088234 | -2.234315 | 0.02546238 | 0.288396818 |
| *ENSG00000272758* | 0.780669 | 0.349446 | 2.234021 | 0.02548171 | NA |
| *LINC01473* | 0.440806 | 0.197337 | 2.233774 | 0.02549792 | NA |
| *LINC00662* | 0.323213 | 0.144876 | 2.23097 | 0.02568311 | 0.29022505 |
| *ATP6V0A1* | 0.444768 | 0.200004 | 2.223793 | 0.0261624 | 0.294806271 |
| *CDC42SE1* | -0.183925 | 0.082734 | -2.223101 | 0.02620902 | 0.294806271 |
| *TERF2IP* | 0.335644 | 0.151043 | 2.222183 | 0.02627091 | 0.294824684 |
| *BARD1* | 0.375982 | 0.169276 | 2.221109 | 0.02634357 | NA |
| *ENSG00000246528* | -0.910668 | 0.410901 | -2.216271 | 0.02667292 | NA |
| *DBNL* | -0.245929 | 0.110986 | -2.215865 | 0.02670076 | 0.298221121 |
| *THUMPD3.AS1* | -0.221265 | 0.099858 | -2.215805 | 0.02670486 | 0.298221121 |
| *CEBPZOS* | 0.251411 | 0.113501 | 2.215053 | 0.0267564 | 0.298221121 |

| | | | | | |
|---|---|---|---|---|---|
| *NUDT9* | 0.391853 | 0.176984 | 2.214066 | 0.02682424 | NA |
| *HNRNPDL* | -0.1963 | 0.088693 | -2.213255 | 0.02688006 | 0.298918461 |
| *JAK3* | -0.447233 | 0.202231 | -2.211493 | 0.02700172 | 0.299590519 |
| *CEMIP2* | 0.626002 | 0.283166 | 2.210726 | 0.0270548 | NA |
| *ZNF202* | -0.597289 | 0.270448 | -2.208522 | 0.0272079 | NA |
| *RIMKLB* | -0.433035 | 0.196092 | -2.208323 | 0.0272218 | NA |
| *LAPTM5* | -0.193666 | 0.087748 | -2.207066 | 0.02730942 | 0.30163334 |
| *CDC37L1.DT* | -0.766702 | 0.347538 | -2.206095 | 0.02737734 | NA |
| *UROD* | -0.323647 | 0.146724 | -2.205816 | 0.02739687 | 0.30163334 |
| *WDR45* | -0.248931 | 0.112886 | -2.205157 | 0.0274431 | 0.30163334 |
| *ATF4* | -0.224464 | 0.1018 | -2.204952 | 0.0274575 | 0.30163334 |
| *GM2A* | 0.588649 | 0.26703 | 2.204431 | 0.02749407 | 0.30163334 |
| *ENSG00000229043* | -1.046519 | 0.475002 | -2.203185 | 0.02758168 | NA |
| *GID8* | -0.208247 | 0.094578 | -2.20186 | 0.02767523 | 0.302941574 |
| *TIGD7* | 0.620404 | 0.281895 | 2.200832 | 0.02774789 | NA |
| *PIGX* | -0.442382 | 0.201094 | -2.199872 | 0.02781595 | NA |
| *NACA* | 0.123785 | 0.056272 | 2.199757 | 0.02782413 | 0.303755203 |
| *SMARCE1* | -0.162353 | 0.073855 | -2.198284 | 0.02792888 | 0.303755203 |
| *AAK1* | -0.774193 | 0.352276 | -2.197691 | 0.02797115 | NA |
| *ZNF441* | -0.400555 | 0.182268 | -2.197618 | 0.02797637 | 0.303755203 |
| *GLS* | 0.322473 | 0.146803 | 2.196642 | 0.02804602 | 0.303755203 |
| *GNB5* | 0.21374 | 0.097355 | 2.195471 | 0.0281298 | 0.303755203 |
| *MS4A1* | -0.22316 | 0.101683 | -2.194677 | 0.02818678 | 0.303755203 |
| *RNF26* | -0.288738 | 0.131593 | -2.19418 | 0.02822249 | 0.303755203 |
| *DEF6* | 0.292517 | 0.133335 | 2.19385 | 0.02824619 | 0.303755203 |
| *APOBEC3D* | -0.315532 | 0.143874 | -2.193108 | 0.02829963 | NA |
| *FKBP5* | 0.816423 | 0.372422 | 2.192201 | 0.02836503 | NA |
| *DHFR2* | 0.485969 | 0.221753 | 2.191487 | 0.02841659 | NA |
| *VPS8* | 0.315313 | 0.14389 | 2.191344 | 0.02842693 | 0.305028478 |
| *GPR137* | -0.399115 | 0.182239 | -2.190062 | 0.02851977 | 0.305355017 |
| *RGS3* | 0.875666 | 0.400127 | 2.18847 | 0.02863537 | NA |
| *CHCHD4* | 0.417232 | 0.190671 | 2.188233 | 0.02865264 | NA |
| *LINC00513* | 0.397351 | 0.181706 | 2.186782 | 0.02875843 | 0.306493047 |
| *NOP53* | 0.162702 | 0.074492 | 2.184145 | 0.02895158 | 0.306493047 |
| *PPP1CB* | -0.25084 | 0.114846 | -2.18414 | 0.02895198 | 0.306493047 |
| *NFKBID* | -0.239415 | 0.109631 | -2.183818 | 0.02897559 | 0.306493047 |
| *INTS13* | 0.457411 | 0.209482 | 2.183539 | 0.02899612 | NA |
| *CTNS* | -0.572562 | 0.2623 | -2.18285 | 0.02904688 | NA |
| *RBM5* | 0.202909 | 0.092969 | 2.182534 | 0.02907017 | 0.306493047 |
| *ZNF580* | -0.227377 | 0.104184 | -2.182461 | 0.0290755 | 0.306493047 |
| *XPO4* | -0.389606 | 0.178599 | -2.181451 | 0.02915008 | NA |
| *NECAP2* | -0.291372 | 0.133576 | -2.181325 | 0.02915939 | 0.306493047 |
| *TGIF1* | 0.322906 | 0.148051 | 2.18105 | 0.02917973 | 0.306493047 |
| *ACSF3* | 0.394953 | 0.181095 | 2.180914 | 0.02918981 | 0.306493047 |
| *TFAP4* | 0.338841 | 0.155546 | 2.178399 | 0.02937636 | NA |
| *PFN1* | -0.113685 | 0.052193 | -2.178163 | 0.02939391 | 0.306835784 |
| *RPLP1* | 0.194731 | 0.089417 | 2.177775 | 0.02942281 | 0.306835784 |

| | | | | | |
|---|---|---|---|---|---|
| *FAM91A1* | 0.348292 | 0.159976 | 2.177145 | 0.02946974 | 0.306835784 |
| *WDR13* | -0.313403 | 0.143954 | -2.177098 | 0.02947329 | 0.306835784 |
| *SYNGAP1.AS1* | 0.747139 | 0.343272 | 2.176519 | 0.02951644 | NA |
| *UBE3A* | 0.243435 | 0.111894 | 2.175593 | 0.02958572 | 0.307352291 |
| *EXOC3.AS1* | -0.475859 | 0.218915 | -2.173718 | 0.02972633 | NA |
| *AGMAT* | 0.577239 | 0.266013 | 2.169967 | 0.03000932 | NA |
| *PSMC3* | -0.262537 | 0.121005 | -2.169646 | 0.03003364 | 0.311344458 |
| *FANCL* | 0.486886 | 0.224458 | 2.169163 | 0.03007033 | NA |
| *ATXN7L2* | -0.881663 | 0.406456 | -2.169145 | 0.0300717 | NA |
| *C14orf119* | 0.209356 | 0.09657 | 2.167919 | 0.03016485 | 0.312043549 |
| *MRPL35* | 0.328154 | 0.151539 | 2.165476 | 0.03035126 | NA |
| *C1GALT1* | 0.339615 | 0.156834 | 2.165441 | 0.03035393 | 0.31333709 |
| *SIVA1* | 0.171268 | 0.079123 | 2.164569 | 0.0304207 | 0.313365211 |
| *PAN3* | 0.196186 | 0.090725 | 2.162428 | 0.03058518 | 0.314118511 |
| *SLC4A7* | 0.408706 | 0.189045 | 2.161948 | 0.03062222 | 0.314118511 |
| *MICU3* | 0.655942 | 0.303574 | 2.160732 | 0.03071602 | NA |
| *BLOC1S1* | -0.187987 | 0.087022 | -2.160235 | 0.03075448 | 0.314815187 |
| *ALDH3A2* | 0.903694 | 0.418366 | 2.160059 | 0.03076811 | NA |
| *CRYZ* | 0.641427 | 0.297035 | 2.159431 | 0.03081673 | NA |
| *VTI1A* | -0.330815 | 0.153198 | -2.159392 | 0.03081981 | NA |
| *BAD* | -0.280777 | 0.130108 | -2.158025 | 0.03092592 | 0.315909234 |
| *LINC00672* | 0.60809 | 0.281817 | 2.157751 | 0.03094716 | NA |
| *PYHIN1* | 1.014773 | 0.470429 | 2.157122 | 0.03099616 | NA |
| *NPHP3* | 0.408426 | 0.189413 | 2.156272 | 0.03106242 | 0.316233297 |
| *TTC21A* | 0.280964 | 0.13032 | 2.155959 | 0.0310869 | 0.316233297 |
| *MIR34AHG* | -1.868089 | 0.86687 | -2.154983 | 0.03116316 | NA |
| *AUP1* | -0.198407 | 0.092244 | -2.150904 | 0.03148374 | 0.318971913 |
| *ICAM3* | 0.23375 | 0.108718 | 2.150063 | 0.03155023 | 0.318971913 |
| *MANF* | 0.280768 | 0.130625 | 2.149416 | 0.03160142 | 0.318971913 |
| *DAAM1* | -0.325029 | 0.151269 | -2.148677 | 0.03166 | 0.318971913 |
| *TMEM138* | -0.308735 | 0.143705 | -2.148399 | 0.03168207 | 0.318971913 |
| *OAT* | 0.421171 | 0.19604 | 2.148391 | 0.03168272 | NA |
| *GRAP* | 0.393 | 0.183012 | 2.147396 | 0.03176173 | 0.31911738 |
| *L1TD1* | -0.767604 | 0.357539 | -2.146911 | 0.0318004 | NA |
| *NFS1* | 0.898964 | 0.418836 | 2.146339 | 0.03184598 | NA |
| *TMSB4Y* | -1.259929 | 0.587045 | -2.146223 | 0.03185517 | NA |
| *SSU72* | -0.215261 | 0.100372 | -2.144645 | 0.03198122 | 0.320664146 |
| *USP53* | 0.990879 | 0.462225 | 2.143713 | 0.03205587 | NA |
| *NFU1* | -0.229121 | 0.106919 | -2.14293 | 0.03211868 | 0.32138387 |
| *PDE3B* | -0.465589 | 0.217781 | -2.137882 | 0.03252631 | 0.324672224 |
| *TEX30* | 0.335917 | 0.157175 | 2.137217 | 0.03258038 | NA |
| *ARID4B* | 0.141121 | 0.06606 | 2.136267 | 0.03265766 | 0.324672224 |
| *ZMAT3* | -0.306232 | 0.143374 | -2.135893 | 0.03268813 | 0.324672224 |
| *LYRM2* | -0.184382 | 0.086338 | -2.135591 | 0.03271273 | 0.324672224 |
| *C12orf76* | 0.283776 | 0.132961 | 2.134275 | 0.03282032 | 0.325080567 |
| *FYN* | 0.541641 | 0.253856 | 2.133653 | 0.03287119 | NA |
| *DCTN4* | -0.467705 | 0.219244 | -2.133257 | 0.03290367 | NA |

| | | | | | |
|---|---|---|---|---|---|
| *OGT* | 0.361733 | 0.169584 | 2.133058 | 0.03291998 | 0.325408992 |
| *SNHG9* | -0.335347 | 0.157346 | -2.131268 | 0.03306704 | 0.325777956 |
| *MARCHF6* | 0.162642 | 0.076322 | 2.130984 | 0.03309047 | 0.325777956 |
| *ITM2A* | 0.903738 | 0.424297 | 2.129969 | 0.03317421 | NA |
| *ZC3H18* | 0.266024 | 0.124934 | 2.129319 | 0.03322785 | 0.325850903 |
| *PLPBP* | 0.237014 | 0.111312 | 2.12928 | 0.03323107 | 0.325850903 |
| *BAG2* | 0.415234 | 0.195204 | 2.127177 | 0.0334054 | NA |
| *CDCA4* | 0.455255 | 0.214039 | 2.126974 | 0.03342225 | NA |
| *ZMIZ2* | -0.387629 | 0.182345 | -2.125798 | 0.0335201 | 0.327630066 |
| *ABCB4* | 0.249472 | 0.117372 | 2.125482 | 0.03354643 | 0.327630066 |
| *GTF2IRD2* | -0.407263 | 0.191798 | -2.12339 | 0.03372121 | 0.328680999 |
| *ENSG00000273466* | 1.453839 | 0.684786 | 2.123054 | 0.03374934 | NA |
| *C6orf203* | 0.396694 | 0.186902 | 2.122477 | 0.03379774 | NA |
| *PSMG3* | -0.431164 | 0.203437 | -2.119402 | 0.03405651 | 0.331218411 |
| *GABARAP* | -0.301618 | 0.142361 | -2.118687 | 0.03411692 | 0.331218411 |
| *SHKBP1* | -0.321995 | 0.152074 | -2.117358 | 0.03422944 | 0.331652777 |
| *MAP3K8* | -0.446497 | 0.211197 | -2.114123 | 0.03450478 | 0.332962665 |
| *SYNC* | -0.416678 | 0.197196 | -2.113013 | 0.03459964 | 0.332962665 |
| *COL19A1* | -0.362375 | 0.171503 | -2.112931 | 0.03460665 | 0.332962665 |
| *ATP5ME* | -0.135501 | 0.064146 | -2.112393 | 0.03465271 | 0.332962665 |
| *TMEM99* | 0.477762 | 0.226228 | 2.111855 | 0.03469888 | NA |
| *FCRLA* | 0.333993 | 0.158157 | 2.111785 | 0.03470488 | 0.332962665 |
| *NKILA* | 0.811093 | 0.384564 | 2.109126 | 0.03493368 | NA |
| *TRMT112* | 0.187818 | 0.089066 | 2.108745 | 0.03496664 | 0.334817561 |
| *ARHGAP18* | 0.801295 | 0.380135 | 2.107925 | 0.03503752 | NA |
| *FADS1* | -0.462327 | 0.21937 | -2.107519 | 0.03507264 | NA |
| *NDUFA3* | -0.256226 | 0.121599 | -2.107142 | 0.03510529 | 0.335302133 |
| *FARSA* | 0.254352 | 0.120742 | 2.106577 | 0.0351543 | 0.335302133 |
| *SRRM2* | -0.122441 | 0.058215 | -2.103238 | 0.03544494 | 0.336055695 |
| *YAE1* | 0.605166 | 0.287763 | 2.102999 | 0.0354659 | NA |
| *JUNB* | -0.249411 | 0.118602 | -2.102927 | 0.03547212 | 0.336055695 |
| *FAM216A* | 0.383834 | 0.182535 | 2.102797 | 0.03548348 | NA |
| *RPL10A* | 0.199442 | 0.094869 | 2.102281 | 0.03552864 | 0.336055695 |
| *EPB41L2* | -0.405236 | 0.192767 | -2.102204 | 0.03553541 | 0.336055695 |
| *ATG4C* | 0.303164 | 0.144295 | 2.101005 | 0.03564051 | 0.336055695 |
| *LTBP3* | 0.268363 | 0.127734 | 2.10095 | 0.03564539 | 0.336055695 |
| *CCL5* | -1.23367 | 0.587379 | -2.100294 | 0.03570295 | NA |
| *ENSG00000270820* | 0.613259 | 0.292182 | 2.098889 | 0.03582665 | NA |
| *CRY1* | -0.526017 | 0.250629 | -2.098787 | 0.03583565 | NA |
| *PRDM8* | -0.588434 | 0.280392 | -2.098614 | 0.03585095 | NA |
| *SAR1B* | 0.392969 | 0.187261 | 2.098511 | 0.03586005 | 0.337141063 |
| *SLC38A11* | 1.210203 | 0.576858 | 2.097923 | 0.03591194 | NA |
| *CFLAR* | 0.263405 | 0.125578 | 2.097544 | 0.03594545 | 0.337141063 |
| *IDH2* | -0.254533 | 0.121363 | -2.097298 | 0.03596723 | 0.337141063 |
| *MRPS11* | -0.207918 | 0.099215 | -2.095621 | 0.03611583 | 0.337419276 |
| *DDX6* | -0.271711 | 0.12967 | -2.095407 | 0.03613483 | 0.337419276 |
| *HIST1H2BN* | 0.723943 | 0.345629 | 2.094566 | 0.03620962 | NA |

| | | | | | |
|---|---|---|---|---|---|
| *STK16* | -0.229098 | 0.109468 | -2.092832 | 0.0363642 | 0.337790199 |
| *SLC2A1* | -0.27136 | 0.129728 | -2.091755 | 0.03646046 | 0.337790199 |
| *CNIH4* | 0.275036 | 0.131552 | 2.090705 | 0.03655455 | 0.337790199 |
| *AQR* | 0.260539 | 0.124642 | 2.090295 | 0.03659131 | 0.337790199 |
| *TRIM44* | 0.20624 | 0.098668 | 2.09023 | 0.03659711 | 0.337790199 |
| *ERP29* | 0.13541 | 0.064784 | 2.090168 | 0.03660275 | 0.337790199 |
| *CCDC107* | -0.200192 | 0.095808 | -2.089515 | 0.03666135 | 0.337790199 |
| *MAPRE2* | 0.339325 | 0.162468 | 2.088563 | 0.03674708 | 0.337790199 |
| *TFDP2* | -0.261709 | 0.125338 | -2.088022 | 0.03679587 | 0.337790199 |
| *SRP19* | -0.172345 | 0.082585 | -2.086884 | 0.03689864 | 0.338099296 |
| *RASSF1* | 0.218847 | 0.10492 | 2.085833 | 0.03699374 | 0.338337138 |
| *NCBP1* | -0.504233 | 0.241808 | -2.085261 | 0.03704557 | NA |
| *FBXO36* | -1.145428 | 0.549595 | -2.084131 | 0.0371483 | NA |
| *ENSG00000277007* | 0.760875 | 0.365099 | 2.084023 | 0.03715808 | NA |
| *PCNX4* | 0.322904 | 0.154948 | 2.083948 | 0.03716492 | 0.338755039 |
| *SERP1* | 0.112403 | 0.053941 | 2.083805 | 0.0371779 | 0.338755039 |
| *WSB2* | -0.60955 | 0.29268 | -2.082654 | 0.03728278 | NA |
| *PNPLA7* | 0.556353 | 0.26729 | 2.081459 | 0.03739195 | NA |
| *GGCX* | -0.368852 | 0.177306 | -2.080305 | 0.03749753 | NA |
| *WDR11* | -0.277185 | 0.133407 | -2.077741 | 0.03773319 | 0.342774417 |
| *MED25* | -0.38109 | 0.183494 | -2.076855 | 0.03781496 | 0.342774417 |
| *ZNF667.AS1* | 0.400422 | 0.192816 | 2.076701 | 0.03782918 | 0.342774417 |
| *RPL27* | 0.133333 | 0.064259 | 2.074934 | 0.03799267 | 0.343619444 |
| *SNED1* | -1.123316 | 0.541385 | -2.074894 | 0.03799636 | NA |
| *AKTIP* | 0.521581 | 0.251456 | 2.074244 | 0.03805669 | NA |
| *NAB2* | 0.79764 | 0.384972 | 2.071941 | 0.0382709 | NA |
| *PTGES3* | 0.171657 | 0.082849 | 2.071937 | 0.03827134 | 0.345501236 |
| *THRB* | 2.002988 | 0.966801 | 2.07177 | 0.03828691 | NA |
| *YTHDC1* | 0.175121 | 0.084628 | 2.069289 | 0.038519 | 0.347096649 |
| *CCDC157* | -0.7678 | 0.371183 | -2.068518 | 0.03859134 | NA |
| *WASF1* | -0.544075 | 0.263085 | -2.068055 | 0.03863482 | NA |
| *TUBB4B* | 0.427345 | 0.206764 | 2.066824 | 0.03875073 | 0.347673644 |
| *ATP8A1* | -0.310724 | 0.150357 | -2.066574 | 0.03877433 | 0.347673644 |
| *ENSG00000258297* | -0.334702 | 0.161963 | -2.066529 | 0.03877852 | NA |
| *SLC43A2* | -0.214103 | 0.103614 | -2.066342 | 0.0387962 | 0.347673644 |
| *IRAK1* | -0.461991 | 0.223597 | -2.066173 | 0.03881217 | NA |
| *N6AMT1* | 0.593487 | 0.287371 | 2.06523 | 0.03890129 | NA |
| *EIF5* | 0.128413 | 0.062195 | 2.064669 | 0.03895433 | 0.348175087 |
| *PTRH1* | -0.359785 | 0.174269 | -2.064545 | 0.03896607 | NA |
| *ARL11* | 0.486967 | 0.235905 | 2.064245 | 0.03899447 | 0.348175087 |
| *WBP1L* | -0.326554 | 0.158211 | -2.064045 | 0.03901348 | NA |
| *TMEM177* | -0.608039 | 0.294746 | -2.062928 | 0.03911951 | NA |
| *SYNJ2BP* | 0.225364 | 0.10932 | 2.061506 | 0.03925479 | 0.349860969 |
| *WDR89* | 0.368041 | 0.178542 | 2.061373 | 0.03926751 | NA |
| *LINGO3* | -0.488141 | 0.236809 | -2.061333 | 0.03927131 | NA |
| *WDR34* | -0.307586 | 0.149274 | -2.060554 | 0.03934562 | 0.350032955 |
| *LINC00562* | -0.876489 | 0.42553 | -2.059758 | 0.03942167 | NA |

| | | | | | |
|---|---|---|---|---|---|
| *CHD1* | 0.199472 | 0.096873 | 2.059102 | 0.03948451 | 0.350537667 |
| *LINC01184* | 0.30932 | 0.150308 | 2.057907 | 0.03959902 | NA |
| *FAM117B* | -0.390653 | 0.189837 | -2.05784 | 0.03960545 | 0.350537667 |
| *AMD1* | 0.219065 | 0.106464 | 2.057643 | 0.03962438 | 0.350537667 |
| *OR2A1.AS1* | -0.903568 | 0.439145 | -2.05756 | 0.03963242 | NA |
| *ENSG00000244459* | 0.454549 | 0.220972 | 2.057042 | 0.03968215 | NA |
| *MRNIP* | 0.249678 | 0.121386 | 2.056903 | 0.0396956 | 0.350537667 |
| *PCGF1* | -0.619752 | 0.301409 | -2.056184 | 0.03976477 | NA |
| *MYNN* | 0.342994 | 0.166841 | 2.055813 | 0.03980058 | NA |
| *ACCS* | 0.704202 | 0.342662 | 2.05509 | 0.03987029 | NA |
| *MFSD13A* | -0.595192 | 0.289625 | -2.055046 | 0.03987457 | NA |
| *ZNF318* | 0.424584 | 0.206615 | 2.054948 | 0.03988402 | 0.350537667 |
| *PRCD* | 1.005907 | 0.489512 | 2.054917 | 0.03988701 | NA |
| *ENSG00000267939* | -1.514017 | 0.736942 | -2.05446 | 0.03993119 | NA |
| *PCF11* | 0.173448 | 0.084427 | 2.054408 | 0.03993623 | 0.350537667 |
| *CISD2* | 0.226925 | 0.110466 | 2.054257 | 0.0399508 | 0.350537667 |
| *CIAO2B* | 0.135959 | 0.066205 | 2.053605 | 0.04001391 | 0.350537667 |
| *PSMA4* | -0.153836 | 0.074923 | -2.053263 | 0.04004712 | 0.350537667 |
| *MTMR6* | -0.357768 | 0.174268 | -2.05297 | 0.04007547 | NA |
| *MBNL1* | -0.155684 | 0.075887 | -2.051531 | 0.04021529 | 0.351191722 |
| *MED28* | 0.150678 | 0.073465 | 2.051016 | 0.04026539 | 0.351191722 |
| *TMX2* | 0.453229 | 0.221065 | 2.050211 | 0.04034386 | 0.351250049 |
| *OSGIN2* | 0.556272 | 0.27141 | 2.049563 | 0.04040713 | NA |
| *SNAP23* | -0.185228 | 0.090423 | -2.048462 | 0.04051478 | 0.352111603 |
| *CCDC71* | -0.336216 | 0.16432 | -2.046107 | 0.04074586 | NA |
| *LAMTOR3* | 0.301822 | 0.147561 | 2.045407 | 0.04081478 | 0.353803136 |
| *ENSG00000274265* | -0.291366 | 0.142476 | -2.045008 | 0.04085403 | 0.353803136 |
| *ENSG00000226644* | -1.948206 | 0.953322 | -2.043596 | 0.04099346 | NA |
| *SNRPA1* | -0.193406 | 0.094693 | -2.042456 | 0.04110637 | 0.355274877 |
| *SETDB2* | -0.373755 | 0.183112 | -2.041126 | 0.04123831 | 0.355274877 |
| *STMN1* | -0.300985 | 0.147463 | -2.041091 | 0.0412418 | 0.355274877 |
| *GPR137B* | 0.724951 | 0.355292 | 2.040434 | 0.04130708 | NA |
| *ENSG00000276509* | 0.438521 | 0.215074 | 2.038931 | 0.04145688 | NA |
| *ADCK1* | -0.918081 | 0.450328 | -2.038695 | 0.0414805 | NA |
| *ASAH1* | 0.255154 | 0.125184 | 2.038236 | 0.04152637 | 0.357097544 |
| *SWAP70* | 0.187523 | 0.092085 | 2.036417 | 0.0417085 | 0.357779075 |
| *ANAPC15* | -0.322941 | 0.158617 | -2.035985 | 0.04175186 | 0.357779075 |
| *ZKSCAN4* | 0.58723 | 0.288507 | 2.035412 | 0.04180939 | NA |
| *CLEC16A* | -0.483199 | 0.237403 | -2.035348 | 0.04181585 | NA |
| *PPP5C* | -0.513872 | 0.252595 | -2.034373 | 0.04191404 | NA |
| *TMSB10* | -0.169801 | 0.083503 | -2.033466 | 0.04200549 | 0.359323192 |
| *ENSG00000261669* | 0.789415 | 0.388459 | 2.032174 | 0.04213604 | NA |
| *S100A13* | -0.719974 | 0.354324 | -2.031963 | 0.0421574 | NA |
| *KLHDC7B* | -0.68298 | 0.336208 | -2.031421 | 0.04221227 | NA |
| *LSR* | 0.774424 | 0.381431 | 2.030312 | 0.04232483 | NA |
| *VASH1* | -1.561613 | 0.769268 | -2.029998 | 0.04235672 | NA |
| *ASB16.AS1* | 0.324484 | 0.159889 | 2.029427 | 0.04241479 | NA |

| | | | | | |
|---|---|---|---|---|---|
| *DDX55* | 0.352222 | 0.173683 | 2.027953 | 0.04256507 | NA |
| *RASA2* | 0.285232 | 0.14068 | 2.02752 | 0.04260929 | 0.362790343 |
| *JTB* | -0.100343 | 0.049505 | -2.02695 | 0.04266748 | 0.362790343 |
| *EIF3J.DT* | 0.222961 | 0.110011 | 2.026714 | 0.04269162 | 0.362790343 |
| *UNC45A* | -0.302401 | 0.149219 | -2.02656 | 0.04270739 | 0.362790343 |
| *TAPBPL* | 0.394635 | 0.194955 | 2.024241 | 0.04294541 | 0.363968582 |
| *TENT4B* | 0.426684 | 0.210818 | 2.023949 | 0.0429754 | NA |
| *MPV17* | 0.284802 | 0.140798 | 2.022771 | 0.04309678 | 0.363968582 |
| *HIF1A* | 0.212589 | 0.105106 | 2.02261 | 0.04311333 | 0.363968582 |
| *USF1* | -0.655907 | 0.324307 | -2.022485 | 0.04312623 | NA |
| *SLC2A14* | -3.245464 | 1.604725 | -2.022442 | 0.04313069 | NA |
| *TMX4* | -0.307133 | 0.15194 | -2.021409 | 0.04323746 | 0.363968582 |
| *CD3G* | -2.604577 | 1.288512 | -2.021385 | 0.04323998 | NA |
| *RPS6KB2* | -0.209756 | 0.103778 | -2.021194 | 0.04325969 | 0.363968582 |
| *XRRA1* | -0.587078 | 0.290507 | -2.020878 | 0.0432924 | 0.363968582 |
| *GRK2* | 0.240396 | 0.119003 | 2.02009 | 0.04337402 | 0.364029253 |
| *MRPS36* | 0.249804 | 0.123859 | 2.016843 | 0.04371187 | 0.366236594 |
| *PSMD9* | -0.236012 | 0.117099 | -2.015497 | 0.04385265 | 0.366788069 |
| *RNF144B* | 0.705911 | 0.350565 | 2.013638 | 0.04404757 | NA |
| *IGHA2* | -0.877752 | 0.435938 | -2.013482 | 0.04406398 | NA |
| *DNASE1* | 0.288262 | 0.14317 | 2.013429 | 0.04406952 | 0.367972987 |
| *C15orf41* | -0.748959 | 0.37202 | -2.013223 | 0.04409114 | NA |
| *THG1L* | 0.428212 | 0.212839 | 2.011903 | 0.04423013 | 0.368684879 |
| *TMBIM6* | -0.127894 | 0.063604 | -2.010791 | 0.04434752 | 0.369034717 |
| *ZNF610* | 1.261166 | 0.627364 | 2.010261 | 0.04440358 | NA |
| *CEP85L* | 0.639366 | 0.318148 | 2.009649 | 0.0444684 | NA |
| *CLDN11* | -1.201339 | 0.598005 | -2.008911 | 0.04454654 | NA |
| *IDH1* | 0.237243 | 0.118102 | 2.008803 | 0.04455803 | 0.370156968 |
| *PRRT3* | 0.553075 | 0.275667 | 2.006314 | 0.04482279 | NA |
| *KYAT3* | 0.435636 | 0.217196 | 2.005726 | 0.04488553 | 0.372245622 |
| *H3F3C* | -0.979094 | 0.488413 | -2.004645 | 0.04500105 | NA |
| *HINT1* | 0.184085 | 0.091861 | 2.003942 | 0.04507624 | 0.372521123 |
| *AKAP9* | 0.159574 | 0.079639 | 2.003722 | 0.04509981 | 0.372521123 |
| *PLAC8* | 0.198523 | 0.099129 | 2.002675 | 0.04521213 | 0.372521123 |
| *PLEKHF1* | 0.523083 | 0.261284 | 2.001973 | 0.04528764 | NA |
| *RALGPS2* | 0.275388 | 0.137567 | 2.001845 | 0.04530146 | 0.372521123 |
| *ERLIN1* | 0.571408 | 0.285494 | 2.001472 | 0.04534158 | NA |
| *BAX* | -0.339753 | 0.169805 | -2.000845 | 0.04540905 | 0.372521123 |
| *PHF14* | 0.191864 | 0.095923 | 2.000194 | 0.04547928 | 0.372521123 |
| *SF1* | -0.15519 | 0.077605 | -1.999741 | 0.0455282 | 0.372521123 |
| *CEBPZ* | 0.213787 | 0.107002 | 1.997962 | 0.04572081 | 0.372521123 |
| *IGKC* | -0.318196 | 0.159269 | -1.997848 | 0.04573311 | 0.372521123 |
| *CERS4* | -0.291214 | 0.145811 | -1.997197 | 0.04580376 | 0.372521123 |
| *ZNF562* | -0.236458 | 0.118396 | -1.997188 | 0.04580478 | 0.372521123 |
| *RPF2* | 0.271361 | 0.135919 | 1.996493 | 0.04588028 | 0.372521123 |
| *CXorf21* | 0.288648 | 0.144596 | 1.996234 | 0.04590849 | 0.372521123 |
| *USP5* | -0.383487 | 0.192188 | -1.99537 | 0.04600253 | NA |

| | | | | | |
|---|---|---|---|---|---|
| *SELENOW* | -0.277616 | 0.139145 | -1.995163 | 0.04602506 | 0.372610949 |
| *HVCN1* | -0.201188 | 0.100869 | -1.994549 | 0.04609209 | 0.372610949 |
| *PLSCR1* | -0.297184 | 0.149084 | -1.993406 | 0.04621705 | 0.372610949 |
| *TACC3* | 0.312183 | 0.156641 | 1.992987 | 0.04626282 | 0.372610949 |
| *C1orf52* | 0.224118 | 0.112472 | 1.992645 | 0.04630032 | 0.372610949 |
| *SBNO2* | -0.621366 | 0.311922 | -1.992054 | 0.04636511 | NA |
| *FCRL3* | -1.05357 | 0.529041 | -1.991471 | 0.04642917 | 0.37303437 |
| *SLC7A6* | 0.355313 | 0.178457 | 1.991029 | 0.04647771 | NA |
| *WAC* | 0.1818 | 0.091381 | 1.989472 | 0.04664918 | 0.374187581 |
| *LRRC7* | -0.626442 | 0.314908 | -1.989287 | 0.04666958 | NA |
| *ENSG00000258634* | -1.012905 | 0.509289 | -1.988863 | 0.04671633 | NA |
| *GPAA1* | -0.247995 | 0.124737 | -1.988141 | 0.0467961 | 0.374518013 |
| *GUK1* | -0.187762 | 0.094518 | -1.986531 | 0.04697437 | 0.374518013 |
| *FBXO25* | -0.329246 | 0.165742 | -1.986499 | 0.04697799 | 0.374518013 |
| *MIB2* | 0.286571 | 0.144272 | 1.986331 | 0.04699654 | 0.374518013 |
| *ZNF584* | 0.55274 | 0.278427 | 1.985227 | 0.04711925 | NA |
| *SLC2A5* | -0.43702 | 0.220161 | -1.985005 | 0.04714393 | 0.37500056 |
| *KXD1* | -0.219631 | 0.110678 | -1.984408 | 0.04721037 | 0.37500056 |
| *C12orf49* | -0.508407 | 0.25629 | -1.983716 | 0.04728752 | 0.375004586 |
| *CLTA* | 0.151105 | 0.076236 | 1.982064 | 0.04747204 | 0.375858695 |
| *MRPL38* | -0.488016 | 0.246318 | -1.981245 | 0.0475638 | NA |
| *ZNF830* | 0.229563 | 0.115879 | 1.98106 | 0.04758452 | 0.376140607 |
| *ENTPD1* | 0.368491 | 0.186099 | 1.980075 | 0.04769515 | NA |
| *LY9* | 0.361945 | 0.182832 | 1.979657 | 0.0477421 | 0.376228172 |
| *PSMD13* | -0.215961 | 0.109094 | -1.979592 | 0.04774938 | 0.376228172 |
| *C17orf58* | -0.828588 | 0.418616 | -1.97935 | 0.04777663 | NA |
| *ATP6V0E2* | -0.471573 | 0.238454 | -1.977628 | 0.04797066 | NA |
| *GORASP1* | 0.359941 | 0.182258 | 1.974897 | 0.04827979 | NA |
| *MTRF1L* | 0.45072 | 0.228277 | 1.974444 | 0.04833132 | 0.379491388 |
| *NCF1* | -0.15496 | 0.07853 | -1.973248 | 0.04846736 | 0.379491388 |
| *MOAP1* | -0.306584 | 0.155462 | -1.972075 | 0.04860109 | 0.379491388 |
| *FKBP3* | -0.195745 | 0.099341 | -1.970434 | 0.04878866 | 0.379491388 |
| *SPOCK2* | -0.185321 | 0.094077 | -1.969889 | 0.04885109 | 0.379491388 |
| *ENSG00000272927* | -0.575274 | 0.292074 | -1.969613 | 0.04888276 | NA |
| *PTPRC* | 0.164171 | 0.083377 | 1.969021 | 0.04895071 | 0.379491388 |
| *NFE2L2* | 0.246302 | 0.125106 | 1.968746 | 0.04898221 | 0.379491388 |
| *C10orf88* | 0.527299 | 0.267947 | 1.967921 | 0.04907707 | NA |
| *TMED2* | -0.194692 | 0.098941 | -1.967753 | 0.04909643 | 0.379491388 |
| *PMF1* | -0.233724 | 0.118784 | -1.967644 | 0.04910899 | 0.379491388 |
| *NSMCE1* | -0.266333 | 0.135418 | -1.966742 | 0.04921302 | 0.379491388 |
| *RHOBTB2* | 0.823117 | 0.418569 | 1.966504 | 0.04924037 | NA |
| *IRAK4* | -0.235962 | 0.119998 | -1.966384 | 0.04925426 | 0.379491388 |
| *PRDX1* | 0.163638 | 0.083238 | 1.965916 | 0.04930827 | 0.379491388 |
| *PLEKHG1* | -0.327123 | 0.166446 | -1.965333 | 0.04937576 | 0.379491388 |
| *TIMM23* | -0.426228 | 0.216883 | -1.965241 | 0.0493864 | NA |
| *PSMB4* | -0.153177 | 0.077946 | -1.965166 | 0.04939501 | 0.379491388 |
| *NRAS* | -0.322054 | 0.163923 | -1.964672 | 0.04945225 | 0.379491388 |

| | | | | | |
|---|---|---|---|---|---|
| *MGAT4A* | -0.473059 | 0.240792 | -1.964597 | 0.04946088 | NA |
| *FAM222B* | 0.635691 | 0.323577 | 1.964574 | 0.04946362 | NA |
| *SNAPC5* | -0.326219 | 0.166129 | -1.963642 | 0.04957167 | 0.379491388 |
| *TNFRSF13B* | 0.837147 | 0.426341 | 1.963563 | 0.04958077 | 0.379491388 |
| *IER3* | -0.675373 | 0.344026 | -1.963146 | 0.04962922 | NA |
| *MDH2* | -0.169608 | 0.086426 | -1.962459 | 0.04970908 | 0.379491388 |
| *FCMR* | 0.220933 | 0.112603 | 1.962045 | 0.04975726 | 0.379491388 |
| *RSRC2* | 0.237868 | 0.121265 | 1.961553 | 0.04981453 | 0.379491388 |
| *MTPN* | 0.169356 | 0.086342 | 1.961454 | 0.04982603 | 0.379491388 |
| *CPT1A* | -0.445588 | 0.22718 | -1.961385 | 0.04983417 | NA |
| *PEX13* | 0.363368 | 0.185266 | 1.961337 | 0.04983968 | NA |
| *SBNO1* | 0.205894 | 0.105003 | 1.960846 | 0.04989694 | 0.379491388 |
| *TMC6* | -0.194462 | 0.099195 | -1.960414 | 0.04994736 | 0.379491388 |

| Cluster 5 | log2FC | lfcSE | stat | pvalue | padj |
|---|---|---|---|---|---|
| *NFKBIA* | -0.71892 | 0.066472 | -10.815443 | < 2.22e-16 | < 2.22e-16 |
| *LTB* | -1.092821 | 0.119951 | -9.110537 | < 2.22e-16 | 4.67E-16 |
| *IL13RA1* | -1.515699 | 0.167368 | -9.056083 | < 2.22e-16 | 5.13E-16 |
| *NCR3* | -1.651037 | 0.186618 | -8.847142 | < 2.22e-16 | 2.56E-15 |
| *MARCKSL1* | -1.022467 | 0.13046 | -7.83741 | 4.60E-15 | 1.05E-11 |
| *NCF4* | -0.707989 | 0.091842 | -7.708787 | 1.27E-14 | 2.41E-11 |
| *ENSG00000266844* | -1.79307 | 0.233662 | -7.673767 | 1.67E-14 | 2.72E-11 |
| *NT5C* | -0.624744 | 0.089057 | -7.015135 | 2.30E-12 | 3.27E-09 |
| *COCH* | -1.311671 | 0.189993 | -6.903799 | 5.06E-12 | 5.85E-09 |
| *FNBP1* | -0.690901 | 0.100104 | -6.901803 | 5.13E-12 | 5.85E-09 |
| *CTSH* | -0.596976 | 0.089192 | -6.69319 | 2.18E-11 | 2.26E-08 |
| *COL18A1* | -2.216711 | 0.345756 | -6.411198 | 1.44E-10 | 1.37E-07 |
| *SEC62* | 0.36247 | 0.05667 | 6.396209 | 1.59E-10 | 1.40E-07 |
| *TEX9* | -1.715713 | 0.268807 | -6.382699 | 1.74E-10 | 1.40E-07 |
| *TAGLN2* | -0.566962 | 0.088958 | -6.373399 | 1.85E-10 | 1.40E-07 |
| *SYNGR2* | -0.681794 | 0.108771 | -6.268169 | 3.65E-10 | 2.46E-07 |
| *MAP3K7CL* | -1.180419 | 0.188346 | -6.267298 | 3.67E-10 | 2.46E-07 |
| *BLNK* | 0.453643 | 0.074061 | 6.125238 | 9.05E-10 | 5.73E-07 |
| *JPT1* | -0.733566 | 0.120115 | -6.10718 | 1.01E-09 | 6.08E-07 |
| *GLIPR1* | 0.525834 | 0.086481 | 6.08031 | 1.20E-09 | 6.83E-07 |
| *MARCKS* | -0.936965 | 0.155214 | -6.036604 | 1.57E-09 | 8.53E-07 |
| *IL23A* | -1.374582 | 0.228167 | -6.024458 | 1.70E-09 | 8.78E-07 |
| *PDE4D* | -1.415602 | 0.236425 | -5.987523 | 2.13E-09 | 1.05E-06 |
| *CR1* | -1.349354 | 0.231741 | -5.822691 | 5.79E-09 | 2.61E-06 |
| *CYTIP* | 0.428279 | 0.073625 | 5.817029 | 5.99E-09 | 2.61E-06 |
| *TCTN1* | -1.000957 | 0.172158 | -5.814185 | 6.09E-09 | 2.61E-06 |
| *CPNE5* | -0.878492 | 0.151155 | -5.811852 | 6.18E-09 | 2.61E-06 |
| *CD40* | -0.44205 | 0.076466 | -5.780992 | 7.43E-09 | 3.02E-06 |
| *GPSM3* | -0.271399 | 0.047247 | -5.744309 | 9.23E-09 | 3.62E-06 |
| *PDE4B* | 0.597773 | 0.104822 | 5.702758 | 1.18E-08 | 4.47E-06 |
| *MFSD10* | -0.679551 | 0.120057 | -5.660214 | 1.51E-08 | 5.55E-06 |
| *LMO4* | 0.658842 | 0.116833 | 5.63918 | 1.71E-08 | 6.02E-06 |
| *MAL* | -2.549903 | 0.452477 | -5.635434 | 1.75E-08 | 6.02E-06 |
| *PSMB10* | -0.488912 | 0.086967 | -5.62182 | 1.89E-08 | 6.33E-06 |
| *RSRP1* | 0.463248 | 0.08278 | 5.596138 | 2.19E-08 | 7.13E-06 |
| *ZEB2* | 0.909023 | 0.163524 | 5.558962 | 2.71E-08 | 8.58E-06 |
| *KLF3* | 0.735405 | 0.132995 | 5.52957 | 3.21E-08 | 9.88E-06 |
| *AK8* | -1.940642 | 0.354341 | -5.476766 | 4.33E-08 | 1.30E-05 |
| *SAMD12* | -2.164757 | 0.39563 | -5.471666 | 4.46E-08 | 1.30E-05 |
| *RNGTT* | -1.123604 | 0.206787 | -5.433616 | 5.52E-08 | 1.57E-05 |
| *GDPD5* | -1.60711 | 0.296093 | -5.427726 | 5.71E-08 | 1.58E-05 |
| *ENSG00000167414* | -2.354116 | 0.434019 | -5.423992 | 5.83E-08 | 1.58E-05 |
| *LINC00926* | -0.569602 | 0.105633 | -5.392254 | 6.96E-08 | 1.84E-05 |
| *ATF5* | -1.472486 | 0.274234 | -5.369446 | 7.90E-08 | 2.04E-05 |
| *LYPLAL1* | -0.841422 | 0.157202 | -5.352487 | 8.68E-08 | 2.20E-05 |
| *ENSG00000125726* | -1.319104 | 0.246981 | -5.340903 | 9.25E-08 | 2.25E-05 |

| | | | | | |
|---|---|---|---|---|---|
| *PPM1K* | 0.438424 | 0.082098 | 5.340248 | 9.28E-08 | 2.25E-05 |
| *C9orf16* | -0.34711 | 0.065287 | -5.316661 | 1.06E-07 | 2.51E-05 |
| *MIF* | -0.464604 | 0.087502 | -5.30965 | 1.10E-07 | 2.55E-05 |
| *IGHE* | -1.649871 | 0.31125 | -5.300791 | 1.15E-07 | 2.63E-05 |
| *RNF130* | 1.336643 | 0.25236 | 5.296573 | 1.18E-07 | 2.63E-05 |
| *P2RY8* | -0.60655 | 0.114982 | -5.275156 | 1.33E-07 | 2.90E-05 |
| *PASK* | -1.079442 | 0.205172 | -5.261158 | 1.43E-07 | 3.04E-05 |
| *ATXN1* | 1.194377 | 0.227221 | 5.256467 | 1.47E-07 | 3.04E-05 |
| *FAM177B* | 1.398392 | 0.266041 | 5.256303 | 1.47E-07 | 3.04E-05 |
| *RABAC1* | 0.301912 | 0.05756 | 5.245185 | 1.56E-07 | 3.17E-05 |
| *TENT5A* | 1.638367 | 0.312694 | 5.239527 | 1.61E-07 | 3.22E-05 |
| *ITGAL* | -0.619498 | 0.119484 | -5.184784 | 2.16E-07 | 4.25E-05 |
| *MRPL54* | -0.4646 | 0.089773 | -5.175268 | 2.28E-07 | 4.39E-05 |
| *DUSP10* | 1.031837 | 0.200699 | 5.141221 | 2.73E-07 | 5.18E-05 |
| *PFN1* | -0.242586 | 0.047381 | -5.119873 | 3.06E-07 | 5.71E-05 |
| *MYL12A* | -0.28048 | 0.054951 | -5.104216 | 3.32E-07 | 6.10E-05 |
| *MSC* | -1.828122 | 0.358846 | -5.094445 | 3.50E-07 | 6.32E-05 |
| *ETV7* | -1.89343 | 0.372694 | -5.080388 | 3.77E-07 | 6.70E-05 |
| *ZBED2* | 1.789543 | 0.354388 | 5.049672 | 4.43E-07 | 7.75E-05 |
| *HMGB2* | -0.437343 | 0.086687 | -5.045097 | 4.53E-07 | 7.82E-05 |
| *CD84* | 1.04783 | 0.208393 | 5.028146 | 4.95E-07 | 8.39E-05 |
| *ATP6V0E2* | -0.861012 | 0.171313 | -5.025964 | 5.01E-07 | 8.39E-05 |
| *TSC22D3* | 0.46229 | 0.092505 | 4.997438 | 5.81E-07 | 9.59E-05 |
| *CLK1* | 0.490127 | 0.098466 | 4.977616 | 6.44E-07 | 0.000104706 |
| *PDE3B* | -1.263605 | 0.255067 | -4.954014 | 7.27E-07 | 0.000115178 |
| *GATD3A* | 5.051733 | 1.0198 | 4.953651 | 7.28E-07 | 0.000115178 |
| *CAPN2* | -0.779804 | 0.158235 | -4.928151 | 8.30E-07 | 0.000129475 |
| *SQOR* | -0.647609 | 0.131992 | -4.906427 | 9.28E-07 | 0.000141929 |
| *LINC02397* | 0.407962 | 0.083175 | 4.90487 | 9.35E-07 | 0.000141929 |
| *CYCS* | -0.349476 | 0.071587 | -4.881852 | 1.05E-06 | 0.000157447 |
| *EPS15* | -0.541886 | 0.111121 | -4.876557 | 1.08E-06 | 0.000159631 |
| *TAGAP* | -0.601184 | 0.124245 | -4.838689 | 1.31E-06 | 0.000190786 |
| *TENT5C* | 0.897257 | 0.185677 | 4.832345 | 1.35E-06 | 0.000193421 |
| *FAS* | -1.17531 | 0.243289 | -4.830924 | 1.36E-06 | 0.000193421 |
| *UBE2I* | -0.281176 | 0.058348 | -4.818924 | 1.44E-06 | 0.000202888 |
| *DAP* | -0.620915 | 0.12912 | -4.808813 | 1.52E-06 | 0.000210821 |
| *TFEC* | 1.090121 | 0.227233 | 4.797378 | 1.61E-06 | 0.000220527 |
| *BIK* | -1.666123 | 0.347762 | -4.790987 | 1.66E-06 | 0.000224959 |
| *STX7* | 0.609687 | 0.127858 | 4.768482 | 1.86E-06 | 0.000244199 |
| *MIIP* | -0.483322 | 0.101365 | -4.768128 | 1.86E-06 | 0.000244199 |
| *NECAP2* | -0.42313 | 0.088754 | -4.767429 | 1.87E-06 | 0.000244199 |
| *TGIF1* | 0.737375 | 0.155423 | 4.744311 | 2.09E-06 | 0.000270698 |
| *MIR3142HG* | -2.031154 | 0.429 | -4.734629 | 2.19E-06 | 0.000280756 |
| *TTC32* | -0.950531 | 0.200956 | -4.730046 | 2.24E-06 | 0.000283979 |
| *BBX* | 0.403946 | 0.08565 | 4.716256 | 2.40E-06 | 0.000300571 |
| *BPTF* | 0.36769 | 0.078193 | 4.70234 | 2.57E-06 | 0.000318309 |
| *OPTN* | -0.830357 | 0.177035 | -4.690354 | 2.73E-06 | 0.000333196 |

| | | | | | |
|---|---|---|---|---|---|
| *EHD1* | -0.550067 | 0.11732 | -4.688602 | 2.75E-06 | 0.000333196 |
| *MVP* | -0.635706 | 0.135665 | -4.68585 | 2.79E-06 | 0.000334149 |
| *LCN8* | 4.670297 | 0.999921 | 4.670665 | 3.00E-06 | 0.000356081 |
| *EBI3* | -1.048703 | 0.225459 | -4.651419 | 3.30E-06 | 0.000384235 |
| *NSF* | 1.28705 | 0.27674 | 4.65076 | 3.31E-06 | 0.000384235 |
| *MRPL36* | -0.441451 | 0.095013 | -4.646196 | 3.38E-06 | 0.000388862 |
| *CDC42* | -0.3975 | 0.085799 | -4.632905 | 3.61E-06 | 0.000410544 |
| *IL6R* | -1.572316 | 0.341134 | -4.609082 | 4.04E-06 | 0.000450328 |
| *C12orf75* | -1.095261 | 0.237659 | -4.608532 | 4.06E-06 | 0.000450328 |
| *SLC38A2* | 0.499819 | 0.108477 | 4.607584 | 4.07E-06 | 0.000450328 |
| *PRKCB* | -0.435596 | 0.094686 | -4.600411 | 4.22E-06 | 0.000461635 |
| *TESC* | -1.903574 | 0.41514 | -4.585375 | 4.53E-06 | 0.000491411 |
| *ANXA7* | -0.531131 | 0.116476 | -4.559994 | 5.12E-06 | 0.000549482 |
| *CDKN1A* | -0.605111 | 0.132857 | -4.554592 | 5.25E-06 | 0.000558524 |
| *PLAAT3* | -1.588497 | 0.349143 | -4.549707 | 5.37E-06 | 0.000566355 |
| *CIRBP* | -0.20639 | 0.045417 | -4.544288 | 5.51E-06 | 0.00057579 |
| *NMB* | 1.413277 | 0.311416 | 4.53823 | 5.67E-06 | 0.000587191 |
| *LRIF1* | 0.620261 | 0.136837 | 4.532836 | 5.82E-06 | 0.000596966 |
| *HRK* | -0.962769 | 0.212544 | -4.529732 | 5.91E-06 | 0.000600395 |
| *TIMM10* | -0.506074 | 0.111813 | -4.526085 | 6.01E-06 | 0.000605436 |
| *TRADD* | -0.545352 | 0.120647 | -4.520236 | 6.18E-06 | 0.000616948 |
| *JAM3* | 2.20131 | 0.487352 | 4.516881 | 6.28E-06 | 0.000621351 |
| *IER3* | -1.676992 | 0.372002 | -4.508016 | 6.54E-06 | 0.000641362 |
| *PRELID1* | -0.401076 | 0.088999 | -4.506502 | 6.59E-06 | 0.000641362 |
| *NDUFB8* | -0.275559 | 0.061319 | -4.493899 | 6.99E-06 | 0.000674772 |
| *SUB1* | -0.516588 | 0.116001 | -4.453326 | 8.46E-06 | 0.000808983 |
| *UBE2N* | -0.480593 | 0.108116 | -4.445176 | 8.78E-06 | 0.00083183 |
| *PRDX2* | -0.529166 | 0.119115 | -4.442485 | 8.89E-06 | 0.00083183 |
| *CD48* | 0.441313 | 0.09935 | 4.441992 | 8.91E-06 | 0.00083183 |
| *TELO2* | -0.948111 | 0.214533 | -4.419408 | 9.90E-06 | 0.000909001 |
| *LGALS3* | -0.705779 | 0.159713 | -4.419054 | 9.91E-06 | 0.000909001 |
| *CERS6* | 1.83683 | 0.415796 | 4.41762 | 9.98E-06 | 0.000909001 |
| *ENSG00000070423* | -0.471663 | 0.10684 | -4.414664 | 1.01E-05 | 0.000914198 |
| *ISCU* | -0.351577 | 0.079695 | -4.411513 | 1.03E-05 | 0.000920299 |
| *BIN2* | -0.695872 | 0.157885 | -4.407465 | 1.05E-05 | 0.000928049 |
| *ITGB1* | -0.845121 | 0.191863 | -4.404812 | 1.06E-05 | 0.000928049 |
| *LINC02384* | -1.561015 | 0.354402 | -4.404639 | 1.06E-05 | 0.000928049 |
| *RGS2* | 1.30378 | 0.296885 | 4.391533 | 1.13E-05 | 0.000978278 |
| *RNF207* | -1.108342 | 0.253097 | -4.379129 | 1.19E-05 | 0.001027798 |
| *RORA* | -1.253165 | 0.286475 | -4.37443 | 1.22E-05 | 0.00104229 |
| *DHTKD1* | -0.539133 | 0.123935 | -4.350122 | 1.36E-05 | 0.001155896 |
| *REL* | -0.372591 | 0.085682 | -4.348534 | 1.37E-05 | 0.001155896 |
| *SRSF5* | -0.277474 | 0.063988 | -4.336367 | 1.45E-05 | 0.001211658 |
| *RPP25* | 1.088914 | 0.251194 | 4.334954 | 1.46E-05 | 0.001211658 |
| *HNRNPDL* | -0.247814 | 0.057206 | -4.331965 | 1.48E-05 | 0.001219326 |
| *PCDH9* | 1.132925 | 0.262148 | 4.321706 | 1.55E-05 | 0.001268251 |
| *AEN* | -0.561603 | 0.130215 | -4.3129 | 1.61E-05 | 0.00131042 |

| | | | | | |
|---|---|---|---|---|---|
| *SLC25A39* | -0.355271 | 0.082594 | -4.301436 | 1.70E-05 | 0.001370317 |
| *DOK2* | -1.648405 | 0.383593 | -4.297273 | 1.73E-05 | 0.001376875 |
| *RAB24* | -0.743928 | 0.173117 | -4.297256 | 1.73E-05 | 0.001376875 |
| *LY86* | -0.406567 | 0.094985 | -4.280315 | 1.87E-05 | 0.001473152 |
| *ENSG00000279483* | -0.390649 | 0.091291 | -4.279154 | 1.88E-05 | 0.001473152 |
| *GBGT1* | 1.213508 | 0.28372 | 4.277137 | 1.89E-05 | 0.001476377 |
| *ENSG00000227468* | -2.268529 | 0.530628 | -4.275174 | 1.91E-05 | 0.001479314 |
| *CD72* | 0.772487 | 0.180785 | 4.272959 | 1.93E-05 | 0.001483994 |
| *HLA.C* | -0.489519 | 0.114809 | -4.263762 | 2.01E-05 | 0.001536067 |
| *GPR183* | 0.380819 | 0.089458 | 4.256963 | 2.07E-05 | 0.001572961 |
| *SPART* | 0.755645 | 0.178577 | 4.231489 | 2.32E-05 | 0.001750499 |
| *JCHAIN* | -0.993386 | 0.23489 | -4.229163 | 2.35E-05 | 0.001757054 |
| *ENSG00000254397* | 1.959934 | 0.465181 | 4.213275 | 2.52E-05 | 0.001870693 |
| *RUNX1* | 1.255564 | 0.298125 | 4.211536 | 2.54E-05 | 0.001870693 |
| *RAC2* | -0.386493 | 0.091803 | -4.210016 | 2.55E-05 | 0.001870693 |
| *NME1* | -0.640687 | 0.152212 | -4.209175 | 2.56E-05 | 0.001870693 |
| *LTA* | -1.038379 | 0.2468 | -4.207372 | 2.58E-05 | 0.00187367 |
| *SLC2A1* | -0.610823 | 0.145305 | -4.203714 | 2.63E-05 | 0.001892167 |
| *MME* | -2.93398 | 0.701418 | -4.182924 | 2.88E-05 | NA |
| *CCDC167* | -0.587973 | 0.140669 | -4.179833 | 2.92E-05 | 0.002089029 |
| *CTSB* | 0.483123 | 0.116286 | 4.154622 | 3.26E-05 | 0.002318657 |
| *PCNX4* | 0.505502 | 0.121714 | 4.153187 | 3.28E-05 | 0.002318758 |
| *MGAT4A* | -0.653717 | 0.157736 | -4.144375 | 3.41E-05 | 0.002394872 |
| *IL15* | -0.730688 | 0.176544 | -4.138835 | 3.49E-05 | 0.002438377 |
| *BTG2* | 0.320984 | 0.077654 | 4.133534 | 3.57E-05 | 0.002480112 |
| *ATAD3C* | -1.492718 | 0.362228 | -4.120936 | 3.77E-05 | 0.00260193 |
| *INPP5K* | -0.584151 | 0.141794 | -4.119714 | 3.79E-05 | 0.00260193 |
| *MTERF4* | -0.397817 | 0.096646 | -4.116223 | 3.85E-05 | 0.002625812 |
| *CEMIP2* | 1.335617 | 0.325398 | 4.104567 | 4.05E-05 | 0.002740642 |
| *PRKCE* | 0.806417 | 0.196515 | 4.10359 | 4.07E-05 | 0.002740642 |
| *MSMO1* | -0.449146 | 0.109586 | -4.098581 | 4.16E-05 | 0.002771349 |
| *TCAF2* | -0.913114 | 0.222804 | -4.09829 | 4.16E-05 | 0.002771349 |
| *PSME2* | -0.39363 | 0.096446 | -4.081339 | 4.48E-05 | 0.00296413 |
| *SNED1* | -1.73388 | 0.425159 | -4.078188 | 4.54E-05 | 0.002972762 |
| *ENSG00000254614* | -1.274299 | 0.312483 | -4.077976 | 4.54E-05 | 0.002972762 |
| *BAIAP3* | -1.864171 | 0.457757 | -4.072402 | 4.65E-05 | 0.003018214 |
| *AKT3* | 0.822682 | 0.202045 | 4.071785 | 4.67E-05 | 0.003018214 |
| *CLEC2B* | 0.66769 | 0.164375 | 4.061981 | 4.87E-05 | 0.003130062 |
| *LCK* | -1.067728 | 0.263622 | -4.050217 | 5.12E-05 | 0.003273159 |
| *TRIM4* | 0.574367 | 0.14199 | 4.04512 | 5.23E-05 | 0.003326516 |
| *ALDH3A2* | 1.552086 | 0.383882 | 4.043132 | 5.27E-05 | 0.003336222 |
| *TRAF1* | -0.848377 | 0.210254 | -4.035001 | 5.46E-05 | 0.003434797 |
| *IFNG.AS1* | -1.743474 | 0.433316 | -4.023564 | 5.73E-05 | 0.00358622 |
| *RASGEF1B* | 0.646113 | 0.160652 | 4.021824 | 5.77E-05 | 0.003593069 |
| *TPI1* | -0.195976 | 0.048749 | -4.020068 | 5.82E-05 | 0.003600281 |
| *MRPL20* | -0.354438 | 0.088215 | -4.017885 | 5.87E-05 | 0.003614145 |
| *NDUFA1* | -0.214758 | 0.053543 | -4.010911 | 6.05E-05 | 0.003702589 |

| | | | | | |
|---|---|---|---|---|---|
| *DHCR24* | -1.035768 | 0.25853 | -4.00637 | 6.17E-05 | 0.003753542 |
| *BEX3* | -1.603684 | 0.400405 | -4.005156 | 6.20E-05 | 0.003753542 |
| *VCPKMT* | -0.742482 | 0.186039 | -3.991003 | 6.58E-05 | 0.003963681 |
| *UNC119* | -0.502939 | 0.126081 | -3.989002 | 6.64E-05 | 0.003976228 |
| *ARPC1B* | -0.308308 | 0.077318 | -3.987531 | 6.68E-05 | 0.003980005 |
| *ARID5A* | -0.532797 | 0.133995 | -3.976259 | 7.00E-05 | 0.004151605 |
| *PARM1* | -1.706577 | 0.429904 | -3.969666 | 7.20E-05 | 0.004246057 |
| *PPIA* | -0.204724 | 0.051666 | -3.962419 | 7.42E-05 | 0.004343208 |
| *LYRM2* | -0.323613 | 0.081683 | -3.961812 | 7.44E-05 | 0.004343208 |
| *SMAGP* | -0.547338 | 0.138442 | -3.953564 | 7.70E-05 | 0.004459315 |
| *LILRB1* | 0.722806 | 0.182847 | 3.95307 | 7.72E-05 | 0.004459315 |
| *ADAMTS6* | 1.113715 | 0.282015 | 3.949132 | 7.84E-05 | 0.004489947 |
| *PLEKHB1* | -1.113053 | 0.281856 | -3.949015 | 7.85E-05 | 0.004489947 |
| *RIN3* | 1.063832 | 0.270059 | 3.939262 | 8.17E-05 | 0.004653036 |
| *KYNU* | -0.523665 | 0.133077 | -3.935041 | 8.32E-05 | 0.004712024 |
| *DGKG* | 2.023743 | 0.514488 | 3.933511 | 8.37E-05 | 0.004718652 |
| *HSD17B11* | 0.323231 | 0.08231 | 3.926971 | 8.60E-05 | 0.004814067 |
| *UQCRC1* | -0.393809 | 0.1003 | -3.926328 | 8.63E-05 | 0.004814067 |
| *CLDND1* | -0.467855 | 0.11923 | -3.923965 | 8.71E-05 | 0.004824222 |
| *CCDC88C* | -0.461745 | 0.117688 | -3.923468 | 8.73E-05 | 0.004824222 |
| *TRAF4* | -0.759875 | 0.193732 | -3.922306 | 8.77E-05 | 0.004824222 |
| *CKS2* | 0.610199 | 0.156108 | 3.908817 | 9.27E-05 | 0.005077127 |
| *BMPR2* | 0.872665 | 0.223853 | 3.898387 | 9.68E-05 | 0.005275452 |
| *ARHGAP24* | 0.496399 | 0.127716 | 3.886756 | 0.00010159 | 0.005508253 |
| *PSMB3* | -0.337931 | 0.087067 | -3.881275 | 0.00010391 | 0.005607212 |
| *GSTK1* | -0.344393 | 0.088799 | -3.878332 | 0.00010518 | 0.005643686 |
| *ENSG00000273748* | -1.197174 | 0.308757 | -3.877403 | 0.00010558 | 0.005643686 |
| *ATP5MG* | -0.184278 | 0.047551 | -3.8754 | 0.00010645 | 0.00566372 |
| *ENSG00000180953* | -0.570995 | 0.14754 | -3.87011 | 0.00010879 | 0.005761126 |
| *ENSG00000272211* | -0.989803 | 0.25609 | -3.865064 | 0.00011106 | 0.005854296 |
| *RAB30* | 0.38834 | 0.100516 | 3.863478 | 0.00011178 | 0.005865311 |
| *HTR3A* | -0.948673 | 0.245704 | -3.861043 | 0.0001129 | 0.005896911 |
| *SH2B3* | 0.858501 | 0.222596 | 3.856773 | 0.00011489 | 0.005973423 |
| *JSRP1* | -1.187316 | 0.308355 | -3.850485 | 0.00011788 | 0.00610103 |
| *KRCC1* | 0.369182 | 0.095964 | 3.84709 | 0.00011953 | 0.006158193 |
| *CHCHD5* | -0.421246 | 0.109591 | -3.84381 | 0.00012114 | 0.006210258 |
| *TIAM2* | -0.896584 | 0.233357 | -3.842115 | 0.00012198 | 0.006210258 |
| *TSPAN5* | -1.020831 | 0.265723 | -3.841718 | 0.00012218 | 0.006210258 |
| *SEPTIN7* | 0.33402 | 0.086994 | 3.839595 | 0.00012324 | 0.006236354 |
| *ENSG00000237773* | -1.286031 | 0.335887 | -3.828766 | 0.00012879 | 0.00648838 |
| *FARP2* | 0.943484 | 0.246557 | 3.826635 | 0.00012991 | 0.006515959 |
| *ARL17A* | -0.627796 | 0.164263 | -3.821903 | 0.00013243 | 0.006613151 |
| *NCBP3* | 0.993391 | 0.260259 | 3.816934 | 0.00013512 | 0.006718258 |
| *H2AFZ* | 0.364179 | 0.09549 | 3.813814 | 0.00013684 | 0.006774108 |
| *PSME1* | -0.203441 | 0.053565 | -3.79804 | 0.00014585 | 0.007188703 |
| *FCGR2B* | 0.580801 | 0.153105 | 3.793481 | 0.00014855 | 0.007290466 |
| *TUBB6* | 1.021008 | 0.269288 | 3.791505 | 0.00014974 | 0.00731721 |

| | | | | | |
|---|---|---|---|---|---|
| *SESN3* | 0.409815 | 0.108323 | 3.783275 | 0.00015478 | 0.007531211 |
| *C9orf139* | -0.773585 | 0.204571 | -3.781496 | 0.00015589 | 0.007552987 |
| *IFNLR1* | 1.012013 | 0.268006 | 3.776081 | 0.00015932 | 0.007686278 |
| *POU2AF1* | -0.429136 | 0.114032 | -3.763289 | 0.00016769 | 0.00805634 |
| *CD80* | -1.253557 | 0.333409 | -3.759814 | 0.00017004 | 0.008134645 |
| *NUPL2* | -0.451331 | 0.120135 | -3.756849 | 0.00017207 | 0.008134645 |
| *UROS* | -0.448588 | 0.119409 | -3.756742 | 0.00017214 | 0.008134645 |
| *THAP2* | 0.706335 | 0.188021 | 3.756683 | 0.00017218 | 0.008134645 |
| *LDLR* | -1.468282 | 0.390974 | -3.75545 | 0.00017303 | 0.008140992 |
| *RBM3* | -0.418467 | 0.111536 | -3.751854 | 0.00017553 | 0.008221095 |
| *HLA.A* | -0.401759 | 0.107109 | -3.750935 | 0.00017618 | 0.008221095 |
| *AKAP9* | 0.293421 | 0.078263 | 3.749181 | 0.00017741 | 0.008244991 |
| *HLA.DOB* | -0.56457 | 0.150696 | -3.746413 | 0.00017938 | 0.008302565 |
| *DUSP4* | 1.530337 | 0.409203 | 3.739796 | 0.00018417 | 0.008489702 |
| *TAPBP* | -0.322112 | 0.086181 | -3.737642 | 0.00018575 | 0.008528211 |
| *ZNF593* | -0.469603 | 0.126333 | -3.717194 | 0.00020145 | 0.009198088 |
| *CDC42SE1* | -0.34082 | 0.091703 | -3.716552 | 0.00020196 | 0.009198088 |
| *MYO1G* | -0.466258 | 0.125596 | -3.712353 | 0.00020534 | 0.009313513 |
| *PITPNC1* | -0.77814 | 0.209663 | -3.711382 | 0.00020613 | 0.009313513 |
| *C17orf49* | -0.734792 | 0.198045 | -3.71022 | 0.00020708 | 0.009319386 |
| *RFX5* | -0.386112 | 0.104284 | -3.702504 | 0.00021348 | 0.009569697 |
| *AP1S2* | -0.325022 | 0.087819 | -3.701029 | 0.00021473 | 0.009587785 |
| *FASTK* | -0.404668 | 0.109532 | -3.694528 | 0.0002203 | 0.009776019 |
| *PROCA1* | 0.940262 | 0.254563 | 3.69363 | 0.00022108 | 0.009776019 |
| *CDC42EP3* | -0.401857 | 0.108812 | -3.693121 | 0.00022152 | 0.009776019 |
| *ENSA* | -0.241307 | 0.065431 | -3.687958 | 0.00022606 | 0.009904687 |
| *TBX21* | 1.705429 | 0.462448 | 3.687831 | 0.00022617 | 0.009904687 |
| *PSAP* | 0.453544 | 0.123301 | 3.678359 | 0.00023474 | 0.010240385 |
| *GTF3C6* | -0.336782 | 0.091611 | -3.676223 | 0.00023671 | 0.01028708 |
| *SRGN* | 0.835118 | 0.227646 | 3.668487 | 0.00024399 | 0.010563022 |
| *IZUMO4* | -0.64371 | 0.175542 | -3.666976 | 0.00024544 | 0.010585359 |
| *SMIM4* | -0.504616 | 0.137788 | -3.662253 | 0.00025001 | 0.010741791 |
| *HLA.DQB2* | -2.740139 | 0.749243 | -3.65721 | 0.00025498 | 0.010914091 |
| *JAZF1* | 0.526904 | 0.144158 | 3.655048 | 0.00025713 | 0.010922764 |
| *PNPLA8* | 0.388121 | 0.106191 | 3.654954 | 0.00025723 | 0.010922764 |
| *PPP1R15A* | -0.548325 | 0.150089 | -3.653339 | 0.00025885 | 0.010922764 |
| *IGHEP1* | -1.803207 | 0.493871 | -3.651173 | 0.00026105 | 0.010922764 |
| *FDFT1* | -0.409269 | 0.112099 | -3.650959 | 0.00026126 | 0.010922764 |
| *S1PR2* | -0.933203 | 0.255645 | -3.650389 | 0.00026184 | 0.010922764 |
| *HSBP1* | 0.303662 | 0.083187 | 3.65034 | 0.00026189 | 0.010922764 |
| *HSD17B10* | -0.394918 | 0.108263 | -3.647765 | 0.00026453 | 0.010992533 |
| *SNX11* | -0.54135 | 0.149281 | -3.626387 | 0.00028741 | 0.011899993 |
| *UBC* | -0.363209 | 0.100212 | -3.624415 | 0.00028962 | 0.011947729 |
| *XYLT1* | 0.904889 | 0.249954 | 3.620221 | 0.00029435 | 0.012099217 |
| *NFKB2* | -0.622232 | 0.17235 | -3.610292 | 0.00030585 | 0.012442281 |
| *PPIB* | -0.228191 | 0.063206 | -3.610241 | 0.00030591 | 0.012442281 |
| *IER2* | -0.324222 | 0.089817 | -3.609817 | 0.00030641 | 0.012442281 |

| | | | | | |
|---|---|---|---|---|---|
| *SLA* | 0.954793 | 0.26454 | 3.609263 | 0.00030707 | 0.012442281 |
| *SLC25A29* | 1.465537 | 0.406432 | 3.605863 | 0.00031112 | 0.012561622 |
| *ZBTB38* | -0.465897 | 0.129257 | -3.604426 | 0.00031284 | 0.0125867 |
| *GPR137* | -0.461154 | 0.128109 | -3.599705 | 0.00031858 | 0.012772318 |
| *PLPP5* | -0.521692 | 0.145216 | -3.592525 | 0.00032749 | 0.013083462 |
| *BBC3* | -0.900152 | 0.250709 | -3.590419 | 0.00033015 | 0.013143361 |
| *FRG1* | -0.340532 | 0.094868 | -3.589513 | 0.0003313 | 0.013143361 |
| *POU2F2* | 0.419557 | 0.116931 | 3.588068 | 0.00033314 | 0.01317051 |
| *CDCA7* | -1.062675 | 0.296639 | -3.582386 | 0.00034047 | 0.013413833 |
| *GMFG* | -0.276363 | 0.07726 | -3.577036 | 0.00034751 | 0.013644073 |
| *S100A11* | 0.27984 | 0.078346 | 3.571842 | 0.00035448 | 0.013869743 |
| *MKNK2* | -0.367468 | 0.102974 | -3.568537 | 0.00035898 | 0.013997784 |
| *NOP10* | 0.191403 | 0.053745 | 3.561286 | 0.00036904 | 0.014341053 |
| *CYBC1* | -0.282816 | 0.079436 | -3.560304 | 0.00037043 | 0.014345789 |
| *SPIB* | -0.312259 | 0.087733 | -3.559177 | 0.00037202 | 0.014348554 |
| *MRPL32* | 0.373592 | 0.104987 | 3.558473 | 0.00037302 | 0.014348554 |
| *GDAP1* | -1.097204 | 0.308418 | -3.557515 | 0.00037438 | 0.01435246 |
| *JUNB* | -0.328895 | 0.092565 | -3.55311 | 0.00038071 | 0.014546006 |
| *PRAG1* | 1.039112 | 0.293124 | 3.544956 | 0.00039268 | 0.014953326 |
| *DBI* | -0.272757 | 0.076967 | -3.543821 | 0.00039437 | 0.014967743 |
| *ARHGDIA* | -0.261219 | 0.073755 | -3.541689 | 0.00039757 | 0.015039133 |
| *XRN2* | -0.324589 | 0.091984 | -3.528739 | 0.00041755 | 0.015742283 |
| *SP100* | -0.210048 | 0.059607 | -3.523883 | 0.00042527 | 0.015980657 |
| *SNX9* | 0.504959 | 0.143775 | 3.512144 | 0.00044451 | 0.016648566 |
| *COPS3* | -0.438486 | 0.125042 | -3.506716 | 0.00045367 | 0.016936113 |
| *ETFDH* | -0.686295 | 0.195832 | -3.504511 | 0.00045745 | 0.017021221 |
| *ZNF92* | -0.418735 | 0.119718 | -3.497688 | 0.00046931 | 0.017405724 |
| *DHRS4L2* | 0.469958 | 0.134427 | 3.496011 | 0.00047227 | 0.017405841 |
| *SMC6* | 0.406123 | 0.116189 | 3.495363 | 0.00047342 | 0.017405841 |
| *ARHGAP11A* | -1.324187 | 0.378948 | -3.494373 | 0.00047518 | 0.017405841 |
| *SF1* | -0.201141 | 0.057564 | -3.494232 | 0.00047543 | 0.017405841 |
| *CSGALNACT2* | -0.592531 | 0.169657 | -3.492518 | 0.00047849 | 0.017461779 |
| *SRGAP2B* | 0.800993 | 0.229587 | 3.488849 | 0.00048511 | 0.017646671 |
| *TMEM131* | 0.548611 | 0.157307 | 3.487515 | 0.00048753 | 0.017678473 |
| *RSBN1L* | 0.294878 | 0.084577 | 3.486501 | 0.00048938 | 0.017689286 |
| *LPP* | 0.516929 | 0.148315 | 3.485352 | 0.00049149 | 0.017709217 |
| *CYB5A* | -0.510126 | 0.14665 | -3.478517 | 0.0005042 | 0.018097196 |
| *ANXA6* | -0.455104 | 0.130858 | -3.477858 | 0.00050544 | 0.018097196 |
| *ELOB* | -0.164296 | 0.047302 | -3.473362 | 0.00051398 | 0.018299125 |
| *PYHIN1* | 0.91661 | 0.263909 | 3.4732 | 0.00051429 | 0.018299125 |
| *OSBPL8* | 0.359701 | 0.10363 | 3.471009 | 0.00051851 | 0.018309299 |
| *GBP2* | 0.567638 | 0.163539 | 3.470958 | 0.00051861 | 0.018309299 |
| *MRPL23* | -0.276036 | 0.079537 | -3.470546 | 0.0005194 | 0.018309299 |
| *EPG5* | 0.897097 | 0.258633 | 3.468615 | 0.00052315 | 0.018368884 |
| *VPREB3* | -0.377534 | 0.108883 | -3.467324 | 0.00052567 | 0.018368884 |
| *CBLB* | 0.471282 | 0.135933 | 3.467003 | 0.0005263 | 0.018368884 |
| *RNH1* | -0.458489 | 0.132268 | -3.466366 | 0.00052755 | 0.018368884 |

| | | | | | |
|---|---|---|---|---|---|
| *SLCO5A1* | -1.501682 | 0.433405 | -3.464847 | 0.00053053 | 0.018416594 |
| *AKAP2* | -0.844598 | 0.24445 | -3.455089 | 0.00055011 | 0.019001571 |
| *TBC1D9* | 0.437045 | 0.126504 | 3.45479 | 0.00055072 | 0.019001571 |
| *UBE2J2* | -0.3264 | 0.094519 | -3.453254 | 0.00055387 | 0.019052363 |
| *ETF1* | -0.519935 | 0.150822 | -3.44735 | 0.00056611 | 0.019405843 |
| *TMSB10* | -0.33219 | 0.09638 | -3.446665 | 0.00056755 | 0.019405843 |
| *MLST8* | -0.456734 | 0.13257 | -3.445235 | 0.00057056 | 0.019450406 |
| *SIPA1L1* | 0.391054 | 0.113936 | 3.432216 | 0.00059867 | 0.020347616 |
| *ICAM1* | -0.964375 | 0.281056 | -3.431252 | 0.0006008 | 0.020359328 |
| *B4GALT1* | 0.448101 | 0.130639 | 3.430075 | 0.00060342 | 0.020387202 |
| *TOMM5* | -0.341442 | 0.099658 | -3.426135 | 0.00061224 | 0.020579794 |
| *CHORDC1* | 0.519388 | 0.151614 | 3.425732 | 0.00061314 | 0.020579794 |
| *PLGRKT* | -0.606904 | 0.177192 | -3.425116 | 0.00061454 | 0.020579794 |
| *ENSG00000265218* | -1.892601 | 0.553822 | -3.417347 | 0.00063235 | 0.02111406 |
| *IKBKE* | -0.678227 | 0.198657 | -3.414055 | 0.00064004 | 0.021308339 |
| *LRP5* | -1.142803 | 0.335676 | -3.404488 | 0.00066288 | 0.022004607 |
| *ANXA2* | -0.440372 | 0.129495 | -3.400687 | 0.00067217 | 0.022114415 |
| *ENSG00000263264* | -0.92534 | 0.272148 | -3.400131 | 0.00067354 | 0.022114415 |
| *ST13* | 0.204294 | 0.060085 | 3.400093 | 0.00067363 | 0.022114415 |
| *PSMA4* | -0.256203 | 0.075355 | -3.399959 | 0.00067396 | 0.022114415 |
| *SLC9A3R1* | -0.49888 | 0.146787 | -3.398658 | 0.00067717 | 0.022156028 |
| *ZNF107* | 0.89779 | 0.264276 | 3.397167 | 0.00068087 | 0.022213246 |
| *COX7B* | -0.214087 | 0.06319 | -3.387977 | 0.0007041 | 0.022905404 |
| *GSAP* | 0.528 | 0.155893 | 3.38694 | 0.00070677 | 0.022926695 |
| *ECH1* | -0.321083 | 0.09494 | -3.381965 | 0.00071969 | 0.023238509 |
| *NDUFS6* | -0.269139 | 0.079588 | -3.381671 | 0.00072046 | 0.023238509 |
| *ENSG00000225885* | -0.765194 | 0.226331 | -3.380859 | 0.0007226 | 0.023241437 |
| *CRK* | 0.455281 | 0.134755 | 3.378579 | 0.00072862 | 0.023369039 |
| *PHF23* | 0.475035 | 0.140706 | 3.376088 | 0.00073524 | 0.023506135 |
| *ZNF667.AS1* | 1.160232 | 0.343729 | 3.375425 | 0.00073702 | 0.023506135 |
| *CCND3* | 0.433057 | 0.128696 | 3.36496 | 0.00076555 | 0.024347873 |
| *PIK3R5* | -0.650399 | 0.193354 | -3.363778 | 0.00076883 | 0.024384231 |
| *ARL3* | 0.52282 | 0.155575 | 3.360561 | 0.00077784 | 0.024601464 |
| *LNPEP* | 0.399213 | 0.119061 | 3.353023 | 0.00079934 | 0.025211341 |
| *IL2RG* | -0.378966 | 0.113136 | -3.349656 | 0.00080912 | 0.02537616 |
| *BEX2* | -0.53007 | 0.158265 | -3.349254 | 0.0008103 | 0.02537616 |
| *ENSG00000225205* | -1.188039 | 0.354752 | -3.348927 | 0.00081125 | 0.02537616 |
| *IQGAP2* | 1.244571 | 0.371821 | 3.347233 | 0.00081623 | 0.025461803 |
| *MAP1LC3A* | -0.902166 | 0.269606 | -3.346239 | 0.00081916 | 0.025466083 |
| *SLC2A14* | -4.275329 | 1.277715 | -3.346074 | 0.00081964 | NA |
| *BMS1* | -0.473971 | 0.141667 | -3.345671 | 0.00082084 | 0.025466083 |
| *PHF20L1* | 0.426236 | 0.127438 | 3.344644 | 0.00082388 | 0.02549109 |
| *ZMIZ2* | -0.487947 | 0.145943 | -3.343412 | 0.00082755 | 0.025535163 |
| *E2F5* | 0.522862 | 0.156691 | 3.336895 | 0.0008472 | 0.02605688 |
| *VPREB1* | -1.733113 | 0.519473 | -3.336294 | 0.00084903 | 0.02605688 |
| *RNF144B* | 1.166389 | 0.350067 | 3.331899 | 0.00086256 | 0.026400685 |
| *GSN* | -0.661515 | 0.198725 | -3.328791 | 0.00087224 | 0.026625509 |

| | | | | | |
|---|---|---|---|---|---|
| *COX7A2* | -0.195227 | 0.058689 | -3.326473 | 0.00087953 | 0.026776121 |
| *ARHGDIB* | -0.256681 | 0.077208 | -3.32453 | 0.00088568 | 0.026891548 |
| *TNFAIP8L2* | 0.791866 | 0.238307 | 3.322884 | 0.00089092 | 0.026978754 |
| *GPX1* | -0.309029 | 0.093034 | -3.321661 | 0.00089483 | 0.027017656 |
| *MGME1* | 0.493475 | 0.148592 | 3.321002 | 0.00089695 | 0.027017656 |
| *CD247* | -1.377388 | 0.414854 | -3.320175 | 0.00089961 | 0.027026334 |
| *LARP4B* | 0.684876 | 0.206651 | 3.314175 | 0.00091914 | 0.027540365 |
| *LINC01215* | -0.614931 | 0.185652 | -3.312271 | 0.00092542 | 0.0275413 |
| *LILRA4* | 2.046829 | 0.618135 | 3.311297 | 0.00092865 | 0.0275413 |
| *ACCS* | 1.931609 | 0.583368 | 3.311133 | 0.00092919 | 0.0275413 |
| *SPN* | -1.28746 | 0.388879 | -3.310696 | 0.00093064 | 0.0275413 |
| *PILRB* | 0.763414 | 0.230603 | 3.310508 | 0.00093127 | 0.0275413 |
| *MCRIP2* | -0.488978 | 0.147761 | -3.309236 | 0.00093551 | 0.027595114 |
| *CALM1* | -0.228988 | 0.069218 | -3.308229 | 0.00093888 | 0.027622951 |
| *BRI3BP* | 0.750604 | 0.227153 | 3.304398 | 0.00095181 | 0.0279311 |
| *APPBP2* | 0.625292 | 0.189389 | 3.301626 | 0.00096126 | 0.028136033 |
| *ACSF3* | -0.476407 | 0.144358 | -3.300163 | 0.00096629 | 0.028210572 |
| *IMPDH2* | -0.473379 | 0.143541 | -3.297864 | 0.00097423 | 0.028369891 |
| *CDC37* | -0.242775 | 0.073733 | -3.292608 | 0.00099263 | 0.028831751 |
| *NCF2* | -0.903088 | 0.274462 | -3.290394 | 0.00100047 | 0.02891928 |
| *NEK6* | -0.566778 | 0.172256 | -3.290324 | 0.00100072 | 0.02891928 |
| *TRMT1L* | 0.689834 | 0.209749 | 3.288849 | 0.00100598 | 0.028941489 |
| *RAB3GAP2* | -0.426221 | 0.129602 | -3.288683 | 0.00100657 | 0.028941489 |
| *TOP1* | 0.350661 | 0.106731 | 3.285459 | 0.00101816 | 0.029200984 |
| *SLC11A1* | 1.645782 | 0.501606 | 3.281025 | 0.00103431 | 0.029537407 |
| *HMGB3* | -1.157207 | 0.352744 | -3.280587 | 0.00103591 | 0.029537407 |
| *TRAK1* | 0.540801 | 0.164873 | 3.280108 | 0.00103768 | 0.029537407 |
| *PECAM1* | 0.95561 | 0.2914 | 3.279372 | 0.00104039 | 0.029540693 |
| *CYTH1* | -0.326908 | 0.099812 | -3.27525 | 0.00105569 | 0.029900601 |
| *LPAR5* | 0.474235 | 0.145007 | 3.270422 | 0.00107387 | 0.030279412 |
| *PPBP* | 3.518736 | 1.076368 | 3.269082 | 0.00107897 | 0.030279412 |
| *UBE2F* | 0.437476 | 0.133823 | 3.26906 | 0.00107905 | 0.030279412 |
| *TNRC6B* | 0.228769 | 0.069984 | 3.268891 | 0.0010797 | 0.030279412 |
| *COX6C* | -0.20603 | 0.063053 | -3.267562 | 0.00108478 | 0.030347193 |
| *VAV2* | -0.472982 | 0.144785 | -3.266786 | 0.00108776 | 0.030355946 |
| *ABI2* | 0.503394 | 0.154252 | 3.263447 | 0.00110066 | 0.030640813 |
| *TIGIT* | -1.440586 | 0.441662 | -3.261742 | 0.0011073 | 0.030715445 |
| *TTC39C* | -0.544721 | 0.167022 | -3.261375 | 0.00110873 | 0.030715445 |
| *ENSG00000228037* | -0.86523 | 0.265862 | -3.254433 | 0.00113619 | 0.031376644 |
| *ERG28* | -0.334099 | 0.102675 | -3.253953 | 0.00113811 | 0.031376644 |
| *ENSG00000266088* | -2.765873 | 0.850836 | -3.25077 | 0.00115093 | 0.031653298 |
| *HIPK2* | -0.69548 | 0.214074 | -3.248789 | 0.00115897 | 0.031797775 |
| *SEPTIN9* | -0.206281 | 0.063552 | -3.245875 | 0.0011709 | 0.032047849 |
| *VARS* | -0.7707 | 0.237528 | -3.244665 | 0.00117589 | 0.032107118 |
| *DICER1* | 0.47189 | 0.145505 | 3.243119 | 0.00118229 | 0.032204678 |
| *GCC2* | 0.263941 | 0.081584 | 3.235187 | 0.00121563 | 0.033033846 |
| *HECA* | 0.52866 | 0.163499 | 3.233423 | 0.00122316 | 0.03315938 |

| | | | | | |
|---|---|---|---|---|---|
| *NUDT3* | 0.410734 | 0.127065 | 3.232463 | 0.00122728 | 0.033191945 |
| *ARPC3* | -0.223343 | 0.069117 | -3.23137 | 0.00123199 | 0.03324025 |
| *GFOD1* | 0.642739 | 0.199025 | 3.229435 | 0.00124035 | 0.033386835 |
| *HSPB1* | 0.644217 | 0.199795 | 3.224397 | 0.00126238 | 0.03389974 |
| *MIAT* | -0.946602 | 0.293811 | -3.221806 | 0.00127385 | 0.034127236 |
| *C1GALT1* | 0.463656 | 0.144112 | 3.217327 | 0.00129391 | 0.03458322 |
| *CSF2RB* | -0.779514 | 0.242349 | -3.216498 | 0.00129765 | 0.034602045 |
| *ENO1* | -0.268627 | 0.08367 | -3.210559 | 0.00132477 | 0.035159365 |
| *AFF3* | 0.35331 | 0.110053 | 3.210356 | 0.00132571 | 0.035159365 |
| *ENSG00000256092* | -0.725202 | 0.225927 | -3.209899 | 0.00132782 | 0.035159365 |
| *BHLHE41* | 0.876037 | 0.273015 | 3.208754 | 0.00133312 | 0.035217772 |
| *TERF2IP* | 0.289011 | 0.090155 | 3.205706 | 0.00134732 | 0.035510489 |
| *ZSCAN2* | -0.822131 | 0.256933 | -3.199782 | 0.00137532 | 0.036164762 |
| *ZNF397* | 0.367951 | 0.115085 | 3.197216 | 0.00138761 | 0.036403988 |
| *KRTCAP2* | -0.25686 | 0.080371 | -3.195915 | 0.00139388 | 0.036484404 |
| *GGCX* | -0.552087 | 0.172815 | -3.194668 | 0.00139992 | 0.036558373 |
| *APOBEC3H* | -1.612023 | 0.504789 | -3.193459 | 0.00140579 | 0.036572935 |
| *TIMM22* | -0.423203 | 0.132535 | -3.193136 | 0.00140737 | 0.036572935 |
| *ODF3B* | -0.577629 | 0.180929 | -3.192574 | 0.00141011 | 0.036572935 |
| *ENSG00000089127* | 0.815878 | 0.255743 | 3.190222 | 0.00142164 | 0.036788092 |
| *ZNF568* | 0.632978 | 0.19859 | 3.187368 | 0.00143574 | 0.037068839 |
| *TCF7* | -0.731877 | 0.229794 | -3.184932 | 0.00144788 | 0.037238047 |
| *BTNL9* | -1.368689 | 0.429765 | -3.184741 | 0.00144884 | 0.037238047 |
| *ZBTB24* | -0.399384 | 0.125471 | -3.183092 | 0.00145711 | 0.037366376 |
| *GRAMD1C* | -0.730446 | 0.229819 | -3.178347 | 0.00148117 | 0.037842992 |
| *ATP5MC3* | -0.215307 | 0.06775 | -3.177934 | 0.00148329 | 0.037842992 |
| *RHOH* | 0.270367 | 0.085089 | 3.177469 | 0.00148567 | 0.037842992 |
| *GALM* | 1.009233 | 0.317747 | 3.176216 | 0.0014921 | 0.037903715 |
| *ATP5PD* | -0.204509 | 0.064398 | -3.175709 | 0.00149471 | 0.037903715 |
| *BTK* | 0.325831 | 0.102645 | 3.174334 | 0.00150181 | 0.037999086 |
| *S100PBP* | 0.554882 | 0.17516 | 3.167861 | 0.00153565 | 0.038769207 |
| *RASSF6* | -1.35162 | 0.426887 | -3.166223 | 0.00154432 | 0.038901936 |
| *FBXO21* | -0.364587 | 0.115284 | -3.162506 | 0.00156418 | 0.039315004 |
| *HDAC1* | -0.335387 | 0.106125 | -3.160301 | 0.00157606 | 0.039357016 |
| *EIF4E3* | -1.778117 | 0.56269 | -3.160026 | 0.00157755 | 0.039357016 |
| *PEX6* | 0.983579 | 0.311264 | 3.159947 | 0.00157798 | 0.039357016 |
| *TMEM138* | -0.359424 | 0.113755 | -3.159634 | 0.00157967 | 0.039357016 |
| *ADA* | -0.461264 | 0.146165 | -3.15577 | 0.00160075 | 0.039795011 |
| *FCRL2* | 0.629622 | 0.199653 | 3.15359 | 0.00161276 | 0.040006199 |
| *CCDC137* | 0.438063 | 0.139016 | 3.151165 | 0.00162621 | 0.040219553 |
| *NOP14.AS1* | -0.847462 | 0.26897 | -3.150767 | 0.00162842 | 0.040219553 |
| *MINDY2* | 0.496228 | 0.15754 | 3.149849 | 0.00163355 | 0.040258786 |
| *ENSG00000282988* | 1.153982 | 0.366578 | 3.147989 | 0.00164398 | 0.040422313 |
| *PSMD12* | -0.372519 | 0.118358 | -3.147402 | 0.00164728 | 0.040422313 |
| *ABI1* | 0.34905 | 0.110982 | 3.145097 | 0.00166032 | 0.040575571 |
| *ITGB3BP* | -0.474249 | 0.150822 | -3.144421 | 0.00166416 | 0.040575571 |
| *KLHL8* | 0.855985 | 0.272224 | 3.14441 | 0.00166422 | 0.040575571 |

| | | | | | |
|---|---|---|---|---|---|
| *PPCS* | -0.281687 | 0.089653 | -3.141954 | 0.00167824 | 0.040830061 |
| *SPTLC2* | 0.58733 | 0.187033 | 3.14024 | 0.00168809 | 0.040904716 |
| *TLE3* | -0.332787 | 0.105977 | -3.14017 | 0.0016885 | 0.040904716 |
| *IGLC2* | -0.591701 | 0.188616 | -3.137058 | 0.00170652 | 0.041253614 |
| *ATP5MGL* | -0.727537 | 0.232167 | -3.133681 | 0.00172629 | 0.041642982 |
| *KIAA0040* | 0.520626 | 0.166203 | 3.132468 | 0.00173343 | 0.04172702 |
| *DHRS4* | -0.75256 | 0.24034 | -3.131239 | 0.00174071 | 0.041740335 |
| *TRIP10* | 0.845749 | 0.270109 | 3.131135 | 0.00174132 | 0.041740335 |
| *ENSG00000267892* | 1.375582 | 0.43971 | 3.128385 | 0.0017577 | 0.042044429 |
| *ATP1B3* | -0.338561 | 0.108375 | -3.123973 | 0.00178427 | 0.042590497 |
| *NDUFA6* | -0.289766 | 0.092857 | -3.120546 | 0.00180516 | 0.042976725 |
| *GPR18* | 0.731535 | 0.234473 | 3.119907 | 0.00180908 | 0.042976725 |
| *GRK6* | -0.571606 | 0.183254 | -3.119203 | 0.00181341 | 0.042976725 |
| *PSMB1* | -0.219841 | 0.070498 | -3.118381 | 0.00181848 | 0.042976725 |
| *INSIG1* | -0.299885 | 0.096171 | -3.118244 | 0.00181932 | 0.042976725 |
| *RHOF* | -0.334095 | 0.10717 | -3.117429 | 0.00182436 | 0.043006593 |
| *UGT8* | 0.822748 | 0.264019 | 3.116246 | 0.00183169 | 0.043090166 |
| *PRICKLE1* | 1.081873 | 0.347291 | 3.115178 | 0.00183834 | 0.043157417 |
| *PRDM2* | -0.23076 | 0.074137 | -3.112608 | 0.00185442 | 0.04344541 |
| *SPOP* | 0.353455 | 0.113648 | 3.110076 | 0.00187039 | 0.043627046 |
| *CD82* | -0.294294 | 0.094637 | -3.10971 | 0.00187271 | 0.043627046 |
| *CEP104* | 0.585137 | 0.188179 | 3.109466 | 0.00187426 | 0.043627046 |
| *ZNF557* | 0.648745 | 0.20867 | 3.108955 | 0.0018775 | 0.043627046 |
| *TBC1D5* | 0.396361 | 0.127634 | 3.105457 | 0.00189985 | 0.044056503 |
| *PRKAG1* | -0.348097 | 0.112122 | -3.104618 | 0.00190525 | 0.04407319 |
| *NAA10* | -0.297051 | 0.095695 | -3.104142 | 0.00190832 | 0.04407319 |
| *JHY* | -1.440905 | 0.464432 | -3.102508 | 0.00191888 | 0.044227506 |
| *MXD4* | 0.587444 | 0.189399 | 3.101619 | 0.00192466 | 0.044270961 |
| *CCSER2* | -0.379902 | 0.122975 | -3.089269 | 0.0020065 | 0.04596086 |
| *ARID4A* | 0.380401 | 0.123148 | 3.088959 | 0.00200859 | 0.04596086 |
| *HLA.DRA* | -0.278408 | 0.090137 | -3.088716 | 0.00201023 | 0.04596086 |
| *PRDX6* | -0.242524 | 0.078535 | -3.088104 | 0.00201438 | 0.045963413 |
| *MPP6* | 0.987857 | 0.320037 | 3.086695 | 0.00202395 | 0.046089365 |
| *ANO9* | -0.732422 | 0.237331 | -3.086085 | 0.00202811 | 0.046091833 |
| *ORMDL3* | -0.46271 | 0.150016 | -3.084393 | 0.00203968 | 0.046262606 |
| *PRDM8* | -1.004486 | 0.326211 | -3.079251 | 0.00207522 | 0.046975054 |
| *HIKESHI* | -0.291716 | 0.094784 | -3.077703 | 0.00208603 | 0.047125973 |
| *SNX17* | -0.249171 | 0.081039 | -3.074716 | 0.00210703 | 0.047506214 |
| *TMEM123* | -0.344383 | 0.112123 | -3.07147 | 0.00213007 | 0.047930861 |
| *PSME3* | -0.45979 | 0.149789 | -3.069577 | 0.00214362 | 0.048140598 |
| *NEAT1* | 0.405704 | 0.132205 | 3.068761 | 0.00214948 | 0.048177177 |
| *ENSG00000272316* | 0.665408 | 0.216949 | 3.067126 | 0.00216128 | 0.048277013 |
| *ABI3* | 0.378228 | 0.123323 | 3.066968 | 0.00216242 | 0.048277013 |
| *SLC3A2* | -0.329094 | 0.107344 | -3.065791 | 0.00217095 | 0.048372662 |
| *GCSAM* | -0.866975 | 0.282989 | -3.06364 | 0.00218662 | 0.048626681 |
| *ACAT1* | -0.578819 | 0.189 | -3.062534 | 0.00219471 | 0.048654094 |
| *SLC38A7* | -0.797824 | 0.260531 | -3.062304 | 0.0021964 | 0.048654094 |

| | | | | | |
|---|---|---|---|---|---|
| *ENSG00000130313* | -0.214832 | 0.070193 | -3.060583 | 0.00220906 | 0.048839567 |
| *TMEM159* | 0.518281 | 0.169631 | 3.055337 | 0.00224808 | 0.049543481 |
| *SMS* | -0.338219 | 0.110711 | -3.054975 | 0.00225079 | 0.049543481 |
| *SIDT1* | 0.684582 | 0.224119 | 3.054554 | 0.00225395 | 0.049543481 |
| *PWWP3A* | 0.426623 | 0.1397 | 3.053847 | 0.00225928 | 0.049564768 |
| *PRR11* | -1.184217 | 0.387931 | -3.052648 | 0.00226832 | 0.049667439 |
| *TNS3* | 0.923576 | 0.302722 | 3.050907 | 0.00228151 | 0.049775167 |
| *SQLE* | -0.380744 | 0.1248 | -3.050845 | 0.00228198 | 0.049775167 |
| *PPP1R7* | -0.281687 | 0.092361 | -3.049863 | 0.00228946 | 0.049842808 |
| *FBXL8* | -1.027488 | 0.336971 | -3.049191 | 0.00229459 | 0.049859113 |
| *SPINK2* | -1.844774 | 0.605083 | -3.048796 | 0.00229761 | NA |
| *TBC1D4* | -2.118487 | 0.695017 | -3.048111 | 0.00230285 | 0.049943366 |
| *TYROBP* | 1.486227 | 0.48788 | 3.046297 | 0.00231679 | 0.05011008 |
| *MSH6* | -0.589508 | 0.193537 | -3.045966 | 0.00231934 | 0.05011008 |
| *DNAJB11* | -0.398755 | 0.130955 | -3.044976 | 0.00232699 | 0.050180045 |
| *HIST1H4C* | 0.309291 | 0.101688 | 3.041562 | 0.00235354 | 0.050592485 |
| *BARD1* | 0.658194 | 0.216413 | 3.041376 | 0.002355 | 0.050592485 |
| *ATP5IF1* | -0.212162 | 0.069777 | -3.040583 | 0.00236121 | 0.050630363 |
| *COA6.AS1* | -0.780178 | 0.256944 | -3.036367 | 0.00239448 | 0.051247195 |
| *CLIP2* | -0.678081 | 0.22355 | -3.033239 | 0.00241944 | 0.051684223 |
| *ATP5ME* | -0.168229 | 0.055502 | -3.031019 | 0.0024373 | 0.051968398 |
| *GADD45GIP1* | -0.19177 | 0.063294 | -3.029846 | 0.00244679 | 0.05203339 |
| *ARNTL2* | -1.247441 | 0.411763 | -3.029512 | 0.00244949 | 0.05203339 |
| *FLNA* | -0.425828 | 0.140617 | -3.028286 | 0.00245946 | 0.052080578 |
| *THEMIS2* | 0.471911 | 0.155843 | 3.028113 | 0.00246086 | 0.052080578 |
| *SNHG15* | -0.360456 | 0.119065 | -3.027388 | 0.00246677 | 0.052108858 |
| *CWC15* | -0.288097 | 0.095257 | -3.024432 | 0.002491 | 0.052523216 |
| *FCRLA* | 0.419826 | 0.138908 | 3.022322 | 0.00250843 | 0.052690508 |
| *EBP* | -0.510441 | 0.168902 | -3.022112 | 0.00251017 | 0.052690508 |
| *CSNK1G3* | -0.321272 | 0.106318 | -3.021794 | 0.00251282 | 0.052690508 |
| *ENSG00000268400* | -1.566919 | 0.51868 | -3.020976 | 0.00251961 | 0.052735815 |
| *DUSP18* | 0.503377 | 0.16674 | 3.01894 | 0.00253661 | 0.052965177 |
| *PTPRS* | 1.28675 | 0.426281 | 3.01855 | 0.00253987 | 0.052965177 |
| *DDX6* | -0.335708 | 0.111327 | -3.01551 | 0.00256547 | 0.053401206 |
| *KLF9* | 0.939208 | 0.31194 | 3.010864 | 0.00260506 | 0.054126247 |
| *IL4I1* | -2.298578 | 0.76377 | -3.009515 | 0.00261665 | NA |
| *MARCHF1* | 0.244377 | 0.081286 | 3.006383 | 0.00264376 | 0.054701061 |
| *ALDH2* | 1.171226 | 0.389603 | 3.006201 | 0.00264534 | 0.054701061 |
| *TCOF1* | -0.319405 | 0.106256 | -3.005995 | 0.00264714 | 0.054701061 |
| *ZNF322* | 0.613197 | 0.20406 | 3.004988 | 0.00265591 | 0.054782938 |
| *KDM4B* | 0.545755 | 0.181734 | 3.003036 | 0.002673 | 0.055029185 |
| *CLCF1* | 0.624362 | 0.207946 | 3.002523 | 0.00267751 | 0.055029185 |
| *CBR1* | 0.478737 | 0.159509 | 3.001321 | 0.00268811 | 0.055049355 |
| *FAM53C* | 0.496628 | 0.16547 | 3.001315 | 0.00268817 | 0.055049355 |
| *THOC3* | -0.472203 | 0.157434 | -2.999377 | 0.00270532 | 0.055197168 |
| *NFE2L1* | -0.440317 | 0.146821 | -2.998997 | 0.0027087 | 0.055197168 |
| *LST1* | -1.043603 | 0.348 | -2.998859 | 0.00270993 | 0.055197168 |

| | | | | | |
|---|---|---|---|---|---|
| *PHPT1* | -0.400367 | 0.133559 | -2.997672 | 0.0027205 | 0.055289573 |
| *ROCK1* | 0.236526 | 0.078914 | 2.997261 | 0.00272417 | 0.055289573 |
| *UQCRFS1* | -0.206894 | 0.069044 | -2.996552 | 0.00273052 | 0.055319654 |
| *PWP1* | -0.303678 | 0.101418 | -2.99432 | 0.00275058 | 0.055568655 |
| *CRYBG1* | -0.440193 | 0.14702 | -2.994099 | 0.00275257 | 0.055568655 |
| *CCDC15* | -1.513863 | 0.505808 | -2.992958 | 0.00276288 | 0.05567816 |
| *MRPL47* | -0.31048 | 0.10376 | -2.992296 | 0.00276888 | 0.055700369 |
| *PFKP* | -0.889787 | 0.297563 | -2.990244 | 0.00278754 | 0.055977004 |
| *CHMP6* | -0.463733 | 0.155111 | -2.989694 | 0.00279257 | 0.055979281 |
| *TRAC* | -1.063878 | 0.356124 | -2.987385 | 0.00281375 | 0.056304707 |
| *JDP2* | 0.795309 | 0.266737 | 2.981627 | 0.00286721 | 0.057273733 |
| *IDH2* | -0.374895 | 0.125799 | -2.980115 | 0.0028814 | 0.057456462 |
| *PLIN3* | -0.418571 | 0.140527 | -2.978579 | 0.00289588 | 0.057644216 |
| *APOBEC3B* | -1.890287 | 0.634768 | -2.977918 | 0.00290214 | 0.057655006 |
| *ADD3* | 0.333771 | 0.112099 | 2.977452 | 0.00290655 | 0.057655006 |
| *DMAC2L* | -0.344806 | 0.115988 | -2.972765 | 0.0029513 | 0.058440871 |
| *SHLD2* | -0.505355 | 0.170053 | -2.971748 | 0.00296109 | 0.05853302 |
| *CD200R1* | 1.302202 | 0.438373 | 2.970532 | 0.00297285 | 0.058663498 |
| *ENSG00000271204* | 0.470375 | 0.158381 | 2.969899 | 0.00297898 | 0.058682718 |
| *SIGLEC5* | 1.38168 | 0.465603 | 2.967508 | 0.00300224 | 0.058992494 |
| *NOL4L* | -1.221195 | 0.411562 | -2.96722 | 0.00300506 | 0.058992494 |
| *SH3PXD2A* | 0.875675 | 0.29529 | 2.965475 | 0.00302216 | 0.059225906 |
| *GHITM* | -0.269114 | 0.090793 | -2.964053 | 0.00303616 | 0.059398214 |
| *ZNF487* | -1.133078 | 0.382412 | -2.962978 | 0.00304678 | 0.059461606 |
| *CARD6* | 1.627546 | 0.549442 | 2.962181 | 0.00305468 | 0.059461606 |
| *SARS* | -0.297979 | 0.100604 | -2.961899 | 0.00305748 | 0.059461606 |
| *ACP1* | -0.227657 | 0.076869 | -2.961616 | 0.00306029 | 0.059461606 |
| *BMERB1* | 1.329343 | 0.448973 | 2.960853 | 0.00306789 | 0.059507599 |
| *TUBGCP5* | -0.743719 | 0.25128 | -2.959728 | 0.00307911 | 0.059623749 |
| *NDUFA13* | -0.253407 | 0.085655 | -2.958472 | 0.00309169 | 0.059692493 |
| *IGHM* | 0.388805 | 0.13143 | 2.95826 | 0.00309381 | 0.059692493 |
| *ZC3H7A* | -0.401955 | 0.135897 | -2.957804 | 0.00309839 | 0.059692493 |
| *GLS* | 0.379668 | 0.128426 | 2.956324 | 0.0031133 | 0.059878347 |
| *RAB37* | -0.445779 | 0.150818 | -2.955747 | 0.00311913 | 0.059889444 |
| *NUMA1* | 0.482943 | 0.163555 | 2.952792 | 0.00314914 | 0.060363869 |
| *MYL9* | -1.078571 | 0.365363 | -2.952056 | 0.00315666 | 0.060406177 |
| *ALG14* | -0.747938 | 0.25342 | -2.95138 | 0.00316357 | 0.060436973 |
| *SURF2* | -0.350091 | 0.118686 | -2.949731 | 0.00318051 | 0.060603768 |
| *MORN3* | 1.611547 | 0.546422 | 2.949272 | 0.00318523 | 0.060603768 |
| *RFTN1* | -0.547239 | 0.185569 | -2.948978 | 0.00318827 | 0.060603768 |
| *SMAP2* | -0.300108 | 0.101838 | -2.946927 | 0.00320949 | 0.060905369 |
| *JUP* | -0.980592 | 0.333205 | -2.942913 | 0.00325139 | 0.061569496 |
| *RHOB* | 0.688492 | 0.234074 | 2.941346 | 0.00326789 | 0.061569496 |
| *CBL* | 0.517882 | 0.176072 | 2.941312 | 0.00326825 | 0.061569496 |
| *FLNB* | -0.770206 | 0.261873 | -2.941146 | 0.00327001 | 0.061569496 |
| *CD27* | -0.650417 | 0.221155 | -2.941002 | 0.00327152 | 0.061569496 |
| *LINC00662* | 0.461305 | 0.156886 | 2.940381 | 0.00327809 | 0.061591256 |

| | | | | | |
|---|---|---|---|---|---|
| *TGFBR2* | 0.354239 | 0.120599 | 2.937339 | 0.00331043 | 0.062096372 |
| *RANBP1* | -0.26969 | 0.091927 | -2.933741 | 0.00334903 | 0.062717239 |
| *KIF20B* | 0.40725 | 0.13886 | 2.932808 | 0.00335912 | 0.062802844 |
| *NT5E* | -0.738595 | 0.251917 | -2.931896 | 0.003369 | 0.06287666 |
| *CPTP* | -0.478205 | 0.163131 | -2.931425 | 0.00337411 | 0.06287666 |
| *ENSG00000256950* | -1.659156 | 0.566319 | -2.929721 | 0.00339267 | 0.062999382 |
| *BACH2* | 0.402904 | 0.137538 | 2.929401 | 0.00339616 | 0.062999382 |
| *ELP3* | -0.548845 | 0.187364 | -2.929297 | 0.0033973 | 0.062999382 |
| *TNFRSF25* | -1.269731 | 0.433729 | -2.927479 | 0.00341722 | 0.063265797 |
| *VASP* | -0.270589 | 0.092467 | -2.926328 | 0.0034299 | 0.063397389 |
| *YPEL2* | 0.465132 | 0.159013 | 2.925128 | 0.00344315 | 0.063539155 |
| *EID1* | -0.285194 | 0.097577 | -2.922763 | 0.0034694 | 0.063878302 |
| *EIF3J.DT* | 0.424513 | 0.145259 | 2.922463 | 0.00347274 | 0.063878302 |
| *SELL* | -0.706237 | 0.241737 | -2.921509 | 0.0034834 | 0.063933475 |
| *DND1* | 1.633138 | 0.559066 | 2.92119 | 0.00348697 | 0.063933475 |
| *KPNB1* | 0.21022 | 0.072127 | 2.914571 | 0.00356178 | 0.065200029 |
| *ATP5F1EP2* | -0.332282 | 0.114033 | -2.91392 | 0.00356922 | 0.065231343 |
| *ZNF761* | -0.676865 | 0.232377 | -2.912787 | 0.00358219 | 0.065358413 |
| *HACD4* | -0.525487 | 0.180436 | -2.912311 | 0.00358765 | 0.065358413 |
| *CDK6* | -1.1203 | 0.384875 | -2.910816 | 0.00360486 | 0.065566968 |
| *SCLT1* | -0.810927 | 0.27874 | -2.909261 | 0.00362284 | 0.065788908 |
| *DEF8* | -0.547669 | 0.188315 | -2.908261 | 0.00363445 | 0.065894717 |
| *FAM120A* | 0.336518 | 0.115747 | 2.907369 | 0.00364483 | 0.06597784 |
| *ZNF829* | 0.772097 | 0.265852 | 2.904242 | 0.00368143 | 0.066534612 |
| *PGM3* | 0.861649 | 0.296755 | 2.903574 | 0.0036893 | 0.066571097 |
| *NAPA* | -0.348671 | 0.120191 | -2.90098 | 0.00371998 | 0.067018455 |
| *NME4* | 0.831776 | 0.286822 | 2.899976 | 0.00373191 | 0.067127294 |
| *ITPRIPL2* | -0.79593 | 0.275407 | -2.890016 | 0.00385223 | 0.069182076 |
| *ZNF318* | 0.599624 | 0.207595 | 2.888429 | 0.00387172 | 0.069422617 |
| *CCSAP* | 1.15084 | 0.398558 | 2.88751 | 0.00388305 | 0.069516319 |
| *PTP4A3* | -1.023515 | 0.354577 | -2.886583 | 0.0038945 | 0.069611928 |
| *GGA2* | 0.334712 | 0.116154 | 2.881624 | 0.00395632 | 0.070605783 |
| *NDUFS8* | -0.244765 | 0.084958 | -2.881025 | 0.00396384 | 0.070605783 |
| *TMEM179B* | -0.249462 | 0.0866 | -2.880638 | 0.00396871 | 0.070605783 |
| *MYBL2* | -1.489436 | 0.517182 | -2.87991 | 0.00397789 | 0.070658782 |
| *ADAP2* | 1.015507 | 0.352998 | 2.87681 | 0.00401717 | 0.071245304 |
| *BDP1* | 0.292432 | 0.101731 | 2.874562 | 0.00404589 | 0.071643014 |
| *ANXA5* | -0.46511 | 0.161872 | -2.873312 | 0.00406193 | 0.071781437 |
| *ENSG00000235078* | 0.896536 | 0.312059 | 2.872971 | 0.00406631 | 0.071781437 |
| *EPHA4* | -0.976376 | 0.340005 | -2.871648 | 0.00408337 | 0.071970983 |
| *MKKS* | -0.326334 | 0.113717 | -2.869707 | 0.00410853 | 0.07230243 |
| *AUP1* | -0.233903 | 0.081583 | -2.867057 | 0.00414309 | 0.072798179 |
| *POLE3* | -0.264611 | 0.092483 | -2.861173 | 0.00422076 | 0.07396129 |
| *EMC9* | 0.694872 | 0.242872 | 2.86106 | 0.00422228 | 0.07396129 |
| *BLCAP* | -0.297613 | 0.104074 | -2.859634 | 0.00424131 | 0.074180543 |
| *RUNX3* | -0.317306 | 0.11101 | -2.858363 | 0.00425833 | 0.07426587 |
| *SERPINB9* | -0.299445 | 0.104763 | -2.858296 | 0.00425923 | 0.07426587 |

| | | | | | |
|---|---|---|---|---|---|
| *PFDN4* | -0.208099 | 0.072836 | -2.857097 | 0.00427535 | 0.074432993 |
| *PXMP2* | -0.661705 | 0.231829 | -2.854281 | 0.00431344 | 0.074981422 |
| *NFE2L2* | 0.293004 | 0.102713 | 2.85265 | 0.00433563 | 0.075252237 |
| *RAB40B* | 1.239021 | 0.434735 | 2.85006 | 0.0043711 | 0.075644013 |
| *DUSP14* | 0.679096 | 0.238277 | 2.850032 | 0.00437149 | 0.075644013 |
| *ABCB4* | 0.519646 | 0.182436 | 2.848366 | 0.00439444 | 0.075824917 |
| *DGCR6* | -1.482061 | 0.520331 | -2.848306 | 0.00439526 | 0.075824917 |
| *NOC3L* | 0.414398 | 0.14567 | 2.844782 | 0.00444418 | 0.076552833 |
| *TUBA1C* | -0.597921 | 0.21024 | -2.843995 | 0.00445517 | 0.076626248 |
| *RPL8* | 0.270633 | 0.095288 | 2.84017 | 0.00450895 | 0.077434275 |
| *LEPROT* | 0.417224 | 0.14695 | 2.839224 | 0.00452234 | 0.077487244 |
| *PIGC* | -0.366835 | 0.129213 | -2.838991 | 0.00452565 | 0.077487244 |
| *CDC40* | 0.267333 | 0.09419 | 2.838236 | 0.00453637 | 0.077546963 |
| *PURA* | 0.442038 | 0.155769 | 2.837786 | 0.00454276 | 0.077546963 |
| *CD58* | -0.361033 | 0.12729 | -2.8363 | 0.00456396 | 0.077792298 |
| *KCNK6* | 0.477573 | 0.168457 | 2.834987 | 0.00458275 | 0.077833143 |
| *MAPRE2* | 0.423115 | 0.149248 | 2.834972 | 0.00458297 | 0.077833143 |
| *SOX5* | 1.079063 | 0.380662 | 2.834701 | 0.00458687 | 0.077833143 |
| *ASF1A* | 0.378515 | 0.133596 | 2.833286 | 0.00460722 | 0.077952786 |
| *N4BP3* | -0.668229 | 0.235852 | -2.833259 | 0.00460761 | 0.077952786 |
| *PPP2R5C* | -0.261105 | 0.092224 | -2.831216 | 0.00463713 | 0.078229583 |
| *RENBP* | 0.730978 | 0.25821 | 2.830942 | 0.00464111 | 0.078229583 |
| *LY6E* | 0.378336 | 0.133654 | 2.830703 | 0.00464458 | 0.078229583 |
| *S1PR1* | 0.426963 | 0.15087 | 2.830015 | 0.00465459 | 0.078282292 |
| *EIF3A* | -0.209387 | 0.074005 | -2.829343 | 0.00466437 | 0.078331167 |
| *PTPN18* | 0.502422 | 0.17767 | 2.827829 | 0.00468648 | 0.078586581 |
| *ERP29* | 0.158063 | 0.05591 | 2.827112 | 0.00469699 | 0.078647005 |
| *CYP3A5* | 0.738678 | 0.261374 | 2.826136 | 0.00471133 | 0.078771198 |
| *TRA2B* | -0.200956 | 0.07112 | -2.82558 | 0.00471951 | 0.078792346 |
| *SLC19A1* | -1.281506 | 0.454236 | -2.821234 | 0.00478393 | 0.079750793 |
| *ROR1* | 1.204433 | 0.42699 | 2.820752 | 0.00479112 | 0.079753901 |
| *KCNC3* | 0.979689 | 0.347521 | 2.81908 | 0.00481615 | 0.08005357 |
| *EIF4G3* | 0.528955 | 0.187743 | 2.817439 | 0.00484083 | 0.080346447 |
| *CORO2B* | -2.23774 | 0.794929 | -2.81502 | 0.00487742 | NA |
| *CAPZB* | -0.25638 | 0.091134 | -2.813231 | 0.00490464 | 0.081287135 |
| *CXorf21* | 0.445883 | 0.158623 | 2.810956 | 0.00493945 | 0.08174501 |
| *PPIL4* | 0.431917 | 0.153757 | 2.809095 | 0.0049681 | 0.081977063 |
| *GATAD1* | 0.437772 | 0.155866 | 2.808647 | 0.00497501 | 0.081977063 |
| *C12orf74.PLEKHG7* | -0.654418 | 0.233002 | -2.808644 | 0.00497507 | 0.081977063 |
| *CCL4* | -1.301935 | 0.463658 | -2.807961 | 0.00498563 | 0.082032299 |
| *FUT11* | 0.604936 | 0.215496 | 2.807182 | 0.0049977 | 0.082112202 |
| *TAOK3* | 0.237267 | 0.084559 | 2.805922 | 0.00501729 | 0.082242478 |
| *IFIT5* | 0.545451 | 0.194405 | 2.805743 | 0.00502007 | 0.082242478 |
| *FAHD2B* | -0.823504 | 0.293674 | -2.804143 | 0.00504505 | 0.082446893 |
| *ENSG00000237513* | 0.917276 | 0.327129 | 2.804017 | 0.00504703 | 0.082446893 |
| *ENSG00000213865* | 0.703823 | 0.251072 | 2.803269 | 0.00505874 | 0.082499164 |
| *FBRS* | -0.382159 | 0.136345 | -2.802888 | 0.00506472 | 0.082499164 |

| | | | | | |
|---|---|---|---|---|---|
| *ENSG00000180448* | -0.319561 | 0.114058 | -2.801755 | 0.00508255 | 0.082576588 |
| *PRKCH* | 2.404249 | 0.85815 | 2.801664 | 0.00508398 | 0.082576588 |
| *ENSG00000272948* | -2.418471 | 0.863395 | -2.801117 | 0.0050926 | NA |
| *PDIK1L* | -0.819898 | 0.292754 | -2.800636 | 0.00510021 | 0.082722141 |
| *DUSP5* | -1.538144 | 0.549351 | -2.799929 | 0.00511138 | 0.082785504 |
| *ZNF22* | 0.285182 | 0.102064 | 2.794141 | 0.00520377 | 0.084095142 |
| *DRG2* | -0.369571 | 0.132276 | -2.79394 | 0.00520702 | 0.084095142 |
| *RNFT1* | 0.478531 | 0.171355 | 2.792621 | 0.0052283 | 0.08420548 |
| *GTF2I* | 0.25347 | 0.090765 | 2.7926 | 0.00522864 | 0.08420548 |
| *COX5B* | -0.169738 | 0.060849 | -2.789476 | 0.00527934 | 0.084805431 |
| *RFX2* | 0.817857 | 0.293203 | 2.789387 | 0.00528079 | 0.084805431 |
| *CFL1* | -0.138171 | 0.049547 | -2.788701 | 0.00529199 | 0.084865555 |
| *MSMP* | -1.024571 | 0.367502 | -2.787933 | 0.00530454 | 0.084947309 |
| *GRHPR* | -0.376078 | 0.134974 | -2.786292 | 0.00533148 | 0.085258722 |
| *PACSIN2* | 0.679518 | 0.244232 | 2.782264 | 0.00539812 | 0.086203333 |
| *HSP90B1* | -0.27065 | 0.097298 | -2.781672 | 0.00540797 | 0.086239697 |
| *CST7* | 2.35645 | 0.848701 | 2.776538 | 0.00549412 | 0.087457233 |
| *CREM* | 0.56478 | 0.203436 | 2.776209 | 0.00549968 | 0.087457233 |
| *CORO7* | -0.48968 | 0.176564 | -2.773381 | 0.00554772 | 0.088098032 |
| *ATP5MF* | -0.176374 | 0.063642 | -2.771361 | 0.00558225 | 0.08852296 |
| *FBLN2* | 2.142063 | 0.773242 | 2.770238 | 0.00560154 | NA |
| *NMT2* | 0.535575 | 0.193371 | 2.769672 | 0.00561128 | 0.088859515 |
| *PIK3C2A* | 0.561815 | 0.203004 | 2.767501 | 0.00564878 | 0.089226146 |
| *LAMTOR3* | 0.462681 | 0.167188 | 2.767425 | 0.0056501 | 0.089226146 |
| *YBX3* | 0.901049 | 0.325982 | 2.764104 | 0.00570794 | 0.090014609 |
| *TSNARE1* | -1.40518 | 0.508566 | -2.763024 | 0.00572685 | 0.090166741 |
| *PIGF* | 0.525227 | 0.190117 | 2.76265 | 0.00573342 | 0.090166741 |
| *NDUFB7* | -0.199012 | 0.072094 | -2.760456 | 0.00577208 | 0.090649528 |
| *TTC3* | 0.256506 | 0.092959 | 2.759339 | 0.00579184 | 0.090834475 |
| *RPS9* | 0.319199 | 0.11579 | 2.756716 | 0.0058385 | 0.091440391 |
| *TMEM141* | -0.344733 | 0.125177 | -2.753951 | 0.00588806 | 0.091983152 |
| *PTPN12* | 0.58 | 0.210612 | 2.753881 | 0.00588931 | 0.091983152 |
| *DVL3* | 0.694076 | 0.252656 | 2.747126 | 0.00601201 | 0.093571053 |
| *ZFYVE21* | 0.547785 | 0.19942 | 2.746895 | 0.00601625 | 0.093571053 |
| *GUK1* | -0.235536 | 0.08575 | -2.746769 | 0.00601854 | 0.093571053 |
| *ENSG00000179094* | -0.76847 | 0.279802 | -2.74648 | 0.00602385 | 0.093571053 |
| *BACH1.IT2* | -1.871399 | 0.681466 | -2.746138 | 0.00603014 | NA |
| *CBWD3* | -0.36329 | 0.132463 | -2.742589 | 0.00609569 | 0.094557875 |
| *DCP1B* | -0.554224 | 0.202213 | -2.740794 | 0.00612908 | 0.09475783 |
| *TXK* | -1.473584 | 0.537797 | -2.740039 | 0.00614319 | 0.09475783 |
| *WAC* | 0.246755 | 0.090056 | 2.740008 | 0.00614376 | 0.09475783 |
| *DHCR7* | -0.66866 | 0.244066 | -2.739667 | 0.00615015 | 0.09475783 |
| *AP4M1* | -0.519086 | 0.189471 | -2.739665 | 0.00615019 | 0.09475783 |
| *ENSG00000260267* | -1.066361 | 0.389331 | -2.738954 | 0.0061635 | 0.094779485 |
| *C12orf49* | -0.563148 | 0.205626 | -2.738701 | 0.00616824 | 0.094779485 |
| *PSMD7* | -0.259057 | 0.094649 | -2.737047 | 0.00619935 | 0.095010493 |
| *FAM131A* | 0.941206 | 0.343892 | 2.736922 | 0.0062017 | 0.095010493 |

| | | | | | |
|---|---|---|---|---|---|
| *FKBP11* | -0.360427 | 0.131707 | -2.736572 | 0.00620831 | 0.095010493 |
| *PNP* | 0.687393 | 0.251228 | 2.736127 | 0.0062167 | 0.095011252 |
| *ENSG00000278158* | 0.507077 | 0.18544 | 2.734457 | 0.00624832 | 0.095366431 |
| *HERC4* | 0.319266 | 0.116908 | 2.730914 | 0.0063159 | 0.096268778 |
| *BBS7* | -0.753597 | 0.276137 | -2.729074 | 0.00635125 | 0.096678305 |
| *PKM* | -0.264695 | 0.097078 | -2.726606 | 0.00639894 | 0.097151693 |
| *SINHCAF* | -0.291957 | 0.107078 | -2.726581 | 0.00639942 | 0.097151693 |
| *FBXO8* | 0.573721 | 0.210734 | 2.722494 | 0.00647913 | 0.098230806 |
| *CALM3* | -0.228814 | 0.084076 | -2.721519 | 0.00649827 | 0.098389973 |
| *CCL3L1.CCL3L3* | -2.532956 | 0.93181 | -2.718319 | 0.00656145 | NA |
| *TRIM27* | 0.385003 | 0.141655 | 2.717904 | 0.00656969 | 0.0993393 |
| *NOTCH2* | 0.431454 | 0.158799 | 2.716977 | 0.00658811 | 0.099485692 |
| *TMEM255B* | 1.06475 | 0.391973 | 2.716386 | 0.00659988 | 0.099531476 |
| *TULP3* | 0.742892 | 0.273559 | 2.715651 | 0.00661455 | 0.099612713 |
| *ADRB2* | 0.819895 | 0.30196 | 2.71524 | 0.00662277 | 0.099612713 |
| *ENSG00000274422* | 0.385719 | 0.142099 | 2.714446 | 0.00663867 | 0.099720229 |
| *ENSG00000273319* | 0.439711 | 0.162084 | 2.712848 | 0.00667076 | 0.100070188 |
| *VNN2* | 0.418644 | 0.15435 | 2.712301 | 0.00668179 | 0.100103749 |
| *SECISBP2* | 0.300997 | 0.111 | 2.711698 | 0.00669396 | 0.100120443 |
| *SSBP4* | -0.310863 | 0.114651 | -2.711375 | 0.00670049 | 0.100120443 |
| *GCNT1* | 1.11086 | 0.409851 | 2.7104 | 0.00672022 | 0.100283587 |
| *TSPAN13* | -0.532371 | 0.196501 | -2.709257 | 0.0067434 | 0.100497898 |
| *ENSG00000261079* | -1.857196 | 0.686175 | -2.706593 | 0.00679776 | NA |
| *TUNAR* | -3.452219 | 1.275772 | -2.705983 | 0.00681025 | NA |
| *TIMM8B* | -0.242499 | 0.089659 | -2.704673 | 0.00683717 | 0.101762178 |
| *TNFAIP3* | -0.507204 | 0.187611 | -2.703492 | 0.0068615 | 0.101990949 |
| *ATP5F1E* | -0.105418 | 0.03904 | -2.700286 | 0.00692798 | 0.102813694 |
| *ST6GAL1* | 0.336509 | 0.124635 | 2.699954 | 0.00693491 | 0.102813694 |
| *CHML* | 0.655343 | 0.242834 | 2.698727 | 0.00696052 | 0.102893298 |
| *NLN* | 1.294372 | 0.479678 | 2.698415 | 0.00696705 | 0.102893298 |
| *PEX13* | 0.537678 | 0.199264 | 2.698324 | 0.00696895 | 0.102893298 |
| *CENPU* | -1.367091 | 0.506712 | -2.697967 | 0.00697643 | 0.102893298 |
| *HOXB2* | 1.048754 | 0.388978 | 2.696178 | 0.00701403 | 0.103224726 |
| *ZNF48* | 0.804788 | 0.298515 | 2.695975 | 0.00701829 | 0.103224726 |
| *ABLIM1* | 0.409096 | 0.151764 | 2.695605 | 0.0070261 | 0.103224726 |
| *DDX17* | 0.189026 | 0.070137 | 2.695116 | 0.00703642 | 0.103243076 |
| *CMIP* | 0.448417 | 0.166441 | 2.694144 | 0.00705697 | 0.10341134 |
| *SMIM10* | -2.798896 | 1.039366 | -2.692889 | 0.00708359 | NA |
| *SLAMF6* | 0.528455 | 0.196282 | 2.692321 | 0.00709566 | 0.103844714 |
| *RAP1B* | -0.308644 | 0.114673 | -2.691525 | 0.00711262 | 0.103959248 |
| *CLEC11A* | -1.107656 | 0.412151 | -2.687498 | 0.00719896 | 0.105086288 |
| *PNO1* | -0.47615 | 0.177207 | -2.686971 | 0.00721032 | 0.10511746 |
| *PELO* | 0.727722 | 0.270961 | 2.685711 | 0.00723756 | 0.105379624 |
| *SDHAF2* | -0.277749 | 0.103456 | -2.684719 | 0.00725908 | 0.105558026 |
| *UBE2L3* | -0.192982 | 0.071905 | -2.68383 | 0.00727842 | 0.105657906 |
| *PABPC4* | 0.463426 | 0.172697 | 2.683466 | 0.00728633 | 0.105657906 |
| *CDK19* | 0.501663 | 0.18697 | 2.683124 | 0.00729379 | 0.105657906 |

| | | | | | |
|---|---|---|---|---|---|
| *STK17B* | -0.242186 | 0.090367 | -2.680022 | 0.00736174 | 0.106506639 |
| *RNF149* | 0.341267 | 0.127396 | 2.678796 | 0.00738874 | 0.10676172 |
| *DNASE1* | 0.517607 | 0.193319 | 2.677469 | 0.00741806 | 0.107044876 |
| *ABCA2* | 1.441407 | 0.538438 | 2.677015 | 0.00742813 | 0.107044876 |
| *ERICH1* | 0.236719 | 0.088439 | 2.676636 | 0.00743655 | 0.107044876 |
| *ZNF667* | 1.308454 | 0.488923 | 2.676196 | 0.0074463 | 0.107049932 |
| *MARCHF2* | -0.939616 | 0.351351 | -2.674299 | 0.00748856 | 0.107521692 |
| *TRIM8* | 0.368379 | 0.137812 | 2.673057 | 0.00751636 | 0.10778492 |
| *RABGGTA* | -0.379138 | 0.141917 | -2.67155 | 0.00755019 | 0.108133926 |
| *ADORA2A* | -1.971242 | 0.738127 | -2.670601 | 0.00757156 | NA |
| *FAM129A* | -1.000973 | 0.374838 | -2.670414 | 0.00757577 | 0.108363952 |
| *HACE1* | 1.675857 | 0.627893 | 2.669015 | 0.0076074 | 0.108555958 |
| *EEA1* | 0.41094 | 0.153969 | 2.668977 | 0.00760826 | 0.108555958 |
| *DEK* | 0.19408 | 0.072749 | 2.667815 | 0.00763462 | 0.108795692 |
| *ENSG00000256325* | -1.399703 | 0.524894 | -2.666641 | 0.00766135 | 0.109040196 |
| *CMPK1* | -0.389707 | 0.146176 | -2.66602 | 0.00767551 | 0.109105316 |
| *SH3KBP1* | 0.199329 | 0.074825 | 2.663957 | 0.00772275 | 0.109428175 |
| *APOOL* | 0.639128 | 0.239944 | 2.663658 | 0.00772961 | 0.109428175 |
| *RCC1* | 0.395603 | 0.148531 | 2.663444 | 0.00773454 | 0.109428175 |
| *SRP19* | -0.290333 | 0.10901 | -2.663351 | 0.00773667 | 0.109428175 |
| *TRIM5* | 0.639091 | 0.240016 | 2.662701 | 0.00775163 | 0.109474182 |
| *PPP1R10* | 0.371091 | 0.139383 | 2.662375 | 0.00775915 | 0.109474182 |
| *PCYOX1* | 0.582378 | 0.218843 | 2.661173 | 0.0077869 | 0.109633441 |
| *DNMT1* | -0.322282 | 0.121122 | -2.660799 | 0.00779555 | 0.109633441 |
| *TOMM22* | -0.208136 | 0.078228 | -2.660636 | 0.00779932 | 0.109633441 |
| *TBC1D8B* | 1.432247 | 0.538539 | 2.659505 | 0.00782555 | 0.109843189 |
| *CCDC50* | -0.333078 | 0.125257 | -2.659162 | 0.00783354 | 0.109843189 |
| *RANGRF* | -0.276407 | 0.103966 | -2.658624 | 0.00784604 | 0.109883219 |
| *BCL11A* | 0.344791 | 0.129775 | 2.65683 | 0.00788793 | 0.11033411 |
| *FBXL16* | -1.481948 | 0.55848 | -2.653538 | 0.00796527 | 0.111279241 |
| *GMCL1* | 0.361447 | 0.136317 | 2.651522 | 0.00801299 | 0.111808746 |
| *LINC01869* | 0.802164 | 0.302622 | 2.650712 | 0.00803222 | 0.111939873 |
| *PMEPA1* | 0.495276 | 0.186879 | 2.650251 | 0.0080432 | 0.111955859 |
| *LINC00539* | -1.49929 | 0.565975 | -2.649037 | 0.00807215 | 0.112221539 |
| *SLC25A30* | 1.135296 | 0.428677 | 2.648371 | 0.00808808 | 0.112235292 |
| *KSR1* | -0.754238 | 0.284815 | -2.648171 | 0.00809285 | 0.112235292 |
| *ABHD5* | -0.610194 | 0.230573 | -2.646427 | 0.0081347 | 0.112678418 |
| *EVI2A* | 0.441211 | 0.166791 | 2.645286 | 0.0081622 | 0.112921971 |
| *NARF* | -0.234858 | 0.088847 | -2.643398 | 0.00820785 | 0.113415763 |
| *SLC1A5* | -0.516191 | 0.195363 | -2.642218 | 0.00823649 | 0.11367357 |
| *FADS2* | -1.576383 | 0.597338 | -2.639014 | 0.00831476 | 0.114614871 |
| *DTX1* | 1.160814 | 0.440631 | 2.634436 | 0.00842773 | 0.115934978 |
| *MAP3K1* | 0.267826 | 0.101668 | 2.634308 | 0.00843089 | 0.115934978 |
| *P2RX4* | -0.462592 | 0.17572 | -2.632557 | 0.00847447 | 0.116390005 |
| *ENSG00000261386* | 0.575547 | 0.218663 | 2.632121 | 0.00848536 | 0.116390005 |
| *TCF4* | 0.187653 | 0.071316 | 2.631276 | 0.0085065 | 0.116390005 |
| *LINC02422* | 0.735465 | 0.279523 | 2.631145 | 0.00850978 | 0.116390005 |

| | | | | | |
|---|---|---|---|---|---|
| *ARHGEF10* | 1.409668 | 0.535805 | 2.630932 | 0.0085151 | 0.116390005 |
| *HLA.G* | -1.32891 | 0.505293 | -2.629977 | 0.00853907 | 0.116517999 |
| *P3H4* | 1.720472 | 0.654269 | 2.629611 | 0.00854827 | 0.116517999 |
| *ZNF90* | -0.390805 | 0.148646 | -2.629107 | 0.00856095 | 0.116517999 |
| *TTC33* | 0.706755 | 0.268872 | 2.628596 | 0.00857381 | 0.116517999 |
| *ENSG00000258168* | -0.941235 | 0.358085 | -2.628524 | 0.00857563 | 0.116517999 |
| *RDX* | -0.416159 | 0.158379 | -2.627611 | 0.00859867 | 0.116691834 |
| *DDX39A* | -0.224438 | 0.085471 | -2.625896 | 0.00864212 | 0.117141823 |
| *DDI2* | 0.510449 | 0.19446 | 2.624964 | 0.00866582 | 0.117323479 |
| *ZNF283* | 0.951637 | 0.362675 | 2.623943 | 0.00869184 | 0.117536035 |
| *MYO15B* | 0.892875 | 0.340746 | 2.620355 | 0.00878382 | 0.118638934 |
| *CTSC* | 0.38143 | 0.145711 | 2.617718 | 0.008852 | 0.119218884 |
| *RNF41* | 0.278123 | 0.106248 | 2.617688 | 0.00885277 | 0.119218884 |
| *GLMN* | -0.428749 | 0.163809 | -2.617365 | 0.00886116 | 0.119218884 |
| *CRTC3* | 0.369411 | 0.141156 | 2.617052 | 0.00886927 | 0.119218884 |
| *NUFIP2* | 0.363108 | 0.138767 | 2.616674 | 0.00887912 | 0.119218884 |
| *SCRN3* | -0.725137 | 0.277226 | -2.615686 | 0.00890485 | 0.119423596 |
| *MPST* | 0.282447 | 0.10805 | 2.614032 | 0.00894808 | 0.119862098 |
| *ZSCAN18* | 0.579553 | 0.221774 | 2.613262 | 0.00896826 | 0.119991357 |
| *TRIM13* | 0.266487 | 0.102006 | 2.612468 | 0.00898912 | 0.120129284 |
| *PHF2* | 0.985914 | 0.377447 | 2.612062 | 0.0089998 | 0.120131027 |
| *ENSG00000251661* | -1.499405 | 0.574249 | -2.61107 | 0.00902595 | 0.120338984 |
| *TRIO* | -0.496699 | 0.19032 | -2.609805 | 0.00905939 | 0.120496473 |
| *SMARCB1* | -0.321647 | 0.123257 | -2.60957 | 0.00906561 | 0.120496473 |
| *TNFAIP8* | -0.289231 | 0.110841 | -2.609423 | 0.00906951 | 0.120496473 |
| *HS3ST1* | 1.059962 | 0.406344 | 2.608537 | 0.00909302 | 0.120625135 |
| *ATP5PF* | -0.15767 | 0.06045 | -2.60826 | 0.00910039 | 0.120625135 |
| *TIPRL* | -0.414066 | 0.158823 | -2.607092 | 0.00913149 | 0.120896653 |
| *LRRC37B* | 0.580232 | 0.222636 | 2.606194 | 0.00915547 | 0.120963985 |
| *NEMP1* | 0.774818 | 0.297323 | 2.605982 | 0.00916113 | 0.120963985 |
| *APAF1* | 0.561014 | 0.215328 | 2.605398 | 0.00917677 | 0.120963985 |
| *YTHDC1* | 0.24269 | 0.093161 | 2.605056 | 0.00918594 | 0.120963985 |
| *MYO1D* | -1.15546 | 0.443569 | -2.604915 | 0.00918969 | 0.120963985 |
| *RGS1* | 0.724749 | 0.278294 | 2.604259 | 0.00920731 | 0.121055947 |
| *TRIAP1* | -0.287115 | 0.110273 | -2.603664 | 0.00922331 | 0.121126351 |
| *FCGR2A* | 1.165545 | 0.448014 | 2.601583 | 0.00927947 | 0.121723491 |
| *SERINC1* | 0.344523 | 0.132493 | 2.600303 | 0.00931415 | 0.121947627 |
| *TOPORS* | 0.312494 | 0.120183 | 2.600162 | 0.00931799 | 0.121947627 |
| *NDUFS7* | -0.211248 | 0.081256 | -2.599768 | 0.00932868 | 0.121947627 |
| *DPH7* | -0.380471 | 0.146384 | -2.599127 | 0.00934611 | 0.121957932 |
| *ENSG00000167807* | 0.879136 | 0.338341 | 2.59837 | 0.00936674 | 0.121957932 |
| *CBX6* | -0.263882 | 0.101572 | -2.597968 | 0.00937772 | 0.121957932 |
| *RGCC* | 0.975829 | 0.37564 | 2.597776 | 0.00938296 | 0.121957932 |
| *C1orf174* | 0.358216 | 0.137894 | 2.597764 | 0.00938331 | 0.121957932 |
| *SIAH1* | 0.392431 | 0.151087 | 2.597382 | 0.00939374 | 0.121957932 |
| *ARMCX2* | 0.674595 | 0.259835 | 2.596241 | 0.00942498 | 0.122224214 |
| *ATP2B4* | -0.857822 | 0.330636 | -2.594463 | 0.0094739 | 0.122580402 |

| | | | | | |
|---|---|---|---|---|---|
| *GM2A* | 0.487714 | 0.187984 | 2.594448 | 0.00947428 | 0.122580402 |
| *DEDD2* | 0.356817 | 0.137551 | 2.594069 | 0.00948475 | 0.122580402 |
| *CLMN* | 0.83361 | 0.321617 | 2.591937 | 0.00954373 | 0.123202867 |
| *CLN8* | 0.367003 | 0.141703 | 2.589943 | 0.0095992 | 0.123755465 |
| *WSB1* | 0.42969 | 0.165928 | 2.589617 | 0.00960828 | 0.123755465 |
| *CXCR5* | -0.318655 | 0.123139 | -2.58777 | 0.00965995 | 0.12428044 |
| *CD99* | -0.268465 | 0.103786 | -2.58672 | 0.00968944 | 0.124519181 |
| *RHBDD2* | -0.388854 | 0.15039 | -2.585641 | 0.00971981 | 0.12476857 |
| *SH2D3C* | 0.49743 | 0.192485 | 2.584251 | 0.00975908 | 0.125131575 |
| *BOLA2B* | -0.439858 | 0.170252 | -2.583567 | 0.00977844 | 0.125238787 |
| *IL12A* | 0.956266 | 0.3704 | 2.58171 | 0.00983121 | 0.125773226 |
| *DAP3* | -0.234434 | 0.090893 | -2.579244 | 0.00990168 | 0.126477082 |
| *UQCRC2* | -0.205572 | 0.07971 | -2.579008 | 0.00990845 | 0.126477082 |
| *CCPG1* | 0.399349 | 0.154878 | 2.578475 | 0.00992374 | 0.126530416 |
| *SNRPA1* | -0.295757 | 0.114755 | -2.577291 | 0.00995781 | 0.126822863 |
| *RHOBTB3* | 1.643786 | 0.638544 | 2.574273 | 0.01004509 | 0.127791525 |
| *RASGRP3* | -0.430791 | 0.167394 | -2.57351 | 0.01006726 | 0.127930615 |
| *CAMK2D* | 0.54122 | 0.21034 | 2.573073 | 0.01008 | 0.127949653 |
| *IFT57* | 0.431118 | 0.167623 | 2.571953 | 0.01011268 | 0.128221513 |
| *SPPL2A* | -0.315104 | 0.122557 | -2.571076 | 0.0101383 | 0.128228008 |
| *TMED8* | 0.332389 | 0.129293 | 2.570828 | 0.01014556 | 0.128228008 |
| *ENSG00000238142* | 0.662041 | 0.257557 | 2.570469 | 0.01015611 | 0.128228008 |
| *HADHA* | 0.175487 | 0.068272 | 2.570396 | 0.01015824 | 0.128228008 |
| *GNAS* | 0.190397 | 0.074085 | 2.569964 | 0.0101709 | 0.128245669 |
| *ACBD3* | -0.45862 | 0.178482 | -2.56955 | 0.01018308 | 0.128257241 |
| *SKIL* | 0.355162 | 0.138263 | 2.568733 | 0.01020711 | 0.128337902 |
| *DNAJC9* | -0.342619 | 0.133389 | -2.568566 | 0.01021203 | 0.128337902 |
| *BZW1* | -0.227494 | 0.088594 | -2.567829 | 0.01023375 | 0.128469153 |
| *CCDC28B* | -0.552706 | 0.215288 | -2.567283 | 0.01024989 | 0.12853006 |
| *ARPC5* | -0.147067 | 0.057299 | -2.566663 | 0.01026823 | 0.128618364 |
| *ENSG00000272933* | -0.964709 | 0.375973 | -2.565898 | 0.0102909 | 0.128760702 |
| *THAP9.AS1* | -0.484961 | 0.189095 | -2.564649 | 0.01032804 | 0.129037918 |
| *PSMD3* | -0.326374 | 0.127272 | -2.56439 | 0.01033573 | 0.129037918 |
| *ARID4B* | 0.252234 | 0.098389 | 2.563647 | 0.01035789 | 0.129172919 |
| *VSIG10L* | 1.053735 | 0.411097 | 2.563226 | 0.01037045 | 0.129188099 |
| *ZNRD2* | -0.321223 | 0.125344 | -2.562735 | 0.01038512 | 0.12922943 |
| *TANK* | -0.224398 | 0.087688 | -2.559048 | 0.01049593 | 0.130373672 |
| *ENSG00000271553* | -1.12411 | 0.439292 | -2.558914 | 0.01049997 | 0.130373672 |
| *PLEKHA2* | 0.347948 | 0.136034 | 2.557793 | 0.01053387 | 0.130548493 |
| *GDF11* | 0.473946 | 0.185302 | 2.557691 | 0.01053698 | 0.130548493 |
| *MIR210HG* | -1.221008 | 0.477591 | -2.556597 | 0.01057015 | 0.130694639 |
| *COBLL1* | -0.236052 | 0.092333 | -2.556545 | 0.01057173 | 0.130694639 |
| *ICAM2* | -0.481076 | 0.188243 | -2.555608 | 0.01060024 | 0.130812519 |
| *POLR2G* | -0.226152 | 0.088524 | -2.554695 | 0.01062809 | 0.130812519 |
| *TMC6* | -0.252843 | 0.098976 | -2.554596 | 0.0106311 | 0.130812519 |
| *CD53* | -0.270667 | 0.105953 | -2.554594 | 0.01063117 | 0.130812519 |
| *GORASP2* | -0.357618 | 0.140004 | -2.554347 | 0.01063871 | 0.130812519 |

| | | | | | |
|---|---|---|---|---|---|
| *LCP1* | -0.243043 | 0.095177 | -2.553589 | 0.01066191 | 0.1309563 |
| *PAFAH1B2* | 0.231839 | 0.090809 | 2.553051 | 0.01067838 | 0.131017229 |
| *ENSG00000249249* | -0.815392 | 0.319656 | -2.55084 | 0.01074638 | 0.131709658 |
| *ENSG00000273669* | -1.480241 | 0.580659 | -2.549241 | 0.01079576 | NA |
| *PLEKHB2* | 0.346566 | 0.136026 | 2.547782 | 0.01084102 | 0.13236327 |
| *LDLRAD4* | 0.441163 | 0.173168 | 2.547599 | 0.0108467 | 0.13236327 |
| *PRMT1* | -0.263529 | 0.103447 | -2.54748 | 0.01085042 | 0.13236327 |
| *SLC2A5* | -0.828871 | 0.32539 | -2.54732 | 0.01085539 | 0.13236327 |
| *METTL15* | -0.411096 | 0.161392 | -2.547182 | 0.01085967 | 0.13236327 |
| *ARRDC5* | 0.815549 | 0.320246 | 2.546635 | 0.01087671 | 0.13236327 |
| *ORAI2* | 0.215828 | 0.084766 | 2.546159 | 0.01089156 | 0.13236327 |
| *HYOU1* | -0.648803 | 0.25482 | -2.546122 | 0.01089271 | 0.13236327 |
| *TCL1B* | -1.367421 | 0.537501 | -2.544032 | 0.0109581 | 0.132918173 |
| *ENSG00000246528* | -1.402147 | 0.551177 | -2.543917 | 0.01096172 | 0.132918173 |
| *ENSG00000260708* | -0.706028 | 0.277634 | -2.543018 | 0.01098997 | 0.133118881 |
| *ZFP1* | -0.756616 | 0.297703 | -2.541511 | 0.01103744 | 0.133551852 |
| *ENSG00000227070* | -1.954816 | 0.769294 | -2.541051 | 0.01105199 | NA |
| *ATXN3* | 0.261252 | 0.102877 | 2.539472 | 0.01110201 | 0.134190507 |
| *ENSG00000272462* | 2.029764 | 0.799679 | 2.538222 | 0.01114172 | NA |
| *CNNM4* | 1.675273 | 0.66012 | 2.537829 | 0.01115424 | 0.134678849 |
| *H2AFV* | -0.199594 | 0.078693 | -2.536374 | 0.01120071 | 0.135096637 |
| *ARF1* | -0.202138 | 0.079712 | -2.535867 | 0.01121693 | 0.135149178 |
| *ENSG00000260000* | 1.291269 | 0.509401 | 2.534879 | 0.0112486 | 0.135348235 |
| *ENSG00000253948* | -0.72045 | 0.284261 | -2.534471 | 0.01126173 | 0.135348235 |
| *VAT1* | 0.491344 | 0.193897 | 2.534047 | 0.01127535 | 0.135348235 |
| *NINJ1* | -0.507495 | 0.200284 | -2.533872 | 0.011281 | 0.135348235 |
| *ARPC5L* | -0.228321 | 0.090154 | -2.532559 | 0.01132333 | 0.13564763 |
| *SELENOT* | -0.182295 | 0.071988 | -2.532279 | 0.01133237 | 0.13564763 |
| *SAP18* | -0.128338 | 0.050686 | -2.531991 | 0.0113417 | 0.13564763 |
| *PUF60* | -0.316794 | 0.125238 | -2.529529 | 0.01142156 | 0.136396464 |
| *BFAR* | -0.259771 | 0.102704 | -2.529323 | 0.01142827 | 0.136396464 |
| *SLC15A3* | 0.75757 | 0.299614 | 2.528489 | 0.01145546 | 0.136453361 |
| *COMMD2* | -0.3126 | 0.123649 | -2.528124 | 0.01146738 | 0.136453361 |
| *PSMC1* | -0.215822 | 0.08537 | -2.528075 | 0.01146899 | 0.136453361 |
| *PLTP* | -1.071969 | 0.424094 | -2.527666 | 0.01148235 | 0.136469779 |
| *FOXK1* | -0.323861 | 0.128195 | -2.526323 | 0.01152634 | 0.13668222 |
| *ZNF552* | 0.613265 | 0.242777 | 2.526037 | 0.01153575 | 0.13668222 |
| *KLHL28* | 0.427284 | 0.169153 | 2.526022 | 0.01153624 | 0.13668222 |
| *CCDC117* | 0.678214 | 0.268638 | 2.524635 | 0.01158185 | 0.137079995 |
| *SNRNP35* | -0.321174 | 0.127239 | -2.52417 | 0.01159718 | 0.137118906 |
| *ENSG00000259804* | -1.517804 | 0.601632 | -2.52281 | 0.01164212 | 0.137507431 |
| *CD63* | 0.414322 | 0.164324 | 2.52138 | 0.01168955 | 0.137924541 |
| *NEK7* | 0.557774 | 0.221352 | 2.519849 | 0.0117405 | 0.138382365 |
| *POLR3K* | 0.243603 | 0.096696 | 2.519273 | 0.01175974 | 0.13846572 |
| *SARAF* | 0.214722 | 0.085271 | 2.518122 | 0.01179825 | 0.138775656 |
| *MRPL28* | -0.259968 | 0.103284 | -2.517027 | 0.01183498 | 0.139043187 |
| *SNX5* | 0.280093 | 0.111293 | 2.516716 | 0.01184542 | 0.139043187 |

| | | | | | |
|---|---|---|---|---|---|
| *KIDINS220* | 0.591536 | 0.235113 | 2.515965 | 0.01187071 | 0.139196649 |
| *C2orf15* | -1.074437 | 0.427228 | -2.514901 | 0.0119066 | 0.139469387 |
| *ADRM1* | -0.267624 | 0.106444 | -2.514219 | 0.01192964 | 0.139469387 |
| *PGK1* | -0.27207 | 0.108221 | -2.514023 | 0.01193626 | 0.139469387 |
| *ARL4D* | 0.795118 | 0.316298 | 2.513825 | 0.01194297 | 0.139469387 |
| *FGF9* | -1.367257 | 0.544195 | -2.512439 | 0.01198998 | 0.139874859 |
| *RIPOR1* | 0.330336 | 0.131514 | 2.511791 | 0.01201201 | 0.139988435 |
| *CCDC66* | 0.343794 | 0.136934 | 2.510644 | 0.01205113 | 0.140300724 |
| *DDB2* | -0.28235 | 0.112481 | -2.510209 | 0.01206597 | 0.140330004 |
| *PCNP* | -0.191736 | 0.076429 | -2.50869 | 0.01211798 | 0.140791192 |
| *C17orf51* | 1.61038 | 0.642079 | 2.508072 | 0.0121392 | 0.1408939 |
| *RTL10* | 1.143342 | 0.456019 | 2.507223 | 0.01216838 | 0.14108878 |
| *GTF2F1* | 0.270745 | 0.108018 | 2.506488 | 0.01219373 | 0.141238842 |
| *ETNK1* | 0.355822 | 0.142032 | 2.505223 | 0.01223741 | 0.141600753 |
| *ENSG00000233461* | -0.451357 | 0.180215 | -2.504544 | 0.01226093 | 0.141728842 |
| *COMMD8* | -0.292786 | 0.116975 | -2.502975 | 0.01231544 | 0.142194018 |
| *TP53TG1* | -0.275218 | 0.10997 | -2.502667 | 0.01232615 | 0.142194018 |
| *HIGD1A* | -0.327919 | 0.131106 | -2.501181 | 0.012378 | 0.14249983 |
| *BAK1* | -0.442414 | 0.176907 | -2.500836 | 0.01239006 | 0.14249983 |
| *DDX5* | -0.260237 | 0.104082 | -2.500312 | 0.01240838 | 0.14249983 |
| *ISOC2* | -0.400832 | 0.160318 | -2.500233 | 0.01241116 | 0.14249983 |
| *CXCR4* | 0.279198 | 0.111674 | 2.500117 | 0.01241523 | 0.14249983 |
| *GABARAPL1* | 1.143316 | 0.457479 | 2.499167 | 0.01244857 | 0.142738545 |
| *SDSL* | 2.264314 | 0.906227 | 2.498617 | 0.0124679 | 0.142816367 |
| *FAM234B* | 1.367705 | 0.547985 | 2.495879 | 0.01256453 | 0.14374506 |
| *ZNF91* | 0.320561 | 0.12845 | 2.495606 | 0.01257422 | 0.14374506 |
| *ALOX5AP* | 0.395889 | 0.158699 | 2.494588 | 0.01261033 | 0.143940115 |
| *WDR74* | 0.279534 | 0.112068 | 2.494334 | 0.01261938 | 0.143940115 |
| *BOP1* | -0.469722 | 0.188336 | -2.494057 | 0.01262921 | 0.143940115 |
| *SUDS3* | 0.27288 | 0.109453 | 2.493121 | 0.01266258 | 0.144176158 |
| *CXorf56* | -0.494044 | 0.198193 | -2.49274 | 0.01267616 | 0.144186587 |
| *KIF22* | -0.444467 | 0.17843 | -2.490984 | 0.01273899 | 0.144756643 |
| *PSMD6.AS2* | 1.687168 | 0.677472 | 2.49039 | 0.0127603 | 0.144854193 |
| *SH3TC1* | 0.421906 | 0.169595 | 2.487729 | 0.01285615 | 0.145796978 |
| *ENSG00000257275* | -1.147503 | 0.461617 | -2.485833 | 0.01292487 | 0.146430421 |
| *ENSG00000259888* | 2.325056 | 0.935727 | 2.484759 | 0.01296392 | NA |
| *TNKS1BP1* | -1.538451 | 0.619315 | -2.484116 | 0.01298734 | NA |
| *GGT1* | -1.101101 | 0.443284 | -2.483964 | 0.0129929 | 0.146784332 |
| *PTAFR* | -0.885852 | 0.356676 | -2.483632 | 0.013005 | 0.146784332 |
| *IDH3G* | -0.268289 | 0.108026 | -2.483558 | 0.01300771 | 0.146784332 |
| *ESF1* | 0.251714 | 0.101353 | 2.483536 | 0.01300853 | 0.146784332 |
| *LRRK2* | 0.504547 | 0.203184 | 2.483206 | 0.01302057 | 0.146784332 |
| *RP2* | 0.654292 | 0.263539 | 2.482718 | 0.01303843 | 0.146815235 |
| *NBPF9* | 0.809397 | 0.326093 | 2.482104 | 0.01306093 | 0.146815235 |
| *COX14* | -0.176766 | 0.071217 | -2.482075 | 0.01306199 | 0.146815235 |
| *RGL1* | -2.725948 | 1.098491 | -2.481538 | 0.01308166 | NA |
| *ENSG00000092330* | -0.224482 | 0.090489 | -2.480762 | 0.01311017 | 0.14704952 |

| | | | | | |
|---|---|---|---|---|---|
| *NDUFAF3* | -0.202375 | 0.081579 | -2.480712 | 0.01311204 | 0.14704952 |
| *LIMD2* | -0.22391 | 0.09027 | -2.480452 | 0.01312158 | 0.14704952 |
| *SFT2D2* | 0.424082 | 0.171094 | 2.478653 | 0.01318794 | 0.14764783 |
| *SLC27A4* | -1.028986 | 0.415266 | -2.4779 | 0.01321583 | 0.147814759 |
| *KLK2* | 1.76354 | 0.712005 | 2.476864 | 0.01325423 | 0.148098744 |
| *TMC8* | -0.272341 | 0.110034 | -2.475062 | 0.01332131 | 0.148337646 |
| *ZNF628* | -0.751506 | 0.30364 | -2.47499 | 0.01332401 | 0.148337646 |
| *TMED2* | -0.179019 | 0.072332 | -2.474957 | 0.01332523 | 0.148337646 |
| *CERS4* | -0.265973 | 0.107469 | -2.47489 | 0.01332772 | 0.148337646 |
| *BLZF1* | 0.588301 | 0.237898 | 2.472908 | 0.01340186 | 0.149017184 |
| *TUBB4B* | -0.328808 | 0.133097 | -2.470437 | 0.01349483 | 0.149904487 |
| *SCAF11* | 0.146606 | 0.059358 | 2.469864 | 0.01351642 | 0.149998058 |
| *NUP50.DT* | -0.813324 | 0.329397 | -2.469126 | 0.01354434 | 0.150161442 |
| *C1orf109* | 0.512984 | 0.207917 | 2.467249 | 0.01361556 | 0.150804258 |
| *DNAJB1* | 0.440152 | 0.178426 | 2.466863 | 0.01363025 | 0.150820232 |
| *AHCYL1* | 0.443945 | 0.180099 | 2.465 | 0.01370133 | 0.151328987 |
| *ENSG00000131408* | 0.213464 | 0.086599 | 2.464961 | 0.01370281 | 0.151328987 |
| *RBM39* | 0.134528 | 0.054586 | 2.464534 | 0.01371916 | 0.151362702 |
| *METRNL* | 1.217334 | 0.494956 | 2.459478 | 0.01391391 | 0.153362848 |
| *TLR4* | -2.163997 | 0.879934 | -2.459273 | 0.01392188 | NA |
| *ENSG00000268403* | -0.623928 | 0.253778 | -2.458552 | 0.01394986 | 0.153504085 |
| *UPP1* | 0.708443 | 0.288166 | 2.458453 | 0.01395369 | 0.153504085 |
| *RGS10* | 0.298272 | 0.121392 | 2.457108 | 0.01400607 | 0.15393162 |
| *SMAD4* | 0.503826 | 0.205477 | 2.451986 | 0.01420701 | 0.155927653 |
| *SNRPG* | -0.143429 | 0.0585 | -2.451782 | 0.01421508 | 0.155927653 |
| *GKAP1* | 0.463908 | 0.189275 | 2.450972 | 0.01424708 | 0.156128293 |
| *MTHFR* | 0.446973 | 0.182405 | 2.450448 | 0.01426786 | 0.156160096 |
| *ZC3H6* | 0.457652 | 0.186781 | 2.450207 | 0.01427742 | 0.156160096 |
| *PDK2* | -0.457412 | 0.186758 | -2.449219 | 0.01431665 | 0.15643899 |
| *BROX* | 0.260914 | 0.10662 | 2.447126 | 0.01440006 | 0.157199502 |
| *ENSG00000237976* | -0.926679 | 0.378781 | -2.44648 | 0.01442588 | 0.157330544 |
| *ENSG00000226571* | 1.030504 | 0.421294 | 2.446045 | 0.0144433 | 0.157369776 |
| *PTPRJ* | 1.245054 | 0.509611 | 2.443146 | 0.01455985 | 0.158487975 |
| *SPATA13* | 0.707886 | 0.289879 | 2.442004 | 0.01460598 | 0.158838281 |
| *GSDME* | -1.199555 | 0.491323 | -2.441478 | 0.01462729 | 0.158918269 |
| *POLH* | 0.759813 | 0.311351 | 2.44037 | 0.01467224 | 0.159244057 |
| *CNIH1* | 0.282838 | 0.115921 | 2.439915 | 0.01469071 | 0.159244057 |
| *GPR146* | 0.783709 | 0.321241 | 2.43963 | 0.01470231 | 0.159244057 |
| *CD55* | -0.274463 | 0.112514 | -2.439362 | 0.01471322 | 0.159244057 |
| *TMEM109* | 0.34701 | 0.142335 | 2.437977 | 0.0147697 | 0.159703478 |
| *REV1* | 0.390671 | 0.160286 | 2.43734 | 0.01479578 | 0.159752584 |
| *PCOLCE* | 1.042704 | 0.427832 | 2.43718 | 0.0148023 | 0.159752584 |
| *FABP5* | 0.502648 | 0.206321 | 2.436244 | 0.01484066 | 0.159913057 |
| *NOTCH1* | 0.739119 | 0.303542 | 2.434982 | 0.01489252 | 0.159913057 |
| *FANCB* | -1.014061 | 0.416505 | -2.434691 | 0.0149045 | 0.159913057 |
| *SOS1* | 0.49467 | 0.203187 | 2.434558 | 0.01490997 | 0.159913057 |
| *ENSG00000132481* | -0.90414 | 0.371428 | -2.434226 | 0.01492368 | 0.159913057 |

| | | | | | |
|---|---|---|---|---|---|
| *ZDHHC4* | 0.301876 | 0.124032 | 2.433852 | 0.01493909 | 0.159913057 |
| *MID1IP1* | 0.492142 | 0.202222 | 2.433676 | 0.01494636 | 0.159913057 |
| *IGFBP4* | -0.963446 | 0.395913 | -2.433479 | 0.01495449 | 0.159913057 |
| *RAPGEF2* | 0.976854 | 0.401431 | 2.433429 | 0.01495656 | 0.159913057 |
| *RGS17* | -1.355582 | 0.557072 | -2.433404 | 0.01495762 | 0.159913057 |
| *GTSF1* | -0.625603 | 0.25745 | -2.429996 | 0.01509899 | 0.161273089 |
| *HSPA1A* | 1.312088 | 0.540033 | 2.429642 | 0.01511376 | 0.1612795 |
| *MPZL1* | 0.528297 | 0.217516 | 2.428773 | 0.01515001 | 0.161503723 |
| *SLF1* | 0.409225 | 0.168512 | 2.428459 | 0.01516314 | 0.161503723 |
| *MDH2* | -0.207088 | 0.0853 | -2.427769 | 0.01519203 | 0.161660208 |
| *BABAM2* | -0.393799 | 0.16228 | -2.426664 | 0.01523836 | 0.162001862 |
| *BET1* | 0.35046 | 0.144469 | 2.425853 | 0.01527247 | 0.162006405 |
| *CCDC180* | -0.974523 | 0.401736 | -2.425782 | 0.01527545 | 0.162006405 |
| *ENSG00000257354* | -0.748517 | 0.308585 | -2.425639 | 0.01528148 | 0.162006405 |
| *RNF24* | 0.643748 | 0.265587 | 2.423864 | 0.01535634 | 0.162405383 |
| *BCR* | 0.785793 | 0.324199 | 2.423797 | 0.01535917 | 0.162405383 |
| *PDK4* | 1.598314 | 0.659443 | 2.423733 | 0.0153619 | 0.162405383 |
| *IGHA2* | -1.263211 | 0.521442 | -2.422534 | 0.01541267 | 0.162790923 |
| *THOC7* | -0.204978 | 0.084666 | -2.421029 | 0.01547663 | 0.163230182 |
| *FAM184B* | -1.640939 | 0.677827 | -2.420881 | 0.01548293 | 0.163230182 |
| *ABTB2* | 1.988283 | 0.821869 | 2.419222 | 0.01555376 | NA |
| *H2AFJ* | -0.416085 | 0.172092 | -2.417806 | 0.01561439 | 0.164256333 |
| *MAP4K3.DT* | 0.546221 | 0.225927 | 2.417682 | 0.0156197 | 0.164256333 |
| *CCNT2* | 0.385008 | 0.159253 | 2.417593 | 0.01562354 | 0.164256333 |
| *POLR2B* | -0.227998 | 0.094344 | -2.416666 | 0.0156634 | 0.164405064 |
| *PHF6* | 0.367796 | 0.152217 | 2.416255 | 0.01568107 | 0.164405064 |
| *UPF2* | -0.158293 | 0.065519 | -2.415985 | 0.01569271 | 0.164405064 |
| *WIPF2* | 0.336398 | 0.139268 | 2.415477 | 0.01571464 | 0.164405064 |
| *MAP3K2* | 0.320777 | 0.132805 | 2.415403 | 0.01571781 | 0.164405064 |
| *KCTD2* | 1.056069 | 0.43725 | 2.415252 | 0.01572432 | 0.164405064 |
| *BACH1* | 0.383233 | 0.158845 | 2.412622 | 0.01583822 | 0.165398483 |
| *SLC35B2* | -0.342115 | 0.141816 | -2.412388 | 0.01584839 | 0.165398483 |
| *BCL11B* | 2.162671 | 0.896872 | 2.411349 | 0.01589365 | 0.16569344 |
| *ENSG00000077463* | -0.374809 | 0.155466 | -2.410874 | 0.01591435 | 0.16569344 |
| *GPX7* | 0.389983 | 0.161769 | 2.410737 | 0.01592031 | 0.16569344 |
| *IL4R* | -0.456911 | 0.189717 | -2.408387 | 0.0160232 | 0.166612061 |
| *BCDIN3D* | 0.543507 | 0.225814 | 2.406881 | 0.01608941 | 0.167147795 |
| *ENSG00000271133* | -1.471358 | 0.612158 | -2.403561 | 0.01623627 | 0.168519798 |
| *CCNG2* | 0.43342 | 0.180389 | 2.402692 | 0.0162749 | 0.1687272 |
| *ZNF250* | 0.719861 | 0.299668 | 2.402196 | 0.01629695 | 0.1687272 |
| *EIF2AK3* | -0.542762 | 0.225952 | -2.402112 | 0.01630071 | 0.1687272 |
| *TNFRSF14* | 0.207249 | 0.086314 | 2.401097 | 0.01634599 | 0.169042135 |
| *ADAP1* | 1.006077 | 0.419223 | 2.39986 | 0.01640134 | 0.169455031 |
| *SNAP23* | -0.209907 | 0.087478 | -2.39954 | 0.01641568 | 0.169455031 |
| *ELK3* | 0.224984 | 0.093789 | 2.398831 | 0.0164475 | 0.16962974 |
| *FAIM* | 0.35266 | 0.147078 | 2.39778 | 0.01649476 | 0.169785964 |
| *MRPL12* | -0.268821 | 0.112119 | -2.397639 | 0.01650111 | 0.169785964 |

| | | | | | |
|---|---|---|---|---|---|
| *RASGRP2* | -0.225822 | 0.0942 | -2.397272 | 0.01651764 | 0.169785964 |
| *ISCA2* | -0.218416 | 0.091123 | -2.396944 | 0.01653247 | 0.169785964 |
| *NDUFA4* | -0.119837 | 0.049998 | -2.396839 | 0.01653721 | 0.169785964 |
| *TMF1* | 0.268353 | 0.112002 | 2.395972 | 0.01657635 | 0.170034479 |
| *CCNG1* | -0.216404 | 0.090343 | -2.395358 | 0.01660416 | 0.170166494 |
| *CMTM7* | -0.335936 | 0.14031 | -2.394245 | 0.01665461 | 0.170530006 |
| *C1orf56* | -0.286223 | 0.119573 | -2.393716 | 0.01667865 | 0.170622778 |
| *HIST1H3E* | 1.206618 | 0.504391 | 2.392225 | 0.01674658 | 0.171163876 |
| *SLC35G5* | 1.205195 | 0.503866 | 2.391895 | 0.01676163 | 0.171164081 |
| *SRSF3* | -0.138268 | 0.057822 | -2.391279 | 0.0167898 | 0.171298085 |
| *TMEM156* | -0.315642 | 0.132025 | -2.390768 | 0.01681316 | 0.171382872 |
| *ETFRF1* | -0.365199 | 0.152882 | -2.38876 | 0.01690534 | 0.172017957 |
| *IGLC3* | -0.448147 | 0.187664 | -2.388026 | 0.01693916 | 0.172017957 |
| *SEZ6* | 0.609776 | 0.255366 | 2.387851 | 0.01694721 | 0.172017957 |
| *MRTO4* | -0.369884 | 0.154921 | -2.387572 | 0.01696008 | 0.172017957 |
| *LSM7* | 0.24735 | 0.103626 | 2.386949 | 0.01698883 | 0.172017957 |
| *MAST4* | 0.912281 | 0.382216 | 2.386819 | 0.01699488 | 0.172017957 |
| *ARID1A* | 0.267873 | 0.112238 | 2.386656 | 0.01700238 | 0.172017957 |
| *NR5A2* | -1.400758 | 0.587064 | -2.386037 | 0.01703102 | 0.172017957 |
| *KRT10* | -0.211233 | 0.08853 | -2.385997 | 0.01703288 | 0.172017957 |
| *CLCN7* | -0.339923 | 0.142476 | -2.385834 | 0.01704045 | 0.172017957 |
| *LIMK1* | 0.451287 | 0.189174 | 2.38557 | 0.01705266 | 0.172017957 |
| *PXMP4* | -0.609007 | 0.255297 | -2.385482 | 0.01705676 | 0.172017957 |
| *IDS* | 0.201988 | 0.084686 | 2.38515 | 0.01707218 | 0.172021105 |
| *GRAP2* | 0.989957 | 0.415313 | 2.383639 | 0.0171424 | 0.172575937 |
| *ARID3B* | 0.512164 | 0.214932 | 2.382914 | 0.01717621 | 0.172763585 |
| *PHETA2* | 1.821283 | 0.764359 | 2.382757 | 0.01718353 | NA |
| *KLK1* | 1.287474 | 0.540597 | 2.381579 | 0.01723859 | 0.173237922 |
| *STRAP* | -0.262982 | 0.110464 | -2.380693 | 0.01728012 | 0.173502117 |
| *APP* | 0.556629 | 0.233884 | 2.379936 | 0.01731567 | 0.173622242 |
| *TLR7* | 0.577798 | 0.242794 | 2.379788 | 0.01732258 | 0.173622242 |
| *POLR2I* | -0.250013 | 0.105087 | -2.3791 | 0.01735497 | 0.17377433 |
| *GPR65* | 0.390253 | 0.164053 | 2.378817 | 0.01736828 | 0.17377433 |
| *NINJ2* | -0.437966 | 0.184208 | -2.377559 | 0.01742766 | 0.174069734 |
| *SLC31A2* | 1.300688 | 0.54709 | 2.377466 | 0.01743206 | 0.174069734 |
| *ATRX* | 0.158209 | 0.066563 | 2.376819 | 0.01746264 | 0.174069734 |
| *RHNO1* | -0.525028 | 0.220904 | -2.376729 | 0.01746693 | 0.174069734 |
| *ENSG00000263394* | 0.970841 | 0.408504 | 2.376574 | 0.01747424 | 0.174069734 |
| *SNRPD3* | -0.197425 | 0.08311 | -2.375469 | 0.01752668 | 0.174439466 |
| *ENSG00000260641* | -2.019112 | 0.850769 | -2.373278 | 0.01763097 | NA |
| *TMEM101* | 0.31414 | 0.132517 | 2.37057 | 0.01776069 | 0.176614192 |
| *ELOA* | -0.297595 | 0.125581 | -2.369737 | 0.01780074 | 0.176665439 |
| *FOLR2* | -0.718149 | 0.303054 | -2.369705 | 0.01780226 | 0.176665439 |
| *FGFR1OP2* | 0.249787 | 0.105418 | 2.369495 | 0.0178124 | 0.176665439 |
| *HIST1H2AG* | 1.691772 | 0.714206 | 2.368746 | 0.01784852 | 0.176869644 |
| *TENT4B* | 0.550037 | 0.232257 | 2.368221 | 0.01787388 | 0.176966951 |
| *CIB1* | 0.317358 | 0.134025 | 2.367898 | 0.01788946 | 0.176967328 |

| | | | | | |
|---|---|---|---|---|---|
| *TTC7A* | -0.437313 | 0.184773 | -2.366755 | 0.01794482 | 0.177360864 |
| *GSDMD* | -0.268794 | 0.113598 | -2.366195 | 0.01797199 | 0.177475366 |
| *SLC25A42* | -0.398135 | 0.168295 | -2.36569 | 0.01799653 | 0.177563638 |
| *MCM6* | 0.922178 | 0.390156 | 2.363611 | 0.01809781 | 0.178408354 |
| *KLK4* | 1.799422 | 0.761511 | 2.362963 | 0.0181295 | 0.178480235 |
| *GADD45A* | -0.548106 | 0.232031 | -2.362213 | 0.01816619 | 0.178480235 |
| *ADAM28* | 0.215418 | 0.091197 | 2.362123 | 0.0181706 | 0.178480235 |
| *SFXN4* | -0.61273 | 0.259414 | -2.361976 | 0.01817782 | 0.178480235 |
| *CBLN3* | -0.652289 | 0.276201 | -2.36165 | 0.01819379 | 0.178480235 |
| *UBE2D3* | -0.126196 | 0.053453 | -2.360883 | 0.01823147 | 0.178480235 |
| *BYSL* | 0.933433 | 0.395375 | 2.360882 | 0.01823152 | 0.178480235 |
| *NKILA* | -1.680653 | 0.711923 | -2.360721 | 0.01823942 | NA |
| *MRPL2* | -0.262369 | 0.11114 | -2.360719 | 0.01823954 | 0.178480235 |
| *PIM1* | -0.409211 | 0.173352 | -2.360584 | 0.01824618 | 0.178480235 |
| *ZFP69B* | 1.946399 | 0.824684 | 2.360174 | 0.01826638 | NA |
| *QRSL1* | -0.337304 | 0.142934 | -2.359848 | 0.01828245 | 0.178681476 |
| *AZIN1.AS1* | -1.298493 | 0.5504 | -2.359178 | 0.01831548 | 0.178848006 |
| *HDX* | 1.483382 | 0.628854 | 2.358865 | 0.0183309 | 0.178848006 |
| *UQCRH* | -0.163911 | 0.069497 | -2.358523 | 0.01834783 | 0.17885989 |
| *SERPINB1* | -0.250558 | 0.10627 | -2.35774 | 0.01838658 | 0.17896099 |
| *C1orf122* | -0.305764 | 0.129698 | -2.357505 | 0.01839821 | 0.17896099 |
| *GTF3A* | -0.167083 | 0.070886 | -2.357047 | 0.01842092 | 0.17896099 |
| *ATP10D* | 0.547785 | 0.232415 | 2.35693 | 0.01842673 | 0.17896099 |
| *BEND4* | 0.903296 | 0.383302 | 2.356616 | 0.01844229 | 0.17896099 |
| *MXI1* | 0.596754 | 0.25327 | 2.356198 | 0.01846308 | 0.17896099 |
| *ENSG00000224610* | -2.132759 | 0.905228 | -2.356045 | 0.01847067 | NA |
| *FNIP1* | -0.337602 | 0.143303 | -2.355865 | 0.01847962 | 0.17896099 |
| *KCTD6* | 0.518775 | 0.220271 | 2.35517 | 0.01851425 | 0.17896099 |
| *BAX* | -0.37787 | 0.16047 | -2.354775 | 0.01853392 | 0.17896099 |
| *PIK3R1* | 0.389995 | 0.165644 | 2.354409 | 0.01855217 | 0.17896099 |
| *PPIE* | -0.294909 | 0.125258 | -2.354406 | 0.01855231 | 0.17896099 |
| *FAM217B* | 0.382042 | 0.162278 | 2.354249 | 0.01856016 | 0.17896099 |
| *C6orf203* | 0.44492 | 0.18899 | 2.354202 | 0.01856253 | 0.17896099 |
| *MRPL41* | -0.178551 | 0.075888 | -2.352827 | 0.01863129 | 0.179471924 |
| *DCXR* | -0.267061 | 0.113549 | -2.351947 | 0.01867545 | 0.179669744 |
| *POGZ* | 0.307733 | 0.13085 | 2.351789 | 0.01868338 | 0.179669744 |
| *TCL1A* | -0.627904 | 0.267167 | -2.350229 | 0.01876189 | 0.180176691 |
| *AGRN* | -0.964904 | 0.410578 | -2.350112 | 0.01876775 | 0.180176691 |
| *LAPTM5* | -0.203404 | 0.086585 | -2.349184 | 0.01881462 | 0.180430728 |
| *PAFAH1B1* | 0.225548 | 0.09602 | 2.348961 | 0.0188259 | 0.180430728 |
| *CCDC138* | -0.721319 | 0.307141 | -2.348494 | 0.01884952 | 0.180505117 |
| *KDM6B* | 0.419681 | 0.178778 | 2.347503 | 0.01889972 | 0.180737596 |
| *SEC61G* | -0.184324 | 0.078523 | -2.347387 | 0.0189056 | 0.180737596 |
| *JAK3* | -0.461986 | 0.196848 | -2.346913 | 0.01892969 | 0.180737596 |
| *AUTS2* | 1.201411 | 0.511944 | 2.346763 | 0.01893729 | 0.180737596 |
| *ENSG00000245904* | 0.351593 | 0.149876 | 2.345901 | 0.01898113 | 0.181004312 |
| *BRK1* | -0.136614 | 0.058246 | -2.345445 | 0.01900437 | 0.181074226 |

| | | | | | |
|---|---|---|---|---|---|
| *GLRX3* | -0.240696 | 0.10266 | -2.344601 | 0.01904747 | 0.181333217 |
| *LRRCC1* | 0.95291 | 0.406526 | 2.344032 | 0.01907655 | 0.18145833 |
| *NFKBIB* | -0.354354 | 0.151222 | -2.343266 | 0.01911575 | 0.181679355 |
| *CELF1* | 0.209135 | 0.089275 | 2.342592 | 0.01915032 | 0.181762878 |
| *ZMYND8* | 0.360727 | 0.154003 | 2.342336 | 0.01916346 | 0.181762878 |
| *CNOT3* | 0.388151 | 0.165723 | 2.342161 | 0.01917242 | 0.181762878 |
| *NDUFAF6* | -0.329319 | 0.1407 | -2.340577 | 0.01925397 | 0.182384067 |
| *TMEM243* | 0.163285 | 0.069783 | 2.33988 | 0.01928993 | 0.182421174 |
| *ABCB1* | 1.01428 | 0.433475 | 2.33988 | 0.01928991 | 0.182421174 |
| *ELK1* | -0.413381 | 0.17682 | -2.337856 | 0.01939474 | 0.183139447 |
| *DNA2* | -1.172159 | 0.501396 | -2.337792 | 0.01939805 | 0.183139447 |
| *NUDT13* | 1.357627 | 0.580864 | 2.337254 | 0.01942597 | 0.183206221 |
| *SNHG32* | 0.178381 | 0.076328 | 2.337036 | 0.0194373 | 0.183206221 |
| *CASP1* | 0.505716 | 0.216421 | 2.336725 | 0.01945347 | 0.183206963 |
| *PSMD6* | -0.266797 | 0.11424 | -2.335403 | 0.01952236 | 0.183703753 |
| *ENG* | 0.79487 | 0.340436 | 2.334858 | 0.01955082 | 0.183819652 |
| *OAS3* | 0.960993 | 0.411754 | 2.333898 | 0.01960105 | 0.184139921 |
| *SEC61A2* | -0.674673 | 0.289139 | -2.333388 | 0.01962778 | 0.184181145 |
| *TARBP2* | -0.404685 | 0.173447 | -2.333197 | 0.01963779 | 0.184181145 |
| *CARD8* | -0.262014 | 0.112357 | -2.331978 | 0.01970186 | 0.184629915 |
| *MRPL34* | -0.203932 | 0.08747 | -2.331451 | 0.0197296 | 0.184737837 |
| *EPRS* | -0.305223 | 0.131011 | -2.329742 | 0.01981979 | 0.185429883 |
| *YWHAG* | 0.331089 | 0.142141 | 2.329295 | 0.01984344 | 0.185498725 |
| *LTBP3* | 0.278726 | 0.119695 | 2.32863 | 0.01987869 | 0.18556955 |
| *SYNPO* | -0.82446 | 0.354068 | -2.328537 | 0.01988362 | 0.18556955 |
| *ZNF267* | -0.338326 | 0.145332 | -2.327959 | 0.0199143 | 0.185591152 |
| *ENSG00000280433* | -1.028761 | 0.441931 | -2.327879 | 0.01991853 | 0.185591152 |
| *SIGIRR* | -0.264875 | 0.113842 | -2.326686 | 0.01998199 | 0.185920243 |
| *PRDX3* | 0.295373 | 0.126955 | 2.326601 | 0.01998651 | 0.185920243 |
| *C16orf74* | -0.39101 | 0.168128 | -2.325669 | 0.02003621 | 0.186084331 |
| *CENPC* | -0.289117 | 0.124316 | -2.325657 | 0.02003683 | 0.186084331 |
| *INPP5F* | 0.753375 | 0.324125 | 2.324334 | 0.0201076 | 0.186589355 |
| *PHTF2* | -0.356323 | 0.153349 | -2.323615 | 0.02014614 | 0.186703791 |
| *RBX1* | -0.131757 | 0.056707 | -2.323492 | 0.02015273 | 0.186703791 |
| *ITGAM* | -0.590877 | 0.25439 | -2.322725 | 0.02019392 | 0.18693334 |
| *BAG6* | -0.24655 | 0.106174 | -2.322143 | 0.02022524 | 0.187071177 |
| *S100A10* | -0.41006 | 0.176691 | -2.320779 | 0.0202988 | 0.187599132 |
| *KDELR2* | -0.282172 | 0.121619 | -2.320134 | 0.0203336 | 0.187768371 |
| *PIP4P2* | 0.533153 | 0.229922 | 2.318847 | 0.02040335 | 0.188259732 |
| *SBDS* | -0.248096 | 0.107046 | -2.317665 | 0.02046753 | 0.188698987 |
| *USF1* | -0.729659 | 0.314891 | -2.317178 | 0.02049406 | 0.188790741 |
| *FANCL* | 0.735084 | 0.317353 | 2.316297 | 0.02054204 | 0.189079773 |
| *CBWD2* | -0.463302 | 0.200069 | -2.315717 | 0.02057373 | 0.189218466 |
| *ENSG00000007237* | 1.078939 | 0.466266 | 2.314002 | 0.02066762 | 0.189928621 |
| *PVT1* | -0.545624 | 0.235979 | -2.312175 | 0.02076806 | 0.190405223 |
| *FDPS* | -0.183729 | 0.079467 | -2.31201 | 0.02077713 | 0.190405223 |
| *ROMO1* | -0.160351 | 0.069366 | -2.311649 | 0.02079706 | 0.190405223 |

| | | | | | |
|---|---|---|---|---|---|
| *SRP72* | -0.220015 | 0.095187 | -2.311404 | 0.02081055 | 0.190405223 |
| *DCTD* | -0.277439 | 0.120039 | -2.311232 | 0.02082002 | 0.190405223 |
| *CCNJ* | -0.872721 | 0.377614 | -2.311144 | 0.02082491 | 0.190405223 |
| *ABHD15* | 0.420509 | 0.181969 | 2.310881 | 0.02083942 | 0.190405223 |
| *YWHAE* | 0.183046 | 0.07922 | 2.31059 | 0.02085554 | 0.190405223 |
| *DCUN1D3* | -1.330407 | 0.575852 | -2.310328 | 0.02086999 | 0.190405223 |
| *ARL6IP6* | 0.315613 | 0.136683 | 2.309091 | 0.02093855 | 0.190797073 |
| *TAX1BP1* | -0.216731 | 0.093865 | -2.308948 | 0.02094646 | 0.190797073 |
| *DYNLL1* | 0.207553 | 0.089918 | 2.308251 | 0.02098518 | 0.190935036 |
| *GPR137B* | 0.883456 | 0.382797 | 2.307898 | 0.02100482 | 0.190935036 |
| *PIM3* | -0.79338 | 0.343786 | -2.30777 | 0.02101191 | 0.190935036 |
| *PSMD9* | -0.232882 | 0.100975 | -2.306336 | 0.02109185 | 0.191315359 |
| *KLF10* | -0.580772 | 0.25182 | -2.306299 | 0.02109391 | 0.191315359 |
| *ENSG00000275964* | -0.666843 | 0.289163 | -2.306116 | 0.02110417 | 0.191315359 |
| *PPA2* | -0.296557 | 0.128786 | -2.302716 | 0.02129481 | 0.192796347 |
| *POLR2L* | -0.15613 | 0.067806 | -2.302599 | 0.02130141 | 0.192796347 |
| *PSMB6* | -0.153238 | 0.066591 | -2.301195 | 0.02138063 | 0.193359704 |
| *SPPL2B* | 0.22714 | 0.098722 | 2.300799 | 0.02140302 | 0.19337465 |
| *PSMB4* | -0.205259 | 0.089235 | -2.300207 | 0.02143648 | 0.19337465 |
| *PHF3* | 0.24518 | 0.1066 | 2.300005 | 0.02144793 | 0.19337465 |
| *MYO7B* | -1.359557 | 0.591121 | -2.299965 | 0.02145022 | 0.19337465 |
| *ENSG00000224521* | -1.706002 | 0.742631 | -2.297242 | 0.02160499 | NA |
| *PSMC3* | -0.216174 | 0.094126 | -2.296656 | 0.0216384 | 0.194912544 |
| *POLA2* | -0.704975 | 0.306996 | -2.296365 | 0.02165505 | 0.194912544 |
| *NUDT5* | -0.253367 | 0.110409 | -2.294809 | 0.02174405 | 0.195497115 |
| *TTYH3* | 0.726925 | 0.316794 | 2.29463 | 0.02175433 | 0.195497115 |
| *RER1* | -0.229109 | 0.099918 | -2.292966 | 0.02184998 | 0.1962018 |
| *CRNDE* | -1.601689 | 0.699228 | -2.290653 | 0.02198349 | NA |
| *KMT2A* | 0.213925 | 0.093393 | 2.290586 | 0.02198738 | 0.19718489 |
| *PTTG1IP* | 0.529582 | 0.231216 | 2.290427 | 0.02199661 | 0.19718489 |
| *LGMN* | 0.663175 | 0.2896 | 2.289965 | 0.02202332 | 0.19718489 |
| *AIDA* | 0.213548 | 0.093272 | 2.289532 | 0.02204846 | 0.19718489 |
| *BMI1* | 0.31702 | 0.138468 | 2.289486 | 0.0220511 | 0.19718489 |
| *WDR13* | -0.315277 | 0.137719 | -2.289275 | 0.02206337 | 0.19718489 |
| *IFIT3* | 0.695749 | 0.304213 | 2.287046 | 0.02219316 | 0.198189272 |
| *GCHFR* | -0.239236 | 0.104652 | -2.28601 | 0.0222537 | 0.198574135 |
| *STX11* | -0.447319 | 0.195744 | -2.285221 | 0.0222999 | 0.198830562 |
| *ENSG00000272277* | -0.964178 | 0.421991 | -2.284834 | 0.02232256 | 0.198876851 |
| *EAPP* | -0.199872 | 0.087489 | -2.284535 | 0.0223401 | 0.198877531 |
| *A2M.AS1* | -2.405551 | 1.054999 | -2.280145 | 0.02259909 | NA |
| *ENSG00000267121* | 0.32664 | 0.143282 | 2.279695 | 0.02262581 | 0.201263629 |
| *IGHV1.2* | 2.071262 | 0.908779 | 2.279171 | 0.02265688 | NA |
| *MARK2* | 0.393841 | 0.172852 | 2.278484 | 0.02269776 | 0.201746028 |
| *UBR1* | 0.450211 | 0.197669 | 2.277598 | 0.02275056 | 0.202057628 |
| *POLR2J3* | 0.30915 | 0.135755 | 2.277267 | 0.02277029 | 0.202075233 |
| *HFE* | -0.875356 | 0.384518 | -2.276501 | 0.02281607 | 0.202323795 |
| *ACSF2* | -0.413586 | 0.181724 | -2.275905 | 0.02285172 | 0.20248221 |

| | | | | | |
|---|---|---|---|---|---|
| *SMYD5* | -0.636399 | 0.279751 | -2.274879 | 0.0229132 | 0.202766037 |
| *MSH2* | -0.4843 | 0.2129 | -2.274776 | 0.02291937 | 0.202766037 |
| *ATP5F1B* | -0.223676 | 0.098376 | -2.273681 | 0.02298517 | 0.203055458 |
| *CKLF* | 0.350155 | 0.154025 | 2.273363 | 0.02300431 | 0.203055458 |
| *RAB33B* | 0.543319 | 0.239025 | 2.273069 | 0.02302204 | 0.203055458 |
| *ZNF74* | 1.189331 | 0.523232 | 2.273046 | 0.02302341 | 0.203055458 |
| *ENSG00000259943* | -0.582878 | 0.256588 | -2.271649 | 0.02310775 | 0.203435893 |
| *SPTY2D1* | 0.497001 | 0.218788 | 2.27161 | 0.02311008 | 0.203435893 |
| *ENSG00000276216* | -1.361777 | 0.599521 | -2.271443 | 0.02312015 | 0.203435893 |
| *RGMB* | 0.917524 | 0.404016 | 2.27101 | 0.02314638 | 0.203494225 |
| *CD74* | -0.243202 | 0.107111 | -2.27056 | 0.0231736 | 0.203494225 |
| *PDE7B* | 1.0918 | 0.480917 | 2.270247 | 0.02319257 | 0.203494225 |
| *VBP1* | -0.216407 | 0.095327 | -2.270154 | 0.02319827 | 0.203494225 |
| *PODXL2* | -0.914576 | 0.403001 | -2.269415 | 0.02324308 | 0.203711648 |
| *SMARCA4* | -0.330171 | 0.145504 | -2.269156 | 0.02325884 | 0.203711648 |
| *PGD* | -0.281383 | 0.124047 | -2.26835 | 0.02330785 | 0.203984035 |
| *GARS* | -0.365817 | 0.161326 | -2.267565 | 0.02335572 | 0.204245904 |
| *TLCD1* | -1.652176 | 0.728694 | -2.267312 | 0.02337116 | NA |
| *GPRIN3* | 1.21321 | 0.53527 | 2.26654 | 0.02341836 | 0.204636542 |
| *ANAPC15* | -0.295997 | 0.130735 | -2.26411 | 0.02356736 | 0.205780636 |
| *HSH2D* | -0.246179 | 0.10882 | -2.262256 | 0.02368162 | 0.206619843 |
| *RPA3* | 0.187696 | 0.082992 | 2.261598 | 0.02372224 | 0.206815761 |
| *PSMC5* | -0.230315 | 0.101877 | -2.260729 | 0.02377607 | 0.20690735 |
| *ZNF70* | 1.02847 | 0.45493 | 2.26072 | 0.02377659 | 0.20690735 |
| *PSMB2* | -0.244604 | 0.108205 | -2.260548 | 0.02378726 | 0.20690735 |
| *TM2D3* | 0.303671 | 0.134371 | 2.259955 | 0.02382406 | 0.20706923 |
| *GNG11* | 1.161354 | 0.51413 | 2.258874 | 0.02389123 | 0.207391852 |
| *FADS1* | -0.659297 | 0.291883 | -2.258771 | 0.0238976 | 0.207391852 |
| *APBB1* | 0.861901 | 0.381663 | 2.258276 | 0.02392844 | 0.207501291 |
| *DENND6B* | -0.427136 | 0.189216 | -2.257397 | 0.02398325 | 0.207818363 |
| *SPATA20* | 0.875647 | 0.387999 | 2.25683 | 0.0240187 | 0.207967244 |
| *SLC25A5* | -0.165817 | 0.0735 | -2.256008 | 0.0240701 | 0.208207282 |
| *PTPN2* | 0.247248 | 0.109605 | 2.255803 | 0.024083 | 0.208207282 |
| *TYW3* | 0.349464 | 0.154982 | 2.254865 | 0.02414179 | 0.208451352 |
| *FAM234A* | -0.421327 | 0.18686 | -2.254769 | 0.02414784 | 0.208451352 |
| *POLE2* | 1.198928 | 0.532018 | 2.253549 | 0.02422456 | 0.208955203 |
| *PHKG2* | -0.332109 | 0.147401 | -2.253106 | 0.02425247 | 0.209037584 |
| *SOD1* | -0.137932 | 0.061231 | -2.252628 | 0.02428263 | 0.209139214 |
| *BANP* | 0.336349 | 0.14935 | 2.252088 | 0.02431671 | 0.209274443 |
| *COPS2* | -0.304603 | 0.135304 | -2.251251 | 0.02436962 | 0.209571382 |
| *FCGBP* | -1.362771 | 0.60572 | -2.249838 | 0.02445922 | 0.209653344 |
| *SP110* | 0.176791 | 0.078589 | 2.249552 | 0.02447742 | 0.209653344 |
| *LACTB2.AS1* | -0.83193 | 0.369832 | -2.249479 | 0.02448204 | 0.209653344 |
| *ITPR2* | 0.377368 | 0.16777 | 2.249308 | 0.02449288 | 0.209653344 |
| *NAGK* | 0.276758 | 0.123076 | 2.248666 | 0.02453377 | 0.209653344 |
| *STK38L* | -0.483356 | 0.214958 | -2.248602 | 0.02453782 | 0.209653344 |
| *TNIP2* | -0.317087 | 0.141019 | -2.248546 | 0.02454137 | 0.209653344 |

| | | | | | |
|---|---|---|---|---|---|
| *ZFP36L2* | 0.385855 | 0.171604 | 2.248514 | 0.02454341 | 0.209653344 |
| *NSUN5* | -0.219434 | 0.097591 | -2.248491 | 0.02454487 | 0.209653344 |
| *ENSG00000279278* | 0.430342 | 0.19144 | 2.247927 | 0.02458087 | 0.209654966 |
| *CYTH2* | 0.331784 | 0.14761 | 2.247702 | 0.0245952 | 0.209654966 |
| *TMEM170A* | 0.271363 | 0.120745 | 2.247417 | 0.0246134 | 0.209654966 |
| *GCFC2* | -0.474439 | 0.211112 | -2.247334 | 0.02461872 | 0.209654966 |
| *CYB5R4* | -0.223178 | 0.099366 | -2.246032 | 0.02470198 | 0.210163531 |
| *WNT16* | 1.250037 | 0.556605 | 2.245823 | 0.02471535 | 0.210163531 |
| *PWWP2A* | 0.422079 | 0.188041 | 2.244611 | 0.02479314 | 0.210667694 |
| *ENSG00000272758* | 0.780125 | 0.347687 | 2.243758 | 0.02484795 | 0.210975974 |
| *MGLL* | -1.073254 | 0.478434 | -2.243264 | 0.02487977 | 0.211088737 |
| *ZC3H12A* | -0.362815 | 0.161863 | -2.241492 | 0.0249942 | 0.211730606 |
| *EMP3* | -0.174021 | 0.077643 | -2.241303 | 0.02500643 | 0.211730606 |
| *TRAF2* | -0.370818 | 0.165453 | -2.24123 | 0.02501121 | 0.211730606 |
| *ZNF528* | 0.64461 | 0.287665 | 2.240839 | 0.02503651 | 0.211787294 |
| *PARP15* | 0.403233 | 0.179997 | 2.240216 | 0.02507693 | 0.211964379 |
| *TRAF3* | -0.334428 | 0.149302 | -2.239942 | 0.02509468 | 0.211964379 |
| *MFGE8* | -0.752129 | 0.335896 | -2.239172 | 0.02514471 | 0.212229523 |
| *NASP* | 0.237224 | 0.105957 | 2.238861 | 0.02516493 | 0.212242914 |
| *USP10* | 0.302694 | 0.135221 | 2.23851 | 0.02518782 | 0.21227868 |
| *OLA1* | -0.217316 | 0.097095 | -2.238179 | 0.02520938 | 0.212303224 |
| *ENTPD1.AS1* | 0.376512 | 0.168252 | 2.237783 | 0.02523524 | 0.21236393 |
| *ENSG00000255856* | -1.807333 | 0.80839 | -2.23572 | 0.02537009 | NA |
| *TOP2A* | -1.299239 | 0.58125 | -2.235251 | 0.02540086 | 0.213465257 |
| *ZNF677* | 0.586399 | 0.262346 | 2.235209 | 0.0254036 | 0.213465257 |
| *RMI1* | 0.76999 | 0.344595 | 2.23448 | 0.02545152 | 0.21355142 |
| *NPC2* | 0.29162 | 0.130519 | 2.23432 | 0.02546201 | 0.21355142 |
| *POU2F1* | 0.378774 | 0.169535 | 2.234197 | 0.02547012 | 0.21355142 |
| *DAGLB* | 0.666695 | 0.298449 | 2.233868 | 0.02549176 | 0.213575551 |
| *HEXB* | 0.28376 | 0.12705 | 2.233455 | 0.02551899 | 0.213646442 |
| *SUGP2* | 0.285073 | 0.127751 | 2.231483 | 0.02564916 | 0.214421653 |
| *ENSG00000268439* | 1.462558 | 0.655429 | 2.231453 | 0.02565114 | 0.214421653 |
| *RFX1* | 0.349659 | 0.156714 | 2.231197 | 0.02566808 | 0.214421653 |
| *MIR762HG* | -0.621393 | 0.278625 | -2.230211 | 0.02573345 | 0.214810141 |
| *ADGRB2* | -1.867646 | 0.837677 | -2.229553 | 0.02577712 | NA |
| *LSP1* | -0.139924 | 0.062766 | -2.2293 | 0.02579394 | 0.215017994 |
| *SLC35A3* | 0.475366 | 0.213253 | 2.229121 | 0.02580589 | 0.215017994 |
| *BCCIP* | -0.304763 | 0.136727 | -2.228984 | 0.025815 | 0.215017994 |
| *ZBTB44* | 0.234083 | 0.105051 | 2.228282 | 0.0258617 | 0.215103594 |
| *ARL6IP4* | -0.152667 | 0.068514 | -2.228262 | 0.02586306 | 0.215103594 |
| *S100A4* | -0.296967 | 0.133322 | -2.227436 | 0.02591815 | 0.215374119 |
| *SUPV3L1* | -0.43627 | 0.195882 | -2.227207 | 0.02593342 | 0.215374119 |
| *SNX18* | 0.614176 | 0.275846 | 2.226516 | 0.02597965 | 0.215600786 |
| *CRLF3* | 0.244343 | 0.109795 | 2.225459 | 0.02605044 | 0.21603081 |
| *MIS18A* | 0.45893 | 0.206449 | 2.222974 | 0.02621752 | 0.21725817 |
| *SHROOM3* | -2.338417 | 1.052335 | -2.222121 | 0.0262751 | NA |
| *AEBP2* | 0.382027 | 0.17194 | 2.221867 | 0.02629229 | 0.21763259 |

| | | | | | |
|---|---|---|---|---|---|
| *MICU2* | -0.34956 | 0.157336 | -2.221739 | 0.02630094 | 0.21763259 |
| *CD2BP2* | -0.205576 | 0.092563 | -2.220931 | 0.02635564 | 0.217926877 |
| *OAS2* | 0.517154 | 0.232892 | 2.220571 | 0.02638 | 0.217970004 |
| *DHRS4.AS1* | 0.456606 | 0.205667 | 2.220119 | 0.02641068 | 0.21806529 |
| *PRRG4* | 2.478839 | 1.116834 | 2.219524 | 0.02645112 | NA |
| *DNAAF4* | -0.837609 | 0.377467 | -2.219023 | 0.02648516 | 0.218521738 |
| *INSIG2* | 0.36663 | 0.165243 | 2.218734 | 0.02650485 | 0.218525848 |
| *MINDY1* | 1.326998 | 0.598437 | 2.217439 | 0.0265931 | 0.218906033 |
| *C19orf48* | 0.409683 | 0.184762 | 2.217357 | 0.02659872 | 0.218906033 |
| *TSC22D2* | 0.422742 | 0.190664 | 2.217212 | 0.02660864 | 0.218906033 |
| *LINC01857* | 0.811925 | 0.366487 | 2.215426 | 0.02673084 | 0.219752576 |
| *SCAMP2* | -0.217361 | 0.098153 | -2.214521 | 0.02679298 | 0.220104556 |
| *TLK2* | 0.471465 | 0.212928 | 2.214201 | 0.02681494 | 0.220126076 |
| *ICA1* | -1.305978 | 0.589942 | -2.213738 | 0.02684677 | NA |
| *PPCDC* | -0.439927 | 0.198815 | -2.212745 | 0.02691526 | 0.220790424 |
| *SFR1* | -0.345329 | 0.156142 | -2.211635 | 0.02699192 | 0.221259882 |
| *ENSG00000272669* | 1.634385 | 0.739218 | 2.210967 | 0.02703815 | NA |
| *RN7SL832P* | 1.648246 | 0.74566 | 2.210453 | 0.02707372 | NA |
| *SLC25A19* | 0.383003 | 0.173271 | 2.210429 | 0.02707542 | 0.221784689 |
| *BCL2* | -0.305653 | 0.138325 | -2.20968 | 0.0271274 | 0.222050765 |
| *CYHR1* | 0.350716 | 0.15877 | 2.208949 | 0.02717819 | 0.222306668 |
| *PTGES2* | -0.3482 | 0.157663 | -2.208509 | 0.02720881 | 0.222397322 |
| *HPRT1* | -0.294771 | 0.133504 | -2.207948 | 0.02724789 | 0.222557033 |
| *RAB1A* | 0.259573 | 0.117698 | 2.20542 | 0.0274246 | 0.223783587 |
| *MAP3K14* | 0.416541 | 0.188887 | 2.205238 | 0.02743737 | 0.223783587 |
| *ZNF284* | 0.816042 | 0.370137 | 2.2047 | 0.02747519 | 0.223790224 |
| *TMUB1* | -0.230523 | 0.104562 | -2.204667 | 0.02747749 | 0.223790224 |
| *TNFSF12* | 0.32817 | 0.148951 | 2.203215 | 0.02757959 | 0.224184409 |
| *HMOX1* | 0.412472 | 0.187222 | 2.203122 | 0.02758613 | 0.224184409 |
| *SHPRH* | 0.348331 | 0.158126 | 2.202875 | 0.02760353 | 0.224184409 |
| *ING2* | 0.459429 | 0.20856 | 2.20286 | 0.02760465 | 0.224184409 |
| *CPM* | 1.007934 | 0.457682 | 2.202257 | 0.02764717 | 0.224369696 |
| *UBXN11* | -0.501471 | 0.227772 | -2.201636 | 0.02769102 | 0.224565479 |
| *PDIA3* | -0.215986 | 0.098118 | -2.201284 | 0.02771594 | 0.224607645 |
| *EBLN3P* | 0.24689 | 0.112185 | 2.200726 | 0.0277554 | 0.224767417 |
| *TNFRSF13C* | -0.165249 | 0.07512 | -2.199809 | 0.02782048 | 0.225041641 |
| *CDK14* | -0.480868 | 0.218607 | -2.199691 | 0.02782881 | 0.225041641 |
| *RERE* | 0.376818 | 0.171335 | 2.199306 | 0.02785618 | 0.225041641 |
| *N4BP2L2* | 0.130936 | 0.05954 | 2.199135 | 0.02786832 | 0.225041641 |
| *ZFAND5* | 0.303727 | 0.138172 | 2.198177 | 0.02793647 | 0.225104968 |
| *PELP1* | 0.469655 | 0.213696 | 2.197771 | 0.02796546 | 0.225104968 |
| *SETBP1* | 0.356051 | 0.162026 | 2.197491 | 0.02798537 | 0.225104968 |
| *RRBP1* | 0.569926 | 0.259367 | 2.197375 | 0.02799369 | 0.225104968 |
| *PPP1CA* | -0.157882 | 0.07185 | -2.197371 | 0.02799396 | 0.225104968 |
| *NR4A2* | 0.784038 | 0.356809 | 2.197359 | 0.02799479 | 0.225104968 |
| *STK38* | -0.300485 | 0.136799 | -2.196548 | 0.02805277 | 0.22525649 |
| *TRMT6* | -0.415359 | 0.189115 | -2.196328 | 0.02806849 | 0.22525649 |

| | | | | | |
|---|---|---|---|---|---|
| *TXNIP* | -0.283634 | 0.129144 | -2.196256 | 0.02807364 | 0.22525649 |
| *ZNF439* | 0.255371 | 0.11629 | 2.195989 | 0.02809276 | 0.22525649 |
| *LINC00638* | 1.186105 | 0.540373 | 2.194975 | 0.02816537 | 0.225348133 |
| *THEM4* | -0.368639 | 0.167951 | -2.194923 | 0.02816915 | 0.225348133 |
| *HNRNPA1L2* | -0.37756 | 0.172022 | -2.19484 | 0.02817505 | 0.225348133 |
| *ENSG00000230709* | 0.551812 | 0.251427 | 2.194725 | 0.02818336 | 0.225348133 |
| *CCL5* | -0.683932 | 0.311943 | -2.192488 | 0.02834427 | 0.226475658 |
| *CLIP1* | 0.293829 | 0.13405 | 2.191935 | 0.0283842 | 0.226635716 |
| *CYSLTR1* | -0.389707 | 0.177893 | -2.190678 | 0.02847509 | 0.227202074 |
| *LENG8* | -0.232388 | 0.106124 | -2.189781 | 0.02854013 | 0.227561535 |
| *ENSG00000277369* | -1.376315 | 0.628656 | -2.189296 | 0.02857536 | NA |
| *LIX1L* | 0.350239 | 0.159987 | 2.18917 | 0.02858451 | 0.227755943 |
| *MALAT1* | 0.143718 | 0.0657 | 2.187496 | 0.02870637 | 0.228566943 |
| *ENSG00000263470* | -2.087438 | 0.954293 | -2.187419 | 0.02871197 | NA |
| *SPIN2B* | 0.514065 | 0.2351 | 2.186579 | 0.02877327 | 0.228910328 |
| *APOL3* | -0.367487 | 0.168082 | -2.186354 | 0.02878971 | 0.228910328 |
| *RANBP9* | -0.446245 | 0.20416 | -2.185761 | 0.02883309 | 0.229040496 |
| *UBAP2L* | 0.328844 | 0.150472 | 2.185419 | 0.02885815 | 0.229040496 |
| *CRIP2* | -1.052476 | 0.481615 | -2.185306 | 0.02886643 | 0.229040496 |
| *CAST* | -0.215801 | 0.098773 | -2.184814 | 0.0289025 | 0.229098041 |
| *PARP11* | 0.492646 | 0.225503 | 2.184658 | 0.02891392 | 0.229098041 |
| *XPO4* | -0.393963 | 0.18043 | -2.183464 | 0.02900164 | 0.229633249 |
| *HELZ2* | 0.523949 | 0.240016 | 2.182974 | 0.02903774 | 0.229759355 |
| *CHRNA6* | -1.558914 | 0.714684 | -2.181263 | 0.02916397 | NA |
| *IL2RA* | -0.601376 | 0.275787 | -2.180581 | 0.02921444 | 0.23099693 |
| *NUDT12* | 1.398805 | 0.641585 | 2.180235 | 0.02924004 | 0.231038906 |
| *ADGRE2* | 1.505991 | 0.691151 | 2.17896 | 0.02933466 | 0.231494384 |
| *UFC1* | -0.15603 | 0.071609 | -2.17891 | 0.02933835 | 0.231494384 |
| *ENSG00000273253* | 0.617049 | 0.283295 | 2.178119 | 0.02939719 | 0.231798088 |
| *APBB1IP* | 0.235163 | 0.107987 | 2.177692 | 0.02942897 | 0.23187823 |
| *UBE2T* | 0.535619 | 0.245986 | 2.177436 | 0.02944809 | 0.23187823 |
| *KCNA3* | -0.859755 | 0.394916 | -2.177055 | 0.02947643 | 0.231941031 |
| *IFIT1* | 1.804023 | 0.828771 | 2.176744 | 0.02949968 | 0.231963626 |
| *CALHM6* | 0.404419 | 0.185874 | 2.175768 | 0.02957264 | 0.23237684 |
| *NIFK.AS1* | 0.425726 | 0.195797 | 2.174328 | 0.02968048 | 0.233063371 |
| *NCBP2AS2* | -0.267557 | 0.123116 | -2.173203 | 0.02976508 | 0.233566604 |
| *GXYLT1* | 0.454379 | 0.209126 | 2.172751 | 0.02979907 | 0.233672322 |
| *TMEM189* | 0.44266 | 0.203904 | 2.170921 | 0.02993718 | 0.234403832 |
| *TEX264* | -0.254463 | 0.117247 | -2.170316 | 0.02998293 | 0.234403832 |
| *ZNF780A* | 0.403257 | 0.185806 | 2.170312 | 0.02998323 | 0.234403832 |
| *TBCD* | 0.64583 | 0.29759 | 2.170202 | 0.02999156 | 0.234403832 |
| *ENSG00000273117* | -1.182932 | 0.545092 | -2.170153 | 0.02999529 | 0.234403832 |
| *UBE2V2* | -0.19763 | 0.0911 | -2.16938 | 0.03005383 | 0.234700187 |
| *CLDND2* | 0.712202 | 0.32871 | 2.166658 | 0.03026096 | 0.236155783 |
| *CARS* | -0.33331 | 0.153921 | -2.16546 | 0.03035246 | 0.236707571 |
| *MICB* | 0.522348 | 0.241264 | 2.165048 | 0.03038403 | 0.236736849 |
| *PSMD4* | -0.178976 | 0.082673 | -2.164868 | 0.03039779 | 0.236736849 |

| | | | | | |
|---|---|---|---|---|---|
| *ENSG00000267939* | -1.822026 | 0.842196 | -2.163423 | 0.03050865 | NA |
| *ANKRD10* | -0.326094 | 0.150799 | -2.162435 | 0.03058465 | 0.237984591 |
| *ENOX2* | -0.420838 | 0.194631 | -2.162238 | 0.03059981 | 0.237984591 |
| *SNX29* | -0.275376 | 0.127421 | -2.161149 | 0.03068386 | 0.238475362 |
| *SNU13* | -0.130817 | 0.06055 | -2.16046 | 0.03073712 | 0.23868825 |
| *USP34* | 0.188619 | 0.087313 | 2.160252 | 0.03075318 | 0.23868825 |
| *CCDC144A* | 0.924674 | 0.428391 | 2.158481 | 0.03089047 | 0.239590559 |
| *CD59* | -0.310102 | 0.143704 | -2.157922 | 0.03093388 | 0.239763906 |
| *TRERF1* | -0.885593 | 0.410495 | -2.157381 | 0.03097597 | 0.239926797 |
| *ACAT2* | -0.266527 | 0.123628 | -2.155879 | 0.03109309 | 0.240670198 |
| *COPS7B* | -0.258643 | 0.120009 | -2.155186 | 0.03114725 | 0.240925694 |
| *PSMA3* | -0.201136 | 0.093343 | -2.154797 | 0.03117769 | 0.240997397 |
| *COX4I1* | -0.116744 | 0.054206 | -2.153699 | 0.03126381 | 0.241244245 |
| *CDC37L1* | 0.386088 | 0.179271 | 2.153655 | 0.03126723 | 0.241244245 |
| *NAGPA.AS1* | 1.785399 | 0.829207 | 2.153139 | 0.03130776 | NA |
| *EDEM1* | -0.274988 | 0.127718 | -2.153093 | 0.03131141 | 0.241244245 |
| *KLHL26* | -0.860925 | 0.399955 | -2.152556 | 0.03135359 | 0.241244245 |
| *DOCK7* | 1.09096 | 0.506852 | 2.152421 | 0.03136424 | 0.241244245 |
| *BCL6* | 0.538152 | 0.250089 | 2.151843 | 0.03140973 | 0.241244245 |
| *ZNF518A* | 0.257354 | 0.119599 | 2.151819 | 0.0314116 | 0.241244245 |
| *HTRA2* | -0.295581 | 0.137363 | -2.151818 | 0.03141166 | 0.241244245 |
| *ING3* | -0.308914 | 0.143566 | -2.151714 | 0.03141987 | 0.241244245 |
| *ZNF296* | -0.316787 | 0.147261 | -2.1512 | 0.03146039 | 0.241244245 |
| *TRAPPC8* | 0.396644 | 0.18439 | 2.151118 | 0.03146686 | 0.241244245 |
| *NUAK2* | 0.275202 | 0.127947 | 2.150907 | 0.03148354 | 0.241244245 |
| *PAQR7* | -0.693332 | 0.322348 | -2.150878 | 0.0314858 | 0.241244245 |
| *ELP6* | -0.295612 | 0.137454 | -2.150619 | 0.03150625 | 0.241244245 |
| *ZNF516* | 0.991761 | 0.461264 | 2.150094 | 0.03154776 | 0.241399715 |
| *IARS* | -0.261243 | 0.121539 | -2.149456 | 0.03159824 | 0.241623625 |
| *PDLIM1* | -0.138065 | 0.06425 | -2.148855 | 0.03164593 | 0.241812066 |
| *PIP4K2A* | -0.231342 | 0.107671 | -2.14861 | 0.03166536 | 0.241812066 |
| *NFATC2* | 0.397581 | 0.185152 | 2.147321 | 0.03176773 | 0.242431202 |
| *PM20D2* | 0.519012 | 0.241842 | 2.146079 | 0.0318667 | 0.242981161 |
| *SIDT1.AS1* | -0.567314 | 0.264374 | -2.145881 | 0.03188247 | 0.242981161 |
| *COPZ1* | -0.21088 | 0.09833 | -2.144613 | 0.03198379 | 0.243590233 |
| *SNHG30* | -0.292229 | 0.13632 | -2.143696 | 0.03205722 | 0.243986266 |
| *FHOD1* | 0.499112 | 0.232931 | 2.142742 | 0.03213378 | 0.244405637 |
| *CCDC112* | 0.377288 | 0.1761 | 2.142458 | 0.03215668 | 0.244416527 |
| *HEBP2* | -0.501527 | 0.234167 | -2.141751 | 0.03221354 | 0.244685376 |
| *C12orf42* | 0.573585 | 0.267867 | 2.141308 | 0.03224919 | 0.244792851 |
| *RSU1* | -0.263633 | 0.12316 | -2.140579 | 0.03230797 | 0.244913898 |
| *GFPT1* | 0.524898 | 0.245214 | 2.140577 | 0.03230816 | 0.244913898 |
| *MRPL45* | 0.26962 | 0.125989 | 2.140024 | 0.03235283 | 0.245089375 |
| *GYPC* | 0.291776 | 0.136408 | 2.138995 | 0.03243609 | 0.245556749 |
| *RETREG2* | 0.247172 | 0.115573 | 2.138666 | 0.03246273 | 0.245595096 |
| *RNF146* | 0.323124 | 0.151168 | 2.137521 | 0.03255564 | 0.246134452 |
| *IL6ST* | 0.879379 | 0.411517 | 2.136918 | 0.0326047 | 0.246182787 |

| | | | | | |
|---|---|---|---|---|---|
| *ENSG00000276728* | 0.542341 | 0.253828 | 2.136648 | 0.03262665 | 0.246182787 |
| *NHSL2* | -0.329325 | 0.154146 | -2.136445 | 0.03264314 | 0.246182787 |
| *NDUFV2* | 0.16864 | 0.078937 | 2.136379 | 0.03264852 | 0.246182787 |
| *ZFYVE16* | 0.405641 | 0.190138 | 2.133406 | 0.0328914 | 0.247850113 |
| *TERF1* | 0.248214 | 0.116419 | 2.13208 | 0.03300025 | 0.248368679 |
| *MIR29B2CHG* | 0.375195 | 0.17598 | 2.132036 | 0.03300385 | 0.248368679 |
| *UTP14C* | 0.634967 | 0.297945 | 2.131153 | 0.03307654 | 0.248751315 |
| *ARL5A* | 0.186054 | 0.087358 | 2.129777 | 0.03319003 | 0.249440055 |
| *ANKRD40* | -0.335545 | 0.157653 | -2.128376 | 0.03330595 | 0.250146164 |
| *ITGA6.AS1* | -0.645551 | 0.303626 | -2.126141 | 0.03349155 | 0.251171464 |
| *ZNF232* | 0.643168 | 0.302523 | 2.12601 | 0.03350242 | 0.251171464 |
| *DEGS2* | 1.476816 | 0.694666 | 2.125935 | 0.03350865 | 0.251171464 |
| *NDUFAB1* | -0.201612 | 0.094886 | -2.124777 | 0.03360527 | 0.2514239 |
| *CCT7* | -0.186237 | 0.087651 | -2.12475 | 0.03360752 | 0.2514239 |
| *OSBPL9* | -0.283322 | 0.133345 | -2.124737 | 0.03360857 | 0.2514239 |
| *ALOX5* | -0.276892 | 0.130368 | -2.123937 | 0.03367539 | 0.251758325 |
| *FTH1* | 0.247894 | 0.116764 | 2.12303 | 0.03375135 | 0.251996807 |
| *PLEKHA1* | 0.402688 | 0.189696 | 2.122803 | 0.03377034 | 0.251996807 |
| *FMO5* | 0.830732 | 0.391344 | 2.122763 | 0.03377368 | 0.251996807 |
| *PBXIP1* | -0.258336 | 0.121717 | -2.122435 | 0.03380119 | 0.25203689 |
| *MTHFD2* | -0.413098 | 0.194688 | -2.121844 | 0.03385081 | 0.252241704 |
| *IGF1R* | 1.420201 | 0.669974 | 2.119786 | 0.03402413 | 0.253027663 |
| *MEF2A* | 0.339319 | 0.160118 | 2.119182 | 0.03407508 | 0.253027663 |
| *FAM98C* | 0.313264 | 0.147842 | 2.118915 | 0.03409762 | 0.253027663 |
| *TRIM35* | 0.839711 | 0.396298 | 2.118888 | 0.03409996 | 0.253027663 |
| *ENSG00000270562* | 1.15662 | 0.545944 | 2.118567 | 0.03412707 | 0.253027663 |
| *OTUB1* | -0.195456 | 0.092261 | -2.118509 | 0.03413195 | 0.253027663 |
| *TMEM237* | 1.674042 | 0.790212 | 2.118473 | 0.03413506 | NA |
| *ARHGAP5* | 0.568218 | 0.268238 | 2.118336 | 0.03414663 | 0.253027663 |
| *CD200* | 0.770205 | 0.36359 | 2.118331 | 0.03414708 | 0.253027663 |
| *POLR3D* | 0.421808 | 0.199139 | 2.118161 | 0.03416146 | 0.253027663 |
| *SCARB1* | -1.060393 | 0.500694 | -2.117845 | 0.03418823 | 0.253027663 |
| *MRPL3* | -0.27013 | 0.127558 | -2.117697 | 0.03420074 | 0.253027663 |
| *TAMM41* | 0.588141 | 0.277873 | 2.11658 | 0.03429554 | 0.253203433 |
| *PIP5K1B* | 0.610286 | 0.288338 | 2.116563 | 0.03429697 | 0.253203433 |
| *PPP6C* | -0.219526 | 0.103723 | -2.116474 | 0.03430452 | 0.253203433 |
| *GTF2H5* | -0.24674 | 0.116596 | -2.116203 | 0.03432758 | 0.253203433 |
| *BLVRA* | -0.432603 | 0.204433 | -2.116107 | 0.03433568 | 0.253203433 |
| *SET* | -0.152402 | 0.072036 | -2.115627 | 0.03437653 | 0.25321578 |
| *SDHB* | -0.185193 | 0.087538 | -2.115565 | 0.03438184 | 0.25321578 |
| *FAM216A* | 0.44976 | 0.212664 | 2.114883 | 0.03443991 | 0.25331665 |
| *NAGS* | 1.237353 | 0.585111 | 2.114733 | 0.03445273 | 0.25331665 |
| *TMEM106A* | 0.335589 | 0.158699 | 2.114621 | 0.03446228 | 0.25331665 |
| *MANBA* | 0.337927 | 0.159831 | 2.11428 | 0.03449137 | 0.253366902 |
| *LARP7* | -0.200502 | 0.094889 | -2.113023 | 0.03459884 | 0.253992488 |
| *KCNN1* | 1.698386 | 0.803881 | 2.112732 | 0.03462368 | NA |
| *VAPA* | -0.154568 | 0.073187 | -2.111961 | 0.03468976 | 0.254495905 |

| | | | | | |
|---|---|---|---|---|---|
| *AHI1* | 0.351184 | 0.166322 | 2.111464 | 0.03473248 | 0.254645212 |
| *ENSG00000272871* | 0.937503 | 0.444131 | 2.110871 | 0.03478337 | 0.254854241 |
| *LINC01250* | -3.131219 | 1.483587 | -2.110574 | 0.03480898 | NA |
| *MDS2* | -1.182526 | 0.560301 | -2.110519 | 0.03481372 | 0.254912512 |
| *CEP170* | 0.312034 | 0.147869 | 2.110207 | 0.03484052 | 0.254944843 |
| *SH3RF1* | -0.83762 | 0.397012 | -2.109808 | 0.03487489 | 0.255032449 |
| *MED28* | 0.191446 | 0.09076 | 2.109364 | 0.0349132 | 0.255148708 |
| *RNASEH2C* | -0.138185 | 0.065548 | -2.108163 | 0.03501689 | 0.255742373 |
| *FAM133B* | 0.149338 | 0.070861 | 2.107468 | 0.035077 | 0.256017148 |
| *ENSG00000261770* | 1.289838 | 0.612316 | 2.10649 | 0.03516179 | 0.256271086 |
| *LRRC7* | -0.763264 | 0.362352 | -2.106413 | 0.03516848 | 0.256271086 |
| *UTP4* | -0.411346 | 0.195294 | -2.106288 | 0.03517932 | 0.256271086 |
| *RSL1D1* | -0.155223 | 0.073713 | -2.105788 | 0.0352228 | 0.256423759 |
| *KLHL34* | 1.657041 | 0.787324 | 2.104649 | 0.03532184 | NA |
| *ASAH1* | 0.355673 | 0.169035 | 2.104142 | 0.03536607 | 0.257302284 |
| *ENSG00000271746* | -1.427301 | 0.678623 | -2.103231 | 0.03544557 | NA |
| *MED8* | -0.269606 | 0.128214 | -2.102778 | 0.03548518 | 0.257879742 |
| *DPH5* | 0.28164 | 0.133941 | 2.102714 | 0.03549074 | 0.257879742 |
| *ZNF485* | 0.970984 | 0.46192 | 2.102063 | 0.03554778 | 0.258129471 |
| *LPAR2* | -0.589229 | 0.2804 | -2.101387 | 0.03560699 | 0.258394656 |
| *B3GALNT2* | 0.515933 | 0.245696 | 2.09988 | 0.0357394 | 0.259100316 |
| *PEX16* | -0.225549 | 0.107438 | -2.099346 | 0.03578638 | 0.259100316 |
| *TACO1* | -0.353594 | 0.168433 | -2.099309 | 0.03578966 | 0.259100316 |
| *DMAP1* | -0.287324 | 0.13687 | -2.099246 | 0.03579526 | 0.259100316 |
| *BOLA1* | 0.349672 | 0.166595 | 2.098941 | 0.03582211 | 0.259129967 |
| *INO80B* | 0.417098 | 0.198746 | 2.098649 | 0.03584786 | 0.25915155 |
| *FOXP4* | 0.663615 | 0.316394 | 2.097435 | 0.03595505 | 0.259761555 |
| *AHR* | -0.574787 | 0.27413 | -2.096767 | 0.03601419 | 0.259955273 |
| *NDUFV3* | 0.264824 | 0.12631 | 2.096617 | 0.03602753 | 0.259955273 |
| *DUT* | -0.146186 | 0.069759 | -2.095586 | 0.03611892 | 0.260374241 |
| *MRPS35* | -0.274286 | 0.130896 | -2.095446 | 0.03613133 | 0.260374241 |
| *DNAJC3.DT* | 0.956413 | 0.456522 | 2.094997 | 0.03617128 | 0.260497295 |
| *ZNF385A* | 0.940442 | 0.44904 | 2.094341 | 0.03622965 | 0.260752724 |
| *KDM2B* | -0.247263 | 0.118128 | -2.093167 | 0.03633428 | 0.261246345 |
| *PFDN2* | 0.166002 | 0.079311 | 2.093056 | 0.03634413 | 0.261246345 |
| *MTAP* | 0.478886 | 0.228846 | 2.092618 | 0.03638329 | 0.261362863 |
| *VAMP3* | -0.292446 | 0.139824 | -2.091526 | 0.03648097 | 0.261826983 |
| *AZIN1* | -0.243781 | 0.116565 | -2.091381 | 0.03649389 | 0.261826983 |
| *CSDE1* | -0.142548 | 0.068195 | -2.090312 | 0.03658975 | 0.262331591 |
| *NAV2* | -1.135472 | 0.543266 | -2.090083 | 0.0366103 | 0.262331591 |
| *MAVS* | 0.307204 | 0.147092 | 2.08851 | 0.03675189 | 0.263180481 |
| *PDPR* | -0.616332 | 0.295193 | -2.087897 | 0.03680713 | 0.263407653 |
| *TAF9* | -0.199268 | 0.095451 | -2.087645 | 0.03682988 | 0.263407653 |
| *RARA.AS1* | 0.703981 | 0.337316 | 2.087005 | 0.03688764 | 0.263511664 |
| *ENSG00000254531* | 0.485694 | 0.232727 | 2.086972 | 0.03689071 | 0.263511664 |
| *SNX10* | -0.296366 | 0.142033 | -2.086604 | 0.03692391 | 0.26351546 |
| *RACGAP1* | -0.745042 | 0.357122 | -2.086241 | 0.03695682 | 0.26351546 |

| | | | | | |
|---|---|---|---|---|---|
| *BLOC1S3* | 0.414578 | 0.198724 | 2.086198 | 0.03696067 | 0.26351546 |
| *RARA* | -0.273034 | 0.130903 | -2.085774 | 0.0369991 | 0.263624378 |
| *METTL23* | -0.167994 | 0.080571 | -2.085052 | 0.03706461 | 0.263925998 |
| *FCER2* | -0.763355 | 0.366157 | -2.084776 | 0.03708967 | 0.263939382 |
| *SNHG25* | -0.33541 | 0.160937 | -2.084104 | 0.03715074 | 0.264208849 |
| *WWP1* | 0.511222 | 0.245403 | 2.083197 | 0.03723331 | 0.264630725 |
| *POR* | -0.388654 | 0.186719 | -2.081487 | 0.03738936 | 0.265540092 |
| *CLNS1A* | 0.283795 | 0.136366 | 2.081118 | 0.0374231 | 0.265540092 |
| *ALDH6A1* | 0.502928 | 0.241698 | 2.080813 | 0.03745101 | 0.265540092 |
| *LZTFL1* | 0.651398 | 0.313069 | 2.080686 | 0.03746261 | 0.265540092 |
| *ENSG00000272189* | 1.001026 | 0.481145 | 2.080507 | 0.03747905 | 0.265540092 |
| *FBXL12* | -0.370068 | 0.177895 | -2.080266 | 0.03750118 | 0.265540092 |
| *FAM177A1* | -0.212141 | 0.102012 | -2.079564 | 0.03756556 | 0.265718383 |
| *SCAMP3* | -0.285409 | 0.13725 | -2.079482 | 0.03757304 | 0.265718383 |
| *SRA1* | -0.242225 | 0.116516 | -2.078898 | 0.03762673 | 0.265932917 |
| *SLTM* | 0.158493 | 0.076262 | 2.078265 | 0.03768497 | 0.266179286 |
| *DERL1* | -0.246153 | 0.118486 | -2.077483 | 0.03775705 | 0.266459882 |
| *DCK* | -0.144862 | 0.069746 | -2.077001 | 0.03780146 | 0.266459882 |
| *ARF4* | -0.298756 | 0.143849 | -2.076869 | 0.03781364 | 0.266459882 |
| *PCK2* | -0.424394 | 0.204348 | -2.076819 | 0.0378183 | 0.266459882 |
| *ENSG00000228434* | 0.382708 | 0.184312 | 2.076411 | 0.03785593 | 0.266560063 |
| *PATL2* | 0.364449 | 0.175623 | 2.075173 | 0.03797048 | 0.267201432 |
| *CNKSR3* | -1.158748 | 0.558542 | -2.074594 | 0.03802418 | 0.267414005 |
| *ZNF506* | 0.325285 | 0.156819 | 2.07427 | 0.03805425 | 0.267460285 |
| *FAM41C* | -0.538853 | 0.25986 | -2.073625 | 0.03811419 | 0.267646681 |
| *ARL11* | 0.467167 | 0.225321 | 2.073342 | 0.03814044 | 0.267646681 |
| *SKA2* | -0.272679 | 0.131524 | -2.073226 | 0.03815129 | 0.267646681 |
| *ZNF430* | 0.215546 | 0.104003 | 2.072499 | 0.03821895 | 0.267956234 |
| *GATAD2B* | 0.308534 | 0.148992 | 2.070816 | 0.03837601 | 0.268712649 |
| *BCL2A1* | -0.381107 | 0.184051 | -2.070656 | 0.038391 | 0.268712649 |
| *NEB* | 1.193679 | 0.576494 | 2.070585 | 0.03839764 | 0.268712649 |
| *DCTN3* | -0.211833 | 0.102357 | -2.069556 | 0.03849398 | 0.269221417 |
| *TLE4* | -0.389209 | 0.188115 | -2.068996 | 0.03854649 | 0.269423193 |
| *LYN* | 0.237825 | 0.115035 | 2.067416 | 0.03869496 | 0.270189617 |
| *H2AFY* | -0.198238 | 0.095891 | -2.067324 | 0.03870361 | 0.270189617 |
| *SCCPDH* | 0.369098 | 0.178686 | 2.065621 | 0.0388643 | 0.271145164 |
| *CEP126* | 0.62897 | 0.304563 | 2.065154 | 0.03890843 | 0.271286855 |
| *ATF3* | -0.79091 | 0.383123 | -2.064375 | 0.0389822 | 0.271634859 |
| *GLRX2* | 0.333318 | 0.161684 | 2.061534 | 0.0392521 | 0.273348285 |
| *TWNK* | -0.676239 | 0.328126 | -2.060913 | 0.03931137 | 0.273416929 |
| *JTB* | -0.103045 | 0.050001 | -2.060865 | 0.03931593 | 0.273416929 |
| *CD2* | -2.306236 | 1.119233 | -2.060551 | 0.03934591 | NA |
| *ENSG00000229255* | -1.387686 | 0.673481 | -2.060468 | 0.03935381 | NA |
| *RARS* | 0.292035 | 0.141732 | 2.060466 | 0.03935397 | 0.273416929 |
| *TFRC* | -0.530245 | 0.257347 | -2.060424 | 0.03935801 | 0.273416929 |
| *ABHD4* | 0.600663 | 0.291609 | 2.059822 | 0.03941556 | 0.273649706 |
| *CSTF3* | 0.366746 | 0.178264 | 2.057318 | 0.03965562 | 0.27514864 |

| | | | | | |
|---|---|---|---|---|---|
| *MED23* | -0.421329 | 0.2049 | -2.056269 | 0.03975661 | 0.275670492 |
| *LAMC1* | 1.117449 | 0.543529 | 2.055916 | 0.03979066 | 0.275670492 |
| *PLCB2* | 0.557548 | 0.27121 | 2.055783 | 0.03980347 | 0.275670492 |
| *PPP3CA* | 0.183586 | 0.08933 | 2.055152 | 0.03986433 | 0.275697549 |
| *CD1D* | 0.484891 | 0.23595 | 2.055062 | 0.03987304 | 0.275697549 |
| *GUCD1* | -0.228831 | 0.111354 | -2.05499 | 0.03988002 | 0.275697549 |
| *RAB11FIP3* | -1.045402 | 0.509018 | -2.053762 | 0.03999876 | 0.276350662 |
| *HCFC1* | 0.356511 | 0.173692 | 2.052542 | 0.04011703 | 0.276999721 |
| *STK26* | 0.375227 | 0.182873 | 2.051843 | 0.04018496 | 0.277300566 |
| *MRPS24* | 0.814401 | 0.396982 | 2.051481 | 0.04022009 | 0.277374885 |
| *ABHD14A* | 0.316385 | 0.154321 | 2.050179 | 0.04034699 | 0.278081635 |
| *SLC5A6* | -0.450174 | 0.219668 | -2.04934 | 0.04042889 | 0.278429381 |
| *C21orf62.AS1* | 0.844339 | 0.412079 | 2.048973 | 0.04046472 | 0.278429381 |
| *PCCB* | -0.578038 | 0.28212 | -2.048911 | 0.04047081 | 0.278429381 |
| *RNPS1* | -0.124504 | 0.060787 | -2.048202 | 0.04054019 | 0.278524931 |
| *LINC02202* | -1.180736 | 0.576504 | -2.048097 | 0.0405505 | 0.278524931 |
| *EIF4H* | -0.161303 | 0.07876 | -2.04802 | 0.04055808 | 0.278524931 |
| *CASP7* | 0.398385 | 0.194695 | 2.0462 | 0.04073668 | 0.279582762 |
| *SPATA2L* | 0.473648 | 0.231548 | 2.04557 | 0.04079873 | 0.279685886 |
| *SEC13* | -0.293678 | 0.143569 | -2.045548 | 0.04080083 | 0.279685886 |
| *DCTPP1* | -0.206192 | 0.100904 | -2.043444 | 0.04100854 | 0.280888401 |
| *PDCL3* | -0.270704 | 0.132485 | -2.043271 | 0.04102559 | 0.280888401 |
| *GNA11* | 1.395906 | 0.683466 | 2.042394 | 0.04111245 | 0.281313892 |
| *WFS1* | -1.40508 | 0.688079 | -2.042035 | 0.0411481 | NA |
| *ATP5F1C* | -0.193839 | 0.094989 | -2.040648 | 0.04128579 | 0.282330341 |
| *PTP4A2* | -0.172707 | 0.084653 | -2.040168 | 0.04133355 | 0.282382106 |
| *YDJC* | -0.239327 | 0.117313 | -2.040074 | 0.04134296 | 0.282382106 |
| *MBOAT2* | -1.688592 | 0.827902 | -2.039603 | 0.0413899 | NA |
| *NPHP3* | 0.381052 | 0.186855 | 2.039293 | 0.04142075 | 0.282743791 |
| *PTPN20* | -1.34917 | 0.66206 | -2.037837 | 0.04156629 | 0.283567246 |
| *GLI4* | -0.600314 | 0.294661 | -2.037303 | 0.04161972 | 0.283677815 |
| *ALDOA* | -0.12106 | 0.059425 | -2.037177 | 0.04163232 | 0.283677815 |
| *COX8A* | -0.108718 | 0.053379 | -2.036702 | 0.04167988 | 0.283831987 |
| *APBB2* | -1.215692 | 0.597039 | -2.036202 | 0.04173005 | 0.284003808 |
| *TTC28* | 0.62354 | 0.306372 | 2.035236 | 0.04182714 | 0.284494486 |
| *CHP1* | -0.232745 | 0.114385 | -2.034745 | 0.04187655 | 0.284518238 |
| *RPRD2* | 0.298063 | 0.14649 | 2.034705 | 0.04188061 | 0.284518238 |
| *CEP85L* | 0.43149 | 0.212154 | 2.033851 | 0.04196659 | 0.284750739 |
| *ENSG00000237491* | -0.58088 | 0.285658 | -2.033484 | 0.04200369 | 0.284750739 |
| *MTX3* | -0.539546 | 0.265338 | -2.033426 | 0.04200955 | 0.284750739 |
| *CYFIP1* | -0.953944 | 0.469144 | -2.033373 | 0.04201486 | 0.284750739 |
| *ZMYM6* | 0.447637 | 0.220211 | 2.032764 | 0.04207635 | 0.284874621 |
| *IL18BP* | -0.747213 | 0.367597 | -2.032697 | 0.04208318 | 0.284874621 |
| *POLR1B* | -0.565726 | 0.278373 | -2.032259 | 0.04212746 | 0.285004897 |
| *EHD3* | -0.521672 | 0.256756 | -2.031784 | 0.04217552 | 0.285031944 |
| *LY96* | 0.282265 | 0.138943 | 2.031514 | 0.04220288 | 0.285031944 |
| *GOPC* | 0.345331 | 0.169999 | 2.031366 | 0.04221792 | 0.285031944 |

| | | | | | |
|---|---|---|---|---|---|
| *DPYD* | 0.74109 | 0.364848 | 2.031231 | 0.04223159 | 0.285031944 |
| *OAZ2* | 0.18981 | 0.09347 | 2.030703 | 0.04228517 | 0.285224467 |
| *PSMD8* | -0.152346 | 0.075053 | -2.029846 | 0.04237221 | 0.285507644 |
| *ZNF626* | 0.341341 | 0.168165 | 2.029796 | 0.0423773 | 0.285507644 |
| *TPP1* | 0.272744 | 0.134395 | 2.029419 | 0.04241567 | 0.285594958 |
| *GDPD1* | 1.18851 | 0.585711 | 2.029175 | 0.04244042 | 0.285594958 |
| *DYRK1B* | 0.467209 | 0.230286 | 2.02882 | 0.04247666 | 0.285669983 |
| *ENSG00000258377* | -1.338801 | 0.660165 | -2.02798 | 0.04256226 | NA |
| *LINC02413* | -0.721535 | 0.355917 | -2.027259 | 0.0426359 | 0.286571648 |
| *MMAB* | -0.259223 | 0.12792 | -2.026452 | 0.04271845 | 0.286901033 |
| *MTHFD2L* | -0.380884 | 0.187986 | -2.026136 | 0.04275084 | 0.286901033 |
| *ENSG00000260979* | -0.728284 | 0.359461 | -2.026042 | 0.0427605 | 0.286901033 |
| *ENSG00000269983* | 1.325527 | 0.654377 | 2.025634 | 0.04280228 | 0.287012226 |
| *TOB1* | 0.354859 | 0.175231 | 2.02509 | 0.0428581 | 0.287217366 |
| *WDR76* | -0.410969 | 0.203046 | -2.024022 | 0.04296793 | 0.287784031 |
| *AP5S1* | -0.49991 | 0.247037 | -2.023623 | 0.04300893 | 0.287889312 |
| *SLFN13* | -0.788378 | 0.389718 | -2.022948 | 0.04307852 | 0.288185702 |
| *IGHG3* | 0.498481 | 0.246529 | 2.022 | 0.04317631 | 0.28860915 |
| *FBXL17* | 0.513996 | 0.254221 | 2.021844 | 0.04319252 | 0.28860915 |
| *CRIPT* | 0.237104 | 0.117327 | 2.020878 | 0.0432924 | 0.289099281 |
| *ZNF420* | 0.436299 | 0.215945 | 2.02042 | 0.04333987 | 0.289099281 |
| *ZNF257* | 0.673246 | 0.333224 | 2.020399 | 0.04334204 | 0.289099281 |
| *FAM43A* | 0.621517 | 0.307691 | 2.019941 | 0.04338954 | 0.289190044 |
| *ZC3H14* | 0.334401 | 0.165565 | 2.019756 | 0.04340872 | 0.289190044 |
| *PPP3R1* | 0.416049 | 0.206014 | 2.019516 | 0.0434336 | 0.289190044 |
| *KMT2E* | 0.105626 | 0.05231 | 2.019245 | 0.04346177 | 0.289190044 |
| *BRD4* | 0.192904 | 0.095542 | 2.019044 | 0.04348264 | 0.289190044 |
| *SH2B2* | -1.159086 | 0.574213 | -2.018564 | 0.04353251 | 0.289299552 |
| *CBR4* | 0.274482 | 0.13599 | 2.018397 | 0.04354992 | 0.289299552 |
| *LYZ* | 1.38644 | 0.68697 | 2.018195 | 0.04357094 | NA |
| *ZNF33A* | 0.214248 | 0.10622 | 2.017027 | 0.04369273 | 0.289916275 |
| *CTNNBL1* | -0.228751 | 0.11342 | -2.016855 | 0.0437107 | 0.289916275 |
| *DMD* | 0.478684 | 0.237352 | 2.016774 | 0.04371915 | 0.289916275 |
| *KCNG1* | 1.420877 | 0.704649 | 2.016433 | 0.04375475 | 0.289983489 |
| *ENSG00000224934* | -1.971117 | 0.977909 | -2.015644 | 0.04383725 | NA |
| *R3HDM4* | -0.180717 | 0.089657 | -2.015636 | 0.04383809 | 0.290140342 |
| *TAF1A* | 0.53062 | 0.263272 | 2.015482 | 0.04385412 | 0.290140342 |
| *AMIGO2* | 0.847007 | 0.420254 | 2.015463 | 0.04385615 | 0.290140342 |
| *CDK5RAP1* | -0.302392 | 0.150067 | -2.01505 | 0.04389946 | 0.290140342 |
| *NPAT* | -0.29448 | 0.146145 | -2.014989 | 0.04390583 | 0.290140342 |
| *GPRASP1* | 1.541166 | 0.765042 | 2.014487 | 0.04395847 | NA |
| *CAPN10.DT* | 0.425877 | 0.211414 | 2.014419 | 0.04396558 | 0.290366629 |
| *EFNA3* | 0.849456 | 0.421762 | 2.014063 | 0.04400297 | 0.290445097 |
| *GATD1* | 0.368069 | 0.182858 | 2.012871 | 0.04412822 | 0.291103063 |
| *TRIR* | 0.104451 | 0.051913 | 2.012026 | 0.04421725 | 0.291521457 |
| *RFESD* | -1.104588 | 0.549298 | -2.01091 | 0.044335 | 0.292022499 |
| *NMNAT1* | 0.486839 | 0.24211 | 2.010819 | 0.04434454 | 0.292022499 |

| | | | | | |
|---|---|---|---|---|---|
| *SGPP1* | 0.444953 | 0.221374 | 2.00996 | 0.04443543 | 0.292451916 |
| *PRRG2* | -0.652407 | 0.324763 | -2.008868 | 0.0445511 | 0.293043769 |
| *MAN1A1* | 0.537891 | 0.267817 | 2.008426 | 0.04459803 | 0.293183146 |
| *ZNF500* | 0.432052 | 0.215234 | 2.007361 | 0.04471123 | 0.29375765 |
| *ZNF441* | -0.363664 | 0.181244 | -2.006492 | 0.04480383 | 0.293972422 |
| *TRMT112* | 0.137446 | 0.068505 | 2.006375 | 0.04481625 | 0.293972422 |
| *SLC37A2* | 0.804213 | 0.400838 | 2.006327 | 0.04482137 | 0.293972422 |
| *SPAG1* | 0.518556 | 0.258512 | 2.005929 | 0.04486385 | 0.294046562 |
| *ENSG00000255284* | 0.477979 | 0.238346 | 2.005397 | 0.04492067 | 0.294046562 |
| *POLR2J* | -0.172268 | 0.085938 | -2.004559 | 0.04501025 | 0.294046562 |
| *ENO1.AS1* | -1.292785 | 0.644934 | -2.004523 | 0.04501404 | NA |
| *GAB1* | 1.167674 | 0.582523 | 2.004512 | 0.04501521 | 0.294046562 |
| *GALK1* | 0.329624 | 0.164466 | 2.004205 | 0.04504806 | 0.294046562 |
| *ABTB1* | -0.228451 | 0.114016 | -2.003676 | 0.04510475 | 0.294046562 |
| *TSN* | -0.211148 | 0.105385 | -2.003591 | 0.04511386 | 0.294046562 |
| *CNN2* | -0.286875 | 0.143182 | -2.003563 | 0.04511687 | 0.294046562 |
| *SMCO4* | 0.551804 | 0.275414 | 2.003542 | 0.04511911 | 0.294046562 |
| *ZNF589* | -0.53517 | 0.267115 | -2.003523 | 0.04512117 | 0.294046562 |
| *NDUFV2.AS1* | -0.438963 | 0.219109 | -2.003403 | 0.04513405 | 0.294046562 |
| *SEC11C* | -0.218902 | 0.109279 | -2.00314 | 0.04516227 | 0.294046562 |
| *STOML1* | -0.875596 | 0.437215 | -2.002667 | 0.04521304 | 0.294046562 |
| *ETV3* | 0.658882 | 0.329005 | 2.002652 | 0.04521467 | 0.294046562 |
| *HMGN2* | 0.113676 | 0.056764 | 2.002602 | 0.04522005 | 0.294046562 |
| *STAM2* | 0.589683 | 0.294526 | 2.002145 | 0.04526911 | 0.29419755 |
| *LINC00944* | -0.697183 | 0.348391 | -2.001152 | 0.04537605 | 0.294724289 |
| *EPN2* | 0.672325 | 0.336121 | 2.000248 | 0.04547351 | 0.295188909 |
| *TMEM208* | -0.212369 | 0.1062 | -1.99971 | 0.04553155 | 0.295310525 |
| *ELF2* | 0.194883 | 0.097461 | 1.999594 | 0.04554411 | 0.295310525 |
| *XPC* | -0.254643 | 0.127366 | -1.999302 | 0.04557574 | 0.295319034 |
| *IQGAP1* | 0.171581 | 0.085829 | 1.999102 | 0.0455973 | 0.295319034 |
| *PDS5A* | 0.270788 | 0.135492 | 1.998549 | 0.04565722 | 0.295423861 |
| *PIGH* | -0.32172 | 0.160983 | -1.998473 | 0.04566538 | 0.295423861 |
| *SMPD2* | -0.335815 | 0.16806 | -1.99818 | 0.04569714 | 0.295461435 |
| *KRAS* | 0.205385 | 0.102814 | 1.997633 | 0.04575644 | 0.295557365 |
| *SCP2* | -0.166532 | 0.083368 | -1.997565 | 0.04576389 | 0.295557365 |
| *GIMAP7* | -2.669102 | 1.33678 | -1.996666 | 0.04586152 | NA |
| *SREBF2.AS1* | -0.615028 | 0.308068 | -1.996403 | 0.04589008 | 0.296204332 |
| *MFSD2B* | -0.751783 | 0.376733 | -1.995535 | 0.04598459 | 0.296554147 |
| *ZBTB32* | -0.472615 | 0.236849 | -1.995427 | 0.04599637 | 0.296554147 |
| *SLC25A25* | 0.515894 | 0.258589 | 1.995034 | 0.04603917 | 0.296662122 |
| *PPP1R37* | 0.412794 | 0.206947 | 1.994683 | 0.04607743 | 0.296709433 |
| *THAP3* | 0.33196 | 0.166459 | 1.994242 | 0.04612564 | 0.296709433 |
| *SYBU* | -1.012427 | 0.507695 | -1.994165 | 0.04613402 | 0.296709433 |
| *BAD* | -0.209413 | 0.105022 | -1.993989 | 0.0461533 | 0.296709433 |
| *PLCL2* | -0.413922 | 0.207607 | -1.993774 | 0.04617681 | 0.296709433 |
| *ZSWIM6* | 0.606091 | 0.30407 | 1.993266 | 0.04623236 | 0.29689882 |
| *RPL7L1* | 0.198218 | 0.09953 | 1.991533 | 0.04642231 | 0.29795063 |

| | | | | | |
|---|---|---|---|---|---|
| *AFTPH* | 0.20394 | 0.102418 | 1.991259 | 0.04645247 | 0.29797623 |
| *DHRS11* | -0.837461 | 0.420728 | -1.990506 | 0.0465352 | 0.298226612 |
| *DTX3L* | 0.347655 | 0.174676 | 1.990279 | 0.04656024 | 0.298226612 |
| *ETV6* | -0.280018 | 0.140699 | -1.990189 | 0.04657008 | 0.298226612 |
| *RBM5* | 0.225611 | 0.113407 | 1.989393 | 0.04665785 | 0.298620731 |
| *ATG4C* | 0.306685 | 0.154275 | 1.987908 | 0.04682187 | 0.299502162 |
| *ENSG00000268352* | 1.589442 | 0.799589 | 1.987825 | 0.04683099 | NA |
| *TMEM230* | -0.174561 | 0.08785 | -1.98703 | 0.0469191 | 0.299955573 |
| *RPP25L* | -0.258737 | 0.130249 | -1.986485 | 0.04697954 | 0.300173397 |
| *ZNF395* | 0.293244 | 0.147661 | 1.985934 | 0.04704071 | 0.30039567 |
| *DTD2* | -0.5111 | 0.257428 | -1.985413 | 0.04709855 | 0.30059649 |
| *TRIM38* | 0.167652 | 0.084487 | 1.984363 | 0.04721544 | 0.301005022 |
| *SAE1* | -0.281942 | 0.142082 | -1.984363 | 0.04721542 | 0.301005022 |
| *LYRM1* | -0.251057 | 0.126573 | -1.983485 | 0.04731324 | 0.30134174 |
| *COX6A1* | -0.11696 | 0.058969 | -1.983414 | 0.04732119 | 0.30134174 |
| *OTUD1* | 0.617889 | 0.311605 | 1.982925 | 0.04737586 | 0.301370606 |
| *CD47* | 0.158388 | 0.079877 | 1.982899 | 0.04737866 | 0.301370606 |
| *RTRAF* | -0.119793 | 0.06044 | -1.982021 | 0.04747685 | 0.301647974 |
| *ZNF292* | 0.194634 | 0.098204 | 1.981928 | 0.04748731 | 0.301647974 |
| *CRACR2B* | -0.624437 | 0.31513 | -1.98152 | 0.04753303 | 0.301647974 |
| *CDKN2A* | -0.774931 | 0.391133 | -1.981248 | 0.04756342 | 0.301647974 |
| *RBM42* | -0.236386 | 0.119318 | -1.981143 | 0.04757526 | 0.301647974 |
| *PRMT6* | 0.527092 | 0.266072 | 1.981017 | 0.04758932 | 0.301647974 |
| *PDCD7* | 0.391464 | 0.197624 | 1.980853 | 0.04760771 | 0.301647974 |
| *HSPA5* | -0.319645 | 0.161442 | -1.979934 | 0.04771091 | 0.302133699 |
| *MST1* | 1.330508 | 0.672121 | 1.979567 | 0.04775218 | 0.302187597 |
| *AHRR* | -1.310865 | 0.662275 | -1.979337 | 0.04777811 | 0.302187597 |
| *TARBP1* | -0.444572 | 0.224628 | -1.979151 | 0.04779904 | 0.302187597 |
| *MRRF* | 0.384764 | 0.194463 | 1.978592 | 0.04786191 | 0.302417132 |
| *ENSG00000224220* | -0.67378 | 0.340635 | -1.978013 | 0.0479272 | 0.302578412 |
| *TXLNB* | -0.725263 | 0.366705 | -1.977785 | 0.047953 | 0.302578412 |
| *VPS35* | 0.199472 | 0.100884 | 1.97724 | 0.04801452 | 0.302578412 |
| *CXorf65* | -0.838996 | 0.424338 | -1.977189 | 0.04802031 | 0.302578412 |
| *LRCH4* | -0.172122 | 0.087057 | -1.97711 | 0.04802924 | 0.302578412 |
| *GRAPL* | 0.881017 | 0.445646 | 1.976943 | 0.04804806 | 0.302578412 |
| *PWWP2B* | 0.562526 | 0.284592 | 1.976603 | 0.04808653 | 0.302578412 |
| *SNX21* | 1.264245 | 0.639675 | 1.976387 | 0.04811093 | 0.302578412 |
| *TXNDC15* | 0.294988 | 0.149288 | 1.975962 | 0.04815911 | 0.302578412 |
| *CHRNE* | 1.489995 | 0.754122 | 1.975802 | 0.04817724 | 0.302578412 |
| *DHRS1* | -0.289412 | 0.146491 | -1.975636 | 0.04819604 | 0.302578412 |
| *DONSON* | 0.937354 | 0.474549 | 1.975252 | 0.04823951 | 0.302578412 |
| *LZTS2* | 0.532498 | 0.269636 | 1.974875 | 0.04828235 | 0.302578412 |
| *ENSG00000262049* | -0.394189 | 0.199606 | -1.974832 | 0.04828723 | 0.302578412 |
| *LTO1* | -0.392347 | 0.198681 | -1.974756 | 0.0482959 | 0.302578412 |
| *KARS* | -0.196168 | 0.099345 | -1.974608 | 0.04831263 | 0.302578412 |
| *ENSG00000272053* | -0.648341 | 0.328429 | -1.974069 | 0.04837388 | 0.302795499 |
| *PXK* | 0.29097 | 0.147449 | 1.973358 | 0.04845475 | 0.302973321 |

| | | | | | |
|---|---|---|---|---|---|
| *LINC00672* | 0.650568 | 0.329691 | 1.973264 | 0.04846553 | 0.302973321 |
| *PTCD1* | -0.514881 | 0.260965 | -1.972989 | 0.04849684 | 0.302973321 |
| *KLHL5* | -0.325922 | 0.165201 | -1.972884 | 0.04850873 | 0.302973321 |
| *ERF* | 0.507834 | 0.257565 | 1.971674 | 0.04864689 | 0.303669656 |
| *EIF3E* | 0.168396 | 0.085429 | 1.971173 | 0.04870407 | 0.303859993 |
| *HMGCS1* | -0.308212 | 0.15653 | -1.969021 | 0.04895068 | 0.305015115 |
| *ADAM15* | 0.765381 | 0.388722 | 1.968965 | 0.04895706 | 0.305015115 |
| *TMEM59* | -0.117538 | 0.059699 | -1.968856 | 0.04896959 | 0.305015115 |
| *WDR11* | -0.239539 | 0.1217 | -1.968274 | 0.04903658 | 0.305015115 |
| *PARP12* | -0.441403 | 0.224299 | -1.967926 | 0.04907658 | 0.305015115 |
| *SOGA1* | 0.52001 | 0.26427 | 1.967722 | 0.04910006 | 0.305015115 |
| *MAD2L1* | -0.652494 | 0.331633 | -1.967518 | 0.04912353 | 0.305015115 |
| *H3F3B* | -0.111063 | 0.056455 | -1.967268 | 0.04915229 | 0.305015115 |
| *PDPK1* | 0.307101 | 0.156106 | 1.967259 | 0.04915333 | 0.305015115 |
| *APC* | 0.348693 | 0.177263 | 1.9671 | 0.04917164 | 0.305015115 |
| *LMCD1* | -1.181356 | 0.600589 | -1.966994 | 0.04918389 | 0.305015115 |
| *C4orf36* | 0.713331 | 0.362742 | 1.966496 | 0.04924136 | 0.3052053 |
| *NCK1* | 0.225183 | 0.114566 | 1.965529 | 0.04935308 | 0.30556225 |
| *CYB5R1* | -0.336082 | 0.170999 | -1.965405 | 0.04936737 | 0.30556225 |
| *PRR7* | -0.585461 | 0.29792 | -1.965161 | 0.04939565 | 0.30556225 |
| *SLC25A17* | 0.45408 | 0.231076 | 1.965069 | 0.0494063 | 0.30556225 |
| *CD226* | -0.982828 | 0.50031 | -1.964437 | 0.04947942 | 0.30575812 |
| *DNMT3A* | 0.466579 | 0.237545 | 1.964172 | 0.04951015 | 0.30575812 |
| *BRD3OS* | 0.385572 | 0.196328 | 1.963922 | 0.04953915 | 0.30575812 |
| *LRPAP1* | -0.188928 | 0.096211 | -1.963678 | 0.04956749 | 0.30575812 |
| *ENPP4* | -0.639681 | 0.325792 | -1.963465 | 0.04959221 | 0.30575812 |
| *ERLIN1* | 0.619619 | 0.31561 | 1.963243 | 0.049618 | 0.30575812 |
| *NCKAP5L* | 0.759262 | 0.386797 | 1.962945 | 0.04965257 | 0.30575812 |
| *IFNAR1* | -0.186854 | 0.095191 | -1.962943 | 0.0496528 | 0.30575812 |
| *FBXL5* | -0.26722 | 0.136189 | -1.962126 | 0.04974776 | 0.306177302 |
| *AP1S3* | -0.30821 | 0.157184 | -1.960826 | 0.04989929 | 0.3066074 |
| *DGAT2* | 0.67582 | 0.34467 | 1.960773 | 0.04990548 | 0.3066074 |
| *INTS2* | 0.764816 | 0.390065 | 1.960742 | 0.04990913 | 0.3066074 |
| *CEP19* | 0.464908 | 0.237153 | 1.960373 | 0.04995219 | 0.3066074 |
| *HUWE1* | 0.295783 | 0.150881 | 1.960372 | 0.04995229 | 0.3066074 |

| Cluster 6 | log2FC | lfcSE | stat | pvalue | padj |
|---|---|---|---|---|---|
| *RPL8* | 0.291116 | 0.078616 | 3.702999 | 0.000213066 | 0.999975 |
| *RPS5* | 0.217438 | 0.064063 | 3.394151 | 0.000688418 | 0.999975 |
| *ENSG00000272211* | -1.924375 | 0.586996 | -3.278343 | 0.001044186 | 0.999975 |
| *RPS9* | 0.324713 | 0.111146 | 2.921486 | 0.003483661 | 0.999975 |
| *GATD3A* | 3.793876 | 1.342504 | 2.825971 | 0.004713757 | 0.999975 |
| *ZBED2* | 2.666112 | 0.957528 | 2.784371 | 0.005363168 | 0.999975 |
| *IGHA1* | -1.029965 | 0.372817 | -2.762654 | 0.005733349 | 0.999975 |
| *ARRDC3* | -1.260022 | 0.483171 | -2.607816 | 0.009112182 | 0.999975 |
| *MT.ND2* | -0.306245 | 0.124422 | -2.461346 | 0.013841672 | 0.999975 |
| *MT.ND3* | -0.342426 | 0.140316 | -2.440388 | 0.014671479 | 0.999975 |
| *CKS2* | 1.249205 | 0.527946 | 2.366163 | 0.017973543 | 0.999975 |
| *H3F3B* | -0.451573 | 0.194619 | -2.320289 | 0.02032526 | 0.999975 |
| *CCDC191* | -1.459879 | 0.642009 | -2.273921 | 0.022970712 | 0.999975 |
| *FKBP5* | 1.523412 | 0.684697 | 2.224943 | 0.026085098 | 0.999975 |
| *ENSG00000279483* | -0.50764 | 0.23418 | -2.167736 | 0.030178767 | 0.999975 |
| *RGCC* | 2.272594 | 1.080785 | 2.102726 | 0.035489745 | 0.999975 |
| *USP53* | 1.261884 | 0.605639 | 2.08356 | 0.037200171 | 0.999975 |
| *ZNF107* | 1.405976 | 0.680553 | 2.06593 | 0.038835044 | 0.999975 |
| *HLA.C* | -0.577522 | 0.282867 | -2.041673 | 0.041183964 | 0.999975 |
| *FCGR2B* | 1.173359 | 0.577103 | 2.033187 | 0.042033681 | 0.999975 |
| *HLA.A* | -0.478721 | 0.237665 | -2.014272 | 0.043980934 | 0.999975 |
| *MT.CO1* | 0.170556 | 0.084934 | 2.00809 | 0.04463373 | 0.999975 |
| *RPS6KA2* | 1.484232 | 0.739227 | 2.007816 | 0.044662896 | 0.999975 |
| *RPS8* | 0.139498 | 0.070415 | 1.981086 | 0.047581593 | 0.999975 |
| *TCL1B* | -1.198105 | 0.609515 | -1.965671 | 0.04933664 | 0.999975 |

| Cluster 7 | log2FC | lfcSE | stat | pvalue | padj |
|---|---|---|---|---|---|
| *NFKBIA* | -0.589599 | 0.108085 | -5.454954 | 4.90E-08 | 0.000153717 |
| *CD83* | -0.596054 | 0.114193 | -5.219688 | 1.79E-07 | 0.000219207 |
| *ENSG00000272211* | -1.304309 | 0.251281 | -5.190645 | 2.10E-07 | 0.000219207 |
| *RSRP1* | 0.510122 | 0.111103 | 4.591446 | 4.40E-06 | 0.003453258 |
| *RPL8* | 0.392014 | 0.088595 | 4.424784 | 9.65E-06 | 0.006058769 |
| *IL4R* | -0.603296 | 0.147597 | -4.087463 | 4.36E-05 | 0.022808867 |
| *CKS2* | 1.314245 | 0.332853 | 3.948426 | 7.87E-05 | NA |
| *DHTKD1* | -0.720035 | 0.183639 | -3.920932 | 8.82E-05 | NA |
| *ARRDC3* | -0.922866 | 0.238396 | -3.871148 | 0.000108324 | 0.048560153 |
| *HHEX* | 0.438362 | 0.115716 | 3.788267 | 0.000151702 | 0.059505099 |
| *TNFAIP8L2* | 1.098924 | 0.291262 | 3.772975 | 0.000161313 | NA |
| *LMO4* | 0.839182 | 0.228088 | 3.679203 | 0.000233964 | NA |
| *UCP2* | -0.468388 | 0.128721 | -3.638776 | 0.000273937 | 0.095512832 |
| *CYB5A* | -0.644184 | 0.181004 | -3.558953 | 0.000372336 | 0.099575711 |
| *PFKFB3* | -0.663773 | 0.18778 | -3.534837 | 0.000408027 | 0.099575711 |
| *SPINT2* | -0.474838 | 0.135254 | -3.510707 | 0.000446917 | 0.099575711 |
| *SYNGR2* | -0.742773 | 0.211756 | -3.507677 | 0.000452037 | 0.099575711 |
| *HLA.A* | -0.527609 | 0.150451 | -3.506836 | 0.000453468 | 0.099575711 |
| *VSIR* | -1.593327 | 0.455605 | -3.497167 | 0.000470227 | NA |
| *GATD3A* | 2.725647 | 0.783069 | 3.480721 | 0.000500065 | NA |
| *YARS* | -0.822365 | 0.236412 | -3.478522 | 0.000504188 | 0.099575711 |
| *ENSG00000273319* | 0.68188 | 0.196131 | 3.476652 | 0.000507716 | 0.099575711 |
| *THAP5* | 0.568686 | 0.166482 | 3.415895 | 0.000635727 | 0.112460356 |
| *SLC25A26* | -0.578526 | 0.170352 | -3.396068 | 0.000683615 | 0.112460356 |
| *SRSF5* | -0.338015 | 0.099678 | -3.391053 | 0.000696246 | 0.112460356 |
| *FTL* | 0.437397 | 0.129289 | 3.383085 | 0.000716765 | 0.112460356 |
| *ALDH2* | -1.031022 | 0.30603 | -3.369021 | 0.000754357 | NA |
| *S1PR4* | 0.631195 | 0.191337 | 3.298861 | 0.000970778 | 0.145061986 |
| *SGK1* | -1.427222 | 0.436534 | -3.269442 | 0.001077599 | NA |
| *RPS5* | 0.171608 | 0.053141 | 3.229312 | 0.001240885 | 0.176995282 |
| *TMEM170A* | 0.7315 | 0.228467 | 3.201776 | 0.00136583 | NA |
| *ATP6V1C2* | -2.601158 | 0.819725 | -3.173208 | 0.001507645 | NA |
| *CCDC191* | -1.198102 | 0.378156 | -3.168275 | 0.001533463 | NA |
| *EGLN2* | 0.428155 | 0.136386 | 3.139285 | 0.001693604 | 0.223077602 |
| *UBE2J1* | 0.533853 | 0.170173 | 3.137124 | 0.001706138 | 0.223077602 |
| *SMIM10* | -3.105246 | 0.992002 | -3.130281 | 0.00174639 | NA |
| *SPATA13* | 1.458187 | 0.470059 | 3.102137 | 0.001921288 | NA |
| *LRIF1* | 0.669042 | 0.215724 | 3.101373 | 0.001926257 | NA |
| *POLM* | -0.485731 | 0.157549 | -3.083037 | 0.002048995 | 0.241153095 |
| *ERGIC2* | 0.545944 | 0.177197 | 3.081 | 0.002063065 | 0.241153095 |
| *PTPN18* | 0.525551 | 0.170673 | 3.079292 | 0.002074931 | 0.241153095 |
| *SOX4* | -1.160561 | 0.38091 | -3.046815 | 0.002312798 | NA |
| *GALNT2* | -0.902337 | 0.29659 | -3.042372 | 0.002347219 | 0.258279367 |
| *IL21R* | -0.991515 | 0.326304 | -3.038622 | 0.002376632 | NA |
| *CLEC2B* | 0.454505 | 0.14964 | 3.037322 | 0.002386903 | 0.258279367 |
| *LTBP3* | 0.618855 | 0.205083 | 3.01758 | 0.002548016 | 0.266522484 |

| | | | | | |
|---|---|---|---|---|---|
| *ZNF384* | -0.90725 | 0.302328 | -3.000878 | 0.002692023 | NA |
| *SLC38A2* | 0.493172 | 0.164858 | 2.991489 | 0.002776203 | 0.27727229 |
| *ZMAT2* | -0.392499 | 0.131451 | -2.985894 | 0.002827506 | 0.27727229 |
| *BLZF1* | 0.962376 | 0.322434 | 2.984717 | 0.002838405 | NA |
| *TTC32* | -0.900678 | 0.301862 | -2.983738 | 0.002847509 | NA |
| *TNFRSF13B* | 1.188875 | 0.400008 | 2.972126 | 0.002957449 | 0.281226501 |
| *ATP5F1EP2* | -0.611148 | 0.207654 | -2.943114 | 0.003249289 | 0.288292273 |
| *TIMM10* | -0.545754 | 0.185467 | -2.942594 | 0.003254751 | 0.288292273 |
| *PNPLA8* | 0.539085 | 0.183511 | 2.937625 | 0.003307368 | 0.288292273 |
| *NUFIP2* | 0.618614 | 0.211229 | 2.928645 | 0.003404427 | NA |
| *RPS9* | 0.268888 | 0.09216 | 2.91762 | 0.003527141 | 0.291458824 |
| *HLA.C* | -0.569329 | 0.195148 | -2.917415 | 0.003529457 | 0.291458824 |
| *PSMB10* | -0.376336 | 0.12979 | -2.899568 | 0.003736774 | 0.300666567 |
| *FNBP1* | -0.437726 | 0.1515 | -2.88928 | 0.003861248 | 0.302914882 |
| *ENSG00000166927* | -1.011916 | 0.351132 | -2.881865 | 0.003953289 | NA |
| *ZNF213.AS1* | -1.064327 | 0.369396 | -2.881259 | 0.003960901 | NA |
| *UNC119* | -0.521173 | 0.181803 | -2.866691 | 0.004147879 | 0.317464465 |
| *ZBED2* | 1.360155 | 0.476039 | 2.857236 | 0.004273474 | NA |
| *TSC22D1* | -1.089259 | 0.381678 | -2.853868 | 0.004319047 | NA |
| *RPS15A* | 0.21629 | 0.076095 | 2.842354 | 0.004478168 | 0.329987664 |
| *ZFP36L2* | 0.633505 | 0.223871 | 2.829776 | 0.004658059 | 0.329987664 |
| *ENSG00000257275* | -0.774008 | 0.275222 | -2.812309 | 0.004918727 | 0.329987664 |
| *CIRBP* | -0.279231 | 0.099329 | -2.811177 | 0.004936062 | 0.329987664 |
| *LINC00662* | 0.539275 | 0.19185 | 2.810915 | 0.00494009 | 0.329987664 |
| *COL19A1* | -0.704555 | 0.250692 | -2.810444 | 0.004947324 | 0.329987664 |
| *TPP1* | 0.476629 | 0.169983 | 2.803979 | 0.005047612 | 0.329987664 |
| *HSH2D* | -0.353241 | 0.126405 | -2.794515 | 0.00519776 | 0.332868817 |
| *NSMAF* | 0.864556 | 0.310467 | 2.784693 | 0.00535784 | NA |
| *RNF130* | 0.729023 | 0.262284 | 2.779523 | 0.00544388 | 0.34165788 |
| *SLC7A7* | -1.472032 | 0.532204 | -2.765916 | 0.005676308 | NA |
| *DDX6* | -0.409017 | 0.148651 | -2.751526 | 0.005931836 | 0.364982375 |
| *ZNF267* | -0.471753 | 0.171855 | -2.745064 | 0.006049904 | 0.365088445 |
| *XRRA1* | -0.847882 | 0.30968 | -2.737927 | 0.006182773 | 0.36606681 |
| *UNC50* | 0.556374 | 0.203385 | 2.735569 | 0.006227258 | NA |
| *CHP1* | -0.465002 | 0.170259 | -2.731141 | 0.006311541 | 0.366770658 |
| *RPLP2* | 0.206919 | 0.076733 | 2.696614 | 0.007004847 | 0.399658381 |
| *SLC50A1* | -0.606358 | 0.22551 | -2.688826 | 0.00717037 | 0.401796824 |
| *IGHG1* | 1.300883 | 0.490485 | 2.652238 | 0.007996008 | 0.42838818 |
| *ATP5MGL* | -0.977288 | 0.368573 | -2.651545 | 0.008012438 | NA |
| *NOP10* | 0.387292 | 0.146104 | 2.650796 | 0.008030222 | 0.42838818 |
| *NDUFA7* | -0.527851 | 0.199574 | -2.644893 | 0.008171678 | 0.42838818 |
| *RBM3* | -0.299262 | 0.11335 | -2.640162 | 0.008286646 | 0.42838818 |
| *CLEC2D* | -0.298164 | 0.113005 | -2.638495 | 0.008327495 | 0.42838818 |
| *CLK1* | 0.488751 | 0.186085 | 2.626495 | 0.008626926 | 0.436633762 |
| *SFXN1* | 0.602102 | 0.229929 | 2.618643 | 0.008828026 | NA |
| *RUBCNL* | -0.313043 | 0.119763 | -2.61386 | 0.008952568 | 0.445923143 |
| *ITGB3BP* | -0.518411 | 0.199336 | -2.600686 | 0.009303757 | 0.455216371 |

| | | | | | |
|---|---|---|---|---|---|
| *CRIPT* | 0.526297 | 0.202727 | 2.596085 | 0.009429275 | 0.455216371 |
| *NAB2* | 1.108011 | 0.428529 | 2.585614 | 0.009720574 | NA |
| *RASGRP1* | -1.129173 | 0.436989 | -2.583982 | 0.0097667 | NA |
| *DMAC2L* | -0.552195 | 0.214137 | -2.5787 | 0.009917275 | NA |
| *ASAH1* | 0.465621 | 0.180623 | 2.577862 | 0.009941357 | 0.467786817 |
| *NUP88* | -0.406019 | 0.158046 | -2.568996 | 0.010199376 | 0.467786817 |
| *SYNJ2BP* | 0.452199 | 0.176202 | 2.566362 | 0.010277149 | 0.467786817 |
| *SKAP2* | 0.379343 | 0.14783 | 2.566065 | 0.010285943 | 0.467786817 |
| *MIR155HG* | 0.73704 | 0.287723 | 2.561631 | 0.010418206 | NA |
| *PPM1A* | 0.757275 | 0.295797 | 2.560116 | 0.010463711 | NA |
| *MRPL54* | -0.31938 | 0.12502 | -2.554639 | 0.010629787 | 0.469893581 |
| *ENSG00000245904* | 0.295859 | 0.115815 | 2.554575 | 0.010631754 | 0.469893581 |
| *GNPDA1* | 1.057631 | 0.415654 | 2.544498 | 0.010943481 | NA |
| *RPL38* | 0.121179 | 0.047641 | 2.543613 | 0.010971248 | 0.478163552 |
| *CHRAC1* | -0.492513 | 0.194327 | -2.534456 | 0.011262199 | 0.484120268 |
| *ENSG00000226571* | 1.431657 | 0.564893 | 2.534386 | 0.011264451 | NA |
| *ZCCHC10* | 0.368837 | 0.146648 | 2.51512 | 0.011899177 | 0.497621402 |
| *TOMM20* | 0.255903 | 0.101759 | 2.514795 | 0.011910159 | 0.497621402 |
| *IZUMO4* | -0.630715 | 0.25137 | -2.509109 | 0.012103602 | 0.497621402 |
| *MIF* | -0.369093 | 0.147284 | -2.505999 | 0.012210595 | 0.497621402 |
| *NRP2* | -1.104423 | 0.440829 | -2.50533 | 0.012233704 | NA |
| *TMEM99* | 0.804338 | 0.322419 | 2.494698 | 0.012606431 | NA |
| *GPM6B* | 0.589582 | 0.236571 | 2.492202 | 0.012695368 | NA |
| *EIF5* | 0.302731 | 0.121666 | 2.488223 | 0.012838317 | 0.516495366 |
| *CD5* | -1.680969 | 0.676053 | -2.486446 | 0.012902616 | NA |
| *ZNF138* | 0.605276 | 0.243672 | 2.483984 | 0.01299217 | NA |
| *FKBP5* | 0.816051 | 0.32929 | 2.478216 | 0.013204103 | NA |
| *FAM41C* | -1.068826 | 0.431298 | -2.478161 | 0.01320617 | NA |
| *RNF149* | 0.47673 | 0.19248 | 2.476778 | 0.013257446 | NA |
| *LY96* | 0.523387 | 0.211341 | 2.476509 | 0.013267445 | 0.526695745 |
| *UBE2J2* | -0.35492 | 0.143563 | -2.472224 | 0.013427552 | 0.526695745 |
| *CAPZB* | -0.290764 | 0.118008 | -2.463938 | 0.013741998 | 0.532375187 |
| *RIN3* | 0.62335 | 0.253603 | 2.457973 | 0.013972368 | 0.534698661 |
| *RND1* | -1.217249 | 0.495818 | -2.455032 | 0.014087212 | NA |
| *RPS28* | 0.112823 | 0.046269 | 2.438443 | 0.014750667 | 0.554477763 |
| *MGAT1* | 0.649552 | 0.266753 | 2.435035 | 0.014890355 | 0.554477763 |
| *DBI* | -0.275258 | 0.113186 | -2.431913 | 0.015019315 | 0.554477763 |
| *GLIDR* | -1.07206 | 0.440917 | -2.431432 | 0.015039286 | NA |
| *TELO2* | -0.780202 | 0.321514 | -2.42665 | 0.015238967 | NA |
| *ZBTB1* | 0.636054 | 0.263347 | 2.415275 | 0.015723333 | 0.573633941 |
| *TRIM14* | 1.04446 | 0.432959 | 2.412375 | 0.015848967 | NA |
| *GYPC* | 0.244222 | 0.101309 | 2.410669 | 0.015923291 | 0.573633941 |
| *MCRS1* | -0.582877 | 0.242152 | -2.40707 | 0.016081067 | NA |
| *BOD1* | 0.622027 | 0.25843 | 2.406945 | 0.016086611 | 0.573633941 |
| *KCTD18* | -0.687714 | 0.286056 | -2.404122 | 0.016211382 | NA |
| *FNIP1* | -0.488311 | 0.203415 | -2.400561 | 0.01636996 | 0.577179056 |
| *ENSG00000277654* | -0.639811 | 0.266727 | -2.398746 | 0.016451311 | NA |

| | | | | | |
|---|---|---|---|---|---|
| *PDE4D* | -1.437387 | 0.59925 | -2.398642 | 0.016455992 | NA |
| *IGHM* | -0.462739 | 0.193859 | -2.386984 | 0.016987208 | 0.589008319 |
| *MAPKAPK2* | -0.486684 | 0.204057 | -2.385034 | 0.017077551 | NA |
| *SESN3* | 0.367948 | 0.154278 | 2.384962 | 0.017080866 | 0.589008319 |
| *NFKBIB* | -0.601624 | 0.252923 | -2.378687 | 0.01737443 | NA |
| *UBB* | -0.259926 | 0.109355 | -2.376897 | 0.017458972 | 0.595391791 |
| *ENSG00000237491* | -1.049311 | 0.441935 | -2.374353 | 0.017579734 | NA |
| *DNAJA1* | 0.35214 | 0.148396 | 2.372975 | 0.017645455 | 0.595391791 |
| *APH1A* | -0.276539 | 0.117055 | -2.362475 | 0.018153369 | 0.606013541 |
| *TRMT61B* | -0.668557 | 0.283415 | -2.358937 | 0.018327358 | NA |
| *KIAA0040* | 0.556219 | 0.236294 | 2.353922 | 0.018576502 | 0.610470541 |
| *SLC35A4* | -0.488628 | 0.207633 | -2.353325 | 0.018606364 | NA |
| *FXYD5* | 0.316651 | 0.134634 | 2.351936 | 0.018675963 | 0.610470541 |
| *PLTP* | -1.870831 | 0.797371 | -2.34625 | 0.018963379 | NA |
| *SCAND1* | -0.330193 | 0.141114 | -2.339908 | 0.019288476 | 0.623992129 |
| *IL13RA1* | -0.996821 | 0.426562 | -2.33687 | 0.019445944 | NA |
| *CERS4* | -0.413139 | 0.176959 | -2.334653 | 0.019561564 | 0.624242488 |
| *SLC17A5* | 0.907611 | 0.388898 | 2.333798 | 0.01960628 | NA |
| *HCK* | -1.128729 | 0.483904 | -2.332549 | 0.019671836 | NA |
| *DUSP14* | 1.041223 | 0.446394 | 2.332522 | 0.019673258 | NA |
| *IGHD* | -0.4356 | 0.186792 | -2.332008 | 0.019700296 | 0.624242488 |
| *CTDSP2* | 0.382699 | 0.164364 | 2.32836 | 0.019893005 | 0.624242488 |
| *FXYD7* | 1.007824 | 0.433147 | 2.326747 | 0.019978715 | NA |
| *ENSG00000258377* | -2.091176 | 0.899501 | -2.324817 | 0.02008176 | NA |
| *ARL11* | 0.654361 | 0.282232 | 2.31852 | 0.020421094 | 0.633328028 |
| *RHBDD2* | -0.599664 | 0.258932 | -2.315911 | 0.020563122 | NA |
| *IQSEC1* | -0.425062 | 0.183573 | -2.315489 | 0.020586188 | 0.633328028 |
| *HTR3A* | -1.22267 | 0.528102 | -2.315214 | 0.020601196 | NA |
| *PIM1* | -0.355945 | 0.154162 | -2.308907 | 0.020948719 | 0.638224085 |
| *PDPK1* | 0.587186 | 0.254604 | 2.306273 | 0.021095373 | NA |
| *FBXL3* | 0.727554 | 0.316108 | 2.301599 | 0.021357807 | 0.644430758 |
| *RPS14* | 0.090191 | 0.039419 | 2.28801 | 0.022136922 | 0.657517493 |
| *HIST1H2BH* | 1.754652 | 0.767736 | 2.285489 | 0.022284151 | NA |
| *INKA1* | 0.636941 | 0.279259 | 2.280824 | 0.022558884 | NA |
| *RPL23A* | 0.152884 | 0.067062 | 2.279758 | 0.022622039 | 0.657517493 |
| *PTGIR* | -1.256031 | 0.551964 | -2.275567 | 0.022871959 | NA |
| *PRDM8* | -0.685827 | 0.302545 | -2.26686 | 0.023398781 | NA |
| *H3F3B* | -0.197658 | 0.087278 | -2.264697 | 0.023531298 | 0.657517493 |
| *PSMC3* | -0.343157 | 0.151526 | -2.264676 | 0.023532587 | 0.657517493 |
| *CCNG2* | 0.595164 | 0.262929 | 2.263592 | 0.023599221 | NA |
| *SLA* | 0.390024 | 0.172726 | 2.258043 | 0.023942953 | 0.657517493 |
| *MRPL38* | -0.881614 | 0.390504 | -2.257632 | 0.023968581 | NA |
| *YWHAE* | 0.34078 | 0.151022 | 2.25649 | 0.024039941 | 0.657517493 |
| *MYLIP* | -0.703816 | 0.312333 | -2.25342 | 0.024232695 | NA |
| *C7orf50* | -0.502197 | 0.223136 | -2.250633 | 0.024408789 | 0.657517493 |
| *TRMT112* | 0.257489 | 0.11442 | 2.250383 | 0.024424661 | 0.657517493 |
| *A4GALT* | 1.054016 | 0.468546 | 2.249547 | 0.024477693 | NA |

| | | | | | |
|---|---|---|---|---|---|
| *FCRL3* | -1.056591 | 0.469706 | -2.249472 | 0.024482472 | 0.657517493 |
| *ATP1B3* | -0.495195 | 0.220176 | -2.249091 | 0.024506676 | 0.657517493 |
| *AKIRIN2* | 0.374651 | 0.166624 | 2.248477 | 0.02454582 | 0.657517493 |
| *H2AFZ* | 0.325059 | 0.14465 | 2.24721 | 0.024626626 | 0.657517493 |
| *RPS3* | 0.140592 | 0.062602 | 2.245831 | 0.024714856 | 0.657517493 |
| *GSTO1* | 0.306363 | 0.136424 | 2.245672 | 0.024725004 | 0.657517493 |
| *GBGT1* | 0.866527 | 0.386301 | 2.243137 | 0.024888003 | NA |
| *HYI* | -0.982748 | 0.438532 | -2.240997 | 0.025026246 | NA |
| *UBE2L3* | -0.331356 | 0.148028 | -2.238479 | 0.025189867 | 0.662078259 |
| *ZNF224* | 0.366937 | 0.164067 | 2.236509 | 0.02531848 | 0.662078259 |
| *IRF5* | 0.494119 | 0.221047 | 2.235353 | 0.025394205 | NA |
| *PRICKLE1* | 0.797476 | 0.357057 | 2.233467 | 0.025518145 | NA |
| *IGLC7* | -3.737968 | 1.67412 | -2.232796 | NA | NA |
| *SRP72* | -0.305074 | 0.136804 | -2.23 | 0.02574744 | 0.666809765 |
| *RBM23* | -0.244954 | 0.110088 | -2.22507 | 0.026076534 | 0.666809765 |
| *CD1D* | 0.353763 | 0.159074 | 2.22389 | 0.026155875 | 0.666809765 |
| *U2AF1* | -0.505831 | 0.227694 | -2.221541 | 0.026314327 | NA |
| *IDE* | -0.785005 | 0.353973 | -2.2177 | 0.026575263 | NA |
| *LBH* | -0.33423 | 0.150746 | -2.217169 | 0.026611528 | 0.666809765 |
| *SCRN1* | 1.008914 | 0.455228 | 2.216281 | 0.026672285 | NA |
| *ENSG00000250850* | 0.908368 | 0.41014 | 2.214778 | 0.02677529 | NA |
| *HIST1H2AG* | 1.388216 | 0.628361 | 2.209264 | 0.027156295 | NA |
| *HLX* | 1.491108 | 0.675678 | 2.206832 | 0.027325771 | NA |
| *CYB561A3* | -0.306831 | 0.139175 | -2.204637 | 0.027479596 | 0.666809765 |
| *SVBP* | -0.281622 | 0.127882 | -2.202207 | 0.027650676 | 0.666809765 |
| *AKAP13* | 0.268659 | 0.122082 | 2.200655 | 0.02776045 | 0.666809765 |
| *FAM91A1* | 0.575982 | 0.262191 | 2.196804 | 0.028034418 | NA |
| *FLCN* | 0.512732 | 0.233412 | 2.196679 | 0.028043402 | 0.666809765 |
| *PFDN5* | 0.134653 | 0.061299 | 2.196647 | 0.028045695 | 0.666809765 |
| *SLC15A2* | 1.208311 | 0.550179 | 2.196216 | 0.028076506 | NA |
| *CD79B* | -0.239211 | 0.109007 | -2.194452 | 0.028202959 | 0.666809765 |
| *GSTP1* | -0.282609 | 0.128905 | -2.192378 | 0.028352192 | 0.666809765 |
| *CYTH1* | -0.237575 | 0.108412 | -2.191402 | 0.028422746 | 0.666809765 |
| *THG1L* | 0.485503 | 0.221637 | 2.190537 | 0.028485289 | NA |
| *RASGRP3* | -0.517549 | 0.23646 | -2.188734 | 0.028616163 | 0.666809765 |
| *SPN* | -1.00512 | 0.459407 | -2.187865 | 0.028679413 | NA |
| *RBM39* | 0.181451 | 0.082971 | 2.18693 | 0.02874766 | 0.666809765 |
| *ATG4B* | -0.40115 | 0.183538 | -2.185654 | 0.028840911 | 0.666809765 |
| *SRRT* | -0.415245 | 0.190056 | -2.184857 | 0.02889934 | 0.666809765 |
| *GOLIM4* | -0.800833 | 0.367021 | -2.181978 | 0.029111137 | NA |
| *NBPF3* | -1.83386 | 0.841469 | -2.179356 | 0.029305234 | NA |
| *CAPN12* | 0.561631 | 0.258242 | 2.174824 | 0.029643341 | NA |
| *GRHPR* | -0.531943 | 0.245067 | -2.170604 | 0.029961139 | 0.681518633 |
| *VPS26A* | 0.483213 | 0.222631 | 2.170471 | 0.029971183 | 0.681518633 |
| *BBX* | 0.317405 | 0.146727 | 2.163235 | 0.030523081 | 0.68496208 |
| *CD74* | -0.261628 | 0.120969 | -2.162766 | 0.030559175 | 0.68496208 |
| *CRK* | 0.603806 | 0.280116 | 2.155553 | 0.031118548 | NA |

| | | | | | |
|---|---|---|---|---|---|
| *IFI16* | 0.269488 | 0.125049 | 2.155061 | 0.031157092 | 0.692431746 |
| *RPS8* | 0.129299 | 0.060061 | 2.152808 | 0.03133375 | 0.692431746 |
| *SLC16A1.AS1* | 0.470025 | 0.218349 | 2.15263 | 0.031347749 | NA |
| *PSMD9* | -0.35387 | 0.164857 | -2.146519 | 0.031831558 | 0.698513479 |
| *SEL1L3* | 0.489558 | 0.228431 | 2.143134 | 0.032102302 | 0.698983806 |
| *RNF43* | -0.894971 | 0.418165 | -2.140232 | 0.032336057 | NA |
| *ZMYND8* | 0.464683 | 0.217397 | 2.137488 | 0.032558311 | NA |
| *LYST* | -0.560405 | 0.262617 | -2.133927 | 0.032848776 | 0.698983806 |
| *RPL30* | 0.091023 | 0.042655 | 2.133915 | 0.032849702 | 0.698983806 |
| *ZDHHC4* | 0.37938 | 0.177907 | 2.132463 | 0.032968793 | 0.698983806 |
| *PLGRKT* | -0.575713 | 0.270106 | -2.131431 | 0.033053622 | 0.698983806 |
| *KDM6B* | 0.501702 | 0.235467 | 2.130664 | 0.033116805 | NA |
| *DNAH1* | -1.07023 | 0.502309 | -2.130623 | 0.033120196 | NA |
| *CMTM7* | -0.355945 | 0.167258 | -2.128114 | 0.033327602 | 0.698983806 |
| *LINC00926* | -0.360153 | 0.169317 | -2.127095 | 0.033412228 | 0.698983806 |
| *LY9* | 0.823538 | 0.38766 | 2.124383 | 0.033638117 | 0.699049087 |
| *COPS2* | 0.508754 | 0.239593 | 2.123406 | 0.033719822 | NA |
| *ATP6V1C1* | 0.538654 | 0.253728 | 2.122956 | 0.033757554 | NA |
| *USP31* | -1.315533 | 0.619971 | -2.121926 | 0.033843959 | NA |
| *ENSG00000233184* | 0.724912 | 0.341725 | 2.121329 | 0.033894142 | NA |
| *ATP6V0E2* | -0.727399 | 0.343179 | -2.119589 | 0.03404073 | NA |
| *ZBTB20* | 0.421772 | 0.199106 | 2.11833 | 0.034147139 | 0.704958699 |
| *WDR76* | -0.702221 | 0.331709 | -2.116977 | 0.034261838 | NA |
| *HSBP1* | 0.348161 | 0.16466 | 2.114427 | 0.034478825 | 0.705977608 |
| *DGLUCY* | 0.459021 | 0.217131 | 2.114024 | 0.034513219 | NA |
| *CCT8* | 0.50686 | 0.239937 | 2.112467 | 0.034646447 | 0.705977608 |
| *TM7SF2* | -0.718787 | 0.34039 | -2.111659 | 0.034715689 | NA |
| *SYPL1* | 0.307146 | 0.145706 | 2.107992 | 0.035031668 | 0.70853716 |
| *A1BG.AS1* | -0.779226 | 0.369975 | -2.10616 | 0.035190421 | NA |
| *CD82* | -0.281067 | 0.13348 | -2.105691 | 0.03523122 | 0.70853716 |
| *RCHY1* | 0.532909 | 0.253371 | 2.103271 | 0.035442104 | NA |
| *GPR137* | -0.394081 | 0.187373 | -2.103187 | 0.035449437 | 0.70853716 |
| *ZBED4* | 0.964253 | 0.459219 | 2.099769 | 0.035749138 | NA |
| *KLHL24* | 0.401475 | 0.19137 | 2.0979 | 0.035914002 | 0.713279364 |
| *KCNK6* | 0.647335 | 0.309418 | 2.092103 | 0.03642926 | NA |
| *SCAI* | 0.433206 | 0.207132 | 2.091451 | 0.036487654 | NA |
| *AKAP17A* | 0.417894 | 0.199831 | 2.091239 | 0.036506633 | 0.714258789 |
| *BHLHE40* | -0.340186 | 0.162738 | -2.090391 | 0.036582658 | 0.714258789 |
| *JADE1* | 0.429627 | 0.205594 | 2.089684 | 0.036646165 | 0.714258789 |
| *EXOC3.AS1* | -0.733095 | 0.351173 | -2.087561 | 0.036837486 | NA |
| *GKAP1* | 0.542437 | 0.259904 | 2.087067 | 0.036882099 | NA |
| *STK17A* | 0.271012 | 0.130178 | 2.081853 | 0.037355925 | 0.723598096 |
| *ENSG00000279483* | -0.321927 | 0.154885 | -2.078482 | 0.037664943 | 0.724331578 |
| *NIFK* | 0.398885 | 0.191947 | 2.078097 | 0.037700446 | NA |
| *PTPMT1* | -0.39142 | 0.188356 | -2.078081 | 0.037701893 | NA |
| *TMED10* | -0.347678 | 0.167445 | -2.076369 | 0.037859879 | 0.724331578 |
| *ODF3B* | -0.593033 | 0.285878 | -2.074425 | 0.038039827 | NA |

| | | | | | |
|---|---|---|---|---|---|
| *XBP1* | 0.612489 | 0.295352 | 2.073762 | 0.038101391 | NA |
| *ATAD2B* | -0.718112 | 0.346591 | -2.071932 | 0.03827175 | NA |
| *C9orf40* | 1.266525 | 0.611473 | 2.071269 | 0.038333655 | NA |
| *KRCC1* | 0.374971 | 0.181345 | 2.067725 | 0.038665915 | 0.724331578 |
| *TMEM70* | 0.380905 | 0.184318 | 2.066565 | 0.038775143 | 0.724331578 |
| *RPL32* | 0.104419 | 0.050537 | 2.066198 | 0.038809787 | 0.724331578 |
| *SNAPC4* | -0.950698 | 0.461235 | -2.061203 | 0.039283663 | NA |
| *UBE2N* | 0.248752 | 0.120712 | 2.060714 | 0.039330347 | 0.724331578 |
| *TMEM179B* | -0.263375 | 0.127834 | -2.060298 | 0.039370057 | 0.724331578 |
| *TRAF1* | -0.812658 | 0.394496 | -2.059991 | 0.039399369 | NA |
| *EIF4H* | -0.237889 | 0.115493 | -2.059769 | 0.03942061 | 0.724331578 |
| *ARIH2* | 0.391899 | 0.190457 | 2.057677 | 0.03962116 | NA |
| *MSANTD2* | 0.994508 | 0.483476 | 2.056997 | 0.039686557 | NA |
| *RPS16* | 0.122027 | 0.059374 | 2.055239 | 0.039855947 | 0.724331578 |
| *HINT2* | -0.290594 | 0.141521 | -2.053359 | 0.040037786 | 0.724331578 |
| *PKM* | -0.28655 | 0.139625 | -2.052281 | 0.040142399 | 0.724331578 |
| *ATG16L2* | -0.428955 | 0.20903 | -2.052124 | 0.040157659 | NA |
| *LDHA* | -0.383699 | 0.186982 | -2.052061 | 0.040163701 | 0.724331578 |
| *C2orf69* | 0.504298 | 0.245864 | 2.051128 | 0.040254441 | NA |
| *FOXO3* | 0.563727 | 0.274849 | 2.051043 | 0.040262726 | NA |
| *NDUFAF7* | -0.569285 | 0.277944 | -2.0482 | 0.040540368 | NA |
| *MSL2* | 0.529244 | 0.258426 | 2.047953 | 0.040564635 | NA |
| *MESD* | 0.358592 | 0.175158 | 2.047248 | 0.040633708 | NA |
| *GM2A* | 0.596525 | 0.291818 | 2.044165 | 0.040937212 | 0.734062701 |
| *TSPYL1* | -0.319823 | 0.156779 | -2.039957 | 0.04135462 | 0.734741573 |
| *CLEC4A* | -1.280321 | 0.628309 | -2.037725 | 0.04157748 | NA |
| *RPL11* | 0.096874 | 0.047564 | 2.036726 | 0.041677501 | 0.734741573 |
| *LSR* | 0.851386 | 0.418164 | 2.036008 | 0.041749599 | NA |
| *ENSG00000241490* | 0.735201 | 0.361432 | 2.034132 | 0.04193833 | NA |
| *NUP210* | 0.371336 | 0.18256 | 2.034047 | 0.04194682 | NA |
| *MARCHF5* | 0.53725 | 0.264358 | 2.032279 | 0.042125404 | NA |
| *C17orf49* | -0.493849 | 0.243047 | -2.031912 | 0.042162551 | NA |
| *LYRM7* | 0.327975 | 0.161412 | 2.031911 | 0.04216266 | 0.734741573 |
| *HMGB2* | -0.287986 | 0.141806 | -2.030843 | 0.042270934 | 0.734741573 |
| *EIF3J.DT* | 0.322452 | 0.158914 | 2.029091 | 0.042449009 | 0.734741573 |
| *RGS10* | 0.525015 | 0.258774 | 2.028854 | 0.042473145 | 0.734741573 |
| *CBWD3* | -0.532876 | 0.262812 | -2.02759 | 0.042602075 | NA |
| *SAP18* | -0.179306 | 0.088483 | -2.02644 | 0.042719666 | 0.734741573 |
| *CLMN* | 0.77198 | 0.381218 | 2.025034 | 0.042863893 | NA |
| *SEC13* | -0.425312 | 0.210056 | -2.024758 | 0.042892263 | NA |
| *AMD1* | 0.357958 | 0.176883 | 2.023691 | 0.043001953 | 0.734741573 |
| *LIMD2* | -0.20878 | 0.103208 | -2.022911 | 0.043082361 | 0.734741573 |
| *DYNLT1* | 0.361132 | 0.178802 | 2.019724 | 0.043412011 | NA |
| *RPL24* | 0.098783 | 0.048944 | 2.018303 | 0.043559756 | 0.73766782 |
| *PCM1* | 0.367383 | 0.182168 | 2.016726 | 0.043724096 | 0.73766782 |
| *ENSG00000170161* | -0.582861 | 0.289239 | -2.015152 | 0.043888704 | NA |
| *ZNF12* | 0.67141 | 0.333183 | 2.015136 | 0.043890379 | NA |

| | | | | | |
|---|---|---|---|---|---|
| *DYNLL1* | 0.267539 | 0.133164 | 2.009091 | 0.044527454 | 0.747204011 |
| *MGME1* | 0.425643 | 0.212231 | 2.005569 | 0.044902285 | NA |
| *FLAD1* | -0.579028 | 0.288858 | -2.00454 | 0.045012237 | NA |
| *NR6A1* | -1.515757 | 0.756448 | -2.003783 | 0.045093298 | NA |
| *PPFIBP1* | -0.908897 | 0.453633 | -2.003595 | 0.045113512 | NA |
| *NELFE* | -0.266902 | 0.133322 | -2.001933 | 0.045291981 | 0.748357928 |
| *KCTD5* | 0.573497 | 0.286638 | 2.00077 | 0.045417199 | NA |
| *IQCH.AS1* | 0.823908 | 0.411829 | 2.000605 | 0.045434984 | NA |
| *PRAG1* | 1.086219 | 0.543807 | 1.997435 | 0.045777983 | NA |
| *ZFPL1* | -0.477726 | 0.239302 | -1.99633 | 0.045898048 | NA |
| *PCNX4* | 0.355185 | 0.178018 | 1.995216 | 0.046019357 | 0.748357928 |
| *LNPEP* | 0.307306 | 0.154099 | 1.994214 | 0.046128687 | 0.748357928 |
| *TMPO.AS1* | -1.348536 | 0.676641 | -1.992984 | 0.046263146 | NA |
| *UBE2A* | -0.28647 | 0.143789 | -1.992292 | 0.046339029 | 0.748357928 |
| *RGS19* | 0.349943 | 0.175747 | 1.991177 | 0.046461394 | 0.748357928 |
| *RPS18* | 0.17632 | 0.088567 | 1.990814 | 0.046501301 | 0.748357928 |
| *JAK3* | -0.48513 | 0.24383 | -1.989624 | 0.046632361 | 0.748357928 |
| *GNB5* | 0.287794 | 0.144663 | 1.989404 | 0.046656613 | 0.748357928 |
| *THEM4* | -0.761094 | 0.382941 | -1.987497 | 0.04686735 | NA |
| *SAMD10* | -1.004783 | 0.506011 | -1.985693 | 0.047067422 | NA |
| *MYO1G* | 0.340083 | 0.171322 | 1.985048 | 0.047139186 | 0.748357928 |
| *GPS2* | -0.396858 | 0.199946 | -1.98483 | 0.047163381 | 0.748357928 |
| *STK16* | -0.331904 | 0.16756 | -1.980812 | 0.047612306 | 0.748357928 |
| *PRCP* | -0.425816 | 0.215063 | -1.979956 | 0.047708528 | NA |
| *ZNF775* | -1.231133 | 0.62186 | -1.97976 | 0.047730509 | NA |
| *CD48* | 0.248769 | 0.125692 | 1.979192 | 0.047794424 | 0.748357928 |
| *PTPN2* | 0.286286 | 0.144693 | 1.97858 | 0.047863262 | 0.748357928 |
| *H1FX* | -0.870562 | 0.440459 | -1.976489 | 0.048099418 | NA |
| *RPS27A* | 0.104657 | 0.052961 | 1.976121 | 0.048141022 | 0.748357928 |
| *TMEM156* | 0.462051 | 0.233992 | 1.974643 | 0.048308711 | 0.748357928 |
| *TMEM251* | 0.507835 | 0.257214 | 1.974366 | 0.048340107 | NA |
| *EXOSC8* | 0.329322 | 0.166852 | 1.973735 | 0.048411937 | 0.748357928 |
| *MOV10* | 0.834246 | 0.422751 | 1.973376 | 0.048452762 | NA |
| *ERAP1* | 0.523951 | 0.265538 | 1.973163 | 0.048476979 | NA |
| *ENSG00000280007* | -1.04824 | 0.531819 | -1.971048 | 0.048718379 | NA |
| *PTP4A3* | 1.212435 | 0.615729 | 1.969106 | 0.048940952 | NA |
| *AK6* | 0.325493 | 0.165349 | 1.968523 | 0.049007916 | 0.753857068 |
| *S100A13* | -0.887426 | 0.451977 | -1.96343 | 0.049596211 | NA |
| *FCGR2B* | 0.735124 | 0.374712 | 1.961836 | 0.049781586 | NA |
| *NTHL1* | -0.505914 | 0.257899 | -1.961676 | 0.049800174 | NA |

| Cluster 8 | log2FC | lfcSE | stat | pvalue | padj |
|---|---|---|---|---|---|
| *ARRDC3* | -1.217494 | 0.178462 | -6.822133 | 8.97E-12 | 3.94E-08 |
| *GPR183* | 0.795786 | 0.127664 | 6.233426 | 4.56E-10 | 1.00E-06 |
| *CKS2* | 1.291314 | 0.212905 | 6.0652 | 1.32E-09 | 1.93E-06 |
| *RPL8* | 0.404548 | 0.068476 | 5.907912 | 3.46E-09 | 3.80E-06 |
| *RPS5* | 0.359067 | 0.064754 | 5.54507 | 2.94E-08 | 2.16E-05 |
| *RPL19* | 0.184563 | 0.033289 | 5.544213 | 2.95E-08 | 2.16E-05 |
| *NPM1* | 0.262409 | 0.048778 | 5.379614 | 7.46E-08 | 4.68E-05 |
| *IL4R* | -0.645521 | 0.12056 | -5.354341 | 8.59E-08 | 4.71E-05 |
| *RENBP* | 1.070374 | 0.206719 | 5.17792 | 2.24E-07 | 0.000109444 |
| *NOP10* | 0.39864 | 0.077639 | 5.134518 | 2.83E-07 | 0.000124179 |
| *ENSG00000271204* | 0.588149 | 0.116237 | 5.059895 | 4.19E-07 | 0.000167413 |
| *ENSG00000272211* | -1.425394 | 0.283833 | -5.021947 | 5.12E-07 | 0.000187125 |
| *RPS9* | 0.393657 | 0.08104 | 4.85753 | 1.19E-06 | 0.000401378 |
| *ZNF107* | 1.251603 | 0.259418 | 4.824653 | 1.40E-06 | 0.000439775 |
| *UQCRH* | 0.274445 | 0.057058 | 4.809932 | 1.51E-06 | 0.000441872 |
| *CD48* | 0.417354 | 0.089388 | 4.669027 | 3.03E-06 | 0.00083034 |
| *PHF21A* | -0.634812 | 0.138881 | -4.570911 | 4.86E-06 | 0.001254015 |
| *GATM* | 2.07213 | 0.455871 | 4.545434 | 5.48E-06 | NA |
| *CD79B* | -0.384521 | 0.085029 | -4.52221 | 6.12E-06 | 0.001492535 |
| *RPS23* | 0.212636 | 0.047223 | 4.502789 | 6.71E-06 | 0.001528853 |
| *RPL24* | 0.258941 | 0.05761 | 4.49475 | 6.97E-06 | 0.001528853 |
| *RPL9* | 0.238295 | 0.053381 | 4.464038 | 8.04E-06 | 0.001681356 |
| *BLVRB* | 0.986088 | 0.221522 | 4.451422 | 8.53E-06 | 0.00170219 |
| *RPL30* | 0.200286 | 0.045132 | 4.43775 | 9.09E-06 | 0.001735081 |
| *NFKBIA* | -0.460984 | 0.105277 | -4.378765 | 1.19E-05 | 0.002183183 |
| *ENSG00000259888* | 2.25305 | 0.526238 | 4.281426 | 1.86E-05 | NA |
| *HNRNPDL* | -0.305029 | 0.071915 | -4.241538 | 2.22E-05 | 0.003898211 |
| *CIRBP* | -0.322846 | 0.076738 | -4.207118 | 2.59E-05 | 0.004143345 |
| *RPS15* | 0.154384 | 0.036701 | 4.20652 | 2.59E-05 | 0.004143345 |
| *RPL34* | 0.185431 | 0.044196 | 4.195705 | 2.72E-05 | 0.004143345 |
| *TIMM10* | -0.623214 | 0.148586 | -4.194308 | 2.74E-05 | 0.004143345 |
| *ZBED2* | 2.386924 | 0.573599 | 4.161311 | 3.16E-05 | NA |
| *HRK* | -1.99613 | 0.48311 | -4.131832 | 3.60E-05 | 0.005266291 |
| *FAM102A* | 0.995495 | 0.244181 | 4.076869 | 4.56E-05 | 0.006464078 |
| *C16orf74* | -0.706452 | 0.174051 | -4.058885 | 4.93E-05 | 0.006764375 |
| *DEF8* | -0.710954 | 0.175525 | -4.050451 | 5.11E-05 | 0.006800381 |
| *RPL38* | 0.232586 | 0.057815 | 4.022916 | 5.75E-05 | 0.007421941 |
| *BHLHE41* | 2.403203 | 0.599118 | 4.011237 | 6.04E-05 | NA |
| *GLUL* | -0.932084 | 0.233066 | -3.999229 | 6.35E-05 | 0.007775757 |
| *RAB9A* | 0.840757 | 0.210272 | 3.998427 | 6.38E-05 | 0.007775757 |
| *COX17* | 0.386403 | 0.097603 | 3.958947 | 7.53E-05 | 0.008750432 |
| *RNF130* | 0.771157 | 0.194861 | 3.957482 | 7.57E-05 | 0.008750432 |
| *RPL36* | 0.161432 | 0.040922 | 3.944843 | 7.99E-05 | 0.00897768 |
| *EEF1B2* | 0.350161 | 0.088968 | 3.935797 | 8.29E-05 | 0.00897768 |
| *DYNLL1* | 0.585873 | 0.148958 | 3.933132 | 8.38E-05 | 0.00897768 |
| *RPS18* | 0.232238 | 0.05933 | 3.91433 | 9.07E-05 | 0.009223867 |

| | | | | | |
|---|---|---|---|---|---|
| *TMEM204* | -2.749617 | 0.702557 | -3.913731 | 9.09E-05 | NA |
| *RPL32* | 0.16719 | 0.042732 | 3.912527 | 9.13E-05 | 0.009223867 |
| *RPL26* | 0.195267 | 0.049946 | 3.909601 | 9.24E-05 | 0.009223867 |
| *LPIN1* | 0.766975 | 0.199106 | 3.8521 | 0.00011711 | 0.011424634 |
| *MIF* | -0.398733 | 0.103734 | -3.843795 | 0.00012115 | 0.011561584 |
| *CYTIP* | 0.246049 | 0.064642 | 3.806321 | 0.00014105 | 0.013174581 |
| *RACK1* | 0.199933 | 0.052678 | 3.795376 | 0.00014742 | 0.013482777 |
| *RPS16* | 0.261779 | 0.069298 | 3.77758 | 0.00015836 | 0.014187756 |
| *ZNF385A* | -1.756664 | 0.46519 | -3.77623 | 0.00015922 | NA |
| *RPL37* | 0.158753 | 0.04252 | 3.733619 | 0.00018875 | 0.01657204 |
| *CDC42* | -0.299827 | 0.081159 | -3.694338 | 0.00022046 | 0.018630438 |
| *GSTO1* | 0.524447 | 0.141969 | 3.694086 | 0.00022068 | 0.018630438 |
| *MHENCR* | -0.445944 | 0.121253 | -3.677786 | 0.00023527 | 0.01948719 |
| *HSPE1* | 0.327608 | 0.089297 | 3.668766 | 0.00024372 | 0.019813871 |
| *ARL17A* | -0.827737 | 0.226323 | -3.65733 | 0.00025486 | NA |
| *COMMD6* | 0.192184 | 0.052588 | 3.654518 | 0.00025767 | 0.0205664 |
| *RPSA* | 0.32729 | 0.089872 | 3.641736 | 0.00027081 | 0.02122921 |
| *RHBDD2* | -0.76753 | 0.212399 | -3.613624 | 0.00030195 | NA |
| *RPL13* | 0.212496 | 0.058924 | 3.606294 | 0.0003106 | 0.023921725 |
| *ATP6V1C1* | 0.914417 | 0.253945 | 3.600845 | 0.00031719 | NA |
| *KDM4B* | 0.887889 | 0.24667 | 3.599501 | 0.00031883 | 0.024131978 |
| *SLC50A1* | -0.391264 | 0.109143 | -3.584883 | 0.00033723 | 0.025092155 |
| *GATD3A* | 4.009907 | 1.119135 | 3.583042 | 0.00033962 | NA |
| *DDX6* | -0.447103 | 0.125117 | -3.573476 | 0.00035227 | 0.025716228 |
| *IGHD* | -0.482851 | 0.135262 | -3.569742 | 0.00035733 | 0.025716228 |
| *H3F3B* | -0.326028 | 0.091991 | -3.544112 | 0.00039394 | 0.02768146 |
| *RPL23A* | 0.202605 | 0.057202 | 3.541903 | 0.00039725 | 0.02768146 |
| *NSMAF* | 1.222799 | 0.346017 | 3.533932 | 0.00040943 | 0.027990041 |
| *RPS12* | 0.249997 | 0.070807 | 3.530669 | 0.00041451 | 0.027990041 |
| *RPS3* | 0.223737 | 0.063441 | 3.526679 | 0.00042081 | 0.027990041 |
| *CLEC2B* | 0.391386 | 0.111302 | 3.516414 | 0.00043742 | 0.028660695 |
| *HVCN1* | -0.303504 | 0.086506 | -3.508481 | 0.00045067 | 0.029094991 |
| *IGHG1* | 1.38147 | 0.396309 | 3.485838 | 0.0004906 | NA |
| *ENSG00000273319* | 0.609357 | 0.175188 | 3.478297 | 0.00050461 | 0.031370701 |
| *RPS15A* | 0.210148 | 0.060423 | 3.477931 | 0.0005053 | 0.031370701 |
| *CHORDC1* | 0.731069 | 0.210268 | 3.476839 | 0.00050736 | 0.031370701 |
| *DDX58* | -0.817431 | 0.23527 | -3.474433 | 0.00051193 | NA |
| *RPS13* | 0.166302 | 0.048221 | 3.44873 | 0.00056323 | 0.034341307 |
| *TNFRSF13B* | 1.086378 | 0.31639 | 3.433667 | 0.00059548 | 0.035810116 |
| *KANK2* | -2.061716 | 0.60185 | -3.425633 | 0.00061337 | NA |
| *RPL13A* | 0.186535 | 0.054478 | 3.424043 | 0.00061697 | 0.03660121 |
| *ANP32B* | 0.255031 | 0.074645 | 3.416593 | 0.0006341 | 0.037115961 |
| *SRSF6* | -0.427 | 0.125787 | -3.394631 | 0.00068721 | 0.03969545 |
| *STX7* | 0.271701 | 0.080448 | 3.377325 | 0.00073195 | 0.041584408 |
| *SLC5A3* | -0.654081 | 0.19396 | -3.372255 | 0.00074556 | 0.041584408 |
| *MT.CO1* | 0.246372 | 0.073142 | 3.368392 | 0.00075608 | 0.041584408 |
| *RPL7A* | 0.24526 | 0.072826 | 3.367765 | 0.0007578 | 0.041584408 |

| | | | | | |
|---|---|---|---|---|---|
| *EYA3* | 1.064287 | 0.316477 | 3.362923 | 0.00077122 | NA |
| *FAM177B* | 0.864073 | 0.257921 | 3.350138 | 0.00080771 | 0.043409957 |
| *ZNF580* | -0.448836 | 0.134052 | -3.348216 | 0.00081334 | 0.043409957 |
| *ENSG00000282988* | 1.03827 | 0.310122 | 3.347944 | 0.00081413 | NA |
| *ERP29* | 0.209545 | 0.062631 | 3.345706 | 0.00082074 | 0.043409957 |
| *TULP4* | -0.478669 | 0.144348 | -3.316087 | 0.00091287 | 0.047708478 |
| *BCL7A* | -0.758484 | 0.228985 | -3.312381 | 0.00092506 | 0.047776398 |
| *ADPRH* | 1.80564 | 0.545422 | 3.31054 | 0.00093116 | NA |
| *IGHG3* | 1.795687 | 0.544381 | 3.298585 | 0.00097174 | NA |
| *RPS6* | 0.213225 | 0.064681 | 3.296589 | 0.00097867 | 0.049254035 |
| *ATP5IF1* | -0.250812 | 0.076091 | -3.296203 | 0.00098001 | 0.049254035 |
| *RPL35* | 0.166079 | 0.050504 | 3.288404 | 0.00100757 | 0.049254035 |
| *MGAT1* | 0.516778 | 0.157222 | 3.286935 | 0.00101284 | 0.049254035 |
| *RPL3* | 0.1851 | 0.056326 | 3.286259 | 0.00101528 | 0.049254035 |
| *RPS8* | 0.266669 | 0.081186 | 3.28468 | 0.00102098 | 0.049254035 |
| *LNPEP* | 0.483267 | 0.147617 | 3.273787 | 0.00106117 | 0.050636029 |
| *RPL11* | 0.148564 | 0.045437 | 3.269665 | 0.00107675 | 0.050827271 |
| *IFI44L* | 1.927422 | 0.590733 | 3.262762 | 0.00110332 | NA |
| *MINDY2* | 0.618274 | 0.1897 | 3.259221 | 0.00111719 | 0.052174963 |
| *ATP6V0E2* | -1.165972 | 0.358835 | -3.249325 | 0.00115679 | 0.053071878 |
| *TOMM7* | 0.28808 | 0.088684 | 3.248398 | 0.00116057 | 0.053071878 |
| *NT5DC4* | 2.493167 | 0.769642 | 3.239385 | 0.00119788 | NA |
| *RPL10* | 0.216421 | 0.066888 | 3.23558 | 0.00121396 | 0.054819577 |
| *HEXA* | 0.481334 | 0.148869 | 3.233283 | 0.00122376 | 0.054819577 |
| *RPS3A* | 0.15488 | 0.047989 | 3.227412 | 0.00124915 | 0.054866855 |
| *HHEX* | 0.366127 | 0.113448 | 3.227261 | 0.00124981 | 0.054866855 |
| *CREBL2* | 0.857545 | 0.266345 | 3.219676 | 0.00128336 | NA |
| *RPL37A* | 0.19417 | 0.060577 | 3.205318 | 0.00134913 | 0.058597288 |
| *NCF1* | -0.271932 | 0.084937 | -3.201585 | 0.00136674 | 0.058597288 |
| *UCP2* | -0.472651 | 0.147709 | -3.199883 | 0.00137483 | 0.058597288 |
| *NDUFB11* | 0.305684 | 0.095813 | 3.190407 | 0.00142072 | 0.059592037 |
| *RPL7* | 0.19423 | 0.060912 | 3.188716 | 0.00142906 | 0.059592037 |
| *TMEM140* | 0.645271 | 0.202487 | 3.186733 | 0.0014389 | 0.059592037 |
| *HSD17B4* | 0.708932 | 0.222796 | 3.18198 | 0.00146272 | 0.060012565 |
| *H2AFZ* | 0.230241 | 0.072522 | 3.174781 | 0.0014995 | 0.060951735 |
| *KRTCAP2* | -0.291761 | 0.092105 | -3.16771 | 0.00153645 | 0.061880803 |
| *ENSG00000225885* | -1.094481 | 0.34566 | -3.166354 | 0.00154363 | NA |
| *TAGAP* | -0.589948 | 0.187219 | -3.151111 | 0.00162651 | 0.064912434 |
| *CLK1* | 0.458962 | 0.1458 | 3.147896 | 0.0016445 | 0.065039323 |
| *NAP1L1* | 0.232863 | 0.074402 | 3.129787 | 0.00174933 | 0.068567591 |
| *THRAP3* | -0.322731 | 0.103254 | -3.125599 | 0.00177443 | 0.068935845 |
| *PDE4DIP* | -0.780491 | 0.250517 | -3.115519 | 0.00183622 | NA |
| *MAML3* | -1.554957 | 0.501649 | -3.099689 | 0.00193724 | NA |
| *CERS4* | -0.461364 | 0.148962 | -3.097186 | 0.00195367 | 0.074956823 |
| *SNHG30* | -0.518163 | 0.167382 | -3.095689 | 0.00196356 | 0.074956823 |
| *SLC2A1* | -0.557297 | 0.180877 | -3.081085 | 0.00206248 | 0.078054236 |
| *TCL1B* | -1.154096 | 0.374997 | -3.077618 | 0.00208662 | NA |

| | | | | | |
|---|---|---|---|---|---|
| *FAM117B* | -0.771052 | 0.250642 | -3.076303 | 0.00209585 | 0.078113118 |
| *FAU* | 0.108294 | 0.035209 | 3.075766 | 0.00209962 | 0.078113118 |
| *DDX5* | -0.37573 | 0.122465 | -3.068061 | 0.00215452 | 0.079046774 |
| *RPL21* | 0.170037 | 0.055437 | 3.067201 | 0.00216073 | 0.079046774 |
| *RPL22* | 0.151871 | 0.049592 | 3.06238 | 0.00219585 | 0.079667471 |
| *LINC00847* | 0.644298 | 0.210917 | 3.054747 | 0.00225251 | 0.081053349 |
| *LAMC1* | 1.415897 | 0.463668 | 3.053688 | 0.00226047 | NA |
| *DNAJA1* | 0.450464 | 0.148032 | 3.043022 | 0.00234215 | 0.083593939 |
| *EIF3K* | 0.255737 | 0.08477 | 3.016825 | 0.00255437 | 0.08838612 |
| *HMGB2* | -0.428466 | 0.14208 | -3.015662 | 0.00256419 | 0.08838612 |
| *CFLAR* | 0.390029 | 0.129349 | 3.015325 | 0.00256704 | 0.08838612 |
| *ITM2C* | 0.52249 | 0.173295 | 3.015026 | 0.00256957 | 0.08838612 |
| *SLC16A7* | 0.588206 | 0.195149 | 3.01414 | 0.00257709 | 0.08838612 |
| *C15orf61* | 0.6099 | 0.202569 | 3.010828 | 0.00260537 | 0.088448968 |
| *THUMPD3.AS1* | -0.285959 | 0.095028 | -3.009217 | 0.00261922 | 0.088448968 |
| *ZNF318* | 0.5283 | 0.176139 | 2.999342 | 0.00270564 | 0.090669754 |
| *USF1* | -1.014657 | 0.338345 | -2.998886 | 0.00270969 | NA |
| *ENSG00000273448* | -0.798308 | 0.266559 | -2.994866 | 0.00274566 | NA |
| *SLC38A2* | 0.409797 | 0.13726 | 2.985558 | 0.00283061 | 0.092949339 |
| *MRPS36* | 0.450947 | 0.151134 | 2.983754 | 0.00284736 | 0.092949339 |
| *ST13* | 0.194851 | 0.065321 | 2.982995 | 0.00285443 | 0.092949339 |
| *RPL18* | 0.16029 | 0.053777 | 2.980645 | 0.00287642 | 0.092949339 |
| *RNF146* | 0.595573 | 0.199835 | 2.980315 | 0.00287952 | 0.092949339 |
| *PCNX4* | 0.488015 | 0.164161 | 2.972785 | 0.00295111 | 0.093243854 |
| *THOC3* | -0.61551 | 0.207049 | -2.972776 | 0.0029512 | 0.093243854 |
| *CRK* | 0.619961 | 0.208555 | 2.972654 | 0.00295237 | 0.093243854 |
| *APBB2* | -0.927301 | 0.312349 | -2.968801 | 0.00298965 | NA |
| *ENSG00000254397* | 1.284079 | 0.433249 | 2.963836 | 0.0030383 | NA |
| *RPS28* | 0.220297 | 0.074359 | 2.962602 | 0.00305051 | 0.095655295 |
| *L3HYPDH* | -0.8042 | 0.271691 | -2.959984 | 0.00307655 | NA |
| *PET100* | -0.292324 | 0.099029 | -2.951893 | 0.00315832 | 0.095989574 |
| *HSH2D* | -0.460622 | 0.156103 | -2.950767 | 0.00316986 | 0.095989574 |
| *TRA2B* | -0.272252 | 0.092323 | -2.948894 | 0.00318914 | 0.095989574 |
| *TTC32* | -0.974175 | 0.330421 | -2.948281 | 0.00319546 | 0.095989574 |
| *UBE2J2* | -0.47339 | 0.160605 | -2.947536 | 0.00320317 | 0.095989574 |
| *UBA52* | 0.113328 | 0.038449 | 2.947507 | 0.00320348 | 0.095989574 |
| *TCTN1* | -0.799977 | 0.271503 | -2.946471 | 0.00321423 | 0.095989574 |
| *CREBRF* | 0.363545 | 0.123567 | 2.942083 | 0.00326012 | 0.096702219 |
| *RPLP2* | 0.21293 | 0.072486 | 2.937526 | 0.00330843 | 0.097476517 |
| *NAB2* | 1.188394 | 0.405107 | 2.933532 | 0.00335129 | NA |
| *SNHG9* | -0.402351 | 0.13729 | -2.930659 | 0.00338244 | 0.097726963 |
| *RPL31* | 0.170144 | 0.058059 | 2.930531 | 0.00338383 | 0.097726963 |
| *IL2RG* | -0.473417 | 0.16172 | -2.927376 | 0.00341835 | 0.097726963 |
| *TBCA* | 0.261583 | 0.08937 | 2.926983 | 0.00342268 | 0.097726963 |
| *TMED2* | -0.349307 | 0.119361 | -2.926478 | 0.00342824 | 0.097726963 |
| *FBXO6* | -1.11263 | 0.380435 | -2.924628 | 0.00344868 | NA |
| *RPL5* | 0.213283 | 0.072967 | 2.922999 | 0.00346678 | 0.098188071 |

| | | | | | |
|---|---|---|---|---|---|
| *HLA.A* | -0.448438 | 0.15394 | -2.913073 | 0.00357891 | 0.099666718 |
| *ZCCHC7* | -0.399721 | 0.137289 | -2.911522 | 0.00359673 | 0.099666718 |
| *RPL10A* | 0.303686 | 0.104379 | 2.909454 | 0.00362061 | 0.099666718 |
| *FLCN* | 0.534245 | 0.183707 | 2.908137 | 0.00363589 | 0.099666718 |
| *ERGIC2* | 0.427973 | 0.147209 | 2.907241 | 0.00364632 | 0.099666718 |
| *NCBP2AS2* | -0.528589 | 0.181977 | -2.904704 | 0.00367601 | 0.099666718 |
| *ZFAS1* | 0.235 | 0.080908 | 2.904542 | 0.00367791 | 0.099666718 |
| *RPL29* | 0.195076 | 0.067213 | 2.902366 | 0.00370356 | 0.099746089 |
| *RPL28* | 0.200354 | 0.069233 | 2.893898 | 0.00380492 | 0.101851258 |
| *CPM* | -0.98715 | 0.342269 | -2.884133 | 0.00392493 | NA |
| *FXYD7* | 0.788078 | 0.273311 | 2.883453 | 0.00393341 | 0.104652547 |
| *CLECL1* | 1.122528 | 0.389918 | 2.878883 | 0.00399087 | NA |
| *MX1* | 0.940284 | 0.327446 | 2.871569 | 0.0040844 | NA |
| *LRWD1* | 0.670971 | 0.233741 | 2.870575 | 0.00409726 | NA |
| *CYB5A* | -0.548135 | 0.190966 | -2.870327 | 0.00410048 | 0.108374098 |
| *ENSG00000007237* | -1.526777 | 0.532067 | -2.869519 | 0.00411096 | NA |
| *NAA10* | -0.479825 | 0.167267 | -2.868621 | 0.00412266 | 0.108374098 |
| *CCDC25* | 0.449436 | 0.156808 | 2.866161 | 0.00415482 | 0.10856951 |
| *MIR29B2CHG* | 0.623473 | 0.21779 | 2.862731 | 0.00420007 | 0.109102435 |
| *SLA* | 0.635674 | 0.222539 | 2.856462 | 0.00428392 | 0.110625846 |
| *SNAP23* | -0.328977 | 0.115454 | -2.849428 | 0.00437979 | 0.112440123 |
| *FKBP5* | 1.043427 | 0.366638 | 2.845933 | 0.00442816 | NA |
| *CCNG2* | 0.638149 | 0.224242 | 2.845804 | 0.00442995 | 0.113066714 |
| *C17orf58* | 1.951667 | 0.686577 | 2.842604 | 0.00447467 | NA |
| *LINC00926* | -0.351679 | 0.123933 | -2.837659 | 0.00454457 | 0.115321805 |
| *C7orf50* | -0.412248 | 0.14543 | -2.834681 | 0.00458715 | 0.115733162 |
| *NUDT14* | 0.578658 | 0.204755 | 2.826096 | 0.00471191 | 0.118201596 |
| *CCDC144A* | 2.017015 | 0.715657 | 2.818412 | 0.00482618 | NA |
| *SMIM20* | 0.434261 | 0.154236 | 2.815557 | 0.00486927 | 0.119148619 |
| *DENND2D* | 0.560654 | 0.199188 | 2.814693 | 0.00488238 | 0.119148619 |
| *SKAP2* | 0.40548 | 0.144069 | 2.814479 | 0.00488564 | 0.119148619 |
| *CAPG* | 0.376022 | 0.133609 | 2.814347 | 0.00488765 | 0.119148619 |
| *IGHA1* | -0.440108 | 0.156493 | -2.812312 | 0.00491868 | 0.119148619 |
| *PDE7B* | -0.888183 | 0.315847 | -2.812071 | 0.00492237 | NA |
| *YWHAE* | 0.378263 | 0.134556 | 2.811185 | 0.00493594 | 0.119148619 |
| *RPL35A* | 0.182555 | 0.064944 | 2.810943 | 0.00493965 | 0.119148619 |
| *RPS27A* | 0.117945 | 0.04205 | 2.804847 | 0.00503405 | 0.120762097 |
| *NOTCH2* | 0.48643 | 0.17366 | 2.801055 | 0.00509358 | 0.121526177 |
| *CD22* | 0.565926 | 0.202896 | 2.789244 | 0.00528312 | 0.125217652 |
| *RPS20* | 0.294541 | 0.10565 | 2.787884 | 0.00530535 | 0.125217652 |
| *ZNF606* | -0.631708 | 0.227066 | -2.782042 | 0.00540181 | 0.126812611 |
| *SAR1A* | -0.414871 | 0.149576 | -2.773643 | 0.00554324 | 0.129440628 |
| *ZBTB16* | 1.896645 | 0.684417 | 2.771183 | 0.0055853 | NA |
| *KLK4* | 2.097178 | 0.757052 | 2.770191 | 0.00560234 | NA |
| *MCL1* | -0.387814 | 0.140408 | -2.762051 | 0.00574396 | 0.133417822 |
| *CTNNAL1* | -1.309362 | 0.47423 | -2.761026 | 0.005762 | NA |
| *RPL15* | 0.125514 | 0.045484 | 2.759512 | 0.00578877 | 0.133751077 |

| | | | | | |
|---|---|---|---|---|---|
| *EIF3E* | 0.205428 | 0.074616 | 2.753137 | 0.00590272 | 0.135669821 |
| *AK6* | 0.488757 | 0.177665 | 2.751009 | 0.0059412 | 0.135776419 |
| *ST3GAL5* | 0.542655 | 0.197367 | 2.749468 | 0.00596921 | 0.135776419 |
| *RBM26.AS1* | -1.062394 | 0.386637 | -2.747784 | 0.00599995 | NA |
| *HLA.C* | -0.489031 | 0.178099 | -2.74584 | 0.00603561 | 0.136264219 |
| *CDKN1A* | -0.625944 | 0.228038 | -2.744911 | 0.00605274 | 0.136264219 |
| *ARIH2OS* | -0.997916 | 0.36407 | -2.740998 | 0.00612529 | NA |
| *CBWD2* | -0.850094 | 0.311903 | -2.725508 | 0.00642027 | NA |
| *CD83* | -0.293883 | 0.107879 | -2.724194 | 0.00644586 | 0.144374 |
| *BATF* | 1.361252 | 0.49975 | 2.723867 | 0.00645226 | NA |
| *EPB42* | -0.825033 | 0.303266 | -2.720496 | 0.00651841 | NA |
| *RSRP1* | 0.34076 | 0.125352 | 2.718437 | 0.00655911 | 0.145453486 |
| *LBH* | -0.344096 | 0.126581 | -2.718376 | 0.00656032 | 0.145453486 |
| *ENSG00000167414* | -1.570447 | 0.577844 | -2.717773 | 0.0065723 | NA |
| *KMT2A* | 0.411723 | 0.151575 | 2.716302 | 0.00660157 | 0.145632644 |
| *SENP2* | -0.677381 | 0.249493 | -2.715025 | 0.00662708 | NA |
| *RPL27A* | 0.143485 | 0.052942 | 2.710238 | 0.0067235 | 0.147580761 |
| *FAM32A* | -0.331268 | 0.122434 | -2.70568 | 0.00681647 | 0.148877059 |
| *IFI27L2* | 0.328382 | 0.121505 | 2.702621 | 0.00687951 | 0.149510138 |
| *CDC42SE1* | -0.384156 | 0.142248 | -2.700616 | 0.00692112 | 0.149673539 |
| *CLEC2D* | -0.323012 | 0.119843 | -2.695293 | 0.00703267 | 0.150842777 |
| *FAM184B* | -1.705132 | 0.632758 | -2.694761 | 0.00704391 | 0.150842777 |
| *SDSL* | 1.619495 | 0.601376 | 2.692983 | 0.00708159 | NA |
| *NDUFA13* | -0.266138 | 0.099167 | -2.683734 | 0.0072805 | 0.154762438 |
| *SCAND1* | -0.32201 | 0.120021 | -2.682956 | 0.00729745 | 0.154762438 |
| *WDR11* | -0.653822 | 0.244423 | -2.674959 | 0.00747383 | 0.157740984 |
| *DOK3* | 0.966309 | 0.361262 | 2.674814 | 0.00747707 | NA |
| *LGALS1* | 1.509988 | 0.564889 | 2.67307 | 0.00751605 | 0.157873067 |
| *NPHP3* | 0.580063 | 0.217557 | 2.666262 | 0.00767 | NA |
| *RPS14* | 0.220589 | 0.082841 | 2.6628 | 0.00774935 | 0.1612456 |
| *NMT2* | -0.403234 | 0.151461 | -2.662289 | 0.00776112 | 0.1612456 |
| *PRKCB* | -0.51205 | 0.192438 | -2.660862 | 0.00779409 | 0.1612456 |
| *RPS27* | 0.131641 | 0.049503 | 2.659272 | 0.00783098 | 0.1612456 |
| *TAGLN2* | -0.291718 | 0.10975 | -2.658014 | 0.00786026 | 0.1612456 |
| *LARP1* | 0.450068 | 0.169612 | 2.653514 | 0.00796585 | 0.161568993 |
| *SRSF5* | -0.280454 | 0.105692 | -2.653513 | 0.00796588 | 0.161568993 |
| *CNN3* | -0.764477 | 0.288195 | -2.652643 | 0.00798644 | 0.161568993 |
| *TTC37* | 0.519351 | 0.196378 | 2.644648 | 0.00817759 | 0.164058004 |
| *LYRM7* | 0.377395 | 0.142716 | 2.644374 | 0.00818422 | 0.164058004 |
| *GCSH* | -0.592392 | 0.224302 | -2.641051 | 0.00826493 | NA |
| *KDSR* | 0.567778 | 0.21532 | 2.636903 | 0.00836667 | 0.166953072 |
| *APLP2* | -0.486868 | 0.184777 | -2.634892 | 0.00841642 | 0.167185854 |
| *ADAMTS6* | 0.909157 | 0.345764 | 2.629415 | 0.0085532 | NA |
| *RDH5* | 0.489907 | 0.18637 | 2.628682 | 0.00857165 | 0.169502502 |
| *IZUMO4* | -0.665777 | 0.253458 | -2.626775 | 0.00861983 | 0.169690741 |
| *UGT8* | 1.185548 | 0.451563 | 2.625431 | 0.00865394 | NA |
| *LMO4* | 0.535233 | 0.204148 | 2.621786 | 0.00874704 | NA |

| | | | | | |
|---|---|---|---|---|---|
| *MME* | -2.927514 | 1.117051 | -2.620753 | 0.00877359 | NA |
| *CBLN3* | -0.895808 | 0.341863 | -2.620375 | 0.00878332 | NA |
| *NEIL1* | -0.652103 | 0.249121 | -2.617619 | 0.00885457 | 0.173533753 |
| *RHOF* | 0.377704 | 0.144594 | 2.61217 | 0.00899696 | 0.175540709 |
| *NREP* | -0.776085 | 0.297286 | -2.610569 | 0.00903916 | NA |
| *LSM2* | 0.265121 | 0.102139 | 2.59569 | 0.00944012 | 0.183372272 |
| *WASF1* | -1.154465 | 0.444884 | -2.594981 | 0.0094596 | NA |
| *HIP1R* | -0.493956 | 0.190455 | -2.593557 | 0.00949888 | 0.183700872 |
| *TMEM191B* | -0.762152 | 0.29419 | -2.590675 | 0.0095788 | NA |
| *RPL23* | 0.13102 | 0.050612 | 2.588713 | 0.00963353 | 0.18548766 |
| *ENSG00000251364* | 1.27111 | 0.491199 | 2.587768 | 0.00966001 | NA |
| *TMEM205* | -0.529083 | 0.204727 | -2.584343 | 0.00975648 | 0.187034608 |
| *ENSG00000187186* | 1.408872 | 0.545715 | 2.581701 | 0.00983147 | NA |
| *SWAP70* | 0.367357 | 0.14237 | 2.5803 | 0.00987146 | 0.187705356 |
| *VDAC2* | 0.247381 | 0.095934 | 2.578656 | 0.00991854 | 0.187705356 |
| *SESN3* | 0.380322 | 0.147491 | 2.578615 | 0.00991974 | 0.187705356 |
| *HCST* | 1.012116 | 0.392653 | 2.577632 | 0.009948 | NA |
| *CCT3* | 0.276578 | 0.107401 | 2.575198 | 0.01001828 | 0.188350167 |
| *ENSG00000279483* | -0.256777 | 0.09974 | -2.574462 | 0.01003962 | 0.188350167 |
| *IL23A* | -0.92928 | 0.361176 | -2.572927 | 0.01008425 | NA |
| *ATP9B* | -0.684426 | 0.26608 | -2.572255 | 0.01010386 | NA |
| *PSMA4* | -0.255906 | 0.099537 | -2.570952 | 0.01014195 | 0.189460247 |
| *TMEM14A* | 0.611083 | 0.238183 | 2.565604 | 0.01029962 | 0.191064685 |
| *CXCR4* | 0.144937 | 0.056504 | 2.565091 | 0.01031488 | 0.191064685 |
| *RGS10* | 0.907023 | 0.354312 | 2.559953 | 0.01046863 | 0.192311524 |
| *RPLP1* | 0.219036 | 0.085565 | 2.559885 | 0.01047068 | 0.192311524 |
| *RNASEK* | -0.554511 | 0.216678 | -2.559146 | 0.01049298 | NA |
| *LIMD1* | -0.917506 | 0.35885 | -2.556798 | 0.01056405 | 0.192311524 |
| *EIF4B* | 0.192 | 0.07513 | 2.555563 | 0.01060161 | 0.192311524 |
| *C9orf16* | -0.276647 | 0.108311 | -2.554188 | 0.01064358 | 0.192311524 |
| *RPL18A* | 0.163712 | 0.064156 | 2.551768 | 0.01071779 | 0.192311524 |
| *EIF5* | 0.225671 | 0.088464 | 2.551006 | 0.01074125 | 0.192311524 |
| *ASF1A* | 0.555118 | 0.217666 | 2.550313 | 0.01076262 | 0.192311524 |
| *ACYP2* | -0.421799 | 0.16542 | -2.549865 | 0.01077645 | 0.192311524 |
| *HCK* | -1.01233 | 0.397034 | -2.549729 | 0.01078066 | NA |
| *ENSG00000268400* | -2.224688 | 0.873754 | -2.546125 | 0.01089262 | NA |
| *MRPL28* | -0.337067 | 0.132445 | -2.544967 | 0.01092882 | 0.193208439 |
| *C19orf48* | 0.650353 | 0.255635 | 2.544073 | 0.01095681 | 0.193208439 |
| *TCL1A* | -0.492815 | 0.193716 | -2.544012 | 0.01095875 | 0.193208439 |
| *PPIL3* | 0.542645 | 0.213793 | 2.538181 | 0.01114303 | 0.195289652 |
| *ENSG00000224505* | -0.485315 | 0.19126 | -2.537468 | 0.01116576 | 0.195289652 |
| *ENSG00000262049* | -0.672482 | 0.265604 | -2.531893 | 0.01134487 | NA |
| *IFIT5* | 0.791867 | 0.312843 | 2.531196 | 0.01136742 | NA |
| *DUSP10* | 1.016273 | 0.401519 | 2.531073 | 0.01137141 | 0.198097192 |
| *FAAP100* | -0.807209 | 0.319194 | -2.528901 | 0.01144203 | NA |
| *RPLP0* | 0.296406 | 0.117433 | 2.524049 | 0.01160116 | 0.201300801 |
| *ADA* | -0.487317 | 0.193329 | -2.520663 | 0.0117134 | 0.20244807 |

| | | | | | |
|---|---|---|---|---|---|
| *NACA* | 0.145762 | 0.058015 | 2.512468 | 0.01198901 | 0.205265105 |
| *LSM6* | 0.372828 | 0.148429 | 2.511824 | 0.01201089 | 0.205265105 |
| *RPS21* | 0.13942 | 0.055509 | 2.511655 | 0.01201666 | 0.205265105 |
| *MDFIC* | 0.972304 | 0.387211 | 2.511043 | 0.01203751 | NA |
| *PIGF* | 0.6144 | 0.24475 | 2.510311 | 0.0120625 | NA |
| *ACP1* | -0.23363 | 0.093083 | -2.509917 | 0.01207597 | 0.205384579 |
| *BIN2* | -0.614188 | 0.244822 | -2.508712 | 0.01211722 | 0.205384579 |
| *NSMCE4A* | -0.626845 | 0.250077 | -2.506611 | 0.01218947 | 0.205814493 |
| *SLC26A6* | -1.129513 | 0.450804 | -2.50555 | 0.01222612 | NA |
| *HIPK2* | 1.015212 | 0.405386 | 2.504311 | 0.01226902 | NA |
| *LINC01006* | 0.928601 | 0.371288 | 2.501025 | 0.01238344 | NA |
| *XKR8* | 0.750656 | 0.300255 | 2.500065 | 0.01241706 | NA |
| *LIMD2* | -0.220668 | 0.088332 | -2.49816 | 0.01248398 | 0.209979628 |
| *EEF1A1* | 0.221435 | 0.08871 | 2.496179 | 0.01255393 | 0.210087649 |
| *PSMC4* | -0.450975 | 0.180732 | -2.49527 | 0.01258612 | 0.210087649 |
| *SEM1* | 0.23522 | 0.094339 | 2.49335 | 0.01265441 | 0.210427541 |
| *VARS* | -1.116425 | 0.448749 | -2.487863 | 0.01285131 | NA |
| *TARBP1* | -0.930259 | 0.373959 | -2.487594 | 0.01286103 | NA |
| *TLE2* | -1.396397 | 0.561393 | -2.487379 | 0.01286884 | NA |
| *ACCS* | 1.217358 | 0.489421 | 2.487343 | 0.01287014 | NA |
| *TCEAL4* | 0.396263 | 0.159435 | 2.485414 | 0.01294007 | 0.214365705 |
| *THAP8* | 1.194755 | 0.481127 | 2.48324 | 0.01301935 | NA |
| *WARS* | 1.276757 | 0.514197 | 2.483011 | 0.01302771 | NA |
| *MT1F* | 1.36643 | 0.551174 | 2.479127 | 0.01317043 | NA |
| *LGALS3BP* | -1.302909 | 0.525588 | -2.478956 | 0.01317674 | NA |
| *ENSG00000225938* | -0.448517 | 0.181176 | -2.475584 | 0.01330186 | NA |
| *GLS* | 0.517679 | 0.209926 | 2.466002 | 0.01366305 | NA |
| *PRDX3* | 0.390982 | 0.158584 | 2.465457 | 0.01368386 | 0.225752816 |
| *JUND* | -0.300382 | 0.121899 | -2.464197 | 0.01373208 | 0.225752816 |
| *HINT1* | 0.18137 | 0.073641 | 2.462903 | 0.01378172 | 0.225752816 |
| *RPL27* | 0.132237 | 0.053758 | 2.459856 | 0.01389928 | 0.22683203 |
| *CCT4* | 0.284231 | 0.115718 | 2.456246 | 0.01403971 | 0.228275355 |
| *PPP1R14A* | -0.769508 | 0.313512 | -2.454476 | 0.01410899 | 0.228555211 |
| *LILRA4* | 1.713189 | 0.698207 | 2.453697 | 0.0141396 | NA |
| *NEK6* | 1.664151 | 0.678484 | 2.452749 | 0.01417691 | NA |
| *TCF7* | -0.565165 | 0.230878 | -2.447898 | 0.01436925 | 0.231915407 |
| *AIP* | 0.29976 | 0.122742 | 2.442192 | 0.01459837 | 0.234750381 |
| *ENSG00000270659* | -1.266716 | 0.519042 | -2.44049 | 0.01466734 | NA |
| *SMARCE1* | -0.383929 | 0.157416 | -2.438944 | 0.01473027 | 0.23600682 |
| *WDR46* | -0.407025 | 0.167102 | -2.435786 | 0.01485948 | 0.237211398 |
| *ADRB2* | 1.207859 | 0.496598 | 2.432267 | 0.01500464 | NA |
| *MTX2* | 0.762875 | 0.313726 | 2.431659 | 0.01502985 | NA |
| *ARF5* | -0.345778 | 0.142363 | -2.428854 | 0.01514664 | 0.240686674 |
| *SELENOF* | -0.21497 | 0.088542 | -2.427893 | 0.01518684 | 0.240686674 |
| *RAB33A* | 1.876969 | 0.773413 | 2.426866 | 0.01522986 | NA |
| *PDPK1* | 0.526584 | 0.216995 | 2.426716 | 0.01523616 | NA |
| *HOXB2* | 1.224795 | 0.50473 | 2.426635 | 0.01523959 | NA |

| | | | | | |
|---|---|---|---|---|---|
| *RPS4X* | 0.228274 | 0.094194 | 2.423436 | 0.01537447 | 0.24278385 |
| *IFNG.AS1* | -1.729449 | 0.713862 | -2.422664 | 0.01540716 | NA |
| *C11orf1* | 0.732996 | 0.302584 | 2.422457 | 0.01541597 | NA |
| *ZFPL1* | -0.505832 | 0.209051 | -2.419655 | 0.01553522 | 0.244443083 |
| *DNAJB1* | 0.551825 | 0.228294 | 2.417164 | 0.01564195 | NA |
| *MFAP1* | 0.572747 | 0.237069 | 2.415952 | 0.01569412 | NA |
| *YBX3* | 0.510069 | 0.211191 | 2.4152 | 0.01572657 | 0.246570082 |
| *SMIM10* | -2.96597 | 1.228452 | -2.414396 | 0.01576131 | NA |
| *TCEAL8* | 0.340345 | 0.141182 | 2.410679 | 0.01592284 | 0.248758909 |
| *TMEM71* | -0.69673 | 0.289153 | -2.409555 | 0.01597197 | NA |
| *PTPN7* | -0.613514 | 0.254705 | -2.408722 | 0.01600851 | 0.249210453 |
| *NSUN7* | -1.10649 | 0.459596 | -2.407526 | 0.01606102 | NA |
| *LGALS3* | 1.462824 | 0.607674 | 2.407252 | 0.0160731 | NA |
| *BLNK* | 0.228004 | 0.094741 | 2.406597 | 0.01610195 | 0.249779287 |
| *EXOG* | -0.42336 | 0.176019 | -2.40519 | 0.01616408 | 0.249860263 |
| *DOCK9* | -1.226726 | 0.510929 | -2.40097 | 0.01635168 | NA |
| *ENSG00000275457* | -1.153587 | 0.480496 | -2.400824 | 0.01635818 | NA |
| *VSIR* | -0.617321 | 0.257164 | -2.400498 | 0.01637277 | NA |
| *CETN3* | 0.555939 | 0.231612 | 2.4003 | 0.01638165 | NA |
| *RPL39* | 0.181223 | 0.075579 | 2.397784 | 0.01649458 | 0.254074409 |
| *RGS16* | -1.075882 | 0.448982 | -2.396271 | 0.01656285 | NA |
| *SYNGR2* | -0.424975 | 0.177423 | -2.395268 | 0.0166082 | 0.254930082 |
| *ACTG1* | 0.444112 | 0.185528 | 2.393781 | 0.01667572 | 0.255074606 |
| *PSMB3* | -0.337922 | 0.141268 | -2.392052 | 0.01675445 | 0.255389041 |
| *PRAG1* | 1.214425 | 0.507775 | 2.391661 | 0.01677234 | NA |
| *CHMP6* | -0.68359 | 0.285939 | -2.390684 | 0.01681702 | NA |
| *POLE3* | -0.349142 | 0.14609 | -2.38992 | 0.01685207 | 0.25585425 |
| *MRPL32* | 0.346513 | 0.145055 | 2.388843 | 0.01690153 | 0.25585425 |
| *PTPRS* | 1.786998 | 0.749132 | 2.385425 | 0.01705939 | NA |
| *ENSG00000125726* | -1.460738 | 0.613357 | -2.381544 | 0.01724023 | NA |
| *LRIF1* | 0.413023 | 0.173658 | 2.378368 | 0.01738946 | 0.262315137 |
| *ZNF680* | 0.443364 | 0.186513 | 2.377116 | 0.01744858 | 0.262315137 |
| *ZNF844* | -0.708838 | 0.298272 | -2.376482 | 0.01747861 | NA |
| *LILRB1* | 0.537768 | 0.226345 | 2.375871 | 0.01750759 | 0.262315137 |
| *SREBF2.AS1* | -1.032379 | 0.435088 | -2.372806 | 0.01765354 | NA |
| *RPS19* | 0.177518 | 0.075032 | 2.36591 | 0.01798581 | 0.268137769 |
| *MTMR10* | -1.242421 | 0.525384 | -2.364786 | 0.01804051 | NA |
| *PDCD4* | 0.464537 | 0.196464 | 2.364486 | 0.01805509 | 0.268137769 |
| *GUCD1* | -0.38043 | 0.160927 | -2.363987 | 0.01807945 | 0.268137769 |
| *HELZ2* | 0.672957 | 0.284812 | 2.362814 | 0.01813678 | NA |
| *C21orf62* | -0.994212 | 0.420839 | -2.362451 | 0.01815455 | NA |
| *PIGM* | 0.749959 | 0.31751 | 2.362002 | 0.01817655 | NA |
| *CD37* | -0.151398 | 0.064111 | -2.361489 | 0.01820173 | 0.26825635 |
| *ARPC3* | -0.197925 | 0.083819 | -2.361327 | 0.01820966 | 0.26825635 |
| *ATP6AP2* | -0.387576 | 0.164378 | -2.357829 | 0.01838218 | 0.269892209 |
| *ZNF217* | 0.6834 | 0.290003 | 2.35653 | 0.01844659 | NA |
| *PGBD4* | -1.133658 | 0.481218 | -2.355812 | 0.01848228 | NA |

| | | | | | |
|---|---|---|---|---|---|
| *ERI3* | -0.517547 | 0.219967 | -2.35284 | 0.01863066 | NA |
| *DNAJC15* | 0.458541 | 0.194977 | 2.351765 | 0.0186846 | 0.272790138 |
| *CCT8* | 0.343638 | 0.146143 | 2.351382 | 0.01870383 | 0.272790138 |
| *KLK2* | 1.646026 | 0.701556 | 2.346251 | 0.01896332 | NA |
| *KCTD10* | -0.83192 | 0.3547 | -2.345417 | 0.01900582 | NA |
| *CLTA* | 0.223657 | 0.095409 | 2.344198 | 0.01906803 | 0.276869121 |
| *WDFY1* | 0.756786 | 0.322946 | 2.343385 | 0.01910965 | 0.276869121 |
| *ENSG00000275799* | 0.639039 | 0.27277 | 2.342775 | 0.01914093 | NA |
| *RCSD1* | -0.388272 | 0.166159 | -2.33675 | 0.01945217 | 0.279847398 |
| *ZNF638* | -0.387972 | 0.16606 | -2.33634 | 0.01947352 | 0.279847398 |
| *TCIRG1* | -0.492072 | 0.210733 | -2.335047 | 0.01954096 | 0.279847398 |
| *HNRNPA0* | -0.174913 | 0.074954 | -2.333587 | 0.01961738 | 0.279847398 |
| *ENSA* | -0.320771 | 0.137477 | -2.333271 | 0.01963394 | 0.279847398 |
| *MAP3K1* | 0.407206 | 0.174618 | 2.331979 | 0.01970179 | 0.279905691 |
| *RBM18* | 0.521686 | 0.223741 | 2.331651 | 0.01971903 | NA |
| *THEMIS2* | 0.644877 | 0.276819 | 2.329596 | 0.01982754 | NA |
| *RNF34* | 0.438395 | 0.188264 | 2.328619 | 0.01987928 | 0.281516201 |
| *PSMB10* | -0.280321 | 0.120491 | -2.3265 | 0.01999192 | 0.282201017 |
| *PTPRE* | -0.712445 | 0.306401 | -2.325205 | 0.02006101 | NA |
| *ISG20* | -0.311248 | 0.133961 | -2.323415 | 0.02015684 | 0.283617115 |
| *CLEC4A* | -1.327424 | 0.571381 | -2.323184 | 0.02016927 | NA |
| *DIS3L* | 0.69288 | 0.298886 | 2.318211 | 0.02043784 | NA |
| *FAM162A* | 0.366733 | 0.158208 | 2.318051 | 0.02044654 | 0.28437404 |
| *GPSM3* | -0.162552 | 0.070155 | -2.31705 | 0.02050099 | 0.28437404 |
| *NPM3* | 0.511422 | 0.220835 | 2.315859 | 0.02056596 | NA |
| *TNFAIP8* | 0.226098 | 0.097631 | 2.315841 | 0.02056697 | 0.28437404 |
| *PDE3B* | -0.595308 | 0.257153 | -2.314993 | 0.02061334 | 0.28437404 |
| *DPP9* | -0.577639 | 0.249615 | -2.314124 | 0.02066092 | NA |
| *FUT11* | 0.566357 | 0.244795 | 2.313602 | 0.02068955 | NA |
| *PPCS* | -0.259587 | 0.112288 | -2.311791 | 0.02078919 | 0.28437404 |
| *YWHAG* | 0.49031 | 0.212174 | 2.310883 | 0.02083934 | NA |
| *CYB561A3* | -0.407153 | 0.176298 | -2.309459 | 0.02091811 | 0.28437404 |
| *HMOX1* | 0.922811 | 0.399632 | 2.309153 | 0.02093511 | NA |
| *USP34* | 0.290042 | 0.12561 | 2.309068 | 0.0209398 | 0.28437404 |
| *CHPT1* | 0.342234 | 0.148251 | 2.308478 | 0.02097259 | 0.28437404 |
| *TAOK3* | 0.383847 | 0.166289 | 2.308314 | 0.0209817 | 0.28437404 |
| *TOP2A* | -1.798637 | 0.779319 | -2.307959 | 0.02100144 | NA |
| *ITGA5* | -1.202907 | 0.521337 | -2.30735 | 0.02103532 | NA |
| *MCM7* | -0.42612 | 0.184703 | -2.307057 | 0.02105165 | 0.28437404 |
| *ENSG00000089009* | 0.15901 | 0.068923 | 2.307055 | 0.02105175 | 0.28437404 |
| *RAD9A* | -0.306003 | 0.132715 | -2.305711 | 0.02112676 | 0.28437404 |
| *ZKSCAN1* | -0.295495 | 0.128166 | -2.305564 | 0.02113498 | 0.28437404 |
| *BMS1* | -0.473172 | 0.205259 | -2.305243 | 0.02115294 | 0.28437404 |
| *LINC.PINT* | 0.282542 | 0.122593 | 2.304719 | 0.0211823 | 0.28437404 |
| *RBM4* | -0.355773 | 0.154513 | -2.302538 | 0.02130484 | 0.285147097 |
| *PTCD2* | 1.022943 | 0.444712 | 2.300235 | 0.0214349 | NA |
| *UNC119* | -0.400876 | 0.174291 | -2.300043 | 0.02144576 | 0.285202512 |

| | | | | | |
|---|---|---|---|---|---|
| *PTPRC* | 0.240985 | 0.10478 | 2.299905 | 0.02145361 | 0.285202512 |
| *AFF3* | 0.277665 | 0.120775 | 2.299019 | 0.02150388 | 0.285202512 |
| *S1PR2* | -0.817213 | 0.355848 | -2.296521 | 0.02164611 | NA |
| *ALDH9A1* | 0.48389 | 0.211014 | 2.293162 | 0.02183871 | 0.286101378 |
| *MT.CO3* | 0.216848 | 0.094571 | 2.292972 | 0.02184963 | 0.286101378 |
| *CAMTA1* | -0.369718 | 0.161255 | -2.292747 | 0.02186257 | 0.286101378 |
| *DDT* | 0.283884 | 0.12383 | 2.292529 | 0.02187511 | 0.286101378 |
| *SEPTIN6* | 0.233544 | 0.101952 | 2.290729 | 0.02197908 | 0.286101378 |
| *CD1D* | 1.039622 | 0.45402 | 2.289815 | 0.02203206 | NA |
| *ZNF197* | -1.07089 | 0.468226 | -2.287124 | 0.02218857 | NA |
| *ENSG00000223881* | -1.639721 | 0.716968 | -2.287022 | 0.02219455 | NA |
| *TOMM5* | -0.342731 | 0.149893 | -2.286502 | 0.02222493 | 0.286101378 |
| *AKAP9* | 0.195426 | 0.085493 | 2.28587 | 0.02226185 | 0.286101378 |
| *NUAK2* | -0.338909 | 0.148276 | -2.28566 | 0.02227415 | 0.286101378 |
| *CDC25B* | -0.532614 | 0.233048 | -2.285421 | 0.02228818 | 0.286101378 |
| *IGKC* | -0.38846 | 0.169988 | -2.285226 | 0.02229959 | 0.286101378 |
| *PDCD4.AS1* | 0.655938 | 0.287063 | 2.284994 | 0.0223132 | NA |
| *AKIRIN2* | 0.324191 | 0.141889 | 2.284828 | 0.02232292 | 0.286101378 |
| *GLRX2* | 0.584377 | 0.25597 | 2.282993 | 0.02243081 | NA |
| *ARHGDIB* | -0.210431 | 0.09218 | -2.282836 | 0.02244003 | 0.286101378 |
| *EVI2A* | 0.72716 | 0.318658 | 2.281944 | 0.02249262 | 0.286101378 |
| *RBIS* | 0.223301 | 0.097879 | 2.281403 | 0.02252461 | 0.286101378 |
| *CSKMT* | -0.3101 | 0.13595 | -2.280987 | 0.02254922 | 0.286101378 |
| *LDB1* | 0.981602 | 0.430629 | 2.279462 | 0.02263965 | NA |
| *STK4* | 0.23334 | 0.102505 | 2.276378 | 0.02282339 | 0.288745517 |
| *SOAT1* | 0.715767 | 0.314662 | 2.274722 | 0.02292263 | NA |
| *MRPL42* | 0.393101 | 0.172951 | 2.272902 | 0.02303209 | 0.290548499 |
| *ITGA6.AS1* | -0.77291 | 0.341209 | -2.265208 | 0.02349991 | NA |
| *A4GALT* | 1.479946 | 0.653592 | 2.264328 | 0.02355396 | NA |
| *ZC3H12A* | -0.593941 | 0.262434 | -2.263201 | 0.02362332 | NA |
| *GYPC* | 0.276182 | 0.122065 | 2.262587 | 0.02366116 | 0.297179097 |
| *ENSG00000257275* | -0.562004 | 0.248447 | -2.26207 | 0.02369309 | 0.297179097 |
| *GNLY* | -2.494631 | 1.102895 | -2.261893 | 0.02370401 | NA |
| *TMUB1* | -0.281321 | 0.124585 | -2.258063 | 0.02394171 | 0.298927988 |
| *PARP14* | 0.293207 | 0.129908 | 2.257031 | 0.02400611 | 0.298927988 |
| *VOPP1* | -0.34828 | 0.154342 | -2.25654 | 0.02403681 | 0.298927988 |
| *RB1* | -0.322782 | 0.143313 | -2.25228 | 0.02430461 | 0.299993012 |
| *NR6A1* | -1.504241 | 0.667894 | -2.252215 | 0.0243087 | NA |
| *CNIH4* | 0.457404 | 0.203154 | 2.251514 | 0.02435301 | 0.299993012 |
| *ZDHHC14* | -0.817393 | 0.363104 | -2.251125 | 0.02437762 | 0.299993012 |
| *ENSG00000267364* | 1.006415 | 0.447094 | 2.251016 | 0.02438452 | NA |
| *GTF2F2* | 0.407047 | 0.180843 | 2.250838 | 0.02439579 | 0.299993012 |
| *SNHG10* | -0.495528 | 0.220175 | -2.250606 | 0.02441048 | NA |
| *WASHC2C* | 0.604669 | 0.26875 | 2.249934 | 0.02445312 | NA |
| *PIM1* | -0.344468 | 0.153187 | -2.248682 | 0.02453271 | 0.300806621 |
| *ZNF33A* | 0.369942 | 0.164591 | 2.247643 | 0.02459899 | 0.300806621 |
| *ENSG00000254614* | -0.801353 | 0.356579 | -2.247336 | 0.02461853 | NA |

| | | | | | |
|---|---|---|---|---|---|
| *ENSG00000273253* | 0.717714 | 0.319435 | 2.246825 | 0.02465121 | NA |
| *UXT* | 0.208344 | 0.092753 | 2.246224 | 0.02468966 | 0.301076732 |
| *IL21R* | -0.836003 | 0.372442 | -2.244654 | 0.02479038 | NA |
| *FCGR2B* | 0.556411 | 0.24807 | 2.24296 | 0.02489937 | 0.302792871 |
| *LACTB2* | 0.503431 | 0.224637 | 2.241086 | 0.02502048 | NA |
| *ZNF266* | -0.747711 | 0.333757 | -2.240283 | 0.02507253 | NA |
| *ENSG00000263394* | 0.986304 | 0.440487 | 2.239121 | 0.02514805 | NA |
| *RNF185* | 0.609168 | 0.272114 | 2.23865 | 0.02517867 | NA |
| *RBM19* | -0.5211 | 0.232941 | -2.237041 | 0.02528365 | NA |
| *RSBN1L* | 0.360431 | 0.161303 | 2.234493 | 0.02545065 | 0.308641797 |
| *CCDC90B* | 0.322839 | 0.144662 | 2.231673 | 0.02563657 | 0.309217094 |
| *AGPAT2* | -0.340226 | 0.152456 | -2.231637 | 0.02563896 | 0.309217094 |
| *GNPDA1* | 0.984259 | 0.441507 | 2.229317 | 0.02579283 | NA |
| *ENSG00000258056* | 1.010512 | 0.453299 | 2.229238 | 0.02579806 | NA |
| *LAPTM5* | -0.179307 | 0.080496 | -2.227521 | 0.0259125 | 0.311659978 |
| *IFI30* | 0.469818 | 0.211162 | 2.224915 | 0.02608695 | 0.31278158 |
| *HDX* | 1.714021 | 0.770907 | 2.223384 | 0.02618992 | NA |
| *ENSG00000278834* | -1.36409 | 0.61385 | -2.222189 | 0.02627053 | NA |
| *ZBTB25* | 0.462361 | 0.208073 | 2.222113 | 0.02627567 | 0.31278158 |
| *RPS29* | 0.128578 | 0.057867 | 2.221964 | 0.02628574 | 0.31278158 |
| *ACAT1* | -0.533146 | 0.239975 | -2.22167 | 0.02630561 | 0.31278158 |
| *KIAA0040* | 0.338496 | 0.152418 | 2.220837 | 0.026362 | 0.31278158 |
| *MRPL41* | -0.368541 | 0.16604 | -2.219593 | 0.02644638 | 0.312937001 |
| *EPS8L2* | 0.813793 | 0.366738 | 2.219007 | 0.02648624 | NA |
| *C1QBP* | 0.346505 | 0.156242 | 2.217745 | 0.0265722 | 0.313576828 |
| *NFE2L2* | 0.307081 | 0.138586 | 2.215821 | 0.02670378 | 0.313576828 |
| *CRIPT* | 0.383884 | 0.173303 | 2.215107 | 0.02675274 | 0.313576828 |
| *NBPF15* | 0.920714 | 0.415662 | 2.215054 | 0.02675635 | NA |
| *EIF4A2* | 0.158604 | 0.071619 | 2.214559 | 0.02679035 | 0.313576828 |
| *ENSG00000273748* | -0.855945 | 0.386679 | -2.213581 | 0.02685761 | 0.313576828 |
| *ZNHIT3* | 0.268025 | 0.121394 | 2.207885 | 0.02725226 | 0.317340681 |
| *KLHL6* | 0.655891 | 0.297544 | 2.204354 | 0.02749946 | 0.318784124 |
| *SPPL2A* | -0.365946 | 0.166035 | -2.20403 | 0.02752226 | 0.318784124 |
| *TOE1* | -0.536737 | 0.243534 | -2.203954 | 0.02752759 | NA |
| *ENSG00000180448* | -0.286637 | 0.13019 | -2.201682 | 0.02768781 | 0.318784124 |
| *PHF3* | 0.273361 | 0.124208 | 2.200838 | 0.02774747 | 0.318784124 |
| *LY6E* | 0.352685 | 0.16026 | 2.200712 | 0.02775644 | 0.318784124 |
| *DLGAP1.AS1* | -0.345767 | 0.157172 | -2.199929 | 0.02781192 | 0.318784124 |
| *ZNF441* | -0.479249 | 0.217866 | -2.199749 | 0.02782473 | NA |
| *SETBP1* | 0.57817 | 0.262964 | 2.198662 | 0.02790197 | NA |
| *APBB1* | 0.783933 | 0.356676 | 2.197889 | 0.02795703 | NA |
| *DGCR6* | -1.642796 | 0.747816 | -2.196791 | 0.0280354 | NA |
| *ATF7IP2* | -0.495903 | 0.225758 | -2.196615 | 0.02804793 | 0.320652083 |
| *JARID2* | -0.329517 | 0.15038 | -2.191226 | 0.02843543 | 0.324237753 |
| *THG1L* | 0.513257 | 0.234379 | 2.18986 | 0.02853438 | NA |
| *P4HA1* | -0.563782 | 0.257526 | -2.189221 | 0.02858075 | NA |
| *TXN* | 0.36722 | 0.167969 | 2.18624 | 0.02879804 | 0.327521726 |

| | | | | | |
|---|---|---|---|---|---|
| *FAM78A* | 0.565669 | 0.259389 | 2.180772 | 0.02920032 | NA |
| *ATP2B1.AS1* | -0.354655 | 0.16286 | -2.177665 | 0.029431 | 0.33385547 |
| *ENSG00000272182* | -1.415776 | 0.650777 | -2.175518 | 0.02959134 | NA |
| *KDELR2* | -0.325908 | 0.149855 | -2.174828 | 0.02964298 | 0.335393453 |
| *PRICKLE1* | 0.63481 | 0.29195 | 2.174381 | 0.0296765 | NA |
| *SPON1* | -0.582867 | 0.268228 | -2.173031 | 0.02977797 | NA |
| *IRS2* | 0.441792 | 0.203327 | 2.172811 | 0.02979452 | 0.336241457 |
| *GLMP* | 0.727388 | 0.334801 | 2.172598 | 0.02981059 | NA |
| *PILRB* | 0.581353 | 0.267728 | 2.171435 | 0.02989828 | 0.336547255 |
| *MBNL2* | -0.579997 | 0.267245 | -2.170285 | 0.0299853 | NA |
| *GPRIN3* | 1.270375 | 0.585449 | 2.169917 | 0.03001312 | NA |
| *USP53* | 0.908881 | 0.418899 | 2.169688 | 0.03003047 | NA |
| *RP9* | 0.511163 | 0.235692 | 2.168775 | 0.03009981 | NA |
| *TMEM243* | 0.248947 | 0.114863 | 2.167344 | 0.03020866 | 0.339021816 |
| *STAM2* | 0.770944 | 0.355713 | 2.167322 | 0.03021028 | NA |
| *MAVS* | 0.391034 | 0.180491 | 2.166506 | 0.03027256 | 0.339021816 |
| *BAG2* | 0.666968 | 0.307911 | 2.166104 | 0.03030325 | NA |
| *FAM160B1* | -0.841211 | 0.38839 | -2.165893 | 0.03031938 | NA |
| *TLE3* | -0.345045 | 0.159495 | -2.163364 | 0.03051318 | 0.340403987 |
| *COL9A3* | -0.738666 | 0.341521 | -2.162872 | 0.03055106 | 0.340403987 |
| *FCRL4* | 1.767295 | 0.817246 | 2.162502 | 0.03057951 | NA |
| *PSMC3* | -0.319856 | 0.148083 | -2.159982 | 0.03077409 | 0.34202085 |
| *IL3RA* | -2.152479 | 0.997293 | -2.158321 | 0.03090285 | NA |
| *IMPDH2* | 0.254463 | 0.117992 | 2.156604 | 0.03103652 | 0.344066507 |
| *OPTN* | -0.649321 | 0.301236 | -2.155524 | 0.03112087 | 0.344132549 |
| *NAIP* | 0.841034 | 0.390213 | 2.15532 | 0.0311368 | NA |
| *ENSG00000259943* | -0.75118 | 0.34864 | -2.154601 | 0.0311931 | NA |
| *ENSG00000232698* | -1.97676 | 0.917809 | -2.15378 | 0.03125742 | NA |
| *GBGT1* | 0.823322 | 0.38292 | 2.150116 | 0.03154604 | NA |
| *ARPC1B* | -0.169744 | 0.079043 | -2.147507 | 0.03175295 | 0.349596089 |
| *S100A4* | 0.557542 | 0.259655 | 2.147239 | 0.03177422 | 0.349596089 |
| *CD40* | -0.296761 | 0.13832 | -2.145465 | 0.03191565 | 0.3502743 |
| *C6orf120* | 0.788246 | 0.367557 | 2.144554 | 0.03198852 | NA |
| *PEG10* | 1.186232 | 0.553142 | 2.144533 | 0.03199017 | NA |
| *MGAT4A* | -0.652293 | 0.304207 | -2.144238 | 0.0320138 | NA |
| *MGST2* | -1.55322 | 0.724552 | -2.143699 | 0.03205703 | NA |
| *C9orf72* | -0.648897 | 0.30283 | -2.142774 | 0.03213121 | NA |
| *CASZ1* | -1.197984 | 0.559388 | -2.141597 | 0.03222592 | NA |
| *IL10RB* | 0.357049 | 0.166726 | 2.141528 | 0.03223151 | 0.352858627 |
| *POLR1D* | 0.322755 | 0.150836 | 2.139773 | 0.03237311 | 0.353371728 |
| *CD55* | -0.308352 | 0.14416 | -2.138954 | 0.03243936 | 0.353371728 |
| *WASH9P* | -0.500355 | 0.234059 | -2.137732 | 0.03253848 | 0.353574085 |
| *TRIP10* | 0.689658 | 0.323003 | 2.135144 | 0.03274923 | NA |
| *HLA.DMA* | -0.253601 | 0.11882 | -2.134329 | 0.03281587 | 0.354384105 |
| *TLK1* | 0.284064 | 0.133108 | 2.13409 | 0.03283538 | 0.354384105 |
| *KRCC1* | 0.286483 | 0.134257 | 2.133848 | 0.0328552 | 0.354384105 |
| *SNHG32* | 0.208281 | 0.097662 | 2.132662 | 0.03295242 | 0.354561603 |

| | | | | | |
|---|---|---|---|---|---|
| *RBM11* | -1.772798 | 0.831416 | -2.132263 | 0.03298526 | NA |
| *MED6* | -0.31004 | 0.145548 | -2.130157 | 0.03315865 | 0.355908244 |
| *FBXO44* | -0.432052 | 0.202856 | -2.129849 | 0.03318412 | NA |
| *SAG* | -2.092472 | 0.983331 | -2.127943 | 0.03334184 | NA |
| *LINC01215* | -0.39059 | 0.183588 | -2.127536 | 0.03337557 | 0.356425413 |
| *NOC3L* | 0.409471 | 0.192475 | 2.127399 | 0.03338696 | 0.356425413 |
| *QARS* | 0.279366 | 0.131365 | 2.126635 | 0.0334504 | 0.356425413 |
| *GPR160* | 0.812755 | 0.382669 | 2.123913 | 0.03367746 | NA |
| *RPS11* | 0.225168 | 0.106081 | 2.122613 | 0.03378628 | 0.358705594 |
| *ZFAND2A* | -0.457092 | 0.215432 | -2.121745 | 0.03385914 | 0.358705594 |
| *MBTPS1* | -0.460218 | 0.217045 | -2.12038 | 0.03397397 | 0.358705594 |
| *HSP90AB1* | 0.227383 | 0.107253 | 2.120055 | 0.03400137 | 0.358705594 |
| *SMIM10L1* | -0.31695 | 0.149577 | -2.118976 | 0.03409252 | 0.358705594 |
| *SDR39U1* | -0.351727 | 0.166116 | -2.117363 | 0.03422905 | 0.358705594 |
| *TMEM102* | 0.723851 | 0.341877 | 2.117285 | 0.03423567 | NA |
| *TSN* | -0.326394 | 0.154157 | -2.117277 | 0.03423637 | 0.358705594 |
| *MYL9* | 1.289513 | 0.609178 | 2.116809 | 0.03427606 | NA |
| *ENSG00000228106* | -0.618851 | 0.292382 | -2.116583 | 0.03429529 | NA |
| *MRPL45* | 0.401627 | 0.189942 | 2.114473 | 0.03447487 | 0.360344457 |
| *TPP1* | 0.368899 | 0.174591 | 2.112933 | 0.03460652 | 0.360662686 |
| *JMY* | 0.765115 | 0.362226 | 2.112259 | 0.03466425 | NA |
| *RGS2* | 0.528935 | 0.250419 | 2.112196 | 0.03466963 | 0.360662686 |
| *ENSG00000258017* | -1.531335 | 0.725274 | -2.111387 | 0.03473907 | NA |
| *KLF2* | -0.587745 | 0.278651 | -2.109251 | 0.03492295 | 0.36243913 |
| *AMZ2* | -0.333879 | 0.15844 | -2.107294 | 0.03509214 | 0.36333609 |
| *CENPB* | -0.663766 | 0.314991 | -2.107252 | 0.03509576 | NA |
| *PYCR3* | -0.59008 | 0.280041 | -2.107118 | 0.03510732 | NA |
| *ENSG00000275964* | -0.784723 | 0.372477 | -2.10677 | 0.03513756 | NA |
| *LAS1L* | -0.574022 | 0.272503 | -2.106475 | 0.03516311 | NA |
| *SLC16A3* | 0.578274 | 0.274603 | 2.105853 | 0.03521714 | NA |
| *KLHL5* | -0.514584 | 0.244527 | -2.104405 | 0.03534317 | 0.364326957 |
| *TMEM179B* | -0.318569 | 0.151391 | -2.104282 | 0.03535382 | 0.364326957 |
| *SMARCC1* | 0.377925 | 0.17972 | 2.102852 | 0.03547869 | 0.364332788 |
| *FAM210B* | 0.916268 | 0.435771 | 2.102637 | 0.03549751 | NA |
| *PPP1R16B* | 0.475143 | 0.226003 | 2.102376 | 0.03552037 | 0.364332788 |
| *CYBB* | -0.494413 | 0.23537 | -2.100579 | 0.03567795 | 0.365096032 |
| *KLHL21* | 0.828682 | 0.394538 | 2.100384 | 0.03569506 | NA |
| *PARP15* | 0.705882 | 0.336386 | 2.09843 | 0.03586721 | 0.366179214 |
| *ATP5MC2* | 0.159432 | 0.076017 | 2.097321 | 0.0359652 | 0.366327624 |
| *PLEK* | -0.655757 | 0.312826 | -2.096235 | 0.03606131 | NA |
| *BIRC3* | -0.389781 | 0.18598 | -2.095822 | 0.03609797 | 0.366828923 |
| *C17orf49* | -0.63357 | 0.302733 | -2.092832 | 0.03636413 | NA |
| *PTTG1IP* | 0.478212 | 0.228641 | 2.091539 | 0.03647976 | NA |
| *CD180* | 0.430647 | 0.20604 | 2.090111 | 0.0366078 | 0.368871265 |
| *RAB11B* | -0.305894 | 0.146353 | -2.090107 | 0.03660815 | 0.368871265 |
| *GUK1* | -0.218209 | 0.104404 | -2.090045 | 0.03661378 | 0.368871265 |
| *NSMCE2* | 0.414374 | 0.198283 | 2.089808 | 0.03663505 | 0.368871265 |

| | | | | | |
|---|---|---|---|---|---|
| *RBBP8* | -0.686223 | 0.328688 | -2.087764 | 0.03681914 | NA |
| *KRR1* | 0.392701 | 0.188288 | 2.085639 | 0.03701136 | 0.370563313 |
| *NBPF12* | -0.633476 | 0.303744 | -2.085558 | 0.03701865 | NA |
| *ABI1* | 0.317746 | 0.152402 | 2.084921 | 0.03707643 | 0.370563313 |
| *ENSG00000281706* | 1.320804 | 0.633579 | 2.084671 | 0.03709916 | NA |
| *MRPL44* | 0.602743 | 0.289194 | 2.084218 | 0.03714032 | 0.370563313 |
| *SCAI* | 0.356894 | 0.171261 | 2.083918 | 0.03716764 | 0.370563313 |
| *LINC01003* | -1.289205 | 0.618905 | -2.083042 | 0.03724739 | NA |
| *WDSUB1* | 0.558743 | 0.268267 | 2.082784 | 0.03727088 | 0.370563313 |
| *TMBIM4* | -0.187969 | 0.090267 | -2.08236 | 0.03730956 | 0.370563313 |
| *SMOX* | -1.60638 | 0.77227 | -2.080077 | 0.0375185 | NA |
| *ANXA6* | -0.302652 | 0.145537 | -2.079547 | 0.03756711 | 0.372279016 |
| *INTS6L* | 0.749556 | 0.360514 | 2.079134 | 0.037605 | NA |
| *RAB3A* | 1.1057 | 0.532193 | 2.077628 | 0.03774367 | NA |
| *NOL12* | -0.45346 | 0.218446 | -2.075843 | 0.0379085 | 0.373265392 |
| *ENSG00000272583* | -1.039383 | 0.500889 | -2.075078 | 0.03797927 | NA |
| *HNRNPH1* | -0.193177 | 0.093106 | -2.0748 | 0.03800512 | 0.373265392 |
| *PFDN5* | 0.104459 | 0.050347 | 2.074785 | 0.03800649 | 0.373265392 |
| *SCRN2* | 0.449362 | 0.216583 | 2.074782 | 0.03800675 | 0.373265392 |
| *ARFIP1* | -0.638555 | 0.3078 | -2.074575 | 0.0380259 | NA |
| *CYBC1* | -0.227367 | 0.109653 | -2.073516 | 0.03812432 | 0.373584304 |
| *HIST3H2A* | 1.32284 | 0.638403 | 2.072109 | 0.03825528 | NA |
| *ATP5PB* | 0.254334 | 0.122955 | 2.068505 | 0.03859255 | 0.377330316 |
| *NHLRC2* | 0.609431 | 0.294637 | 2.06841 | 0.03860147 | NA |
| *YTHDC2* | 0.481696 | 0.233067 | 2.066769 | 0.03875594 | NA |
| *VAT1* | 0.517763 | 0.250687 | 2.06538 | 0.03888708 | 0.379365102 |
| *NAA40* | -0.708357 | 0.343184 | -2.064074 | 0.03901071 | NA |
| *ETNK1* | 0.412258 | 0.199752 | 2.063848 | 0.03903214 | 0.37993593 |
| *GPBP1L1* | -0.285501 | 0.138401 | -2.062856 | 0.03912628 | 0.380009644 |
| *COCH* | -0.941859 | 0.456965 | -2.061117 | 0.03929188 | NA |
| *C2CD3* | 1.030966 | 0.500687 | 2.059102 | 0.03948444 | NA |
| *EAF2* | -0.487194 | 0.236648 | -2.058728 | 0.03952034 | 0.381719331 |
| *NCBP2* | -0.278977 | 0.135567 | -2.057854 | 0.0396042 | 0.381719331 |
| *SOD2* | -0.390598 | 0.189865 | -2.057244 | 0.03966275 | 0.381719331 |
| *HSD17B11* | 0.175638 | 0.08539 | 2.0569 | 0.03969588 | 0.381719331 |
| *SH3GLB1* | 0.262898 | 0.127839 | 2.056472 | 0.03973707 | 0.381719331 |
| *ENSG00000260261* | -0.79762 | 0.388498 | -2.053088 | 0.04006409 | NA |
| *ATP13A3* | 0.695154 | 0.338696 | 2.052446 | 0.04012637 | NA |
| *C1orf56* | -0.378251 | 0.184359 | -2.051713 | 0.04019752 | 0.384365647 |
| *ANKRD17* | 0.437805 | 0.213416 | 2.051417 | 0.04022635 | 0.384365647 |
| *TRMT13* | 0.338925 | 0.165305 | 2.050298 | 0.04033535 | 0.384365647 |
| *LINC01857* | 0.506938 | 0.247285 | 2.050017 | 0.04036277 | 0.384365647 |
| *POLE2* | 1.281485 | 0.625184 | 2.049772 | 0.04038666 | NA |
| *HSBP1* | 0.332448 | 0.162267 | 2.048771 | 0.04048451 | 0.384638321 |
| *SPCS2* | -0.194972 | 0.095204 | -2.047932 | 0.04056664 | 0.384638321 |
| *IRF1* | -0.233434 | 0.114061 | -2.04657 | 0.04070031 | 0.3847556 |
| *SRSF2* | -0.194361 | 0.094995 | -2.046021 | 0.0407543 | 0.3847556 |

| | | | | | |
|---|---|---|---|---|---|
| *LY96* | 0.451443 | 0.220901 | 2.043648 | 0.04098833 | 0.385395946 |
| *NDUFS5* | 0.187255 | 0.091632 | 2.043553 | 0.0409977 | 0.385395946 |
| *EXD3* | -0.549349 | 0.268977 | -2.042367 | 0.04111509 | NA |
| *E2F4* | -0.414634 | 0.203065 | -2.04188 | 0.04116346 | 0.386100663 |
| *ZNF263* | -0.777316 | 0.380907 | -2.040699 | 0.04128074 | 0.386100663 |
| *KCNMB3* | -0.751939 | 0.368482 | -2.040642 | 0.04128646 | NA |
| *ANK1* | -1.354002 | 0.663748 | -2.039933 | 0.04135705 | NA |
| *IKBIP* | 0.406458 | 0.199285 | 2.039579 | 0.04139227 | 0.386100663 |
| *ZFYVE21* | 0.589182 | 0.288913 | 2.039305 | 0.04141963 | NA |
| *PRR4* | 0.388028 | 0.190279 | 2.039256 | 0.04142447 | 0.386100663 |
| *ZNF827* | 0.918001 | 0.450179 | 2.03919 | 0.04143105 | NA |
| *RPS6KB2.AS1* | -1.156091 | 0.567064 | -2.038732 | 0.04147678 | NA |
| *RASSF5* | -0.277228 | 0.136155 | -2.03612 | 0.04173835 | 0.387840009 |
| *LINC00667* | -0.395366 | 0.194223 | -2.035628 | 0.04178777 | 0.387840009 |
| *MYC* | 0.342591 | 0.168486 | 2.03335 | 0.04201718 | 0.388641884 |
| *EIF3L* | 0.188132 | 0.09254 | 2.032975 | 0.04205508 | 0.388641884 |
| *CD47* | 0.179202 | 0.088184 | 2.032137 | 0.04213976 | 0.388641884 |
| *OPA3* | 0.660306 | 0.324967 | 2.031919 | 0.0421619 | NA |
| *CD151* | -0.601159 | 0.295995 | -2.030973 | 0.04225776 | NA |
| *TTN.AS1* | 0.801579 | 0.394692 | 2.030897 | 0.04226543 | NA |
| *ALG11* | -1.023927 | 0.504594 | -2.02921 | 0.04243695 | NA |
| *CERS2* | 0.46276 | 0.228349 | 2.026549 | 0.04270854 | 0.392673635 |
| *KCTD5* | 0.456834 | 0.225437 | 2.026436 | 0.04272008 | NA |
| *ZNF706* | 0.19809 | 0.09777 | 2.026088 | 0.04275581 | 0.392673635 |
| *DHX57* | -0.537466 | 0.265514 | -2.024251 | 0.04294434 | 0.393213066 |
| *THOC2* | 0.31543 | 0.155862 | 2.023771 | 0.04299368 | 0.393213066 |
| *VASP* | -0.269599 | 0.133289 | -2.022658 | 0.04310842 | 0.393350934 |
| *RPRML* | -2.029939 | 1.003659 | -2.022538 | 0.0431208 | NA |
| *CHTF18* | -0.738699 | 0.365331 | -2.022 | 0.04317631 | NA |
| *BNIP2* | -0.271892 | 0.134474 | -2.021888 | 0.04318796 | 0.393350934 |
| *CDK11A* | -0.385768 | 0.191042 | -2.019285 | 0.04345764 | 0.394284009 |
| *SNAPC2* | 0.429861 | 0.212891 | 2.019165 | 0.04347004 | 0.394284009 |
| *TOB1* | 0.829636 | 0.411005 | 2.018558 | 0.04353322 | NA |
| *EPC1* | -0.207882 | 0.103024 | -2.017797 | 0.04361242 | 0.394759882 |
| *DIPK1B* | -1.177899 | 0.584412 | -2.015529 | 0.04384927 | NA |
| *CHD7* | 0.477435 | 0.237062 | 2.013964 | 0.04401331 | 0.397568778 |
| *ENSG00000268584* | -0.884426 | 0.439285 | -2.01333 | 0.04407994 | NA |
| *IK* | -0.236393 | 0.117473 | -2.012312 | 0.04418705 | 0.397995581 |
| *FAM122A* | 0.435463 | 0.216455 | 2.011792 | 0.04424188 | 0.397995581 |
| *ENSG00000258308* | -1.873531 | 0.93178 | -2.010701 | 0.04435702 | NA |
| *ARL3* | 0.451558 | 0.224701 | 2.009593 | 0.04447427 | NA |
| *SIMC1* | -0.44894 | 0.223401 | -2.009574 | 0.04447624 | NA |
| *RIN3* | 0.533268 | 0.265394 | 2.009342 | 0.04450083 | 0.399506463 |
| *FOXO3* | 0.668193 | 0.332546 | 2.009326 | 0.04450262 | NA |
| *CTBP1.AS* | -1.410965 | 0.702423 | -2.008712 | 0.04456771 | NA |
| *PAPOLG* | 0.706824 | 0.351918 | 2.008489 | 0.04459134 | NA |
| *GNA12* | 0.522306 | 0.260161 | 2.007627 | 0.04468296 | NA |

| | | | | | |
|---|---|---|---|---|---|
| *RASSF1.AS1* | -1.074224 | 0.535339 | -2.006624 | 0.04478976 | NA |
| *ARMC1* | 0.425427 | 0.212137 | 2.005435 | 0.04491661 | 0.40241615 |
| *ZNF117* | -1.05132 | 0.524478 | -2.004508 | 0.04501564 | NA |
| *ATP2A3* | -0.31653 | 0.157919 | -2.00438 | 0.04502933 | 0.402604392 |
| *SGPP1* | 0.841284 | 0.420054 | 2.002799 | 0.04519886 | NA |
| *GTF2F1* | 0.253829 | 0.126752 | 2.002561 | 0.04522443 | 0.403029689 |
| *GPATCH4* | 0.456565 | 0.228029 | 2.002225 | 0.04526051 | 0.403029689 |
| *GAB1* | 0.93316 | 0.466128 | 2.001938 | 0.04529144 | NA |
| *EXOSC5* | 0.486519 | 0.243065 | 2.001596 | 0.0453282 | NA |
| *ZCCHC9* | 0.394145 | 0.197119 | 1.999534 | 0.04555064 | 0.404492737 |
| *AZIN1* | -0.367393 | 0.183789 | -1.998993 | 0.04560909 | 0.404492737 |
| *TSC22D1* | -0.722011 | 0.361538 | -1.997051 | 0.04581959 | NA |
| *LAMB1* | -0.892231 | 0.447048 | -1.995828 | 0.04595266 | NA |
| *DARS* | 0.262559 | 0.131632 | 1.994651 | 0.04608091 | 0.407853204 |
| *NFATC2* | 0.58603 | 0.293811 | 1.99458 | 0.04608874 | NA |
| *PCLAF* | -1.416759 | 0.71072 | -1.993413 | 0.04621628 | NA |
| *SHKBP1* | -0.285051 | 0.143043 | -1.992756 | 0.04628821 | 0.408863667 |
| *ENSG00000271737* | -0.925611 | 0.465032 | -1.990424 | 0.04654429 | NA |
| *KDM5B* | 0.627232 | 0.315302 | 1.98931 | 0.04666705 | NA |
| *PTGES2* | -0.39006 | 0.196131 | -1.988773 | 0.04672631 | 0.411904589 |
| *FOXRED1* | -0.66109 | 0.332668 | -1.987238 | 0.046896 | NA |
| *CCDC66* | 0.34872 | 0.175483 | 1.987206 | 0.0468996 | 0.412603652 |
| *LINC02422* | 0.50562 | 0.254467 | 1.986979 | 0.04692477 | NA |
| *PSMC1* | -0.280645 | 0.141309 | -1.986035 | 0.04702949 | 0.412918902 |
| *SIGLEC14* | 0.727423 | 0.36633 | 1.985704 | 0.04706618 | NA |
| *IFITM3* | -1.254177 | 0.631868 | -1.984873 | 0.04715866 | NA |
| *SENP5* | -0.384632 | 0.193935 | -1.983307 | 0.04733319 | 0.414569198 |
| *SMPD2* | -0.44758 | 0.225747 | -1.98266 | 0.04740539 | NA |
| *NRBF2* | 0.343806 | 0.173407 | 1.982652 | 0.04740632 | 0.414569198 |
| *ENSG00000266088* | -2.201717 | 1.110794 | -1.982111 | 0.04746684 | NA |
| *HOOK1* | 1.051888 | 0.531104 | 1.98057 | 0.04763951 | NA |
| *RPRD1A* | -0.37395 | 0.18882 | -1.980459 | 0.04765203 | 0.41588948 |
| *POU2F2* | 0.22034 | 0.111348 | 1.978832 | 0.04783493 | 0.416657411 |
| *SEPSECS.AS1* | 0.763464 | 0.386022 | 1.977775 | 0.04795415 | NA |
| *IDH2* | -0.303586 | 0.15351 | -1.977621 | 0.04797152 | 0.41678685 |
| *WRNIP1* | 0.700629 | 0.354386 | 1.977023 | 0.04803899 | NA |
| *KAT2A* | -0.380379 | 0.1924 | -1.977017 | 0.04803967 | 0.41678685 |
| *FAAP24* | -0.830852 | 0.420424 | -1.976222 | 0.04812962 | NA |
| *KHDC4* | -0.255416 | 0.129316 | -1.975129 | 0.04825348 | 0.417816127 |
| *POU2AF1* | -0.201423 | 0.102065 | -1.973481 | 0.0484408 | 0.418447872 |
| *ATP6V1A* | 0.442154 | 0.224124 | 1.972811 | 0.04851708 | 0.418447872 |
| *ARMH1* | -0.874963 | 0.444209 | -1.969713 | 0.04887131 | NA |
| *LHFPL4* | 0.937625 | 0.476065 | 1.96953 | 0.04889231 | NA |
| *TRIM4* | 0.448559 | 0.22776 | 1.969434 | 0.04890325 | 0.419928263 |
| *HLA.F* | -0.213038 | 0.108186 | -1.969189 | 0.04893139 | 0.419928263 |
| *MICOS10* | 0.200228 | 0.101701 | 1.968803 | 0.04897569 | 0.419928263 |
| *GCNT2* | 0.631227 | 0.320629 | 1.968716 | 0.04898575 | NA |

| | | | | | |
|---|---|---|---|---|---|
| *MYH3* | -1.247331 | 0.63374 | -1.968205 | 0.04904451 | NA |
| *ENSG00000225342* | -0.676192 | 0.344048 | -1.965399 | 0.04936807 | NA |
| *MT.ND6* | 0.420339 | 0.214127 | 1.96304 | 0.04964157 | 0.424807975 |
| *ENSG00000280011* | -1.106075 | 0.563746 | -1.962009 | 0.04976142 | NA |
| *LTB* | -0.342365 | 0.174502 | -1.961961 | 0.049767 | 0.424920632 |
| *DDX11* | -0.859189 | 0.438054 | -1.961378 | 0.04983494 | NA |
| *RND1* | -1.145074 | 0.583838 | -1.961286 | 0.04984561 | NA |
| *LYST* | -0.389073 | 0.198393 | -1.961121 | 0.04986487 | 0.424920632 |
| *SEPTIN7* | 0.195252 | 0.099596 | 1.960434 | 0.04994511 | 0.424920632 |

| Cluster 9 | log2FC | lfcSE | stat | pvalue | padj |
|---|---|---|---|---|---|
| *DHTKD1* | -1.036891 | 0.13692 | -7.572973 | 3.65E-14 | 3.61E-10 |
| *ARL17A* | -1.124521 | 0.17256 | -6.516703 | 7.19E-11 | 3.55E-07 |
| *CKS2* | 0.734163 | 0.117336 | 6.25691 | 3.93E-10 | 1.29E-06 |
| *TMEM70* | 0.594117 | 0.09883 | 6.011529 | 1.84E-09 | 3.70E-06 |
| *CHORDC1* | 0.699586 | 0.116432 | 6.00852 | 1.87E-09 | 3.70E-06 |
| *SRSF7* | 0.386678 | 0.065027 | 5.946443 | 2.74E-09 | 4.52E-06 |
| *TNFRSF12A* | 1.411121 | 0.239721 | 5.886518 | 3.94E-09 | 5.57E-06 |
| *SDCBP* | 0.640324 | 0.109344 | 5.856048 | 4.74E-09 | 5.86E-06 |
| *ARRDC3* | -0.602893 | 0.105291 | -5.72598 | 1.03E-08 | 1.13E-05 |
| *KDM6B* | 0.611342 | 0.112803 | 5.419565 | 5.97E-08 | 5.66E-05 |
| *ENSG00000273748* | -1.210698 | 0.223786 | -5.410064 | 6.30E-08 | 5.66E-05 |
| *RHOF* | -0.500456 | 0.092829 | -5.391149 | 7.00E-08 | 5.77E-05 |
| *ENSG00000130313* | -0.417909 | 0.079602 | -5.249987 | 1.52E-07 | 0.000115674 |
| *DUSP14* | 0.72054 | 0.140978 | 5.111018 | 3.20E-07 | 0.000226268 |
| *RIN3* | 1.240795 | 0.245838 | 5.04721 | 4.48E-07 | 0.000281731 |
| *HNRNPH3* | 0.398148 | 0.078935 | 5.043971 | 4.56E-07 | 0.000281731 |
| *ALOX5* | -0.588493 | 0.117962 | -4.988837 | 6.07E-07 | 0.000353243 |
| *FLYWCH2* | -0.55958 | 0.112548 | -4.971946 | 6.63E-07 | 0.000364047 |
| *IZUMO4* | -0.729663 | 0.147306 | -4.95338 | 7.29E-07 | 0.000379493 |
| *PIGX* | -0.830526 | 0.168058 | -4.941902 | 7.74E-07 | 0.000382411 |
| *ENSG00000254614* | -1.593047 | 0.323019 | -4.931746 | 8.15E-07 | 0.000383661 |
| *PSME1* | -0.249706 | 0.051184 | -4.878609 | 1.07E-06 | 0.000480084 |
| *MACROD2* | -1.300687 | 0.268543 | -4.843488 | 1.28E-06 | 0.000536692 |
| *MRPL39* | 0.486389 | 0.100541 | 4.837707 | 1.31E-06 | 0.000536692 |
| *GATD3A* | 5.27088 | 1.091011 | 4.831189 | 1.36E-06 | 0.000536692 |
| *RDX* | 0.529223 | 0.109924 | 4.814448 | 1.48E-06 | 0.000561249 |
| *TTC32* | -0.768015 | 0.161168 | -4.765318 | 1.89E-06 | 0.000690393 |
| *ENSG00000272211* | -1.296262 | 0.273841 | -4.733635 | 2.21E-06 | 0.000778641 |
| *PFKL* | -0.458855 | 0.097263 | -4.71765 | 2.39E-06 | 0.000789912 |
| *HMOX1* | 1.756464 | 0.372393 | 4.716695 | 2.40E-06 | 0.000789912 |
| *ARL4C* | -1.24735 | 0.266228 | -4.685268 | 2.80E-06 | 0.000867381 |
| *CD48* | 0.506911 | 0.108373 | 4.677459 | 2.90E-06 | 0.000867381 |
| *HSPH1* | 0.782082 | 0.167249 | 4.676141 | 2.92E-06 | 0.000867381 |
| *TRADD* | -0.488355 | 0.104528 | -4.67198 | 2.98E-06 | 0.000867381 |
| *GALNT2* | -0.762319 | 0.164446 | -4.635677 | 3.56E-06 | 0.0010049 |
| *LMO4* | 0.813322 | 0.179415 | 4.533195 | 5.81E-06 | 0.001559153 |
| *C17orf49* | -0.730199 | 0.161111 | -4.532268 | 5.84E-06 | 0.001559153 |
| *POU2F2* | 0.402547 | 0.089293 | 4.508146 | 6.54E-06 | 0.001668709 |
| *RFTN1* | -0.541688 | 0.120294 | -4.503046 | 6.70E-06 | 0.001668709 |
| *FCRL5* | 1.556078 | 0.34569 | 4.501366 | 6.75E-06 | 0.001668709 |
| *MIR181A1HG* | -1.029829 | 0.229113 | -4.494856 | 6.96E-06 | 0.001678617 |
| *SEC14L1* | -0.455058 | 0.101435 | -4.486218 | 7.25E-06 | 0.001706479 |
| *BHLHE41* | 1.486264 | 0.332854 | 4.465207 | 8.00E-06 | 0.001839059 |
| *BRIX1* | 0.487664 | 0.109992 | 4.43364 | 9.27E-06 | 0.002081801 |
| *UPP1* | 0.624087 | 0.140979 | 4.426807 | 9.56E-06 | 0.002101065 |
| *SLCO4A1* | 1.048956 | 0.238372 | 4.400497 | 1.08E-05 | 0.002321131 |

| | | | | | |
|---|---|---|---|---|---|
| *SIPA1L3* | -0.66239 | 0.150891 | -4.389863 | 1.13E-05 | 0.002364021 |
| *GNPDA1* | 1.305779 | 0.297718 | 4.385957 | 1.15E-05 | 0.002364021 |
| *UTP4* | 0.643434 | 0.146809 | 4.382783 | 1.17E-05 | 0.002364021 |
| *DIRAS1* | -1.094427 | 0.250166 | -4.374807 | 1.22E-05 | 0.002403084 |
| *GTF2IRD2* | -0.611267 | 0.1401 | -4.363086 | 1.28E-05 | 0.002485857 |
| *PPIL4* | 0.498588 | 0.114766 | 4.344399 | 1.40E-05 | 0.002604967 |
| *FADS1* | -0.986989 | 0.227201 | -4.344112 | 1.40E-05 | 0.002604967 |
| *SOCS3* | 1.304265 | 0.300501 | 4.340297 | 1.42E-05 | 0.002604967 |
| *NR4A3* | 1.020945 | 0.235658 | 4.332322 | 1.48E-05 | 0.002652048 |
| *POLR2J* | -0.283007 | 0.06558 | -4.31543 | 1.59E-05 | 0.002731389 |
| *ENSG00000272831* | -0.681885 | 0.158173 | -4.311003 | 1.63E-05 | 0.002731389 |
| *GTF2B* | 0.564457 | 0.130951 | 4.310448 | 1.63E-05 | 0.002731389 |
| *KDM4B* | 0.557955 | 0.12946 | 4.309872 | 1.63E-05 | 0.002731389 |
| *GNG11* | 1.953506 | 0.453606 | 4.306613 | 1.66E-05 | 0.002731389 |
| *NR4A1* | 1.072268 | 0.249681 | 4.294555 | 1.75E-05 | 0.00283686 |
| *TAF1A* | 0.742445 | 0.173161 | 4.287593 | 1.81E-05 | 0.002843904 |
| *RHEB* | 0.33249 | 0.077621 | 4.283495 | 1.84E-05 | 0.002843904 |
| *TCL1B* | -2.212826 | 0.516612 | -4.28334 | 1.84E-05 | 0.002843904 |
| *CMTM7* | -0.767365 | 0.179637 | -4.271764 | 1.94E-05 | 0.002905385 |
| *C2orf68* | -0.423452 | 0.099129 | -4.271725 | 1.94E-05 | 0.002905385 |
| *MED21* | 0.435666 | 0.102441 | 4.25283 | 2.11E-05 | 0.003114621 |
| *ASF1A* | 0.564089 | 0.134735 | 4.186671 | 2.83E-05 | 0.004106238 |
| *SRSF11* | 0.27241 | 0.06511 | 4.183862 | 2.87E-05 | 0.004106238 |
| *DNTTIP2* | 0.368259 | 0.088709 | 4.151335 | 3.31E-05 | 0.004668201 |
| *PHKG1* | -0.779234 | 0.188066 | -4.143414 | 3.42E-05 | 0.004684519 |
| *HSPD1* | 0.668359 | 0.161329 | 4.142825 | 3.43E-05 | 0.004684519 |
| *AK6* | 0.42589 | 0.102852 | 4.140802 | 3.46E-05 | 0.004684519 |
| *KBTBD8* | 1.100314 | 0.265918 | 4.1378 | 3.51E-05 | 0.004684519 |
| *DNAJB9* | 0.442506 | 0.108046 | 4.095545 | 4.21E-05 | 0.005487885 |
| *ATF3* | 0.696111 | 0.169984 | 4.095153 | 4.22E-05 | 0.005487885 |
| *CALM3* | -0.413642 | 0.101244 | -4.085609 | 4.40E-05 | 0.005644192 |
| *TNFRSF1B* | 0.916502 | 0.224657 | 4.079558 | 4.51E-05 | 0.00571886 |
| *PPIF* | 0.671223 | 0.164852 | 4.071674 | 4.67E-05 | 0.005782479 |
| *KLF10* | 0.723687 | 0.177762 | 4.071092 | 4.68E-05 | 0.005782479 |
| *TSPAN33* | -0.518677 | 0.127767 | -4.059535 | 4.92E-05 | 0.00600124 |
| *UCP2* | -0.416064 | 0.102741 | -4.04965 | 5.13E-05 | 0.006067688 |
| *ZNRD1ASP* | -0.764972 | 0.188936 | -4.048843 | 5.15E-05 | 0.006067688 |
| *DUSP22* | -0.478055 | 0.118083 | -4.048457 | 5.16E-05 | 0.006067688 |
| *SPAG9* | 0.520383 | 0.12888 | 4.037717 | 5.40E-05 | 0.006264011 |
| *HDAC2* | 0.420517 | 0.104205 | 4.035476 | 5.45E-05 | 0.006264011 |
| *NAF1* | 0.696883 | 0.173411 | 4.018678 | 5.85E-05 | 0.006626298 |
| *RAB8B* | 0.426532 | 0.106186 | 4.01684 | 5.90E-05 | 0.006626298 |
| *PTGIR* | -1.621316 | 0.403962 | -4.013538 | 5.98E-05 | 0.006644205 |
| *C22orf39* | -0.368901 | 0.092102 | -4.005364 | 6.19E-05 | 0.006801786 |
| *MFSD10* | -0.315581 | 0.078939 | -3.997784 | 6.39E-05 | 0.006875458 |
| *HSPE1* | 0.434901 | 0.10879 | 3.997616 | 6.40E-05 | 0.006875458 |
| *LINC02256* | -0.737371 | 0.184864 | -3.988716 | 6.64E-05 | 0.00695771 |

| | | | | | |
|---|---|---|---|---|---|
| *ADD3* | 0.66307 | 0.166336 | 3.986319 | 6.71E-05 | 0.00695771 |
| *SLC9A7* | -0.599321 | 0.15052 | -3.981684 | 6.84E-05 | 0.00695771 |
| *FKBP5* | 0.842042 | 0.211505 | 3.98119 | 6.86E-05 | 0.00695771 |
| *WBP1L* | -0.632431 | 0.158859 | -3.981076 | 6.86E-05 | 0.00695771 |
| *CYB5A* | -0.72591 | 0.182398 | -3.979805 | 6.90E-05 | 0.00695771 |
| *ZBTB10* | 0.507817 | 0.127805 | 3.973363 | 7.09E-05 | 0.007076457 |
| *FKBP4* | 0.467207 | 0.118047 | 3.957796 | 7.56E-05 | 0.007477743 |
| *IKZF2* | -1.174491 | 0.296937 | -3.955349 | 7.64E-05 | 0.007477743 |
| *MREG* | -0.827408 | 0.209307 | -3.953077 | 7.72E-05 | 0.007477743 |
| *HHIP.AS1* | -3.01619 | 0.764353 | -3.946071 | 7.94E-05 | NA |
| *SPHK1* | 0.976011 | 0.247709 | 3.940149 | 8.14E-05 | 0.007815796 |
| *NCBP3* | 0.973206 | 0.247359 | 3.934385 | 8.34E-05 | 0.007906437 |
| *LDLRAD4* | 0.825748 | 0.210023 | 3.931704 | 8.43E-05 | 0.007906437 |
| *DDX21* | 0.563809 | 0.143445 | 3.930485 | 8.48E-05 | 0.007906437 |
| *PNPLA6* | -0.671466 | 0.171039 | -3.925796 | 8.64E-05 | 0.007948085 |
| *BRK1* | -0.247383 | 0.063032 | -3.924724 | 8.68E-05 | 0.007948085 |
| *IGFBP4* | -0.870549 | 0.222294 | -3.916213 | 9.00E-05 | 0.008158297 |
| *SLC8B1* | -0.974201 | 0.249295 | -3.907826 | 9.31E-05 | 0.00836988 |
| *MPHOSPH10* | 0.408308 | 0.10459 | 3.903899 | 9.47E-05 | 0.008430274 |
| *SRSF5* | -0.339977 | 0.08737 | -3.891237 | 9.97E-05 | 0.008652686 |
| *ENSG00000237181* | 0.853165 | 0.219318 | 3.890077 | 0.000100213 | 0.008652686 |
| *ARNTL2* | -1.118711 | 0.287736 | -3.887982 | 0.000101081 | 0.008652686 |
| *BUD23* | 0.380294 | 0.097901 | 3.884484 | 0.000102547 | 0.008652686 |
| *MRPL44* | 0.418568 | 0.107788 | 3.883244 | 0.000103072 | 0.008652686 |
| *EIF5* | 0.335682 | 0.086451 | 3.882894 | 0.00010322 | 0.008652686 |
| *HOMER2* | -1.03943 | 0.267744 | -3.882177 | 0.000103526 | 0.008652686 |
| *HIST1H4C* | 0.480682 | 0.123904 | 3.879464 | 0.000104687 | 0.008652686 |
| *SGPL1* | -0.473697 | 0.122129 | -3.878669 | 0.00010503 | 0.008652686 |
| *NFE2L2* | 0.320994 | 0.082971 | 3.868761 | 0.00010939 | 0.008937418 |
| *NDUFA7* | -0.441474 | 0.114345 | -3.860877 | 0.000112981 | 0.009155148 |
| *SMIM10* | -3.542224 | 0.918152 | -3.857994 | 0.000114321 | NA |
| *COMMD2* | -0.34333 | 0.089115 | -3.85266 | 0.000116842 | 0.009391039 |
| *C16orf74* | -0.742968 | 0.193033 | -3.84892 | 0.00011864 | 0.009458667 |
| *PPM1F* | -0.801184 | 0.208463 | -3.843288 | 0.000121397 | 0.009601027 |
| *ENSG00000267002* | -0.595631 | 0.155071 | -3.841026 | 0.000122521 | 0.009613034 |
| *MAP2K3* | 0.282386 | 0.07356 | 3.838858 | 0.000123608 | 0.009621937 |
| *FNBP1* | -0.300945 | 0.078446 | -3.836358 | 0.000124872 | 0.009644448 |
| *CCDC115* | -0.388487 | 0.101412 | -3.830769 | 0.000127743 | 0.009789707 |
| *IL32* | -4.463641 | 1.166944 | -3.82507 | 0.000130735 | NA |
| *PDE6D* | -0.554787 | 0.145111 | -3.823187 | 0.000131738 | 0.01001816 |
| *SLC7A6* | 0.77807 | 0.203619 | 3.821197 | 0.000132805 | 0.010022242 |
| *SNHG30* | -0.510018 | 0.13367 | -3.815507 | 0.000135903 | 0.010178345 |
| *NFKBIE* | 0.48957 | 0.12866 | 3.805153 | 0.000141717 | 0.010474609 |
| *DMAC2L* | -0.473061 | 0.124336 | -3.804696 | 0.000141978 | 0.010474609 |
| *GLS* | 0.441147 | 0.116473 | 3.787558 | 0.000152135 | 0.011140783 |
| *LRRC7* | -1.34822 | 0.356164 | -3.785394 | 0.000153465 | 0.011155571 |
| *CLECL1* | 0.921328 | 0.24429 | 3.771454 | 0.000162299 | 0.011711587 |

| | | | | | |
|---|---|---|---|---|---|
| *SEPTIN9* | -0.324157 | 0.086078 | -3.765849 | 0.000165984 | 0.011890706 |
| *CDC42* | -0.255972 | 0.068113 | -3.758019 | 0.000171264 | 0.012116529 |
| *SUPV3L1* | 0.488269 | 0.129952 | 3.757314 | 0.000171747 | 0.012116529 |
| *BLVRB* | 0.903571 | 0.240582 | 3.755765 | 0.000172813 | 0.012116529 |
| *NPW* | 2.367068 | 0.630913 | 3.751811 | 0.000175562 | NA |
| *NUAK2* | -0.519425 | 0.138455 | -3.751564 | 0.000175735 | 0.012234593 |
| *DDX24* | 0.250263 | 0.06697 | 3.73693 | 0.00018628 | 0.012878102 |
| *CCT8* | 0.383085 | 0.102607 | 3.733512 | 0.000188828 | 0.012963556 |
| *PER2* | 0.72825 | 0.195617 | 3.722845 | 0.00019699 | 0.013430658 |
| *SERTAD3* | 0.495462 | 0.13321 | 3.719407 | 0.000199691 | 0.013521553 |
| *PPM1D* | 0.587289 | 0.15818 | 3.712796 | 0.000204982 | 0.013785371 |
| *CDC42EP3* | 0.752321 | 0.203027 | 3.705518 | 0.00021096 | 0.014091527 |
| *KLHL2* | -0.693865 | 0.187502 | -3.700566 | 0.000215119 | 0.014156218 |
| *ENSG00000055483* | 0.733255 | 0.198198 | 3.699606 | 0.000215935 | 0.014156218 |
| *BTLA* | -0.694274 | 0.187679 | -3.699266 | 0.000216224 | 0.014156218 |
| *MYO1C* | -0.489504 | 0.132524 | -3.693698 | 0.000221016 | 0.014374796 |
| *COMMD3* | -0.373967 | 0.101508 | -3.6841 | 0.000229512 | 0.014829801 |
| *B3GNT7* | 0.684245 | 0.186015 | 3.678441 | 0.000234664 | 0.015064221 |
| *C12orf29* | 0.461219 | 0.125471 | 3.675915 | 0.000236999 | 0.015115938 |
| *VPS28* | -0.281936 | 0.076754 | -3.67322 | 0.000239513 | 0.01517836 |
| *EGR3* | 1.586717 | 0.432478 | 3.668899 | 0.000243597 | 0.01526504 |
| *GADD45A* | 0.893827 | 0.243649 | 3.668509 | 0.000243969 | 0.01526504 |
| *COQ10B* | 0.315722 | 0.08615 | 3.664784 | 0.000247548 | 0.015391546 |
| *CLEC2D* | -0.451514 | 0.123362 | -3.660069 | 0.000252147 | 0.015579539 |
| *CD5* | -2.278785 | 0.622824 | -3.658793 | 0.000253406 | NA |
| *XRRA1* | -1.08298 | 0.296433 | -3.653374 | 0.000258816 | 0.015892287 |
| *GLRX2* | 0.456387 | 0.125056 | 3.649451 | 0.000262801 | 0.016037353 |
| *PRAG1* | 1.710855 | 0.46922 | 3.64617 | 0.000266178 | 0.016134923 |
| *PUS7* | 0.75419 | 0.206926 | 3.644738 | 0.000267664 | 0.016134923 |
| *ENSG00000283013* | -0.676218 | 0.185723 | -3.640999 | 0.000271582 | 0.016201082 |
| *ENSG00000224505* | -0.789046 | 0.216875 | -3.63826 | 0.000274486 | 0.016201082 |
| *ESF1* | 0.320033 | 0.088 | 3.636722 | 0.00027613 | 0.016201082 |
| *SORL1* | -0.542158 | 0.149135 | -3.635358 | 0.000277595 | 0.016201082 |
| *COMMD8* | -0.350362 | 0.096387 | -3.634967 | 0.000278016 | 0.016201082 |
| *DHX57* | -0.867011 | 0.238555 | -3.634431 | 0.000278594 | 0.016201082 |
| *ENSG00000277654* | -0.770691 | 0.212343 | -3.629471 | 0.000284002 | 0.016337385 |
| *C1GALT1* | 0.48618 | 0.134027 | 3.627473 | 0.000286209 | 0.016337385 |
| *CDC37L1* | 0.449611 | 0.123975 | 3.626622 | 0.000287153 | 0.016337385 |
| *AKIRIN1* | 0.291757 | 0.080457 | 3.626267 | 0.000287549 | 0.016337385 |
| *NDUFB10* | -0.282556 | 0.078016 | -3.621782 | 0.00029258 | 0.016475267 |
| *ENSG00000257275* | -0.982007 | 0.27131 | -3.619498 | 0.000295175 | 0.016475267 |
| *SNX29* | -0.66782 | 0.184532 | -3.618991 | 0.000295754 | 0.016475267 |
| *TERF2IP* | 0.339884 | 0.093937 | 3.618215 | 0.000296641 | 0.016475267 |
| *PEX6* | 1.354965 | 0.374752 | 3.615626 | 0.000299623 | 0.016547873 |
| *CYREN* | -0.382263 | 0.105887 | -3.610115 | 0.000306061 | 0.016803429 |
| *TYW3* | 0.552146 | 0.153001 | 3.608772 | 0.000307649 | 0.016803429 |
| *LCN8* | 3.870969 | 1.072845 | 3.608136 | 0.000308405 | NA |

| | | | | | |
|---|---|---|---|---|---|
| *HMGXB4* | -0.350484 | 0.097223 | -3.604971 | 0.000312188 | 0.016888766 |
| *GNL3* | 0.485154 | 0.134593 | 3.604605 | 0.000312628 | 0.016888766 |
| *BLOC1S1* | -0.319289 | 0.088778 | -3.596473 | 0.000322561 | 0.017298732 |
| *ABTB1* | -0.46612 | 0.129732 | -3.592957 | 0.000326947 | 0.017298732 |
| *RRN3* | 0.450646 | 0.125425 | 3.592953 | 0.000326952 | 0.017298732 |
| *FCRL4* | 1.866408 | 0.519494 | 3.592742 | 0.000327217 | 0.017298732 |
| *ADGRE5* | 0.50164 | 0.139887 | 3.586035 | 0.000335744 | 0.017655159 |
| *MXD1* | 0.724635 | 0.202191 | 3.583917 | 0.000338479 | 0.017704798 |
| *RCL1* | 0.571303 | 0.160129 | 3.567772 | 0.00036003 | 0.018662088 |
| *TTPAL* | -0.437013 | 0.122502 | -3.567389 | 0.000360556 | 0.018662088 |
| *JMJD6* | 0.38856 | 0.109021 | 3.564072 | 0.000365145 | 0.018801188 |
| *YWHAE* | 0.346754 | 0.097386 | 3.560629 | 0.000369967 | 0.018950754 |
| *PILRB* | 0.694769 | 0.195277 | 3.55787 | 0.000373874 | 0.019052178 |
| *EIF3A* | 0.293979 | 0.082811 | 3.549988 | 0.000385248 | 0.019531104 |
| *CCDC28B* | -0.587041 | 0.165665 | -3.54355 | 0.000394779 | 0.019837359 |
| *C15orf65* | 1.904351 | 0.537795 | 3.541035 | 0.00039856 | 0.019837359 |
| *NRDE2* | -0.460799 | 0.130138 | -3.540849 | 0.000398841 | 0.019837359 |
| *NOP56* | 0.439343 | 0.124089 | 3.540536 | 0.000399316 | 0.019837359 |
| *NDUFAF4* | 0.55782 | 0.157828 | 3.534363 | 0.00040876 | 0.020191321 |
| *FOSL1* | 1.748144 | 0.494773 | 3.533223 | 0.000410526 | 0.020191321 |
| *PRMT5* | 0.480171 | 0.136187 | 3.525831 | 0.000422157 | 0.020660591 |
| *ENSG00000273015* | -0.444634 | 0.126354 | -3.518963 | 0.000433237 | 0.021069704 |
| *CEP95* | 0.455904 | 0.129591 | 3.518021 | 0.000434778 | 0.021069704 |
| *RFK* | 0.412542 | 0.117535 | 3.509963 | 0.000448168 | 0.021612647 |
| *EIPR1* | -0.461538 | 0.131549 | -3.508486 | 0.000450665 | 0.021627543 |
| *PRMT3* | 0.87902 | 0.250795 | 3.504938 | 0.000456714 | 0.021645685 |
| *CHRNB1* | -0.591314 | 0.168716 | -3.504786 | 0.000456974 | 0.021645685 |
| *TSTD1* | -0.486489 | 0.138822 | -3.504415 | 0.000457612 | 0.021645685 |
| *HSP90AA1* | 0.326743 | 0.093291 | 3.502401 | 0.000461085 | 0.021706105 |
| *NFRKB* | -0.615612 | 0.175864 | -3.500502 | 0.000464383 | 0.021757763 |
| *H3F3B* | 0.21481 | 0.061444 | 3.496023 | 0.000472249 | 0.021944933 |
| *NUCB2* | -0.920566 | 0.263342 | -3.495702 | 0.000472817 | 0.021944933 |
| *BYSL* | 0.874043 | 0.250316 | 3.491762 | 0.000479846 | 0.022167088 |
| *IGHD* | -0.66644 | 0.191637 | -3.477607 | 0.000505912 | 0.023262524 |
| *SLC1A5* | 0.674009 | 0.194041 | 3.473538 | 0.000513644 | 0.023451031 |
| *BAG2* | 0.564351 | 0.162537 | 3.472132 | 0.000516341 | 0.023451031 |
| *COMMD10* | 0.722497 | 0.208109 | 3.471724 | 0.000517128 | 0.023451031 |
| *DNAJB4* | 0.618887 | 0.178428 | 3.468547 | 0.00052328 | 0.023621688 |
| *LRRC61* | -1.038709 | 0.299999 | -3.462374 | 0.000535432 | 0.024060374 |
| *FAM91A1* | 0.491471 | 0.142085 | 3.458994 | 0.000542196 | 0.024184367 |
| *HLX* | 1.115258 | 0.322523 | 3.457915 | 0.000544372 | 0.024184367 |
| *GYG1* | 0.583577 | 0.168816 | 3.456885 | 0.000546458 | 0.024184367 |
| *CXCR3* | 2.251813 | 0.651649 | 3.455562 | 0.000549146 | 0.024184367 |
| *SLAMF7* | 1.083111 | 0.313497 | 3.454936 | 0.000550423 | 0.024184367 |
| *ENSG00000229043* | -1.680558 | 0.48645 | -3.45474 | 0.000550824 | NA |
| *TMEM273* | 1.475328 | 0.427213 | 3.45338 | 0.000553608 | 0.024216682 |
| *HSPA8* | 0.485957 | 0.140771 | 3.452108 | 0.000556225 | 0.024223974 |

| | | | | | |
|---|---|---|---|---|---|
| *USE1* | -0.296884 | 0.086133 | -3.446806 | 0.000567255 | 0.024588172 |
| *ATF6B* | -0.310929 | 0.090236 | -3.44571 | 0.000569562 | 0.024588172 |
| *COX4I1* | -0.181102 | 0.052584 | -3.444063 | 0.000573043 | 0.024630875 |
| *KRT10* | -0.212414 | 0.061922 | -3.43037 | 0.000602759 | 0.025691031 |
| *TTC19* | -0.366355 | 0.106811 | -3.429917 | 0.000603767 | 0.025691031 |
| *RB1* | -0.361462 | 0.105409 | -3.429137 | 0.000605504 | 0.025691031 |
| *PDPR* | -0.924242 | 0.269646 | -3.427609 | 0.000608921 | 0.025725614 |
| *MRPL28* | -0.279958 | 0.081816 | -3.421811 | 0.000622055 | 0.026126205 |
| *MIDN* | 0.440615 | 0.128845 | 3.419717 | 0.000626863 | 0.026126205 |
| *ZBED1* | -0.428249 | 0.125244 | -3.419327 | 0.000627762 | 0.026126205 |
| *ERGIC1* | -0.609334 | 0.17823 | -3.418802 | 0.000628974 | 0.026126205 |
| *RNGTT* | -0.512928 | 0.150255 | -3.413709 | 0.00064085 | 0.026481618 |
| *UROD* | -0.398973 | 0.116903 | -3.412844 | 0.000642888 | 0.026481618 |
| *CENPH* | -0.644743 | 0.189429 | -3.403608 | 0.00066502 | 0.027173437 |
| *SMIM27* | -0.403514 | 0.118558 | -3.40351 | 0.000665261 | 0.027173437 |
| *MPP6* | 0.647434 | 0.190286 | 3.402416 | 0.000667929 | 0.027173437 |
| *ZNF627* | -0.706108 | 0.207965 | -3.395323 | 0.000685478 | 0.027773088 |
| *WDR13* | -0.342481 | 0.100976 | -3.391721 | 0.00069455 | 0.02801477 |
| *NIFK* | 0.555046 | 0.163696 | 3.390713 | 0.00069711 | 0.02801477 |
| *KLF6* | 0.310981 | 0.091747 | 3.389538 | 0.000700104 | 0.028021176 |
| *TFB2M* | 0.601725 | 0.177911 | 3.38217 | 0.000719155 | 0.028667625 |
| *EPS15* | -0.439301 | 0.130001 | -3.379222 | 0.000726913 | 0.028756964 |
| *RAB10* | 0.306805 | 0.090795 | 3.379089 | 0.000727264 | 0.028756964 |
| *ORMDL3* | -0.534946 | 0.158361 | -3.37801 | 0.000730123 | 0.028756964 |
| *ALOX5AP* | -0.724149 | 0.214465 | -3.376535 | 0.000734051 | 0.028796951 |
| *TDG* | 0.343715 | 0.101867 | 3.374146 | 0.00074045 | 0.028933162 |
| *GNL2* | 0.56509 | 0.167681 | 3.370042 | 0.000751568 | 0.029104937 |
| *USF1* | -1.035801 | 0.307448 | -3.369023 | 0.000754352 | 0.029104937 |
| *ENSG00000275964* | -1.026401 | 0.304739 | -3.368129 | 0.000756803 | 0.029104937 |
| *RPF2* | 0.493952 | 0.146664 | 3.367907 | 0.000757412 | 0.029104937 |
| *LAMTOR4* | -0.288598 | 0.085711 | -3.367124 | 0.000759566 | 0.029104937 |
| *TEX30* | 0.416212 | 0.123669 | 3.365527 | 0.000763976 | 0.029160887 |
| *NMT2* | -0.729419 | 0.217101 | -3.359818 | 0.000779939 | 0.02961263 |
| *TAGLN2* | -0.307227 | 0.091459 | -3.359158 | 0.000781802 | 0.02961263 |
| *HDDC2* | -0.401636 | 0.119721 | -3.354776 | 0.000794292 | 0.029970877 |
| *PRAF2* | -0.505899 | 0.151111 | -3.347872 | 0.000814346 | 0.030610731 |
| *EIF4A3* | 0.439458 | 0.131369 | 3.345205 | 0.000822217 | 0.030789527 |
| *ARL8B* | 0.413481 | 0.123685 | 3.34302 | 0.000828719 | 0.030857504 |
| *OXLD1* | -0.402846 | 0.120524 | -3.342468 | 0.000830368 | 0.030857504 |
| *STIP1* | 0.410878 | 0.122985 | 3.340872 | 0.000835158 | 0.030857504 |
| *METTL26* | -0.358694 | 0.10738 | -3.34042 | 0.000836517 | 0.030857504 |
| *NABP1* | 0.413595 | 0.123987 | 3.335801 | 0.000850541 | 0.031193507 |
| *UGT8* | 0.926004 | 0.27772 | 3.334306 | 0.000855125 | 0.031193507 |
| *ENSG00000259321* | -0.796999 | 0.239153 | -3.332593 | 0.000860407 | 0.031193507 |
| *NEDD4L* | 1.750494 | 0.525278 | 3.332509 | 0.000860667 | 0.031193507 |
| *TXN* | 0.378739 | 0.11368 | 3.33164 | 0.000863358 | 0.031193507 |
| *NR1D2* | 0.651331 | 0.195552 | 3.33073 | 0.000866187 | 0.031193507 |

| | | | | | |
|---|---|---|---|---|---|
| *PFKFB3* | -0.461751 | 0.138654 | -3.330239 | 0.000867713 | 0.031193507 |
| *EEA1* | 0.406694 | 0.122288 | 3.325715 | 0.00088192 | 0.031544912 |
| *TXLNG* | 0.494112 | 0.148601 | 3.325099 | 0.00088387 | 0.031544912 |
| *DBI* | -0.41222 | 0.124034 | -3.323449 | 0.000889118 | 0.031618051 |
| *SLC50A1* | -0.374451 | 0.112718 | -3.322023 | 0.000893672 | 0.031666114 |
| *MED23* | -0.618517 | 0.18628 | -3.32036 | 0.000899016 | 0.031741683 |
| *NDE1* | -0.408583 | 0.123263 | -3.314718 | 0.000917355 | 0.032273933 |
| *RBM25* | 0.250439 | 0.075616 | 3.311994 | 0.000926334 | 0.032474255 |
| *SLC35F6* | -0.540641 | 0.163615 | -3.304347 | 0.000951978 | 0.033150549 |
| *IGKC* | -0.437217 | 0.132324 | -3.304148 | 0.000952656 | 0.033150549 |
| *ELOVL5* | 0.311669 | 0.094352 | 3.303258 | 0.000955685 | 0.033150549 |
| *TBC1D2B* | -0.91796 | 0.278332 | -3.298081 | 0.00097348 | 0.033346802 |
| *IGKV3.11* | -1.313108 | 0.398171 | -3.297853 | 0.000974273 | 0.033346802 |
| *MME* | -3.059808 | 0.927958 | -3.297358 | 0.000975991 | NA |
| *NCBP2AS2* | -0.333481 | 0.101158 | -3.296631 | 0.00097852 | 0.033346802 |
| *AKAP1* | 0.735645 | 0.223187 | 3.296092 | 0.000980398 | 0.033346802 |
| *DESI1* | 0.457862 | 0.138922 | 3.295827 | 0.000981323 | 0.033346802 |
| *NHLRC3* | -0.542112 | 0.164488 | -3.295753 | 0.000981582 | 0.033346802 |
| *HDAC1* | -0.34021 | 0.103291 | -3.293696 | 0.000988794 | 0.033392489 |
| *SMARCE1* | -0.257639 | 0.078228 | -3.293443 | 0.000989682 | 0.033392489 |
| *IL13RA1* | -0.516074 | 0.157287 | -3.281107 | 0.001034005 | 0.034762575 |
| *RAB1A* | 0.342263 | 0.104342 | 3.280204 | 0.001037321 | 0.034762575 |
| *LSM3* | -0.179841 | 0.054954 | -3.27258 | 0.001065708 | 0.03544463 |
| *ENSG00000273272* | 2.400283 | 0.733847 | 3.270821 | 0.001072357 | 0.03544463 |
| *PIKFYVE* | 0.489691 | 0.149718 | 3.270757 | 0.0010726 | 0.03544463 |
| *ZPR1* | 0.351946 | 0.107604 | 3.270739 | 0.001072669 | 0.03544463 |
| *ALG12* | -0.602471 | 0.184244 | -3.269967 | 0.001075601 | 0.03544463 |
| *CNOT6L* | -0.572825 | 0.175465 | -3.264611 | 0.001096146 | 0.03600166 |
| *PEX13* | 0.572294 | 0.175387 | 3.263044 | 0.001102224 | 0.036081409 |
| *LPIN1* | 0.61914 | 0.189905 | 3.260269 | 0.001113067 | 0.036316113 |
| *SYNGR2* | -0.309925 | 0.095151 | -3.257209 | 0.001125135 | 0.036589108 |
| *MTMR4* | -0.596503 | 0.183267 | -3.254826 | 0.001134617 | 0.036776474 |
| *ERV3.1* | -0.373353 | 0.114788 | -3.252549 | 0.001143747 | 0.036928072 |
| *ENSG00000279483* | -0.393798 | 0.121102 | -3.2518 | 0.001146765 | 0.036928072 |
| *HNRNPA2B1* | 0.233781 | 0.071936 | 3.249843 | 0.001154687 | 0.037009148 |
| *IER5* | 0.348513 | 0.107257 | 3.249331 | 0.00115677 | 0.037009148 |
| *COA7* | 0.775145 | 0.238669 | 3.247788 | 0.00116306 | 0.037090372 |
| *COPS3* | 0.395851 | 0.122002 | 3.244631 | 0.001176029 | 0.037339413 |
| *NOP16* | 0.476449 | 0.146869 | 3.244052 | 0.001178424 | 0.037339413 |
| *STRBP* | -0.322054 | 0.099338 | -3.241989 | 0.001186985 | 0.037441768 |
| *FLCN* | 0.605237 | 0.186718 | 3.241451 | 0.001189229 | 0.037441768 |
| *EIF2AK4* | -0.541861 | 0.167304 | -3.23878 | 0.001200422 | 0.037674185 |
| *FAM30A* | -0.381707 | 0.117979 | -3.235385 | 0.001214787 | 0.038004374 |
| *CDK5R1* | 2.59867 | 0.803798 | 3.232987 | 0.001225031 | NA |
| *PTPN22* | 0.88927 | 0.27521 | 3.231242 | 0.001232534 | 0.038332738 |
| *VAMP8* | -0.226835 | 0.070203 | -3.231126 | 0.001233038 | 0.038332738 |
| *SVBP* | -0.326746 | 0.101228 | -3.227821 | 0.001247371 | 0.038564219 |

| | | | | | |
|---|---|---|---|---|---|
| *POLR3D* | 0.534481 | 0.165601 | 3.227527 | 0.001248654 | 0.038564219 |
| *VDAC1* | 0.424042 | 0.13142 | 3.226624 | 0.001252599 | 0.038564219 |
| *ZDHHC12* | -0.382687 | 0.118664 | -3.224958 | 0.00125991 | 0.038564219 |
| *NDUFB7* | -0.348012 | 0.107942 | -3.224067 | 0.001263838 | 0.038564219 |
| *ERF* | 0.613262 | 0.190241 | 3.223611 | 0.001265853 | 0.038564219 |
| *TNFRSF10B* | 0.392048 | 0.121634 | 3.223173 | 0.00126779 | 0.038564219 |
| *ARL6IP1* | 0.255948 | 0.079448 | 3.221585 | 0.001274838 | 0.038659645 |
| *DYNLL1* | 0.341547 | 0.106118 | 3.218568 | 0.001288326 | 0.038865253 |
| *CTNND1* | -0.666803 | 0.20719 | -3.218311 | 0.00128948 | 0.038865253 |
| *ENSG00000224610* | -2.788744 | 0.867077 | -3.21626 | 0.001298731 | NA |
| *ARCN1* | 0.45143 | 0.14067 | 3.209137 | 0.00133134 | 0.039669698 |
| *H1FX* | -0.852359 | 0.265655 | -3.208521 | 0.001334196 | 0.039669698 |
| *P2RY8* | -0.499522 | 0.155718 | -3.207862 | 0.001337259 | 0.039669698 |
| *AKIRIN2* | 0.271007 | 0.084503 | 3.207057 | 0.001341006 | 0.039669698 |
| *SNX17* | -0.237403 | 0.07403 | -3.206855 | 0.001341948 | 0.039669698 |
| *KRAS* | 0.263672 | 0.082225 | 3.206702 | 0.001342662 | 0.039669698 |
| *SLC35B2* | 0.753584 | 0.235028 | 3.206359 | 0.001344259 | 0.039669698 |
| *TMEM219* | -0.334469 | 0.104475 | -3.201436 | 0.001367447 | 0.040132837 |
| *GLA* | 0.715302 | 0.223441 | 3.201304 | 0.001368073 | 0.040132837 |
| *SCRN1* | 0.995163 | 0.311043 | 3.199437 | 0.001376963 | 0.040274119 |
| *CCDC71* | -0.477753 | 0.14962 | -3.193109 | 0.0014075 | 0.040925833 |
| *SLC16A3* | 0.641545 | 0.200916 | 3.193104 | 0.001407524 | 0.040925833 |
| *C15orf62* | -0.863952 | 0.27078 | -3.190607 | 0.001419743 | 0.041160047 |
| *RCC1* | 0.379239 | 0.119014 | 3.186516 | 0.001439974 | 0.041624513 |
| *ENSG00000225205* | -1.118996 | 0.351378 | -3.184589 | 0.001449595 | 0.041780451 |
| *NOP58* | 0.36613 | 0.115005 | 3.183586 | 0.001454631 | 0.041803714 |
| *ARF5* | -0.414133 | 0.130193 | -3.180906 | 0.001468155 | 0.042070086 |
| *SNHG25* | -0.660432 | 0.207843 | -3.177552 | 0.001485239 | 0.042350325 |
| *PYGO2* | -0.421619 | 0.132709 | -3.177008 | 0.001488028 | 0.042350325 |
| *ZNF605* | -0.891716 | 0.280743 | -3.176269 | 0.001491823 | 0.042350325 |
| *SLC6A6* | -0.375541 | 0.118257 | -3.175639 | 0.00149507 | 0.042350325 |
| *NAMPT* | 0.645435 | 0.203405 | 3.173159 | 0.001507897 | 0.04252758 |
| *ATP5IF1* | -0.394101 | 0.124214 | -3.172768 | 0.001509931 | 0.04252758 |
| *BDP1* | 0.227355 | 0.071715 | 3.170282 | 0.001522909 | 0.042771256 |
| *DDAH2* | 0.441207 | 0.139439 | 3.164161 | 0.001555307 | 0.04349661 |
| *ODF3B* | -0.582021 | 0.183968 | -3.163711 | 0.001557713 | 0.04349661 |
| *PPP2R2A* | 0.337608 | 0.106754 | 3.162474 | 0.001564349 | 0.04349661 |
| *ATP1A1* | 0.470394 | 0.14876 | 3.162104 | 0.001566336 | 0.04349661 |
| *CYBC1* | -0.287712 | 0.091133 | -3.157074 | 0.001593607 | 0.044075452 |
| *MOB4* | 0.276373 | 0.087553 | 3.156619 | 0.001596097 | 0.044075452 |
| *ITGAX* | 2.297958 | 0.728047 | 3.156333 | 0.001597663 | NA |
| *MDK* | -1.083175 | 0.343258 | -3.155574 | 0.001601828 | 0.044110516 |
| *RPP25* | 0.83756 | 0.265612 | 3.153324 | 0.001614224 | 0.044328386 |
| *LARP4* | 0.466593 | 0.148064 | 3.151288 | 0.001625521 | 0.044514965 |
| *MIS12* | 0.38037 | 0.120778 | 3.149327 | 0.001636469 | 0.044690965 |
| *DHX15* | 0.348862 | 0.110845 | 3.147309 | 0.001647806 | 0.04475536 |
| *SYNC* | -0.484734 | 0.154016 | -3.147296 | 0.001647881 | 0.04475536 |

| | | | | | |
|---|---|---|---|---|---|
| *WDR33* | 0.340708 | 0.108331 | 3.145068 | 0.00166048 | 0.044870024 |
| *COL9A3* | -1.767209 | 0.561921 | -3.144945 | 0.00166118 | 0.044870024 |
| *ZMAT2* | -0.230714 | 0.073398 | -3.14334 | 0.001670319 | 0.044993946 |
| *CHD1* | 0.296982 | 0.094715 | 3.13555 | 0.001715323 | 0.046080659 |
| *ENSG00000278834* | -1.455694 | 0.465069 | -3.130061 | 0.001747699 | NA |
| *TLN1* | -0.316443 | 0.101143 | -3.12867 | 0.001755994 | 0.047045418 |
| *RNF168* | -0.434804 | 0.139113 | -3.125541 | 0.001774781 | 0.047350316 |
| *RSL1D1* | 0.335578 | 0.107379 | 3.125181 | 0.001776954 | 0.047350316 |
| *TEX10* | 0.460703 | 0.147707 | 3.119027 | 0.001814495 | 0.048220683 |
| *ADAMTS6* | 1.72907 | 0.554428 | 3.118653 | 0.001816796 | NA |
| *C1orf122* | -0.390939 | 0.125582 | -3.113011 | 0.001851893 | 0.048985376 |
| *HLA.G* | -1.671651 | 0.537039 | -3.112717 | 0.001853736 | 0.048985376 |
| *ACTR1B* | -0.460377 | 0.147946 | -3.111796 | 0.001859528 | 0.048985376 |
| *PPCS* | -0.269473 | 0.086632 | -3.110565 | 0.001867298 | 0.048985376 |
| *ZMAT3* | -0.494285 | 0.158911 | -3.110447 | 0.001868044 | 0.048985376 |
| *NCL* | 0.339941 | 0.109485 | 3.104898 | 0.001903447 | 0.049781686 |
| *ZNF428* | -0.311832 | 0.100523 | -3.102102 | 0.00192152 | 0.050121767 |
| *FANCB* | -0.869468 | 0.280523 | -3.099458 | 0.001938753 | 0.050359414 |
| *ADSS* | 0.378201 | 0.122034 | 3.099142 | 0.001940819 | 0.050359414 |
| *UQCR10* | -0.175256 | 0.056623 | -3.095171 | 0.001966994 | 0.050784457 |
| *TAF6* | -0.407319 | 0.131601 | -3.095099 | 0.001967474 | 0.050784457 |
| *GEMIN4* | 0.644821 | 0.208756 | 3.088874 | 0.002009164 | 0.051501488 |
| *STAT3* | 0.329695 | 0.106785 | 3.087473 | 0.00201866 | 0.051501488 |
| *PRPF38B* | 0.203355 | 0.065865 | 3.087459 | 0.002018757 | 0.051501488 |
| *RNF26* | -0.441348 | 0.142955 | -3.087332 | 0.002019622 | 0.051501488 |
| *AGTRAP* | -0.728439 | 0.235963 | -3.087085 | 0.002021301 | 0.051501488 |
| *RFFL* | -0.348922 | 0.113141 | -3.08396 | 0.00204265 | 0.051792051 |
| *PUS1* | 0.475695 | 0.15428 | 3.083323 | 0.002047027 | 0.051792051 |
| *FIBP* | -0.28184 | 0.091414 | -3.083121 | 0.002048421 | 0.051792051 |
| *DCTN1* | -0.457377 | 0.148417 | -3.081711 | 0.002058147 | 0.051905201 |
| *TMEM123* | -0.403772 | 0.13107 | -3.080589 | 0.002065913 | 0.051968484 |
| *RCHY1* | 0.325138 | 0.105604 | 3.078835 | 0.002078121 | 0.05198793 |
| *UTP6* | 0.368739 | 0.119774 | 3.078619 | 0.002079621 | 0.05198793 |
| *DNAJA2* | 0.28549 | 0.092749 | 3.078087 | 0.002083339 | 0.05198793 |
| *APBB2* | -1.510385 | 0.490789 | -3.077461 | 0.002087721 | 0.05198793 |
| *MOB3A* | -0.293442 | 0.095383 | -3.07647 | 0.00209467 | 0.05202993 |
| *DDX6* | -0.259188 | 0.084273 | -3.075585 | 0.002100902 | 0.052053921 |
| *IQSEC1* | -0.378106 | 0.123 | -3.074046 | 0.002111769 | 0.052192372 |
| *EIF4E* | 0.37061 | 0.120631 | 3.072254 | 0.002124487 | 0.052310772 |
| *AP2M1* | -0.23329 | 0.075944 | -3.071881 | 0.002127142 | 0.052310772 |
| *NFAT5* | 0.424549 | 0.138336 | 3.068965 | 0.002148019 | 0.052685745 |
| *HSP90AB1* | 0.448568 | 0.14623 | 3.067559 | 0.002158146 | 0.052685745 |
| *CHCHD5* | -0.463726 | 0.151193 | -3.067106 | 0.002161422 | 0.052685745 |
| *IVNS1ABP* | 0.508949 | 0.165955 | 3.06679 | 0.002163708 | 0.052685745 |
| *TFEC* | 2.378492 | 0.775583 | 3.066714 | NA | NA |
| *GCSAM* | -0.918239 | 0.299668 | -3.064192 | 0.002182591 | 0.053014973 |
| *JTB* | -0.193223 | 0.063076 | -3.063353 | 0.00218872 | 0.053033552 |

| | | | | | |
|---|---|---|---|---|---|
| *ENSG00000273156* | -1.148535 | 0.376102 | -3.053786 | 0.00225973 | 0.05462027 |
| *HCG18* | 0.388026 | 0.12713 | 3.052201 | 0.002271702 | 0.05477573 |
| *RAB11B* | -0.265055 | 0.086866 | -3.051322 | 0.002278362 | 0.054802646 |
| *ALKBH4* | -0.470894 | 0.154419 | -3.049449 | 0.002292617 | 0.055011684 |
| *QPCT* | 0.794921 | 0.260762 | 3.048448 | 0.002300265 | 0.055061556 |
| *UPF2* | -0.221209 | 0.072634 | -3.045551 | 0.002322542 | 0.055432884 |
| *NECAP2* | -0.231867 | 0.076147 | -3.044976 | 0.002326992 | 0.055432884 |
| *HNRNPU* | 0.279532 | 0.091981 | 3.039006 | 0.002373602 | 0.056291085 |
| *RNF19A* | 0.517102 | 0.170161 | 3.038904 | 0.002374406 | 0.056291085 |
| *TXNL1* | 0.283732 | 0.093392 | 3.038081 | 0.002380899 | 0.056309969 |
| *FAM210A* | 0.335889 | 0.110598 | 3.037011 | 0.002389367 | 0.056375375 |
| *LINC01116* | -0.589463 | 0.194328 | -3.033336 | 0.002418663 | 0.05693071 |
| *NDUFB8* | -0.293736 | 0.096931 | -3.030356 | 0.002442658 | 0.057358947 |
| *APOBEC3D* | -0.554345 | 0.183017 | -3.028931 | 0.002454204 | 0.057493519 |
| *ENSG00000277969* | 0.599455 | 0.197976 | 3.027925 | 0.002462394 | 0.05754901 |
| *FLOT2* | -0.409079 | 0.135313 | -3.023214 | 0.002501055 | 0.058016699 |
| *ENSG00000230709* | 1.208414 | 0.399717 | 3.023171 | 0.002501405 | 0.058016699 |
| *KRTCAP2* | -0.231463 | 0.076568 | -3.022971 | 0.002503065 | 0.058016699 |
| *KCTD6* | 0.661969 | 0.219011 | 3.022542 | 0.00250661 | 0.058016699 |
| *BAG4* | 0.497818 | 0.164825 | 3.020277 | 0.002525436 | 0.058016699 |
| *PDE4D* | -0.828446 | 0.274351 | -3.019653 | 0.002530644 | 0.058016699 |
| *CDK4* | 0.353245 | 0.116994 | 3.019344 | 0.00253323 | 0.058016699 |
| *MYCBP2* | 0.227035 | 0.07521 | 3.018695 | 0.002538663 | 0.058016699 |
| *C19orf25* | -0.227423 | 0.075344 | -3.018444 | 0.002540765 | 0.058016699 |
| *MPPE1* | -0.482281 | 0.15978 | -3.018405 | 0.002541092 | 0.058016699 |
| *SNRPB2* | 0.273956 | 0.090822 | 3.016411 | 0.002557864 | 0.058265072 |
| *RGCC* | 2.043628 | 0.678184 | 3.013382 | 0.002583539 | 0.058714634 |
| *ICOSLG* | -0.581677 | 0.193105 | -3.012236 | 0.002593312 | 0.058801566 |
| *UQCR11* | -0.187073 | 0.062126 | -3.011206 | 0.002602121 | 0.05886629 |
| *RRP15* | 0.339734 | 0.112874 | 3.009859 | 0.002613688 | 0.058992965 |
| *NEDD8* | -0.169892 | 0.056467 | -3.008665 | 0.00262398 | 0.059090356 |
| *CLK1* | 0.396913 | 0.132029 | 3.006243 | 0.002644977 | 0.059251537 |
| *LSP1* | -0.407743 | 0.135634 | -3.006194 | 0.002645402 | 0.059251537 |
| *ATF7IP2* | -0.475064 | 0.158051 | -3.005767 | 0.002649118 | 0.059251537 |
| *TANK* | 0.272297 | 0.090712 | 3.001785 | 0.002684016 | 0.059804658 |
| *PCMTD2* | -0.653287 | 0.217701 | -3.000852 | 0.002692253 | 0.059804658 |
| *LINC00339* | 1.024025 | 0.341256 | 3.000756 | 0.002693102 | 0.059804658 |
| *BTN2A2* | -0.46377 | 0.15458 | -3.000198 | 0.002698045 | 0.059804658 |
| *TEAD2* | -2.70088 | 0.900407 | -2.999622 | 0.002703151 | NA |
| *ESRRA* | 0.47024 | 0.156811 | 2.99877 | 0.002710723 | 0.059951246 |
| *LYAR* | 0.520003 | 0.173564 | 2.996025 | 0.002735237 | 0.060358377 |
| *PTPRS* | 2.217846 | 0.74034 | 2.995713 | 0.002738042 | NA |
| *IRF2* | -0.253658 | 0.084772 | -2.992251 | 0.002769284 | 0.060967185 |
| *BIN1* | 0.651319 | 0.217722 | 2.991524 | 0.002775889 | 0.060967185 |
| *SNAP23* | -0.369511 | 0.123552 | -2.990719 | 0.002783219 | 0.060967185 |
| *RGS10* | 0.606567 | 0.202919 | 2.989202 | 0.002797068 | 0.060967185 |
| *DNAJC25* | 0.619816 | 0.207391 | 2.988636 | 0.002802258 | 0.060967185 |

| | | | | | |
|---|---|---|---|---|---|
| *NAA15* | 0.321976 | 0.107749 | 2.9882 | 0.002806263 | 0.060967185 |
| *WDR70* | -0.341121 | 0.11418 | -2.987575 | 0.002812002 | 0.060967185 |
| *CREBRF* | 0.334403 | 0.111943 | 2.987246 | 0.002815028 | 0.060967185 |
| *RRP9* | 0.552456 | 0.184961 | 2.986888 | 0.002818329 | 0.060967185 |
| *MIIP* | -0.488848 | 0.163931 | -2.982036 | 0.002863381 | 0.06180651 |
| *ZNF558* | -0.898206 | 0.301422 | -2.979891 | 0.002883512 | 0.061951821 |
| *FADS3* | -0.485954 | 0.163102 | -2.979453 | 0.002887632 | 0.061951821 |
| *MRPS30* | 0.309159 | 0.103783 | 2.978908 | 0.002892772 | 0.061951821 |
| *SRPRB* | 0.417839 | 0.140278 | 2.978654 | 0.002895179 | 0.061951821 |
| *CYTH1* | -0.221695 | 0.074478 | -2.976638 | 0.002914282 | 0.062124684 |
| *GID8* | -0.285094 | 0.095795 | -2.976095 | 0.002919443 | 0.062124684 |
| *CEP162* | -0.505404 | 0.169837 | -2.975815 | 0.00292211 | 0.062124684 |
| *DNAJB1* | 0.76453 | 0.257071 | 2.974001 | 0.002939442 | 0.062359071 |
| *NEU1* | 0.33784 | 0.113624 | 2.97331 | 0.002946063 | 0.062365692 |
| *HLA.DQB2* | -3.149047 | 1.060136 | -2.970419 | 0.002973937 | NA |
| *SNRNP48* | 0.412657 | 0.139004 | 2.968676 | 0.002990855 | 0.063165199 |
| *MAD2L2* | -0.42312 | 0.142556 | -2.968086 | 0.002996609 | 0.063165199 |
| *IDH2* | -0.450589 | 0.15188 | -2.966744 | 0.003009714 | 0.063240703 |
| *PSMB8.AS1* | -0.379979 | 0.128107 | -2.966096 | 0.003016061 | 0.063240703 |
| *LMBRD1* | -0.274783 | 0.092652 | -2.965758 | 0.003019382 | 0.063240703 |
| *NAPB* | -0.554397 | 0.187555 | -2.955917 | 0.003117411 | 0.065155859 |
| *ZNF79* | 0.7582 | 0.256755 | 2.953007 | 0.003146944 | 0.065458301 |
| *NTMT1* | 0.308503 | 0.104544 | 2.950925 | 0.003168235 | 0.065458301 |
| *EIF4A1* | 0.311876 | 0.105689 | 2.950879 | 0.003168707 | 0.065458301 |
| *SAR1B* | 0.312182 | 0.1058 | 2.95067 | 0.003170858 | 0.065458301 |
| *SERTAD1* | 0.393964 | 0.133572 | 2.949451 | 0.003183388 | 0.065458301 |
| *GEM* | 1.733493 | 0.58794 | 2.948419 | 0.003194033 | 0.065458301 |
| *GPR137* | -0.293155 | 0.09943 | -2.94836 | 0.003194653 | 0.065458301 |
| *PMAIP1* | 0.348358 | 0.118158 | 2.948241 | 0.003195876 | 0.065458301 |
| *GCH1* | 0.536203 | 0.181897 | 2.947835 | 0.003200078 | 0.065458301 |
| *EXOG* | -0.464753 | 0.157668 | -2.947665 | 0.003201837 | 0.065458301 |
| *CHTOP* | 0.192429 | 0.065296 | 2.947007 | 0.003208656 | 0.065458301 |
| *WASF1* | -0.631796 | 0.214418 | -2.946564 | 0.003213255 | 0.065458301 |
| *FAM53B* | -0.697857 | 0.236874 | -2.946112 | 0.003217958 | 0.065458301 |
| *NFATC2* | 0.650542 | 0.220897 | 2.945006 | 0.003229482 | 0.065557831 |
| *VAMP5* | -0.716502 | 0.243403 | -2.943692 | 0.003243222 | 0.065684398 |
| *RBM39* | 0.15217 | 0.051703 | 2.943141 | 0.003249006 | 0.065684398 |
| *ATXN1* | 0.988741 | 0.336073 | 2.942047 | 0.003260502 | 0.06578229 |
| *MIF* | -0.353688 | 0.120276 | -2.940641 | 0.003275342 | 0.065860191 |
| *PHLDA2* | 1.203851 | 0.409415 | 2.940419 | 0.003277687 | 0.065860191 |
| *VKORC1* | -0.260467 | 0.088653 | -2.938048 | 0.003302864 | 0.066231465 |
| *APH1A* | -0.219315 | 0.074714 | -2.935377 | 0.003331425 | 0.066627876 |
| *TXNDC11* | 0.417379 | 0.142211 | 2.934941 | 0.003336112 | 0.066627876 |
| *ATP5ME* | -0.212985 | 0.072662 | -2.931182 | 0.003376754 | 0.067303609 |
| *ARL5B* | 0.657384 | 0.224524 | 2.927903 | 0.003412563 | 0.067880488 |
| *ARID5A* | -0.365629 | 0.125004 | -2.92493 | 0.00344534 | 0.068394839 |
| *ANAPC15* | -0.4625 | 0.158221 | -2.92312 | 0.003465432 | 0.068655836 |

| | | | | | |
|---|---|---|---|---|---|
| *ADAMTS7* | -0.555737 | 0.190273 | -2.920741 | 0.003492 | 0.069043818 |
| *NHSL2* | -0.457172 | 0.156591 | -2.919529 | 0.003505603 | 0.069174434 |
| *ELL2* | 0.529804 | 0.181681 | 2.91612 | 0.003544145 | 0.069719585 |
| *MED4* | -0.203253 | 0.069707 | -2.915839 | 0.003547335 | 0.069719585 |
| *RAN* | 0.369069 | 0.126616 | 2.914877 | 0.003558282 | 0.06975689 |
| *TMOD2* | -0.520623 | 0.178636 | -2.914434 | 0.003563345 | 0.06975689 |
| *BCAR3* | 0.682126 | 0.234188 | 2.912725 | 0.003582894 | 0.069910761 |
| *CCM2* | -0.260263 | 0.08936 | -2.912512 | 0.003585349 | 0.069910761 |
| *GM2A* | 0.77757 | 0.267041 | 2.911799 | 0.003593538 | 0.069932517 |
| *ARID1A* | 0.343499 | 0.118072 | 2.909243 | 0.00362305 | 0.070368307 |
| *PHKB* | -0.313624 | 0.107905 | -2.906487 | 0.003655117 | 0.07079506 |
| *TMX2* | 0.311017 | 0.107021 | 2.906126 | 0.003659344 | 0.07079506 |
| *HEXIM1* | 0.436515 | 0.150286 | 2.904563 | 0.003677656 | 0.070804082 |
| *NPHP3* | 0.635524 | 0.218832 | 2.90417 | 0.003682279 | 0.070804082 |
| *GPN2* | 0.331638 | 0.114235 | 2.903121 | 0.003694641 | 0.070804082 |
| *PCIF1* | -0.390841 | 0.13465 | -2.902656 | 0.003700132 | 0.070804082 |
| *NAB2* | 0.724602 | 0.249691 | 2.901993 | 0.003707974 | 0.070804082 |
| *NRAS* | -0.290094 | 0.099968 | -2.901864 | 0.003709501 | 0.070804082 |
| *DUSP4* | 1.24312 | 0.428434 | 2.90154 | 0.003713329 | 0.070804082 |
| *MTRR* | 0.410652 | 0.141545 | 2.901222 | 0.003717107 | 0.070804082 |
| *CNIH1* | 0.269906 | 0.093092 | 2.89934 | 0.003739496 | 0.071093572 |
| *FADS2* | -1.683043 | 0.581389 | -2.894865 | 0.003793222 | NA |
| *FAM117B* | -0.651686 | 0.225291 | -2.892642 | 0.003820162 | 0.07248775 |
| *TELO2* | -0.482583 | 0.167038 | -2.889057 | 0.003863985 | 0.073178847 |
| *ACSL5* | 0.639744 | 0.221925 | 2.882697 | 0.003942866 | 0.074529964 |
| *CCDC86* | 0.507131 | 0.175963 | 2.882029 | 0.003951228 | 0.074545499 |
| *DEDD2* | 0.434284 | 0.150724 | 2.881327 | 0.003960045 | 0.074569528 |
| *SCOC* | -0.25528 | 0.088627 | -2.880395 | 0.003971775 | 0.074590121 |
| *FOSB* | 0.864587 | 0.300199 | 2.880041 | 0.003976228 | 0.074590121 |
| *VPS37A* | 0.416985 | 0.144905 | 2.87765 | 0.004006492 | 0.0750155 |
| *WDR3* | 0.654003 | 0.227746 | 2.871631 | 0.004083596 | 0.076314611 |
| *ICA1L* | -0.594894 | 0.207212 | -2.870942 | 0.004092503 | 0.076324809 |
| *FOXO3* | 0.530844 | 0.184938 | 2.870396 | 0.004099583 | 0.076324809 |
| *PHF21A* | -0.437675 | 0.152686 | -2.866509 | 0.004150258 | 0.07708713 |
| *THEMIS2* | 1.219728 | 0.425576 | 2.866062 | 0.004156124 | 0.07708713 |
| *PPP1CA* | -0.184288 | 0.064321 | -2.865133 | 0.004168337 | 0.077168871 |
| *RGMB* | 0.934051 | 0.326172 | 2.863678 | 0.004187531 | 0.077379309 |
| *ALKBH1* | 0.473921 | 0.165528 | 2.863087 | 0.004195359 | 0.077379324 |
| *DUSP10* | 0.650248 | 0.227225 | 2.861692 | 0.004213857 | 0.077575767 |
| *ZNF239* | 1.555979 | 0.543988 | 2.860317 | 0.004232173 | NA |
| *PRPF18* | 0.316758 | 0.110845 | 2.857661 | 0.004267758 | 0.07842203 |
| *NDUFB2* | -0.206868 | 0.072429 | -2.856152 | 0.004288103 | 0.0786497 |
| *ENSG00000159882* | -0.451199 | 0.158026 | -2.855221 | 0.004300692 | 0.078734512 |
| *LRP10* | -0.271785 | 0.095232 | -2.85393 | 0.004318198 | 0.078908881 |
| *ING1* | 0.437437 | 0.153312 | 2.853253 | 0.004327422 | 0.078931541 |
| *WARS2* | 0.562595 | 0.197383 | 2.850268 | 0.004368245 | 0.079529418 |
| *ELOC* | 0.264366 | 0.092818 | 2.848208 | 0.004396623 | 0.079898917 |

| | | | | | |
|---|---|---|---|---|---|
| *PVT1* | 0.65944 | 0.231803 | 2.844831 | 0.004443499 | 0.080602631 |
| *SMAP2* | -0.272068 | 0.095665 | -2.843956 | 0.004455713 | 0.080676158 |
| *ENOPH1* | 0.363002 | 0.127729 | 2.841975 | 0.004483494 | 0.081030758 |
| *IPO7* | 0.370151 | 0.130315 | 2.840432 | 0.004505247 | 0.081275313 |
| *ZNF773* | 0.603116 | 0.212393 | 2.839624 | 0.004516677 | 0.081290884 |
| *TAPBP* | -0.294156 | 0.103623 | -2.838723 | 0.004529442 | 0.081290884 |
| *TFG* | 0.344006 | 0.121231 | 2.837616 | 0.004545179 | 0.081290884 |
| *PLIN2* | 1.075425 | 0.379044 | 2.8372 | 0.004551105 | 0.081290884 |
| *PDE3B* | -1.045072 | 0.368403 | -2.836766 | 0.004557306 | 0.081290884 |
| *AKAP2* | -0.911994 | 0.321554 | -2.836205 | 0.004565307 | 0.081290884 |
| *CENPC* | -0.331602 | 0.116938 | -2.835707 | 0.004572432 | 0.081290884 |
| *DNAJA1* | 0.299999 | 0.105802 | 2.835487 | 0.004575588 | 0.081290884 |
| *SLC9A9* | -0.762309 | 0.26891 | -2.834811 | 0.004585284 | 0.081290884 |
| *ANP32A* | -0.211992 | 0.074787 | -2.834598 | 0.004588338 | 0.081290884 |
| *FUZ* | -0.584742 | 0.206467 | -2.832141 | 0.004623749 | 0.081771696 |
| *RPL7L1* | 0.243354 | 0.085982 | 2.830294 | 0.004650521 | 0.082098298 |
| *GPR65* | 0.614853 | 0.217286 | 2.829695 | 0.004659247 | 0.08210573 |
| *ENSG00000254397* | 1.210117 | 0.428194 | 2.826096 | 0.004711912 | 0.082861453 |
| *C15orf40* | -0.30973 | 0.109628 | -2.825283 | 0.004723889 | 0.082861453 |
| *SLC1A4* | -0.78387 | 0.277471 | -2.825053 | 0.004727277 | 0.082861453 |
| *NUDT2* | -0.567116 | 0.200831 | -2.823853 | 0.004745014 | 0.082877208 |
| *TAF4B* | 0.606491 | 0.214792 | 2.823613 | 0.004748574 | 0.082877208 |
| *MED17* | 0.287497 | 0.101858 | 2.82253 | 0.004764639 | 0.082877208 |
| *FAM184B* | -1.89945 | 0.673259 | -2.821276 | 0.004783296 | 0.082877208 |
| *ABI1* | 0.250594 | 0.088827 | 2.821125 | 0.004785549 | 0.082877208 |
| *SNHG9* | -0.297303 | 0.105385 | -2.821109 | 0.004785789 | 0.082877208 |
| *FABP5* | 0.622852 | 0.220816 | 2.820682 | 0.004792175 | 0.082877208 |
| *SLC2A5* | -0.872799 | 0.309451 | -2.820476 | 0.004795242 | 0.082877208 |
| *ENSG00000256940* | -0.673614 | 0.238961 | -2.818929 | 0.004818411 | 0.083034689 |
| *MRPL11* | -0.210822 | 0.074793 | -2.818747 | 0.004821152 | 0.083034689 |
| *HSPA1B* | 2.218248 | 0.787566 | 2.816586 | 0.004853701 | 0.083389307 |
| *C9orf16* | -0.363381 | 0.12903 | -2.816261 | 0.004858612 | 0.083389307 |
| *FTSJ1* | 0.410095 | 0.14571 | 2.81447 | 0.004885769 | 0.083603917 |
| *FAM53C* | 0.326594 | 0.116047 | 2.814322 | 0.00488803 | 0.083603917 |
| *C1orf52* | 0.272247 | 0.096792 | 2.812706 | 0.00491266 | 0.083768918 |
| *WDTC1* | -0.408472 | 0.145231 | -2.812577 | 0.004914624 | 0.083768918 |
| *NAA50* | 0.201961 | 0.071855 | 2.810678 | 0.004943726 | 0.084030849 |
| *VDAC2* | 0.245703 | 0.087427 | 2.810371 | 0.004948438 | 0.084030849 |
| *GTF3C6* | -0.170288 | 0.060603 | -2.809888 | 0.00495588 | 0.084030849 |
| *TAF1D* | 0.210482 | 0.074922 | 2.809361 | 0.004963991 | 0.084030849 |
| *TM2D3* | 0.40144 | 0.142936 | 2.808535 | 0.004976744 | 0.08410272 |
| *CABLES1* | -1.093528 | 0.389769 | -2.805579 | 0.005022632 | 0.084733339 |
| *VAPA* | 0.166715 | 0.059455 | 2.804043 | 0.005046621 | 0.084953554 |
| *SMC6* | -0.25994 | 0.09272 | -2.803485 | 0.005055362 | 0.084953554 |
| *SRBD1* | -0.568067 | 0.202657 | -2.803096 | 0.005061465 | 0.084953554 |
| *FIP1L1* | 0.238076 | 0.084964 | 2.80208 | 0.005077433 | 0.084960037 |
| *SURF6* | 0.277328 | 0.098989 | 2.801596 | 0.005085051 | 0.084960037 |

| | | | | | |
|---|---|---|---|---|---|
| *BORCS6* | 0.332316 | 0.118631 | 2.801259 | 0.005090358 | 0.084960037 |
| *ZC2HC1A* | 0.92669 | 0.330856 | 2.800888 | 0.005096227 | 0.084960037 |
| *TGIF2* | 0.29133 | 0.104049 | 2.799916 | 0.005111584 | 0.085005136 |
| *CEBPZ* | 0.290611 | 0.103814 | 2.799348 | 0.005120588 | 0.085005136 |
| *ABCF2* | 0.369309 | 0.131956 | 2.798725 | 0.005130486 | 0.085005136 |
| *IL4R* | -0.263518 | 0.094162 | -2.798546 | 0.005133327 | 0.085005136 |
| *MAML3* | -1.367324 | 0.488761 | -2.797529 | 0.005149518 | 0.085130661 |
| *PAXIP1.AS1* | -0.482965 | 0.172772 | -2.79538 | 0.005183869 | 0.085555468 |
| *FRY.AS1* | -1.807199 | 0.64659 | -2.79497 | 0.00519045 | NA |
| *ZEB1.AS1* | -0.84766 | 0.303419 | -2.793696 | 0.005210938 | 0.085825223 |
| *COASY* | -0.321992 | 0.115294 | -2.792804 | 0.005225332 | 0.085825223 |
| *UNC50* | 0.279436 | 0.100058 | 2.792747 | 0.005226258 | 0.085825223 |
| *ZNF24* | 0.240567 | 0.086196 | 2.790929 | 0.005255705 | 0.086165677 |
| *IER5L* | 2.356979 | 0.845089 | 2.78903 | 0.005286622 | NA |
| *ATP13A3* | 0.606864 | 0.217658 | 2.788157 | 0.005300889 | 0.086711475 |
| *ZNRD2* | -0.301882 | 0.108286 | -2.787812 | 0.005306539 | 0.086711475 |
| *C12orf60* | 1.052553 | 0.377742 | 2.786437 | 0.005329104 | 0.0869365 |
| *TVP23A* | 1.230851 | 0.44198 | 2.784854 | 0.005355186 | 0.087123914 |
| *RPL22L1* | 0.471179 | 0.169205 | 2.78467 | 0.005358218 | 0.087123914 |
| *GATM* | 1.714832 | 0.616043 | 2.783622 | 0.005375558 | NA |
| *USP10* | 0.280527 | 0.10085 | 2.781619 | 0.005408849 | 0.087709605 |
| *PAGR1* | -0.536689 | 0.192954 | -2.781431 | 0.005411983 | 0.087709605 |
| *ZNF670* | 0.606432 | 0.218313 | 2.777814 | 0.005472587 | 0.088546639 |
| *RUBCNL* | -0.442257 | 0.159299 | -2.776265 | 0.005498738 | 0.088824389 |
| *CARD8* | -0.468791 | 0.169009 | -2.773757 | 0.00554131 | 0.08936605 |
| *ZNF654* | 0.517065 | 0.186452 | 2.773183 | 0.005551087 | 0.089377917 |
| *PSMA5* | 0.257746 | 0.092989 | 2.771792 | 0.005574864 | 0.089614811 |
| *MAPRE2* | 0.303878 | 0.109848 | 2.766362 | 0.005668562 | 0.090973063 |
| *LRRC8B* | 0.708769 | 0.256552 | 2.762676 | 0.005732971 | 0.091730066 |
| *MCRIP1* | -0.283667 | 0.102702 | -2.76204 | 0.005744153 | 0.091730066 |
| *EGR1* | 1.55872 | 0.564353 | 2.761961 | 0.005745526 | 0.091730066 |
| *ENSG00000233178* | -0.853011 | 0.308904 | -2.761415 | 0.005755145 | 0.091730066 |
| *ZFHX2* | -0.721479 | 0.261309 | -2.761019 | 0.005762125 | 0.091730066 |
| *HSPA9* | 0.455797 | 0.165155 | 2.75981 | 0.005783498 | 0.091922288 |
| *HLA.C* | -0.503593 | 0.182514 | -2.759196 | 0.005794377 | 0.091947369 |
| *CD37* | -0.347918 | 0.126133 | -2.758346 | 0.005809475 | 0.092039212 |
| *MRPL54* | -0.322248 | 0.117015 | -2.753914 | 0.005888726 | 0.09314551 |
| *PDGFA* | 2.062811 | 0.749119 | 2.75365 | 0.005893481 | NA |
| *LY9* | 0.847642 | 0.30791 | 2.752883 | 0.005907294 | 0.093289952 |
| *HERC1* | -0.499666 | 0.181798 | -2.748462 | 0.005987558 | 0.0944067 |
| *KANK2* | -2.111799 | 0.768417 | -2.748245 | 0.005991529 | NA |
| *SLC38A2* | 0.197532 | 0.072003 | 2.743368 | 0.006081255 | 0.09573135 |
| *RPP21* | -0.44084 | 0.160802 | -2.741519 | 0.006115575 | 0.096118562 |
| *ILF2* | 0.235994 | 0.086168 | 2.738787 | 0.006166627 | 0.096728316 |
| *ARL14EPL* | 1.328684 | 0.485231 | 2.73825 | 0.006176703 | 0.096728316 |
| *ACSL4* | 0.566347 | 0.206968 | 2.736399 | 0.006211559 | 0.096728316 |
| *TMEM64* | 0.582235 | 0.212775 | 2.736388 | 0.006211782 | 0.096728316 |

| | | | | | |
|---|---|---|---|---|---|
| *SQSTM1* | 0.320471 | 0.117123 | 2.73619 | 0.006215521 | 0.096728316 |
| *CWC22* | 0.249239 | 0.091093 | 2.736101 | 0.006217202 | 0.096728316 |
| *SRD5A1* | -0.519806 | 0.190001 | -2.735801 | 0.006222862 | 0.096728316 |
| *TMEM256* | -0.303465 | 0.111172 | -2.729698 | 0.006339243 | 0.098382656 |
| *TVP23B* | 0.464608 | 0.170243 | 2.729082 | 0.006351092 | 0.098412065 |
| *RMND5B* | -0.310795 | 0.114002 | -2.726221 | 0.006406412 | 0.099062864 |
| *GRPEL1* | 0.431148 | 0.158208 | 2.725205 | 0.006426164 | 0.099062864 |
| *GRHPR* | -0.659806 | 0.242122 | -2.725099 | 0.006428229 | 0.099062864 |
| *SPIN2B* | 0.685981 | 0.251751 | 2.724845 | 0.006433174 | 0.099062864 |
| *MID1IP1* | -0.56089 | 0.205945 | -2.723491 | 0.006459604 | 0.099315148 |
| *MRPL32* | 0.25943 | 0.095313 | 2.721858 | 0.006491602 | 0.099530729 |
| *TOP1* | 0.277788 | 0.102062 | 2.721748 | 0.006493761 | 0.099530729 |
| *NOCT* | -0.982614 | 0.361301 | -2.719654 | 0.006535027 | NA |
| *POMGNT1* | -0.403796 | 0.148548 | -2.718287 | 0.006562094 | 0.100422389 |
| *ANKRD46* | 0.540382 | 0.198894 | 2.716941 | 0.006588836 | 0.1005846 |
| *POLH* | 0.543295 | 0.199993 | 2.71657 | 0.006596217 | 0.1005846 |
| *ARHGAP18* | 0.998862 | 0.36774 | 2.716219 | 0.006603217 | 0.1005846 |
| *MBNL2* | -0.633747 | 0.233429 | -2.71495 | 0.006628577 | 0.100815561 |
| *MBTPS1* | -0.38271 | 0.140994 | -2.714365 | 0.006640303 | 0.100838761 |
| *ATG101* | 0.240928 | 0.08884 | 2.711943 | 0.006689014 | 0.101422691 |
| *TIMMDC1* | 0.291736 | 0.10767 | 2.709554 | 0.006737378 | 0.101999566 |
| *PSENEN* | -0.247882 | 0.091626 | -2.70537 | 0.006822837 | 0.103085183 |
| *VOPP1* | -0.282891 | 0.10458 | -2.705024 | 0.006829941 | 0.103085183 |
| *PSMD9* | -0.247665 | 0.091598 | -2.703823 | 0.006854675 | 0.103300789 |
| *HCG11* | -0.568683 | 0.210428 | -2.702502 | 0.006881986 | 0.103554506 |
| *AEBP2* | 0.383502 | 0.142063 | 2.699519 | 0.00694397 | 0.104328401 |
| *EIF5B* | 0.278584 | 0.103265 | 2.697764 | 0.006980695 | 0.104563683 |
| *METTL8* | 0.507618 | 0.188193 | 2.697324 | 0.006989921 | 0.104563683 |
| *TENT5A* | 0.553342 | 0.205178 | 2.69689 | 0.00699904 | 0.104563683 |
| *HIST1H2BJ* | 0.647225 | 0.240054 | 2.69617 | 0.007014177 | 0.104563683 |
| *ZNF441* | -0.614422 | 0.227919 | -2.695786 | 0.007022278 | 0.104563683 |
| *CYTOR* | 1.089933 | 0.404316 | 2.695747 | 0.007023092 | 0.104563683 |
| *ENSG00000267633* | -1.785993 | 0.662784 | -2.694683 | 0.007045571 | NA |
| *INSIG2* | 0.687706 | 0.255297 | 2.693748 | 0.007065351 | 0.105034675 |
| *YWHAG* | 0.317044 | 0.117806 | 2.69125 | 0.00711848 | 0.105665604 |
| *CBX6* | -0.276887 | 0.10298 | -2.688753 | 0.007171953 | 0.106299739 |
| *CERS4* | -0.317425 | 0.11817 | -2.686181 | 0.007227394 | 0.106961099 |
| *OPTN* | -0.386366 | 0.143912 | -2.684741 | 0.007258613 | 0.107262549 |
| *KCTD5* | 0.369916 | 0.137917 | 2.682156 | 0.007314942 | 0.107798354 |
| *CD3EAP* | 0.624532 | 0.232854 | 2.682076 | 0.00731668 | 0.107798354 |
| *SPAG4* | 0.929045 | 0.346698 | 2.679695 | 0.007368926 | 0.108277314 |
| *PPP4R2* | 0.277838 | 0.103688 | 2.679573 | 0.007371608 | 0.108277314 |
| *RIOK3* | 0.279736 | 0.104414 | 2.679099 | 0.007382046 | 0.108277314 |
| *PPA2* | 0.429166 | 0.160259 | 2.677944 | 0.007407559 | 0.108490567 |
| *TMEM216* | 0.301797 | 0.112741 | 2.676898 | 0.007430718 | 0.108534582 |
| *BLOC1S2* | -0.253664 | 0.094763 | -2.676817 | 0.007432522 | 0.108534582 |
| *CPNE3* | -0.338336 | 0.126463 | -2.675381 | 0.007464428 | 0.108839725 |

| | | | | | |
|---|---|---|---|---|---|
| *TMEM141* | -0.310514 | 0.116113 | -2.674244 | 0.007489797 | 0.109042941 |
| *ZBED2* | 1.468844 | 0.549534 | 2.672891 | 0.007520064 | 0.109042941 |
| *ENSG00000253982* | -0.473082 | 0.176997 | -2.672827 | 0.007521495 | 0.109042941 |
| *DDX31* | 0.568723 | 0.212838 | 2.672096 | 0.007537898 | 0.109042941 |
| *UTP18* | 0.31668 | 0.118538 | 2.671546 | 0.007550275 | 0.109042941 |
| *HDX* | 1.688083 | 0.631898 | 2.671448 | 0.007552487 | NA |
| *LARP1* | 0.332754 | 0.124583 | 2.67093 | 0.007564139 | 0.109042941 |
| *DDB2* | -0.254943 | 0.095457 | -2.670768 | 0.007567783 | 0.109042941 |
| *MED22* | -0.392309 | 0.146905 | -2.670497 | 0.007573894 | 0.109042941 |
| *NOP2* | 0.384951 | 0.144159 | 2.670332 | 0.007577635 | 0.109042941 |
| *DHRS7B* | -0.329447 | 0.12351 | -2.667378 | 0.007644557 | 0.109846056 |
| *FAM41C* | -0.472334 | 0.177173 | -2.665944 | 0.007677257 | 0.110155825 |
| *PEX12* | 1.120397 | 0.42057 | 2.663998 | 0.007721803 | 0.110634413 |
| *CCNL1* | 0.244577 | 0.091876 | 2.662044 | 0.007766772 | 0.111117668 |
| *CCL22* | 2.092216 | 0.786351 | 2.660664 | 0.007798671 | 0.111412804 |
| *PSMG3* | -0.319581 | 0.120139 | -2.660089 | 0.007812 | 0.111442188 |
| *ENSG00000260793* | -0.84039 | 0.316158 | -2.658135 | 0.007857443 | 0.11192893 |
| *TRIM27* | 0.24935 | 0.093827 | 2.657559 | 0.007870878 | 0.111958993 |
| *TRMT1L* | 0.484249 | 0.182619 | 2.651694 | 0.008008911 | 0.113758752 |
| *UCK2* | 0.589643 | 0.222824 | 2.64622 | 0.008139694 | 0.115450521 |
| *RHOA* | 0.184031 | 0.069574 | 2.645104 | 0.008166573 | 0.115647139 |
| *NBDY* | -0.371228 | 0.14037 | -2.644636 | 0.008177878 | 0.115647139 |
| *NOM1* | 0.335832 | 0.127008 | 2.644191 | 0.00818865 | 0.115647139 |
| *DHRSX* | -0.405468 | 0.153548 | -2.640664 | 0.008274383 | 0.116691233 |
| *PKIG* | -0.537533 | 0.203668 | -2.639269 | 0.008308509 | 0.116916338 |
| *RBM38* | -0.322821 | 0.122326 | -2.639011 | 0.008314835 | 0.116916338 |
| *PPP2R5C* | -0.246451 | 0.093408 | -2.638423 | 0.008329267 | 0.116916338 |
| *CENPM* | 0.616754 | 0.233789 | 2.638082 | 0.008337651 | 0.116916338 |
| *EIF4A2* | 0.202075 | 0.076672 | 2.635585 | 0.008399245 | 0.117613224 |
| *L3HYPDH* | -0.766639 | 0.29099 | -2.634587 | 0.008423979 | 0.11764915 |
| *RUNX3* | 0.249165 | 0.094577 | 2.634521 | 0.008425612 | 0.11764915 |
| *CMC2* | 0.252355 | 0.09581 | 2.633915 | 0.008440673 | 0.117693216 |
| *PEX10* | -0.592394 | 0.224961 | -2.633317 | 0.008455531 | 0.117734337 |
| *CPNE2* | -1.888523 | 0.718534 | -2.6283 | 0.008581275 | NA |
| *ZNF302* | -0.263681 | 0.100335 | -2.627999 | 0.008588872 | 0.119422772 |
| *HRK* | -1.441138 | 0.548651 | -2.626696 | 0.008621843 | 0.11971284 |
| *LPP.AS2* | -1.387343 | 0.528364 | -2.625733 | 0.008646272 | NA |
| *VPS25* | -0.283334 | 0.107923 | -2.625328 | 0.008656554 | 0.12002622 |
| *NPM1* | 0.326466 | 0.124439 | 2.623507 | 0.008702958 | 0.120500615 |
| *PSD3* | 1.435757 | 0.547332 | 2.623192 | 0.008711017 | NA |
| *ASB1* | 0.355522 | 0.135579 | 2.622244 | 0.008735282 | 0.120693255 |
| *DOCK10* | -0.447107 | 0.170521 | -2.62201 | 0.008741288 | 0.120693255 |
| *TGM2* | 2.23817 | 0.853756 | 2.621558 | 0.008752879 | NA |
| *HEATR1* | 0.418826 | 0.159802 | 2.620912 | 0.008769479 | 0.120758776 |
| *HGSNAT* | -0.49343 | 0.188269 | -2.620874 | 0.008770463 | 0.120758776 |
| *YBEY* | -0.294974 | 0.112696 | -2.617436 | 0.008859318 | 0.121671211 |
| *RRAGC* | 0.268095 | 0.102435 | 2.617219 | 0.008864936 | 0.121671211 |

| | | | | | |
|---|---|---|---|---|---|
| *FTL* | 0.252897 | 0.096641 | 2.616884 | 0.008873654 | 0.121671211 |
| *WDR76* | -0.569838 | 0.217828 | -2.615997 | 0.008896741 | 0.12177956 |
| *TFEB* | 0.350023 | 0.133819 | 2.615634 | 0.008906193 | 0.12177956 |
| *LINC02166* | 1.259314 | 0.481775 | 2.613905 | 0.008951389 | NA |
| *ZNF692* | -0.4675 | 0.178909 | -2.61306 | 0.008973542 | 0.122530994 |
| *PIGF* | 0.639347 | 0.244903 | 2.610619 | 0.009037863 | 0.123239053 |
| *ENSG00000224746* | -1.375713 | 0.527094 | -2.609996 | 0.009054321 | NA |
| *PET100* | -0.234241 | 0.089782 | -2.609011 | 0.009080431 | 0.123648959 |
| *ENSG00000237499* | -0.905875 | 0.347275 | -2.608521 | 0.009093451 | 0.123655922 |
| *MGAT5B* | -1.124268 | 0.431137 | -2.607684 | 0.009115698 | NA |
| *TUT4* | 0.256567 | 0.098448 | 2.60611 | 0.009157698 | 0.124358521 |
| *OGFRL1* | -0.46376 | 0.178085 | -2.604153 | 0.00921017 | 0.124899506 |
| *KIAA1586* | -0.519872 | 0.199735 | -2.602815 | 0.00924619 | 0.125162789 |
| *ELF2* | -0.227534 | 0.087429 | -2.602492 | 0.009254906 | 0.125162789 |
| *HLA.A* | -0.452282 | 0.173929 | -2.600387 | 0.009311858 | 0.125723231 |
| *EMC2* | 0.372337 | 0.14323 | 2.599576 | 0.009333909 | 0.125723231 |
| *FUOM* | -0.492584 | 0.189488 | -2.599554 | 0.009334498 | 0.125723231 |
| *RNASEH2B* | 0.33054 | 0.127262 | 2.597326 | 0.009395284 | 0.126369768 |
| *SELENOF* | -0.13418 | 0.05168 | -2.596386 | 0.009421011 | 0.126486822 |
| *TICAM1* | 0.427995 | 0.164884 | 2.595737 | 0.00943884 | 0.126486822 |
| *MRPL43* | -0.191647 | 0.073842 | -2.59537 | 0.009448921 | 0.126486822 |
| *THRAP3* | -0.223144 | 0.085985 | -2.595143 | 0.009455165 | 0.126486822 |
| *GNG3* | -2.363786 | 0.910903 | -2.594992 | 0.009459308 | NA |
| *CTNNBIP1* | -0.89522 | 0.345223 | -2.593164 | 0.00950975 | 0.12704512 |
| *VMP1* | 0.226869 | 0.087529 | 2.591934 | 0.009543801 | 0.127327961 |
| *PBX4* | 1.170384 | 0.451642 | 2.591395 | 0.009558769 | 0.127355788 |
| *ENSG00000279927* | -1.240121 | 0.478555 | -2.591388 | 0.00955896 | NA |
| *NR4A2* | 1.254824 | 0.484443 | 2.590241 | 0.009590874 | 0.127434772 |
| *NBPF9* | 0.855254 | 0.330233 | 2.589848 | 0.009601834 | 0.127434772 |
| *GPS1* | -0.362204 | 0.139858 | -2.589793 | 0.009603369 | 0.127434772 |
| *USP14* | 0.299088 | 0.11551 | 2.589268 | 0.009618018 | 0.127458084 |
| *TOB1* | 0.524218 | 0.202564 | 2.587914 | 0.009655898 | 0.127788766 |
| *TUNAR* | -1.864856 | 0.720847 | -2.587035 | 0.00968058 | NA |
| *CCDC159* | -0.563557 | 0.217903 | -2.586278 | 0.009701857 | 0.128109597 |
| *ZNF296* | 0.5271 | 0.203818 | 2.586129 | 0.009706058 | 0.128109597 |
| *ADNP2* | 0.736761 | 0.28508 | 2.5844 | 0.009754866 | 0.128582143 |
| *MCOLN2* | 1.290717 | 0.499698 | 2.582994 | 0.009794713 | 0.128935469 |
| *G2E3* | 0.404072 | 0.156533 | 2.581387 | 0.009840424 | 0.129364942 |
| *DEGS1* | 0.305943 | 0.118555 | 2.580599 | 0.009862897 | 0.129488175 |
| *NUDT16* | -0.486751 | 0.188655 | -2.580112 | 0.009876829 | 0.129499116 |
| *ENSG00000260879* | -1.66763 | 0.646537 | -2.579327 | 0.009899289 | NA |
| *KDM5B* | 0.423766 | 0.164391 | 2.577796 | 0.009943256 | 0.130197383 |
| *HLA.DMB* | -0.453577 | 0.176052 | -2.576377 | 0.009984178 | 0.130560292 |
| *TOX2* | 2.387925 | 0.926855 | 2.576375 | 0.009984226 | NA |
| *CD99* | -0.239714 | 0.093123 | -2.574152 | 0.010048626 | 0.131128541 |
| *ENSG00000272195* | -1.597807 | 0.620729 | -2.574083 | 0.010050623 | NA |
| *ZNF827* | 0.750003 | 0.291381 | 2.573961 | 0.010054161 | 0.131128541 |

| | | | | | |
|---|---|---|---|---|---|
| *XPO6* | 0.37905 | 0.14735 | 2.572448 | 0.0100982 | 0.131508112 |
| *MIR4435.2HG* | 1.700101 | 0.661031 | 2.571892 | 0.010114431 | 0.131508112 |
| *TCTN3* | 0.356638 | 0.138684 | 2.571593 | 0.010123171 | 0.131508112 |
| *FBXO25* | -0.283036 | 0.110155 | -2.569437 | 0.010186387 | 0.13215567 |
| *MIR155HG* | 0.543112 | 0.21153 | 2.567543 | 0.010242216 | 0.132622582 |
| *GPX1* | -0.465096 | 0.181171 | -2.567164 | 0.010253419 | 0.132622582 |
| *CHRNE* | 1.572247 | 0.612545 | 2.566747 | 0.010265743 | NA |
| *TFDP2* | -0.50127 | 0.195311 | -2.566525 | 0.010272319 | 0.132622582 |
| *ZBTB20* | 0.469308 | 0.182893 | 2.566031 | 0.010286977 | 0.132622582 |
| *LACTB2* | 0.421564 | 0.164297 | 2.565858 | 0.010292091 | 0.132622582 |
| *KLK1* | 1.702505 | 0.663617 | 2.565495 | 0.010302867 | 0.132622582 |
| *ENSG00000272155* | 1.311911 | 0.511385 | 2.565407 | 0.01030547 | NA |
| *MICOS13* | -0.180787 | 0.070482 | -2.564991 | 0.010317855 | 0.132642804 |
| *TSEN15* | -0.462504 | 0.180418 | -2.563519 | 0.010361712 | 0.133033613 |
| *TMEM220.AS1* | -1.51187 | 0.590302 | -2.56118 | 0.010431737 | NA |
| *TAGAP* | -0.548219 | 0.214084 | -2.560764 | 0.010444238 | 0.133919246 |
| *DLAT* | -0.409439 | 0.160151 | -2.556585 | 0.010570539 | 0.135363149 |
| *MCRS1* | -0.259547 | 0.101666 | -2.552933 | 0.010682014 | 0.136613702 |
| *PPBP* | 3.467389 | 1.358769 | 2.551862 | 0.010714909 | NA |
| *MGME1* | 0.394043 | 0.154432 | 2.55157 | 0.01072387 | 0.136971806 |
| *FAM228B* | -0.471274 | 0.184807 | -2.550089 | 0.010769531 | 0.137377526 |
| *HM13* | 0.281178 | 0.110295 | 2.549323 | 0.010793232 | 0.13744886 |
| *JAM3* | 0.737115 | 0.289177 | 2.54901 | 0.01080293 | 0.13744886 |
| *IGHM* | -0.401936 | 0.157735 | -2.548176 | 0.010828786 | 0.137451266 |
| *RBMS1* | -0.442909 | 0.173821 | -2.548072 | 0.010832014 | 0.137451266 |
| *BRD2* | 0.157584 | 0.061854 | 2.547659 | 0.01084483 | 0.137451266 |
| *HHEX* | 0.359748 | 0.141257 | 2.54676 | 0.010872818 | 0.137629551 |
| *ENSG00000261335* | -2.046899 | 0.804467 | -2.544415 | 0.010946084 | NA |
| *MIR22HG* | 0.526243 | 0.206864 | 2.543908 | 0.010961993 | 0.1385809 |
| *HIST1H2BH* | 1.582929 | 0.622708 | 2.542007 | 0.01102179 | NA |
| *LINC01857* | 0.499478 | 0.196504 | 2.541822 | 0.011027621 | 0.139232524 |
| *RCSD1* | -0.527754 | 0.207812 | -2.539573 | 0.01109879 | 0.139389701 |
| *ARHGAP1* | -0.458251 | 0.180445 | -2.539566 | 0.011099006 | 0.139389701 |
| *STAM2* | 0.624845 | 0.246066 | 2.539341 | 0.011106159 | 0.139389701 |
| *PDCD5* | 0.290694 | 0.114477 | 2.539318 | 0.011106875 | 0.139389701 |
| *SGF29* | 0.386486 | 0.152238 | 2.538698 | 0.011126581 | 0.139389701 |
| *BIN3* | -0.377949 | 0.148876 | -2.53869 | 0.011126823 | 0.139389701 |
| *XIAP* | 0.311603 | 0.122771 | 2.53809 | 0.011145938 | 0.139389701 |
| *ZNF74* | 1.548173 | 0.610006 | 2.537963 | 0.011149971 | NA |
| *ATP6V0E2* | -1.026929 | 0.404642 | -2.537872 | 0.011152868 | 0.139389701 |
| *LIMS1* | 0.362431 | 0.142978 | 2.534872 | 0.011248827 | 0.140342673 |
| *ABHD17B* | 0.372709 | 0.147048 | 2.534602 | 0.01125751 | 0.140342673 |
| *TOP2A* | -1.11912 | 0.441611 | -2.534174 | 0.011271281 | NA |
| *DIPK1B* | -1.949302 | 0.769575 | -2.53296 | 0.011310372 | NA |
| *GSDMD* | -0.460588 | 0.181851 | -2.532782 | 0.011316133 | 0.140895833 |
| *TMEM205* | -0.260204 | 0.102811 | -2.530901 | 0.01137701 | 0.141475622 |
| *PYHIN1* | 1.142996 | 0.452152 | 2.5279 | 0.011474689 | NA |

| | | | | | |
|---|---|---|---|---|---|
| *CD68* | 0.809709 | 0.320581 | 2.525751 | 0.011545141 | 0.143231586 |
| *PSMD6* | 0.26107 | 0.103366 | 2.525688 | 0.011547195 | 0.143231586 |
| *RNF114* | 0.226575 | 0.08973 | 2.525069 | 0.011567555 | 0.143304323 |
| *HVCN1* | -0.513658 | 0.203563 | -2.523336 | 0.011624733 | 0.143749607 |
| *ITGB3BP* | -0.610933 | 0.242147 | -2.522983 | 0.011636393 | 0.143749607 |
| *PITPNB* | 0.259126 | 0.102719 | 2.522659 | 0.011647121 | 0.143749607 |
| *CTSC* | 0.710801 | 0.28204 | 2.520216 | 0.011728298 | 0.14455658 |
| *UTP14A* | 0.377726 | 0.149902 | 2.519812 | 0.011741749 | 0.14455658 |
| *KCNRG* | 1.415555 | 0.561848 | 2.519463 | 0.011753409 | NA |
| *ATP6V0D1* | 0.26645 | 0.105766 | 2.519248 | 0.011760563 | 0.144608114 |
| *NUDT1* | -0.354408 | 0.140708 | -2.518746 | 0.011777343 | 0.144634548 |
| *RAD51AP1* | -1.210683 | 0.481144 | -2.516261 | 0.011860719 | NA |
| *TMC8* | -0.412218 | 0.163857 | -2.51572 | 0.011878944 | 0.145491569 |
| *METAP2* | 0.192302 | 0.076444 | 2.515579 | 0.011883706 | 0.145491569 |
| *JADE1* | 0.397977 | 0.158231 | 2.51517 | 0.011897512 | 0.145491569 |
| *PRDM8* | -0.515733 | 0.205069 | -2.514918 | 0.011905996 | 0.145491569 |
| *SQOR* | -0.394754 | 0.157008 | -2.514221 | 0.011929582 | 0.145599806 |
| *SLC35D1* | 0.745652 | 0.296982 | 2.510761 | 0.012047122 | 0.146640929 |
| *ZNF207* | 0.189697 | 0.075561 | 2.510511 | 0.012055664 | 0.146640929 |
| *ECH1* | -0.230842 | 0.091954 | -2.510402 | 0.012059385 | 0.146640929 |
| *TPCN1* | -0.50691 | 0.202001 | -2.509442 | 0.012092201 | 0.146859334 |
| *ADA* | -0.9045 | 0.360721 | -2.50748 | 0.012159561 | 0.147360194 |
| *NETO2* | 1.013812 | 0.40435 | 2.50726 | 0.012167131 | 0.147360194 |
| *CD79B* | -0.383823 | 0.153112 | -2.506814 | 0.012182491 | 0.147360194 |
| *UBA2* | 0.189006 | 0.075406 | 2.506507 | 0.012193065 | 0.147360194 |
| *RGS2* | 0.868266 | 0.34653 | 2.505604 | 0.012224257 | 0.147404178 |
| *IFI30* | 0.441351 | 0.17615 | 2.505538 | 0.012226525 | 0.147404178 |
| *TERF1* | 0.228977 | 0.091483 | 2.502938 | 0.012316717 | 0.148019196 |
| *TCEAL3* | -0.415313 | 0.165936 | -2.502855 | 0.012319616 | 0.148019196 |
| *C7orf50* | -0.397556 | 0.158846 | -2.502773 | 0.012322456 | 0.148019196 |
| *MFSD6* | 0.82442 | 0.32969 | 2.500595 | 0.01239849 | 0.148751789 |
| *CNKSR3* | -1.554084 | 0.621947 | -2.498741 | 0.012463547 | NA |
| *MAD2L1BP* | 0.303341 | 0.12145 | 2.497655 | 0.012501774 | 0.149809136 |
| *DGAT2* | 0.562929 | 0.225589 | 2.495373 | 0.01258248 | 0.150567651 |
| *HEXIM2* | 0.529806 | 0.212347 | 2.495005 | 0.012595534 | 0.150567651 |
| *ORAI3* | -0.339965 | 0.136314 | -2.493984 | 0.012631842 | 0.150819306 |
| *PEG10* | 1.401822 | 0.562252 | 2.493225 | 0.012658871 | NA |
| *ASCL2* | 1.242845 | 0.498822 | 2.491559 | 0.012718386 | NA |
| *ENSG00000279529* | -0.571196 | 0.229375 | -2.49023 | 0.012766046 | 0.15215436 |
| *EVI2B* | -0.235565 | 0.094604 | -2.489996 | 0.01277444 | 0.15215436 |
| *JRK* | -0.609964 | 0.245135 | -2.488272 | 0.012836534 | 0.152709962 |
| *EMC10* | -0.235438 | 0.094666 | -2.487039 | 0.012881146 | 0.153056498 |
| *NDUFA1* | -0.157091 | 0.063192 | -2.485919 | 0.012921738 | 0.153354509 |
| *CIRBP* | -0.176718 | 0.071106 | -2.485286 | 0.012944738 | 0.153443267 |
| *NT5C2* | -0.485568 | 0.195413 | -2.484832 | 0.012961245 | 0.153454934 |
| *OAZ2* | -0.221656 | 0.089229 | -2.484128 | 0.012986906 | 0.153574825 |
| *NDUFS8* | -0.25388 | 0.102308 | -2.481521 | 0.013082293 | 0.154211537 |

| | | | | | |
|---|---|---|---|---|---|
| *ANAPC16* | -0.149422 | 0.060216 | -2.481444 | 0.013085129 | 0.154211537 |
| *PELI1* | 0.365525 | 0.147307 | 2.481378 | 0.013087546 | 0.154211537 |
| *TTC39B* | -1.653687 | 0.667369 | -2.477921 | 0.013215024 | NA |
| *ENSG00000237950* | -1.626822 | 0.656752 | -2.477071 | 0.013246551 | NA |
| *SERPINB9* | 0.359955 | 0.14537 | 2.476133 | 0.013281406 | 0.156186709 |
| *CDCA7L* | -0.42427 | 0.171354 | -2.475989 | 0.013286771 | 0.156186709 |
| *ALKBH6* | -0.646057 | 0.261082 | -2.474538 | 0.013340863 | 0.156636308 |
| *CHP1* | -0.298569 | 0.12071 | -2.473433 | 0.013382178 | 0.156731376 |
| *ZC3H12D* | -0.481398 | 0.194653 | -2.473114 | 0.013394137 | 0.156731376 |
| *TOMM5* | -0.235099 | 0.095064 | -2.47305 | 0.013396522 | 0.156731376 |
| *TNFRSF13B* | 0.992687 | 0.40152 | 2.472325 | 0.013423757 | 0.15686437 |
| *GPSM3* | -0.221702 | 0.089724 | -2.470941 | 0.013475817 | 0.157203795 |
| *C1orf162* | -0.410279 | 0.166057 | -2.470707 | 0.013484606 | 0.157203795 |
| *WARS* | 0.740969 | 0.300256 | 2.467786 | 0.013595145 | 0.15830578 |
| *DHRS3* | 2.089275 | 0.846891 | 2.466995 | 0.013625234 | 0.158469482 |
| *SH3BGRL* | -0.336224 | 0.136313 | -2.466565 | 0.013641603 | 0.15847343 |
| *RSL24D1* | 0.191742 | 0.077756 | 2.465954 | 0.013664875 | 0.158485212 |
| *ACAT1* | -0.540452 | 0.219188 | -2.465697 | 0.01367468 | 0.158485212 |
| *SELENOK* | 0.208901 | 0.084828 | 2.462635 | 0.013792014 | 0.159657904 |
| *DDIT4* | 0.440672 | 0.178993 | 2.461949 | 0.013818419 | 0.159663473 |
| *AFDN* | 0.879367 | 0.357246 | 2.461519 | 0.013835024 | 0.159663473 |
| *ACP2* | -0.714483 | 0.290342 | -2.460832 | 0.013861513 | 0.159663473 |
| *SLBP* | 0.438191 | 0.178072 | 2.460754 | 0.013864541 | 0.159663473 |
| *L1CAM* | -0.772048 | 0.313773 | -2.460529 | 0.013873247 | 0.159663473 |
| *TUBB4B* | 0.397162 | 0.161487 | 2.459406 | 0.013916696 | 0.159977278 |
| *SSH2* | -0.292756 | 0.119089 | -2.458285 | 0.013960251 | 0.16016657 |
| *NFKBID* | 0.252663 | 0.102786 | 2.458148 | 0.013965566 | 0.16016657 |
| *ENSG00000265218* | -1.178396 | 0.479506 | -2.45752 | 0.01399001 | NA |
| *HK2* | 0.8673 | 0.353068 | 2.456467 | 0.014031067 | 0.160576103 |
| *RABGGTB* | 0.233927 | 0.095232 | 2.456398 | 0.01403376 | 0.160576103 |
| *IQCG* | -0.630361 | 0.256685 | -2.455775 | 0.0140581 | 0.160668646 |
| *KIF21A* | 1.301007 | 0.529847 | 2.455438 | 0.014071319 | NA |
| *SPINT2* | -0.308985 | 0.12596 | -2.453036 | 0.01416562 | 0.161310725 |
| *POLR2B* | -0.26232 | 0.10694 | -2.452977 | 0.014167945 | 0.161310725 |
| *TP53TG1* | -0.393333 | 0.160372 | -2.452632 | 0.014181526 | 0.161310725 |
| *SUB1* | 0.2079 | 0.084774 | 2.452415 | 0.014190105 | 0.161310725 |
| *CCDC191* | -0.669602 | 0.273065 | -2.452174 | 0.014199602 | 0.161310725 |
| *RGL1* | -1.377406 | 0.561763 | -2.451934 | 0.014209077 | NA |
| *DCAF11* | -0.266933 | 0.10887 | -2.451855 | 0.014212183 | 0.161310725 |
| *MKRN2* | 0.383121 | 0.156334 | 2.450665 | 0.014259254 | 0.161659388 |
| *GPR146* | -0.764396 | 0.312029 | -2.449756 | 0.014295294 | 0.161882328 |
| *CIITA* | 0.674999 | 0.275612 | 2.449087 | 0.014321869 | 0.161913742 |
| *BABAM1* | -0.257303 | 0.105071 | -2.448862 | 0.014330824 | 0.161913742 |
| *NOL6* | 0.696388 | 0.284472 | 2.448005 | 0.014364983 | 0.162114404 |
| *LMF2* | -0.301854 | 0.123413 | -2.445889 | 0.014449564 | 0.162882995 |
| *SLC16A1.AS1* | 0.346484 | 0.141841 | 2.442768 | 0.014575117 | 0.164111171 |
| *PTPN12* | 0.519693 | 0.212793 | 2.442248 | 0.014596106 | 0.164160527 |

| | | | | | |
|---|---|---|---|---|---|
| *XRN2* | 0.248282 | 0.101749 | 2.440141 | 0.014681519 | 0.164933523 |
| *B9D2* | -0.343694 | 0.140895 | -2.439366 | 0.014713055 | 0.165100181 |
| *C6orf226* | -0.582658 | 0.238931 | -2.438606 | 0.01474405 | 0.16526041 |
| *HDAC6* | -0.362506 | 0.148699 | -2.43785 | 0.014774895 | 0.165418583 |
| *ST13* | 0.143587 | 0.058912 | 2.437321 | 0.014796543 | 0.165473553 |
| *UBAC1* | -0.506203 | 0.207825 | -2.435716 | 0.014862344 | 0.166021623 |
| *ADD1* | -0.224485 | 0.092211 | -2.43446 | 0.01491404 | 0.166411058 |
| *CASP10* | 0.592484 | 0.243694 | 2.43126 | 0.015046397 | 0.167698621 |
| *KLHDC10* | -0.547345 | 0.225214 | -2.430334 | 0.015084906 | 0.167938486 |
| *TIAL1* | 0.202117 | 0.083202 | 2.429248 | 0.015130195 | 0.168253213 |
| *RHOBTB2* | 1.519136 | 0.62557 | 2.428404 | 0.015165452 | NA |
| *PRPF19* | 0.285078 | 0.117394 | 2.42838 | 0.015166453 | 0.168466911 |
| *ENSG00000278158* | -0.376959 | 0.155295 | -2.427367 | 0.015208843 | 0.168562717 |
| *EXOSC3* | 0.224857 | 0.092634 | 2.427359 | 0.015209179 | 0.168562717 |
| *METRNL* | 1.006883 | 0.414913 | 2.426733 | 0.015235484 | 0.16866517 |
| *ALDH4A1* | -1.296345 | 0.53459 | -2.424935 | 0.015311149 | NA |
| *FCRL2* | 0.521189 | 0.214958 | 2.424605 | 0.015325061 | 0.169467059 |
| *PTPN20* | -1.406811 | 0.580374 | -2.423974 | 0.015351686 | NA |
| *PLEKHO2* | 0.511239 | 0.210922 | 2.423829 | 0.015357841 | 0.16963979 |
| *ZNF165* | 0.497921 | 0.205484 | 2.423163 | 0.015386024 | 0.169639955 |
| *ZNF213.AS1* | -0.616592 | 0.254473 | -2.423018 | 0.015392175 | 0.169639955 |
| *SLC25A36* | 0.360501 | 0.148862 | 2.421703 | 0.015447988 | 0.170065488 |
| *IDI1* | 0.332979 | 0.137595 | 2.41999 | 0.015520928 | 0.17067841 |
| *ZKSCAN7* | -1.727758 | 0.714859 | -2.416922 | 0.015652355 | NA |
| *TRMT2B* | -0.431399 | 0.178497 | -2.41684 | 0.015655908 | 0.171751961 |
| *PRKAR1B* | 0.714768 | 0.295782 | 2.41654 | 0.015668804 | 0.171751961 |
| *ZNF28* | -0.471522 | 0.195181 | -2.415817 | 0.01569995 | 0.171751961 |
| *WDR47* | 0.65126 | 0.269587 | 2.415769 | 0.015702001 | 0.171751961 |
| *DLGAP1.AS1* | -0.410454 | 0.169912 | -2.41569 | 0.015705419 | 0.171751961 |
| *SRGAP2* | 0.653255 | 0.27065 | 2.413654 | 0.015793439 | 0.172523691 |
| *MMUT* | 0.577441 | 0.239399 | 2.412044 | 0.01586335 | 0.173096107 |
| *ACSL1* | 0.504578 | 0.209256 | 2.411297 | 0.015895912 | 0.173260186 |
| *MKKS* | -0.287554 | 0.119296 | -2.410423 | 0.015934039 | 0.173484485 |
| *ADAM15* | 0.724463 | 0.300737 | 2.40896 | 0.015998057 | 0.173989868 |
| *RSRC2* | 0.35086 | 0.14571 | 2.407931 | 0.016043194 | 0.174145601 |
| *UQCRQ* | -0.179256 | 0.074447 | -2.407831 | 0.016047607 | 0.174145601 |
| *SLC39A13* | -0.514898 | 0.213923 | -2.406931 | 0.016087189 | 0.17429812 |
| *PSMD14* | 0.336576 | 0.139849 | 2.406711 | 0.016096923 | 0.17429812 |
| *ENSG00000259529* | -0.472326 | 0.196298 | -2.406168 | 0.016120867 | 0.174366398 |
| *CYP2R1* | -0.451009 | 0.187481 | -2.405624 | 0.016144872 | 0.174378088 |
| *ENSA* | -0.128826 | 0.053558 | -2.405345 | 0.016157225 | 0.174378088 |
| *ATP2B1* | -0.255261 | 0.106156 | -2.404592 | 0.016190549 | 0.174547183 |
| *CEP164* | -0.393797 | 0.16382 | -2.403835 | 0.016224115 | 0.174718519 |
| *PAK1IP1* | 0.446433 | 0.185755 | 2.403338 | 0.016246177 | 0.174765726 |
| *SETBP1* | 0.919371 | 0.382878 | 2.401208 | 0.016341023 | 0.17559495 |
| *THG1L* | 0.518422 | 0.215951 | 2.400642 | 0.016366329 | 0.175675927 |
| *NDUFA4* | -0.142319 | 0.059299 | -2.400003 | 0.016394921 | 0.17572204 |

| | | | | | |
|---|---|---|---|---|---|
| *VTI1A* | -0.311034 | 0.129612 | -2.399743 | 0.016406589 | 0.17572204 |
| *CNOT9* | -0.242906 | 0.101238 | -2.399356 | 0.01642395 | 0.17572204 |
| *COMMD9* | -0.212928 | 0.088807 | -2.397639 | 0.016501137 | 0.175810936 |
| *RAD17* | -0.335083 | 0.13976 | -2.397569 | 0.016504254 | 0.175810936 |
| *LRWD1* | 0.408834 | 0.170541 | 2.397285 | 0.016517069 | 0.175810936 |
| *DDX20* | 0.335802 | 0.140103 | 2.396826 | 0.01653776 | 0.175810936 |
| *LMNA* | 0.500817 | 0.208951 | 2.396814 | 0.016538305 | 0.175810936 |
| *GABPB1* | -0.232146 | 0.096857 | -2.3968 | 0.016538961 | 0.175810936 |
| *TMEM11* | 0.207678 | 0.086685 | 2.395778 | 0.016585144 | 0.176083082 |
| *ZNF506* | 0.368328 | 0.153778 | 2.39519 | 0.016611771 | 0.176083082 |
| *ENSG00000236255* | 0.988413 | 0.412682 | 2.395096 | 0.016616014 | NA |
| *TRIP11* | 0.231037 | 0.096464 | 2.395052 | 0.016617997 | 0.176083082 |
| *TMCO3* | -0.350324 | 0.146321 | -2.394226 | 0.016655504 | 0.176291554 |
| *ZNF330* | 0.315122 | 0.131711 | 2.39252 | 0.016733102 | 0.176603013 |
| *SDAD1* | 0.235691 | 0.098518 | 2.392373 | 0.016739823 | 0.176603013 |
| *MCL1* | 0.232196 | 0.097067 | 2.392108 | 0.016751899 | 0.176603013 |
| *ENSG00000237188* | -0.786321 | 0.328728 | -2.39201 | 0.016756385 | 0.176603013 |
| *NOLC1* | 0.340005 | 0.142168 | 2.391565 | 0.016776732 | 0.176629153 |
| *DENND5A* | 0.423112 | 0.177027 | 2.390105 | 0.016843552 | 0.177131192 |
| *UPF3A* | -0.299758 | 0.125435 | -2.389741 | 0.016860252 | 0.177131192 |
| *FBXO11* | 0.298268 | 0.124853 | 2.388955 | 0.016896372 | 0.177322223 |
| *SERPINB6* | 0.465288 | 0.194859 | 2.387817 | 0.016948761 | 0.177683401 |
| *EYA3* | 0.519395 | 0.217608 | 2.386836 | 0.016994061 | 0.177888909 |
| *ANKAR* | -0.533763 | 0.223649 | -2.386614 | 0.017004352 | 0.177888909 |
| *YY1* | 0.151154 | 0.063353 | 2.38592 | 0.017036439 | 0.178036191 |
| *ASPH* | 1.41203 | 0.592035 | 2.385043 | 0.017077106 | NA |
| *ZNF254* | -0.446518 | 0.187216 | -2.385043 | 0.017077146 | 0.178105438 |
| *PRCP* | -0.363937 | 0.152594 | -2.385001 | 0.017079097 | 0.178105438 |
| *ATP5PD* | -0.201685 | 0.084615 | -2.383559 | 0.017146149 | 0.178509367 |
| *RRP1B* | 0.382715 | 0.160576 | 2.383391 | 0.017153945 | 0.178509367 |
| *RRM2B* | -0.308042 | 0.129293 | -2.38251 | 0.017195047 | 0.178748935 |
| *LRP5L* | -0.796074 | 0.334191 | -2.382096 | 0.017214411 | 0.178762252 |
| *MS4A1* | -0.295127 | 0.123916 | -2.381681 | 0.017233805 | 0.178775858 |
| *SPRY1* | -0.76141 | 0.319774 | -2.381086 | 0.017261699 | 0.178802457 |
| *PLEKHG1* | -0.247168 | 0.103823 | -2.380675 | 0.017280967 | 0.178802457 |
| *CDCA4* | 0.33896 | 0.142399 | 2.380354 | 0.017296021 | 0.178802457 |
| *MBOAT2* | -1.438501 | 0.604468 | -2.379781 | 0.017322927 | NA |
| *SESTD1* | -0.424307 | 0.17835 | -2.379072 | 0.017356303 | 0.178802457 |
| *UQCRHL* | -0.543099 | 0.228298 | -2.378899 | 0.017364444 | 0.178802457 |
| *WDR4* | 0.437226 | 0.183806 | 2.378743 | 0.017371766 | 0.178802457 |
| *ANKRD36C* | -0.499521 | 0.209998 | -2.378696 | 0.017373984 | 0.178802457 |
| *DIMT1* | 0.2895 | 0.121729 | 2.378228 | 0.017396062 | 0.178802457 |
| *MLYCD* | -0.692493 | 0.291188 | -2.378163 | 0.017399147 | 0.178802457 |
| *FARSA* | 0.237011 | 0.09969 | 2.377472 | 0.01743176 | 0.178951584 |
| *CARD11* | 0.478485 | 0.201302 | 2.376947 | 0.017456574 | 0.179020428 |
| *PWWP2B* | 0.63189 | 0.265884 | 2.37656 | 0.017474924 | 0.179022898 |
| *MYL12B* | 0.208715 | 0.087838 | 2.376138 | 0.017494939 | 0.17904241 |

| | | | | | |
|---|---|---|---|---|---|
| *BCL2A1* | 0.447838 | 0.188545 | 2.37523 | 0.017538033 | 0.179297821 |
| *SDC3* | 1.888946 | 0.79551 | 2.374508 | 0.017572365 | NA |
| *CYCS* | 0.196277 | 0.082691 | 2.373618 | 0.017614761 | 0.179896207 |
| *ZNF530* | -0.499648 | 0.210556 | -2.372989 | 0.017644768 | 0.180016698 |
| *SRP9* | -0.190323 | 0.080251 | -2.371614 | 0.017710582 | 0.18047462 |
| *POLG2* | 0.421913 | 0.177927 | 2.371271 | 0.017727022 | 0.18047462 |
| *RNF13* | -0.275173 | 0.116064 | -2.370872 | 0.017746169 | 0.18047462 |
| *MRPL13* | 0.193986 | 0.081845 | 2.370163 | 0.017780252 | 0.18047462 |
| *PUM3* | 0.380084 | 0.160364 | 2.37013 | 0.017781844 | 0.18047462 |
| *COPS2* | 0.290993 | 0.122794 | 2.369769 | 0.017799186 | 0.18047462 |
| *TMA16* | 0.349813 | 0.14775 | 2.367603 | 0.017903759 | 0.181158 |
| *EIF5A2* | 0.443864 | 0.187545 | 2.366711 | 0.017946947 | 0.181158 |
| *EDF1* | -0.130519 | 0.055151 | -2.366586 | 0.017952998 | 0.181158 |
| *TENT4B* | 0.541709 | 0.228906 | 2.366512 | 0.017956588 | 0.181158 |
| *PURA* | 0.389557 | 0.164615 | 2.366479 | 0.017958208 | 0.181158 |
| *IDH3A* | 0.277269 | 0.117191 | 2.365966 | 0.017983087 | 0.181161947 |
| *WAC* | 0.171262 | 0.072393 | 2.365716 | 0.017995249 | 0.181161947 |
| *TMEM170A* | 0.269041 | 0.113829 | 2.363557 | 0.018100452 | 0.182035675 |
| *PI4KB* | -0.296391 | 0.125435 | -2.362905 | 0.018132338 | 0.182112542 |
| *PPFIBP1* | -1.025557 | 0.434071 | -2.362647 | 0.018144938 | 0.182112542 |
| *MDN1* | 0.47586 | 0.201515 | 2.36141 | 0.018205614 | 0.182536203 |
| *WDR6* | -0.269062 | 0.113973 | -2.36074 | 0.01823851 | 0.182680756 |
| *STARD10* | -0.337411 | 0.143005 | -2.359433 | 0.018302869 | 0.183139842 |
| *PYM1* | -0.263031 | 0.111529 | -2.358405 | 0.018353638 | 0.1833362 |
| *ZNF148* | -0.228828 | 0.097031 | -2.358285 | 0.018359583 | 0.1833362 |
| *COX8A* | -0.156719 | 0.06648 | -2.357391 | 0.018403864 | 0.183592936 |
| *RIOK1* | 0.367892 | 0.156179 | 2.355585 | 0.018493574 | 0.184035056 |
| *ETNK1* | 0.251465 | 0.106758 | 2.355473 | 0.018499161 | 0.184035056 |
| *RUFY2* | -0.412754 | 0.175239 | -2.355375 | 0.018504031 | 0.184035056 |
| *DTX3* | 1.806038 | 0.766959 | 2.354803 | 0.018532504 | NA |
| *EXOSC5* | 0.386856 | 0.164434 | 2.352651 | 0.018640098 | 0.185202014 |
| *TNS3* | 0.704811 | 0.299747 | 2.351351 | 0.018705371 | 0.185663949 |
| *ACSM3* | -0.671889 | 0.285846 | -2.350528 | 0.018746781 | 0.185758891 |
| *PTRHD1* | -0.317259 | 0.13498 | -2.350414 | 0.018752516 | 0.185758891 |
| *CLCF1* | 0.521319 | 0.221949 | 2.348828 | 0.018832618 | 0.186365625 |
| *ITFG1* | -0.323176 | 0.137634 | -2.348083 | 0.018870327 | 0.186552052 |
| *SERF2* | -0.161412 | 0.068755 | -2.34765 | 0.018892257 | 0.186582266 |
| *SYNE2* | -0.546962 | 0.23311 | -2.34637 | 0.018957255 | 0.186871929 |
| *WRAP53* | -0.491841 | 0.209649 | -2.346026 | 0.018974773 | 0.186871929 |
| *SDF2L1* | -0.26074 | 0.111145 | -2.345957 | 0.018978294 | 0.186871929 |
| *PLTP* | -1.358493 | 0.579182 | -2.34554 | 0.01899956 | NA |
| *NDUFA13* | -0.186108 | 0.079352 | -2.345358 | 0.019008817 | 0.18698623 |
| *TAF5* | 0.550428 | 0.234893 | 2.343319 | 0.019113046 | 0.187824628 |
| *HACD4* | -1.026091 | 0.438092 | -2.342183 | 0.019171301 | 0.188210016 |
| *FXYD2* | -0.922957 | 0.394394 | -2.340194 | 0.019273702 | 0.18869438 |
| *SRSF10* | 0.176496 | 0.075422 | 2.340127 | 0.019277164 | 0.18869438 |
| *BRMS1L* | 0.476087 | 0.203466 | 2.339879 | 0.019289989 | 0.18869438 |

| | | | | | |
|---|---|---|---|---|---|
| *CCND2* | 0.542867 | 0.23202 | 2.339744 | 0.019296988 | 0.18869438 |
| *FCHSD2* | -0.368947 | 0.157753 | -2.338767 | 0.019347497 | 0.18891499 |
| *SPATC1L* | -0.588951 | 0.25188 | -2.338218 | 0.019375941 | 0.18891499 |
| *LARP4B* | 0.432595 | 0.185012 | 2.3382 | 0.019376876 | 0.18891499 |
| *IFITM3* | -1.801063 | 0.770318 | -2.338077 | 0.019383243 | NA |
| *SREK1* | 0.179513 | 0.076811 | 2.337073 | 0.019435402 | 0.189298901 |
| *NEURL1* | 1.718641 | 0.735687 | 2.336105 | 0.019485779 | NA |
| *ESS2* | -0.412108 | 0.176434 | -2.335757 | 0.019503905 | 0.189779136 |
| *ERGIC2* | 0.262424 | 0.112406 | 2.334604 | 0.019564137 | 0.19017803 |
| *ENSG00000233903* | 1.018254 | 0.436287 | 2.333911 | 0.019600362 | NA |
| *PRDX1* | 0.26727 | 0.114529 | 2.333654 | 0.019613857 | 0.190298871 |
| *ID3* | 0.26893 | 0.115268 | 2.333081 | 0.019643898 | 0.190298871 |
| *YPEL1* | -0.589462 | 0.252672 | -2.332918 | 0.019652455 | 0.190298871 |
| *TSC22D2* | 0.346273 | 0.14843 | 2.332897 | 0.019653565 | 0.190298871 |
| *SIK3* | 0.488601 | 0.209501 | 2.332208 | 0.01968974 | 0.190462591 |
| *OTULINL* | -1.165627 | 0.499897 | -2.331732 | 0.019714772 | NA |
| *EML4* | 0.297757 | 0.127703 | 2.331644 | 0.019719438 | 0.190563409 |
| *ANKRD36* | -0.5209 | 0.223465 | -2.331009 | 0.019752877 | 0.190700142 |
| *PLCB2* | 1.227244 | 0.526724 | 2.329959 | 0.01980831 | NA |
| *C18orf25* | 0.36881 | 0.158295 | 2.329893 | 0.019811827 | 0.190882327 |
| *CDKN1B* | 0.314229 | 0.13487 | 2.329856 | 0.019813764 | 0.190882327 |
| *SRD5A3* | 0.779584 | 0.334649 | 2.329555 | 0.019829673 | 0.190882327 |
| *ZCCHC7* | -0.349529 | 0.150081 | -2.328945 | 0.019861971 | 0.191007239 |
| *NMD3* | 0.283293 | 0.121677 | 2.328241 | 0.01989931 | 0.191180353 |
| *SERP2* | -1.894184 | 0.813741 | -2.327748 | 0.019925499 | NA |
| *PTGER4* | 0.57722 | 0.248128 | 2.326302 | 0.020002449 | 0.191689817 |
| *NAP1L1* | 0.26669 | 0.114654 | 2.326039 | 0.020016467 | 0.191689817 |
| *ARPP19* | 0.153126 | 0.065832 | 2.326008 | 0.020018143 | 0.191689817 |
| *IFI16* | 0.229701 | 0.098763 | 2.325787 | 0.020029899 | 0.191689817 |
| *MIAT* | -0.994781 | 0.427791 | -2.325388 | 0.020051201 | 0.191693069 |
| *SMIM14* | -0.499389 | 0.214786 | -2.325055 | 0.020069019 | 0.191693069 |
| *ITGB1BP1* | -0.39392 | 0.169497 | -2.32405 | 0.020122798 | 0.192021218 |
| *EPB41L2* | -0.387114 | 0.166626 | -2.323248 | 0.020165851 | 0.192114794 |
| *ATP5F1E* | -0.128896 | 0.055486 | -2.323036 | 0.020177221 | 0.192114794 |
| *APOBEC3C* | -0.34238 | 0.147401 | -2.322781 | 0.020190903 | 0.192114794 |
| *CRYZ* | 0.742194 | 0.319793 | 2.320857 | 0.020294546 | 0.192915269 |
| *MBOAT7* | 0.643383 | 0.277272 | 2.3204 | 0.020319224 | 0.192964309 |
| *POLE3* | -0.159428 | 0.068745 | -2.319108 | 0.020389204 | 0.19344306 |
| *ARPC5* | -0.203922 | 0.087981 | -2.3178 | 0.020460196 | 0.19393049 |
| *PSMB10* | -0.255317 | 0.110176 | -2.317359 | 0.020484178 | 0.193971821 |
| *ADGRB2* | -1.280403 | 0.552588 | -2.317101 | 0.020498208 | NA |
| *ECE2* | 0.542106 | 0.23402 | 2.316497 | 0.020531133 | 0.194230414 |
| *IMPACT* | -0.517473 | 0.223521 | -2.315102 | 0.020607328 | 0.194764857 |
| *ALKBH7* | -0.240569 | 0.103954 | -2.314188 | 0.02065741 | 0.194991887 |
| *PIK3AP1* | -0.291207 | 0.125849 | -2.313944 | 0.020670797 | 0.194991887 |
| *ADAP1* | 1.568436 | 0.678165 | 2.312765 | 0.020735557 | NA |
| *NCF1* | -0.333274 | 0.144107 | -2.312688 | 0.020739801 | 0.195456317 |

| | | | | | |
|---|---|---|---|---|---|
| *TIMELESS* | -0.582487 | 0.251942 | -2.31199 | 0.020778222 | 0.195631908 |
| *PNKD* | -0.280993 | 0.121589 | -2.311005 | 0.020832586 | 0.195957135 |
| *PRKAG1* | -0.254027 | 0.109987 | -2.309602 | 0.020910192 | 0.196352006 |
| *ETFDH* | -0.383697 | 0.166162 | -2.309171 | 0.020934111 | 0.196352006 |
| *MFSD1* | 0.46342 | 0.200687 | 2.30917 | 0.020934151 | 0.196352006 |
| *KIAA1324L* | 1.321439 | 0.57239 | 2.308636 | 0.02096381 | NA |
| *HUS1* | 0.3462 | 0.150001 | 2.30798 | 0.021000249 | 0.196785267 |
| *AP3B1* | -0.48765 | 0.211714 | -2.303345 | 0.02125942 | 0.199025217 |
| *MMADHC* | 0.190724 | 0.082874 | 2.30138 | 0.021370149 | 0.199872557 |
| *TMEM230* | -0.156168 | 0.067883 | -2.300545 | 0.021417365 | 0.19992941 |
| *USP24* | -0.330338 | 0.143606 | -2.300305 | 0.021430958 | 0.19992941 |
| *ZNF322* | 0.395146 | 0.171855 | 2.299301 | 0.021487825 | 0.19992941 |
| *NUP210* | 0.359576 | 0.156391 | 2.29921 | 0.021493039 | 0.19992941 |
| *GTF2H1* | 0.230625 | 0.100325 | 2.298776 | 0.021517641 | 0.19992941 |
| *C12orf43* | -0.294141 | 0.127993 | -2.298105 | 0.021555819 | 0.19992941 |
| *SSNA1* | -0.214349 | 0.093325 | -2.2968 | 0.0216302 | 0.19992941 |
| *DNAJC16* | 0.557873 | 0.242952 | 2.296229 | 0.02166278 | 0.19992941 |
| *WDR34* | -0.423157 | 0.184288 | -2.296168 | 0.021666252 | 0.19992941 |
| *EZH2* | 0.368938 | 0.160676 | 2.296165 | 0.021666446 | 0.19992941 |
| *KLHDC2* | -0.266019 | 0.115865 | -2.295934 | 0.021679663 | 0.19992941 |
| *GTF2H5* | -0.277771 | 0.120993 | -2.295766 | 0.021689265 | 0.19992941 |
| *DNAL1* | -0.628855 | 0.273921 | -2.295758 | 0.021689748 | 0.19992941 |
| *BORCS7* | -0.22212 | 0.096762 | -2.295517 | 0.021703514 | 0.19992941 |
| *HECA* | 0.326496 | 0.142239 | 2.295406 | 0.021709871 | 0.19992941 |
| *FKBPL* | 0.515661 | 0.224699 | 2.294901 | 0.021738795 | 0.19992941 |
| *MYO1E* | 0.5553 | 0.241997 | 2.294653 | 0.021752991 | 0.19992941 |
| *IFRD2* | 0.254329 | 0.110859 | 2.294164 | 0.021781107 | 0.19992941 |
| *TADA2A* | 0.463716 | 0.202171 | 2.293677 | 0.021809038 | 0.19992941 |
| *ACTL6A* | 0.365343 | 0.159286 | 2.29362 | 0.021812334 | 0.19992941 |
| *GAR1* | 0.385429 | 0.168046 | 2.293589 | 0.021814121 | 0.19992941 |
| *JAM2* | -0.766354 | 0.33416 | -2.293377 | 0.021826337 | 0.19992941 |
| *MRM1* | 0.510331 | 0.222564 | 2.292963 | 0.021850113 | 0.19992941 |
| *ZNF33B* | -0.371611 | 0.162085 | -2.292689 | 0.021865893 | 0.19992941 |
| *RAB1B* | -0.261098 | 0.113897 | -2.292413 | 0.021881815 | 0.19992941 |
| *MAGEH1* | -0.458892 | 0.200217 | -2.291968 | 0.021907475 | 0.199979041 |
| *C8orf82* | -0.947703 | 0.413559 | -2.291577 | 0.021930072 | NA |
| *MSMP* | -0.847985 | 0.370129 | -2.291054 | 0.021960275 | 0.200276089 |
| *BBS12* | -0.861729 | 0.376216 | -2.290515 | 0.021991454 | 0.200375591 |
| *ENSG00000235501* | -1.880291 | 0.82133 | -2.289326 | 0.022060446 | NA |
| *APOO* | 0.613507 | 0.267996 | 2.289237 | 0.022065568 | 0.200620931 |
| *COLGALT1* | -0.458769 | 0.200415 | -2.289091 | 0.022074055 | 0.200620931 |
| *SRPK1* | -0.205804 | 0.089954 | -2.287863 | 0.022145486 | 0.200620931 |
| *BMI1* | 0.28568 | 0.124888 | 2.287495 | 0.022166946 | 0.200620931 |
| *MRPL57* | -0.159107 | 0.06956 | -2.287345 | 0.022175683 | 0.200620931 |
| *TFAP4* | 0.634835 | 0.277549 | 2.287288 | 0.02217903 | 0.200620931 |
| *WDR90* | 0.973195 | 0.425538 | 2.286976 | 0.022197214 | 0.200620931 |
| *SUPT16H* | -0.245186 | 0.107225 | -2.286644 | 0.022216587 | 0.200620931 |

| | | | | | |
|---|---|---|---|---|---|
| *TMEM192* | 0.353201 | 0.154481 | 2.286377 | 0.02223224 | 0.200620931 |
| *ELP6* | -0.362182 | 0.158428 | -2.286104 | 0.022248175 | 0.200620931 |
| *ANAPC7* | -0.328574 | 0.143742 | -2.285863 | 0.022262281 | 0.200620931 |
| *ENSG00000224086* | -0.511738 | 0.223875 | -2.285821 | 0.022264728 | 0.200620931 |
| *RAB14* | 0.172591 | 0.075515 | 2.285523 | 0.022282195 | 0.200620931 |
| *CCDC69* | -0.359428 | 0.15739 | -2.283671 | 0.022390846 | 0.200985042 |
| *SEMA7A* | 0.265266 | 0.116163 | 2.283557 | 0.022397574 | 0.200985042 |
| *HES1* | 0.787996 | 0.345076 | 2.283546 | 0.022398242 | 0.200985042 |
| *RPS6KB2.AS1* | -1.036145 | 0.453761 | -2.28346 | 0.022403259 | NA |
| *DNAJA3* | 0.276691 | 0.121172 | 2.283449 | 0.022403957 | 0.200985042 |
| *ROR1* | -1.508746 | 0.660998 | -2.282529 | 0.022458148 | 0.201288528 |
| *TRMT11* | 0.363256 | 0.159262 | 2.280863 | 0.022556533 | 0.201811084 |
| *ENSG00000267458* | -0.698436 | 0.306217 | -2.280851 | 0.022557278 | 0.201811084 |
| *NXPE3* | -0.535066 | 0.234747 | -2.279328 | 0.022647561 | 0.202435617 |
| *TSN* | -0.187849 | 0.082435 | -2.278763 | 0.022681187 | 0.202507585 |
| *CRYZL1* | -0.263477 | 0.115636 | -2.278504 | 0.022696581 | 0.202507585 |
| *KLK4* | 2.292028 | 1.006234 | 2.277828 | 0.022736841 | NA |
| *IGHE* | -1.034939 | 0.454354 | -2.277823 | 0.022737099 | 0.202686171 |
| *MBNL1* | -0.233556 | 0.102575 | -2.276929 | 0.022790483 | 0.202713135 |
| *MAST3* | 0.577398 | 0.253603 | 2.276783 | 0.022799164 | 0.202713135 |
| *BCL7A* | -0.569198 | 0.250005 | -2.276742 | 0.022801639 | 0.202713135 |
| *TTC14* | -0.238261 | 0.104725 | -2.275113 | 0.022899129 | 0.202900531 |
| *RP9* | 0.399979 | 0.175812 | 2.275042 | 0.022903423 | 0.202900531 |
| *HIP1R* | -0.37381 | 0.164326 | -2.274806 | 0.022917549 | 0.202900531 |
| *SLC9B2* | 0.570332 | 0.250783 | 2.274205 | 0.022953665 | 0.202900531 |
| *EHD4* | -0.292361 | 0.128562 | -2.274081 | 0.022961088 | 0.202900531 |
| *MYD88* | -0.337537 | 0.148433 | -2.274007 | 0.022965583 | 0.202900531 |
| *GSTO1* | 0.228665 | 0.100557 | 2.273993 | 0.022966386 | 0.202900531 |
| *FOSL2* | 0.788304 | 0.346949 | 2.272107 | 0.023080064 | 0.203722783 |
| *EBNA1BP2* | 0.483369 | 0.212957 | 2.269802 | 0.023219575 | 0.20477138 |
| *AKAP17A* | 0.242813 | 0.106993 | 2.269438 | 0.023241718 | 0.204783977 |
| *FXYD7* | 0.828837 | 0.365568 | 2.267258 | 0.023374457 | 0.205770153 |
| *NOC3L* | 0.205926 | 0.090876 | 2.266021 | 0.023450121 | 0.205949378 |
| *MDM4* | -0.193829 | 0.085539 | -2.265983 | 0.023452452 | 0.205949378 |
| *HIF1A* | 0.21887 | 0.096593 | 2.265903 | 0.023457313 | 0.205949378 |
| *XRCC1* | -0.375908 | 0.16596 | -2.265057 | 0.023509181 | 0.206221618 |
| *LINC00623* | -0.394895 | 0.174396 | -2.264362 | 0.023551862 | 0.206412864 |
| *ENSG00000268858* | -0.861421 | 0.380443 | -2.264257 | 0.023558338 | NA |
| *BACE2* | 1.561015 | 0.689556 | 2.263798 | 0.023586535 | NA |
| *CCNH* | 0.265573 | 0.117324 | 2.263591 | 0.023599295 | 0.206645373 |
| *ZMAT5* | -0.301637 | 0.133286 | -2.263079 | 0.023630828 | 0.206738381 |
| *AKAP12* | -2.061008 | 0.910778 | -2.26291 | 0.023641258 | NA |
| *CHIC2* | 0.305521 | 0.135043 | 2.262391 | 0.023673283 | 0.206926677 |
| *WDFY1* | 0.34956 | 0.154546 | 2.261844 | 0.023707039 | 0.206965742 |
| *CDK6* | -0.549082 | 0.242792 | -2.261535 | 0.023726145 | 0.206965742 |
| *KIAA0040* | 0.352709 | 0.155976 | 2.261302 | 0.023740557 | 0.206965742 |
| *MCM6* | 0.836724 | 0.370267 | 2.259783 | 0.023834718 | 0.207577592 |

| | | | | | |
|---|---|---|---|---|---|
| *APBB1IP* | 0.436508 | 0.193189 | 2.259493 | 0.023852736 | 0.207577592 |
| *ITPKB* | -0.322861 | 0.142916 | -2.259104 | 0.023876947 | 0.207605537 |
| *OVCH1.AS1* | -1.513783 | 0.670118 | -2.258979 | 0.02388469 | NA |
| *ADAM10* | -0.2896 | 0.128216 | -2.258681 | 0.023903226 | 0.207651397 |
| *ENSG00000272335* | 1.339095 | 0.592957 | 2.258334 | 0.023924814 | NA |
| *BBS9* | -0.72371 | 0.320681 | -2.256791 | 0.024021134 | 0.20849248 |
| *LIPT2* | -0.877108 | 0.388686 | -2.2566 | 0.024033107 | NA |
| *PPRC1* | 0.471031 | 0.2088 | 2.255894 | 0.024077289 | 0.208598628 |
| *CEP68* | -0.527971 | 0.234057 | -2.255741 | 0.02408688 | 0.208598628 |
| *CCDC130* | 0.268828 | 0.119223 | 2.254833 | 0.024143795 | 0.208598628 |
| *PNPT1* | 0.385747 | 0.171133 | 2.254077 | 0.024191342 | 0.208598628 |
| *SKAP2* | 0.303963 | 0.134853 | 2.254034 | 0.024194049 | 0.208598628 |
| *S100A11* | 0.271057 | 0.12026 | 2.253923 | 0.024200989 | 0.208598628 |
| *PHF6* | 0.464225 | 0.205976 | 2.253786 | 0.024209619 | 0.208598628 |
| *APBB1* | 0.60904 | 0.270279 | 2.253376 | 0.024235447 | 0.208598628 |
| *MRPS2* | 0.340367 | 0.151048 | 2.253371 | 0.02423576 | 0.208598628 |
| *BAZ2B* | -0.463529 | 0.20574 | -2.252985 | 0.02426008 | 0.208598628 |
| *FXYD1* | 0.857365 | 0.380561 | 2.2529 | 0.024265469 | 0.208598628 |
| *MT.ATP8* | -0.497061 | 0.22072 | -2.252002 | 0.024322159 | 0.208904311 |
| *RANBP2* | 0.334196 | 0.148445 | 2.251307 | 0.024366076 | 0.209085104 |
| *TMC6* | -0.318735 | 0.141615 | -2.250721 | 0.024403189 | 0.209085104 |
| *PITPNA.AS1* | -0.404405 | 0.179692 | -2.250544 | 0.024414409 | 0.209085104 |
| *RBBP8* | 0.37643 | 0.16729 | 2.250164 | 0.024438549 | 0.209085104 |
| *MANF* | 0.226393 | 0.100619 | 2.25 | 0.024448956 | 0.209085104 |
| *MAD1L1* | 0.403781 | 0.179498 | 2.249509 | 0.024480155 | 0.209170972 |
| *UBE2S* | 0.317481 | 0.141264 | 2.24742 | 0.024613181 | 0.210125998 |
| *GNG7* | -0.318255 | 0.141697 | -2.246034 | 0.024701807 | 0.210422703 |
| *SNAPC1* | 0.302371 | 0.134629 | 2.245954 | 0.024706936 | 0.210422703 |
| *CDKN2AIP* | 0.230542 | 0.102691 | 2.245014 | 0.024767225 | 0.210422703 |
| *AHSA1* | 0.272119 | 0.121212 | 2.244979 | 0.024769494 | 0.210422703 |
| *ENSG00000274422* | 0.42953 | 0.191346 | 2.244784 | 0.024782025 | 0.210422703 |
| *DOP1B* | -0.47116 | 0.209903 | -2.244661 | 0.024789924 | 0.210422703 |
| *NAT10* | 0.474598 | 0.211444 | 2.244552 | 0.02479693 | 0.210422703 |
| *HTR3A* | -0.949032 | 0.422982 | -2.243671 | 0.024853576 | NA |
| *SLC25A38* | 0.201028 | 0.08964 | 2.242605 | 0.024922286 | 0.211305077 |
| *R3HDM2* | -0.368116 | 0.164185 | -2.242074 | 0.024956583 | 0.211393241 |
| *PIGQ* | -0.455492 | 0.203183 | -2.241782 | 0.024975451 | 0.211393241 |
| *NET1* | 1.625391 | 0.725063 | 2.241723 | 0.024979257 | NA |
| *SRP68* | 0.265977 | 0.118694 | 2.240871 | 0.025034441 | 0.211711277 |
| *PSMA6* | -0.269392 | 0.120295 | -2.239429 | 0.025128033 | 0.21227619 |
| *ENSG00000256092* | -0.428999 | 0.191595 | -2.239092 | 0.025149944 | 0.21227619 |
| *ABL1* | -0.626343 | 0.279761 | -2.23885 | 0.025165658 | 0.21227619 |
| *NRP2* | -0.454591 | 0.203085 | -2.238422 | 0.025193522 | 0.212330062 |
| *UAP1L1* | 0.978226 | 0.43708 | 2.238094 | 0.02521492 | NA |
| *MPV17L2* | 0.385681 | 0.172384 | 2.237339 | 0.025264166 | 0.212561378 |
| *C11orf68* | -0.396517 | 0.177277 | -2.236713 | 0.025305134 | 0.212561378 |
| *SNU13* | 0.182631 | 0.081663 | 2.236393 | 0.025326019 | 0.212561378 |

| | | | | | |
|---|---|---|---|---|---|
| *TRIM4* | 0.308705 | 0.138047 | 2.236229 | 0.025336792 | 0.212561378 |
| *HILPDA* | -0.405234 | 0.181228 | -2.236042 | 0.025349048 | 0.212561378 |
| *ATP6V1C2* | -1.144509 | 0.511882 | -2.235886 | 0.025359253 | NA |
| *ECHS1* | -0.194186 | 0.086852 | -2.235827 | 0.025363132 | 0.212561378 |
| *ARL6IP6* | 0.205287 | 0.091832 | 2.235461 | 0.025387094 | 0.212561378 |
| *LUCAT1* | 0.691547 | 0.309393 | 2.235171 | 0.025406112 | 0.212561378 |
| *RNF7* | -0.146008 | 0.065327 | -2.235038 | 0.025414841 | 0.212561378 |
| *GRWD1* | 0.256998 | 0.115002 | 2.234716 | 0.025435981 | 0.212561378 |
| *GPBP1L1* | -0.201619 | 0.090244 | -2.234148 | 0.025473325 | 0.21269366 |
| *AFF4* | 0.250708 | 0.112301 | 2.232454 | 0.025584972 | 0.213445599 |
| *KATNAL1* | -0.551881 | 0.247305 | -2.231583 | 0.025642535 | 0.213678967 |
| *TMEM242* | -0.313826 | 0.140661 | -2.231079 | 0.025675908 | 0.213678967 |
| *GXYLT1* | 0.414175 | 0.185662 | 2.230805 | 0.025694051 | 0.213678967 |
| *NASP* | 0.204543 | 0.091693 | 2.230724 | 0.025699402 | 0.213678967 |
| *RIOX2* | 0.402433 | 0.180468 | 2.229943 | 0.025751217 | 0.213869357 |
| *RRP1* | 0.273363 | 0.122599 | 2.229727 | 0.025765568 | 0.213869357 |
| *DNAJC2* | 0.250912 | 0.112617 | 2.228006 | 0.025880127 | 0.214542254 |
| *ZNF516* | 0.809183 | 0.363211 | 2.227857 | 0.025890037 | 0.214542254 |
| *ARHGAP19* | -0.993988 | 0.446428 | -2.226539 | 0.025978081 | NA |
| *GLRX3* | 0.230897 | 0.103734 | 2.225865 | 0.026023212 | 0.215465224 |
| *RHBDD2* | -0.235033 | 0.105644 | -2.224769 | 0.026096732 | 0.215893135 |
| *PTTG1* | 0.418449 | 0.188142 | 2.224108 | 0.026141149 | 0.216037931 |
| *DCAF16* | -0.329282 | 0.148083 | -2.223639 | 0.026172752 | 0.216037931 |
| *SUZ12* | 0.239167 | 0.107562 | 2.223534 | 0.026179794 | 0.216037931 |
| *JAML* | -1.518141 | 0.682818 | -2.223346 | 0.026192462 | NA |
| *UBR5.AS1* | -0.41512 | 0.186735 | -2.223044 | 0.026212826 | 0.216130105 |
| *MARCHF3* | -0.449193 | 0.202127 | -2.222329 | 0.026261104 | 0.216347727 |
| *DHX30* | -0.247802 | 0.11158 | -2.220845 | 0.026361486 | 0.216993884 |
| *ARHGEF26* | -0.50225 | 0.226209 | -2.220288 | 0.026399236 | 0.217065202 |
| *KYNU* | 0.677413 | 0.305177 | 2.219736 | 0.026436701 | 0.217065202 |
| *RND1* | -0.587313 | 0.264605 | -2.219581 | 0.026447194 | 0.217065202 |
| *NUDT8* | -0.736347 | 0.331774 | -2.219423 | 0.026457978 | 0.217065202 |
| *ITGA6.AS1* | -1.076428 | 0.485052 | -2.219204 | 0.026472861 | NA |
| *SPATA33* | -0.369488 | 0.166504 | -2.219096 | 0.026480224 | 0.217067578 |
| *MARCKSL1* | -0.334529 | 0.150783 | -2.218617 | 0.026512789 | 0.217074311 |
| *RBBP9* | -0.454814 | 0.205043 | -2.218141 | 0.026545213 | 0.217074311 |
| *TMA7* | -0.15179 | 0.068432 | -2.218116 | 0.026546919 | 0.217074311 |
| *MFSD13A* | -0.666137 | 0.300374 | -2.217694 | 0.026575728 | 0.217130288 |
| *GPR157* | 0.805301 | 0.363225 | 2.217084 | 0.026617344 | 0.217290723 |
| *PFDN2* | 0.220127 | 0.099305 | 2.216669 | 0.026645689 | 0.21734264 |
| *ZKSCAN1* | -0.255832 | 0.115495 | -2.215085 | 0.026754262 | 0.218048337 |
| *CD84* | 0.595609 | 0.26896 | 2.214485 | 0.026795417 | 0.218203866 |
| *PLAAT3* | -0.879036 | 0.397118 | -2.21354 | 0.026860431 | 0.21855327 |
| *RPRD1B* | 0.346731 | 0.156706 | 2.212623 | 0.026923635 | 0.218887379 |
| *ENSG00000236514* | 1.355805 | 0.61307 | 2.2115 | 0.027001198 | NA |
| *SLC35E4* | 0.67018 | 0.303122 | 2.210922 | 0.027041222 | 0.219662715 |
| *ICAM1* | 0.371017 | 0.167948 | 2.209113 | 0.027166765 | 0.220329225 |

| | | | | | |
|---|---|---|---|---|---|
| *SRI* | 0.214177 | 0.096952 | 2.209098 | 0.027167846 | 0.220329225 |
| *RIC3* | 0.590801 | 0.267573 | 2.207996 | 0.027244521 | 0.220769947 |
| *CCDC144A* | 1.621383 | 0.734555 | 2.207301 | 0.027293031 | NA |
| *FKBP8* | -0.186224 | 0.084369 | -2.207255 | 0.027296264 | 0.220827863 |
| *LINC01678* | -0.486828 | 0.220558 | -2.207254 | 0.027296343 | 0.220827863 |
| *RUNX1* | -0.296405 | 0.134362 | -2.206017 | 0.027382787 | 0.221346058 |
| *UXS1* | -0.303255 | 0.137508 | -2.205367 | 0.02742836 | 0.221533306 |
| *STX10* | -0.370611 | 0.168115 | -2.204507 | 0.027488714 | 0.221641433 |
| *DOCK11* | -0.318264 | 0.144394 | -2.204129 | 0.027515279 | 0.221641433 |
| *DKC1* | 0.3126 | 0.141829 | 2.204059 | 0.027520169 | 0.221641433 |
| *HLA.DQA1* | 0.492753 | 0.223583 | 2.203899 | 0.027531426 | 0.221641433 |
| *RSU1* | -0.321072 | 0.145725 | -2.203273 | 0.027575493 | 0.221759209 |
| *NT5E* | 0.877692 | 0.398404 | 2.203019 | 0.027593375 | 0.221759209 |
| *ATP5F1B* | 0.220339 | 0.10003 | 2.202736 | 0.027613351 | 0.221759209 |
| *ENSG00000256116* | -1.296161 | 0.589735 | -2.19787 | 0.027958378 | NA |
| *NAA38* | -0.256133 | 0.116577 | -2.19711 | 0.028012606 | 0.224680679 |
| *PNO1* | 0.490547 | 0.223284 | 2.19697 | 0.028022585 | 0.224680679 |
| *TMEM50B* | -0.395893 | 0.180253 | -2.196322 | 0.028068938 | 0.224869955 |
| *DUSP16* | 1.301333 | 0.592599 | 2.195976 | 0.02809366 | NA |
| *ENSG00000259038* | -0.570205 | 0.25973 | -2.195376 | 0.028136649 | 0.22492712 |
| *LYST* | -0.332156 | 0.151303 | -2.195302 | 0.028141973 | 0.22492712 |
| *MTG1* | -0.373842 | 0.170294 | -2.195269 | 0.02814433 | 0.22492712 |
| *SNHG15* | 0.319504 | 0.145568 | 2.194875 | 0.028172575 | 0.224970987 |
| *AMD1* | 0.261053 | 0.118956 | 2.194539 | 0.028196689 | 0.224981816 |
| *SNRNP40* | 0.194141 | 0.088504 | 2.193576 | 0.028265908 | 0.22535223 |
| *LYRM2* | -0.202502 | 0.092332 | -2.193193 | 0.028293474 | 0.225390241 |
| *CMTM8* | -1.159208 | 0.528627 | -2.192867 | 0.028316997 | NA |
| *ITGAE* | -0.255645 | 0.116617 | -2.192184 | 0.028366234 | 0.225787911 |
| *GBF1* | 0.460848 | 0.210321 | 2.191165 | 0.028439821 | 0.226030992 |
| *DNPH1* | -0.353121 | 0.161171 | -2.190968 | 0.028454134 | 0.226030992 |
| *TMEM209* | -0.433734 | 0.197978 | -2.190813 | 0.028465364 | 0.226030992 |
| *LY86* | -0.376052 | 0.171708 | -2.190067 | 0.02851941 | 0.2262784 |
| *ATIC* | 0.423083 | 0.193233 | 2.189501 | 0.028560447 | 0.226422278 |
| *PLCL2* | -0.465945 | 0.213077 | -2.186745 | 0.028761108 | 0.227794693 |
| *TET2* | 0.325822 | 0.149016 | 2.186492 | 0.028779645 | 0.227794693 |
| *STK33* | -1.098065 | 0.502463 | -2.185363 | 0.028862234 | NA |
| *KMO* | 0.380217 | 0.174026 | 2.184821 | 0.028901999 | 0.22845423 |
| *LPAR5* | 0.416692 | 0.19073 | 2.184723 | 0.028909189 | 0.22845423 |
| *C17orf99* | -0.925237 | 0.423708 | -2.183667 | 0.028986714 | NA |
| *P2RY11* | 0.668255 | 0.306187 | 2.182508 | 0.02907204 | 0.229557655 |
| *AFTPH* | 0.192576 | 0.088249 | 2.182182 | 0.029096103 | 0.229564302 |
| *BZW2* | 0.391993 | 0.179716 | 2.181181 | 0.029170014 | 0.22996392 |
| *LGALS3* | 0.567566 | 0.260382 | 2.17974 | 0.029276714 | 0.230289344 |
| *IAH1* | -0.151116 | 0.069328 | -2.17973 | 0.029277466 | 0.230289344 |
| *TGS1* | 0.337502 | 0.15484 | 2.17968 | 0.029281176 | 0.230289344 |
| *HAUS4* | -1.024393 | 0.47029 | -2.178217 | 0.029389878 | 0.23096052 |
| *RUNDC3B* | 1.907377 | 0.876085 | 2.177159 | 0.029468706 | NA |

| | | | | | |
|---|---|---|---|---|---|
| *C17orf75* | 0.770171 | 0.353845 | 2.176579 | 0.029511968 | 0.231735757 |
| *SLC16A1* | 0.415385 | 0.190974 | 2.175093 | 0.029623182 | 0.232424422 |
| *FAM129C* | -0.462288 | 0.212656 | -2.17388 | 0.029714115 | 0.232869887 |
| *ENSG00000170161* | -0.445115 | 0.204772 | -2.173708 | 0.029727068 | 0.232869887 |
| *PLLP* | -1.29517 | 0.595971 | -2.173209 | 0.029764554 | NA |
| *MNAT1* | 0.431393 | 0.19851 | 2.173159 | 0.029768373 | 0.233008816 |
| *INTS13* | 0.395343 | 0.182048 | 2.171646 | 0.0298824 | 0.233600481 |
| *ENSG00000258056* | 0.644702 | 0.296889 | 2.171529 | 0.029891221 | 0.233600481 |
| *BTG3* | 0.236603 | 0.108994 | 2.170785 | 0.029947433 | 0.233854913 |
| *ZMYM5* | 0.288818 | 0.133075 | 2.170341 | 0.029981024 | 0.233932442 |
| *UBTD2* | 0.699621 | 0.322442 | 2.169761 | 0.030024936 | 0.23409031 |
| *AFF3* | -0.377304 | 0.173961 | -2.168897 | 0.030090536 | 0.234416898 |
| *ENSG00000238009* | -1.430163 | 0.659431 | -2.168784 | 0.030099092 | NA |
| *NEK6* | 0.957212 | 0.441477 | 2.168204 | 0.030143143 | 0.234641817 |
| *EZH1* | -0.245726 | 0.113437 | -2.166183 | 0.030297176 | 0.235655294 |
| *UBXN2A* | -0.268215 | 0.123842 | -2.165781 | 0.030327944 | 0.235709162 |
| *CA2* | 0.394123 | 0.182023 | 2.165245 | 0.030368965 | 0.235842565 |
| *ASB2* | 0.857734 | 0.396742 | 2.161942 | 0.030622617 | 0.237505856 |
| *ENSG00000263089* | 0.564757 | 0.26124 | 2.161831 | 0.030631192 | 0.237505856 |
| *NUDT9* | 0.307236 | 0.14215 | 2.161346 | 0.030668604 | 0.237507533 |
| *METTL2A* | 0.299058 | 0.138375 | 2.161206 | 0.030679458 | 0.237507533 |
| *ENSG00000265100* | -1.213359 | 0.561497 | -2.160934 | 0.030700454 | NA |
| *RPS6KB1* | 0.281966 | 0.130501 | 2.160649 | 0.030722438 | 0.237654165 |
| *WDR45* | -0.190061 | 0.087995 | -2.159906 | 0.030779948 | 0.237912876 |
| *GLI4* | -0.864597 | 0.400445 | -2.159092 | 0.030843041 | 0.238075055 |
| *CXorf56* | -0.529915 | 0.245443 | -2.159014 | 0.030849094 | 0.238075055 |
| *H3F3A* | -0.167041 | 0.077444 | -2.156932 | 0.03101095 | 0.239059979 |
| *TRIM3* | -0.659902 | 0.305971 | -2.156751 | 0.031025081 | 0.239059979 |
| *RPL39L* | 0.760094 | 0.352516 | 2.156193 | 0.031068575 | 0.239208668 |
| *MARS2* | 0.735022 | 0.341149 | 2.154551 | 0.031196973 | 0.239378298 |
| *ADH5* | -0.288371 | 0.133865 | -2.15419 | 0.031225311 | 0.239378298 |
| *DCAF7* | 0.191894 | 0.089092 | 2.153877 | 0.031249819 | 0.239378298 |
| *ZNF587B* | 0.582211 | 0.270366 | 2.153415 | 0.031286116 | 0.239378298 |
| *KPNB1* | 0.160381 | 0.074479 | 2.153383 | 0.031288616 | 0.239378298 |
| *ZBTB21* | 0.336598 | 0.156314 | 2.153338 | 0.031292158 | 0.239378298 |
| *PUM1* | 0.339384 | 0.15761 | 2.153316 | 0.031293873 | 0.239378298 |
| *MRPL55* | -0.272642 | 0.126622 | -2.153193 | 0.031303487 | 0.239378298 |
| *MYL6* | -0.155426 | 0.072193 | -2.152913 | 0.031325495 | 0.239378298 |
| *ARHGAP5* | 0.355533 | 0.165165 | 2.152597 | 0.031350392 | 0.239378298 |
| *BLZF1* | 0.347341 | 0.161376 | 2.152366 | 0.031368566 | 0.239378298 |
| *RHOT2* | -0.223972 | 0.104066 | -2.152205 | 0.031381173 | 0.239378298 |
| *CD2* | -3.331441 | 1.548248 | -2.151748 | 0.03141719 | NA |
| *RPL17* | -0.242804 | 0.112853 | -2.151502 | 0.031436575 | 0.239613703 |
| *ASB8* | -0.251178 | 0.116762 | -2.151199 | 0.031460508 | 0.239613703 |
| *RNF146* | 0.255696 | 0.119035 | 2.148067 | 0.031708408 | 0.241150196 |
| *RFXANK* | -0.265693 | 0.123691 | -2.148034 | 0.031711031 | 0.241150196 |
| *AEBP1* | -0.51387 | 0.239297 | -2.147412 | 0.031760513 | 0.241340841 |

| | | | | | |
|---|---|---|---|---|---|
| *ENSG00000273221* | 0.457166 | 0.213137 | 2.144938 | 0.031957803 | 0.242485049 |
| *NIPSNAP2* | -0.207438 | 0.096735 | -2.144401 | 0.032000805 | 0.242485049 |
| *SIDT1.AS1* | -0.723752 | 0.33751 | -2.144384 | 0.032002132 | 0.242485049 |
| *FDPS* | -0.200057 | 0.093297 | -2.144296 | 0.032009204 | 0.242485049 |
| *HYLS1* | -0.33524 | 0.156504 | -2.142061 | 0.03218855 | 0.24365697 |
| *POLA2* | -0.551953 | 0.257736 | -2.14154 | 0.032230481 | 0.243787707 |
| *GDI2* | 0.169113 | 0.079 | 2.140656 | 0.032301816 | 0.244140484 |
| *ENSG00000255135* | -0.665127 | 0.310785 | -2.14015 | 0.032342637 | 0.244262271 |
| *DCAF6* | -0.270711 | 0.126527 | -2.13955 | 0.03239116 | 0.24433274 |
| *PRMT2* | -0.216212 | 0.101061 | -2.139423 | 0.032401398 | 0.24433274 |
| *PM20D2* | 0.561045 | 0.262296 | 2.138977 | 0.032437491 | 0.244418472 |
| *ENSG00000250616* | -1.079588 | 0.504817 | -2.138572 | 0.032470373 | NA |
| *ENSG00000262202* | 0.669232 | 0.312977 | 2.138276 | 0.032494344 | 0.244660382 |
| *PDCL3* | -0.285507 | 0.13356 | -2.137667 | 0.032543758 | 0.244845963 |
| *DPY19L4* | 1.212873 | 0.567893 | 2.135741 | 0.032700527 | NA |
| *CDKN2D* | 0.369095 | 0.172828 | 2.135623 | 0.032710185 | 0.245910942 |
| *ZDHHC3* | -0.290585 | 0.136173 | -2.133945 | 0.032847323 | 0.246616993 |
| *INKA1* | 0.571947 | 0.268034 | 2.133863 | 0.032853993 | 0.246616993 |
| *ENSG00000271895* | -1.357098 | 0.636147 | -2.133308 | 0.032899446 | NA |
| *TUBA1C* | 0.347349 | 0.162936 | 2.131817 | 0.033021933 | 0.247554483 |
| *ZNF17* | 0.39326 | 0.184479 | 2.131731 | 0.033028967 | 0.247554483 |
| *CCDC138* | -0.63308 | 0.297339 | -2.129151 | 0.033241802 | 0.248820643 |
| *PTP4A1* | 0.376844 | 0.177043 | 2.128548 | 0.03329171 | 0.248820643 |
| *IWS1* | 0.243433 | 0.11438 | 2.128291 | 0.033312966 | 0.248820643 |
| *NT5DC3* | 0.720833 | 0.338736 | 2.128008 | 0.033336447 | 0.248820643 |
| *C19orf48* | 0.434159 | 0.204028 | 2.127938 | 0.033342201 | 0.248820643 |
| *FUNDC1* | -0.444212 | 0.20876 | -2.127857 | 0.033348913 | 0.248820643 |
| *THRB* | 1.461888 | 0.687361 | 2.126811 | 0.033435753 | NA |
| *TMSB10* | -0.221581 | 0.104212 | -2.126255 | 0.033482018 | 0.249415253 |
| *PHTF2* | -0.383778 | 0.180497 | -2.126228 | 0.033484257 | 0.249415253 |
| *WDR43* | 0.377304 | 0.177511 | 2.125522 | 0.033543096 | 0.249415253 |
| *CTR9* | 0.303726 | 0.142901 | 2.12543 | 0.03355073 | 0.249415253 |
| *CCDC82* | 0.229567 | 0.10802 | 2.125218 | 0.033568464 | 0.249415253 |
| *SYNGR3* | 1.036381 | 0.48769 | 2.125079 | 0.033579982 | 0.249415253 |
| *LRRC8C* | 0.567693 | 0.267221 | 2.124433 | 0.033633938 | 0.249481875 |
| *H2AFZ* | 0.203285 | 0.095692 | 2.124368 | 0.033639423 | 0.249481875 |
| *RAB31* | 0.854649 | 0.402483 | 2.123443 | 0.033716724 | 0.249684979 |
| *MMD* | -0.623475 | 0.293616 | -2.123436 | 0.033717322 | 0.249684979 |
| *TTC28* | -0.757998 | 0.357288 | -2.121531 | 0.033877167 | 0.250544936 |
| *NAA35* | 0.364412 | 0.171775 | 2.121448 | 0.033884137 | 0.250544936 |
| *XPNPEP3* | -0.381739 | 0.180005 | -2.12071 | 0.033946204 | 0.250816271 |
| *PCYOX1L* | -1.618157 | 0.763252 | -2.120083 | 0.03399902 | NA |
| *ADORA2A.AS1* | 0.815942 | 0.384994 | 2.119364 | 0.034059696 | 0.251466879 |
| *MT1X* | -0.691752 | 0.326596 | -2.118067 | 0.034169398 | 0.252088558 |
| *ELL3* | -1.261002 | 0.595606 | -2.117175 | 0.034244992 | NA |
| *GCLC* | 0.502537 | 0.237364 | 2.117155 | 0.034246711 | 0.252211434 |
| *KLF13* | -0.255664 | 0.120764 | -2.117046 | 0.034255969 | 0.252211434 |

| | | | | | |
|---|---|---|---|---|---|
| *THAP9.AS1* | -0.302344 | 0.14282 | -2.116968 | 0.034262589 | 0.252211434 |
| *ATP5MF* | -0.130522 | 0.061664 | -2.116662 | 0.03428857 | 0.252214887 |
| *EIF2S1* | 0.272091 | 0.128593 | 2.115905 | 0.034352916 | 0.252447918 |
| *LSM8* | -0.132575 | 0.062663 | -2.115688 | 0.034371323 | 0.252447918 |
| *SGSM3* | -0.283764 | 0.134188 | -2.114667 | 0.034458361 | 0.252802127 |
| *UPF1* | 0.367735 | 0.173909 | 2.114522 | 0.034470693 | 0.252802127 |
| *PA2G4* | 0.212031 | 0.100298 | 2.113999 | 0.034515377 | 0.252942194 |
| *RAB40C* | -0.529741 | 0.250747 | -2.112653 | 0.034630453 | 0.25332361 |
| *TMSB4Y* | -1.737448 | 0.822506 | -2.112383 | 0.034653609 | NA |
| *POU2AF1* | -0.251377 | 0.119005 | -2.112315 | 0.03465947 | 0.25332361 |
| *PLEKHF1* | 0.58284 | 0.275945 | 2.112158 | 0.034672934 | 0.25332361 |
| *ENSG00000070423* | 0.214965 | 0.101787 | 2.111909 | 0.03469425 | 0.25332361 |
| *CCDC112* | -0.41366 | 0.195872 | -2.111891 | 0.034695797 | 0.25332361 |
| *SDHAF1* | -0.25617 | 0.121338 | -2.111217 | 0.034753669 | 0.25332361 |
| *HSF2* | 0.458244 | 0.217081 | 2.11094 | 0.034777428 | 0.25332361 |
| *STMP1* | -0.248632 | 0.117793 | -2.110761 | 0.034792849 | 0.25332361 |
| *MTMR12* | -0.428076 | 0.202812 | -2.110701 | 0.034798044 | 0.25332361 |
| *ZBTB26* | -0.617633 | 0.29278 | -2.109546 | 0.034897454 | 0.253545672 |
| *CELF1* | 0.216121 | 0.102453 | 2.109468 | 0.03490418 | 0.253545672 |
| *TRBC2* | -0.19133 | 0.090701 | -2.109453 | 0.034905488 | 0.253545672 |
| *VKORC1L1* | 0.482584 | 0.22889 | 2.108367 | 0.034999266 | 0.253854013 |
| *TYK2* | -0.268946 | 0.127564 | -2.108317 | 0.03500359 | 0.253854013 |
| *PPP1R10* | 0.267538 | 0.126927 | 2.107805 | 0.035047829 | 0.253854013 |
| *MRPS26* | 0.229357 | 0.108854 | 2.107021 | 0.035115799 | 0.253854013 |
| *SDSL* | 2.136029 | 1.013772 | 2.107012 | 0.035116572 | NA |
| *CBLL1* | 0.263162 | 0.124925 | 2.106557 | 0.035155955 | 0.253854013 |
| *ZNHIT3* | 0.191115 | 0.090727 | 2.106491 | 0.035161757 | 0.253854013 |
| *MRTFA* | -0.386516 | 0.183498 | -2.106383 | 0.035171075 | 0.253854013 |
| *TACC1* | 0.274098 | 0.130147 | 2.106061 | 0.035199093 | 0.253854013 |
| *JMY* | 0.569959 | 0.270658 | 2.10583 | 0.03521915 | 0.253854013 |
| *MRPL47* | 0.184077 | 0.087418 | 2.105713 | 0.035229284 | 0.253854013 |
| *RPRD1A* | -0.2463 | 0.116968 | -2.1057 | 0.035230397 | 0.253854013 |
| *ZNF704* | -1.313527 | 0.623846 | -2.105531 | 0.035245071 | NA |
| *IGLL5* | -0.718624 | 0.341364 | -2.105157 | 0.035277636 | 0.254009256 |
| *GPATCH4* | 0.296096 | 0.140714 | 2.104236 | 0.035357841 | 0.254274577 |
| *ENSG00000266962* | -0.316508 | 0.150421 | -2.104144 | 0.035365926 | 0.254274577 |
| *CNFN* | -0.498149 | 0.236909 | -2.102705 | 0.035491583 | 0.254951352 |
| *ANKRD37* | 0.334599 | 0.15915 | 2.102408 | 0.035517584 | 0.254951352 |
| *ZBTB22* | -0.569427 | 0.270875 | -2.102181 | 0.035537423 | 0.254951352 |
| *EIF3J* | 0.224287 | 0.106719 | 2.101659 | 0.035583176 | 0.255026177 |
| *PPP3CA* | -0.235126 | 0.111899 | -2.101238 | 0.035620111 | 0.255026177 |
| *DDX39A* | 0.244645 | 0.116458 | 2.100715 | 0.03566595 | 0.255026177 |
| *USP39* | 0.260361 | 0.123954 | 2.100457 | 0.035688702 | 0.255026177 |
| *ELF4* | 0.421215 | 0.200545 | 2.100354 | 0.035697732 | 0.255026177 |
| *ABI3* | 0.605785 | 0.288428 | 2.100298 | 0.035702633 | 0.255026177 |
| *PTPMT1* | -0.353258 | 0.168264 | -2.099433 | 0.035778723 | 0.255385168 |
| *RRP8* | 0.284861 | 0.135792 | 2.097773 | 0.035925176 | 0.255900562 |

| | | | | | |
|---|---|---|---|---|---|
| *ENSG00000259436* | -0.426695 | 0.20341 | -2.097716 | 0.035930268 | 0.255900562 |
| *PTPN9* | -0.499903 | 0.23831 | -2.097704 | 0.03593126 | 0.255900562 |
| *RBM41* | 0.362569 | 0.172862 | 2.097442 | 0.035954469 | 0.255900562 |
| *ENSG00000227070* | -0.963493 | 0.459513 | -2.096769 | 0.03601402 | 0.256067104 |
| *ILF3* | 0.174508 | 0.083246 | 2.096285 | 0.036056958 | 0.256067104 |
| *ZNF394* | 0.231915 | 0.110655 | 2.095841 | 0.036096336 | 0.256067104 |
| *DYNC1LI2* | 0.318083 | 0.151772 | 2.095799 | 0.036100069 | 0.256067104 |
| *LAMTOR2* | -0.262631 | 0.125349 | -2.095197 | 0.036153511 | 0.256067104 |
| *NFKB1* | 0.258468 | 0.123377 | 2.094939 | 0.03617644 | 0.256067104 |
| *ZBTB8OS* | -0.156862 | 0.074877 | -2.094936 | 0.03617671 | 0.256067104 |
| *ZNF585A* | 0.536705 | 0.256204 | 2.094835 | 0.036185657 | 0.256067104 |
| *CCT2* | 0.315041 | 0.15041 | 2.09455 | 0.036210986 | 0.256067104 |
| *DYNLT1* | 0.264086 | 0.126123 | 2.093875 | 0.036271105 | 0.256308895 |
| *LRIF1* | 0.286921 | 0.137152 | 2.091991 | 0.036439339 | 0.257313791 |
| *DENND4A* | -0.290781 | 0.139041 | -2.091341 | 0.036497513 | 0.257382914 |
| *AASDHPPT* | 0.278913 | 0.133368 | 2.0913 | 0.036501198 | 0.257382914 |
| *ENSG00000273271* | -1.222557 | 0.584681 | -2.090982 | 0.036529688 | NA |
| *GADD45B* | 0.278201 | 0.133069 | 2.090649 | 0.036559534 | 0.257387685 |
| *HDAC8* | -0.439944 | 0.21046 | -2.090394 | 0.03658242 | 0.257387685 |
| *TNFAIP8L2* | 0.522456 | 0.249937 | 2.090348 | 0.036586527 | 0.257387685 |
| *POLB* | 0.544117 | 0.260358 | 2.089875 | 0.036628995 | 0.257387685 |
| *AKAP8L* | 0.273338 | 0.130807 | 2.089624 | 0.036651573 | 0.257387685 |
| *SLC25A53* | -0.326731 | 0.156364 | -2.089552 | 0.036658088 | 0.257387685 |
| *PITPNM2* | -1.006653 | 0.481774 | -2.089469 | 0.036665525 | NA |
| *PF4* | 3.99046 | 1.911254 | 2.087876 | 0.036809037 | NA |
| *SRSF3* | 0.140006 | 0.06706 | 2.087794 | 0.036816443 | 0.258316078 |
| *HAUS1* | -0.262684 | 0.125907 | -2.086339 | 0.036947942 | 0.259054862 |
| *RNF149* | 0.216502 | 0.103789 | 2.085987 | 0.036979794 | 0.25909443 |
| *ARFGAP3* | 0.176499 | 0.084628 | 2.085597 | 0.037015166 | 0.259158588 |
| *CLNS1A* | 0.35781 | 0.171656 | 2.084465 | 0.037117857 | 0.259693653 |
| *SYNGR1* | 1.032011 | 0.495351 | 2.083392 | 0.037215534 | 0.260192905 |
| *EFNA4* | 0.423789 | 0.203529 | 2.082209 | 0.037323415 | 0.260762741 |
| *NXT1* | 0.176074 | 0.084593 | 2.081423 | 0.037395228 | 0.260778355 |
| *CIB1* | 0.166744 | 0.080122 | 2.081127 | 0.037422268 | 0.260778355 |
| *RARA.AS1* | -0.529721 | 0.254542 | -2.081074 | 0.037427169 | 0.260778355 |
| *MAFK* | 0.478782 | 0.23007 | 2.08103 | 0.037431164 | 0.260778355 |
| *LRMP* | 0.298334 | 0.143423 | 2.080101 | 0.037516227 | 0.261186913 |
| *ELOB* | -0.170842 | 0.082165 | -2.079259 | 0.037593529 | 0.261367276 |
| *TP53RK* | 0.28936 | 0.139187 | 2.078926 | 0.037624179 | 0.261367276 |
| *FCGR2B* | 0.600635 | 0.288947 | 2.078701 | 0.037644805 | 0.261367276 |
| *RNASEK* | -0.307572 | 0.147966 | -2.078668 | 0.037647886 | 0.261367276 |
| *SUSD3* | 0.733858 | 0.353137 | 2.07811 | 0.037699267 | 0.26145725 |
| *TRAPPC1* | -0.14131 | 0.068004 | -2.077952 | 0.03771374 | 0.26145725 |
| *ENSG00000166927* | -0.659751 | 0.317697 | -2.07667 | 0.037831996 | 0.262093282 |
| *LITAF* | 0.273098 | 0.131535 | 2.076239 | 0.037871862 | 0.262185736 |
| *NDEL1* | 0.294826 | 0.142057 | 2.0754 | 0.037949509 | 0.262361642 |
| *RBM7* | 0.210594 | 0.101478 | 2.075264 | 0.03796208 | 0.262361642 |

| | | | | | |
|---|---|---|---|---|---|
| *MICOS10* | 0.126242 | 0.060844 | 2.074845 | 0.038000936 | 0.262361642 |
| *AGPAT2* | -0.26798 | 0.129158 | -2.074818 | 0.038003426 | 0.262361642 |
| *YTHDF1* | 0.395035 | 0.190511 | 2.073557 | 0.038120484 | 0.26292579 |
| *TPT1.AS1* | -0.28914 | 0.139459 | -2.073299 | 0.038144435 | 0.26292579 |
| *CD274* | 1.258626 | 0.607133 | 2.073065 | 0.038166203 | NA |
| *ERCC1* | 0.456646 | 0.220325 | 2.072599 | 0.038209606 | 0.26292579 |
| *SF3A3* | 0.217368 | 0.104878 | 2.072577 | 0.038211699 | 0.26292579 |
| *ADIPOR2* | 0.291118 | 0.140466 | 2.072508 | 0.038218123 | 0.26292579 |
| *FAM216A* | 0.483647 | 0.233434 | 2.071876 | 0.038277045 | 0.26314803 |
| *ENSG00000263531* | -0.993429 | 0.479552 | -2.071577 | 0.038304934 | NA |
| *POLR2G* | -0.193344 | 0.093339 | -2.071408 | 0.038320683 | 0.263264955 |
| *PLD6* | 1.448186 | 0.699232 | 2.07111 | 0.038348538 | NA |
| *CWC25* | 0.19946 | 0.096327 | 2.070668 | 0.038389839 | 0.26329839 |
| *RAB11A* | -0.149824 | 0.072367 | -2.070345 | 0.038420088 | 0.26329839 |
| *LGALS3BP* | -2.088372 | 1.008837 | -2.070079 | 0.038444972 | NA |
| *MIR29B2CHG* | 0.399485 | 0.193006 | 2.069805 | 0.038470589 | 0.26329839 |
| *IKBKB* | -0.278738 | 0.134677 | -2.06968 | 0.038482293 | 0.26329839 |
| *HGS* | -0.243697 | 0.117748 | -2.069649 | 0.038485198 | 0.26329839 |
| *MRTFB* | -0.589349 | 0.284758 | -2.069648 | 0.03848535 | 0.26329839 |
| *CTPS1* | 0.413225 | 0.199689 | 2.069339 | 0.038514271 | 0.263314028 |
| *QDPR* | 0.746213 | 0.360702 | 2.068778 | 0.03856696 | 0.263362109 |
| *TRNT1* | 0.235321 | 0.113753 | 2.068696 | 0.038574584 | 0.263362109 |
| *C11orf98* | -0.508485 | 0.245852 | -2.068257 | 0.038615835 | 0.263461797 |
| *ENSG00000260274* | -1.125561 | 0.54438 | -2.067601 | 0.038677531 | NA |
| *IL21R* | -0.333394 | 0.161267 | -2.067335 | 0.038702618 | 0.263741433 |
| *ECE1* | -0.316057 | 0.152887 | -2.067255 | 0.038710178 | 0.263741433 |
| *NOB1* | 0.269662 | 0.130493 | 2.066491 | 0.038782131 | 0.263938602 |
| *CAPZB* | -0.133255 | 0.06449 | -2.066268 | 0.038803137 | 0.263938602 |
| *G3BP1* | 0.212739 | 0.102967 | 2.066098 | 0.038819212 | 0.263938602 |
| *SNX2* | -0.205286 | 0.099412 | -2.065001 | 0.03892287 | 0.26446151 |
| *NOL8* | 0.271769 | 0.131628 | 2.064668 | 0.0389544 | 0.264493955 |
| *SRP54* | 0.218617 | 0.105905 | 2.064281 | 0.038991087 | 0.264561352 |
| *PALLD* | -1.497115 | 0.725548 | -2.063427 | 0.039072095 | NA |
| *KIF5B* | 0.193042 | 0.093587 | 2.062715 | 0.039139757 | 0.265387955 |
| *PIK3CB* | -0.613522 | 0.29748 | -2.062397 | 0.039169959 | 0.265396516 |
| *RANGRF* | -0.256484 | 0.124378 | -2.062137 | 0.039194711 | 0.265396516 |
| *ZNF43* | -0.286593 | 0.139016 | -2.061588 | 0.039246976 | 0.265412336 |
| *RBM4* | -0.226407 | 0.109824 | -2.061548 | 0.039250742 | 0.265412336 |
| *LINC01535* | -2.126877 | 1.032261 | -2.060406 | 0.039359763 | NA |
| *GABARAP* | -0.290407 | 0.140956 | -2.060275 | 0.03937222 | 0.265951563 |
| *PCTP* | -0.523009 | 0.25387 | -2.060149 | 0.03938429 | 0.265951563 |
| *CNKSR1* | 1.376373 | 0.668249 | 2.059671 | 0.039430027 | NA |
| *SH2D2A* | 1.18506 | 0.575609 | 2.058792 | 0.039514186 | 0.26664658 |
| *PXMP2* | -0.429745 | 0.208913 | -2.057054 | 0.039681072 | 0.267590093 |
| *NDUFA3* | -0.307345 | 0.149485 | -2.056027 | 0.039779959 | 0.268074077 |
| *FUS* | 0.15948 | 0.077591 | 2.055403 | 0.039840121 | 0.268286744 |
| *MPC2* | -0.212119 | 0.103214 | -2.055137 | 0.039865793 | 0.268286744 |

| | | | | | |
|---|---|---|---|---|---|
| *FAHD2B* | -1.119184 | 0.5451 | -2.053171 | 0.040055974 | NA |
| *CFL1* | -0.117452 | 0.057207 | -2.053087 | 0.040064139 | 0.269438148 |
| *DHX33* | 0.439714 | 0.21428 | 2.052056 | 0.040164201 | 0.269927457 |
| *LAMTOR1* | -0.165291 | 0.080578 | -2.051335 | 0.040234327 | 0.270215051 |
| *ENSG00000077463* | -0.219468 | 0.107028 | -2.050554 | 0.04031039 | 0.270475321 |
| *ENSG00000272221* | 0.812066 | 0.396173 | 2.049776 | 0.040386285 | 0.270475321 |
| *SLC41A3* | -0.356236 | 0.1738 | -2.049695 | 0.040394245 | 0.270475321 |
| *LGMN* | 0.383443 | 0.187093 | 2.049483 | 0.040414874 | 0.270475321 |
| *B3GNT2* | 0.382399 | 0.1866 | 2.049298 | 0.040432956 | 0.270475321 |
| *PREX1* | 0.497337 | 0.242692 | 2.049254 | 0.040437237 | 0.270475321 |
| *HESX1* | -2.001203 | 0.976604 | -2.049145 | 0.040447957 | NA |
| *SACS* | 0.411861 | 0.201031 | 2.04875 | 0.040486581 | 0.270524681 |
| *ENSG00000130749* | -0.277767 | 0.135587 | -2.048619 | 0.040499345 | 0.270524681 |
| *LSM10* | -0.202204 | 0.098725 | -2.048164 | 0.040543972 | 0.270639907 |
| *NR6A1* | -0.486724 | 0.237718 | -2.047483 | 0.040610696 | 0.270878667 |
| *FEM1C* | 0.460038 | 0.224711 | 2.04724 | 0.04063454 | 0.270878667 |
| *NSMAF* | 0.507071 | 0.24775 | 2.046706 | 0.040686949 | 0.270881224 |
| *IFNAR1* | -0.166318 | 0.081263 | -2.046656 | 0.040691914 | 0.270881224 |
| *FCGRT* | -0.401164 | 0.196065 | -2.046081 | 0.040748437 | 0.270881224 |
| *GART* | 0.301032 | 0.147129 | 2.046036 | 0.040752796 | 0.270881224 |
| *MRPL2* | -0.192711 | 0.094207 | -2.045613 | 0.040794509 | 0.270881224 |
| *SOCS4* | 0.378165 | 0.184871 | 2.045564 | 0.040799327 | 0.270881224 |
| *LINC02158* | -0.964832 | 0.471747 | -2.045231 | 0.040832072 | NA |
| *HIC1* | 0.756974 | 0.370353 | 2.043924 | 0.040961067 | 0.271768741 |
| *ENSG00000230138* | -0.677373 | 0.331452 | -2.043652 | 0.040987982 | 0.271768741 |
| *MOSPD3* | -0.365622 | 0.17902 | -2.042353 | 0.041116529 | 0.272438341 |
| *TUFT1* | 0.847785 | 0.415197 | 2.041886 | 0.041162871 | 0.272562724 |
| *SUPT7L* | -0.309734 | 0.151715 | -2.041558 | 0.041195384 | 0.272595426 |
| *ATF4* | 0.232126 | 0.113803 | 2.039715 | 0.04137875 | 0.273356143 |
| *TCL1A* | -0.688422 | 0.337529 | -2.039595 | 0.041390661 | 0.273356143 |
| *SPAG7* | -0.152642 | 0.074841 | -2.039569 | 0.041393298 | 0.273356143 |
| *ARPC3* | -0.120761 | 0.059232 | -2.038801 | 0.041469917 | 0.273679304 |
| *CCS* | -0.253197 | 0.124246 | -2.037869 | 0.041563058 | 0.273785188 |
| *EXOC2* | -0.337752 | 0.165739 | -2.03785 | 0.041564965 | 0.273785188 |
| *IKZF1* | -0.35819 | 0.175772 | -2.037809 | 0.041569044 | 0.273785188 |
| *MFSD2A* | 0.658919 | 0.323526 | 2.036682 | 0.041681923 | 0.274335444 |
| *PSMA4* | -0.128825 | 0.06326 | -2.036421 | 0.041708089 | 0.274335444 |
| *ZNF564* | -0.534714 | 0.26265 | -2.035844 | 0.041766007 | 0.274533737 |
| *TFAM* | 0.244306 | 0.12003 | 2.035382 | 0.041812491 | 0.274656666 |
| *GTF2IRD2B* | -0.571457 | 0.280834 | -2.03486 | 0.041864916 | 0.274818433 |
| *SFI1* | -0.303165 | 0.149019 | -2.0344 | 0.041911312 | 0.274940431 |
| *TBRG4* | 0.258954 | 0.127317 | 2.033928 | 0.041958901 | 0.275070087 |
| *SPN* | -1.161106 | 0.570936 | -2.033688 | 0.041983087 | NA |
| *ENSG00000262580* | 1.571579 | 0.772968 | 2.033175 | 0.042034799 | NA |
| *AHI1* | -0.309027 | 0.152054 | -2.032358 | 0.042117437 | 0.275926433 |
| *SYNGAP1.AS1* | 1.074637 | 0.528952 | 2.031635 | 0.042190598 | NA |
| *PADI4* | -1.42105 | 0.700473 | -2.028702 | 0.042488628 | NA |

| | | | | | |
|---|---|---|---|---|---|
| *PAQR4* | -1.847922 | 0.911077 | -2.028283 | 0.04253139 | NA |
| *PLEC* | -0.435024 | 0.214483 | -2.028248 | 0.042534916 | 0.278476939 |
| *ARFRP1* | -0.216381 | 0.106716 | -2.027636 | 0.04259745 | 0.278542235 |
| *NMB* | 0.825288 | 0.407027 | 2.027599 | 0.04260124 | 0.278542235 |
| *NAA20* | 0.149495 | 0.073745 | 2.027189 | 0.042643074 | 0.27863148 |
| *FKBP3* | -0.166794 | 0.082301 | -2.02664 | 0.042699263 | 0.278811339 |
| *KLHDC3* | -0.234971 | 0.11596 | -2.026316 | 0.042732402 | 0.278811339 |
| *PIK3CG* | -0.647883 | 0.31977 | -2.026093 | 0.042755208 | 0.278811339 |
| *NKRF* | 0.432512 | 0.213617 | 2.024711 | 0.04289709 | 0.279552166 |
| *PFKP* | -0.453646 | 0.224135 | -2.023987 | 0.042971479 | 0.27972708 |
| *C4orf46* | -0.457214 | 0.225936 | -2.023645 | 0.043006648 | 0.27972708 |
| *ZBTB1* | 0.401665 | 0.198488 | 2.023624 | 0.043008817 | 0.27972708 |
| *CCT6A* | 0.229614 | 0.113492 | 2.023175 | 0.043055069 | 0.279843796 |
| *MKNK2* | -0.33837 | 0.167289 | -2.022672 | 0.043107018 | 0.279997357 |
| *MAPKAP1* | -0.239579 | 0.118473 | -2.022215 | 0.043154132 | 0.280016468 |
| *SMAD4* | 0.329805 | 0.163101 | 2.022094 | 0.043166609 | 0.280016468 |
| *TSPYL1* | -0.237818 | 0.117656 | -2.021298 | 0.043248916 | 0.280366417 |
| *EWSR1* | 0.251324 | 0.124484 | 2.018921 | 0.043495467 | 0.281779939 |
| *NUP35* | 0.324255 | 0.160636 | 2.018571 | 0.043531781 | 0.281830507 |
| *SLC11A1* | 2.009457 | 0.995867 | 2.017796 | 0.043612555 | NA |
| *NOP14* | 0.376801 | 0.186751 | 2.017668 | 0.043625835 | 0.282254585 |
| *COL19A1* | -0.429471 | 0.21295 | -2.016769 | 0.043719615 | 0.282676334 |
| *DOCK7* | -0.646593 | 0.320662 | -2.016431 | 0.043754964 | 0.282718437 |
| *TAF2* | 0.311128 | 0.154317 | 2.016159 | 0.043783323 | 0.282718437 |
| *ENSG00000130811* | 0.119271 | 0.059196 | 2.014841 | 0.043921363 | 0.283312042 |
| *VAPB* | 0.260044 | 0.129088 | 2.014468 | 0.043960378 | 0.283312042 |
| *DDX60L* | -0.454591 | 0.225674 | -2.014371 | 0.043970578 | 0.283312042 |
| *MCTS1* | 0.333683 | 0.165666 | 2.014187 | 0.043989883 | 0.283312042 |
| *CEP89* | -0.303305 | 0.150612 | -2.013819 | 0.044028551 | 0.283352037 |
| *S100PBP* | -0.34294 | 0.170353 | -2.013112 | 0.044102825 | 0.283352037 |
| *STK17A* | 0.256427 | 0.127405 | 2.012696 | 0.044146598 | 0.283352037 |
| *ENSG00000268713* | 0.661715 | 0.328803 | 2.0125 | 0.044167214 | 0.283352037 |
| *UBE4B* | 0.263975 | 0.131181 | 2.012292 | 0.04418912 | 0.283352037 |
| *HSPA1A* | 1.405214 | 0.698318 | 2.012282 | 0.04419023 | 0.283352037 |
| *CBLN3* | -0.778164 | 0.386719 | -2.01222 | 0.044196727 | 0.283352037 |
| *PRSS21* | 1.392564 | 0.692175 | 2.011868 | 0.044233819 | NA |
| *ZBTB11* | 0.308675 | 0.153476 | 2.011228 | 0.044301421 | 0.283644296 |
| *ABLIM1* | -0.263731 | 0.131145 | -2.010989 | 0.044326648 | 0.283644296 |
| *DOLPP1* | 0.728258 | 0.36217 | 2.010816 | 0.044344875 | 0.283644296 |
| *MAP3K12* | -0.663104 | 0.329788 | -2.010701 | 0.044357079 | 0.283644296 |
| *ABCB1* | 0.468912 | 0.233448 | 2.008634 | 0.044576 | 0.284859947 |
| *KLHL5* | -0.396431 | 0.197425 | -2.008006 | 0.044642661 | 0.284975708 |
| *ZNF493* | -0.301566 | 0.150188 | -2.00792 | 0.044651767 | 0.284975708 |
| *MAPKAPK5* | 0.233694 | 0.11645 | 2.00682 | 0.04476887 | 0.285538744 |
| *C16orf86* | -0.916028 | 0.456608 | -2.00616 | 0.044839153 | NA |
| *TMEM183A* | -0.172913 | 0.086203 | -2.005892 | 0.044867799 | 0.285985207 |
| *ZBTB24* | -0.251594 | 0.125473 | -2.005164 | 0.044945563 | 0.286038783 |

| | | | | | |
|---|---|---|---|---|---|
| *DAZAP1* | -0.319052 | 0.159116 | -2.005149 | 0.044947123 | 0.286038783 |
| *ACADVL* | -0.229658 | 0.114545 | -2.004959 | 0.044967438 | 0.286038783 |
| *PELI3* | -0.879604 | 0.438785 | -2.004635 | 0.0450021 | NA |
| *EBPL* | -0.684647 | 0.341546 | -2.004551 | 0.045011095 | 0.286038783 |
| *TMEM67* | -0.500505 | 0.249702 | -2.004412 | 0.045025988 | 0.286038783 |
| *SECISBP2* | 0.245747 | 0.122617 | 2.004189 | 0.045049806 | 0.286038783 |
| *CARMIL1* | -1.354252 | 0.675874 | -2.003705 | 0.045101699 | 0.286083149 |
| *SOX5* | 1.532821 | 0.765016 | 2.003645 | 0.045108089 | NA |
| *AGO1* | -0.279586 | 0.139545 | -2.003558 | 0.045117401 | 0.286083149 |
| *GCHFR* | -0.329644 | 0.164558 | -2.003205 | 0.045155295 | 0.286083149 |
| *POMC* | -0.497997 | 0.248642 | -2.002864 | 0.04519191 | 0.286083149 |
| *VARS* | 0.466805 | 0.233102 | 2.002578 | 0.045222643 | 0.286083149 |
| *FUBP1* | 0.267605 | 0.133635 | 2.002505 | 0.045230423 | 0.286083149 |
| *COLEC12* | -0.852368 | 0.425932 | -2.001182 | 0.045372782 | NA |
| *LY6E* | 0.375423 | 0.187622 | 2.000959 | 0.045396802 | 0.286883279 |
| *DTWD1* | 0.231429 | 0.115669 | 2.000791 | 0.045414964 | 0.286883279 |
| *ABCD1* | -0.915219 | 0.457446 | -2.000715 | 0.045423151 | NA |
| *TMEM115* | 0.210536 | 0.105251 | 2.000329 | 0.045464798 | 0.287014684 |
| *TXNL4B* | 0.289599 | 0.144804 | 1.999941 | 0.045506679 | 0.287095743 |
| *RBM43* | -0.739303 | 0.369841 | -1.998976 | 0.045610902 | 0.287569757 |
| *ENSG00000274265* | -0.359474 | 0.179885 | -1.998355 | 0.045678141 | 0.287810136 |
| *CD7* | -3.958443 | 1.981178 | -1.998024 | NA | NA |
| *ATP5F1EP2* | -0.328432 | 0.164478 | -1.996812 | 0.045845617 | 0.288557208 |
| *KPTN* | -0.600094 | 0.300539 | -1.996725 | 0.045855085 | 0.288557208 |
| *GLO1* | 0.243746 | 0.122112 | 1.996091 | 0.045924025 | 0.288710651 |
| *ZNF226* | -0.262834 | 0.131696 | -1.995765 | 0.045959551 | 0.288710651 |
| *SGO2* | -0.4494 | 0.225185 | -1.995695 | 0.045967081 | 0.288710651 |
| *ENSG00000277825* | -1.240192 | 0.621797 | -1.994528 | 0.046094394 | NA |
| *SELENOI* | 0.521369 | 0.261445 | 1.994181 | 0.046132327 | 0.289181026 |
| *ASH1L.AS1* | -0.556324 | 0.279018 | -1.993865 | 0.046166801 | 0.289181026 |
| *ENSG00000277511* | -0.583822 | 0.292814 | -1.993837 | 0.046169909 | 0.289181026 |
| *ALKBH2* | 0.446685 | 0.224037 | 1.993796 | 0.046174322 | 0.289181026 |
| *BNIP2* | -0.226939 | 0.11383 | -1.993669 | 0.04618823 | 0.289181026 |
| *PCNX4* | 0.280813 | 0.140877 | 1.993313 | 0.046227198 | 0.289241825 |
| *GTPBP4* | 0.367056 | 0.184218 | 1.992511 | 0.046315017 | 0.289608005 |
| *KLHL8* | 0.411593 | 0.206643 | 1.991811 | 0.046391845 | 0.289905047 |
| *HNRNPD* | 0.203728 | 0.102314 | 1.991198 | 0.046459145 | 0.289972299 |
| *NUP155* | -0.374686 | 0.188173 | -1.991178 | 0.046461271 | 0.289972299 |
| *TLR1* | -0.338097 | 0.169891 | -1.990077 | 0.046582443 | 0.290545131 |
| *ENSG00000273759* | -1.214043 | 0.610187 | -1.989625 | 0.04663227 | NA |
| *IFI35* | -0.31581 | 0.158747 | -1.989388 | 0.046658414 | 0.290638546 |
| *NOL10* | 0.235213 | 0.118238 | 1.989313 | 0.046666708 | 0.290638546 |
| *CEP128* | -0.784995 | 0.39464 | -1.989141 | 0.046685617 | 0.290638546 |
| *PRPF40B* | -0.687368 | 0.345712 | -1.988268 | 0.046782051 | 0.290874955 |
| *ENSG00000273619* | -1.212648 | 0.609902 | -1.988267 | 0.046782132 | NA |
| *SLC17A9* | 0.668022 | 0.335982 | 1.988265 | 0.046782438 | 0.290874955 |
| *SCML1* | 0.549742 | 0.276603 | 1.987476 | 0.046869668 | 0.291061619 |

| | | | | | |
|---|---|---|---|---|---|
| *MSANTD4* | 0.334078 | 0.168093 | 1.987461 | 0.046871343 | 0.291061619 |
| *C2orf76* | -0.302783 | 0.152368 | -1.987184 | 0.046902025 | 0.291069318 |
| *ZNF132* | 1.219827 | 0.613987 | 1.986731 | 0.046952224 | NA |
| *DYNLT3* | -0.201425 | 0.101387 | -1.986694 | 0.046956279 | 0.291223195 |
| *IKZF3* | 0.16951 | 0.085344 | 1.9862 | 0.047011089 | 0.29138033 |
| *TPM1* | 0.580774 | 0.292498 | 1.985565 | 0.047081643 | 0.291634787 |
| *GGCX* | -0.352396 | 0.17756 | -1.98466 | 0.047182268 | 0.291959526 |
| *WEE1* | 0.49893 | 0.251415 | 1.98449 | 0.04720121 | 0.291959526 |
| *SNHG10* | -0.441687 | 0.222602 | -1.984202 | 0.047233376 | 0.291959526 |
| *SAMD9* | -0.260885 | 0.131492 | -1.984033 | 0.047252199 | 0.291959526 |
| *CCDC28A* | -0.287178 | 0.144773 | -1.983646 | 0.047295278 | 0.292043175 |
| *SHROOM3* | -2.279327 | 1.149289 | -1.98325 | 0.04733948 | NA |
| *CRLF3* | 0.234174 | 0.11812 | 1.982505 | 0.047422752 | 0.292504102 |
| *MGST2* | -1.204259 | 0.6075 | -1.982317 | 0.047443739 | NA |
| *ZNF502* | -0.603565 | 0.304552 | -1.981814 | 0.047500102 | 0.292504102 |
| *GRN* | -0.310413 | 0.156636 | -1.981749 | 0.047507326 | 0.292504102 |
| *DELE1* | -0.321433 | 0.162209 | -1.981593 | 0.047524849 | 0.292504102 |
| *PLEKHB2* | 0.213172 | 0.107584 | 1.981454 | 0.047540377 | 0.292504102 |
| *FAM200B* | -0.328176 | 0.165646 | -1.981193 | 0.047569689 | 0.292504102 |
| *C11orf71* | -0.446279 | 0.225265 | -1.981127 | 0.047577038 | 0.292504102 |
| *MAP4K5* | 0.323114 | 0.163183 | 1.980076 | 0.047694996 | 0.293040121 |
| *RPA2* | 0.146453 | 0.073973 | 1.979822 | 0.047723507 | 0.293040121 |
| *BORCS5* | 0.40652 | 0.205405 | 1.979115 | 0.047803008 | 0.293346084 |
| *C1orf109* | 0.302175 | 0.152732 | 1.978467 | 0.047876029 | 0.293611923 |
| *POLR3F* | 0.368517 | 0.18631 | 1.977971 | 0.047931993 | 0.293772895 |
| *PLEKHG2* | 0.872735 | 0.441437 | 1.977031 | 0.048038087 | 0.294240725 |
| *CCT4* | 0.20185 | 0.102123 | 1.976546 | 0.04809298 | 0.294394553 |
| *AP1S1* | -0.289188 | 0.146365 | -1.975797 | 0.048177785 | 0.294628952 |
| *UBE2T* | 0.563556 | 0.285246 | 1.975681 | 0.048190877 | 0.294628952 |
| *LINC01597* | -1.192254 | 0.603574 | -1.975323 | 0.048231529 | NA |
| *UQCRB* | -0.120694 | 0.061112 | -1.974965 | 0.048272094 | 0.294836402 |
| *HLA.F* | -0.250229 | 0.126707 | -1.974856 | 0.048284456 | 0.294836402 |
| *LCMT1.AS1* | 1.481025 | 0.750222 | 1.974116 | 0.048368547 | NA |
| *C1orf112* | -0.763788 | 0.386903 | -1.97411 | 0.048369258 | 0.295162932 |
| *U2AF1L4* | -0.263336 | 0.133412 | -1.97386 | 0.048397644 | 0.295162932 |
| *KCMF1* | 0.315641 | 0.159992 | 1.972859 | 0.048511609 | 0.295675564 |
| *ENSG00000236935* | 0.899521 | 0.456305 | 1.971317 | 0.048687621 | 0.296461654 |
| *HNRNPF* | 0.228374 | 0.115869 | 1.970965 | 0.048727833 | 0.296461654 |
| *MGAT1* | 0.19165 | 0.097238 | 1.970942 | 0.048730547 | 0.296461654 |
| *IL2RA* | -0.456854 | 0.231984 | -1.969332 | 0.048914985 | 0.297312802 |
| *METTL15* | -0.288796 | 0.146689 | -1.968761 | 0.048980516 | 0.297312802 |
| *MRPL34* | -0.22649 | 0.115047 | -1.968683 | 0.048989562 | 0.297312802 |
| *RSRP1* | 0.264484 | 0.134347 | 1.968672 | 0.04899075 | 0.297312802 |
| *PLCE1* | -0.282948 | 0.143832 | -1.967205 | 0.049159569 | 0.297988604 |
| *RPP38* | 0.202456 | 0.102917 | 1.967181 | 0.049162393 | 0.297988604 |
| *ENSG00000231609* | -1.015367 | 0.516528 | -1.965753 | 0.049327114 | NA |
| *CD247* | -2.32587 | 1.183413 | -1.965392 | 0.049368863 | NA |

| | | | | | |
|---|---|---|---|---|---|
| *HTATIP2* | 0.224313 | 0.114136 | 1.96532 | 0.049377247 | 0.299107515 |
| *RASSF5* | -0.195172 | 0.099325 | -1.964979 | 0.049416665 | 0.299121516 |
| *KIF16B* | -0.553866 | 0.281908 | -1.964705 | 0.049448375 | 0.299121516 |
| *RSBN1* | 0.22981 | 0.116982 | 1.964491 | 0.049473175 | 0.299121516 |
| *CFAP54* | 1.699606 | 0.86528 | 1.964226 | 0.04950389 | NA |
| *YPEL3* | -0.250508 | 0.127541 | -1.964144 | 0.049513389 | 0.299121516 |
| *KATNA1* | 0.269825 | 0.137386 | 1.963994 | 0.049530844 | 0.299121516 |
| *POLR1C* | 0.331428 | 0.168829 | 1.963093 | 0.049635381 | 0.299569828 |
| *ENSG00000268400* | -0.967054 | 0.492741 | -1.962602 | 0.049692476 | 0.299731432 |
| *MNDA* | 1.736916 | 0.885673 | 1.961127 | 0.049864212 | NA |
| *ENSG00000230555* | -1.026782 | 0.523691 | -1.960663 | 0.04991832 | NA |

| Cluster 10 | log2FC | lfcSE | stat | pvalue | padj |
|---|---|---|---|---|---|
| *LEF1* | -3.277376 | 0.470469 | -6.966191 | 3.26E-12 | 3.54E-08 |
| *SMIM10* | -2.838875 | 0.413627 | -6.863377 | 6.73E-12 | 3.66E-08 |
| *POU2F2* | 0.4511 | 0.071072 | 6.347059 | 2.19E-10 | 6.77E-07 |
| *SESN3* | 0.564108 | 0.089628 | 6.293883 | 3.10E-10 | 6.77E-07 |
| *TMEM123* | -0.644281 | 0.102377 | -6.293245 | 3.11E-10 | 6.77E-07 |
| *CST7* | -4.156998 | 0.664327 | -6.257454 | 3.91E-10 | 7.10E-07 |
| *ACAP1* | 0.563093 | 0.091739 | 6.138009 | 8.36E-10 | 1.06E-06 |
| *MIF* | -0.612702 | 0.099896 | -6.133411 | 8.60E-10 | 1.06E-06 |
| *ARL4C* | -1.322753 | 0.215755 | -6.130801 | 8.74E-10 | 1.06E-06 |
| *DNAJB1* | 0.8305 | 0.139241 | 5.964473 | 2.45E-09 | 2.67E-06 |
| *BIN1* | 0.599304 | 0.105109 | 5.701719 | 1.19E-08 | 1.17E-05 |
| *KLK4* | 2.735244 | 0.481551 | 5.680067 | 1.35E-08 | 1.22E-05 |
| *RAB31* | 1.79837 | 0.322444 | 5.577306 | 2.44E-08 | 2.05E-05 |
| *FAM102A* | 1.04648 | 0.190936 | 5.480778 | 4.23E-08 | 3.08E-05 |
| *MACROD2* | -0.983326 | 0.179427 | -5.480355 | 4.24E-08 | 3.08E-05 |
| *KRCC1* | 0.473217 | 0.08688 | 5.446771 | 5.13E-08 | 3.20E-05 |
| *ATP6V1C2* | -1.827083 | 0.33565 | -5.443419 | 5.23E-08 | 3.20E-05 |
| *CDC42EP3* | 1.038097 | 0.19078 | 5.441325 | 5.29E-08 | 3.20E-05 |
| *FCRL4* | 1.588379 | 0.293878 | 5.404894 | 6.48E-08 | 3.71E-05 |
| *APBB2* | -1.928416 | 0.363193 | -5.30962 | 1.10E-07 | 5.70E-05 |
| *GATD3A* | 4.160742 | 0.783649 | 5.309447 | 1.10E-07 | 5.70E-05 |
| *SLC50A1* | -0.565696 | 0.110325 | -5.127539 | 2.94E-07 | 0.000143736 |
| *ZNF322* | 0.846105 | 0.165219 | 5.12111 | 3.04E-07 | 0.000143736 |
| *HHIP.AS1* | -3.065624 | 0.600415 | -5.105843 | 3.29E-07 | 0.000149348 |
| *PCIF1* | -0.608118 | 0.119519 | -5.088036 | 3.62E-07 | 0.000157509 |
| *HOMER2* | -1.276196 | 0.252408 | -5.056087 | 4.28E-07 | 0.000177221 |
| *CIRBP* | -0.344716 | 0.068325 | -5.045268 | 4.53E-07 | 0.000177221 |
| *SRGN* | -0.574961 | 0.114095 | -5.039311 | 4.67E-07 | 0.000177221 |
| *ENSG00000272211* | -1.017528 | 0.202 | -5.037278 | 4.72E-07 | 0.000177221 |
| *RSRP1* | 0.501517 | 0.099776 | 5.026419 | 5.00E-07 | 0.000181299 |
| *FTL* | 0.359395 | 0.072357 | 4.966944 | 6.80E-07 | 0.000238803 |
| *SCRN1* | 1.270594 | 0.25674 | 4.94895 | 7.46E-07 | 0.000253784 |
| *MRPL54* | -0.427416 | 0.087079 | -4.908344 | 9.18E-07 | 0.000302933 |
| *SWAP70* | 0.299062 | 0.061665 | 4.849815 | 1.24E-06 | 0.000395591 |
| *ADA* | -0.64187 | 0.134141 | -4.785045 | 1.71E-06 | 0.000531602 |
| *TTC32* | -0.83229 | 0.177982 | -4.676253 | 2.92E-06 | 0.00086433 |
| *PHF23* | 0.502801 | 0.107549 | 4.675088 | 2.94E-06 | 0.00086433 |
| *RUBCNL* | -0.513987 | 0.110385 | -4.656326 | 3.22E-06 | 0.000921994 |
| *GART* | -0.489767 | 0.105712 | -4.633011 | 3.60E-06 | 0.001005753 |
| *LPIN1* | 0.611573 | 0.132172 | 4.627105 | 3.71E-06 | 0.00100898 |
| *CBFA2T3* | 1.217315 | 0.264884 | 4.595652 | 4.31E-06 | 0.001145202 |
| *LDHA* | -0.410841 | 0.089721 | -4.579085 | 4.67E-06 | 0.001185692 |
| *TMEM170A* | 0.454331 | 0.099288 | 4.575866 | 4.74E-06 | 0.001185692 |
| *CCDC191* | -1.220477 | 0.267116 | -4.569091 | 4.90E-06 | 0.001185692 |
| *TNFRSF13B* | 1.421134 | 0.311057 | 4.56873 | 4.91E-06 | 0.001185692 |
| *HMOX1* | 1.113125 | 0.244073 | 4.560632 | 5.10E-06 | 0.001185692 |

| | | | | | |
|---|---|---|---|---|---|
| *NCBP3* | 0.81316 | 0.178464 | 4.556426 | 5.20E-06 | 0.001185692 |
| *BHLHE41* | 1.787357 | 0.392362 | 4.555381 | 5.23E-06 | 0.001185692 |
| *SYNGR2* | -0.401018 | 0.088213 | -4.546019 | 5.47E-06 | 0.001214343 |
| *CRK* | 0.552476 | 0.121901 | 4.532191 | 5.84E-06 | 0.001270709 |
| *SERTAD3* | 0.604573 | 0.134151 | 4.506662 | 6.59E-06 | 0.00140543 |
| *RCC1* | 0.441141 | 0.097979 | 4.502409 | 6.72E-06 | 0.001406286 |
| *CAV1* | -0.758524 | 0.168976 | -4.488943 | 7.16E-06 | 0.001469902 |
| *KIAA0040* | 0.633006 | 0.143127 | 4.422684 | 9.75E-06 | 0.001916346 |
| *APAF1* | 1.126218 | 0.254767 | 4.420581 | 9.84E-06 | 0.001916346 |
| *DEDD2* | 0.468402 | 0.105992 | 4.419223 | 9.91E-06 | 0.001916346 |
| *DBI* | -0.360214 | 0.081629 | -4.412792 | 1.02E-05 | 0.001916346 |
| *ID3* | 0.396926 | 0.089952 | 4.412634 | 1.02E-05 | 0.001916346 |
| *ABRACL* | -0.497587 | 0.113155 | -4.397384 | 1.10E-05 | 0.002013896 |
| *NAB2* | 0.627813 | 0.142863 | 4.394516 | 1.11E-05 | 0.002013896 |
| *PLGRKT* | -0.586997 | 0.133919 | -4.383226 | 1.17E-05 | 0.002070129 |
| *JMJD6* | 0.589244 | 0.134488 | 4.381392 | 1.18E-05 | 0.002070129 |
| *PARK7* | -0.25979 | 0.059791 | -4.344949 | 1.39E-05 | 0.002398771 |
| *GID8* | -0.354723 | 0.081692 | -4.342217 | 1.41E-05 | 0.002398771 |
| *IL27RA* | 0.592806 | 0.136702 | 4.336498 | 1.45E-05 | 0.002424136 |
| *CHCHD5* | -0.406058 | 0.093832 | -4.327499 | 1.51E-05 | 0.00245418 |
| *KCTD7* | 0.959004 | 0.221627 | 4.327115 | 1.51E-05 | 0.00245418 |
| *ENSG00000253701* | 1.078406 | 0.249945 | 4.314579 | 1.60E-05 | 0.002559455 |
| *MAML3* | -1.585732 | 0.370282 | -4.282496 | 1.85E-05 | 0.002915146 |
| *MIR29B2CHG* | 0.656564 | 0.153586 | 4.274897 | 1.91E-05 | 0.002973288 |
| *RNF43* | -1.124725 | 0.264223 | -4.256729 | 2.07E-05 | 0.003179959 |
| *HIP1R* | -0.451171 | 0.106575 | -4.23337 | 2.30E-05 | 0.003480084 |
| *MT1X* | -0.733507 | 0.173469 | -4.228454 | 2.35E-05 | 0.003508269 |
| *MAP3K8* | -0.574588 | 0.136166 | -4.21976 | 2.45E-05 | 0.003539832 |
| *HPRT1* | -0.363316 | 0.086108 | -4.2193 | 2.45E-05 | 0.003539832 |
| *H1FX* | -1.31288 | 0.311304 | -4.217362 | 2.47E-05 | 0.003539832 |
| *KRTCAP2* | -0.301749 | 0.071689 | -4.209137 | 2.56E-05 | 0.003579605 |
| *LMBRD1* | -0.392894 | 0.093347 | -4.208975 | 2.57E-05 | 0.003579605 |
| *ADI1* | -0.357616 | 0.085524 | -4.181494 | 2.90E-05 | 0.003989874 |
| *GLUL* | -0.596497 | 0.143014 | -4.170889 | 3.03E-05 | 0.004127944 |
| *IL21R* | -0.751451 | 0.180553 | -4.161935 | 3.16E-05 | 0.004212585 |
| *SUSD3* | 0.881508 | 0.211869 | 4.160626 | 3.17E-05 | 0.004212585 |
| *SNHG9* | -0.419864 | 0.101178 | -4.149768 | 3.33E-05 | 0.004277747 |
| *SIAH2* | -0.571151 | 0.137719 | -4.14723 | 3.37E-05 | 0.004277747 |
| *CFAP73* | -1.173217 | 0.283068 | -4.144647 | 3.40E-05 | 0.004277747 |
| *TMEM120A* | -0.642795 | 0.155276 | -4.139697 | 3.48E-05 | 0.004277747 |
| *CHMP6* | -0.48437 | 0.117006 | -4.139686 | 3.48E-05 | 0.004277747 |
| *ASPSCR1* | -0.457915 | 0.110651 | -4.13838 | 3.50E-05 | 0.004277747 |
| *ENSG00000226571* | 1.310699 | 0.31672 | 4.13836 | 3.50E-05 | 0.004277747 |
| *FIS1* | -0.321783 | 0.078016 | -4.124578 | 3.71E-05 | 0.004440432 |
| *ASF1A* | 0.528484 | 0.128161 | 4.123595 | 3.73E-05 | 0.004440432 |
| *ARID5A* | -0.533026 | 0.129347 | -4.120906 | 3.77E-05 | 0.004440432 |
| *WASHC2C* | 0.603819 | 0.14657 | 4.119668 | 3.79E-05 | 0.004440432 |

| | | | | | |
|---|---|---|---|---|---|
| *FCGR2B* | 1.048985 | 0.25579 | 4.100957 | 4.11E-05 | 0.00476401 |
| *STX7* | 0.340396 | 0.083491 | 4.077064 | 4.56E-05 | 0.005225232 |
| *LACC1* | -1.428536 | 0.350698 | -4.073403 | 4.63E-05 | 0.005252788 |
| *CD5* | -2.145451 | 0.528118 | -4.062445 | 4.86E-05 | 0.005410952 |
| *RNF34* | 0.513099 | 0.126327 | 4.061682 | 4.87E-05 | 0.005410952 |
| *DUS4L* | -0.699471 | 0.173469 | -4.032244 | 5.52E-05 | 0.006073805 |
| *GALNT2* | -0.727716 | 0.180724 | -4.02668 | 5.66E-05 | 0.006157082 |
| *SP110* | 0.41821 | 0.104009 | 4.020908 | 5.80E-05 | 0.006229706 |
| *TOB2* | 0.401639 | 0.099929 | 4.019257 | 5.84E-05 | 0.006229706 |
| *LCN8* | 4.124227 | 1.027611 | 4.013411 | 5.98E-05 | 0.006324082 |
| *NINJ1* | -0.642947 | 0.160316 | -4.010508 | 6.06E-05 | 0.006340799 |
| *GBGT1* | 0.844982 | 0.212169 | 3.982598 | 6.82E-05 | 0.007065908 |
| *ENSG00000283013* | -0.686143 | 0.173259 | -3.960222 | 7.49E-05 | 0.00766786 |
| *EIF2AK2* | 0.398931 | 0.100775 | 3.958626 | 7.54E-05 | 0.00766786 |
| *DDAH2* | 0.563955 | 0.143222 | 3.937636 | 8.23E-05 | 0.008292817 |
| *GRHPR* | -0.517961 | 0.131772 | -3.930745 | 8.47E-05 | 0.008455868 |
| *TIMM10* | -0.521002 | 0.132812 | -3.922841 | 8.75E-05 | 0.00861942 |
| *TPP1* | 0.373897 | 0.095339 | 3.921759 | 8.79E-05 | 0.00861942 |
| *ATF2* | 0.597069 | 0.152735 | 3.909192 | 9.26E-05 | 0.008999243 |
| *MED21* | 0.507331 | 0.130924 | 3.874996 | 0.00010663 | 0.010270095 |
| *ENSG00000273319* | 0.723996 | 0.187195 | 3.867594 | 0.00010991 | 0.010493919 |
| *LINC00638* | 2.615354 | 0.677791 | 3.858646 | 0.00011402 | NA |
| *EIF4A2* | 0.172071 | 0.044775 | 3.843006 | 0.00012154 | 0.011410479 |
| *PPP1R18* | 0.352145 | 0.091636 | 3.842855 | 0.00012161 | 0.011410479 |
| *ATP1B3* | -0.362639 | 0.094586 | -3.833941 | 0.00012611 | 0.011731121 |
| *WDFY1* | 0.571664 | 0.149572 | 3.822012 | 0.00013237 | 0.01220922 |
| *LRIF1* | 0.55601 | 0.145769 | 3.814312 | 0.00013656 | 0.012389798 |
| *MX2* | 1.128416 | 0.295843 | 3.814241 | 0.0001366 | 0.012389798 |
| *CYB5R3* | 0.376574 | 0.098913 | 3.807101 | 0.00014061 | 0.01264753 |
| *ARL17A* | -0.507676 | 0.13344 | -3.804526 | 0.00014208 | 0.012675022 |
| *CDCA4* | 0.473937 | 0.124731 | 3.799662 | 0.00014489 | 0.012821316 |
| *RPL8* | 0.222603 | 0.05868 | 3.793478 | 0.00014855 | 0.013039018 |
| *CYB5A* | -0.441237 | 0.116508 | -3.787192 | 0.00015236 | 0.013266257 |
| *MRPL23* | -0.310776 | 0.082139 | -3.783528 | 0.00015462 | 0.013356279 |
| *NME1* | -0.297206 | 0.078669 | -3.777931 | 0.00015814 | 0.013552427 |
| *SRP14.AS1* | 0.64525 | 0.171287 | 3.767078 | 0.00016517 | 0.013969781 |
| *ZNRD2* | -0.339556 | 0.090152 | -3.766468 | 0.00016557 | 0.013969781 |
| *GCSH* | -0.455329 | 0.121068 | -3.760919 | 0.00016929 | 0.014173523 |
| *STRAP* | -0.276573 | 0.073596 | -3.757989 | 0.00017128 | 0.014202314 |
| *TGOLN2* | 0.307672 | 0.081905 | 3.756453 | 0.00017234 | 0.014202314 |
| *CPM* | -1.184342 | 0.315429 | -3.7547 | 0.00017355 | 0.014202314 |
| *SERP2* | -1.646412 | 0.439616 | -3.745113 | 0.00018031 | 0.0146457 |
| *DOCK8* | 0.335985 | 0.08981 | 3.741086 | 0.00018323 | 0.014772158 |
| *ORAI2* | 0.28864 | 0.077262 | 3.735874 | 0.00018706 | 0.014880448 |
| *HIST1H4C* | 0.278735 | 0.074617 | 3.735551 | 0.0001873 | 0.014880448 |
| *PIP5K1B* | -0.735708 | 0.19707 | -3.733231 | 0.00018904 | 0.014909403 |
| *CHORDC1* | 0.414949 | 0.111266 | 3.729355 | 0.00019197 | 0.014950666 |

| | | | | | |
|---|---|---|---|---|---|
| *SGPP1* | 1.006149 | 0.269824 | 3.728911 | 0.00019231 | 0.014950666 |
| *ZNF217* | 0.538212 | 0.144879 | 3.714909 | 0.00020328 | 0.015691236 |
| *IMPACT* | -0.596928 | 0.161575 | -3.694435 | 0.00022038 | 0.016723228 |
| *HMCES* | -0.895007 | 0.242267 | -3.694293 | 0.0002205 | 0.016723228 |
| *BCAT1* | -0.751113 | 0.203365 | -3.693423 | 0.00022126 | 0.016723228 |
| *ZBTB20* | 0.45887 | 0.124583 | 3.683253 | 0.00023028 | 0.017285 |
| *ELOB* | -0.171579 | 0.04661 | -3.681157 | 0.00023218 | 0.01730842 |
| *OSBPL8* | 0.366561 | 0.099728 | 3.675599 | 0.00023729 | 0.017569304 |
| *SPDL1* | 0.719033 | 0.196592 | 3.657483 | 0.0002547 | 0.018731067 |
| *NAGK* | 0.41231 | 0.11284 | 3.653938 | 0.00025825 | 0.018864304 |
| *PIGQ* | -0.574912 | 0.158039 | -3.637793 | 0.00027498 | 0.01995284 |
| *HLA.A* | -0.487569 | 0.13451 | -3.624786 | 0.0002892 | 0.020845414 |
| *SRPK1* | -0.328468 | 0.09082 | -3.616686 | 0.0002984 | 0.021366967 |
| *SNX8* | -0.446884 | 0.123711 | -3.612317 | 0.00030347 | 0.021588277 |
| *TCF4* | 0.539482 | 0.149453 | 3.609699 | 0.00030655 | 0.021665686 |
| *PDE7B* | -1.258658 | 0.349355 | -3.602808 | 0.0003148 | 0.021875174 |
| *SERPINB6* | 0.691573 | 0.191977 | 3.602371 | 0.00031533 | 0.021875174 |
| *ARF5* | -0.376263 | 0.104454 | -3.602191 | 0.00031555 | 0.021875174 |
| *NFKBIA* | -0.399197 | 0.111018 | -3.595777 | 0.00032343 | 0.022279456 |
| *LILRB1* | 0.569778 | 0.158618 | 3.592144 | 0.00032797 | 0.022445818 |
| *MLST8* | -0.405681 | 0.112985 | -3.590563 | 0.00032996 | 0.022445818 |
| *NTHL1* | -0.46736 | 0.13037 | -3.584868 | 0.00033725 | 0.022798878 |
| *XXYLT1.AS2* | 1.671067 | 0.466732 | 3.580359 | 0.00034312 | 0.023052748 |
| *RDX* | 0.507969 | 0.142349 | 3.568466 | 0.00035908 | 0.023976644 |
| *PRCP* | -0.475806 | 0.133711 | -3.558462 | 0.00037303 | 0.024756644 |
| *PHGDH* | -1.138611 | 0.320125 | -3.556773 | 0.00037544 | 0.024765287 |
| *RUVBL2* | -0.357687 | 0.101009 | -3.541153 | 0.00039838 | 0.026120408 |
| *ENSG00000279483* | -0.37212 | 0.105171 | -3.538237 | 0.00040281 | 0.026252489 |
| *PMS2* | -0.496263 | 0.140516 | -3.531733 | 0.00041285 | 0.026596437 |
| *NDUFB10* | -0.276813 | 0.078381 | -3.531652 | 0.00041297 | 0.026596437 |
| *CLCF1* | 0.552388 | 0.156611 | 3.527139 | 0.00042008 | 0.026894775 |
| *RGS19* | 0.367913 | 0.104401 | 3.52403 | 0.00042504 | 0.027053211 |
| *SOX4* | -0.844678 | 0.240823 | -3.507457 | 0.00045241 | 0.028465332 |
| *DAZAP1* | -0.393787 | 0.112277 | -3.507279 | 0.00045272 | 0.028465332 |
| *SKAP2* | 0.337699 | 0.096338 | 3.505351 | 0.00045601 | 0.028465332 |
| *POLR2I* | -0.282294 | 0.080555 | -3.504373 | 0.00045768 | 0.028465332 |
| *WDR11* | -0.366921 | 0.104836 | -3.499941 | 0.00046536 | 0.028778341 |
| *SSPN* | 2.104394 | 0.60192 | 3.496138 | 0.00047205 | 0.029026752 |
| *MRPL44* | 0.406948 | 0.116504 | 3.492992 | 0.00047764 | 0.029205869 |
| *LINC02245* | -0.648651 | 0.186277 | -3.482182 | 0.00049735 | 0.030082548 |
| *LRWD1* | 0.636316 | 0.182739 | 3.482095 | 0.00049751 | 0.030082548 |
| *CD53* | 0.267677 | 0.076965 | 3.477892 | 0.00050537 | 0.030389463 |
| *CARMIL1* | -1.613174 | 0.464413 | -3.473574 | 0.00051358 | 0.030712923 |
| *PAM* | -1.173035 | 0.33791 | -3.471438 | 0.00051768 | 0.030789153 |
| *ENSG00000263394* | 1.324308 | 0.382038 | 3.466428 | 0.00052742 | 0.031198237 |
| *ADAMTS6* | 1.001567 | 0.28911 | 3.464315 | 0.00053158 | 0.031274391 |
| *ZC3H6* | 0.695359 | 0.200941 | 3.460506 | 0.00053916 | 0.031431822 |

| | | | | | |
|---|---|---|---|---|---|
| *SNAPC1* | 0.542533 | 0.156798 | 3.46007 | 0.00054004 | 0.031431822 |
| *IKZF3* | 0.283912 | 0.082131 | 3.456808 | 0.00054662 | 0.031645515 |
| *HDDC2* | -0.27251 | 0.078931 | -3.452498 | 0.00055542 | 0.031985264 |
| *TUNAR* | -2.844977 | 0.824074 | -3.452331 | 0.00055577 | NA |
| *KLHL6* | 0.59585 | 0.172759 | 3.449024 | 0.00056262 | 0.032158821 |
| *LGALS1* | 1.081508 | 0.313724 | 3.447325 | 0.00056617 | 0.032158821 |
| *NRP2* | -0.882205 | 0.25595 | -3.446785 | 0.0005673 | 0.032158821 |
| *TNFRSF12A* | 1.229474 | 0.357137 | 3.442579 | 0.0005762 | 0.032422967 |
| *ARRDC3* | -0.398424 | 0.115761 | -3.441772 | 0.00057792 | 0.032422967 |
| *PALLD* | -2.006014 | 0.582944 | -3.441179 | 0.00057919 | NA |
| *PDGFA* | 1.164959 | 0.340099 | 3.425353 | 0.000614 | 0.034270724 |
| *UCP2* | -0.380483 | 0.111185 | -3.422052 | 0.0006215 | 0.034512471 |
| *METRNL* | 1.146097 | 0.33545 | 3.416594 | 0.0006341 | 0.03503308 |
| *ACADVL* | -0.260936 | 0.076431 | -3.413995 | 0.00064018 | 0.035190361 |
| *MIR22HG* | 0.82831 | 0.242999 | 3.408693 | 0.00065275 | 0.035701158 |
| *ZBTB16* | 2.984238 | 0.876424 | 3.405018 | 0.0006616 | NA |
| *UBE2E3* | -0.29012 | 0.085287 | -3.401702 | 0.00066968 | 0.036189787 |
| *BOLA2B* | -0.550829 | 0.161954 | -3.401143 | 0.00067105 | 0.036189787 |
| *CKS2* | 0.384342 | 0.113012 | 3.400894 | 0.00067166 | 0.036189787 |
| *MYO1F* | 0.721188 | 0.212621 | 3.391896 | 0.00069411 | 0.037215075 |
| *PNPLA8* | 0.41256 | 0.121725 | 3.389274 | 0.00070078 | 0.037388715 |
| *RBM18* | 0.426689 | 0.125995 | 3.386546 | 0.00070779 | 0.0375782 |
| *TCL1B* | -1.454291 | 0.430222 | -3.380326 | 0.000724 | 0.038252431 |
| *CLEC2D* | -0.367385 | 0.108847 | -3.375242 | 0.00073751 | 0.038449568 |
| *MME* | -2.427719 | 0.719397 | -3.37466 | 0.00073907 | 0.038449568 |
| *ENSG00000273748* | -0.970034 | 0.287806 | -3.370443 | 0.00075047 | 0.038449568 |
| *ENSG00000271133* | -1.172809 | 0.347985 | -3.370291 | 0.00075089 | 0.038449568 |
| *RASSF1* | 0.433225 | 0.128557 | 3.369913 | 0.00075192 | 0.038449568 |
| *HIKESHI* | -0.300314 | 0.089118 | -3.369828 | 0.00075215 | 0.038449568 |
| *GIT2* | 0.292208 | 0.086716 | 3.369715 | 0.00075246 | 0.038449568 |
| *POLE3* | -0.325387 | 0.096655 | -3.366495 | 0.0007613 | 0.038719584 |
| *MED31* | 0.485503 | 0.144505 | 3.359761 | 0.0007801 | 0.039340533 |
| *DYNLT1* | 0.366542 | 0.109127 | 3.358856 | 0.00078266 | 0.039340533 |
| *RANBP1* | -0.384017 | 0.11435 | -3.358258 | 0.00078435 | 0.039340533 |
| *BLVRB* | 0.601073 | 0.179239 | 3.353475 | 0.00079804 | 0.039843198 |
| *B3GNT2* | 0.636379 | 0.18996 | 3.350069 | 0.00080791 | 0.040152238 |
| *PHKG1* | -0.510924 | 0.152745 | -3.344948 | 0.00082298 | 0.04071511 |
| *ABI1* | 0.281673 | 0.084272 | 3.342406 | 0.00083056 | 0.040903934 |
| *CD180* | 0.688933 | 0.206233 | 3.340556 | 0.00083611 | 0.040991837 |
| *TTYH3* | -0.762129 | 0.228448 | -3.33612 | 0.00084957 | 0.041408734 |
| *MAN1B1* | 0.496102 | 0.148754 | 3.335053 | 0.00085283 | 0.041408734 |
| *TSPAN18* | 1.658886 | 0.497605 | 3.333738 | 0.00085687 | 0.041408734 |
| *VANGL2* | -2.408766 | 0.72262 | -3.333377 | 0.00085799 | NA |
| *CLDN15* | 0.603652 | 0.181126 | 3.33278 | 0.00085983 | 0.041408734 |
| *AEBP1* | -0.773865 | 0.232792 | -3.324276 | 0.00088648 | 0.04242464 |
| *TSEN15* | -0.393745 | 0.11847 | -3.323574 | 0.00088872 | 0.04242464 |
| *PFKL* | -0.329829 | 0.099313 | -3.321126 | 0.00089655 | 0.042552718 |

| | | | | | |
|---|---|---|---|---|---|
| *LMO4* | 0.628586 | 0.18935 | 3.319706 | 0.00090112 | 0.042552718 |
| *GINM1* | 0.382032 | 0.115102 | 3.319084 | 0.00090313 | 0.042552718 |
| *ENSG00000170161* | -0.590855 | 0.178117 | -3.317223 | 0.00090917 | 0.042652669 |
| *IFI30* | 0.489469 | 0.147711 | 3.313682 | 0.00092076 | 0.043010955 |
| *TSN* | -0.314937 | 0.095091 | -3.311965 | 0.00092643 | 0.04309094 |
| *ADRB2* | 1.444305 | 0.436454 | 3.30918 | 0.0009357 | 0.043336653 |
| *CREBRF* | 0.428199 | 0.129588 | 3.304317 | 0.00095208 | 0.043908671 |
| *PAQR8* | 0.965309 | 0.292722 | 3.297704 | 0.00097479 | 0.044766182 |
| *FLCN* | 0.515815 | 0.156663 | 3.292513 | 0.00099296 | 0.045409284 |
| *UBE2J2* | -0.279344 | 0.084905 | -3.29007 | 0.00100162 | 0.045613699 |
| *ENSG00000228463* | 0.588979 | 0.179232 | 3.286123 | 0.00101577 | 0.046064998 |
| *MRPL40* | -0.286567 | 0.087246 | -3.284598 | 0.00102128 | 0.046122867 |
| *ENSG00000244459* | 0.916276 | 0.279618 | 3.276889 | 0.00104958 | 0.047204926 |
| *PPM1K* | 0.36092 | 0.110531 | 3.265319 | 0.00109341 | 0.048605485 |
| *IDH1* | 0.371797 | 0.113886 | 3.264642 | 0.00109603 | 0.048605485 |
| *TMEM273* | 1.328272 | 0.406976 | 3.263762 | 0.00109944 | 0.048605485 |
| *AIM2* | 1.829987 | 0.560822 | 3.263044 | 0.00110223 | 0.048605485 |
| *ATM* | 0.265042 | 0.081242 | 3.262391 | 0.00110477 | 0.048605485 |
| *PTPMT1* | -0.332222 | 0.101856 | -3.261687 | 0.00110751 | 0.048605485 |
| *GCFC2* | -0.419894 | 0.128798 | -3.260106 | 0.0011137 | 0.048680951 |
| *ZNF580* | -0.329866 | 0.101218 | -3.258964 | 0.0011182 | 0.048681818 |
| *ANAPC15* | -0.32864 | 0.100949 | -3.255513 | 0.00113188 | 0.049001128 |
| *ARL4D* | 1.222175 | 0.375494 | 3.254847 | 0.00113454 | 0.049001128 |
| *SPINT2* | -0.333962 | 0.102747 | -3.250333 | 0.0011527 | 0.049588817 |
| *RALGPS2* | 0.340866 | 0.104958 | 3.247634 | 0.00116369 | 0.0498645 |
| *IGHG3* | 1.294635 | 0.399006 | 3.244648 | 0.00117596 | 0.050192756 |
| *ZNF506* | 0.531242 | 0.163899 | 3.241284 | 0.00118993 | 0.05059045 |
| *DDB2* | -0.357473 | 0.110355 | -3.239301 | 0.00119823 | 0.050620779 |
| *CXCR5* | 0.303144 | 0.093595 | 3.238894 | 0.00119994 | 0.050620779 |
| *LRRC61* | -0.823771 | 0.254506 | -3.236751 | 0.00120899 | 0.050770718 |
| *HCST* | 0.647922 | 0.20029 | 3.234917 | 0.00121678 | 0.050770718 |
| *NHP2* | -0.219483 | 0.067852 | -3.234751 | 0.00121749 | 0.050770718 |
| *TERF2IP* | 0.305621 | 0.094535 | 3.232888 | 0.00122546 | 0.050889493 |
| *GAPT* | 0.353617 | 0.109415 | 3.231903 | 0.00122969 | 0.050889493 |
| *FKBP11* | -0.345394 | 0.107002 | -3.22792 | 0.00124694 | 0.051407894 |
| *TBXAS1* | 1.475507 | 0.458056 | 3.221238 | 0.00127638 | 0.052423073 |
| *PEMT* | -0.558684 | 0.173535 | -3.219437 | 0.00128443 | 0.052555225 |
| *TSEN2* | -0.54239 | 0.168845 | -3.212354 | 0.00131652 | 0.053666745 |
| *TSC22D4* | 0.288565 | 0.089883 | 3.210458 | 0.00132523 | 0.053820342 |
| *TOP2B* | -0.326787 | 0.102057 | -3.202018 | 0.00136468 | 0.055216454 |
| *GLS* | 0.352735 | 0.11037 | 3.195945 | 0.00139373 | 0.056182983 |
| *CAPG* | 0.629508 | 0.197314 | 3.190387 | 0.00142082 | 0.0569359 |
| *NAALADL1* | 0.538607 | 0.168844 | 3.18997 | 0.00142287 | 0.0569359 |
| *IKZF2* | -1.147974 | 0.359997 | -3.188845 | 0.00142842 | 0.056948606 |
| *MLX* | 0.317424 | 0.099643 | 3.185616 | 0.00144446 | 0.057377906 |
| *CENPS.CORT* | -1.597934 | 0.501643 | -3.185398 | 0.00144555 | NA |
| *GAMT* | -0.430922 | 0.135383 | -3.182982 | 0.00145767 | 0.057474981 |

| | | | | | |
|---|---|---|---|---|---|
| *MAN1A1* | -0.498369 | 0.156612 | -3.182181 | 0.0014617 | 0.057474981 |
| *GPR18* | 0.69022 | 0.216916 | 3.181974 | 0.00146275 | 0.057474981 |
| *LYST* | -0.680412 | 0.21396 | -3.180095 | 0.00147227 | 0.057640917 |
| *EIF2AK4* | -0.569315 | 0.179108 | -3.178606 | 0.00147985 | 0.0577301 |
| *APOL6* | 0.928926 | 0.292481 | 3.17602 | 0.00149311 | 0.05803926 |
| *INKA1* | 0.816122 | 0.257101 | 3.174321 | 0.00150188 | 0.058172297 |
| *KLK1* | 1.738649 | 0.548122 | 3.172013 | 0.00151386 | 0.058428696 |
| *TMED8* | 0.383947 | 0.121319 | 3.164769 | 0.00155206 | 0.059691241 |
| *LINC01869* | 1.102404 | 0.348789 | 3.16066 | 0.00157412 | 0.060326547 |
| *POLD2* | -0.321993 | 0.102009 | -3.156502 | 0.00159674 | 0.060978649 |
| *SRSF5* | -0.316044 | 0.100177 | -3.154871 | 0.00160569 | 0.061105999 |
| *ENSG00000173821* | 0.327994 | 0.104018 | 3.153242 | 0.00161468 | 0.061233921 |
| *FKBP1A* | -0.242199 | 0.07684 | -3.151983 | 0.00162166 | 0.061285205 |
| *DAPL1* | -0.646932 | 0.205442 | -3.148979 | 0.00163842 | 0.061338813 |
| *SEPTIN7* | 0.215355 | 0.06839 | 3.148917 | 0.00163877 | 0.061338813 |
| *G2E3* | 0.478877 | 0.152087 | 3.1487 | 0.00163999 | 0.061338813 |
| *CHD1* | 0.280883 | 0.08933 | 3.144343 | 0.0016646 | 0.062046336 |
| *FBXO10* | 0.766051 | 0.243824 | 3.141814 | 0.00167905 | 0.062371085 |
| *CD200R1* | 2.968525 | 0.945343 | 3.140155 | 0.00168858 | NA |
| *USF1* | -0.737922 | 0.23521 | -3.137286 | 0.0017052 | 0.063127147 |
| *CCND3* | 0.255384 | 0.081476 | 3.134457 | 0.00172173 | 0.063303322 |
| *PDCD4* | 0.504245 | 0.160875 | 3.134391 | 0.00172211 | 0.063303322 |
| *TRIM4* | 0.397666 | 0.126908 | 3.13349 | 0.00172741 | 0.063303322 |
| *PLEKHF2* | 0.349142 | 0.111505 | 3.131173 | 0.00174109 | 0.063590793 |
| *XIAP* | 0.388681 | 0.124293 | 3.127137 | 0.00176518 | 0.064089999 |
| *IRF5* | 0.536665 | 0.171665 | 3.126241 | 0.00177057 | 0.064089999 |
| *SLC39A8* | -0.622535 | 0.199152 | -3.125932 | 0.00177243 | 0.064089999 |
| *MED4* | -0.243699 | 0.078083 | -3.121022 | 0.00180224 | 0.064952412 |
| *GPR146* | -1.109994 | 0.356254 | -3.115739 | 0.00183485 | 0.065590426 |
| *PRDX2* | -0.325956 | 0.104645 | -3.114887 | 0.00184015 | 0.065590426 |
| *NAE1* | -0.332721 | 0.106863 | -3.113526 | 0.00184867 | 0.065590426 |
| *MRPS36* | 0.1952 | 0.062709 | 3.112763 | 0.00185345 | 0.065590426 |
| *ZNF391* | 1.813362 | 0.5827 | 3.111999 | 0.00185825 | 0.065590426 |
| *CDKN1B* | 0.344224 | 0.110633 | 3.111396 | 0.00186205 | 0.065590426 |
| *NDUFB5* | 0.214915 | 0.069074 | 3.111383 | 0.00186213 | 0.065590426 |
| *SIGLEC10* | 0.907879 | 0.292722 | 3.101501 | 0.00192542 | 0.067601032 |
| *ENSG00000261448* | 1.015254 | 0.328428 | 3.091257 | 0.00199311 | 0.069729316 |
| *ENSG00000282988* | 1.043358 | 0.337654 | 3.090021 | 0.00200143 | 0.069729316 |
| *BAZ2B* | -0.539688 | 0.174724 | -3.088807 | 0.00200962 | 0.069729316 |
| *CGAS* | 0.434233 | 0.140596 | 3.088504 | 0.00201167 | 0.069729316 |
| *ING5* | -0.392356 | 0.127091 | -3.087215 | 0.00202041 | 0.069810085 |
| *ATP6V0E2* | -0.758153 | 0.245928 | -3.082822 | 0.00205048 | 0.070447527 |
| *CPNE2* | -1.007628 | 0.326873 | -3.082629 | 0.00205181 | 0.070447527 |
| *MXD1* | 0.923766 | 0.299921 | 3.080027 | 0.00206982 | 0.070842454 |
| *LINC01588* | 0.753389 | 0.244699 | 3.078839 | 0.00207809 | 0.070902684 |
| *GEMIN8* | -0.508848 | 0.16535 | -3.077389 | 0.00208823 | 0.070920494 |
| *SOCS3* | 0.841574 | 0.273513 | 3.076901 | 0.00209165 | 0.070920494 |

| | | | | | |
|---|---|---|---|---|---|
| *MORC3* | 0.416089 | 0.135307 | 3.075137 | 0.00210406 | 0.071085225 |
| *ITM2A* | 1.106756 | 0.359996 | 3.074357 | 0.00210957 | 0.071085225 |
| *YWHAG* | 0.339718 | 0.110673 | 3.069563 | 0.00214372 | 0.071449468 |
| *GTF2B* | 0.414608 | 0.135113 | 3.068604 | 0.00215062 | 0.071449468 |
| *NRAS* | -0.384654 | 0.125391 | -3.067645 | 0.00215752 | 0.071449468 |
| *GFOD2* | 0.49015 | 0.159807 | 3.067131 | 0.00216124 | 0.071449468 |
| *TMEM208* | -0.229162 | 0.074717 | -3.067081 | 0.0021616 | 0.071449468 |
| *FCRL2* | 0.558254 | 0.182021 | 3.066979 | 0.00216234 | 0.071449468 |
| *DHX57* | -0.509999 | 0.166353 | -3.065771 | 0.00217109 | 0.071449468 |
| *PTPRC* | 0.259718 | 0.084722 | 3.065523 | 0.00217289 | 0.071449468 |
| *SEL1L3* | 0.678428 | 0.221527 | 3.062509 | 0.0021949 | 0.071955562 |
| *NUFIP2* | 0.459699 | 0.150241 | 3.059751 | 0.00221521 | 0.072046605 |
| *SPN* | -0.760901 | 0.248706 | -3.05944 | 0.00221751 | 0.072046605 |
| *CAPN12* | 0.545266 | 0.178224 | 3.059437 | 0.00221753 | 0.072046605 |
| *CDK2AP1* | -1.018562 | 0.33343 | -3.054798 | 0.00225212 | 0.072872947 |
| *AHDC1* | 0.605646 | 0.198297 | 3.054234 | 0.00225636 | 0.072872947 |
| *CASP3* | -0.328064 | 0.10762 | -3.048363 | 0.00230092 | 0.073939094 |
| *TPI1* | -0.227463 | 0.074625 | -3.048097 | 0.00230295 | 0.073939094 |
| *IFI27L2* | 0.384482 | 0.12635 | 3.042984 | 0.00234245 | 0.074817482 |
| *TXNDC17* | -0.309225 | 0.10165 | -3.042046 | 0.00234976 | 0.074817482 |
| *IL2RA* | -0.610469 | 0.200687 | -3.041895 | 0.00235094 | 0.074817482 |
| *CD109* | -1.099535 | 0.361595 | -3.040794 | 0.00235955 | 0.074872838 |
| *LINC02422* | 0.876445 | 0.288589 | 3.037003 | 0.00238943 | 0.075600573 |
| *ACP5* | 0.346548 | 0.114318 | 3.031449 | 0.00243383 | 0.076485254 |
| *DRAM1* | -0.682549 | 0.225172 | -3.031234 | 0.00243556 | 0.076485254 |
| *IGHG2* | 0.827481 | 0.273017 | 3.030873 | 0.00243848 | 0.076485254 |
| *POLR2B* | -0.247149 | 0.081612 | -3.028343 | 0.00245899 | 0.076906989 |
| *SNAI3* | 0.82452 | 0.272359 | 3.027323 | 0.0024673 | 0.076945782 |
| *RNF146* | 0.421161 | 0.139355 | 3.022214 | 0.00250933 | 0.078033114 |
| *BAZ1A* | 0.220574 | 0.073044 | 3.019731 | 0.00252999 | 0.078451317 |
| *PHLPP1* | -0.722619 | 0.23943 | -3.018078 | 0.00254383 | 0.078656382 |
| *SLAMF7* | 0.56021 | 0.185883 | 3.013772 | 0.00258022 | 0.07955546 |
| *CPSF6* | -0.285945 | 0.095043 | -3.008598 | 0.00262456 | 0.080694196 |
| *MRNIP* | 0.435426 | 0.144773 | 3.00764 | 0.00263285 | 0.080721022 |
| *POLR2J* | -0.190177 | 0.063279 | -3.005357 | 0.0026527 | 0.081100924 |
| *AKAP12* | -1.417331 | 0.471873 | -3.003626 | 0.00266784 | 0.081335332 |
| *ENSG00000237499* | -0.541161 | 0.180339 | -3.000796 | 0.00269275 | 0.081865695 |
| *HNRNPDL* | -0.162754 | 0.054351 | -2.994498 | 0.00274897 | 0.083341944 |
| *YIPF5* | 0.347507 | 0.116211 | 2.990312 | 0.00278693 | 0.084258044 |
| *CTSC* | 0.547221 | 0.183199 | 2.98703 | 0.00281702 | 0.084403678 |
| *KMT2C* | 0.288281 | 0.096535 | 2.986296 | 0.00282379 | 0.084403678 |
| *MPRIP.AS1* | 0.922887 | 0.309181 | 2.98494 | 0.00283634 | 0.084403678 |
| *NRROS* | -0.434109 | 0.145441 | -2.984786 | 0.00283777 | 0.084403678 |
| *MNDA* | 2.512035 | 0.842142 | 2.98291 | 0.00285522 | 0.084403678 |
| *ENSG00000277654* | -0.70333 | 0.235832 | -2.982339 | 0.00286055 | 0.084403678 |
| *TGM2* | 1.665306 | 0.55849 | 2.981802 | 0.00286557 | 0.084403678 |
| *SLFN11* | 0.921867 | 0.309177 | 2.981683 | 0.00286669 | 0.084403678 |

| | | | | | |
|---|---|---|---|---|---|
| *CTSA* | 0.264505 | 0.088714 | 2.981538 | 0.00286805 | 0.084403678 |
| *GPR157* | 0.984421 | 0.330187 | 2.981405 | 0.00286929 | 0.084403678 |
| *ENSG00000231856* | -0.941988 | 0.316086 | -2.98016 | 0.00288098 | 0.084519132 |
| *TM6SF1* | -1.325592 | 0.44526 | -2.97712 | 0.0029097 | NA |
| *KCNK6* | 0.444685 | 0.149402 | 2.976434 | 0.00291622 | 0.085322816 |
| *MTSS1* | -0.483069 | 0.162377 | -2.974985 | 0.00293003 | 0.085467544 |
| *HMGXB4* | -0.324587 | 0.109132 | -2.974265 | 0.00293691 | 0.085467544 |
| *NT5DC2* | -0.875488 | 0.294435 | -2.97345 | 0.00294472 | 0.085467544 |
| *ALOX5AP* | -0.844113 | 0.284061 | -2.971588 | 0.00296264 | 0.085758838 |
| *MAGOH* | -0.259345 | 0.087347 | -2.969119 | 0.00298654 | 0.086111307 |
| *CD68* | 0.98782 | 0.332745 | 2.968699 | 0.00299064 | 0.086111307 |
| *NOTCH1* | 0.714165 | 0.240775 | 2.966104 | 0.00301599 | 0.086329364 |
| *FLOT1* | 0.441415 | 0.14882 | 2.966088 | 0.00301614 | 0.086329364 |
| *GNG11* | 0.861992 | 0.290687 | 2.965361 | 0.00302328 | 0.086329364 |
| *PRKAG1* | -0.271966 | 0.091735 | -2.964684 | 0.00302994 | 0.086329364 |
| *PAPOLA* | -0.185725 | 0.062674 | -2.963358 | 0.00304302 | 0.086475895 |
| *FANCE* | 1.313977 | 0.443847 | 2.960431 | 0.00307209 | NA |
| *PILRB* | 0.651346 | 0.22003 | 2.960257 | 0.00307383 | 0.087123716 |
| *CABLES1* | -1.028597 | 0.347762 | -2.957759 | 0.00309884 | 0.087511366 |
| *ASAH1* | 0.29492 | 0.099727 | 2.957288 | 0.00310358 | 0.087511366 |
| *NDUFB8* | -0.211612 | 0.071594 | -2.955725 | 0.00311935 | 0.087517515 |
| *STX12* | 0.344468 | 0.116545 | 2.955672 | 0.00311988 | 0.087517515 |
| *ENSG00000256092* | -0.486923 | 0.164898 | -2.952872 | 0.00314832 | 0.088074005 |
| *SNRPD3* | -0.212404 | 0.07195 | -2.95213 | 0.0031559 | 0.088074005 |
| *GATM* | 1.053087 | 0.356923 | 2.950465 | 0.00317296 | 0.088323643 |
| *ENSG00000237596* | -1.533408 | 0.520178 | -2.94785 | 0.00319993 | NA |
| *PSMA4* | -0.236314 | 0.080196 | -2.946683 | 0.00321202 | 0.089182798 |
| *ADAM28* | 0.391702 | 0.13306 | 2.943787 | 0.00324223 | 0.089354836 |
| *PPP2R5B* | 0.628186 | 0.213431 | 2.943277 | 0.00324757 | 0.089354836 |
| *ENSG00000007237* | -0.720346 | 0.244755 | -2.943132 | 0.00324909 | 0.089354836 |
| *CLPX* | -0.360856 | 0.122617 | -2.942945 | 0.00325106 | 0.089354836 |
| *AIP* | 0.333967 | 0.113669 | 2.938064 | 0.00330268 | 0.090545117 |
| *SPTSSA* | -0.28894 | 0.098393 | -2.936594 | 0.00331838 | 0.090717158 |
| *ACADM* | 0.327859 | 0.111672 | 2.935917 | 0.00332563 | 0.090717158 |
| *SNHG25* | -0.494429 | 0.168579 | -2.932914 | 0.00335797 | 0.091370258 |
| *ZNHIT3* | 0.322919 | 0.110232 | 2.92944 | 0.00339573 | 0.092164034 |
| *TNKS1BP1* | -1.460891 | 0.498878 | -2.928351 | 0.00340765 | NA |
| *TICAM1* | 0.49031 | 0.167458 | 2.927961 | 0.00341193 | 0.092164034 |
| *GPX1* | -0.241373 | 0.082439 | -2.927905 | 0.00341254 | 0.092164034 |
| *HNRNPA1L2* | -0.400936 | 0.136994 | -2.926662 | 0.00342621 | 0.092304005 |
| *ABI3* | 0.493622 | 0.1688 | 2.924302 | 0.0034523 | 0.092558662 |
| *ATP6V1A* | 0.359967 | 0.123096 | 2.924269 | 0.00345267 | 0.092558662 |
| *CD55* | -0.40689 | 0.13918 | -2.923477 | 0.00346146 | 0.092566287 |
| *SLC16A1.AS1* | 0.345749 | 0.118336 | 2.921749 | 0.00348072 | 0.092752793 |
| *ENSG00000130520* | -0.152287 | 0.052129 | -2.921324 | 0.00348547 | 0.092752793 |
| *ARHGAP44* | 1.281321 | 0.438929 | 2.919197 | 0.00350935 | 0.093160285 |
| *NCBP2AS2* | -0.266981 | 0.091513 | -2.917403 | 0.0035296 | 0.093469883 |

| | | | | | |
|---|---|---|---|---|---|
| *MRPL12* | -0.310519 | 0.106477 | -2.916303 | 0.00354207 | 0.09357248 |
| *CRTC3* | 0.332833 | 0.114193 | 2.914648 | 0.0035609 | 0.09384225 |
| *TYROBP* | 1.34313 | 0.462303 | 2.905306 | 0.00366895 | 0.096456106 |
| *LINC01857* | 0.598692 | 0.20627 | 2.902462 | 0.00370242 | 0.097101491 |
| *TOE1* | -0.450609 | 0.155332 | -2.900938 | 0.00372047 | 0.097340479 |
| *MARCHF1* | 0.59172 | 0.204314 | 2.89613 | 0.00377796 | 0.098607422 |
| *OSTM1* | 0.476005 | 0.164472 | 2.894148 | 0.00380189 | 0.098890163 |
| *THAP3* | 0.414824 | 0.143419 | 2.892389 | 0.00382325 | 0.098890163 |
| *RPRD1A* | -0.299454 | 0.103564 | -2.891481 | 0.0038343 | 0.098890163 |
| *SH3BP1* | 0.473008 | 0.163592 | 2.891385 | 0.00383549 | 0.098890163 |
| *ENSG00000173727* | 0.389867 | 0.134855 | 2.891019 | 0.00383995 | 0.098890163 |
| *ENSG00000268205* | 0.416456 | 0.144095 | 2.890156 | 0.00385051 | 0.098890163 |
| *CYB561D2* | -0.356395 | 0.12338 | -2.888593 | 0.0038697 | 0.098890163 |
| *HSPA1B* | 1.028903 | 0.356252 | 2.888128 | 0.00387542 | 0.098890163 |
| *ITSN2* | 0.236258 | 0.081814 | 2.887759 | 0.00387997 | 0.098890163 |
| *EBNA1BP2* | -0.300412 | 0.104056 | -2.887034 | 0.00388893 | 0.098890163 |
| *CD52* | 0.337835 | 0.117029 | 2.886757 | 0.00389235 | 0.098890163 |
| *FKBP3* | -0.248586 | 0.086126 | -2.886315 | 0.00389782 | 0.098890163 |
| *BCL7A* | -0.603221 | 0.209155 | -2.88409 | 0.00392546 | 0.099359863 |
| *NSUN7* | -1.205148 | 0.418289 | -2.881135 | 0.00396246 | 0.100063667 |
| *SLC39A4* | -0.304596 | 0.105835 | -2.878039 | 0.00400156 | 0.100730092 |
| *HHEX* | 0.294764 | 0.102434 | 2.877582 | 0.00400736 | 0.100730092 |
| *TFDP2* | -0.431927 | 0.150176 | -2.876133 | 0.0040258 | 0.100907058 |
| *BCAS4* | 0.460764 | 0.160274 | 2.874849 | 0.00404221 | 0.100907058 |
| *PTGIR* | -1.166665 | 0.405818 | -2.874848 | 0.00404222 | 0.100907058 |
| *CALHM2* | 1.194381 | 0.415861 | 2.872068 | 0.00407796 | 0.101566263 |
| *LPAR5* | 0.527067 | 0.183561 | 2.871343 | 0.00408732 | 0.101567053 |
| *GPR171* | -2.560546 | 0.891961 | -2.870692 | 0.00409575 | NA |
| *ENSG00000274422* | 0.538295 | 0.187595 | 2.869454 | 0.00411181 | 0.101943001 |
| *LY9* | 0.335801 | 0.117149 | 2.866442 | 0.00415114 | 0.102424692 |
| *FIBP* | -0.215919 | 0.075328 | -2.866385 | 0.00415189 | 0.102424692 |
| *IL13RA1* | -0.552111 | 0.192655 | -2.865807 | 0.00415947 | 0.102424692 |
| *ENSG00000277007* | 1.396793 | 0.487463 | 2.865433 | 0.0041644 | NA |
| *ZNF827* | 0.650009 | 0.227053 | 2.862811 | 0.00419901 | 0.103164869 |
| *SIGLEC5* | 1.531081 | 0.535134 | 2.861117 | 0.00422151 | 0.103484064 |
| *CREM* | -0.333704 | 0.1168 | -2.857058 | 0.00427588 | 0.104230081 |
| *CNR1* | 1.055477 | 0.369436 | 2.856995 | 0.00427672 | 0.104230081 |
| *YARS* | -0.300193 | 0.105098 | -2.856311 | 0.00428595 | 0.104230081 |
| *BCL10* | 0.295582 | 0.103495 | 2.855993 | 0.00429025 | 0.104230081 |
| *C12orf65* | 0.275893 | 0.096637 | 2.854946 | 0.00430442 | 0.104341431 |
| *TRIR* | 0.144698 | 0.050707 | 2.853594 | 0.00432278 | 0.104553613 |
| *PARPBP* | -0.775915 | 0.271983 | -2.852806 | 0.00433351 | 0.104580738 |
| *ADSL* | -0.234819 | 0.082366 | -2.850938 | 0.00435905 | 0.104964272 |
| *SCRN2* | -0.385334 | 0.135368 | -2.846559 | 0.00441946 | 0.105963639 |
| *IFI16* | 0.24285 | 0.085315 | 2.846518 | 0.00442002 | 0.105963639 |
| *ITGB7* | 0.625871 | 0.21995 | 2.845511 | 0.00443403 | 0.105965257 |
| *FABP5* | -0.345447 | 0.121443 | -2.844523 | 0.0044478 | 0.105965257 |

| | | | | | |
|---|---|---|---|---|---|
| *CHRAC1* | -0.249343 | 0.087665 | -2.844269 | 0.00445134 | 0.105965257 |
| *CALM3* | -0.345769 | 0.12164 | -2.842564 | 0.00447523 | 0.105965257 |
| *ENSG00000197302* | 0.346292 | 0.121843 | 2.842129 | 0.00448133 | 0.105965257 |
| *AKAP9* | 0.27198 | 0.095697 | 2.842102 | 0.00448172 | 0.105965257 |
| *LMCD1* | 2.059185 | 0.724589 | 2.841868 | NA | NA |
| *LGALS9* | -0.437763 | 0.154053 | -2.841638 | 0.00448824 | 0.105965257 |
| *CLK1* | 0.384679 | 0.135437 | 2.840288 | 0.00450728 | 0.106184573 |
| *DNAJC5B* | 1.817025 | 0.640212 | 2.838162 | 0.00453741 | 0.106663402 |
| *GTF3C6* | -0.250734 | 0.088416 | -2.83584 | 0.00457053 | 0.107210509 |
| *HAUS3* | 0.522862 | 0.184663 | 2.831444 | 0.00463383 | 0.108461621 |
| *HIST3H2A* | 1.037825 | 0.366772 | 2.829618 | 0.00466036 | 0.108706066 |
| *GALE* | 0.587611 | 0.207684 | 2.829351 | 0.00466425 | 0.108706066 |
| *CYTOR* | 0.802714 | 0.283994 | 2.82652 | 0.00470568 | 0.109437233 |
| *NDUFV3* | 0.306736 | 0.108627 | 2.82374 | 0.00474669 | 0.109914953 |
| *SENP2* | -0.376443 | 0.133344 | -2.823094 | 0.00475627 | 0.109914953 |
| *IGHG4* | 1.675629 | 0.593547 | 2.823077 | 0.00475652 | 0.109914953 |
| *SRFBP1* | 0.299359 | 0.106068 | 2.822328 | 0.00476763 | 0.109938416 |
| *TOMM5* | -0.290584 | 0.103002 | -2.821154 | 0.00478513 | 0.110108495 |
| *HOXB7* | 3.212599 | 1.139324 | 2.819741 | 0.00480625 | NA |
| *TMEM60* | 0.353257 | 0.125283 | 2.819662 | 0.00480743 | 0.11038824 |
| *SNAP23* | -0.263468 | 0.093478 | -2.81851 | 0.00482472 | 0.110551988 |
| *RGMB* | 0.969383 | 0.344159 | 2.816673 | 0.0048524 | 0.110754271 |
| *SERINC1* | 0.428549 | 0.152173 | 2.816196 | 0.0048596 | 0.110754271 |
| *SMIM4* | -0.36521 | 0.129696 | -2.815901 | 0.00486407 | 0.110754271 |
| *ADCK2* | 0.528263 | 0.187681 | 2.814689 | 0.00488245 | 0.110940692 |
| *CNIH4* | 0.284952 | 0.101408 | 2.809958 | 0.0049548 | 0.112350111 |
| *PEX10* | -0.452887 | 0.161222 | -2.809094 | 0.00496811 | 0.112417631 |
| *C1GALT1* | 0.429176 | 0.15283 | 2.808189 | 0.0049821 | 0.112500458 |
| *COX7A2L* | 0.171392 | 0.061073 | 2.806357 | 0.00501052 | 0.112712825 |
| *SKIL* | 0.383409 | 0.136627 | 2.806247 | 0.00501222 | 0.112712825 |
| *SYBU* | 2.823791 | 1.006601 | 2.805274 | 0.00502738 | NA |
| *ZNF79* | 1.20111 | 0.428472 | 2.80324 | 0.0050592 | 0.113300316 |
| *ZC3H12D* | -0.519966 | 0.185492 | -2.803168 | 0.00506033 | 0.113300316 |
| *MRPL1* | -0.232849 | 0.083122 | -2.80128 | 0.00509004 | 0.113300316 |
| *NFATC2* | 0.493118 | 0.176033 | 2.801277 | 0.00509008 | 0.113300316 |
| *PODXL2* | -0.804734 | 0.287276 | -2.801257 | 0.0050904 | 0.113300316 |
| *IVD* | 0.516398 | 0.184417 | 2.80017 | 0.00510756 | 0.11345047 |
| *IGHA1* | 0.532557 | 0.190668 | 2.793107 | 0.00522045 | 0.115721699 |
| *SNHG30* | -0.373464 | 0.133752 | -2.792212 | 0.00523491 | 0.115806451 |
| *NSMCE4A* | -0.267356 | 0.095814 | -2.790357 | 0.00526499 | 0.116107018 |
| *ARL6IP6* | 0.392486 | 0.140673 | 2.79006 | 0.00526983 | 0.116107018 |
| *HERC4* | 0.318632 | 0.11434 | 2.786698 | 0.0053248 | 0.117081122 |
| *MTHFR* | 0.442247 | 0.158862 | 2.783839 | 0.00537197 | 0.117880079 |
| *TMEM147* | -0.190314 | 0.068412 | -2.781882 | 0.00540447 | 0.118296406 |
| *MIDN* | 0.392968 | 0.141285 | 2.781389 | 0.00541268 | 0.118296406 |
| *FGFBP2* | -2.639702 | 0.949386 | -2.780432 | 0.00542867 | NA |
| *TYW3* | 0.360485 | 0.129706 | 2.779253 | 0.0054484 | 0.118838489 |

| | | | | | |
|---|---|---|---|---|---|
| *VPREB1* | -2.015525 | 0.725765 | -2.777103 | 0.00548457 | 0.119388158 |
| *HNRNPA0* | -0.179594 | 0.064749 | -2.773687 | 0.0055425 | 0.120310316 |
| *RIMKLB* | -0.595519 | 0.214733 | -2.773303 | 0.00554904 | 0.120310316 |
| *RENBP* | 0.489896 | 0.176702 | 2.772445 | 0.00556368 | 0.120387954 |
| *ZNF189* | 0.73835 | 0.266486 | 2.770685 | 0.00559385 | 0.120781503 |
| *AKAP13* | 0.266755 | 0.096298 | 2.770091 | 0.00560407 | 0.120781503 |
| *SLC2A14* | -3.578463 | 1.293368 | -2.766779 | 0.00566131 | NA |
| *MDH2* | -0.205997 | 0.074465 | -2.766375 | 0.00566833 | 0.121925109 |
| *SSR4* | -0.127114 | 0.045965 | -2.765474 | 0.00568401 | 0.122021135 |
| *SLC9B2* | -0.356629 | 0.129055 | -2.763381 | 0.00572059 | 0.122564767 |
| *DTD2* | -0.421648 | 0.1527 | -2.761293 | 0.00575729 | 0.123108772 |
| *ENSG00000228106* | -0.292012 | 0.105807 | -2.759866 | 0.00578252 | 0.123185072 |
| *ZMAT2* | -0.205612 | 0.074502 | -2.75981 | 0.0057835 | 0.123185072 |
| *UBXN6* | -0.245202 | 0.088879 | -2.758836 | 0.00580077 | 0.12322371 |
| *MICAL1* | -0.356528 | 0.12925 | -2.758431 | 0.00580795 | 0.12322371 |
| *ARHGAP25* | 0.342627 | 0.124274 | 2.757022 | 0.00583304 | 0.123289566 |
| *RNASEH2C* | -0.175438 | 0.063634 | -2.756984 | 0.00583371 | 0.123289566 |
| *CTBP1* | 0.259199 | 0.094044 | 2.756161 | 0.00584843 | 0.12336102 |
| *SMIM29* | 0.254558 | 0.092399 | 2.754995 | 0.00586931 | 0.12348711 |
| *TRIM52* | 0.424453 | 0.154123 | 2.753993 | 0.00588731 | 0.12348711 |
| *CLK3* | 0.320354 | 0.116364 | 2.753031 | 0.00590462 | 0.12348711 |
| *IL7* | 0.539462 | 0.195966 | 2.752843 | 0.00590802 | 0.12348711 |
| *ALDH16A1* | -0.400819 | 0.145634 | -2.752229 | 0.00591911 | 0.12348711 |
| *RASAL1* | -0.752294 | 0.273363 | -2.751999 | 0.00592327 | 0.12348711 |
| *FAM91A1* | 0.399644 | 0.145319 | 2.750121 | 0.00595733 | 0.12348711 |
| *RBM8A* | -0.148003 | 0.053822 | -2.749868 | 0.00596193 | 0.12348711 |
| *MTHFD2L* | -0.398312 | 0.144864 | -2.749562 | 0.0059675 | 0.12348711 |
| *IGHD* | -0.561346 | 0.20416 | -2.749542 | 0.00596786 | 0.12348711 |
| *PF4* | 3.819515 | 1.389779 | 2.74829 | 0.0059907 | 0.123660509 |
| *NDUFB7* | -0.177497 | 0.064595 | -2.747838 | 0.00599897 | 0.123660509 |
| *PPIL4* | 0.380577 | 0.138782 | 2.742272 | 0.00610158 | 0.125538036 |
| *ENTPD1* | 0.611725 | 0.22344 | 2.737757 | 0.00618598 | 0.127034284 |
| *GLIPR2* | 0.517707 | 0.189151 | 2.737012 | 0.00620001 | 0.127082645 |
| *WWOX* | -0.59747 | 0.218417 | -2.735459 | 0.00622934 | 0.127443865 |
| *MRPS34* | -0.215028 | 0.078736 | -2.730988 | 0.00631448 | 0.128779821 |
| *ITGB3BP* | -0.353334 | 0.129389 | -2.730788 | 0.0063183 | 0.128779821 |
| *A4GALT* | 1.04391 | 0.382719 | 2.727614 | 0.00637943 | 0.12978254 |
| *FXR1* | 0.206351 | 0.075721 | 2.72516 | 0.00642703 | 0.130507141 |
| *ACTN1* | 1.784665 | 0.655084 | 2.724329 | 0.00644322 | NA |
| *EIF3B* | -0.273384 | 0.100349 | -2.724325 | 0.00644331 | 0.130593963 |
| *EVI2A* | 0.392817 | 0.144243 | 2.723298 | 0.00646337 | 0.130757083 |
| *LY96* | 0.395527 | 0.145303 | 2.722085 | 0.00648714 | 0.130783268 |
| *LINC01781* | 2.015826 | 0.740566 | 2.722006 | 0.0064887 | 0.130783268 |
| *ZBTB32* | 1.428501 | 0.52543 | 2.718725 | 0.00655341 | 0.131843415 |
| *ZNF704* | -1.065388 | 0.392214 | -2.716347 | 0.00660067 | 0.132386591 |
| *GRN* | 0.249027 | 0.091711 | 2.715338 | 0.00662082 | 0.132386591 |
| *DMAC2L* | -0.244325 | 0.090001 | -2.714707 | 0.00663345 | 0.132386591 |

| | | | | | |
|---|---|---|---|---|---|
| *PIGF* | 0.440787 | 0.162438 | 2.713579 | 0.00665607 | 0.132386591 |
| *NHSL2* | -0.441899 | 0.162883 | -2.712982 | 0.00666808 | 0.132386591 |
| *ENSG00000268403* | -0.548348 | 0.202149 | -2.712598 | 0.0066758 | 0.132386591 |
| *ARID1A* | 0.252002 | 0.092903 | 2.712516 | 0.00667746 | 0.132386591 |
| *HLA.C* | -0.426295 | 0.157159 | -2.712503 | 0.00667771 | 0.132386591 |
| *CRIP2* | 2.000822 | 0.738468 | 2.709422 | 0.00674005 | NA |
| *SAMD4A* | 0.930207 | 0.343438 | 2.708512 | 0.00675857 | 0.133482607 |
| *ZC3H7B* | -0.404322 | 0.149327 | -2.707633 | 0.00677649 | 0.133482607 |
| *GTF3C2* | -0.322704 | 0.119189 | -2.707501 | 0.00677919 | 0.133482607 |
| *GIHCG* | -0.377141 | 0.139302 | -2.707361 | 0.00678206 | 0.133482607 |
| *ENSG00000273141* | -2.346237 | 0.866645 | -2.707263 | 0.00678404 | NA |
| *ZNF557* | 0.388199 | 0.143459 | 2.705994 | 0.00681004 | 0.133791411 |
| *PDCD7* | 0.411573 | 0.152174 | 2.704626 | 0.00683813 | 0.134064575 |
| *TNRC6C* | 0.57742 | 0.213571 | 2.703637 | 0.00685852 | 0.134064575 |
| *CREB3L2* | -0.461694 | 0.170775 | -2.703522 | 0.00686089 | 0.134064575 |
| *IFIT2* | 0.855645 | 0.316613 | 2.702492 | 0.00688218 | 0.13423958 |
| *ENSG00000272758* | 0.798884 | 0.2959 | 2.699842 | 0.00693725 | 0.135071558 |
| *PTGS1* | 0.726321 | 0.26913 | 2.698768 | 0.00695967 | 0.135266156 |
| *MRPS26* | -0.194726 | 0.072191 | -2.697386 | 0.00698862 | 0.135586747 |
| *UPP1* | 0.777786 | 0.288414 | 2.696768 | 0.00700161 | 0.135597054 |
| *OGFRL1* | -0.472763 | 0.175403 | -2.695294 | 0.00703266 | 0.135956428 |
| *COL9A3* | -0.963205 | 0.357762 | -2.692309 | 0.00709591 | 0.136935923 |
| *TIMM8B* | -0.218384 | 0.08115 | -2.691104 | 0.00712159 | 0.137188282 |
| *HDAC2* | 0.235091 | 0.08744 | 2.688594 | 0.00717536 | 0.137979878 |
| *GLIDR* | -0.609097 | 0.226713 | -2.686639 | 0.0072175 | 0.138545345 |
| *ENSG00000277476* | 1.682267 | 0.626571 | 2.68488 | 0.00725559 | NA |
| *ENSG00000275964* | -0.691639 | 0.25795 | -2.681288 | 0.00733393 | 0.140532604 |
| *PSMB3* | -0.191027 | 0.071303 | -2.679103 | 0.00738197 | 0.14108733 |
| *CORO1A* | 0.2819 | 0.105234 | 2.678793 | 0.00738881 | 0.14108733 |
| *MYBBP1A* | -0.416186 | 0.155507 | -2.676323 | 0.00744349 | 0.141882498 |
| *LETM1* | -0.23693 | 0.088604 | -2.674048 | 0.00749417 | 0.142598765 |
| *HMGB3* | -0.691481 | 0.258729 | -2.672606 | 0.00752647 | 0.142804944 |
| *TLR7* | 0.830027 | 0.310597 | 2.672362 | 0.00753194 | 0.142804944 |
| *CBWD2* | -0.469113 | 0.17559 | -2.671638 | 0.0075482 | 0.142804944 |
| *BEND4* | 0.62042 | 0.23226 | 2.671226 | 0.00755748 | 0.142804944 |
| *STMP1* | -0.208163 | 0.07795 | -2.670482 | 0.00757425 | 0.142847251 |
| *VPS13C* | 0.272 | 0.101891 | 2.669512 | 0.00759616 | 0.142847251 |
| *ZNF567* | 0.390269 | 0.146278 | 2.667988 | 0.0076307 | 0.142847251 |
| *TMEM177* | -0.55307 | 0.207393 | -2.666767 | 0.00765846 | 0.142847251 |
| *SRSF11* | 0.2335 | 0.087566 | 2.666546 | 0.0076635 | 0.142847251 |
| *IGHG1* | 1.230969 | 0.461675 | 2.666308 | 0.00766894 | 0.142847251 |
| *RAB3GAP2* | 0.41596 | 0.15601 | 2.666248 | 0.0076703 | 0.142847251 |
| *NUDCD3* | -0.308357 | 0.115656 | -2.666154 | 0.00767246 | 0.142847251 |
| *APRT* | -0.145978 | 0.054757 | -2.665918 | 0.00767784 | 0.142847251 |
| *ARMT1* | 0.507397 | 0.190419 | 2.664644 | 0.007707 | 0.143144955 |
| *PSMB1* | -0.211928 | 0.079578 | -2.663142 | 0.00774147 | 0.143540311 |
| *FOXP4* | 0.579676 | 0.217949 | 2.659691 | 0.00782125 | 0.144703821 |

| | | | | | |
|---|---|---|---|---|---|
| *ENSG00000258056* | 0.74439 | 0.280005 | 2.658487 | 0.00784924 | 0.144703821 |
| *MSI2* | -0.470468 | 0.17697 | -2.658464 | 0.00784978 | 0.144703821 |
| *KLF6* | 0.330515 | 0.124363 | 2.657661 | 0.0078685 | 0.144703821 |
| *TMEM179B* | -0.235551 | 0.088634 | -2.657567 | 0.0078707 | 0.144703821 |
| *SLC9A3R1* | 0.39983 | 0.150523 | 2.656282 | 0.00790075 | 0.145011483 |
| *WDR70* | -0.315216 | 0.118757 | -2.654295 | 0.00794744 | 0.145143499 |
| *STXBP4* | -0.751793 | 0.283259 | -2.654085 | 0.00795237 | 0.145143499 |
| *ZBTB25* | 0.357084 | 0.13456 | 2.653721 | 0.00796097 | 0.145143499 |
| *KIF5B* | 0.247898 | 0.093416 | 2.653707 | 0.00796129 | 0.145143499 |
| *PARP15* | 0.395097 | 0.148937 | 2.652777 | 0.00798325 | 0.145300567 |
| *VSIG10L* | 1.944709 | 0.733584 | 2.650969 | 0.00802613 | NA |
| *ODF3B* | -0.479896 | 0.181115 | -2.649676 | 0.0080569 | 0.146381567 |
| *AKIRIN2* | 0.241015 | 0.090988 | 2.648868 | 0.00807618 | 0.146381567 |
| *ATP5MC3* | -0.26071 | 0.098434 | -2.648583 | 0.008083 | 0.146381567 |
| *CDCA7L* | -0.408438 | 0.154433 | -2.644754 | 0.00817504 | 0.147581605 |
| *ZNF350* | -0.340718 | 0.128831 | -2.644698 | 0.00817638 | 0.147581605 |
| *TCTN1* | -0.879453 | 0.332747 | -2.643011 | 0.00821725 | 0.147971301 |
| *NDUFAF8* | -0.200264 | 0.075785 | -2.64254 | 0.00822867 | 0.147971301 |
| *BCL2L13* | -0.326805 | 0.12369 | -2.642125 | 0.00823876 | 0.147971301 |
| *RASSF3* | 0.305845 | 0.115905 | 2.638753 | 0.00832116 | 0.149205093 |
| *CD1D* | 0.632491 | 0.239821 | 2.637347 | 0.00835574 | 0.149292306 |
| *TUT7* | 0.344687 | 0.130709 | 2.637053 | 0.00836298 | 0.149292306 |
| *HES6* | -0.728287 | 0.276192 | -2.636883 | 0.00836717 | 0.149292306 |
| *UFD1* | -0.262211 | 0.099492 | -2.635488 | 0.00840163 | 0.1496618 |
| *ARL11* | 0.586558 | 0.223055 | 2.629651 | 0.00854725 | 0.151913356 |
| *PHF19* | -0.57885 | 0.220153 | -2.629306 | 0.00855594 | 0.151913356 |
| *RCBTB1* | -0.678867 | 0.258331 | -2.627893 | 0.00859156 | 0.152297272 |
| *ACP1* | -0.157244 | 0.059856 | -2.627037 | 0.0086132 | 0.152432582 |
| *ZNHIT2* | -0.363008 | 0.138295 | -2.624886 | 0.00866781 | 0.152705037 |
| *ENSG00000268400* | -1.205398 | 0.459248 | -2.624723 | 0.00867195 | 0.152705037 |
| *COX17* | 0.327034 | 0.124616 | 2.624339 | 0.00868172 | 0.152705037 |
| *PRICKLE1* | 0.771523 | 0.294001 | 2.624222 | 0.00868471 | 0.152705037 |
| *EPS15* | -0.312013 | 0.118935 | -2.623399 | 0.00870572 | 0.152827566 |
| *IKZF1* | -0.251169 | 0.09588 | -2.619609 | 0.00880307 | 0.154287582 |
| *ENSG00000274536* | -0.607384 | 0.232023 | -2.617773 | 0.00885056 | 0.154870589 |
| *IL4R* | -0.418254 | 0.159937 | -2.615115 | 0.00891974 | 0.155664241 |
| *ZNF747* | -0.422674 | 0.161659 | -2.614598 | 0.00893324 | 0.155664241 |
| *TPM1* | 0.601327 | 0.230007 | 2.614385 | 0.00893882 | 0.155664241 |
| *PTPRS* | 1.299226 | 0.497218 | 2.612991 | 0.00897537 | 0.155773287 |
| *SUMO1* | -0.150843 | 0.057742 | -2.612363 | 0.00899187 | 0.155773287 |
| *RPLP2* | 0.134009 | 0.051339 | 2.610298 | 0.00904633 | 0.155773287 |
| *PYURF* | -0.186765 | 0.071565 | -2.609727 | 0.00906145 | 0.155773287 |
| *XRRA1* | -0.716451 | 0.274586 | -2.60921 | 0.00907515 | 0.155773287 |
| *RASSF2* | 0.338692 | 0.129834 | 2.608644 | 0.00909018 | 0.155773287 |
| *INTS6* | -0.237649 | 0.091101 | -2.608639 | 0.0090903 | 0.155773287 |
| *FAAP24* | -0.654178 | 0.250775 | -2.608629 | 0.00909057 | 0.155773287 |
| *ZNF749* | -1.343054 | 0.514938 | -2.608189 | 0.00910228 | 0.155773287 |

| | | | | | |
|---|---|---|---|---|---|
| *NDUFC2* | -0.260392 | 0.099838 | -2.608148 | 0.00910337 | 0.155773287 |
| *CNP* | 0.323403 | 0.124009 | 2.607904 | 0.00910986 | 0.155773287 |
| *HIVEP3* | -0.54982 | 0.21085 | -2.607642 | 0.00911683 | 0.155773287 |
| *C16orf74* | -0.489808 | 0.187907 | -2.606654 | 0.00914317 | 0.155978496 |
| *PM20D2* | 0.505336 | 0.193991 | 2.604944 | 0.00918893 | 0.156480562 |
| *ATXN2* | 0.35045 | 0.13461 | 2.603453 | 0.00922898 | 0.156480562 |
| *H3F3B* | -0.137925 | 0.052987 | -2.602994 | 0.00924135 | 0.156480562 |
| *FRG1* | -0.183836 | 0.070633 | -2.602677 | 0.00924992 | 0.156480562 |
| *ENSG00000264443* | 0.699829 | 0.268965 | 2.601931 | 0.00927004 | 0.156480562 |
| *SAMD9L* | 0.427152 | 0.164178 | 2.601751 | 0.00927491 | 0.156480562 |
| *U2SURP* | -0.171928 | 0.066109 | -2.600676 | 0.00930403 | 0.156480562 |
| *MAP3K1* | 0.332755 | 0.127992 | 2.599817 | 0.00932734 | 0.156480562 |
| *LACTB2* | 0.380433 | 0.146333 | 2.599769 | 0.00932864 | 0.156480562 |
| *ENSG00000254397* | 1.062596 | 0.408762 | 2.599547 | 0.00933468 | 0.156480562 |
| *ITFG1* | -0.317718 | 0.122239 | -2.599156 | 0.00934532 | 0.156480562 |
| *NIPAL2* | 0.731787 | 0.281583 | 2.598827 | 0.00935428 | 0.156480562 |
| *CHTOP* | 0.167985 | 0.064655 | 2.598182 | 0.00937188 | 0.156480562 |
| *MVB12A* | -0.29026 | 0.11172 | -2.598109 | 0.00937388 | 0.156480562 |
| *SDSL* | 1.217314 | 0.469015 | 2.595468 | 0.00944623 | 0.157446755 |
| *ENSG00000070423* | -0.261132 | 0.100637 | -2.594802 | 0.00946454 | 0.157510803 |
| *PWWP2B* | 0.951105 | 0.366785 | 2.593085 | 0.00951193 | 0.158027513 |
| *ADPRH* | 0.794367 | 0.306395 | 2.592626 | 0.00952463 | 0.158027513 |
| *C17orf49* | -0.368065 | 0.142015 | -2.591734 | 0.00954936 | 0.158196654 |
| *NEMP1* | 0.605112 | 0.233603 | 2.590346 | 0.00958796 | 0.158594786 |
| *METTL1* | -0.390984 | 0.151053 | -2.588384 | 0.00964275 | 0.159259076 |
| *CD3D* | -2.441079 | 0.94353 | -2.587178 | 0.00967657 | NA |
| *DNAJB4* | 0.564918 | 0.218426 | 2.586308 | 0.00970102 | 0.159835816 |
| *SMKR1* | -0.500141 | 0.193396 | -2.586094 | 0.00970705 | 0.159835816 |
| *RFX1* | 0.373275 | 0.14447 | 2.583754 | 0.00977315 | 0.160461946 |
| *CEP104* | 0.414367 | 0.160377 | 2.583704 | 0.00977456 | 0.160461946 |
| *ARID3B* | 0.582522 | 0.225513 | 2.583091 | 0.00979195 | 0.160505469 |
| *HLA.DQA1* | 0.395143 | 0.153045 | 2.581874 | 0.00982653 | 0.160830038 |
| *ELP5* | -0.38953 | 0.15094 | -2.580698 | 0.00986007 | 0.161136564 |
| *PCNX4* | 0.342688 | 0.132864 | 2.579246 | 0.00990164 | 0.161573302 |
| *ARMCX1* | -1.459737 | 0.566256 | -2.577877 | 0.00994094 | NA |
| *RUNX2* | 1.307329 | 0.507237 | 2.577354 | 0.00995599 | 0.162205817 |
| *OXSM* | 0.548403 | 0.212818 | 2.576861 | 0.0099702 | 0.162205817 |
| *SPOP* | 0.284479 | 0.110593 | 2.572306 | 0.01010237 | 0.164083645 |
| *UAP1L1* | 0.887678 | 0.345152 | 2.571846 | 0.01011578 | 0.164083645 |
| *IER3IP1* | -0.199029 | 0.077492 | -2.568384 | 0.01021738 | 0.165322787 |
| *ERLIN2* | -0.513128 | 0.1998 | -2.568209 | 0.01022255 | 0.165322787 |
| *PLCB2* | 0.70612 | 0.275093 | 2.566844 | 0.01026288 | 0.165728691 |
| *MIR181A1HG* | -0.666649 | 0.25985 | -2.565515 | 0.01030228 | 0.166107227 |
| *RPA3* | -0.194609 | 0.075884 | -2.564558 | 0.01033074 | 0.166107227 |
| *PSMA6* | -0.267148 | 0.104171 | -2.564512 | 0.0103321 | 0.166107227 |
| *HPS3* | 0.42328 | 0.165101 | 2.56376 | 0.01035452 | 0.166222042 |
| *DNAJC30* | -0.27515 | 0.10735 | -2.563113 | 0.01037384 | 0.166287043 |

| | | | | | |
|---|---|---|---|---|---|
| *ENSG00000272277* | -1.170768 | 0.456902 | -2.562407 | 0.01039495 | 0.166380292 |
| *CCDC121* | 2.502368 | 0.977453 | 2.56009 | 0.01046452 | NA |
| *FOLR2* | 1.769788 | 0.691378 | 2.5598 | 0.01047325 | NA |
| *ARMCX3* | 0.27105 | 0.106045 | 2.555993 | 0.01058852 | 0.169025683 |
| *MTRF1* | 0.486485 | 0.190368 | 2.5555 | 0.01060354 | 0.169025683 |
| *ENSG00000275464* | -1.29125 | 0.505304 | -2.555392 | 0.01060681 | 0.169025683 |
| *ENSG00000229728* | -1.544646 | 0.604631 | -2.554691 | 0.01062822 | NA |
| *C8orf82* | -0.920965 | 0.360542 | -2.554395 | 0.01063725 | 0.169262943 |
| *STYX* | 0.412077 | 0.161397 | 2.553187 | 0.01067422 | 0.169603239 |
| *DHTKD1* | -0.368052 | 0.144309 | -2.550447 | 0.01075848 | 0.170692914 |
| *CFD* | 1.735498 | 0.680604 | 2.549938 | 0.01077419 | NA |
| *PBX4* | 1.088538 | 0.427084 | 2.548767 | 0.01081046 | 0.17102892 |
| *ICMT* | -0.577014 | 0.226391 | -2.548746 | 0.01081109 | 0.17102892 |
| *CENPU* | -0.658069 | 0.258281 | -2.547884 | 0.01083786 | 0.171203508 |
| *RPS28* | 0.083048 | 0.032607 | 2.54693 | 0.01086754 | 0.171423561 |
| *PPP2R5A* | 0.397861 | 0.156309 | 2.545344 | 0.01091701 | 0.171954697 |
| *GCSAM* | -0.650631 | 0.256039 | -2.541137 | 0.01104926 | 0.173786324 |
| *BCAP31* | -0.222655 | 0.0877 | -2.538834 | 0.01112226 | 0.17468213 |
| *RPGRIP1L* | -0.638574 | 0.251738 | -2.536665 | 0.0111914 | 0.175514688 |
| *TOB1* | 0.497788 | 0.196313 | 2.53568 | 0.01122291 | 0.175613187 |
| *JMJD4* | -0.3381 | 0.133359 | -2.535255 | 0.01123657 | 0.175613187 |
| *NCOA3* | 0.280505 | 0.110655 | 2.534958 | 0.01124609 | 0.175613187 |
| *MIR155HG* | 0.403296 | 0.159224 | 2.532895 | 0.0113125 | 0.176397195 |
| *FAM184B* | -1.730781 | 0.683497 | -2.532244 | 0.01133351 | 0.176471962 |
| *KLK2* | 2.049721 | 0.810396 | 2.529281 | 0.01142964 | NA |
| *PSMB10* | -0.228253 | 0.090244 | -2.529274 | 0.01142986 | 0.177703874 |
| *ZNF668* | 0.509937 | 0.201652 | 2.528801 | 0.01144528 | 0.177703874 |
| *PLEKHO2* | 0.818926 | 0.323978 | 2.527719 | 0.01148062 | 0.177982281 |
| *FLI1* | 0.339322 | 0.134278 | 2.527015 | 0.01150366 | 0.177982281 |
| *CMSS1* | -0.235849 | 0.093341 | -2.526752 | 0.01151227 | 0.177982281 |
| *RFTN1* | -0.272714 | 0.107958 | -2.526106 | 0.01153346 | 0.178056921 |
| *LY86.AS1* | 2.248861 | 0.890618 | 2.525056 | 0.011568 | NA |
| *PDE4B* | 0.274566 | 0.108765 | 2.524391 | 0.01158989 | 0.178674657 |
| *TAF1A* | 0.519586 | 0.206023 | 2.52198 | 0.01166965 | 0.179649878 |
| *PPBP* | 2.500724 | 0.991709 | 2.521632 | 0.01168117 | NA |
| *MSMP* | -0.933922 | 0.370388 | -2.521472 | 0.0116865 | 0.179655202 |
| *BTG2* | 0.23137 | 0.09181 | 2.520084 | 0.0117327 | 0.179800265 |
| *INTS10* | -0.242339 | 0.096183 | -2.519552 | 0.01175045 | 0.179800265 |
| *NAB1* | 0.535384 | 0.212496 | 2.519504 | 0.01175204 | 0.179800265 |
| *BAG1* | -0.205895 | 0.08173 | -2.519205 | 0.01176202 | 0.179800265 |
| *SPIB* | 0.176736 | 0.070174 | 2.518546 | 0.01178405 | 0.179884464 |
| *PIAS2* | -0.296739 | 0.11793 | -2.516223 | 0.01186202 | 0.180591 |
| *UST* | 0.887116 | 0.352594 | 2.515968 | 0.01187061 | 0.180591 |
| *BBX* | 0.210842 | 0.083811 | 2.515686 | 0.01188011 | 0.180591 |
| *NADSYN1* | -0.287444 | 0.114305 | -2.514717 | 0.01191279 | 0.180835125 |
| *NUBP2* | -0.189601 | 0.075469 | -2.512309 | 0.01199441 | 0.181218555 |
| *MIF.AS1* | -0.461719 | 0.183801 | -2.512052 | 0.01200315 | 0.181218555 |

| | | | | | |
|---|---|---|---|---|---|
| *ASPH* | 0.740717 | 0.294891 | 2.511835 | 0.01201053 | 0.181218555 |
| *HMGB2* | -0.253368 | 0.100877 | -2.511651 | 0.01201678 | 0.181218555 |
| *PDE4DIP* | -0.387581 | 0.154333 | -2.51133 | 0.01202773 | 0.181218555 |
| *MYL6B* | -0.276933 | 0.110287 | -2.51103 | 0.01203795 | 0.181218555 |
| *NUDT8* | -0.608676 | 0.242535 | -2.509639 | 0.01208546 | 0.181682561 |
| *TMEM154* | 0.465664 | 0.185643 | 2.508379 | 0.01212867 | 0.181920139 |
| *GADD45G* | -0.842329 | 0.335933 | -2.507428 | 0.01216134 | 0.181920139 |
| *TMEM187* | 0.771817 | 0.307899 | 2.506724 | 0.01218557 | 0.181920139 |
| *RHOBTB2* | 1.076577 | 0.429561 | 2.506224 | 0.01220283 | 0.181920139 |
| *ZBTB7B* | 0.442231 | 0.176455 | 2.506192 | 0.01220395 | 0.181920139 |
| *ZSCAN22* | 0.938886 | 0.374656 | 2.505994 | 0.01221076 | 0.181920139 |
| *WDR89* | 0.385545 | 0.153862 | 2.505777 | 0.01221827 | 0.181920139 |
| *FAM76B* | 0.338578 | 0.135173 | 2.504785 | 0.0122526 | 0.182182021 |
| *ENSG00000249684* | 2.060176 | 0.82273 | 2.504074 | 0.01227724 | NA |
| *HMMR* | 1.087422 | 0.434282 | 2.503953 | 0.01228143 | NA |
| *MGME1* | 0.337666 | 0.135005 | 2.501137 | 0.01237952 | 0.183678529 |
| *ENSG00000167414* | -0.969877 | 0.387808 | -2.500923 | 0.01238699 | 0.183678529 |
| *RHOBTB1* | -2.221979 | 0.888472 | -2.5009 | 0.01238781 | NA |
| *SMARCE1* | -0.184485 | 0.073818 | -2.499192 | 0.01244768 | 0.184327259 |
| *ASB2* | 1.436665 | 0.575031 | 2.498411 | 0.01247514 | 0.184482953 |
| *CHMP5* | 0.23546 | 0.094318 | 2.496456 | 0.01254414 | 0.185251532 |
| *MBLAC1* | -1.224827 | 0.491214 | -2.493469 | 0.01265016 | 0.186564115 |
| *RSBN1* | 0.276603 | 0.110993 | 2.492079 | 0.01269979 | 0.187042609 |
| *ZNF420* | 0.432136 | 0.17362 | 2.488975 | 0.01281118 | 0.188428222 |
| *TTI2* | 0.495703 | 0.199415 | 2.485793 | 0.01292632 | 0.189865162 |
| *IRF2BPL* | 0.399072 | 0.160607 | 2.484783 | 0.01296305 | 0.190148089 |
| *EXOG* | -0.315479 | 0.127108 | -2.481965 | 0.01306602 | 0.191400422 |
| *SLC9A9* | -0.611432 | 0.246452 | -2.480935 | 0.01310382 | 0.191586162 |
| *CENPB* | -0.445141 | 0.179445 | -2.480661 | 0.0131139 | 0.191586162 |
| *DGCR6* | -1.505 | 0.607114 | -2.478941 | 0.01317731 | NA |
| *GCNT2* | 0.55493 | 0.223922 | 2.478222 | 0.01320389 | 0.192642204 |
| *AUP1* | -0.146328 | 0.059137 | -2.474378 | 0.01334683 | 0.194368899 |
| *REEP4* | 0.555059 | 0.224355 | 2.474026 | 0.01335998 | 0.194368899 |
| *ZNF616* | 0.868461 | 0.351113 | 2.473454 | 0.01338141 | 0.194368899 |
| *PPP1R16B* | 0.403422 | 0.163124 | 2.473107 | 0.0133944 | 0.194368899 |
| *TAF13* | 0.459115 | 0.185686 | 2.472529 | 0.01341606 | 0.194368899 |
| *AKAP17A* | 0.248973 | 0.100716 | 2.472023 | 0.0134351 | 0.194368899 |
| *ARL4A* | -0.383464 | 0.155158 | -2.471447 | 0.01345677 | 0.194368899 |
| *BLM* | 0.525023 | 0.212454 | 2.471225 | 0.0134651 | 0.194368899 |
| *C9orf16* | -0.228832 | 0.092637 | -2.470196 | 0.01350391 | 0.194670874 |
| *CFAP298* | -0.268028 | 0.108554 | -2.469067 | 0.0135466 | 0.195028009 |
| *RACGAP1* | 0.765263 | 0.31025 | 2.466606 | 0.01364003 | 0.196113778 |
| *ENSG00000278932* | 1.923096 | 0.780236 | 2.464763 | 0.0137104 | NA |
| *RIN3* | 0.555621 | 0.225458 | 2.464407 | 0.01372403 | 0.196920389 |
| *KCNN4* | 0.731511 | 0.296857 | 2.46419 | 0.01373232 | 0.196920389 |
| *C6orf226* | -0.433474 | 0.175951 | -2.463605 | 0.01375475 | 0.196982562 |
| *AHI1* | -0.374095 | 0.15192 | -2.462442 | 0.01379943 | 0.197362715 |

| | | | | | |
|---|---|---|---|---|---|
| *TRMT6* | -0.396373 | 0.161065 | -2.460946 | 0.01385714 | 0.197927974 |
| *ATP5ME* | -0.179212 | 0.072858 | -2.459743 | 0.01390364 | 0.198294587 |
| *FBXL17* | 0.507753 | 0.206459 | 2.459341 | 0.01391925 | 0.198294587 |
| *PPCS* | -0.191214 | 0.077786 | -2.458195 | 0.01396372 | 0.198418986 |
| *TTC9C* | 0.285951 | 0.116327 | 2.458177 | 0.01396444 | 0.198418986 |
| *BACE2* | 1.621129 | 0.659657 | 2.457533 | 0.01398949 | NA |
| *ENSG00000273338* | -0.441985 | 0.179969 | -2.455889 | 0.01405365 | 0.199426185 |
| *BAG4* | 0.406111 | 0.165453 | 2.454541 | 0.01410647 | 0.19991506 |
| *ENSG00000176320* | 2.192084 | 0.893158 | 2.454308 | 0.01411561 | NA |
| *ZNF284* | 0.723975 | 0.295122 | 2.453138 | 0.01416159 | 0.200280014 |
| *ENSG00000188242* | -0.693042 | 0.282534 | -2.45295 | 0.01416902 | 0.200280014 |
| *HDX* | 1.097874 | 0.447907 | 2.451121 | 0.01424122 | 0.200951434 |
| *CERS4* | -0.273239 | 0.111489 | -2.450812 | 0.01425345 | 0.200951434 |
| *METAP2* | -0.160181 | 0.065394 | -2.44948 | 0.01430628 | 0.201435394 |
| *NDFIP2* | -0.816799 | 0.333983 | -2.44563 | 0.01445994 | 0.203335925 |
| *SDHAF3* | -0.223449 | 0.091416 | -2.444324 | 0.01451239 | 0.203738151 |
| *SLC25A5* | -0.169442 | 0.069341 | -2.443594 | 0.0145418 | 0.203738151 |
| *CD27* | 1.249254 | 0.511252 | 2.443522 | 0.0145447 | 0.203738151 |
| *CRLF3* | 0.244995 | 0.100307 | 2.442458 | 0.01458762 | 0.204076722 |
| *MPP1* | -0.647211 | 0.2651 | -2.441387 | 0.01463099 | 0.20440473 |
| *TMEM156* | 0.373798 | 0.153136 | 2.440951 | 0.01464863 | 0.20440473 |
| *CNBP* | -0.198843 | 0.081497 | -2.439873 | 0.01469244 | 0.204492364 |
| *MTCH2* | -0.250632 | 0.102724 | -2.439872 | 0.01469249 | 0.204492364 |
| *C7orf26* | -0.409975 | 0.168176 | -2.437774 | 0.014778 | 0.205419798 |
| *COL4A4* | 1.989942 | 0.816578 | 2.436928 | 0.01481265 | NA |
| *ENSG00000262089* | 0.873271 | 0.358428 | 2.436391 | 0.01483465 | 0.205944267 |
| *CHTF18* | -0.549726 | 0.225677 | -2.435895 | 0.01485501 | 0.20596423 |
| *FSD1* | -1.19947 | 0.492519 | -2.435376 | 0.01487633 | NA |
| *NFE2L1* | 0.3092 | 0.126982 | 2.434997 | 0.01489191 | 0.2061739 |
| *GCC2* | 0.246813 | 0.101401 | 2.434033 | 0.01493164 | 0.2061739 |
| *SLF2* | 0.310426 | 0.127551 | 2.433747 | 0.01494343 | 0.2061739 |
| *FCGR2A* | 2.511286 | 1.031876 | 2.43371 | 0.01494498 | NA |
| *SMC5* | -0.242846 | 0.099814 | -2.43298 | 0.01497512 | 0.2061739 |
| *GNPDA1* | 0.340383 | 0.139906 | 2.432944 | 0.01497663 | 0.2061739 |
| *ENSG00000265778* | -0.999918 | 0.41102 | -2.432771 | 0.01498379 | 0.2061739 |
| *CD48* | 0.224889 | 0.09249 | 2.431493 | 0.01503674 | 0.206641206 |
| *YWHAQ* | -0.191062 | 0.078668 | -2.428704 | 0.01515289 | 0.207061635 |
| *ARHGAP9* | -0.247544 | 0.101942 | -2.428273 | 0.01517091 | 0.207061635 |
| *TOMM7* | 0.166739 | 0.068679 | 2.427821 | 0.01518985 | 0.207061635 |
| *PIGT* | -0.262534 | 0.108156 | -2.42737 | 0.01520875 | 0.207061635 |
| *POLD4* | 0.23882 | 0.098389 | 2.4273 | 0.01521167 | 0.207061635 |
| *RPS9* | 0.154049 | 0.063466 | 2.427277 | 0.01521263 | 0.207061635 |
| *RASGEF1B* | -0.493497 | 0.20337 | -2.426593 | 0.01524134 | 0.207061635 |
| *PSMC3IP* | 0.571455 | 0.235524 | 2.42631 | 0.01525323 | 0.207061635 |
| *LGALS3* | 0.782886 | 0.322672 | 2.42626 | 0.01525534 | 0.207061635 |
| *APBB1* | 0.828546 | 0.341499 | 2.426207 | 0.01525757 | 0.207061635 |
| *ERP29* | 0.145845 | 0.060132 | 2.425402 | 0.01529145 | 0.207201175 |

| | | | | | |
|---|---|---|---|---|---|
| *TMEM134* | 0.165877 | 0.068401 | 2.425058 | 0.01530593 | 0.207201175 |
| *ITGA10* | 0.868764 | 0.358464 | 2.423574 | 0.0153686 | 0.207674025 |
| *TAGAP* | -0.514927 | 0.212488 | -2.423328 | 0.01537902 | 0.207674025 |
| *SLCO3A1* | -1.545305 | 0.63786 | -2.42264 | 0.01540818 | NA |
| *MCM7* | -0.309483 | 0.127855 | -2.420569 | 0.01549625 | 0.208997812 |
| *CTSB* | 0.26225 | 0.108458 | 2.417979 | 0.01560698 | 0.210026276 |
| *GLI4* | -0.953225 | 0.39426 | -2.417754 | 0.01561662 | 0.210026276 |
| *VAT1* | 0.387218 | 0.160192 | 2.417215 | 0.01563976 | 0.210026276 |
| *TRAM1* | -0.168327 | 0.069643 | -2.416984 | 0.0156497 | 0.210026276 |
| *RGL1* | -1.234665 | 0.511177 | -2.415339 | 0.01572056 | NA |
| *PCBP4* | 0.606615 | 0.251647 | 2.410576 | 0.01592735 | 0.213489314 |
| *RFC4* | 0.343542 | 0.142549 | 2.40999 | 0.01595298 | 0.213568428 |
| *SNAP29* | 0.286317 | 0.118826 | 2.409543 | 0.0159725 | 0.213568428 |
| *ZNF667.AS1* | 0.324697 | 0.134822 | 2.408337 | 0.0160254 | 0.214004281 |
| *SRD5A3* | 0.479742 | 0.199237 | 2.407903 | 0.01604442 | 0.214004281 |
| *ENSG00000237491* | -0.583535 | 0.2424 | -2.407324 | 0.0160699 | 0.214081746 |
| *LINC01250* | -2.766859 | 1.149611 | -2.406779 | 0.01609389 | NA |
| *CHCHD3* | -0.265076 | 0.110215 | -2.405078 | 0.01616903 | 0.215139084 |
| *ENSG00000276449* | 1.048697 | 0.436256 | 2.403858 | 0.01622307 | 0.215530326 |
| *SLC29A2* | -0.588059 | 0.244687 | -2.403315 | 0.0162472 | 0.215530326 |
| *ENSG00000089127* | 0.66073 | 0.275035 | 2.402348 | 0.01629021 | 0.215530326 |
| *GLRX* | 0.385678 | 0.160614 | 2.401277 | 0.01633798 | 0.215530326 |
| *ING3* | -0.326934 | 0.136152 | -2.401252 | 0.0163391 | 0.215530326 |
| *SEPTIN6* | 0.152384 | 0.06346 | 2.401244 | 0.01633946 | 0.215530326 |
| *LY6E* | 0.252344 | 0.105137 | 2.400151 | 0.01638829 | 0.215530326 |
| *PCGF5* | -0.251931 | 0.104969 | -2.400061 | 0.01639232 | 0.215530326 |
| *ITM2C* | -0.369723 | 0.154048 | -2.400046 | 0.01639302 | 0.215530326 |
| *STT3B* | -0.338552 | 0.141092 | -2.399514 | 0.01641684 | 0.215530326 |
| *ZFAND6* | 0.351161 | 0.14639 | 2.398807 | 0.01644859 | 0.215530326 |
| *HPSE* | -2.055749 | 0.857031 | -2.398686 | 0.016454 | NA |
| *ETFRF1* | 0.313782 | 0.130823 | 2.39853 | 0.01646101 | 0.215530326 |
| *SAR1B* | 0.265649 | 0.110757 | 2.398498 | 0.01646248 | 0.215530326 |
| *RBM38* | -0.315403 | 0.131516 | -2.398204 | 0.01647567 | 0.215530326 |
| *ENSG00000262429* | -1.087831 | 0.453698 | -2.397698 | 0.01649848 | NA |
| *MAGEH1* | -0.407146 | 0.169937 | -2.395863 | 0.01658129 | 0.216465864 |
| *PTTG1IP* | 0.398566 | 0.1664 | 2.39522 | 0.01661041 | 0.216465864 |
| *NDUFA7* | -0.279841 | 0.116859 | -2.394699 | 0.01663401 | 0.216465864 |
| *EML4* | 0.219004 | 0.091458 | 2.394578 | 0.01663951 | 0.216465864 |
| *RANGRF* | -0.231037 | 0.09649 | -2.394421 | 0.01664663 | 0.216465864 |
| *ANGPTL6* | 1.474415 | 0.616004 | 2.393518 | 0.01668768 | NA |
| *ZNF43* | -0.308936 | 0.129092 | -2.393143 | 0.01670471 | 0.216722989 |
| *LENG1* | 0.271828 | 0.113593 | 2.392994 | 0.0167115 | 0.216722989 |
| *FUT7* | 1.705627 | 0.712839 | 2.392726 | 0.01672374 | NA |
| *BLOC1S2* | -0.177242 | 0.074077 | -2.392673 | 0.01672614 | 0.216722989 |
| *LINC02193* | -0.732803 | 0.306491 | -2.390942 | 0.01680519 | 0.217488288 |
| *B4GALT1.AS1* | -0.981338 | 0.411189 | -2.386585 | 0.01700567 | 0.219759516 |
| *TMEM161A* | -0.386871 | 0.162125 | -2.386252 | 0.01702107 | 0.219759516 |

| | | | | | |
|---|---|---|---|---|---|
| *DUSP12* | -0.250611 | 0.105057 | -2.385484 | 0.01705665 | 0.219910123 |
| *TESMIN* | -2.337267 | 0.979865 | -2.385295 | 0.01706545 | NA |
| *TXLNB* | 1.014102 | 0.425213 | 2.384928 | 0.01708247 | 0.219910123 |
| *ZNF879* | -0.529627 | 0.222094 | -2.384694 | 0.01709335 | 0.219910123 |
| *GNL3L* | -0.293324 | 0.123197 | -2.380931 | 0.01726894 | 0.221671608 |
| *GANC* | 0.446424 | 0.187503 | 2.380887 | 0.017271 | 0.221671608 |
| *ACSL3* | -0.310835 | 0.130619 | -2.379715 | 0.01732602 | 0.222115854 |
| *MMAB* | -0.15846 | 0.066643 | -2.377734 | 0.0174194 | 0.222683932 |
| *SAR1A* | -0.245148 | 0.103107 | -2.377613 | 0.0174251 | 0.222683932 |
| *DUSP14* | 0.446502 | 0.187805 | 2.377473 | 0.01743171 | 0.222683932 |
| *STAG3* | 0.458965 | 0.193149 | 2.376225 | 0.0174908 | 0.223176886 |
| *OTUD6B.AS1* | 0.246687 | 0.103836 | 2.375733 | 0.01751415 | 0.223213163 |
| *HLA.G* | -1.345857 | 0.56667 | -2.375027 | 0.01754766 | 0.223378643 |
| *AHCYL1* | 0.253389 | 0.106737 | 2.37396 | 0.01759847 | 0.223763675 |
| *PPP1R16A* | -0.411149 | 0.173227 | -2.373465 | 0.01762207 | 0.223802374 |
| *PELI3* | -1.04999 | 0.44261 | -2.372269 | 0.01767921 | 0.223912874 |
| *COX11* | -0.212351 | 0.089519 | -2.372139 | 0.01768546 | 0.223912874 |
| *HINT2* | -0.210999 | 0.088954 | -2.371992 | 0.01769249 | 0.223912874 |
| *FBXL3* | 0.307428 | 0.129645 | 2.371311 | 0.0177251 | 0.224065082 |
| *ZNF532* | 0.60182 | 0.253838 | 2.370876 | 0.01774596 | 0.224068503 |
| *RNF19A* | 0.36518 | 0.154109 | 2.369622 | 0.0178063 | 0.224433154 |
| *LPCAT2* | -1.852675 | 0.782028 | -2.369064 | 0.01783317 | NA |
| *HEXA* | 0.266488 | 0.112488 | 2.369039 | 0.01783437 | 0.224433154 |
| *NUP210* | 0.259321 | 0.109465 | 2.368991 | 0.0178367 | 0.224433154 |
| *RASA4* | 0.443264 | 0.18722 | 2.367609 | 0.01790343 | 0.225012602 |
| *C19orf48* | 0.33371 | 0.141009 | 2.366579 | 0.01795335 | 0.225379803 |
| *SAP25* | 0.70138 | 0.296496 | 2.365562 | 0.01800272 | 0.225531429 |
| *RING1* | -0.251801 | 0.106448 | -2.365477 | 0.01800687 | 0.225531429 |
| *SULT1A3* | 0.598288 | 0.253082 | 2.364014 | 0.01807815 | 0.225648098 |
| *FAM129C* | -0.387281 | 0.163835 | -2.363844 | 0.01808645 | 0.225648098 |
| *PACSIN2* | 0.311162 | 0.13164 | 2.363734 | 0.01809177 | 0.225648098 |
| *SEMA4A* | -1.066171 | 0.451314 | -2.36237 | 0.0181585 | 0.225648098 |
| *IL7R* | -1.205665 | 0.510428 | -2.362068 | 0.01817332 | 0.225648098 |
| *ENSG00000271646* | -0.375553 | 0.158995 | -2.362042 | 0.01817459 | 0.225648098 |
| *ADGRE2* | 1.090221 | 0.461563 | 2.36202 | 0.01817566 | 0.225648098 |
| *HAUS4* | 0.507005 | 0.214661 | 2.36189 | 0.01818205 | 0.225648098 |
| *STMN1* | -0.457091 | 0.193624 | -2.360717 | 0.01823962 | 0.226104824 |
| *FUS* | 0.111086 | 0.047089 | 2.359061 | 0.01832125 | 0.226858294 |
| *RPS15A* | 0.128049 | 0.054299 | 2.358214 | 0.01836309 | 0.22688864 |
| *DAXX* | 0.234096 | 0.09927 | 2.358168 | 0.01836539 | 0.22688864 |
| *SKP2* | -0.426967 | 0.181123 | -2.357337 | 0.01840653 | 0.227139097 |
| *ZNF574* | 0.494825 | 0.210048 | 2.355766 | 0.01848454 | 0.227791444 |
| *CHD2* | 0.236758 | 0.10053 | 2.355095 | 0.01851796 | 0.227791444 |
| *MAVS* | 0.233871 | 0.099308 | 2.355011 | 0.01852218 | 0.227791444 |
| *CTSS* | 0.180807 | 0.076865 | 2.352282 | 0.01865861 | 0.229149592 |
| *NUAK2* | -0.341641 | 0.145258 | -2.351961 | 0.01867472 | 0.229149592 |
| *PSME2* | -0.163232 | 0.069432 | -2.350954 | 0.01872537 | 0.22928596 |

| | | | | | |
|---|---|---|---|---|---|
| *TUBB4A* | 1.734952 | 0.738156 | 2.350387 | 0.0187539 | NA |
| *HACD2* | -0.316085 | 0.134486 | -2.350319 | 0.01875734 | 0.22928596 |
| *GRK2* | 0.253267 | 0.107773 | 2.349996 | 0.01877359 | 0.22928596 |
| *TMEM87B* | 0.504156 | 0.214545 | 2.349882 | 0.01877936 | 0.22928596 |
| *BLOC1S1* | -0.124437 | 0.05296 | -2.349648 | 0.01879117 | 0.22928596 |
| *SNRPD1* | -0.170706 | 0.072668 | -2.349113 | 0.01881821 | 0.229315436 |
| *TRIM5* | 0.590703 | 0.251495 | 2.348766 | 0.01883572 | 0.229315436 |
| *USP40* | 0.623864 | 0.265705 | 2.347957 | 0.01887671 | 0.229557675 |
| *DNPEP* | -0.186351 | 0.079404 | -2.346873 | 0.01893173 | 0.229872145 |
| *ATP6V0E1* | 0.177714 | 0.075732 | 2.346615 | 0.01894481 | 0.229872145 |
| *HAUS6* | 0.369096 | 0.157326 | 2.346066 | 0.01897272 | 0.229943531 |
| *C10orf143* | 1.018612 | 0.434252 | 2.345669 | 0.01899295 | 0.229943531 |
| *ZNF785* | 0.393869 | 0.16802 | 2.344182 | 0.01906885 | 0.230575085 |
| *ENSG00000272086* | -0.839239 | 0.358145 | -2.343295 | 0.01911428 | 0.230575085 |
| *CCT8* | 0.244129 | 0.104187 | 2.343184 | 0.01911993 | 0.230575085 |
| *SUDS3* | 0.225493 | 0.096242 | 2.342991 | 0.01912985 | 0.230575085 |
| *TBX21* | 1.511667 | 0.645304 | 2.342566 | 0.01915165 | NA |
| *SAT2* | 0.270239 | 0.115361 | 2.342554 | 0.01915229 | 0.230590125 |
| *ENSG00000185527* | -0.919756 | 0.392797 | -2.341557 | 0.0192035 | 0.230951257 |
| *ZNF540* | 0.909221 | 0.388522 | 2.340205 | 0.01927315 | 0.231293146 |
| *RPS27A* | 0.109318 | 0.046713 | 2.34018 | 0.01927443 | 0.231293146 |
| *DEF8* | -0.347402 | 0.148515 | -2.339168 | 0.01932672 | 0.231665245 |
| *PRDM1* | 0.886535 | 0.379193 | 2.337952 | 0.01938976 | 0.232113258 |
| *TNS3* | 0.539321 | 0.230713 | 2.337624 | 0.01940675 | 0.232113258 |
| *GATD3B* | -1.148076 | 0.49139 | -2.336385 | 0.0194712 | 0.232628421 |
| *MRPL38* | -0.482878 | 0.206767 | -2.335379 | 0.01952366 | 0.23299941 |
| *EIF4B* | 0.148084 | 0.063448 | 2.333919 | 0.01959996 | 0.233653846 |
| *PGBD2* | 0.366129 | 0.156938 | 2.332946 | 0.019651 | 0.233792846 |
| *WDR55* | -0.265462 | 0.113792 | -2.332877 | 0.01965458 | 0.233792846 |
| *DDX17* | 0.155255 | 0.066579 | 2.331889 | 0.01970654 | 0.234154988 |
| *HIST1H3E* | 0.866868 | 0.371949 | 2.330608 | 0.01977404 | 0.234185309 |
| *GGCT* | -0.199812 | 0.085737 | -2.330527 | 0.01977831 | 0.234185309 |
| *STAT3* | 0.263556 | 0.113122 | 2.329827 | 0.01981531 | 0.234185309 |
| *PTPDC1* | 0.737376 | 0.316498 | 2.329797 | 0.0198169 | 0.234185309 |
| *TNFAIP3* | -0.360999 | 0.154988 | -2.329206 | 0.01984817 | 0.234185309 |
| *ENSG00000271204* | 0.491627 | 0.211072 | 2.329194 | 0.01984878 | 0.234185309 |
| *SLC25A43* | 0.417865 | 0.179419 | 2.328988 | 0.01985971 | 0.234185309 |
| *ZNF341.AS1* | -1.726044 | 0.741849 | -2.326679 | 0.01998237 | NA |
| *SRSF6* | -0.284116 | 0.122174 | -2.325496 | 0.02004545 | 0.236062291 |
| *ZNF81* | -0.662846 | 0.285073 | -2.325181 | 0.02006226 | 0.236062291 |
| *ACTR3B* | -0.524815 | 0.225776 | -2.324487 | 0.0200994 | 0.236243958 |
| *RAB6A* | 0.227252 | 0.097794 | 2.323788 | 0.02013688 | 0.236397749 |
| *CDC42* | -0.154902 | 0.06668 | -2.323083 | 0.02017467 | 0.236397749 |
| *RPP21* | -0.28946 | 0.124616 | -2.32282 | 0.02018882 | 0.236397749 |
| *SYTL1* | 0.278021 | 0.119718 | 2.322291 | 0.02021728 | 0.236397749 |
| *CYP4V2* | 0.744595 | 0.320717 | 2.32166 | 0.02025126 | 0.236397749 |
| *MRPL57* | -0.154827 | 0.066702 | -2.321167 | 0.02027784 | 0.236397749 |

| | | | | | |
|---|---|---|---|---|---|
| *TBC1D10C* | 0.216359 | 0.093217 | 2.32102 | 0.02028578 | 0.236397749 |
| *PDE6B* | -0.429582 | 0.185108 | -2.320702 | 0.02030295 | 0.236397749 |
| *NCK2* | 0.367291 | 0.158301 | 2.320214 | 0.02032932 | 0.236397749 |
| *FAM13A* | -0.824861 | 0.355512 | -2.320207 | 0.02032969 | 0.236397749 |
| *TAF12* | 0.193221 | 0.083303 | 2.319503 | 0.0203678 | 0.236565601 |
| *ENSG00000228506* | -1.41462 | 0.609927 | -2.319328 | 0.02037724 | NA |
| *UXT* | 0.094432 | 0.040719 | 2.319137 | 0.02038759 | 0.236565601 |
| *CCDC126* | 0.497156 | 0.214489 | 2.317862 | 0.02045681 | 0.23679632 |
| *NOL9* | -0.345235 | 0.14895 | -2.317789 | 0.02046077 | 0.23679632 |
| *NIPAL3* | 0.33358 | 0.143935 | 2.317569 | 0.02047274 | 0.23679632 |
| *ENSG00000259976* | 0.653086 | 0.282147 | 2.3147 | 0.02062935 | 0.238107566 |
| *DSTN* | -0.165202 | 0.071373 | -2.314634 | 0.02063299 | 0.238107566 |
| *RESF1* | 0.214164 | 0.092567 | 2.313614 | 0.0206889 | 0.238107566 |
| *SRP9* | -0.165677 | 0.071611 | -2.313557 | 0.02069205 | 0.238107566 |
| *TUBGCP5* | -0.592635 | 0.256165 | -2.313494 | 0.02069549 | 0.238107566 |
| *C3orf14* | 1.505504 | 0.651199 | 2.311894 | 0.02078355 | 0.238592996 |
| *FAR1* | -0.394609 | 0.170705 | -2.311645 | 0.02079724 | 0.238592996 |
| *ZNF397* | 0.267439 | 0.115697 | 2.311533 | 0.02080345 | 0.238592996 |
| *C20orf27* | -0.225854 | 0.097747 | -2.310611 | 0.02085438 | 0.238925292 |
| *DCLRE1C* | 0.205137 | 0.088806 | 2.309928 | 0.02089215 | 0.239068425 |
| *ROCK1* | 0.159337 | 0.068997 | 2.30932 | 0.02092583 | 0.239068425 |
| *ELMO1* | 0.293779 | 0.127221 | 2.309195 | 0.02093276 | 0.239068425 |
| *TFEC* | 0.87407 | 0.378623 | 2.308549 | 0.02096862 | 0.239083173 |
| *MGAT5B* | -1.185703 | 0.513652 | -2.30838 | 0.02097799 | 0.239083173 |
| *C2orf42* | 0.478185 | 0.207202 | 2.307817 | 0.02100931 | 0.239189625 |
| *SUZ12* | 0.227188 | 0.098552 | 2.305257 | 0.0211522 | 0.240564835 |
| *CPNE3* | -0.227969 | 0.098938 | -2.30416 | 0.02121366 | 0.240912648 |
| *PRPSAP2* | 0.297002 | 0.128911 | 2.303921 | 0.02122705 | 0.240912648 |
| *TOLLIP.AS1* | -1.484193 | 0.644298 | -2.30358 | 0.02124623 | NA |
| *ENSG00000262580* | 1.215657 | 0.527778 | 2.303349 | 0.02125921 | NA |
| *ENSG00000238142* | 0.453114 | 0.196777 | 2.302674 | 0.02129722 | 0.24145718 |
| *MAP2K3* | -0.226434 | 0.098369 | -2.301876 | 0.02134215 | 0.241714861 |
| *TLE6* | -2.663642 | 1.1576 | -2.301004 | 0.02139139 | NA |
| *NDRG3* | 0.375065 | 0.163132 | 2.299156 | 0.02149609 | 0.24320526 |
| *PIEZO1* | 0.445128 | 0.193654 | 2.298578 | 0.02152892 | 0.243323712 |
| *KLHDC3* | -0.208757 | 0.090845 | -2.297949 | 0.02156467 | 0.243414673 |
| *NUCB2* | -0.64808 | 0.282062 | -2.29765 | 0.02158169 | 0.243414673 |
| *SIDT1.AS1* | -0.719195 | 0.313252 | -2.295898 | 0.02168173 | 0.24417482 |
| *BLNK* | 0.255481 | 0.111288 | 2.295684 | 0.02169396 | 0.24417482 |
| *ADARB1* | -0.438637 | 0.191115 | -2.295149 | 0.0217246 | 0.24426708 |
| *RAF1* | 0.247854 | 0.108048 | 2.293912 | 0.02179553 | 0.244811702 |
| *GOLPH3L* | 0.457726 | 0.199667 | 2.29244 | 0.02188029 | 0.245510366 |
| *ERGIC1* | -0.223546 | 0.097536 | -2.291938 | 0.02190924 | 0.245535904 |
| *LPCAT1* | 0.419561 | 0.183085 | 2.291618 | 0.02192768 | 0.245535904 |
| *GABPB1* | -0.231923 | 0.101246 | -2.290696 | 0.02198102 | 0.245864101 |
| *PTPRN2* | 0.448723 | 0.195921 | 2.29033 | 0.02200217 | 0.245864101 |
| *PSME4* | -0.277673 | 0.121331 | -2.288556 | 0.02210518 | 0.246761829 |

| | | | | | |
|---|---|---|---|---|---|
| *SENP1* | -0.469863 | 0.205346 | -2.288151 | 0.02212871 | 0.246771378 |
| *TRA2B* | -0.182702 | 0.079879 | -2.28725 | 0.02218126 | 0.246847181 |
| *SGCE* | 0.893512 | 0.390653 | 2.287226 | 0.02218264 | 0.246847181 |
| *IL9R* | 1.197403 | 0.523653 | 2.286633 | 0.02221728 | 0.246847181 |
| *ARHGAP18* | 0.519619 | 0.227257 | 2.286479 | 0.02222623 | 0.246847181 |
| *THOC3* | -0.416684 | 0.182376 | -2.284752 | 0.02232737 | 0.247717688 |
| *PIM1* | -0.502911 | 0.220187 | -2.284022 | 0.02237027 | 0.247913827 |
| *ABHD11* | -0.39269 | 0.172002 | -2.283057 | 0.02242702 | 0.247913827 |
| *SDAD1* | -0.203163 | 0.089004 | -2.282626 | 0.02245243 | 0.247913827 |
| *C2orf76* | -0.339066 | 0.148558 | -2.28238 | 0.02246692 | 0.247913827 |
| *MSMO1* | 0.224658 | 0.09844 | 2.282183 | 0.02247854 | 0.247913827 |
| *DTX3L* | 0.27911 | 0.122302 | 2.282129 | 0.02248171 | 0.247913827 |
| *ZSWIM7* | -0.312498 | 0.136973 | -2.281467 | 0.02252083 | 0.248093836 |
| *BANK1* | 0.382661 | 0.167783 | 2.280686 | 0.02256706 | 0.24835178 |
| *RASA3* | 0.634116 | 0.278242 | 2.279013 | 0.02266631 | 0.24896572 |
| *TTC17* | -0.249566 | 0.109525 | -2.278632 | 0.02268896 | 0.24896572 |
| *LEPROTL1* | -0.329022 | 0.144404 | -2.278489 | 0.02269745 | 0.24896572 |
| *DUS3L* | -0.392644 | 0.172373 | -2.277882 | 0.02273362 | 0.24896572 |
| *RPP40* | -0.449814 | 0.197498 | -2.277558 | 0.02275295 | 0.24896572 |
| *MAN2B1* | -0.286357 | 0.125736 | -2.277438 | 0.0227601 | 0.24896572 |
| *ERICH1* | 0.170391 | 0.074871 | 2.275784 | 0.02285893 | 0.249538639 |
| *PSMG4* | -0.239379 | 0.105198 | -2.275514 | 0.02287514 | 0.249538639 |
| *LINC01684* | -0.695435 | 0.30563 | -2.275412 | 0.02288125 | 0.249538639 |
| *LTBP4* | 0.468703 | 0.20606 | 2.274596 | 0.0229302 | 0.249643967 |
| *ALG5* | -0.224343 | 0.098635 | -2.274486 | 0.02293679 | 0.249643967 |
| *USP54* | -0.92714 | 0.407818 | -2.273415 | 0.02300116 | 0.25009448 |
| *PPFIBP2* | 0.366772 | 0.161406 | 2.272361 | 0.02306472 | 0.250535358 |
| *MYO9B* | 0.277595 | 0.122301 | 2.269764 | 0.0232219 | 0.251922842 |
| *FAM241A* | -0.482224 | 0.212486 | -2.269437 | 0.02324178 | 0.251922842 |
| *LINC01480* | -0.512778 | 0.225982 | -2.269106 | 0.02326189 | 0.251922842 |
| *CD83* | -0.349592 | 0.154093 | -2.268712 | 0.02328586 | 0.251931654 |
| *RNF167* | 0.178144 | 0.0786 | 2.266455 | 0.02342356 | 0.253169815 |
| *FASTKD2* | -0.342749 | 0.151279 | -2.265681 | 0.02347094 | 0.253181499 |
| *OSBPL1A* | -1.090993 | 0.481533 | -2.265666 | 0.02347183 | 0.253181499 |
| *DUSP18* | 0.430548 | 0.190063 | 2.265298 | 0.02349442 | 0.253181499 |
| *HNRNPK* | -0.120148 | 0.053052 | -2.264729 | 0.0235293 | 0.253214203 |
| *ERI1* | 0.436387 | 0.192709 | 2.264484 | 0.02354435 | 0.253214203 |
| *TEX30* | 0.29607 | 0.130784 | 2.263801 | 0.02358633 | 0.253214203 |
| *C1orf216* | 0.36741 | 0.162335 | 2.263292 | 0.02361769 | 0.253214203 |
| *SMAD2* | 0.271318 | 0.11989 | 2.263051 | 0.02363255 | 0.253214203 |
| *AKT3* | 0.468315 | 0.206946 | 2.262978 | 0.02363705 | 0.253214203 |
| *SPRYD4* | -0.378497 | 0.16746 | -2.260225 | 0.02380728 | 0.254541243 |
| *PXDC1* | -0.664004 | 0.293823 | -2.259882 | 0.02382859 | 0.254541243 |
| *CNKSR2* | 0.849597 | 0.376065 | 2.259173 | 0.02387261 | 0.254541243 |
| *ZNF318* | 0.352542 | 0.156049 | 2.259169 | 0.02387289 | 0.254541243 |
| *IER2* | -0.153674 | 0.068032 | -2.258839 | 0.02389342 | 0.254541243 |
| *MAP2K1* | 0.270838 | 0.119908 | 2.258713 | 0.02390125 | 0.254541243 |

| | | | | | |
|---|---|---|---|---|---|
| *MZB1* | -0.401087 | 0.177841 | -2.255317 | 0.02411344 | 0.256550013 |
| *ELP6* | -0.304793 | 0.135172 | -2.254852 | 0.02414265 | 0.256609963 |
| *SLC35D2* | 1.273117 | 0.56478 | 2.254184 | 0.02418461 | NA |
| *ICAM3* | 0.170265 | 0.075605 | 2.252022 | 0.02432091 | 0.25820965 |
| *ZNF860* | 0.636377 | 0.282684 | 2.251197 | 0.02437307 | 0.25820965 |
| *RNF123* | -0.498106 | 0.22128 | -2.251018 | 0.02438443 | 0.25820965 |
| *ZFC3H1* | 0.288466 | 0.128153 | 2.25096 | 0.02438805 | 0.25820965 |
| *MRFAP1L1* | 0.273865 | 0.121715 | 2.25006 | 0.02444511 | 0.258382079 |
| *ZBTB4* | 0.414285 | 0.18413 | 2.249955 | 0.02445181 | 0.258382079 |
| *SRSF2* | -0.173019 | 0.076941 | -2.248727 | 0.02452984 | 0.258730444 |
| *RBM26.AS1* | -0.642821 | 0.285891 | -2.248482 | 0.02454546 | 0.258730444 |
| *MAPK9* | 0.450942 | 0.200569 | 2.248315 | 0.0245561 | 0.258730444 |
| *EYA3* | 0.575063 | 0.255901 | 2.247212 | 0.02462649 | 0.259037028 |
| *LPP* | 0.334735 | 0.148975 | 2.24692 | 0.02464515 | 0.259037028 |
| *ZNF16* | -0.365768 | 0.162799 | -2.246741 | 0.02465659 | 0.259037028 |
| *TMEM101* | 0.27397 | 0.121989 | 2.245849 | 0.0247137 | 0.259386623 |
| *DGKG* | 1.814856 | 0.808119 | 2.245776 | 0.02471833 | NA |
| *EID1* | -0.149974 | 0.066795 | -2.245297 | 0.02474908 | 0.259507737 |
| *TENT4B* | 0.420864 | 0.187516 | 2.244424 | 0.02480515 | 0.259726917 |
| *SIGMAR1* | -0.314892 | 0.140312 | -2.244228 | 0.02481771 | 0.259726917 |
| *ENSG00000267364* | 1.207889 | 0.538403 | 2.243467 | 0.02486672 | 0.25998981 |
| *PPP1R13L* | 0.762148 | 0.339908 | 2.242218 | 0.02494729 | 0.260505171 |
| *POMGNT1* | -0.293187 | 0.130773 | -2.241961 | 0.02496388 | 0.260505171 |
| *KLHDC7B* | -0.644262 | 0.287528 | -2.240692 | 0.02504601 | 0.261111858 |
| *SEPHS1* | -0.407634 | 0.181971 | -2.240107 | 0.02508397 | 0.261257359 |
| *ENSG00000267939* | -1.613054 | 0.720179 | -2.239796 | 0.02510418 | NA |
| *ARL5A* | 0.177759 | 0.079373 | 2.239555 | 0.02511982 | 0.261380659 |
| *SH2D2A* | -0.625106 | 0.2794 | -2.237313 | 0.02526589 | 0.262649451 |
| *ZNF362* | -0.536652 | 0.23993 | -2.236707 | 0.02530551 | 0.262775796 |
| *CMTM8* | -1.163331 | 0.520183 | -2.236389 | 0.02532633 | 0.262775796 |
| *PHOSPHO2* | 0.520158 | 0.232737 | 2.234962 | 0.02541985 | 0.263494875 |
| *PRR4* | 0.426181 | 0.190768 | 2.234025 | 0.02548144 | 0.263882029 |
| *ENSG00000268568* | -0.818675 | 0.366762 | -2.232171 | 0.02560367 | 0.264895723 |
| *FAM160B2* | -0.476232 | 0.213483 | -2.230774 | 0.02569611 | 0.265599676 |
| *TCF3* | 0.217504 | 0.097572 | 2.229157 | 0.02580349 | 0.266456565 |
| *ZCCHC10* | 0.212387 | 0.09531 | 2.228393 | 0.02585431 | 0.266567859 |
| *PARS2* | 0.7457 | 0.334667 | 2.228187 | 0.02586807 | 0.266567859 |
| *NOTCH2* | 0.277182 | 0.124415 | 2.227892 | 0.02588775 | 0.266567859 |
| *NME2* | -0.327136 | 0.146889 | -2.227101 | 0.02594055 | 0.266625977 |
| *NT5DC4* | 0.780724 | 0.350561 | 2.227073 | 0.02594238 | 0.266625977 |
| *ENSG00000262165* | -1.705363 | 0.765746 | -2.227061 | 0.02594321 | NA |
| *AIMP2* | -0.319103 | 0.143414 | -2.225045 | 0.02607824 | 0.267640928 |
| *CBX3* | -0.145456 | 0.065377 | -2.224865 | 0.02609032 | 0.267640928 |
| *BCL2L11* | -0.338002 | 0.15199 | -2.223839 | 0.02615927 | 0.268095575 |
| *SIGLEC14* | 0.835957 | 0.376022 | 2.223159 | 0.02620507 | 0.268312297 |
| *TGFBR2* | 0.198349 | 0.089262 | 2.222101 | 0.02627646 | 0.268554337 |
| *PPIL2* | -0.338824 | 0.152481 | -2.222078 | 0.02627806 | 0.268554337 |

| | | | | | |
|---|---|---|---|---|---|
| *P4HB* | -0.153552 | 0.069139 | -2.220914 | 0.02635681 | 0.269071758 |
| *KDM1B* | 0.724752 | 0.326377 | 2.220599 | 0.02637813 | 0.269071758 |
| *IGFBP4* | -0.841472 | 0.379211 | -2.219009 | 0.0264861 | 0.269920176 |
| *FBXO8* | 0.427447 | 0.192764 | 2.217468 | 0.02659111 | 0.2705651 |
| *CYP3A5* | 0.536829 | 0.242107 | 2.217319 | 0.02660133 | 0.2705651 |
| *NFE2L2* | 0.176376 | 0.079557 | 2.216987 | 0.02662396 | 0.2705651 |
| *SH2D3C* | 0.416807 | 0.188051 | 2.216454 | 0.02666043 | 0.270682906 |
| *ZNF285* | 0.802175 | 0.362041 | 2.215701 | 0.02671196 | 0.270953345 |
| *AGFG2* | 0.79636 | 0.359832 | 2.213142 | 0.02688786 | 0.272301643 |
| *C6orf203* | 0.331907 | 0.149978 | 2.21304 | 0.02689492 | 0.272301643 |
| *TMSB4Y* | -1.220146 | 0.551932 | -2.210679 | 0.02705807 | 0.273698957 |
| *FAM214B* | 0.728775 | 0.330134 | 2.207509 | 0.02727849 | 0.274984419 |
| *HAUS1* | -0.205091 | 0.09291 | -2.207416 | 0.02728504 | 0.274984419 |
| *UCK2* | -0.373644 | 0.169293 | -2.207083 | 0.02730827 | 0.274984419 |
| *TRMT1L* | 0.390759 | 0.177052 | 2.207029 | 0.02731205 | 0.274984419 |
| *ENSG00000245869* | 0.501748 | 0.227355 | 2.206896 | 0.0273213 | 0.274984419 |
| *MRPS15* | -0.180286 | 0.081706 | -2.206529 | 0.027347 | 0.274984419 |
| *H2AFY* | -0.171287 | 0.077664 | -2.205495 | 0.02741939 | 0.274984419 |
| *IGKV1.12* | -0.721682 | 0.327235 | -2.205395 | 0.02742641 | 0.274984419 |
| *CHID1* | -0.268407 | 0.121706 | -2.205367 | 0.02742832 | 0.274984419 |
| *NRBF2* | 0.239368 | 0.108545 | 2.205232 | 0.02743781 | 0.274984419 |
| *ARMC7* | 0.408997 | 0.185543 | 2.204323 | 0.02750167 | 0.275370865 |
| *SLC27A3* | 0.820377 | 0.372311 | 2.203474 | 0.02756133 | 0.275714579 |
| *HIPK1* | 0.37359 | 0.169694 | 2.201555 | 0.02769677 | 0.276815106 |
| *P4HA1* | -0.36953 | 0.167943 | -2.200333 | 0.02778331 | 0.27716654 |
| *ARID3A* | 0.452174 | 0.205535 | 2.199987 | 0.02780782 | 0.27716654 |
| *ZNF430* | 0.232573 | 0.105748 | 2.199304 | 0.02785628 | 0.27716654 |
| *ENSG00000251364* | 1.016704 | 0.46229 | 2.199279 | 0.02785809 | 0.27716654 |
| *RABEPK* | -0.273592 | 0.124402 | -2.199263 | 0.02785926 | 0.27716654 |
| *PITPNM2* | -0.788615 | 0.358693 | -2.19858 | 0.0279078 | 0.277242904 |
| *HMG20B* | -0.161939 | 0.073672 | -2.198118 | 0.02794067 | 0.277242904 |
| *QTRT1* | -0.20914 | 0.095147 | -2.198081 | 0.02794335 | 0.277242904 |
| *NCAPH2* | -0.321046 | 0.146161 | -2.196528 | 0.02805417 | 0.277756874 |
| *FHOD1* | 0.383185 | 0.174473 | 2.196245 | 0.02807443 | 0.277756874 |
| *PRKACA* | 0.391523 | 0.178286 | 2.196039 | 0.02808916 | 0.277756874 |
| *DIS3L* | 0.449165 | 0.204544 | 2.195926 | 0.02809724 | 0.277756874 |
| *SSU72* | -0.1819 | 0.082874 | -2.1949 | 0.02817077 | 0.278116949 |
| *ZNF417* | 0.687691 | 0.313375 | 2.194467 | 0.02820184 | 0.278116949 |
| *AHCTF1* | 0.359915 | 0.164019 | 2.194349 | 0.02821032 | 0.278116949 |
| *POR* | -0.309238 | 0.140959 | -2.193812 | 0.02824893 | 0.278245531 |
| *MRFAP1* | 0.171336 | 0.078135 | 2.192831 | 0.02831954 | 0.278688866 |
| *THRB* | 1.643647 | 0.749602 | 2.192694 | 0.02832947 | NA |
| *ZNF225* | 0.649641 | 0.296801 | 2.188813 | 0.02861041 | 0.281296881 |
| *SNRPA1* | -0.213985 | 0.097785 | -2.188325 | 0.02864593 | 0.281391976 |
| *C15orf62* | -0.580335 | 0.265295 | -2.187509 | 0.02870537 | 0.281721573 |
| *RMND5B* | -0.291806 | 0.133426 | -2.187021 | 0.028741 | 0.281817119 |
| *SIAE* | 1.404104 | 0.642149 | 2.18657 | 0.02877394 | NA |

| | | | | | |
|---|---|---|---|---|---|
| *DCBLD2* | -0.969631 | 0.443716 | -2.185251 | 0.02887045 | 0.282540603 |
| *JRKL* | 0.526143 | 0.240823 | 2.184769 | 0.02890579 | 0.282540603 |
| *ENSG00000253140* | 1.373869 | 0.628843 | 2.184759 | 0.02890654 | 0.282540603 |
| *USP44* | 0.942871 | 0.431686 | 2.184161 | 0.02895039 | 0.282540603 |
| *UROS* | -0.293434 | 0.134354 | -2.184037 | 0.02895953 | 0.282540603 |
| *URB1.AS1* | -0.576235 | 0.263857 | -2.183887 | 0.02897054 | 0.282540603 |
| *DUSP2* | -0.454766 | 0.208342 | -2.182785 | 0.02905165 | 0.28291859 |
| *GABARAPL2* | 0.156619 | 0.071756 | 2.182654 | 0.02906128 | 0.28291859 |
| *HIST2H2BF* | 1.443723 | 0.661525 | 2.182418 | 0.02907873 | NA |
| *SERAC1* | -0.680878 | 0.312133 | -2.181369 | 0.02915611 | 0.283588122 |
| *SIRT7* | 0.221833 | 0.101769 | 2.179781 | 0.02927367 | 0.284307116 |
| *ELMSAN1* | 0.30635 | 0.140549 | 2.179665 | 0.02928228 | 0.284307116 |
| *LINC01678* | -0.398703 | 0.182998 | -2.178723 | 0.02935228 | 0.28458321 |
| *FAM120A* | 0.208974 | 0.095922 | 2.178578 | 0.02936301 | 0.28458321 |
| *AFF2* | -1.539875 | 0.707048 | -2.177894 | 0.02941396 | NA |
| *PDP2* | 0.664169 | 0.305141 | 2.176599 | 0.0295105 | 0.285754654 |
| *NADK2* | -0.48566 | 0.223163 | -2.176253 | 0.02953638 | 0.285754654 |
| *MRPL19* | -0.202806 | 0.093218 | -2.175601 | 0.02958508 | 0.2859716 |
| *TMF1* | 0.235138 | 0.108106 | 2.175074 | 0.02962458 | 0.285983262 |
| *ENSG00000259153* | 0.825537 | 0.379682 | 2.174286 | 0.02968365 | 0.285983262 |
| *ECHDC2* | 0.678637 | 0.31214 | 2.174146 | 0.02969416 | 0.285983262 |
| *SGPL1* | -0.286972 | 0.132004 | -2.173968 | 0.02970754 | 0.285983262 |
| *MCM8* | 0.555549 | 0.255562 | 2.173833 | 0.02971767 | 0.285983262 |
| *BRD3OS* | 0.320661 | 0.147662 | 2.171586 | 0.02988688 | 0.287319492 |
| *SUN1* | -0.224307 | 0.103306 | -2.171289 | 0.02990932 | 0.287319492 |
| *CMTM3* | -0.788146 | 0.363091 | -2.170656 | 0.02995717 | 0.287525452 |
| *ZNF107* | 0.415916 | 0.191796 | 2.168538 | 0.03011781 | 0.288812563 |
| *FANCL* | 0.535299 | 0.246892 | 2.168154 | 0.03014697 | 0.288837727 |
| *CYREN* | -0.219751 | 0.101437 | -2.166382 | 0.030282 | 0.289312401 |
| *SEC24D* | 0.513469 | 0.237064 | 2.165949 | 0.03031507 | 0.289312401 |
| *USP30* | -0.751546 | 0.347002 | -2.165825 | 0.03032456 | 0.289312401 |
| *PTMA* | -0.132137 | 0.061029 | -2.165153 | 0.03037594 | 0.289312401 |
| *CALR* | -0.176128 | 0.081347 | -2.165149 | 0.03037625 | 0.289312401 |
| *RBM19* | -0.260749 | 0.120438 | -2.165006 | 0.03038723 | 0.289312401 |
| *COG3* | -0.374917 | 0.173192 | -2.164745 | 0.03040722 | 0.289312401 |
| *HADHA* | 0.166765 | 0.077049 | 2.164393 | 0.03043417 | 0.289312401 |
| *KCP* | 1.043722 | 0.482228 | 2.164373 | 0.03043575 | 0.289312401 |
| *RNF32* | 1.082795 | 0.500489 | 2.163476 | 0.03050456 | NA |
| *NAP1L1* | 0.158312 | 0.073175 | 2.163461 | 0.03050575 | 0.289724757 |
| *TRIM32* | 0.780535 | 0.361279 | 2.160478 | 0.0307357 | 0.291362447 |
| *ZBTB37* | 0.439344 | 0.2034 | 2.160003 | 0.03077244 | 0.291362447 |
| *SLC4A7* | 0.382898 | 0.177278 | 2.159872 | 0.03078256 | 0.291362447 |
| *ARFGEF2* | 0.334734 | 0.154981 | 2.159837 | 0.03078526 | 0.291362447 |
| *ASAP1* | 0.43818 | 0.202954 | 2.159015 | 0.030849 | 0.291711961 |
| *ENSG00000260274* | -0.727068 | 0.336951 | -2.157784 | 0.03094466 | 0.292166368 |
| *SPATC1L* | -0.403092 | 0.186815 | -2.157705 | 0.03095074 | 0.292166368 |
| *PMF1* | -0.203886 | 0.094539 | -2.15663 | 0.0310345 | 0.292368533 |

| | | | | | |
|---|---|---|---|---|---|
| *MEST* | -0.404967 | 0.187784 | -2.156562 | 0.03103981 | 0.292368533 |
| *OSBPL3* | 0.616572 | 0.285927 | 2.156396 | 0.03105274 | 0.292368533 |
| *NUDT16L1* | -0.231286 | 0.107278 | -2.155941 | 0.03108825 | 0.292449903 |
| *SNHG19* | -0.272469 | 0.126435 | -2.155007 | 0.03116131 | 0.292884061 |
| *NAA16* | 0.438021 | 0.203326 | 2.154279 | 0.03121831 | 0.293166614 |
| *TMEM242* | -0.249928 | 0.116037 | -2.153866 | 0.0312507 | 0.293217789 |
| *KIAA1958* | -0.691241 | 0.321252 | -2.151712 | 0.03142003 | 0.294552583 |
| *MSANTD4* | 0.398821 | 0.185384 | 2.151321 | 0.03145089 | 0.294588209 |
| *NDUFS6* | -0.158064 | 0.073503 | -2.150437 | 0.03152063 | 0.294598454 |
| *S100A11* | 0.375498 | 0.174638 | 2.150153 | 0.03154308 | 0.294598454 |
| *RIPOR2* | 0.191997 | 0.089298 | 2.150076 | 0.03154919 | 0.294598454 |
| *BAZ2A* | 0.220993 | 0.102791 | 2.149922 | 0.03156139 | 0.294598454 |
| *NDUFAF4* | 0.228758 | 0.106419 | 2.149594 | 0.03158732 | 0.294598454 |
| *KLF13* | 0.25062 | 0.116648 | 2.148521 | 0.03167242 | 0.29499381 |
| *SLC38A2* | 0.20449 | 0.095191 | 2.148214 | 0.03169676 | 0.29499381 |
| *IFRD2* | -0.19669 | 0.091567 | -2.148034 | 0.03171102 | 0.29499381 |
| *LINC00339* | 0.788101 | 0.367112 | 2.146759 | 0.03181245 | 0.295396108 |
| *TMEM121* | -0.817756 | 0.380979 | -2.14646 | 0.03183629 | 0.295396108 |
| *KDM4D* | -1.509386 | 0.703333 | -2.146047 | 0.03186925 | NA |
| *ZNF384* | -0.362672 | 0.169017 | -2.145773 | 0.03189108 | 0.295396108 |
| *DPM1* | 0.21498 | 0.100192 | 2.145675 | 0.0318989 | 0.295396108 |
| *EIF2AK3* | -0.43099 | 0.200875 | -2.145563 | 0.03190784 | 0.295396108 |
| *LRRC56* | 0.791657 | 0.368994 | 2.145447 | 0.03191711 | 0.295396108 |
| *FAM43A* | 0.693924 | 0.323515 | 2.144952 | 0.03195665 | 0.295510799 |
| *SBNO1* | 0.177258 | 0.082665 | 2.144299 | 0.03200893 | 0.295742908 |
| *MOSPD3* | -0.265407 | 0.123805 | -2.143751 | 0.03205287 | 0.295897696 |
| *ARCN1* | 0.287918 | 0.134379 | 2.142576 | 0.03214714 | 0.296259619 |
| *DAGLB* | 0.515282 | 0.240555 | 2.142055 | 0.03218906 | 0.296259619 |
| *DNTTIP1* | 0.267349 | 0.12481 | 2.142045 | 0.03218989 | 0.296259619 |
| *UBR5* | 0.268147 | 0.125191 | 2.141907 | 0.03220095 | 0.296259619 |
| *MTRNR2L12* | -0.300746 | 0.140517 | -2.140281 | 0.03233209 | 0.296992567 |
| *FLYWCH1* | 0.416308 | 0.194514 | 2.140242 | 0.03233519 | 0.296992567 |
| *CD320* | -0.228405 | 0.106774 | -2.139158 | 0.03242285 | 0.29754662 |
| *TMSB10* | -0.125138 | 0.058527 | -2.138135 | 0.03250581 | 0.297880265 |
| *BAG6* | -0.211753 | 0.099041 | -2.138035 | 0.03251394 | 0.297880265 |
| *ATRAID* | -0.160855 | 0.075349 | -2.134811 | 0.03277646 | 0.299763754 |
| *CYTIP* | 0.249963 | 0.117091 | 2.134778 | 0.03277912 | 0.299763754 |
| *ENSG00000265206* | 0.25093 | 0.117559 | 2.134497 | 0.03280215 | 0.299763754 |
| *NAA10* | -0.171707 | 0.080469 | -2.133814 | 0.03285802 | 0.300022379 |
| *CHKB* | -0.355581 | 0.166745 | -2.132485 | 0.03296702 | 0.300574865 |
| *SERTAD1* | 0.372615 | 0.174739 | 2.132403 | 0.03297376 | 0.300574865 |
| *MYLK.AS1* | -0.604498 | 0.283568 | -2.131759 | 0.03302665 | 0.300794734 |
| *TRIM16* | -0.43312 | 0.203246 | -2.131018 | 0.0330877 | 0.300794734 |
| *TM9SF4* | -0.355591 | 0.166871 | -2.130937 | 0.03309433 | 0.300794734 |
| *TMX2* | 0.200479 | 0.094088 | 2.130766 | 0.03310842 | 0.300794734 |
| *TENT5A* | 0.442287 | 0.207883 | 2.127574 | 0.03337239 | 0.302768837 |
| *PDPR* | -0.542707 | 0.255096 | -2.127466 | 0.03338135 | 0.302768837 |

| | | | | | |
|---|---|---|---|---|---|
| *ISCA1* | -0.189378 | 0.08904 | -2.126881 | 0.03342994 | 0.302957071 |
| *HIST1H1B* | 1.429306 | 0.672256 | 2.126133 | 0.03349217 | NA |
| *VWCE* | -0.669377 | 0.314964 | -2.125253 | 0.0335655 | 0.303871832 |
| *NR6A1* | -0.514664 | 0.242195 | -2.124999 | 0.03358672 | 0.303871832 |
| *PDIA3* | -0.169025 | 0.079557 | -2.124585 | 0.03362126 | 0.303884195 |
| *RPS5* | 0.127103 | 0.059833 | 2.124296 | 0.03364538 | 0.303884195 |
| *MT.CO1* | 0.142723 | 0.067196 | 2.12398 | 0.03367184 | 0.303884195 |
| *ARF4.AS1* | -0.684189 | 0.322293 | -2.12288 | 0.03376388 | 0.304197039 |
| *ZNF468* | -0.413884 | 0.194967 | -2.122839 | 0.03376731 | 0.304197039 |
| *MDFIC* | 0.426475 | 0.200924 | 2.122565 | 0.03379036 | 0.304197039 |
| *JMJD8* | -0.311155 | 0.146645 | -2.121827 | 0.03385227 | 0.304502535 |
| *PHF13* | 0.680196 | 0.320723 | 2.120822 | 0.03393679 | 0.304881309 |
| *FAM168B* | 0.291286 | 0.137378 | 2.120327 | 0.03397851 | 0.304881309 |
| *GPSM3* | 0.147442 | 0.069541 | 2.120209 | 0.03398843 | 0.304881309 |
| *RNF26* | -0.241934 | 0.11412 | -2.119996 | 0.03400642 | 0.304881309 |
| *ZNF2* | 0.71944 | 0.339653 | 2.11816 | 0.03416156 | 0.30602009 |
| *SAMD4B* | 0.24813 | 0.117216 | 2.116864 | 0.03427137 | 0.306751314 |
| *FGFR1OP2* | 0.221093 | 0.104481 | 2.116119 | 0.03433466 | 0.306981429 |
| *SPRYD7* | 0.28116 | 0.13288 | 2.115898 | 0.03435349 | 0.306981429 |
| *POMP* | -0.152288 | 0.071998 | -2.115153 | 0.03441688 | 0.307040192 |
| *FAF2* | -0.281803 | 0.133269 | -2.114544 | 0.03446885 | 0.307040192 |
| *HDAC8* | -0.304031 | 0.143783 | -2.114508 | 0.03447188 | 0.307040192 |
| *ENSG00000260806* | -0.715066 | 0.338173 | -2.114496 | 0.03447291 | 0.307040192 |
| *ENSG00000188206* | 0.375135 | 0.177441 | 2.114137 | 0.03450358 | 0.307062087 |
| *SHLD1* | -0.284198 | 0.134498 | -2.113017 | 0.03459927 | 0.307662162 |
| *LYRM4* | -0.142288 | 0.067352 | -2.112603 | 0.03463475 | 0.307726226 |
| *SDF2L1* | -0.259178 | 0.122764 | -2.111185 | 0.03475641 | 0.308550389 |
| *EEF1E1* | -0.196266 | 0.092983 | -2.110768 | 0.03479227 | 0.308550389 |
| *ABHD3* | 0.326659 | 0.154776 | 2.110532 | 0.03481256 | 0.308550389 |
| *STK10* | 0.263097 | 0.12483 | 2.107649 | 0.03506139 | 0.310503005 |
| *WDR45* | -0.231757 | 0.11003 | -2.106298 | 0.03517845 | 0.311286387 |
| *LINC00937* | -1.35238 | 0.642501 | -2.104868 | 0.03530281 | NA |
| *ZNF276* | 0.315368 | 0.149838 | 2.104726 | 0.03531516 | 0.311929232 |
| *VPS37A* | 0.289076 | 0.137349 | 2.104683 | 0.03531888 | 0.311929232 |
| *ZFAND4* | 0.827942 | 0.39342 | 2.104474 | 0.03533708 | 0.311929232 |
| *HLA.DPA1* | 0.212132 | 0.100833 | 2.103791 | 0.03539671 | 0.312202409 |
| *MYO1D* | 0.928425 | 0.4414 | 2.103363 | 0.03543406 | NA |
| *ENSG00000006025* | 0.552209 | 0.262562 | 2.103155 | 0.03545226 | 0.312439155 |
| *FBXO25* | -0.296229 | 0.140906 | -2.102309 | 0.0355262 | 0.312837546 |
| *SQSTM1* | 0.191245 | 0.090991 | 2.101786 | 0.03557206 | 0.312988153 |
| *KPNB1* | 0.142089 | 0.067619 | 2.10131 | 0.03561377 | 0.313101994 |
| *TCOF1* | -0.199932 | 0.095193 | -2.100282 | 0.03570402 | 0.313642101 |
| *FERMT3* | 0.241829 | 0.115163 | 2.099882 | 0.03573924 | 0.313698273 |
| *POLR2E* | -0.16088 | 0.076626 | -2.09954 | 0.03576932 | 0.313709326 |
| *CXCR3* | 1.530395 | 0.72909 | 2.099049 | 0.03581259 | 0.313835969 |
| *SGPP2* | 1.202238 | 0.572893 | 2.098539 | 0.03585752 | 0.313970976 |
| *MAPKAPK2* | -0.265552 | 0.126561 | -2.09822 | 0.03588569 | 0.313970976 |

| | | | | | |
|---|---|---|---|---|---|
| *NAP1L6* | -1.315932 | 0.627204 | -2.098093 | 0.03589697 | NA |
| *ARHGEF1* | 0.222297 | 0.105968 | 2.09777 | 0.03592547 | 0.314066487 |
| *BLOC1S3* | 0.397707 | 0.189758 | 2.095857 | 0.03609484 | 0.315104456 |
| *MOB4* | 0.21293 | 0.101605 | 2.09566 | 0.03611233 | 0.315104456 |
| *WASH9P* | -0.320799 | 0.153093 | -2.09545 | 0.03613105 | 0.315104456 |
| *ZNF654* | 0.448546 | 0.214106 | 2.094967 | 0.03617391 | 0.315225683 |
| *RNF6* | 0.222172 | 0.106086 | 2.094258 | 0.03623697 | 0.315284403 |
| *UBXN8* | -0.401322 | 0.191631 | -2.09424 | 0.03623859 | 0.315284403 |
| *CEP295* | 0.383743 | 0.183275 | 2.093814 | 0.03627652 | 0.315362305 |
| *TWISTNB* | -0.259132 | 0.123801 | -2.093131 | 0.0363375 | 0.315597774 |
| *CLTC* | 0.218343 | 0.104327 | 2.092861 | 0.0363616 | 0.315597774 |
| *ZFYVE21* | 0.325943 | 0.155809 | 2.091933 | 0.0364445 | 0.316065303 |
| *ENSG00000255224* | -1.034855 | 0.495075 | -2.090299 | 0.03659099 | NA |
| *ZBTB21* | 0.4563 | 0.218372 | 2.089552 | 0.03665809 | 0.317664488 |
| *COL4A3* | 2.086439 | 0.998766 | 2.089017 | 0.03670622 | NA |
| *NDUFV2* | -0.128396 | 0.061467 | -2.088874 | 0.03671908 | 0.317737306 |
| *RAB33A* | 1.081183 | 0.517607 | 2.08881 | 0.03672487 | 0.317737306 |
| *SLC43A3* | -0.268852 | 0.128744 | -2.088272 | 0.03677333 | 0.317903839 |
| *MUS81* | 0.258835 | 0.123979 | 2.087728 | 0.03682239 | 0.318075328 |
| *PITPNM1* | 0.393947 | 0.18876 | 2.087024 | 0.03688595 | 0.318371695 |
| *SLC11A2* | 0.513263 | 0.246138 | 2.085268 | 0.03704498 | 0.319490928 |
| *CBWD6* | -0.729618 | 0.350072 | -2.084198 | 0.0371422 | 0.319840705 |
| *OFD1* | 0.260841 | 0.125153 | 2.084174 | 0.03714431 | 0.319840705 |
| *UGT8* | 0.526486 | 0.252764 | 2.082917 | 0.03725877 | 0.320572708 |
| *TOMM20* | 0.128609 | 0.061754 | 2.082591 | 0.03728855 | 0.32057546 |
| *TMCC1.AS1* | 1.091859 | 0.524354 | 2.082294 | 0.0373156 | NA |
| *ZNF354B* | 0.477306 | 0.229269 | 2.081864 | 0.03735491 | 0.32089251 |
| *TIMM50* | -0.232472 | 0.111697 | -2.081279 | 0.03740834 | 0.321098087 |
| *NAGPA* | 0.299106 | 0.143789 | 2.080175 | 0.0375095 | 0.321528531 |
| *SFT2D1* | -0.231577 | 0.11133 | -2.080087 | 0.03751757 | 0.321528531 |
| *HSD17B11* | 0.236607 | 0.113805 | 2.079062 | 0.03761163 | 0.321660172 |
| *WAC* | 0.201373 | 0.096861 | 2.078979 | 0.03761926 | 0.321660172 |
| *ZNF234* | 0.375102 | 0.180428 | 2.078954 | 0.03762159 | 0.321660172 |
| *DEF6* | 0.291164 | 0.140102 | 2.078234 | 0.03768778 | 0.321879798 |
| *STK17A* | 0.279152 | 0.134335 | 2.078032 | 0.03770643 | 0.321879798 |
| *MBOAT2* | -1.328035 | 0.639194 | -2.077672 | 0.03773961 | NA |
| *CCL4* | -0.836865 | 0.402818 | -2.077525 | 0.0377531 | 0.322025646 |
| *KPNA2* | -0.2304 | 0.110918 | -2.0772 | 0.03778307 | 0.32202892 |
| *SMIM12* | -0.150522 | 0.072544 | -2.074899 | 0.0379959 | 0.323588278 |
| *MRPL53* | -0.210799 | 0.10161 | -2.07458 | 0.03802549 | 0.323588278 |
| *BLCAP* | -0.258237 | 0.124579 | -2.072879 | 0.03818357 | 0.324337964 |
| *CHD7* | 0.240452 | 0.116004 | 2.072792 | 0.03819165 | 0.324337964 |
| *EP300* | 0.23007 | 0.11102 | 2.072325 | 0.03823518 | 0.324337964 |
| *USP53* | 0.574689 | 0.277324 | 2.072264 | 0.03824081 | 0.324337964 |
| *RAB7A* | -0.150477 | 0.072623 | -2.072031 | 0.03826258 | 0.324337964 |
| *DSCAML1* | -1.466393 | 0.707846 | -2.071627 | 0.03830028 | NA |
| *CNOT4* | 0.213743 | 0.10323 | 2.070543 | 0.03840157 | 0.324607027 |

| | | | | | |
|---|---|---|---|---|---|
| *CCDC106* | -0.306168 | 0.147877 | -2.070421 | 0.03841298 | 0.324607027 |
| *ZNF593* | -0.220241 | 0.106392 | -2.070087 | 0.03844422 | 0.324607027 |
| *MTF1* | 0.342291 | 0.165367 | 2.069891 | 0.03846256 | 0.324607027 |
| *CNN2* | 0.213138 | 0.10299 | 2.069497 | 0.03849952 | 0.324607027 |
| *TRMT112* | 0.225275 | 0.108856 | 2.069477 | 0.03850133 | 0.324607027 |
| *RBM17* | -0.153119 | 0.07399 | -2.069458 | 0.03850309 | 0.324607027 |
| *DDX18* | -0.117823 | 0.056948 | -2.068968 | 0.03854908 | 0.324743169 |
| *IGHV3.23* | -0.816429 | 0.394682 | -2.068572 | 0.0385863 | 0.324805331 |
| *ENSG00000272030* | -1.06707 | 0.51585 | -2.068566 | 0.03858688 | NA |
| *NAV2.AS3* | -1.574985 | 0.761396 | -2.068549 | 0.03858842 | NA |
| *SEC23A* | 0.265365 | 0.128333 | 2.067786 | 0.03866019 | 0.325175842 |
| *SMYD5* | -0.389446 | 0.188394 | -2.067194 | 0.03871586 | 0.325392584 |
| *COPB1* | 0.210727 | 0.10199 | 2.06615 | 0.03881428 | 0.325968032 |
| *MAD2L1BP* | 0.328359 | 0.15904 | 2.064629 | 0.0389581 | 0.326923608 |
| *RAPGEF5* | -1.458862 | 0.706863 | -2.063854 | 0.03903157 | NA |
| *OSBP* | 0.386787 | 0.187426 | 2.063681 | 0.03904798 | 0.32742539 |
| *CD79B* | -0.274297 | 0.133002 | -2.062356 | 0.03917386 | 0.327520276 |
| *MANBA* | -0.256172 | 0.124232 | -2.062046 | 0.03920334 | 0.327520276 |
| *RPN1* | -0.222503 | 0.107919 | -2.061766 | 0.03923004 | 0.327520276 |
| *MEA1* | -0.165622 | 0.080337 | -2.061591 | 0.03924665 | 0.327520276 |
| *TMEM183A* | -0.153282 | 0.074356 | -2.061467 | 0.0392585 | 0.327520276 |
| *GNA13* | 0.276344 | 0.134056 | 2.061414 | 0.03926353 | 0.327520276 |
| *SREBF1* | -0.441295 | 0.214081 | -2.061347 | 0.03926993 | 0.327520276 |
| *WBP1L* | -0.316554 | 0.153597 | -2.060938 | 0.03930895 | 0.327594654 |
| *MPHOSPH8* | 0.198649 | 0.096483 | 2.0589 | 0.0395038 | 0.328711855 |
| *DHX30* | -0.201792 | 0.098013 | -2.058821 | 0.03951138 | 0.328711855 |
| *ZNF354A* | -0.281657 | 0.136821 | -2.058581 | 0.0395344 | 0.328711855 |
| *BET1* | 0.239065 | 0.116148 | 2.058274 | 0.03956381 | 0.328711855 |
| *HLA.DQB2* | -1.655289 | 0.804389 | -2.057822 | 0.03960723 | NA |
| *SLC15A2* | 0.867654 | 0.4217 | 2.057513 | 0.03963689 | 0.329067789 |
| *ENSG00000183308* | -1.475592 | 0.71734 | -2.057034 | 0.03968296 | NA |
| *AP2S1* | -0.20105 | 0.097746 | -2.056861 | 0.03969956 | 0.329222621 |
| *CD40* | -0.142075 | 0.069095 | -2.056229 | 0.03976045 | 0.329222621 |
| *SNAPC4* | -0.340751 | 0.165727 | -2.056098 | 0.03977307 | 0.329222621 |
| *IAH1* | 0.132634 | 0.064523 | 2.055613 | 0.03981985 | 0.329222621 |
| *SYNC* | -0.331044 | 0.161063 | -2.055368 | 0.03984351 | 0.329222621 |
| *TP53INP1* | 0.496324 | 0.241478 | 2.055365 | 0.03984377 | 0.329222621 |
| *RANBP2* | 0.226541 | 0.110232 | 2.055121 | 0.03986727 | 0.329222621 |
| *GM2A* | 0.346106 | 0.168564 | 2.053262 | 0.0400472 | 0.330244592 |
| *REL* | -0.238591 | 0.116203 | -2.053215 | 0.04005172 | 0.330244592 |
| *ATP6V1G1* | 0.164226 | 0.080041 | 2.051764 | 0.04019258 | 0.33051498 |
| *ENSG00000254281* | -1.124641 | 0.548217 | -2.051453 | 0.04022287 | NA |
| *PEX6* | 0.696786 | 0.339672 | 2.051348 | 0.0402331 | 0.33051498 |
| *EEA1* | 0.283789 | 0.138378 | 2.050834 | 0.04028311 | 0.33051498 |
| *ENSG00000231609* | -0.751287 | 0.366347 | -2.05075 | 0.04029129 | 0.33051498 |
| *WDR18* | -0.224129 | 0.109296 | -2.050667 | 0.04029943 | 0.33051498 |
| *ZNF451* | 0.27914 | 0.136159 | 2.050098 | 0.0403549 | 0.33051498 |

| | | | | | |
|---|---|---|---|---|---|
| *LINC01006* | 0.45486 | 0.221875 | 2.05007 | 0.04035756 | 0.33051498 |
| *FKBP5* | 0.446788 | 0.217964 | 2.049821 | 0.04038194 | 0.33051498 |
| *PCMTD2* | -0.401156 | 0.195737 | -2.049469 | 0.0404163 | 0.33051498 |
| *DCAF8* | -0.223977 | 0.109287 | -2.049446 | 0.04041849 | 0.33051498 |
| *MAST3* | 0.395336 | 0.192899 | 2.049446 | 0.04041855 | 0.33051498 |
| *PPP4C* | -0.106648 | 0.052057 | -2.048698 | 0.04049167 | 0.330864334 |
| *RAB23* | 1.149433 | 0.561107 | 2.04851 | 0.04051007 | NA |
| *CLIP1* | 0.423952 | 0.207032 | 2.04776 | 0.04058348 | 0.331365788 |
| *LINC00158* | 0.621479 | 0.303674 | 2.046535 | 0.04070376 | 0.33154185 |
| *NSMF* | -0.467396 | 0.228425 | -2.046165 | 0.04074011 | 0.33154185 |
| *ZNF714* | 0.452005 | 0.220945 | 2.045785 | 0.04077751 | 0.33154185 |
| *CEP78* | -0.301331 | 0.147332 | -2.045249 | 0.04083036 | 0.33154185 |
| *RALB* | 0.261558 | 0.127913 | 2.044802 | 0.04087437 | 0.33154185 |
| *DOK1* | 0.342455 | 0.167496 | 2.044561 | 0.04089815 | 0.33154185 |
| *TAPBP* | -0.168321 | 0.082328 | -2.04453 | 0.04090121 | 0.33154185 |
| *EGR2* | -0.485173 | 0.237361 | -2.044027 | 0.04095086 | 0.33154185 |
| *TNFSF12* | 0.406039 | 0.198662 | 2.043871 | 0.04096628 | 0.33154185 |
| *ATP5PF* | -0.145813 | 0.071358 | -2.043407 | 0.04101215 | 0.33154185 |
| *FAM49B* | 0.136804 | 0.06695 | 2.043388 | 0.04101403 | 0.33154185 |
| *ENSG00000266962* | 0.32115 | 0.157174 | 2.043277 | 0.04102499 | 0.33154185 |
| *PDE7A* | 0.230438 | 0.112785 | 2.043168 | 0.04103584 | 0.33154185 |
| *SCAND1* | -0.168606 | 0.082574 | -2.041861 | 0.04116529 | 0.33154185 |
| *AP2M1* | -0.150334 | 0.073636 | -2.041588 | 0.04119242 | 0.33154185 |
| *ENSG00000262049* | -0.375738 | 0.184081 | -2.041155 | 0.04123545 | 0.33154185 |
| *TCIRG1* | 0.337821 | 0.165528 | 2.040868 | 0.04126396 | 0.33154185 |
| *SLC2A4RG* | 0.889086 | 0.43573 | 2.040451 | 0.04130547 | 0.33154185 |
| *TBC1D14* | -0.296489 | 0.145318 | -2.04028 | 0.04132245 | 0.33154185 |
| *DEGS1* | 0.287689 | 0.141008 | 2.040229 | 0.04132753 | 0.33154185 |
| *IL2RB* | 1.05746 | 0.518333 | 2.040117 | 0.04133869 | 0.33154185 |
| *GTF2IRD2* | -0.382595 | 0.18758 | -2.039639 | 0.04138629 | 0.33154185 |
| *IL12RB1* | 0.427373 | 0.209548 | 2.039498 | 0.04140033 | 0.33154185 |
| *SHF* | -0.519376 | 0.254671 | -2.039399 | 0.04141018 | 0.33154185 |
| *SP140* | 0.270877 | 0.132826 | 2.039341 | 0.041416 | 0.33154185 |
| *CIAO1* | -0.204919 | 0.100486 | -2.039271 | 0.04142296 | 0.33154185 |
| *ENSG00000179094* | -0.506302 | 0.248281 | -2.039226 | 0.0414275 | 0.33154185 |
| *GPR65* | 0.222614 | 0.109208 | 2.038432 | 0.04150678 | 0.331932217 |
| *ENSG00000260018* | -0.771665 | 0.378845 | -2.036891 | 0.04166096 | 0.332865555 |
| *LNPEP* | 0.248281 | 0.121906 | 2.036655 | 0.04168465 | 0.332865555 |
| *EAPP* | -0.214149 | 0.105196 | -2.035721 | 0.04177837 | 0.333296344 |
| *IGHV3.15* | -1.256005 | 0.616991 | -2.035695 | 0.041781 | NA |
| *CLEC11A* | -0.67367 | 0.330976 | -2.035405 | 0.04181018 | 0.333296344 |
| *ATOX1* | -0.263233 | 0.129354 | -2.034982 | 0.04185268 | 0.333296344 |
| *PPP2R5C* | -0.165233 | 0.0812 | -2.034898 | 0.04186109 | 0.333296344 |
| *JOSD1* | -0.301589 | 0.148274 | -2.033991 | 0.04195254 | 0.33378032 |
| *TBX6* | 1.081285 | 0.53179 | 2.033295 | 0.04202276 | NA |
| *NMT2* | -0.46911 | 0.230782 | -2.032696 | 0.04208322 | 0.334412741 |
| *CELF1* | 0.153868 | 0.0757 | 2.032595 | 0.04209348 | 0.334412741 |

| | | | | | |
|---|---|---|---|---|---|
| *CAB39* | 0.312177 | 0.153662 | 2.03158 | 0.0421962 | 0.334984305 |
| *CD58* | -0.306539 | 0.150948 | -2.030764 | 0.04227894 | 0.335157169 |
| *CASZ1* | -0.572284 | 0.28184 | -2.030532 | 0.04230253 | 0.335157169 |
| *CWC15* | -0.13922 | 0.068575 | -2.030171 | 0.04233921 | 0.335157169 |
| *FNIP1* | -0.26044 | 0.128286 | -2.030151 | 0.04234115 | 0.335157169 |
| *CHMP2B* | 0.227296 | 0.112022 | 2.029025 | 0.04245575 | 0.335820008 |
| *UQCC2* | -0.21831 | 0.107717 | -2.026704 | 0.04269266 | 0.337304564 |
| *DCXR* | -0.262029 | 0.129296 | -2.02658 | 0.04270541 | 0.337304564 |
| *ZC3H11A* | 0.242236 | 0.119583 | 2.025682 | 0.04279738 | 0.337728274 |
| *TMEM14B* | -0.21476 | 0.106036 | -2.02536 | 0.04283036 | 0.337728274 |
| *NINJ2* | -0.492289 | 0.243088 | -2.025148 | 0.04285215 | 0.337728274 |
| *IKBKG* | 0.256981 | 0.12692 | 2.024749 | 0.04289317 | 0.337807005 |
| *PKNOX1* | -0.301903 | 0.149142 | -2.024263 | 0.0429431 | 0.337955678 |
| *LYPLAL1* | 0.297315 | 0.14694 | 2.023373 | 0.04303473 | 0.338432048 |
| *GSEC* | -1.47115 | 0.727157 | -2.023154 | 0.04305724 | NA |
| *DCAF15* | 0.386229 | 0.190913 | 2.023064 | 0.04306659 | 0.338438077 |
| *RNF4* | 0.22615 | 0.111806 | 2.022699 | 0.04310415 | 0.338488876 |
| *GIN1* | 0.463803 | 0.229363 | 2.022138 | 0.04316208 | 0.338699397 |
| *ENSG00000279576* | -0.490346 | 0.242573 | -2.021437 | 0.0432345 | 0.339023273 |
| *IRF4* | -0.221806 | 0.10978 | -2.020463 | 0.04333542 | 0.339406472 |
| *HRK* | -0.761769 | 0.377144 | -2.019835 | 0.04340048 | 0.339406472 |
| *NUTF2* | -0.137446 | 0.068049 | -2.019794 | 0.04340472 | 0.339406472 |
| *PPP1R12A* | 0.207362 | 0.102667 | 2.019756 | 0.04340872 | 0.339406472 |
| *RTL8A* | -0.301348 | 0.149249 | -2.019089 | 0.04347801 | 0.339406472 |
| *ZNF823* | 0.834231 | 0.413205 | 2.018927 | 0.04349477 | 0.339406472 |
| *PLEKHG2* | 0.72793 | 0.360565 | 2.018861 | 0.04350166 | 0.339406472 |
| *G6PC3* | -0.403898 | 0.20011 | -2.018375 | 0.04355227 | 0.339557918 |
| *CA11* | -0.279549 | 0.138555 | -2.017607 | 0.04363218 | 0.339804013 |
| *OSTC* | -0.144861 | 0.071803 | -2.017472 | 0.04364627 | 0.339804013 |
| *JAML* | -1.519208 | 0.753529 | -2.016125 | 0.04378688 | 0.340621284 |
| *RBM15B* | -0.314018 | 0.155795 | -2.015588 | 0.0438431 | 0.340621284 |
| *HPS1* | 0.207494 | 0.102946 | 2.015568 | 0.04384513 | 0.340621284 |
| *ALG1L2* | 1.465484 | 0.728132 | 2.012663 | 0.0441501 | 0.342745813 |
| *UBE2W* | 0.226182 | 0.112407 | 2.012172 | 0.04420185 | 0.342903014 |
| *KCTD18* | -0.308384 | 0.153367 | -2.010758 | 0.04435107 | 0.343558126 |
| *XPO4* | -0.297325 | 0.147868 | -2.010748 | 0.04435211 | 0.343558126 |
| *ZBTB1* | 0.326096 | 0.162221 | 2.010199 | 0.04441017 | 0.343558126 |
| *CENPX* | 0.15826 | 0.07873 | 2.010176 | 0.04441256 | 0.343558126 |
| *GTF2H2C* | -0.336497 | 0.167498 | -2.008963 | 0.0445411 | 0.34430777 |
| *RBM4* | -0.218645 | 0.108861 | -2.008474 | 0.044593 | 0.344464293 |
| *VRK2* | 0.323793 | 0.161381 | 2.006393 | 0.04481434 | 0.345650708 |
| *MRPS7* | -0.150362 | 0.07495 | -2.006158 | 0.04483936 | 0.345650708 |
| *RPL30* | 0.07545 | 0.03761 | 2.006135 | 0.04484186 | 0.345650708 |
| *MEGF6* | -0.991337 | 0.494381 | -2.005208 | 0.04494085 | NA |
| *MRPL34* | -0.12671 | 0.063226 | -2.004073 | 0.04506224 | 0.347103609 |
| *ENSG00000272008* | 1.082952 | 0.540495 | 2.003629 | 0.04510983 | NA |
| *ATF7IP2* | -0.259694 | 0.12965 | -2.003038 | 0.04517317 | 0.347445105 |

| | | | | | |
|---|---|---|---|---|---|
| *UBA6.AS1* | -0.466892 | 0.233097 | -2.002997 | 0.04517766 | 0.347445105 |
| *ZNF587* | 0.326344 | 0.162947 | 2.002767 | 0.04520234 | 0.347445105 |
| *TFEB* | 0.347029 | 0.173337 | 2.002048 | 0.04527953 | 0.347557653 |
| *CSRNP2* | 0.59878 | 0.299085 | 2.002036 | 0.04528085 | 0.347557653 |
| *ENSG00000224086* | -0.406208 | 0.202967 | -2.001349 | 0.04535481 | 0.347880011 |
| *TLR10* | 0.229825 | 0.114862 | 2.000882 | 0.04540509 | 0.348020389 |
| *TRAF5* | 0.245004 | 0.122484 | 2.000285 | 0.04546953 | 0.348269074 |
| *TATDN2* | 0.424612 | 0.212471 | 1.998445 | 0.04566841 | 0.348906735 |
| *ZFYVE1* | 0.535515 | 0.267968 | 1.998428 | 0.04567031 | 0.348906735 |
| *LARP4B* | 0.282928 | 0.141593 | 1.998184 | 0.04569675 | 0.348906735 |
| *CAMK2D* | -0.318462 | 0.159383 | -1.99809 | 0.04570688 | 0.348906735 |
| *KCTD3* | -0.964063 | 0.482493 | -1.998085 | 0.0457074 | NA |
| *LRRC14* | -0.375261 | 0.187815 | -1.998033 | 0.04571307 | 0.348906735 |
| *NOP58* | -0.134273 | 0.067218 | -1.99757 | 0.04576334 | 0.348939233 |
| *RASSF7* | 0.258881 | 0.129609 | 1.997403 | 0.04578144 | 0.348939233 |
| *PURA* | 0.297059 | 0.14877 | 1.996762 | 0.04585103 | 0.349148619 |
| *MAMLD1* | -1.386903 | 0.694614 | -1.996652 | 0.04586305 | NA |
| *RTCA* | 0.28336 | 0.141924 | 1.996559 | 0.04587307 | 0.349148619 |
| *ENSG00000203546* | -0.68703 | 0.344287 | -1.995514 | 0.04598681 | 0.349705126 |
| *ENSG00000225302* | 1.479244 | 0.741306 | 1.995458 | 0.04599295 | NA |
| *RPL39L* | 0.456604 | 0.228899 | 1.994782 | 0.04606664 | 0.349705126 |
| *GRAP* | 0.371712 | 0.186349 | 1.994703 | 0.0460753 | 0.349705126 |
| *GFM2* | 0.432767 | 0.216996 | 1.994358 | 0.04611289 | 0.349705126 |
| *CFLAR* | 0.260042 | 0.130405 | 1.994111 | 0.04613993 | 0.349705126 |
| *POLR2H* | -0.185951 | 0.093253 | -1.994056 | 0.04614594 | 0.349705126 |
| *ZC3H8* | -0.244595 | 0.122676 | -1.993826 | 0.0461711 | 0.349705126 |
| *OTUD1* | 0.54634 | 0.27423 | 1.99227 | 0.04634143 | 0.350330631 |
| *MRPL28* | -0.156484 | 0.078555 | -1.992037 | 0.04636699 | 0.350330631 |
| *SH3YL1* | -0.295353 | 0.148274 | -1.991938 | 0.04637789 | 0.350330631 |
| *FANCM* | -0.522481 | 0.262303 | -1.991896 | 0.04638244 | 0.350330631 |
| *PMF1.BGLAP* | -1.711529 | 0.859358 | -1.991636 | 0.046411 | NA |
| *RTN4* | 0.153864 | 0.07729 | 1.99074 | 0.04650944 | 0.350885153 |
| *CBX7* | 0.295546 | 0.148469 | 1.990627 | 0.04652186 | 0.350885153 |
| *POLR3A* | -0.346302 | 0.174015 | -1.99007 | 0.04658327 | 0.350885153 |
| *FAM49A* | 0.375152 | 0.188558 | 1.989586 | 0.04663659 | 0.350885153 |
| *DTWD1* | 0.228358 | 0.11478 | 1.989527 | 0.04664303 | 0.350885153 |
| *ENSG00000263884* | -1.56198 | 0.785146 | -1.989414 | 0.04665558 | NA |
| *GSTO2* | 1.769103 | 0.889352 | 1.989204 | 0.04667864 | NA |
| *CCDC66* | 0.314885 | 0.158297 | 1.989202 | 0.04667893 | 0.350885153 |
| *NECTIN1* | -0.864675 | 0.434689 | -1.989178 | 0.04668152 | 0.350885153 |
| *METTL26* | -0.196644 | 0.098877 | -1.988779 | 0.04672555 | 0.350973673 |
| *ACSL5* | 0.342347 | 0.172173 | 1.988389 | 0.04676863 | 0.35105155 |
| *MFSD13A* | -0.491484 | 0.247215 | -1.988085 | 0.0468023 | 0.35105155 |
| *ATP6V0A2* | -0.287689 | 0.144727 | -1.98781 | 0.04683268 | 0.35105155 |
| *IL6ST* | -0.443596 | 0.223218 | -1.987275 | 0.04689193 | 0.35108654 |
| *RYBP* | 0.383279 | 0.192875 | 1.987185 | 0.04690186 | 0.35108654 |
| *C9orf72* | -0.333596 | 0.167948 | -1.986308 | 0.0469991 | 0.351572622 |

| | | | | | |
|---|---|---|---|---|---|
| *ZNF439* | 0.272964 | 0.137529 | 1.984776 | 0.04716936 | 0.35245554 |
| *PRR3* | -0.254217 | 0.128106 | -1.984427 | 0.04720821 | 0.35245554 |
| *NBPF9* | 0.59719 | 0.300973 | 1.984199 | 0.0472337 | 0.35245554 |
| *CETP* | -0.618396 | 0.311698 | -1.983955 | 0.04726088 | 0.35245554 |
| *ENSG00000272221* | 0.843618 | 0.425255 | 1.983792 | 0.04727904 | 0.35245554 |
| *IMP3* | -0.162517 | 0.081941 | -1.983352 | 0.04732808 | 0.352579629 |
| *IFT22* | -0.276538 | 0.139454 | -1.983009 | 0.04736646 | 0.352624155 |
| *ENSG00000262292* | 0.81881 | 0.41303 | 1.982445 | 0.04742946 | 0.352838191 |
| *RBAK.RBAKDN* | 0.268423 | 0.13545 | 1.981716 | 0.04751108 | 0.352838191 |
| *SERTAD2* | 0.289154 | 0.145914 | 1.981674 | 0.04751578 | 0.352838191 |
| *ENSG00000273329* | -0.441899 | 0.223002 | -1.981592 | 0.04752488 | 0.352838191 |
| *DDX5* | -0.191589 | 0.096754 | -1.980172 | 0.04768424 | 0.353779996 |
| *TACC1* | 0.27281 | 0.137824 | 1.979407 | 0.04777017 | 0.354066456 |
| *CYP1B1* | 1.335432 | 0.674768 | 1.979099 | 0.04780484 | NA |
| *ENSG00000261386* | 0.530764 | 0.268235 | 1.978729 | 0.04784649 | 0.354066456 |
| *MTFP1* | -0.215783 | 0.109053 | -1.978692 | 0.04785074 | 0.354066456 |
| *HVCN1* | -0.280421 | 0.141734 | -1.978507 | 0.04787155 | 0.354066456 |
| *MED11* | 0.220799 | 0.111625 | 1.97805 | 0.04792311 | 0.354066456 |
| *PCTP* | -0.383395 | 0.193829 | -1.978009 | 0.04792775 | 0.354066456 |
| *SNRNP200* | -0.175482 | 0.088726 | -1.977806 | 0.04795057 | 0.354066456 |
| *CDC5L* | 0.188314 | 0.095288 | 1.976263 | 0.04812496 | 0.355113225 |
| *MYL12A* | 0.153001 | 0.077435 | 1.975859 | 0.04817075 | 0.355188817 |
| *QPRT* | -0.740432 | 0.374789 | -1.975597 | 0.04820047 | 0.355188817 |
| *KIF1C* | 0.56593 | 0.286601 | 1.974626 | 0.04831061 | 0.355759582 |
| *GLRX2* | 0.223504 | 0.113245 | 1.973633 | 0.0484235 | 0.35603483 |
| *PET100* | -0.136227 | 0.069025 | -1.973592 | 0.04842812 | 0.35603483 |
| *FAM98B* | -0.334184 | 0.169361 | -1.973207 | 0.048472 | 0.35603483 |
| *NUP50.DT* | -0.400603 | 0.203028 | -1.973147 | 0.04847883 | 0.35603483 |
| *CYTH3* | -0.64716 | 0.328071 | -1.972624 | 0.04853843 | 0.356223109 |
| *CAPN2* | 0.833107 | 0.422394 | 1.972348 | 0.04856993 | 0.356223109 |
| *INPP5D* | 0.20252 | 0.102716 | 1.971649 | 0.04864968 | 0.356567785 |
| *VAMP5* | -0.35476 | 0.180058 | -1.97025 | 0.04880973 | 0.357344465 |
| *STAT5A* | -0.258068 | 0.130989 | -1.970149 | 0.04882132 | 0.357344465 |
| *LBX2.AS1* | 1.453918 | 0.738623 | 1.968417 | 0.04902011 | NA |
| *MFSD4B* | 0.443491 | 0.225421 | 1.967391 | 0.04913816 | 0.359416393 |
| *ZNF91* | 0.229166 | 0.116499 | 1.967111 | 0.04917044 | 0.359416393 |
| *FAN1* | 0.580656 | 0.295352 | 1.965979 | 0.04930101 | 0.360128977 |
| *FUNDC2* | -0.190464 | 0.0969 | -1.965582 | 0.04934692 | 0.360222556 |
| *CPPED1* | 0.64959 | 0.330544 | 1.965217 | 0.04938917 | 0.360289362 |
| *FAM136A* | -0.164219 | 0.083589 | -1.964588 | 0.04946198 | 0.360578842 |
| *ENSG00000240859* | -0.989368 | 0.503729 | -1.964089 | 0.04951974 | NA |
| *CNST* | 0.339213 | 0.172708 | 1.964081 | 0.04952067 | 0.360765041 |
| *SPIN3* | 1.842443 | 0.938139 | 1.963933 | 0.04953783 | NA |
| *RSBN1L* | 0.178039 | 0.090686 | 1.963261 | 0.04961589 | 0.361136521 |
| *BTF3L4* | -0.184792 | 0.094146 | -1.962821 | 0.04966698 | 0.361136521 |
| *CREBBP* | 0.194204 | 0.098943 | 1.962785 | 0.0496712 | 0.361136521 |
| *ATAT1* | 0.325775 | 0.166026 | 1.962198 | 0.04973949 | 0.361391583 |

| | | | | | |
|---|---|---|---|---|---|
| *PRR7* | -0.411719 | 0.209936 | -1.961162 | 0.04986009 | 0.36184936 |
| *QPCTL* | -0.804294 | 0.410127 | -1.961086 | 0.04986899 | 0.36184936 |
| *EAF1* | -0.311075 | 0.15866 | -1.960644 | 0.04992056 | 0.361982234 |
