## Supplemental Table 6 for "Single-cell Landscape Analysis Unravels Molecular Programming of the Human B Cell Compartment in Chronic GVHD"

**Supplemental Table 6. All DEGs by cluster in the blood B cell scRNA-Seq dataset for ATRA-treated B cells.** Log2FC values represent the change in expression for the gene of interest in ATRA-treated B cells when compared to untreated B cells from all 8 allo-HCT patients combined, along with associated statistical data.

| Cluster 1 | log2FC | lfcSE | stat | pvalue | padj |
|---|---|---|---|---|---|
| *VPREB3* | 3.239544 | 0.091617 | 35.359826 | < 2.22e-16 | < 2.22e-16 |
| *AHI1* | 1.715068 | 0.138638 | 12.370799 | < 2.22e-16 | < 2.22e-16 |
| *PLAAT4* | 2.070436 | 0.176793 | 11.711066 | < 2.22e-16 | < 2.22e-16 |
| *RRBP1* | 1.952289 | 0.168521 | 11.584832 | < 2.22e-16 | < 2.22e-16 |
| *DUS2* | 1.59709 | 0.151949 | 10.510684 | < 2.22e-16 | < 2.22e-16 |
| *CD38* | 3.185154 | 0.336321 | 9.470584 | < 2.22e-16 | < 2.22e-16 |
| *DENND6B* | 2.261815 | 0.245054 | 9.22987 | < 2.22e-16 | < 2.22e-16 |
| *CD1D* | 3.322237 | 0.371023 | 8.954262 | < 2.22e-16 | 5.40E-16 |
| *SLC12A4* | 2.522336 | 0.289223 | 8.721086 | < 2.22e-16 | 3.87E-15 |
| *DPEP2* | 1.010012 | 0.121526 | 8.311097 | < 2.22e-16 | 1.20E-13 |
| *RARA.AS1* | 2.080188 | 0.261336 | 7.95983 | 1.72E-15 | 1.98E-12 |
| *LCAT* | 2.443899 | 0.311124 | 7.855068 | 4.00E-15 | 4.21E-12 |
| *AGO1* | 1.045363 | 0.13458 | 7.767574 | 8.00E-15 | 7.78E-12 |
| *RAB37* | 1.342816 | 0.176705 | 7.599198 | 2.98E-14 | 2.69E-11 |
| *VPREB1* | 4.622933 | 0.622501 | 7.426388 | 1.12E-13 | 9.40E-11 |
| *MRPS6* | 0.957351 | 0.131557 | 7.277101 | 3.41E-13 | 2.69E-10 |
| *C16orf54* | 1.165205 | 0.162778 | 7.158264 | 8.17E-13 | 6.07E-10 |
| *SLC5A3* | 1.030773 | 0.148455 | 6.943318 | 3.83E-12 | 2.69E-09 |
| *P2RX5* | 1.038367 | 0.15263 | 6.803172 | 1.02E-11 | 6.81E-09 |
| *SIGIRR* | 0.78388 | 0.117274 | 6.684187 | 2.32E-11 | 1.47E-08 |
| *RMDN2* | 1.974119 | 0.296347 | 6.661511 | 2.71E-11 | 1.63E-08 |
| *PNOC* | 1.665266 | 0.250408 | 6.65021 | 2.93E-11 | 1.68E-08 |
| *AGPAT5* | 0.888394 | 0.135542 | 6.55438 | 5.59E-11 | 3.07E-08 |
| *FAM3C* | 0.801948 | 0.123515 | 6.492692 | 8.43E-11 | 4.44E-08 |
| *MPEG1* | 1.614321 | 0.25443 | 6.344853 | 2.23E-10 | 1.13E-07 |
| *SEPTIN9* | 0.568186 | 0.090558 | 6.274284 | 3.51E-10 | 1.71E-07 |
| *FCRL1* | 1.113825 | 0.182636 | 6.098604 | 1.07E-09 | 5.01E-07 |
| *GPX1* | 0.568352 | 0.093355 | 6.088055 | 1.14E-09 | 5.16E-07 |
| *GRN* | 0.862574 | 0.142076 | 6.071216 | 1.27E-09 | 5.53E-07 |
| *MS4A1* | -0.60819 | 0.100407 | -6.05726 | 1.38E-09 | 5.83E-07 |
| *MAPKAPK2* | 1.028439 | 0.178178 | 5.771964 | 7.84E-09 | 3.19E-06 |
| *P2RX5.TAX1BP3* | 1.623417 | 0.288252 | 5.631932 | 1.78E-08 | 7.04E-06 |
| *GBP4* | 0.775912 | 0.142806 | 5.433327 | 5.53E-08 | 2.12E-05 |
| *SEL1L3* | 0.986761 | 0.183249 | 5.384804 | 7.25E-08 | 2.70E-05 |
| *XYLT1* | 1.234467 | 0.233752 | 5.281092 | 1.28E-07 | 4.64E-05 |
| *APOL3* | 0.821357 | 0.157845 | 5.203578 | 1.95E-07 | 6.86E-05 |
| *MYL2* | 4.501103 | 0.881083 | 5.108602 | 3.25E-07 | 0.00011083 |
| *FAM117A* | 0.697754 | 0.137884 | 5.060428 | 4.18E-07 | 0.00013909 |
| *IRAK2* | 1.101905 | 0.218317 | 5.047276 | 4.48E-07 | 0.00014519 |
| *ASB2* | 4.8577 | 0.97452 | 4.984712 | 6.21E-07 | 0.000196014 |
| *ENSG00000214797* | 2.590847 | 0.521524 | 4.967835 | 6.77E-07 | 0.000208646 |

| | | | | | |
|---|---|---|---|---|---|
| *CD72* | 0.972445 | 0.197063 | 4.934683 | 8.03E-07 | 0.000241512 |
| *RABGAP1L* | 0.787875 | 0.160196 | 4.918192 | 8.73E-07 | 0.000256659 |
| *DBNL* | 0.583976 | 0.120595 | 4.84246 | 1.28E-06 | 0.000368258 |
| *PLAAT3* | 3.245782 | 0.68321 | 4.750783 | 2.03E-06 | 0.000568941 |
| *PPP1R18* | 0.534823 | 0.115085 | 4.647187 | 3.36E-06 | 0.000924256 |
| *ICAM2* | 0.810193 | 0.175718 | 4.610751 | 4.01E-06 | 0.001078589 |
| *CHST15* | 0.876363 | 0.190651 | 4.596685 | 4.29E-06 | 0.001129952 |
| *TRABD* | 0.437682 | 0.095562 | 4.580065 | 4.65E-06 | 0.001198602 |
| *SPRY1* | 0.889388 | 0.195263 | 4.55483 | 5.24E-06 | 0.001324857 |
| *CD79B* | 0.281374 | 0.062569 | 4.497007 | 6.89E-06 | 0.00170738 |
| *MCL1* | 0.502184 | 0.113125 | 4.439203 | 9.03E-06 | 0.002193938 |
| *ATXN10* | 0.57549 | 0.130103 | 4.423342 | 9.72E-06 | 0.002316874 |
| *ETFB* | 0.547662 | 0.124261 | 4.407367 | 1.05E-05 | 0.002448264 |
| *SMC6* | -0.575894 | 0.131498 | -4.379475 | 1.19E-05 | 0.002732963 |
| *TNFRSF13B* | -1.642074 | 0.380392 | -4.316788 | 1.58E-05 | 0.003572002 |
| *ST3GAL1* | 0.879332 | 0.204035 | 4.309722 | 1.63E-05 | 0.003623368 |
| *LHFPL2* | 1.250313 | 0.291873 | 4.283752 | 1.84E-05 | 0.003947746 |
| *GPR183* | -0.615168 | 0.143628 | -4.283057 | 1.84E-05 | 0.003947746 |
| *SREBF1* | 1.408247 | 0.329841 | 4.269477 | 1.96E-05 | 0.004125996 |
| *ZNF563* | 1.007111 | 0.236517 | 4.258092 | 2.06E-05 | 0.00427062 |
| *PPP2R5C* | 0.432457 | 0.101785 | 4.248723 | 2.15E-05 | 0.00438135 |
| *PXK* | 0.504086 | 0.119392 | 4.222119 | 2.42E-05 | 0.004853766 |
| *SNX18* | 0.965772 | 0.230012 | 4.198787 | 2.68E-05 | 0.005297792 |
| *SLC2A5* | 1.180447 | 0.281795 | 4.189021 | 2.80E-05 | 0.005445882 |
| *GDPGP1* | 1.365762 | 0.329251 | 4.14809 | 3.35E-05 | 0.006418204 |
| *GLRX* | -0.497023 | 0.119937 | -4.144032 | 3.41E-05 | 0.006435413 |
| *SPOCK2* | 0.588104 | 0.143118 | 4.109215 | 3.97E-05 | 0.007376723 |
| *CABLES1* | -1.920071 | 0.468108 | -4.101773 | 4.10E-05 | 0.007507689 |
| *PUS1* | 0.899317 | 0.220029 | 4.087256 | 4.37E-05 | 0.00787892 |
| *NRM* | 0.645795 | 0.16107 | 4.009416 | 6.09E-05 | 0.010832141 |
| *PLEKHA2* | 0.38904 | 0.097543 | 3.9884 | 6.65E-05 | 0.011673421 |
| *CCL2* | 4.592547 | 1.154323 | 3.978563 | 6.93E-05 | 0.0120003 |
| *ENSG00000273319* | 0.69243 | 0.175228 | 3.951598 | 7.76E-05 | 0.013254993 |
| *SLC5A5* | 3.627945 | 0.920823 | 3.939892 | 8.15E-05 | 0.0137331 |
| *BTBD7* | 0.510624 | 0.12976 | 3.935144 | 8.31E-05 | 0.013823163 |
| *IRF4* | 0.900479 | 0.231307 | 3.893004 | 9.90E-05 | 0.01624674 |
| *NEURL2* | 1.992856 | 0.515316 | 3.867248 | 0.00011007 | 0.017462707 |
| *NCF1* | 0.279332 | 0.072236 | 3.866939 | 0.00011021 | 0.017462707 |
| *ENSG00000275799* | 0.608916 | 0.157499 | 3.866149 | 0.000110567 | 0.017462707 |
| *C12orf65* | 0.551271 | 0.142983 | 3.85551 | 0.000115489 | 0.018014788 |
| *CD82* | -0.558455 | 0.145518 | -3.837713 | 0.000124185 | 0.019124953 |
| *AP3B1* | 0.549058 | 0.143175 | 3.834866 | 0.000125633 | 0.019124953 |
| *NMT2* | -0.492979 | 0.128716 | -3.829968 | 0.00012816 | 0.019257731 |
| *HCK* | -0.970205 | 0.253495 | -3.827306 | 0.000129553 | 0.019257731 |
| *CD79A* | 0.424419 | 0.11279 | 3.762925 | 0.000167938 | 0.024673174 |
| *TAF4B* | 0.957342 | 0.255662 | 3.74456 | 0.00018071 | 0.026244523 |
| *ADAM19* | 0.675578 | 0.181298 | 3.726341 | 0.00019428 | 0.027690576 |

| | | | | | |
|---|---|---|---|---|---|
| *PWP1* | 0.393781 | 0.105703 | 3.725342 | 0.00019505 | 0.027690576 |
| *SMAD3* | 0.69559 | 0.188467 | 3.690778 | 0.000223569 | 0.031386638 |
| *ANAPC16* | 0.28018 | 0.076185 | 3.677639 | 0.000235403 | 0.032684785 |
| *FCER2* | 1.329128 | 0.36175 | 3.674161 | 0.000238633 | 0.03277307 |
| *IRF2BPL* | 1.159442 | 0.317642 | 3.650152 | 0.000262085 | 0.035460423 |
| *ENSG00000226380* | 0.809307 | 0.221821 | 3.648464 | 0.000263813 | 0.035460423 |
| *PSD3* | -1.23821 | 0.340215 | -3.639488 | 0.000273181 | 0.036333093 |
| *CLEC17A* | 0.804011 | 0.221566 | 3.628759 | 0.000284787 | 0.037482161 |
| *SLC25A5* | -0.261237 | 0.072351 | -3.610706 | 0.000305364 | 0.039776046 |
| *SIDT1* | -0.694796 | 0.192758 | -3.604495 | 0.000312761 | 0.040323775 |
| *H3F3A* | 0.311838 | 0.086677 | 3.597704 | 0.000321039 | 0.040972978 |
| *CALHM6* | 0.467378 | 0.130364 | 3.585167 | 0.000336862 | 0.042562567 |
| *PAOX* | 0.793202 | 0.221514 | 3.580819 | 0.000342519 | 0.042848803 |
| *CD180* | 0.729607 | 0.204661 | 3.564947 | 0.000363929 | 0.045080856 |
| *IFI30* | 0.62223 | 0.178034 | 3.495007 | 0.00047405 | 0.058056203 |
| *ENSG00000089127* | 0.683138 | 0.195581 | 3.492866 | 0.000477867 | 0.058056203 |
| *ARHGEF1* | 0.344339 | 0.099399 | 3.464226 | 0.000531759 | 0.063988347 |
| *PLPP3* | 3.212139 | 0.930052 | 3.453719 | 0.000552912 | 0.065906108 |
| *RASSF3* | 0.650355 | 0.188736 | 3.445853 | 0.000569261 | 0.067220664 |
| *CTNNB1* | 0.479561 | 0.13937 | 3.440933 | 0.000579712 | 0.067820985 |
| *IL1B* | 3.239503 | 0.942287 | 3.437916 | 0.00058621 | 0.067951984 |
| *CIB1* | 0.28996 | 0.08454 | 3.429843 | 0.00060393 | 0.069369612 |
| *ENSG00000261448* | 1.426513 | 0.416719 | 3.423197 | 0.000618892 | 0.070447717 |
| *CHL1* | 1.625927 | 0.475561 | 3.418963 | 0.000628603 | 0.070914292 |
| *LINC00513* | 0.539292 | 0.158102 | 3.41103 | 0.000647179 | 0.072363758 |
| *TRANK1* | -0.5732 | 0.16894 | -3.392915 | 0.00069153 | 0.076644563 |
| *PALLD* | 2.910568 | 0.858589 | 3.389944 | 0.000699069 | 0.07680638 |
| *MMACHC* | 1.436586 | 0.424087 | 3.38748 | 0.000705378 | 0.076831517 |
| *MOGS* | -0.825814 | 0.244105 | -3.383023 | 0.000716926 | 0.077421837 |
| *CPNE5* | 0.929721 | 0.275187 | 3.378509 | 0.0007288 | 0.078037158 |
| *ENSG00000227486* | 0.600087 | 0.177822 | 3.374646 | 0.000739107 | 0.078475794 |
| *MAP3K1* | 0.434953 | 0.129105 | 3.369001 | 0.000754411 | 0.079433194 |
| *NFATC3* | 0.393448 | 0.117273 | 3.354977 | 0.000793717 | 0.082881061 |
| *AKAP12* | 2.973588 | 0.891141 | 3.336832 | 0.000847393 | 0.087760738 |
| *FXYD5* | -0.23377 | 0.070442 | -3.318616 | 0.000904647 | 0.092928574 |
| *SMIM3* | 3.085858 | 0.930725 | 3.31554 | 0.000914661 | 0.093199532 |
| *ZFP36L1* | -0.350535 | 0.105899 | -3.310096 | 0.00093264 | 0.094271229 |
| *GABPB1* | 0.286211 | 0.08673 | 3.300024 | 0.000966765 | 0.096945016 |
| *RINL* | 0.468097 | 0.142199 | 3.291838 | 0.000995349 | 0.099025497 |
| *ZNF3* | 0.60168 | 0.182972 | 3.28837 | 0.001007692 | 0.099470182 |
| *MBD6* | 0.796908 | 0.242657 | 3.284087 | 0.001023132 | 0.099808849 |
| *SLC45A3* | 1.001573 | 0.305075 | 3.283045 | 0.001026921 | 0.099808849 |
| *CERK* | 0.677761 | 0.206673 | 3.27939 | 0.001040318 | 0.100339071 |
| *LMNA* | -2.649884 | 0.810167 | -3.270786 | 0.001072491 | 0.102658488 |
| *TSPAN18* | 3.381633 | 1.035753 | 3.264903 | 0.001095015 | 0.104026384 |
| *ACP5* | 0.858481 | 0.264079 | 3.250852 | 0.001150596 | 0.107761145 |
| *C9orf16* | -0.265397 | 0.081644 | -3.250657 | 0.001151385 | 0.107761145 |

| | | | | | |
|---|---|---|---|---|---|
| *ERAP1* | 0.573238 | 0.17693 | 3.239914 | 0.001195659 | 0.111081971 |
| *NUP210* | 0.378681 | 0.117395 | 3.225711 | 0.001256602 | 0.115891755 |
| *TNFRSF18* | -1.254858 | 0.38961 | -3.220808 | 0.001278297 | 0.117038323 |
| *PLXNB2* | 1.598995 | 0.50009 | 3.197417 | 0.001386645 | 0.125072093 |
| *A1BG* | -0.350948 | 0.10983 | -3.195369 | 0.001396521 | 0.125072093 |
| *CYTIP* | -0.191769 | 0.06003 | -3.194545 | 0.001400514 | 0.125072093 |
| *MEF2A* | 0.557722 | 0.174643 | 3.193491 | 0.001405638 | 0.125072093 |
| *DDIT4* | 1.094364 | 0.345667 | 3.165946 | 0.001545796 | 0.136581347 |
| *GLIPR2* | 0.514372 | 0.162644 | 3.162559 | 0.001563889 | 0.137220376 |
| *ENSG00000261766* | 0.722994 | 0.22896 | 3.157724 | 0.001590058 | 0.138554331 |
| *C8orf58* | -1.504791 | 0.478008 | -3.148042 | 0.001643679 | 0.142197975 |
| *LCK* | 0.68596 | 0.218032 | 3.146145 | 0.001654381 | 0.142197975 |
| *POU2AF1* | 0.281908 | 0.090007 | 3.132072 | 0.001735774 | 0.148185837 |
| *ADD1* | 0.349358 | 0.111919 | 3.121532 | 0.001799126 | 0.152563443 |
| *TIPARP* | -0.811544 | 0.260505 | -3.115278 | 0.001837715 | 0.154796898 |
| *LY86* | -0.340526 | 0.1094 | -3.112663 | 0.001854078 | 0.155140887 |
| *LINC02576* | -1.2031 | 0.389879 | -3.085827 | 0.002029872 | 0.167963121 |
| *BIN2* | 0.65178 | 0.211258 | 3.085237 | 0.002033902 | 0.167963121 |
| *USP8* | 0.274906 | 0.089377 | 3.075788 | 0.002099467 | 0.171665307 |
| *ZNF26* | 0.42254 | 0.137417 | 3.074875 | 0.002105906 | 0.171665307 |
| *SCIMP* | 1.09172 | 0.355341 | 3.072315 | 0.002124056 | 0.172034932 |
| *CD83* | -0.339848 | 0.111081 | -3.059454 | 0.00221741 | 0.177331465 |
| *DAAM1* | -0.680209 | 0.222331 | -3.059439 | 0.002217521 | 0.177331465 |
| *LRRC61* | 0.746927 | 0.244587 | 3.053829 | 0.002259406 | 0.178834861 |
| *CTSS* | 0.274396 | 0.089874 | 3.053137 | 0.002264628 | 0.178834861 |
| *EML6* | -1.88302 | 0.617265 | -3.050583 | 0.002283972 | 0.179242171 |
| *CD99* | -0.536373 | 0.175987 | -3.047802 | 0.002305215 | 0.179792531 |
| *KANSL3* | 0.769244 | 0.253388 | 3.035839 | 0.002398678 | 0.18593433 |
| *FCMR* | 0.389535 | 0.128777 | 3.024889 | 0.002487243 | 0.191623905 |
| *JDP2* | 1.1989 | 0.397545 | 3.015759 | 0.002563373 | 0.196292198 |
| *RNF207* | 2.091936 | 0.695054 | 3.009747 | 0.002614655 | 0.197836104 |
| *VPS26B* | -1.765548 | 0.586615 | -3.009724 | 0.00261485 | 0.197836104 |
| *TMEM41B* | 0.539439 | 0.179554 | 3.004335 | 0.002661625 | 0.200176353 |
| *ENSG00000240859* | 2.230564 | 0.743793 | 2.998903 | 0.002709534 | 0.202573711 |
| *CD5* | -1.499014 | 0.502599 | -2.982521 | 0.002858848 | 0.212479679 |
| *LINC02453* | 1.283003 | 0.431427 | 2.973858 | 0.002940813 | 0.217293411 |
| *IK* | 0.25904 | 0.087237 | 2.969396 | 0.002983859 | 0.218666894 |
| *AHNAK* | -0.997392 | 0.336119 | -2.967373 | 0.003003564 | 0.218666894 |
| *FOXN3* | 0.351252 | 0.118445 | 2.965521 | 0.003021704 | 0.218666894 |
| *PTPN6* | 0.514237 | 0.173446 | 2.964817 | 0.003028627 | 0.218666894 |
| *ISG20* | -0.322367 | 0.109012 | -2.957182 | 0.003104653 | 0.222882312 |
| *PLEKHO1* | 0.455326 | 0.15422 | 2.952438 | 0.003152751 | 0.224621733 |
| *CLCN4* | 1.536938 | 0.520767 | 2.951296 | 0.003164438 | 0.224621733 |
| *FCRL2* | 0.606028 | 0.206891 | 2.929207 | 0.003398279 | 0.239872927 |
| *VHL* | 0.377488 | 0.129243 | 2.920772 | 0.003491652 | 0.245094578 |
| *NDUFB9* | 0.261051 | 0.089684 | 2.910774 | 0.003605345 | 0.25167701 |
| *MLXIP* | 0.434494 | 0.149919 | 2.898197 | 0.003753152 | 0.26055539 |

| | | | | | |
|---|---|---|---|---|---|
| *GBP7* | 1.029267 | 0.355378 | 2.896258 | 0.003776417 | 0.260737883 |
| *CYBB* | 0.509676 | 0.176108 | 2.89412 | 0.003802231 | 0.261093386 |
| *CDC26* | -0.303667 | 0.105364 | -2.882069 | 0.003950727 | 0.269823947 |
| *SLC2A3* | -0.661164 | 0.229751 | -2.877747 | 0.004005262 | 0.271966012 |
| *WNT10A* | -1.001444 | 0.348185 | -2.876185 | 0.00402514 | 0.271966012 |
| *ENSG00000271204* | 0.530962 | 0.185095 | 2.868591 | 0.004123041 | 0.276036758 |
| *AK1* | 0.796151 | 0.277838 | 2.86552 | 0.00416325 | 0.276036758 |
| *PDCD4* | 0.545849 | 0.190519 | 2.865069 | 0.004169183 | 0.276036758 |
| *SLCO4A1* | -1.052977 | 0.367557 | -2.864796 | 0.004172776 | 0.276036758 |
| *HTR3A* | 1.228382 | 0.429412 | 2.860616 | 0.004228187 | 0.278245559 |
| *GCHFR* | -0.50638 | 0.177908 | -2.846303 | 0.004423003 | 0.289557724 |
| *SLC1A4* | -1.380221 | 0.485606 | -2.842266 | 0.004479412 | 0.291739039 |
| *ATG2A* | 0.611877 | 0.215487 | 2.839505 | 0.004518354 | 0.292766152 |
| *OTUD7A* | 1.829475 | 0.644723 | 2.837614 | 0.004545213 | 0.293003883 |
| *SLC43A2* | 0.295343 | 0.104212 | 2.834064 | 0.004596013 | 0.294774735 |
| *WDFY2* | -0.471378 | 0.166765 | -2.826605 | 0.004704428 | 0.29851522 |
| *CTSA* | 0.465178 | 0.1646 | 2.82611 | 0.004711712 | 0.29851522 |
| *ICA1* | 2.948529 | 1.045229 | 2.82094 | 0.004788319 | 0.29851522 |
| *TP53I11* | -0.685886 | 0.243157 | -2.820758 | 0.004791027 | 0.29851522 |
| *ENSG00000270019* | 0.839659 | 0.297783 | 2.819698 | 0.004806886 | 0.29851522 |
| *COA1* | 0.413426 | 0.146807 | 2.816121 | 0.004860741 | 0.29851522 |
| *FADS3* | -0.390184 | 0.138588 | -2.815417 | 0.0048714 | 0.29851522 |
| *ENSG00000279278* | 0.650736 | 0.231171 | 2.814953 | 0.00487844 | 0.29851522 |
| *SRPRB* | -0.675226 | 0.239918 | -2.814404 | 0.004886778 | 0.29851522 |
| *NANP* | 1.271546 | 0.45184 | 2.814153 | 0.004890594 | 0.29851522 |
| *TIMM23B* | 0.871122 | 0.309796 | 2.811923 | 0.004924637 | 0.299148003 |
| *CTDSP1* | 0.342076 | 0.121884 | 2.806563 | 0.005007313 | 0.30271482 |
| *AP3S1* | 0.387119 | 0.138436 | 2.796381 | 0.005167842 | 0.307947269 |
| *ENSG00000258377* | 1.668491 | 0.596931 | 2.795114 | 0.005188143 | 0.307947269 |
| *DNMT3A* | 0.900307 | 0.32224 | 2.793901 | 0.005207649 | 0.307947269 |
| *SEMA4A* | 1.567559 | 0.56124 | 2.793027 | 0.005221736 | 0.307947269 |
| *ZDBF2* | -1.903363 | 0.681745 | -2.7919 | 0.005239953 | 0.307947269 |
| *CD22* | 0.492349 | 0.176351 | 2.79187 | 0.005240441 | 0.307947269 |
| *CRX* | 1.549056 | 0.55514 | 2.790389 | 0.005264472 | 0.307947269 |
| *ABCB1* | 0.642207 | 0.230693 | 2.783819 | 0.005372296 | 0.312806264 |
| *CENPM* | 0.483589 | 0.173891 | 2.780996 | 0.005419244 | 0.313097887 |
| *ARHGAP4* | 0.387309 | 0.139376 | 2.77888 | 0.005454662 | 0.313097887 |
| *IZUMO4* | -0.551379 | 0.198513 | -2.777549 | 0.005477066 | 0.313097887 |
| *ARID3A* | 0.557055 | 0.20066 | 2.776115 | 0.005501266 | 0.313097887 |
| *KLF8* | 0.466097 | 0.167912 | 2.775839 | 0.005505954 | 0.313097887 |
| *MYADM* | 0.62661 | 0.225833 | 2.774658 | 0.005525986 | 0.313097887 |
| *VSIG10L* | 1.898372 | 0.68531 | 2.770093 | 0.005604028 | 0.316102206 |
| *MANF* | 0.386559 | 0.139775 | 2.765591 | 0.005681981 | 0.319074819 |
| *FAM167A* | 1.387309 | 0.502111 | 2.762955 | 0.005728074 | 0.319203412 |
| *LSP1* | 0.326205 | 0.11808 | 2.762572 | 0.005734798 | 0.319203412 |
| *ST13* | -0.150864 | 0.054643 | -2.76091 | 0.005764048 | 0.319424348 |
| *MYC* | 0.336197 | 0.121878 | 2.758467 | 0.005807322 | 0.320417098 |

| | | | | | |
|---|---|---|---|---|---|
| *GPANK1* | 0.313919 | 0.114017 | 2.753273 | 0.005900272 | 0.32413018 |
| *IL21R* | 0.663703 | 0.241482 | 2.748454 | 0.005987701 | 0.327278568 |
| *POLD4* | 0.263775 | 0.096016 | 2.747209 | 0.006010477 | 0.327278568 |
| *TP53* | 0.375057 | 0.13659 | 2.745858 | 0.006035291 | 0.327278568 |
| *ROR1* | -1.690949 | 0.616916 | -2.740971 | 0.006125786 | 0.329213979 |
| *GEN1* | -0.932234 | 0.340134 | -2.740787 | 0.006129214 | 0.329213979 |
| *TBC1D9* | 0.794872 | 0.290129 | 2.73972 | 0.006149149 | 0.329213979 |
| *WDR97* | 1.699877 | 0.621382 | 2.735638 | 0.00622595 | 0.331919296 |
| *SERPINB6* | 0.596341 | 0.218109 | 2.734143 | 0.006254279 | 0.332028615 |
| *RAB3IP* | -0.525092 | 0.192537 | -2.727223 | 0.006386984 | 0.336325938 |
| *SEC61B* | -0.330612 | 0.12123 | -2.727147 | 0.006388463 | 0.336325938 |
| *ENSG00000272644* | 1.707326 | 0.626865 | 2.723596 | 0.006457553 | 0.338230306 |
| *RNF7* | -0.272583 | 0.100156 | -2.721577 | 0.006497125 | 0.338230306 |
| *TTBK2* | -0.805726 | 0.296268 | -2.719582 | 0.006536458 | 0.338230306 |
| *PPP1CA* | -0.181757 | 0.066842 | -2.719187 | 0.00654426 | 0.338230306 |
| *CDC42SE1* | 0.52517 | 0.193186 | 2.718469 | 0.006558482 | 0.338230306 |
| *ENSG00000260136* | -1.402884 | 0.51684 | -2.714348 | 0.006640639 | 0.34107509 |
| *ENSG00000261884* | 0.639585 | 0.235975 | 2.710395 | 0.006720309 | 0.342487575 |
| *APOBEC3D* | 0.472068 | 0.17425 | 2.709135 | 0.006745881 | 0.342487575 |
| *CECR7* | 1.534627 | 0.5665 | 2.708959 | 0.006749458 | 0.342487575 |
| *SLC25A45* | 0.605708 | 0.224269 | 2.700814 | 0.006916993 | 0.348983406 |
| *NKILA* | 1.264259 | 0.468234 | 2.700059 | 0.006932713 | 0.348983406 |
| *TOP1MT* | -0.603868 | 0.224366 | -2.691434 | 0.007114553 | 0.356715793 |
| *ENSG00000130520* | -0.287846 | 0.107007 | -2.689978 | 0.007145663 | 0.356859512 |
| *TBL1X* | -0.615529 | 0.229058 | -2.687214 | 0.007205076 | 0.358409974 |
| *HNRNPK* | 0.192157 | 0.071782 | 2.676955 | 0.007429462 | 0.367678199 |
| *HIVEP2* | 0.742035 | 0.277287 | 2.676048 | 0.007449594 | 0.367678199 |
| *BRD3OS* | -0.711749 | 0.266378 | -2.671948 | 0.007541243 | 0.369681575 |
| *BHLHE40* | 0.672632 | 0.251934 | 2.669878 | 0.007587878 | 0.369681575 |
| *C7orf50* | -0.246975 | 0.09254 | -2.668834 | 0.007611517 | 0.369681575 |
| *ZCCHC17* | 0.335489 | 0.12579 | 2.66705 | 0.007652029 | 0.369681575 |
| *ENSG00000259768* | 1.272001 | 0.476976 | 2.6668 | 0.007657716 | 0.369681575 |
| *METTL7A* | 0.345447 | 0.129634 | 2.664791 | 0.007703613 | 0.369681575 |
| *TCTN3* | -0.47027 | 0.176527 | -2.664006 | 0.007721622 | 0.369681575 |
| *ENSG00000249476* | -1.971213 | 0.740165 | -2.663208 | 0.007739954 | 0.369681575 |
| *TENT5C* | -0.359334 | 0.135079 | -2.660178 | 0.007809932 | 0.369681575 |
| *LBR* | 0.309841 | 0.116477 | 2.660105 | 0.007811626 | 0.369681575 |
| *HINT2* | 0.287688 | 0.10815 | 2.660088 | 0.007812029 | 0.369681575 |
| *AGAP3* | 2.592503 | 0.975285 | 2.658199 | 0.007855942 | 0.37037249 |
| *CNOT7* | 0.208678 | 0.078559 | 2.656304 | 0.007900233 | 0.371075981 |
| *TRBC2* | -0.221709 | 0.083612 | -2.651635 | 0.008010312 | 0.372658303 |
| *P2RY8* | 0.351964 | 0.132881 | 2.648723 | 0.008079653 | 0.372658303 |
| *ENSG00000273445* | 3.330759 | 1.257797 | 2.64809 | 0.008094809 | NA |
| *CARD11* | 0.561688 | 0.212226 | 2.646645 | 0.008129469 | 0.372658303 |
| *CYBC1* | -0.231295 | 0.087393 | -2.646614 | 0.008130218 | 0.372658303 |
| *CARNMT1* | 0.654125 | 0.247161 | 2.64655 | 0.008131743 | 0.372658303 |
| *ARL3* | -0.514534 | 0.194482 | -2.645662 | 0.008153118 | 0.372658303 |

| | | | | | |
|---|---|---|---|---|---|
| *PLCXD1* | 0.696559 | 0.263318 | 2.645317 | 0.008161433 | 0.372658303 |
| *FUNDC1* | 0.522282 | 0.197501 | 2.644454 | 0.008182292 | 0.372658303 |
| *GTF2H5* | 0.30721 | 0.11622 | 2.643359 | 0.008208804 | 0.372658303 |
| *RBX1* | -0.209763 | 0.07938 | -2.642532 | 0.008228862 | 0.372658303 |
| *NIPAL4* | 1.37713 | 0.521728 | 2.639553 | 0.008301536 | 0.374606811 |
| *CCDC12* | 0.23257 | 0.088202 | 2.636802 | 0.008369173 | 0.37520945 |
| *DHFR2* | 0.681154 | 0.258352 | 2.636533 | 0.0083758 | 0.37520945 |
| *SH3KBP1* | -0.275976 | 0.10476 | -2.63435 | 0.00842985 | 0.37520945 |
| *IQSEC1* | -0.371486 | 0.141025 | -2.634196 | 0.008433675 | 0.37520945 |
| *GMEB1* | 0.429146 | 0.163149 | 2.630385 | 0.008528813 | 0.375931114 |
| *ENSG00000234484* | 1.965427 | 0.74722 | 2.630318 | 0.008530491 | 0.375931114 |
| *CTSZ* | 0.342007 | 0.130061 | 2.629583 | 0.008548978 | 0.375931114 |
| *JCHAIN* | 1.128514 | 0.429337 | 2.628501 | 0.008576206 | 0.375931114 |
| *MPV17L* | 1.107449 | 0.421568 | 2.62698 | 0.008614645 | 0.375931114 |
| *JADE1* | 0.360064 | 0.137092 | 2.626436 | 0.008628415 | 0.375931114 |
| *EMC6* | -0.381964 | 0.145747 | -2.62074 | 0.008773915 | 0.3786344 |
| *FKBP5* | 0.96305 | 0.367573 | 2.620027 | 0.008792273 | 0.3786344 |
| *DDX28* | 0.474302 | 0.181073 | 2.619391 | 0.008808703 | 0.3786344 |
| *EIF4G3* | 0.513396 | 0.196003 | 2.619328 | 0.00881033 | 0.3786344 |
| *KLF3* | 0.393256 | 0.150359 | 2.615448 | 0.008911061 | 0.380108621 |
| *HDAC9* | 0.538837 | 0.206045 | 2.615146 | 0.00891892 | 0.380108621 |
| *TMEM175* | 0.678585 | 0.259543 | 2.614536 | 0.008934884 | 0.380108621 |
| *GPR157* | 0.867439 | 0.332227 | 2.610981 | 0.009028283 | 0.382082665 |
| *CYLD* | -0.299242 | 0.114632 | -2.610471 | 0.009041766 | 0.382082665 |
| *ENSG00000270589* | 1.720093 | 0.659846 | 2.606811 | 0.009138976 | 0.384527372 |
| *RALA* | -0.390998 | 0.150037 | -2.606006 | 0.009160486 | 0.384527372 |
| *PTPN12* | -0.784838 | 0.301782 | -2.600678 | 0.009303975 | 0.38900645 |
| *SLCO5A1* | 1.279357 | 0.492105 | 2.599765 | 0.009328766 | 0.38900645 |
| *SELL* | 0.569664 | 0.219339 | 2.597178 | 0.009399309 | 0.390319449 |
| *DPEP3* | 1.616677 | 0.622673 | 2.596349 | 0.009422037 | 0.390319449 |
| *INVS* | 0.89976 | 0.346835 | 2.594205 | 0.009480991 | 0.391478158 |
| *RASA3* | 0.75459 | 0.291538 | 2.588305 | 0.009644963 | 0.3969515 |
| *FUCA1* | -0.68137 | 0.263622 | -2.584646 | 0.009747899 | 0.399885384 |
| *MMP7* | 1.879867 | 0.728215 | 2.581472 | 0.009837996 | 0.402275332 |
| *MOSPD3* | -0.468293 | 0.182502 | -2.565965 | 0.010288931 | 0.419356899 |
| *AFF3* | -0.253559 | 0.098878 | -2.564363 | 0.010336548 | 0.419943048 |
| *TNK2* | 0.423042 | 0.165314 | 2.559026 | 0.010496598 | 0.425078567 |
| *PCBP4* | -1.001591 | 0.391846 | -2.556084 | 0.010585762 | 0.427319807 |
| *ADCK1* | 1.205057 | 0.472055 | 2.552789 | 0.010686434 | 0.427679351 |
| *SH3BGRL3* | -0.185185 | 0.072644 | -2.549208 | 0.010796774 | 0.427679351 |
| *DMD* | -0.611274 | 0.239794 | -2.549161 | 0.010798232 | 0.427679351 |
| *SVBP* | -0.269944 | 0.105896 | -2.549135 | 0.010799062 | 0.427679351 |
| *MMP11* | 0.71074 | 0.278834 | 2.548967 | 0.010804249 | 0.427679351 |
| *KCTD10* | 0.624265 | 0.245024 | 2.547777 | 0.01084117 | 0.427679351 |
| *CCDC106* | -0.532162 | 0.208965 | -2.546663 | 0.010875854 | 0.427679351 |
| *ZNF337.AS1* | 0.934388 | 0.366969 | 2.546231 | 0.010889297 | 0.427679351 |
| *ENSG00000180448* | 0.288126 | 0.113172 | 2.545911 | 0.010899308 | 0.427679351 |

| | | | | | |
|---|---|---|---|---|---|
| *PSMB10* | 0.342761 | 0.134739 | 2.543881 | 0.01096283 | 0.428545269 |
| *SEC23IP* | -0.564106 | 0.221823 | -2.543042 | 0.01098921 | 0.428545269 |
| *ENSG00000228106* | 0.449616 | 0.176998 | 2.54024 | 0.011077648 | 0.430664882 |
| *SRGN* | 0.543289 | 0.214103 | 2.537515 | 0.011164252 | 0.43270038 |
| *TELO2* | 0.557883 | 0.220199 | 2.533547 | 0.011291457 | 0.436292224 |
| *GTF3A* | 0.201389 | 0.07956 | 2.531282 | 0.011364629 | 0.43662599 |
| *ENSG00000263884* | 1.563218 | 0.617594 | 2.531141 | 0.011369209 | 0.43662599 |
| *ENSG00000253535* | 0.789119 | 0.312334 | 2.526526 | 0.011519682 | 0.440414568 |
| *ENSG00000166927* | -0.835111 | 0.330609 | -2.525981 | 0.011537572 | 0.440414568 |
| *SP110* | 0.340164 | 0.134758 | 2.524267 | 0.011593983 | 0.440759899 |
| *CLIC3* | 1.503333 | 0.595927 | 2.522681 | 0.011646399 | 0.440759899 |
| *ZNF239* | 1.365026 | 0.541133 | 2.522534 | 0.011651271 | 0.440759899 |
| *FBP1* | 1.124717 | 0.446415 | 2.519445 | 0.011753992 | 0.44321587 |
| *HSD11B1L* | -1.075723 | 0.4272 | -2.518076 | 0.011799788 | 0.44321587 |
| *PLA2G7* | 3.754507 | 1.491405 | 2.517431 | 0.011821428 | 0.44321587 |
| *ENSG00000176320* | 2.680585 | 1.065726 | 2.515266 | 0.011894276 | 0.444627727 |
| *ENSG00000261786* | 2.637035 | 1.049446 | 2.512788 | 0.011978135 | 0.44563712 |
| *DNAJA2* | 0.307701 | 0.122504 | 2.511769 | 0.012012782 | 0.44563712 |
| *KDM2B* | 0.384072 | 0.152935 | 2.511349 | 0.012027088 | 0.44563712 |
| *METTL21A* | -0.315669 | 0.125912 | -2.50706 | 0.012173991 | 0.449761338 |
| *MATR3* | 0.958311 | 0.382632 | 2.504524 | 0.012261629 | 0.451152774 |
| *ENSG00000258181* | 0.650882 | 0.25998 | 2.503583 | 0.012294267 | 0.451152774 |
| *ZBTB25* | 0.366285 | 0.146369 | 2.502478 | 0.012332719 | 0.451152774 |
| *ENSG00000278002* | 0.519955 | 0.207902 | 2.500962 | 0.01238565 | 0.451152774 |
| *SBF2.AS1* | -0.652 | 0.2608 | -2.500001 | 0.012419288 | 0.451152774 |
| *MLH1* | -0.741818 | 0.296862 | -2.498864 | 0.012459225 | 0.451152774 |
| *ECHS1* | 0.332092 | 0.132901 | 2.498796 | 0.0124616 | 0.451152774 |
| *IKZF1* | 0.32648 | 0.130777 | 2.496465 | 0.012543819 | 0.452831876 |
| *GALK1* | -0.57284 | 0.229713 | -2.493718 | 0.012641284 | 0.455050207 |
| *GYPC* | 0.23282 | 0.093439 | 2.491682 | 0.012713988 | 0.455574482 |
| *SLF2* | -0.391315 | 0.157112 | -2.490667 | 0.012750348 | 0.455574482 |
| *IER3.AS1* | 1.417004 | 0.569012 | 2.490286 | 0.012764018 | 0.455574482 |
| *BSCL2* | 1.074307 | 0.431607 | 2.489084 | 0.012807257 | 0.45561258 |
| *PTPA* | -0.811897 | 0.326292 | -2.488254 | 0.012837204 | 0.45561258 |
| *ENDOD1* | 0.649393 | 0.261832 | 2.480192 | 0.013131165 | 0.462544168 |
| *C1orf56* | 0.562825 | 0.226929 | 2.480181 | 0.013131558 | 0.462544168 |
| *MOB4* | 0.293409 | 0.118315 | 2.479889 | 0.013142331 | 0.462544168 |
| *MAP3K2* | -0.324779 | 0.131303 | -2.473503 | 0.013379568 | 0.468920828 |
| *RNF122* | 1.039932 | 0.420579 | 2.472617 | 0.013412798 | 0.468920828 |
| *FBXO10* | 0.776173 | 0.313982 | 2.472029 | 0.013434851 | 0.468920828 |
| *SLC35E3* | 0.291658 | 0.118571 | 2.459767 | 0.013902727 | 0.48050934 |
| *CNST* | 0.484397 | 0.196931 | 2.459727 | 0.013904257 | 0.48050934 |
| *REL* | -0.253587 | 0.103101 | -2.459588 | 0.01390967 | 0.48050934 |
| *UHRF2* | 0.426281 | 0.173331 | 2.459346 | 0.01391905 | 0.48050934 |
| *DPPA4* | -1.351698 | 0.549836 | -2.458368 | 0.013957018 | 0.48050934 |
| *TNFSF10* | 0.497905 | 0.202852 | 2.45452 | 0.014107269 | 0.483508011 |
| *ENSG00000260793* | 0.617817 | 0.251741 | 2.454179 | 0.014120653 | 0.483508011 |

| | | | | | |
|---|---|---|---|---|---|
| *PI4K2A* | -1.771477 | 0.72385 | -2.447297 | 0.014393213 | 0.491508772 |
| *ISCA1* | 0.288784 | 0.118099 | 2.445267 | 0.014474489 | 0.492701428 |
| *MICU1* | 0.457542 | 0.187174 | 2.44448 | 0.014506128 | 0.492701428 |
| *ABI3* | 0.422575 | 0.173356 | 2.437621 | 0.014784274 | 0.500802403 |
| *SERPINB1* | 0.265955 | 0.109199 | 2.435505 | 0.014871029 | 0.502394269 |
| *IRF1* | 0.289272 | 0.118826 | 2.434422 | 0.014915581 | 0.502555643 |
| *ENSG00000283013* | 0.340998 | 0.140291 | 2.430648 | 0.015071832 | 0.505600014 |
| *UBE2G1* | 0.360361 | 0.148331 | 2.429437 | 0.015122281 | 0.505600014 |
| *GABPB2* | 0.516299 | 0.212562 | 2.428933 | 0.015143332 | 0.505600014 |
| *ENSG00000226571* | -1.528334 | 0.629361 | -2.428391 | 0.015166 | 0.505600014 |
| *ZFP36* | 0.205473 | 0.084776 | 2.423734 | 0.015361837 | 0.510240671 |
| *PLP2* | -0.478333 | 0.197618 | -2.420491 | 0.015499571 | 0.510240671 |
| *HSH2D* | 0.310247 | 0.128189 | 2.420235 | 0.015510497 | 0.510240671 |
| *ZNF660* | -1.269792 | 0.524732 | -2.419886 | 0.01552536 | 0.510240671 |
| *FLNA* | -0.49394 | 0.204173 | -2.419219 | 0.015553886 | 0.510240671 |
| *LTBP3* | -0.42503 | 0.175717 | -2.418837 | 0.015570235 | 0.510240671 |
| *ENSG00000273247* | 0.261598 | 0.108189 | 2.417977 | 0.015607056 | 0.510240671 |
| *MFSD14B* | -0.878911 | 0.363703 | -2.416563 | 0.015667809 | 0.510240671 |
| *CCDC157* | 0.905777 | 0.374939 | 2.415797 | 0.015700808 | 0.510240671 |
| *ENSG00000234773* | 1.947449 | 0.806431 | 2.414899 | 0.015739593 | 0.510240671 |
| *CCDC71L* | -1.091306 | 0.452113 | -2.413788 | 0.015787634 | 0.510240671 |
| *ZNF587* | 0.494212 | 0.20475 | 2.413738 | 0.015789798 | 0.510240671 |
| *MDP1* | 0.53899 | 0.223621 | 2.41029 | 0.015939842 | 0.513775275 |
| *CYTH2* | -0.399349 | 0.165786 | -2.408816 | 0.01600436 | 0.5145422 |
| *RPS6KA2* | -1.114612 | 0.46337 | -2.405447 | 0.01615271 | 0.515196363 |
| *RSF1* | 0.24001 | 0.099798 | 2.404952 | 0.016174586 | 0.515196363 |
| *PARP12* | -0.869379 | 0.361499 | -2.404927 | 0.0161757 | 0.515196363 |
| *KDM3A* | 0.573482 | 0.238488 | 2.404653 | 0.016187808 | 0.515196363 |
| *C15orf39* | 0.784129 | 0.326348 | 2.402735 | 0.016272963 | 0.515313967 |
| *RUBCN* | -0.334316 | 0.139143 | -2.402675 | 0.016275638 | 0.515313967 |
| *RHBDF2* | 0.274379 | 0.114238 | 2.401817 | 0.016313857 | 0.515313967 |
| *CR2* | 0.87206 | 0.363309 | 2.400324 | 0.01638056 | 0.516044636 |
| *PHGDH* | 2.173084 | 0.90565 | 2.399473 | 0.016418674 | 0.516044636 |
| *UBE2O* | 0.430057 | 0.179473 | 2.396225 | 0.016564929 | 0.519349575 |
| *TM2D3* | 0.394095 | 0.164571 | 2.39468 | 0.016634858 | 0.520251066 |
| *SESN3* | -0.297235 | 0.124201 | -2.393177 | 0.016703167 | 0.521097579 |
| *AP4M1* | 0.454816 | 0.190154 | 2.391833 | 0.016764487 | 0.521722384 |
| *LINC02352* | 0.774256 | 0.324087 | 2.389036 | 0.016892666 | 0.524419736 |
| *SERPINF2* | -1.645888 | 0.689512 | -2.387034 | 0.016984907 | 0.52472111 |
| *NCF4* | -0.354058 | 0.148354 | -2.38658 | 0.017005885 | 0.52472111 |
| *CDCA7L* | 0.359273 | 0.150585 | 2.38585 | 0.017039689 | 0.52472111 |
| *LINC.PINT* | 0.34669 | 0.14544 | 2.38373 | 0.017138188 | 0.52472111 |
| *FMN1* | 1.519336 | 0.637388 | 2.383691 | 0.017140006 | 0.52472111 |
| *TBC1D2B* | -0.911031 | 0.382274 | -2.383189 | 0.017163392 | 0.52472111 |
| *CHP1* | -0.295747 | 0.124171 | -2.381769 | 0.017229695 | 0.52472111 |
| *GRINA* | 0.340936 | 0.143156 | 2.381564 | 0.017239303 | 0.52472111 |
| *CHMP1B* | 0.244881 | 0.102879 | 2.38028 | 0.017299479 | 0.52472111 |

| | | | | | |
|---|---|---|---|---|---|
| *PARP14* | -0.24377 | 0.102442 | -2.379599 | 0.017331484 | 0.52472111 |
| *TBCD* | -0.933958 | 0.392583 | -2.37901 | 0.017359195 | 0.52472111 |
| *PHKG1* | 0.474791 | 0.199984 | 2.374145 | 0.017589651 | 0.530418225 |
| *PEAK3* | 1.446709 | 0.610007 | 2.371625 | 0.017710048 | 0.532777272 |
| *LARGE1* | 1.21848 | 0.514004 | 2.370567 | 0.017760842 | 0.533036196 |
| *PHLDB1* | 1.587751 | 0.670127 | 2.369327 | 0.017820482 | 0.533558749 |
| *PRICKLE1* | 0.55251 | 0.233377 | 2.367456 | 0.01791085 | 0.534330983 |
| *EPHB6* | -0.724934 | 0.306359 | -2.366292 | 0.017967289 | 0.534330983 |
| *MICALL1* | -1.027208 | 0.434325 | -2.365068 | 0.018026766 | 0.534330983 |
| *KCNN1* | 1.256269 | 0.531438 | 2.363906 | 0.018083416 | 0.534330983 |
| *TMC8* | 0.336725 | 0.142446 | 2.36388 | 0.018084662 | 0.534330983 |
| *ITGB2* | 0.627966 | 0.265686 | 2.363566 | 0.018100013 | 0.534330983 |
| *TGFBR2* | -0.259453 | 0.109853 | -2.361819 | 0.018185507 | 0.535603454 |
| *LIPT2* | 0.725518 | 0.307354 | 2.360531 | 0.0182488 | 0.536217646 |
| *DAPP1* | -0.257548 | 0.109195 | -2.358594 | 0.018344323 | 0.537773833 |
| *SNHG21* | -0.515024 | 0.218572 | -2.356313 | 0.018457349 | 0.539764755 |
| *TTC37* | -0.489557 | 0.207835 | -2.355502 | 0.018497676 | 0.539764755 |
| *IER3* | 1.033816 | 0.439935 | 2.349928 | 0.018777041 | 0.546654183 |
| *TRAF7* | -0.58744 | 0.250202 | -2.347864 | 0.018881399 | 0.548428674 |
| *FTH1* | -0.151138 | 0.0644 | -2.346858 | 0.018932456 | 0.548650417 |
| *MAP9* | -0.502275 | 0.214131 | -2.345638 | 0.018994569 | 0.549190805 |
| *FYN* | 0.589602 | 0.251795 | 2.341592 | 0.019201676 | 0.552844767 |
| *ERP27* | -1.782987 | 0.761622 | -2.341039 | 0.019230149 | 0.552844767 |
| *SHF* | 0.835387 | 0.357094 | 2.339402 | 0.019314651 | 0.552844767 |
| *APOBEC3C* | 0.195566 | 0.083612 | 2.33897 | 0.019337009 | 0.552844767 |
| *ENSG00000235609* | 1.424145 | 0.608891 | 2.338917 | 0.019339722 | 0.552844767 |
| *MLH3* | 0.436529 | 0.186809 | 2.336764 | 0.019451452 | 0.553899544 |
| *FOXO3* | 0.628405 | 0.26895 | 2.336517 | 0.019464297 | 0.553899544 |
| *ANKRD44* | 0.232766 | 0.099678 | 2.33517 | 0.019534522 | 0.554648737 |
| *RASSF1* | 0.28674 | 0.123043 | 2.330409 | 0.019784562 | 0.560313006 |
| *SPN* | 1.109225 | 0.476372 | 2.328487 | 0.019886244 | 0.560313006 |
| *ZSWIM9* | 0.576346 | 0.247532 | 2.328366 | 0.019892682 | 0.560313006 |
| *BICDL1* | 1.328377 | 0.570859 | 2.32698 | 0.019966323 | 0.560313006 |
| *ITGB2.AS1* | 2.162025 | 0.929189 | 2.326787 | 0.019976582 | 0.560313006 |
| *INKA1* | -0.416588 | 0.179074 | -2.326346 | 0.020000092 | 0.560313006 |
| *LNPEP* | 0.255904 | 0.110181 | 2.322577 | 0.020201881 | 0.56471409 |
| *AGPAT2* | -0.326441 | 0.14082 | -2.31815 | 0.020441189 | 0.570128543 |
| *LINC00894* | 1.831305 | 0.790265 | 2.317329 | 0.020485822 | 0.570128543 |
| *ZNF665* | 0.584575 | 0.252475 | 2.315376 | 0.02059236 | 0.571834 |
| *MIPEP* | 0.796687 | 0.344389 | 2.313338 | 0.020704078 | 0.572436104 |
| *ENSG00000228835* | 0.595745 | 0.257574 | 2.31291 | 0.02072761 | 0.572436104 |
| *HNRNPL* | 0.326013 | 0.140987 | 2.312365 | 0.02075756 | 0.572436104 |
| *RIMS2* | 1.343742 | 0.581283 | 2.311681 | 0.020795265 | 0.572436104 |
| *ENSG00000251364* | 0.887372 | 0.384102 | 2.31025 | 0.020874337 | 0.573363572 |
| *ZCCHC14* | -1.997077 | 0.865786 | -2.306665 | 0.021073526 | 0.576349343 |
| *WDR82* | 0.390823 | 0.169433 | 2.306651 | 0.02107427 | 0.576349343 |
| *TIMP1* | -0.798901 | 0.346511 | -2.305557 | 0.021135404 | 0.576772843 |

| | | | | | |
|---|---|---|---|---|---|
| *ARHGAP15* | 0.231316 | 0.100462 | 2.30252 | 0.021305887 | 0.578001421 |
| *MDK* | 0.834485 | 0.36256 | 2.301649 | 0.021354953 | 0.578001421 |
| *ABRAXAS1* | -0.305005 | 0.132576 | -2.300611 | 0.021413636 | 0.578001421 |
| *PLAA* | -0.694919 | 0.302134 | -2.300034 | 0.021446321 | 0.578001421 |
| *RHOT1* | -0.483248 | 0.21012 | -2.299865 | 0.02145586 | 0.578001421 |
| *CD1C* | 0.866353 | 0.376823 | 2.299098 | 0.021499396 | 0.578001421 |
| *CHAF1A* | 1.148004 | 0.499446 | 2.298554 | 0.021530297 | 0.578001421 |
| *TOMM70* | 0.445433 | 0.193812 | 2.298271 | 0.021546393 | 0.578001421 |
| *TTC7A* | -0.584736 | 0.254707 | -2.295724 | 0.021691675 | 0.579830668 |
| *MSRB1* | -0.452415 | 0.197146 | -2.29482 | 0.021743468 | 0.579830668 |
| *CCNA2* | 1.203582 | 0.524723 | 2.293749 | 0.021804893 | 0.579830668 |
| *OLA1* | -0.256756 | 0.111962 | -2.293254 | 0.021833383 | 0.579830668 |
| *CCT6A* | -0.25735 | 0.112232 | -2.293019 | 0.021846905 | 0.579830668 |
| *CFL1* | -0.198969 | 0.086818 | -2.291787 | 0.021917958 | 0.579830668 |
| *CCNDBP1* | 0.190834 | 0.08328 | 2.291477 | 0.021935818 | 0.579830668 |
| *TP53TG1* | -0.35641 | 0.155658 | -2.289699 | 0.022038757 | 0.581335477 |
| *SLC25A46* | 0.387271 | 0.16925 | 2.288156 | 0.022128447 | 0.582485259 |
| *FSD1L* | -0.759424 | 0.332187 | -2.286133 | 0.022246487 | 0.584374973 |
| *RHOBTB2* | 0.73562 | 0.322398 | 2.281713 | 0.022506306 | 0.589973394 |
| *TOGARAM1* | 0.35358 | 0.155335 | 2.276248 | 0.022831157 | 0.597249821 |
| *HACD1* | -2.012375 | 0.884652 | -2.274765 | 0.022920054 | 0.598336526 |
| *PLEKHF1* | 0.614144 | 0.270265 | 2.272374 | 0.023063951 | 0.600049424 |
| *GNAZ* | -1.379817 | 0.607288 | -2.272097 | 0.023080651 | 0.600049424 |
| *FXYD1* | -0.519569 | 0.228852 | -2.270323 | 0.023187988 | 0.6016021 |
| *SYVN1* | 0.352891 | 0.155509 | 2.269269 | 0.023251955 | 0.602025521 |
| *FLI1* | 0.308315 | 0.135976 | 2.267423 | 0.023364401 | 0.602967907 |
| *PRDX2* | -0.332808 | 0.146799 | -2.267105 | 0.023383797 | 0.602967907 |
| *FAM32A* | -0.25181 | 0.111152 | -2.265454 | 0.023484812 | 0.604339302 |
| *ACAD10* | 0.547299 | 0.242238 | 2.259341 | 0.023862158 | 0.612801568 |
| *NME6* | 0.458722 | 0.203331 | 2.256034 | 0.024068532 | 0.61684768 |
| *IGLC7* | -2.028622 | 0.899738 | -2.25468 | 0.024153429 | 0.617770394 |
| *AVP* | -2.71625 | 1.205579 | -2.253066 | 0.024254962 | 0.618513991 |
| *DNAJC7* | 0.196142 | 0.087071 | 2.252663 | 0.024280407 | 0.618513991 |
| *LINC01679* | 1.730186 | 0.768524 | 2.251309 | 0.024365954 | 0.619326689 |
| *DVL3* | -0.612676 | 0.272292 | -2.250068 | 0.024444623 | 0.619326689 |
| *UBL7.AS1* | -0.441842 | 0.196389 | -2.249836 | 0.02445936 | 0.619326689 |
| *ENSG00000261187* | 1.27818 | 0.569061 | 2.246121 | 0.024696282 | 0.623052696 |
| *NOC3L* | 0.386443 | 0.17212 | 2.245195 | 0.02475563 | 0.623052696 |
| *RASSF6* | 2.579071 | 1.150004 | 2.242663 | 0.024918567 | 0.623052696 |
| *CCDC127* | -0.607329 | 0.270847 | -2.242334 | 0.024939792 | 0.623052696 |
| *ACACA* | 0.836169 | 0.372915 | 2.24225 | 0.024945209 | 0.623052696 |
| *KLHL5* | -0.45924 | 0.204923 | -2.241037 | 0.025023666 | 0.623052696 |
| *SUFU* | 0.943795 | 0.421142 | 2.241036 | 0.02502372 | 0.623052696 |
| *AMFR* | 0.649628 | 0.289956 | 2.240437 | 0.025062588 | 0.623052696 |
| *NFE2L3* | 1.500876 | 0.669943 | 2.240303 | 0.025071236 | 0.623052696 |
| *ZNF132* | 1.418344 | 0.633227 | 2.239866 | 0.02509963 | 0.623052696 |
| *GPR160* | 0.742987 | 0.332109 | 2.237178 | 0.025274709 | 0.623962637 |

| | | | | | |
|---|---|---|---|---|---|
| *ENSG00000260228* | 1.45073 | 0.648473 | 2.237147 | 0.025276764 | 0.623962637 |
| *CLPB* | -1.607606 | 0.71885 | -2.236358 | 0.025328331 | 0.623962637 |
| *FAM204A* | 0.238035 | 0.106452 | 2.236085 | 0.025346216 | 0.623962637 |
| *SLC25A19* | 0.438057 | 0.195953 | 2.23552 | 0.025383205 | 0.623962637 |
| *GGA2* | 0.24689 | 0.110536 | 2.233578 | 0.025510854 | 0.624713394 |
| *STK17A* | 0.276436 | 0.123765 | 2.233551 | 0.025512632 | 0.624713394 |
| *NADSYN1* | -0.378716 | 0.169628 | -2.232624 | 0.025573773 | 0.624838294 |
| *SC5D* | -0.441357 | 0.197743 | -2.231975 | 0.025616639 | 0.624838294 |
| *PIP5K1B* | 0.732446 | 0.328736 | 2.22807 | 0.025875875 | 0.628737691 |
| *LINC01781* | 1.074229 | 0.482135 | 2.228067 | 0.025876027 | 0.628737691 |
| *ZNF254* | -0.760826 | 0.341653 | -2.226894 | 0.025954374 | 0.629430941 |
| *TECPR2* | -1.399737 | 0.628882 | -2.225755 | 0.026030572 | 0.6300695 |
| *CLCF1* | -0.513196 | 0.230864 | -2.222936 | 0.026220103 | 0.633443602 |
| *IKBKB* | 0.358594 | 0.161415 | 2.221562 | 0.02631289 | 0.63447206 |
| *DERL1* | 0.28979 | 0.130529 | 2.220124 | 0.02641034 | 0.635231347 |
| *ABCD1* | 1.220562 | 0.549898 | 2.219615 | 0.02644493 | 0.635231347 |
| *TRIM66* | 0.725483 | 0.327087 | 2.218012 | 0.026554002 | 0.636641006 |
| *TBCB* | -0.235903 | 0.106423 | -2.216653 | 0.026646824 | 0.637203402 |
| *SLC37A1* | -0.550708 | 0.248493 | -2.216192 | 0.026678322 | 0.637203402 |
| *ENSG00000233912* | 1.217653 | 0.549904 | 2.214302 | 0.026808024 | 0.639093168 |
| *SLC25A23* | -1.462637 | 0.660946 | -2.212945 | 0.026901425 | 0.640112068 |
| *C15orf62* | -0.76582 | 0.346279 | -2.211568 | 0.026996526 | 0.64061435 |
| *WBP2NL* | 0.551434 | 0.249385 | 2.211172 | 0.027023937 | 0.64061435 |
| *C5orf56* | 0.406402 | 0.184053 | 2.208066 | 0.027239645 | 0.643511481 |
| *ADSL* | -0.377702 | 0.171138 | -2.207005 | 0.027313681 | 0.643511481 |
| *GPR146* | -0.59958 | 0.271692 | -2.206838 | 0.027325386 | 0.643511481 |
| *LSM11* | 0.872381 | 0.395492 | 2.205811 | 0.027397236 | 0.643511481 |
| *PPP1R3E* | 0.337426 | 0.152975 | 2.20576 | 0.027400805 | 0.643511481 |
| *UXT* | -0.141656 | 0.064256 | -2.20456 | 0.027484957 | 0.643886047 |
| *HEATR6* | -0.75212 | 0.34124 | -2.204081 | 0.027518676 | 0.643886047 |
| *PLD4* | -1.053973 | 0.478647 | -2.201987 | 0.027666253 | 0.644575006 |
| *HCG11* | -0.638142 | 0.289849 | -2.201639 | 0.027690794 | 0.644575006 |
| *ARHGEF12* | -1.018518 | 0.462825 | -2.200655 | 0.027760475 | 0.644575006 |
| *LINC02256* | 0.415055 | 0.18863 | 2.200367 | 0.027780901 | 0.644575006 |
| *DCP1A* | 0.302748 | 0.137632 | 2.199692 | 0.027828778 | 0.644575006 |
| *CLEC16A* | -0.689991 | 0.313763 | -2.199084 | 0.027871929 | 0.644575006 |
| *SLX4* | -1.147494 | 0.522145 | -2.197652 | 0.027973913 | 0.644575006 |
| *CD44* | -0.264719 | 0.120502 | -2.196811 | 0.02803393 | 0.644575006 |
| *FMC1* | 0.261218 | 0.118963 | 2.195793 | 0.028106749 | 0.644575006 |
| *MPC2* | -0.278091 | 0.126709 | -2.19472 | 0.028183682 | 0.644575006 |
| *DHRS3* | 2.518512 | 1.14779 | 2.194227 | 0.028219068 | 0.644575006 |
| *BCAP31* | 0.233643 | 0.106491 | 2.194015 | 0.028234355 | 0.644575006 |
| *TAF12* | 0.261137 | 0.119024 | 2.19398 | 0.028236829 | 0.644575006 |
| *RAB20* | 1.155632 | 0.526871 | 2.193389 | 0.028279355 | 0.644575006 |
| *ENSG00000281325* | -0.768133 | 0.350279 | -2.192917 | 0.028313346 | 0.644575006 |
| *HLA.DOB* | 0.436499 | 0.199126 | 2.192073 | 0.028374239 | 0.644799471 |
| *SHISAL2A* | 0.313166 | 0.142957 | 2.19064 | 0.02847786 | 0.645657205 |

| | | | | | |
|---|---|---|---|---|---|
| *RHOBTB3* | 1.373052 | 0.62714 | 2.189387 | 0.028568691 | 0.645657205 |
| *CNP* | 0.3339 | 0.152525 | 2.189151 | 0.028585834 | 0.645657205 |
| *GPR18* | 0.532041 | 0.243082 | 2.188731 | 0.028616386 | 0.645657205 |
| *SEC22A* | 0.56845 | 0.25994 | 2.186848 | 0.028753653 | 0.647597861 |
| *TRAPPC6A* | 0.238975 | 0.109368 | 2.185052 | 0.028885012 | 0.649319341 |
| *COA4* | 0.259064 | 0.118642 | 2.183575 | 0.028993492 | 0.649319341 |
| *MTX3* | -1.015116 | 0.465076 | -2.182688 | 0.029058829 | 0.649319341 |
| *FLOT1* | 0.3034 | 0.139024 | 2.182365 | 0.029082572 | 0.649319341 |
| *ZBTB41* | -0.828926 | 0.379967 | -2.181574 | 0.029141012 | 0.649319341 |
| *SNAPC3* | 0.336785 | 0.154381 | 2.181518 | 0.0291451 | 0.649319341 |
| *ENSG00000251034* | 1.012695 | 0.46449 | 2.180229 | 0.029240466 | 0.649319341 |
| *ENSG00000274292* | 1.246521 | 0.571741 | 2.180219 | 0.029241211 | 0.649319341 |
| *UTP20* | 0.594243 | 0.272766 | 2.17858 | 0.029362895 | 0.650877512 |
| *LMAN1* | 0.387071 | 0.177753 | 2.17758 | 0.029437292 | 0.651383866 |
| *TLR6* | -0.573843 | 0.263732 | -2.175853 | 0.02956626 | 0.651542107 |
| *ZSCAN18* | 0.250816 | 0.11528 | 2.175702 | 0.029577564 | 0.651542107 |
| *TTC30B* | 1.062807 | 0.488592 | 2.175244 | 0.029611822 | 0.651542107 |
| *TCEAL9* | -1.706381 | 0.785055 | -2.173582 | 0.029736532 | 0.651542107 |
| *TSPYL2* | 0.421014 | 0.193703 | 2.173499 | 0.029742744 | 0.651542107 |
| *L1CAM* | -1.423521 | 0.654989 | -2.173352 | 0.029753842 | 0.651542107 |
| *LUC7L* | -0.340324 | 0.15668 | -2.172093 | 0.02984866 | 0.652487579 |
| *SNAP47* | 0.289037 | 0.133183 | 2.170232 | 0.029989299 | 0.653883832 |
| *HLA.DOA* | 0.307293 | 0.141618 | 2.169879 | 0.030016037 | 0.653883832 |
| *GPATCH2* | -0.567211 | 0.261523 | -2.168874 | 0.03009226 | 0.654416024 |
| *MTERF4* | 0.278899 | 0.128697 | 2.167099 | 0.03022732 | 0.655915529 |
| *MBNL3* | 0.590262 | 0.272458 | 2.16643 | 0.030278332 | 0.655915529 |
| *PRDM4* | 0.534088 | 0.246587 | 2.165925 | 0.03031695 | 0.655915529 |
| *ENSG00000268027* | 0.43425 | 0.200802 | 2.162577 | 0.030573707 | 0.660339815 |
| *BBS12* | -1.468608 | 0.679452 | -2.161459 | 0.03065989 | 0.661071185 |
| *QDPR* | -0.987717 | 0.457362 | -2.159595 | 0.030804056 | 0.662082344 |
| *CTNND1* | -0.490595 | 0.227224 | -2.159079 | 0.03084405 | 0.662082344 |
| *RBM28* | 0.372015 | 0.172323 | 2.158822 | 0.030863989 | 0.662082344 |
| *PREPL* | 0.368373 | 0.17072 | 2.157755 | 0.030946887 | 0.662103571 |
| *SPIN3* | 1.879501 | 0.871224 | 2.15731 | 0.030981484 | 0.662103571 |
| *ZNF92* | 0.25114 | 0.116442 | 2.156788 | 0.031022186 | 0.662103571 |
| *SUMF2* | 0.225372 | 0.104617 | 2.154254 | 0.031220227 | 0.665206686 |
| *CD24* | -0.239238 | 0.111133 | -2.152718 | 0.031340876 | 0.666653139 |
| *SERINC2* | 1.44335 | 0.670836 | 2.15157 | 0.031431231 | 0.667126857 |
| *VPS50* | 0.553901 | 0.257497 | 2.151094 | 0.031468746 | 0.667126857 |
| *TRIM14* | 0.625171 | 0.291253 | 2.14649 | 0.031833924 | 0.672821883 |
| *BTG2* | -0.198453 | 0.09246 | -2.146365 | 0.031843885 | 0.672821883 |
| *NPM3* | -0.490237 | 0.228479 | -2.145655 | 0.031900511 | 0.67289309 |
| *SH3D19* | 0.82711 | 0.386655 | 2.139142 | 0.032424137 | 0.682798291 |
| *ZNF699* | 0.509133 | 0.2382 | 2.137419 | 0.032563964 | 0.682911044 |
| *TUT1* | 0.489672 | 0.22921 | 2.136349 | 0.032650972 | 0.682911044 |
| *FCHO1* | 0.48166 | 0.225565 | 2.135351 | 0.032732322 | 0.682911044 |
| *ACTA2* | 1.483712 | 0.694979 | 2.134901 | 0.03276911 | 0.682911044 |

| | | | | | |
|---|---|---|---|---|---|
| *AIMP1* | -0.254614 | 0.119272 | -2.13474 | 0.032782261 | 0.682911044 |
| *REST* | 0.256922 | 0.12036 | 2.134619 | 0.032792122 | 0.682911044 |
| *CHD1* | 0.25635 | 0.120102 | 2.134427 | 0.032807836 | 0.682911044 |
| *LAIR1* | 0.508902 | 0.238712 | 2.131866 | 0.033017838 | 0.686118431 |
| *SF3A1* | 0.357428 | 0.167763 | 2.130554 | 0.03312593 | 0.686118431 |
| *PRKACB* | 0.353013 | 0.165731 | 2.130041 | 0.033168201 | 0.686118431 |
| *ABHD15* | 0.341429 | 0.160302 | 2.129909 | 0.033179134 | 0.686118431 |
| *RIN3* | -0.40897 | 0.192174 | -2.128123 | 0.033326857 | 0.688047117 |
| *ZFP36L2* | -0.383412 | 0.180348 | -2.125949 | 0.03350752 | 0.689671883 |
| *GSN* | -0.699171 | 0.328888 | -2.125862 | 0.033514724 | 0.689671883 |
| *KDELR1* | -0.279378 | 0.131502 | -2.124513 | 0.03362723 | 0.690861865 |
| *S100A11* | -0.388031 | 0.183057 | -2.119728 | 0.03402901 | 0.696649329 |
| *SP100* | 0.180055 | 0.084959 | 2.119305 | 0.034064734 | 0.696649329 |
| *TMEM256* | -0.176429 | 0.083253 | -2.119191 | 0.03407434 | 0.696649329 |
| *SH3GLB1* | -0.236761 | 0.111856 | -2.116659 | 0.034288828 | 0.698608424 |
| *ERAL1* | -0.518651 | 0.245183 | -2.11536 | 0.034399239 | 0.698608424 |
| *ZFYVE19* | 0.365614 | 0.172841 | 2.115325 | 0.034402255 | 0.698608424 |
| *YPEL3* | 0.263204 | 0.124456 | 2.114837 | 0.03444384 | 0.698608424 |
| *SLC38A1* | -0.235704 | 0.111474 | -2.114425 | 0.034478998 | 0.698608424 |
| *PLCE1* | 0.431497 | 0.204099 | 2.114156 | 0.034501912 | 0.698608424 |
| *SFXN1* | -0.433175 | 0.204962 | -2.113444 | 0.034562825 | 0.698722073 |
| *IFT172* | 0.701767 | 0.332651 | 2.109619 | 0.034891211 | 0.70423395 |
| *NUP88* | 0.309631 | 0.146876 | 2.108115 | 0.035021041 | 0.704438711 |
| *NME7* | 0.43872 | 0.208128 | 2.107931 | 0.035036942 | 0.704438711 |
| *CERS4* | -0.300979 | 0.142809 | -2.107565 | 0.035068615 | 0.704438711 |
| *ENSG00000250222* | 0.962686 | 0.457249 | 2.105387 | 0.035257666 | 0.707112086 |
| *KLHDC3* | 0.237298 | 0.112776 | 2.104166 | 0.035363952 | 0.7080658 |
| *ENSG00000282988* | 0.579271 | 0.275452 | 2.102988 | 0.035466847 | 0.7080658 |
| *ENSG00000124593* | -1.082636 | 0.514827 | -2.102913 | 0.03547334 | 0.7080658 |
| *CACTIN* | 0.674938 | 0.32116 | 2.101561 | 0.035591755 | 0.708892787 |
| *ZNF219* | -1.297419 | 0.617478 | -2.101159 | 0.035626982 | 0.708892787 |
| *ATP11A* | -0.706412 | 0.336717 | -2.097943 | 0.035910209 | 0.711931657 |
| *SH3BP1* | -0.540626 | 0.257863 | -2.096564 | 0.036032224 | 0.711931657 |
| *SOBP* | 1.498224 | 0.714731 | 2.096207 | 0.036063837 | 0.711931657 |
| *ENSG00000130749* | 0.396124 | 0.189012 | 2.095757 | 0.036103724 | 0.711931657 |
| *ZNF341* | 0.792604 | 0.378299 | 2.095181 | 0.036154891 | 0.711931657 |
| *CNNM3* | 0.753244 | 0.359681 | 2.094203 | 0.03624186 | 0.711931657 |
| *ATP6V0A1* | 0.43907 | 0.209665 | 2.09415 | 0.036246642 | 0.711931657 |
| *ADAM10* | -0.351164 | 0.167703 | -2.093956 | 0.03626391 | 0.711931657 |
| *TALDO1* | -0.180999 | 0.086449 | -2.093699 | 0.036286821 | 0.711931657 |
| *NEK5* | -0.694951 | 0.332047 | -2.092931 | 0.036355354 | 0.712170391 |
| *CXCR4* | 0.102829 | 0.049158 | 2.091806 | 0.03645585 | 0.713033542 |
| *SRSF8* | 0.175442 | 0.083906 | 2.090932 | 0.036534182 | 0.713461187 |
| *PACRGL* | -1.322773 | 0.633279 | -2.088769 | 0.036728525 | 0.714167985 |
| *GALT* | 0.285382 | 0.136685 | 2.08789 | 0.036807729 | 0.714167985 |
| *NT5E* | 0.546445 | 0.261723 | 2.087879 | 0.036808733 | 0.714167985 |
| *DDB2* | -0.317375 | 0.152018 | -2.087738 | 0.036821444 | 0.714167985 |

| | | | | | |
|---|---|---|---|---|---|
| *RAPGEF5* | -1.279098 | 0.612774 | -2.087389 | 0.03685299 | 0.714167985 |
| *CDC42SE2* | 0.233307 | 0.111837 | 2.086141 | 0.036965815 | 0.715257377 |
| *ENSG00000133961* | 0.245553 | 0.117744 | 2.085477 | 0.03702599 | 0.715326271 |
| *KDM3B* | 0.356287 | 0.17093 | 2.084409 | 0.037123 | 0.716105495 |
| *UQCR11* | 0.169444 | 0.081347 | 2.082966 | 0.037254287 | 0.717542551 |
| *WARS2* | -0.932644 | 0.448177 | -2.080971 | 0.037436564 | 0.719955842 |
| *NOC2L* | 0.251269 | 0.120836 | 2.079421 | 0.037578664 | 0.720615042 |
| *HACD2* | -0.438897 | 0.211092 | -2.079173 | 0.037601417 | 0.720615042 |
| *ATP2A1.AS1* | -1.63471 | 0.786397 | -2.078733 | 0.037641941 | 0.720615042 |
| *SPOP* | 0.243103 | 0.117016 | 2.077523 | 0.03775336 | 0.721654625 |
| *ENSG00000277825* | 0.992944 | 0.478287 | 2.076041 | 0.037890155 | 0.723175387 |
| *SLC12A6* | 0.307395 | 0.148144 | 2.074983 | 0.037988131 | 0.723951784 |
| *HIC1* | 2.343222 | 1.129301 | 2.074931 | 0.03799297 | NA |
| *EXD3* | -0.564914 | 0.272402 | -2.073821 | 0.038095947 | 0.724913081 |
| *CD37* | -0.087913 | 0.042425 | -2.072222 | 0.038244759 | 0.726650414 |
| *PCF11.AS1* | 0.815818 | 0.393884 | 2.071215 | 0.038338689 | 0.727341346 |
| *ENSG00000233038* | 3.800671 | 1.836036 | 2.070042 | 0.038448405 | NA |
| *ENSG00000272994* | 0.399917 | 0.193314 | 2.068745 | 0.038569983 | 0.728381965 |
| *SMPD1* | 0.478095 | 0.231145 | 2.06838 | 0.038604349 | 0.728381965 |
| *ZFYVE16* | -0.433854 | 0.209803 | -2.067908 | 0.038648682 | 0.728381965 |
| *INPP5B* | 0.421953 | 0.204068 | 2.067711 | 0.038667203 | 0.728381965 |
| *CDK13* | 0.345609 | 0.167176 | 2.067343 | 0.038701825 | 0.728381965 |
| *ICOSLG* | -0.289957 | 0.140334 | -2.066195 | 0.038810116 | 0.728381965 |
| *SYTL1* | -0.296991 | 0.143742 | -2.066139 | 0.038815391 | 0.728381965 |
| *ATP5MG* | -0.139275 | 0.067422 | -2.065722 | 0.038854725 | 0.728381965 |
| *PPP1R3F* | 1.352746 | 0.655104 | 2.064934 | 0.038929235 | 0.728697608 |
| *CYB561* | 1.160127 | 0.562254 | 2.063349 | 0.0390795 | 0.729784367 |
| *RCC1L* | 0.551677 | 0.267401 | 2.063103 | 0.039102811 | 0.729784367 |
| *KCNH4* | 1.781503 | 0.864952 | 2.059656 | 0.039431425 | 0.73420698 |
| *TLR10* | -0.456937 | 0.221879 | -2.059399 | 0.039455998 | 0.73420698 |
| *MOB1B* | -1.025403 | 0.498178 | -2.058306 | 0.039560727 | 0.734301946 |
| *LPCAT4* | -0.445491 | 0.216504 | -2.057656 | 0.03962316 | 0.734301946 |
| *ICA1L* | 0.356813 | 0.173452 | 2.057128 | 0.039673898 | 0.734301946 |
| *HERPUD1* | -0.196352 | 0.095459 | -2.056924 | 0.039693568 | 0.734301946 |
| *CPLANE2* | 1.07898 | 0.525378 | 2.053721 | 0.04000267 | 0.738938211 |
| *ARHGDIB* | 0.136773 | 0.066649 | 2.052142 | 0.040155833 | 0.739000925 |
| *MYO1C* | -0.555726 | 0.270812 | -2.052075 | 0.040162393 | 0.739000925 |
| *EMC7* | 0.255531 | 0.12459 | 2.050983 | 0.040268632 | 0.739000925 |
| *ARMC2* | 1.271583 | 0.620013 | 2.050898 | 0.04027685 | 0.739000925 |
| *TMEM154* | 0.329512 | 0.160685 | 2.050676 | 0.040298507 | 0.739000925 |
| *RASA4* | 0.601647 | 0.29358 | 2.049346 | 0.040428269 | 0.740306056 |
| *ENSG00000262429* | 0.937988 | 0.4586 | 2.04533 | 0.040822294 | 0.743590377 |
| *FCRLB* | -1.487255 | 0.72729 | -2.044927 | 0.040862071 | 0.743590377 |
| *UBLCP1* | -0.428732 | 0.209658 | -2.04491 | 0.040863764 | 0.743590377 |
| *SERPINF1* | -0.839981 | 0.411 | -2.04375 | 0.040978289 | 0.743590377 |
| *ENSG00000273188* | 0.831819 | 0.407032 | 2.043621 | 0.040991024 | 0.743590377 |
| *EHD1* | 0.290315 | 0.14211 | 2.042895 | 0.041062799 | 0.743590377 |

| | | | | | |
|---|---|---|---|---|---|
| *VOPP1* | -0.258127 | 0.126357 | -2.042846 | 0.041067733 | 0.743590377 |
| *FGD2* | 0.242216 | 0.118574 | 2.042737 | 0.041078439 | 0.743590377 |
| *DNAH1* | 0.99446 | 0.487061 | 2.041756 | 0.041175748 | 0.744285511 |
| *DDX24* | 0.195025 | 0.095587 | 2.040278 | 0.04132261 | 0.745873118 |
| *NAA30* | -0.50431 | 0.247411 | -2.038346 | 0.041515383 | 0.748283678 |
| *FAM81A* | -1.704392 | 0.836882 | -2.036599 | 0.041690239 | 0.750364906 |
| *TMEM65* | 0.636569 | 0.31275 | 2.035396 | 0.041811031 | 0.751468529 |
| *SUGP2* | -0.332255 | 0.163319 | -2.03439 | 0.041912259 | 0.752217892 |
| *OGFOD3* | -0.529047 | 0.260384 | -2.031797 | 0.042174184 | 0.755845121 |
| *MYBL2* | -1.465411 | 0.721794 | -2.030233 | 0.04233281 | 0.756548845 |
| *ABLIM2* | -0.645798 | 0.318091 | -2.03023 | 0.042333204 | 0.756548845 |
| *ENSG00000257275* | -0.288028 | 0.141951 | -2.029061 | 0.042452064 | 0.757005681 |
| *CD27* | -1.119643 | 0.552008 | -2.028311 | 0.042528478 | 0.757005681 |
| *RESF1* | -0.243124 | 0.11993 | -2.027218 | 0.042640159 | 0.757005681 |
| *HNRNPH2* | 0.267236 | 0.131891 | 2.026185 | 0.042745826 | 0.757005681 |
| *TGIF2* | -0.355489 | 0.175477 | -2.025842 | 0.042781016 | 0.757005681 |
| *ATP2B1* | -0.176809 | 0.087277 | -2.025838 | 0.042781353 | 0.757005681 |
| *SLC9A9* | 0.474443 | 0.234225 | 2.025588 | 0.042806983 | 0.757005681 |
| *HMGN1* | -0.124636 | 0.06154 | -2.025285 | 0.042838074 | 0.757005681 |
| *RAB4A* | -0.289522 | 0.143073 | -2.023605 | 0.043010769 | 0.758495733 |
| *PTP4A3* | 1.258874 | 0.622664 | 2.021755 | 0.043201708 | 0.758495733 |
| *TMEM30A* | -0.38882 | 0.192363 | -2.02128 | 0.043250811 | 0.758495733 |
| *EAF2* | 0.301543 | 0.149207 | 2.020969 | 0.043282976 | 0.758495733 |
| *CIZ1* | 0.333585 | 0.165114 | 2.02033 | 0.043349151 | 0.758495733 |
| *SYNGAP1* | 0.674213 | 0.333836 | 2.019591 | 0.043425816 | 0.758495733 |
| *EIF4ENIF1* | 0.55961 | 0.277104 | 2.019493 | 0.043435976 | 0.758495733 |
| *ISCU* | -0.164427 | 0.081456 | -2.01861 | 0.043527727 | 0.758495733 |
| *CCR7* | 0.205454 | 0.101879 | 2.016647 | 0.043732366 | 0.758495733 |
| *ENSG00000179743* | 0.378122 | 0.187524 | 2.016387 | 0.043759479 | 0.758495733 |
| *DNAL1* | 0.540311 | 0.26798 | 2.016231 | 0.043775798 | 0.758495733 |
| *QSOX2* | -0.481164 | 0.238749 | -2.015354 | 0.043867569 | 0.758495733 |
| *MTRR* | -0.601987 | 0.298807 | -2.014634 | 0.04394301 | 0.758495733 |
| *MAP3K7CL* | 0.912051 | 0.453039 | 2.013185 | 0.044095152 | 0.758495733 |
| *MR1* | 0.388923 | 0.193249 | 2.01255 | 0.044161986 | 0.758495733 |
| *ENSG00000273702* | 0.617712 | 0.307071 | 2.011627 | 0.04425924 | 0.758495733 |
| *FASTK* | 0.233334 | 0.116001 | 2.011473 | 0.044275569 | 0.758495733 |
| *GOLGA2* | -0.430175 | 0.21392 | -2.010911 | 0.044334909 | 0.758495733 |
| *MCM5* | 0.345623 | 0.171879 | 2.01085 | 0.044341311 | 0.758495733 |
| *KNL1* | 0.644069 | 0.320452 | 2.009877 | 0.04444423 | 0.758495733 |
| *ENSG00000276728* | 0.447097 | 0.222451 | 2.009866 | 0.044445357 | 0.758495733 |
| *PATJ* | 0.566934 | 0.282083 | 2.009815 | 0.044450775 | 0.758495733 |
| *WWP2* | -0.519236 | 0.25836 | -2.009737 | 0.044459074 | 0.758495733 |
| *TYW3* | -0.382347 | 0.190284 | -2.009345 | 0.044500576 | 0.758495733 |
| *CBY1* | -0.756685 | 0.376628 | -2.009105 | 0.044525994 | 0.758495733 |
| *CLN8* | -0.348765 | 0.173658 | -2.008347 | 0.044606436 | 0.758495733 |
| *CWC27* | 0.307716 | 0.153234 | 2.008153 | 0.044627045 | 0.758495733 |
| *RBIS* | -0.191284 | 0.095255 | -2.008121 | 0.044630416 | 0.758495733 |

| | | | | | |
|---|---|---|---|---|---|
| *FLII* | -0.354527 | 0.176574 | -2.007812 | 0.044663302 | 0.758495733 |
| *SKIL* | 0.265892 | 0.132516 | 2.006484 | 0.044804596 | 0.759071263 |
| *GID8* | -0.248046 | 0.12366 | -2.005877 | 0.044869426 | 0.759071263 |
| *MAGEF1* | 0.329518 | 0.164282 | 2.005802 | 0.044877423 | 0.759071263 |
| *SNN* | 0.403436 | 0.201253 | 2.00462 | 0.045003695 | 0.760189425 |
| *TXLNB* | -1.284842 | 0.642049 | -2.001158 | 0.045375354 | 0.762899353 |
| *CDKN1A* | 0.362049 | 0.180946 | 2.000872 | 0.045406215 | 0.762899353 |
| *LINC01800* | -1.431797 | 0.71568 | -2.000612 | 0.045434225 | 0.762899353 |
| *RAB5IF* | -0.225589 | 0.11279 | -2.000088 | 0.045490769 | 0.762899353 |
| *CPTP* | -0.469557 | 0.23484 | -1.99948 | 0.045556467 | 0.762899353 |
| *ATM* | 0.218922 | 0.109499 | 1.99931 | 0.045574812 | 0.762899353 |
| *MARK4* | -0.843457 | 0.421897 | -1.999199 | 0.045586784 | 0.762899353 |
| *MCM9* | 0.482461 | 0.241467 | 1.998042 | 0.045712136 | 0.763794243 |
| *TCTN1* | -0.598982 | 0.299852 | -1.99759 | 0.045761159 | 0.763794243 |
| *RABAC1* | 0.166206 | 0.083371 | 1.993556 | 0.046200616 | 0.770111849 |
| *CLIC4* | -0.420929 | 0.211216 | -1.992883 | 0.046274287 | 0.770323612 |
| *KCTD20* | -0.334211 | 0.167882 | -1.990752 | 0.046508139 | 0.773197807 |
| *HSD17B14* | -1.507572 | 0.757755 | -1.989523 | 0.04664348 | 0.774428877 |
| *MRNIP* | 0.283273 | 0.142448 | 1.988607 | 0.046744586 | 0.775089042 |
| *CCR9* | 2.068483 | 1.040811 | 1.987377 | 0.04688067 | 0.776326697 |
| *BCAS4* | 0.327796 | 0.165015 | 1.986457 | 0.04698256 | 0.776995616 |
| *ARHGEF9* | 0.420172 | 0.21162 | 1.985501 | 0.047088787 | 0.777734403 |
| *LEPROTL1* | 0.345382 | 0.174029 | 1.984621 | 0.047186667 | 0.778333594 |
| *NQO1* | 1.17675 | 0.593213 | 1.983688 | 0.04729064 | 0.779031607 |
| *SGMS1* | 0.463294 | 0.233819 | 1.981423 | 0.047543863 | 0.781354689 |
| *HIP1R* | 0.276783 | 0.139727 | 1.98088 | 0.047604729 | 0.781354689 |
| *TERF2IP* | 0.238609 | 0.120494 | 1.980254 | 0.047674958 | 0.781354689 |
| *S100A6* | -0.444495 | 0.22453 | -1.979671 | 0.04774049 | 0.781354689 |
| *L3MBTL1* | -1.064023 | 0.537475 | -1.979668 | 0.047740864 | 0.781354689 |
| *PACS1* | 0.277686 | 0.140382 | 1.978078 | 0.047919937 | 0.783200105 |
| *ENSG00000282393* | 0.921454 | 0.465953 | 1.977567 | 0.047977593 | 0.783200105 |
| *PLSCR1* | -0.380906 | 0.192667 | -1.977017 | 0.048039728 | 0.783202533 |
| *ARHGEF3* | 0.457083 | 0.231327 | 1.975916 | 0.048164266 | 0.784221006 |
| *TACC3* | 0.379616 | 0.192209 | 1.975023 | 0.048265574 | 0.784367271 |
| *ENTPD1* | -0.395833 | 0.200448 | -1.974742 | 0.048297407 | 0.784367271 |
| *MREG* | -1.043232 | 0.52844 | -1.974172 | 0.048362222 | 0.784411642 |
| *GRAP* | 0.406186 | 0.205906 | 1.972674 | 0.048532713 | 0.785201836 |
| *LINC01857* | 0.408394 | 0.207028 | 1.972652 | 0.04853523 | 0.785201836 |
| *NAPA* | -0.375885 | 0.19064 | -1.971704 | 0.048643368 | 0.785588294 |
| *GABARAPL1* | 0.614577 | 0.311754 | 1.971353 | 0.048683469 | 0.785588294 |
| *ARL16* | -0.377823 | 0.191777 | -1.970118 | 0.0488248 | 0.785909779 |
| *GARS* | 0.35652 | 0.180966 | 1.970092 | 0.048827794 | 0.785909779 |
| *SLC51A* | 1.010721 | 0.513342 | 1.968905 | 0.048964038 | 0.787100028 |
| *MAP4* | -0.258308 | 0.131281 | -1.967593 | 0.04911492 | 0.788522256 |
| *ZFAND2B* | -0.259967 | 0.132169 | -1.966935 | 0.049190696 | 0.788736604 |
| *ENSG00000265206* | -0.210333 | 0.107078 | -1.964296 | 0.049495804 | 0.791826839 |

| Cluster 2 | log2FC | lfcSE | stat | pvalue | padj |
|---|---|---|---|---|---|
| *VPREB3* | 3.034294 | 0.09097 | 33.35473 | < 2.22e-16 | < 2.22e-16 |
| *CD38* | 3.051433 | 0.245412 | 12.433914 | < 2.22e-16 | < 2.22e-16 |
| *PLAAT4* | 2.141232 | 0.191363 | 11.189352 | < 2.22e-16 | < 2.22e-16 |
| *RARA.AS1* | 2.480806 | 0.3052 | 8.128454 | 4.35E-16 | 9.66E-13 |
| *AHI1* | 1.631033 | 0.201404 | 8.098314 | 5.57E-16 | 9.90E-13 |
| *DUS2* | 1.54804 | 0.1959 | 7.902192 | 2.74E-15 | 4.06E-12 |
| *SLC12A4* | 2.387136 | 0.30835 | 7.741645 | 9.81E-15 | 1.25E-11 |
| *RRBP1* | 1.550321 | 0.20328 | 7.626543 | 2.41E-14 | 2.68E-11 |
| *FCRL1* | 1.141723 | 0.153163 | 7.454285 | 9.04E-14 | 8.92E-11 |
| *P2RX5.TAX1BP3* | 1.854819 | 0.255359 | 7.263568 | 3.77E-13 | 3.35E-10 |
| *LCAT* | 2.825853 | 0.39723 | 7.113893 | 1.13E-12 | NA |
| *RAB37* | 1.279916 | 0.185126 | 6.913766 | 4.72E-12 | 3.81E-09 |
| *TRABD* | 0.633739 | 0.092243 | 6.870337 | 6.41E-12 | 4.74E-09 |
| *CD1D* | 2.367338 | 0.346709 | 6.828033 | 8.61E-12 | 5.88E-09 |
| *CD72* | 0.992145 | 0.146633 | 6.766193 | 1.32E-11 | 8.39E-09 |
| *P2RX5* | 1.032489 | 0.153563 | 6.723568 | 1.77E-11 | 1.05E-08 |
| *SPRY1* | 1.113203 | 0.170928 | 6.512705 | 7.38E-11 | 4.10E-08 |
| *FAM3C* | 0.701706 | 0.109645 | 6.399772 | 1.56E-10 | 8.13E-08 |
| *SEPTIN9* | 0.600936 | 0.094237 | 6.376852 | 1.81E-10 | 8.92E-08 |
| *ICAM2* | 0.911121 | 0.14622 | 6.231178 | 4.63E-10 | 2.17E-07 |
| *DENND6B* | 1.960659 | 0.315262 | 6.219136 | 5.00E-10 | 2.22E-07 |
| *AGPAT5* | 0.788069 | 0.127548 | 6.178608 | 6.47E-10 | 2.74E-07 |
| *MRPS6* | 0.867285 | 0.145842 | 5.946723 | 2.74E-09 | 1.10E-06 |
| *AGO1* | 1.077876 | 0.182502 | 5.906115 | 3.50E-09 | 1.35E-06 |
| *DPEP2* | 0.984316 | 0.167034 | 5.892897 | 3.79E-09 | 1.41E-06 |
| *PNOC* | 1.384339 | 0.242749 | 5.702745 | 1.18E-08 | 4.19E-06 |
| *SIGIRR* | 0.770474 | 0.137781 | 5.592008 | 2.24E-08 | 7.67E-06 |
| *RMDN2* | 1.974604 | 0.357137 | 5.528988 | 3.22E-08 | 1.06E-05 |
| *DBNL* | 0.547936 | 0.100017 | 5.478442 | 4.29E-08 | 1.36E-05 |
| *LINC.PINT* | 0.581551 | 0.109126 | 5.329166 | 9.87E-08 | 3.02E-05 |
| *MPEG1* | 1.15014 | 0.21742 | 5.289932 | 1.22E-07 | 3.62E-05 |
| *PLAAT3* | 2.976095 | 0.575361 | 5.172568 | 2.31E-07 | NA |
| *CD79B* | 0.31698 | 0.061772 | 5.131428 | 2.88E-07 | 8.24E-05 |
| *MS4A1* | -0.511386 | 0.100792 | -5.073662 | 3.90E-07 | 0.000108362 |
| *ANAPC16* | 0.329325 | 0.065126 | 5.056728 | 4.27E-07 | 0.000114848 |
| *GRN* | 0.717523 | 0.146776 | 4.888571 | 1.02E-06 | 0.000265457 |
| *ATXN10* | 0.636677 | 0.137254 | 4.638691 | 3.51E-06 | 0.000890181 |
| *CD180* | 0.806356 | 0.17532 | 4.599329 | 4.24E-06 | 0.001046215 |
| *C12orf65* | 0.638003 | 0.141541 | 4.507559 | 6.56E-06 | 0.001574925 |
| *H3F3A* | 0.377295 | 0.085155 | 4.430708 | 9.39E-06 | 0.002196346 |
| *CLEC17A* | 0.901524 | 0.203849 | 4.422517 | 9.76E-06 | 0.002222817 |
| *FAM117A* | 0.690666 | 0.157203 | 4.393468 | 1.12E-05 | 0.002478231 |
| *NCF1* | 0.315745 | 0.072588 | 4.349795 | 1.36E-05 | 0.002953296 |
| *SLC5A3* | 0.760359 | 0.175875 | 4.32329 | 1.54E-05 | 0.003252263 |
| *SMAD3* | 0.725938 | 0.168691 | 4.303366 | 1.68E-05 | 0.003424573 |
| *GBP4* | 0.648942 | 0.150861 | 4.301597 | 1.70E-05 | 0.003424573 |

| | | | | | |
|---|---|---|---|---|---|
| *MRPS36* | 0.4364 | 0.101629 | 4.294071 | 1.75E-05 | 0.003464082 |
| *CD83* | -0.307799 | 0.073406 | -4.193126 | 2.75E-05 | 0.005314908 |
| *RABGAP1L* | 0.662136 | 0.158571 | 4.175634 | 2.97E-05 | 0.005618159 |
| *PTK2B* | 0.61848 | 0.149258 | 4.143692 | 3.42E-05 | 0.00620956 |
| *PXK* | 0.524563 | 0.126607 | 4.143253 | 3.42E-05 | 0.00620956 |
| *JAK1* | 0.345714 | 0.083953 | 4.117949 | 3.82E-05 | 0.006793517 |
| *FOXO3* | 1.06231 | 0.258502 | 4.10949 | 3.97E-05 | 0.006909032 |
| *C16orf54* | 0.727527 | 0.177723 | 4.09361 | 4.25E-05 | 0.007257618 |
| *CD22* | 0.692782 | 0.170625 | 4.060254 | 4.90E-05 | 0.008164017 |
| *FCER2* | 1.426549 | 0.351588 | 4.057445 | 4.96E-05 | 0.008164017 |
| *SPOCK2* | 0.589008 | 0.145329 | 4.052919 | 5.06E-05 | 0.008172292 |
| *PLEKHO1* | 0.548176 | 0.135903 | 4.033581 | 5.49E-05 | 0.008716723 |
| *CXCR4* | 0.167131 | 0.041666 | 4.011164 | 6.04E-05 | 0.009419179 |
| *CD79A* | 0.467968 | 0.116992 | 3.999995 | 6.33E-05 | 0.009594201 |
| *RINL* | 0.57086 | 0.142763 | 3.99866 | 6.37E-05 | 0.009594201 |
| *ENSG00000225489* | 1.243173 | 0.313387 | 3.966893 | 7.28E-05 | 0.010784007 |
| *RUBCN* | -0.620071 | 0.157412 | -3.939163 | 8.18E-05 | 0.01191106 |
| *IRAK2* | 0.943004 | 0.239851 | 3.931624 | 8.44E-05 | 0.012092674 |
| *CHST15* | 0.796998 | 0.203098 | 3.924198 | 8.70E-05 | 0.012273864 |
| *ZNF595* | 1.278442 | 0.32695 | 3.910205 | 9.22E-05 | 0.012645263 |
| *WWP2* | 0.901898 | 0.230696 | 3.909471 | 9.25E-05 | 0.012645263 |
| *CPNE5* | 1.003339 | 0.256921 | 3.905251 | 9.41E-05 | 0.012673004 |
| *ATP6V0A1* | 0.824013 | 0.213807 | 3.854012 | 0.0001162 | 0.015410946 |
| *HSH2D* | 0.352237 | 0.09204 | 3.827006 | 0.00012971 | 0.016950204 |
| *LINC02397* | 0.377386 | 0.09976 | 3.782944 | 0.00015499 | 0.019831511 |
| *NUP210* | 0.456328 | 0.120691 | 3.780961 | 0.00015622 | 0.019831511 |
| *NAGPA.AS1* | 2.091888 | 0.554679 | 3.77135 | 0.00016237 | NA |
| *DPEP3* | 2.176361 | 0.578113 | 3.764595 | 0.00016682 | NA |
| *C8orf89* | 2.214185 | 0.58949 | 3.756103 | 0.00017258 | NA |
| *SLC25A45* | 0.892522 | 0.23864 | 3.740032 | 0.000184 | 0.0230281 |
| *KLF8* | 0.641582 | 0.171802 | 3.734421 | 0.00018815 | 0.023220539 |
| *SERPINB6* | 0.770903 | 0.206916 | 3.725674 | 0.00019479 | 0.023711477 |
| *NFATC3* | 0.526412 | 0.141463 | 3.721197 | 0.00019828 | 0.023809719 |
| *GPR183* | -0.573418 | 0.154236 | -3.717788 | 0.00020098 | 0.023811467 |
| *SLC2A5* | 1.11038 | 0.298976 | 3.713938 | 0.00020406 | 0.02385875 |
| *DCK* | 0.300162 | 0.08092 | 3.70936 | 0.00020778 | 0.023892997 |
| *VNN2* | 0.687887 | 0.185564 | 3.707 | 0.00020973 | 0.023892997 |
| *LINC00513* | 0.64305 | 0.173747 | 3.701079 | 0.00021469 | 0.024147948 |
| *TNFRSF13B* | -1.719854 | 0.466864 | -3.683844 | 0.00022974 | 0.025449635 |
| *LY86* | -0.368345 | 0.100057 | -3.681369 | 0.00023199 | 0.025449635 |
| *TNFRSF18* | -2.210402 | 0.603909 | -3.660155 | 0.00025206 | 0.027314967 |
| *TGFBR2* | -0.479643 | 0.131921 | -3.635837 | 0.00027708 | 0.029664207 |
| *DDIT4* | 1.044095 | 0.288007 | 3.625242 | 0.00028869 | 0.030396426 |
| *IRF4* | 0.830838 | 0.229298 | 3.623396 | 0.00029076 | 0.030396426 |
| *ST3GAL1* | 0.722354 | 0.199544 | 3.620025 | 0.00029457 | 0.030437048 |
| *MAPKAPK2* | 0.739362 | 0.20576 | 3.593326 | 0.00032648 | 0.033346417 |
| *ENSG00000263394* | 0.863715 | 0.24097 | 3.584331 | 0.00033794 | 0.034124546 |

| | | | | | |
|---|---|---|---|---|---|
| *ICA1* | 3.909597 | 1.103795 | 3.541959 | 0.00039717 | NA |
| *ZFP36L1* | -0.331665 | 0.09443 | -3.512286 | 0.00044427 | 0.044357121 |
| *ATP6V1E1* | 0.453004 | 0.129498 | 3.498151 | 0.0004685 | 0.0462562 |
| *BTBD7* | 0.564849 | 0.162038 | 3.485898 | 0.00049049 | 0.047895408 |
| *ZNF563* | 0.885071 | 0.255207 | 3.468046 | 0.00052426 | 0.050636435 |
| *POLM* | 0.453001 | 0.131241 | 3.451675 | 0.00055712 | 0.053231787 |
| *SLC43A2* | 0.455163 | 0.13226 | 3.441422 | 0.00057867 | 0.054702305 |
| *CD82* | -0.492381 | 0.143432 | -3.432846 | 0.00059728 | 0.05586772 |
| *CDKN1A* | 0.583798 | 0.170294 | 3.428168 | 0.00060767 | 0.056247438 |
| *NAPSA* | 1.538831 | 0.449544 | 3.423095 | 0.00061913 | NA |
| *FCMR* | 0.435349 | 0.12748 | 3.41505 | 0.0006377 | 0.056985812 |
| *TMEM154* | 0.483157 | 0.141506 | 3.414389 | 0.00063925 | 0.056985812 |
| *SNX18* | 0.792416 | 0.232295 | 3.411253 | 0.00064665 | 0.056985812 |
| *IRF2BPL* | 0.982681 | 0.288097 | 3.410937 | 0.0006474 | 0.056985812 |
| *XYLT1* | 0.788127 | 0.231068 | 3.410806 | 0.00064771 | 0.056985812 |
| *NMT2* | -0.535934 | 0.157922 | -3.393663 | 0.00068965 | 0.060080327 |
| *ADD1* | 0.492168 | 0.145328 | 3.386614 | 0.00070761 | 0.061046617 |
| *SMARCB1* | 0.335671 | 0.099216 | 3.383247 | 0.00071634 | 0.061205906 |
| *VPREB1* | 2.942504 | 0.873941 | 3.366938 | 0.00076008 | 0.064324317 |
| *LINC02132* | 1.439941 | 0.427899 | 3.365144 | 0.00076504 | NA |
| *S100A4* | -0.955883 | 0.284539 | -3.359414 | 0.00078108 | 0.065477983 |
| *CENPQ* | 0.829763 | 0.247661 | 3.350393 | 0.00080697 | 0.067016196 |
| *ZNF594* | 0.89897 | 0.269653 | 3.333805 | 0.00085667 | 0.070484568 |
| *PCBP1* | -0.283969 | 0.085288 | -3.329518 | 0.00086996 | 0.070821831 |
| *SNRPD3* | 0.332836 | 0.10003 | 3.327368 | 0.00087671 | 0.070821831 |
| *CSTF1* | 0.635894 | 0.192474 | 3.303795 | 0.00095386 | 0.076360032 |
| *CABLES1* | -1.757833 | 0.534508 | -3.288696 | 0.00100653 | 0.07985718 |
| *AKAP12* | 2.524871 | 0.770353 | 3.27755 | 0.00104712 | NA |
| *SEL1L3* | 0.569252 | 0.174335 | 3.265266 | 0.00109361 | 0.085998498 |
| *CIB1* | 0.31545 | 0.096709 | 3.261859 | 0.00110684 | 0.086275488 |
| *PIGX* | -1.144981 | 0.353234 | -3.241423 | 0.00118935 | 0.091900174 |
| *CALHM6* | 0.40424 | 0.124858 | 3.237601 | 0.00120539 | 0.092337292 |
| *CHP1* | -0.477359 | 0.147973 | -3.225978 | 0.00125543 | 0.095348431 |
| *P2RY8* | 0.493458 | 0.153215 | 3.22069 | 0.00127883 | 0.096302025 |
| *CCNG1* | -0.409235 | 0.127266 | -3.215594 | 0.00130175 | 0.096562933 |
| *GMFG* | 0.220015 | 0.068432 | 3.215093 | 0.00130402 | 0.096562933 |
| *PPP1R15B* | 0.713021 | 0.222123 | 3.21003 | 0.00132721 | 0.097467707 |
| *FCHO1* | 0.698151 | 0.217973 | 3.202923 | 0.0013604 | 0.098762607 |
| *NKAPL* | 1.728687 | 0.539817 | 3.202356 | 0.00136308 | NA |
| *SNAP23* | 0.317389 | 0.099137 | 3.201515 | 0.00136707 | 0.098762607 |
| *MTG1* | 0.65184 | 0.203817 | 3.198161 | 0.00138307 | 0.099112725 |
| *ETFB* | 0.492788 | 0.154321 | 3.193273 | 0.0014067 | 0.099999538 |
| *UROS* | 0.509674 | 0.159764 | 3.190176 | 0.00142186 | 0.100275034 |
| *ZNF3* | 0.558199 | 0.175475 | 3.181083 | 0.00146726 | 0.102661612 |
| *LINC02649* | 1.32698 | 0.417498 | 3.178408 | 0.00148086 | NA |
| *ENSG00000214797* | 1.716383 | 0.540774 | 3.173937 | 0.00150386 | NA |
| *ENSG00000231760* | 1.899841 | 0.598717 | 3.173187 | 0.00150775 | NA |

| | | | | | |
|---|---|---|---|---|---|
| *GPX1* | 0.321333 | 0.101854 | 3.154853 | 0.00160579 | 0.111010085 |
| *ENSG00000262292* | 0.869054 | 0.275557 | 3.153807 | 0.00161156 | 0.111010085 |
| *SCN4A* | 2.275882 | 0.723333 | 3.146383 | 0.00165303 | NA |
| *EMC6* | -0.455152 | 0.145744 | -3.122961 | 0.00179042 | 0.122381819 |
| *CD47* | -0.285346 | 0.091687 | -3.11218 | 0.00185711 | 0.125971707 |
| *PSMB10* | 0.362142 | 0.116465 | 3.109448 | 0.00187437 | 0.126031766 |
| *PELI1* | -1.065683 | 0.342932 | -3.107564 | 0.00188636 | 0.126031766 |
| *ENSG00000277602* | 1.495227 | 0.482609 | 3.098213 | 0.00194692 | NA |
| *DNAJC7* | 0.325543 | 0.105174 | 3.095292 | 0.00196619 | 0.130384826 |
| *MFSD10* | -0.418778 | 0.135428 | -3.092248 | 0.00198647 | 0.13065308 |
| *MARCKS* | -0.784003 | 0.253699 | -3.090285 | 0.00199964 | 0.13065308 |
| *ATF6* | -0.74151 | 0.240761 | -3.079863 | 0.00207096 | 0.134325088 |
| *AP3B1* | 0.58747 | 0.191602 | 3.066091 | 0.00216877 | 0.138756114 |
| *SIDT1* | -0.697752 | 0.227588 | -3.065852 | 0.0021705 | 0.138756114 |
| *ENSG00000271204* | 0.54742 | 0.17905 | 3.057353 | 0.00223301 | 0.141732498 |
| *SFXN2* | 1.435299 | 0.470387 | 3.051315 | 0.00227842 | NA |
| *SNAPC2* | -0.78357 | 0.25796 | -3.037565 | 0.00238498 | 0.150304588 |
| *GCHFR* | -0.630412 | 0.209066 | -3.015379 | 0.00256658 | 0.159947013 |
| *ENSG00000273319* | 0.451729 | 0.149852 | 3.014506 | 0.00257398 | 0.159947013 |
| *CENPM* | 0.568498 | 0.188962 | 3.008537 | 0.00262509 | 0.161990065 |
| *SPINT2* | -0.308891 | 0.10275 | -3.006234 | 0.00264505 | 0.162096034 |
| *ARHGAP15* | 0.305227 | 0.102058 | 2.99071 | 0.0027833 | 0.169025782 |
| *PAOX* | 0.671463 | 0.224622 | 2.9893 | 0.00279617 | 0.169025782 |
| *MAPK1* | -0.470702 | 0.157934 | -2.980377 | 0.00287893 | 0.172852755 |
| *FAM86C1* | 1.354672 | 0.455096 | 2.976671 | 0.00291397 | NA |
| *TFG* | 0.467798 | 0.158542 | 2.950626 | 0.00317131 | 0.187535564 |
| *COA1* | 0.400695 | 0.135852 | 2.949499 | 0.00318289 | 0.187535564 |
| *INPP5B* | 0.694187 | 0.235388 | 2.949121 | 0.0031868 | 0.187535564 |
| *CXCR5* | 0.440628 | 0.149564 | 2.94608 | 0.00321829 | 0.188143029 |
| *SH2B2* | 1.685222 | 0.572054 | 2.945912 | 0.00322004 | NA |
| *ENSG00000254777* | 2.094552 | 0.715085 | 2.929094 | 0.00339951 | NA |
| *EWSR1* | 0.386306 | 0.131894 | 2.928921 | 0.00340141 | 0.197548355 |
| *ZNF44* | 0.489026 | 0.167242 | 2.924056 | 0.00345502 | 0.199359236 |
| *MCL1* | 0.35031 | 0.120005 | 2.919137 | 0.00351002 | 0.200392536 |
| *RESF1* | -0.385416 | 0.132063 | -2.918426 | 0.00351803 | 0.200392536 |
| *KCNN1* | 1.972787 | 0.678895 | 2.905879 | 0.00366223 | NA |
| *GPER1* | -2.128222 | 0.736459 | -2.889802 | 0.00385484 | NA |
| *FDFT1* | 0.339286 | 0.117431 | 2.889244 | 0.00386169 | 0.21856695 |
| *ERAP1* | 0.620592 | 0.215271 | 2.88284 | 0.00394108 | 0.22026481 |
| *LHFPL2* | 1.223049 | 0.424254 | 2.882825 | 0.00394127 | 0.22026481 |
| *IFNAR2* | 0.43595 | 0.151788 | 2.872093 | 0.00407763 | 0.226461374 |
| *MAP3K2* | -0.51615 | 0.179882 | -2.869382 | 0.00411275 | 0.226847466 |
| *ENSG00000273783* | 1.875864 | 0.653948 | 2.868521 | 0.00412396 | NA |
| *RNPEPL1* | 0.498494 | 0.173835 | 2.867626 | 0.00413564 | 0.226847466 |
| *ARHGEF3* | 0.639045 | 0.223016 | 2.865463 | 0.004164 | 0.227001588 |
| *HIVEP3* | 1.353874 | 0.472499 | 2.865347 | 0.00416553 | NA |
| *TSPAN18* | 2.510452 | 0.87804 | 2.859154 | 0.00424772 | NA |

| | | | | | |
|---|---|---|---|---|---|
| *ANKRD36C* | -0.718136 | 0.251199 | -2.858828 | 0.00425209 | 0.230390513 |
| *FADS3* | -0.369165 | 0.129286 | -2.855411 | 0.00429811 | 0.230759865 |
| *NRM* | 0.516376 | 0.180901 | 2.854472 | 0.00431084 | 0.230759865 |
| *ENSG00000253535* | 0.895944 | 0.315156 | 2.84286 | 0.00447108 | 0.236567569 |
| *MYC* | 0.395322 | 0.139216 | 2.839628 | 0.00451662 | 0.236567569 |
| *ATP5F1B* | 0.290086 | 0.10222 | 2.837852 | 0.00454182 | 0.236567569 |
| *FLOT1* | 0.383436 | 0.13521 | 2.835859 | 0.00457026 | 0.236567569 |
| *ZNF581* | -0.32674 | 0.115232 | -2.835499 | 0.00457541 | 0.236567569 |
| *SLC44A2* | 0.315237 | 0.111185 | 2.835244 | 0.00457907 | 0.236567569 |
| *FGF9* | 2.99957 | 1.05902 | 2.832403 | 0.00461996 | NA |
| *MZB1* | -0.559387 | 0.197594 | -2.830985 | 0.00464049 | 0.238354846 |
| *APOL3* | 0.495284 | 0.175483 | 2.822402 | 0.00476655 | 0.240899862 |
| *EFCAB5* | 1.247835 | 0.442375 | 2.820763 | 0.00479096 | 0.240899862 |
| *SLC27A5* | -0.988445 | 0.350448 | -2.820516 | 0.00479465 | 0.240899862 |
| *HOXB3* | 2.793942 | 0.991072 | 2.819112 | 0.00481567 | NA |
| *ZNF780A* | 0.641343 | 0.227503 | 2.819056 | 0.00481651 | 0.240899862 |
| *SELL* | 0.58601 | 0.20799 | 2.817496 | 0.00483997 | 0.240899862 |
| *MYO9B* | 0.409979 | 0.145555 | 2.816653 | 0.0048527 | 0.240899862 |
| *EIF2AK4* | -0.783045 | 0.278371 | -2.812958 | 0.0049088 | 0.241298667 |
| *LINC02062* | 1.597238 | 0.567814 | 2.812957 | 0.00490882 | NA |
| *C7orf50* | -0.320533 | 0.114035 | -2.810838 | 0.00494127 | 0.241298667 |
| *POP7* | -0.534824 | 0.190298 | -2.810452 | 0.0049472 | 0.241298667 |
| *PSMC6* | 0.320808 | 0.114207 | 2.809014 | 0.00496935 | 0.241298667 |
| *STMN3* | 0.435974 | 0.155477 | 2.804099 | 0.00504574 | 0.243676164 |
| *TCIRG1* | -0.613782 | 0.219326 | -2.798488 | 0.00513425 | 0.246610727 |
| *DMD* | -0.635901 | 0.227852 | -2.790848 | 0.00525702 | 0.251149959 |
| *ENSG00000283013* | 0.401087 | 0.143879 | 2.787674 | 0.00530879 | 0.251525614 |
| *ENSG00000251364* | 1.165921 | 0.418594 | 2.785323 | 0.00534744 | NA |
| *KDM2B* | 0.450505 | 0.161748 | 2.78523 | 0.00534898 | 0.251525614 |
| *CAPRIN2* | -0.995901 | 0.357572 | -2.78518 | 0.0053498 | 0.251525614 |
| *USF3* | -0.440345 | 0.158205 | -2.783378 | 0.00537961 | 0.251595712 |
| *UBE2E1* | 0.379504 | 0.136444 | 2.781383 | 0.00541279 | 0.251822086 |
| *MCM7* | 0.465233 | 0.16746 | 2.77817 | 0.0054666 | 0.251874484 |
| *ENSG00000227486* | 0.413467 | 0.14884 | 2.777932 | 0.0054706 | 0.251874484 |
| *CCDC154* | 1.246366 | 0.449052 | 2.775546 | 0.00551092 | NA |
| *TMEM38B* | 0.566989 | 0.204503 | 2.772516 | 0.00556248 | 0.254784475 |
| *C1orf159* | -1.512527 | 0.545962 | -2.770386 | 0.00559899 | NA |
| *ENSG00000230709* | -1.846118 | 0.66723 | -2.766841 | 0.00566024 | NA |
| *ENSG00000259802* | 1.930093 | 0.701928 | 2.7497 | 0.00596498 | NA |
| *FCGRT* | 0.364299 | 0.132491 | 2.74961 | 0.00596663 | 0.271894637 |
| *TENT5C* | -0.415571 | 0.151488 | -2.743262 | 0.00608321 | 0.275792631 |
| *MAP3K1* | 0.423318 | 0.154406 | 2.741587 | 0.00611433 | 0.275796412 |
| *UBB* | 0.214956 | 0.078454 | 2.739879 | 0.00614619 | 0.275833405 |
| *GPR146* | -0.804629 | 0.294723 | -2.730122 | 0.00633109 | 0.281446068 |
| *LINC00926* | 0.220367 | 0.08073 | 2.7297 | 0.0063392 | 0.281446068 |
| *MOB3C* | 0.776791 | 0.284717 | 2.728295 | 0.00636627 | 0.281446068 |
| *HTR3A* | 1.272951 | 0.467018 | 2.725697 | 0.00641659 | NA |

| | | | | | |
|---|---|---|---|---|---|
| *TBC1D22B* | 0.830079 | 0.304612 | 2.725032 | 0.00642952 | 0.282835297 |
| *CYBA* | 0.230959 | 0.084831 | 2.722578 | 0.00647747 | 0.283541055 |
| *TLE3* | -0.411522 | 0.15138 | -2.718459 | 0.00655868 | 0.285688442 |
| *PAXIP1.AS2* | 0.676074 | 0.250911 | 2.694474 | 0.00704997 | 0.305590517 |
| *GTPBP8* | -0.778239 | 0.289022 | -2.692662 | 0.00708842 | 0.30576543 |
| *EPHB6* | -0.719319 | 0.267679 | -2.687248 | 0.00720435 | 0.309264875 |
| *CLDND2* | 1.011259 | 0.377235 | 2.68071 | 0.00734661 | 0.313855824 |
| *SLC38A2* | -0.399758 | 0.150721 | -2.652309 | 0.00799434 | 0.338718053 |
| *PCBP4* | -1.321227 | 0.498239 | -2.651795 | 0.0080065 | 0.338718053 |
| *HLA.DMB* | 0.25139 | 0.094872 | 2.649788 | 0.00805423 | 0.338718053 |
| *ENSG00000224505* | 0.426816 | 0.161235 | 2.647171 | 0.00811682 | 0.338718053 |
| *PLP2* | -0.491261 | 0.18562 | -2.646596 | 0.00813065 | 0.338718053 |
| *LGALS1* | -0.970207 | 0.366836 | -2.644796 | 0.00817402 | 0.338718053 |
| *ABI3* | 0.47154 | 0.178349 | 2.643912 | 0.00819541 | 0.338718053 |
| *MEF2A* | 0.453422 | 0.171716 | 2.640542 | 0.00827736 | 0.340521563 |
| *KCTD21.AS1* | 1.120297 | 0.424467 | 2.639303 | 0.00830768 | NA |
| *SPIB* | 0.293631 | 0.111291 | 2.638402 | 0.00832978 | 0.341098651 |
| *RUNX2* | 3.85235 | 1.462854 | 2.633447 | 0.0084523 | NA |
| *ZNF419* | 0.614211 | 0.233534 | 2.630072 | 0.00853669 | 0.34612604 |
| *KEAP1* | 0.975044 | 0.370889 | 2.628941 | 0.00856513 | 0.34612604 |
| *FTH1* | -0.129098 | 0.049147 | -2.626785 | 0.00861958 | 0.34612604 |
| *FCRL2* | 0.52605 | 0.200312 | 2.626155 | 0.00863556 | 0.34612604 |
| *DNMT3A* | 0.854599 | 0.325598 | 2.624706 | 0.00867238 | 0.34612604 |
| *GYPC* | 0.251402 | 0.095844 | 2.623036 | 0.00871501 | 0.34612604 |
| *PGK1* | 0.326832 | 0.12462 | 2.622637 | 0.00872521 | 0.34612604 |
| *PLXNB2* | 1.530281 | 0.584206 | 2.619419 | 0.00880796 | NA |
| *NIPAL4* | 1.428364 | 0.545399 | 2.618933 | 0.00882052 | NA |
| *VHL* | 0.311016 | 0.118799 | 2.618009 | 0.00884444 | 0.346257513 |
| *IKZF1* | 0.372212 | 0.142221 | 2.617147 | 0.00886681 | 0.346257513 |
| *SNX2* | 0.22646 | 0.086557 | 2.616324 | 0.00888822 | 0.346257513 |
| *GABPB1* | 0.250804 | 0.095882 | 2.615766 | 0.00890275 | 0.346257513 |
| *PHF6* | -0.606867 | 0.232193 | -2.613631 | 0.00895858 | 0.346257513 |
| *GIT2* | 0.287888 | 0.110155 | 2.613488 | 0.00896233 | 0.346257513 |
| *HLA.DMA* | 0.248027 | 0.094997 | 2.61089 | 0.00903068 | 0.347388027 |
| *SHISAL2A* | 0.302659 | 0.11619 | 2.604852 | 0.00919141 | 0.350651463 |
| *MIEF2* | 0.738812 | 0.28375 | 2.603739 | 0.00922131 | 0.350651463 |
| *PRDM4* | 0.684546 | 0.262956 | 2.603271 | 0.0092339 | 0.350651463 |
| *LRPAP1* | 0.317244 | 0.121959 | 2.601228 | 0.00928906 | 0.351245008 |
| *ARID4B* | 0.257501 | 0.099055 | 2.599575 | 0.00933394 | 0.351446627 |
| *PUS1* | 0.649057 | 0.250024 | 2.595983 | 0.00943207 | 0.352762648 |
| *LSP1* | 0.248013 | 0.095611 | 2.593992 | 0.00948688 | 0.352762648 |
| *PPP1R18* | 0.343389 | 0.132447 | 2.592651 | 0.00952395 | 0.352762648 |
| *ENSG00000089127* | 0.493801 | 0.190487 | 2.592306 | 0.0095335 | 0.352762648 |
| *OS9* | 0.311454 | 0.120202 | 2.591085 | 0.00956739 | 0.352762648 |
| *CARS2* | 0.61878 | 0.239069 | 2.588287 | 0.00964544 | 0.352948343 |
| *YARS2* | -0.650445 | 0.251384 | -2.587455 | 0.00966878 | 0.352948343 |
| *NIFK* | 0.348849 | 0.134885 | 2.586274 | 0.00970198 | 0.352948343 |

| | | | | | |
|---|---|---|---|---|---|
| *GALK1* | -0.762558 | 0.295166 | -2.583489 | 0.00978067 | 0.352948343 |
| *WDR11* | 0.44031 | 0.170446 | 2.583281 | 0.00978657 | 0.352948343 |
| *SLC25A5* | -0.199945 | 0.077425 | -2.582429 | 0.00981074 | 0.352948343 |
| *KRAS* | -0.402493 | 0.156521 | -2.571499 | 0.01012593 | 0.361266309 |
| *AARS* | 0.586217 | 0.228026 | 2.57083 | 0.01014551 | 0.361266309 |
| *ZKSCAN4* | 1.012114 | 0.393788 | 2.570202 | 0.01016392 | 0.361266309 |
| *HSPA4L* | 1.33966 | 0.521281 | 2.569939 | 0.01017163 | NA |
| *ENSG00000271937* | 1.651796 | 0.642791 | 2.569724 | 0.01017796 | NA |
| *SMC6* | -0.337741 | 0.131612 | -2.566195 | 0.01028209 | 0.363816256 |
| *POU2AF1* | 0.279311 | 0.108928 | 2.564179 | 0.01034203 | 0.363816256 |
| *STAU2* | 0.486674 | 0.189838 | 2.563627 | 0.01035849 | 0.363816256 |
| *GPT2* | 0.843466 | 0.329474 | 2.560034 | 0.01046619 | 0.365202267 |
| *ZNF555* | 0.943131 | 0.368472 | 2.559571 | 0.01048015 | 0.365202267 |
| *ALKBH2* | 0.560683 | 0.219481 | 2.554581 | 0.01063158 | 0.369032017 |
| *C22orf15* | 1.738637 | 0.682307 | 2.548174 | 0.01082884 | NA |
| *CTNNA1* | -0.855451 | 0.336559 | -2.541753 | 0.01102981 | 0.379982132 |
| *DAPK3* | 0.539287 | 0.212179 | 2.541666 | 0.01103257 | 0.379982132 |
| *ILVBL* | 0.635741 | 0.250395 | 2.538947 | 0.01111868 | 0.3813939 |
| *SC5D* | -0.483658 | 0.190592 | -2.537667 | 0.0111594 | 0.3813939 |
| *CHL1* | 1.69234 | 0.667188 | 2.536526 | 0.01119583 | NA |
| *TRMT10C* | 0.366775 | 0.144801 | 2.53296 | 0.01131038 | 0.383660249 |
| *CENPS* | -0.995926 | 0.393195 | -2.532908 | 0.01131206 | 0.383660249 |
| *ZNF827* | 0.8948 | 0.353365 | 2.532224 | 0.01133414 | NA |
| *GHRL* | 0.621355 | 0.24549 | 2.531083 | 0.0113711 | 0.383967059 |
| *MMP11* | 0.734328 | 0.290305 | 2.529505 | 0.01142235 | 0.383967059 |
| *SHISA5* | 0.296411 | 0.117222 | 2.528634 | 0.01145074 | 0.383967059 |
| *ETV6* | -0.682143 | 0.270087 | -2.52564 | 0.01154879 | 0.385787423 |
| *WDR5B* | 0.679952 | 0.269359 | 2.524331 | 0.01159186 | 0.385787423 |
| *TBC1D9* | 0.596108 | 0.236281 | 2.522884 | 0.01163969 | 0.385933936 |
| *SH3KBP1* | -0.299272 | 0.118762 | -2.519926 | 0.01173796 | 0.386591191 |
| *ADAMTS7* | -1.210683 | 0.48107 | -2.516649 | 0.01184769 | 0.386591191 |
| *TECR* | 0.312332 | 0.124113 | 2.516511 | 0.01185232 | 0.386591191 |
| *CCDC22* | 0.551429 | 0.219126 | 2.51649 | 0.01185303 | 0.386591191 |
| *FXYD5* | -0.225537 | 0.089649 | -2.515777 | 0.01187704 | 0.386591191 |
| *ARL14EPL* | 3.348189 | 1.331915 | 2.513816 | 0.01194328 | NA |
| *SEMA4A* | 1.685778 | 0.671139 | 2.511815 | 0.01201122 | NA |
| *ISG20* | -0.276371 | 0.110052 | -2.511268 | 0.01202984 | 0.390135508 |
| *PTP4A3* | 1.526123 | 0.607924 | 2.510384 | 0.01205999 | NA |
| *ZNF202* | 0.874463 | 0.348662 | 2.508058 | 0.01213968 | 0.392266158 |
| *U2SURP* | 0.25854 | 0.103205 | 2.505101 | 0.01224165 | 0.394127967 |
| *TMEM263* | 0.490411 | 0.195942 | 2.502839 | 0.01232016 | 0.39522346 |
| *XPO4* | 0.501553 | 0.200749 | 2.49841 | 0.01247518 | 0.397390593 |
| *GRK5* | -0.541747 | 0.216888 | -2.497816 | 0.01249609 | 0.397390593 |
| *PDIA3* | 0.36076 | 0.144472 | 2.497086 | 0.01252187 | 0.397390593 |
| *PARP14* | -0.312246 | 0.125252 | -2.49293 | 0.01266939 | 0.400641302 |
| *PWP1* | 0.306952 | 0.123206 | 2.491378 | 0.01272488 | 0.40096913 |
| *EPN1* | 0.387759 | 0.155724 | 2.490033 | 0.01277314 | 0.401067418 |

| | | | | | |
|---|---|---|---|---|---|
| *CCDC57* | -0.43046 | 0.173047 | -2.487538 | 0.01286308 | 0.402469378 |
| *LINC00996* | 2.294011 | 0.923728 | 2.483427 | 0.01301251 | NA |
| *ENDOD1* | 0.874248 | 0.352338 | 2.48128 | 0.01309113 | 0.405752013 |
| *BANK1* | -0.259879 | 0.104744 | -2.481093 | 0.01309802 | 0.405752013 |
| *SLC12A2* | -0.729747 | 0.294146 | -2.480904 | 0.01310498 | 0.405752013 |
| *ANKAR* | 0.499334 | 0.201877 | 2.473453 | 0.01338144 | 0.411480079 |
| *ENSG00000236514* | 0.845073 | 0.341661 | 2.473422 | 0.0133826 | 0.411480079 |
| *C15orf62* | -1.088993 | 0.441134 | -2.468622 | 0.01356343 | 0.415519324 |
| *RASGEF1B* | -0.466615 | 0.189107 | -2.467461 | 0.01360749 | 0.415519324 |
| *ENSG00000282393* | 1.231029 | 0.499614 | 2.463958 | 0.01374121 | NA |
| *FUT2* | 1.801679 | 0.731345 | 2.463517 | 0.01375815 | NA |
| *PNMA1* | -1.047758 | 0.425494 | -2.462448 | 0.01379923 | 0.419808282 |
| *RBM7* | 0.339964 | 0.138122 | 2.461327 | 0.01384243 | 0.419808282 |
| *ZNF791* | 0.333792 | 0.135957 | 2.45513 | 0.01408336 | 0.423486902 |
| *C16orf74* | 0.372945 | 0.151927 | 2.45476 | 0.01409787 | 0.423486902 |
| *KLHDC7B* | 0.828158 | 0.337399 | 2.454535 | 0.0141067 | 0.423486902 |
| *INSIG1* | 0.316616 | 0.129323 | 2.44826 | 0.0143548 | 0.428431493 |
| *ENO1* | 0.261875 | 0.106978 | 2.447933 | 0.01436784 | 0.428431493 |
| *RNF34* | -0.489195 | 0.200076 | -2.445041 | 0.01448357 | 0.430438013 |
| *SLC16A1* | 1.073688 | 0.439156 | 2.444893 | 0.01448953 | NA |
| *ENSG00000281571* | 0.821669 | 0.33659 | 2.441156 | 0.01464031 | 0.433645975 |
| *MKNK2* | -0.360033 | 0.147647 | -2.438472 | 0.01474948 | 0.435428201 |
| *PIM2* | -0.27375 | 0.112431 | -2.434834 | 0.01489862 | 0.436711981 |
| *ABCC4* | 2.534362 | 1.040917 | 2.43474 | 0.01490248 | NA |
| *ENSG00000166927* | -0.889464 | 0.365376 | -2.434378 | 0.01491741 | 0.436711981 |
| *ANKZF1* | -0.350784 | 0.144129 | -2.43382 | 0.01494041 | 0.436711981 |
| *ENSG00000223881* | 1.150294 | 0.473363 | 2.430045 | 0.01509694 | NA |
| *PBRM1* | -0.474302 | 0.195232 | -2.429428 | 0.01512266 | 0.440572674 |
| *LZTR1* | -1.237756 | 0.509728 | -2.428266 | 0.01517121 | NA |
| *SNRNP35* | 0.391987 | 0.161428 | 2.428256 | 0.01517165 | 0.440572674 |
| *RFC3* | 0.797418 | 0.328658 | 2.426288 | 0.01525417 | 0.441526321 |
| *CALCOCO1* | 0.326282 | 0.134665 | 2.422918 | 0.01539638 | 0.44300078 |
| *CR2* | 0.810639 | 0.334599 | 2.422719 | 0.01540482 | 0.44300078 |
| *SF3A3* | 0.387419 | 0.16003 | 2.420917 | 0.01548141 | 0.443767048 |
| *FYTTD1* | -0.316992 | 0.131049 | -2.418886 | 0.01556814 | 0.444818261 |
| *IRF2BP2* | -0.560265 | 0.232293 | -2.411892 | 0.01586999 | 0.451989593 |
| *TACC1* | -0.506115 | 0.210328 | -2.406312 | 0.01611449 | 0.457045692 |
| *CDK13* | 0.420209 | 0.174687 | 2.405499 | 0.01615039 | 0.457045692 |
| *VTI1A* | -0.575207 | 0.239603 | -2.400669 | 0.01636514 | 0.459352878 |
| *SDHAF3* | -0.475127 | 0.197945 | -2.4003 | 0.01638165 | 0.459352878 |
| *LAMTOR5* | -0.178771 | 0.074485 | -2.400096 | 0.01639078 | 0.459352878 |
| *SLC45A3* | 0.840447 | 0.350328 | 2.399027 | 0.01643869 | 0.459352878 |
| *UTP23* | 0.413543 | 0.172722 | 2.39427 | 0.0166535 | 0.463896532 |
| *ABTB2* | 3.064802 | 1.282388 | 2.389918 | 0.01685212 | NA |
| *KANK1* | 1.314955 | 0.551143 | 2.38587 | 0.01703878 | NA |
| *MAMSTR* | 1.091971 | 0.457817 | 2.385171 | 0.01707119 | NA |
| *DNAJC17* | -0.645973 | 0.271057 | -2.383164 | 0.01716456 | 0.474861359 |

| | | | | | |
|---|---|---|---|---|---|
| *TRUB1* | 0.76824 | 0.322417 | 2.382757 | 0.01718353 | 0.474861359 |
| *VPS13A* | -0.595074 | 0.249984 | -2.380448 | 0.01729159 | 0.474861359 |
| *COMMD4* | -0.492049 | 0.206717 | -2.380299 | 0.01729861 | 0.474861359 |
| *TM7SF3* | 0.475231 | 0.199803 | 2.378495 | 0.01738347 | 0.474861359 |
| *MFNG* | 0.384512 | 0.161708 | 2.377807 | 0.01741594 | 0.474861359 |
| *SLC12A6* | 0.363736 | 0.152978 | 2.377696 | 0.0174212 | 0.474861359 |
| *PDE7A* | 0.260916 | 0.109863 | 2.374924 | 0.01755254 | 0.476978292 |
| *COPS7A* | -0.627235 | 0.264474 | -2.371633 | 0.01770969 | 0.479781396 |
| *DAP3* | -0.309069 | 0.130431 | -2.369595 | 0.0178076 | 0.480500582 |
| *SNX9* | 0.356089 | 0.150359 | 2.368261 | 0.0178719 | 0.480500582 |
| *ENSG00000255031* | -1.318319 | 0.557231 | -2.365837 | 0.01798936 | NA |
| *GLIPR2* | 0.450073 | 0.190281 | 2.365309 | 0.01801501 | 0.480500582 |
| *VIM* | -0.5173 | 0.218704 | -2.365292 | 0.01801585 | 0.480500582 |
| *CTSS* | 0.225162 | 0.095228 | 2.36444 | 0.01805735 | 0.480500582 |
| *LBR* | 0.319211 | 0.135009 | 2.364372 | 0.01806068 | 0.480500582 |
| *RNF207* | 1.732224 | 0.733224 | 2.362475 | 0.01815338 | NA |
| *B3GNTL1* | 1.125878 | 0.476798 | 2.361332 | 0.01820941 | NA |
| *AGBL5* | -1.236885 | 0.523809 | -2.361328 | 0.01820964 | NA |
| *HSD17B1* | 1.094302 | 0.464091 | 2.357948 | 0.01837628 | NA |
| *UBE2J2* | 0.284817 | 0.120793 | 2.357889 | 0.01837918 | 0.484061438 |
| *SESN3* | -0.330321 | 0.140097 | -2.357808 | 0.0183832 | 0.484061438 |
| *UBE2Z* | -0.52662 | 0.223446 | -2.35681 | 0.01843268 | 0.484061438 |
| *RAD50* | -0.349128 | 0.148164 | -2.356362 | 0.0184549 | 0.484061438 |
| *BTG2* | -0.245463 | 0.104181 | -2.356121 | 0.0184669 | 0.484061438 |
| *SATB1* | 0.389716 | 0.165505 | 2.354701 | 0.01853761 | 0.484485824 |
| *ENSG00000226571* | -1.905699 | 0.80968 | -2.353644 | 0.01859042 | NA |
| *SNAI3.AS1* | -1.507628 | 0.640973 | -2.352091 | 0.01866822 | NA |
| *ACTN1* | 2.129071 | 0.906536 | 2.34858 | 0.01884515 | NA |
| *SFT2D3* | 1.190923 | 0.507385 | 2.34718 | 0.01891614 | NA |
| *WASF2* | 0.217863 | 0.092941 | 2.344103 | 0.01907291 | 0.497014354 |
| *YIPF4* | 0.218657 | 0.093378 | 2.341631 | 0.01919971 | 0.49885565 |
| *CARD8.AS1* | 0.401919 | 0.172155 | 2.334637 | 0.0195624 | 0.506797218 |
| *C1orf50* | 0.469816 | 0.201335 | 2.333502 | 0.0196218 | 0.506858345 |
| *ENSG00000253106* | 1.769714 | 0.758832 | 2.332155 | 0.01969256 | NA |
| *COMMD6* | -0.154394 | 0.066234 | -2.331024 | 0.0197521 | 0.508745396 |
| *CD99* | -0.458725 | 0.196888 | -2.329877 | 0.01981263 | 0.508776722 |
| *CYTH4* | -0.396739 | 0.17036 | -2.328835 | 0.01986783 | 0.508776722 |
| *R3HDM1* | -0.557268 | 0.239509 | -2.326707 | 0.01998087 | 0.510201118 |
| *ARL2BP* | -0.328278 | 0.141163 | -2.325523 | 0.02004401 | 0.510346966 |
| *MORC2* | -1.425361 | 0.613413 | -2.323655 | 0.02014398 | NA |
| *RNF215* | 1.241696 | 0.534851 | 2.321574 | 0.02025589 | NA |
| *PFDN6* | 0.275056 | 0.11848 | 2.321535 | 0.02025799 | 0.514321462 |
| *ENSG00000272871* | -1.733007 | 0.74684 | -2.320452 | 0.02031644 | NA |
| *ENSG00000238198* | 1.713099 | 0.738266 | 2.320437 | 0.02031724 | NA |
| *ISYNA1* | 0.707331 | 0.305038 | 2.31883 | 0.02040425 | 0.515457894 |
| *GDPGP1* | 0.967447 | 0.417261 | 2.318563 | 0.02041877 | 0.515457894 |
| *CKLF* | -0.466515 | 0.201355 | -2.316871 | 0.02051075 | 0.516313061 |

| | | | | | |
|---|---|---|---|---|---|
| *HNRNPK* | 0.172969 | 0.074706 | 2.315321 | 0.02059535 | 0.516978257 |
| *NUMBL* | -1.128584 | 0.487522 | -2.314942 | 0.02061612 | NA |
| *CD63* | -0.41276 | 0.178563 | -2.311572 | 0.02080125 | 0.520675889 |
| *HSD3B7* | 2.214864 | 0.959043 | 2.309452 | 0.0209185 | NA |
| *CLIC1* | -0.212251 | 0.091974 | -2.307718 | 0.0210148 | 0.52154266 |
| *TOX2* | 1.860684 | 0.806374 | 2.307472 | 0.02102854 | NA |
| *STK17B* | -0.275227 | 0.119295 | -2.307121 | 0.02104807 | 0.52154266 |
| *MRNIP* | 0.31072 | 0.134709 | 2.306602 | 0.02107701 | 0.52154266 |
| *CD58* | -0.719132 | 0.311901 | -2.30564 | 0.02113074 | 0.52154266 |
| *UNC79* | 1.805713 | 0.783373 | 2.305048 | 0.02116389 | NA |
| *ORMDL2* | -0.36344 | 0.157685 | -2.304841 | 0.0211755 | 0.52154266 |
| *WDPCP* | 0.725749 | 0.314996 | 2.30399 | 0.02122318 | 0.52154266 |
| *HNRNPH2* | 0.300727 | 0.130548 | 2.303571 | 0.02124673 | 0.52154266 |
| *EBNA1BP2* | -0.493921 | 0.214585 | -2.301748 | 0.02134941 | 0.522619356 |
| *C12orf42* | -0.46036 | 0.200278 | -2.29861 | 0.02152708 | 0.525520868 |
| *ENSG00000276476* | 1.625532 | 0.707445 | 2.297749 | 0.02157609 | NA |
| *ENSG00000248559* | 0.795527 | 0.34655 | 2.295563 | 0.0217009 | 0.528312905 |
| *TNNI2* | 1.007175 | 0.439022 | 2.294135 | 0.02178275 | NA |
| *ENSG00000279278* | 0.610151 | 0.265987 | 2.293914 | 0.02179541 | 0.529164062 |
| *CD3D* | 2.458991 | 1.072589 | 2.292577 | 0.02187239 | NA |
| *RAB5IF* | -0.286447 | 0.125073 | -2.290244 | 0.0220072 | 0.532323324 |
| *EIF2S3* | -0.238107 | 0.104018 | -2.289094 | 0.02207386 | 0.532323324 |
| *RASA3* | 0.65652 | 0.286871 | 2.288554 | 0.02210526 | 0.532323324 |
| *DLGAP1.AS1* | -0.319341 | 0.139622 | -2.287178 | 0.02218541 | 0.532809697 |
| *PAG1* | -1.016597 | 0.44471 | -2.28598 | 0.02225542 | 0.533050306 |
| *SLC26A2* | -0.813646 | 0.356499 | -2.282322 | 0.02247035 | 0.534298316 |
| *HIP1* | -0.998881 | 0.437696 | -2.282133 | 0.02248147 | 0.534298316 |
| *SLCO3A1* | 3.093415 | 1.356913 | 2.279745 | 0.02262284 | NA |
| *ZNF821* | 0.404687 | 0.177604 | 2.278596 | 0.02269107 | 0.534298316 |
| *CD200R1* | 3.360459 | 1.475364 | 2.277715 | 0.02274357 | NA |
| *AP1G2* | -0.291607 | 0.12806 | -2.277105 | 0.02277997 | 0.534298316 |
| *ULK1* | 0.77476 | 0.340262 | 2.276954 | 0.02278895 | 0.534298316 |
| *KIF2A* | 0.321561 | 0.141232 | 2.276829 | 0.02279647 | 0.534298316 |
| *PIK3IP1* | 0.28821 | 0.126607 | 2.276405 | 0.02282177 | 0.534298316 |
| *ECHS1* | 0.320503 | 0.140809 | 2.276159 | 0.02283652 | 0.534298316 |
| *KATNB1* | 0.7838 | 0.344445 | 2.275544 | 0.02287331 | 0.534298316 |
| *GBP7* | 0.897925 | 0.394701 | 2.274952 | 0.02290881 | 0.534298316 |
| *ENSG00000251136* | 0.670324 | 0.294802 | 2.273814 | 0.02297718 | 0.5344901 |
| *AP3S1* | 0.316169 | 0.13926 | 2.270346 | 0.02318663 | 0.534870118 |
| *LMCD1* | 2.773108 | 1.221566 | 2.270126 | 0.02319997 | NA |
| *RRM2B* | -0.478278 | 0.210752 | -2.269392 | 0.02324449 | 0.534870118 |
| *RPA3* | 0.199451 | 0.087898 | 2.269132 | 0.02326032 | 0.534870118 |
| *TMUB2* | 0.314655 | 0.138734 | 2.268041 | 0.02332673 | 0.534870118 |
| *NOTCH1* | 0.780018 | 0.34404 | 2.267233 | 0.02337598 | 0.534870118 |
| *UBE2V1* | 0.574914 | 0.253757 | 2.265613 | 0.0234751 | 0.534870118 |
| *DTX3L* | 0.470538 | 0.207704 | 2.265426 | 0.02348658 | 0.534870118 |
| *S1PR4* | 0.285693 | 0.126111 | 2.265404 | 0.02348789 | 0.534870118 |

| | | | | | |
|---|---|---|---|---|---|
| *KHNYN* | -0.554106 | 0.244678 | -2.264632 | 0.02353525 | 0.534870118 |
| *ENSG00000272583* | -1.173289 | 0.519029 | -2.260545 | 0.02378744 | NA |
| *OTUD7B* | 2.654478 | 1.175232 | 2.258683 | 0.02390309 | NA |
| *CISH* | 1.833614 | 0.812138 | 2.257763 | 0.02396047 | NA |
| *RAP2C.AS1* | -1.824414 | 0.808291 | -2.257126 | 0.02400022 | NA |
| *TMCO4* | 0.708119 | 0.313902 | 2.255861 | 0.02407935 | 0.545839615 |
| *SLC9A6* | -1.524543 | 0.676251 | -2.254405 | 0.02417068 | NA |
| *MKLN1* | -0.48689 | 0.215977 | -2.254365 | 0.02417324 | 0.546573618 |
| *DNASE2* | 0.333829 | 0.148255 | 2.251725 | 0.02433965 | 0.548939469 |
| *ABTB1* | 0.252579 | 0.112379 | 2.247574 | 0.02460336 | 0.553482167 |
| *RORA* | 1.586803 | 0.707419 | 2.243089 | 0.0248911 | NA |
| *UTP4* | 0.572378 | 0.255341 | 2.241619 | 0.02498599 | 0.560125051 |
| *SETBP1* | 0.587245 | 0.262043 | 2.241021 | 0.02502472 | 0.560125051 |
| *SLCO5A1* | 1.513022 | 0.675188 | 2.240891 | 0.02503316 | NA |
| *FAM135A* | -0.689457 | 0.307928 | -2.239021 | 0.02515457 | 0.561616927 |
| *UEVLD* | -1.240096 | 0.554651 | -2.235813 | 0.02536403 | NA |
| *MCCC1* | 0.48929 | 0.219065 | 2.233534 | 0.02551372 | 0.565745108 |
| *LPCAT1* | -0.795328 | 0.356116 | -2.233338 | 0.02552666 | 0.565745108 |
| *CTBP2* | 2.965935 | 1.328123 | 2.233178 | 0.02553722 | NA |
| *ARIH2* | 0.312235 | 0.139853 | 2.232593 | 0.02557578 | 0.565745108 |
| *ENSG00000268027* | 0.405466 | 0.181635 | 2.232315 | 0.02559414 | 0.565745108 |
| *TAF4B* | 0.724828 | 0.325841 | 2.224486 | 0.02611579 | 0.575843346 |
| *CIC* | 0.508508 | 0.228983 | 2.220722 | 0.02636982 | 0.579738079 |
| *CCL4* | 1.239672 | 0.558426 | 2.219939 | 0.0264229 | 0.579738079 |
| *ENSG00000267152* | 0.955316 | 0.430573 | 2.218708 | 0.02650658 | NA |
| *WDR27* | -1.180727 | 0.532244 | -2.218394 | 0.02652798 | NA |
| *ZNF710* | -1.567938 | 0.707382 | -2.216536 | 0.0266548 | NA |
| *SMIM20* | 0.399323 | 0.180218 | 2.21577 | 0.02670726 | 0.584533698 |
| *ENSG00000268798* | 0.963653 | 0.435278 | 2.213879 | 0.02683712 | NA |
| *DPY19L1* | -1.667804 | 0.753592 | -2.213139 | 0.02688804 | NA |
| *ENSG00000259001* | 1.053146 | 0.476207 | 2.21153 | 0.02699914 | NA |
| *CENPT* | -0.403481 | 0.182463 | -2.211302 | 0.02701497 | 0.589815673 |
| *POLB* | -0.834733 | 0.377787 | -2.209532 | 0.02713768 | 0.590227874 |
| *CTNNBL1* | 0.319472 | 0.144615 | 2.209114 | 0.02716669 | 0.590227874 |
| *ALKBH1* | 0.729601 | 0.330544 | 2.207272 | 0.02729507 | 0.591570737 |
| *REC8* | 0.592677 | 0.269173 | 2.201846 | 0.02767619 | 0.598371272 |
| *ZNF525* | -0.965896 | 0.438899 | -2.200723 | 0.02775563 | 0.598632301 |
| *HMGB3* | -1.679684 | 0.76378 | -2.199173 | 0.02786561 | NA |
| *TPRN* | -1.564819 | 0.711589 | -2.19905 | 0.02787436 | NA |
| *ARHGAP4* | 0.296491 | 0.134832 | 2.198962 | 0.0278806 | 0.598962369 |
| *C1orf122* | -0.437903 | 0.199182 | -2.1985 | 0.0279135 | 0.598962369 |
| *JUN* | 0.237378 | 0.108014 | 2.197663 | 0.02797315 | 0.598962369 |
| *ENSG00000272909* | 1.602725 | 0.72986 | 2.195934 | 0.02809668 | NA |
| *DMAC1* | 0.252192 | 0.114888 | 2.195105 | 0.0281561 | 0.60143044 |
| *RUBCNL* | 0.205292 | 0.093682 | 2.19138 | 0.02842431 | 0.603162241 |
| *PDE4DIP* | -0.558289 | 0.254824 | -2.190883 | 0.02846023 | 0.603162241 |
| *SERHL2* | -1.37776 | 0.628906 | -2.190725 | 0.0284717 | NA |

| | | | | | |
|---|---|---|---|---|---|
| *TMEM184C* | -0.85459 | 0.390106 | -2.190661 | 0.02847632 | 0.603162241 |
| *AATF* | 0.316719 | 0.144746 | 2.188097 | 0.02866253 | 0.603162241 |
| *CYB5R4* | 0.28147 | 0.128692 | 2.187152 | 0.02873142 | 0.603162241 |
| *KATNBL1* | -0.326255 | 0.149196 | -2.186757 | 0.02876029 | 0.603162241 |
| *SLC25A32* | 0.459155 | 0.210017 | 2.186279 | 0.02879518 | 0.603162241 |
| *TBL1X* | -0.589699 | 0.269739 | -2.186182 | 0.0288023 | 0.603162241 |
| *LINC02576* | -0.95578 | 0.437575 | -2.184265 | 0.02894281 | 0.603162241 |
| *ZNF699* | 0.560028 | 0.256438 | 2.183869 | 0.02897188 | 0.603162241 |
| *CDC42BPB* | -1.004036 | 0.459815 | -2.183564 | 0.02899427 | NA |
| *TET3* | 0.787919 | 0.360953 | 2.182887 | 0.02904412 | 0.603162241 |
| *HMGB2* | 0.214939 | 0.09847 | 2.182784 | 0.0290517 | 0.603162241 |
| *ENSG00000250541* | 1.753766 | 0.803642 | 2.182272 | 0.02908943 | NA |
| *HARBI1* | 0.930962 | 0.426692 | 2.181812 | 0.02912339 | NA |
| *NFXL1* | -1.030394 | 0.472962 | -2.178598 | 0.02936152 | NA |
| *SNAPC3* | 0.34851 | 0.159996 | 2.178245 | 0.02938777 | 0.608717311 |
| *COMMD7* | -0.463307 | 0.212828 | -2.176909 | 0.02948735 | 0.609359443 |
| *SLC41A1* | 0.660721 | 0.303913 | 2.174048 | 0.02970151 | 0.612361032 |
| *IL4R* | 0.384996 | 0.177303 | 2.171401 | 0.02990091 | 0.614752003 |
| *PPP2R5C* | 0.230384 | 0.106162 | 2.170116 | 0.02999806 | 0.614752003 |
| *PLCL2* | -0.606107 | 0.279343 | -2.16976 | 0.03002503 | 0.614752003 |
| *ACVR2A* | -1.664037 | 0.767425 | -2.168339 | 0.03013287 | NA |
| *SPN* | 1.081333 | 0.499704 | 2.163947 | 0.03046839 | NA |
| *KLF2* | 0.422931 | 0.19572 | 2.160895 | 0.03070348 | 0.62668642 |
| *ITGB2* | 0.751286 | 0.347768 | 2.160306 | 0.03074896 | 0.62668642 |
| *CENPJ* | -0.800443 | 0.370744 | -2.15902 | 0.03084865 | 0.627279318 |
| *PTP4A2* | -0.296579 | 0.137505 | -2.156863 | 0.0310163 | 0.629248561 |
| *NRDE2* | 0.352676 | 0.163947 | 2.151161 | 0.03146351 | 0.635433559 |
| *LYPLAL1* | 0.333166 | 0.154878 | 2.151152 | 0.03146419 | 0.635433559 |
| *ZFP36L2* | -0.444329 | 0.206719 | -2.149433 | 0.0316001 | 0.636731185 |
| *CTNND1* | -0.491109 | 0.228671 | -2.147669 | 0.03174006 | 0.63692922 |
| *ENSG00000255046* | 0.730054 | 0.340087 | 2.146666 | 0.03181991 | 0.63692922 |
| *PDLIM2* | -0.492249 | 0.229499 | -2.144885 | 0.03196205 | 0.63692922 |
| *PTPA* | -0.740542 | 0.345329 | -2.144456 | 0.03199634 | 0.63692922 |
| *ATG13* | -0.586446 | 0.273488 | -2.14432 | 0.03200724 | 0.63692922 |
| *DHTKD1* | -0.398354 | 0.185856 | -2.143344 | 0.03208549 | 0.63692922 |
| *ZNF844* | 0.543806 | 0.253757 | 2.143018 | 0.03211167 | 0.63692922 |
| *UBL4A* | -0.423356 | 0.197802 | -2.140305 | 0.03233012 | 0.639330234 |
| *AMPD3* | -0.598599 | 0.279755 | -2.13973 | 0.03237662 | 0.639330234 |
| *SH2D4A* | 1.637538 | 0.765504 | 2.139164 | 0.03242239 | NA |
| *ENSG00000274605* | 0.880701 | 0.411777 | 2.138782 | 0.03245335 | NA |
| *ENSG00000268093* | 1.472849 | 0.690128 | 2.134167 | 0.03282908 | NA |
| *TMEM222* | -0.413552 | 0.19393 | -2.132475 | 0.03296785 | 0.648235682 |
| *DDX20* | 0.466635 | 0.218892 | 2.131806 | 0.03302278 | 0.648235682 |
| *DIS3L* | 0.491174 | 0.230563 | 2.130325 | 0.03314479 | 0.648235682 |
| *ARRDC2* | -0.314701 | 0.147746 | -2.13001 | 0.03317078 | 0.648235682 |
| *ENSG00000272009* | 0.367123 | 0.172379 | 2.129749 | 0.03319235 | 0.648235682 |
| *ENSG00000278831* | 1.498052 | 0.703585 | 2.129169 | 0.03324025 | NA |

| | | | | | |
|---|---|---|---|---|---|
| *SGK1* | -1.235664 | 0.580868 | -2.127272 | 0.03339747 | 0.650178266 |
| *SH3BGRL3* | -0.165652 | 0.077889 | -2.126783 | 0.03343816 | 0.650178266 |
| *POMP* | 0.164002 | 0.077152 | 2.1257 | 0.03352823 | 0.65050618 |
| *COX16* | 0.433455 | 0.204145 | 2.123272 | 0.03373107 | 0.651622013 |
| *USP12* | -0.807625 | 0.380424 | -2.12296 | 0.03375721 | 0.651622013 |
| *GPSM1* | 1.321748 | 0.622876 | 2.122007 | 0.03383715 | NA |
| *RNF149* | -0.412711 | 0.194511 | -2.121789 | 0.03385545 | 0.651622013 |
| *BCAS4* | 0.319505 | 0.150603 | 2.121508 | 0.03387907 | 0.651622013 |
| *LINC00894* | 1.614172 | 0.76122 | 2.120508 | 0.03396322 | NA |
| *TRANK1* | -0.386252 | 0.18226 | -2.119237 | 0.03407047 | 0.653004745 |
| *PEAK1* | 0.604203 | 0.285148 | 2.118912 | 0.03409793 | 0.653004745 |
| *TBC1D19* | -1.108959 | 0.523369 | -2.118888 | 0.03409993 | NA |
| *TMEM159* | -0.523949 | 0.247604 | -2.11608 | 0.03433799 | 0.656187838 |
| *CDK12* | -0.346851 | 0.16412 | -2.113395 | 0.03456694 | 0.658744737 |
| *TFB2M* | 0.596375 | 0.282363 | 2.112083 | 0.03467933 | 0.658744737 |
| *KIAA1586* | 0.56627 | 0.268132 | 2.11191 | 0.03469419 | 0.658744737 |
| *SUMO4* | 0.965728 | 0.457332 | 2.111655 | 0.03471608 | NA |
| *LILRA2* | 1.687359 | 0.800287 | 2.108441 | 0.03499288 | NA |
| *HCK* | -0.586759 | 0.278342 | -2.108049 | 0.03502675 | 0.663533123 |
| *LRRC42* | -1.015187 | 0.481605 | -2.107924 | 0.03503753 | NA |
| *PATL2* | 0.401757 | 0.190654 | 2.107252 | 0.03509572 | 0.663533123 |
| *BTBD3* | -1.427998 | 0.678323 | -2.105188 | 0.03527496 | NA |
| *TRADD* | -0.334564 | 0.158982 | -2.104421 | 0.03534174 | 0.666765883 |
| *STK38* | 0.448444 | 0.213441 | 2.101026 | 0.03563868 | 0.668143221 |
| *EVL* | 0.24623 | 0.11728 | 2.099503 | 0.0357726 | 0.668143221 |
| *WDFY2* | -0.407922 | 0.194384 | -2.098535 | 0.03585792 | 0.668143221 |
| *TMC8* | 0.318986 | 0.152016 | 2.098365 | 0.03587291 | 0.668143221 |
| *ANKH* | -0.795206 | 0.378967 | -2.098351 | 0.03587411 | 0.668143221 |
| *NSMCE1* | 0.251369 | 0.119824 | 2.097814 | 0.03592155 | 0.668143221 |
| *SCPEP1* | 0.448259 | 0.213702 | 2.097593 | 0.03594108 | 0.668143221 |
| *SREBF1* | 0.859335 | 0.41019 | 2.09497 | 0.03617365 | NA |
| *SEC61B* | -0.26419 | 0.12612 | -2.094751 | 0.03619312 | 0.670889638 |
| *AP2S1* | -0.25512 | 0.121823 | -2.09419 | 0.03624309 | 0.670889638 |
| *SELENOW* | -0.29627 | 0.141527 | -2.093379 | 0.03631532 | 0.670889638 |
| *COQ8A* | 0.428399 | 0.204806 | 2.091724 | 0.03646323 | 0.6722246 |
| *ABRAXAS2* | 0.460359 | 0.220228 | 2.090372 | 0.03658443 | 0.673027589 |
| *CD81* | 0.359844 | 0.172211 | 2.08955 | 0.03665827 | 0.673027589 |
| *MPP7* | -1.445318 | 0.691909 | -2.088885 | 0.03671807 | NA |
| *PRDM2* | -0.231193 | 0.110774 | -2.087069 | 0.0368819 | 0.675737256 |
| *ECE2* | -1.427793 | 0.684793 | -2.085 | 0.03706934 | NA |
| *AUH* | 0.545157 | 0.261486 | 2.084841 | 0.03708373 | 0.676106133 |
| *DYNC1I2* | -0.233778 | 0.112173 | -2.084076 | 0.03715328 | 0.676106133 |
| *RHOG* | -0.279282 | 0.134011 | -2.08402 | 0.03715836 | 0.676106133 |
| *TSC22D1* | -0.719538 | 0.345392 | -2.08325 | 0.03722841 | 0.676106133 |
| *TMEM156* | 0.329436 | 0.15818 | 2.082657 | 0.03728247 | 0.676106133 |
| *CCR7* | 0.177221 | 0.085137 | 2.081595 | 0.03737947 | 0.676484632 |
| *TTC3* | -0.188894 | 0.090834 | -2.079562 | 0.03756569 | 0.677724336 |

| | | | | | |
|---|---|---|---|---|---|
| *WDR81* | -0.718024 | 0.34534 | -2.079183 | 0.03760051 | 0.677724336 |
| *ENSG00000249141* | 1.563952 | 0.752657 | 2.077909 | 0.03771772 | NA |
| *SCML4* | 0.963599 | 0.464079 | 2.076369 | 0.03785985 | NA |
| *ZNF211* | -0.609282 | 0.293464 | -2.076169 | 0.0378783 | 0.680096389 |
| *EIF4B* | -0.147729 | 0.071157 | -2.076095 | 0.03788518 | 0.680096389 |
| *CCNDBP1* | 0.170529 | 0.082196 | 2.074655 | 0.03801849 | 0.681113496 |
| *ZMAT2* | 0.201125 | 0.097027 | 2.072878 | 0.03818366 | 0.68269622 |
| *ENSG00000260261* | -1.152807 | 0.556179 | -2.072728 | 0.0381976 | NA |
| *CD68* | -1.144494 | 0.552323 | -2.072145 | 0.03825194 | NA |
| *ZNF682* | 0.676604 | 0.326737 | 2.070794 | 0.03837808 | 0.684497804 |
| *LINC01991* | -1.698018 | 0.820375 | -2.069807 | 0.03847046 | NA |
| *ENSG00000251661* | -1.580089 | 0.763529 | -2.069456 | 0.03850332 | NA |
| *RHEX* | -0.77452 | 0.374299 | -2.069255 | 0.03852214 | 0.684497804 |
| *UBE2G1* | 0.347468 | 0.167977 | 2.068539 | 0.03858938 | 0.684497804 |
| *UBTF* | 0.310753 | 0.150257 | 2.068142 | 0.03862671 | 0.684497804 |
| *LUC7L* | -0.399969 | 0.193504 | -2.06698 | 0.03873603 | 0.684497804 |
| *ZFYVE27* | 0.346623 | 0.16772 | 2.06668 | 0.03876435 | 0.684497804 |
| *GUCD1* | 0.281474 | 0.136279 | 2.065428 | 0.03888256 | 0.684497804 |
| *CYTIP* | -0.151744 | 0.073475 | -2.065236 | 0.03890067 | 0.684497804 |
| *TAX1BP3* | -0.425046 | 0.205982 | -2.063513 | 0.03906393 | 0.686012029 |
| *NEB* | 1.265365 | 0.61321 | 2.063511 | 0.03906412 | NA |
| *ENSG00000257497* | 1.088646 | 0.528107 | 2.061413 | 0.03926368 | NA |
| *GABARAPL2* | 0.175382 | 0.085088 | 2.061185 | 0.03928537 | 0.687500692 |
| *ENSG00000272112* | -1.577666 | 0.765724 | -2.06036 | 0.03936414 | NA |
| *COPA* | -0.350062 | 0.169914 | -2.060232 | 0.03937633 | 0.687500692 |
| *PTPRE* | -0.643914 | 0.312552 | -2.060186 | 0.03938081 | 0.687500692 |
| *LINC00158* | 2.985874 | 1.450108 | 2.05907 | 0.0394875 | NA |
| *SOCS7* | -1.294861 | 0.629251 | -2.057781 | 0.03961118 | NA |
| *ASCC1* | 0.511005 | 0.248339 | 2.057695 | 0.0396194 | 0.689415317 |
| *NCBP2* | -0.253837 | 0.123376 | -2.057422 | 0.03964565 | 0.689415317 |
| *LMNA* | -1.7363 | 0.843935 | -2.057385 | 0.03964919 | NA |
| *ENSG00000278730* | -1.003273 | 0.488139 | -2.0553 | 0.03985007 | NA |
| *SLK* | -0.657208 | 0.319929 | -2.054231 | 0.03995332 | 0.692434773 |
| *TFB1M* | 0.437085 | 0.212797 | 2.054006 | 0.03997513 | 0.692434773 |
| *GLT8D1* | 0.445273 | 0.217038 | 2.051596 | 0.04020899 | 0.693979782 |
| *FUOM* | 0.541198 | 0.263809 | 2.051477 | 0.04022053 | 0.693979782 |
| *PALLD* | 2.711564 | 1.322488 | 2.05035 | 0.04033025 | NA |
| *DNAJC11* | 0.616504 | 0.300774 | 2.049721 | 0.04039163 | 0.694060698 |
| *PLEKHB1* | 0.877162 | 0.427969 | 2.04959 | 0.04040441 | 0.694060698 |
| *FAM89A* | 1.223155 | 0.596973 | 2.04893 | 0.04046895 | NA |
| *BRK1* | 0.16727 | 0.081645 | 2.04875 | 0.04048653 | 0.694060698 |
| *ENAH* | 1.864233 | 0.91014 | 2.048293 | 0.04053129 | NA |
| *IRF8* | 0.261305 | 0.127593 | 2.047953 | 0.04056461 | 0.694060698 |
| *TBKBP1* | 1.764625 | 0.861863 | 2.047454 | 0.04061352 | NA |
| *BIN2* | 0.511444 | 0.24998 | 2.045937 | 0.04076256 | 0.694060698 |
| *FALEC* | 0.523721 | 0.256008 | 2.045721 | 0.04078388 | 0.694060698 |
| *MYL2* | 2.370648 | 1.159078 | 2.045288 | 0.0408265 | NA |

| | | | | | |
|---|---|---|---|---|---|
| *ZNF430* | 0.254532 | 0.124458 | 2.045125 | 0.0408425 | 0.694060698 |
| *H2AFY* | 0.298912 | 0.146164 | 2.045049 | 0.04085007 | 0.694060698 |
| *AGAP3* | 1.787074 | 0.873999 | 2.04471 | 0.04088345 | NA |
| *QPRT* | 2.141693 | 1.047514 | 2.044549 | 0.04089939 | NA |
| *MAN2A2* | -0.93186 | 0.456006 | -2.043523 | 0.04100067 | 0.695029737 |
| *PLEKHB2* | 0.326656 | 0.159899 | 2.042888 | 0.04106354 | 0.695029737 |
| *CPEB4* | 0.505881 | 0.247961 | 2.04016 | 0.04133438 | 0.698187465 |
| *RAB13* | -0.871541 | 0.427346 | -2.039429 | 0.04140725 | 0.698187465 |
| *SLC5A5* | 2.477055 | 1.215873 | 2.037265 | 0.04162351 | NA |
| *ATPSCKMT* | 0.875822 | 0.430469 | 2.034576 | 0.04189359 | NA |
| *NIN* | -0.345018 | 0.169578 | -2.034567 | 0.0418945 | 0.70399374 |
| *MOB1B* | -1.225211 | 0.602497 | -2.033555 | 0.04199648 | NA |
| *TRBC2* | -0.148396 | 0.07303 | -2.031988 | 0.04215485 | 0.70399374 |
| *PLOD1* | -1.537878 | 0.756964 | -2.03164 | 0.04219009 | NA |
| *PORCN* | 0.697619 | 0.343496 | 2.030936 | 0.04226151 | 0.70399374 |
| *ICA1L* | 0.376887 | 0.185596 | 2.03068 | 0.04228746 | 0.70399374 |
| *GTPBP10* | 0.465368 | 0.229186 | 2.030523 | 0.04230341 | 0.70399374 |
| *MDM4* | -0.155543 | 0.076607 | -2.030413 | 0.04231459 | 0.70399374 |
| *COL9A2* | -1.549929 | 0.763537 | -2.029935 | 0.04236318 | NA |
| *CLCN7* | 0.419922 | 0.207016 | 2.028451 | 0.04251426 | 0.70399374 |
| *ENSG00000269983* | 1.136111 | 0.560102 | 2.0284 | 0.04251943 | NA |
| *GEN1* | -0.854771 | 0.421467 | -2.028084 | 0.04255168 | 0.70399374 |
| *RFX1* | -0.447451 | 0.220745 | -2.027003 | 0.04266207 | 0.70399374 |
| *CHMP2A* | -0.252295 | 0.124506 | -2.026366 | 0.0427273 | 0.70399374 |
| *ENSG00000268746* | 1.786184 | 0.881563 | 2.026155 | 0.04274889 | NA |
| *JTB* | -0.137319 | 0.06783 | -2.024456 | 0.0429233 | 0.70399374 |
| *SRPRA* | 0.386249 | 0.190823 | 2.024114 | 0.04295839 | 0.70399374 |
| *RFFL* | 0.501083 | 0.247623 | 2.023574 | 0.04301405 | 0.70399374 |
| *SP4* | -0.648675 | 0.320776 | -2.022204 | 0.04315525 | 0.70399374 |
| *UAP1* | -0.522621 | 0.258469 | -2.021989 | 0.04317752 | 0.70399374 |
| *ZNF793.AS1* | -1.573025 | 0.778043 | -2.021771 | 0.04320003 | NA |
| *RBM14* | 0.365542 | 0.180937 | 2.020269 | 0.04335553 | 0.70399374 |
| *EMC2* | 0.462688 | 0.2291 | 2.019589 | 0.04342602 | 0.70399374 |
| *FAM174B* | 1.367019 | 0.677194 | 2.018651 | 0.04352348 | NA |
| *CDC42SE1* | 0.279777 | 0.138606 | 2.018514 | 0.04353778 | 0.70399374 |
| *COBLL1* | -0.43267 | 0.214379 | -2.018244 | 0.04356584 | 0.70399374 |
| *RIC1* | -0.755823 | 0.374738 | -2.016936 | 0.04370224 | 0.70399374 |
| *BARD1* | 0.414328 | 0.205468 | 2.016512 | 0.04374649 | 0.70399374 |
| *PSMB3* | 0.209246 | 0.103808 | 2.015695 | 0.04383183 | 0.70399374 |
| *HADHA* | -0.23802 | 0.11812 | -2.015074 | 0.04389686 | 0.70399374 |
| *UBA2* | 0.248664 | 0.123408 | 2.01498 | 0.04390678 | 0.70399374 |
| *ADAM10* | -0.361002 | 0.179161 | -2.014953 | 0.04390953 | 0.70399374 |
| *COPS9* | -0.242521 | 0.120377 | -2.014675 | 0.04393875 | 0.70399374 |
| *TM9SF2* | 0.286711 | 0.142354 | 2.01407 | 0.04400221 | 0.70399374 |
| *PYGM* | 0.975405 | 0.484308 | 2.014017 | 0.04400773 | NA |
| *PDCD2L* | 0.686914 | 0.34111 | 2.013762 | 0.04403449 | 0.70399374 |
| *RCC1* | 0.398581 | 0.197942 | 2.013623 | 0.04404912 | 0.70399374 |

| | | | | | |
|---|---|---|---|---|---|
| *ENSG00000231212* | 1.492263 | 0.741091 | 2.013604 | 0.04405114 | NA |
| *FAM129A* | 2.030821 | 1.009271 | 2.012166 | 0.04420244 | NA |
| *TBC1D8* | 1.094406 | 0.543993 | 2.011802 | 0.04424077 | NA |
| *C8orf88* | -0.857199 | 0.426312 | -2.010732 | 0.04435378 | 0.706649081 |
| *KLHDC10* | -0.593551 | 0.29522 | -2.010538 | 0.04437432 | 0.706649081 |
| *ITM2B* | 0.230838 | 0.114893 | 2.009163 | 0.0445198 | 0.707423971 |
| *ENSG00000255856* | 1.597027 | 0.794913 | 2.00906 | 0.04453076 | NA |
| *MYO15B* | -0.764663 | 0.380699 | -2.008575 | 0.0445822 | 0.707423971 |
| *GCSH* | -0.49434 | 0.246272 | -2.007295 | 0.04471828 | 0.708117478 |
| *STYXL1* | 0.509049 | 0.253679 | 2.006666 | 0.04478528 | 0.708117478 |
| *ENSG00000273329* | 0.647006 | 0.322631 | 2.005408 | 0.04491949 | 0.708977872 |
| *GOSR2* | 0.39963 | 0.199457 | 2.003591 | 0.04511387 | 0.709337577 |
| *ENSG00000231856* | -1.007585 | 0.502987 | -2.003205 | 0.04515534 | NA |
| *ANKRD44* | 0.19768 | 0.098714 | 2.002558 | 0.04522473 | 0.709337577 |
| *TAGAP* | 0.24493 | 0.12233 | 2.002206 | 0.04526258 | 0.709337577 |
| *ENSG00000272758* | 0.628261 | 0.313857 | 2.001741 | 0.04531265 | 0.709337577 |
| *ZNF658* | 1.106835 | 0.552984 | 2.001569 | 0.04533112 | NA |
| *RELA* | 0.313001 | 0.156385 | 2.001473 | 0.04534141 | 0.709337577 |
| *FAM50A* | -0.462897 | 0.231522 | -1.999367 | 0.04556862 | 0.711639235 |
| *ADAM19* | 0.363067 | 0.181821 | 1.996836 | 0.04584305 | 0.714668942 |
| *IGLV2.14* | -1.070371 | 0.53695 | -1.993426 | 0.04621483 | NA |
| *ENSG00000277959* | -1.486169 | 0.745839 | -1.992615 | 0.04630362 | NA |
| *STK3* | -0.707039 | 0.354831 | -1.992608 | 0.04630438 | 0.718949978 |
| *S100A11* | -0.427628 | 0.214677 | -1.991957 | 0.04637582 | 0.718949978 |
| *ANKRD36* | -0.58821 | 0.295387 | -1.991318 | 0.04644593 | 0.718949978 |
| *PAXBP1* | -0.784999 | 0.394297 | -1.990881 | 0.04649397 | 0.718949978 |
| *MPPE1* | 0.330866 | 0.166212 | 1.990624 | 0.0465222 | 0.718949978 |
| *TPCN2* | 0.75472 | 0.379572 | 1.988344 | 0.04677369 | NA |
| *LAPTM5* | -0.16618 | 0.08358 | -1.988267 | 0.04678214 | 0.720460663 |
| *ZBTB8OS* | 0.240033 | 0.120791 | 1.987169 | 0.04690368 | 0.720460663 |
| *UTP14A* | -0.537342 | 0.270586 | -1.985843 | 0.04705071 | 0.720460663 |
| *SUCLG2* | 0.330778 | 0.166682 | 1.984488 | 0.04720148 | 0.720460663 |
| *RAB7A* | 0.204762 | 0.103182 | 1.984472 | 0.04720323 | 0.720460663 |
| *SYVN1* | 0.322361 | 0.162443 | 1.984458 | 0.04720482 | 0.720460663 |
| *APH1A* | -0.201953 | 0.101769 | -1.984417 | 0.04720942 | 0.720460663 |
| *TARDBP* | -0.265617 | 0.133887 | -1.983886 | 0.04726858 | 0.720460663 |
| *IL10RB.DT* | -1.408163 | 0.709867 | -1.983702 | 0.04728912 | NA |
| *ZMIZ1* | -1.641262 | 0.82745 | -1.983519 | 0.04730944 | NA |
| *ANKRD34A* | 1.402839 | 0.708359 | 1.980407 | 0.04765776 | NA |
| *ABCB4* | 0.296939 | 0.149959 | 1.980137 | 0.0476881 | 0.724010686 |
| *CYREN* | -0.416447 | 0.210339 | -1.979882 | 0.04771676 | 0.724010686 |
| *BTBD19* | 3.874803 | 1.957147 | 1.979822 | 0.04772348 | NA |
| *DNAAF4* | -1.347148 | 0.680462 | -1.979754 | 0.04773114 | NA |
| *CIR1* | 0.219098 | 0.110677 | 1.979623 | 0.04774592 | 0.724010686 |
| *LITAF* | 0.188483 | 0.09528 | 1.978202 | 0.04790594 | 0.725199614 |
| *MAPK14* | -0.549451 | 0.277899 | -1.977163 | 0.04802325 | 0.725739175 |
| *YPEL2* | -0.283695 | 0.143546 | -1.976339 | 0.04811641 | 0.72579273 |

| | | | | | |
|---|---|---|---|---|---|
| *ITPR1* | 0.378465 | 0.191561 | 1.975688 | 0.04819015 | 0.72579273 |
| *SDR42E1* | -1.513508 | 0.766472 | -1.974643 | 0.04830869 | NA |
| *SLC9A3R1* | 0.393943 | 0.199542 | 1.974238 | 0.04835463 | 0.727037679 |
| *KDM4D* | 1.687461 | 0.855017 | 1.973599 | 0.04842741 | NA |
| *ENSG00000260572* | 1.570798 | 0.795933 | 1.973531 | 0.04843514 | NA |
| *HIP1R* | 0.274686 | 0.139304 | 1.971835 | 0.04862842 | 0.727704943 |
| *PPP2R5B* | -0.531331 | 0.269482 | -1.971674 | 0.04864687 | 0.727704943 |
| *PAGR1* | 0.454342 | 0.230465 | 1.97141 | 0.04867695 | 0.727704943 |
| *CYBC1* | -0.182722 | 0.092706 | -1.970976 | 0.04872659 | 0.727704943 |
| *ENSG00000232010* | 1.116885 | 0.566882 | 1.970224 | 0.04881272 | NA |
| *DMXL1* | -0.393243 | 0.199607 | -1.970086 | 0.0488285 | 0.72800338 |
| *ZNF616* | -1.084889 | 0.550723 | -1.969937 | 0.04884563 | NA |
| *ZBTB32* | 2.39419 | 1.216582 | 1.967965 | 0.04907208 | NA |
| *MAF1* | 0.196153 | 0.09976 | 1.966258 | 0.04926882 | 0.731476049 |
| *IER3* | 1.209917 | 0.615375 | 1.966147 | 0.04928164 | NA |
| *SP100* | 0.164572 | 0.083708 | 1.966031 | 0.04929499 | 0.731476049 |
| *MCM3* | 0.390211 | 0.198488 | 1.965916 | 0.04930837 | 0.731476049 |

| Cluster 3 | log2FC | lfcSE | stat | pvalue | padj |
|---|---|---|---|---|---|
| *VPREB3* | 3.887109 | 0.113796 | 34.15865 | < 2.22e-16 | < 2.22e-16 |
| *DENND6B* | 3.462668 | 0.157059 | 22.046985 | < 2.22e-16 | < 2.22e-16 |
| *GRN* | 2.005959 | 0.102087 | 19.649487 | < 2.22e-16 | < 2.22e-16 |
| *CD1D* | 5.570262 | 0.292138 | 19.067205 | < 2.22e-16 | < 2.22e-16 |
| *CD38* | 4.41865 | 0.241855 | 18.269795 | < 2.22e-16 | < 2.22e-16 |
| *PLAAT4* | 2.769813 | 0.163928 | 16.896484 | < 2.22e-16 | < 2.22e-16 |
| *NRIP1* | 2.270827 | 0.14061 | 16.1498 | < 2.22e-16 | < 2.22e-16 |
| *RABGAP1L* | 1.649413 | 0.117939 | 13.985332 | < 2.22e-16 | < 2.22e-16 |
| *P2RX5* | 1.592606 | 0.119737 | 13.300881 | < 2.22e-16 | < 2.22e-16 |
| *RNF207* | 3.360265 | 0.253871 | 13.236125 | < 2.22e-16 | < 2.22e-16 |
| *AHI1* | 1.619582 | 0.124304 | 13.029232 | < 2.22e-16 | < 2.22e-16 |
| *SLC2A5* | 3.260712 | 0.250436 | 13.020164 | < 2.22e-16 | < 2.22e-16 |
| *RARA.AS1* | 2.30394 | 0.184471 | 12.489433 | < 2.22e-16 | < 2.22e-16 |
| *IRAK2* | 1.543204 | 0.126304 | 12.218144 | < 2.22e-16 | < 2.22e-16 |
| *ASB2* | 8.698217 | 0.743805 | 11.694221 | < 2.22e-16 | < 2.22e-16 |
| *ENDOD1* | 1.92285 | 0.166119 | 11.575132 | < 2.22e-16 | < 2.22e-16 |
| *APOL3* | 1.200093 | 0.104814 | 11.449705 | < 2.22e-16 | < 2.22e-16 |
| *SLC12A4* | 2.492814 | 0.219336 | 11.365275 | < 2.22e-16 | < 2.22e-16 |
| *LCAT* | 2.663433 | 0.235088 | 11.329504 | < 2.22e-16 | < 2.22e-16 |
| *AGO1* | 1.175843 | 0.104094 | 11.296007 | < 2.22e-16 | < 2.22e-16 |
| *C12orf65* | 1.139716 | 0.101061 | 11.277446 | < 2.22e-16 | < 2.22e-16 |
| *DUS2* | 1.622667 | 0.147321 | 11.014527 | < 2.22e-16 | < 2.22e-16 |
| *P2RX5.TAX1BP3* | 1.95049 | 0.179825 | 10.846608 | < 2.22e-16 | < 2.22e-16 |
| *SLC5A3* | 1.500122 | 0.144479 | 10.382947 | < 2.22e-16 | < 2.22e-16 |
| *RMDN2* | 2.307724 | 0.227744 | 10.13297 | < 2.22e-16 | < 2.22e-16 |
| *SEPTIN9* | 0.708708 | 0.072207 | 9.814982 | < 2.22e-16 | < 2.22e-16 |
| *ZNF563* | 1.534761 | 0.158476 | 9.684476 | < 2.22e-16 | < 2.22e-16 |
| *PLAAT3* | 3.583579 | 0.376796 | 9.510663 | < 2.22e-16 | < 2.22e-16 |
| *MS4A1* | -0.808967 | 0.086907 | -9.30845 | < 2.22e-16 | < 2.22e-16 |
| *GBP4* | 1.026648 | 0.110676 | 9.276144 | < 2.22e-16 | < 2.22e-16 |
| *SMAD3* | 1.037043 | 0.112778 | 9.195401 | < 2.22e-16 | < 2.22e-16 |
| *FAM117A* | 0.993384 | 0.111553 | 8.905053 | < 2.22e-16 | < 2.22e-16 |
| *CD72* | 1.337239 | 0.150967 | 8.857796 | < 2.22e-16 | 2.94E-16 |
| *MRPS6* | 1.35474 | 0.153951 | 8.799831 | < 2.22e-16 | 4.79E-16 |
| *RRBP1* | 1.693871 | 0.192593 | 8.795065 | < 2.22e-16 | 4.86E-16 |
| *ABCB1* | 1.708441 | 0.199898 | 8.546577 | < 2.22e-16 | 4.19E-15 |
| *PNOC* | 1.961278 | 0.231614 | 8.467879 | < 2.22e-16 | 8.03E-15 |
| *GPX1* | 0.891985 | 0.106989 | 8.33715 | < 2.22e-16 | 2.38E-14 |
| *DPEP2* | 1.082573 | 0.131339 | 8.242603 | < 2.22e-16 | 5.14E-14 |
| *RARA* | 0.885835 | 0.107989 | 8.203043 | 2.34E-16 | 6.97E-14 |
| *LGMN* | 1.648739 | 0.204954 | 8.044434 | 8.66E-16 | 2.51E-13 |
| *CD24* | -0.863758 | 0.109604 | -7.880722 | 3.25E-15 | 9.21E-13 |
| *DDX24* | 0.49968 | 0.063984 | 7.809511 | 5.74E-15 | 1.59E-12 |
| *FXYD5* | -0.579303 | 0.074307 | -7.796118 | 6.38E-15 | 1.72E-12 |
| *TENT5C* | -0.812347 | 0.104363 | -7.783891 | 7.03E-15 | 1.86E-12 |
| *CNP* | 0.857932 | 0.110331 | 7.775982 | 7.49E-15 | 1.93E-12 |

| | | | | | |
|---|---|---|---|---|---|
| *PAOX* | 0.994067 | 0.128896 | 7.712134 | 1.24E-14 | 3.13E-12 |
| *IFI30* | 1.195293 | 0.161045 | 7.422108 | 1.15E-13 | 2.85E-11 |
| *CDKN1A* | 0.82084 | 0.113982 | 7.201474 | 5.96E-13 | 1.45E-10 |
| *KCNN1* | 2.347863 | 0.328489 | 7.147466 | 8.84E-13 | 2.10E-10 |
| *VPREB1* | 4.315472 | 0.606483 | 7.115575 | 1.11E-12 | 2.60E-10 |
| *TSPAN18* | 5.109549 | 0.719588 | 7.100655 | 1.24E-12 | 2.84E-10 |
| *CD1C* | 1.96068 | 0.277103 | 7.075638 | 1.49E-12 | 3.34E-10 |
| *FCRL2* | 1.156541 | 0.164273 | 7.040356 | 1.92E-12 | 4.22E-10 |
| *TRABD* | 0.585181 | 0.083228 | 7.031081 | 2.05E-12 | 4.43E-10 |
| *RASGEF1B* | -0.819782 | 0.11725 | -6.991718 | 2.72E-12 | 5.76E-10 |
| *SIGIRR* | 0.866777 | 0.125167 | 6.924974 | 4.36E-12 | 9.09E-10 |
| *HIPK2* | 1.217363 | 0.17947 | 6.783093 | 1.18E-11 | 2.41E-09 |
| *IRF2BPL* | 0.998802 | 0.148002 | 6.74859 | 1.49E-11 | 3.01E-09 |
| *CACYBP* | 0.494565 | 0.073622 | 6.717666 | 1.85E-11 | 3.66E-09 |
| *CTSZ* | 0.650209 | 0.097794 | 6.648787 | 2.96E-11 | 5.76E-09 |
| *SLC16A3* | 0.972008 | 0.146426 | 6.638216 | 3.18E-11 | 6.09E-09 |
| *RAB20* | 2.148246 | 0.324046 | 6.629455 | 3.37E-11 | 6.36E-09 |
| *PLXNB2* | 1.744811 | 0.26476 | 6.59015 | 4.39E-11 | 8.16E-09 |
| *AP3B1* | 0.926273 | 0.140724 | 6.582198 | 4.64E-11 | 8.48E-09 |
| *SPRY1* | 1.236919 | 0.188046 | 6.577747 | 4.78E-11 | 8.60E-09 |
| *ARID5B* | -0.623276 | 0.095484 | -6.527532 | 6.69E-11 | 1.19E-08 |
| *FAM3C* | 0.7931 | 0.122298 | 6.485003 | 8.87E-11 | 1.55E-08 |
| *FADS3* | -0.614174 | 0.095564 | -6.426854 | 1.30E-10 | 2.24E-08 |
| *DBNL* | 0.575332 | 0.089881 | 6.401055 | 1.54E-10 | 2.62E-08 |
| *XYLT1* | 0.917143 | 0.143874 | 6.374613 | 1.83E-10 | 3.07E-08 |
| *MCL1* | 0.499427 | 0.078583 | 6.355433 | 2.08E-10 | 3.43E-08 |
| *CABLES1* | -1.585296 | 0.249685 | -6.349177 | 2.16E-10 | 3.52E-08 |
| *BHLHE40* | 0.906383 | 0.144304 | 6.281049 | 3.36E-10 | 5.40E-08 |
| *SLC43A2* | 0.555293 | 0.088521 | 6.273019 | 3.54E-10 | 5.61E-08 |
| *LY9* | -0.823716 | 0.132355 | -6.223547 | 4.86E-10 | 7.60E-08 |
| *BTBD7* | 0.640597 | 0.103166 | 6.209358 | 5.32E-10 | 8.21E-08 |
| *IER3* | 1.555287 | 0.250562 | 6.207199 | 5.39E-10 | 8.22E-08 |
| *SLC45A3* | 1.174046 | 0.190212 | 6.172315 | 6.73E-10 | 1.01E-07 |
| *SLC5A5* | 4.358378 | 0.710893 | 6.130854 | 8.74E-10 | 1.30E-07 |
| *NMT2* | -0.793189 | 0.130464 | -6.079739 | 1.20E-09 | 1.77E-07 |
| *SPOCK2* | 0.696679 | 0.114845 | 6.066238 | 1.31E-09 | 1.90E-07 |
| *GPR183* | -0.989255 | 0.163507 | -6.050223 | 1.45E-09 | 2.07E-07 |
| *ZNF3* | 0.693639 | 0.114886 | 6.037652 | 1.56E-09 | 2.21E-07 |
| *AGPAT5* | 0.857699 | 0.143752 | 5.966536 | 2.42E-09 | 3.39E-07 |
| *SIGLEC10* | 1.270609 | 0.213582 | 5.949059 | 2.70E-09 | 3.73E-07 |
| *VHL* | 0.563169 | 0.09558 | 5.892136 | 3.81E-09 | 5.21E-07 |
| *HIC1* | 3.729863 | 0.635065 | 5.873197 | 4.27E-09 | 5.77E-07 |
| *LHFPL2* | 1.271461 | 0.216828 | 5.863923 | 4.52E-09 | 6.04E-07 |
| *FOXO3* | 0.998689 | 0.170848 | 5.845497 | 5.05E-09 | 6.67E-07 |
| *NUP210* | 0.520615 | 0.089294 | 5.830369 | 5.53E-09 | 7.22E-07 |
| *CLEC17A* | 0.92956 | 0.160111 | 5.805734 | 6.41E-09 | 8.28E-07 |
| *RHOBTB3* | 2.743626 | 0.475744 | 5.767027 | 8.07E-09 | 1.03E-06 |

| | | | | | |
|---|---|---|---|---|---|
| *SREBF1* | 1.180648 | 0.206016 | 5.73085 | 9.99E-09 | 1.26E-06 |
| *HDAC9* | 0.690904 | 0.121547 | 5.684244 | 1.31E-08 | 1.64E-06 |
| *ENSG00000166927* | -1.402074 | 0.247638 | -5.661797 | 1.50E-08 | 1.85E-06 |
| *CHST15* | 0.921738 | 0.163098 | 5.65142 | 1.59E-08 | 1.95E-06 |
| *FXYD7* | -1.496551 | 0.266737 | -5.610589 | 2.02E-08 | 2.45E-06 |
| *LBR* | 0.449293 | 0.081131 | 5.537885 | 3.06E-08 | 3.68E-06 |
| *ATXN10* | 0.633054 | 0.114949 | 5.507249 | 3.64E-08 | 4.33E-06 |
| *FXYD1* | -1.319706 | 0.24028 | -5.49237 | 3.97E-08 | 4.67E-06 |
| *KANSL3* | 0.990913 | 0.180556 | 5.488131 | 4.06E-08 | 4.73E-06 |
| *GDPGP1* | 1.312121 | 0.240043 | 5.466191 | 4.60E-08 | 5.31E-06 |
| *PLEKHO1* | 0.515916 | 0.094459 | 5.461812 | 4.71E-08 | 5.39E-06 |
| *HCK* | -1.30449 | 0.239268 | -5.451998 | 4.98E-08 | 5.64E-06 |
| *SMC6* | -0.449224 | 0.082717 | -5.430869 | 5.61E-08 | 6.29E-06 |
| *ARL14EPL* | 4.386995 | 0.813118 | 5.395276 | 6.84E-08 | 7.60E-06 |
| *RESF1* | -0.40539 | 0.075352 | -5.379974 | 7.45E-08 | 8.20E-06 |
| *MDM4* | -0.379953 | 0.070893 | -5.359512 | 8.34E-08 | 9.10E-06 |
| *CASP7* | 0.70752 | 0.132085 | 5.356556 | 8.48E-08 | 9.17E-06 |
| *ENSG00000214797* | 2.313623 | 0.432797 | 5.345741 | 9.00E-08 | 9.64E-06 |
| *GBP7* | 1.449089 | 0.271768 | 5.332079 | 9.71E-08 | 1.03E-05 |
| *PUS1* | 0.843946 | 0.158514 | 5.324117 | 1.01E-07 | 1.07E-05 |
| *SEL1L3* | 0.694927 | 0.130842 | 5.311199 | 1.09E-07 | 1.14E-05 |
| *ARHGEF3* | 0.611048 | 0.115355 | 5.297122 | 1.18E-07 | 1.21E-05 |
| *PPCDC* | 0.996469 | 0.188125 | 5.296859 | 1.18E-07 | 1.21E-05 |
| *SLCO5A1* | 1.79338 | 0.342955 | 5.229194 | 1.70E-07 | 1.73E-05 |
| *FCRL1* | 0.98629 | 0.190106 | 5.188096 | 2.12E-07 | 2.14E-05 |
| *JUNB* | 0.43347 | 0.083864 | 5.168719 | 2.36E-07 | 2.35E-05 |
| *MAPKAPK2* | 0.764263 | 0.147987 | 5.164405 | 2.41E-07 | 2.39E-05 |
| *NDUFB9* | 0.362113 | 0.070294 | 5.151397 | 2.59E-07 | 2.54E-05 |
| *ENSG00000261448* | 1.760381 | 0.343043 | 5.131662 | 2.87E-07 | 2.78E-05 |
| *MZB1* | -1.040178 | 0.202719 | -5.131127 | 2.88E-07 | 2.78E-05 |
| *JCHAIN* | 2.189688 | 0.427454 | 5.12263 | 3.01E-07 | 2.89E-05 |
| *ZFP36L1* | -0.592864 | 0.116713 | -5.079669 | 3.78E-07 | 3.60E-05 |
| *SH3KBP1* | -0.431457 | 0.085466 | -5.04828 | 4.46E-07 | 4.21E-05 |
| *C15orf39* | 0.933854 | 0.185365 | 5.037927 | 4.71E-07 | 4.40E-05 |
| *GGA2* | 0.528325 | 0.105271 | 5.01871 | 5.20E-07 | 4.83E-05 |
| *CPEB4* | 0.916013 | 0.184396 | 4.967634 | 6.78E-07 | 6.25E-05 |
| *BRK1* | 0.270058 | 0.05446 | 4.958795 | 7.09E-07 | 6.49E-05 |
| *CIB1* | 0.340434 | 0.068888 | 4.941834 | 7.74E-07 | 7.02E-05 |
| *LBH* | -0.456258 | 0.092623 | -4.925966 | 8.39E-07 | 7.56E-05 |
| *REC8* | 0.773535 | 0.15715 | 4.922261 | 8.56E-07 | 7.65E-05 |
| *GPR146* | -1.34108 | 0.272539 | -4.920685 | 8.62E-07 | 7.65E-05 |
| *ICAM1* | 1.242197 | 0.252793 | 4.913899 | 8.93E-07 | 7.86E-05 |
| *PALLD* | 2.421305 | 0.497314 | 4.86876 | 1.12E-06 | 9.82E-05 |
| *HTR3A* | 1.748883 | 0.359481 | 4.865027 | 1.14E-06 | 9.93E-05 |
| *PIP5K1B* | 1.084064 | 0.223207 | 4.85677 | 1.19E-06 | 0.000102776 |
| *CD180* | 0.859127 | 0.176978 | 4.854435 | 1.21E-06 | 0.000103246 |
| *CHST7* | 2.046558 | 0.421798 | 4.851984 | 1.22E-06 | 0.000103784 |

| | | | | | |
|---|---|---|---|---|---|
| *SERPINF1* | -1.742716 | 0.361004 | -4.827419 | 1.38E-06 | 0.000116605 |
| *FOXP1* | -0.208179 | 0.043243 | -4.814116 | 1.48E-06 | 0.00012377 |
| *CYP2R1* | -0.669923 | 0.139855 | -4.790121 | 1.67E-06 | 0.000138555 |
| *ENSG00000198106* | -1.016351 | 0.213417 | -4.762288 | 1.91E-06 | 0.000158006 |
| *APOBEC3H* | 1.639532 | 0.344562 | 4.758307 | 1.95E-06 | 0.000160043 |
| *ISCA1* | 0.382806 | 0.080669 | 4.745377 | 2.08E-06 | 0.000169445 |
| *MPEG1* | 1.521988 | 0.321243 | 4.73781 | 2.16E-06 | 0.000174699 |
| *KLF13* | 0.397485 | 0.084114 | 4.725568 | 2.29E-06 | 0.000184307 |
| *NFAT5* | 0.643041 | 0.136231 | 4.720228 | 2.36E-06 | 0.000187943 |
| *C16orf54* | 0.625699 | 0.13261 | 4.718336 | 2.38E-06 | 0.000188434 |
| *FYTTD1* | -0.408813 | 0.086794 | -4.710147 | 2.48E-06 | 0.000194866 |
| *SYTL1* | -0.612436 | 0.1303 | -4.700194 | 2.60E-06 | 0.000203263 |
| *GLIPR2* | 0.636961 | 0.136359 | 4.671204 | 2.99E-06 | 0.000232643 |
| *DDX28* | 0.621814 | 0.133314 | 4.664294 | 3.10E-06 | 0.000239036 |
| *STK17B* | -0.489648 | 0.105323 | -4.649039 | 3.33E-06 | 0.000255751 |
| *PSMB10* | 0.418783 | 0.090864 | 4.608915 | 4.05E-06 | 0.000308434 |
| *KDM2B* | 0.4484 | 0.097353 | 4.605927 | 4.11E-06 | 0.000310903 |
| *SPN* | 1.556909 | 0.339755 | 4.582447 | 4.60E-06 | 0.000345752 |
| *ANAPC16* | 0.369124 | 0.080692 | 4.574482 | 4.77E-06 | 0.000355927 |
| *NANP* | 1.05869 | 0.231471 | 4.573746 | 4.79E-06 | 0.000355927 |
| *H3F3A* | 0.385573 | 0.085612 | 4.503712 | 6.68E-06 | 0.000493027 |
| *BCL11A* | -0.469864 | 0.104836 | -4.481899 | 7.40E-06 | 0.000542852 |
| *ENSG00000274536* | -1.142875 | 0.255221 | -4.477977 | 7.54E-06 | 0.000549527 |
| *CD79B* | 0.369764 | 0.082616 | 4.475711 | 7.62E-06 | 0.000550738 |
| *ICAM2* | 0.671641 | 0.150091 | 4.474902 | 7.64E-06 | 0.000550738 |
| *LINC02576* | -1.993854 | 0.447288 | -4.457654 | 8.29E-06 | 0.000593358 |
| *EHD1* | 0.499822 | 0.112779 | 4.431882 | 9.34E-06 | 0.000664919 |
| *NXT1* | 0.394871 | 0.089313 | 4.42121 | 9.81E-06 | 0.000694467 |
| *MIR181A2HG* | 1.507331 | 0.3412 | 4.41773 | 9.97E-06 | 0.000701565 |
| *RUNX2* | 3.096334 | 0.701543 | 4.413604 | 1.02E-05 | 0.000710868 |
| *CERK* | 0.838742 | 0.190491 | 4.40304 | 1.07E-05 | 0.000741664 |
| *ITGB7* | 0.52488 | 0.11924 | 4.401883 | 1.07E-05 | 0.000741664 |
| *PCBP2* | -0.202665 | 0.046247 | -4.38228 | 1.17E-05 | 0.000803604 |
| *TGFBR2* | -0.400988 | 0.091509 | -4.381934 | 1.18E-05 | 0.000803604 |
| *BCAR3* | 1.770814 | 0.406444 | 4.356843 | 1.32E-05 | 0.000896294 |
| *ST3GAL1* | 0.670778 | 0.154233 | 4.349124 | 1.37E-05 | 0.000923152 |
| *CLIC3* | 2.296545 | 0.528866 | 4.342393 | 1.41E-05 | 0.000946523 |
| *CHRAC1* | -0.406758 | 0.093805 | -4.336227 | 1.45E-05 | 0.000966781 |
| *CTNND1* | -0.575463 | 0.13274 | -4.335268 | 1.46E-05 | 0.000966781 |
| *SMIM3* | 1.888098 | 0.435993 | 4.330571 | 1.49E-05 | 0.000982153 |
| *SELENOF* | 0.216076 | 0.050068 | 4.315686 | 1.59E-05 | 0.001044924 |
| *PPFIBP2* | 0.770316 | 0.178979 | 4.303951 | 1.68E-05 | 0.001090747 |
| *SLC2A3* | -0.647438 | 0.150435 | -4.303765 | 1.68E-05 | 0.001090747 |
| *CPNE5* | 1.279915 | 0.298616 | 4.286152 | 1.82E-05 | 0.001174452 |
| *GAK* | 0.457713 | 0.107467 | 4.259102 | 2.05E-05 | 0.001318813 |
| *RINL* | 0.446452 | 0.10489 | 4.256377 | 2.08E-05 | 0.001327807 |
| *MDM2* | 0.496295 | 0.117238 | 4.233224 | 2.30E-05 | 0.001464354 |

| | | | | | |
|---|---|---|---|---|---|
| *KMT2E* | -0.233903 | 0.055368 | -4.224508 | 2.39E-05 | 0.001514099 |
| *C9orf16* | -0.326691 | 0.07739 | -4.221366 | 2.43E-05 | 0.001524221 |
| *CHP1* | -0.375552 | 0.08898 | -4.220622 | 2.44E-05 | 0.001524221 |
| *PSME2* | 0.338298 | 0.080194 | 4.218486 | 2.46E-05 | 0.001530672 |
| *PPP1R18* | 0.458979 | 0.109026 | 4.20983 | 2.56E-05 | 0.001576997 |
| *RPS4X* | -0.393031 | 0.09337 | -4.209404 | 2.56E-05 | 0.001576997 |
| *NRM* | 0.58498 | 0.139311 | 4.199101 | 2.68E-05 | 0.001641976 |
| *UBTF* | 0.448848 | 0.107412 | 4.178747 | 2.93E-05 | 0.001783428 |
| *MCM5* | 0.531499 | 0.127213 | 4.178017 | 2.94E-05 | 0.001783428 |
| *SNAP23* | 0.279611 | 0.067384 | 4.149509 | 3.33E-05 | 0.002010467 |
| *CTSB* | 0.475908 | 0.115365 | 4.125243 | 3.70E-05 | 0.002218475 |
| *NIN* | -0.441364 | 0.107008 | -4.124591 | 3.71E-05 | 0.002218475 |
| *DMD* | -0.764948 | 0.185744 | -4.118288 | 3.82E-05 | 0.002268617 |
| *TELO2* | 0.67223 | 0.163607 | 4.108815 | 3.98E-05 | 0.00235194 |
| *POU2AF1* | 0.356068 | 0.086794 | 4.102451 | 4.09E-05 | 0.002405621 |
| *TFAM* | 0.423631 | 0.103424 | 4.096071 | 4.20E-05 | 0.002460679 |
| *MAST4* | -1.418296 | 0.346938 | -4.088038 | 4.35E-05 | 0.002534941 |
| *SIGLEC14* | 1.092079 | 0.267623 | 4.080658 | 4.49E-05 | 0.002597933 |
| *TOE1* | 0.592041 | 0.145106 | 4.080072 | 4.50E-05 | 0.002597933 |
| *NPM1* | -0.294616 | 0.072288 | -4.075587 | 4.59E-05 | 0.002635728 |
| *EBF1* | -0.63628 | 0.156918 | -4.054863 | 5.02E-05 | 0.002866807 |
| *ERAP1* | 0.525501 | 0.129917 | 4.044899 | 5.23E-05 | 0.002977191 |
| *LSP1* | 0.421493 | 0.104366 | 4.038617 | 5.38E-05 | 0.00304224 |
| *MYC* | 0.531569 | 0.131655 | 4.037598 | 5.40E-05 | 0.00304224 |
| *SMARCB1* | 0.349729 | 0.086748 | 4.031542 | 5.54E-05 | 0.003106996 |
| *PTPRE* | -0.96363 | 0.239299 | -4.026878 | 5.65E-05 | 0.003154366 |
| *FAM167A* | 1.447748 | 0.360402 | 4.017034 | 5.89E-05 | 0.003264198 |
| *SESN3* | -0.357581 | 0.089025 | -4.016617 | 5.90E-05 | 0.003264198 |
| *PAG1* | -1.064004 | 0.265862 | -4.002095 | 6.28E-05 | 0.003455163 |
| *ZNF44* | 0.458085 | 0.114726 | 3.992874 | 6.53E-05 | 0.003575811 |
| *C7orf50* | -0.70333 | 0.176779 | -3.978584 | 6.93E-05 | 0.003771726 |
| *ENSG00000261766* | 0.760216 | 0.191104 | 3.978031 | 6.95E-05 | 0.003771726 |
| *NCF1* | 0.390785 | 0.098364 | 3.972861 | 7.10E-05 | 0.003837039 |
| *ATP6V1E1* | 0.423805 | 0.107201 | 3.953361 | 7.71E-05 | 0.004144908 |
| *A4GALT* | 1.504609 | 0.3813 | 3.946001 | 7.95E-05 | 0.004255088 |
| *TTC3* | -0.336015 | 0.085185 | -3.944507 | 8.00E-05 | 0.004262493 |
| *GTF2E2* | 0.429522 | 0.108978 | 3.941351 | 8.10E-05 | 0.004299692 |
| *ABI3* | 0.577494 | 0.146729 | 3.935791 | 8.29E-05 | 0.004380921 |
| *RUBCN* | -0.60453 | 0.153684 | -3.933582 | 8.37E-05 | 0.004401828 |
| *PHF21A* | -0.454951 | 0.11595 | -3.923682 | 8.72E-05 | 0.004566595 |
| *TOP1* | 0.335956 | 0.085666 | 3.921709 | 8.79E-05 | 0.004583955 |
| *CD48* | -0.427216 | 0.109237 | -3.910896 | 9.20E-05 | 0.00477319 |
| *DDIT4* | 1.288359 | 0.329721 | 3.907419 | 9.33E-05 | 0.004821324 |
| *CCNDBP1* | 0.281604 | 0.072423 | 3.888339 | 0.000100933 | 0.005193876 |
| *TNFSF4* | 0.742124 | 0.190963 | 3.886221 | 0.000101817 | 0.005216793 |
| *DPEP3* | 1.860634 | 0.478944 | 3.88487 | 0.000102385 | 0.005223378 |
| *RGS3* | 0.84847 | 0.219478 | 3.865852 | 0.000110702 | 0.005623579 |

| | | | | | |
|---|---|---|---|---|---|
| *NEURL2* | 1.574146 | 0.407316 | 3.864683 | 0.000111234 | 0.005626527 |
| *TP53I11* | -0.705275 | 0.182836 | -3.857421 | 0.00011459 | 0.005771739 |
| *ATP13A2* | 0.709937 | 0.184337 | 3.851304 | 0.00011749 | 0.00589286 |
| *ENSG00000233038* | 4.054184 | 1.053725 | 3.847478 | 0.00011934 | NA |
| *WDR45* | 0.296611 | 0.077112 | 3.846503 | 0.000119816 | 0.005984249 |
| *ATP11A* | -1.066278 | 0.278027 | -3.835157 | 0.000125484 | 0.006241133 |
| *ZFP36L2* | -0.6403 | 0.16724 | -3.828631 | 0.000128858 | 0.00638222 |
| *ULK1* | 0.699476 | 0.183189 | 3.818336 | 0.000134355 | 0.006626868 |
| *ZCCHC7* | 0.362161 | 0.094948 | 3.814312 | 0.000136563 | 0.006707955 |
| *MACROD2* | -0.873483 | 0.229806 | -3.800954 | 0.00014414 | 0.007050996 |
| *ATF7IP2* | 0.510117 | 0.134535 | 3.791718 | 0.000149609 | 0.007288512 |
| *PWP1* | 0.298863 | 0.078848 | 3.790366 | 0.000150425 | 0.007298392 |
| *CR2* | 1.175512 | 0.310321 | 3.788057 | 0.00015183 | 0.007336612 |
| *P2RY8* | 0.42613 | 0.112742 | 3.779678 | 0.000157032 | 0.007557223 |
| *CCR7* | 0.414306 | 0.109779 | 3.774016 | 0.000160641 | 0.007699739 |
| *ZBED2* | -1.220586 | 0.323577 | -3.772169 | 0.000161835 | 0.007725824 |
| *SSR2* | -0.294638 | 0.078214 | -3.767091 | 0.000165161 | 0.00785307 |
| *LMAN1* | 0.359306 | 0.095471 | 3.763519 | 0.000167539 | 0.007934415 |
| *ENSG00000278158* | -0.417806 | 0.111101 | -3.760587 | 0.000169515 | 0.007973008 |
| *ARHGEF1* | 0.420163 | 0.111736 | 3.760321 | 0.000169696 | 0.007973008 |
| *RHOA* | 0.250468 | 0.066837 | 3.747457 | 0.000178636 | 0.008360038 |
| *RPL12* | -0.591133 | 0.157959 | -3.742307 | 0.000182338 | 0.008499827 |
| *TMEM65* | 0.663069 | 0.177567 | 3.73418 | 0.000188327 | 0.008711839 |
| *SNX18* | 0.785195 | 0.210274 | 3.734147 | 0.000188352 | 0.008711839 |
| *ATP2B1* | -0.324333 | 0.086918 | -3.731488 | 0.000190352 | 0.008770229 |
| *PSAP* | 0.249245 | 0.066958 | 3.722401 | 0.000197337 | 0.009052631 |
| *EIF3E* | -0.308708 | 0.082951 | -3.721548 | 0.000198005 | 0.009052631 |
| *PFDN5* | -0.22885 | 0.061639 | -3.712768 | 0.000205005 | 0.009336757 |
| *FTH1* | -0.250061 | 0.06744 | -3.707917 | 0.000208971 | 0.009481078 |
| *SIDT1* | -0.537254 | 0.145023 | -3.704609 | 0.000211717 | 0.009569117 |
| *ST13* | -0.24678 | 0.066734 | -3.697962 | 0.000217337 | 0.00978595 |
| *ZNF830* | 0.339915 | 0.091946 | 3.696909 | 0.000218241 | 0.009789542 |
| *ETFRF1* | 0.445182 | 0.120517 | 3.693922 | 0.000220821 | 0.009868054 |
| *SNRPB2* | 0.259412 | 0.070352 | 3.687325 | 0.000226623 | 0.010089411 |
| *SLC25A5* | -0.303777 | 0.08243 | -3.685273 | 0.000228458 | 0.010133128 |
| *PDCD4* | 0.704872 | 0.191356 | 3.683553 | 0.000230005 | 0.010163833 |
| *REL* | -0.442769 | 0.120399 | -3.677517 | 0.000235515 | 0.010368765 |
| *RGS1* | -1.571377 | 0.42802 | -3.671268 | 0.00024135 | 0.010586459 |
| *PXK* | 0.474109 | 0.129178 | 3.67019 | 0.00024237 | 0.010592124 |
| *MAP3K1* | 0.392712 | 0.107101 | 3.666732 | 0.00024567 | 0.010697011 |
| *GPER1* | -2.612246 | 0.713151 | -3.66296 | 0.000249317 | 0.010774617 |
| *HLA.DPB1* | -0.51473 | 0.140525 | -3.662913 | 0.000249363 | 0.010774617 |
| *CLCN7* | 0.497676 | 0.1359 | 3.662084 | 0.000250172 | 0.010774617 |
| *LYRM9* | -0.617622 | 0.168779 | -3.659348 | 0.000252858 | 0.010850989 |
| *TAGLN2* | 0.255376 | 0.069996 | 3.648445 | 0.000263832 | 0.011281207 |
| *AHRR* | -2.236515 | 0.613565 | -3.645113 | 0.000267274 | 0.011364525 |
| *CD79A* | 0.474686 | 0.13024 | 3.64471 | 0.000267693 | 0.011364525 |

| | | | | | |
|---|---|---|---|---|---|
| *SLA* | 0.468441 | 0.129454 | 3.618602 | 0.000296198 | 0.012529928 |
| *ENSG00000237513* | -0.641704 | 0.178684 | -3.591286 | 0.00032905 | 0.013870266 |
| *MAN2A2* | -0.696302 | 0.194421 | -3.581414 | 0.00034174 | 0.014354298 |
| *RPL10* | -0.456714 | 0.12773 | -3.575612 | 0.000349409 | 0.014624733 |
| *IER2* | 0.239844 | 0.067111 | 3.573859 | 0.000351759 | 0.014671424 |
| *NFKBIA* | 0.283904 | 0.079462 | 3.572837 | 0.000353135 | 0.014677328 |
| *ENSG00000251364* | 1.376933 | 0.385592 | 3.570954 | 0.000355684 | 0.01469197 |
| *SDE2* | 0.667279 | 0.186917 | 3.569922 | 0.000357088 | 0.01469197 |
| *TBC1D9* | 0.603621 | 0.169123 | 3.569133 | 0.000358164 | 0.01469197 |
| *PISD* | 0.529688 | 0.148416 | 3.568938 | 0.000358431 | 0.01469197 |
| *SRP14* | 0.13772 | 0.038664 | 3.561922 | 0.000368149 | 0.015038451 |
| *VSIG10L* | 2.0132 | 0.566504 | 3.553725 | 0.000379816 | 0.015461874 |
| *MDK* | 0.8345 | 0.234977 | 3.551414 | 0.000383168 | 0.015545101 |
| *SUSD3* | -0.728447 | 0.205927 | -3.537413 | 0.000404067 | 0.016337235 |
| *ENSG00000131408* | 0.245936 | 0.069705 | 3.528225 | 0.000418356 | 0.016857627 |
| *BATF2* | 1.774538 | 0.503394 | 3.52515 | 0.000423243 | 0.016996908 |
| *WDR11* | 0.353432 | 0.100286 | 3.524254 | 0.000424677 | 0.016997098 |
| *LPP* | -0.51529 | 0.146475 | -3.517929 | 0.000434928 | 0.017348966 |
| *UXT* | -0.205126 | 0.058496 | -3.506633 | 0.000453815 | 0.018041787 |
| *TSG101* | 0.382985 | 0.109899 | 3.484884 | 0.000492351 | 0.019508582 |
| *MYADM* | 0.517799 | 0.148705 | 3.482052 | 0.000497586 | 0.019643464 |
| *CD44* | -0.329777 | 0.094742 | -3.480803 | 0.000499912 | 0.019643464 |
| *ECHS1* | 0.381139 | 0.109511 | 3.480375 | 0.000500713 | 0.019643464 |
| *TOP1MT* | -0.638538 | 0.183694 | -3.476103 | 0.000508757 | 0.019870504 |
| *OTUD7A* | 1.758607 | 0.505997 | 3.475531 | 0.000509843 | 0.019870504 |
| *PXDC1* | 1.747081 | 0.503367 | 3.470791 | 0.000518928 | 0.020158468 |
| *AFF3* | -0.348175 | 0.100367 | -3.469029 | 0.000522342 | 0.020225028 |
| *EML2* | 0.532355 | 0.153551 | 3.466961 | 0.000526378 | 0.020282028 |
| *GTF2B* | 0.457466 | 0.131989 | 3.465927 | 0.000528407 | 0.020282028 |
| *ST6GAL1* | -0.390586 | 0.112735 | -3.464629 | 0.000530964 | 0.020282028 |
| *MTSS1* | -0.418186 | 0.120703 | -3.464577 | 0.000531066 | 0.020282028 |
| *TAF12* | 0.29561 | 0.08534 | 3.46393 | 0.000532346 | 0.020282028 |
| *DNAJC7* | 0.267624 | 0.077343 | 3.46023 | 0.000539715 | 0.020497091 |
| *ACP5* | 1.041597 | 0.301122 | 3.459054 | 0.000542076 | 0.020521193 |
| *CHCHD10* | 1.556237 | 0.450444 | 3.454896 | 0.000550505 | 0.020774134 |
| *FCGRT* | -0.462273 | 0.13388 | -3.452877 | 0.000554642 | 0.020864005 |
| *DPH5* | -0.409872 | 0.1188 | -3.450115 | 0.000560349 | 0.021012186 |
| *FCER2* | 1.437275 | 0.41746 | 3.442901 | 0.00057551 | 0.021512848 |
| *C1orf162* | -0.270686 | 0.078776 | -3.436123 | 0.000590102 | 0.021989174 |
| *SIGLEC5* | 1.368259 | 0.399086 | 3.428478 | 0.000606975 | 0.022547223 |
| *OLA1* | -0.352954 | 0.10325 | -3.41846 | 0.000629765 | 0.023320937 |
| *TMEM134* | -0.288398 | 0.0844 | -3.417047 | 0.000633044 | 0.023369531 |
| *DTX4* | 0.640149 | 0.187533 | 3.413535 | 0.00064126 | 0.023599568 |
| *ENTPD1* | -0.724426 | 0.212318 | -3.411982 | 0.000644923 | 0.023610967 |
| *ENSG00000089127* | 0.555286 | 0.162758 | 3.411721 | 0.000645543 | 0.023610967 |
| *TMEM59* | 0.206948 | 0.060705 | 3.409096 | 0.000651785 | 0.023766176 |
| *FAU* | -0.251635 | 0.073846 | -3.407573 | 0.000655434 | 0.023826113 |

| | | | | | |
|---|---|---|---|---|---|
| *C8orf58* | -0.819553 | 0.240834 | -3.402979 | 0.000666554 | 0.02415648 |
| *STK17A* | 0.362104 | 0.106459 | 3.401344 | 0.000670554 | 0.024227586 |
| *COA1* | 0.415377 | 0.122158 | 3.400325 | 0.000673057 | 0.024244336 |
| *SLC25A19* | 0.422716 | 0.124392 | 3.398246 | 0.000678194 | 0.024355554 |
| *CXXC5* | -0.374829 | 0.110388 | -3.395559 | 0.000684885 | 0.024521783 |
| *FAM129C* | -0.36877 | 0.108753 | -3.390899 | 0.000696636 | 0.02486762 |
| *NAP1L1* | -0.357927 | 0.105596 | -3.389575 | 0.000700011 | 0.024913274 |
| *LARGE1* | 1.432508 | 0.424098 | 3.37778 | 0.000730735 | 0.025845512 |
| *RAB7A* | 0.267524 | 0.079218 | 3.37707 | 0.000732624 | 0.025845512 |
| *ZNF581* | -0.404375 | 0.119765 | -3.376417 | 0.000734365 | 0.025845512 |
| *CARHSP1* | 0.251882 | 0.074605 | 3.376216 | 0.000734902 | 0.025845512 |
| *NUBP1* | 0.343553 | 0.101803 | 3.374681 | 0.000739012 | 0.025913379 |
| *EMC6* | -0.312578 | 0.092802 | -3.36822 | 0.000756552 | 0.026450407 |
| *RAB37* | 0.699093 | 0.207711 | 3.365706 | 0.00076348 | 0.026603435 |
| *CYBA* | 0.28184 | 0.083756 | 3.365011 | 0.000765405 | 0.026603435 |
| *MRPL39* | 0.407429 | 0.121244 | 3.360393 | 0.000778316 | 0.02697329 |
| *ZNF337.AS1* | 0.908178 | 0.270832 | 3.353289 | 0.000798574 | 0.027594895 |
| *NDUFA2* | 0.249105 | 0.07439 | 3.348648 | 0.000812069 | 0.027979891 |
| *TMEM159* | -0.527198 | 0.157798 | -3.340962 | 0.000834888 | 0.028682987 |
| *CCNI* | -0.21217 | 0.063589 | -3.336587 | 0.000848139 | 0.029054244 |
| *ZNF280B* | -1.187887 | 0.357429 | -3.323418 | 0.000889217 | 0.030373904 |
| *POLM* | 0.364002 | 0.109771 | 3.316008 | 0.000913132 | 0.031101436 |
| *NCOA4* | 0.338916 | 0.102242 | 3.314826 | 0.000917001 | 0.031143967 |
| *ENSG00000225489* | 1.196544 | 0.361518 | 3.309775 | 0.000933709 | 0.031621069 |
| *GFOD1* | -0.749875 | 0.226689 | -3.307943 | 0.00093984 | 0.031738287 |
| *AP3S1* | 0.502581 | 0.152018 | 3.306065 | 0.000946161 | 0.031861238 |
| *ALOX5* | 0.323351 | 0.09785 | 3.30457 | 0.000951221 | 0.03194115 |
| *TBL1X* | -0.531599 | 0.161086 | -3.300086 | 0.000966552 | 0.032364511 |
| *ETFB* | 0.392031 | 0.118828 | 3.299151 | 0.000969779 | 0.032381355 |
| *SNU13* | 0.160653 | 0.048714 | 3.297906 | 0.000974089 | 0.032434163 |
| *PPP3CA* | -0.250518 | 0.076008 | -3.29592 | 0.000980999 | 0.032573013 |
| *LARGE2* | -0.453981 | 0.137818 | -3.294052 | 0.000987543 | 0.032644875 |
| *KEAP1* | 0.539933 | 0.163927 | 3.293735 | 0.000988656 | 0.032644875 |
| *NEIL1* | -0.868306 | 0.263767 | -3.29194 | 0.000994989 | 0.032762975 |
| *HERC2* | -0.449128 | 0.136529 | -3.289622 | 0.001003219 | 0.032942715 |
| *CD22* | 0.524729 | 0.159597 | 3.287841 | 0.001009588 | 0.032969805 |
| *PCNA* | 0.46281 | 0.140764 | 3.28784 | 0.001009591 | 0.032969805 |
| *CD47* | -0.272887 | 0.083387 | -3.272539 | 0.001065863 | 0.034712088 |
| *MRPS22* | 0.349795 | 0.107271 | 3.260857 | 0.001110761 | 0.036075442 |
| *HERPUD1* | -0.300123 | 0.092135 | -3.257419 | 0.001124304 | 0.036415821 |
| *UBE2D1* | 0.303751 | 0.093443 | 3.250669 | 0.001151338 | 0.037115388 |
| *NPIPB11* | -1.576177 | 0.484907 | -3.250469 | 0.001152148 | 0.037115388 |
| *MMP7* | 1.794069 | 0.552361 | 3.248 | 0.001162192 | 0.037337763 |
| *RPL3* | -0.379696 | 0.11703 | -3.244423 | 0.001176887 | 0.037632703 |
| *CD37* | -0.261224 | 0.08052 | -3.244226 | 0.001177704 | 0.037632703 |
| *RACK1* | -0.392914 | 0.121211 | -3.241571 | 0.001188728 | 0.037883128 |
| *LMNA* | -1.581539 | 0.488632 | -3.236667 | 0.001209346 | 0.038390519 |

| | | | | | |
|---|---|---|---|---|---|
| *EXOSC5* | 0.471437 | 0.145703 | 3.235604 | 0.001213855 | 0.038390519 |
| *EHD4* | -0.396358 | 0.122503 | -3.235491 | 0.001214338 | 0.038390519 |
| *CYTH1* | 0.205042 | 0.063414 | 3.233366 | 0.001223409 | 0.038574695 |
| *HIGD1A* | 0.386785 | 0.1197 | 3.231302 | 0.001232275 | 0.038751454 |
| *PIM1* | -0.355713 | 0.11012 | -3.230214 | 0.001236977 | 0.038796686 |
| *LINC02202* | -2.359982 | 0.73082 | -3.229223 | 0.00124127 | NA |
| *GPR155* | -0.766562 | 0.237664 | -3.225395 | 0.001257991 | 0.039351949 |
| *WDR92* | -0.604733 | 0.187594 | -3.223631 | 0.001265766 | 0.03949122 |
| *ERP29* | -0.263559 | 0.081811 | -3.221568 | 0.001274912 | 0.039672443 |
| *PSD3* | -0.838979 | 0.260795 | -3.217013 | 0.001295329 | 0.040202557 |
| *DMXL1* | -0.373384 | 0.116384 | -3.208202 | 0.001335677 | 0.041346851 |
| *POLR2G* | 0.337568 | 0.105353 | 3.204154 | 0.001354598 | 0.041823647 |
| *ENSG00000188206* | -0.420189 | 0.131245 | -3.201568 | 0.001366816 | 0.042091561 |
| *PTP4A3* | 1.569688 | 0.490933 | 3.197357 | 0.001386933 | 0.042600708 |
| *BCL2L11* | -0.701156 | 0.219591 | -3.193014 | 0.00140796 | 0.043135111 |
| *RPL11* | -0.280579 | 0.087966 | -3.189617 | 0.001424615 | 0.043533152 |
| *GABARAPL2* | 0.179016 | 0.056207 | 3.184928 | 0.001447901 | 0.044114224 |
| *SNN* | 0.587842 | 0.184607 | 3.184299 | 0.001451053 | 0.044114224 |
| *CAMK2D* | -0.328817 | 0.103534 | -3.17592 | 0.001493622 | 0.045155277 |
| *COQ8A* | 0.4684 | 0.147504 | 3.17551 | 0.001495734 | 0.045155277 |
| *PTK2B* | 0.424935 | 0.133824 | 3.175324 | 0.001496692 | 0.045155277 |
| *TCF4* | -0.249942 | 0.078851 | -3.1698 | 0.001525439 | 0.045906053 |
| *TERF2IP* | 0.265009 | 0.083764 | 3.163776 | 0.001557368 | 0.046725143 |
| *C1orf131* | -0.3944 | 0.124685 | -3.163187 | 0.001560518 | 0.046725143 |
| *PCBP1* | -0.208612 | 0.066016 | -3.160031 | 0.001577521 | 0.046975543 |
| *DALRD3* | -0.318211 | 0.10071 | -3.159681 | 0.001579421 | 0.046975543 |
| *RAPGEF5* | -1.731895 | 0.548166 | -3.159438 | 0.001580737 | 0.046975543 |
| *CRYL1* | -0.954199 | 0.302314 | -3.156318 | 0.001597744 | 0.047362542 |
| *NUP88* | 0.305764 | 0.097027 | 3.151327 | 0.001625305 | 0.048059705 |
| *BPTF* | -0.25094 | 0.079757 | -3.146319 | 0.001653395 | 0.048768996 |
| *RPS9* | -0.562421 | 0.178848 | -3.144686 | 0.001662654 | 0.048920706 |
| *IMPA1* | 0.352955 | 0.112379 | 3.140751 | 0.001685151 | 0.049460221 |
| *MIDN* | 0.395264 | 0.125895 | 3.13962 | 0.00169167 | 0.04952925 |
| *MYBL2* | -1.827875 | 0.582548 | -3.137723 | 0.001702658 | 0.049728497 |
| *TXLNG* | 0.488621 | 0.155891 | 3.134369 | 0.001722239 | 0.050152605 |
| *ENSG00000270022* | 1.343041 | 0.428567 | 3.133794 | 0.001725618 | 0.050152605 |
| *SLC25A6* | -0.366016 | 0.116939 | -3.129967 | 0.001748257 | 0.050686651 |
| *SPTAN1* | -0.328742 | 0.105085 | -3.128343 | 0.001757949 | 0.050843649 |
| *TNFRSF18* | -0.909015 | 0.290709 | -3.126896 | 0.001766622 | 0.050892908 |
| *CTSA* | 0.44194 | 0.141347 | 3.126631 | 0.001768215 | 0.050892908 |
| *PRDM2* | -0.313101 | 0.100261 | -3.12285 | 0.00179109 | 0.05142677 |
| *OXA1L* | -0.256881 | 0.082285 | -3.121834 | 0.001797281 | 0.051480184 |
| *EFCAB5* | 0.606405 | 0.195011 | 3.109591 | 0.001873468 | 0.053533444 |
| *QPRT* | 1.607041 | 0.516981 | 3.108508 | 0.001880344 | 0.053601091 |
| *NEDD9* | 0.47518 | 0.152925 | 3.107272 | 0.001888228 | 0.053697062 |
| *CHPT1* | -0.457603 | 0.147309 | -3.106414 | 0.001893711 | 0.053724446 |
| *P4HTM* | -0.466559 | 0.150431 | -3.101475 | 0.001925589 | 0.054498757 |

| | | | | | |
|---|---|---|---|---|---|
| *SRD5A1* | -0.495908 | 0.16021 | -3.095362 | 0.001965726 | 0.055502567 |
| *RAP2B* | -0.490272 | 0.158447 | -3.09423 | 0.001973247 | 0.055582909 |
| *RPL29* | -0.286758 | 0.092809 | -3.089751 | 0.002003247 | 0.056294554 |
| *RASA2* | -0.401102 | 0.129947 | -3.086656 | 0.002024218 | 0.056617672 |
| *ELOA* | 0.3405 | 0.110314 | 3.086648 | 0.002024271 | 0.056617672 |
| *WARS* | 1.084846 | 0.351558 | 3.085829 | 0.002029856 | 0.05664061 |
| *AGO2* | -0.329806 | 0.106914 | -3.08479 | 0.002036959 | 0.056705684 |
| *MEF2A* | 0.416359 | 0.135034 | 3.083354 | 0.002046815 | 0.056787286 |
| *FKBP2* | 0.262469 | 0.085135 | 3.082972 | 0.002049444 | 0.056787286 |
| *ENSG00000272800* | -3.001255 | 0.975002 | -3.078204 | 0.002082526 | NA |
| *BTG2* | -0.310344 | 0.100823 | -3.078109 | 0.002083188 | 0.057588048 |
| *DHRS3* | 3.509207 | 1.140687 | 3.076397 | 0.002095189 | 0.057707179 |
| *CYBC1* | -0.26465 | 0.086047 | -3.075636 | 0.002100539 | 0.057707179 |
| *IL1B* | 2.475638 | 0.804975 | 3.07542 | 0.002102062 | 0.057707179 |
| *METTL7A* | 0.320272 | 0.104199 | 3.073662 | 0.00211449 | 0.057894003 |
| *LIPA* | 0.34807 | 0.113283 | 3.072563 | 0.002122293 | 0.057894003 |
| *SPATA2* | 0.605993 | 0.197281 | 3.071729 | 0.002128229 | 0.057894003 |
| *RPL22L1* | -0.337668 | 0.109928 | -3.071712 | 0.002128349 | 0.057894003 |
| *LYN* | -0.292462 | 0.09527 | -3.069826 | 0.002141835 | 0.058127825 |
| *NSL1* | 0.273961 | 0.089267 | 3.069 | 0.002147766 | 0.058156016 |
| *NIPAL4* | 0.903536 | 0.29455 | 3.067514 | 0.002158475 | 0.058313152 |
| *ING1* | 0.398925 | 0.13022 | 3.063478 | 0.002187801 | 0.058971419 |
| *RPS23* | -0.27427 | 0.089564 | -3.062289 | 0.00219651 | 0.059053826 |
| *VPS51* | -0.316043 | 0.103224 | -3.061706 | 0.002200795 | 0.059053826 |
| *UBA52* | -0.19416 | 0.063454 | -3.059841 | 0.002214544 | 0.059288941 |
| *NSUN2* | 0.427026 | 0.139759 | 3.055451 | 0.002247221 | 0.060028577 |
| *TOM1* | 0.420389 | 0.137767 | 3.051459 | 0.002277318 | 0.060696138 |
| *ASCC1* | 0.509801 | 0.167112 | 3.050664 | 0.002283357 | 0.060720952 |
| *EIF3F* | -0.321336 | 0.105387 | -3.049115 | 0.002295165 | 0.060898726 |
| *EMG1* | 0.263463 | 0.086429 | 3.048307 | 0.002301345 | 0.060926686 |
| *IQCD* | 0.711534 | 0.233547 | 3.046646 | 0.0023141 | 0.061128225 |
| *LINC01588* | -0.364961 | 0.119836 | -3.04551 | 0.002322863 | 0.061223674 |
| *KCNC3* | 0.706594 | 0.232312 | 3.041571 | 0.002353468 | 0.061893084 |
| *EVI2B* | 0.200371 | 0.065923 | 3.039487 | 0.002369816 | 0.062185431 |
| *FEM1A* | 0.349582 | 0.115058 | 3.038314 | 0.002379062 | 0.062290552 |
| *WDR12* | 0.515445 | 0.16991 | 3.03363 | 0.002416304 | 0.063126606 |
| *DPM2* | -0.257888 | 0.08515 | -3.028648 | 0.002456509 | 0.064036227 |
| *RASAL1* | -2.291322 | 0.756794 | -3.027669 | 0.002464477 | NA |
| *SLC1A4* | -0.89242 | 0.295217 | -3.022929 | 0.002503411 | 0.065031676 |
| *MRPS14* | 0.24243 | 0.080204 | 3.02266 | 0.002505637 | 0.065031676 |
| *GEN1* | -0.718159 | 0.237654 | -3.021869 | 0.002512196 | 0.065059855 |
| *SHISAL2A* | 0.346368 | 0.114695 | 3.019903 | 0.002528557 | 0.065341207 |
| *TARS2* | 0.496246 | 0.164396 | 3.018596 | 0.002539486 | 0.065481286 |
| *IQGAP1* | -0.271574 | 0.089992 | -3.017768 | 0.002546434 | 0.065518324 |
| *SC5D* | -0.4982 | 0.165192 | -3.015878 | 0.002562367 | 0.065540282 |
| *FKBP5* | 0.730085 | 0.242113 | 3.015472 | 0.002565795 | 0.065540282 |
| *TIMP1* | -0.521446 | 0.172938 | -3.015211 | 0.002568001 | 0.065540282 |

| | | | | | |
|---|---|---|---|---|---|
| *MICU2* | 0.31925 | 0.105885 | 3.015053 | 0.002569342 | 0.065540282 |
| *GNG2* | 0.383926 | 0.127466 | 3.011996 | 0.002595361 | 0.066062221 |
| *TSTD1* | -0.325043 | 0.108011 | -3.00935 | 0.00261807 | 0.066497865 |
| *CCDC50* | -0.329307 | 0.109481 | -3.007908 | 0.002630525 | 0.066671743 |
| *ECH1* | 0.265036 | 0.088229 | 3.003935 | 0.002665124 | 0.067302502 |
| *LPIN1* | -0.650132 | 0.216461 | -3.003463 | 0.002669257 | 0.067302502 |
| *RPL34* | -0.332353 | 0.11067 | -3.003106 | 0.002672397 | 0.067302502 |
| *AKR1B1* | 0.243817 | 0.081282 | 2.999627 | 0.002703104 | 0.067871892 |
| *ENSG00000166928* | -1.946723 | 0.649069 | -2.999253 | 0.002706425 | 0.067871892 |
| *BUD31* | 0.206981 | 0.069028 | 2.998509 | 0.002713037 | 0.067894473 |
| *WDFY4* | -0.373188 | 0.124496 | -2.997597 | 0.002721173 | 0.067955013 |
| *CCDC86* | 0.615744 | 0.205573 | 2.995266 | 0.002742058 | 0.068333001 |
| *SLAMF7* | -1.020792 | 0.341101 | -2.992637 | 0.002765783 | 0.068670447 |
| *VBP1* | 0.239724 | 0.080122 | 2.99199 | 0.00277165 | 0.068670447 |
| *TNFRSF13B* | -1.091315 | 0.364763 | -2.991849 | 0.00277293 | 0.068670447 |
| *CMTM7* | 0.425682 | 0.142442 | 2.988468 | 0.002803802 | 0.069290631 |
| *SERPINE2* | 0.871463 | 0.291881 | 2.985679 | 0.002829498 | 0.069780593 |
| *GSTK1* | -0.24018 | 0.080572 | -2.980949 | 0.002873569 | 0.070720726 |
| *ENSG00000271857* | 3.285174 | 1.104337 | 2.974794 | 0.002931858 | NA |
| *HSD17B11* | -0.233257 | 0.07843 | -2.974065 | 0.00293883 | 0.072177419 |
| *TNFRSF13C* | -0.293122 | 0.098598 | -2.972904 | 0.002949971 | 0.072301662 |
| *IL2RA* | -0.987686 | 0.33249 | -2.970573 | 0.002972449 | 0.072702671 |
| *GSTP1* | 0.255974 | 0.086271 | 2.967088 | 0.003006345 | 0.073380753 |
| *LSM2* | -0.24628 | 0.08306 | -2.965066 | 0.003026179 | 0.073686507 |
| *SNW1* | 0.286778 | 0.096744 | 2.964313 | 0.003033595 | 0.073686507 |
| *NME4* | -0.574872 | 0.193981 | -2.963555 | 0.003041076 | 0.073686507 |
| *BCCIP* | 0.32558 | 0.109871 | 2.963293 | 0.003043667 | 0.073686507 |
| *HOXB3* | 1.886742 | 0.637197 | 2.961001 | 0.003066407 | 0.074086142 |
| *EIF6* | 0.209246 | 0.070696 | 2.959812 | 0.003078267 | 0.074221825 |
| *LAIR1* | 0.695603 | 0.235313 | 2.956078 | 0.003115782 | 0.074974296 |
| *EIF4B* | -0.246113 | 0.083297 | -2.954647 | 0.003130265 | 0.075170622 |
| *SYNE3* | -0.47506 | 0.160836 | -2.953689 | 0.00314 | 0.075252387 |
| *LDHB* | -0.247669 | 0.083883 | -2.952531 | 0.003151806 | 0.075383343 |
| *TRIM25* | 0.518148 | 0.175691 | 2.949207 | 0.003185901 | 0.076045802 |
| *CHCHD6* | -0.701889 | 0.238191 | -2.946745 | 0.003211383 | 0.076500424 |
| *HIVEP2* | 0.519871 | 0.176513 | 2.945223 | 0.003227221 | 0.076723944 |
| *PPIF* | -0.547311 | 0.185878 | -2.944471 | 0.003235075 | 0.076757158 |
| *RPS18* | -0.30767 | 0.104552 | -2.942743 | 0.003253179 | 0.07703294 |
| *SPA17* | 1.700093 | 0.578064 | 2.941009 | 0.003271445 | 0.07716779 |
| *DNAJA2* | 0.263263 | 0.089528 | 2.940584 | 0.003275946 | 0.07716779 |
| *RIMKLB* | -0.458217 | 0.155838 | -2.940341 | 0.003278516 | 0.07716779 |
| *NEK3* | -1.524734 | 0.518662 | -2.939743 | 0.003284841 | 0.07716779 |
| *ARL4C* | -0.68056 | 0.23167 | -2.937627 | 0.003307344 | 0.077543188 |
| *IGBP1* | -0.355638 | 0.121191 | -2.934522 | 0.003340614 | 0.078169062 |
| *ADD1* | 0.290829 | 0.099134 | 2.933687 | 0.003349613 | 0.078225644 |
| *STAT4* | -0.718723 | 0.245243 | -2.930654 | 0.003382498 | 0.07879336 |
| *RPS28* | -0.271987 | 0.092821 | -2.930224 | 0.00338718 | 0.07879336 |

| | | | | | |
|---|---|---|---|---|---|
| *HGSNAT* | -0.593403 | 0.202716 | -2.927259 | 0.003419639 | 0.079364194 |
| *EPHB6* | -0.504089 | 0.172234 | -2.926765 | 0.003425072 | 0.079364194 |
| *NDUFB3* | 0.228263 | 0.078024 | 2.925537 | 0.003438628 | 0.079464836 |
| *DUSP4* | -1.714408 | 0.58609 | -2.925161 | 0.003442785 | 0.079464836 |
| *RPL19* | -0.219826 | 0.075184 | -2.923819 | 0.003457654 | 0.079653365 |
| *PLD4* | -0.896546 | 0.306795 | -2.922293 | 0.003474641 | 0.079889868 |
| *ZDHHC21* | -0.508914 | 0.174349 | -2.918943 | 0.003512207 | 0.080487988 |
| *UTP3* | 0.30794 | 0.105503 | 2.918766 | 0.003514198 | 0.080487988 |
| *ATF4* | 0.294738 | 0.101014 | 2.917809 | 0.003524997 | 0.080580086 |
| *PDE3B* | -0.76197 | 0.261281 | -2.916283 | 0.003542295 | 0.08082008 |
| *TOB1* | -0.435213 | 0.149383 | -2.913411 | 0.00357504 | 0.08136333 |
| *CPEB2* | -1.273 | 0.437074 | -2.912549 | 0.003584923 | 0.08136333 |
| *WBP2* | 0.330261 | 0.113398 | 2.912399 | 0.00358664 | 0.08136333 |
| *GNAZ* | -0.894725 | 0.307281 | -2.911746 | 0.003594143 | 0.081378247 |
| *ENSG00000274213* | 0.999664 | 0.343473 | 2.910457 | 0.003609006 | 0.08155943 |
| *CAMK2G* | -0.409461 | 0.140948 | -2.905045 | 0.003672001 | 0.082825562 |
| *PSMB4* | 0.282266 | 0.09722 | 2.90338 | 0.00369158 | 0.083109491 |
| *TMEM140* | 0.538252 | 0.185453 | 2.902365 | 0.003703571 | 0.083221821 |
| *SLC35B2* | -0.499909 | 0.172302 | -2.901348 | 0.003715613 | 0.083257706 |
| *TMEM175* | 0.615363 | 0.212118 | 2.901047 | 0.003719176 | 0.083257706 |
| *UBE2T* | 0.507186 | 0.174946 | 2.899107 | 0.003742271 | 0.083559092 |
| *TAF7* | 0.177683 | 0.061297 | 2.898736 | 0.003746698 | 0.083559092 |
| *SSBP1* | 0.184829 | 0.063903 | 2.892352 | 0.003823692 | 0.085116527 |
| *C14orf28* | -0.444455 | 0.153756 | -2.890645 | 0.003844526 | 0.085420342 |
| *ABHD14B* | -0.249603 | 0.086421 | -2.88823 | 0.003874164 | 0.085755467 |
| *ZDHHC23* | -0.67241 | 0.232829 | -2.887999 | 0.003877012 | 0.085755467 |
| *ENSG00000259436* | -0.460383 | 0.159431 | -2.887655 | 0.003881252 | 0.085755467 |
| *ENSA* | 0.184622 | 0.063972 | 2.885982 | 0.003901945 | 0.085940265 |
| *FMNL3* | 0.395544 | 0.137065 | 2.88581 | 0.003904075 | 0.085940265 |
| *RDH10* | 2.048292 | 0.710157 | 2.884282 | 0.003923072 | NA |
| *COX6A1* | 0.208533 | 0.072358 | 2.881954 | 0.003952172 | 0.086838214 |
| *COMMD6* | -0.161615 | 0.056107 | -2.880502 | 0.003970423 | 0.086972109 |
| *GADD45GIP1* | 0.201776 | 0.070054 | 2.880306 | 0.003972899 | 0.086972109 |
| *SNRPC* | 0.224752 | 0.078232 | 2.872876 | 0.004067537 | 0.088880159 |
| *DDX21* | 0.228829 | 0.079729 | 2.87007 | 0.00410381 | 0.089360257 |
| *IRF1* | 0.329123 | 0.114687 | 2.869756 | 0.004107887 | 0.089360257 |
| *FLOT1* | 0.27693 | 0.09652 | 2.86914 | 0.004115892 | 0.089360257 |
| *SPCS2* | 0.188049 | 0.065548 | 2.868857 | 0.004119578 | 0.089360257 |
| *CMC2* | 0.301254 | 0.105059 | 2.867484 | 0.004137492 | 0.089585377 |
| *RABAC1* | 0.220316 | 0.076996 | 2.861404 | 0.004217691 | 0.090999671 |
| *TMEM135* | 0.944355 | 0.330036 | 2.861372 | 0.004218122 | 0.090999671 |
| *TACO1* | 0.392032 | 0.137146 | 2.858499 | 0.004256511 | 0.091513746 |
| *POP4* | 0.288604 | 0.100966 | 2.858436 | 0.004257348 | 0.091513746 |
| *CD5* | -1.085725 | 0.380142 | -2.8561 | 0.004288804 | 0.091922953 |
| *CHEK1* | -1.29734 | 0.454271 | -2.855874 | 0.004291852 | 0.091922953 |
| *HIVEP3* | 0.774619 | 0.27137 | 2.854472 | 0.004310843 | 0.092163645 |
| *GPANK1* | 0.248032 | 0.087011 | 2.850594 | 0.004363772 | 0.093008659 |

| | | | | | |
|---|---|---|---|---|---|
| *SMS* | 0.269483 | 0.094547 | 2.850244 | 0.004368567 | 0.093008659 |
| *HSPA8* | -0.368022 | 0.129137 | -2.849861 | 0.00437384 | 0.093008659 |
| *EEF1E1* | 0.274095 | 0.096366 | 2.844306 | 0.004450828 | 0.094476782 |
| *FBXO15* | -0.897055 | 0.315792 | -2.840652 | 0.004502137 | 0.095395546 |
| *CLYBL* | -0.751729 | 0.264708 | -2.839837 | 0.004513665 | 0.095469641 |
| *CEP170* | 0.339907 | 0.119745 | 2.838601 | 0.004531176 | 0.095669775 |
| *KLHL14* | -0.525911 | 0.185549 | -2.834354 | 0.00459184 | 0.096778719 |
| *SDHD* | 0.246952 | 0.087148 | 2.833714 | 0.004601049 | 0.096801185 |
| *PABPN1* | -0.222454 | 0.078551 | -2.831963 | 0.004626324 | 0.097160976 |
| *CEP68* | -0.372755 | 0.131661 | -2.831177 | 0.004637704 | 0.097228203 |
| *GTDC1* | -0.648396 | 0.229187 | -2.829118 | 0.004667656 | 0.097683841 |
| *RPL32* | -0.265698 | 0.093953 | -2.827988 | 0.004684152 | 0.0978568 |
| *IKZF2* | -0.706443 | 0.24988 | -2.827125 | 0.004696802 | 0.097948928 |
| *SMIM20* | 0.341121 | 0.120711 | 2.825941 | 0.004714193 | 0.098139435 |
| *RHOBTB2* | -1.200366 | 0.425214 | -2.82297 | 0.004758098 | 0.098880272 |
| *C12orf42* | -0.483092 | 0.171175 | -2.822211 | 0.004769375 | 0.098937553 |
| *RPL30* | -0.237906 | 0.084325 | -2.821294 | 0.004783041 | 0.098937553 |
| *ENSG00000230709* | -0.943923 | 0.334613 | -2.820938 | 0.004788344 | 0.098937553 |
| *MADCAM1* | -1.121647 | 0.39767 | -2.82055 | 0.004794147 | 0.098937553 |
| *HSBP1* | 0.246655 | 0.087535 | 2.817777 | 0.004835736 | 0.099622867 |
| *ENSG00000277511* | -0.544062 | 0.193253 | -2.815283 | 0.004873432 | 0.100225764 |
| *FDFT1* | 0.305941 | 0.108705 | 2.814413 | 0.004886638 | 0.100323771 |
| *PRDX6* | 0.209221 | 0.074399 | 2.812146 | 0.004921217 | 0.100730989 |
| *ENO1* | 0.309995 | 0.110299 | 2.810501 | 0.004946441 | 0.100730989 |
| *REM2* | 0.870621 | 0.309819 | 2.810095 | 0.004952686 | 0.100730989 |
| *ENSG00000253535* | 0.722683 | 0.257188 | 2.809943 | 0.004955034 | 0.100730989 |
| *TECPR1* | 0.567744 | 0.202058 | 2.809803 | 0.004957181 | 0.100730989 |
| *RPL41* | -0.246876 | 0.087872 | -2.809493 | 0.004961962 | 0.100730989 |
| *ITGAE* | 0.277516 | 0.098787 | 2.809245 | 0.004965791 | 0.100730989 |
| *ZNF785* | 0.400138 | 0.142801 | 2.80207 | 0.005077584 | 0.10282324 |
| *NFATC3* | 0.326743 | 0.116637 | 2.801377 | 0.005088502 | 0.102841123 |
| *RPS6* | -0.296592 | 0.105898 | -2.800721 | 0.005098855 | 0.102841123 |
| *ACP1* | 0.173264 | 0.061872 | 2.800369 | 0.005104422 | 0.102841123 |
| *HHIP.AS1* | -2.005427 | 0.716714 | -2.798084 | 0.005140679 | NA |
| *PADI4* | -1.219189 | 0.436038 | -2.796062 | 0.005172944 | 0.104045314 |
| *SCAF4* | -0.283876 | 0.101572 | -2.794816 | 0.005192926 | 0.104084557 |
| *SHQ1* | 0.599835 | 0.21463 | 2.794738 | 0.005194185 | 0.104084557 |
| *GPR18* | 0.588343 | 0.210551 | 2.794304 | 0.005201163 | 0.104084557 |
| *PDIA3* | 0.309279 | 0.110732 | 2.793039 | 0.005221546 | 0.104316828 |
| *MOSPD1* | 0.465103 | 0.166656 | 2.790799 | 0.005257807 | 0.104760154 |
| *UBB* | 0.261478 | 0.0937 | 2.79058 | 0.005261362 | 0.104760154 |
| *POU2F2* | -0.20246 | 0.072612 | -2.788236 | 0.005299593 | 0.105238241 |
| *ARHGAP27* | 0.362506 | 0.130023 | 2.788023 | 0.00530308 | 0.105238241 |
| *ATF3* | 0.723982 | 0.25978 | 2.7869 | 0.005321497 | 0.105427719 |
| *GPR137* | 0.227145 | 0.081523 | 2.786248 | 0.005332198 | 0.105463957 |
| *ENSG00000276136* | -0.752841 | 0.270413 | -2.784046 | 0.005368547 | 0.106006506 |
| *EPB41L4A.AS1* | -0.226878 | 0.081511 | -2.783416 | 0.005378987 | 0.10603651 |

| | | | | | |
|---|---|---|---|---|---|
| *ZMYM3* | -0.526635 | 0.189312 | -2.781834 | 0.005405273 | 0.106378279 |
| *TNIP1* | 0.225298 | 0.081109 | 2.777741 | 0.005473831 | 0.107549474 |
| *TXNDC12* | 0.254275 | 0.091776 | 2.770611 | 0.005595115 | 0.109751037 |
| *ANXA7* | 0.294487 | 0.106367 | 2.768596 | 0.005629842 | 0.110250303 |
| *HNRNPC* | 0.217481 | 0.078618 | 2.766286 | 0.005669883 | 0.110851798 |
| *DAD1* | 0.227143 | 0.082151 | 2.76496 | 0.005692974 | 0.111012271 |
| *SLFN11* | -0.716388 | 0.259116 | -2.764743 | 0.005696768 | 0.111012271 |
| *TTC39C* | -0.557497 | 0.201724 | -2.763657 | 0.005715766 | 0.111200181 |
| *ZW10* | 0.640998 | 0.232029 | 2.762584 | 0.005734586 | 0.111308981 |
| *PLEK* | 1.002769 | 0.363024 | 2.762271 | 0.005740086 | 0.111308981 |
| *ANKRD37* | 0.476764 | 0.172778 | 2.759402 | 0.005790721 | 0.112107979 |
| *ZNF652* | -0.381399 | 0.138285 | -2.758072 | 0.005814337 | 0.112284752 |
| *TSPAN14* | -0.349589 | 0.126763 | -2.757824 | 0.005818744 | 0.112284752 |
| *ITGB2* | 0.642127 | 0.233004 | 2.75587 | 0.005853624 | 0.11277476 |
| *RSRC1* | -0.260374 | 0.094573 | -2.753162 | 0.005902264 | 0.113173044 |
| *KATNBL1* | -0.26527 | 0.096354 | -2.753092 | 0.005903529 | 0.113173044 |
| *ENSG00000274184* | -0.680999 | 0.247441 | -2.752166 | 0.005920246 | 0.113173044 |
| *KBTBD6* | -0.400964 | 0.14572 | -2.751606 | 0.005930376 | 0.113173044 |
| *ENSG00000235609* | 1.456059 | 0.529191 | 2.751482 | 0.005932631 | 0.113173044 |
| *RPL38* | -0.233734 | 0.084959 | -2.751125 | 0.005939103 | 0.113173044 |
| *EBLN2* | 0.43511 | 0.158186 | 2.750627 | 0.00594814 | 0.113173044 |
| *FBXO11* | -0.358919 | 0.130492 | -2.750499 | 0.005950463 | 0.113173044 |
| *TEX264* | 0.252923 | 0.092012 | 2.748808 | 0.005981247 | 0.113479027 |
| *CWC15* | 0.212266 | 0.077228 | 2.748567 | 0.005985644 | 0.113479027 |
| *LGALS3* | -0.64217 | 0.233757 | -2.747168 | 0.006011236 | 0.113571799 |
| *NECAP2* | 0.202153 | 0.073589 | 2.74705 | 0.006013393 | 0.113571799 |
| *HNRNPH3* | 0.212001 | 0.077183 | 2.746734 | 0.0060192 | 0.113571799 |
| *HMGN1* | -0.191737 | 0.069866 | -2.744342 | 0.006063239 | 0.114221439 |
| *CD83* | -0.200914 | 0.073228 | -2.743665 | 0.006075754 | 0.114276095 |
| *SPIDR* | -0.438614 | 0.159942 | -2.742336 | 0.006100395 | 0.114558288 |
| *FANCG* | -0.667994 | 0.24385 | -2.739369 | 0.006155723 | 0.115414954 |
| *CAMKMT* | -0.504749 | 0.184324 | -2.738384 | 0.006174187 | 0.11551968 |
| *MAGOH* | 0.211515 | 0.07726 | 2.737703 | 0.006186996 | 0.11551968 |
| *MDH1* | 0.315649 | 0.115305 | 2.737519 | 0.006190463 | 0.11551968 |
| *ARHGEF40* | -1.997523 | 0.729908 | -2.736679 | 0.00620628 | NA |
| *IL6* | -1.471908 | 0.538368 | -2.734018 | 0.006256656 | 0.116376659 |
| *SERPINB1* | 0.316514 | 0.115782 | 2.733708 | 0.006262564 | 0.116376659 |
| *PIM2* | -0.208933 | 0.076437 | -2.733393 | 0.006268547 | 0.116376659 |
| *ZNF277* | -0.331538 | 0.121337 | -2.732384 | 0.006287781 | 0.116376659 |
| *LRPAP1* | 0.246572 | 0.090253 | 2.732 | 0.006295108 | 0.116376659 |
| *ZMIZ1* | -0.976177 | 0.357401 | -2.731318 | 0.006308154 | 0.116376659 |
| *RNF41* | -0.223343 | 0.081773 | -2.731242 | 0.006309612 | 0.116376659 |
| *NFKBIB* | 0.331435 | 0.121361 | 2.730976 | 0.006314709 | 0.116376659 |
| *RNF167* | 0.222799 | 0.081708 | 2.726775 | 0.006395664 | 0.117686157 |
| *LTA4H* | -0.303548 | 0.11138 | -2.725333 | 0.006423666 | 0.118018736 |
| *CKLF* | -0.471538 | 0.173067 | -2.724589 | 0.00643816 | 0.118102486 |
| *MRM2* | 0.262039 | 0.096216 | 2.723449 | 0.006460416 | 0.118328148 |

| | | | | | |
|---|---|---|---|---|---|
| *MAN1B1* | -0.391724 | 0.143944 | -2.72137 | 0.006501193 | 0.1187172 |
| *EEF1D* | -0.266816 | 0.098046 | -2.721348 | 0.006501632 | 0.1187172 |
| *IRF4* | 0.425386 | 0.156661 | 2.715336 | 0.006620859 | 0.120397354 |
| *PDIA6* | 0.253323 | 0.093295 | 2.715297 | 0.006621632 | 0.120397354 |
| *SEC61B* | -0.272077 | 0.100206 | -2.715177 | 0.006624032 | 0.120397354 |
| *SNRPB* | 0.211932 | 0.07811 | 2.713262 | 0.00666245 | 0.120910748 |
| *TRIM33* | -0.302707 | 0.111688 | -2.710296 | 0.006722325 | 0.121811402 |
| *BANK1* | -0.337276 | 0.124513 | -2.708763 | 0.006753456 | 0.122189241 |
| *PSME1* | 0.152106 | 0.056247 | 2.704242 | 0.006846037 | 0.123676051 |
| *SAP18* | 0.167265 | 0.061867 | 2.703597 | 0.006859352 | 0.123728559 |
| *ZKSCAN1* | -0.253914 | 0.093974 | -2.70195 | 0.006893404 | 0.124154389 |
| *PRPF19* | 0.38301 | 0.141788 | 2.701287 | 0.006907177 | 0.124214232 |
| *SERPINB6* | 0.450539 | 0.166836 | 2.700484 | 0.006923872 | 0.124326384 |
| *TANK* | 0.177057 | 0.065583 | 2.699751 | 0.006939129 | 0.124412404 |
| *ENSG00000176320* | 2.490283 | 0.922711 | 2.698877 | 0.00695739 | NA |
| *CCDC28A* | 0.242878 | 0.090062 | 2.696794 | 0.007001047 | 0.125333494 |
| *IK* | 0.205563 | 0.076313 | 2.69368 | 0.007066793 | 0.126320259 |
| *PTBP1* | 0.231591 | 0.08604 | 2.691678 | 0.007109357 | 0.126890273 |
| *TSC22D1* | -0.524518 | 0.194923 | -2.690901 | 0.007125928 | 0.126995355 |
| *MSI2* | -0.335271 | 0.124621 | -2.690319 | 0.007138383 | 0.127026875 |
| *TTC21A* | -0.384677 | 0.143181 | -2.686648 | 0.007217291 | 0.128239077 |
| *FCHSD2* | -0.373494 | 0.139045 | -2.686131 | 0.007228474 | 0.128246068 |
| *RASGRF1* | -1.522952 | 0.567407 | -2.684055 | 0.007273514 | 0.128736724 |
| *GRAP* | 0.457636 | 0.170514 | 2.683859 | 0.007277789 | 0.128736724 |
| *CENPT* | -0.321645 | 0.119946 | -2.681587 | 0.00732739 | 0.129361403 |
| *EIF3L* | -0.347213 | 0.129497 | -2.681245 | 0.007334869 | 0.129361403 |
| *EIF3H* | -0.225828 | 0.08428 | -2.679484 | 0.007373566 | 0.129851236 |
| *FBXO28* | 0.410914 | 0.153526 | 2.676521 | 0.007439087 | 0.130694141 |
| *ESR2* | -0.792838 | 0.296241 | -2.676326 | 0.00744342 | 0.130694141 |
| *HIST1H4H* | 0.597245 | 0.22326 | 2.675113 | 0.00747041 | 0.130940296 |
| *CGAS* | 0.335266 | 0.125347 | 2.674706 | 0.00747947 | 0.130940296 |
| *DHRS1* | -0.304448 | 0.113904 | -2.672848 | 0.007521038 | 0.131421773 |
| *MYL2* | 2.924506 | 1.094339 | 2.672395 | 0.007531189 | NA |
| *CNIH4* | 0.259232 | 0.097029 | 2.67169 | 0.007547029 | 0.131421773 |
| *ZNF506* | 0.361326 | 0.135249 | 2.671557 | 0.007550024 | 0.131421773 |
| *GPR132* | 0.362305 | 0.135618 | 2.671505 | 0.007551196 | 0.131421773 |
| *UBXN1* | -0.249598 | 0.093475 | -2.67022 | 0.007580159 | 0.131732965 |
| *CRIPT* | 0.281271 | 0.105371 | 2.669327 | 0.007600331 | 0.131796502 |
| *NUP62* | 0.288274 | 0.108005 | 2.669078 | 0.00760599 | 0.131796502 |
| *IGHA1* | -0.458329 | 0.171769 | -2.668289 | 0.007623865 | 0.131913957 |
| *JAML* | -2.462621 | 0.923713 | -2.666002 | 0.007675912 | 0.132621468 |
| *RNF181* | 0.221979 | 0.083296 | 2.664957 | 0.007699824 | 0.132841514 |
| *INKA1* | -0.387966 | 0.14566 | -2.663499 | 0.007733256 | 0.13312008 |
| *ANAPC10* | 0.377426 | 0.141715 | 2.663277 | 0.007738368 | 0.13312008 |
| *CCNG1* | -0.219009 | 0.082354 | -2.659357 | 0.007828985 | 0.134484312 |
| *CXCR5* | 0.265282 | 0.099821 | 2.657567 | 0.007870699 | 0.134792798 |
| *FLNA* | -0.315869 | 0.118858 | -2.657522 | 0.007871735 | 0.134792798 |

| | | | | | |
|---|---|---|---|---|---|
| *PURA* | -0.360907 | 0.135828 | -2.657091 | 0.007881814 | 0.134792798 |
| *LY86* | -0.452301 | 0.170277 | -2.656266 | 0.007901136 | 0.134792798 |
| *MANF* | 0.301765 | 0.11361 | 2.656159 | 0.007903641 | 0.134792798 |
| *LINC00342* | -0.789288 | 0.297328 | -2.654602 | 0.007940197 | 0.13522224 |
| *RPL36* | -0.152903 | 0.057624 | -2.653466 | 0.007966989 | 0.1354844 |
| *VIM* | -0.426281 | 0.160767 | -2.651549 | 0.008012351 | 0.13606116 |
| *SRSF8* | 0.245487 | 0.092634 | 2.650081 | 0.008047255 | 0.136274599 |
| *AARS* | 0.379159 | 0.143076 | 2.650056 | 0.008047848 | 0.136274599 |
| *PABPC1* | -0.248246 | 0.093765 | -2.647532 | 0.008108178 | 0.137100874 |
| *CUL1* | 0.355378 | 0.134271 | 2.646729 | 0.008127453 | 0.137231583 |
| *RPL13A* | -0.253197 | 0.095718 | -2.645243 | 0.008163226 | 0.137474913 |
| *SF3B6* | 0.184311 | 0.069678 | 2.64517 | 0.008164994 | 0.137474913 |
| *DAPK2* | 0.715183 | 0.270663 | 2.642338 | 0.008233585 | 0.138241999 |
| *DNAJA1* | 0.263185 | 0.099603 | 2.642329 | 0.008233813 | 0.138241999 |
| *ECSIT* | 0.239639 | 0.090778 | 2.63984 | 0.008294516 | 0.139064758 |
| *DAAM1* | -0.511532 | 0.193904 | -2.638072 | 0.008337892 | 0.139595103 |
| *POLR2C* | 0.233272 | 0.088463 | 2.636946 | 0.008365617 | 0.139862287 |
| *RPS14* | -0.28326 | 0.107487 | -2.635309 | 0.008406064 | 0.14030083 |
| *PRCD* | -0.818879 | 0.310789 | -2.634836 | 0.008417794 | 0.14030083 |
| *EAPP* | 0.266378 | 0.101121 | 2.634239 | 0.008432621 | 0.14030083 |
| *ZBTB17* | 0.425062 | 0.161376 | 2.633979 | 0.008439059 | 0.14030083 |
| *DPP8* | 0.271766 | 0.103198 | 2.633448 | 0.008452289 | 0.140324527 |
| *LGALS1* | -0.71027 | 0.269871 | -2.631892 | 0.008491097 | 0.140772196 |
| *RIMS3* | -1.462898 | 0.55613 | -2.630498 | 0.008525991 | 0.141019768 |
| *RPL39* | -0.240046 | 0.09126 | -2.630348 | 0.008529756 | 0.141019768 |
| *MKNK2* | -0.41495 | 0.157898 | -2.627968 | 0.008589656 | 0.14174608 |
| *RASGRP1* | -0.670919 | 0.25533 | -2.627656 | 0.008597537 | 0.14174608 |
| *YTHDF2* | 0.284465 | 0.108326 | 2.626019 | 0.008638997 | 0.142232345 |
| *SLC39A7* | 0.368201 | 0.140303 | 2.624326 | 0.008682065 | 0.142681685 |
| *DNAJC4* | -0.300978 | 0.114702 | -2.624003 | 0.008690295 | 0.142681685 |
| *DNAAF4* | -0.686537 | 0.261762 | -2.622755 | 0.008722187 | 0.143007768 |
| *MORF4L1* | 0.14004 | 0.053429 | 2.621041 | 0.008766174 | 0.143530999 |
| *MPHOSPH9* | 0.388393 | 0.148304 | 2.618895 | 0.008821519 | 0.144110574 |
| *RCBTB1* | -0.548472 | 0.209442 | -2.618728 | 0.008825818 | 0.144110574 |
| *ASB16.AS1* | -0.454821 | 0.17385 | -2.616167 | 0.00889231 | 0.1449971 |
| *LRWD1* | 0.443358 | 0.169525 | 2.615296 | 0.008915004 | 0.145168023 |
| *PCBP4* | -0.576958 | 0.220665 | -2.614635 | 0.008932281 | 0.145250369 |
| *MED11* | 0.200789 | 0.076832 | 2.613363 | 0.008965614 | 0.145593249 |
| *PDLIM2* | -0.457294 | 0.17508 | -2.611912 | 0.009003741 | 0.146012926 |
| *MAT2B* | 0.240793 | 0.092229 | 2.610811 | 0.009032781 | 0.146284288 |
| *XBP1* | 0.388073 | 0.148695 | 2.609862 | 0.009057877 | 0.146318104 |
| *GOLGB1* | -0.226719 | 0.086872 | -2.609801 | 0.009059487 | 0.146318104 |
| *B4GALT1* | -0.464908 | 0.178261 | -2.608023 | 0.00910669 | 0.146717936 |
| *PSMD4* | 0.234063 | 0.089769 | 2.607379 | 0.009123818 | 0.146717936 |
| *NDUFAB1* | 0.179358 | 0.068794 | 2.607176 | 0.00912925 | 0.146717936 |
| *PCDH9* | -0.681491 | 0.261407 | -2.607012 | 0.009133614 | 0.146717936 |
| *MCM3* | 0.341512 | 0.131101 | 2.604956 | 0.009188595 | 0.147401931 |

| | | | | | |
|---|---|---|---|---|---|
| *SLC25A13* | -0.481118 | 0.184776 | -2.603782 | 0.009220128 | 0.147708443 |
| *TOR3A* | 0.234973 | 0.090272 | 2.602954 | 0.009242444 | 0.147866671 |
| *PSMB3* | 0.207689 | 0.079818 | 2.602027 | 0.009267455 | 0.148067528 |
| *RDH5* | -0.421758 | 0.162189 | -2.600407 | 0.009311316 | 0.148568616 |
| *FAM133B* | 0.149455 | 0.057605 | 2.594498 | 0.00947293 | 0.150944663 |
| *CAPRIN2* | -0.491516 | 0.189582 | -2.592633 | 0.009524433 | 0.151504696 |
| *SQSTM1* | 0.312109 | 0.120415 | 2.591946 | 0.009543462 | 0.151504696 |
| *ABCG1* | 0.747906 | 0.288562 | 2.591844 | 0.009546313 | 0.151504696 |
| *RPL35* | -0.236621 | 0.091347 | -2.59036 | 0.009587552 | 0.15195631 |
| *AHSA2P* | -0.3981 | 0.153882 | -2.587052 | 0.009680098 | 0.153218811 |
| *NDUFB4* | 0.148902 | 0.057589 | 2.585571 | 0.009721783 | 0.153559482 |
| *MIR497HG* | 1.214799 | 0.469874 | 2.58537 | 0.009727458 | 0.153559482 |
| *SLC25A25.AS1* | 0.602762 | 0.233248 | 2.584208 | 0.009760303 | 0.153873638 |
| *FCHO1* | 0.531592 | 0.205781 | 2.58329 | 0.009786315 | 0.15404053 |
| *HAX1* | 0.200348 | 0.077567 | 2.58292 | 0.009796807 | 0.15404053 |
| *GPKOW* | 0.449726 | 0.174307 | 2.580083 | 0.009877658 | 0.155106626 |
| *TMX1* | 0.25956 | 0.100675 | 2.578188 | 0.009932004 | 0.15575426 |
| *TAF4B* | 0.62023 | 0.240854 | 2.575129 | 0.010020266 | 0.156931366 |
| *PPP1R11* | 0.211273 | 0.082115 | 2.572882 | 0.010085562 | 0.157495146 |
| *PSMC6* | 0.266102 | 0.103454 | 2.572179 | 0.010106077 | 0.157495146 |
| *CHMP2B* | 0.232795 | 0.090517 | 2.571843 | 0.010115888 | 0.157495146 |
| *PAXX* | 0.463056 | 0.180051 | 2.571797 | 0.010117219 | 0.157495146 |
| *SIPA1L1* | -0.308187 | 0.119842 | -2.571608 | 0.010122734 | 0.157495146 |
| *CEP19* | 0.443037 | 0.172319 | 2.571032 | 0.010139582 | 0.157495146 |
| *SLC39A1* | 0.226381 | 0.088062 | 2.570711 | 0.01014901 | 0.157495146 |
| *NDUFA4* | 0.133212 | 0.051968 | 2.563367 | 0.010366242 | 0.160656479 |
| *CHMP1B* | 0.210397 | 0.082133 | 2.561671 | 0.010417009 | 0.161233047 |
| *RPS21* | -0.226139 | 0.088297 | -2.561101 | 0.010434102 | 0.161287614 |
| *NINJ1* | -0.568232 | 0.222065 | -2.558851 | 0.010501876 | 0.162124419 |
| *TPRA1* | 0.421008 | 0.164571 | 2.558218 | 0.010521004 | 0.162209051 |
| *CASZ1* | -0.638242 | 0.249593 | -2.557131 | 0.010553957 | 0.162218692 |
| *IKZF3* | -0.247823 | 0.096924 | -2.55689 | 0.010561272 | 0.162218692 |
| *RAB4B* | -0.262665 | 0.10273 | -2.556847 | 0.01056257 | 0.162218692 |
| *ENSG00000245869* | 0.592735 | 0.231961 | 2.555328 | 0.010608776 | 0.162718094 |
| *UVSSA* | -0.981497 | 0.384245 | -2.554355 | 0.010638459 | 0.162779798 |
| *ARID1B* | -0.274844 | 0.1076 | -2.554299 | 0.010640187 | 0.162779798 |
| *GPT2* | 0.759581 | 0.297652 | 2.551905 | 0.010713568 | 0.163691756 |
| *TMEM41B* | 0.336225 | 0.13187 | 2.549664 | 0.010782686 | 0.164536311 |
| *ZNF250* | -0.519641 | 0.20391 | -2.548382 | 0.010822394 | 0.164780363 |
| *GAB2* | -0.630348 | 0.247378 | -2.548122 | 0.010830469 | 0.164780363 |
| *ENSG00000257275* | -0.462642 | 0.181584 | -2.547806 | 0.010840266 | 0.164780363 |
| *ENSG00000238045* | 0.589099 | 0.231283 | 2.547093 | 0.010862452 | 0.164906726 |
| *FHIT* | -0.64913 | 0.255063 | -2.544982 | 0.010928344 | 0.165695445 |
| *APEX1* | -0.244665 | 0.09616 | -2.544343 | 0.01094834 | 0.165787162 |
| *SORBS3* | -0.76852 | 0.302125 | -2.543719 | 0.010967915 | 0.165872269 |
| *PRDX5* | 0.214936 | 0.084521 | 2.542981 | 0.010991112 | 0.166011884 |
| *KBTBD7* | -0.360935 | 0.141969 | -2.54235 | 0.011010973 | 0.166100807 |

| | | | | | |
|---|---|---|---|---|---|
| *DPY30* | 0.193387 | 0.076099 | 2.541245 | 0.011045849 | 0.166415718 |
| *EIF2S3* | -0.214612 | 0.084492 | -2.540028 | 0.011084354 | 0.166580232 |
| *ZNF780A* | 0.36155 | 0.142355 | 2.539781 | 0.011092178 | 0.166580232 |
| *JUND* | -0.295547 | 0.116376 | -2.539572 | 0.011098809 | 0.166580232 |
| *SLCO4A1* | -0.581439 | 0.22904 | -2.538591 | 0.01112998 | 0.166703408 |
| *BRF2* | 0.504368 | 0.198693 | 2.538431 | 0.011135064 | 0.166703408 |
| *RPS7* | -0.260886 | 0.102815 | -2.537427 | 0.011167053 | 0.166972025 |
| *CARS2* | 0.370402 | 0.146045 | 2.536214 | 0.011205826 | 0.167341267 |
| *DNASE2* | 0.272774 | 0.107712 | 2.532434 | 0.011327368 | 0.168851097 |
| *MRM3* | 0.396128 | 0.156437 | 2.532187 | 0.011335339 | 0.168851097 |
| *SIDT2* | -0.438522 | 0.173218 | -2.53162 | 0.01135371 | 0.168913086 |
| *HMGXB4* | 0.244336 | 0.096547 | 2.530745 | 0.011382072 | 0.169123369 |
| *TMX3* | -0.402711 | 0.159191 | -2.529743 | 0.011414603 | 0.169394986 |
| *MMD* | -0.509315 | 0.201407 | -2.528785 | 0.011445814 | 0.169646366 |
| *LIN7B* | -0.568516 | 0.224927 | -2.527558 | 0.01148588 | 0.17002821 |
| *IGKC* | 0.34177 | 0.135382 | 2.524486 | 0.011586769 | 0.171308366 |
| *LY6E* | -0.297529 | 0.118027 | -2.520866 | 0.011706649 | 0.172865759 |
| *MT.ATP8* | -0.515926 | 0.204851 | -2.518542 | 0.011784167 | 0.17362088 |
| *THEMIS2* | -0.791189 | 0.314188 | -2.518204 | 0.011795493 | 0.17362088 |
| *CXCR4* | 0.20855 | 0.082823 | 2.518022 | 0.011801604 | 0.17362088 |
| *TRA2A* | -0.210958 | 0.083804 | -2.517283 | 0.01182638 | 0.17365076 |
| *CLCN3* | 0.361581 | 0.14365 | 2.51709 | 0.011832852 | 0.17365076 |
| *SERPINB9* | -0.266946 | 0.106116 | -2.515603 | 0.011882908 | 0.174170325 |
| *APIP* | 0.261792 | 0.10409 | 2.51505 | 0.011901554 | 0.174228791 |
| *PHF12* | -0.352557 | 0.140243 | -2.513891 | 0.011940739 | 0.174587415 |
| *C15orf65* | -0.965435 | 0.384562 | -2.510482 | 0.01205663 | 0.176065312 |
| *GNG11* | -1.031079 | 0.410898 | -2.509332 | 0.012095985 | 0.176423282 |
| *MTMR10* | 0.600918 | 0.239596 | 2.508045 | 0.01214012 | 0.176668457 |
| *MTMR4* | -0.323624 | 0.129038 | -2.507975 | 0.01214252 | 0.176668457 |
| *ACTR10* | 0.320043 | 0.127648 | 2.507238 | 0.012167887 | 0.176680966 |
| *MAP3K2* | -0.261201 | 0.104185 | -2.507086 | 0.012173106 | 0.176680966 |
| *ENSG00000257194* | -1.931388 | 0.770451 | -2.506829 | 0.01218195 | NA |
| *MCTP2* | -0.525529 | 0.209667 | -2.506487 | 0.012193738 | 0.176764591 |
| *ALDH1B1* | -1.017225 | 0.40593 | -2.505915 | 0.012213503 | 0.176835458 |
| *PSMB1* | 0.159037 | 0.063511 | 2.504093 | 0.012276565 | 0.177334245 |
| *RPS5* | -0.272657 | 0.108886 | -2.504058 | 0.012277789 | 0.177334245 |
| *SAFB* | 0.248087 | 0.099104 | 2.503303 | 0.012304013 | 0.177497338 |
| *ENSG00000247134* | -1.075747 | 0.429988 | -2.501806 | 0.012356145 | 0.178033329 |
| *SEC23B* | 0.454513 | 0.181756 | 2.500679 | 0.012395552 | 0.178384901 |
| *RPS27L* | 0.318426 | 0.127382 | 2.499776 | 0.012427177 | 0.17862376 |
| *ZCCHC2* | -0.548618 | 0.219881 | -2.49507 | 0.012593217 | 0.180791753 |
| *RFX3* | -0.539938 | 0.216515 | -2.493768 | 0.012639502 | 0.181161201 |
| *PPHLN1* | 0.176395 | 0.070742 | 2.49349 | 0.012649432 | 0.181161201 |
| *SVBP* | -0.186794 | 0.074957 | -2.491999 | 0.01270264 | 0.181615279 |
| *LINC02422* | -0.508918 | 0.204242 | -2.491746 | 0.012711694 | 0.181615279 |
| *MRNIP* | 0.2567 | 0.103079 | 2.490319 | 0.012762864 | 0.182127453 |
| *LENG8* | -0.270098 | 0.108521 | -2.488906 | 0.012813693 | 0.182633536 |

| | | | | | |
|---|---|---|---|---|---|
| *CCT7* | 0.256036 | 0.103001 | 2.485769 | 0.0129272 | 0.184030688 |
| *TPT1* | -0.19304 | 0.077715 | -2.483958 | 0.012993098 | 0.184747556 |
| *PAFAH1B3* | -0.379415 | 0.152786 | -2.483315 | 0.013016589 | 0.184860442 |
| *ENO2* | -0.481993 | 0.194197 | -2.481974 | 0.013065673 | 0.1853361 |
| *OGFR* | 0.224823 | 0.090643 | 2.480327 | 0.013126207 | 0.185972851 |
| *LACTB2.AS1* | 0.769314 | 0.310306 | 2.479209 | 0.013167416 | 0.186334613 |
| *CNR1* | 0.766783 | 0.309557 | 2.477036 | 0.013247871 | 0.186763507 |
| *MIB2* | -0.294535 | 0.118915 | -2.476845 | 0.013254947 | 0.186763507 |
| *ATP5MF* | 0.190488 | 0.076917 | 2.476531 | 0.013266615 | 0.186763507 |
| *QARS* | -0.368184 | 0.148682 | -2.476326 | 0.013274213 | 0.186763507 |
| *SEPTIN1* | -0.18124 | 0.073201 | -2.475934 | 0.013288832 | 0.186763507 |
| *TENT4A* | -0.697663 | 0.281832 | -2.475454 | 0.013306697 | 0.186763507 |
| *FKBP1B* | -0.770049 | 0.311204 | -2.474417 | 0.013345394 | 0.186763507 |
| *PTRHD1* | -0.231526 | 0.093574 | -2.474254 | 0.013351474 | 0.186763507 |
| *TMEM230* | 0.218436 | 0.088284 | 2.47424 | 0.013351999 | 0.186763507 |
| *NBEAL1* | -0.442563 | 0.178874 | -2.474164 | 0.01335484 | 0.186763507 |
| *POLR3D* | 0.367579 | 0.148679 | 2.472305 | 0.013424478 | 0.187516771 |
| *TUT1* | 0.40791 | 0.165048 | 2.471461 | 0.013456212 | 0.187739426 |
| *GPR160* | 0.558237 | 0.226023 | 2.46982 | 0.013518111 | 0.188381931 |
| *SPINK2* | -1.932957 | 0.782649 | -2.469763 | 0.013520274 | NA |
| *C16orf72* | 0.288018 | 0.116686 | 2.468328 | 0.013574584 | 0.188947406 |
| *RPL27A* | -0.213881 | 0.086706 | -2.466729 | 0.013635354 | 0.189571283 |
| *ENSG00000273188* | 0.790962 | 0.320877 | 2.465002 | 0.013701265 | 0.190265112 |
| *WWC3* | -0.271033 | 0.110022 | -2.463437 | 0.013761211 | 0.190874586 |
| *COMMD7* | -0.33689 | 0.136796 | -2.462713 | 0.013789012 | 0.19103728 |
| *PRDX3* | 0.224744 | 0.091319 | 2.461094 | 0.013851421 | 0.191678513 |
| *LSR* | -0.706768 | 0.287365 | -2.459483 | 0.013913745 | 0.192197244 |
| *IL21R* | 0.628993 | 0.255762 | 2.459289 | 0.013921244 | 0.192197244 |
| *ZNF524* | -0.251126 | 0.102163 | -2.458081 | 0.013968185 | 0.192537965 |
| *MRPL13* | 0.249373 | 0.101474 | 2.457501 | 0.013990749 | 0.192537965 |
| *NUDT1* | 0.213084 | 0.086711 | 2.457404 | 0.013994515 | 0.192537965 |
| *TNFRSF14* | -0.179769 | 0.073176 | -2.456681 | 0.014022698 | 0.19258561 |
| *RNF122* | 0.811948 | 0.330532 | 2.456485 | 0.014030381 | 0.19258561 |
| *TGS1* | 0.283984 | 0.115633 | 2.455896 | 0.014053389 | 0.192678933 |
| *ICA1* | 2.4801 | 1.009863 | 2.455878 | 0.014054067 | NA |
| *GNG7* | -0.236735 | 0.096422 | -2.455202 | 0.014080549 | 0.192828901 |
| *SPATS2L* | -1.867155 | 0.760513 | -2.455126 | 0.014083527 | NA |
| *WDFY2* | -0.288072 | 0.117401 | -2.453755 | 0.014137307 | 0.193383399 |
| *FNBP1* | 0.245859 | 0.100218 | 2.453232 | 0.014157911 | 0.193442633 |
| *SERPINF2* | -1.311971 | 0.534937 | -2.452572 | 0.014183903 | 0.19355012 |
| *LINC00996* | 1.403227 | 0.57223 | 2.452206 | 0.014198343 | 0.19355012 |
| *PLIN2* | 0.565355 | 0.230718 | 2.450412 | 0.014269285 | 0.194294378 |
| *PKNOX1* | 0.350039 | 0.143019 | 2.447496 | 0.014385274 | 0.195649602 |
| *MIIP* | -0.237377 | 0.097079 | -2.445194 | 0.014477406 | 0.196550619 |
| *TARS* | 0.281691 | 0.11521 | 2.445015 | 0.014484592 | 0.196550619 |
| *MARCKS* | -0.565564 | 0.231631 | -2.441661 | 0.014619886 | 0.198160308 |
| *KLB* | -1.789308 | 0.732838 | -2.441615 | 0.014621747 | NA |

| | | | | | |
|---|---|---|---|---|---|
| *RPL26* | -0.252971 | 0.10369 | -2.439687 | 0.014699985 | 0.199019049 |
| *ECE1* | 0.402117 | 0.164878 | 2.438881 | 0.014732828 | 0.199236774 |
| *RTN3* | 0.282992 | 0.116088 | 2.43774 | 0.014779419 | 0.19963972 |
| *PPP6R1* | -0.297433 | 0.122092 | -2.436144 | 0.014844764 | 0.200294784 |
| *UBE2L3* | 0.153553 | 0.063043 | 2.43567 | 0.014864232 | 0.200330071 |
| *CHMP5* | 0.202224 | 0.083074 | 2.434252 | 0.014922601 | 0.200888969 |
| *GALK1* | -0.33158 | 0.136436 | -2.430308 | 0.015086016 | 0.202679855 |
| *FCMR* | 0.325837 | 0.134077 | 2.430218 | 0.015089734 | 0.202679855 |
| *ENSG00000255328* | -1.555154 | 0.640056 | -2.429717 | 0.015110599 | 0.202725616 |
| *IL7R* | -0.936904 | 0.385674 | -2.429266 | 0.015129433 | 0.202725616 |
| *MYO1E* | -0.632013 | 0.260204 | -2.42891 | 0.015144304 | 0.202725616 |
| *CCDC154* | 0.817866 | 0.336951 | 2.427255 | 0.015213558 | 0.203423582 |
| *HIST1H2BD* | 0.49501 | 0.204209 | 2.424037 | 0.015349032 | 0.204994931 |
| *OTUD1* | -0.572286 | 0.236126 | -2.423646 | 0.015365566 | 0.204994931 |
| *ENSG00000234915* | -1.871117 | 0.772083 | -2.423465 | 0.015373225 | NA |
| *ABHD15* | 0.346534 | 0.143087 | 2.421845 | 0.015441936 | 0.205750019 |
| *VNN2* | 0.372078 | 0.153656 | 2.421496 | 0.015456782 | 0.205750019 |
| *KMT2D* | -0.376037 | 0.155467 | -2.418762 | 0.015573429 | 0.206959764 |
| *LINC02397* | 0.245584 | 0.101542 | 2.41855 | 0.015582484 | 0.206959764 |
| *NRBP2* | -1.247203 | 0.515943 | -2.417327 | 0.015634951 | 0.207424848 |
| *PSMC5* | 0.187526 | 0.077606 | 2.41639 | 0.015675263 | 0.207727816 |
| *TTC21B* | -0.517184 | 0.214177 | -2.414754 | 0.015745843 | 0.208430781 |
| *RPS15A* | -0.243503 | 0.100935 | -2.412474 | 0.015844678 | 0.209274333 |
| *EEF2* | -0.333373 | 0.138195 | -2.412342 | 0.015850427 | 0.209274333 |
| *CACTIN* | -0.594788 | 0.246588 | -2.412067 | 0.015862385 | 0.209274333 |
| *DEGS2* | 1.091716 | 0.453123 | 2.409315 | 0.015982517 | 0.21062547 |
| *FAM219B* | -0.252067 | 0.10484 | -2.404305 | 0.016203231 | 0.213297679 |
| *HACL1* | 0.32781 | 0.136409 | 2.403132 | 0.016255319 | 0.213746649 |
| *ATF5* | 0.578022 | 0.240689 | 2.401531 | 0.016326605 | 0.214446805 |
| *SNHG32* | -0.209919 | 0.087432 | -2.400924 | 0.016353727 | 0.214565951 |
| *SBDS* | 0.220084 | 0.091781 | 2.397932 | 0.016487914 | 0.216088014 |
| *SELENON* | -1.081719 | 0.451363 | -2.39656 | 0.016549801 | 0.21666023 |
| *MARCHF1* | -0.431321 | 0.180051 | -2.395544 | 0.016595716 | 0.216857518 |
| *PSMB6* | 0.168884 | 0.070503 | 2.39542 | 0.016601358 | 0.216857518 |
| *GFI1* | 1.734235 | 0.724818 | 2.392649 | 0.016727217 | NA |
| *AVPI1* | -0.977045 | 0.408382 | -2.392476 | 0.016735126 | 0.218152014 |
| *PLP2* | -0.326303 | 0.136402 | -2.392218 | 0.016746909 | 0.218152014 |
| *METTL9* | -0.174028 | 0.072758 | -2.391874 | 0.01676258 | 0.218152014 |
| *AIF1* | -2.185122 | 0.91358 | -2.391824 | 0.016764884 | NA |
| *RBM22* | 0.255631 | 0.10689 | 2.391527 | 0.016778445 | 0.218152014 |
| *NIP7* | 0.267919 | 0.112043 | 2.391226 | 0.016792218 | 0.218152014 |
| *PDHA1* | 0.296607 | 0.124209 | 2.387969 | 0.016941753 | 0.21985439 |
| *GRAPL* | 0.835323 | 0.349976 | 2.386803 | 0.016995586 | 0.220312464 |
| *SLC2A4RG* | -0.947281 | 0.397034 | -2.385892 | 0.017037763 | 0.220448138 |
| *MAGOHB* | 0.269031 | 0.112765 | 2.385776 | 0.017043143 | 0.220448138 |
| *TGFBR1* | -0.448955 | 0.188412 | -2.382832 | 0.017180032 | 0.221475901 |
| *CWF19L2* | 0.206137 | 0.086512 | 2.38275 | 0.017183848 | 0.221475901 |

| | | | | | |
|---|---|---|---|---|---|
| *PPDPF* | -0.227682 | 0.095569 | -2.382397 | 0.017200326 | 0.221475901 |
| *WNT10A* | -0.632878 | 0.265662 | -2.382265 | 0.017206505 | 0.221475901 |
| *AEN* | 0.335975 | 0.141043 | 2.382067 | 0.017215759 | 0.221475901 |
| *CYTIP* | -0.178062 | 0.074781 | -2.3811 | 0.017261021 | 0.221818121 |
| *FGF9* | 1.252731 | 0.526208 | 2.380678 | 0.017280816 | 0.221832675 |
| *PPT1* | 0.244372 | 0.102737 | 2.378623 | 0.017377436 | 0.222773188 |
| *CCDC88A* | -0.514317 | 0.216278 | -2.378041 | 0.017404898 | 0.222773188 |
| *IL17RA* | -0.410946 | 0.172817 | -2.377926 | 0.017410305 | 0.222773188 |
| *AP1S2* | 0.207884 | 0.087491 | 2.376063 | 0.017498481 | 0.223474027 |
| *ENSG00000277283* | -0.7807 | 0.328611 | -2.375758 | 0.017512954 | 0.223474027 |
| *RPA2* | 0.206293 | 0.086839 | 2.375578 | 0.017521477 | 0.223474027 |
| *RAP2C.AS1* | -1.023337 | 0.431233 | -2.373049 | 0.017641921 | 0.224769043 |
| *MID1IP1* | -0.218677 | 0.092221 | -2.371221 | 0.017729418 | 0.225397929 |
| *TTC7A* | -0.333471 | 0.140637 | -2.371153 | 0.017732696 | 0.225397929 |
| *RPL4* | -0.221468 | 0.093413 | -2.370831 | 0.017748167 | 0.225397929 |
| *ENSG00000274712* | -0.652868 | 0.275657 | -2.368406 | 0.017864903 | 0.226420207 |
| *SMAP2* | -0.318827 | 0.134619 | -2.368368 | 0.017866758 | 0.226420207 |
| *TOX4* | 0.211169 | 0.08923 | 2.36656 | 0.017954285 | 0.227062298 |
| *PRRC2B* | -0.253429 | 0.107089 | -2.366532 | 0.017955629 | 0.227062298 |
| *ENSG00000259274* | -1.272175 | 0.53772 | -2.365869 | 0.0179878 | 0.227227399 |
| *TOMM20* | -0.1915 | 0.080965 | -2.365228 | 0.018018972 | 0.227379532 |
| *DNAH1* | 0.915823 | 0.387316 | 2.364537 | 0.018052612 | 0.227525172 |
| *LRRC14* | -0.547581 | 0.231613 | -2.364205 | 0.018068795 | 0.227525172 |
| *ELOF1* | 0.2498 | 0.105693 | 2.363452 | 0.018105571 | 0.227747002 |
| *FIP1L1* | 0.256143 | 0.10848 | 2.361192 | 0.018216313 | 0.228662012 |
| *ENSG00000258572* | -1.006725 | 0.426365 | -2.361182 | 0.018216785 | 0.228662012 |
| *KCNC4* | -0.605366 | 0.256601 | -2.359169 | 0.018315888 | 0.22966346 |
| *CLDN12* | -0.724859 | 0.307451 | -2.357638 | 0.018391606 | 0.23036988 |
| *SCAMP2* | 0.221941 | 0.094181 | 2.356537 | 0.018446229 | 0.230699106 |
| *CEP70* | 0.904942 | 0.384048 | 2.356326 | 0.018456705 | 0.230699106 |
| *LINC01089* | -0.259801 | 0.110285 | -2.355711 | 0.018487288 | 0.230838646 |
| *WDR74* | 0.251037 | 0.106633 | 2.354214 | 0.018561915 | 0.231259629 |
| *ENSG00000261884* | 0.346708 | 0.147279 | 2.354088 | 0.018568214 | 0.231259629 |
| *CEP78* | -0.367962 | 0.156365 | -2.353226 | 0.018611329 | 0.231259629 |
| *EIF5B* | 0.153016 | 0.065024 | 2.35322 | 0.018611639 | 0.231259629 |
| *PFKL* | -0.23344 | 0.099206 | -2.353087 | 0.018618278 | 0.231259629 |
| *BICD1* | -0.340544 | 0.144779 | -2.352157 | 0.018664889 | 0.231535901 |
| *LAMTOR5* | -0.153106 | 0.0651 | -2.351867 | 0.018679476 | 0.231535901 |
| *HHEX* | -0.200039 | 0.08508 | -2.351185 | 0.018713704 | 0.231718536 |
| *ARHGAP15* | 0.190404 | 0.081066 | 2.348756 | 0.018836244 | 0.232993162 |
| *RNPS1* | 0.146399 | 0.062387 | 2.346624 | 0.018944368 | 0.234087005 |
| *SINHCAF* | 0.191663 | 0.081736 | 2.344916 | 0.019031375 | 0.234917923 |
| *HACD4* | -0.540741 | 0.230698 | -2.343934 | 0.01908155 | 0.235202961 |
| *KIFC2* | -0.723404 | 0.308683 | -2.343516 | 0.01910296 | 0.235202961 |
| *RPL35A* | -0.247177 | 0.105482 | -2.343304 | 0.019113827 | 0.235202961 |
| *ABTB1* | 0.190569 | 0.081358 | 2.342357 | 0.019162389 | 0.235335762 |
| *DNASE1L3* | -1.145063 | 0.488858 | -2.342321 | 0.019164214 | 0.235335762 |

| | | | | | |
|---|---|---|---|---|---|
| *TFEB* | 0.473086 | 0.202024 | 2.341735 | 0.019194343 | 0.235454541 |
| *PPP2R5C* | 0.173217 | 0.073981 | 2.341362 | 0.019213503 | 0.235454541 |
| *RGS2* | -0.88454 | 0.378022 | -2.339918 | 0.019287955 | 0.236123503 |
| *LINC00672* | -0.677297 | 0.289554 | -2.339101 | 0.019330195 | 0.236397148 |
| *EIF2AK3* | -0.373163 | 0.159615 | -2.337892 | 0.019392826 | 0.236784405 |
| *METTL21A* | -0.237608 | 0.101648 | -2.337566 | 0.01940979 | 0.236784405 |
| *CYTOR* | -0.841631 | 0.360123 | -2.337069 | 0.019435611 | 0.236784405 |
| *SLC23A2* | 0.390221 | 0.166989 | 2.336803 | 0.019449444 | 0.236784405 |
| *PFDN2* | 0.162779 | 0.069677 | 2.336199 | 0.019480872 | 0.236784405 |
| *RPS6KA1* | 0.279148 | 0.119489 | 2.336189 | 0.019481379 | 0.236784405 |
| *SLC6A16* | -0.472243 | 0.202214 | -2.335358 | 0.019524752 | 0.237069186 |
| *C1orf43* | 0.174801 | 0.074866 | 2.334856 | 0.019550934 | 0.237109968 |
| *SMOX* | 1.054663 | 0.451767 | 2.33453 | 0.019568005 | 0.237109968 |
| *LCMT1* | -0.259724 | 0.111316 | -2.333204 | 0.019637467 | 0.237535214 |
| *FAM177B* | -0.517393 | 0.22178 | -2.33291 | 0.019652895 | 0.237535214 |
| *TMEM154* | 0.311973 | 0.133738 | 2.332716 | 0.019663048 | 0.237535214 |
| *ATP23* | -0.456181 | 0.195685 | -2.331199 | 0.019742891 | 0.238257608 |
| *SESTD1* | -0.313004 | 0.134322 | -2.330251 | 0.01979288 | 0.238618625 |
| *FAM111A* | -0.291789 | 0.125291 | -2.328899 | 0.019864427 | 0.239173531 |
| *EIF3K* | -0.229759 | 0.098667 | -2.328621 | 0.019879149 | 0.239173531 |
| *ENSG00000233461* | 0.493031 | 0.211926 | 2.326434 | 0.019995435 | 0.240237034 |
| *EEF1B2* | -0.335836 | 0.144383 | -2.326008 | 0.020018134 | 0.240237034 |
| *PSMA5* | 0.173857 | 0.074753 | 2.325748 | 0.020031999 | 0.240237034 |
| *HSPA5* | 0.324701 | 0.13963 | 2.325441 | 0.020048384 | 0.240237034 |
| *RPL37* | -0.183246 | 0.078826 | -2.324688 | 0.02008864 | 0.240477007 |
| *IFNAR1* | 0.213178 | 0.091736 | 2.323813 | 0.02013553 | 0.240795825 |
| *GFM1* | 0.330769 | 0.142389 | 2.323004 | 0.020178926 | 0.241072252 |
| *ZNF169* | -0.36109 | 0.155537 | -2.321571 | 0.020256062 | 0.241467499 |
| *EPM2AIP1* | -0.17611 | 0.075858 | -2.321562 | 0.020256547 | 0.241467499 |
| *DDAH2* | -0.277642 | 0.119608 | -2.321257 | 0.020272951 | 0.241467499 |
| *IARS* | 0.270708 | 0.116658 | 2.320526 | 0.020312444 | 0.241695722 |
| *HSD17B8* | -0.627331 | 0.270445 | -2.319624 | 0.020361203 | 0.242033624 |
| *CDV3* | -0.287191 | 0.123832 | -2.319194 | 0.020384506 | 0.242068555 |
| *KHDRBS2* | -0.639885 | 0.275996 | -2.31846 | 0.020424329 | 0.242299402 |
| *ENSG00000225889* | -0.798578 | 0.344679 | -2.316873 | 0.020510624 | 0.243080551 |
| *LIMS2* | -1.466188 | 0.632887 | -2.316667 | 0.020521879 | NA |
| *HIST1H2BC* | 0.59423 | 0.256567 | 2.316084 | 0.020553687 | 0.243326062 |
| *LINC02035* | -0.998813 | 0.431314 | -2.315743 | 0.02057228 | 0.243326062 |
| *ERGIC1* | -0.242214 | 0.104731 | -2.312733 | 0.020737349 | 0.245034663 |
| *FSIP2* | 0.671352 | 0.29038 | 2.311981 | 0.020778721 | 0.245279694 |
| *SYK* | -0.223658 | 0.096757 | -2.311532 | 0.020803463 | 0.245328136 |
| *PAFAH1B2* | -0.171707 | 0.074351 | -2.309403 | 0.020921249 | 0.246472627 |
| *LCK* | 0.507652 | 0.219922 | 2.308329 | 0.020980855 | 0.246798296 |
| *BCL9* | -1.212189 | 0.525176 | -2.308157 | 0.020990416 | 0.246798296 |
| *MOB4* | 0.229295 | 0.09939 | 2.307022 | 0.021053583 | 0.247296384 |
| *RTKN* | -0.777793 | 0.337265 | -2.30618 | 0.021100558 | 0.247410377 |
| *SLC9A3R1* | 0.338124 | 0.146621 | 2.306102 | 0.021104915 | 0.247410377 |

| | | | | | |
|---|---|---|---|---|---|
| *KARS* | 0.202393 | 0.087782 | 2.305625 | 0.021131568 | 0.247478768 |
| *EEF1A1* | -0.331651 | 0.143943 | -2.304043 | 0.021220211 | 0.247501918 |
| *PPP1R8* | 0.268472 | 0.116534 | 2.30381 | 0.021233296 | 0.247501918 |
| *HIST1H1D* | -0.354896 | 0.154055 | -2.303692 | 0.021239948 | 0.247501918 |
| *ATP1B3* | 0.288205 | 0.125116 | 2.303502 | 0.021250607 | 0.247501918 |
| *ETFA* | 0.195814 | 0.085009 | 2.303447 | 0.02125368 | 0.247501918 |
| *ZEB1* | -0.317379 | 0.137809 | -2.303029 | 0.021277212 | 0.247501918 |
| *PLEKHJ1* | -0.213917 | 0.092898 | -2.302699 | 0.021295764 | 0.247501918 |
| *NACA* | -0.1797 | 0.078045 | -2.302536 | 0.021304946 | 0.247501918 |
| *TIMMDC1* | 0.262431 | 0.114006 | 2.301901 | 0.021340771 | 0.247501918 |
| *ENSG00000273002* | 0.638803 | 0.277513 | 2.301883 | 0.021341757 | 0.247501918 |
| *UEVLD* | -0.537364 | 0.233538 | -2.300969 | 0.021393391 | 0.247858903 |
| *DHRS12* | -0.3379 | 0.146966 | -2.299177 | 0.021494879 | 0.248792238 |
| *SRP72* | 0.200252 | 0.08717 | 2.297265 | 0.021603676 | 0.249222744 |
| *SHMT2* | -0.208475 | 0.090754 | -2.29715 | 0.021610209 | 0.249222744 |
| *NDUFAF4* | 0.251626 | 0.109556 | 2.296775 | 0.021631632 | 0.249222744 |
| *HMOX1* | 0.797512 | 0.34727 | 2.296521 | 0.021646117 | 0.249222744 |
| *C11orf80* | -0.378831 | 0.164985 | -2.296148 | 0.021667418 | 0.249222744 |
| *ATP5MG* | -0.132663 | 0.057778 | -2.296091 | 0.02167068 | 0.249222744 |
| *ATP5PF* | 0.139408 | 0.060727 | 2.295665 | 0.021695039 | 0.249222744 |
| *SH3BP5* | -0.342827 | 0.149342 | -2.295582 | 0.021699801 | 0.249222744 |
| *ATP6AP2* | 0.265611 | 0.115764 | 2.294412 | 0.021766857 | 0.249751568 |
| *LY75* | -0.470279 | 0.205022 | -2.293799 | 0.021802022 | 0.249913824 |
| *RNPC3* | -0.337367 | 0.147116 | -2.293214 | 0.021835714 | 0.250058893 |
| *GPR65* | -0.261475 | 0.114082 | -2.291991 | 0.021906166 | 0.250624245 |
| *C22orf15* | 1.348436 | 0.588399 | 2.291704 | 0.021922704 | NA |
| *PITPNM1* | -0.455087 | 0.19874 | -2.289858 | 0.022029526 | 0.251793251 |
| *KANK2* | -1.189264 | 0.51959 | -2.288852 | 0.022087959 | 0.252042725 |
| *ENSG00000251136* | 0.456376 | 0.199436 | 2.288334 | 0.022118106 | 0.252042725 |
| *ATXN1L* | 0.317574 | 0.138779 | 2.288334 | 0.022118074 | 0.252042725 |
| *RPL8* | -0.250023 | 0.109287 | -2.287758 | 0.022151637 | 0.252042725 |
| *MYL5* | -0.459129 | 0.200706 | -2.287576 | 0.02216222 | 0.252042725 |
| *NOP10* | 0.15878 | 0.069418 | 2.287296 | 0.022178572 | 0.252042725 |
| *HIST1H4C* | -0.297505 | 0.130098 | -2.286774 | 0.022209046 | 0.252147971 |
| *CINP* | 0.258065 | 0.112973 | 2.284312 | 0.022353225 | 0.253492713 |
| *SLC4A1AP* | 0.315968 | 0.138368 | 2.28353 | 0.022399143 | 0.253492713 |
| *RPS15* | -0.182294 | 0.079838 | -2.283307 | 0.022412309 | 0.253492713 |
| *DDX18* | 0.151706 | 0.066443 | 2.283259 | 0.022415094 | 0.253492713 |
| *GADD45A* | 0.460148 | 0.20156 | 2.282936 | 0.022434116 | 0.253492713 |
| *SMYD4* | -0.319452 | 0.140027 | -2.28135 | 0.02252774 | 0.254308872 |
| *S100A4* | -0.488687 | 0.214316 | -2.280216 | 0.02259491 | 0.254825134 |
| *LINC02362* | -1.581621 | 0.693868 | -2.279426 | 0.022641739 | NA |
| *PTPMT1* | -0.222825 | 0.097829 | -2.277693 | 0.022744852 | 0.256104495 |
| *RPL7A* | -0.250804 | 0.110119 | -2.277583 | 0.022751438 | 0.256104495 |
| *TINCR* | 1.274262 | 0.559699 | 2.276693 | 0.022804547 | NA |
| *EIF1* | 0.14463 | 0.06353 | 2.276569 | 0.022811973 | 0.256542969 |
| *COX7B* | 0.149991 | 0.065904 | 2.275887 | 0.022852802 | 0.256759221 |

| | | | | | |
|---|---|---|---|---|---|
| *C4orf3* | 0.178263 | 0.078348 | 2.275272 | 0.022889614 | 0.256929975 |
| *PCSK7* | -0.222168 | 0.097661 | -2.27488 | 0.02291312 | 0.25695119 |
| *ARL6IP4* | -0.152881 | 0.067271 | -2.272621 | 0.023049032 | 0.258231707 |
| *DIS3L2* | -0.31694 | 0.13954 | -2.27132 | 0.023127624 | 0.258647294 |
| *MCPH1* | 0.252793 | 0.1113 | 2.271286 | 0.023129644 | 0.258647294 |
| *COA4* | 0.207274 | 0.091285 | 2.270619 | 0.02317007 | 0.258855842 |
| *LPP.AS2* | -0.852202 | 0.375675 | -2.268452 | 0.023301661 | 0.259905069 |
| *RPS27* | -0.184624 | 0.081391 | -2.268353 | 0.023307715 | 0.259905069 |
| *TLR7* | -0.599306 | 0.264246 | -2.26799 | 0.023329795 | 0.25990747 |
| *ICAM3* | -0.242982 | 0.107173 | -2.2672 | 0.023377996 | 0.260200599 |
| *ENSG00000260257* | -0.404185 | 0.17838 | -2.26587 | 0.023459332 | 0.260861629 |
| *KAT6A* | 0.206541 | 0.091168 | 2.265488 | 0.02348275 | 0.260877987 |
| *PSMC3* | 0.200548 | 0.088552 | 2.264738 | 0.02352875 | 0.261144955 |
| *GHITM* | 0.198451 | 0.087643 | 2.264318 | 0.023554552 | 0.261187461 |
| *NT5C2* | 0.300373 | 0.132709 | 2.263394 | 0.023611436 | 0.261574223 |
| *LZTS3* | -1.161672 | 0.513433 | -2.26256 | 0.023662848 | 0.261814993 |
| *MLXIP* | 0.282571 | 0.124903 | 2.262327 | 0.02367722 | 0.261814993 |
| *TPI1* | 0.133827 | 0.059168 | 2.261805 | 0.023709475 | 0.261928003 |
| *SPPL2B* | -0.250488 | 0.110809 | -2.260537 | 0.023787966 | 0.262551114 |
| *TRIOBP* | -0.361662 | 0.160112 | -2.25881 | 0.023895226 | 0.263490303 |
| *UBTD2* | -0.690358 | 0.305768 | -2.257787 | 0.02395894 | 0.263948026 |
| *MBD6* | 0.407128 | 0.180429 | 2.256452 | 0.02404232 | 0.264621349 |
| *HNRNPF* | 0.228983 | 0.101505 | 2.255877 | 0.024078362 | 0.264772882 |
| *TCF3* | -0.321965 | 0.142839 | -2.254048 | 0.02419316 | 0.265789365 |
| *JUN* | 0.277184 | 0.12304 | 2.252788 | 0.024272531 | 0.26641512 |
| *BOLA2.SMG1P6* | 0.424942 | 0.1887 | 2.251943 | 0.024325885 | 0.26675442 |
| *SLC25A3* | -0.220642 | 0.098034 | -2.250664 | 0.024406822 | 0.267395294 |
| *LINC01003* | -0.722452 | 0.321431 | -2.247607 | 0.02460127 | 0.269108307 |
| *VANGL2* | -1.413905 | 0.629125 | -2.247416 | 0.024613471 | 0.269108307 |
| *ENSG00000265206* | -0.214659 | 0.095534 | -2.246925 | 0.024644816 | 0.269108307 |
| *PSMA3* | 0.20632 | 0.091829 | 2.246786 | 0.024653735 | 0.269108307 |
| *ARHGAP5* | -0.253563 | 0.112894 | -2.246032 | 0.024701938 | 0.269387096 |
| *LINC01184* | -0.360545 | 0.160581 | -2.24525 | 0.024752059 | 0.269686271 |
| *TKT* | -0.304539 | 0.13569 | -2.244371 | 0.024808569 | 0.270054447 |
| *NPM3* | -0.412401 | 0.183857 | -2.243051 | 0.024893556 | 0.270460488 |
| *RPF1* | 0.223169 | 0.099515 | 2.242562 | 0.024925099 | 0.270460488 |
| *ZCCHC17* | 0.204127 | 0.091028 | 2.242479 | 0.024930436 | 0.270460488 |
| *STARD9* | -0.481844 | 0.2149 | -2.242171 | 0.024950352 | 0.270460488 |
| *TPD52L2* | 0.231255 | 0.103145 | 2.242027 | 0.024959633 | 0.270460488 |
| *SLX4* | -0.747464 | 0.334027 | -2.237734 | 0.025238435 | 0.273232497 |
| *LRP10* | 0.184993 | 0.082684 | 2.237344 | 0.025263873 | 0.273259013 |
| *ENSG00000254281* | -1.178622 | 0.526894 | -2.236924 | 0.025291307 | 0.273307062 |
| *DDX17* | -0.161857 | 0.07242 | -2.234963 | 0.025419759 | 0.274348948 |
| *HS2ST1* | -0.481022 | 0.215253 | -2.234679 | 0.025438454 | 0.274348948 |
| *TMED9* | 0.197633 | 0.088462 | 2.23411 | 0.025475805 | 0.274348948 |
| *CNTRL* | -0.169519 | 0.075906 | -2.233266 | 0.025531426 | 0.274348948 |
| *TSPYL2* | 0.373364 | 0.16719 | 2.233172 | 0.025537578 | 0.274348948 |

| | | | | | |
|---|---|---|---|---|---|
| *PSMA6* | 0.291871 | 0.130705 | 2.233044 | 0.025546043 | 0.274348948 |
| *TMEM219* | -0.171505 | 0.076817 | -2.232633 | 0.025573162 | 0.274348948 |
| *LPXN* | 0.374483 | 0.167778 | 2.232006 | 0.025614572 | 0.274348948 |
| *RPL15* | -0.184318 | 0.082587 | -2.231816 | 0.025627113 | 0.274348948 |
| *SLC38A1* | -0.199369 | 0.089334 | -2.231728 | 0.02563296 | 0.274348948 |
| *AHCY* | -0.363336 | 0.162814 | -2.231597 | 0.025641598 | 0.274348948 |
| *UCK2* | 0.583459 | 0.26152 | 2.231028 | 0.025679295 | 0.274505193 |
| *LINC00924* | -1.846145 | 0.827579 | -2.230779 | 0.025695789 | NA |
| *FAM76A* | -0.238027 | 0.10681 | -2.228514 | 0.025846274 | 0.276041924 |
| *PLEKHB2* | 0.257552 | 0.115634 | 2.227302 | 0.0259271 | 0.276585262 |
| *STRADB* | -0.358282 | 0.160877 | -2.227054 | 0.025943684 | 0.276585262 |
| *TTC33* | 0.363744 | 0.163369 | 2.226517 | 0.02597958 | 0.276719776 |
| *CDC123* | 0.220312 | 0.098988 | 2.225635 | 0.02603861 | 0.277040716 |
| *AGBL5* | -0.590292 | 0.265255 | -2.225371 | 0.026056324 | 0.277040716 |
| *PRPSAP2* | 0.296274 | 0.133159 | 2.224953 | 0.026084406 | 0.277065055 |
| *ENSG00000227373* | 1.361434 | 0.611979 | 2.224643 | 0.026105229 | 0.277065055 |
| *RBM7* | 0.235222 | 0.105774 | 2.223825 | 0.026160217 | 0.277400977 |
| *TOMM5* | 0.189793 | 0.085381 | 2.222879 | 0.026223972 | 0.277486127 |
| *UBE2O* | 0.315573 | 0.141976 | 2.222719 | 0.026234761 | 0.277486127 |
| *MAP4K4* | -0.235724 | 0.106077 | -2.222198 | 0.026269924 | 0.277486127 |
| *KIAA1109* | -0.303818 | 0.136773 | -2.221337 | 0.026328164 | 0.277486127 |
| *PIP4K2A* | -0.401734 | 0.180855 | -2.221306 | 0.026330229 | 0.277486127 |
| *CLIC4* | -0.254941 | 0.114784 | -2.221047 | 0.026347783 | 0.277486127 |
| *MRPS18A* | 0.214946 | 0.096787 | 2.220805 | 0.026364146 | 0.277486127 |
| *BOD1L1* | 0.180001 | 0.081053 | 2.22077 | 0.026366576 | 0.277486127 |
| *CALM3* | -0.220639 | 0.09936 | -2.220596 | 0.02637834 | 0.277486127 |
| *TIGIT* | -1.919363 | 0.864645 | -2.219828 | 0.026430422 | NA |
| *ZFP14* | -0.225698 | 0.101742 | -2.21834 | 0.026531691 | 0.278852531 |
| *MGMT* | -0.270996 | 0.122281 | -2.216167 | 0.02668004 | 0.27987236 |
| *KANSL1L* | -0.552306 | 0.249219 | -2.216146 | 0.026681522 | 0.27987236 |
| *FAM30A* | -0.202499 | 0.091395 | -2.215657 | 0.026714986 | 0.27987236 |
| *FAM209B* | -1.754859 | 0.792243 | -2.215051 | 0.026756543 | NA |
| *SGO1* | 0.333485 | 0.150571 | 2.214804 | 0.026773551 | 0.27987236 |
| *DTX2* | 0.392021 | 0.17701 | 2.214683 | 0.026781808 | 0.27987236 |
| *RPL18* | -0.198471 | 0.089625 | -2.214465 | 0.026796775 | 0.27987236 |
| *RPL5* | -0.24085 | 0.108772 | -2.214267 | 0.0268104 | 0.27987236 |
| *FAR1* | -0.410079 | 0.185207 | -2.21417 | 0.026817079 | 0.27987236 |
| *LENG1* | 0.242459 | 0.109617 | 2.211863 | 0.02697615 | 0.281107896 |
| *TOR1AIP1* | 0.284498 | 0.128656 | 2.211302 | 0.027014929 | 0.281107896 |
| *PRMT1* | 0.187039 | 0.084589 | 2.211163 | 0.027024532 | 0.281107896 |
| *METTL1* | 0.653393 | 0.295508 | 2.211083 | 0.02703006 | 0.281107896 |
| *SERP1* | -0.146606 | 0.066338 | -2.20997 | 0.02710723 | 0.281239742 |
| *ZC3H14* | -0.263676 | 0.119315 | -2.20991 | 0.027111426 | 0.281239742 |
| *TFIP11* | 0.331356 | 0.149943 | 2.209877 | 0.027113716 | 0.281239742 |
| *AK2* | 0.222251 | 0.100681 | 2.207481 | 0.027280439 | 0.282722384 |
| *HOMER2* | -0.691802 | 0.313522 | -2.206549 | 0.027345599 | 0.282906524 |
| *RPS8* | -0.28199 | 0.127797 | -2.206546 | 0.027345806 | 0.282906524 |

| | | | | | |
|---|---|---|---|---|---|
| *ACYP1* | 0.229107 | 0.10387 | 2.205715 | 0.027403963 | 0.283261659 |
| *IL6ST* | -0.449563 | 0.203862 | -2.205232 | 0.027437805 | 0.283365063 |
| *TMSB4Y* | -1.622997 | 0.736086 | -2.204903 | 0.027460915 | NA |
| *CORO2B* | -1.653715 | 0.750177 | -2.204434 | 0.027493863 | 0.283681546 |
| *ZBTB18* | -0.312808 | 0.14192 | -2.204116 | 0.027516179 | 0.283681546 |
| *IFI27L1* | 0.403311 | 0.183052 | 2.203256 | 0.027576733 | 0.283943174 |
| *RBM42* | 0.300124 | 0.136232 | 2.203036 | 0.02759223 | 0.283943174 |
| *CCR9* | 1.846828 | 0.838381 | 2.20285 | 0.027605316 | NA |
| *SPIB* | 0.160954 | 0.07307 | 2.202738 | 0.027613217 | 0.283943174 |
| *INPP5B* | 0.321446 | 0.146032 | 2.201195 | 0.027722254 | 0.284818009 |
| *MPZL3* | -1.052623 | 0.478642 | -2.199184 | 0.027864808 | 0.286035386 |
| *LINC00630* | -1.70836 | 0.777645 | -2.196839 | 0.02803193 | NA |
| *AIMP1* | 0.164146 | 0.074754 | 2.195815 | 0.028105171 | 0.288247034 |
| *ENSG00000260030* | 2.714015 | 1.236601 | 2.194739 | 0.02818234 | NA |
| *NUCKS1* | -0.18145 | 0.082695 | -2.194214 | 0.028220054 | 0.288247034 |
| *PTPN18* | -0.380619 | 0.173466 | -2.194197 | 0.028221257 | 0.288247034 |
| *OPA3* | 0.290335 | 0.132345 | 2.193768 | 0.028252087 | 0.288247034 |
| *BCAS2* | 0.231604 | 0.10558 | 2.19364 | 0.028261268 | 0.288247034 |
| *RUNX3* | 0.212531 | 0.096887 | 2.193592 | 0.028264743 | 0.288247034 |
| *ENSG00000231856* | -0.65629 | 0.299196 | -2.193514 | 0.028270379 | 0.288247034 |
| *INPP5E* | -0.562215 | 0.256314 | -2.19346 | 0.028274253 | 0.288247034 |
| *CASP4* | -0.150796 | 0.068763 | -2.192976 | 0.028309104 | 0.288355025 |
| *RUNX1* | 0.291417 | 0.132979 | 2.191447 | 0.028419468 | 0.289231347 |
| *SEC61A2* | -0.440864 | 0.201209 | -2.191079 | 0.028446043 | 0.289254159 |
| *HDAC1* | 0.21931 | 0.100123 | 2.1904 | 0.028495244 | 0.289506806 |
| *SLC7A5* | -1.187776 | 0.542477 | -2.189542 | 0.028557498 | 0.289891523 |
| *RAB11A* | 0.155457 | 0.071043 | 2.188217 | 0.028653834 | 0.290621265 |
| *TPRKB* | 0.153221 | 0.07004 | 2.187627 | 0.028696794 | 0.290691999 |
| *FAM49B* | 0.170715 | 0.078043 | 2.18745 | 0.028709717 | 0.290691999 |
| *RPL7* | -0.280586 | 0.128322 | -2.186578 | 0.028773352 | 0.29108837 |
| *UBE2G1* | 0.240739 | 0.110144 | 2.185682 | 0.028838865 | 0.291318186 |
| *ENSG00000133961* | 0.211768 | 0.096893 | 2.185597 | 0.028845083 | 0.291318186 |
| *GTPBP1* | 0.27914 | 0.127824 | 2.183785 | 0.02897805 | 0.292412633 |
| *KPNA2* | -0.325365 | 0.149074 | -2.182574 | 0.029067232 | 0.29306377 |
| *MPP1* | -0.601845 | 0.275854 | -2.181754 | 0.029127658 | 0.293186466 |
| *ABLIM2* | -0.496415 | 0.227532 | -2.18174 | 0.02912873 | 0.293186466 |
| *ANKLE1* | 0.609294 | 0.27936 | 2.181036 | 0.029180756 | 0.29328622 |
| *SLC2A1* | 0.258138 | 0.118361 | 2.180938 | 0.029187987 | 0.29328622 |
| *AFF2* | -1.551188 | 0.711357 | -2.180604 | 0.02921269 | NA |
| *CLASP1* | -0.42242 | 0.193737 | -2.180376 | 0.02922962 | 0.293456493 |
| *GRK3* | 0.256981 | 0.11788 | 2.180033 | 0.029255032 | 0.293463772 |
| *TSPAN33* | 0.294479 | 0.135123 | 2.179341 | 0.02930632 | 0.293730378 |
| *TMEM8A* | -0.458771 | 0.210563 | -2.17878 | 0.029347984 | 0.293900158 |
| *QSOX2* | -0.478023 | 0.219931 | -2.173513 | 0.029741758 | 0.297592826 |
| *ENSG00000261732* | 0.374614 | 0.172392 | 2.173043 | 0.029777078 | 0.297695646 |
| *SELENOT* | 0.16132 | 0.074327 | 2.17042 | 0.029975019 | 0.299422729 |
| *ENSG00000267416* | -1.689397 | 0.778397 | -2.170355 | 0.029979991 | NA |

| | | | | | |
|---|---|---|---|---|---|
| *ATP5F1B* | 0.198203 | 0.091382 | 2.168944 | 0.03008695 | 0.300288476 |
| *FIBP* | -0.204635 | 0.094364 | -2.168579 | 0.030114672 | 0.300313004 |
| *NUDT14* | -0.455513 | 0.210135 | -2.167714 | 0.030180429 | 0.300512029 |
| *ZNF93* | -0.415104 | 0.191499 | -2.167652 | 0.030185191 | 0.300512029 |
| *SEPTIN6* | -0.160795 | 0.074191 | -2.167315 | 0.030210866 | 0.300515958 |
| *ENSG00000230155* | 0.400134 | 0.184752 | 2.165788 | 0.030327379 | 0.301422708 |
| *ENSG00000261684* | -1.670038 | 0.771214 | -2.165467 | 0.030351928 | NA |
| *TWF2* | 0.189627 | 0.08759 | 2.16495 | 0.030391529 | 0.301807939 |
| *AGAP4* | -0.627402 | 0.289865 | -2.164466 | 0.030428591 | 0.301923761 |
| *ALG2* | -0.345791 | 0.159798 | -2.163929 | 0.030469797 | 0.302080467 |
| *RPL7L1* | 0.159567 | 0.073757 | 2.163408 | 0.030509793 | 0.302224925 |
| *SEC22C* | -0.23394 | 0.108194 | -2.162221 | 0.030601139 | 0.302672153 |
| *ZFC3H1* | -0.245091 | 0.113355 | -2.16216 | 0.030605866 | 0.302672153 |
| *HLCS* | -0.883918 | 0.409011 | -2.161112 | 0.030686657 | 0.303058577 |
| *HLA.DPA1* | -0.292825 | 0.135505 | -2.160992 | 0.030695931 | 0.303058577 |
| *DOP1B* | -0.380207 | 0.176099 | -2.159055 | 0.030845884 | 0.304063381 |
| *CD59* | 0.252338 | 0.116876 | 2.159017 | 0.030848863 | 0.304063381 |
| *CYLD* | -0.207409 | 0.096094 | -2.158398 | 0.030896917 | 0.304284713 |
| *MRPS28* | 0.260188 | 0.120614 | 2.157191 | 0.030990787 | 0.304943418 |
| *C5orf56* | 0.343315 | 0.159195 | 2.156568 | 0.031039353 | 0.304943418 |
| *ARMC7* | 0.426967 | 0.197986 | 2.15655 | 0.031040762 | 0.304943418 |
| *FAM13A* | 0.576891 | 0.267607 | 2.155742 | 0.031103844 | 0.305003406 |
| *C3orf33* | -0.948159 | 0.439833 | -2.155725 | 0.031105125 | 0.305003406 |
| *PSD4* | -0.357542 | 0.165876 | -2.155486 | 0.031123844 | 0.305003406 |
| *MAN2A1* | -0.338244 | 0.157036 | -2.153933 | 0.031245422 | 0.305397625 |
| *ENSG00000229728* | 1.089926 | 0.506147 | 2.153381 | 0.031288782 | 0.305397625 |
| *BIRC6* | -0.189383 | 0.08795 | -2.153312 | 0.031294171 | 0.305397625 |
| *ZNRD2* | 0.219641 | 0.102013 | 2.15307 | 0.031313154 | 0.305397625 |
| *SCOC.AS1* | 0.502352 | 0.233321 | 2.153055 | 0.031314317 | 0.305397625 |
| *TBL1XR1* | -0.22139 | 0.102828 | -2.153006 | 0.031318222 | 0.305397625 |
| *MTG1* | 0.291312 | 0.135394 | 2.151592 | 0.031429491 | 0.306231445 |
| *FAM172A* | -0.274565 | 0.127679 | -2.150435 | 0.031520802 | 0.306869597 |
| *RPL28* | -0.263577 | 0.122608 | -2.149748 | 0.031575181 | 0.30701592 |
| *SEC62* | -0.15366 | 0.071483 | -2.149592 | 0.031587488 | 0.30701592 |
| *DNAJC10* | -0.195491 | 0.090985 | -2.148603 | 0.03166587 | 0.307526307 |
| *NFYC* | 0.251775 | 0.117218 | 2.147923 | 0.031719909 | 0.307799638 |
| *NDUFS4* | -0.311012 | 0.14483 | -2.147425 | 0.031759478 | 0.307932234 |
| *ENSG00000253891* | 0.902322 | 0.420332 | 2.146692 | 0.031817797 | 0.308246257 |
| *ARID4B* | 0.164176 | 0.076495 | 2.146231 | 0.031854543 | 0.308350943 |
| *INSL3* | 1.015962 | 0.473508 | 2.145608 | 0.031904292 | 0.308581223 |
| *SLC27A5* | -0.468355 | 0.218338 | -2.145088 | 0.03194581 | 0.308731577 |
| *CDK7* | 0.370387 | 0.172697 | 2.144722 | 0.031975073 | 0.308763356 |
| *POLR3GL* | -0.16475 | 0.076856 | -2.143619 | 0.032063423 | 0.309261925 |
| *DIMT1* | 0.294396 | 0.137361 | 2.14322 | 0.032095422 | 0.309261925 |
| *R3HCC1L* | 0.404618 | 0.1888 | 2.143104 | 0.032104754 | 0.309261925 |
| *PIKFYVE* | 0.282316 | 0.131813 | 2.141794 | 0.032210074 | 0.309826581 |
| *SLC26A1* | 1.239409 | 0.578777 | 2.141428 | 0.0322395 | 0.309826581 |

| | | | | | |
|---|---|---|---|---|---|
| *GNPDA2* | 0.269302 | 0.12576 | 2.141403 | 0.032241565 | 0.309826581 |
| *RXRB* | 0.264809 | 0.1238 | 2.139011 | 0.032434811 | 0.311311519 |
| *SUCLG1* | 0.235742 | 0.11022 | 2.138842 | 0.032448471 | 0.311311519 |
| *PEG10* | -1.067023 | 0.49906 | -2.138066 | 0.032511424 | 0.311663951 |
| *NOP53* | -0.193527 | 0.090594 | -2.136209 | 0.032662376 | 0.312757336 |
| *CNEP1R1* | -0.37629 | 0.176172 | -2.13592 | 0.032685943 | 0.312757336 |
| *TMEM50A* | 0.185324 | 0.086775 | 2.135679 | 0.032705564 | 0.312757336 |
| *ENSG00000280206* | 0.46866 | 0.219484 | 2.135281 | 0.032738098 | 0.312757336 |
| *CSK* | 0.205589 | 0.096292 | 2.135049 | 0.032757036 | 0.312757336 |
| *TCTA* | 0.329409 | 0.1544 | 2.133487 | 0.032884825 | 0.313725452 |
| *U2AF1L4* | -0.223919 | 0.104978 | -2.133016 | 0.032923414 | 0.313841716 |
| *PQLC3* | -0.298234 | 0.139963 | -2.130796 | 0.033105957 | 0.315288519 |
| *SCAF11* | -0.121643 | 0.057096 | -2.130526 | 0.033128238 | 0.315288519 |
| *ZNF432* | -0.566098 | 0.265787 | -2.129892 | 0.033180528 | 0.315533545 |
| *LMO2* | -1.841243 | 0.864648 | -2.12947 | 0.033215399 | NA |
| *H6PD* | -0.393771 | 0.184967 | -2.128875 | 0.033264582 | 0.316064021 |
| *ANK1* | -0.919506 | 0.431982 | -2.128574 | 0.033289489 | 0.316064021 |
| *SH2B3* | 0.421989 | 0.198303 | 2.127994 | 0.033337562 | 0.316267835 |
| *SDR42E1* | -0.955591 | 0.449375 | -2.126487 | 0.033462759 | 0.317173015 |
| *UBE2V2* | 0.15699 | 0.073836 | 2.126203 | 0.033486341 | 0.317173015 |
| *USP4* | 0.251674 | 0.118451 | 2.124706 | 0.033611135 | 0.318011383 |
| *WDR81* | -0.423523 | 0.199352 | -2.1245 | 0.033628359 | 0.318011383 |
| *NT5DC1* | -0.284363 | 0.133948 | -2.122946 | 0.033758366 | 0.318888407 |
| *ITPA* | 0.238906 | 0.112575 | 2.122188 | 0.03382197 | 0.318888407 |
| *UBL7.AS1* | -0.347004 | 0.163516 | -2.122137 | 0.033826264 | 0.318888407 |
| *NOTCH2* | 0.263917 | 0.124369 | 2.122057 | 0.033832927 | 0.318888407 |
| *PPARA* | -0.345469 | 0.162819 | -2.121792 | 0.033855234 | 0.318888407 |
| *LAMA5* | -0.767504 | 0.361894 | -2.120799 | 0.033938712 | 0.319421587 |
| *DCAF5* | -0.279734 | 0.131978 | -2.119541 | 0.03404475 | 0.32016609 |
| *STK4* | -0.141427 | 0.066783 | -2.117697 | 0.034200716 | 0.321260197 |
| *BHLHE40.AS1* | 1.117988 | 0.527968 | 2.117527 | 0.034215143 | 0.321260197 |
| *SAR1B* | 0.235107 | 0.111054 | 2.11705 | 0.034255585 | 0.321386065 |
| *CCDC59* | 0.15085 | 0.071274 | 2.116482 | 0.034303831 | 0.32158489 |
| *CCDC141* | -0.733807 | 0.346769 | -2.116124 | 0.034334261 | 0.321616517 |
| *KLC2* | -0.415114 | 0.196292 | -2.114779 | 0.034448802 | 0.322369133 |
| *PHB2* | -0.219721 | 0.103909 | -2.114544 | 0.034468846 | 0.322369133 |
| *JARID2* | -0.260249 | 0.123166 | -2.112992 | 0.034601419 | 0.323354617 |
| *WDR54* | -0.219666 | 0.104018 | -2.111805 | 0.034703193 | 0.324050948 |
| *ZNF616* | 0.465721 | 0.220643 | 2.110743 | 0.034794416 | 0.324647744 |
| *TLE1* | 0.644997 | 0.305782 | 2.109338 | 0.034915387 | 0.325297269 |
| *ENSG00000180448* | 0.256264 | 0.121501 | 2.109152 | 0.034931503 | 0.325297269 |
| *MICAL3* | -0.222643 | 0.105573 | -2.108895 | 0.034953687 | 0.325297269 |
| *LTV1* | 0.279383 | 0.132507 | 2.108437 | 0.034993234 | 0.325297269 |
| *PDHB* | 0.187156 | 0.088773 | 2.10826 | 0.035008481 | 0.325297269 |
| *ERCC5* | 0.309739 | 0.146933 | 2.108032 | 0.035028225 | 0.325297269 |
| *SHISA5* | 0.201499 | 0.095674 | 2.106097 | 0.035195913 | 0.326570234 |
| *PSENEN* | 0.196835 | 0.093472 | 2.105817 | 0.035220244 | 0.326570234 |

| | | | | | |
|---|---|---|---|---|---|
| *CDKN2D* | 0.242431 | 0.115188 | 2.10465 | 0.035321774 | 0.327256373 |
| *HIST1H4E* | 0.630377 | 0.299649 | 2.103716 | 0.035403251 | 0.327755799 |
| *AP1S3* | -0.470892 | 0.223874 | -2.10338 | 0.03543259 | 0.327772135 |
| *PRKRA.AS1* | -0.466277 | 0.221763 | -2.102593 | 0.03550138 | 0.32815311 |
| *MAGED1* | -0.590218 | 0.280874 | -2.101366 | 0.035608872 | 0.328709533 |
| *ZMYND11* | -0.460651 | 0.21928 | -2.100743 | 0.035663544 | 0.328709533 |
| *HSP90B1* | 0.194311 | 0.092499 | 2.100689 | 0.035668236 | 0.328709533 |
| *PARP10* | 0.281275 | 0.133899 | 2.100644 | 0.035672188 | 0.328709533 |
| *ZNF736* | -0.409225 | 0.194866 | -2.100038 | 0.035725466 | 0.328945478 |
| *EIF4EBP3* | 0.610618 | 0.290842 | 2.099482 | 0.035774412 | 0.329044957 |
| *UTRN* | -0.328975 | 0.156737 | -2.098902 | 0.03582551 | 0.329044957 |
| *MYO5A* | -0.376894 | 0.179586 | -2.098685 | 0.035844691 | 0.329044957 |
| *LINC00426* | -0.591129 | 0.28167 | -2.098659 | 0.035846994 | 0.329044957 |
| *ENSG00000272211* | -0.608261 | 0.289981 | -2.097592 | 0.035941213 | 0.329628698 |
| *SLC9A7* | 0.272433 | 0.129896 | 2.097311 | 0.035966049 | 0.329628698 |
| *RIPK2* | 0.258272 | 0.123187 | 2.096575 | 0.036031181 | 0.329971228 |
| *SF3B5* | 0.160961 | 0.076817 | 2.09539 | 0.036136316 | 0.33047394 |
| *DARS* | 0.156958 | 0.074908 | 2.09533 | 0.036141678 | 0.33047394 |
| *ERGIC3* | -0.201142 | 0.096061 | -2.093885 | 0.036270234 | 0.331274425 |
| *CCNT2* | -0.253883 | 0.121259 | -2.09372 | 0.036284958 | 0.331274425 |
| *NSMCE2* | 0.241816 | 0.115645 | 2.091014 | 0.036526809 | 0.333085423 |
| *ZNF689* | 0.330075 | 0.157865 | 2.090874 | 0.036539362 | 0.333085423 |
| *LINC02245* | -0.688339 | 0.329309 | -2.090253 | 0.036595056 | 0.333337495 |
| *UPP1* | 0.502321 | 0.240426 | 2.089291 | 0.03668158 | 0.333665521 |
| *ENSG00000266378* | -1.305069 | 0.624713 | -2.089068 | 0.036701562 | 0.333665521 |
| *CCNJ* | -0.718751 | 0.344079 | -2.088916 | 0.036715277 | 0.333665521 |
| *IFI35* | 0.263997 | 0.126423 | 2.088198 | 0.036779961 | 0.333998008 |
| *LARP7* | 0.146255 | 0.070086 | 2.086797 | 0.036906533 | 0.334891574 |
| *PIK3IP1* | 0.276289 | 0.132428 | 2.08633 | 0.036948748 | 0.335008068 |
| *CLK4* | -0.242209 | 0.116125 | -2.085758 | 0.037000529 | 0.335008068 |
| *EXOSC3* | 0.210881 | 0.101121 | 2.085439 | 0.037029511 | 0.335008068 |
| *COX7A2L* | -0.121577 | 0.058299 | -2.08541 | 0.037032102 | 0.335008068 |
| *RTL6* | -0.68457 | 0.328426 | -2.084395 | 0.037124224 | 0.335586049 |
| *STX12* | 0.192456 | 0.092362 | 2.083718 | 0.037185784 | 0.335887097 |
| *APOBEC3B* | 1.099998 | 0.528045 | 2.083152 | 0.037237344 | 0.335927213 |
| *KCTD20* | -0.245413 | 0.117859 | -2.082255 | 0.037319222 | 0.335927213 |
| *SLC25A11* | 0.20035 | 0.096226 | 2.082071 | 0.037336028 | 0.335927213 |
| *PPARD* | -0.398691 | 0.191513 | -2.081791 | 0.037361537 | 0.335927213 |
| *ENSG00000228172* | 1.188341 | 0.570851 | 2.081701 | 0.037369804 | 0.335927213 |
| *RAE1* | 0.300161 | 0.144202 | 2.081534 | 0.03738503 | 0.335927213 |
| *GDE1* | -0.270803 | 0.1301 | -2.081501 | 0.037388046 | 0.335927213 |
| *AKIRIN1* | -0.171988 | 0.082649 | -2.080944 | 0.037438987 | 0.336130842 |
| *CHRNA6* | -1.263293 | 0.607253 | -2.080342 | 0.037494182 | 0.336340198 |
| *ENSG00000269549* | 0.893758 | 0.429731 | 2.079806 | 0.037543318 | 0.336340198 |
| *PSMB2* | 0.163 | 0.078374 | 2.079764 | 0.03754719 | 0.336340198 |
| *PEX1* | -0.38731 | 0.186317 | -2.078765 | 0.037638912 | 0.336907943 |
| *PRCC* | 0.203321 | 0.097832 | 2.078257 | 0.037685652 | 0.337072495 |

| | | | | | |
|---|---|---|---|---|---|
| *PARP15* | -0.276276 | 0.13301 | -2.077107 | 0.037791688 | 0.337700274 |
| *ZBTB8OS* | 0.196256 | 0.094495 | 2.07688 | 0.037812658 | 0.337700274 |
| *GOLGA8B* | -0.286326 | 0.1379 | -2.076324 | 0.037864015 | 0.337905066 |
| *AHNAK* | -0.468356 | 0.22568 | -2.075312 | 0.037957619 | 0.338267044 |
| *QSOX1* | 0.534408 | 0.257512 | 2.07527 | 0.03796149 | 0.338267044 |
| *TXLNA* | -0.308238 | 0.148568 | -2.07472 | 0.03801248 | 0.338467681 |
| *ENSG00000233184* | -0.436776 | 0.210573 | -2.074228 | 0.038058133 | 0.338552517 |
| *IMPA2* | -0.598862 | 0.288747 | -2.074004 | 0.03807897 | 0.338552517 |
| *ENSG00000267136* | 1.215255 | 0.586191 | 2.073137 | 0.038159498 | NA |
| *HES6* | -0.515911 | 0.248873 | -2.072989 | 0.038173286 | 0.339137406 |
| *SDHB* | 0.229595 | 0.110814 | 2.071886 | 0.038276082 | 0.339796708 |
| *CETP* | -0.876725 | 0.423253 | -2.0714 | 0.03832146 | 0.33980827 |
| *RPL14* | -0.186989 | 0.090292 | -2.070936 | 0.038364823 | 0.33980827 |
| *C12orf10* | -0.240165 | 0.115987 | -2.070616 | 0.038394727 | 0.33980827 |
| *SORD* | -0.446479 | 0.215664 | -2.070248 | 0.038429114 | 0.33980827 |
| *ATP2C1* | -0.32316 | 0.156112 | -2.070051 | 0.038447529 | 0.33980827 |
| *GTF2F1* | -0.185731 | 0.089724 | -2.070037 | 0.038448904 | 0.33980827 |
| *ANP32B* | -0.133121 | 0.064328 | -2.069391 | 0.03850938 | 0.339962398 |
| *ZEB1.AS1* | -0.470035 | 0.227173 | -2.069064 | 0.038540113 | 0.339962398 |
| *BRD3* | -0.216064 | 0.104433 | -2.068935 | 0.038552142 | 0.339962398 |
| *TK2* | -1.306241 | 0.631555 | -2.068293 | 0.038612437 | 0.340241688 |
| *TRIAP1* | 0.233044 | 0.112715 | 2.067549 | 0.038682452 | 0.340606156 |
| *SLIRP* | 0.138357 | 0.066944 | 2.066767 | 0.038756083 | 0.341001894 |
| *KCNH4* | 1.383732 | 0.669898 | 2.065586 | 0.03886763 | NA |
| *SSR3* | -0.170902 | 0.082743 | -2.065448 | 0.038880663 | 0.341681158 |
| *DNAJC28* | -0.815473 | 0.394837 | -2.065341 | 0.038890772 | 0.341681158 |
| *GNAI2* | -0.186525 | 0.09033 | -2.064938 | 0.038928849 | 0.341763094 |
| *PCTP* | 0.350667 | 0.169884 | 2.064157 | 0.039002808 | 0.342159691 |
| *SLC37A1* | -0.332956 | 0.161394 | -2.062993 | 0.039113251 | 0.342875525 |
| *PKN1* | 0.238791 | 0.115781 | 2.062442 | 0.039165706 | 0.343082353 |
| *ZNF675* | 0.306525 | 0.148651 | 2.062053 | 0.039202689 | 0.343153438 |
| *FAM111B* | -0.325856 | 0.158076 | -2.061389 | 0.039265903 | 0.343453854 |
| *MRPS10* | 0.167472 | 0.081255 | 2.061059 | 0.039297437 | 0.343476934 |
| *TGOLN2* | -0.160973 | 0.07815 | -2.059796 | 0.039418015 | 0.344277693 |
| *OSTM1* | 0.258631 | 0.125604 | 2.059109 | 0.039483834 | 0.344599368 |
| *CENPJ* | -0.376578 | 0.182955 | -2.058314 | 0.039560033 | 0.344686459 |
| *HNRNPR* | 0.181934 | 0.0884 | 2.058079 | 0.03958251 | 0.344686459 |
| *MEF2D* | -0.318054 | 0.154553 | -2.057892 | 0.039600508 | 0.344686459 |
| *FUOM* | 0.400455 | 0.194606 | 2.057774 | 0.03961179 | 0.344686459 |
| *LIG1* | -0.347329 | 0.168812 | -2.057493 | 0.039638798 | 0.344686459 |
| *SLC35F5* | -0.465916 | 0.226575 | -2.056341 | 0.039749648 | 0.345275544 |
| *RYK* | -0.359689 | 0.174944 | -2.056024 | 0.03978017 | 0.345275544 |
| *C15orf62* | -0.458929 | 0.223262 | -2.055559 | 0.039825015 | 0.345275544 |
| *TRAPPC4* | 0.217752 | 0.105937 | 2.055487 | 0.039831951 | 0.345275544 |
| *ENSG00000268713* | -0.666326 | 0.324202 | -2.055282 | 0.039851775 | 0.345275544 |
| *CYB5R4* | 0.212227 | 0.103297 | 2.05453 | 0.039924467 | 0.345453989 |
| *TMCO1* | 0.170405 | 0.08295 | 2.0543 | 0.039946673 | 0.345453989 |

| | | | | | |
|---|---|---|---|---|---|
| *ENSG00000272871* | -1.041617 | 0.507075 | -2.054167 | 0.039959555 | 0.345453989 |
| *ARF4* | 0.242423 | 0.118109 | 2.052537 | 0.040117526 | 0.346253356 |
| *KLHDC1* | -0.693823 | 0.338073 | -2.052288 | 0.04014172 | 0.346253356 |
| *EAF2* | 0.291498 | 0.142051 | 2.052065 | 0.04016335 | 0.346253356 |
| *CCNG2* | -0.340182 | 0.16578 | -2.052012 | 0.040168535 | 0.346253356 |
| *TCEAL8* | -0.224727 | 0.109539 | -2.051564 | 0.040212072 | 0.346377461 |
| *BRD3OS* | -0.404626 | 0.197309 | -2.050728 | 0.040293471 | 0.346827292 |
| *QDPR* | -0.573674 | 0.279976 | -2.049009 | 0.040461231 | 0.347956971 |
| *NKX6.3* | -1.90625 | 0.930609 | -2.048389 | 0.040521889 | NA |
| *OCIAD1* | 0.159066 | 0.077663 | 2.048155 | 0.040544815 | 0.347956971 |
| *CELF2* | -0.224732 | 0.109726 | -2.048125 | 0.040547756 | 0.347956971 |
| *DIPK1A* | 0.380963 | 0.186009 | 2.048094 | 0.040550827 | 0.347956971 |
| *AKR7A2* | -0.208159 | 0.101646 | -2.047887 | 0.040571074 | 0.347956971 |
| *ATP2B1.AS1* | -0.28007 | 0.136793 | -2.047403 | 0.040618514 | 0.348112673 |
| *ZC3H15* | 0.122926 | 0.0601 | 2.045369 | 0.040818481 | 0.349574413 |
| *RIOK1* | 0.286957 | 0.140317 | 2.045067 | 0.040848279 | 0.34957775 |
| *NDUFS7* | 0.180358 | 0.088236 | 2.044043 | 0.040949279 | 0.350009595 |
| *RBM8A* | 0.121729 | 0.059563 | 2.043705 | 0.040982701 | 0.350009595 |
| *NTPCR* | 0.222213 | 0.108764 | 2.043072 | 0.041045282 | 0.350009595 |
| *HIVEP1* | -0.381666 | 0.186817 | -2.042992 | 0.041053253 | 0.350009595 |
| *TEX2* | -0.906359 | 0.44376 | -2.042453 | 0.041106597 | 0.350009595 |
| *IFNAR2* | 0.209682 | 0.102665 | 2.042395 | 0.041112413 | 0.350009595 |
| *EP400P1* | -0.386763 | 0.189385 | -2.042208 | 0.041130875 | 0.350009595 |
| *ERN1* | -0.918301 | 0.449668 | -2.042174 | 0.041134298 | 0.350009595 |
| *WDR60* | -0.42057 | 0.206002 | -2.041585 | 0.041192687 | 0.350255699 |
| *ESYT1* | -0.256525 | 0.125725 | -2.040368 | 0.041313707 | 0.351033622 |
| *DIP2C* | -0.977516 | 0.479331 | -2.039334 | 0.041416749 | 0.351140548 |
| *DHRS4.AS1* | -0.361526 | 0.177304 | -2.039016 | 0.041448436 | 0.351140548 |
| *MAMSTR* | 0.648895 | 0.318305 | 2.038595 | 0.041490481 | 0.351140548 |
| *TCTN1* | -0.618864 | 0.303608 | -2.038365 | 0.041513425 | 0.351140548 |
| *TRIB1* | 0.608794 | 0.298687 | 2.038234 | 0.04152654 | 0.351140548 |
| *TASP1* | 0.351618 | 0.172535 | 2.037951 | 0.041554823 | 0.351140548 |
| *EHMT1* | -0.286459 | 0.140594 | -2.037493 | 0.041600671 | 0.351140548 |
| *SLC12A6* | 0.214551 | 0.105311 | 2.037315 | 0.041618464 | 0.351140548 |
| *SOGA1* | -0.455843 | 0.223796 | -2.036868 | 0.041663303 | 0.351140548 |
| *USF3* | -0.269251 | 0.132194 | -2.03679 | 0.041671106 | 0.351140548 |
| *RPS2* | -0.368212 | 0.180789 | -2.036692 | 0.041680904 | 0.351140548 |
| *ZC3H6* | -0.428493 | 0.210391 | -2.03665 | 0.041685079 | 0.351140548 |
| *AMT* | 0.798641 | 0.392203 | 2.036296 | 0.041720655 | 0.351140548 |
| *COX16* | 0.25764 | 0.126536 | 2.036105 | 0.04173985 | 0.351140548 |
| *TRIB2* | -0.733113 | 0.36014 | -2.035632 | 0.041787351 | 0.35129154 |
| *ENSG00000272004* | 1.124426 | 0.552502 | 2.035152 | 0.041835627 | NA |
| *CAPZB* | 0.146661 | 0.072067 | 2.035057 | 0.041845131 | 0.351528671 |
| *OAT* | 0.405567 | 0.199347 | 2.034478 | 0.041903447 | 0.351769969 |
| *AZIN1.AS1* | -1.095852 | 0.53899 | -2.033157 | 0.042036646 | 0.352503521 |
| *SUCLG2* | -0.253852 | 0.124864 | -2.033024 | 0.042050138 | 0.352503521 |
| *NAPB* | 0.376976 | 0.185483 | 2.032404 | 0.042112819 | 0.352780182 |

| | | | | | |
|---|---|---|---|---|---|
| *RNASE6* | -0.989079 | 0.4868 | -2.031797 | 0.042174255 | 0.352978843 |
| *LINC01869* | -1.154229 | 0.568143 | -2.031583 | 0.042195923 | 0.352978843 |
| *SLC25A32* | 0.285056 | 0.140366 | 2.030798 | 0.042275524 | 0.353396031 |
| *ZNF79* | -0.807044 | 0.397462 | -2.030494 | 0.042306362 | 0.353405291 |
| *BAG6* | 0.211423 | 0.104176 | 2.029485 | 0.042408957 | 0.353850809 |
| *ENSG00000246528* | 0.895395 | 0.441268 | 2.02914 | 0.042444037 | 0.353850809 |
| *RBM41* | -0.245168 | 0.120865 | -2.028445 | 0.042514891 | 0.353850809 |
| *IKZF4* | -1.263898 | 0.623105 | -2.028386 | 0.042520891 | 0.353850809 |
| *ENSG00000228106* | 0.248537 | 0.122537 | 2.028262 | 0.042533562 | 0.353850809 |
| *TP53INP1* | -0.328271 | 0.161852 | -2.028215 | 0.042538303 | 0.353850809 |
| *TBCK* | -0.375747 | 0.185389 | -2.026807 | 0.042682143 | 0.354214083 |
| *TAP1* | 0.212508 | 0.104853 | 2.02672 | 0.042691048 | 0.354214083 |
| *ENSG00000238260* | 1.101881 | 0.543707 | 2.026608 | 0.042702556 | 0.354214083 |
| *CFAP298* | 0.223407 | 0.110237 | 2.026604 | 0.042702905 | 0.354214083 |
| *CREM* | -0.463104 | 0.228573 | -2.026068 | 0.042757862 | 0.354214083 |
| *ERO1A* | -0.326673 | 0.161239 | -2.026024 | 0.042762295 | 0.354214083 |
| *PRX* | 1.190932 | 0.587836 | 2.025958 | 0.042769101 | NA |
| *HLA.DRA* | -0.306756 | 0.151438 | -2.025622 | 0.042803556 | 0.354214083 |
| *TRPS1* | -0.388151 | 0.191636 | -2.025458 | 0.042820361 | 0.354214083 |
| *ABRAXAS2* | 0.260172 | 0.128517 | 2.024407 | 0.042928262 | 0.354266099 |
| *EIF2S1* | 0.247317 | 0.122179 | 2.024228 | 0.042946735 | 0.354266099 |
| *NDUFS1* | 0.310983 | 0.153634 | 2.02418 | 0.042951618 | 0.354266099 |
| *ARMH1* | -0.716618 | 0.354113 | -2.023697 | 0.043001363 | 0.354266099 |
| *TAPT1* | -0.451551 | 0.223151 | -2.023522 | 0.04301932 | 0.354266099 |
| *NSMF* | -0.594375 | 0.293812 | -2.022975 | 0.043075715 | 0.354266099 |
| *GOLGA6L9* | -0.936386 | 0.462911 | -2.022823 | 0.043091418 | 0.354266099 |
| *ZDHHC13* | -0.331561 | 0.163963 | -2.022174 | 0.043158403 | 0.354266099 |
| *ODF2L* | -0.193263 | 0.095581 | -2.021982 | 0.043178205 | 0.354266099 |
| *RPL21* | -0.188583 | 0.093268 | -2.021953 | 0.043181162 | 0.354266099 |
| *POGK* | -0.230507 | 0.114003 | -2.021929 | 0.043183638 | 0.354266099 |
| *SND1* | -0.373334 | 0.184643 | -2.021923 | 0.043184283 | 0.354266099 |
| *CA13* | -0.589883 | 0.291959 | -2.020433 | 0.043338545 | 0.354878137 |
| *YJEFN3* | 0.533388 | 0.264002 | 2.020393 | 0.043342615 | 0.354878137 |
| *EGLN2* | -0.173496 | 0.085881 | -2.020191 | 0.043363564 | 0.354878137 |
| *ERCC4* | -0.354318 | 0.175432 | -2.019684 | 0.043416143 | 0.354878137 |
| *NBN* | 0.260381 | 0.128943 | 2.019349 | 0.043450907 | 0.354878137 |
| *CAMSAP1* | -0.452306 | 0.224041 | -2.018853 | 0.043502492 | 0.354878137 |
| *MRPL33* | 0.159692 | 0.079103 | 2.018782 | 0.043509868 | 0.354878137 |
| *ATF7IP* | -0.292289 | 0.144787 | -2.018752 | 0.04351299 | 0.354878137 |
| *KCTD17* | -0.346442 | 0.171624 | -2.018612 | 0.043527578 | 0.354878137 |
| *DHX15* | 0.251035 | 0.124488 | 2.01654 | 0.043743541 | 0.35639443 |
| *SLC33A1* | -0.341214 | 0.169432 | -2.013871 | 0.044023097 | 0.358426409 |
| *MRPL24* | 0.279542 | 0.138851 | 2.013255 | 0.044087862 | 0.358569801 |
| *CCDC92* | -0.360833 | 0.17924 | -2.013129 | 0.044101039 | 0.358569801 |
| *ENSG00000226266* | -0.95921 | 0.476573 | -2.012725 | 0.044143543 | 0.358670055 |
| *MICAL1* | -0.294233 | 0.146217 | -2.012304 | 0.044187904 | 0.358785253 |
| *CRNKL1* | 0.27215 | 0.135358 | 2.010587 | 0.044369135 | 0.360010861 |

| | | | | | |
|---|---|---|---|---|---|
| *ENSG00000254802* | 0.62573 | 0.311311 | 2.009984 | 0.044432885 | 0.360026991 |
| *MIR222HG* | 0.940383 | 0.467911 | 2.009748 | 0.044457844 | 0.360026991 |
| *NUFIP2* | -0.247482 | 0.12316 | -2.009441 | 0.044490353 | 0.360026991 |
| *FOS* | -0.865222 | 0.430582 | -2.009423 | 0.044492273 | 0.360026991 |
| *SLC25A1* | -0.314987 | 0.156807 | -2.008754 | 0.044563273 | 0.360356208 |
| *USP15* | -0.159488 | 0.079456 | -2.007239 | 0.044724259 | 0.361192081 |
| *C1GALT1C1* | 0.375574 | 0.187113 | 2.007209 | 0.044727412 | 0.361192081 |
| *GUCY2C* | -0.441528 | 0.220146 | -2.005609 | 0.044898007 | 0.362323567 |
| *IDS* | -0.157463 | 0.078576 | -2.003956 | 0.045074788 | 0.363426646 |
| *OAS3* | 0.557196 | 0.278075 | 2.003759 | 0.045095844 | 0.363426646 |
| *SMARCD1* | -0.30328 | 0.151394 | -2.003254 | 0.04515005 | 0.363616969 |
| *WDR33* | 0.184703 | 0.092262 | 2.00195 | 0.045290078 | 0.364497733 |
| *RPS3A* | -0.168451 | 0.084225 | -2.000004 | 0.045499791 | 0.36539207 |
| *KLHL28* | -0.264663 | 0.132353 | -1.999678 | 0.045535083 | 0.36539207 |
| *ENSG00000279278* | 0.447282 | 0.223683 | 1.999627 | 0.045540598 | 0.36539207 |
| *GBA2* | -0.302281 | 0.151188 | -1.999371 | 0.045568247 | 0.36539207 |
| *ZFYVE16* | -0.26293 | 0.131514 | -1.999256 | 0.045580677 | 0.36539207 |
| *USF2* | -0.206555 | 0.10333 | -1.99898 | 0.045610511 | 0.36539207 |
| *ANKZF1* | -0.184065 | 0.092082 | -1.998926 | 0.045616373 | 0.36539207 |
| *SLC9A9* | 0.456931 | 0.228668 | 1.998224 | 0.045692379 | 0.365716514 |
| *TMEM183A* | -0.158244 | 0.07921 | -1.997765 | 0.045742155 | 0.365716514 |
| *BTG3* | 0.214225 | 0.107236 | 1.9977 | 0.045749176 | 0.365716514 |
| *CD82* | -0.193488 | 0.096885 | -1.99709 | 0.045815358 | 0.365999436 |
| *LY86.AS1* | -1.557779 | 0.780033 | -1.997067 | 0.04581787 | NA |
| *SMAD2* | -0.206986 | 0.103672 | -1.996539 | 0.045875324 | 0.366232352 |
| *UBLCP1* | -0.280951 | 0.140754 | -1.99604 | 0.045929519 | 0.366418923 |
| *ENTPD7* | -2.008404 | 1.006294 | -1.995841 | 0.045951189 | NA |
| *ENSG00000272918* | -0.478537 | 0.239788 | -1.995668 | 0.045970114 | 0.366496812 |
| *SLC48A1* | -0.320201 | 0.160499 | -1.995033 | 0.046039276 | 0.366802195 |
| *XYLT2* | -0.47538 | 0.238345 | -1.994508 | 0.046096618 | 0.367013056 |
| *ENSG00000224905* | -1.401524 | 0.704032 | -1.990712 | 0.046512606 | NA |
| *FAM107B* | 0.19089 | 0.095903 | 1.990437 | 0.04654277 | 0.370317208 |
| *ENSG00000273156* | -0.601679 | 0.302398 | -1.989693 | 0.046624796 | 0.370634809 |
| *MRTO4* | 0.228596 | 0.114901 | 1.989509 | 0.046645047 | 0.370634809 |
| *TLK1* | -0.172144 | 0.08661 | -1.987575 | 0.046858764 | 0.372084253 |
| *GLYCTK* | 0.4753 | 0.239288 | 1.98631 | 0.046998933 | 0.37294814 |
| *TANC2* | -0.779969 | 0.392751 | -1.98591 | 0.047043291 | 0.373051103 |
| *CHD4* | -0.169751 | 0.08553 | -1.984702 | 0.047177649 | 0.373867143 |
| *CCNL2* | -0.191668 | 0.096589 | -1.984363 | 0.047215433 | 0.373917289 |
| *TANGO2* | -0.341974 | 0.172375 | -1.983891 | 0.047268048 | 0.374084741 |
| *XPC* | 0.173153 | 0.087312 | 1.983152 | 0.047350421 | 0.374487326 |
| *ANKMY1* | 0.377909 | 0.19064 | 1.982317 | 0.047443733 | 0.374975835 |
| *HACD1* | -0.976264 | 0.492558 | -1.982027 | 0.047476184 | 0.374982987 |
| *SRP68* | 0.268289 | 0.135444 | 1.980811 | 0.047612427 | 0.375773158 |
| *AGRN* | -0.673912 | 0.340262 | -1.980568 | 0.047639788 | 0.375773158 |
| *ZMYM6* | -0.30643 | 0.15474 | -1.980289 | 0.047671063 | 0.375773158 |
| *ENSG00000237310* | 0.540441 | 0.272971 | 1.97985 | 0.04772034 | 0.375912317 |

| | | | | | |
|---|---|---|---|---|---|
| *AMD1* | -0.24024 | 0.121422 | -1.978546 | 0.047867106 | 0.376818733 |
| *ENSG00000245522* | 0.656337 | 0.331914 | 1.977431 | 0.047992987 | 0.377428826 |
| *LINC01800* | -1.109225 | 0.560981 | -1.977297 | 0.048008108 | 0.377428826 |
| *FAM160B2* | -0.344348 | 0.174202 | -1.976718 | 0.048073533 | 0.377693382 |
| *ABRAXAS1* | -0.182855 | 0.092618 | -1.974289 | 0.048348843 | 0.379312316 |
| *BCLAF3* | -0.521934 | 0.26441 | -1.973956 | 0.048386783 | 0.379312316 |
| *DIAPH1* | -0.258306 | 0.130883 | -1.973568 | 0.048430842 | 0.379312316 |
| *STAG2* | 0.175673 | 0.08902 | 1.973404 | 0.048449517 | 0.379312316 |
| *LFNG* | 0.286662 | 0.145267 | 1.973345 | 0.048456314 | 0.379312316 |
| *PABPC3* | -0.708831 | 0.359227 | -1.973209 | 0.048471702 | 0.379312316 |
| *GLIPR1* | -0.223762 | 0.113419 | -1.972882 | 0.048508988 | 0.379312316 |
| *PHF8* | 0.338162 | 0.171425 | 1.972655 | 0.048534873 | 0.379312316 |
| *RALGPS1* | -0.411049 | 0.208429 | -1.972131 | 0.048594664 | 0.379530075 |
| *CD68* | -0.481184 | 0.244029 | -1.971834 | 0.048628608 | 0.379545803 |
| *ZNF165* | 0.46312 | 0.234903 | 1.971537 | 0.048662538 | 0.379561413 |
| *NDUFA5* | 0.130256 | 0.066082 | 1.971131 | 0.048708916 | 0.379659431 |
| *FAM122A* | 0.210403 | 0.106775 | 1.970533 | 0.048777322 | 0.379659431 |
| *CPM* | 0.477639 | 0.242416 | 1.970329 | 0.048800675 | 0.379659431 |
| *CCDC71L* | -0.451899 | 0.229387 | -1.970033 | 0.048834557 | 0.379659431 |
| *ZNF227* | 0.299428 | 0.151992 | 1.970031 | 0.0488348 | 0.379659431 |
| *ZNF493* | -0.200018 | 0.101578 | -1.969112 | 0.048940236 | 0.380230446 |
| *XPO5* | 0.438253 | 0.222609 | 1.968712 | 0.04898619 | 0.380332696 |
| *LMF2* | -0.196861 | 0.100019 | -1.968229 | 0.049041704 | 0.380332696 |
| *RBM28* | 0.230145 | 0.116934 | 1.968162 | 0.049049384 | 0.380332696 |
| *MED29* | 0.188213 | 0.095649 | 1.967743 | 0.049097563 | 0.380458106 |
| *GSTM4* | -0.501431 | 0.255025 | -1.966203 | 0.04927522 | 0.381586016 |
| *ZBTB3* | 0.489601 | 0.24905 | 1.965873 | 0.04931333 | 0.381632523 |
| *LRRFIP1* | -0.206278 | 0.104952 | -1.965456 | 0.049361455 | 0.381756415 |

| Cluster 4 | log2FC | lfcSE | stat | pvalue | padj |
|---|---|---|---|---|---|
| *VPREB3* | 2.019106 | 0.072711 | 27.769031 | < 2.22e-16 | < 2.22e-16 |
| *PLAAT4* | 1.838477 | 0.168514 | 10.909953 | < 2.22e-16 | < 2.22e-16 |
| *CD38* | 2.041542 | 0.242906 | 8.404655 | < 2.22e-16 | 1.63E-13 |
| *AHI1* | 1.6348 | 0.213267 | 7.665507 | 1.78E-14 | 5.07E-11 |
| *MS4A1* | -0.681183 | 0.102817 | -6.625227 | 3.47E-11 | 7.90E-08 |
| *DENND6B* | 1.924781 | 0.295721 | 6.508777 | 7.58E-11 | 1.31E-07 |
| *RRBP1* | 1.118218 | 0.172054 | 6.499241 | 8.07E-11 | 1.31E-07 |
| *MRPS6* | 0.98125 | 0.151477 | 6.477872 | 9.30E-11 | 1.32E-07 |
| *GRN* | 1.188852 | 0.192162 | 6.18673 | 6.14E-10 | 7.77E-07 |
| *SEPTIN9* | 0.623549 | 0.106302 | 5.865849 | 4.47E-09 | 5.09E-06 |
| *CD1D* | 2.324307 | 0.413927 | 5.615264 | 1.96E-08 | 2.03E-05 |
| *DPEP2* | 1.100664 | 0.200651 | 5.485457 | 4.12E-08 | 3.91E-05 |
| *SIGIRR* | 0.996736 | 0.186979 | 5.330728 | 9.78E-08 | 8.57E-05 |
| *SLC12A4* | 3.593533 | 0.687828 | 5.224465 | 1.75E-07 | 0.000142061 |
| *AGO1* | 1.414221 | 0.272122 | 5.197005 | 2.03E-07 | 0.000153744 |
| *DUS2* | 1.411895 | 0.279878 | 5.044688 | 4.54E-07 | 0.000301683 |
| *DBNL* | 0.727911 | 0.144438 | 5.039591 | 4.67E-07 | 0.000301683 |
| *TRABD* | 0.647435 | 0.128577 | 5.035387 | 4.77E-07 | 0.000301683 |
| *SMAD3* | 1.343852 | 0.268596 | 5.003248 | 5.64E-07 | 0.000337849 |
| *ASB2* | 3.76393 | 0.755296 | 4.983386 | 6.25E-07 | 0.000355737 |
| *BTG2* | -0.355278 | 0.071452 | -4.972278 | 6.62E-07 | 0.000358803 |
| *RARA.AS1* | 3.047223 | 0.621406 | 4.903755 | 9.40E-07 | 0.000486648 |
| *DDIT4* | 1.723666 | 0.352862 | 4.884815 | 1.04E-06 | 0.000512543 |
| *P2RX5* | 0.575582 | 0.131412 | 4.379999 | 1.19E-05 | 0.005630863 |
| *FAM3C* | 0.754908 | 0.173038 | 4.362661 | 1.28E-05 | 0.005757808 |
| *CABLES1* | -1.88776 | 0.433206 | -4.357647 | 1.31E-05 | 0.005757808 |
| *VPREB1* | 3.446085 | 0.796029 | 4.329094 | 1.50E-05 | 0.006314477 |
| *IRAK2* | 1.78763 | 0.414715 | 4.310504 | 1.63E-05 | 0.006579526 |
| *EVI2B* | 0.472179 | 0.109701 | 4.304233 | 1.68E-05 | 0.006579526 |
| *MCM5* | 0.862363 | 0.203026 | 4.247552 | 2.16E-05 | 0.00820317 |
| *RCBTB2* | -2.644794 | 0.628014 | -4.211362 | 2.54E-05 | 0.009323965 |
| *GNG3* | -2.84684 | 0.688089 | -4.137312 | 3.51E-05 | 0.01250427 |
| *APOL3* | 1.053283 | 0.25726 | 4.094232 | 4.24E-05 | 0.014615757 |
| *AGPAT5* | 0.772669 | 0.189625 | 4.074713 | 4.61E-05 | 0.015086889 |
| *SLC5A3* | 0.983941 | 0.241909 | 4.067407 | 4.75E-05 | 0.015086889 |
| *NEIL1* | -0.648514 | 0.159472 | -4.066633 | 4.77E-05 | 0.015086889 |
| *CD79B* | 0.281948 | 0.069999 | 4.027874 | 5.63E-05 | 0.01732164 |
| *FAM117A* | 0.955156 | 0.238447 | 4.005729 | 6.18E-05 | 0.017991356 |
| *DGKD* | -0.658789 | 0.164579 | -4.002875 | 6.26E-05 | 0.017991356 |
| *H3F3A* | 0.353311 | 0.088463 | 3.993899 | 6.50E-05 | 0.017991356 |
| *TTC39C* | -0.947967 | 0.237479 | -3.991789 | 6.56E-05 | 0.017991356 |
| *CHST15* | 1.032448 | 0.258825 | 3.988974 | 6.64E-05 | 0.017991356 |
| *MCTP2* | -0.936432 | 0.235139 | -3.982456 | 6.82E-05 | 0.018062122 |
| *GSTP1* | 0.320295 | 0.080715 | 3.96823 | 7.24E-05 | 0.018739005 |
| *RAB37* | 1.052529 | 0.268594 | 3.918661 | 8.90E-05 | 0.022531676 |
| *PLAAT3* | 2.539781 | 0.652036 | 3.895155 | 9.81E-05 | 0.024232958 |

| | | | | | |
|---|---|---|---|---|---|
| *STK17B* | -0.421192 | 0.108261 | -3.890539 | 0.00010002 | 0.024232958 |
| *PNOC* | 1.045482 | 0.27215 | 3.841565 | 0.00012225 | 0.028701033 |
| *LINC01353* | -1.776717 | 0.4628 | -3.839063 | 0.00012351 | 0.028701033 |
| *RMDN2* | 2.010232 | 0.529096 | 3.799368 | 0.00014507 | 0.033037293 |
| *SPOCK2* | 0.624823 | 0.166368 | 3.755667 | 0.00017288 | 0.038467374 |
| *TMC8* | 0.861967 | 0.229756 | 3.751663 | 0.00017567 | 0.038467374 |
| *CD24* | -0.498975 | 0.133813 | -3.7289 | 0.00019232 | 0.041319156 |
| *ISG20* | -0.41323 | 0.111768 | -3.697224 | 0.00021797 | 0.045963454 |
| *P2RX5.TAX1BP3* | 2.092047 | 0.567045 | 3.689383 | 0.0002248 | 0.046541448 |
| *HHIP.AS1* | -2.512868 | 0.685064 | -3.668079 | 0.00024438 | 0.049692002 |
| *MAP3K1* | 0.707603 | 0.194228 | 3.643161 | 0.00026931 | 0.053800529 |
| *ZNF3* | 1.02196 | 0.281769 | 3.626944 | 0.0002868 | 0.056070273 |
| *XYLT1* | 1.251459 | 0.345885 | 3.618131 | 0.00029674 | 0.056070273 |
| *ZNF730* | -2.444933 | 0.676102 | -3.616217 | 0.00029894 | 0.056070273 |
| *CYBC1* | -0.359055 | 0.099324 | -3.614983 | 0.00030037 | 0.056070273 |
| *PAOX* | 1.258947 | 0.34892 | 3.60812 | 0.00030842 | 0.056645434 |
| *ZNF684* | -1.603205 | 0.448503 | -3.574569 | 0.00035081 | 0.063406663 |
| *ETFB* | 0.647252 | 0.181402 | 3.568045 | 0.00035966 | 0.063990415 |
| *C7orf50* | -0.343987 | 0.096691 | -3.557585 | 0.00037428 | 0.06556805 |
| *GBP4* | 0.716485 | 0.202228 | 3.542955 | 0.00039567 | 0.066412 |
| *RIMS3* | -2.958722 | 0.835496 | -3.541276 | 0.0003982 | 0.066412 |
| *TRBC2* | -0.295395 | 0.083458 | -3.539423 | 0.000401 | 0.066412 |
| *ENSG00000260778* | -1.781111 | 0.503354 | -3.538488 | 0.00040243 | 0.066412 |
| *RAPGEF5* | -2.043015 | 0.582975 | -3.504463 | 0.00045753 | 0.074426844 |
| *IL2RA* | -1.418319 | 0.407653 | -3.479235 | 0.00050285 | 0.080646883 |
| *CD3G* | -4.41776 | 1.276568 | -3.460653 | 0.00053887 | NA |
| *KMT2E* | -0.281799 | 0.081484 | -3.458323 | 0.00054355 | 0.085963929 |
| *ACP5* | 0.785353 | 0.22794 | 3.445445 | 0.00057012 | 0.088931041 |
| *MIR3142HG* | -3.230923 | 0.940713 | -3.434548 | 0.00059354 | NA |
| *ATP6V0A1* | 1.119726 | 0.327725 | 3.416666 | 0.00063393 | 0.097547961 |
| *AP3S1* | 0.540275 | 0.158578 | 3.406998 | 0.00065682 | 0.09878158 |
| *PPP1R18* | 0.430009 | 0.126364 | 3.402938 | 0.00066665 | 0.09878158 |
| *DEF8* | -0.797763 | 0.234471 | -3.402399 | 0.00066797 | 0.09878158 |
| *RUBCN* | -0.492578 | 0.145013 | -3.396788 | 0.00068182 | 0.099536627 |
| *FOXO3* | 1.425621 | 0.420656 | 3.389044 | 0.00070137 | 0.101094586 |
| *PFKL* | -0.453355 | 0.133915 | -3.385407 | 0.00071073 | 0.101163287 |
| *MAPKAPK2* | 1.000857 | 0.297038 | 3.369452 | 0.00075318 | 0.105882048 |
| *POU2AF1* | 0.429354 | 0.128636 | 3.337752 | 0.00084459 | 0.117284695 |
| *ZNF638* | -0.43517 | 0.130751 | -3.328229 | 0.000874 | 0.11990662 |
| *GPER1* | -1.914 | 0.578057 | -3.311094 | 0.00092932 | 0.125978078 |
| *MPEG1* | 0.775383 | 0.234481 | 3.306814 | 0.00094364 | 0.12641398 |
| *ENSG00000257815* | -0.72811 | 0.220488 | -3.302265 | 0.00095908 | 0.126988273 |
| *DCAF12* | -2.5553 | 0.777234 | -3.287685 | 0.00101015 | 0.132213048 |
| *PARP2* | -0.977608 | 0.297773 | -3.283065 | 0.00102685 | 0.132872257 |
| *MDH2* | 0.469474 | 0.143337 | 3.27531 | 0.00105546 | 0.135039774 |
| *USP46.AS1* | -2.661087 | 0.812849 | -3.273778 | 0.0010612 | NA |
| *CERS4* | -0.54055 | 0.165506 | -3.266036 | 0.00109064 | 0.137990711 |

| | | | | | |
|---|---|---|---|---|---|
| *LCAT* | 1.894977 | 0.585315 | 3.237531 | 0.00120569 | 0.150870253 |
| *SNAP23* | 0.454782 | 0.140879 | 3.228169 | 0.00124585 | 0.153235778 |
| *GALNT2* | 0.625639 | 0.194185 | 3.221864 | 0.0012736 | 0.153235778 |
| *MPZL1* | -0.935655 | 0.290427 | -3.221651 | 0.00127454 | 0.153235778 |
| *SPRY1* | 0.590263 | 0.183267 | 3.22078 | 0.00127842 | 0.153235778 |
| *RNF207* | 1.794268 | 0.56012 | 3.203361 | 0.00135833 | 0.160685825 |
| *LSM7* | -0.394039 | 0.123093 | -3.201151 | 0.0013688 | 0.160685825 |
| *ABI3* | 0.737404 | 0.230826 | 3.194638 | 0.00140007 | 0.162574673 |
| *RPS19* | 0.269826 | 0.084535 | 3.191891 | 0.00141345 | 0.162574673 |
| *ENSG00000183308* | -2.16586 | 0.679947 | -3.185339 | 0.00144585 | 0.164638386 |
| *NKX6.3* | -1.891098 | 0.59524 | -3.177034 | 0.0014879 | 0.167090843 |
| *SLC38A6* | -1.359711 | 0.428213 | -3.175317 | 0.00149673 | 0.167090843 |
| *TNFRSF4* | -3.254171 | 1.026025 | -3.171629 | 0.00151587 | NA |
| *UBAP2L* | -0.628907 | 0.198788 | -3.163703 | 0.00155776 | 0.172215367 |
| *SNX18* | 1.386266 | 0.438825 | 3.159039 | 0.0015829 | 0.173312614 |
| *RABGAP1L* | 0.495643 | 0.157317 | 3.150612 | 0.00162929 | 0.176692366 |
| *BCLAF1* | -0.357172 | 0.113515 | -3.146482 | 0.00165248 | 0.177516512 |
| *RNGTT* | -0.845863 | 0.270113 | -3.131513 | 0.00173908 | 0.183292494 |
| *GPX1* | 0.289211 | 0.092413 | 3.129533 | 0.00175084 | 0.183292494 |
| *RINL* | 0.698354 | 0.223206 | 3.12874 | 0.00175557 | 0.183292494 |
| *GPT2* | 1.218762 | 0.389851 | 3.12623 | 0.00177063 | 0.183292494 |
| *TOX2* | -2.205178 | 0.705397 | -3.126151 | 0.00177111 | NA |
| *NOB1* | 0.643095 | 0.206111 | 3.120133 | 0.0018077 | 0.184807357 |
| *ENSG00000275580* | -2.236424 | 0.717147 | -3.118503 | 0.00181772 | 0.184807357 |
| *MARCHF1* | -0.41155 | 0.132125 | -3.114853 | 0.00184036 | 0.185453279 |
| *TMEM88* | -2.481371 | 0.79668 | -3.114641 | 0.00184169 | NA |
| *SLC25A5* | -0.241472 | 0.078265 | -3.085296 | 0.0020335 | 0.203117672 |
| *ZNF436* | -1.750639 | 0.570301 | -3.069673 | 0.00214293 | 0.212187543 |
| *CD226* | -2.984439 | 0.972486 | -3.068875 | 0.00214866 | NA |
| *ATXN10* | 0.649424 | 0.211973 | 3.063707 | 0.00218613 | 0.214524251 |
| *FOXP1* | -0.216662 | 0.070776 | -3.061242 | 0.00220421 | 0.214524251 |
| *RNF130* | 1.255243 | 0.411304 | 3.05186 | 0.00227429 | 0.219468462 |
| *FAM167A* | 1.831743 | 0.601001 | 3.047822 | 0.00230507 | 0.220569726 |
| *TMEM52* | -1.929527 | 0.634287 | -3.042038 | 0.00234982 | 0.22141415 |
| *PPP3CC* | -0.497331 | 0.163506 | -3.041659 | 0.00235278 | 0.22141415 |
| *TBKBP1* | -2.757435 | 0.912457 | -3.02199 | 0.00251119 | NA |
| *LINC02202* | -2.177106 | 0.720748 | -3.020619 | 0.00252258 | 0.235448087 |
| *SMIM20* | 0.799289 | 0.264915 | 3.01716 | 0.00255155 | 0.23621573 |
| *C15orf39* | 1.30374 | 0.432834 | 3.012102 | 0.00259445 | 0.237085769 |
| *CD79A* | 0.324997 | 0.107931 | 3.011151 | 0.00260259 | 0.237085769 |
| *FAM207A* | 1.174581 | 0.391941 | 2.996831 | 0.00272802 | 0.246539597 |
| *CERK* | 1.040938 | 0.347995 | 2.991248 | 0.0027784 | 0.249115052 |
| *ZNF540* | -0.993128 | 0.333501 | -2.977883 | 0.00290247 | 0.258206527 |
| *PXK* | 0.639143 | 0.215094 | 2.971464 | 0.00296384 | 0.261621821 |
| *ENSG00000272368* | -0.810855 | 0.273699 | -2.962581 | 0.00305072 | 0.267081353 |
| *ATP6V0D1* | -0.472731 | 0.159789 | -2.95847 | 0.0030917 | 0.267081353 |
| *SERPINB1* | 0.494473 | 0.167163 | 2.958037 | 0.00309605 | 0.267081353 |

| | | | | | |
|---|---|---|---|---|---|
| *MYO9B* | 0.728603 | 0.247297 | 2.946271 | 0.0032163 | 0.275368807 |
| *EVL* | 0.35031 | 0.119585 | 2.929381 | 0.00339638 | 0.288616166 |
| *DYNLL2* | -0.817013 | 0.279438 | -2.923774 | 0.00345816 | 0.291689435 |
| *CCNB1* | -1.35297 | 0.464198 | -2.91464 | 0.003561 | 0.294017078 |
| *SYNGAP1* | 1.700978 | 0.583599 | 2.914635 | 0.00356104 | 0.294017078 |
| *TCL1A* | -0.463623 | 0.159078 | -2.914445 | 0.00356322 | 0.294017078 |
| *AGAP3* | -2.844526 | 0.977131 | -2.911099 | 0.0036016 | NA |
| *ST3GAL1* | 0.919136 | 0.315825 | 2.910269 | 0.00361117 | 0.295830491 |
| *TBRG1* | -0.4538 | 0.156137 | -2.906431 | 0.00365578 | 0.297345068 |
| *DDX17* | -0.251451 | 0.086638 | -2.90232 | 0.0037041 | 0.299138998 |
| *QSOX2* | -0.728164 | 0.251368 | -2.896805 | 0.00376984 | 0.301780632 |
| *HSH2D* | 0.344468 | 0.119022 | 2.894152 | 0.00380184 | 0.301780632 |
| *CYTH3* | -1.071464 | 0.37037 | -2.892958 | 0.00381632 | 0.301780632 |
| *LBH* | -0.355331 | 0.123007 | -2.88871 | 0.00386825 | 0.303777898 |
| *HCK* | -0.990814 | 0.343814 | -2.88183 | 0.00395374 | 0.305685353 |
| *CD37* | -0.150251 | 0.052148 | -2.881243 | 0.00396111 | 0.305685353 |
| *ASTE1* | -0.713552 | 0.247736 | -2.880291 | 0.00397308 | 0.305685353 |
| *C3orf52* | -2.244779 | 0.783844 | -2.863809 | 0.0041858 | NA |
| *SUSD6* | -0.55667 | 0.194791 | -2.857787 | 0.00426607 | 0.326025125 |
| *ENSG00000271204* | 0.616583 | 0.218022 | 2.828083 | 0.00468276 | 0.355483992 |
| *RPL18A* | 0.238805 | 0.084637 | 2.821527 | 0.00477956 | 0.360429428 |
| *CD109* | -3.466531 | 1.22973 | -2.818937 | 0.00481829 | NA |
| *SYNGR2* | 0.417417 | 0.148279 | 2.815086 | 0.00487642 | 0.365314349 |
| *AGPAT2* | -0.462852 | 0.164783 | -2.808861 | 0.00497172 | 0.370019207 |
| *FCMR* | 0.34628 | 0.123474 | 2.804481 | 0.00503976 | 0.372647831 |
| *PRR7* | -1.174821 | 0.420613 | -2.793115 | 0.00522031 | 0.383507423 |
| *MPG* | 0.455178 | 0.163256 | 2.78812 | 0.00530149 | 0.38697453 |
| *NFATC3* | 0.487519 | 0.17501 | 2.785663 | 0.00534184 | 0.387436308 |
| *MRNIP* | 0.527187 | 0.190122 | 2.772884 | 0.0055562 | 0.40043301 |
| *ENSG00000275636* | -2.562055 | 0.926709 | -2.764682 | 0.00569783 | 0.40728113 |
| *BRPF1* | -0.83243 | 0.301249 | -2.763258 | 0.00572275 | 0.40728113 |
| *ARNT* | -0.71353 | 0.259353 | -2.751193 | 0.00593787 | 0.417969599 |
| *TADA2A* | -0.859186 | 0.312437 | -2.749948 | 0.00596048 | 0.417969599 |
| *RAB11B* | -0.326395 | 0.118745 | -2.748709 | 0.00598306 | 0.417969599 |
| *ENSG00000258168* | -2.974757 | 1.083558 | -2.745359 | 0.00604447 | NA |
| *MSI2* | -0.454049 | 0.165545 | -2.742758 | 0.00609255 | 0.423023871 |
| *ENSG00000249790* | -2.219031 | 0.809382 | -2.741637 | 0.00611338 | NA |
| *STXBP5* | -1.28726 | 0.46977 | -2.740192 | 0.00614034 | 0.423757558 |
| *ALG2* | -0.660249 | 0.24156 | -2.733266 | 0.00627096 | 0.42861821 |
| *JAML* | -2.666264 | 0.976304 | -2.730976 | 0.0063147 | 0.42861821 |
| *PBX3* | -0.90763 | 0.332403 | -2.730507 | 0.00632369 | 0.42861821 |
| *GRK5* | -0.647486 | 0.237309 | -2.728448 | 0.00636332 | 0.428751874 |
| *LGALS4* | -1.597368 | 0.585869 | -2.726493 | 0.00640113 | 0.428762627 |
| *SLC39A13* | -0.905211 | 0.332757 | -2.720335 | 0.00652157 | 0.434275725 |
| *COL8A1* | -2.078186 | 0.765496 | -2.714823 | 0.00663112 | NA |
| *SIDT2* | -0.66648 | 0.245501 | -2.714773 | 0.00663213 | 0.439070143 |
| *EIF3K* | -0.178443 | 0.065818 | -2.711171 | 0.0067046 | 0.440545333 |

| | | | | | |
|---|---|---|---|---|---|
| *NPW* | -2.355475 | 0.869486 | -2.709044 | 0.00674774 | NA |
| *ENSG00000198106* | -0.40619 | 0.149999 | -2.707946 | 0.0067701 | 0.440545333 |
| *DTX4* | 1.320224 | 0.48754 | 2.707928 | 0.00677048 | 0.440545333 |
| *LINC02362* | -2.428144 | 0.897015 | -2.706915 | 0.00679116 | NA |
| *UMPS* | 1.432644 | 0.530164 | 2.702266 | 0.00688685 | 0.445571622 |
| *RUNX3* | 0.685018 | 0.253816 | 2.698876 | 0.00695741 | 0.446471436 |
| *HIST1H2AK* | -1.880797 | 0.69715 | -2.697836 | 0.00697918 | 0.446471436 |
| *MCL1* | 0.395912 | 0.146855 | 2.695932 | 0.00701921 | 0.446523503 |
| *CR2* | 1.259428 | 0.467527 | 2.693809 | 0.00706406 | 0.446880002 |
| *NIPSNAP1* | 1.005815 | 0.373954 | 2.689675 | 0.00715217 | 0.449559663 |
| *BSPRY* | -2.145912 | 0.797949 | -2.689285 | 0.00716053 | NA |
| *ZWINT* | -1.939284 | 0.721425 | -2.688128 | 0.00718537 | 0.449559663 |
| *TLK1* | -0.368437 | 0.13733 | -2.682863 | 0.0072995 | 0.452800386 |
| *NEK1* | 1.277288 | 0.476355 | 2.681378 | 0.00733197 | 0.452800386 |
| *CNTRL* | -0.371317 | 0.138551 | -2.68001 | 0.007362 | 0.452800386 |
| *ZNF440* | 1.96557 | 0.733844 | 2.678457 | 0.00739623 | 0.452800386 |
| *GGT1* | -2.271364 | 0.8491 | -2.675026 | 0.00747234 | NA |
| *BYSL* | -1.140473 | 0.426432 | -2.674454 | 0.0074851 | 0.455790532 |
| *CHST12* | 0.563546 | 0.210874 | 2.672432 | 0.00753037 | 0.45610811 |
| *ITGB2.AS1* | -2.542449 | 0.951727 | -2.671407 | 0.0075534 | NA |
| *LINC02576* | -1.151109 | 0.431283 | -2.669033 | 0.00760699 | 0.456300385 |
| *TBC1D9* | 0.846599 | 0.317228 | 2.668738 | 0.00761369 | 0.456300385 |
| *SLC29A1* | -2.043063 | 0.766071 | -2.666937 | 0.0076546 | 0.456350475 |
| *SQSTM1* | 0.355099 | 0.13329 | 2.664104 | 0.00771938 | 0.456622353 |
| *PHF20L1* | -0.390421 | 0.146596 | -2.663234 | 0.00773936 | 0.456622353 |
| *TBL1X* | -0.724496 | 0.272609 | -2.657634 | 0.00786913 | 0.457414698 |
| *SMS* | 0.59855 | 0.225335 | 2.656265 | 0.00790115 | 0.457414698 |
| *ENSG00000255224* | -1.713427 | 0.645095 | -2.656084 | 0.00790539 | 0.457414698 |
| *CD72* | 0.463777 | 0.17474 | 2.654104 | 0.00795193 | 0.457414698 |
| *PSMB10* | 0.405134 | 0.152649 | 2.654031 | 0.00795364 | 0.457414698 |
| *CYB5RL* | -1.122885 | 0.424216 | -2.646965 | 0.00812179 | 0.464737579 |
| *SLC2A5* | 0.682062 | 0.257909 | 2.644587 | 0.00817906 | 0.46567457 |
| *PIK3R5* | 1.707172 | 0.645999 | 2.642683 | 0.00822519 | 0.46597124 |
| *JDP2* | 1.799591 | 0.6818 | 2.639469 | 0.00830359 | 0.46777391 |
| *IGHV3.15* | -1.526894 | 0.578803 | -2.63802 | 0.00833917 | 0.46777391 |
| *LINC02210.CRHR1* | -2.097084 | 0.795675 | -2.635603 | 0.0083988 | NA |
| *PPT1* | 0.715946 | 0.272065 | 2.631527 | 0.0085002 | 0.472787049 |
| *CDKN3* | -1.906865 | 0.724748 | -2.631073 | 0.00851158 | 0.472787049 |
| *PMEPA1* | -0.570898 | 0.217207 | -2.628362 | 0.00857971 | 0.474257852 |
| *SLC25A53* | -0.808965 | 0.308331 | -2.623692 | 0.00869824 | 0.478487389 |
| *KNL1* | 1.612216 | 0.61499 | 2.621534 | 0.00875351 | 0.479212516 |
| *MZB1* | -0.509555 | 0.194846 | -2.615173 | 0.00891822 | 0.485893672 |
| *RAB20* | 1.7434 | 0.668046 | 2.609701 | 0.00906213 | 0.490129536 |
| *E2F5* | -0.50735 | 0.194466 | -2.60894 | 0.00908232 | 0.490129536 |
| *PON2* | -1.799118 | 0.690023 | -2.607332 | 0.0091251 | 0.490129536 |
| *MRPL27* | 0.566745 | 0.217529 | 2.605374 | 0.00917741 | 0.490625151 |
| *VHL* | 0.389554 | 0.150106 | 2.595198 | 0.00945364 | 0.499689112 |

| | | | | | |
|---|---|---|---|---|---|
| *GBP2* | 0.567616 | 0.218974 | 2.592169 | 0.00953729 | 0.499689112 |
| *RARA* | 0.552616 | 0.213415 | 2.589398 | 0.00961438 | 0.499689112 |
| *RAMP2.AS1* | -1.744364 | 0.673713 | -2.58918 | 0.00962048 | 0.499689112 |
| *ABHD11* | -0.712902 | 0.275492 | -2.587744 | 0.00966068 | 0.499689112 |
| *ENSG00000263470* | -2.174558 | 0.840526 | -2.587138 | 0.00967769 | NA |
| *TMEM134* | -0.313823 | 0.121344 | -2.586227 | 0.0097033 | 0.499689112 |
| *C2orf42* | -0.782734 | 0.302866 | -2.584427 | 0.0097541 | 0.499689112 |
| *LEPR* | -1.927145 | 0.74612 | -2.582888 | 0.0097977 | 0.499689112 |
| *SCYL2* | -0.73937 | 0.286283 | -2.582653 | 0.00980439 | 0.499689112 |
| *RFC4* | 0.905059 | 0.350529 | 2.581983 | 0.00982344 | 0.499689112 |
| *DNAJB9* | -0.428769 | 0.16617 | -2.580301 | 0.00987143 | 0.499689112 |
| *MYCBP2* | 0.292748 | 0.113531 | 2.578564 | 0.0099212 | 0.499689112 |
| *PIGN* | -1.190063 | 0.461909 | -2.576398 | 0.00998357 | 0.499689112 |
| *ENSG00000271976* | -1.490505 | 0.579244 | -2.573191 | 0.01007657 | 0.499689112 |
| *ACAP2* | -0.384998 | 0.149626 | -2.573067 | 0.01008018 | 0.499689112 |
| *TAF11* | -0.470879 | 0.183045 | -2.572479 | 0.01009731 | 0.499689112 |
| *NMT2* | -0.440777 | 0.171398 | -2.571657 | 0.0101213 | 0.499689112 |
| *NOTCH2* | 0.623468 | 0.242657 | 2.569336 | 0.01018935 | 0.499689112 |
| *HERC6* | -0.877929 | 0.341813 | -2.568448 | 0.01021549 | 0.499689112 |
| *ENSG00000251364* | 1.729416 | 0.673412 | 2.568139 | 0.0102246 | 0.499689112 |
| *ENSG00000260735* | -1.350401 | 0.526626 | -2.56425 | 0.01033989 | 0.501817629 |
| *ANXA2R* | -0.555091 | 0.216618 | -2.562535 | 0.0103911 | 0.501817629 |
| *ZNF438* | -1.33029 | 0.519265 | -2.561873 | 0.01041095 | 0.501817629 |
| *DDX42* | -0.554152 | 0.216402 | -2.560757 | 0.01044444 | 0.501817629 |
| *PATL2* | 0.635849 | 0.248476 | 2.559 | 0.01049737 | 0.502241679 |
| *PLEKHO1* | 0.395129 | 0.154596 | 2.555879 | 0.01059199 | 0.504648264 |
| *SLC35B1* | -0.614564 | 0.240819 | -2.55197 | 0.01071156 | 0.508218837 |
| *AGAP5* | -1.371699 | 0.538335 | -2.54804 | 0.01083302 | 0.511848878 |
| *LOR* | -2.704562 | 1.062101 | -2.546426 | 0.01088322 | NA |
| *WIPF1* | -0.251667 | 0.098894 | -2.544799 | 0.01093407 | 0.513677351 |
| *PI4K2A* | -1.218819 | 0.479113 | -2.54391 | 0.01096194 | 0.513677351 |
| *KCNJ5* | -1.768801 | 0.696222 | -2.54057 | 0.01106721 | 0.516484849 |
| *BCL2L11* | 1.23776 | 0.487804 | 2.537413 | 0.01116752 | 0.517533917 |
| *CARD19* | -0.664605 | 0.262144 | -2.535268 | 0.01123612 | 0.517533917 |
| *CHIT1* | -1.464909 | 0.57783 | -2.535188 | 0.01123869 | 0.517533917 |
| *SNHG10* | -0.621024 | 0.24508 | -2.533959 | 0.01127819 | 0.517533917 |
| *BPGM* | -0.693117 | 0.273661 | -2.532757 | 0.01131694 | 0.517533917 |
| *SRRT* | 0.689045 | 0.272348 | 2.53002 | 0.01140559 | 0.518013266 |
| *NFXL1* | -0.935862 | 0.369961 | -2.529626 | 0.0114184 | 0.518013266 |
| *ENSG00000231105* | -2.215406 | 0.87612 | -2.528658 | 0.01144997 | NA |
| *ST6GALNAC6* | 0.59584 | 0.236148 | 2.523166 | 0.01163036 | 0.525535303 |
| *SIGLEC10* | 1.001494 | 0.397248 | 2.521079 | 0.01169955 | 0.525845192 |
| *SLC43A2* | 0.349748 | 0.138779 | 2.520177 | 0.01172958 | 0.525845192 |
| *GRINA* | 0.572236 | 0.2272 | 2.518641 | 0.01178088 | 0.526074022 |
| *NHEJ1* | -1.883261 | 0.748161 | -2.517186 | 0.01182963 | NA |
| *GZMM* | -3.05829 | 1.215316 | -2.516457 | 0.01185412 | NA |
| *CTBP2* | -2.631201 | 1.04616 | -2.515104 | 0.01189972 | NA |

| | | | | | |
|---|---|---|---|---|---|
| *CYTIP* | -0.180013 | 0.071598 | -2.514224 | 0.01192945 | 0.530627594 |
| *LRRC3* | -1.606058 | 0.639477 | -2.51152 | 0.01202124 | 0.532629995 |
| *NEURL4* | -1.278809 | 0.509902 | -2.507953 | 0.01214329 | 0.53595203 |
| *SLC8B1* | -0.685661 | 0.273596 | -2.506112 | 0.0122067 | 0.5366708 |
| *ANAPC16* | 0.218059 | 0.08713 | 2.502673 | 0.01232593 | 0.537621442 |
| *MTERF3* | -0.691389 | 0.276625 | -2.499373 | 0.01244133 | 0.537621442 |
| *ENSG00000282951* | -1.775605 | 0.710493 | -2.499117 | 0.01245031 | 0.537621442 |
| *CENPE* | -1.85636 | 0.742906 | -2.498783 | 0.01246205 | 0.537621442 |
| *IER3* | 1.783737 | 0.713861 | 2.498717 | 0.0124644 | 0.537621442 |
| *NXPH4* | -2.141789 | 0.857818 | -2.496787 | 0.01253243 | NA |
| *MBD6* | 1.269334 | 0.508608 | 2.495703 | 0.01257078 | 0.540135859 |
| *ENSG00000262049* | -0.662291 | 0.266119 | -2.4887 | 0.01282112 | 0.540135859 |
| *USP13* | -1.255952 | 0.504822 | -2.487912 | 0.01284955 | 0.540135859 |
| *C1QBP* | 0.286962 | 0.115351 | 2.487731 | 0.01285608 | 0.540135859 |
| *LSP1* | 0.327184 | 0.131523 | 2.487652 | 0.01285896 | 0.540135859 |
| *CXXC5* | -0.30879 | 0.124223 | -2.485771 | 0.01292713 | 0.540135859 |
| *TLR1* | -0.770746 | 0.310143 | -2.485131 | 0.01295036 | 0.540135859 |
| *DDX46* | -0.254497 | 0.102471 | -2.483601 | 0.01300613 | 0.540135859 |
| *PPM1B* | -0.448243 | 0.180498 | -2.483371 | 0.01301455 | 0.540135859 |
| *HSPA1A* | -1.885731 | 0.759922 | -2.481479 | 0.01308384 | 0.540135859 |
| *ENSG00000188206* | -0.502815 | 0.202753 | -2.479935 | 0.01314065 | 0.540135859 |
| *LINC01800* | -1.888924 | 0.761697 | -2.479888 | 0.01314236 | 0.540135859 |
| *PBXIP1* | -0.401042 | 0.161799 | -2.478649 | 0.0131881 | 0.540135859 |
| *HDAC9* | 0.882529 | 0.356063 | 2.478574 | 0.01319086 | 0.540135859 |
| *FEZ2* | -0.44044 | 0.17792 | -2.475488 | 0.01330544 | 0.540135859 |
| *EPHX2* | -1.925411 | 0.777863 | -2.475258 | 0.01331399 | 0.540135859 |
| *ZNF780A* | 0.931144 | 0.376242 | 2.474854 | 0.01332908 | 0.540135859 |
| *GIMAP4* | -2.639886 | 1.066905 | -2.47434 | 0.01334825 | NA |
| *PRKCB* | -0.285816 | 0.115605 | -2.472346 | 0.01342295 | 0.540637122 |
| *ZNF791* | 0.342666 | 0.138619 | 2.471988 | 0.0134364 | 0.540637122 |
| *NT5DC4* | -3.109179 | 1.258143 | -2.471244 | 0.0134644 | NA |
| *ENSG00000233184* | -0.655517 | 0.265483 | -2.469152 | 0.01354336 | 0.54302211 |
| *SPRYD7* | 0.97612 | 0.395709 | 2.466765 | 0.01363397 | 0.544736783 |
| *SPAG7* | -0.290466 | 0.117918 | -2.463285 | 0.01376706 | 0.548131245 |
| *ENSG00000267317* | -1.383111 | 0.562315 | -2.459671 | 0.01390645 | 0.549544937 |
| *PRRT3.AS1* | -1.716327 | 0.69817 | -2.458323 | 0.01395875 | 0.549544937 |
| *ZER1* | 1.662651 | 0.676366 | 2.458211 | 0.0139631 | 0.549544937 |
| *CRIPT* | 0.537341 | 0.218665 | 2.457376 | 0.01399561 | 0.549544937 |
| *MIXL1* | -1.824188 | 0.742341 | -2.457344 | 0.01399685 | NA |
| *NDUFB3* | 0.332748 | 0.135837 | 2.449619 | 0.01430075 | 0.556958093 |
| *KCNN1* | 1.92106 | 0.784404 | 2.449069 | 0.01432258 | 0.556958093 |
| *SLX4IP* | -0.706315 | 0.288455 | -2.448611 | 0.01434084 | 0.556958093 |
| *HEATR6* | 1.503821 | 0.614542 | 2.447061 | 0.01440264 | 0.556958093 |
| *SSBP3* | -0.897658 | 0.36693 | -2.446403 | 0.01442897 | 0.556958093 |
| *MAST1* | -1.96018 | 0.801618 | -2.445278 | 0.01447403 | NA |
| *OAZ1* | -0.147406 | 0.060327 | -2.443454 | 0.01454744 | 0.558459605 |
| *MRPL15* | 0.664536 | 0.272237 | 2.441017 | 0.01464595 | 0.558459605 |

| | | | | | |
|---|---|---|---|---|---|
| *ENSG00000258572* | -1.409511 | 0.577487 | -2.440768 | 0.01465606 | 0.558459605 |
| *PPP1R15A* | 0.656213 | 0.268996 | 2.439494 | 0.01470786 | 0.558459605 |
| *GYPC* | 0.241361 | 0.098944 | 2.439365 | 0.01471308 | 0.558459605 |
| *PHF21A* | -0.443835 | 0.182088 | -2.437484 | 0.01478987 | 0.559509206 |
| *ENSG00000257275* | -0.470473 | 0.193314 | -2.433717 | 0.01494467 | 0.562066958 |
| *BCL11A* | -0.278942 | 0.114629 | -2.433438 | 0.0149562 | 0.562066958 |
| *YES1* | -1.868039 | 0.769018 | -2.429123 | 0.0151354 | NA |
| *LINC02391* | -1.714081 | 0.705684 | -2.428963 | 0.01514209 | 0.567180978 |
| *VSIG1* | -2.510084 | 1.03408 | -2.42736 | 0.01520915 | NA |
| *PFDN1* | -0.326213 | 0.13441 | -2.426996 | 0.01522442 | 0.567556563 |
| *D2HGDH* | -0.844451 | 0.348034 | -2.426344 | 0.01525181 | 0.567556563 |
| *ENSG00000268439* | -1.227701 | 0.506693 | -2.42297 | 0.01539421 | 0.57098962 |
| *CACNB1* | -1.934391 | 0.799557 | -2.419329 | 0.01554918 | NA |
| *SYK* | -0.302703 | 0.125331 | -2.415217 | 0.01572583 | 0.579599439 |
| *NANS* | 0.755806 | 0.312942 | 2.415164 | 0.01572813 | 0.579599439 |
| *ENSG00000259802* | -1.923628 | 0.797477 | -2.412142 | 0.01585909 | 0.582540168 |
| *SRSF7* | -0.25287 | 0.104958 | -2.409257 | 0.01598503 | 0.585278215 |
| *PKN3* | -2.042917 | 0.848689 | -2.407144 | 0.01607783 | NA |
| *CD180* | 0.595168 | 0.247401 | 2.405687 | 0.0161421 | 0.587786498 |
| *SEC23B* | 1.289462 | 0.53608 | 2.405355 | 0.01615677 | 0.587786498 |
| *FUZ* | 1.042957 | 0.434016 | 2.403037 | 0.01625954 | 0.588415441 |
| *ARAP2* | -0.506704 | 0.210895 | -2.402635 | 0.01627741 | 0.588415441 |
| *DMXL2* | -2.255961 | 0.939213 | -2.401969 | 0.01630711 | NA |
| *ATMIN* | -0.721158 | 0.300369 | -2.400907 | 0.01635448 | 0.58933053 |
| *ASNS* | -0.769923 | 0.320877 | -2.399435 | 0.0164204 | 0.589839396 |
| *L3MBTL3* | -0.703059 | 0.293314 | -2.396953 | 0.01653205 | 0.591861474 |
| *C21orf62* | -0.880485 | 0.3675 | -2.395877 | 0.01658065 | 0.591861474 |
| *ENSG00000226310* | -1.794859 | 0.749512 | -2.394703 | 0.01663385 | NA |
| *SREK1* | -0.377354 | 0.157848 | -2.390615 | 0.01682018 | 0.595539551 |
| *TAGLN2* | 0.222726 | 0.093249 | 2.38851 | 0.01691683 | 0.595539551 |
| *RANBP2* | -0.530615 | 0.222172 | -2.388309 | 0.0169261 | 0.595539551 |
| *SHARPIN* | -0.376825 | 0.157861 | -2.387068 | 0.01698333 | 0.595539551 |
| *ZNF165* | 1.449335 | 0.607317 | 2.386454 | 0.01701171 | 0.595539551 |
| *BANK1* | -0.299113 | 0.125366 | -2.385909 | 0.01703696 | 0.595539551 |
| *ARMH1* | -1.015319 | 0.425597 | -2.385632 | 0.01704978 | 0.595539551 |
| *ENSG00000260267* | -1.123065 | 0.471287 | -2.382973 | 0.01717346 | 0.597037336 |
| *PRPF38B* | -0.24768 | 0.10396 | -2.382457 | 0.01719753 | 0.597037336 |
| *GPBP1* | -0.228193 | 0.095856 | -2.38057 | 0.01728587 | 0.597117947 |
| *ENSG00000269399* | 1.635743 | 0.687374 | 2.379698 | 0.01732683 | 0.597117947 |
| *RAB39B* | -0.778586 | 0.327267 | -2.379053 | 0.01735717 | 0.597117947 |
| *ENSG00000267811* | -1.800522 | 0.757536 | -2.376812 | 0.01746298 | 0.598948651 |
| *TUSC2* | -0.379092 | 0.159664 | -2.374302 | 0.01758217 | 0.601225685 |
| *SLC16A11* | 1.171932 | 0.493833 | 2.373136 | 0.01763779 | 0.601321783 |
| *VNN2* | 0.705394 | 0.29787 | 2.368126 | 0.01787846 | 0.607707439 |
| *SNTB1* | -2.357717 | 0.995921 | -2.367373 | 0.01791488 | NA |
| *BCDIN3D* | -0.603855 | 0.255254 | -2.365705 | 0.01799576 | 0.608692929 |
| *AHSA2P* | -0.566307 | 0.23942 | -2.365323 | 0.01801436 | 0.608692929 |

| | | | | | |
|---|---|---|---|---|---|
| *YIPF2* | 1.167042 | 0.494045 | 2.362216 | 0.01816607 | 0.610045912 |
| *HES1* | -1.084546 | 0.459317 | -2.361215 | 0.01821518 | 0.610045912 |
| *ENSG00000261386* | -0.545494 | 0.231121 | -2.360205 | 0.01826485 | 0.610045912 |
| *ESCO1* | -0.422674 | 0.17909 | -2.360127 | 0.0182687 | 0.610045912 |
| *GTPBP2* | -0.9543 | 0.404799 | -2.357465 | 0.01840019 | 0.612640203 |
| *SNX9* | 0.60707 | 0.257914 | 2.353773 | 0.01858398 | 0.616705298 |
| *ENSG00000229729* | -0.982885 | 0.417901 | -2.351958 | 0.01867487 | 0.616705298 |
| *ZNF837* | -0.79659 | 0.338959 | -2.350107 | 0.01876802 | 0.616705298 |
| *HAVCR2* | -1.904016 | 0.810777 | -2.348383 | 0.01885513 | NA |
| *VASH1* | -1.959421 | 0.834457 | -2.34814 | 0.01886745 | NA |
| *TMEM115* | 0.797799 | 0.339791 | 2.347909 | 0.01887911 | 0.616705298 |
| *TYW1B* | -1.141883 | 0.486483 | -2.347221 | 0.01891404 | 0.616705298 |
| *ENSG00000273466* | -1.763636 | 0.751815 | -2.345839 | 0.01898429 | 0.616705298 |
| *PGD* | -0.5973 | 0.254629 | -2.345771 | 0.01898776 | 0.616705298 |
| *SNX19* | -0.769842 | 0.328499 | -2.343513 | 0.0191031 | 0.616705298 |
| *MRPS18C* | -0.24835 | 0.106007 | -2.342782 | 0.01914056 | 0.616705298 |
| *APLF* | -1.227165 | 0.523858 | -2.342551 | 0.01915243 | 0.616705298 |
| *NSMCE1* | 0.435625 | 0.186014 | 2.341898 | 0.01918598 | 0.616705298 |
| *JMJD1C.AS1* | -2.180158 | 0.931096 | -2.341496 | 0.01920662 | NA |
| *EMC8* | -0.579988 | 0.247704 | -2.341456 | 0.0192087 | 0.616705298 |
| *SNAI3.AS1* | -1.18126 | 0.504738 | -2.34034 | 0.0192662 | 0.616705298 |
| *TOP1MT* | -0.599403 | 0.256187 | -2.339712 | 0.01929864 | 0.616705298 |
| *CD47* | -0.269569 | 0.115249 | -2.339015 | 0.01933466 | 0.616705298 |
| *DSCR9* | -1.823468 | 0.780233 | -2.337081 | 0.01943496 | NA |
| *ENSG00000259943* | -0.762833 | 0.326944 | -2.333224 | 0.01963641 | 0.621176303 |
| *CHTF8* | -0.368415 | 0.157906 | -2.333125 | 0.01964161 | 0.621176303 |
| *CD5* | -1.09205 | 0.468225 | -2.332318 | 0.01968395 | 0.621176303 |
| *FNBP1* | 0.339 | 0.145425 | 2.331109 | 0.01974763 | 0.621176303 |
| *USF3* | -0.389961 | 0.167305 | -2.33084 | 0.01976182 | 0.621176303 |
| *WHAMM* | 0.90121 | 0.386773 | 2.330076 | 0.01980214 | 0.621176303 |
| *PRXL2A* | 0.769986 | 0.330648 | 2.32872 | 0.01987391 | 0.621714783 |
| *PQLC3* | -0.569706 | 0.244788 | -2.327342 | 0.01994708 | 0.62229416 |
| *TPCN1* | 0.635103 | 0.27331 | 2.323743 | 0.02013926 | 0.624928308 |
| *TSR2* | 0.428168 | 0.184261 | 2.323706 | 0.02014127 | 0.624928308 |
| *TLE4* | -0.506999 | 0.218359 | -2.321862 | 0.02024036 | 0.626296139 |
| *RNF157* | -1.93675 | 0.83446 | -2.320961 | 0.02028895 | NA |
| *DCXR.DT* | -1.560416 | 0.672405 | -2.32065 | 0.02030577 | 0.626617294 |
| *SOX5* | -2.998396 | 1.292286 | -2.320227 | 0.02032862 | NA |
| *ENSG00000268292* | -1.684873 | 0.726772 | -2.318298 | 0.02043315 | NA |
| *MTHFD2L* | -0.562359 | 0.24271 | -2.317005 | 0.02050347 | 0.631008009 |
| *NAA10* | -0.279574 | 0.120804 | -2.314279 | 0.02065243 | 0.632744357 |
| *ARSA* | -0.79951 | 0.345519 | -2.31394 | 0.02067102 | 0.632744357 |
| *LBX2.AS1* | -2.475568 | 1.070149 | -2.313292 | 0.02070657 | NA |
| *SNX21* | -1.937744 | 0.83801 | -2.312315 | 0.0207603 | NA |
| *ENSG00000244198* | -1.486913 | 0.643206 | -2.311722 | 0.02079299 | 0.63477152 |
| *GNG4* | -3.401706 | 1.47199 | -2.310957 | 0.02083524 | NA |
| *DUSP2* | -1.827935 | 0.791022 | -2.310851 | 0.02084106 | NA |

| | | | | | |
|---|---|---|---|---|---|
| *ENSG00000277369* | -1.865815 | 0.808178 | -2.308669 | 0.02096195 | NA |
| *ENSG00000237481* | -1.732902 | 0.750652 | -2.308528 | 0.02096981 | NA |
| *TBC1D16* | -1.351839 | 0.585711 | -2.30803 | 0.02099745 | 0.639299353 |
| *PPP1R14B* | -0.670838 | 0.291082 | -2.304639 | 0.02118681 | 0.643344403 |
| *TMEM206* | -0.732753 | 0.318222 | -2.30265 | 0.02129856 | 0.645017935 |
| *ENSG00000278668* | -1.547737 | 0.672923 | -2.300022 | 0.02144697 | 0.646541705 |
| *PRKAR2A.AS1* | -1.653119 | 0.718785 | -2.299881 | 0.02145499 | NA |
| *PDHA1* | 0.677676 | 0.294676 | 2.299731 | 0.02146347 | 0.646541705 |
| *TNIP2* | 0.58193 | 0.253151 | 2.298749 | 0.02151922 | 0.646541705 |
| *RAB5A* | -0.44363 | 0.193379 | -2.294099 | 0.02178483 | 0.650060218 |
| *LYG1* | -1.62115 | 0.706721 | -2.293904 | 0.02179601 | 0.650060218 |
| *PTMS* | -1.651829 | 0.720303 | -2.293241 | 0.02183415 | 0.650060218 |
| *RBM8A* | -0.214662 | 0.093631 | -2.292646 | 0.0218684 | 0.650060218 |
| *CHRAC1* | -0.422853 | 0.184513 | -2.291721 | 0.02192176 | 0.650060218 |
| *PACS1* | 0.468456 | 0.204753 | 2.287903 | 0.02214318 | 0.654121726 |
| *UTRN* | -0.444459 | 0.194309 | -2.287381 | 0.02217362 | 0.654121726 |
| *NAA40* | 1.110652 | 0.486254 | 2.2841 | 0.02236566 | 0.658082089 |
| *EXD3* | -0.707254 | 0.309828 | -2.282731 | 0.02244622 | 0.658750184 |
| *ZNF563* | 0.84654 | 0.371169 | 2.280743 | 0.02256365 | 0.659488106 |
| *NAPA* | -0.445472 | 0.195421 | -2.279548 | 0.02263453 | 0.659488106 |
| *ACTG1* | 0.427215 | 0.187427 | 2.27937 | 0.02264511 | 0.659488106 |
| *P3H4* | -1.600333 | 0.702678 | -2.277477 | 0.02275777 | 0.659806978 |
| *ENSG00000254427* | -1.477257 | 0.648773 | -2.277001 | 0.02278616 | 0.659806978 |
| *ZNF568* | 0.84888 | 0.372926 | 2.276269 | 0.02282989 | 0.659806978 |
| *PTCD2* | -0.784207 | 0.344726 | -2.274869 | 0.02291377 | 0.659899239 |
| *DENND1A* | -0.967658 | 0.425717 | -2.273007 | 0.02302574 | 0.659899239 |
| *HSPA1B* | -2.214868 | 0.974792 | -2.272144 | 0.02307781 | 0.659899239 |
| *ABCB1* | 0.695688 | 0.306405 | 2.270486 | 0.02317813 | 0.659899239 |
| *ZCCHC18* | -0.555648 | 0.244757 | -2.270205 | 0.02319515 | 0.659899239 |
| *PLEKHN1* | -1.638242 | 0.722099 | -2.268724 | 0.02328514 | 0.659899239 |
| *FLT1* | -1.471556 | 0.650012 | -2.263892 | 0.02358079 | 0.659899239 |
| *LMNB2* | -1.237817 | 0.546824 | -2.263651 | 0.0235956 | 0.659899239 |
| *MTA1* | -0.68586 | 0.303162 | -2.262353 | 0.02367562 | 0.659899239 |
| *NDUFS7* | 0.29793 | 0.131757 | 2.261205 | 0.02374659 | 0.659899239 |
| *ERAP1* | 0.630575 | 0.278879 | 2.261106 | 0.02375267 | 0.659899239 |
| *FXYD5* | -0.216566 | 0.095783 | -2.261009 | 0.02375873 | 0.659899239 |
| *IL27RA* | 0.565003 | 0.25007 | 2.259379 | 0.02385985 | 0.659899239 |
| *NFYC* | -0.411114 | 0.18209 | -2.257753 | 0.02396106 | 0.659899239 |
| *RBM45* | -0.786269 | 0.34827 | -2.257644 | 0.02396786 | 0.659899239 |
| *AP3B1* | 0.490303 | 0.217593 | 2.253298 | 0.02424037 | 0.659899239 |
| *TXLNB* | -1.37567 | 0.610549 | -2.253171 | 0.02424838 | 0.659899239 |
| *RPE* | -0.690506 | 0.306461 | -2.25316 | 0.02424904 | 0.659899239 |
| *LINC01410* | -1.688032 | 0.749791 | -2.251338 | 0.02436414 | 0.659899239 |
| *HSBP1* | -0.283658 | 0.126013 | -2.251023 | 0.02438405 | 0.659899239 |
| *MOB4* | 0.427148 | 0.189798 | 2.250543 | 0.02441449 | 0.659899239 |
| *IFT27* | 0.684514 | 0.30445 | 2.248363 | 0.02455307 | 0.659899239 |
| *MIR4453HG* | -0.833863 | 0.370883 | -2.248319 | 0.02455584 | 0.659899239 |

| | | | | | |
|---|---|---|---|---|---|
| *CADM4* | -1.95502 | 0.869663 | -2.248019 | 0.02457495 | NA |
| *RPTOR* | -1.030826 | 0.458903 | -2.246284 | 0.02468584 | 0.659899239 |
| *PHF13* | -0.938354 | 0.417875 | -2.245538 | 0.02473359 | 0.659899239 |
| *KLF12* | 0.957005 | 0.426221 | 2.245329 | 0.02474705 | 0.659899239 |
| *RPL37A* | 0.118753 | 0.052907 | 2.244569 | 0.0247958 | 0.659899239 |
| *APOLD1* | 0.565597 | 0.252124 | 2.243326 | 0.0248758 | 0.659899239 |
| *ST13* | -0.190455 | 0.084903 | -2.243217 | 0.0248828 | 0.659899239 |
| *THG1L* | 0.726327 | 0.323832 | 2.242917 | 0.02490217 | 0.659899239 |
| *C1GALT1* | -0.438612 | 0.195568 | -2.242763 | 0.02491213 | 0.659899239 |
| *SMG7* | -0.549163 | 0.245144 | -2.240168 | 0.02508005 | 0.659899239 |
| *ENSG00000255320* | -2.056687 | 0.918171 | -2.239981 | 0.02509213 | 0.659899239 |
| *LY96* | 0.665945 | 0.297349 | 2.239607 | 0.02511645 | 0.659899239 |
| *HIPK1* | -0.719453 | 0.321251 | -2.239533 | 0.02512127 | 0.659899239 |
| *CFLAR* | 0.426647 | 0.190517 | 2.23942 | 0.02512862 | 0.659899239 |
| *BOD1L1* | 0.35416 | 0.158182 | 2.238938 | 0.02515992 | 0.659899239 |
| *ZC3H10* | 0.963522 | 0.430448 | 2.238414 | 0.02519404 | 0.659899239 |
| *TIGAR* | -0.801247 | 0.357997 | -2.238137 | 0.0252121 | 0.659899239 |
| *LINC00513* | 0.479778 | 0.214408 | 2.237686 | 0.02524154 | 0.659899239 |
| *ENSG00000272563* | -1.878373 | 0.839427 | -2.237683 | 0.02524172 | NA |
| *FOSB* | -1.723031 | 0.770217 | -2.237072 | 0.02528161 | 0.659899239 |
| *PCDH9* | -0.417456 | 0.18661 | -2.237045 | 0.02528337 | 0.659899239 |
| *TACC1* | -0.472141 | 0.211116 | -2.236409 | 0.02532502 | 0.659899239 |
| *ECHS1* | 0.419097 | 0.187978 | 2.2295 | 0.02578062 | 0.665630252 |
| *SDHC* | 0.312849 | 0.140335 | 2.229303 | 0.02579373 | 0.665630252 |
| *SPATS2* | -0.46722 | 0.209593 | -2.229175 | 0.02580227 | 0.665630252 |
| *TOR4A* | -0.91581 | 0.410866 | -2.228972 | 0.02581574 | 0.665630252 |
| *MIR222HG* | -1.221936 | 0.548488 | -2.227826 | 0.02589215 | 0.665630252 |
| *DOCK6* | -1.738165 | 0.780525 | -2.226917 | 0.02595281 | NA |
| *DBF4* | -0.525369 | 0.236141 | -2.22481 | 0.02609398 | 0.665630252 |
| *IRF2BP2* | 0.6399 | 0.287704 | 2.22416 | 0.0261377 | 0.665630252 |
| *GALC* | -1.036991 | 0.466504 | -2.222897 | 0.02622275 | 0.665630252 |
| *ENSG00000267042* | -1.717358 | 0.77282 | -2.222197 | 0.02626998 | 0.665630252 |
| *KIAA0355* | -0.365163 | 0.164341 | -2.221981 | 0.02628461 | 0.665630252 |
| *PTGR2* | 1.472803 | 0.662854 | 2.221912 | 0.02628928 | 0.665630252 |
| *PKN1* | 0.537444 | 0.241889 | 2.221861 | 0.0262927 | 0.665630252 |
| *BLVRB* | 0.792377 | 0.356657 | 2.221681 | 0.02630488 | 0.665630252 |
| *CFL1* | -0.160823 | 0.072468 | -2.219218 | 0.02647192 | 0.66723237 |
| *DPH1* | -1.565162 | 0.705366 | -2.218936 | 0.02649105 | NA |
| *CENPN* | -1.018731 | 0.459461 | -2.217232 | 0.02660727 | 0.66723237 |
| *GP1BA* | -1.364741 | 0.615647 | -2.216759 | 0.02663954 | 0.66723237 |
| *ENSG00000279278* | 0.739701 | 0.333953 | 2.214987 | 0.02676094 | 0.66723237 |
| *MRM1* | 1.632328 | 0.737005 | 2.214813 | 0.02677291 | 0.66723237 |
| *E2F4* | -0.372898 | 0.168502 | -2.213013 | 0.02689675 | 0.66723237 |
| *TMEM217* | -2.491155 | 1.125856 | -2.212678 | 0.02691988 | NA |
| *DDA1* | -0.43813 | 0.198019 | -2.212561 | 0.02692794 | 0.66723237 |
| *COG3* | -0.650971 | 0.294223 | -2.21251 | 0.02693145 | 0.66723237 |
| *ICAM2* | 0.465149 | 0.210264 | 2.212219 | 0.02695154 | 0.66723237 |

| | | | | | |
|---|---|---|---|---|---|
| *FAM89A* | -1.424404 | 0.644089 | -2.211504 | 0.02700099 | 0.66723237 |
| *RESF1* | -0.275407 | 0.124543 | -2.211334 | 0.02701275 | 0.66723237 |
| *ENSG00000261684* | -1.885435 | 0.853384 | -2.209364 | 0.02714931 | NA |
| *LRP5* | -1.803913 | 0.817058 | -2.207815 | 0.02725719 | NA |
| *ZNF2* | 1.564225 | 0.708727 | 2.20709 | 0.02730775 | 0.671174561 |
| *DGUOK* | -0.237465 | 0.107627 | -2.206375 | 0.02735772 | 0.671174561 |
| *PLP2* | -0.382655 | 0.17346 | -2.206012 | 0.02738314 | 0.671174561 |
| *TAF4B* | 1.207808 | 0.548179 | 2.203309 | 0.02757297 | 0.671174561 |
| *PAXBP1* | -0.707881 | 0.321359 | -2.202774 | 0.02761069 | 0.671174561 |
| *NKTR* | -0.301174 | 0.136726 | -2.202763 | 0.02761149 | 0.671174561 |
| *POMT2* | -1.285093 | 0.583491 | -2.20242 | 0.02763566 | 0.671174561 |
| *PARP16* | 1.173203 | 0.532912 | 2.201496 | 0.02770092 | 0.671174561 |
| *CCR7* | 0.247423 | 0.11243 | 2.20069 | 0.02775798 | 0.671174561 |
| *ZNF721* | -0.397161 | 0.180582 | -2.199336 | 0.02785407 | 0.671174561 |
| *UBL7* | 0.371226 | 0.168794 | 2.199288 | 0.02785747 | 0.671174561 |
| *ENSG00000250155* | -1.903804 | 0.865795 | -2.19891 | 0.02788432 | NA |
| *CAMSAP2* | -2.572131 | 1.169967 | -2.198465 | 0.02791598 | NA |
| *CAPN7* | 0.77386 | 0.352143 | 2.197572 | 0.02797959 | 0.671174561 |
| *POU2F2* | -0.19921 | 0.090718 | -2.195913 | 0.02809814 | 0.671174561 |
| *DDX60L* | -0.617902 | 0.281422 | -2.195642 | 0.02811756 | 0.671174561 |
| *AHCYL2* | -1.360983 | 0.619928 | -2.19539 | 0.02813567 | 0.671174561 |
| *SMARCE1* | -0.261218 | 0.118985 | -2.195378 | 0.02813647 | 0.671174561 |
| *TFEC* | -2.31054 | 1.052605 | -2.195069 | 0.02815868 | NA |
| *GDPD5* | -1.898826 | 0.865128 | -2.19485 | 0.02817436 | 0.671174561 |
| *HP1BP3* | -0.221554 | 0.101048 | -2.192561 | 0.02833904 | 0.672564676 |
| *LYSMD3* | -0.665121 | 0.303381 | -2.19236 | 0.0283535 | 0.672564676 |
| *ENSG00000224934* | -1.704545 | 0.777612 | -2.192025 | 0.02837774 | NA |
| *TAB1* | 1.111974 | 0.507385 | 2.191579 | 0.02840991 | 0.672564676 |
| *TMEM251* | 0.73956 | 0.337781 | 2.189466 | 0.02856302 | 0.674786474 |
| *ENSG00000248734* | -1.999353 | 0.913338 | -2.189061 | 0.02859239 | NA |
| *TMEM79* | 1.054428 | 0.481943 | 2.187869 | 0.02867918 | 0.676127881 |
| *GSEC* | -1.800759 | 0.823425 | -2.186913 | 0.02874891 | NA |
| *CNP* | 0.448747 | 0.205247 | 2.186373 | 0.02878834 | 0.676292736 |
| *NDUFS5* | 0.230537 | 0.105511 | 2.184968 | 0.02889121 | 0.676292736 |
| *PIP4K2A* | -0.382062 | 0.174863 | -2.184922 | 0.02889457 | 0.676292736 |
| *FCRL1* | 0.522213 | 0.239051 | 2.184524 | 0.02892373 | 0.676292736 |
| *LARP7* | -0.333103 | 0.152551 | -2.183556 | 0.0289949 | 0.676567527 |
| *ENSG00000251136* | 1.198885 | 0.54956 | 2.181537 | 0.02914375 | 0.677291172 |
| *AGRN* | -1.331841 | 0.61051 | -2.181521 | 0.02914487 | 0.677291172 |
| *AMDHD1* | -2.050631 | 0.941029 | -2.179136 | 0.02932154 | NA |
| *PCM1* | -0.282266 | 0.129618 | -2.177678 | 0.02943 | 0.681395336 |
| *RPS4Y2* | -1.367754 | 0.628122 | -2.177528 | 0.02944116 | 0.681395336 |
| *TPP1* | 0.407964 | 0.187423 | 2.176705 | 0.02950258 | 0.681431691 |
| *SF3B1* | -0.256732 | 0.117996 | -2.175762 | 0.02957308 | 0.681677406 |
| *FCGR2A* | -2.585582 | 1.188718 | -2.175101 | 0.02962255 | NA |
| *RGS12* | -1.181069 | 0.543229 | -2.174163 | 0.0296929 | 0.683056734 |
| *GNG11* | -1.078414 | 0.496295 | -2.172929 | 0.02978564 | 0.68380865 |

| | | | | | |
|---|---|---|---|---|---|
| *ENSG00000269051* | -1.571159 | 0.723785 | -2.170753 | 0.02994984 | 0.684942119 |
| *EGR1* | -1.739952 | 0.80157 | -2.170681 | 0.02995532 | 0.684942119 |
| *TUBB4A* | -1.962094 | 0.904039 | -2.170364 | 0.02997926 | NA |
| *NEU3* | -1.146811 | 0.528513 | -2.169884 | 0.03001567 | 0.684946708 |
| *SYT1* | -2.398898 | 1.106947 | -2.16713 | 0.03022495 | NA |
| *MAT2A* | -0.320607 | 0.147971 | -2.166687 | 0.03025873 | 0.689112236 |
| *RRS1* | 0.753775 | 0.348235 | 2.164558 | 0.03042159 | 0.689304536 |
| *TNNC2* | -1.364012 | 0.630284 | -2.164124 | 0.03045486 | 0.689304536 |
| *TRIM2* | -1.596485 | 0.737907 | -2.163531 | 0.03050033 | 0.689304536 |
| *ENSG00000278238* | -1.193392 | 0.551741 | -2.162958 | 0.0305444 | 0.689304536 |
| *NUP37* | 0.716347 | 0.331413 | 2.161491 | 0.0306574 | 0.689304536 |
| *TAB3* | 1.369841 | 0.633961 | 2.160767 | 0.03071337 | 0.689304536 |
| *ENSG00000272540* | -1.64009 | 0.759102 | -2.160565 | 0.03072894 | NA |
| *ENSG00000272004* | -1.503122 | 0.696204 | -2.159026 | 0.03084816 | 0.689304536 |
| *CDK5R1* | -2.309481 | 1.069795 | -2.158806 | 0.0308652 | NA |
| *DUSP16* | -2.52136 | 1.167974 | -2.158747 | 0.03086978 | NA |
| *PILRA* | -1.716363 | 0.795219 | -2.158352 | 0.03090045 | NA |
| *SMCHD1* | -0.235404 | 0.109114 | -2.157423 | 0.03097268 | 0.689304536 |
| *YWHAB* | -0.199066 | 0.092271 | -2.157393 | 0.03097505 | 0.689304536 |
| *UBP1* | -0.755285 | 0.350137 | -2.157115 | 0.03099672 | 0.689304536 |
| *DIS3L* | 0.743888 | 0.344863 | 2.157056 | 0.03100133 | 0.689304536 |
| *FAM41C* | -0.826608 | 0.383495 | -2.155461 | 0.0311258 | 0.689304536 |
| *ENSG00000234290* | -1.829402 | 0.848746 | -2.155418 | 0.03112912 | NA |
| *SRPK1* | 0.604246 | 0.280394 | 2.154993 | 0.03116237 | 0.689304536 |
| *ANKDD1A* | -1.54105 | 0.715233 | -2.154614 | 0.03119206 | 0.689304536 |
| *LINC01134* | -1.427557 | 0.662638 | -2.154354 | 0.03121246 | 0.689304536 |
| *L3MBTL1* | -0.99065 | 0.4599 | -2.154057 | 0.03123572 | 0.689304536 |
| *SUSD3* | -0.404371 | 0.187943 | -2.151569 | 0.03143134 | 0.6894829 |
| *TYMP* | 0.503205 | 0.234208 | 2.148536 | 0.03167117 | 0.6894829 |
| *ZNF362* | -0.736137 | 0.342639 | -2.148433 | 0.03167936 | 0.6894829 |
| *RPL22* | 0.118238 | 0.055043 | 2.148109 | 0.03170508 | 0.6894829 |
| *FAM111A* | -0.452441 | 0.210713 | -2.14719 | 0.03177814 | 0.6894829 |
| *USP31* | -1.255672 | 0.585036 | -2.146315 | 0.03184783 | 0.6894829 |
| *RABGGTA* | 0.744612 | 0.346971 | 2.146038 | 0.03186994 | 0.6894829 |
| *SAR1B* | -0.522322 | 0.24341 | -2.145855 | 0.03188458 | 0.6894829 |
| *GMEB1* | 0.578731 | 0.269726 | 2.145624 | 0.031903 | 0.6894829 |
| *PRDM2* | -0.249221 | 0.116186 | -2.145013 | 0.03195184 | 0.6894829 |
| *ARHGAP9* | 0.356961 | 0.166514 | 2.143733 | 0.03205431 | 0.6894829 |
| *ELMO3* | -1.116766 | 0.520956 | -2.143686 | 0.03205809 | 0.6894829 |
| *P2RX1* | 0.712146 | 0.332281 | 2.143203 | 0.03209683 | 0.6894829 |
| *MAATS1* | -2.023426 | 0.944487 | -2.142355 | 0.03216489 | NA |
| *ARID2* | -0.495741 | 0.231432 | -2.142064 | 0.03218837 | 0.6894829 |
| *C1orf21* | -2.971906 | 1.388085 | -2.141011 | 0.03227314 | NA |
| *GDI2* | 0.26631 | 0.124392 | 2.140892 | 0.03228275 | 0.6894829 |
| *LSM12* | -0.418613 | 0.195537 | -2.140835 | 0.03228737 | 0.6894829 |
| *LSM11* | -1.102105 | 0.514856 | -2.140607 | 0.03230571 | 0.6894829 |
| *HSD3B7* | -2.163566 | 1.01075 | -2.140555 | 0.03230995 | NA |

| | | | | | |
|---|---|---|---|---|---|
| *CTSZ* | 0.393507 | 0.183859 | 2.140261 | 0.0323337 | 0.6894829 |
| *COL18A1* | -2.465398 | 1.152334 | -2.139483 | 0.03239655 | NA |
| *ENSG00000236617* | -1.564454 | 0.73159 | -2.138431 | 0.03248175 | NA |
| *MAPK3* | 0.85259 | 0.398937 | 2.137155 | 0.03258537 | 0.693550617 |
| *ENSG00000276216* | -1.651082 | 0.772824 | -2.136426 | 0.03264472 | NA |
| *ZNF653* | -1.235282 | 0.578236 | -2.136294 | 0.03265543 | 0.693745081 |
| *PRR14* | -0.37983 | 0.177967 | -2.134273 | 0.03282048 | 0.693803074 |
| *BMPR2* | 0.896086 | 0.419901 | 2.13404 | 0.03283953 | 0.693803074 |
| *PGM1* | -0.713674 | 0.334427 | -2.134022 | 0.03284095 | 0.693803074 |
| *ERAL1* | -0.561169 | 0.263236 | -2.131814 | 0.03302218 | 0.693895289 |
| *CLCN6* | 0.859149 | 0.403061 | 2.131561 | 0.03304293 | 0.693895289 |
| *FAM3A* | -0.457097 | 0.21445 | -2.131487 | 0.03304904 | 0.693895289 |
| *C1RL.AS1* | 1.557306 | 0.730958 | 2.130499 | 0.03313046 | 0.693895289 |
| *C21orf58* | -1.06669 | 0.500777 | -2.130071 | 0.03316574 | 0.693895289 |
| *IMPDH1* | 0.802704 | 0.376941 | 2.129524 | 0.03321094 | 0.693895289 |
| *EDN1* | -2.011547 | 0.946179 | -2.125968 | 0.03350589 | NA |
| *CIAO1* | -0.460967 | 0.217123 | -2.12307 | 0.03374796 | 0.7038242 |
| *THADA* | -0.582443 | 0.274905 | -2.118708 | 0.03411514 | 0.704196182 |
| *UCP3* | -1.447442 | 0.683279 | -2.118376 | 0.03414325 | 0.704196182 |
| *NT5C* | -0.304044 | 0.143549 | -2.118049 | 0.0341709 | 0.704196182 |
| *ENSG00000269910* | -1.892038 | 0.893297 | -2.118038 | 0.03417186 | NA |
| *TAF1A* | 1.114823 | 0.526601 | 2.117015 | 0.03425862 | 0.704196182 |
| *ENSG00000259888* | -2.640074 | 1.247196 | -2.116808 | 0.03427616 | NA |
| *LTBP4* | 0.935006 | 0.442043 | 2.115192 | 0.03441355 | 0.704196182 |
| *S1PR1* | -0.357955 | 0.169261 | -2.11481 | 0.03444612 | 0.704196182 |
| *FAM81A* | -1.577 | 0.746135 | -2.11356 | 0.03455287 | NA |
| *C17orf67* | -1.006031 | 0.476452 | -2.111503 | 0.03472912 | 0.704196182 |
| *ENSG00000247363* | -1.200022 | 0.568358 | -2.111384 | 0.0347393 | 0.704196182 |
| *ZNF606* | -0.420954 | 0.199446 | -2.110619 | 0.03480507 | 0.704196182 |
| *LINC01521* | -1.055928 | 0.50031 | -2.110549 | 0.0348111 | 0.704196182 |
| *NAT9* | -0.404117 | 0.191512 | -2.110141 | 0.0348462 | 0.704196182 |
| *GIT2* | 0.328878 | 0.155958 | 2.108763 | 0.03496504 | 0.704196182 |
| *DDX47* | -1.235052 | 0.586051 | -2.107413 | 0.03508183 | 0.704196182 |
| *ZNF182* | -0.7312 | 0.347011 | -2.107138 | 0.03510558 | 0.704196182 |
| *ENSG00000232533* | 1.427488 | 0.677481 | 2.107055 | 0.03511285 | 0.704196182 |
| *USP4* | 0.561046 | 0.266285 | 2.106934 | 0.03512333 | 0.704196182 |
| *DOCK5* | -1.663217 | 0.790353 | -2.104399 | 0.03534367 | 0.704196182 |
| *SLF1* | 0.591046 | 0.280942 | 2.103796 | 0.03539624 | 0.704196182 |
| *TMED4* | -0.291818 | 0.138774 | -2.102839 | 0.03547984 | 0.704196182 |
| *SLC30A1* | -1.249182 | 0.594081 | -2.102713 | 0.03549087 | 0.704196182 |
| *GINM1* | -0.437584 | 0.208134 | -2.102412 | 0.0355172 | 0.704196182 |
| *ZNF280B* | -0.822919 | 0.391516 | -2.101879 | 0.03556388 | 0.704196182 |
| *BAZ2A* | -0.259785 | 0.123646 | -2.101039 | 0.03563755 | 0.704196182 |
| *RPS6* | 0.130908 | 0.062341 | 2.099878 | 0.03573955 | 0.704196182 |
| *ARHGAP22* | -1.347964 | 0.641994 | -2.099652 | 0.0357595 | 0.704196182 |
| *MEF2A* | 0.43563 | 0.207533 | 2.099085 | 0.03580939 | 0.704196182 |
| *UAP1* | -0.588977 | 0.280611 | -2.098912 | 0.03582468 | 0.704196182 |

| | | | | | |
|---|---|---|---|---|---|
| *ENSG00000258768* | -1.275899 | 0.60795 | -2.09869 | 0.03584424 | 0.704196182 |
| *SLC17A5* | 1.231152 | 0.586819 | 2.098008 | 0.03590442 | 0.704196182 |
| *MAP3K11* | 0.568674 | 0.271135 | 2.09738 | 0.03595993 | 0.704196182 |
| *PEAK3* | -1.49946 | 0.714971 | -2.097232 | 0.03597301 | 0.704196182 |
| *SMAP1* | -0.352075 | 0.167882 | -2.097153 | 0.03597999 | 0.704196182 |
| *ZNF543* | -0.80181 | 0.382459 | -2.096458 | 0.0360416 | 0.704196182 |
| *ENSG00000273319* | 0.346128 | 0.165117 | 2.096257 | 0.03605936 | 0.704196182 |
| *PRELID2* | -1.095932 | 0.52291 | -2.095833 | 0.03609699 | 0.704196182 |
| *CD3EAP* | 1.257697 | 0.600225 | 2.095376 | 0.03613755 | 0.704196182 |
| *NGLY1* | -0.262321 | 0.125197 | -2.095258 | 0.03614805 | 0.704196182 |
| *ENSG00000223821* | -1.388817 | 0.662876 | -2.095139 | 0.03615861 | 0.704196182 |
| *CPNE5* | 0.614822 | 0.293482 | 2.094925 | 0.03617764 | 0.704196182 |
| *EFCAB13* | -0.837672 | 0.400312 | -2.092547 | 0.03638958 | 0.705192613 |
| *RGP1* | -0.381159 | 0.182182 | -2.092184 | 0.03642209 | 0.705192613 |
| *CLDND2* | 1.261046 | 0.603126 | 2.09085 | 0.03654148 | 0.705192613 |
| *PRDM4* | 0.858006 | 0.410402 | 2.09065 | 0.03655944 | 0.705192613 |
| *C15orf62* | -0.825882 | 0.395072 | -2.090462 | 0.03657635 | 0.705192613 |
| *AP1G2* | -0.300823 | 0.143954 | -2.08971 | 0.03664385 | 0.705192613 |
| *FMO4* | -1.369492 | 0.655415 | -2.089504 | 0.03666234 | 0.705192613 |
| *ATP6V1E1* | 0.433479 | 0.207537 | 2.088688 | 0.03673586 | 0.705415216 |
| *TYSND1* | 0.878365 | 0.420725 | 2.087741 | 0.03682119 | 0.705863384 |
| *LMO7* | -0.78062 | 0.374096 | -2.086685 | 0.03691661 | 0.706503326 |
| *ADAM9* | -1.848007 | 0.885877 | -2.086077 | 0.03697162 | NA |
| *FBXO5* | 1.000934 | 0.479952 | 2.085489 | 0.03702489 | 0.707386682 |
| *MYBL1* | -1.529004 | 0.733891 | -2.083422 | 0.03721276 | 0.709774252 |
| *AKAP17A* | -0.337113 | 0.16196 | -2.081454 | 0.03739239 | 0.709774252 |
| *OSM* | -1.577968 | 0.758301 | -2.080925 | 0.03744077 | 0.709774252 |
| *ZNF146* | -0.377569 | 0.181446 | -2.080893 | 0.03744372 | 0.709774252 |
| *FAM230J* | -1.572793 | 0.755972 | -2.08049 | 0.03748064 | 0.709774252 |
| *C8orf37* | -1.042266 | 0.501085 | -2.080018 | 0.03752385 | 0.709774252 |
| *RHOA* | 0.201274 | 0.096809 | 2.079086 | 0.03760945 | 0.709961027 |
| *ENSG00000251417* | -1.504543 | 0.723844 | -2.078545 | 0.03765916 | NA |
| *ATP2C1* | 0.885978 | 0.426275 | 2.078419 | 0.03767077 | 0.709961027 |
| *SCAND1* | 0.244952 | 0.118013 | 2.075643 | 0.03792701 | 0.709961027 |
| *SLC23A2* | 1.117248 | 0.538267 | 2.075637 | 0.03792755 | 0.709961027 |
| *ELMSAN1* | -0.317193 | 0.152835 | -2.075396 | 0.03794982 | 0.709961027 |
| *ANK1* | -1.327126 | 0.639597 | -2.074942 | 0.03799193 | 0.709961027 |
| *ATP6V1C2* | -1.530569 | 0.738351 | -2.072957 | 0.0381763 | 0.709961027 |
| *RABAC1* | 0.225555 | 0.108817 | 2.072792 | 0.03819161 | 0.709961027 |
| *AZIN1* | 0.55654 | 0.268511 | 2.072693 | 0.03820087 | 0.709961027 |
| *TTC30A* | -1.561307 | 0.753291 | -2.072649 | 0.03820499 | 0.709961027 |
| *YDJC* | -0.323752 | 0.15625 | -2.072016 | 0.03826395 | 0.709961027 |
| *WRNIP1* | 1.108753 | 0.535296 | 2.071291 | 0.03833164 | 0.709961027 |
| *ARF5* | -0.287739 | 0.138927 | -2.071156 | 0.03834426 | 0.709961027 |
| *CASD1* | -0.649313 | 0.313677 | -2.070001 | 0.03845227 | 0.710016833 |
| *CNOT11* | -0.588521 | 0.28437 | -2.069564 | 0.03849319 | 0.710016833 |
| *ZNF561* | 0.646407 | 0.312406 | 2.069125 | 0.03853433 | 0.710016833 |

| | | | | | |
|---|---|---|---|---|---|
| *ANKRD28* | 0.961773 | 0.465639 | 2.065493 | 0.03887638 | 0.714167523 |
| *MAP2K6* | -1.365669 | 0.661241 | -2.06531 | 0.03889368 | 0.714167523 |
| *METTL7A* | 0.411379 | 0.199254 | 2.064599 | 0.03896097 | 0.714167523 |
| *ENSG00000276718* | -1.649323 | 0.798957 | -2.064346 | 0.0389849 | NA |
| *NCF1* | 0.184135 | 0.089243 | 2.063304 | 0.03908378 | 0.714167523 |
| *GGA2* | 0.263789 | 0.127916 | 2.062215 | 0.03918723 | 0.714167523 |
| *TMBIM1* | -0.419115 | 0.203281 | -2.061754 | 0.03923119 | 0.714167523 |
| *BBS4* | 0.885316 | 0.429445 | 2.061538 | 0.03925178 | 0.714167523 |
| *C6orf52* | -1.573911 | 0.763502 | -2.061437 | 0.03926134 | 0.714167523 |
| *HAL* | -1.588908 | 0.771341 | -2.059931 | 0.03940516 | 0.714645898 |
| *TESPA1* | 0.996182 | 0.48362 | 2.059847 | 0.03941316 | 0.714645898 |
| *SCRN2* | -0.466816 | 0.226721 | -2.058984 | 0.03949579 | 0.715005713 |
| *INKA1* | -0.458592 | 0.222916 | -2.057238 | 0.03966338 | 0.716899903 |
| *ARRDC3* | 0.440062 | 0.214088 | 2.05552 | 0.03982881 | 0.717022016 |
| *AP1M1* | -0.393735 | 0.19156 | -2.055412 | 0.03983919 | 0.717022016 |
| *C19orf66* | 0.322227 | 0.156785 | 2.055207 | 0.03985904 | 0.717022016 |
| *MIER1* | -0.350878 | 0.170825 | -2.054017 | 0.039974 | 0.717955812 |
| *ENSG00000258944* | -1.549733 | 0.754791 | -2.053194 | 0.04005376 | NA |
| *TMCC3* | -1.494838 | 0.728073 | -2.053143 | 0.04005869 | NA |
| *ZNF442* | -1.265285 | 0.617119 | -2.050309 | 0.04033431 | 0.721999775 |
| *BCAT1* | -2.447301 | 1.194789 | -2.048312 | 0.04052944 | NA |
| *TOB1* | 0.834748 | 0.407931 | 2.046297 | 0.04072711 | 0.721999775 |
| *DDX59* | 0.755326 | 0.369139 | 2.046186 | 0.04073808 | 0.721999775 |
| *NDUFB6* | 0.411155 | 0.200999 | 2.045562 | 0.04079948 | 0.721999775 |
| *EIF3M* | 0.224352 | 0.109705 | 2.045052 | 0.0408497 | 0.721999775 |
| *COX10* | -0.870564 | 0.425837 | -2.04436 | 0.04091799 | 0.721999775 |
| *ENSG00000203546* | -1.128706 | 0.552248 | -2.043839 | 0.04096947 | 0.721999775 |
| *SYT15* | -1.416541 | 0.693117 | -2.043724 | 0.04098081 | 0.721999775 |
| *NQO1* | -1.385863 | 0.67821 | -2.043412 | 0.04101166 | 0.721999775 |
| *EMC6* | -0.340082 | 0.166437 | -2.043305 | 0.04102227 | 0.721999775 |
| *LBR* | 0.355334 | 0.173946 | 2.042779 | 0.04107428 | 0.721999775 |
| *ITGB1BP1* | 0.508464 | 0.249032 | 2.041758 | 0.04117555 | 0.721999775 |
| *ZNF253* | 0.708087 | 0.346876 | 2.041326 | 0.04121847 | 0.721999775 |
| *IDS* | -0.222205 | 0.108895 | -2.040538 | 0.04129673 | 0.721999775 |
| *ENSG00000179094* | 1.338349 | 0.655912 | 2.04044 | 0.04130656 | 0.721999775 |
| *CHL1* | 1.545735 | 0.757628 | 2.04023 | 0.04132745 | 0.721999775 |
| *RBM7* | 0.421241 | 0.206487 | 2.040034 | 0.04134691 | 0.721999775 |
| *RPL7A* | 0.100561 | 0.0493 | 2.039774 | 0.0413728 | 0.721999775 |
| *SIGLEC14* | 1.224206 | 0.600488 | 2.038685 | 0.04148147 | 0.721999775 |
| *ADAT1* | -0.635607 | 0.31182 | -2.038376 | 0.04151237 | 0.721999775 |
| *ERGIC1* | -0.31111 | 0.15264 | -2.038193 | 0.04153068 | 0.721999775 |
| *ITGAV* | -1.50154 | 0.73697 | -2.03745 | 0.04160495 | 0.722188301 |
| *SNHG21* | -0.496309 | 0.243706 | -2.036504 | 0.04169977 | 0.722732564 |
| *FCHO1* | 0.705826 | 0.347031 | 2.033898 | 0.04196191 | 0.726170622 |
| *ARRDC4* | -1.659617 | 0.816625 | -2.032289 | 0.04212441 | NA |
| *ITPR1* | 0.489385 | 0.241139 | 2.029476 | 0.0424098 | 0.732807855 |
| *DUBR* | -1.627262 | 0.802303 | -2.028239 | 0.04253587 | 0.733136166 |

| | | | | | |
|---|---|---|---|---|---|
| *GNG5* | -0.187496 | 0.092453 | -2.028026 | 0.04255757 | 0.733136166 |
| *CCNA1* | -2.989636 | 1.474168 | -2.028015 | 0.04255868 | NA |
| *ABHD14A* | 0.628427 | 0.310134 | 2.026309 | 0.04273317 | 0.734299645 |
| *C3orf38* | 0.551625 | 0.272259 | 2.026105 | 0.04275408 | 0.734299645 |
| *RFTN1* | 0.438298 | 0.216433 | 2.025095 | 0.04285759 | 0.734968889 |
| *RSPH9* | -2.032226 | 1.004436 | -2.023251 | 0.04304725 | NA |
| *SPPL2B* | -0.27487 | 0.135882 | -2.022854 | 0.04308824 | 0.737813196 |
| *AKIRIN1* | -0.319612 | 0.158065 | -2.022025 | 0.04317381 | 0.738168338 |
| *GLI4* | -1.044792 | 0.517466 | -2.019055 | 0.04348153 | 0.741832366 |
| *UNC119B* | 1.265676 | 0.627409 | 2.017306 | 0.04366363 | 0.741832366 |
| *STT3A* | 1.021139 | 0.506292 | 2.016899 | 0.04370608 | 0.741832366 |
| *UTP3* | 0.469303 | 0.232713 | 2.016662 | 0.04373082 | 0.741832366 |
| *ENSG00000273272* | -1.541436 | 0.764442 | -2.016421 | 0.04375601 | NA |
| *LRRCC1* | 1.406848 | 0.697715 | 2.016364 | 0.04376189 | 0.741832366 |
| *MAP2K4* | 1.037883 | 0.514772 | 2.016201 | 0.04377899 | 0.741832366 |
| *SPATS2L* | -1.852171 | 0.918648 | -2.016193 | 0.04377983 | NA |
| *MIA2* | -0.511335 | 0.25374 | -2.015197 | 0.04388399 | 0.742277696 |
| *VTA1* | 0.748546 | 0.371541 | 2.014704 | 0.04393564 | 0.742277696 |
| *MLX* | -0.321617 | 0.159765 | -2.013065 | 0.04410781 | 0.74408246 |
| *LRRC46* | -1.373395 | 0.682847 | -2.01128 | 0.04429594 | 0.746150648 |
| *KCNRG* | -1.349121 | 0.671376 | -2.009487 | 0.04448556 | 0.747585861 |
| *RMND5B* | -0.390603 | 0.194448 | -2.008774 | 0.04456108 | 0.747585861 |
| *GRK6* | -0.569985 | 0.28377 | -2.008614 | 0.0445781 | 0.747585861 |
| *TNFRSF12A* | -1.733783 | 0.863221 | -2.008503 | 0.04458984 | NA |
| *DTX2* | -0.690866 | 0.344684 | -2.004348 | 0.04503284 | 0.754101454 |
| *OPA3* | -0.555181 | 0.27708 | -2.003681 | 0.04510428 | 0.754188651 |
| *KDM4A.AS1* | -1.170488 | 0.584377 | -2.002966 | 0.04518092 | 0.754362298 |
| *ENSG00000185527* | 1.480781 | 0.739556 | 2.002256 | 0.04525717 | 0.754529201 |
| *RBBP9* | -0.689774 | 0.344674 | -2.001234 | 0.04536715 | 0.75525698 |
| *TFB2M* | 0.969339 | 0.484844 | 1.99928 | 0.04557807 | 0.757660509 |
| *IL4I1* | -2.141782 | 1.072162 | -1.99763 | 0.0457568 | NA |
| *MTRF1* | -0.60353 | 0.30219 | -1.997185 | 0.0458051 | 0.760324573 |
| *MFSD10* | -0.358158 | 0.179513 | -1.995169 | 0.04602443 | 0.761320511 |
| *CALU* | -0.589675 | 0.295683 | -1.99428 | 0.04612149 | 0.761320511 |
| *POLE3* | -0.28727 | 0.144062 | -1.994077 | 0.04614362 | 0.761320511 |
| *DENND5A* | 0.941901 | 0.472352 | 1.994067 | 0.04614472 | 0.761320511 |
| *GLCE* | -0.991229 | 0.497214 | -1.993567 | 0.04619939 | 0.761320511 |
| *LINC01816* | -1.470278 | 0.737836 | -1.99269 | 0.04629537 | 0.761799751 |
| *PERP* | -1.797468 | 0.902063 | -1.992619 | 0.04630318 | NA |
| *TLR10* | 0.522375 | 0.262473 | 1.990199 | 0.04656897 | 0.762238737 |
| *BROX* | -0.30234 | 0.152057 | -1.98834 | 0.04677414 | 0.762238737 |
| *GOLIM4* | 1.124515 | 0.565593 | 1.988206 | 0.04678891 | 0.762238737 |
| *TRPM7* | 0.485001 | 0.243989 | 1.987795 | 0.04683441 | 0.762238737 |
| *EIF4EBP1* | 0.405913 | 0.204235 | 1.987476 | 0.04686965 | 0.762238737 |
| *HIVEP2* | 0.886916 | 0.446638 | 1.985761 | 0.04705989 | 0.762238737 |
| *ENSG00000268713* | -0.986326 | 0.496749 | -1.985561 | 0.04708204 | 0.762238737 |
| *MAFG.DT* | -1.525398 | 0.768356 | -1.985274 | 0.04711402 | 0.762238737 |

| | | | | | |
|---|---|---|---|---|---|
| *DHRS3* | 1.946255 | 0.980503 | 1.984957 | 0.04714928 | 0.762238737 |
| *CLCN7* | 0.483556 | 0.243629 | 1.984804 | 0.04716623 | 0.762238737 |
| *ZNF607* | -1.04922 | 0.528706 | -1.984505 | 0.04719954 | 0.762238737 |
| *ZNF524* | -0.357734 | 0.18033 | -1.983772 | 0.04728125 | 0.762238737 |
| *EXOC1* | 0.599343 | 0.302167 | 1.983485 | 0.04731333 | 0.762238737 |
| *LMO2* | -1.776133 | 0.895823 | -1.982683 | 0.04740285 | NA |
| *IKZF1* | 0.36683 | 0.18504 | 1.982437 | 0.04743034 | 0.762238737 |
| *SLC5A5* | 1.53602 | 0.774984 | 1.982002 | 0.04747903 | 0.762238737 |
| *CIB1* | 0.167983 | 0.08478 | 1.981395 | 0.04754698 | 0.762238737 |
| *ACVR1B* | -1.244522 | 0.628126 | -1.981324 | 0.04755493 | 0.762238737 |
| *SREBF2.AS1* | 1.316854 | 0.664637 | 1.981313 | 0.04755619 | 0.762238737 |
| *TNNI2* | 1.395194 | 0.70431 | 1.980939 | 0.04759808 | 0.762238737 |
| *CASK* | -0.869079 | 0.438977 | -1.979785 | 0.04772766 | 0.762238737 |
| *NRBF2* | -0.374689 | 0.189349 | -1.978831 | 0.04783507 | 0.762238737 |
| *CENPS.CORT* | -1.687807 | 0.853126 | -1.978379 | 0.04788593 | 0.762238737 |
| *C19orf73* | -1.069167 | 0.54064 | -1.977596 | 0.04797425 | 0.762238737 |
| *FGR* | 0.80077 | 0.405187 | 1.976299 | 0.04812087 | 0.762238737 |
| *RBM18* | 0.692092 | 0.350203 | 1.976258 | 0.04812553 | 0.762238737 |
| *USP5* | -0.563911 | 0.285389 | -1.975941 | 0.04816143 | 0.762238737 |
| *MYOM1* | -1.063744 | 0.538392 | -1.975779 | 0.04817982 | 0.762238737 |
| *ENSG00000263335* | -1.227053 | 0.621093 | -1.975633 | 0.04819636 | 0.762238737 |
| *ZFP69B* | -1.605911 | 0.812955 | -1.9754 | 0.04822276 | NA |
| *HPSE* | -1.665608 | 0.843185 | -1.975376 | 0.04822548 | NA |
| *CCDC151* | -1.565717 | 0.79294 | -1.974573 | 0.04831666 | NA |
| *AGAP4* | 1.347108 | 0.682413 | 1.974037 | 0.04837754 | 0.76239884 |
| *EXOC4* | 0.529862 | 0.268514 | 1.973315 | 0.04845964 | 0.76239884 |
| *RCL1* | 0.882282 | 0.447206 | 1.972874 | 0.04850989 | 0.76239884 |
| *LINC02245* | -1.11363 | 0.564486 | -1.972822 | 0.04851583 | 0.76239884 |
| *PUS1* | 0.63322 | 0.321008 | 1.972599 | 0.04854125 | 0.76239884 |
| *KATNBL1* | -0.319536 | 0.162165 | -1.97044 | 0.04878796 | 0.764514148 |
| *KYAT3* | -0.514857 | 0.26136 | -1.969918 | 0.0488478 | 0.764514148 |
| *MPP7* | 1.372286 | 0.696779 | 1.969471 | 0.04889904 | 0.764514148 |
| *ADAP1* | -1.519289 | 0.771575 | -1.969075 | 0.04894448 | 0.764514148 |
| *PEX6* | 1.084709 | 0.551327 | 1.96745 | 0.04913139 | 0.765684782 |
| *CENPB* | 0.999717 | 0.508223 | 1.967086 | 0.04917331 | 0.765684782 |
| *SERGEF* | -0.465176 | 0.23653 | -1.966671 | 0.04922115 | 0.765684782 |
| *ADARB1* | -1.004348 | 0.510855 | -1.966013 | 0.04929708 | 0.765819701 |

| Cluster 5 | log2FC | lfcSE | stat | pvalue | padj |
|---|---|---|---|---|---|
| *VPREB3* | 3.700958 | 0.109845 | 33.692593 | < 2.22e-16 | < 2.22e-16 |
| *PLAAT4* | 2.964256 | 0.13949 | 21.250729 | < 2.22e-16 | < 2.22e-16 |
| *DENND6B* | 3.435999 | 0.198172 | 17.3385 | < 2.22e-16 | < 2.22e-16 |
| *GRN* | 1.851849 | 0.112157 | 16.511196 | < 2.22e-16 | < 2.22e-16 |
| *CD38* | 5.601764 | 0.374535 | 14.956598 | < 2.22e-16 | < 2.22e-16 |
| *CD1D* | 3.892037 | 0.261573 | 14.879352 | < 2.22e-16 | < 2.22e-16 |
| *ASB2* | 4.829368 | 0.326659 | 14.784131 | < 2.22e-16 | < 2.22e-16 |
| *RABGAP1L* | 1.629215 | 0.121193 | 13.443113 | < 2.22e-16 | < 2.22e-16 |
| *CNP* | 1.783749 | 0.152885 | 11.667247 | < 2.22e-16 | < 2.22e-16 |
| *RNF207* | 3.11202 | 0.293551 | 10.601302 | < 2.22e-16 | < 2.22e-16 |
| *C12orf65* | 1.214399 | 0.118834 | 10.219329 | < 2.22e-16 | < 2.22e-16 |
| *SEPTIN9* | 0.642307 | 0.062991 | 10.1968 | < 2.22e-16 | < 2.22e-16 |
| *ENDOD1* | 1.633086 | 0.171849 | 9.503002 | < 2.22e-16 | < 2.22e-16 |
| *P2RX5* | 1.621094 | 0.171127 | 9.473039 | < 2.22e-16 | < 2.22e-16 |
| *PNOC* | 1.076261 | 0.114763 | 9.378134 | < 2.22e-16 | < 2.22e-16 |
| *SLC12A4* | 2.283579 | 0.247878 | 9.212494 | < 2.22e-16 | < 2.22e-16 |
| *DUS2* | 1.338061 | 0.147912 | 9.046335 | < 2.22e-16 | < 2.22e-16 |
| *NRIP1* | 1.94529 | 0.221981 | 8.763306 | < 2.22e-16 | 1.36E-15 |
| *SIGIRR* | 0.945989 | 0.112723 | 8.392181 | < 2.22e-16 | 3.24E-14 |
| *SLC5A3* | 1.217119 | 0.147147 | 8.27148 | < 2.22e-16 | 8.52E-14 |
| *RAB20* | 2.567239 | 0.321534 | 7.984338 | 1.41E-15 | 8.67E-13 |
| *DDIT4* | 1.452503 | 0.186226 | 7.799666 | 6.21E-15 | 3.64E-12 |
| *RAB37* | 1.16335 | 0.149549 | 7.77904 | 7.31E-15 | 4.09E-12 |
| *MAP3K1* | 0.738729 | 0.096115 | 7.685909 | 1.52E-14 | 8.16E-12 |
| *MPEG1* | 1.513519 | 0.200536 | 7.547374 | 4.44E-14 | 2.29E-11 |
| *RARA.AS1* | 2.419415 | 0.324355 | 7.459164 | 8.71E-14 | 4.32E-11 |
| *DNAJC7* | 0.466735 | 0.063728 | 7.323818 | 2.41E-13 | 1.15E-10 |
| *AGPAT5* | 1.146471 | 0.162161 | 7.069943 | 1.55E-12 | 7.08E-10 |
| *CACYBP* | 0.562876 | 0.079659 | 7.066043 | 1.59E-12 | 7.08E-10 |
| *AHI1* | 1.140805 | 0.163239 | 6.988552 | 2.78E-12 | 1.19E-09 |
| *HIC1* | 3.093494 | 0.444076 | 6.966134 | 3.26E-12 | 1.35E-09 |
| *JCHAIN* | 1.6258 | 0.2349 | 6.921252 | 4.48E-12 | 1.80E-09 |
| *P2RX5.TAX1BP3* | 1.49181 | 0.22024 | 6.773567 | 1.26E-11 | 4.91E-09 |
| *IRAK2* | 1.341597 | 0.199227 | 6.734012 | 1.65E-11 | 6.26E-09 |
| *C15orf39* | 0.955626 | 0.142441 | 6.708905 | 1.96E-11 | 7.22E-09 |
| *SIGLEC10* | 0.930928 | 0.139691 | 6.664189 | 2.66E-11 | 9.53E-09 |
| *S100A4* | -0.882047 | 0.133314 | -6.616288 | 3.68E-11 | 1.26E-08 |
| *PPCDC* | 1.300801 | 0.196726 | 6.612239 | 3.79E-11 | 1.26E-08 |
| *CLIC3* | 2.072955 | 0.313544 | 6.611369 | 3.81E-11 | 1.26E-08 |
| *ANAPC16* | 0.299951 | 0.045763 | 6.554477 | 5.58E-11 | 1.80E-08 |
| *SLC2A5* | 2.173665 | 0.335077 | 6.487061 | 8.75E-11 | 2.75E-08 |
| *HTR3A* | 1.607236 | 0.250088 | 6.42667 | 1.30E-10 | 4.00E-08 |
| *MRPS6* | 1.036729 | 0.162541 | 6.378266 | 1.79E-10 | 5.37E-08 |
| *PLAAT3* | 2.236839 | 0.352403 | 6.34739 | 2.19E-10 | 6.41E-08 |
| *TRABD* | 0.501229 | 0.079751 | 6.284896 | 3.28E-10 | 9.39E-08 |
| *NUP210* | 0.513913 | 0.084782 | 6.061546 | 1.35E-09 | 3.78E-07 |

| | | | | | |
|---|---|---|---|---|---|
| *PPP1R18* | 0.514904 | 0.085277 | 6.038049 | 1.56E-09 | 4.28E-07 |
| *CR2* | 1.690965 | 0.280291 | 6.032888 | 1.61E-09 | 4.32E-07 |
| *DPEP2* | 1.009331 | 0.167751 | 6.016857 | 1.78E-09 | 4.68E-07 |
| *GBP4* | 0.917511 | 0.153686 | 5.970052 | 2.37E-09 | 6.09E-07 |
| *PSMB10* | 0.509484 | 0.085378 | 5.96741 | 2.41E-09 | 6.09E-07 |
| *TSPAN18* | 3.947041 | 0.669404 | 5.896354 | 3.72E-09 | 9.21E-07 |
| *SERPINF1* | -1.472244 | 0.251135 | -5.86237 | 4.56E-09 | 1.11E-06 |
| *ITGAL* | 0.691671 | 0.118201 | 5.851628 | 4.87E-09 | 1.16E-06 |
| *MS4A1* | -0.726376 | 0.126051 | -5.76256 | 8.28E-09 | 1.94E-06 |
| *VPREB1* | 3.752328 | 0.654167 | 5.73604 | 9.69E-09 | 2.23E-06 |
| *ZFP36L1* | -0.456848 | 0.080141 | -5.700527 | 1.19E-08 | 2.70E-06 |
| *LCAT* | 1.940799 | 0.341016 | 5.691229 | 1.26E-08 | 2.80E-06 |
| *GPR183* | -0.501253 | 0.088168 | -5.685182 | 1.31E-08 | 2.85E-06 |
| *DHRS3* | 4.017502 | 0.708439 | 5.67092 | 1.42E-08 | 3.02E-06 |
| *CD1C* | 1.589152 | 0.280287 | 5.669726 | 1.43E-08 | 3.02E-06 |
| *SLC16A3* | 0.862125 | 0.152764 | 5.643508 | 1.67E-08 | 3.46E-06 |
| *VHL* | 0.656437 | 0.116673 | 5.626302 | 1.84E-08 | 3.77E-06 |
| *IFI30* | 0.883572 | 0.158404 | 5.577961 | 2.43E-08 | 4.90E-06 |
| *ETFB* | 0.651573 | 0.116903 | 5.573598 | 2.50E-08 | 4.95E-06 |
| *FAM3C* | 0.59137 | 0.106188 | 5.569088 | 2.56E-08 | 5.00E-06 |
| *RMDN2* | 1.60587 | 0.290199 | 5.533687 | 3.14E-08 | 6.03E-06 |
| *GRINA* | 0.653072 | 0.122196 | 5.344454 | 9.07E-08 | 1.72E-05 |
| *EVI2B* | 0.318786 | 0.059889 | 5.322903 | 1.02E-07 | 1.91E-05 |
| *CD72* | 0.944799 | 0.178529 | 5.29213 | 1.21E-07 | 2.23E-05 |
| *RRBP1* | 1.359371 | 0.257466 | 5.279808 | 1.29E-07 | 2.35E-05 |
| *CD24* | -0.821602 | 0.157056 | -5.231266 | 1.68E-07 | 3.01E-05 |
| *PLP2* | -0.428349 | 0.083036 | -5.158613 | 2.49E-07 | 4.39E-05 |
| *ABI3* | 0.587428 | 0.114119 | 5.14751 | 2.64E-07 | 4.60E-05 |
| *MAPKAPK2* | 0.889025 | 0.172959 | 5.140096 | 2.75E-07 | 4.72E-05 |
| *FCRL2* | 1.008846 | 0.197075 | 5.119092 | 3.07E-07 | 5.21E-05 |
| *SEL1L3* | 0.768423 | 0.150693 | 5.099277 | 3.41E-07 | 5.71E-05 |
| *PIP5K1B* | 1.379175 | 0.272721 | 5.05709 | 4.26E-07 | 7.03E-05 |
| *CTSZ* | 0.586082 | 0.11626 | 5.041114 | 4.63E-07 | 7.55E-05 |
| *GLIPR2* | 0.850813 | 0.168928 | 5.036544 | 4.74E-07 | 7.64E-05 |
| *KCNH4* | 3.584812 | 0.713398 | 5.02498 | 5.03E-07 | 8.01E-05 |
| *C5orf56* | 0.788613 | 0.157873 | 4.995247 | 5.88E-07 | 9.23E-05 |
| *NAPSA* | 1.878063 | 0.376612 | 4.986726 | 6.14E-07 | 9.53E-05 |
| *CTSA* | 0.722062 | 0.145153 | 4.97449 | 6.54E-07 | 0.000100357 |
| *GDPGP1* | 1.371444 | 0.280366 | 4.891619 | 1.00E-06 | 0.000151615 |
| *FCRL1* | 0.715618 | 0.146895 | 4.871615 | 1.11E-06 | 0.000165855 |
| *PAXX* | 0.572733 | 0.117693 | 4.86632 | 1.14E-06 | 0.0001684 |
| *ENSG00000233038* | 4.065694 | 0.839693 | 4.841879 | 1.29E-06 | 0.000186682 |
| *VSIG10L* | 1.659909 | 0.342858 | 4.841387 | 1.29E-06 | 0.000186682 |
| *JUN* | 0.463564 | 0.09626 | 4.815748 | 1.47E-06 | 0.00020997 |
| *CARHSP1* | 0.290532 | 0.060496 | 4.802505 | 1.57E-06 | 0.000220952 |
| *ENSG00000260030* | 3.557533 | 0.740974 | 4.801159 | 1.58E-06 | 0.000220952 |
| *LSP1* | 0.296186 | 0.062438 | 4.743671 | 2.10E-06 | 0.00028925 |

| | | | | | |
|---|---|---|---|---|---|
| *ZNF3* | 0.669403 | 0.141147 | 4.742592 | 2.11E-06 | 0.00028925 |
| *TNFRSF13B* | -0.456744 | 0.096504 | -4.732881 | 2.21E-06 | 0.000300251 |
| *GTF2E2* | 0.530131 | 0.112373 | 4.71761 | 2.39E-06 | 0.000320314 |
| *C16orf54* | 0.619489 | 0.132778 | 4.665617 | 3.08E-06 | 0.000408756 |
| *RHOG* | -0.294041 | 0.063137 | -4.657214 | 3.21E-06 | 0.000421448 |
| *XYLT1* | 0.969948 | 0.208439 | 4.653393 | 3.27E-06 | 0.000424999 |
| *CD22* | 0.577297 | 0.124784 | 4.626377 | 3.72E-06 | 0.000475357 |
| *CPNE5* | 0.698905 | 0.151078 | 4.626118 | 3.73E-06 | 0.000475357 |
| *CPEB4* | 0.952137 | 0.206434 | 4.61231 | 3.98E-06 | 0.000503082 |
| *DERL3* | 0.608568 | 0.132198 | 4.603472 | 4.16E-06 | 0.000519826 |
| *RGS1* | -1.23171 | 0.268839 | -4.581596 | 4.61E-06 | 0.000571743 |
| *FCMR* | 0.405685 | 0.088989 | 4.55883 | 5.14E-06 | 0.000629575 |
| *SPOCK2* | 0.788657 | 0.173049 | 4.557408 | 5.18E-06 | 0.000629575 |
| *C16orf72* | 0.535427 | 0.117934 | 4.540056 | 5.62E-06 | 0.000677289 |
| *BTG2* | -0.349145 | 0.07733 | -4.515021 | 6.33E-06 | 0.000755388 |
| *GPX1* | 0.418863 | 0.092829 | 4.512202 | 6.42E-06 | 0.000758476 |
| *BTBD7* | 0.585085 | 0.129956 | 4.502166 | 6.73E-06 | 0.000787973 |
| *PHKB* | 0.53323 | 0.118617 | 4.495381 | 6.94E-06 | 0.000806192 |
| *BICDL1* | 1.344904 | 0.300526 | 4.475172 | 7.63E-06 | 0.000878432 |
| *GPR146* | -1.326492 | 0.297355 | -4.460965 | 8.16E-06 | 0.000930431 |
| *APOL3* | 0.732505 | 0.166252 | 4.405988 | 1.05E-05 | 0.001190292 |
| *ARL14EPL* | 3.188668 | 0.728453 | 4.377313 | 1.20E-05 | 0.001346325 |
| *SMAD3* | 0.817083 | 0.187022 | 4.368918 | 1.25E-05 | 0.001387063 |
| *MCM5* | 0.444662 | 0.101977 | 4.360415 | 1.30E-05 | 0.001429748 |
| *ENSG00000198106* | -0.900354 | 0.206925 | -4.351113 | 1.35E-05 | 0.001479141 |
| *PUS1* | 0.766459 | 0.176676 | 4.338212 | 1.44E-05 | 0.001555488 |
| *AGO1* | 0.706477 | 0.164682 | 4.289954 | 1.79E-05 | 0.001919048 |
| *ULK1* | 0.991701 | 0.231389 | 4.285868 | 1.82E-05 | 0.001938512 |
| *IRF2BPL* | 0.880022 | 0.205709 | 4.277984 | 1.89E-05 | 0.001991983 |
| *S100A6* | -0.531998 | 0.124511 | -4.272709 | 1.93E-05 | 0.002023129 |
| *IRF1* | 0.443977 | 0.10506 | 4.225953 | 2.38E-05 | 0.002472576 |
| *IL1B* | 3.408768 | 0.80965 | 4.210177 | 2.55E-05 | 0.002630504 |
| *LGALS3* | -0.671683 | 0.159683 | -4.206364 | 2.60E-05 | 0.002654025 |
| *RARA* | 0.539695 | 0.128388 | 4.20364 | 2.63E-05 | 0.002665028 |
| *ZNF70* | 1.588848 | 0.379716 | 4.184305 | 2.86E-05 | 0.002879617 |
| *ATXN10* | 0.496026 | 0.118729 | 4.1778 | 2.94E-05 | 0.002940229 |
| *PLXNB2* | 0.981605 | 0.235956 | 4.160124 | 3.18E-05 | 0.003152856 |
| *LHFPL2* | 1.138933 | 0.273967 | 4.157195 | 3.22E-05 | 0.003157567 |
| *PALLD* | 3.451708 | 0.830477 | 4.156294 | 3.23E-05 | 0.003157567 |
| *CHST15* | 1.003222 | 0.242978 | 4.128864 | 3.65E-05 | 0.003532114 |
| *AP3S1* | 0.554941 | 0.134525 | 4.125193 | 3.70E-05 | 0.003554596 |
| *ACSL1* | 0.896184 | 0.217311 | 4.12397 | 3.72E-05 | 0.003554596 |
| *TRIB1* | 0.855719 | 0.208511 | 4.103948 | 4.06E-05 | 0.003848358 |
| *CTSB* | 0.431515 | 0.105846 | 4.076824 | 4.57E-05 | 0.004294235 |
| *CD180* | 0.61154 | 0.151893 | 4.02614 | 5.67E-05 | 0.005294465 |
| *ENSG00000260979* | 1.444956 | 0.360575 | 4.007361 | 6.14E-05 | 0.005655598 |
| *FLNB* | 1.04812 | 0.26156 | 4.007191 | 6.14E-05 | 0.005655598 |

| | | | | | |
|---|---|---|---|---|---|
| *STK17B* | -0.35928 | 0.089767 | -4.002372 | 6.27E-05 | 0.005731123 |
| *KAT6A* | 0.515946 | 0.129016 | 3.999086 | 6.36E-05 | 0.00577035 |
| *ABCB1* | 1.635519 | 0.411374 | 3.975747 | 7.02E-05 | 0.006322144 |
| *HOXB3* | 2.026502 | 0.513363 | 3.947506 | 7.90E-05 | 0.007066667 |
| *TBL1X* | -0.755506 | 0.191676 | -3.941583 | 8.09E-05 | 0.007193553 |
| *CERK* | 0.73534 | 0.187019 | 3.931888 | 8.43E-05 | 0.007392072 |
| *TAF12* | 0.39048 | 0.099314 | 3.931759 | 8.43E-05 | 0.007392072 |
| *RINL* | 0.445053 | 0.113595 | 3.917904 | 8.93E-05 | 0.007777061 |
| *MIDN* | 0.5019 | 0.129679 | 3.870314 | 0.0001087 | 0.009400306 |
| *MCL1* | 0.480371 | 0.124311 | 3.86427 | 0.00011142 | 0.009571882 |
| *CR1* | 0.896373 | 0.232182 | 3.860647 | 0.00011309 | 0.009650598 |
| *VIM* | -0.550273 | 0.143357 | -3.83847 | 0.0001238 | 0.01049561 |
| *LMNA* | -1.583459 | 0.414097 | -3.823882 | 0.00013137 | 0.011064003 |
| *ENSG00000254802* | 1.091236 | 0.288097 | 3.787742 | 0.00015202 | 0.012720518 |
| *HIVEP2* | 0.80081 | 0.211763 | 3.781631 | 0.0001558 | 0.012936621 |
| *PLEKHO1* | 0.560609 | 0.148296 | 3.780342 | 0.00015661 | 0.012936621 |
| *SREBF1* | 1.547743 | 0.410277 | 3.772437 | 0.00016166 | 0.013268555 |
| *TNFRSF18* | -1.410252 | 0.374864 | -3.762036 | 0.00016854 | 0.013745266 |
| *FAM167A* | 1.264201 | 0.336428 | 3.757716 | 0.00017147 | 0.01389672 |
| *ZBTB32* | -0.886804 | 0.236827 | -3.744516 | 0.00018074 | 0.014556472 |
| *RABAC1* | 0.204748 | 0.054789 | 3.737041 | 0.0001862 | 0.014902838 |
| *AHNAK* | -0.491273 | 0.131972 | -3.722555 | 0.00019722 | 0.015687272 |
| *STK17A* | 0.446005 | 0.119965 | 3.717793 | 0.00020097 | 0.01588778 |
| *ZNF337.AS1* | 1.339585 | 0.360815 | 3.71266 | 0.00020509 | 0.016084353 |
| *CLDND2* | 1.136594 | 0.306228 | 3.7116 | 0.00020595 | 0.016084353 |
| *SPRY1* | 1.242648 | 0.335574 | 3.703049 | 0.00021302 | 0.016497535 |
| *CRYL1* | -0.83304 | 0.225017 | -3.702121 | 0.00021381 | 0.016497535 |
| *PDLIM2* | -0.537412 | 0.145352 | -3.697321 | 0.00021789 | 0.016712444 |
| *ABRACL* | -0.442637 | 0.119855 | -3.693095 | 0.00022154 | 0.016892195 |
| *GLYCTK* | 1.017642 | 0.276121 | 3.685494 | 0.00022826 | 0.017302069 |
| *SLC5A5* | 3.381942 | 0.918155 | 3.683409 | 0.00023014 | 0.017342258 |
| *PTK2B* | 0.490465 | 0.133722 | 3.667789 | 0.00024466 | 0.018329341 |
| *ERAP1* | 0.615803 | 0.168106 | 3.66319 | 0.00024909 | 0.018553842 |
| *TBC1D9* | 0.431841 | 0.118025 | 3.658907 | 0.00025329 | 0.018758279 |
| *LGALS1* | -0.805962 | 0.2205 | -3.655157 | 0.00025702 | 0.018870017 |
| *CD5* | -2.155243 | 0.589965 | -3.653173 | 0.00025902 | 0.018870017 |
| *TMEM159* | -0.563205 | 0.154176 | -3.652999 | 0.0002592 | 0.018870017 |
| *CXCR5* | 0.438386 | 0.120147 | 3.648736 | 0.00026353 | 0.019078063 |
| *LIMS2* | -1.485598 | 0.409391 | -3.628798 | 0.00028474 | 0.020412838 |
| *CD79A* | 0.422789 | 0.116521 | 3.628439 | 0.00028514 | 0.020412838 |
| *ACP5* | 0.681978 | 0.189349 | 3.601692 | 0.00031615 | 0.022507958 |
| *ITGAX* | 0.914734 | 0.254354 | 3.596311 | 0.00032276 | 0.022852275 |
| *DPEP3* | 2.249946 | 0.626805 | 3.589548 | 0.00033125 | 0.023325174 |
| *CCNDBP1* | 0.258515 | 0.072366 | 3.572307 | 0.00035385 | 0.024781032 |
| *A4GALT* | 1.174359 | 0.32963 | 3.562664 | 0.00036711 | 0.025570776 |
| *BHLHE40* | 0.506076 | 0.142226 | 3.558258 | 0.00037332 | 0.025863619 |
| *LARGE2* | -0.787548 | 0.221906 | -3.549007 | 0.00038669 | 0.026535056 |

| | | | | | |
|---|---|---|---|---|---|
| *CHCHD10* | 1.594884 | 0.449579 | 3.547505 | 0.0003889 | 0.026535056 |
| *MKNK2* | -0.360593 | 0.101653 | -3.547306 | 0.00038919 | 0.026535056 |
| *NDUFB9* | 0.225383 | 0.063648 | 3.541055 | 0.00039853 | 0.027028754 |
| *MMP11* | 1.001768 | 0.283135 | 3.53813 | 0.00040297 | 0.027186872 |
| *ZNF563* | 0.71897 | 0.203914 | 3.525858 | 0.00042211 | 0.028330001 |
| *FCHO1* | 0.480628 | 0.136486 | 3.521454 | 0.00042919 | 0.028655439 |
| *CD83* | -0.470595 | 0.133879 | -3.515065 | 0.00043965 | 0.029202452 |
| *FXYD1* | -0.800011 | 0.228976 | -3.493859 | 0.00047609 | 0.031409338 |
| *CORO1B* | -0.536899 | 0.15371 | -3.492933 | 0.00047775 | 0.031409338 |
| *DBNL* | 0.407287 | 0.116774 | 3.487821 | 0.00048697 | 0.031853502 |
| *ICAM1* | 0.964852 | 0.277072 | 3.482314 | 0.0004971 | 0.032351682 |
| *PWP1* | 0.343248 | 0.099396 | 3.453332 | 0.00055371 | 0.035854576 |
| *PARP15* | -0.597307 | 0.173215 | -3.448355 | 0.00056401 | 0.036339248 |
| *TMEM65* | 0.777975 | 0.226343 | 3.437155 | 0.00058786 | 0.037687283 |
| *TAF4B* | 1.100883 | 0.320708 | 3.432664 | 0.00059768 | 0.038127456 |
| *CHP1* | -0.383286 | 0.111858 | -3.426532 | 0.00061134 | 0.038806599 |
| *ARL4C* | -0.950825 | 0.278214 | -3.417606 | 0.00063175 | 0.039905227 |
| *ATG2A* | 0.599927 | 0.176231 | 3.404211 | 0.00066355 | 0.041710055 |
| *CASP8AP2* | 0.489981 | 0.144116 | 3.399917 | 0.00067406 | 0.042164964 |
| *RAB13* | -0.946474 | 0.278855 | -3.394144 | 0.00068844 | 0.042855965 |
| *NCF1* | 0.404405 | 0.119447 | 3.38565 | 0.0007101 | 0.043991919 |
| *SYNGAP1* | 0.940269 | 0.278562 | 3.375442 | 0.00073697 | 0.045438338 |
| *ITPRIPL2* | 0.91283 | 0.27087 | 3.369997 | 0.00075169 | 0.046125147 |
| *HSPA4* | -0.396909 | 0.118926 | -3.337452 | 0.0008455 | 0.051635762 |
| *EGR2* | -1.627897 | 0.489528 | -3.32544 | 0.00088279 | 0.053658797 |
| *PPFIBP2* | 0.544933 | 0.164033 | 3.322084 | 0.00089348 | 0.054053405 |
| *ZBED2* | -1.151471 | 0.34723 | -3.316163 | 0.00091262 | 0.054953626 |
| *FAM117A* | 0.531499 | 0.160568 | 3.310113 | 0.00093258 | 0.055894198 |
| *ENSG00000214797* | 2.339041 | 0.70808 | 3.303358 | 0.00095534 | 0.056993366 |
| *DTX4* | 0.934969 | 0.283601 | 3.29677 | 0.00097804 | 0.058078237 |
| *GALK1* | -0.494748 | 0.150178 | -3.294409 | 0.00098629 | 0.058299642 |
| *TMEM41B* | 0.420241 | 0.12787 | 3.286473 | 0.00101451 | 0.059693687 |
| *LAT2* | -0.358 | 0.109306 | -3.275215 | 0.00105582 | 0.061842015 |
| *SP110* | 0.247899 | 0.07581 | 3.270003 | 0.00107546 | 0.062707712 |
| *ST6GAL1* | -0.400251 | 0.12303 | -3.253277 | 0.00114082 | 0.065523983 |
| *NBN* | 0.460487 | 0.141615 | 3.251689 | 0.00114722 | 0.065523983 |
| *SEC62* | -0.17989 | 0.055326 | -3.25146 | 0.00114814 | 0.065523983 |
| *ICAM2* | 0.6064 | 0.186534 | 3.250878 | 0.00115049 | 0.065523983 |
| *ENSG00000203739* | 0.849724 | 0.261392 | 3.250771 | 0.00115093 | 0.065523983 |
| *VNN2* | 0.44994 | 0.138469 | 3.249389 | 0.00115653 | 0.065523983 |
| *IGFBP4* | -1.306893 | 0.402356 | -3.248105 | 0.00116176 | 0.065523983 |
| *INPP5B* | 0.558395 | 0.171949 | 3.24745 | 0.00116444 | 0.065523983 |
| *CYTIP* | -0.235562 | 0.072673 | -3.241405 | 0.00118942 | 0.066638488 |
| *DNAH1* | 1.035113 | 0.320013 | 3.234595 | 0.00121815 | 0.067952909 |
| *FKBP2* | 0.24485 | 0.075845 | 3.228305 | 0.00124526 | 0.069063884 |
| *FXYD2* | -1.141267 | 0.353607 | -3.227496 | 0.00124879 | 0.069063884 |
| *PGPEP1* | 0.700368 | 0.218481 | 3.205623 | 0.00134771 | 0.074215991 |

| | | | | | |
|---|---|---|---|---|---|
| *SLC9A3R1* | 0.462086 | 0.144251 | 3.203344 | 0.00135842 | 0.074411775 |
| *ZNF821* | 0.533151 | 0.166484 | 3.202414 | 0.00136281 | 0.074411775 |
| *GCHFR* | -0.326468 | 0.102037 | -3.199499 | 0.00137667 | 0.074851251 |
| *IL2RA* | -0.880955 | 0.275918 | -3.192818 | 0.00140892 | 0.075967184 |
| *ECHS1* | 0.298873 | 0.093608 | 3.192804 | 0.00140898 | 0.075967184 |
| *GBP7* | 1.239939 | 0.388635 | 3.190493 | 0.0014203 | 0.076258329 |
| *FOXO3* | 0.654952 | 0.205573 | 3.185989 | 0.0014426 | 0.07713412 |
| *EMC6* | -0.396305 | 0.124546 | -3.181994 | 0.00146265 | 0.077882929 |
| *NEURL2* | 1.477558 | 0.467017 | 3.163822 | 0.00155712 | 0.082509962 |
| *EXOSC5* | 0.442381 | 0.139868 | 3.162846 | 0.00156235 | 0.082509962 |
| *DTWD2* | -1.190787 | 0.377199 | -3.15692 | 0.00159445 | 0.083861578 |
| *ENSG00000266088* | -2.900208 | 0.919269 | -3.154909 | 0.00160548 | 0.084098608 |
| *ENSG00000166927* | -0.878213 | 0.278612 | -3.152101 | 0.001621 | 0.084567713 |
| *ENSG00000259436* | -0.719899 | 0.228916 | -3.144811 | 0.00166194 | 0.086353955 |
| *ENSG00000230709* | -0.731747 | 0.232793 | -3.143338 | 0.00167033 | 0.086441271 |
| *MFSD10* | -0.375896 | 0.119803 | -3.137606 | 0.00170334 | 0.087640481 |
| *ING1* | 0.460153 | 0.146688 | 3.136958 | 0.00170711 | 0.087640481 |
| *LSS* | -0.66197 | 0.211256 | -3.133493 | 0.00172739 | 0.088329938 |
| *ENSG00000235609* | 2.322698 | 0.741598 | 3.132019 | 0.00173609 | 0.088423817 |
| *PIM2* | -0.289045 | 0.092343 | -3.130107 | 0.00174743 | 0.088650924 |
| *RUNX3* | 0.34099 | 0.109205 | 3.122473 | 0.00179338 | 0.090625613 |
| *TGM2* | 0.64626 | 0.207289 | 3.117677 | 0.00182283 | 0.091522195 |
| *NFKBIA* | 0.204358 | 0.065557 | 3.117272 | 0.00182533 | 0.091522195 |
| *LPAR2* | 0.839257 | 0.269423 | 3.115016 | 0.00183935 | 0.091867679 |
| *NXT1* | 0.36692 | 0.118118 | 3.106382 | 0.00189392 | 0.094227955 |
| *CIB1* | 0.415448 | 0.133887 | 3.102982 | 0.00191581 | 0.094950511 |
| *UBTF* | 0.41875 | 0.135405 | 3.092582 | 0.00198423 | 0.097964916 |
| *ADCY7* | -0.545548 | 0.176874 | -3.084398 | 0.00203965 | 0.100316466 |
| *EMP3* | -0.238975 | 0.077527 | -3.082477 | 0.00205286 | 0.10058208 |
| *CD99* | -0.316898 | 0.103608 | -3.058611 | 0.00222366 | 0.108048827 |
| *MDK* | 1.396077 | 0.456521 | 3.058077 | 0.00222762 | 0.108048827 |
| *MYADM* | 0.650036 | 0.212589 | 3.057703 | 0.0022304 | 0.108048827 |
| *RHOQ* | 0.502833 | 0.164586 | 3.055147 | 0.00224951 | 0.108566015 |
| *PDIA3* | 0.293888 | 0.096251 | 3.053365 | 0.00226291 | 0.108805221 |
| *SERPINB9* | -0.30481 | 0.099975 | -3.04886 | 0.00229711 | 0.109675769 |
| *SIPA1L1* | -0.318792 | 0.104565 | -3.04874 | 0.00229803 | 0.109675769 |
| *EHD1* | 0.352212 | 0.115968 | 3.037153 | 0.00238825 | 0.113560594 |
| *SNX18* | 0.810992 | 0.267365 | 3.033281 | 0.00241911 | 0.114605128 |
| *ENSG00000089127* | 0.769627 | 0.254597 | 3.022918 | 0.0025035 | 0.118168778 |
| *ATP2B1* | -0.195606 | 0.064741 | -3.021376 | 0.00251628 | 0.118338784 |
| *CARNS1* | -1.404742 | 0.466009 | -3.014407 | 0.00257482 | 0.120651381 |
| *KANSL3* | 0.724137 | 0.24089 | 3.006089 | 0.00264632 | 0.123552336 |
| *RASGEF1B* | -0.445257 | 0.14845 | -2.999375 | 0.00270534 | 0.125801954 |
| *UROS* | 0.342305 | 0.114163 | 2.998398 | 0.00271403 | 0.125801954 |
| *CHRAC1* | -0.338273 | 0.112909 | -2.995979 | 0.00273565 | 0.126349857 |
| *LSM10* | -0.210074 | 0.070332 | -2.986896 | 0.00281826 | 0.12970013 |
| *IRF4* | 0.650003 | 0.217882 | 2.983284 | 0.00285173 | 0.130773676 |

| | | | | | |
|---|---|---|---|---|---|
| *PRDM4* | 0.511776 | 0.171979 | 2.975811 | 0.00292215 | 0.13352763 |
| *APOL1* | 0.631416 | 0.21269 | 2.968717 | 0.00299046 | 0.136088133 |
| *LINC00513* | 0.479824 | 0.161676 | 2.967809 | 0.0029993 | 0.136088133 |
| *GPER1* | -1.584968 | 0.53507 | -2.96217 | 0.00305479 | 0.138119395 |
| *FLOT1* | 0.387399 | 0.130927 | 2.958891 | 0.00308748 | 0.138980184 |
| *CASP7* | 0.521853 | 0.176415 | 2.958102 | 0.0030954 | 0.138980184 |
| *PLEKHO2* | 0.526477 | 0.178767 | 2.945049 | 0.00322903 | 0.144476799 |
| *SGMS1.AS1* | 0.705697 | 0.240139 | 2.938701 | 0.00329591 | 0.146927846 |
| *RESF1* | -0.246297 | 0.08384 | -2.937695 | 0.00330662 | 0.146927846 |
| *RAB8B* | -0.383351 | 0.130628 | -2.934672 | 0.00333901 | 0.147857241 |
| *RBX1* | -0.162737 | 0.055483 | -2.933098 | 0.00335598 | 0.1480998 |
| *DDX24* | 0.284961 | 0.09727 | 2.929577 | 0.00339424 | 0.149276911 |
| *RUFY3* | 0.545002 | 0.186226 | 2.92656 | 0.00342733 | 0.150219578 |
| *PRDX1* | -0.250814 | 0.085869 | -2.920876 | 0.00349049 | 0.152469365 |
| *MAP2K3* | -0.29399 | 0.100727 | -2.918678 | 0.00351519 | 0.153029601 |
| *MPHOSPH9* | 0.546785 | 0.187536 | 2.915631 | 0.0035497 | 0.154011511 |
| *DAAM1* | -0.538234 | 0.18479 | -2.912685 | 0.00358336 | 0.154950423 |
| *CDYL2* | 1.831296 | 0.629965 | 2.906982 | 0.00364934 | 0.15690365 |
| *SH3BGRL3* | -0.239615 | 0.082436 | -2.906678 | 0.00365289 | 0.15690365 |
| *FADS3* | -0.437285 | 0.150561 | -2.904375 | 0.00367987 | 0.157537687 |
| *TRAF3IP2* | -0.718381 | 0.247913 | -2.897714 | 0.00375893 | 0.160389322 |
| *GTF2B* | 0.391244 | 0.135395 | 2.889657 | 0.00385663 | 0.16401477 |
| *GPR132* | 0.402885 | 0.139729 | 2.883331 | 0.00393494 | 0.166794749 |
| *PAOX* | 0.560575 | 0.195057 | 2.873905 | 0.0040543 | 0.171291034 |
| *ENSG00000263264* | 0.763866 | 0.265953 | 2.872184 | 0.00407646 | 0.171664032 |
| *C7orf50* | -0.429702 | 0.149933 | -2.865963 | 0.00415743 | 0.174503728 |
| *HSH2D* | 0.310357 | 0.10838 | 2.863602 | 0.00418853 | 0.174705765 |
| *MIR22HG* | -0.647691 | 0.226185 | -2.86354 | 0.00418936 | 0.174705765 |
| *SMIM20* | 0.408352 | 0.142703 | 2.861546 | 0.00421581 | 0.175241491 |
| *GGA2* | 0.329312 | 0.115195 | 2.858736 | 0.00425333 | 0.176232781 |
| *AHCY* | -0.32637 | 0.114806 | -2.842796 | 0.00447197 | 0.184698132 |
| *RAB3IP* | -0.524639 | 0.184668 | -2.840983 | 0.00449748 | 0.185158073 |
| *HCK* | -0.542546 | 0.191215 | -2.837366 | 0.00454874 | 0.186672164 |
| *PTPA* | -0.659631 | 0.233073 | -2.830151 | 0.00465261 | 0.190029857 |
| *LINC02576* | -1.806206 | 0.638426 | -2.829153 | 0.00466714 | 0.190029857 |
| *SLC25A39* | 0.222704 | 0.078732 | 2.828628 | 0.0046748 | 0.190029857 |
| *BATF2* | 2.352515 | 0.833937 | 2.820976 | 0.00478778 | 0.194010444 |
| *PACS1* | 0.341314 | 0.121096 | 2.818546 | 0.00482417 | 0.194872197 |
| *BANK1* | -0.224634 | 0.079954 | -2.809551 | 0.00496107 | 0.199776221 |
| *ARID5B* | -0.28674 | 0.102286 | -2.803322 | 0.00505792 | 0.201949322 |
| *LARGE1* | 1.030888 | 0.36774 | 2.803303 | 0.00505821 | 0.201949322 |
| *RAB7A* | 0.209617 | 0.074813 | 2.801868 | 0.00508077 | 0.201949322 |
| *PCBP1* | -0.186822 | 0.06668 | -2.801791 | 0.00508197 | 0.201949322 |
| *MYO9B* | 0.303576 | 0.108379 | 2.801067 | 0.0050934 | 0.201949322 |
| *DUSP10* | -0.512705 | 0.183164 | -2.799164 | 0.00512351 | 0.202519898 |
| *ENSG00000261448* | 0.802082 | 0.286698 | 2.797658 | 0.00514747 | 0.202844771 |
| *ENSG00000176320* | 2.292757 | 0.821232 | 2.791851 | 0.00524074 | 0.205702427 |

| | | | | | |
|---|---|---|---|---|---|
| *STAT5A* | 0.417966 | 0.149746 | 2.791163 | 0.00525191 | 0.205702427 |
| *ITGB7* | 0.4283 | 0.153968 | 2.78175 | 0.00540667 | 0.2111222 |
| *CYBA* | 0.224601 | 0.080874 | 2.777185 | 0.00548319 | 0.213463477 |
| *HRK* | -0.588766 | 0.212628 | -2.768994 | 0.00562297 | 0.217800377 |
| *TMCO3* | -0.408087 | 0.147407 | -2.768439 | 0.00563255 | 0.217800377 |
| *GPR18* | 0.599111 | 0.216465 | 2.767702 | 0.0056453 | 0.217800377 |
| *SMYD4* | -0.474857 | 0.171994 | -2.760899 | 0.00576424 | 0.221310625 |
| *CHD7* | 0.313425 | 0.113538 | 2.760538 | 0.00577063 | 0.221310625 |
| *MYO7B* | -1.804508 | 0.654451 | -2.757284 | 0.00582837 | 0.222663067 |
| *RHBDF2* | 0.249735 | 0.090595 | 2.756607 | 0.00584046 | 0.222663067 |
| *KATNBL1* | -0.298804 | 0.108539 | -2.752957 | 0.00590597 | 0.223989853 |
| *PIK3IP1* | 0.33875 | 0.12306 | 2.752732 | 0.00591002 | 0.223989853 |
| *ENSG00000262292* | 0.765643 | 0.278632 | 2.747865 | 0.00599847 | 0.226600494 |
| *ADPGK* | 0.255649 | 0.093064 | 2.747013 | 0.00601408 | 0.226600494 |
| *TLR10* | -0.384397 | 0.140419 | -2.737507 | 0.00619069 | 0.231959891 |
| *CD47* | -0.215769 | 0.078822 | -2.73742 | 0.00619232 | 0.231959891 |
| *ZNF736* | -0.764694 | 0.279766 | -2.733338 | 0.00626959 | 0.233848529 |
| *CDK14* | -0.59391 | 0.217323 | -2.732843 | 0.00627903 | 0.233848529 |
| *SMC5* | -0.299908 | 0.109901 | -2.728902 | 0.00635457 | 0.235979653 |
| *ZNF318* | -0.508355 | 0.186901 | -2.719925 | 0.00652968 | 0.241785773 |
| *HDAC6* | -0.458905 | 0.168808 | -2.718493 | 0.006558 | 0.242138751 |
| *MYO1C* | -0.425446 | 0.156626 | -2.716317 | 0.00660127 | 0.242949273 |
| *GPR155* | -0.609086 | 0.2243 | -2.715496 | 0.00661766 | 0.242949273 |
| *ENSG00000261087* | 0.948387 | 0.350588 | 2.705133 | 0.0068277 | 0.249948215 |
| *CHIT1* | -1.344916 | 0.497541 | -2.703123 | 0.00686912 | 0.250752185 |
| *SGK1* | -0.742648 | 0.27497 | -2.700831 | 0.00691664 | 0.251773444 |
| *CD82* | -0.254069 | 0.094245 | -2.695833 | 0.00702129 | 0.254862925 |
| *PXDC1* | 1.092524 | 0.406294 | 2.689001 | 0.00716661 | 0.259407225 |
| *CFL1* | -0.13263 | 0.049347 | -2.687685 | 0.00719492 | 0.259702338 |
| *SLC2A3* | -0.43105 | 0.16046 | -2.686339 | 0.00722399 | 0.260023092 |
| *RUBCN* | -0.414983 | 0.154642 | -2.683515 | 0.00728527 | 0.261498562 |
| *NAB2* | -0.740443 | 0.2767 | -2.675977 | 0.00745117 | 0.266104529 |
| *PPP3CA* | -0.228362 | 0.085349 | -2.675629 | 0.00745891 | 0.266104529 |
| *SF1* | 0.149492 | 0.055899 | 2.674334 | 0.00748778 | 0.266104529 |
| *DUOX1* | 0.475809 | 0.177942 | 2.673958 | 0.00749619 | 0.266104529 |
| *SLC45A3* | 0.984882 | 0.369036 | 2.668797 | 0.00761234 | 0.268549295 |
| *AP1S3* | -0.405669 | 0.152024 | -2.668449 | 0.00762023 | 0.268549295 |
| *CREM* | -0.537844 | 0.201581 | -2.668125 | 0.00762758 | 0.268549295 |
| *HINT2* | 0.263035 | 0.098847 | 2.661041 | 0.00778994 | 0.273518037 |
| *RAP1A* | -0.180722 | 0.068021 | -2.656852 | 0.00788742 | 0.276068646 |
| *SFXN4* | 0.669188 | 0.251946 | 2.656083 | 0.00790543 | 0.276068646 |
| *POU2AF1* | 0.301961 | 0.113728 | 2.655112 | 0.00792821 | 0.276116056 |
| *SERPINB1* | 0.279179 | 0.105252 | 2.652489 | 0.00799008 | 0.277520629 |
| *ADGRB2* | -2.420612 | 0.913445 | -2.649981 | 0.00804964 | 0.278837856 |
| *RAB30.DT* | -0.323709 | 0.12226 | -2.647697 | 0.00810422 | 0.279915777 |
| *CLIC1* | -0.177311 | 0.066989 | -2.646864 | 0.00812421 | 0.279915777 |
| *NRM* | 0.445585 | 0.168455 | 2.645131 | 0.00816593 | 0.280388388 |

| | | | | | |
|---|---|---|---|---|---|
| *SLC38A2* | -0.267944 | 0.101347 | -2.643823 | 0.00819755 | 0.280388388 |
| *ABLIM2* | -0.868425 | 0.328502 | -2.64359 | 0.0082032 | 0.280388388 |
| *HLA.DOA* | 0.434147 | 0.164402 | 2.640769 | 0.00827182 | 0.281985794 |
| *FYTTD1* | -0.229564 | 0.087014 | -2.638247 | 0.00833358 | 0.282779928 |
| *ANXA2* | -0.340888 | 0.129221 | -2.638027 | 0.008339 | 0.282779928 |
| *TKT* | -0.189066 | 0.071711 | -2.636498 | 0.00837666 | 0.283311324 |
| *ENSG00000261766* | 0.603576 | 0.229409 | 2.631006 | 0.00851326 | 0.287177587 |
| *PSME1* | 0.139398 | 0.053025 | 2.628927 | 0.00856546 | 0.287836121 |
| *ENSG00000273188* | 0.997499 | 0.3795 | 2.628452 | 0.00857745 | 0.287836121 |
| *ITGAM* | 0.654662 | 0.249555 | 2.623315 | 0.00870787 | 0.291453623 |
| *MYOM1* | -1.241343 | 0.47384 | -2.61975 | 0.00879943 | 0.293755116 |
| *LGMN* | 0.733988 | 0.280316 | 2.618435 | 0.00883342 | 0.294114279 |
| *REL* | -0.223481 | 0.085406 | -2.616685 | 0.00887881 | 0.294114279 |
| *EHMT1* | -0.213652 | 0.081665 | -2.61621 | 0.00889119 | 0.294114279 |
| *MYC* | 0.398696 | 0.15244 | 2.615434 | 0.00891142 | 0.294114279 |
| *FXYD5* | -0.330434 | 0.126364 | -2.61494 | 0.00892431 | 0.294114279 |
| *XIAP* | 0.303122 | 0.116017 | 2.612732 | 0.00898218 | 0.295266322 |
| *ARHGEF2* | 0.263398 | 0.100912 | 2.610175 | 0.00904959 | 0.296725161 |
| *MYL2* | 1.96892 | 0.754915 | 2.608136 | 0.00910369 | 0.297741439 |
| *MLXIP* | 0.479515 | 0.183923 | 2.607154 | 0.00912982 | 0.297840253 |
| *BMF* | -0.3873 | 0.148628 | -2.605829 | 0.00916522 | 0.298123304 |
| *MARCKSL1* | -0.339877 | 0.130466 | -2.605099 | 0.00918477 | 0.298123304 |
| *AKR1A1* | -0.219132 | 0.084166 | -2.603556 | 0.00922621 | 0.298715868 |
| *PIN1* | -0.200582 | 0.077191 | -2.598527 | 0.00936247 | 0.302367841 |
| *BRK1* | 0.150187 | 0.057819 | 2.597534 | 0.00938958 | 0.302485282 |
| *SMC6* | -0.293346 | 0.113101 | -2.593654 | 0.0094962 | 0.305157131 |
| *AIM2* | 0.324903 | 0.125414 | 2.590645 | 0.00957964 | 0.305638878 |
| *METTL21A* | -0.266265 | 0.10278 | -2.590636 | 0.00957989 | 0.305638878 |
| *ALDH1B1* | 0.904353 | 0.349097 | 2.590547 | 0.00958235 | 0.305638878 |
| *FCER2* | 0.947455 | 0.365956 | 2.588987 | 0.00962588 | 0.306179444 |
| *ENSG00000261884* | 0.459909 | 0.177692 | 2.588239 | 0.00964682 | 0.306179444 |
| *COMMD7* | -0.25852 | 0.099929 | -2.58704 | 0.00968042 | 0.306491191 |
| *TUBB6* | -0.687056 | 0.265715 | -2.585686 | 0.00971854 | 0.306943855 |
| *LINC01869* | -0.714285 | 0.276629 | -2.582102 | 0.00982006 | 0.30894585 |
| *PIK3R5* | 0.489321 | 0.18953 | 2.581757 | 0.00982988 | 0.30894585 |
| *EHD4* | -0.477297 | 0.185136 | -2.578096 | 0.00993464 | 0.311385737 |
| *RALA* | -0.396995 | 0.154032 | -2.577359 | 0.00995584 | 0.311385737 |
| *PDCD4* | 0.638841 | 0.248044 | 2.575511 | 0.0100092 | 0.312296865 |
| *SYAP1* | -0.410878 | 0.159745 | -2.572081 | 0.01010893 | 0.314646676 |
| *CTSS* | 0.210522 | 0.081903 | 2.570393 | 0.01015832 | 0.315422065 |
| *FXYD7* | -0.855358 | 0.333348 | -2.565962 | 0.01028901 | 0.318032378 |
| *GSAP* | 0.352199 | 0.137263 | 2.56587 | 0.01029175 | 0.318032378 |
| *PBX4* | 0.936722 | 0.365938 | 2.559783 | 0.01047375 | 0.322882171 |
| *POLM* | 0.375847 | 0.147108 | 2.554896 | 0.01062194 | 0.326668954 |
| *KAT2A* | 0.331817 | 0.129994 | 2.552547 | 0.01069386 | 0.328097647 |
| *AK1* | 0.562186 | 0.221203 | 2.541492 | 0.01103806 | 0.337853654 |
| *TRBC2* | -0.170151 | 0.067011 | -2.539143 | 0.01111244 | 0.33904116 |

| | | | | | |
|---|---|---|---|---|---|
| *C19orf38* | 0.625296 | 0.246325 | 2.538499 | 0.01113293 | 0.33904116 |
| *NMT2* | -0.472337 | 0.186122 | -2.537781 | 0.01115579 | 0.33904116 |
| *TBC1D8* | 1.391661 | 0.548571 | 2.536885 | 0.01118436 | 0.339109685 |
| *TMSB10* | -0.244093 | 0.096379 | -2.532641 | 0.01132067 | 0.342437063 |
| *TMEM87B* | -0.499138 | 0.197258 | -2.530377 | 0.01139399 | 0.343847624 |
| *NBEAL2* | 1.021037 | 0.403691 | 2.529253 | 0.01143058 | 0.344145799 |
| *ANKRD37* | -0.560568 | 0.221956 | -2.525575 | 0.01155091 | 0.346958151 |
| *IFNAR2* | 0.32295 | 0.128113 | 2.520823 | 0.01170807 | 0.350860756 |
| *RBMX2* | -0.2888 | 0.11464 | -2.519181 | 0.0117628 | 0.351683205 |
| *SPN* | 0.979166 | 0.389017 | 2.517024 | 0.01183508 | 0.352825411 |
| *SUCLG2* | 0.278775 | 0.110783 | 2.516409 | 0.01185577 | 0.352825411 |
| *NTAN1* | -0.274303 | 0.109462 | -2.505916 | 0.01221347 | 0.362633163 |
| *GAK* | 0.272129 | 0.108774 | 2.501769 | 0.01235747 | 0.365262825 |
| *APOBEC3H* | 1.279193 | 0.511389 | 2.501409 | 0.01237001 | 0.365262825 |
| *ENSG00000272155* | 1.129277 | 0.451545 | 2.500921 | 0.01238708 | 0.365262825 |
| *CSNK2A2* | -0.450696 | 0.180362 | -2.498845 | 0.01245989 | 0.366571111 |
| *SMARCA4* | 0.34137 | 0.136717 | 2.496909 | 0.0125281 | 0.367108152 |
| *WASF1* | -0.64925 | 0.260123 | -2.495934 | 0.01256258 | 0.367108152 |
| *SRR* | 0.814002 | 0.326135 | 2.495905 | 0.01256361 | 0.367108152 |
| *CARD19* | -0.386753 | 0.155079 | -2.493902 | 0.01263474 | 0.368351166 |
| *ESYT1* | -0.34753 | 0.139409 | -2.492891 | 0.01267076 | 0.36856751 |
| *VTI1A* | -0.36552 | 0.147019 | -2.486199 | 0.01291158 | 0.374726654 |
| *IQSEC1* | -0.336108 | 0.135367 | -2.48293 | 0.01303067 | 0.37733301 |
| *SYNGR3* | 0.812296 | 0.327298 | 2.481821 | 0.01307129 | 0.377660581 |
| *USP7* | 0.317171 | 0.127867 | 2.480473 | 0.0131208 | 0.378243109 |
| *OSM* | -1.713731 | 0.69155 | -2.478102 | 0.01320834 | 0.37991667 |
| *ENSG00000262420* | 1.605898 | 0.648981 | 2.474492 | 0.01334256 | 0.382414876 |
| *PLGRKT* | -0.432468 | 0.174847 | -2.473408 | 0.01338311 | 0.382414876 |
| *JDP2* | 0.567481 | 0.229436 | 2.473379 | 0.01338422 | 0.382414876 |
| *TERF2IP* | 0.208095 | 0.084372 | 2.4664 | 0.01364787 | 0.38908502 |
| *ARRDC5* | 0.689183 | 0.279971 | 2.461621 | 0.01383109 | 0.392599222 |
| *IQGAP1* | -0.198456 | 0.080621 | -2.461595 | 0.01383207 | 0.392599222 |
| *ENSG00000249679* | -1.375267 | 0.559305 | -2.458887 | 0.01393686 | 0.394704184 |
| *A1BG* | -0.264237 | 0.10751 | -2.45779 | 0.01397949 | 0.395043082 |
| *PRRT3* | 0.657785 | 0.267888 | 2.455452 | 0.01407074 | 0.396751658 |
| *EIF4A3* | 0.367442 | 0.149696 | 2.454583 | 0.01410482 | 0.396793972 |
| *ENSG00000229127* | -1.202067 | 0.489871 | -2.453844 | 0.01413382 | 0.396793972 |
| *PISD* | 0.58726 | 0.239718 | 2.4498 | 0.01429354 | 0.400116785 |
| *UNKL* | 0.342648 | 0.139897 | 2.449278 | 0.01431428 | 0.400116785 |
| *RHOBTB3* | 1.450464 | 0.592822 | 2.44671 | 0.01441668 | 0.40210673 |
| *SLC25A5* | -0.17857 | 0.073034 | -2.445016 | 0.01448456 | 0.402277281 |
| *LY9* | -0.465548 | 0.190408 | -2.445 | 0.01448523 | 0.402277281 |
| *POLH* | 0.747853 | 0.30623 | 2.442132 | 0.01460081 | 0.404615238 |
| *FAM177A1* | -0.238275 | 0.097618 | -2.440885 | 0.01465133 | 0.405110131 |
| *PLD4* | -0.569548 | 0.233408 | -2.440141 | 0.01468155 | 0.405110131 |
| *BCL2L11* | -0.637324 | 0.261447 | -2.437682 | 0.01478178 | 0.406563864 |
| *SNRNP35* | 0.303067 | 0.124353 | 2.437143 | 0.01480381 | 0.406563864 |

| | | | | | |
|---|---|---|---|---|---|
| *WDR11* | 0.288873 | 0.118559 | 2.436531 | 0.01482889 | 0.406563864 |
| *RB1* | -0.253183 | 0.104114 | -2.431792 | 0.01502431 | 0.411047321 |
| *FBXO10* | 0.638157 | 0.26275 | 2.428762 | 0.01515048 | 0.412881907 |
| *ASCC1* | 0.425653 | 0.175264 | 2.428643 | 0.01515545 | 0.412881907 |
| *SSR3* | -0.193731 | 0.079859 | -2.425898 | 0.01527054 | 0.415139738 |
| *PLEC* | -0.357721 | 0.147537 | -2.424614 | 0.01532466 | 0.415306227 |
| *ATP6V0A1* | 0.434039 | 0.179043 | 2.424224 | 0.01534113 | 0.415306227 |
| *DNAAF4* | -0.92159 | 0.380844 | -2.419863 | 0.01552636 | 0.419439425 |
| *AFF3* | -0.254875 | 0.10541 | -2.417943 | 0.0156085 | 0.420776438 |
| *AP3B1* | 0.290696 | 0.120295 | 2.416527 | 0.01566938 | 0.421133702 |
| *MBD6* | 0.600578 | 0.248604 | 2.415796 | 0.01570087 | 0.421133702 |
| *ENSG00000133961* | 0.296196 | 0.12263 | 2.415357 | 0.0157198 | 0.421133702 |
| *AP1G2* | -0.24231 | 0.100431 | -2.412697 | 0.01583499 | 0.422913437 |
| *MZB1* | -0.545124 | 0.225976 | -2.412308 | 0.01585187 | 0.422913437 |
| *DMPK* | -1.03311 | 0.428755 | -2.409558 | 0.01597188 | 0.424674681 |
| *PLPP5* | -0.34149 | 0.141739 | -2.409285 | 0.0159838 | 0.424674681 |
| *GHRL* | 0.42316 | 0.175773 | 2.407429 | 0.0160653 | 0.425961786 |
| *UPP1* | 0.669754 | 0.278306 | 2.406537 | 0.01610459 | 0.426126827 |
| *SLC25A25* | -0.551863 | 0.229457 | -2.405082 | 0.01616882 | 0.42694966 |
| *TTC13* | -0.747484 | 0.3109 | -2.404261 | 0.0162052 | 0.4270353 |
| *H1FX* | 0.726421 | 0.30232 | 2.402822 | 0.0162691 | 0.427844225 |
| *GABARAPL1* | 0.955856 | 0.397983 | 2.401753 | 0.01631671 | 0.428222191 |
| *CUL2* | 0.442529 | 0.184347 | 2.400517 | 0.01637193 | 0.428798011 |
| *RPS6KC1* | -0.551725 | 0.230063 | -2.398151 | 0.01647808 | 0.430299079 |
| *ISG20* | -0.234119 | 0.097641 | -2.397752 | 0.01649602 | 0.430299079 |
| *LSM2* | -0.169237 | 0.070632 | -2.396042 | 0.01657318 | 0.431438302 |
| *CAMTA1* | -0.247776 | 0.103498 | -2.394006 | 0.01666547 | 0.432966118 |
| *GAPDH* | -0.166369 | 0.069523 | -2.393018 | 0.0167104 | 0.433259991 |
| *PARD6A* | -0.343746 | 0.143743 | -2.391392 | 0.01678461 | 0.43411166 |
| *ARAP1* | -0.438498 | 0.183409 | -2.390824 | 0.01681063 | 0.43411166 |
| *ATP2C1* | -0.418733 | 0.175352 | -2.387954 | 0.01694247 | 0.43664128 |
| *MRNIP* | 0.390127 | 0.164011 | 2.378664 | 0.01737549 | 0.446788713 |
| *TMEM175* | 0.487856 | 0.205152 | 2.378027 | 0.01740555 | 0.446788713 |
| *CYBB* | 0.249476 | 0.104996 | 2.376041 | 0.01749951 | 0.447705914 |
| *USP18* | 1.066879 | 0.44906 | 2.375804 | 0.01751077 | 0.447705914 |
| *C1orf162* | -0.413777 | 0.174399 | -2.372582 | 0.01766422 | 0.450228587 |
| *FLNA* | -0.332372 | 0.140107 | -2.372267 | 0.01767932 | 0.450228587 |
| *NOTCH2* | 0.35843 | 0.151242 | 2.36991 | 0.01779242 | 0.452215162 |
| *LINC02397* | 0.190483 | 0.080456 | 2.367549 | 0.01790635 | 0.45414393 |
| *ITGA3* | 0.94543 | 0.399442 | 2.366879 | 0.01793879 | 0.45414393 |
| *POU2F2* | -0.2758 | 0.116704 | -2.363242 | 0.01811585 | 0.457727252 |
| *SESTD1* | -0.453593 | 0.192274 | -2.359101 | 0.01831928 | 0.461961314 |
| *ACTB* | -0.184134 | 0.078124 | -2.356953 | 0.01842559 | 0.463219487 |
| *GNAS* | 0.166345 | 0.070586 | 2.356641 | 0.01844107 | 0.463219487 |
| *ZNF71* | -0.446702 | 0.189619 | -2.355788 | 0.01848346 | 0.463380951 |
| *SLFN5* | -0.822224 | 0.349303 | -2.353903 | 0.01857746 | 0.464833169 |
| *TNF* | -0.834597 | 0.355 | -2.350977 | 0.01872417 | 0.467596184 |

| | | | | | |
|---|---|---|---|---|---|
| *CHST7* | 0.851677 | 0.362421 | 2.349965 | 0.01877518 | 0.467963074 |
| *H3F3A* | 0.166006 | 0.07078 | 2.345394 | 0.019007 | 0.472666187 |
| *USP15* | -0.210329 | 0.0897 | -2.344801 | 0.01903723 | 0.472666187 |
| *ENSG00000268516* | -0.383899 | 0.164074 | -2.33979 | 0.0192946 | 0.478134948 |
| *NPEPPS* | -0.258783 | 0.110661 | -2.338515 | 0.01936054 | 0.478318615 |
| *MED18* | 0.562273 | 0.24056 | 2.33735 | 0.01942097 | 0.478318615 |
| *CD79B* | 0.246878 | 0.105624 | 2.337332 | 0.01942191 | 0.478318615 |
| *RAB4B* | -0.287394 | 0.122993 | -2.336675 | 0.01945609 | 0.478318615 |
| *UBE2D4* | -0.398102 | 0.170474 | -2.335271 | 0.01952925 | 0.478318615 |
| *BMS1* | 0.320974 | 0.137486 | 2.334594 | 0.01956462 | 0.478318615 |
| *CREBRF* | 0.289654 | 0.12408 | 2.334419 | 0.01957381 | 0.478318615 |
| *DIPK1A* | 0.592258 | 0.253759 | 2.333938 | 0.01959896 | 0.478318615 |
| *DMD* | -0.538248 | 0.230887 | -2.331223 | 0.01974162 | 0.480889521 |
| *HOXB4* | 1.003484 | 0.431193 | 2.327226 | 0.01995322 | 0.485126692 |
| *PPP2R5C* | 0.211944 | 0.09121 | 2.323682 | 0.02014254 | 0.48727933 |
| *TBC1D10C* | -0.157738 | 0.067889 | -2.323475 | 0.02015366 | 0.48727933 |
| *S100A11* | -0.179892 | 0.077425 | -2.323446 | 0.0201552 | 0.48727933 |
| *RBM26* | 0.212685 | 0.091589 | 2.322179 | 0.02022332 | 0.488010556 |
| *HIPK2* | 0.495951 | 0.213811 | 2.319574 | 0.02036395 | 0.488784346 |
| *ANKRD36C* | -0.46125 | 0.198856 | -2.319511 | 0.02036735 | 0.488784346 |
| *PSMB9* | 0.195864 | 0.084448 | 2.319347 | 0.02037623 | 0.488784346 |
| *HNRNPC* | 0.16281 | 0.070214 | 2.318777 | 0.02040711 | 0.488784346 |
| *EVL* | 0.191877 | 0.082844 | 2.316121 | 0.02055164 | 0.491332969 |
| *CD37* | -0.204147 | 0.088187 | -2.314927 | 0.02061692 | 0.491980781 |
| *RBM19* | 0.470923 | 0.203591 | 2.313079 | 0.02071832 | 0.492983008 |
| *SCARB1* | -1.17448 | 0.507825 | -2.312767 | 0.02073543 | 0.492983008 |
| *ABCA2* | 0.965678 | 0.417805 | 2.311314 | 0.02081552 | 0.493264455 |
| *FAM107B* | 0.24349 | 0.105354 | 2.311163 | 0.02082383 | 0.493264455 |
| *SHISAL2A* | 0.358296 | 0.155183 | 2.308866 | 0.02095103 | 0.495366883 |
| *IVNS1ABP* | -0.367613 | 0.159324 | -2.307325 | 0.0210367 | 0.496037741 |
| *MAP4K1* | 0.218734 | 0.094814 | 2.306972 | 0.02105639 | 0.496037741 |
| *SH3BP5* | -0.414427 | 0.17973 | -2.305831 | 0.02112005 | 0.496629494 |
| *HGSNAT* | -0.5211 | 0.226223 | -2.303485 | 0.02125158 | 0.49881219 |
| *CALR* | 0.291645 | 0.126878 | 2.298629 | 0.02152604 | 0.503978282 |
| *GTF3A* | 0.161865 | 0.070431 | 2.298209 | 0.0215499 | 0.503978282 |
| *SACS* | -0.537661 | 0.234242 | -2.295324 | 0.02171456 | 0.506909173 |
| *ENSG00000251364* | 1.142158 | 0.498502 | 2.291182 | 0.02195288 | 0.511222508 |
| *MT1X* | -0.501389 | 0.218908 | -2.290409 | 0.02199762 | 0.511222508 |
| *MFGE8* | 0.764362 | 0.33381 | 2.289812 | 0.0220322 | 0.511222508 |
| *RBM47* | 0.584623 | 0.255393 | 2.289106 | 0.02207317 | 0.511222508 |
| *SS18L2* | -0.130855 | 0.057175 | -2.288684 | 0.0220977 | 0.511222508 |
| *BCL6* | -0.530113 | 0.231814 | -2.286799 | 0.02220758 | 0.512336884 |
| *C12orf74.PLEKHG7* | 0.515004 | 0.225277 | 2.286092 | 0.0222489 | 0.512336884 |
| *HTT* | 0.339849 | 0.148677 | 2.285814 | 0.02226515 | 0.512336884 |
| *KDM4B* | -0.401432 | 0.175678 | -2.285042 | 0.02231039 | 0.512462803 |
| *HP1BP3* | -0.138889 | 0.060821 | -2.283584 | 0.02239597 | 0.513513202 |
| *LUZP1* | -0.513851 | 0.225286 | -2.280884 | 0.02255532 | 0.515346467 |

| | | | | | |
|---|---|---|---|---|---|
| *NAP1L6* | 1.50205 | 0.658542 | 2.280874 | 0.02255591 | 0.515346467 |
| *TCEAL3* | -0.504185 | 0.221161 | -2.279718 | 0.02262442 | 0.515996898 |
| *ZNF496* | 0.773493 | 0.339586 | 2.277754 | 0.02274123 | 0.517744712 |
| *P2RX4* | 0.379534 | 0.16673 | 2.276343 | 0.02282552 | 0.517864211 |
| *SIDT2* | -0.320645 | 0.140861 | -2.27632 | 0.02282686 | 0.517864211 |
| *PTTG1* | -0.596723 | 0.26269 | -2.27159 | 0.02311127 | 0.52078436 |
| *CDC25B* | 0.310526 | 0.136714 | 2.27135 | 0.02312581 | 0.52078436 |
| *MCPH1* | 0.321158 | 0.141415 | 2.271031 | 0.02314512 | 0.52078436 |
| *SVBP* | -0.195813 | 0.086225 | -2.270942 | 0.02315047 | 0.52078436 |
| *ENSG00000230155* | 0.541316 | 0.238378 | 2.270824 | 0.02315765 | 0.52078436 |
| *JUND* | -0.394523 | 0.173793 | -2.270072 | 0.02320322 | 0.520900051 |
| *CIC* | -0.374637 | 0.165151 | -2.268447 | 0.02330201 | 0.52220808 |
| *ENSG00000260349* | 0.590548 | 0.260495 | 2.267022 | 0.02338889 | 0.523245205 |
| *MAST4* | -0.815612 | 0.360135 | -2.264738 | 0.02352874 | 0.52448022 |
| *HOXB7* | 0.669018 | 0.295428 | 2.26457 | 0.02353908 | 0.52448022 |
| *CNR1* | 0.608707 | 0.268848 | 2.264129 | 0.0235662 | 0.52448022 |
| *CARS2* | 0.40128 | 0.17742 | 2.261752 | 0.02371271 | 0.525279656 |
| *TCEA2* | 0.355204 | 0.15706 | 2.261577 | 0.02372358 | 0.525279656 |
| *TSEN15* | -0.37269 | 0.164793 | -2.261563 | 0.02372441 | 0.525279656 |
| *SYBU* | -1.177985 | 0.521159 | -2.260319 | 0.02380147 | 0.525389244 |
| *JAK1* | 0.146553 | 0.064859 | 2.259559 | 0.02384865 | 0.525389244 |
| *DCK* | 0.151574 | 0.067083 | 2.25951 | 0.02385168 | 0.525389244 |
| *OAS3* | 0.913701 | 0.405415 | 2.253741 | 0.02421245 | 0.530689823 |
| *OGG1* | 0.309392 | 0.137289 | 2.253585 | 0.0242223 | 0.530689823 |
| *C9orf72* | -0.4749 | 0.21074 | -2.25349 | 0.02422828 | 0.530689823 |
| *YIF1A* | -0.208551 | 0.092572 | -2.252838 | 0.02426936 | 0.530689823 |
| *LPCAT4* | -0.363824 | 0.161529 | -2.252381 | 0.02429823 | 0.530689823 |
| *CCDC154* | 0.850702 | 0.378265 | 2.248961 | 0.024515 | 0.533654712 |
| *ZNF627* | 0.476021 | 0.211712 | 2.248434 | 0.02454854 | 0.533654712 |
| *TXNDC11* | 0.347263 | 0.154457 | 2.248282 | 0.02455822 | 0.533654712 |
| *TRMT1* | 0.244642 | 0.108907 | 2.246341 | 0.02468215 | 0.535444677 |
| *CD84* | -0.451552 | 0.201086 | -2.245565 | 0.02473189 | 0.535622107 |
| *ZNF706* | 0.178516 | 0.079595 | 2.24281 | 0.02490908 | 0.537776675 |
| *MIPEP* | -0.812592 | 0.362324 | -2.242721 | 0.02491484 | 0.537776675 |
| *MIIP* | -0.221393 | 0.098784 | -2.241188 | 0.0250139 | 0.538622077 |
| *CYB561A3* | -0.321543 | 0.143493 | -2.240822 | 0.02503761 | 0.538622077 |
| *SMARCB1* | 0.275637 | 0.123084 | 2.239426 | 0.02512823 | 0.538957086 |
| *B3GNT2* | -0.41359 | 0.184711 | -2.239126 | 0.0251477 | 0.538957086 |
| *UBB* | 0.163526 | 0.073047 | 2.238651 | 0.02517866 | 0.538957086 |
| *CLIC4* | -0.278977 | 0.124742 | -2.236429 | 0.0253237 | 0.540769931 |
| *DUSP2* | -0.891828 | 0.398838 | -2.236069 | 0.02534728 | 0.540769931 |
| *VAMP1* | -0.419052 | 0.187529 | -2.234599 | 0.0254437 | 0.541378963 |
| *SMIM12* | -0.21221 | 0.094976 | -2.234353 | 0.02545985 | 0.541378963 |
| *DENND6A* | 0.406568 | 0.182165 | 2.231861 | 0.02562414 | 0.543974786 |
| *FAM111B* | -0.326995 | 0.146649 | -2.229781 | 0.02576201 | 0.546001967 |
| *ZNF827* | 0.613148 | 0.275095 | 2.228862 | 0.02582306 | 0.546397383 |
| *TPM4* | -0.200063 | 0.089815 | -2.227492 | 0.02591444 | 0.547431948 |

| | | | | | |
|---|---|---|---|---|---|
| *MFN1* | -0.340681 | 0.153156 | -2.224408 | 0.02612101 | 0.550892529 |
| *POLR2G* | 0.186643 | 0.083958 | 2.223042 | 0.02621296 | 0.55117102 |
| *ABHD15* | 0.386817 | 0.174012 | 2.222941 | 0.02621976 | 0.55117102 |
| *FIP1L1* | 0.254676 | 0.114704 | 2.220292 | 0.02639892 | 0.552546992 |
| *DNPEP* | 0.282895 | 0.127432 | 2.21996 | 0.02642148 | 0.552546992 |
| *ENSG00000271857* | 1.457882 | 0.656796 | 2.219687 | 0.02644003 | 0.552546992 |
| *SLC37A1* | -0.425837 | 0.191867 | -2.219441 | 0.02645674 | 0.552546992 |
| *ENSG00000275441* | -1.126339 | 0.508741 | -2.213973 | 0.02683061 | 0.559448683 |
| *MTG1* | 0.371053 | 0.167835 | 2.210823 | 0.02704807 | 0.56307172 |
| *SAMD1* | -0.558498 | 0.25277 | -2.209511 | 0.02713913 | 0.56396083 |
| *CCDC12* | 0.145771 | 0.066002 | 2.208587 | 0.02720335 | 0.56396083 |
| *RPUSD3* | 0.231998 | 0.105057 | 2.208305 | 0.02722302 | 0.56396083 |
| *USF3* | -0.386615 | 0.175179 | -2.206965 | 0.02731651 | 0.56396083 |
| *ADAM28* | 0.199346 | 0.090332 | 2.206813 | 0.02732716 | 0.56396083 |
| *COMMD5* | -0.315502 | 0.142991 | -2.206438 | 0.02735337 | 0.56396083 |
| *HAUS4* | -0.61031 | 0.276746 | -2.205308 | 0.02743251 | 0.564689071 |
| *BACE2* | -0.611189 | 0.277438 | -2.202974 | 0.02759656 | 0.567051167 |
| *GTF3C5* | 0.321146 | 0.145815 | 2.202425 | 0.02763527 | 0.567051167 |
| *HACD4* | -0.386985 | 0.175889 | -2.200164 | 0.02779525 | 0.569330375 |
| *ENSG00000259772* | -0.751649 | 0.341774 | -2.199258 | 0.02785961 | 0.569330375 |
| *BIN3* | -0.321989 | 0.146426 | -2.198986 | 0.0278789 | 0.569330375 |
| *RUNX2* | 1.212432 | 0.551706 | 2.197604 | 0.02797737 | 0.570437354 |
| *ROMO1* | -0.14705 | 0.066962 | -2.196038 | 0.02808924 | 0.571813421 |
| *CCDC86* | -0.696137 | 0.317095 | -2.195363 | 0.02813759 | 0.571894371 |
| *NEK1* | 0.388509 | 0.177109 | 2.193618 | 0.02826288 | 0.573536098 |
| *PPP1CC* | 0.235584 | 0.107436 | 2.192792 | 0.02832236 | 0.573839535 |
| *CCDC85B* | -0.172474 | 0.078767 | -2.189666 | 0.02854846 | 0.577512465 |
| *SNX8* | -0.308831 | 0.141124 | -2.188372 | 0.0286425 | 0.578055849 |
| *KCNN1* | 1.538863 | 0.703299 | 2.188063 | 0.02866504 | 0.578055849 |
| *FAAP20* | -0.21484 | 0.09833 | -2.184897 | 0.02889638 | 0.581562862 |
| *FURIN* | 0.362469 | 0.16596 | 2.184072 | 0.02895696 | 0.581562862 |
| *PPP1R16B* | 0.376072 | 0.172207 | 2.183835 | 0.02897434 | 0.581562862 |
| *LINS1* | -0.400295 | 0.183529 | -2.181099 | 0.02917606 | 0.583679983 |
| *CTF1* | 1.789331 | 0.820747 | 2.180125 | 0.02924817 | 0.583679983 |
| *TRAF4* | -0.422154 | 0.193638 | -2.180117 | 0.0292488 | 0.583679983 |
| *VWCE* | 1.520267 | 0.697515 | 2.179549 | 0.02929092 | 0.583679983 |
| *C22orf39* | -0.209663 | 0.096205 | -2.179342 | 0.0293063 | 0.583679983 |
| *POLR1C* | 0.447676 | 0.205549 | 2.177956 | 0.02940934 | 0.584473692 |
| *ENSG00000166928* | -1.403549 | 0.644543 | -2.177586 | 0.02943686 | 0.584473692 |
| *NFAT5* | 0.270893 | 0.124513 | 2.175622 | 0.02958354 | 0.584897943 |
| *TFAM* | 0.282103 | 0.129667 | 2.1756 | 0.02958519 | 0.584897943 |
| *MPPED2* | 2.243387 | 1.031216 | 2.175477 | 0.0295944 | 0.584897943 |
| *TP53I11* | -0.506795 | 0.2331 | -2.174157 | 0.02969337 | 0.585602738 |
| *DDAH2* | -0.180063 | 0.082834 | -2.173789 | 0.02972095 | 0.585602738 |
| *RABEP2* | -0.238793 | 0.109915 | -2.172522 | 0.02981634 | 0.586050572 |
| *FAM114A2* | 0.408757 | 0.188173 | 2.172238 | 0.02983768 | 0.586050572 |
| *DAPK2* | 0.458197 | 0.210988 | 2.171676 | 0.02988012 | 0.586050572 |

| | | | | | |
|---|---|---|---|---|---|
| *WDFY2* | -0.289202 | 0.133257 | -2.170263 | 0.02998689 | 0.587250945 |
| *PMM1* | -0.327234 | 0.15115 | -2.16496 | 0.03039078 | 0.59425741 |
| *MPZL1* | -0.455942 | 0.210662 | -2.164334 | 0.03043873 | 0.59429305 |
| *SRP9* | -0.129233 | 0.059754 | -2.16274 | 0.03056122 | 0.59578188 |
| *ATAD2B* | -0.393018 | 0.181957 | -2.159949 | 0.03077661 | 0.598921171 |
| *FAM160B1* | 0.442619 | 0.204968 | 2.159451 | 0.03081521 | 0.598921171 |
| *ALKBH7* | 0.16143 | 0.074845 | 2.156844 | 0.03101782 | 0.601511468 |
| *BCL11A* | -0.276142 | 0.128049 | -2.156536 | 0.03104184 | 0.601511468 |
| *AUH* | 0.416266 | 0.193277 | 2.153727 | 0.03126157 | 0.604859779 |
| *TMEM256* | -0.143269 | 0.066552 | -2.152759 | 0.03133765 | 0.60527463 |
| *TMEM121* | -1.124501 | 0.522475 | -2.152259 | 0.03137696 | 0.60527463 |
| *SERINC5* | -0.681691 | 0.317134 | -2.149537 | 0.03159183 | 0.608138056 |
| *CTNND1* | -0.575661 | 0.267851 | -2.149184 | 0.03161978 | 0.608138056 |
| *SIDT1* | -0.439339 | 0.204499 | -2.148372 | 0.0316842 | 0.608468933 |
| *WDR7* | 0.642788 | 0.299321 | 2.147487 | 0.03175451 | 0.608911535 |
| *PROC* | -1.38314 | 0.644393 | -2.146424 | 0.03183919 | 0.609628315 |
| *PRDM8* | 0.691473 | 0.322446 | 2.144461 | 0.03199597 | 0.611721211 |
| *NECAP2* | 0.183933 | 0.085803 | 2.143655 | 0.03206053 | 0.612047462 |
| *PARP9* | -0.302492 | 0.141154 | -2.142989 | 0.032114 | 0.612161272 |
| *DALRD3* | -0.244122 | 0.113973 | -2.141926 | 0.03219943 | 0.612883063 |
| *C16orf74* | 0.359373 | 0.168007 | 2.139033 | 0.03243301 | 0.6164186 |
| *TRIM38* | 0.176786 | 0.082682 | 2.138146 | 0.0325049 | 0.616875071 |
| *ENSG00000250132* | -1.439858 | 0.674062 | -2.136089 | 0.03267213 | 0.618248412 |
| *ZFAND4* | 0.713954 | 0.334236 | 2.136076 | 0.03267322 | 0.618248412 |
| *UGT8* | 0.476019 | 0.222937 | 2.135217 | 0.03274332 | 0.618666248 |
| *ANKRD13A* | 0.2833 | 0.132744 | 2.134177 | 0.03282826 | 0.619362944 |
| *REC8* | 0.412764 | 0.193487 | 2.133298 | 0.03290025 | 0.619660038 |
| *EBF1* | -0.264867 | 0.124187 | -2.132812 | 0.03294018 | 0.619660038 |
| *PELI1* | -0.356815 | 0.167456 | -2.130803 | 0.0331054 | 0.621860296 |
| *LSM7* | -0.219933 | 0.103463 | -2.125714 | 0.03352712 | 0.625929521 |
| *CEP70* | 0.807045 | 0.379754 | 2.125179 | 0.03357164 | 0.625929521 |
| *SLC43A2* | 0.348229 | 0.163903 | 2.124597 | 0.03362026 | 0.625929521 |
| *ENSG00000273319* | 0.339442 | 0.159796 | 2.124217 | 0.03365196 | 0.625929521 |
| *LSR* | -0.693752 | 0.326647 | -2.123862 | 0.0336817 | 0.625929521 |
| *CSAD* | 0.395833 | 0.186443 | 2.123079 | 0.03374725 | 0.625929521 |
| *IMP3* | 0.142731 | 0.06723 | 2.123038 | 0.03375068 | 0.625929521 |
| *ANKRD39* | 0.344987 | 0.162561 | 2.122193 | 0.03382149 | 0.625929521 |
| *ZNF835* | -0.871493 | 0.410786 | -2.121526 | 0.03387755 | 0.625929521 |
| *SQSTM1* | 0.26439 | 0.124639 | 2.121247 | 0.03390101 | 0.625929521 |
| *NCF2* | -0.580834 | 0.273828 | -2.121164 | 0.03390799 | 0.625929521 |
| *SPATA2L* | -0.426149 | 0.200915 | -2.121048 | 0.03391778 | 0.625929521 |
| *C9orf16* | -0.136338 | 0.064291 | -2.120624 | 0.0339535 | 0.625929521 |
| *LETM2* | -1.143164 | 0.539491 | -2.118969 | 0.03409309 | 0.627604984 |
| *PECAM1* | -0.548738 | 0.259068 | -2.118119 | 0.03416503 | 0.628032148 |
| *TRIM26* | 0.397076 | 0.187533 | 2.117361 | 0.03422923 | 0.628315976 |
| *FAM129C* | -0.254932 | 0.120508 | -2.115482 | 0.03438891 | 0.630227922 |
| *ARHGEF1* | 0.293082 | 0.138608 | 2.114458 | 0.03447619 | 0.630227922 |

| | | | | | |
|---|---|---|---|---|---|
| *SLC2A13* | 0.495667 | 0.234467 | 2.114021 | 0.03451344 | 0.630227922 |
| *UBE2F* | -0.267041 | 0.126333 | -2.113785 | 0.03453365 | 0.630227922 |
| *ZFP36L2* | -0.361548 | 0.171085 | -2.113267 | 0.03457793 | 0.630227922 |
| *PRDM2* | -0.1543 | 0.073063 | -2.11187 | 0.03469764 | 0.631516626 |
| *CLCN3* | 0.348465 | 0.165054 | 2.11122 | 0.03475342 | 0.631639745 |
| *UNC119* | -0.26387 | 0.125023 | -2.110576 | 0.03480875 | 0.631754255 |
| *PIH1D1* | -0.218924 | 0.103766 | -2.109776 | 0.0348777 | 0.632115301 |
| *ADAL* | -0.486977 | 0.230887 | -2.109158 | 0.03493093 | 0.632190999 |
| *KEAP1* | 0.352782 | 0.167442 | 2.106889 | 0.03512717 | 0.634850989 |
| *ATF3* | -0.80584 | 0.382584 | -2.10631 | 0.03517743 | 0.634868824 |
| *TRAPPC1* | -0.143876 | 0.068326 | -2.105721 | 0.03522863 | 0.634903584 |
| *SLC2A11* | 0.437776 | 0.207973 | 2.104972 | 0.03529373 | 0.635111677 |
| *EID1* | 0.203543 | 0.09672 | 2.104455 | 0.03533875 | 0.635111677 |
| *AIFM1* | 0.404115 | 0.192231 | 2.102237 | 0.03553251 | 0.637704597 |
| *UXT* | -0.114008 | 0.054287 | -2.100095 | 0.03572049 | 0.640073576 |
| *CYREN* | -0.27571 | 0.131316 | -2.099602 | 0.03576385 | 0.640073576 |
| *RNF24* | -0.539503 | 0.257196 | -2.097635 | 0.0359374 | 0.642287538 |
| *LY6E* | -0.278954 | 0.133173 | -2.094668 | 0.0362005 | 0.645835775 |
| *ZNF154* | -0.537774 | 0.256829 | -2.093897 | 0.03626912 | 0.645835775 |
| *PSMA5* | 0.192528 | 0.091956 | 2.093705 | 0.03628629 | 0.645835775 |
| *ISOC1* | -0.415295 | 0.198627 | -2.090828 | 0.03654347 | 0.649516 |
| *MRPL24* | 0.308353 | 0.147749 | 2.087013 | 0.036887 | 0.654718787 |
| *C9orf139* | 0.422902 | 0.202803 | 2.085282 | 0.03704375 | 0.656596668 |
| *TFEB* | 0.234093 | 0.112291 | 2.084697 | 0.03709682 | 0.65663405 |
| *INKA2* | -0.681637 | 0.327087 | -2.083961 | 0.03716375 | 0.656916473 |
| *SNAP23* | 0.178698 | 0.085813 | 2.082413 | 0.03730473 | 0.658505183 |
| *RUBCNL* | 0.262113 | 0.125943 | 2.081204 | 0.03741528 | 0.659543107 |
| *PRX* | 1.426858 | 0.685887 | 2.080311 | 0.03749698 | 0.659543107 |
| *DDX28* | 0.395261 | 0.190021 | 2.080092 | 0.03751708 | 0.659543107 |
| *TIAM2* | 0.477791 | 0.229925 | 2.078024 | 0.03770713 | 0.661981098 |
| *RECQL* | -0.260273 | 0.125418 | -2.075249 | 0.03796344 | 0.664702552 |
| *NIFK* | 0.20252 | 0.097589 | 2.075229 | 0.03796532 | 0.664702552 |
| *RPS19* | 0.161766 | 0.078163 | 2.069585 | 0.03849121 | 0.670432373 |
| *KLHL5* | -0.33747 | 0.163073 | -2.06944 | 0.0385048 | 0.670432373 |
| *QSOX2* | -0.386959 | 0.186988 | -2.069432 | 0.03850557 | 0.670432373 |
| *HMGN1* | -0.136599 | 0.066021 | -2.069034 | 0.0385429 | 0.670432373 |
| *HAUS8* | 0.366481 | 0.177136 | 2.068929 | 0.03855272 | 0.670432373 |
| *RUNX1* | -0.533248 | 0.257902 | -2.067643 | 0.03867357 | 0.671021597 |
| *TNFRSF17* | 0.642299 | 0.31067 | 2.067461 | 0.03869075 | 0.671021597 |
| *RCBTB2* | -0.907649 | 0.439344 | -2.065918 | 0.0388362 | 0.671521775 |
| *ZNF133* | 0.795028 | 0.384908 | 2.065501 | 0.03887564 | 0.671521775 |
| *TCAF2* | 0.449746 | 0.217742 | 2.065498 | 0.03887593 | 0.671521775 |
| *PDIA6* | 0.210175 | 0.10188 | 2.062963 | 0.03911617 | 0.674641031 |
| *MSC* | -0.743887 | 0.360674 | -2.062489 | 0.03916122 | 0.674641031 |
| *TPM3* | -0.132926 | 0.064559 | -2.058985 | 0.03949569 | 0.677475415 |
| *SLAMF1* | -0.766392 | 0.37222 | -2.058978 | 0.03949632 | 0.677475415 |
| *PHB2* | -0.1567 | 0.07611 | -2.058851 | 0.03950851 | 0.677475415 |

| | | | | | |
|---|---|---|---|---|---|
| *GDPD5* | -0.618897 | 0.300645 | -2.058564 | 0.03953605 | 0.677475415 |
| *RFX5* | 0.206839 | 0.100516 | 2.057771 | 0.0396121 | 0.677877182 |
| *LINC00894* | 1.995285 | 0.970346 | 2.05626 | 0.03975744 | 0.679461958 |
| *LY75* | -0.424165 | 0.206463 | -2.054435 | 0.0399336 | 0.681568756 |
| *RNMT* | -0.217427 | 0.105911 | -2.052916 | 0.04008077 | 0.683072952 |
| *RASSF5* | -0.247238 | 0.120461 | -2.052431 | 0.04012775 | 0.683072952 |
| *KCNMB4* | -1.832278 | 0.893933 | -2.049681 | 0.04039557 | 0.68551004 |
| *WIPF1* | -0.131474 | 0.064152 | -2.049407 | 0.04042236 | 0.68551004 |
| *HDLBP* | 0.257507 | 0.125655 | 2.049323 | 0.04043052 | 0.68551004 |
| *IMPAD1* | 0.506815 | 0.247615 | 2.046785 | 0.04067919 | 0.68807422 |
| *ENSG00000250303* | -0.553021 | 0.270203 | -2.04669 | 0.04068854 | 0.68807422 |
| *CNTRL* | -0.169076 | 0.082664 | -2.04533 | 0.04082232 | 0.689431737 |
| *NEK7* | -0.416045 | 0.203796 | -2.04148 | 0.0412031 | 0.694951745 |
| *PSME2* | 0.195335 | 0.09574 | 2.040279 | 0.04132257 | 0.696055678 |
| *SIGLEC5* | 0.816391 | 0.400246 | 2.039723 | 0.04137794 | 0.696078442 |
| *POLRMT* | 0.472251 | 0.231744 | 2.037811 | 0.04156886 | 0.698378524 |
| *AGPAT2* | -0.239093 | 0.117376 | -2.036989 | 0.0416511 | 0.69884901 |
| *GADD45G* | -2.001874 | 0.984388 | -2.033624 | 0.04198954 | 0.703611436 |
| *TMEM128* | 0.306624 | 0.150836 | 2.032829 | 0.0420698 | 0.704040758 |
| *FAM219B* | -0.216246 | 0.106438 | -2.031654 | 0.04218869 | 0.705114647 |
| *FAM217B* | 0.306594 | 0.151044 | 2.029836 | 0.04237319 | 0.706420026 |
| *ENSG00000261200* | 1.08251 | 0.53338 | 2.029527 | 0.04240464 | 0.706420026 |
| *WAC.AS1* | 0.246333 | 0.121395 | 2.029184 | 0.04243951 | 0.706420026 |
| *WFS1* | 1.421618 | 0.70086 | 2.028392 | 0.04252027 | 0.706420026 |
| *IFIT1* | -1.572046 | 0.775098 | -2.02819 | 0.04254089 | 0.706420026 |
| *CXCL16* | 0.401324 | 0.197939 | 2.027521 | 0.0426092 | 0.706643691 |
| *PLIN2* | 0.401698 | 0.198431 | 2.024377 | 0.04293137 | 0.711071472 |
| *NAA40* | -0.410233 | 0.202733 | -2.023516 | 0.04301997 | 0.711624248 |
| *STK40* | -0.320393 | 0.158546 | -2.020822 | 0.04329819 | 0.715122244 |
| *AKT3* | -0.335951 | 0.16628 | -2.020395 | 0.04334242 | 0.715122244 |
| *ENSG00000269968* | -0.653615 | 0.323749 | -2.018895 | 0.04349817 | 0.716774265 |
| *LBX2.AS1* | -1.421387 | 0.704578 | -2.017359 | 0.04365803 | 0.718489561 |
| *DDX42* | 0.260185 | 0.129131 | 2.014896 | 0.04391559 | 0.721487695 |
| *SP140* | 0.15833 | 0.078613 | 2.014039 | 0.04400543 | 0.721487695 |
| *LIPA* | 0.244862 | 0.121585 | 2.013927 | 0.04401715 | 0.721487695 |
| *FAM32A* | -0.177174 | 0.087994 | -2.01348 | 0.04406416 | 0.721487695 |
| *RTP4* | 0.459026 | 0.228099 | 2.012402 | 0.0441776 | 0.722427079 |
| *NFIC* | 0.355419 | 0.176779 | 2.010524 | 0.04437575 | 0.724747667 |
| *STMN1* | -0.314457 | 0.156455 | -2.009893 | 0.04444254 | 0.724919693 |
| *HSP90AB1* | -0.10182 | 0.050685 | -2.008882 | 0.04454961 | 0.725747432 |
| *IGLC2* | -0.378776 | 0.188616 | -2.008183 | 0.04462389 | 0.726039714 |
| *ITPKB* | -0.29628 | 0.147685 | -2.006167 | 0.04483843 | 0.728610352 |
| *FNBP1* | 0.200518 | 0.100011 | 2.004955 | 0.04496781 | 0.729792501 |
| *PSD3* | -0.751997 | 0.375245 | -2.004018 | 0.04506818 | 0.730282464 |
| *ENSG00000223881* | 0.800331 | 0.399444 | 2.003615 | 0.04511135 | 0.730282464 |
| *LMAN1* | 0.281933 | 0.140763 | 2.00289 | 0.04518914 | 0.730623872 |
| *IDI1* | -0.178935 | 0.089451 | -2.00037 | 0.04546034 | 0.734087704 |

| | | | | | |
|---|---|---|---|---|---|
| *TENT5C* | -0.335258 | 0.167729 | -1.998805 | 0.04562943 | 0.735316756 |
| *LRRC58* | -0.239469 | 0.119818 | -1.99861 | 0.04565058 | 0.735316756 |
| *U2AF1L4* | -0.242088 | 0.121376 | -1.994529 | 0.04609431 | 0.741537101 |
| *ENSG00000267390* | 0.409055 | 0.205338 | 1.992109 | 0.04635911 | 0.744403888 |
| *UTP3* | 0.255969 | 0.128554 | 1.99114 | 0.04646548 | 0.744403888 |
| *TMEM242* | -0.246229 | 0.123669 | -1.991038 | 0.04647666 | 0.744403888 |
| *RGL2* | -0.405487 | 0.203681 | -1.990794 | 0.04650358 | 0.744403888 |
| *INSIG1* | -0.186129 | 0.09363 | -1.98791 | 0.04682159 | 0.747573029 |
| *CRYM.AS1* | -1.126905 | 0.566907 | -1.987811 | 0.04683258 | 0.747573029 |
| *PGD* | -0.232237 | 0.116863 | -1.987262 | 0.04689339 | 0.747573029 |
| *M6PR* | 0.197287 | 0.099298 | 1.98681 | 0.04694349 | 0.747573029 |
| *ENSG00000272356* | 0.828222 | 0.417036 | 1.985972 | 0.04703648 | 0.747573029 |
| *ABI2* | -0.276494 | 0.139251 | -1.985582 | 0.04707977 | 0.747573029 |
| *ING2* | -0.38468 | 0.193761 | -1.985331 | 0.04710766 | 0.747573029 |
| *ENSG00000267519* | -0.316654 | 0.159746 | -1.982242 | 0.04745217 | 0.752113934 |
| *BACH1.AS1* | 0.957411 | 0.483631 | 1.979632 | 0.04774493 | 0.755824568 |
| *ORAI2* | 0.167358 | 0.0846 | 1.978215 | 0.04790443 | 0.756492178 |
| *MAP3K11* | -0.260701 | 0.131786 | -1.978214 | 0.04790452 | 0.756492178 |
| *CLEC17A* | 0.512395 | 0.259348 | 1.975705 | 0.04818824 | 0.75980879 |
| *PLEKHJ1* | -0.234145 | 0.118545 | -1.975158 | 0.0482502 | 0.75980879 |
| *CD2AP* | 0.307351 | 0.155689 | 1.97413 | 0.04836702 | 0.75980879 |
| *PAN3.AS1* | -1.18562 | 0.600594 | -1.974078 | 0.04837287 | 0.75980879 |
| *COPS6* | 0.151822 | 0.076928 | 1.973559 | 0.04843192 | 0.75980879 |
| *MRM1* | 0.61121 | 0.30975 | 1.973239 | 0.04846832 | 0.75980879 |
| *FAM120A* | -0.207311 | 0.105127 | -1.972003 | 0.04860931 | 0.761093082 |
| *SYPL1* | -0.175085 | 0.088817 | -1.971311 | 0.04868831 | 0.761404858 |
| *ZMAT3* | -0.280928 | 0.142564 | -1.970537 | 0.04877682 | 0.761700739 |
| *PYCARD* | -0.200132 | 0.101584 | -1.970113 | 0.04882546 | 0.761700739 |
| *PIP4K2A* | -0.208149 | 0.105784 | -1.967681 | 0.04910471 | 0.765130884 |
| *ENSG00000275418* | -1.705396 | 0.867139 | -1.966694 | 0.0492185 | 0.765977681 |
| *ITGB1* | 0.376965 | 0.191828 | 1.965122 | 0.04940013 | 0.767877091 |

| Cluster 6 | log2FC | lfcSE | stat | pvalue | padj |
|---|---|---|---|---|---|
| *VPREB3* | 2.398998 | 0.203538 | 11.786475 | < 2.22e-16 | < 2.22e-16 |
| *RIMS3* | -5.167318 | 1.001637 | -5.158873 | 2.48E-07 | 0.00128195 |
| *ENSG00000260526* | -4.125399 | 0.852792 | -4.837518 | 1.31E-06 | 0.00452259 |
| *EDA2R* | -4.021795 | 0.861283 | -4.669537 | 3.02E-06 | 0.00684525 |
| *NRG2* | -4.327405 | 0.930589 | -4.650177 | 3.32E-06 | 0.00684525 |
| *LCA5* | -4.897325 | 1.069226 | -4.58025 | 4.64E-06 | 0.00764376 |
| *ENSG00000269242* | -4.18984 | 0.919395 | -4.557171 | 5.18E-06 | 0.00764376 |
| *ENSG00000272455* | -4.841394 | 1.075238 | -4.502626 | 6.71E-06 | 0.00789628 |
| *ENSG00000254281* | -3.84343 | 0.854632 | -4.497173 | 6.89E-06 | 0.00789628 |
| *STEAP1B* | -3.874637 | 0.872533 | -4.440678 | 8.97E-06 | 0.00925456 |
| *ENSG00000269737* | -4.208938 | 0.966671 | -4.354054 | 1.34E-05 | 0.011862 |
| *EFNB1* | -3.871302 | 0.890542 | -4.34713 | 1.38E-05 | 0.011862 |
| *KLHL17* | -3.704641 | 0.858975 | -4.312861 | 1.61E-05 | 0.01279326 |
| *PRKCD* | -3.803037 | 0.888424 | -4.280655 | 1.86E-05 | 0.01305754 |
| *ENSG00000255882* | -4.022862 | 0.943551 | -4.263534 | 2.01E-05 | 0.01305754 |
| *LINC01550* | -4.34193 | 1.018711 | -4.26218 | 2.02E-05 | 0.01305754 |
| *DNAAF5* | -3.494397 | 0.833306 | -4.193414 | 2.75E-05 | 0.01668119 |
| *ENSG00000250222* | -4.213041 | 1.009901 | -4.171736 | 3.02E-05 | 0.01733114 |
| *MMP14* | -4.860452 | 1.172839 | -4.144175 | 3.41E-05 | 0.0185238 |
| *PEX7* | -3.487203 | 0.84625 | -4.120772 | 3.78E-05 | 0.01871218 |
| *TTLL1* | -4.583909 | 1.11291 | -4.118848 | 3.81E-05 | 0.01871218 |
| *MON1A* | -3.714765 | 0.912946 | -4.068986 | 4.72E-05 | 0.02124004 |
| *LINC02362* | -4.934308 | 1.213878 | -4.064914 | 4.81E-05 | 0.02124004 |
| *TUNAR* | -3.846624 | 0.947802 | -4.058469 | 4.94E-05 | 0.02124004 |
| *DOCK7* | -3.292774 | 0.817924 | -4.025771 | 5.68E-05 | 0.02245931 |
| *DOCK9.DT* | -4.097002 | 1.018956 | -4.020785 | 5.80E-05 | 0.02245931 |
| *TTC30B* | -3.857915 | 0.963479 | -4.004153 | 6.22E-05 | 0.02245931 |
| *LRRC7* | -3.109566 | 0.777362 | -4.000152 | 6.33E-05 | 0.02245931 |
| *RPGRIP1L* | -3.439908 | 0.861063 | -3.994956 | 6.47E-05 | 0.02245931 |
| *BORCS8.MEF2B* | -5.014476 | 1.255869 | -3.992832 | 6.53E-05 | 0.02245931 |
| *KIF9.AS1* | -3.666927 | 0.921242 | -3.980416 | 6.88E-05 | 0.02263599 |
| *ENSG00000262429* | -3.567989 | 0.897462 | -3.975644 | 7.02E-05 | 0.02263599 |
| *SCAT1* | -4.683505 | 1.180855 | -3.966199 | 7.30E-05 | 0.02264674 |
| *C11orf65* | -4.689411 | 1.185613 | -3.955262 | 7.65E-05 | 0.02264674 |
| *UBTD2* | -3.459114 | 0.876161 | -3.948033 | 7.88E-05 | 0.02264674 |
| *TFR2* | -3.944148 | 0.999173 | -3.947413 | 7.90E-05 | 0.02264674 |
| *CFAP53* | -4.527794 | 1.150363 | -3.935969 | 8.29E-05 | 0.02290411 |
| *DOK2* | -3.74273 | 0.95193 | -3.93173 | 8.43E-05 | 0.02290411 |
| *MTCL1* | -3.874723 | 0.98824 | -3.920832 | 8.82E-05 | 0.02335067 |
| *ETS2* | -4.775205 | 1.219986 | -3.914148 | 9.07E-05 | 0.02340676 |
| *ENSG00000261845* | -3.845214 | 0.984296 | -3.906561 | 9.36E-05 | 0.02356461 |
| *MYLK* | -4.546023 | 1.16574 | -3.89969 | 9.63E-05 | 0.02366621 |
| *RUSC1.AS1* | -3.945453 | 1.016404 | -3.881778 | 0.000103696 | 0.02488694 |
| *SUOX* | -4.43226 | 1.144944 | -3.871159 | 0.000108319 | 0.02540577 |
| *LINC01011* | -4.442503 | 1.149527 | -3.864636 | 0.000111255 | 0.02551449 |
| *FITM1* | -4.446148 | 1.153677 | -3.853892 | 0.000116255 | 0.02608149 |

| | | | | | |
|---|---|---|---|---|---|
| *NKX6.3* | -3.905024 | 1.017993 | -3.836002 | 0.000125054 | 0.02689026 |
| *LILRB3* | -3.425206 | 0.892919 | -3.835967 | 0.000125071 | 0.02689026 |
| *FAM66C* | -3.500045 | 0.914734 | -3.8263 | 0.000130084 | 0.02739727 |
| *CCNF* | -2.946367 | 0.771533 | -3.818847 | 0.000134077 | 0.02767351 |
| *ZNF778* | -3.177171 | 0.833927 | -3.809892 | 0.000139027 | 0.02813262 |
| *ENSG00000272323* | -3.141245 | 0.82734 | -3.796799 | 0.000146576 | 0.02908978 |
| *TESMIN* | -3.819005 | 1.008768 | -3.785813 | 0.000153207 | 0.02947101 |
| *FBXL18* | -3.241699 | 0.856642 | -3.784192 | 0.000154209 | 0.02947101 |
| *ENSG00000274922* | -5.552477 | 1.47433 | -3.766102 | 0.000165816 | 0.03110746 |
| *MORN4* | -3.734529 | 0.994498 | -3.755192 | 0.000173209 | 0.03110746 |
| *NLRX1* | -3.212485 | 0.856005 | -3.752882 | 0.000174813 | 0.03110746 |
| *GNMT* | -4.312586 | 1.149147 | -3.75286 | 0.000174829 | 0.03110746 |
| *MPZ* | -2.957587 | 0.789586 | -3.745746 | 0.000179858 | 0.03135851 |
| *NR2F6* | -3.613754 | 0.96662 | -3.738546 | 0.000185088 | 0.03135851 |
| *ENSG00000267731* | -3.383145 | 0.905024 | -3.738182 | 0.000185356 | 0.03135851 |
| *YY2* | -3.61555 | 0.970452 | -3.725637 | 0.000194823 | 0.03194385 |
| *DSP* | -3.107111 | 0.834096 | -3.725123 | 0.00019522 | 0.03194385 |
| *KCTD21* | -4.078626 | 1.096127 | -3.720943 | 0.000198481 | 0.03194385 |
| *FAM214B* | -2.774365 | 0.746635 | -3.715826 | 0.000202541 | 0.03194385 |
| *DNASE1L3* | -3.008675 | 0.810167 | -3.713649 | 0.000204292 | 0.03194385 |
| *STAM* | -2.619392 | 0.708722 | -3.695936 | 0.000219078 | 0.03288635 |
| *CA3.AS1* | -5.642094 | 1.527626 | -3.693373 | 0.000221299 | 0.03288635 |
| *ENSG00000272750* | -2.921729 | 0.791337 | -3.692141 | 0.000222374 | 0.03288635 |
| *PRICKLE3* | -3.155034 | 0.85471 | -3.691351 | 0.000223066 | 0.03288635 |
| *CABP4* | -2.947856 | 0.800761 | -3.681318 | 0.000232031 | 0.03306141 |
| *ENSG00000269973* | -3.479697 | 0.945278 | -3.681136 | 0.000232197 | 0.03306141 |
| *KIF18A* | -3.43613 | 0.933906 | -3.679311 | 0.000233865 | 0.03306141 |
| *TTC22* | -4.808618 | 1.310051 | -3.670558 | 0.000242021 | 0.03375218 |
| *ENSG00000271855* | -3.705484 | 1.014062 | -3.654099 | 0.000258086 | 0.03551268 |
| *FBXL19.AS1* | -2.814771 | 0.771252 | -3.649614 | 0.000262635 | 0.03566307 |
| *ZNF687.AS1* | -3.562818 | 0.977671 | -3.644191 | 0.000268235 | 0.03586075 |
| *RPP40* | -3.089919 | 0.848526 | -3.641513 | 0.000271041 | 0.03586075 |
| *TRAM2.AS1* | -3.20783 | 0.88724 | -3.615517 | 0.000299749 | 0.03905293 |
| *ADNP.AS1* | -3.733385 | 1.033335 | -3.612948 | 0.000302736 | 0.03905293 |
| *PAK1* | -3.064607 | 0.849034 | -3.609523 | NA | NA |
| *ENSG00000228242* | -5.384954 | 1.494191 | -3.603927 | 0.000313445 | 0.03993525 |
| *DCLRE1B* | -3.250855 | 0.903101 | -3.599657 | 0.000318637 | 0.04010162 |
| *EGLN3* | -4.804824 | 1.339552 | -3.586889 | 0.000334647 | 0.04129041 |
| *PRRG2* | -2.822238 | 0.787066 | -3.58577 | 0.000336085 | 0.04129041 |
| *PLAAT4* | 2.053964 | 0.574489 | 3.575291 | 0.000349839 | 0.04168856 |
| *PEAK3* | -4.080305 | 1.141274 | -3.575219 | 0.000349934 | 0.04168856 |
| *ENSG00000233818* | -4.985398 | 1.394871 | -3.574093 | 0.000351444 | 0.04168856 |
| *ENSG00000042317* | -4.130983 | 1.157234 | -3.569703 | 0.000357386 | 0.04191163 |
| *ENSG00000272941* | -3.277561 | 0.919008 | -3.566411 | 0.000361903 | 0.04196451 |
| *ZNF624* | -3.086788 | 0.866695 | -3.561563 | 0.000368654 | 0.04208215 |
| *ENSG00000175773* | -5.239481 | 1.471828 | -3.559845 | 0.000371073 | 0.04208215 |
| *SEC61A2* | -2.63324 | 0.741372 | -3.551845 | 0.00038254 | 0.04291102 |

| | | | | | |
|---|---|---|---|---|---|
| *NPEPL1* | -2.729884 | 0.770632 | -3.542398 | 0.000396507 | 0.04399949 |
| *ENSG00000272821* | -4.440453 | 1.255205 | -3.537631 | 0.000403733 | 0.04432477 |
| *ENSG00000273226* | -3.41913 | 0.96992 | -3.525167 | 0.000423216 | 0.04524716 |
| *SLC35E4* | -4.653346 | 1.320466 | -3.524019 | 0.000425054 | 0.04524716 |
| *FSTL3* | -3.109234 | 0.882334 | -3.523873 | 0.000425288 | 0.04524716 |
| *IQCK* | -3.011326 | 0.855732 | -3.519008 | 0.000433164 | 0.04561477 |
| *ENSG00000261542* | -3.709394 | 1.061484 | -3.494534 | 0.00047489 | 0.04903247 |
| *POMT2* | -3.815747 | 1.091959 | -3.494404 | 0.000475121 | 0.04903247 |
| *ENSG00000270871* | -4.236244 | 1.214806 | -3.487178 | 0.000488147 | 0.04987797 |
| *TRPV1* | -3.532576 | 1.013879 | -3.484217 | 0.000493579 | 0.04993854 |
| *CDCA5* | -4.759384 | 1.367524 | -3.480291 | 0.000500869 | 0.05018413 |
| *LINC00887* | -4.785735 | 1.378655 | -3.471307 | 0.000517932 | 0.05087775 |
| *ENSG00000251034* | -3.429275 | 0.989654 | -3.465127 | 0.000529981 | 0.05087775 |
| *CHEK2* | -2.927888 | 0.845368 | -3.463449 | 0.000533298 | 0.05087775 |
| *ZNF862* | -4.033071 | 1.16489 | -3.46219 | 0.000535799 | 0.05087775 |
| *MSANTD3* | -2.819357 | 0.814344 | -3.462119 | 0.00053594 | 0.05087775 |
| *CHKB.DT* | -3.61049 | 1.043258 | -3.460785 | 0.000538603 | 0.05087775 |
| *LRRC3* | -3.881096 | 1.122047 | -3.458942 | 0.000542302 | 0.05087775 |
| *ENSG00000260879* | -3.183433 | 0.921722 | -3.453788 | 0.000552772 | 0.05130562 |
| *ENKD1* | -2.956688 | 0.857182 | -3.449313 | 0.000562015 | 0.05130562 |
| *PPP1R3D* | -3.012203 | 0.874426 | -3.444777 | 0.000571531 | 0.05130562 |
| *NRSN2* | -3.128599 | 0.908241 | -3.444679 | 0.000571737 | 0.05130562 |
| *POLR1B* | -2.83851 | 0.824239 | -3.443795 | 0.000573611 | 0.05130562 |
| *RAB3A* | -3.499463 | 1.016592 | -3.442346 | 0.000576691 | 0.05130562 |
| *ENSG00000273576* | -3.230707 | 0.93972 | -3.437946 | NA | NA |
| *LMNTD1* | -3.023067 | 0.880233 | -3.434396 | 0.000593876 | 0.05154893 |
| *TBC1D16* | -3.415565 | 0.994953 | -3.432889 | 0.000597186 | 0.05154893 |
| *NPHP4* | -3.901238 | 1.136677 | -3.432142 | 0.000598833 | 0.05154893 |
| *HDX* | -4.180261 | 1.218121 | -3.431728 | 0.000599748 | 0.05154893 |
| *ENSG00000179979* | -3.740624 | 1.091251 | -3.427832 | 0.000608421 | 0.05154893 |
| *ACTL10* | -3.232564 | 0.943154 | -3.427398 | 0.000609396 | 0.05154893 |
| *ENSG00000226853* | -5.030499 | 1.469038 | -3.424349 | 0.000616275 | 0.05170696 |
| *ZSCAN30* | -2.312337 | 0.676597 | -3.417599 | 0.000631762 | 0.05257892 |
| *C16orf46* | -4.210545 | 1.235031 | -3.409261 | 0.000651391 | 0.05370047 |
| *LIG4* | -2.60251 | 0.764163 | -3.405701 | 0.000659943 | 0.05370047 |
| *ADPRH* | -3.2079 | 0.942024 | -3.405327 | 0.000660849 | 0.05370047 |
| *ZMYM1* | -2.464825 | 0.725161 | -3.399003 | 0.000676321 | 0.05452834 |
| *TSPAN32* | -3.26166 | 0.960615 | -3.395387 | 0.000685316 | 0.05466684 |
| *ENSG00000231105* | -4.72428 | 1.391924 | -3.394065 | 0.000688633 | 0.05466684 |
| *ARHGAP19* | -2.523122 | 0.744025 | -3.39118 | 0.000695924 | 0.05470289 |
| *TMEM164* | -3.486981 | 1.028438 | -3.390561 | NA | NA |
| *HIST3H2A* | -3.856289 | 1.137649 | -3.389701 | 0.000699688 | 0.05470289 |
| *POLE2* | -3.679127 | 1.087092 | -3.384376 | 0.000713403 | 0.05513297 |
| *ENSG00000279571* | -4.99802 | 1.477207 | -3.383426 | 0.000715874 | 0.05513297 |
| *TIGD2* | -2.829888 | 0.837385 | -3.379434 | 0.000726353 | 0.05552566 |
| *ENSG00000229729* | -2.699717 | 0.799362 | -3.377338 | 0.000731911 | 0.05553917 |
| *ZDHHC9* | -3.900873 | 1.157008 | -3.371518 | 0.000747551 | 0.05573145 |

| | | | | | |
|---|---|---|---|---|---|
| *SACS* | -2.859752 | 0.848294 | -3.371181 | 0.000748467 | 0.05573145 |
| *APOO* | -2.552369 | 0.757294 | -3.37038 | 0.000750646 | 0.05573145 |
| *ZNF18* | -3.179919 | 0.94519 | -3.364318 | 0.00076733 | 0.05636206 |
| *ATXN7L2* | -3.149872 | 0.936532 | -3.363337 | 0.000770063 | 0.05636206 |
| *KANK1* | -3.557882 | 1.058576 | -3.361007 | NA | NA |
| *AMY2B* | -3.24104 | 0.965542 | -3.356705 | 0.000788771 | 0.05732478 |
| *LY86.AS1* | -3.562673 | 1.062368 | -3.353521 | 0.000797904 | 0.05733628 |
| *MMAA* | -2.691076 | 0.80264 | -3.352781 | 0.000800041 | 0.05733628 |
| *ENSG00000224046* | -2.852037 | 0.851657 | -3.348808 | 0.000811601 | 0.0574775 |
| *KLHL23* | -3.264355 | 0.974935 | -3.348279 | 0.000813151 | 0.0574775 |
| *MAMLD1* | -4.859202 | 1.455732 | -3.337978 | NA | NA |
| *CUBN* | -4.114567 | 1.235128 | -3.331287 | 0.000864454 | 0.06068818 |
| *ENSG00000260081* | -3.048897 | 0.916462 | -3.326812 | 0.000878455 | 0.06125442 |
| *HLCS* | -3.108206 | 0.935158 | -3.323723 | 0.000888245 | 0.0615214 |
| *DLGAP3* | -4.461326 | 1.343549 | -3.320554 | 0.000898389 | 0.06180917 |
| *SDCBP2* | -4.412018 | 1.330759 | -3.315414 | 0.000915074 | 0.06211332 |
| *SEMA4F* | -2.931694 | 0.884712 | -3.313728 | 0.000920609 | 0.06211332 |
| *BNIPL* | -3.729451 | 1.125481 | -3.31365 | 0.000920866 | 0.06211332 |
| *ENSG00000272588* | -3.286454 | 0.993652 | -3.30745 | 0.000941496 | 0.06309249 |
| *KCNMB4* | -3.909978 | 1.18417 | -3.301873 | 0.000960415 | 0.06364056 |
| *ENSG00000249476* | -2.818741 | 0.8538 | -3.301408 | 0.000962009 | 0.06364056 |
| *PART1* | -3.761656 | 1.14281 | -3.291585 | 0.000996247 | 0.06470832 |
| *SCD* | -3.315942 | 1.007757 | -3.290418 | 0.001000386 | 0.06470832 |
| *CEP70* | -3.881799 | 1.179858 | -3.290056 | 0.001001676 | 0.06470832 |
| *LACTB2.AS1* | -2.84584 | 0.865716 | -3.287267 | 0.001011648 | 0.06470832 |
| *SEMA3D* | -4.315024 | 1.312699 | -3.287139 | 0.00101211 | 0.06470832 |
| *MIR22HG* | -3.275872 | 0.996881 | -3.286122 | 0.00101577 | 0.06470832 |
| *ENSG00000234665* | -4.876589 | 1.48745 | -3.278489 | 0.001043645 | 0.06594748 |
| *ENSG00000260219* | -3.60513 | 1.100372 | -3.276284 | 0.001051829 | 0.06594748 |
| *ZNF416* | -2.468212 | 0.753515 | -3.275596 | 0.001054393 | 0.06594748 |
| *KRBA2* | -3.013205 | 0.92151 | -3.269857 | 0.00107602 | 0.06689475 |
| *KDM1B* | -3.337258 | 1.021162 | -3.268098 | 0.001082729 | 0.06690876 |
| *HLA.DQB2* | -3.755159 | 1.150887 | -3.262838 | 0.001103024 | 0.06775718 |
| *ENSG00000278831* | -3.96782 | 1.21784 | -3.258081 | 0.001121684 | 0.06825556 |
| *LINC02604* | -3.68793 | 1.13225 | -3.25717 | 0.00112529 | 0.06825556 |
| *IGIP* | -2.823666 | 0.867845 | -3.253652 | 0.00113932 | 0.06825556 |
| *NIPAL4* | -2.640692 | 0.811742 | -3.253119 | 0.001141459 | 0.06825556 |
| *MMP25.AS1* | -2.570065 | 0.790197 | -3.252435 | 0.001144207 | 0.06825556 |
| *ENSG00000236514* | -2.463099 | 0.758815 | -3.245983 | 0.001170458 | 0.06901608 |
| *HIST1H4E* | -3.023311 | 0.931774 | -3.244685 | 0.001175808 | 0.06901608 |
| *PRKCI* | -2.818437 | 0.869191 | -3.2426 | 0.001184445 | 0.06901608 |
| *PSPH* | -2.510956 | 0.774411 | -3.242409 | 0.001185238 | 0.06901608 |
| *ZNF512B* | -3.77413 | 1.164434 | -3.241172 | 0.001190394 | 0.06901608 |
| *PTPRJ* | -3.921015 | 1.21073 | -3.238555 | 0.001201368 | 0.06926325 |
| *ENSG00000272010* | -3.93619 | 1.216526 | -3.235598 | 0.001213881 | 0.06959587 |
| *NUDT8* | -2.532327 | 0.783881 | -3.2305 | 0.001235741 | 0.06992444 |
| *IFT46* | -2.566647 | 0.794587 | -3.230164 | 0.001237192 | 0.06992444 |

| | | | | | |
|---|---|---|---|---|---|
| *GCNT1* | -2.716194 | 0.841049 | -3.22953 | 0.001239939 | 0.06992444 |
| *ENSG00000238260* | -3.639212 | 1.127459 | -3.227801 | 0.001247458 | 0.06996613 |
| *EXD2* | -3.169497 | 0.983091 | -3.224012 | 0.001264083 | 0.07042853 |
| *ENSG00000240219* | -4.307335 | 1.337035 | -3.221557 | 0.00127496 | 0.07042853 |
| *RHPN1* | -2.758824 | 0.856436 | -3.221284 | 0.001276176 | 0.07042853 |
| *LEKR1* | -3.756345 | 1.167902 | -3.216319 | 0.001298464 | 0.07127737 |
| *PIK3R2* | -2.912027 | 0.90758 | -3.208563 | 0.001333999 | 0.07284056 |
| *MTFR2* | -3.626688 | 1.130934 | -3.206807 | 0.00134217 | 0.072901 |
| *MAP3K10* | -3.695985 | 1.153443 | -3.204307 | 0.001353882 | 0.07315217 |
| *AGBL5* | -2.7487 | 0.858495 | -3.201765 | 0.001365882 | 0.07330167 |
| *CAB39L* | -3.563841 | 1.113878 | -3.199489 | 0.001376715 | 0.07330167 |
| *CLCN4* | -3.349161 | 1.047017 | -3.198766 | 0.00138017 | 0.07330167 |
| *PAG1* | -2.485823 | 0.777513 | -3.197146 | 0.001387946 | 0.07330167 |
| *C17orf58* | -3.868296 | 1.210253 | -3.196271 | 0.001392163 | 0.07330167 |
| *TMPRSS13* | -3.630838 | 1.139109 | -3.187436 | 0.001435402 | 0.07505791 |
| *C1orf159* | -2.768466 | 0.868813 | -3.186491 | 0.001440098 | 0.07505791 |
| *CAPN3* | -3.141899 | 0.986455 | -3.18504 | 0.001447338 | 0.07505791 |
| *ENSG00000267469* | -2.924437 | 0.920091 | -3.178421 | 0.001480795 | 0.07640905 |
| *ENSG00000247765* | -3.492106 | 1.099867 | -3.175026 | 0.00149823 | 0.07670695 |
| *EHBP1L1* | -2.181921 | 0.687348 | -3.174406 | 0.001501434 | 0.07670695 |
| *ZC4H2* | -3.407431 | 1.075152 | -3.169254 | 0.001528307 | 0.07728382 |
| *NDC1* | -2.490428 | 0.786022 | -3.168393 | 0.00153284 | 0.07728382 |
| *ALDH7A1* | -4.604992 | 1.45362 | -3.167948 | 0.001535192 | 0.07728382 |
| *MACROD2* | -3.201606 | 1.011404 | -3.165507 | 0.001548127 | 0.07755666 |
| *DHX32* | -2.873851 | 0.909186 | -3.160904 | 0.001572805 | 0.07811062 |
| *PTGS1* | -3.191974 | 1.010022 | -3.160301 | 0.001576063 | 0.07811062 |
| *CSRNP2* | -2.917432 | 0.923692 | -3.158446 | 0.001586128 | 0.07811062 |
| *TNFSF8* | -4.195958 | 1.329243 | -3.156653 | 0.001595909 | 0.07811062 |
| *ENSG00000278932* | -3.826659 | 1.21233 | -3.156449 | 0.001597029 | 0.07811062 |
| *ENSG00000247121* | -2.713115 | 0.860915 | -3.151433 | 0.001624717 | 0.07908999 |
| *LDLR* | -3.453749 | 1.097736 | -3.146247 | 0.0016538 | 0.08012779 |
| *BBC3* | -2.39947 | 0.763015 | -3.14472 | 0.001662457 | 0.08017085 |
| *SHPK* | -2.367097 | 0.753584 | -3.141118 | 0.00168304 | 0.08078592 |
| *CKAP5* | -2.321548 | 0.74021 | -3.136336 | 0.001710729 | 0.08141824 |
| *SLC35D1* | -2.254322 | 0.718825 | -3.13612 | 0.001711992 | 0.08141824 |
| *ARL6* | -3.821935 | 1.220222 | -3.132163 | 0.001735235 | 0.08214506 |
| *ZNF142* | -2.715292 | 0.868701 | -3.125694 | 0.001773862 | 0.08338511 |
| *HMGN3.AS1* | -4.13163 | 1.322359 | -3.124439 | 0.001781442 | 0.08338511 |
| *ENSG00000280011* | -3.152186 | 1.009106 | -3.123742 | 0.00178567 | 0.08338511 |
| *PARPBP* | -2.485522 | 0.79699 | -3.118636 | 0.0018169 | 0.08409163 |
| *POLE* | -2.798369 | 0.897314 | -3.118605 | 0.001817096 | 0.08409163 |
| *IRAK1BP1* | -2.902662 | 0.931573 | -3.115873 | NA | NA |
| *AHI1* | 2.027739 | 0.651876 | 3.110619 | 0.001866959 | 0.08601346 |
| *ENSG00000258623* | -2.860982 | 0.920162 | -3.109216 | 0.001875843 | 0.08603869 |
| *NOTCH1* | -2.140962 | 0.689154 | -3.106651 | 0.001892194 | 0.08640463 |
| *PKD2* | -3.310615 | 1.066902 | -3.103016 | 0.001915593 | 0.08691742 |
| *ENSG00000268400* | -3.350807 | 1.080106 | -3.102294 | 0.001920269 | 0.08691742 |

| | | | | | |
|---|---|---|---|---|---|
| *TAF1A.AS1* | -3.126288 | 1.008336 | -3.100441 | 0.001932325 | 0.08708119 |
| *LINC02453* | -3.635038 | 1.174494 | -3.094983 | 0.001968241 | 0.08781241 |
| *ENSG00000261351* | -2.736258 | 0.884164 | -3.094739 | 0.001969858 | 0.08781241 |
| *KIAA1324L* | -2.800463 | 0.90586 | -3.091496 | 0.001991507 | 0.08781241 |
| *ADNP2* | -3.074121 | 0.994777 | -3.090263 | 0.001999797 | 0.08781241 |
| *DCBLD1* | -3.178094 | 1.028483 | -3.09008 | 0.002001025 | 0.08781241 |
| *CD38* | 1.779398 | 0.576028 | 3.089081 | 0.002007765 | 0.08781241 |
| *GNE* | -2.497922 | 0.808643 | -3.08903 | 0.002008113 | 0.08781241 |
| *ENSG00000260806* | -4.227174 | 1.370825 | -3.083671 | 0.002044633 | 0.08870691 |
| *BRICD5* | -2.546845 | 0.825957 | -3.083507 | 0.00204576 | 0.08870691 |
| *NUSAP1* | -2.569461 | 0.833899 | -3.081262 | 0.002061249 | 0.08900454 |
| *TNFSF13B* | -3.043429 | 0.988527 | -3.078751 | NA | NA |
| *ARMC5* | -2.968671 | 0.964679 | -3.077368 | 0.002088372 | 0.08979998 |
| *ZNF554* | -3.666642 | 1.192678 | -3.074293 | 0.002110021 | 0.089814 |
| *RAB40C* | -2.836495 | 0.923511 | -3.071424 | 0.002130406 | 0.089814 |
| *HOXB2* | -2.313424 | 0.753701 | -3.069421 | 0.002144744 | 0.089814 |
| *IQGAP2* | -2.24789 | 0.73249 | -3.068834 | 0.002148956 | 0.089814 |
| *ENSG00000276509* | -1.963352 | 0.639802 | -3.068684 | 0.002150039 | 0.089814 |
| *CD1D* | 1.917144 | 0.625299 | 3.065963 | 0.002169703 | 0.089814 |
| *KIFC2* | -2.670815 | 0.871154 | -3.065834 | 0.002170638 | 0.089814 |
| *MPND* | -2.374872 | 0.774754 | -3.065323 | 0.002174347 | 0.089814 |
| *PIGN* | -3.101769 | 1.012309 | -3.064054 | 0.002183593 | 0.089814 |
| *PSEN2* | -2.497214 | 0.815196 | -3.06333 | 0.002188885 | 0.089814 |
| *ENSG00000242349* | -2.651453 | 0.865688 | -3.062826 | 0.002192574 | 0.089814 |
| *MALT1* | -1.834874 | 0.599208 | -3.062166 | 0.002197417 | 0.089814 |
| *ZNF346* | -1.945092 | 0.635483 | -3.060807 | 0.002207414 | 0.089814 |
| *RGS12* | -2.93364 | 0.959114 | -3.058699 | 0.002223002 | 0.089814 |
| *HLX* | -3.598661 | 1.17662 | -3.058473 | 0.002224682 | 0.089814 |
| *TVP23C* | -2.42819 | 0.794036 | -3.058034 | 0.002227944 | 0.089814 |
| *GPR180* | -2.317601 | 0.758391 | -3.055946 | 0.002243519 | 0.09008995 |
| *CCDC152* | -2.445733 | 0.800707 | -3.054467 | 0.00225461 | 0.09018438 |
| *CORO2B* | -3.47848 | 1.139942 | -3.051453 | 0.002277365 | 0.09074288 |
| *NRIP2* | -2.711394 | 0.88984 | -3.047057 | 0.00231094 | 0.0912037 |
| *PEG10* | -3.179617 | 1.043773 | -3.046273 | 0.002316976 | 0.0912037 |
| *EAF1.AS1* | -3.030847 | 0.995279 | -3.045222 | 0.002325083 | 0.0912037 |
| *ENSG00000272058* | -2.359317 | 0.774799 | -3.045068 | 0.002326275 | 0.0912037 |
| *SOX12* | -2.150159 | 0.706317 | -3.044185 | 0.002333118 | 0.0912037 |
| *ENSG00000229043* | -3.114231 | 1.023892 | -3.041562 | NA | NA |
| *IQCN* | -2.459249 | 0.808806 | -3.040591 | 0.002361146 | 0.09195104 |
| *ENSG00000258646* | -3.745214 | 1.232595 | -3.038478 | 0.002377766 | 0.09225018 |
| *FAM174A* | -2.076234 | 0.683849 | -3.036098 | 0.002396618 | 0.09240297 |
| *ACVR1B* | -4.688807 | 1.544415 | -3.035976 | NA | NA |
| *C19orf44* | -3.362897 | 1.107783 | -3.0357 | 0.002399776 | 0.09240297 |
| *BPNT1* | -2.540745 | 0.837685 | -3.033057 | 0.002420902 | 0.09240297 |
| *ENSG00000262151* | -2.867587 | 0.945548 | -3.032725 | 0.002423564 | 0.09240297 |
| *VPS13B* | -1.907605 | 0.629082 | -3.032363 | 0.002426473 | 0.09240297 |
| *ENSG00000275672* | -2.71387 | 0.895572 | -3.030321 | 0.002442941 | 0.09255643 |

| | | | | | |
|---|---|---|---|---|---|
| *CABLES1* | -2.205369 | 0.727931 | -3.029642 | 0.00244844 | 0.09255643 |
| *RNF25* | -2.041613 | 0.674343 | -3.027561 | 0.002465363 | 0.09285602 |
| *ENSG00000111321* | -2.838534 | 0.938003 | -3.026145 | 0.002476935 | 0.09295261 |
| *FMN1* | -4.79826 | 1.587407 | -3.022704 | 0.002505273 | 0.09367543 |
| *ZNF597* | -3.829147 | 1.267532 | -3.020947 | 0.002519851 | 0.09386733 |
| *ZSWIM4* | -3.197723 | 1.058615 | -3.020667 | NA | NA |
| *LAMC1* | -2.536916 | 0.840067 | -3.019898 | 0.002528597 | 0.09386733 |
| *FBXO31* | -2.524307 | 0.836297 | -3.018433 | 0.002540855 | 0.09398433 |
| *FSIP2* | -2.34746 | 0.779963 | -3.009707 | 0.002614994 | 0.09615034 |
| *ENSG00000272716* | -2.599152 | 0.864127 | -3.007838 | 0.002631134 | 0.09615034 |
| *UTP15* | -2.987143 | 0.993123 | -3.007829 | 0.002631212 | 0.09615034 |
| *ZNF614* | -2.569989 | 0.854689 | -3.00693 | 0.002639007 | 0.09615034 |
| *IRS1* | -2.329382 | 0.775213 | -3.004828 | 0.002657312 | 0.09615034 |
| *C2CD2L* | -3.96558 | 1.320108 | -3.003982 | 0.002664715 | 0.09615034 |
| *ANKRD27* | -2.110673 | 0.702718 | -3.003583 | 0.002668205 | 0.09615034 |
| *ZNF726* | -2.51181 | 0.836453 | -3.002929 | 0.002673948 | 0.09615034 |
| *GIMAP1* | -3.102008 | 1.033378 | -3.001815 | 0.002683754 | 0.09616785 |
| *B4GALT5* | -3.015814 | 1.007144 | -2.994423 | 0.002749646 | 0.09818804 |
| *ENSG00000257764* | -2.775421 | 0.927253 | -2.993163 | 0.002761019 | 0.09821684 |
| *PAFAH2* | -2.670179 | 0.892934 | -2.990341 | 0.002786657 | 0.09821684 |
| *PDZD8* | -2.204744 | 0.737329 | -2.990176 | 0.002788167 | 0.09821684 |
| *PYGM* | -2.5112 | 0.839828 | -2.990137 | 0.002788521 | 0.09821684 |
| *HIST1H2AK* | -4.554066 | 1.524851 | -2.986564 | 0.002821315 | 0.0990339 |
| *DIP2C* | -2.687206 | 0.900238 | -2.984994 | 0.002835835 | 0.09920616 |
| *TNRC18* | -3.03931 | 1.018841 | -2.983106 | 0.002853395 | 0.09948325 |
| *PRDM11* | -3.944431 | 1.323463 | -2.980386 | 0.002878855 | 0.09979265 |
| *LINC01481* | -4.620901 | 1.551017 | -2.979272 | 0.002889343 | 0.09979265 |
| *C4orf46* | -2.555922 | 0.858162 | -2.978366 | 0.002897894 | 0.09979265 |
| *ZNF469* | -5.29595 | 1.778332 | -2.978043 | 0.002900949 | 0.09979265 |
| *DGCR6* | -4.150123 | 1.394952 | -2.9751 | 0.002928927 | 0.10042037 |
| *KIF1B* | -2.956675 | 0.994292 | -2.973648 | 0.002942824 | 0.10056272 |
| *LINC00342* | -3.511573 | 1.184142 | -2.965499 | 0.003021919 | 0.10226809 |
| *XXYLT1.AS2* | -2.748232 | 0.926743 | -2.965473 | 0.003022176 | 0.10226809 |
| *MSH5* | -3.0371 | 1.024352 | -2.964899 | 0.003027827 | 0.10226809 |
| *ALDH2* | -2.076013 | 0.700306 | -2.964438 | 0.003032368 | 0.10226809 |
| *SLC35G5* | -3.330036 | 1.124038 | -2.962566 | 0.003050868 | 0.10255687 |
| *TNNT3* | -4.390437 | 1.483427 | -2.959657 | 0.003079813 | 0.10307844 |
| *BBS7* | -2.192699 | 0.741026 | -2.959003 | 0.00308636 | 0.10307844 |
| *PINK1* | -2.459105 | 0.832742 | -2.953021 | 0.003146801 | 0.1045856 |
| *CLDN12* | -3.03685 | 1.028556 | -2.952536 | 0.003151756 | 0.1045856 |
| *ENSG00000272182* | -3.509284 | 1.189724 | -2.949662 | 0.003181221 | 0.10510526 |
| *ENSG00000261188* | -2.656217 | 0.90071 | -2.949025 | 0.003187786 | 0.10510526 |
| *EID2* | -2.924313 | 0.993396 | -2.943755 | 0.003242566 | 0.10633117 |
| *ENSG00000227908* | -5.20051 | 1.76766 | -2.942031 | 0.003260672 | 0.10633117 |
| *LZTR1* | -2.483661 | 0.844344 | -2.941527 | 0.003265984 | 0.10633117 |
| *SPATA2* | -2.326696 | 0.791102 | -2.941084 | 0.003270661 | 0.10633117 |
| *LRRC8A* | -2.86775 | 0.975249 | -2.940533 | 0.003276484 | 0.10633117 |

| | | | | | |
|---|---|---|---|---|---|
| *DCUN1D4* | -2.053278 | 0.698653 | -2.938912 | 0.00329367 | 0.10649917 |
| *ZNF555* | -2.788479 | 0.949075 | -2.9381 | 0.0033023 | 0.10649917 |
| *ZKSCAN4* | -2.770595 | 0.944352 | -2.933859 | 0.003347765 | 0.1075668 |
| *TRIM35* | -3.327027 | 1.13435 | -2.932981 | 0.003357241 | 0.1075668 |
| *TTC21B* | -2.088195 | 0.712363 | -2.931364 | 0.003374768 | 0.1075668 |
| *ENSG00000274712* | -2.025255 | 0.691139 | -2.930313 | 0.003386209 | 0.1075668 |
| *EVI5* | -1.655924 | 0.565125 | -2.930193 | 0.00338752 | 0.1075668 |
| *C1orf198* | -2.976125 | 1.016112 | -2.928934 | 0.003401267 | 0.10767201 |
| *LINC01754* | -4.172716 | 1.425937 | -2.926298 | 0.003430219 | 0.1080245 |
| *PAQR3* | -2.278466 | 0.778692 | -2.926016 | 0.003433337 | 0.1080245 |
| *SEC22A* | -1.944372 | 0.665344 | -2.922354 | 0.00347396 | 0.10897043 |
| *ZNF75D* | -2.691371 | 0.922394 | -2.917811 | 0.003524977 | 0.11023564 |
| *ENSG00000261669* | -2.186248 | 0.749858 | -2.915551 | 0.003550613 | 0.1107019 |
| *RFNG* | -2.383704 | 0.818616 | -2.911869 | 0.003592733 | 0.11123969 |
| *FBRSL1* | -1.99905 | 0.686607 | -2.911491 | 0.003597081 | 0.11123969 |
| *COCH* | -2.2071 | 0.758463 | -2.909965 | 0.003614689 | 0.11123969 |
| *DDX31* | -3.104492 | 1.0669 | -2.909825 | 0.003616313 | 0.11123969 |
| *ZNF354C* | -2.817994 | 0.968697 | -2.909057 | 0.003625202 | 0.11123969 |
| *ZBTB8A* | -3.365899 | 1.157293 | -2.908426 | 0.003632536 | 0.11123969 |
| *WDR81* | -1.914841 | 0.658612 | -2.907391 | 0.003644573 | 0.11127809 |
| *POU5F2* | -2.908996 | 1.001275 | -2.905292 | 0.003669102 | 0.11147516 |
| *COL4A4* | -5.206904 | 1.792399 | -2.904991 | 0.003672631 | 0.11147516 |
| *ENSG00000276136* | -2.286916 | 0.787886 | -2.902599 | 0.003700804 | 0.11200087 |
| *TRIM25* | -1.871778 | 0.645149 | -2.901313 | 0.003716029 | 0.1121328 |
| *RHEBL1* | -2.281659 | 0.786964 | -2.899319 | 0.00373974 | 0.1125193 |
| *LINC01772* | -2.409876 | 0.831492 | -2.898256 | 0.003752438 | 0.11257315 |
| *PHLPP2* | -3.210443 | 1.109391 | -2.893878 | 0.003805165 | 0.11300656 |
| *NEURL4* | -2.584395 | 0.893088 | -2.893776 | 0.003806399 | 0.11300656 |
| *CPNE8* | -2.572049 | 0.888917 | -2.893464 | 0.00381018 | 0.11300656 |
| *FAM234B* | -2.037276 | 0.704106 | -2.893422 | 0.003810686 | 0.11300656 |
| *GTF3C1* | -2.853905 | 0.987238 | -2.890797 | NA | NA |
| *ENSG00000267563* | -2.464181 | 0.853272 | -2.88792 | 0.003877979 | 0.11467263 |
| *FAM8A1* | -2.895748 | 1.0039 | -2.8845 | 0.003920358 | 0.11559456 |
| *BRAF* | -1.690877 | 0.587296 | -2.879087 | 0.003988285 | 0.11726241 |
| *KLHL42* | -2.54441 | 0.88443 | -2.876893 | 0.004016123 | 0.11774543 |
| *ENSG00000276718* | -5.100237 | 1.774677 | -2.873896 | 0.004054426 | 0.11799076 |
| *LLGL1* | -2.721592 | 0.947064 | -2.873714 | 0.004056757 | 0.11799076 |
| *LINC00910* | -2.474587 | 0.861155 | -2.873567 | NA | NA |
| *ENSG00000268093* | -4.656927 | 1.620944 | -2.872971 | 0.00406631 | 0.11799076 |
| *EFL1* | -3.00737 | 1.046891 | -2.872667 | 0.004070224 | 0.11799076 |
| *CECR2* | -2.079626 | 0.724659 | -2.869801 | 0.004107304 | 0.11873215 |
| *C8orf37* | -2.803034 | 0.977263 | -2.868249 | 0.004127501 | 0.1189827 |
| *MAP3K9* | -2.170958 | 0.75762 | -2.865498 | 0.004163543 | 0.11968736 |
| *ENSG00000260101* | -2.554617 | 0.89315 | -2.860233 | 0.004233301 | 0.12106036 |
| *WASHC5* | -2.009818 | 0.702703 | -2.860123 | 0.004234766 | 0.12106036 |
| *ENSG00000272993* | -2.300559 | 0.804941 | -2.858048 | 0.004262565 | 0.12128417 |
| *WDR24* | -2.493559 | 0.87255 | -2.857784 | 0.0042661 | 0.12128417 |

| | | | | | |
|---|---|---|---|---|---|
| *ZNF660* | -2.254017 | 0.78955 | -2.854811 | 0.004306238 | 0.12193111 |
| *DENND4C* | -2.140218 | 0.749809 | -2.854351 | 0.004312486 | 0.12193111 |
| *LSG1* | -1.854053 | 0.650005 | -2.852368 | 0.004339488 | 0.12235933 |
| *EML6* | -2.529797 | 0.887681 | -2.849893 | 0.004373397 | 0.12297946 |
| *ENSG00000273329* | -2.212184 | 0.776751 | -2.847995 | 0.004399564 | 0.12337909 |
| *AGFG2* | -4.940277 | 1.735397 | -2.846771 | NA | NA |
| *SERPINE2* | -2.463554 | 0.865532 | -2.846288 | 0.004423215 | 0.12370618 |
| *ERP27* | -3.668763 | 1.289841 | -2.844352 | 0.004450186 | 0.1241241 |
| *PTP4A3* | -2.185186 | 0.768806 | -2.842313 | 0.004478749 | 0.12458408 |
| *PASK* | -2.158233 | 0.75972 | -2.840828 | 0.004499661 | 0.12482931 |
| *TWNK* | -2.201728 | 0.77611 | -2.836877 | 0.004555715 | 0.12597515 |
| *KANSL1L* | -2.546907 | 0.898 | -2.8362 | 0.004565379 | 0.12597515 |
| *PIGA* | -2.402657 | 0.848008 | -2.833294 | 0.004607101 | 0.12674087 |
| *MDFIC* | -2.261188 | 0.798284 | -2.83256 | 0.004617691 | 0.12674087 |
| *STPG1* | -2.882077 | 1.018851 | -2.828753 | 0.004672969 | 0.12779449 |
| *DONSON* | -3.41808 | 1.208592 | -2.82815 | 0.004681789 | 0.12779449 |
| *MAPK11* | -2.714472 | 0.96007 | -2.827369 | 0.004693228 | 0.12779449 |
| *ENSG00000228037* | -3.512041 | 1.242613 | -2.826335 | 0.004708404 | 0.12787034 |
| *ZFYVE1* | -2.273949 | 0.805157 | -2.824229 | 0.004739451 | 0.12803016 |
| *SDR42E1* | -2.713906 | 0.961305 | -2.823148 | 0.004755462 | 0.12803016 |
| *ZXDC* | -2.224262 | 0.788019 | -2.822598 | 0.004763624 | 0.12803016 |
| *PRMT7* | -1.904927 | 0.674889 | -2.822579 | 0.004763913 | 0.12803016 |
| *LETM2* | -3.779077 | 1.339806 | -2.820615 | 0.004793167 | 0.12815129 |
| *WDPCP* | -2.001736 | 0.709744 | -2.820363 | 0.004796933 | 0.12815129 |
| *ENSG00000280007* | -2.478882 | 0.879105 | -2.819779 | 0.004805673 | 0.12815129 |
| *ENSG00000277511* | -2.388774 | 0.847697 | -2.817957 | 0.004833027 | 0.12854855 |
| *TMCC3* | -4.675816 | 1.661382 | -2.814413 | 0.004886637 | 0.12964035 |
| *DCAKD* | -2.290402 | 0.814194 | -2.813091 | 0.004906782 | 0.12984099 |
| *ZDBF2* | -2.780618 | 0.988902 | -2.811823 | 0.004926153 | 0.12986216 |
| *CD68* | -2.672293 | 0.950523 | -2.811393 | 0.004932749 | 0.12986216 |
| *NOD1* | -2.245561 | 0.799083 | -2.810174 | 0.004951479 | 0.13002358 |
| *FAM71D* | -3.006419 | 1.07051 | -2.808399 | 0.004978855 | 0.13016394 |
| *CAPN2* | -2.160449 | 0.769338 | -2.808192 | 0.00498205 | 0.13016394 |
| *ZNF839* | -2.845793 | 1.015604 | -2.802069 | 0.005077607 | 0.13232552 |
| *MAPK13* | -2.02279 | 0.722392 | -2.800127 | 0.005108249 | 0.13278873 |
| *PPM1F* | -2.047846 | 0.731763 | -2.798511 | 0.005133881 | 0.13311972 |
| *ST3GAL6* | -3.510679 | 1.255256 | -2.796783 | 0.005161411 | 0.13349814 |
| *MMP17* | -2.96331 | 1.060481 | -2.794306 | 0.005201119 | 0.13418886 |
| *C21orf58* | -2.910166 | 1.042326 | -2.791991 | 0.005238476 | 0.13481564 |
| *TKFC* | -1.993677 | 0.714379 | -2.790783 | 0.005258072 | 0.13498334 |
| *SERPINF2* | -3.484994 | 1.249696 | -2.788674 | 0.005292426 | 0.13533341 |
| *ZSWIM3* | -3.028297 | 1.086058 | -2.788337 | 0.005297936 | 0.13533341 |
| *PSMC3IP* | -2.768676 | 0.993697 | -2.786239 | 0.005332356 | 0.13562557 |
| *FHIT* | -1.983024 | 0.711772 | -2.786038 | 0.005335657 | 0.13562557 |
| *ENSG00000270175* | -4.448271 | 1.598613 | -2.782582 | 0.005392827 | 0.13617853 |
| *TCAP* | -3.739004 | 1.343807 | -2.782396 | 0.005395918 | 0.13617853 |
| *ACTR3B* | -2.207652 | 0.793541 | -2.782026 | 0.00540207 | 0.13617853 |

| | | | | | |
|---|---|---|---|---|---|
| *ENSG00000260267* | -2.155431 | 0.774906 | -2.781538 | 0.005410193 | 0.13617853 |
| *GALE* | -2.902035 | 1.043756 | -2.780378 | 0.005429569 | 0.13633371 |
| *TRABD2A* | -2.289072 | 0.823813 | -2.778631 | 0.005458856 | 0.13650732 |
| *HAUS5* | -2.423989 | 0.87266 | -2.777703 | 0.005474469 | 0.13650732 |
| *TRAF3IP2.AS1* | -2.012394 | 0.724508 | -2.777602 | 0.005476166 | 0.13650732 |
| *SCRG1* | -2.566507 | 0.924882 | -2.774957 | 0.005520904 | 0.13702705 |
| *FXR2* | -2.183157 | 0.78678 | -2.7748 | 0.005523571 | 0.13702705 |
| *SGK1* | -3.638223 | 1.312977 | -2.770973 | 0.005588905 | 0.13796671 |
| *ZNF565* | -2.398648 | 0.865693 | -2.770783 | 0.005592166 | 0.13796671 |
| *ZBTB20.AS2* | -3.289027 | 1.187528 | -2.769642 | 0.005611795 | 0.13796671 |
| *ENSG00000142046* | -2.273516 | 0.820924 | -2.76946 | 0.005614924 | 0.13796671 |
| *ZDHHC23* | -2.059221 | 0.743766 | -2.768639 | 0.005629089 | 0.13798621 |
| *ZNF549* | -2.140771 | 0.774448 | -2.764254 | 0.005705309 | 0.1395232 |
| *KCNMB3* | -1.790023 | 0.647919 | -2.762724 | 0.005732116 | 0.13956936 |
| *RWDD2B* | -2.219745 | 0.803498 | -2.762603 | 0.005734245 | 0.13956936 |
| *ENSG00000274341* | -2.983169 | 1.081193 | -2.759145 | 0.005795283 | 0.14062028 |
| *UBOX5* | -2.729669 | 0.990093 | -2.756983 | 0.005833739 | 0.14062028 |
| *PREP* | -2.382649 | 0.864317 | -2.756683 | 0.005839086 | 0.14062028 |
| *AMMECR1L* | -1.905445 | 0.69141 | -2.755883 | 0.005853397 | 0.14062028 |
| *MROH8* | -4.097207 | 1.486843 | -2.755642 | 0.005857701 | 0.14062028 |
| *RNASEL* | -1.949564 | 0.707594 | -2.755202 | 0.005865591 | 0.14062028 |
| *SDHAF4* | -2.180274 | 0.791446 | -2.7548 | 0.005872804 | 0.14062028 |
| *GPSM2* | -2.193564 | 0.796903 | -2.752612 | 0.005912187 | 0.14123558 |
| *ATL1* | -2.789411 | 1.013699 | -2.751715 | 0.005928418 | 0.14129624 |
| *PHC2* | -2.543746 | 0.924966 | -2.750097 | 0.005957758 | 0.14166834 |
| *VWA8* | -3.02578 | 1.101694 | -2.746479 | 0.006023871 | 0.14263735 |
| *FAR1* | -2.343563 | 0.853455 | -2.745971 | 0.006033203 | 0.14263735 |
| *NPIPA1* | -2.298791 | 0.837328 | -2.745388 | 0.006043939 | 0.14263735 |
| *F8A1* | -2.492492 | 0.908555 | -2.74336 | 0.006081401 | 0.14263735 |
| *SRF* | -1.888021 | 0.688231 | -2.743297 | 0.006082563 | 0.14263735 |
| *SNAPC4* | -2.298547 | 0.838196 | -2.742256 | 0.00610188 | 0.14263735 |
| *ZSCAN5A* | -2.280244 | 0.831639 | -2.741867 | 0.006109113 | 0.14263735 |
| *BLM* | -1.744419 | 0.636357 | -2.74126 | 0.0061204 | 0.14263735 |
| *FSD1L* | -2.247186 | 0.819804 | -2.741126 | 0.006122902 | 0.14263735 |
| *ALYREF* | -2.185845 | 0.797655 | -2.740339 | 0.006137592 | 0.14265753 |
| *RPL39L* | -2.323903 | 0.850595 | -2.732093 | 0.006293344 | 0.14567942 |
| *PPME1* | -2.188206 | 0.800965 | -2.731962 | 0.006295835 | 0.14567942 |
| *NPIPB2* | -4.028795 | 1.476905 | -2.727864 | 0.006374588 | 0.14717169 |
| *L3MBTL1* | -2.223445 | 0.81611 | -2.724443 | 0.006441013 | 0.14837333 |
| *WASF1* | -2.040136 | 0.749285 | -2.722778 | 0.006473553 | 0.14879079 |
| *NOP14.AS1* | -2.686564 | 0.987455 | -2.720695 | 0.006514484 | 0.14939883 |
| *ENSG00000261732* | -2.055747 | 0.756156 | -2.718681 | 0.006554269 | 0.14943477 |
| *ZBTB49* | -2.661284 | 0.978974 | -2.718442 | 0.006559011 | 0.14943477 |
| *ENSG00000279467* | -3.071414 | 1.129872 | -2.718373 | 0.00656038 | 0.14943477 |
| *LRR1* | -1.769122 | 0.650966 | -2.717688 | 0.006573971 | 0.14943477 |
| *TRUB1* | -2.026609 | 0.746045 | -2.71647 | 0.006598224 | 0.14965642 |
| *MTHFD1L* | -2.904172 | 1.069923 | -2.714376 | NA | NA |

| | | | | | |
|---|---|---|---|---|---|
| *MTFR1* | -1.863289 | 0.686702 | -2.713385 | 0.00665996 | 0.15072541 |
| *LSM14B* | -2.4738 | 0.911996 | -2.712511 | 0.006677557 | 0.15079298 |
| *DGAT1* | -2.526286 | 0.931934 | -2.710799 | 0.006712123 | 0.15106579 |
| *ENSG00000225205* | -2.397089 | 0.884649 | -2.70965 | 0.006735421 | 0.15106579 |
| *TTLL12* | -2.409049 | 0.889468 | -2.708414 | 0.006760553 | 0.15106579 |
| *SREBF2.AS1* | -2.501486 | 0.923816 | -2.707776 | 0.006773569 | 0.15106579 |
| *COG4* | -1.629328 | 0.601728 | -2.707748 | 0.006774135 | 0.15106579 |
| *MCM2* | -2.748416 | 1.015142 | -2.707419 | 0.006780853 | 0.15106579 |
| *SHLD3* | -2.035156 | 0.75186 | -2.706831 | 0.006792888 | 0.15106579 |
| *MTURN* | -2.563438 | 0.947262 | -2.706154 | 0.006806744 | 0.15106579 |
| *CAMKK1* | -2.695607 | 0.997329 | -2.702827 | 0.006875258 | 0.15225892 |
| *HEATR5A* | -2.288362 | 0.847488 | -2.70017 | 0.006930397 | 0.15302533 |
| *CCDC138* | -2.381702 | 0.882199 | -2.699733 | 0.006939521 | 0.15302533 |
| *ENSG00000227540* | -2.384569 | 0.883801 | -2.698083 | 0.006974015 | 0.15309561 |
| *DNMBP* | -2.080635 | 0.771302 | -2.697562 | 0.006984919 | 0.15309561 |
| *SGSM2* | -1.984638 | 0.736037 | -2.696383 | 0.007009696 | 0.15309561 |
| *LINC00294* | -2.494454 | 0.925161 | -2.696238 | 0.007012749 | 0.15309561 |
| *DZANK1* | -3.520174 | 1.305797 | -2.695805 | 0.00702188 | 0.15309561 |
| *SRD5A1* | -1.731641 | 0.642498 | -2.695169 | 0.007035303 | 0.15309561 |
| *SPINT1.AS1* | -3.430892 | 1.273278 | -2.694535 | 0.007048687 | 0.15309561 |
| *CLHC1* | -2.621487 | 0.973107 | -2.693935 | 0.007061387 | 0.15309561 |
| *SCMH1* | -1.694768 | 0.629551 | -2.692026 | 0.00710194 | 0.15330575 |
| *TRAPPC9* | -2.49637 | 0.92742 | -2.691735 | 0.007108137 | 0.15330575 |
| *GAB1* | -2.278267 | 0.846554 | -2.691226 | 0.007118999 | 0.15330575 |
| *RAD52* | -2.464795 | 0.916046 | -2.690687 | 0.0071305 | 0.15330575 |
| *DOCK9* | -2.744302 | 1.020623 | -2.688849 | 0.007169874 | 0.15360348 |
| *TRAF1* | -2.721323 | 1.012151 | -2.688652 | 0.007174116 | 0.15360348 |
| *AAK1* | -2.693208 | 1.002816 | -2.685646 | NA | NA |
| *LIX1.AS1* | -2.340542 | 0.872156 | -2.683627 | 0.007282838 | 0.15560846 |
| *PPP1R13L* | -2.24596 | 0.838338 | -2.679063 | 0.007382851 | 0.15741946 |
| *FAM149B1* | -2.182696 | 0.815005 | -2.678138 | 0.007403263 | 0.15748993 |
| *ICE1* | -1.937014 | 0.723433 | -2.677532 | 0.007416677 | 0.15748993 |
| *FPGT* | -2.575907 | 0.962336 | -2.676724 | 0.007434575 | 0.15754582 |
| *ETFBKMT* | -1.884376 | 0.704913 | -2.673204 | 0.007513051 | 0.15884199 |
| *KCTD21.AS1* | -2.592371 | 0.970225 | -2.671927 | 0.007541715 | 0.15884199 |
| *ZNF337* | -2.293866 | 0.858615 | -2.671589 | 0.007549295 | 0.15884199 |
| *MTRNR2L3* | -4.918176 | 1.841163 | -2.671233 | 0.007557308 | 0.15884199 |
| *TTC31* | -1.555504 | 0.582678 | -2.669575 | 0.007594727 | 0.15910635 |
| *ENSG00000261220* | -3.73044 | 1.398367 | -2.667713 | 0.007636951 | 0.15910635 |
| *MAMSTR* | -2.821283 | 1.057763 | -2.667218 | 0.007648202 | 0.15910635 |
| *ENSG00000277972* | -1.644373 | 0.616584 | -2.666908 | 0.007655272 | 0.15910635 |
| *ZSCAN32* | -1.78644 | 0.669916 | -2.666663 | 0.00766084 | 0.15910635 |
| *ENSG00000255328* | -3.553741 | 1.332823 | -2.666327 | 0.007668507 | 0.15910635 |
| *ENSG00000278390* | -2.153813 | 0.808067 | -2.665389 | 0.007689933 | 0.15910635 |
| *MYH11* | -2.046721 | 0.76793 | -2.665245 | 0.007693224 | 0.15910635 |
| *HSPA13* | -1.850309 | 0.694995 | -2.662335 | 0.007760068 | 0.16002379 |
| *FECH* | -2.132023 | 0.800921 | -2.661965 | 0.007768597 | 0.16002379 |

| | | | | | |
|---|---|---|---|---|---|
| *ENSG00000253645* | -2.337086 | 0.878481 | -2.660373 | 0.007805413 | 0.16046188 |
| *MRTFA* | -2.040788 | 0.767463 | -2.659134 | 0.007834186 | 0.1607332 |
| *ZNF362* | -2.343427 | 0.881425 | -2.65868 | NA | NA |
| *POFUT1* | -2.350643 | 0.884564 | -2.657403 | 0.00787452 | 0.16124018 |
| *MBTPS2* | -2.099855 | 0.790821 | -2.655285 | 0.007924132 | 0.16166952 |
| *ZNF471* | -2.704981 | 1.01876 | -2.655171 | 0.00792682 | 0.16166952 |
| *CEP97* | -1.882584 | 0.709709 | -2.652613 | 0.00798713 | 0.16257827 |
| *AMT* | -2.855463 | 1.077197 | -2.650827 | 0.0080295 | 0.16311898 |
| *LIN37* | -1.996344 | 0.753964 | -2.647796 | 0.008101829 | 0.16426497 |
| *GTSF1* | -2.256642 | 0.852645 | -2.646637 | 0.008129658 | 0.16450602 |
| *NCDN* | -3.228072 | 1.221229 | -2.643299 | 0.00821025 | 0.1658117 |
| *ENSG00000228989* | -2.570034 | 0.972888 | -2.641654 | 0.008250239 | 0.16629388 |
| *COLGALT1* | -1.933286 | 0.731961 | -2.641242 | NA | NA |
| *PLEKHM3* | -4.795651 | 1.816201 | -2.640485 | 0.008278734 | 0.16654296 |
| *CTDP1* | -3.400033 | 1.288487 | -2.638779 | 0.008320524 | 0.16676871 |
| *C9orf72* | -1.796604 | 0.680865 | -2.638707 | 0.008322276 | 0.16676871 |
| *MOV10* | -2.204912 | 0.836027 | -2.637371 | 0.008355148 | 0.16710295 |
| *KIF21B* | -2.49057 | 0.945289 | -2.634718 | 0.008420714 | 0.16808852 |
| *ITGA5* | -3.203737 | 1.217106 | -2.632257 | 0.008481962 | 0.16876272 |
| *FAM83G* | -2.237982 | 0.850498 | -2.63138 | 0.008503893 | 0.16876272 |
| *TAF5L* | -2.272214 | 0.863586 | -2.631138 | 0.008509941 | 0.16876272 |
| *SPTY2D1* | -1.715862 | 0.652235 | -2.630741 | 0.008519901 | 0.16876272 |
| *PIAS4* | -1.81109 | 0.688679 | -2.629803 | 0.008543444 | 0.16881514 |
| *NIPA1* | -2.793187 | 1.06252 | -2.628833 | 0.008567844 | 0.16881514 |
| *SEPT7.AS1* | -2.616331 | 0.995301 | -2.628683 | 0.008571621 | 0.16881514 |
| *TADA2A* | -1.881105 | 0.716564 | -2.625172 | 0.008660524 | 0.17024116 |
| *CASP10* | -2.333965 | 0.889561 | -2.623726 | 0.008697364 | 0.1706403 |
| *SERPINB8* | -2.257115 | 0.861008 | -2.621479 | 0.008754918 | 0.17144355 |
| *QSOX2* | -1.603941 | 0.612526 | -2.618566 | 0.008830029 | 0.17209907 |
| *NINJ2* | -1.870061 | 0.714414 | -2.617616 | 0.008854632 | 0.17209907 |
| *ZNF608* | -1.805078 | 0.689685 | -2.617251 | 0.008864115 | 0.17209907 |
| *ZNF385A* | -1.952736 | 0.746144 | -2.617104 | 0.008867923 | 0.17209907 |
| *HSDL1* | -1.64867 | 0.629995 | -2.616956 | 0.008871774 | 0.17209907 |
| *SLC27A1* | -4.023208 | 1.538169 | -2.615582 | 0.008907557 | 0.17246902 |
| *FBP1* | -1.894477 | 0.724586 | -2.614563 | 0.00893417 | 0.17266037 |
| *C7orf26* | -1.911894 | 0.732477 | -2.610176 | 0.009049575 | 0.17416752 |
| *DUSP3* | -1.927416 | 0.73852 | -2.609837 | 0.009058536 | 0.17416752 |
| *ENSG00000276075* | -2.917671 | 1.118347 | -2.608914 | 0.009083011 | 0.17416752 |
| *HYI* | -1.878512 | 0.720065 | -2.608811 | 0.009085752 | 0.17416752 |
| *SMCO4* | -2.615153 | 1.002763 | -2.607946 | 0.009108722 | 0.17416752 |
| *NPIPB15* | -2.451242 | 0.939976 | -2.60777 | 0.009113417 | 0.17416752 |
| *PIGK* | -1.931095 | 0.741434 | -2.60454 | 0.00919977 | 0.17549284 |
| *TFB1M* | -1.578768 | 0.606892 | -2.6014 | 0.009284422 | 0.17678087 |
| *ENSG00000215014* | -2.801157 | 1.076984 | -2.600928 | NA | NA |
| *NOL6* | -2.489558 | 0.957523 | -2.599999 | 0.009322398 | 0.17702081 |
| *PAXBP1* | -2.089184 | 0.803634 | -2.599671 | 0.00933133 | 0.17702081 |
| *NUP50.DT* | -2.517683 | 0.969059 | -2.598069 | 0.009374954 | 0.17752207 |

| | | | | | |
|---|---|---|---|---|---|
| *GALC* | -2.280506 | 0.878197 | -2.596805 | 0.009409522 | 0.1778503 |
| *GINS4* | -1.839745 | 0.709068 | -2.594597 | 0.009470185 | 0.17839939 |
| *CRAMP1* | -2.056051 | 0.792517 | -2.594331 | 0.009477523 | 0.17839939 |
| *CNPY4* | -2.443577 | 0.942061 | -2.593863 | 0.009490433 | 0.17839939 |
| *IGHG2* | -2.737026 | 1.056464 | -2.590742 | 0.009576937 | 0.17969817 |
| *MAPRE3* | -3.143797 | 1.213942 | -2.589743 | 0.009604761 | 0.17973139 |
| *MATK* | -4.65357 | 1.797142 | -2.589428 | 0.009613539 | 0.17973139 |
| *ZNF496* | -1.889087 | 0.730766 | -2.585078 | 0.009735702 | 0.1814126 |
| *ENSG00000268858* | -2.298437 | 0.889278 | -2.584611 | 0.009748886 | 0.1814126 |
| *SLC20A2* | -1.7796 | 0.688606 | -2.584353 | 0.009756201 | 0.1814126 |
| *FCRLB* | -2.67651 | 1.036297 | -2.582762 | 0.00980128 | 0.18192303 |
| *IPPK* | -2.300024 | 0.892202 | -2.577918 | 0.009939745 | 0.18416189 |
| *ARHGAP31* | -5.473319 | 2.124305 | -2.576522 | 0.009979987 | 0.18457611 |
| *GINS1* | -4.632642 | 1.799777 | -2.574008 | 0.010052784 | 0.18558986 |
| *WEE1* | -2.260219 | 0.878344 | -2.573273 | 0.01007417 | 0.18565256 |
| *SLC25A23* | -2.617093 | 1.017694 | -2.571592 | 0.010123214 | 0.18590113 |
| *ENSG00000273064* | -2.861206 | 1.112627 | -2.571576 | 0.010123685 | 0.18590113 |
| *FDXACB1* | -2.443004 | 0.950896 | -2.569159 | 0.010194571 | 0.18687028 |
| *MAN1B1.DT* | -2.501584 | 0.974378 | -2.567366 | 0.010247445 | 0.18750644 |
| *TSPAN5* | -2.188559 | 0.853481 | -2.564273 | 0.010339215 | 0.18885079 |
| *KLHL9* | -1.914675 | 0.746871 | -2.563595 | 0.010359424 | 0.18888561 |
| *FURIN* | -1.778086 | 0.694466 | -2.560364 | 0.010456249 | 0.19031479 |
| *PUS10* | -1.905087 | 0.74482 | -2.55778 | 0.010534263 | 0.19119454 |
| *ACTL6A* | -1.569523 | 0.613685 | -2.557537 | 0.010541637 | 0.19119454 |
| *ZHX1* | -2.380432 | 0.932265 | -2.553385 | 0.010668152 | 0.19296921 |
| *ATAD3B* | -1.876136 | 0.734846 | -2.5531 | 0.010676882 | 0.19296921 |
| *GZF1* | -2.0579 | 0.807057 | -2.549881 | 0.010775977 | 0.19441973 |
| *ENSG00000071655* | -1.698338 | 0.666368 | -2.548647 | 0.010814158 | 0.19476808 |
| *HEATR3* | -2.168558 | 0.851391 | -2.547076 | 0.010862974 | 0.19530644 |
| *FYN* | -1.962515 | 0.770593 | -2.546758 | NA | NA |
| *CROT* | -2.168574 | 0.852307 | -2.544357 | 0.010947919 | 0.19649134 |
| *FLNB* | -2.112013 | 0.83083 | -2.542052 | 0.011020364 | 0.19744819 |
| *N6AMT1* | -1.982374 | 0.780658 | -2.539364 | 0.011105437 | 0.19840002 |
| *APEX2* | -1.534189 | 0.604461 | -2.538109 | 0.011145314 | 0.19840002 |
| *ZBED2* | -2.400715 | 0.945896 | -2.538034 | 0.011147717 | 0.19840002 |
| *TAF4* | -2.005696 | 0.790404 | -2.537558 | 0.011162899 | 0.19840002 |
| *FAM220A* | -2.216501 | 0.873681 | -2.536969 | 0.011181685 | 0.19840002 |
| *ZNF781* | -1.894908 | 0.747143 | -2.536204 | 0.011206152 | 0.19840002 |
| *FAM135A* | -1.773748 | 0.699388 | -2.536144 | 0.011208063 | 0.19840002 |
| *DOCK6* | -3.812249 | 1.50465 | -2.533645 | 0.011288315 | 0.19947844 |
| *UBE2Q2* | -1.561904 | 0.617083 | -2.531106 | 0.011370338 | 0.20058442 |
| *GALNT7* | -1.857158 | 0.734144 | -2.529692 | 0.011416255 | 0.20081972 |
| *ENSG00000232010* | -3.10423 | 1.227212 | -2.529498 | 0.011422595 | 0.20081972 |
| *GNGT2* | -2.233776 | 0.88341 | -2.528584 | 0.011452376 | 0.20086926 |
| *DMWD* | -2.014541 | 0.796823 | -2.528217 | 0.011464341 | 0.20086926 |
| *LINC00494* | -1.821989 | 0.720945 | -2.527223 | 0.011496844 | 0.20098192 |
| *SMARCD2* | -1.601646 | 0.633904 | -2.52664 | 0.011515943 | 0.20098192 |

| | | | | | |
|---|---|---|---|---|---|
| *GGACT* | -4.514342 | 1.787396 | -2.525654 | 0.011548332 | 0.20098192 |
| *ENSG00000238045* | -2.096237 | 0.830126 | -2.525202 | 0.011563181 | 0.20098192 |
| *TANC2* | -2.841946 | 1.1255 | -2.525051 | 0.011568145 | 0.20098192 |
| *LIN7B* | -1.949334 | 0.772718 | -2.522699 | 0.011645816 | 0.20181863 |
| *ZNF236* | -1.566845 | 0.621181 | -2.522364 | 0.011656913 | 0.20181863 |
| *TTC33* | -1.627234 | 0.64568 | -2.520185 | 0.01172931 | 0.20181863 |
| *PRR7* | -2.176313 | 0.86357 | -2.520135 | 0.011730997 | 0.20181863 |
| *ENSG00000223473* | -2.254932 | 0.894986 | -2.519518 | 0.011751552 | 0.20181863 |
| *TSBP1.AS1* | -2.795586 | 1.109572 | -2.519517 | 0.011751592 | 0.20181863 |
| *MPZL1* | -1.653972 | 0.656477 | -2.519469 | 0.011753198 | 0.20181863 |
| *MOXD1* | -4.361216 | 1.732506 | -2.517287 | 0.011826248 | 0.20251187 |
| *PCNT* | -1.894491 | 0.752651 | -2.517091 | 0.011832816 | 0.20251187 |
| *PDCL* | -1.439794 | 0.572329 | -2.515676 | 0.01188042 | 0.20276594 |
| *NAIF1* | -1.941734 | 0.771913 | -2.515483 | 0.011886957 | 0.20276594 |
| *EIF3C* | -1.70349 | 0.677532 | -2.514256 | 0.011928395 | 0.20292589 |
| *URB1.AS1* | -2.449908 | 0.974656 | -2.513613 | NA | NA |
| *ELMO2* | -1.923168 | 0.765161 | -2.513417 | 0.011956803 | 0.20292589 |
| *ENSG00000271109* | -1.783643 | 0.709671 | -2.513339 | 0.011959448 | 0.20292589 |
| *PHYH* | -2.323496 | 0.924635 | -2.512881 | 0.011974987 | 0.20292589 |
| *PSTK* | -2.12655 | 0.84676 | -2.511395 | 0.012025502 | 0.20344784 |
| *WDYHV1* | -2.063101 | 0.822066 | -2.509654 | 0.012084956 | 0.20366749 |
| *KLHDC7B* | -2.237573 | 0.891833 | -2.508959 | 0.012108768 | 0.20366749 |
| *HS3ST3B1* | -2.290317 | 0.912981 | -2.508613 | 0.012120605 | 0.20366749 |
| *ZNF280B* | -2.883679 | 1.149621 | -2.508372 | 0.012128875 | 0.20366749 |
| *TCEANC* | -1.728631 | 0.689332 | -2.50769 | 0.012152315 | 0.20366749 |
| *IFIT5* | -1.539161 | 0.61415 | -2.506166 | 0.012204817 | 0.20366749 |
| *ENSG00000268713* | -2.998346 | 1.196655 | -2.505605 | 0.012224201 | 0.20366749 |
| *RAPGEF6* | -1.515322 | 0.60479 | -2.505533 | 0.012226707 | 0.20366749 |
| *IRGQ* | -2.199758 | 0.878111 | -2.505102 | 0.012241602 | 0.20366749 |
| *ZNF354B* | -1.87303 | 0.748365 | -2.50283 | 0.012320487 | 0.20366749 |
| *RCBTB2* | -4.605208 | 1.840134 | -2.502648 | 0.012326815 | 0.20366749 |
| *ATXN7L1* | -1.811107 | 0.723706 | -2.502545 | 0.012330382 | 0.20366749 |
| *WHAMM* | -1.458332 | 0.582757 | -2.502471 | 0.012332967 | 0.20366749 |
| *TBXA2R* | -2.312759 | 0.924348 | -2.502043 | 0.012347881 | 0.20366749 |
| *JADE3* | -2.17026 | 0.867535 | -2.50164 | 0.012361954 | 0.20366749 |
| *STK11IP* | -1.717696 | 0.686698 | -2.501385 | 0.012370847 | 0.20366749 |
| *LINC02576* | -2.669145 | 1.067337 | -2.500751 | 0.012393014 | 0.20366749 |
| *PCNX3* | -2.079896 | 0.831715 | -2.500731 | 0.012393719 | 0.20366749 |
| *SAP130* | -1.88591 | 0.754767 | -2.498666 | 0.012466172 | 0.20426854 |
| *FASTKD1* | -2.161588 | 0.865133 | -2.498561 | 0.012469882 | 0.20426854 |
| *FAM241A* | -2.097998 | 0.839739 | -2.498391 | NA | NA |
| *ABHD16A* | -3.409827 | 1.365392 | -2.497325 | 0.012513421 | 0.20461067 |
| *HIVEP1* | -2.219667 | 0.888989 | -2.496844 | 0.012530421 | 0.20461067 |
| *ENSG00000278743* | -1.639579 | 0.657055 | -2.495345 | 0.012583454 | 0.20487526 |
| *ENSG00000226571* | -2.435371 | 0.975997 | -2.495264 | 0.012586329 | 0.20487526 |
| *ENSG00000258056* | -2.588499 | 1.037666 | -2.494538 | 0.012612111 | 0.20490112 |
| *ACTR8* | -1.778061 | 0.713098 | -2.493431 | 0.012651505 | 0.20490112 |

| | | | | | |
|---|---|---|---|---|---|
| *CHKA* | -3.680544 | 1.476154 | -2.493334 | 0.012654979 | 0.20490112 |
| *UNG* | -1.82293 | 0.731223 | -2.492987 | 0.012667336 | 0.20490112 |
| *BRPF3* | -2.427101 | 0.974534 | -2.490525 | 0.012755463 | 0.20600372 |
| *ARL10* | -1.894874 | 0.762117 | -2.486329 | 0.01290685 | 0.20812295 |
| *BCAS3* | -1.895297 | 0.762871 | -2.484427 | 0.012976027 | 0.20868363 |
| *KDM1A* | -1.54447 | 0.62186 | -2.483631 | 0.013005038 | 0.20868363 |
| *DRAM1* | -2.638802 | 1.062911 | -2.482618 | 0.013042075 | 0.20868363 |
| *ENSG00000257303* | -1.928375 | 0.777204 | -2.481171 | 0.013095162 | 0.20868363 |
| *ENSG00000272189* | -2.83682 | 1.143493 | -2.480838 | 0.013107398 | 0.20868363 |
| *GOLGA1* | -2.195272 | 0.884902 | -2.480808 | 0.013108485 | 0.20868363 |
| *CD27* | -3.194143 | 1.287634 | -2.480628 | 0.013115098 | 0.20868363 |
| *PUDP* | -2.222122 | 0.896008 | -2.480026 | 0.013137281 | 0.20868363 |
| *PTPDC1* | -2.223415 | 0.896549 | -2.479972 | 0.013139273 | 0.20868363 |
| *TMEM169* | -2.674933 | 1.07871 | -2.479752 | 0.013147366 | 0.20868363 |
| *ENSG00000237596* | -2.269146 | 0.915482 | -2.478636 | 0.013188582 | 0.20868363 |
| *ENSG00000267058* | -2.09257 | 0.844285 | -2.478513 | 0.01319312 | 0.20868363 |
| *RAPGEF2* | -1.986694 | 0.801666 | -2.478206 | 0.013204497 | 0.20868363 |
| *ZSWIM1* | -1.965688 | 0.793461 | -2.47736 | 0.013235846 | 0.20885922 |
| *TMEM8A* | -1.939871 | 0.783548 | -2.475751 | 0.013295641 | 0.20948247 |
| *ARRDC3.AS1* | -2.934488 | 1.185911 | -2.474459 | 0.013343802 | 0.20978084 |
| *ANK1* | -2.104216 | 0.850479 | -2.474153 | 0.013355234 | 0.20978084 |
| *SSX2IP* | -2.449766 | 0.990676 | -2.472822 | 0.01340509 | 0.21024396 |
| *ARNT* | -1.644841 | 0.6663 | -2.46862 | 0.013563525 | 0.21240604 |
| *CD24* | -1.189637 | 0.482108 | -2.467572 | 0.013603283 | 0.21270589 |
| *COL4A3* | -5.203833 | 2.110128 | -2.466121 | 0.013658503 | 0.21299641 |
| *RTN4IP1* | -2.124972 | 0.861873 | -2.465527 | 0.013681179 | 0.21299641 |
| *ENSG00000272918* | -1.958428 | 0.794346 | -2.465459 | 0.013683781 | 0.21299641 |
| *ATG4A* | -2.297916 | 0.933399 | -2.46188 | 0.013821085 | 0.2145592 |
| *PELI2* | -2.004812 | 0.814501 | -2.461399 | 0.013839652 | 0.2145592 |
| *RNF144B* | -1.72035 | 0.698983 | -2.46122 | 0.013846553 | 0.2145592 |
| *MTA3* | -1.840617 | 0.748528 | -2.458982 | 0.013933144 | 0.21538984 |
| *GCH1* | -1.733149 | 0.705176 | -2.457751 | 0.013980989 | 0.21538984 |
| *RPS6KA2* | -1.829861 | 0.74453 | -2.457739 | 0.013981486 | 0.21538984 |
| *S100A13* | -2.052986 | 0.835334 | -2.457683 | 0.013983642 | 0.21538984 |
| *CARMIL2* | -1.681087 | 0.684188 | -2.457053 | 0.014008185 | 0.2154463 |
| *LIX1L.AS1* | -3.280673 | 1.335683 | -2.456177 | 0.014042416 | 0.21560464 |
| *MARCHF8* | -1.529544 | 0.62285 | -2.45572 | 0.014060264 | 0.21560464 |
| *ENSG00000272927* | -1.917071 | 0.781393 | -2.453402 | 0.014151226 | 0.21667753 |
| *HIST1H3E* | -1.866497 | 0.762011 | -2.449436 | 0.014308004 | 0.21866242 |
| *CDK7* | -1.787812 | 0.730001 | -2.449053 | 0.014323236 | 0.21866242 |
| *ENSG00000279080* | -1.741648 | 0.711592 | -2.447537 | 0.014383619 | 0.21925989 |
| *PIGW* | -1.885263 | 0.771045 | -2.445076 | 0.014482165 | 0.2204365 |
| *PRKCZ* | -2.697087 | 1.103885 | -2.443269 | 0.014554903 | 0.22083485 |
| *KLK1* | -2.27048 | 0.929429 | -2.442876 | 0.014570732 | 0.22083485 |
| *ZNF446* | -1.821065 | 0.745473 | -2.442832 | 0.014572532 | 0.22083485 |
| *NBPF19* | -1.713519 | 0.702055 | -2.440719 | 0.014658072 | 0.22148442 |
| *ZNF417* | -1.910365 | 0.782726 | -2.440656 | 0.014660623 | 0.22148442 |

| | | | | | |
|---|---|---|---|---|---|
| *MIF.AS1* | -2.51313 | 1.029894 | -2.440184 | 0.014679781 | 0.22148442 |
| *PEX3* | -2.027194 | 0.831965 | -2.436634 | 0.014824685 | 0.22259626 |
| *ARFGAP1* | -1.655582 | 0.679613 | -2.436065 | 0.014848011 | 0.22259626 |
| *ZNF529.AS1* | -2.102203 | 0.863134 | -2.435548 | 0.014869239 | 0.22259626 |
| *CEP128* | -2.002381 | 0.822207 | -2.435372 | 0.014876468 | 0.22259626 |
| *BAP1* | -1.470285 | 0.603756 | -2.43523 | 0.014882316 | 0.22259626 |
| *PLCB3* | -2.817153 | 1.156839 | -2.435216 | 0.014882889 | 0.22259626 |
| *USP12* | -1.741953 | 0.715648 | -2.434092 | 0.014929218 | 0.22277034 |
| *NAB1* | -2.276905 | 0.935502 | -2.433886 | 0.014937701 | 0.22277034 |
| *ZNF771* | -2.635878 | 1.08335 | -2.43308 | 0.014970977 | 0.22294442 |
| *GPR107* | -3.712455 | 1.526613 | -2.431824 | NA | NA |
| *MCU* | -2.469967 | 1.016825 | -2.429098 | 0.015136444 | 0.22424033 |
| *RNF103* | -1.828102 | 0.752684 | -2.428776 | 0.015149869 | 0.22424033 |
| *HELB* | -1.958322 | 0.806314 | -2.428735 | 0.01515159 | 0.22424033 |
| *ZNF384* | -1.501105 | 0.618114 | -2.428524 | 0.015160421 | 0.22424033 |
| *SLC35B4* | -1.923403 | 0.792054 | -2.428375 | 0.015166643 | 0.22424033 |
| *TMEM273* | -2.128324 | 0.877134 | -2.426453 | 0.015247226 | 0.22451229 |
| *C19orf47* | -2.419183 | 0.997214 | -2.425942 | 0.015268694 | 0.22451229 |
| *TDRD7* | -1.972215 | 0.81299 | -2.425878 | 0.015271419 | 0.22451229 |
| *DGKQ* | -1.768343 | 0.729157 | -2.425189 | 0.015300402 | 0.22451229 |
| *CHL1* | -2.650204 | 1.093021 | -2.424659 | 0.015322756 | 0.22451229 |
| *PABPC1L* | -2.140863 | 0.883033 | -2.424443 | 0.015331899 | 0.22451229 |
| *ALMS1* | -2.135024 | 0.880685 | -2.424276 | 0.015338932 | 0.22451229 |
| *SORBS3* | -2.155996 | 0.889556 | -2.423677 | 0.015364245 | 0.22451229 |
| *CCDC51* | -2.21927 | 0.916169 | -2.422337 | 0.015421047 | 0.22451229 |
| *SELENOO* | -1.711069 | 0.70662 | -2.421482 | 0.01545737 | 0.22451229 |
| *AHCTF1* | -1.54466 | 0.638022 | -2.421015 | 0.015477242 | 0.22451229 |
| *PLPP6* | -2.124081 | 0.87737 | -2.420964 | 0.015479421 | 0.22451229 |
| *PCBP4* | -3.105855 | 1.282923 | -2.42092 | 0.015481264 | 0.22451229 |
| *FOXK2* | -2.055671 | 0.849347 | -2.420295 | 0.01550792 | 0.22451229 |
| *SLX1A* | -3.185506 | 1.316835 | -2.419062 | 0.015560581 | 0.22451229 |
| *MATR3* | -2.855896 | 1.180748 | -2.418718 | 0.015575319 | 0.22451229 |
| *MMP28* | -2.909925 | 1.203131 | -2.418626 | 0.015579233 | 0.22451229 |
| *LMAN2L* | -2.013953 | 0.832783 | -2.418341 | 0.01559147 | 0.22451229 |
| *STXBP5* | -2.931519 | 1.212444 | -2.417858 | 0.015612156 | 0.22451229 |
| *FAM185A* | -2.744593 | 1.135221 | -2.417672 | 0.015620138 | 0.22451229 |
| *SERTAD1* | -1.447397 | 0.598911 | -2.416713 | 0.015661352 | 0.22479159 |
| *ADRB2* | -2.751721 | 1.139468 | -2.414919 | 0.01573873 | 0.22519122 |
| *ZHX1.C8orf76* | -3.227096 | 1.336349 | -2.41486 | 0.015741241 | 0.22519122 |
| *CR1* | -1.729495 | 0.716413 | -2.414103 | 0.015773993 | 0.22519122 |
| *MBLAC2* | -2.347232 | 0.972956 | -2.412475 | 0.01584463 | 0.22519122 |
| *STAM2* | -1.959554 | 0.812484 | -2.411807 | 0.015873676 | 0.22519122 |
| *ENSG00000225342* | -1.854898 | 0.769163 | -2.411578 | 0.015883635 | 0.22519122 |
| *BIK* | -3.006029 | 1.246502 | -2.411572 | 0.015883929 | 0.22519122 |
| *ANKHD1* | -1.432713 | 0.59411 | -2.411529 | 0.015885797 | 0.22519122 |
| *LTBP4* | -1.381068 | 0.572757 | -2.411262 | 0.015897419 | 0.22519122 |
| *MACC1* | -3.40061 | 1.410437 | -2.411033 | 0.015907403 | 0.22519122 |

| | | | | | |
|---|---|---|---|---|---|
| *RTP4* | -2.126502 | 0.882303 | -2.410173 | NA | NA |
| *MSANTD2* | -2.852623 | 1.184756 | -2.407773 | 0.01605016 | 0.22673282 |
| *ANAPC1* | -1.623705 | 0.674424 | -2.407544 | 0.016060241 | 0.22673282 |
| *KIAA0232* | -1.607981 | 0.668195 | -2.406454 | 0.016108214 | 0.22709941 |
| *SERPINF1* | -1.723489 | 0.716769 | -2.404523 | 0.016193576 | 0.22799142 |
| *MOB3C* | -2.628118 | 1.093568 | -2.403251 | 0.01625005 | 0.22847482 |
| *LINC00886* | -2.163194 | 0.900978 | -2.400941 | 0.016352971 | 0.22950715 |
| *DRC3* | -2.559104 | 1.066149 | -2.400325 | 0.016380533 | 0.22950715 |
| *GSE1* | -1.63285 | 0.680583 | -2.399193 | 0.016431245 | 0.22950715 |
| *NKX3.1* | -3.656202 | 1.523961 | -2.399143 | 0.016433474 | 0.22950715 |
| *GNPNAT1* | -2.245802 | 0.936095 | -2.399117 | 0.016434669 | 0.22950715 |
| *PSMG3.AS1* | -2.17769 | 0.90809 | -2.3981 | 0.016480384 | 0.22983454 |
| *ENSG00000273002* | -2.435681 | 1.016138 | -2.396998 | 0.016530014 | 0.23021558 |
| *ZBP1* | -2.218969 | 0.925923 | -2.396495 | 0.01655274 | 0.2302214 |
| *RABGEF1* | -2.000525 | 0.835192 | -2.395288 | 0.016607295 | 0.23053622 |
| *LMF1* | -1.802932 | 0.753027 | -2.394247 | 0.016654549 | 0.23053622 |
| *WASIR2* | -2.891518 | 1.207838 | -2.393962 | 0.016667483 | 0.23053622 |
| *DTWD2* | -1.918991 | 0.801857 | -2.393184 | 0.016702848 | 0.23053622 |
| *CCDC61* | -2.915189 | 1.218201 | -2.393029 | 0.01670994 | 0.23053622 |
| *CCHCR1* | -1.538993 | 0.643136 | -2.39295 | 0.01671354 | 0.23053622 |
| *MINDY3* | -1.987659 | 0.83077 | -2.39255 | 0.016731747 | 0.23053622 |
| *FGD4* | -4.884804 | 2.042666 | -2.391386 | 0.016784884 | 0.23077756 |
| *PELP1* | -1.658492 | 0.693585 | -2.391187 | 0.016793987 | 0.23077756 |
| *EMILIN2* | -3.497076 | 1.463615 | -2.389342 | 0.016878593 | 0.23163176 |
| *PATZ1* | -1.647119 | 0.689994 | -2.387148 | 0.01697965 | 0.23222317 |
| *OGDH* | -1.533903 | 0.642763 | -2.386421 | 0.017013245 | 0.23222317 |
| *FTO* | -1.990057 | 0.833932 | -2.386355 | 0.017016308 | 0.23222317 |
| *METTL6* | -1.637201 | 0.686146 | -2.386085 | 0.017028833 | 0.23222317 |
| *SPIN2B* | -1.457851 | 0.61101 | -2.385969 | 0.0170342 | 0.23222317 |
| *PGAP3* | -3.038267 | 1.274235 | -2.384385 | 0.017107704 | 0.23230711 |
| *MYO1B* | -2.911304 | 1.221236 | -2.383899 | 0.017130285 | 0.23230711 |
| *SMOX* | -2.622031 | 1.100072 | -2.383508 | 0.017148525 | 0.23230711 |
| *ZNF212* | -2.134317 | 0.895997 | -2.382058 | 0.017216196 | 0.23230711 |
| *FAF1* | -1.733296 | 0.727675 | -2.381966 | 0.017220491 | 0.23230711 |
| *KIAA0556* | -2.216512 | 0.930755 | -2.381413 | 0.017246372 | 0.23230711 |
| *HMGCS1* | -1.398991 | 0.587601 | -2.380853 | 0.017272606 | 0.23230711 |
| *STX3* | -1.660846 | 0.697587 | -2.380843 | 0.017273053 | 0.23230711 |
| *DHRS4* | -2.077705 | 0.872734 | -2.380685 | 0.017280489 | 0.23230711 |
| *TNKS* | -2.13455 | 0.896616 | -2.380673 | 0.017281052 | 0.23230711 |
| *MCAT* | -1.659715 | 0.697205 | -2.380525 | 0.017287971 | 0.23230711 |
| *CXorf65* | -1.776365 | 0.746579 | -2.379339 | 0.017343727 | 0.23261491 |
| *BRF1* | -2.693167 | 1.132148 | -2.378811 | 0.017368558 | 0.23261491 |
| *ZNF34* | -2.014913 | 0.8471 | -2.378601 | 0.017378498 | 0.23261491 |
| *ZNF564* | -1.866351 | 0.785267 | -2.376709 | 0.017467885 | 0.23326115 |
| *ULK3* | -1.468249 | 0.617788 | -2.376622 | 0.017471983 | 0.23326115 |
| *CBFA2T3* | -1.680113 | 0.707078 | -2.376134 | 0.017495116 | 0.23326821 |
| *PTGIR* | -3.712148 | 1.563495 | -2.374262 | 0.017584064 | 0.23415166 |

| | | | | | |
|---|---|---|---|---|---|
| *TESK2* | -1.511072 | 0.636742 | -2.373129 | 0.017638083 | 0.23432106 |
| *C19orf73* | -3.479416 | 1.466491 | -2.372613 | 0.017662737 | 0.23432106 |
| *BOLA2.SMG1P6* | -1.757713 | 0.740912 | -2.372362 | 0.017674751 | 0.23432106 |
| *DHX34* | -1.988305 | 0.838207 | -2.372094 | 0.017687607 | 0.23432106 |
| *FKBP14* | -3.356703 | 1.417474 | -2.368088 | 0.017880288 | 0.23641031 |
| *DENND6B* | 1.513204 | 0.639059 | 2.367864 | 0.017891129 | 0.23641031 |
| *LCMT2* | -3.051336 | 1.289078 | -2.367068 | 0.017929634 | 0.23661615 |
| *GTPBP2* | -2.084459 | 0.880977 | -2.366077 | 0.017977681 | 0.23694722 |
| *SEMA7A* | -2.154315 | 0.910846 | -2.36518 | 0.018021289 | 0.23721901 |
| *WDR7* | -2.522081 | 1.066609 | -2.36458 | 0.018050525 | 0.23730118 |
| *ENSG00000218018* | -2.224224 | 0.941711 | -2.361898 | 0.018181667 | 0.23872112 |
| *RPS19* | 0.17398 | 0.07369 | 2.360974 | 0.018226997 | 0.23880129 |
| *PLAG1* | -2.734874 | 1.158437 | -2.360831 | 0.018234052 | 0.23880129 |
| *TMEM177* | -2.723671 | 1.154131 | -2.359931 | 0.01827832 | 0.2388704 |
| *F11R* | -2.632141 | 1.115417 | -2.359783 | 0.018285622 | 0.2388704 |
| *KCTD9* | -1.975329 | 0.837472 | -2.35868 | 0.018340058 | 0.23927863 |
| *NAGLU* | -1.922693 | 0.815511 | -2.357655 | 0.018390782 | 0.23963746 |
| *AGTPBP1* | -2.019493 | 0.856897 | -2.356751 | 0.018435583 | 0.23984085 |
| *PIGG* | -1.491115 | 0.632793 | -2.356403 | 0.018452872 | 0.23984085 |
| *SLFN5* | -2.673421 | 1.134932 | -2.355577 | 0.018493969 | 0.23985811 |
| *SLC37A4* | -2.377225 | 1.009248 | -2.355442 | 0.018500684 | 0.23985811 |
| *NEK3* | -2.492585 | 1.059064 | -2.353573 | 0.018593972 | 0.24030449 |
| *CARNS1* | -3.4914 | 1.483452 | -2.353564 | 0.018594382 | 0.24030449 |
| *ENSG00000236935* | -1.577866 | 0.670476 | -2.353353 | 0.01860497 | 0.24030449 |
| *PLEKHG2* | -2.586105 | 1.099359 | -2.352375 | 0.018653952 | 0.24063598 |
| *VPS8* | -1.410492 | 0.599863 | -2.351357 | 0.01870507 | 0.24099415 |
| *RIPOR3* | -2.104461 | 0.895447 | -2.350178 | 0.018764431 | 0.24138612 |
| *ENSG00000263264* | -1.594208 | 0.678437 | -2.349824 | 0.018782273 | 0.24138612 |
| *SSBP4* | -1.22502 | 0.521427 | -2.349358 | 0.018805801 | 0.2413879 |
| *FAM229A* | -1.976707 | 0.841769 | -2.348277 | 0.018860482 | 0.24178903 |
| *ZNF865* | -1.871947 | 0.797802 | -2.346379 | 0.0189568 | 0.24261771 |
| *TAS2R4* | -2.340071 | 0.997439 | -2.346078 | 0.018972141 | 0.24261771 |
| *SNX15* | -3.184988 | 1.358031 | -2.345298 | 0.019011867 | 0.24282483 |
| *MTMR1* | -1.559702 | 0.665346 | -2.344195 | 0.019068213 | 0.24312289 |
| *ENSG00000261505* | -2.286493 | 0.9755 | -2.343919 | 0.01908232 | 0.24312289 |
| *ENSG00000253948* | -2.035863 | 0.869188 | -2.342258 | 0.019167443 | 0.2434254 |
| *MX2* | -1.951512 | 0.833233 | -2.342097 | 0.019175753 | 0.2434254 |
| *DAG1* | -2.787333 | 1.190112 | -2.342076 | 0.019176827 | 0.2434254 |
| *BACH1.AS1* | -2.283163 | 0.975386 | -2.340778 | 0.019243598 | 0.24375585 |
| *RAI1* | -1.992998 | 0.851471 | -2.340652 | 0.019250098 | 0.24375585 |
| *TTC39C* | -1.551198 | 0.66294 | -2.339878 | 0.019290034 | 0.24392747 |
| *REPS1* | -2.057805 | 0.879623 | -2.339417 | 0.019313872 | 0.24392747 |
| *PMM2* | -2.03289 | 0.869307 | -2.338518 | 0.0193604 | 0.24392747 |
| *RGCC* | -2.508969 | 1.07307 | -2.338123 | 0.019380877 | 0.24392747 |
| *FAM120B* | -1.7426 | 0.745617 | -2.337125 | 0.019432691 | 0.24392747 |
| *MYEF2* | -1.570133 | 0.671891 | -2.336885 | 0.019445141 | 0.24392747 |
| *SEC16A* | -1.937716 | 0.829269 | -2.336657 | 0.019457041 | 0.24392747 |

| | | | | | |
|---|---|---|---|---|---|
| *ZNF436* | -3.383804 | 1.448985 | -2.335292 | 0.019528146 | 0.24392747 |
| *IGKV1.12* | -1.928392 | 0.825831 | -2.335093 | 0.01953857 | 0.24392747 |
| *RFX7* | -1.683136 | 0.720855 | -2.334915 | 0.019547864 | 0.24392747 |
| *PXN.AS1* | -1.763744 | 0.75551 | -2.334506 | 0.01956924 | 0.24392747 |
| *SERGEF* | -1.422645 | 0.609448 | -2.334319 | 0.019579034 | 0.24392747 |
| *LDB1* | -1.927114 | 0.825562 | -2.334305 | 0.01957974 | 0.24392747 |
| *OGT* | -1.383662 | 0.59309 | -2.332972 | 0.019649592 | 0.24392747 |
| *ZNF585A* | -1.626782 | 0.697333 | -2.332864 | 0.019655307 | 0.24392747 |
| *RYBP* | -1.747275 | 0.748988 | -2.332849 | 0.019656072 | 0.24392747 |
| *SNX19* | -2.181562 | 0.935254 | -2.332587 | 0.019669833 | 0.24392747 |
| *CRY2* | -1.604478 | 0.688023 | -2.332013 | 0.019700025 | 0.24392747 |
| *ZNF577* | -1.727649 | 0.740944 | -2.331688 | 0.0197171 | 0.24392747 |
| *WWP1* | -1.661864 | 0.712885 | -2.33118 | 0.019743858 | 0.24392747 |
| *SDCBP2.AS1* | -2.023367 | 0.868072 | -2.330874 | 0.019760016 | 0.24392747 |
| *CD3EAP* | -1.692723 | 0.72662 | -2.329584 | 0.01982813 | 0.24447587 |
| *ENSG00000259321* | -1.599098 | 0.686708 | -2.328645 | 0.019877898 | 0.24479703 |
| *MREG* | -3.045476 | 1.308774 | -2.326968 | 0.019966958 | 0.24540102 |
| *NFYA* | -1.621571 | 0.696931 | -2.326731 | 0.019979584 | 0.24540102 |
| *PPFIBP1* | -1.797049 | 0.772594 | -2.325993 | 0.020018926 | 0.24540102 |
| *NBPF15* | -2.075173 | 0.892295 | -2.325657 | 0.020036844 | 0.24540102 |
| *TATDN2* | -1.824585 | 0.784883 | -2.32466 | 0.020090168 | 0.24540102 |
| *LAMA5* | -2.418766 | 1.040482 | -2.324659 | 0.020090216 | 0.24540102 |
| *VPS26B* | -2.794443 | 1.202118 | -2.324599 | 0.020093398 | 0.24540102 |
| *MOSPD2* | -1.820675 | 0.783412 | -2.324033 | 0.020123703 | 0.24540464 |
| *TANGO2* | -1.438519 | 0.619081 | -2.323638 | 0.020144924 | 0.24540464 |
| *VILL* | -1.554429 | 0.669186 | -2.322865 | 0.020186388 | 0.24540464 |
| *HIST2H2BE* | -1.756682 | 0.756271 | -2.32282 | 0.020188812 | 0.24540464 |
| *TRAPPC11* | -1.6031 | 0.690385 | -2.322038 | 0.020230884 | 0.24562674 |
| *GABARAPL1* | -1.844282 | 0.794926 | -2.320069 | 0.02033717 | 0.24650471 |
| *RAP2A* | -2.882886 | 1.242934 | -2.31942 | 0.020372272 | 0.24650471 |
| *ENSG00000269514* | -2.066247 | 0.890865 | -2.319372 | 0.020374856 | 0.24650471 |
| *ITPK1* | -1.682742 | 0.725741 | -2.318653 | 0.020413834 | 0.24668709 |
| *HABP4* | -2.032453 | 0.876765 | -2.318128 | 0.020442349 | 0.24674274 |
| *ENSG00000261353* | -2.935641 | 1.267114 | -2.316793 | 0.020515008 | 0.24707046 |
| *MORC2* | -2.219768 | 0.958222 | -2.316549 | 0.020528294 | 0.24707046 |
| *ENSG00000273437* | -1.752501 | 0.756765 | -2.31578 | 0.020570258 | 0.24707046 |
| *DPYSL2* | -1.793734 | 0.774655 | -2.315525 | 0.020584207 | 0.24707046 |
| *ANKFY1* | -1.507585 | 0.651103 | -2.315434 | 0.020589205 | 0.24707046 |
| *ENSG00000242861* | -1.948453 | 0.841704 | -2.314892 | 0.020618863 | 0.24713899 |
| *ZCCHC4* | -2.33713 | 1.009631 | -2.314837 | NA | NA |
| *CCDC126* | -1.691411 | 0.730799 | -2.314468 | NA | NA |
| *GATD3A* | -3.076753 | 1.329537 | -2.314154 | 0.020659287 | 0.24733624 |
| *IFT27* | -1.543376 | 0.667581 | -2.311892 | 0.020783619 | 0.24853644 |
| *TACC1* | -1.353301 | 0.585712 | -2.310522 | 0.020859245 | 0.2491521 |
| *ENSG00000273674* | -2.84826 | 1.233096 | -2.309846 | NA | NA |
| *DHCR7* | -1.961739 | 0.849698 | -2.308749 | 0.020957516 | 0.24989258 |
| *ADAM15* | -3.432467 | 1.486907 | -2.308461 | 0.02097349 | 0.24989258 |

| | | | | | |
|---|---|---|---|---|---|
| *USP44* | -2.359461 | 1.022255 | -2.308094 | 0.020993882 | 0.24989258 |
| *ITM2A* | -1.811288 | 0.78492 | -2.307607 | 0.021020981 | 0.24992687 |
| *ENSG00000262089* | -2.445852 | 1.060576 | -2.306154 | 0.021102018 | 0.25053696 |
| *WIPI1* | -1.486204 | 0.644631 | -2.305513 | 0.021137866 | 0.25053696 |
| *SLC41A1* | -1.564846 | 0.678779 | -2.305383 | 0.021145125 | 0.25053696 |
| *PIEZO1* | -1.866395 | 0.81061 | -2.302456 | 0.021309481 | 0.25219478 |
| *VMAC* | -1.476889 | 0.642053 | -2.300261 | 0.021433427 | 0.2533711 |
| *TAB1* | -1.556018 | 0.67692 | -2.298674 | 0.02152346 | 0.25412259 |
| *MRPS24* | -2.458222 | 1.069595 | -2.298273 | 0.021546247 | 0.25412259 |
| *COG6* | -1.613498 | 0.702219 | -2.297715 | 0.021578 | 0.25420657 |
| *TTC4* | -2.716051 | 1.183061 | -2.295783 | 0.021688301 | 0.25471042 |
| *DNAJC27.AS1* | -3.823416 | 1.66549 | -2.29567 | 0.021694762 | 0.25471042 |
| *FES* | -3.226407 | 1.405572 | -2.295441 | 0.021707844 | 0.25471042 |
| *ARF4.AS1* | -3.091603 | 1.346964 | -2.295238 | 0.021719494 | 0.25471042 |
| *ENSG00000174171* | -3.045507 | 1.327671 | -2.293871 | 0.02179789 | 0.25523056 |
| *ZNF235* | -1.860442 | 0.811144 | -2.293603 | 0.021813309 | 0.25523056 |
| *UVSSA* | -1.890854 | 0.82485 | -2.29236 | 0.021884885 | 0.25577804 |
| *ENSG00000260136* | -2.419052 | 1.05635 | -2.29001 | 0.022020717 | 0.25703644 |
| *PRKAR1B* | -2.532087 | 1.10589 | -2.289637 | 0.022042369 | 0.25703644 |
| *RHOB* | -2.690393 | 1.1758 | -2.288138 | 0.022129504 | 0.25707543 |
| *GADD45A* | -1.902209 | 0.831439 | -2.287852 | 0.022146155 | 0.25707543 |
| *BAG4* | -1.398784 | 0.611508 | -2.287434 | 0.02217052 | 0.25707543 |
| *SLFN13* | -2.215068 | 0.968598 | -2.28688 | 0.022202847 | 0.25707543 |
| *XXYLT1* | -2.681348 | 1.17265 | -2.286571 | 0.022220853 | 0.25707543 |
| *RUVBL1* | -1.497562 | 0.655068 | -2.286117 | 0.022247389 | 0.25707543 |
| *TNFRSF10D* | -3.759896 | 1.644913 | -2.285773 | 0.022267564 | 0.25707543 |
| *MYH3* | -2.435476 | 1.065619 | -2.285504 | 0.022283317 | 0.25707543 |
| *GPM6A* | -3.001369 | 1.313402 | -2.285187 | 0.022301869 | 0.25707543 |
| *PTER* | -1.416179 | 0.619769 | -2.28501 | 0.022312251 | 0.25707543 |
| *BBS10* | -2.338296 | 1.023377 | -2.284882 | 0.022319728 | 0.25707543 |
| *RNFT2* | -3.044044 | 1.333135 | -2.283372 | 0.02240844 | 0.25780948 |
| *ZNF232* | -1.373398 | 0.602152 | -2.280816 | 0.022559364 | 0.25874063 |
| *KLHL8* | -1.335569 | 0.585684 | -2.280359 | 0.022586425 | 0.25874063 |
| *PGAP2* | -1.48035 | 0.649181 | -2.280336 | 0.022587773 | 0.25874063 |
| *NAA35* | -1.951339 | 0.855759 | -2.280243 | 0.022593276 | 0.25874063 |
| *TMSB4X* | -0.149662 | 0.065654 | -2.27955 | 0.022634372 | 0.25874063 |
| *ZC2HC1A* | -2.045181 | 0.89727 | -2.279337 | 0.022647035 | 0.25874063 |
| *PITPNC1* | -2.582361 | 1.13319 | -2.278841 | 0.022676496 | 0.25874063 |
| *PLEKHF1* | -1.911404 | 0.839023 | -2.278129 | 0.022718879 | 0.25874063 |
| *SLC35B1* | -1.386001 | 0.608483 | -2.277797 | 0.022738678 | 0.25874063 |
| *FDXR* | -2.359017 | 1.036151 | -2.27671 | 0.022803542 | 0.25874063 |
| *CANT1* | -1.321783 | 0.58075 | -2.275994 | 0.022846397 | 0.25874063 |
| *MAP3K7CL* | -2.423621 | 1.064902 | -2.275911 | 0.022851367 | 0.25874063 |
| *ENSG00000264112* | -2.789982 | 1.226132 | -2.275433 | 0.022879944 | 0.25874063 |
| *ZBTB21* | -2.013707 | 0.885133 | -2.275034 | 0.022903879 | 0.25874063 |
| *CHAF1A* | -1.843005 | 0.810351 | -2.274329 | 0.022946196 | 0.25874063 |
| *SLC22A18* | -3.074344 | 1.351902 | -2.274088 | 0.022960688 | 0.25874063 |

| | | | | | |
|---|---|---|---|---|---|
| *CD99L2* | -2.169538 | 0.954028 | -2.274083 | 0.022960975 | 0.25874063 |
| *OTUD6B* | -1.540616 | 0.677481 | -2.274035 | 0.022963883 | 0.25874063 |
| *IKZF2* | -1.960461 | 0.862252 | -2.273651 | 0.022986982 | 0.25874063 |
| *FBXW8* | -2.697832 | 1.186597 | -2.273587 | 0.022990809 | 0.25874063 |
| *ENSG00000260793* | -1.768538 | 0.778456 | -2.271853 | 0.023095396 | 0.25952627 |
| *IMPA2* | -2.467305 | 1.086155 | -2.271596 | 0.023110915 | 0.25952627 |
| *SGCB* | -1.873456 | 0.825046 | -2.27073 | 0.023163337 | 0.25983222 |
| *CCDC7* | -2.608772 | 1.149205 | -2.270067 | 0.023203514 | 0.26000029 |
| *MINDY1* | -2.437091 | 1.073932 | -2.269317 | 0.023249055 | 0.26015206 |
| *FAM91A1* | -1.268636 | 0.559113 | -2.269014 | 0.023267476 | 0.26015206 |
| *NFYB* | -1.437206 | 0.634066 | -2.266651 | 0.02341157 | 0.2611291 |
| *ENSG00000261098* | -1.580047 | 0.697256 | -2.266093 | 0.023445701 | 0.2611291 |
| *GCLM* | -1.580193 | 0.69737 | -2.265933 | 0.023455477 | 0.2611291 |
| *C16orf70* | -1.51678 | 0.669387 | -2.265923 | 0.023456073 | 0.2611291 |
| *DPPA4* | -2.194625 | 0.969762 | -2.263055 | 0.023632284 | 0.26259904 |
| *ZNF782* | -3.477663 | 1.536924 | -2.262742 | 0.023651618 | 0.26259904 |
| *GLI4* | -2.930099 | 1.295052 | -2.262534 | 0.023664448 | 0.26259904 |
| *APOBEC3F* | -1.360346 | 0.601614 | -2.26116 | 0.023749348 | 0.26325808 |
| *LCN8* | -3.552102 | 1.571057 | -2.260964 | 0.023761484 | NA |
| *GIPC1* | -2.626404 | 1.16384 | -2.25667 | 0.02402869 | 0.26577449 |
| *SH3D19* | -2.729542 | 1.209729 | -2.256326 | 0.024050235 | 0.26577449 |
| *AMPD2* | -1.862922 | 0.825681 | -2.256224 | 0.024056626 | 0.26577449 |
| *CNNM3* | -2.773281 | 1.229575 | -2.255479 | 0.024103251 | 0.26577449 |
| *GLRX5* | -1.907465 | 0.845726 | -2.255418 | 0.024107131 | 0.26577449 |
| *PDIK1L* | -2.119767 | 0.940207 | -2.254576 | 0.024159977 | 0.26577449 |
| *GNAQ* | -3.978749 | 1.764753 | -2.254564 | 0.024160682 | NA |
| *AGPS* | -1.548057 | 0.686687 | -2.254383 | 0.024172074 | 0.26577449 |
| *TRIM3* | -3.113505 | 1.381628 | -2.253505 | 0.024227348 | 0.26577449 |
| *FAM118B* | -1.845692 | 0.819068 | -2.253404 | 0.024233704 | 0.26577449 |
| *ENSG00000261474* | -2.176639 | 0.966071 | -2.253084 | 0.024253863 | 0.26577449 |
| *TBC1D2B* | -1.857222 | 0.824354 | -2.252943 | 0.024262724 | 0.26577449 |
| *MTX3* | -2.722536 | 1.208628 | -2.252584 | 0.024285402 | 0.26577449 |
| *ENSG00000267519* | -1.460825 | 0.64903 | -2.250782 | 0.024399347 | 0.26608835 |
| *SAMD1* | -1.652777 | 0.734351 | -2.250663 | 0.024406917 | 0.26608835 |
| *TRAFD1* | -1.456333 | 0.647271 | -2.249958 | 0.024451633 | 0.26608835 |
| *DDHD2* | -1.403212 | 0.623789 | -2.249498 | 0.024480852 | 0.26608835 |
| *LRSAM1* | -2.028943 | 0.901984 | -2.249423 | 0.024485594 | 0.26608835 |
| *GLCE* | -3.020752 | 1.342926 | -2.249381 | 0.024488281 | 0.26608835 |
| *MAN2A2* | -2.100137 | 0.933744 | -2.249158 | 0.024502436 | 0.26608835 |
| *ENSG00000282393* | -2.540584 | 1.129713 | -2.248877 | 0.024520351 | 0.26608835 |
| *WNT10A* | -1.452935 | 0.646616 | -2.246984 | 0.024641076 | 0.2668302 |
| *KMT2B* | -2.141679 | 0.953398 | -2.246364 | 0.024680709 | 0.2668302 |
| *PPARGC1B* | -2.90431 | 1.292956 | -2.246256 | 0.024687585 | 0.2668302 |
| *LINC02018* | -1.884832 | 0.839232 | -2.245899 | 0.024710458 | 0.2668302 |
| *ZNF10* | -3.198486 | 1.42422 | -2.245782 | 0.024717991 | 0.2668302 |
| *ENSG00000237440* | -1.790168 | 0.797484 | -2.24477 | 0.024782883 | 0.26725115 |
| *RORA.AS1* | -2.536834 | 1.130461 | -2.24407 | 0.024827894 | 0.26744039 |

| | | | | | |
|---|---|---|---|---|---|
| *LMBRD2* | -2.391643 | 1.066193 | -2.243162 | 0.024886342 | 0.26744039 |
| *MLLT3* | -2.236234 | 0.996966 | -2.243041 | 0.024894191 | 0.26744039 |
| *BTBD9* | -2.404937 | 1.072565 | -2.24223 | 0.024946503 | 0.26744039 |
| *USP49* | -3.168366 | 1.413093 | -2.242149 | 0.024951747 | 0.26744039 |
| *ANKRD42* | -1.407226 | 0.627642 | -2.242084 | 0.02495592 | 0.26744039 |
| *SGCE* | -2.33882 | 1.043377 | -2.241586 | 0.024988147 | 0.26750797 |
| *RHOBTB2* | -1.795825 | 0.801622 | -2.240238 | 0.025075454 | 0.26800757 |
| *INVS* | -2.185497 | 0.97564 | -2.240064 | 0.025086755 | 0.26800757 |
| *SLC2A8* | -1.578836 | 0.70553 | -2.2378 | 0.025234086 | 0.26874212 |
| *TRIB2* | -2.714738 | 1.213258 | -2.23756 | 0.025249763 | 0.26874212 |
| *PSKH1* | -1.875505 | 0.838319 | -2.237221 | 0.025271928 | 0.26874212 |
| *UBE2M* | -1.719799 | 0.768752 | -2.237132 | 0.025277719 | 0.26874212 |
| *TCFL5* | -1.814223 | 0.811143 | -2.236626 | 0.025310813 | 0.26874212 |
| *MICA* | -1.396274 | 0.624412 | -2.236141 | 0.025342556 | 0.26874212 |
| *NUP58* | -1.387815 | 0.620717 | -2.235823 | 0.02536334 | 0.26874212 |
| *ZNF300* | -3.026006 | 1.353521 | -2.235656 | 0.025374345 | 0.26874212 |
| *METTL25* | -1.800304 | 0.805464 | -2.235115 | 0.025409774 | 0.26874212 |
| *RFX2* | -1.661274 | 0.743292 | -2.235022 | 0.025415922 | 0.26874212 |
| *ITPR3* | -1.673523 | 0.749278 | -2.233515 | 0.025514984 | 0.26878092 |
| *PRKACA* | -1.647618 | 0.737688 | -2.233489 | 0.025516691 | 0.26878092 |
| *PPP2R3A* | -2.879442 | 1.289274 | -2.233382 | 0.025523736 | 0.26878092 |
| *SLC22A5* | -3.006375 | 1.346109 | -2.233382 | 0.02552377 | 0.26878092 |
| *SFXN4* | -2.165588 | 0.96983 | -2.232956 | 0.025551818 | 0.268802 |
| *FRS3* | -3.151601 | 1.411739 | -2.232425 | 0.025586907 | 0.26889703 |
| *IMPACT* | -1.705256 | 0.764115 | -2.231673 | 0.025636554 | 0.269099 |
| *ENSG00000272871* | -2.950537 | 1.322644 | -2.230788 | 0.025695192 | 0.269099 |
| *FBXO2* | -2.957966 | 1.326182 | -2.230438 | 0.025718358 | 0.269099 |
| *CETP* | -2.806043 | 1.258129 | -2.230331 | 0.025725488 | 0.269099 |
| *GPD2* | -2.692988 | 1.20769 | -2.229868 | 0.025756208 | 0.269099 |
| *ENSG00000187951* | -1.936767 | 0.868615 | -2.229718 | NA | NA |
| *ZNF382* | -1.921578 | 0.86182 | -2.229676 | 0.025768989 | 0.269099 |
| *LIG3* | -1.742918 | 0.781795 | -2.22938 | 0.025788654 | 0.269099 |
| *VCPKMT* | -1.441312 | 0.646703 | -2.228709 | 0.025833242 | 0.26910586 |
| *FBXO6* | -1.693789 | 0.760285 | -2.227834 | 0.025891596 | 0.26910586 |
| *CEP85* | -3.026524 | 1.358534 | -2.227787 | 0.025894721 | 0.26910586 |
| *TNFSF12* | -1.262097 | 0.566577 | -2.22758 | 0.025908519 | 0.26910586 |
| *NDRG2* | -3.317205 | 1.489264 | -2.227413 | 0.025919692 | 0.26910586 |
| *ZNF792* | -2.710692 | 1.217298 | -2.22681 | 0.025959944 | 0.26925289 |
| *ENSG00000279816* | -1.659854 | 0.745958 | -2.225132 | 0.026072379 | 0.26975977 |
| *SLC38A5* | -2.281343 | 1.025342 | -2.224958 | 0.026084027 | 0.26975977 |
| *ZCCHC2* | -1.559392 | 0.700947 | -2.224694 | 0.026101771 | 0.26975977 |
| *ENSG00000197813* | -3.159184 | 1.420215 | -2.22444 | 0.026118824 | 0.26975977 |
| *TMEM254* | -2.092639 | 0.940878 | -2.224133 | 0.026139512 | 0.26975977 |
| *RPUSD2* | -1.312322 | 0.59029 | -2.223182 | 0.02620354 | 0.27015038 |
| *NCAPD2* | -1.753094 | 0.78889 | -2.222227 | 0.026267965 | 0.27054431 |
| *HCFC2* | -3.142239 | 1.414359 | -2.22167 | 0.026305634 | 0.27066216 |
| *CERNA1* | -1.683341 | 0.758069 | -2.220564 | 0.026380539 | 0.27095978 |

| | | | | | |
|---|---|---|---|---|---|
| *SLC39A8* | -2.992493 | 1.348182 | -2.21965 | 0.026442534 | 0.27095978 |
| *PPP2R2D* | -1.772723 | 0.79885 | -2.219095 | 0.02648025 | 0.27095978 |
| *ENSG00000230555* | -1.934517 | 0.871783 | -2.219036 | 0.026484304 | 0.27095978 |
| *SLC35A2* | -1.583816 | 0.713856 | -2.218676 | 0.026508779 | 0.27095978 |
| *NMB* | -2.056348 | 0.927051 | -2.21816 | 0.026543957 | 0.27095978 |
| *DDX51* | -2.036351 | 0.918104 | -2.217995 | NA | NA |
| *SHMT1* | -1.68973 | 0.76188 | -2.217843 | 0.026565522 | 0.27095978 |
| *HEATR5B* | -1.711655 | 0.771849 | -2.217603 | 0.026581882 | 0.27095978 |
| *ENSG00000232527* | -2.614381 | 1.179131 | -2.21721 | 0.02660876 | 0.27095978 |
| *CLECL1* | -1.615784 | 0.728826 | -2.216967 | 0.026625326 | 0.27095978 |
| *PLEKHO2* | -2.231032 | 1.006435 | -2.216767 | 0.026639011 | 0.27095978 |
| *ENTR1* | -1.602126 | 0.722812 | -2.216519 | 0.026655997 | 0.27095978 |
| *CHD1L* | -1.34147 | 0.60536 | -2.215986 | 0.026692477 | 0.27095978 |
| *UQCRHL* | -1.641252 | 0.740689 | -2.215845 | 0.026702141 | 0.27095978 |
| *ENSG00000269898* | -2.468251 | 1.114636 | -2.214401 | 0.026801178 | 0.27108887 |
| *ANO10* | -2.644354 | 1.194175 | -2.214377 | 0.026802848 | 0.27108887 |
| *CNN3* | -1.734974 | 0.784 | -2.212977 | 0.026899266 | 0.27108887 |
| *ATAD3A* | -1.666803 | 0.753349 | -2.212524 | 0.026930453 | 0.27108887 |
| *DDX60* | -2.313419 | 1.045891 | -2.211912 | 0.026972723 | 0.27108887 |
| *ABCD3* | -1.744666 | 0.788905 | -2.211504 | 0.027000956 | 0.27108887 |
| *MIR4458HG* | -1.493175 | 0.675193 | -2.211477 | 0.027002835 | 0.27108887 |
| *PTPRK* | -2.083326 | 0.942154 | -2.211238 | 0.027019366 | 0.27108887 |
| *VAV3* | -1.603309 | 0.725133 | -2.211055 | 0.027032009 | 0.27108887 |
| *FCGBP* | -1.675283 | 0.757866 | -2.210526 | 0.027068675 | 0.27108887 |
| *TSTD2* | -1.461404 | 0.661135 | -2.210449 | 0.02707403 | 0.27108887 |
| *NKIRAS1* | -1.480936 | 0.670084 | -2.210075 | 0.027099925 | 0.27108887 |
| *OTUD3* | -2.153337 | 0.974372 | -2.209974 | 0.027106938 | 0.27108887 |
| *LRIG2* | -1.527846 | 0.69143 | -2.20969 | 0.027126719 | 0.27108887 |
| *KIAA0753* | -3.158811 | 1.429688 | -2.209441 | 0.027143987 | 0.27108887 |
| *ZNF584* | -2.089215 | 0.945605 | -2.209395 | 0.027147171 | 0.27108887 |
| *GNA11* | -2.696166 | 1.220432 | -2.20919 | 0.027161424 | 0.27108887 |
| *PNPO* | -2.399969 | 1.087179 | -2.207519 | 0.027277824 | 0.27166948 |
| *COPG2* | -2.787044 | 1.262936 | -2.206798 | 0.027328205 | 0.27166948 |
| *ATP11A* | -1.777783 | 0.805651 | -2.206642 | 0.027339063 | 0.27166948 |
| *RTKN* | -3.736852 | 1.693638 | -2.206405 | 0.027355624 | 0.27166948 |
| *ENSG00000268220* | -3.238035 | 1.467754 | -2.206116 | 0.027375882 | 0.27166948 |
| *MTMR10* | -2.373933 | 1.076081 | -2.206092 | 0.027377545 | 0.27166948 |
| *ATP13A3* | -1.670063 | 0.757615 | -2.204368 | 0.02749848 | 0.27224364 |
| *C20orf96* | -2.065415 | 0.937048 | -2.204171 | 0.027512307 | 0.27224364 |
| *ICMT* | -2.243743 | 1.01815 | -2.203744 | 0.02754234 | 0.27224364 |
| *ABHD6* | -2.257152 | 1.024273 | -2.203663 | 0.02754806 | 0.27224364 |
| *ERLIN1* | -2.244403 | 1.018614 | -2.20339 | 0.027567307 | 0.27224364 |
| *LARGE2* | -1.322934 | 0.600856 | -2.201747 | 0.027683171 | 0.2731265 |
| *INPPL1* | -1.534411 | 0.697397 | -2.200198 | 0.027792881 | 0.27376647 |
| *BPGM* | -1.451905 | 0.660038 | -2.199731 | 0.027825989 | 0.27376647 |
| *THNSL1* | -2.632481 | 1.196764 | -2.199665 | 0.027830642 | 0.27376647 |
| *ZNF436.AS1* | -2.73624 | 1.244298 | -2.199024 | 0.027876241 | 0.27376647 |

| | | | | | |
|---|---|---|---|---|---|
| *AIM2* | -3.297573 | 1.499642 | -2.198907 | 0.027884559 | 0.27376647 |
| *CSTF2* | -2.520481 | 1.146433 | -2.198542 | 0.027910509 | 0.27376647 |
| *RTCA* | -1.260262 | 0.573311 | -2.198216 | 0.02793373 | 0.27376647 |
| *ZKSCAN5* | -2.757501 | 1.254686 | -2.197762 | 0.027966046 | 0.27382315 |
| *NOXA1* | -3.609831 | 1.643451 | -2.196495 | 0.028056562 | NA |
| *AIFM1* | -1.625103 | 0.739869 | -2.196473 | 0.028058118 | 0.27446425 |
| *PLEKHA8* | -2.362213 | 1.075811 | -2.195752 | 0.028109734 | 0.27470876 |
| *C8orf88* | -2.047783 | 0.932927 | -2.19501 | 0.028162895 | 0.27479716 |
| *SCLY* | -1.997053 | 0.909868 | -2.194882 | 0.028172035 | 0.27479716 |
| *C12orf60* | -2.55177 | 1.162963 | -2.194197 | 0.02822127 | 0.27501747 |
| *ENSG00000232412* | -3.281052 | 1.496577 | -2.19237 | 0.028352778 | NA |
| *TBC1D7* | -2.315807 | 1.056327 | -2.192319 | 0.028356466 | 0.27607427 |
| *ITGA3* | -3.081626 | 1.406291 | -2.191314 | 0.028429094 | 0.2765205 |
| *TMEM50B* | -1.212514 | 0.553591 | -2.19027 | 0.028504653 | 0.27682307 |
| *ZSCAN22* | -2.789589 | 1.273702 | -2.190143 | 0.028513849 | 0.27682307 |
| *PILRA* | -3.714045 | 1.69614 | -2.189704 | 0.028545728 | NA |
| *MYLK.AS1* | -1.747186 | 0.798054 | -2.189307 | 0.028574527 | 0.27696459 |
| *TTC39B* | -3.16316 | 1.445141 | -2.188824 | 0.0286096 | 0.27696459 |
| *FAM161B* | -3.043502 | 1.390766 | -2.188364 | 0.028643075 | 0.27696459 |
| *CARD6* | -3.028846 | 1.384182 | -2.188185 | 0.02865611 | 0.27696459 |
| *DUS2* | 1.309778 | 0.598596 | 2.188085 | 0.028663385 | 0.27696459 |
| *IMPDH1* | -1.519631 | 0.694616 | -2.187728 | 0.028689452 | 0.27696459 |
| *SCML4* | -2.479729 | 1.134559 | -2.185633 | 0.028842447 | 0.27790611 |
| *ZNF252P.AS1* | -3.5555 | 1.627079 | -2.185204 | 0.028873923 | 0.27790611 |
| *NHLRC4* | -2.571025 | 1.17659 | -2.18515 | 0.028877834 | 0.27790611 |
| *HIST4H4* | -2.216553 | 1.014587 | -2.184684 | 0.028912004 | 0.27790611 |
| *CENPP* | -3.109799 | 1.423622 | -2.184428 | 0.028930824 | 0.27790611 |
| *GYS1* | -1.53434 | 0.702567 | -2.183907 | 0.028969066 | 0.27790611 |
| *TTC34* | -3.206266 | 1.46846 | -2.18342 | 0.029004891 | 0.27790611 |
| *CLSTN3* | -2.320198 | 1.062719 | -2.183265 | 0.029016319 | 0.27790611 |
| *SESN1* | -1.386268 | 0.635019 | -2.183034 | 0.029033273 | 0.27790611 |
| *ENSG00000271254* | -1.915599 | 0.877652 | -2.182639 | 0.029062387 | 0.27790611 |
| *MORN3* | -3.564706 | 1.633972 | -2.181619 | 0.029137635 | 0.27790611 |
| *ACOX1* | -1.642076 | 0.752781 | -2.181347 | 0.029157772 | 0.27790611 |
| *POT1* | -1.600753 | 0.73384 | -2.181338 | 0.029158404 | 0.27790611 |
| *SPRYD3* | -1.646556 | 0.754979 | -2.180929 | 0.029188694 | 0.27790611 |
| *NUDCD1* | -1.761963 | 0.807907 | -2.180899 | 0.029190913 | 0.27790611 |
| *ENSG00000260495* | -3.349439 | 1.536124 | -2.180448 | 0.029224239 | NA |
| *INTS14* | -1.530894 | 0.702475 | -2.179286 | 0.029310434 | 0.2787868 |
| *CXCL8* | -3.615112 | 1.659318 | -2.178673 | 0.029355938 | 0.27896251 |
| *ZNF570* | -1.516839 | 0.696609 | -2.177462 | 0.029446111 | 0.27921175 |
| *SETD1A* | -1.500032 | 0.688926 | -2.177349 | 0.029454527 | 0.27921175 |
| *SAMD10* | -2.897713 | 1.33102 | -2.177062 | 0.029475959 | 0.27921175 |
| *C9orf64* | -2.456256 | 1.128366 | -2.176825 | 0.029493659 | 0.27921175 |
| *ENSG00000248559* | -1.719844 | 0.790207 | -2.176448 | 0.02952179 | 0.27921175 |
| *DDX11* | -1.463981 | 0.672805 | -2.175937 | 0.029559988 | 0.27921175 |
| *ZNF398* | -2.081741 | 0.956781 | -2.175777 | 0.029571941 | 0.27921175 |

| | | | | | |
|---|---|---|---|---|---|
| *BIRC5* | -3.193004 | 1.468032 | -2.175023 | 0.029628381 | 0.27921175 |
| *ENSG00000231113* | -2.296933 | 1.056218 | -2.174676 | 0.029654412 | 0.27921175 |
| *SH3BP5.AS1* | -1.772796 | 0.81537 | -2.174224 | 0.029688288 | 0.27921175 |
| *RSPRY1* | -1.425553 | 0.655753 | -2.173917 | 0.029711345 | 0.27921175 |
| *CFAP44* | -2.15937 | 0.993327 | -2.173875 | 0.029714499 | 0.27921175 |
| *RCBTB1* | -1.653871 | 0.761259 | -2.172547 | 0.029814426 | 0.27921175 |
| *UCKL1* | -1.198348 | 0.551628 | -2.172385 | 0.029826604 | 0.27921175 |
| *MEI1* | -1.407657 | 0.648811 | -2.169596 | 0.030037462 | 0.27921175 |
| *ETAA1* | -1.604742 | 0.739708 | -2.169428 | 0.030050234 | 0.27921175 |
| *MIR222HG* | -3.095518 | 1.42689 | -2.169417 | 0.030051074 | 0.27921175 |
| *DSE* | -1.71272 | 0.789616 | -2.169054 | 0.030078621 | 0.27921175 |
| *RANGAP1* | -1.562574 | 0.720462 | -2.16885 | 0.030094064 | 0.27921175 |
| *ERLIN2* | -1.689753 | 0.779769 | -2.166991 | 0.030235512 | 0.27921175 |
| *LINC01684* | -2.258449 | 1.042251 | -2.166895 | 0.030242859 | 0.27921175 |
| *B4GAT1* | -3.183429 | 1.469132 | -2.166877 | 0.03024421 | 0.27921175 |
| *FKBP5* | -1.501382 | 0.692961 | -2.166618 | 0.030263964 | 0.27921175 |
| *CSE1L* | -1.402091 | 0.647223 | -2.166318 | 0.030286926 | 0.27921175 |
| *FANCM* | -1.520517 | 0.701939 | -2.166166 | 0.030298482 | 0.27921175 |
| *CCDC71L* | -1.767435 | 0.81629 | -2.165204 | 0.030372034 | 0.27921175 |
| *PDLIM7* | -1.544802 | 0.713484 | -2.165155 | 0.030375852 | 0.27921175 |
| *CEP192* | -1.603365 | 0.740669 | -2.164753 | 0.030406569 | 0.27921175 |
| *SLC9A8* | -2.017821 | 0.932338 | -2.164259 | 0.030444461 | 0.27921175 |
| *SLX4* | -2.980691 | 1.377554 | -2.163757 | 0.030483033 | 0.27921175 |
| *MEGF8* | -2.169296 | 1.002565 | -2.163746 | 0.030483883 | 0.27921175 |
| *ZNF763* | -2.983476 | 1.379178 | -2.163228 | 0.030523639 | 0.27921175 |
| *TMEM94* | -1.765097 | 0.816046 | -2.162989 | 0.030542063 | 0.27921175 |
| *HYAL2* | -2.600008 | 1.20216 | -2.162781 | 0.030557995 | 0.27921175 |
| *LPAR2* | -2.317102 | 1.071738 | -2.162005 | 0.030617797 | 0.27921175 |
| *IGHV3.7* | -4.180724 | 1.933865 | -2.161849 | 0.030629832 | NA |
| *PLEKHB1* | -2.325346 | 1.075782 | -2.161541 | 0.030653599 | 0.27921175 |
| *IL9R* | -2.276614 | 1.053355 | -2.161298 | 0.030672352 | 0.27921175 |
| *GALNT3* | -1.868066 | 0.864488 | -2.160893 | 0.030703605 | 0.27921175 |
| *PLD2* | -2.528779 | 1.170414 | -2.160585 | 0.03072741 | 0.27921175 |
| *CDC42BPB* | -1.942635 | 0.899138 | -2.160552 | 0.030729984 | 0.27921175 |
| *ASXL1* | -1.274605 | 0.59002 | -2.160273 | 0.030751565 | 0.27921175 |
| *ENSG00000237181* | -2.512623 | 1.163635 | -2.159289 | 0.030827745 | 0.27921175 |
| *RELL2* | -1.782038 | 0.825492 | -2.158758 | 0.030868953 | 0.27921175 |
| *SART3* | -1.460847 | 0.676732 | -2.15868 | 0.030875029 | 0.27921175 |
| *ENSG00000228801* | -1.970536 | 0.91288 | -2.158592 | 0.030881834 | 0.27921175 |
| *L2HGDH* | -2.647986 | 1.22677 | -2.158503 | 0.030888739 | 0.27921175 |
| *AASS* | -2.657255 | 1.232089 | -2.156706 | 0.03102856 | 0.27921175 |
| *L1TD1* | -2.130898 | 0.988085 | -2.156593 | 0.03103741 | 0.27921175 |
| *ENSG00000276148* | -2.670167 | 1.2382 | -2.156491 | 0.031045363 | 0.27921175 |
| *SKP2* | -1.786401 | 0.82839 | -2.156475 | 0.031046622 | 0.27921175 |
| *STYX* | -1.45886 | 0.676641 | -2.156033 | 0.031081055 | 0.27921175 |
| *ENSG00000247134* | -3.508108 | 1.627256 | -2.155843 | 0.031095948 | 0.27921175 |
| *ACAP3* | -2.238224 | 1.038843 | -2.154535 | 0.031198219 | 0.27921175 |

| | | | | | |
|---|---|---|---|---|---|
| *HS6ST1* | -2.727869 | 1.266451 | -2.153947 | 0.031244304 | 0.27921175 |
| *MRPS6* | 1.14753 | 0.532899 | 2.153371 | 0.031289507 | 0.27921175 |
| *DFFB* | -2.755885 | 1.279904 | -2.153196 | 0.031303248 | 0.27921175 |
| *RSPH3* | -1.459062 | 0.677646 | -2.153132 | 0.031308287 | 0.27921175 |
| *PEMT* | -1.990224 | 0.924379 | -2.153039 | 0.031315592 | 0.27921175 |
| *GPR89B* | -1.501704 | 0.69758 | -2.152732 | 0.031339726 | 0.27921175 |
| *FBXO4* | -1.393897 | 0.647597 | -2.152415 | 0.031364706 | 0.27921175 |
| *GNAZ* | -2.318842 | 1.077551 | -2.151954 | 0.031400943 | 0.27921175 |
| *NEURL1* | -5.549515 | 2.578839 | -2.151944 | 0.031401798 | 0.27921175 |
| *ACACA* | -2.408222 | 1.119119 | -2.15189 | 0.031406009 | 0.27921175 |
| *ZNF197* | -2.522463 | 1.172246 | -2.151821 | 0.031411486 | 0.27921175 |
| *PIK3R4* | -2.207732 | 1.026018 | -2.151747 | 0.031417284 | 0.27921175 |
| *ANKRD52* | -1.705687 | 0.792737 | -2.151643 | 0.031425493 | 0.27921175 |
| *SUV39H1* | -2.13228 | 0.991008 | -2.151627 | 0.031426779 | 0.27921175 |
| *FASN* | -3.631958 | 1.688033 | -2.151592 | 0.03142953 | NA |
| *NLK* | -1.69867 | 0.789783 | -2.150805 | 0.031491576 | 0.27921175 |
| *ZNF616* | -2.370116 | 1.101988 | -2.150763 | 0.03149491 | 0.27921175 |
| *MAP11* | -2.141329 | 0.99588 | -2.150187 | 0.03154045 | 0.27921175 |
| *CASTOR3* | -1.991885 | 0.926413 | -2.150106 | 0.031546869 | 0.27921175 |
| *CARMIL1* | -2.065739 | 0.960894 | -2.14981 | 0.031570266 | 0.27921175 |
| *TCTEX1D2* | -1.458871 | 0.678612 | -2.149786 | 0.03157213 | 0.27921175 |
| *PRADC1* | -1.6593 | 0.771977 | -2.149416 | 0.031601441 | 0.27921175 |
| *TBC1D10B* | -1.515375 | 0.705037 | -2.149356 | 0.031606198 | 0.27921175 |
| *IGFBP4* | -2.027947 | 0.943514 | -2.149355 | 0.031606292 | 0.27921175 |
| *ZNF35* | -2.427183 | 1.129425 | -2.149043 | 0.031631021 | 0.27921175 |
| *ZNF519* | -1.621937 | 0.754757 | -2.148953 | 0.031638091 | 0.27921175 |
| *TFRC* | -1.763132 | 0.820635 | -2.148497 | 0.031674315 | 0.27921175 |
| *PHRF1* | -1.437132 | 0.669091 | -2.147886 | 0.031722805 | 0.27921175 |
| *MAGED1* | -2.145539 | 0.999158 | -2.147346 | 0.031765712 | 0.27921175 |
| *ALG1L2* | -3.196517 | 1.488626 | -2.147293 | 0.031769925 | 0.27921175 |
| *LINC00526* | -1.817176 | 0.84634 | -2.1471 | 0.031785301 | 0.27921175 |
| *TENT4A* | -2.313154 | 1.077527 | -2.146725 | 0.0318152 | 0.27921175 |
| *CHAC2* | -3.125282 | 1.455941 | -2.146571 | 0.031827449 | 0.27921175 |
| *CPSF1* | -1.779888 | 0.829394 | -2.14601 | 0.031872154 | 0.27921175 |
| *COLEC12* | -1.808663 | 0.843007 | -2.145488 | 0.031913814 | 0.27921175 |
| *ENSG00000229999* | -1.36261 | 0.635264 | -2.144951 | 0.031956733 | 0.27921175 |
| *AGAP6* | -3.179378 | 1.482525 | -2.144569 | NA | NA |
| *AP4B1* | -2.130763 | 0.993607 | -2.144472 | 0.031995089 | 0.27921175 |
| *SNAI3* | -1.796435 | 0.837807 | -2.144212 | 0.032015927 | 0.27921175 |
| *NBPF10* | -1.458569 | 0.68028 | -2.14407 | 0.032027252 | 0.27921175 |
| *TLR7* | -1.980624 | 0.92381 | -2.143973 | 0.032035027 | 0.27921175 |
| *DNAH14* | -2.242476 | 1.046302 | -2.143239 | 0.032093894 | 0.27921175 |
| *ENSG00000271806* | -3.133165 | 1.462076 | -2.142957 | 0.032116586 | 0.27921175 |
| *FICD* | -2.244831 | 1.047674 | -2.14268 | 0.03213877 | 0.27921175 |
| *ZNF251* | -3.463982 | 1.616786 | -2.142512 | 0.032152307 | 0.27921175 |
| *ENSG00000247363* | -2.33024 | 1.087701 | -2.142354 | 0.032165011 | 0.27921175 |
| *PPAT* | -2.030711 | 0.948318 | -2.141382 | 0.032243231 | 0.27921175 |

| | | | | | |
|---|---|---|---|---|---|
| *ASCC1* | -1.337097 | 0.624423 | -2.141332 | 0.032247298 | 0.27921175 |
| *CASP2* | -1.2907 | 0.602884 | -2.140877 | 0.032283949 | 0.27921175 |
| *PGBD4* | -1.667926 | 0.779108 | -2.140816 | 0.032288919 | 0.27921175 |
| *HSPA14* | -1.535337 | 0.717187 | -2.140775 | 0.032292157 | 0.27921175 |
| *KIAA0319L* | -1.298759 | 0.606694 | -2.140714 | 0.032297108 | 0.27921175 |
| *ZC3H18* | -1.33132 | 0.622003 | -2.140374 | 0.032324548 | 0.27921175 |
| *ENSG00000237188* | -2.005334 | 0.937016 | -2.140128 | 0.032344406 | 0.27921175 |
| *RBBP9* | -2.026884 | 0.9471 | -2.140095 | 0.032347083 | 0.27921175 |
| *ENSG00000266385* | -3.104152 | 1.450498 | -2.140059 | 0.032349982 | 0.27921175 |
| *ZNF473* | -1.846959 | 0.863186 | -2.139699 | 0.032379133 | 0.27921175 |
| *ZBTB40* | -1.396699 | 0.6528 | -2.13955 | 0.032391138 | 0.27921175 |
| *TIGD7* | -2.217097 | 1.036522 | -2.138977 | 0.032437501 | 0.27921175 |
| *MAP2K4* | -1.48134 | 0.692588 | -2.138848 | 0.032447941 | 0.27921175 |
| *INKA2* | -2.595036 | 1.213644 | -2.138218 | 0.032499016 | 0.27921175 |
| *ZBED6CL* | -3.181218 | 1.488166 | -2.137677 | 0.032542959 | 0.27921175 |
| *RBM45* | -1.499903 | 0.701666 | -2.137633 | 0.03254657 | 0.27921175 |
| *CTNNBIP1* | -1.852292 | 0.866727 | -2.137111 | 0.032588986 | 0.27921175 |
| *LSM11* | -3.144949 | 1.471692 | -2.136961 | 0.032601169 | 0.27921175 |
| *TRIM39* | -1.386368 | 0.648765 | -2.136934 | 0.032603379 | 0.27921175 |
| *CD5* | -2.234099 | 1.04549 | -2.136891 | 0.03260683 | 0.27921175 |
| *PARP3* | -1.617862 | 0.75723 | -2.136553 | 0.032634394 | 0.27921175 |
| *INTS7* | -1.981348 | 0.927736 | -2.135681 | 0.032705414 | 0.27921175 |
| *WDCP* | -2.345792 | 1.098428 | -2.13559 | 0.032712844 | 0.27921175 |
| *ENSG00000274605* | -2.549495 | 1.194134 | -2.135017 | 0.032759661 | 0.27921175 |
| *PAXIP1* | -2.502523 | 1.172171 | -2.134948 | 0.032765279 | 0.27921175 |
| *PSTPIP1* | -1.584527 | 0.742283 | -2.134668 | 0.032788144 | 0.27921175 |
| *RALGPS1* | -1.53689 | 0.719977 | -2.134638 | 0.032790629 | 0.27921175 |
| *LINC02352* | -1.945725 | 0.911632 | -2.134331 | 0.032815674 | 0.27921175 |
| *ZNF682* | -1.886575 | 0.883932 | -2.1343 | 0.032818203 | 0.27921175 |
| *RAB40B* | -2.668205 | 1.250711 | -2.133349 | 0.032896073 | 0.27961792 |
| *ENSG00000267169* | -2.919149 | 1.368529 | -2.133056 | 0.032920133 | 0.27961792 |
| *ENSG00000224738* | -3.122231 | 1.464259 | -2.132294 | 0.032982687 | 0.27988973 |
| *IL2RA* | -1.939577 | 0.909789 | -2.131898 | 0.033015209 | 0.27988973 |
| *PTTG1* | -3.035039 | 1.423781 | -2.131676 | 0.033033497 | 0.27988973 |
| *CDC14A* | -2.161552 | 1.014192 | -2.131305 | 0.033064029 | 0.2799186 |
| *LINC01597* | -2.292319 | 1.075959 | -2.13049 | 0.033131222 | 0.28025755 |
| *GDPD1* | -3.020591 | 1.418211 | -2.12986 | 0.033183133 | 0.28046677 |
| *TSPAN10* | -2.297506 | 1.079047 | -2.129198 | 0.033237857 | 0.28050697 |
| *PKIA* | -3.508995 | 1.648077 | -2.129145 | 0.03324225 | 0.28050697 |
| *OSBPL10* | -1.300324 | 0.610925 | -2.12845 | 0.033299781 | 0.2805482 |
| *DGCR8* | -1.699695 | 0.798568 | -2.128429 | 0.033301507 | 0.2805482 |
| *RUSC1* | -1.350458 | 0.63472 | -2.127645 | 0.033366487 | 0.28060428 |
| *IGHV1.18* | -3.566494 | 1.676323 | -2.12757 | 0.033372777 | NA |
| *MYD88* | -1.276718 | 0.600146 | -2.127345 | 0.033391451 | 0.28060428 |
| *AZIN1.AS1* | -2.937454 | 1.38097 | -2.127095 | 0.033412153 | 0.28060428 |
| *GOLGA5* | -1.266802 | 0.595789 | -2.12626 | 0.033481631 | 0.28060428 |
| *KCNN4* | -1.599882 | 0.752543 | -2.125968 | 0.033505925 | 0.28060428 |

| | | | | | |
|---|---|---|---|---|---|
| *LINC01569* | -3.577341 | 1.682984 | -2.125594 | 0.033537065 | 0.28060428 |
| *NCOR2* | -1.663002 | 0.782416 | -2.125471 | 0.033547306 | 0.28060428 |
| *ZNF852* | -2.647458 | 1.24561 | -2.125431 | 0.033550668 | 0.28060428 |
| *ZNF155* | -1.748988 | 0.822907 | -2.125378 | 0.033555067 | 0.28060428 |
| *POLA2* | -2.586442 | 1.217381 | -2.124596 | 0.033620355 | 0.28060428 |
| *PDK1* | -1.3016 | 0.612709 | -2.124337 | 0.033641956 | 0.28060428 |
| *RAD51AP1* | -1.772201 | 0.834309 | -2.124153 | 0.033657384 | 0.28060428 |
| *ABCB9* | -4.168407 | 1.962396 | -2.124142 | 0.033658298 | NA |
| *MAP1LC3B2* | -1.910672 | 0.89952 | -2.124102 | 0.033661637 | 0.28060428 |
| *KIF13B* | -2.870324 | 1.352385 | -2.122417 | 0.033802742 | 0.28097348 |
| *XRCC4* | -1.434474 | 0.676059 | -2.121818 | 0.033853041 | 0.28097348 |
| *SLC29A2* | -2.48831 | 1.172746 | -2.12178 | 0.033856195 | 0.28097348 |
| *DUSP14* | -1.699136 | 0.800817 | -2.121754 | 0.033858405 | 0.28097348 |
| *XG* | -3.358979 | 1.583175 | -2.121673 | 0.033865247 | 0.28097348 |
| *SIK2* | -2.362804 | 1.113707 | -2.121567 | 0.033874129 | 0.28097348 |
| *MGAT3* | -2.828482 | 1.333554 | -2.121011 | 0.033920877 | 0.28097348 |
| *TMEM64* | -2.125865 | 1.002535 | -2.120489 | 0.033964801 | 0.28097348 |
| *ZNF93* | -2.507341 | 1.182481 | -2.120407 | 0.033971746 | 0.28097348 |
| *GK5* | -1.911987 | 0.90174 | -2.12033 | 0.033978188 | 0.28097348 |
| *CLSTN1* | -1.325172 | 0.625101 | -2.119934 | 0.034011581 | 0.28102443 |
| *ENSG00000269399* | -1.574415 | 0.742815 | -2.119524 | 0.03404619 | 0.28108534 |
| *GSK3B* | -1.320605 | 0.623189 | -2.119108 | 0.034081301 | 0.2811503 |
| *KPTN* | -2.634794 | 1.243592 | -2.118696 | 0.034116178 | 0.28121322 |
| *NSMF* | -3.009196 | 1.421821 | -2.116438 | 0.03430756 | 0.28245649 |
| *ENSG00000251661* | -2.45966 | 1.162677 | -2.115514 | 0.034386201 | 0.28245649 |
| *STARD5* | -2.053993 | 0.970951 | -2.115445 | 0.034392062 | 0.28245649 |
| *MEAK7* | -2.297898 | 1.086442 | -2.115067 | 0.034424276 | 0.28245649 |
| *VSIR* | -1.312943 | 0.62081 | -2.114887 | 0.034439612 | 0.28245649 |
| *SUV39H2* | -3.005332 | 1.421305 | -2.114488 | 0.034473643 | 0.28245649 |
| *KSR1* | -2.288819 | 1.082481 | -2.11442 | 0.034479439 | 0.28245649 |
| *NAA80* | -1.320795 | 0.624684 | -2.114343 | 0.034485967 | 0.28245649 |
| *RBM25* | -1.121416 | 0.530859 | -2.112454 | 0.034647539 | 0.2835548 |
| *USP30* | -3.891089 | 1.84223 | -2.112163 | 0.034672513 | NA |
| *ZMYM6* | -1.279166 | 0.605828 | -2.111432 | 0.034735162 | 0.28384215 |
| *RMDN1* | -1.404955 | 0.665423 | -2.111372 | 0.034740311 | 0.28384215 |
| *ENSG00000257151* | -2.345973 | 1.111322 | -2.110976 | 0.034774364 | 0.28384215 |
| *ZNF827* | -1.414065 | 0.669931 | -2.110763 | 0.034792667 | 0.28384215 |
| *SLC9B2* | -2.017344 | 0.955976 | -2.110245 | 0.034837229 | 0.2839812 |
| *HERC5* | -1.929931 | 0.915183 | -2.108792 | 0.034962511 | 0.28416534 |
| *SLC2A13* | -1.404182 | 0.665885 | -2.108745 | 0.03496658 | 0.28416534 |
| *TRAK2* | -1.784025 | 0.846124 | -2.108467 | 0.034990649 | 0.28416534 |
| *ZNF440* | -2.600759 | 1.233566 | -2.108325 | 0.035002886 | 0.28416534 |
| *ENSG00000269886* | -2.962546 | 1.405931 | -2.107177 | 0.035102255 | 0.28416534 |
| *TIMM23B* | -1.853796 | 0.879766 | -2.107146 | 0.035104884 | 0.28416534 |
| *ARHGAP18* | -3.206623 | 1.521844 | -2.107065 | 0.035111988 | 0.28416534 |
| *KCTD2* | -2.722274 | 1.292017 | -2.106996 | 0.035117968 | 0.28416534 |
| *ARHGAP22* | -3.282888 | 1.55836 | -2.10663 | 0.035149624 | 0.28416534 |

| | | | | | |
|---|---|---|---|---|---|
| *GALNT6* | -2.671023 | 1.267983 | -2.106513 | 0.035159795 | 0.28416534 |
| *KIAA1841* | -2.97514 | 1.412431 | -2.106396 | 0.035169945 | 0.28416534 |
| *INPP5E* | -2.252208 | 1.06963 | -2.105596 | 0.035239504 | 0.28416534 |
| *DZIP3* | -1.460429 | 0.693777 | -2.105042 | 0.035287683 | 0.28416534 |
| *ENSG00000237773* | -3.019101 | 1.434225 | -2.10504 | 0.035287857 | 0.28416534 |
| *DBF4B* | -2.033716 | 0.966183 | -2.104897 | 0.035300307 | 0.28416534 |
| *COG3* | -1.283755 | 0.609902 | -2.104855 | 0.035303957 | 0.28416534 |
| *KBTBD8* | -2.883484 | 1.370186 | -2.104447 | 0.03533951 | 0.28416534 |
| *PECR* | -1.6611 | 0.789446 | -2.104135 | 0.035366703 | 0.28416534 |
| *CASZ1* | -2.315979 | 1.100805 | -2.103896 | 0.035387548 | 0.28416534 |
| *LRP2BP* | -2.475845 | 1.176938 | -2.103632 | 0.035410525 | 0.28416534 |
| *ZNF879* | -1.565388 | 0.744934 | -2.101378 | 0.035607783 | 0.28552628 |
| *METTL1* | -2.758884 | 1.313104 | -2.10104 | 0.035637437 | 0.28554219 |
| *CDC42EP3* | -1.405964 | 0.669338 | -2.100529 | 0.035682376 | 0.28568047 |
| *ENSG00000273080* | -3.354857 | 1.597591 | -2.099947 | 0.035733489 | NA |
| *GEMIN4* | -2.193471 | 1.044655 | -2.099709 | 0.035754421 | 0.28603537 |
| *CYB5RL* | -3.219008 | 1.533693 | -2.098861 | 0.035829152 | 0.28637307 |
| *LIPT1* | -1.314786 | 0.626506 | -2.0986 | 0.035852133 | 0.28637307 |
| *ENSG00000231025* | -2.388297 | 1.138459 | -2.097834 | 0.035919851 | 0.28658821 |
| *MANEA* | -2.722434 | 1.297961 | -2.09747 | 0.035951975 | 0.28658821 |
| *GSDME* | -1.932204 | 0.921447 | -2.096923 | 0.036000408 | 0.28658821 |
| *PANX1* | -3.825042 | 1.825147 | -2.095745 | 0.036104853 | NA |
| *ZNF749* | -3.417112 | 1.630683 | -2.09551 | 0.036125714 | 0.28658821 |
| *ARMCX2* | -1.673141 | 0.798495 | -2.095369 | 0.036138236 | 0.28658821 |
| *TAF5* | -2.063949 | 0.985028 | -2.09532 | 0.036142525 | 0.28658821 |
| *MMUT* | -2.381482 | 1.136685 | -2.095112 | 0.036161054 | 0.28658821 |
| *ENSG00000271789* | -2.202368 | 1.051368 | -2.094764 | 0.036192018 | 0.28658821 |
| *ZNF599* | -2.642397 | 1.261463 | -2.094707 | 0.036197016 | 0.28658821 |
| *LAMB1* | -2.115379 | 1.009935 | -2.094569 | 0.036209354 | 0.28658821 |
| *PLTP* | -2.906522 | 1.388571 | -2.093175 | 0.036333576 | 0.28658821 |
| *ADORA2A.AS1* | -1.593533 | 0.761328 | -2.093097 | 0.036340498 | 0.28658821 |
| *RRP1B* | -1.410395 | 0.673869 | -2.092981 | 0.03635086 | 0.28658821 |
| *CALHM2* | -1.771256 | 0.84637 | -2.092767 | 0.03636995 | 0.28658821 |
| *FAM200A* | -1.644075 | 0.785826 | -2.092162 | 0.036424069 | 0.28658821 |
| *PCNA* | -1.492774 | 0.71353 | -2.092095 | 0.036429984 | 0.28658821 |
| *YEATS2* | -2.382427 | 1.138842 | -2.091973 | 0.036440924 | 0.28658821 |
| *ENSG00000270562* | -1.865959 | 0.892489 | -2.090737 | NA | NA |
| *SYCE1L* | -1.601403 | 0.765982 | -2.090653 | 0.036559185 | 0.28658821 |
| *AP1AR* | -1.480424 | 0.708118 | -2.090647 | 0.036559737 | 0.28658821 |
| *ARMCX4* | -2.530733 | 1.210539 | -2.090584 | 0.0365654 | 0.28658821 |
| *GBF1* | -2.142781 | 1.025191 | -2.090128 | 0.03660627 | 0.28658821 |
| *GRAP2* | -1.813254 | 0.867628 | -2.089898 | NA | NA |
| *GMIP* | -1.111247 | 0.53184 | -2.08944 | 0.036668158 | 0.28658821 |
| *LINC02656* | -2.82312 | 1.35131 | -2.089173 | 0.036692158 | 0.28658821 |
| *WDR66* | -2.302275 | 1.102075 | -2.089037 | 0.036704437 | 0.28658821 |
| *APTR* | -2.355959 | 1.127847 | -2.088899 | 0.036716802 | 0.28658821 |
| *ACAD9* | -1.85274 | 0.886951 | -2.088886 | 0.036717994 | 0.28658821 |

| | | | | | |
|---|---|---|---|---|---|
| *AP5B1* | -2.179607 | 1.043522 | -2.088702 | 0.03673454 | 0.28658821 |
| *PACS2* | -2.825343 | 1.35295 | -2.088283 | 0.036772283 | 0.28658821 |
| *MIB1* | -1.530265 | 0.732794 | -2.08826 | 0.036774369 | 0.28658821 |
| *ACSM1* | -2.961548 | 1.418254 | -2.088164 | 0.036783 | 0.28658821 |
| *VAV3.AS1* | -2.410185 | 1.154593 | -2.087476 | 0.036845171 | 0.28658821 |
| *CCNG2* | -1.297397 | 0.621529 | -2.08743 | 0.036849309 | 0.28658821 |
| *LPCAT4* | -1.401284 | 0.67133 | -2.087325 | 0.036858773 | 0.28658821 |
| *SEC31B* | -2.028511 | 0.971889 | -2.087184 | 0.036871537 | 0.28658821 |
| *ALG9* | -2.079611 | 0.996426 | -2.087071 | 0.036881736 | 0.28658821 |
| *AUTS2* | -2.258063 | 1.082053 | -2.086833 | 0.036903243 | 0.28658821 |
| *SURF2* | -1.274405 | 0.610711 | -2.086756 | 0.036910224 | 0.28658821 |
| *ENSG00000205664* | -2.025232 | 0.970641 | -2.086489 | 0.036934333 | 0.28658821 |
| *ITGAV* | -3.90844 | 1.873528 | -2.086139 | 0.036966058 | NA |
| *HIST1H2BK* | -2.176652 | 1.043516 | -2.085883 | 0.036989234 | 0.28679857 |
| *ENSG00000275764* | -1.887468 | 0.905229 | -2.085072 | 0.037062771 | 0.287153 |
| *ENSG00000271380* | -3.104498 | 1.489654 | -2.08404 | 0.037156548 | 0.2876636 |
| *HECTD4* | -1.432117 | 0.687642 | -2.082648 | 0.037283339 | 0.28807502 |
| *RP2* | -1.477783 | 0.709578 | -2.082621 | 0.037285754 | 0.28807502 |
| *ENSG00000273156* | -1.575845 | 0.756695 | -2.082537 | 0.037293433 | 0.28807502 |
| *TMEM184B* | -2.131184 | 1.023545 | -2.08216 | 0.037327883 | 0.28812547 |
| *EVA1B* | -2.490347 | 1.196827 | -2.08079 | 0.037453097 | 0.2888759 |
| *NSDHL* | -1.633383 | 0.78523 | -2.080133 | 0.03751332 | 0.28891121 |
| *ANKRD36* | -1.286668 | 0.618704 | -2.079618 | 0.037560543 | 0.28891121 |
| *ENSG00000249141* | -3.743599 | 1.800146 | -2.079608 | 0.037561499 | 0.28891121 |
| *SNAPC1* | -1.304518 | 0.627317 | -2.079519 | 0.037569656 | 0.28891121 |
| *SHROOM1* | -3.701237 | 1.780385 | -2.078897 | 0.037626779 | NA |
| *TEDC1* | -2.702136 | 1.300073 | -2.07845 | 0.037667942 | 0.28945135 |
| *SLC25A16* | -2.166005 | 1.042499 | -2.077704 | 0.037736611 | 0.28976326 |
| *SECISBP2L* | -1.097146 | 0.528286 | -2.076803 | 0.037819789 | 0.28991802 |
| *ENSG00000273759* | -3.259214 | 1.569743 | -2.076272 | 0.037868758 | 0.28991802 |
| *ARL5B* | -1.533169 | 0.738543 | -2.075937 | 0.03789974 | 0.28991802 |
| *CHD3* | -1.334779 | 0.643031 | -2.075761 | 0.037916017 | 0.28991802 |
| *SOCS7* | -2.219978 | 1.06954 | -2.075637 | 0.037927557 | 0.28991802 |
| *ITGA6.AS1* | -1.606196 | 0.773944 | -2.075338 | 0.037955225 | 0.28991802 |
| *ZNF844* | -1.35501 | 0.653044 | -2.074913 | 0.037994618 | 0.28991802 |
| *ENSG00000262049* | -1.275486 | 0.614762 | -2.074765 | 0.038008361 | 0.28991802 |
| *ZNF717* | -1.563199 | 0.753704 | -2.074022 | NA | NA |
| *STK33* | -2.013881 | 0.971014 | -2.073997 | 0.038079554 | 0.28991802 |
| *ENSG00000272764* | -2.086219 | 1.005945 | -2.07389 | 0.038089551 | 0.28991802 |
| *NRSN2.AS1* | -2.630958 | 1.268725 | -2.073703 | 0.038106919 | 0.28991802 |
| *RPL18A* | 0.154175 | 0.074369 | 2.073098 | 0.038163134 | 0.28991802 |
| *STK36* | -2.819617 | 1.360239 | -2.072884 | 0.038183093 | 0.28991802 |
| *TRAPPC12.AS1* | -2.039723 | 0.984265 | -2.072332 | 0.038234493 | 0.28991802 |
| *CRAT* | -1.868533 | 0.901669 | -2.072304 | 0.038237071 | 0.28991802 |
| *MFSD3* | -2.01901 | 0.974346 | -2.072171 | 0.03824954 | 0.28991802 |
| *TBC1D9B* | -1.250736 | 0.603816 | -2.071384 | 0.038322927 | 0.28991802 |
| *FAM76A* | -1.211797 | 0.585026 | -2.071355 | 0.038325627 | 0.28991802 |

| | | | | | |
|---|---|---|---|---|---|
| *POGLUT3* | -3.524144 | 1.701469 | -2.071236 | 0.038336708 | NA |
| *LAT* | -2.530352 | 1.221948 | -2.070753 | 0.038381918 | 0.28991802 |
| *VSTM4* | -2.489895 | 1.202676 | -2.070295 | 0.038424719 | 0.28991802 |
| *ZNF787* | -1.747471 | 0.844308 | -2.069709 | 0.038479568 | 0.28991802 |
| *DIABLO* | -2.490364 | 1.203323 | -2.069573 | 0.038492364 | 0.28991802 |
| *VPS54* | -2.313261 | 1.117842 | -2.069398 | 0.038508747 | 0.28991802 |
| *CUL7* | -2.975183 | 1.437916 | -2.069093 | 0.038537314 | 0.28991802 |
| *ENSG00000266844* | -3.390136 | 1.638581 | -2.068946 | 0.038551153 | NA |
| *LINC00893* | -1.994994 | 0.964426 | -2.068581 | 0.038585416 | 0.28991802 |
| *IFI44* | -2.979831 | 1.440596 | -2.068471 | 0.038595771 | 0.28991802 |
| *C3orf18* | -3.782926 | 1.828906 | -2.068409 | 0.038601537 | NA |
| *RBL1* | -1.606741 | 0.776916 | -2.068102 | 0.038630443 | 0.28991802 |
| *HMGN4* | -1.633237 | 0.789855 | -2.067769 | 0.038661772 | 0.28991802 |
| *CPPED1* | -1.841091 | 0.890432 | -2.067638 | 0.038674046 | 0.28991802 |
| *ENSG00000167912* | -6.197237 | 2.997612 | -2.067391 | NA | NA |
| *URB1* | -2.561106 | 1.239054 | -2.066986 | 0.038735499 | 0.28991802 |
| *KEAP1* | -1.947081 | 0.942389 | -2.066113 | 0.038817838 | 0.28991802 |
| *PIP5K1C* | -1.589616 | 0.769686 | -2.06528 | 0.038896556 | 0.28991802 |
| *ARSG* | -2.419456 | 1.171754 | -2.064816 | 0.038940451 | 0.28991802 |
| *RBM27* | -1.285049 | 0.622467 | -2.064443 | 0.038975751 | 0.28991802 |
| *NEXN* | -3.747824 | 1.8157 | -2.06412 | 0.039006299 | 0.28991802 |
| *ITPKB.AS1* | -1.885655 | 0.913568 | -2.064055 | 0.039012536 | 0.28991802 |
| *WRN* | -1.98827 | 0.963354 | -2.063904 | 0.039026863 | 0.28991802 |
| *ENSG00000261187* | -5.12956 | 2.485391 | -2.063884 | 0.039028683 | 0.28991802 |
| *TSHZ1* | -3.307164 | 1.602485 | -2.063772 | 0.039039311 | NA |
| *SYCP3* | -1.529138 | 0.741001 | -2.06361 | 0.039054668 | 0.28991802 |
| *ENSG00000249592* | -1.943846 | 0.941988 | -2.063556 | 0.03905987 | 0.28991802 |
| *MS4A1* | -0.436323 | 0.211475 | -2.063235 | 0.039090312 | 0.28991802 |
| *ZNF485* | -2.198328 | 1.065481 | -2.063226 | 0.039091201 | 0.28991802 |
| *LDAH* | -1.490725 | 0.722582 | -2.063051 | 0.039107749 | 0.28991802 |
| *RPS6KB1* | -1.194016 | 0.578864 | -2.062687 | 0.03914235 | 0.28991802 |
| *XPO4* | -1.271956 | 0.616756 | -2.062332 | 0.039176146 | 0.28991802 |
| *ACSL4* | -1.612931 | 0.782264 | -2.061876 | 0.039219584 | 0.28991802 |
| *ZNF700* | -1.806351 | 0.876299 | -2.061342 | 0.039270427 | 0.28991802 |
| *ENSG00000275441* | -1.713604 | 0.831477 | -2.060917 | 0.039310946 | 0.28991802 |
| *SLC28A2* | -3.280279 | 1.59197 | -2.060516 | 0.039349255 | 0.28991802 |
| *MIOS* | -1.208637 | 0.586709 | -2.060028 | 0.039395881 | 0.28991802 |
| *ZNF117* | -2.635685 | 1.279472 | -2.059979 | 0.039400533 | 0.28991802 |
| *BBS4* | -1.345742 | 0.65333 | -2.05982 | 0.039415772 | 0.28991802 |
| *CASK* | -2.718444 | 1.319814 | -2.059717 | 0.039425558 | 0.28991802 |
| *PFAS* | -2.450695 | 1.189893 | -2.059593 | 0.039437454 | 0.28991802 |
| *ZNF202* | -1.938747 | 0.9415 | -2.05921 | 0.039474082 | 0.28991802 |
| *HIST1H2AE* | -2.949254 | 1.432537 | -2.058763 | 0.039516962 | 0.28991802 |
| *TMEM220.AS1* | -2.704424 | 1.313857 | -2.058384 | 0.039553244 | 0.28991802 |
| *LNX1* | -2.273956 | 1.104734 | -2.058374 | 0.039554259 | 0.28991802 |
| *CMTR1* | -1.371876 | 0.666726 | -2.057629 | 0.03962572 | 0.28991802 |
| *CLEC16A* | -1.853821 | 0.901132 | -2.057213 | 0.039665729 | 0.28991802 |

| | | | | | |
|---|---|---|---|---|---|
| *ENSG00000258376* | -3.259684 | 1.584595 | -2.057109 | 0.039675767 | 0.28991802 |
| *BRCA2* | -1.448533 | 0.704238 | -2.056881 | 0.039697683 | 0.28991802 |
| *PRR22* | -2.878235 | 1.399503 | -2.056612 | 0.039723533 | 0.28991802 |
| *FAM110A* | -1.247702 | 0.606757 | -2.056344 | 0.039749374 | 0.28991802 |
| *LMNB2* | -3.205213 | 1.558718 | -2.056314 | 0.039752266 | 0.28991802 |
| *CPT2* | -1.340737 | 0.652228 | -2.055625 | 0.039818664 | 0.28991802 |
| *XPNPEP1* | -1.347479 | 0.655556 | -2.055474 | 0.039833268 | 0.28991802 |
| *NBEAL2* | -2.50071 | 1.216687 | -2.055344 | 0.039845827 | 0.28991802 |
| *FBXO33* | -1.230538 | 0.598806 | -2.054987 | 0.039880254 | 0.28991802 |
| *NAGA* | -2.673746 | 1.301456 | -2.054427 | 0.039934418 | 0.28991802 |
| *ERCC4* | -1.455863 | 0.708739 | -2.054161 | 0.039960106 | 0.28991802 |
| *ZNF816* | -1.42362 | 0.693099 | -2.053992 | 0.039976476 | 0.28991802 |
| *ENSG00000259520* | -5.200321 | 2.532083 | -2.053772 | NA | NA |
| *ADHFE1* | -2.621527 | 1.276527 | -2.053639 | 0.040010635 | 0.28991802 |
| *C8orf58* | -1.681655 | 0.81887 | -2.053627 | 0.040011761 | 0.28991802 |
| *PDP1* | -1.586428 | 0.7726 | -2.053363 | 0.040037345 | 0.28991802 |
| *WDR82* | -1.149837 | 0.55998 | -2.053351 | 0.040038527 | 0.28991802 |
| *ENSG00000267787* | -2.436066 | 1.186815 | -2.052609 | 0.040110501 | 0.28991802 |
| *NEIL2* | -1.716965 | 0.836591 | -2.052336 | 0.040137049 | 0.28991802 |
| *MAU2* | -1.23722 | 0.602935 | -2.051994 | 0.040170289 | 0.28991802 |
| *ZNF609* | -1.519703 | 0.740616 | -2.051946 | 0.040174917 | 0.28991802 |
| *FLYWCH1* | -1.31781 | 0.642256 | -2.051844 | 0.04018486 | 0.28991802 |
| *ATPSCKMT* | -2.753117 | 1.34204 | -2.051441 | 0.040223979 | 0.28991802 |
| *SLC1A4* | -3.106136 | 1.514179 | -2.051367 | 0.040231254 | 0.28991802 |
| *CREG1* | -2.904673 | 1.415981 | -2.05135 | 0.040232851 | 0.28991802 |
| *IQCG* | -1.554443 | 0.757793 | -2.051276 | 0.040240058 | 0.28991802 |
| *KIAA0391* | -3.794815 | 1.850004 | -2.051247 | 0.040242905 | NA |
| *NPC1* | -1.454611 | 0.709184 | -2.051105 | 0.040256744 | 0.28991802 |
| *ZNF543* | -1.857633 | 0.905692 | -2.051064 | 0.040260742 | 0.28991802 |
| *WHRN* | -3.125076 | 1.523737 | -2.050929 | 0.040273876 | 0.28991802 |
| *MTOR* | -2.493808 | 1.215977 | -2.050868 | 0.040279799 | 0.28991802 |
| *BRD3OS* | -1.249926 | 0.609657 | -2.050214 | 0.040343593 | 0.28991802 |
| *KDM8* | -2.864574 | 1.397267 | -2.050127 | 0.040352076 | 0.28991802 |
| *ENSG00000124593* | -2.655873 | 1.29547 | -2.050123 | 0.040352405 | 0.28991802 |
| *ELK1* | -1.172311 | 0.571873 | -2.049949 | 0.040369399 | 0.28991802 |
| *CSTF3.DT* | -3.704703 | 1.807501 | -2.049627 | 0.040400813 | NA |
| *ACOX3* | -1.875283 | 0.91534 | -2.04873 | 0.040488539 | 0.29019907 |
| *RRM1* | -1.380433 | 0.673813 | -2.048688 | 0.040492644 | 0.29019907 |
| *TTL* | -1.576569 | 0.769667 | -2.048379 | 0.040522877 | 0.29019907 |
| *ZNF254* | -1.229011 | 0.600039 | -2.048218 | 0.040538672 | 0.29019907 |
| *NEMP2* | -1.962051 | 0.958059 | -2.047943 | 0.040565582 | 0.29019907 |
| *PCGF6* | -1.276388 | 0.623373 | -2.047551 | 0.040603974 | 0.29019907 |
| *NPHP3* | -1.281532 | 0.625953 | -2.047329 | 0.040625805 | 0.29019907 |
| *ZNF287* | -1.633051 | 0.79768 | -2.047251 | 0.040633494 | 0.29019907 |
| *PLBD2* | -3.244654 | 1.584972 | -2.047136 | 0.04064469 | NA |
| *ZYX* | -1.35746 | 0.663637 | -2.045486 | 0.040806962 | 0.29035871 |
| *EPG5* | -1.297722 | 0.634656 | -2.044766 | 0.040877904 | 0.29035871 |

| | | | | | |
|---|---|---|---|---|---|
| *ENSG00000277283* | -1.943193 | 0.950378 | -2.044652 | 0.040889149 | 0.29035871 |
| *GINS2* | -2.362231 | 1.155434 | -2.044453 | 0.04090879 | 0.29035871 |
| *KIAA1143* | -1.088387 | 0.532399 | -2.044305 | 0.040923386 | 0.29035871 |
| *IGHMBP2* | -1.677002 | 0.820372 | -2.044197 | 0.040934105 | 0.29035871 |
| *ENSG00000228427* | -1.935039 | 0.946601 | -2.044196 | 0.040934145 | 0.29035871 |
| *UBLCP1* | -1.374661 | 0.672482 | -2.044162 | 0.040937573 | 0.29035871 |
| *MYO9A* | -1.301067 | 0.636605 | -2.043759 | 0.040977331 | 0.29035871 |
| *B3GNT2* | -1.277757 | 0.625203 | -2.043749 | 0.04097833 | 0.29035871 |
| *MAPK14* | -1.330257 | 0.65092 | -2.043658 | 0.040987333 | 0.29035871 |
| *LRRC34* | -2.899366 | 1.418895 | -2.043397 | 0.04101314 | 0.29035871 |
| *NFYC.AS1* | -2.026785 | 0.992062 | -2.043002 | 0.041052255 | 0.29035871 |
| *ZNF718* | -2.389407 | 1.169558 | -2.042999 | 0.041052515 | 0.29035871 |
| *TONSL* | -2.377811 | 1.164029 | -2.042743 | 0.04107788 | 0.29035871 |
| *ZNF623* | -1.563911 | 0.766038 | -2.041557 | 0.041195466 | 0.29087564 |
| *KRBA1* | -2.376085 | 1.163928 | -2.041437 | 0.041207383 | 0.29087564 |
| *RPP14* | -1.268741 | 0.621809 | -2.040404 | 0.041310142 | 0.29133865 |
| *ENSG00000273456* | -3.553166 | 1.74163 | -2.040139 | 0.041336496 | NA |
| *RND1* | -2.426378 | 1.189452 | -2.039913 | 0.041359036 | 0.29133865 |
| *ADAM12* | -3.331066 | 1.633396 | -2.03935 | 0.041415122 | NA |
| *MEGF6* | -1.886625 | 0.925221 | -2.039107 | 0.041439386 | 0.29133865 |
| *IGLV6.57* | -2.979745 | 1.461645 | -2.038625 | 0.041487469 | 0.29133865 |
| *ERMARD* | -2.130004 | 1.044825 | -2.038622 | 0.041487735 | 0.29133865 |
| *WWP2* | -1.258519 | 0.617475 | -2.03817 | 0.041532975 | 0.29133865 |
| *ENSG00000273270* | -2.138425 | 1.049332 | -2.037893 | 0.041560678 | 0.29133865 |
| *RPE* | -1.346356 | 0.660746 | -2.037628 | 0.0415871 | 0.29133865 |
| *B3GNT7* | -2.228386 | 1.093638 | -2.03759 | 0.041590982 | 0.29133865 |
| *ASB7* | -1.705413 | 0.837029 | -2.037459 | 0.041604051 | 0.29133865 |
| *SLC35G2* | -2.491639 | 1.222948 | -2.037403 | 0.041609663 | 0.29133865 |
| *HMBS* | -1.604177 | 0.787453 | -2.037173 | 0.041632756 | 0.29133865 |
| *RALGDS* | -1.38809 | 0.681405 | -2.0371 | 0.041639972 | 0.29133865 |
| *NBAS* | -1.809157 | 0.888341 | -2.036556 | 0.041694549 | 0.29134263 |
| *INTS2* | -2.362106 | 1.159948 | -2.036389 | 0.041711315 | 0.29134263 |
| *ZMYND19* | -3.417364 | 1.67825 | -2.036267 | 0.041723601 | NA |
| *NPIPB4* | -2.84102 | 1.395435 | -2.035938 | 0.041756557 | 0.29134263 |
| *MINPP1* | -1.610451 | 0.791107 | -2.035693 | 0.041781212 | 0.29134263 |
| *FARP2* | -1.239332 | 0.608906 | -2.035342 | 0.041816523 | 0.29134263 |
| *ZFP30* | -1.865538 | 0.916662 | -2.035143 | 0.04183649 | 0.29134263 |
| *C22orf46* | -1.524151 | 0.749103 | -2.034636 | 0.041887483 | 0.29134263 |
| *MAP1LC3A* | -1.71318 | 0.842023 | -2.0346 | 0.041891186 | 0.29134263 |
| *ROR1* | -1.879798 | 0.923931 | -2.034565 | 0.041894619 | 0.29134263 |
| *KLHDC4* | -1.189946 | 0.585081 | -2.033815 | 0.041970228 | 0.29167189 |
| *PABPC3* | -1.862856 | 0.915963 | -2.033767 | NA | NA |
| *ZP3* | -2.311222 | 1.13665 | -2.033363 | 0.042015864 | 0.29179254 |
| *ACADS* | -2.357753 | 1.159705 | -2.033063 | 0.04204612 | 0.2918063 |
| *SEPSECS.AS1* | -2.090767 | 1.028619 | -2.032596 | 0.042093326 | 0.29193759 |
| *HCFC1R1* | -1.055373 | 0.51931 | -2.032262 | 0.042127187 | 0.29197621 |
| *CNNM2* | -2.170912 | 1.06894 | -2.030902 | 0.042264972 | 0.29271842 |

| | | | | | |
|---|---|---|---|---|---|
| *C2orf15* | -2.686734 | 1.323223 | -2.030446 | 0.042311185 | 0.29271842 |
| *IMMP2L* | -1.625967 | 0.801078 | -2.029724 | 0.042384621 | 0.29271842 |
| *ENSG00000203325* | -2.662513 | 1.311884 | -2.029535 | 0.04240384 | 0.29271842 |
| *ARNTL* | -2.308282 | 1.137467 | -2.029318 | 0.042425871 | 0.29271842 |
| *ENSG00000228835* | -1.593909 | 0.785555 | -2.029021 | 0.042456119 | 0.29271842 |
| *ARHGAP32* | -2.669827 | 1.316669 | -2.027714 | 0.042589426 | 0.29271842 |
| *FAM193B* | -1.445692 | 0.712998 | -2.027624 | 0.042598624 | 0.29271842 |
| *MKS1* | -3.416607 | 1.68513 | -2.027504 | 0.042610939 | NA |
| *DIPK1B* | -2.282881 | 1.126135 | -2.027183 | 0.042643721 | 0.29271842 |
| *MFN2* | -1.666436 | 0.822053 | -2.027165 | 0.042645591 | 0.29271842 |
| *PIK3CB* | -2.269789 | 1.119711 | -2.027121 | 0.042650016 | 0.29271842 |
| *CASP9* | -2.033561 | 1.003244 | -2.026985 | 0.04266399 | 0.29271842 |
| *RAB12* | -2.758517 | 1.360955 | -2.026898 | 0.042672801 | 0.29271842 |
| *TTBK2* | -1.234591 | 0.609207 | -2.026554 | 0.042708058 | 0.29271842 |
| *C20orf194* | -1.928743 | 0.951764 | -2.026493 | 0.042714307 | 0.29271842 |
| *PGM3* | -1.267194 | 0.625332 | -2.026433 | 0.042720461 | 0.29271842 |
| *CPSF4* | -1.189374 | 0.586978 | -2.026267 | 0.042737477 | 0.29271842 |
| *TMEM39A* | -1.424882 | 0.703253 | -2.026131 | 0.042751381 | 0.29271842 |
| *TEF* | -2.573064 | 1.270073 | -2.025918 | 0.042773195 | 0.29271842 |
| *TMEM38A* | -1.860599 | 0.918888 | -2.024838 | 0.04288405 | 0.29328257 |
| *ACP2* | -2.412726 | 1.191941 | -2.0242 | 0.042949625 | 0.2933847 |
| *LYSMD3* | -1.620099 | 0.800379 | -2.024164 | NA | NA |
| *C1RL* | -2.287603 | 1.130296 | -2.023898 | 0.042980654 | 0.2933847 |
| *DDIT4* | 1.274923 | 0.630091 | 2.023394 | 0.043032556 | 0.2933847 |
| *GPR157* | -2.836623 | 1.402167 | -2.023027 | 0.043070379 | 0.2933847 |
| *MSRB2* | -2.050391 | 1.013777 | -2.022525 | 0.043122102 | 0.2933847 |
| *NREP* | -1.29653 | 0.641145 | -2.022211 | 0.043154535 | 0.2933847 |
| *ASPH* | -3.123278 | 1.544562 | -2.022112 | 0.043164806 | 0.2933847 |
| *AKAP5* | -2.346152 | 1.160524 | -2.021632 | 0.04321436 | 0.2933847 |
| *CIP2A* | -1.741551 | 0.861464 | -2.021618 | 0.043215791 | 0.2933847 |
| *HIP1* | -1.738912 | 0.860205 | -2.021509 | 0.043227145 | 0.2933847 |
| *INPP5F* | -2.284918 | 1.130472 | -2.021207 | 0.043258358 | 0.2933847 |
| *MAP3K20* | -1.51476 | 0.749524 | -2.020963 | 0.043283585 | 0.2933847 |
| *GGT7* | -2.569151 | 1.271702 | -2.020246 | 0.043357853 | 0.2933847 |
| *SLC16A13* | -1.451539 | 0.71852 | -2.020178 | 0.043364915 | 0.2933847 |
| *PANK4* | -1.829412 | 0.905578 | -2.020159 | 0.043366854 | 0.2933847 |
| *AP3S2* | -1.661583 | 0.822514 | -2.020128 | 0.043370113 | 0.2933847 |
| *PGGHG* | -1.13419 | 0.561485 | -2.019983 | 0.043385143 | 0.2933847 |
| *GRK5* | -1.241947 | 0.615108 | -2.019071 | 0.043479806 | 0.2933847 |
| *AUH* | -1.255072 | 0.621655 | -2.018921 | 0.043495395 | 0.2933847 |
| *MTM1* | -1.629766 | 0.807278 | -2.018841 | 0.043503756 | 0.2933847 |
| *USP40* | -2.372032 | 1.175364 | -2.018125 | 0.04357824 | 0.2933847 |
| *VTA1* | -1.25579 | 0.62231 | -2.017948 | 0.043596678 | 0.2933847 |
| *SLC38A6* | -2.680845 | 1.328598 | -2.0178 | 0.043612089 | 0.2933847 |
| *SPINT1* | -2.588621 | 1.283107 | -2.017463 | 0.043647177 | 0.2933847 |
| *ZSCAN9* | -1.460657 | 0.724013 | -2.017446 | 0.043648947 | 0.2933847 |
| *ZFYVE21* | -1.186397 | 0.588077 | -2.017417 | 0.043652058 | 0.2933847 |

| | | | | | |
|---|---|---|---|---|---|
| *BCOR* | -1.219746 | 0.60465 | -2.017278 | 0.04366656 | 0.2933847 |
| *PON2* | -3.35896 | 1.665331 | -2.016992 | 0.0436963 | 0.2933935 |
| *KBTBD4* | -1.203012 | 0.596623 | -2.016368 | 0.043761461 | 0.29360618 |
| *RANBP10* | -1.480274 | 0.73421 | -2.016144 | 0.043784875 | 0.29360618 |
| *NINJ1* | -1.979283 | 0.982303 | -2.014942 | 0.043910726 | 0.29425889 |
| *GDPD3* | -3.239654 | 1.608328 | -2.0143 | 0.04397805 | NA |
| *ENSG00000258572* | -3.380427 | 1.679289 | -2.013011 | 0.044113503 | NA |
| *AGER* | -1.656427 | 0.822881 | -2.01296 | 0.044118885 | 0.29485671 |
| *ENSG00000277959* | -2.631599 | 1.307454 | -2.012766 | 0.044139232 | 0.29485671 |
| *DBR1* | -1.732625 | 0.861096 | -2.012115 | 0.044207861 | 0.29485671 |
| *MED26* | -1.512151 | 0.751561 | -2.012014 | 0.044218472 | 0.29485671 |
| *THAP7.AS1* | -1.637574 | 0.813977 | -2.011819 | 0.044239066 | 0.29485671 |
| *IFT74* | -1.390212 | 0.691027 | -2.011806 | 0.044240361 | 0.29485671 |
| *SLC1A5* | -2.582367 | 1.283611 | -2.011799 | 0.044241174 | 0.29485671 |
| *UBXN2B* | -1.5019 | 0.746549 | -2.011791 | 0.044241971 | 0.29485671 |
| *EEF1AKNMT* | -1.269679 | 0.631239 | -2.011409 | 0.044282297 | 0.29485671 |
| *SLC15A2* | -2.119299 | 1.053656 | -2.011377 | 0.044285649 | 0.29485671 |
| *BCO2* | -2.482749 | 1.234735 | -2.010754 | 0.044351448 | 0.29493494 |
| *LYSMD1* | -3.219276 | 1.601052 | -2.010725 | 0.044354558 | 0.29493494 |
| *FBXL22* | -2.167295 | 1.078205 | -2.010096 | 0.044420978 | 0.29501811 |
| *CASP1* | -1.209252 | 0.601612 | -2.01002 | 0.044429069 | 0.29501811 |
| *OR2A1.AS1* | -2.223365 | 1.106306 | -2.00972 | 0.044460807 | 0.29501811 |
| *WNT2B* | -2.253869 | 1.121735 | -2.009271 | 0.044508384 | 0.29501811 |
| *IFT81* | -2.518368 | 1.253384 | -2.009256 | 0.044509999 | 0.29501811 |
| *TPCN2* | -2.389267 | 1.189861 | -2.008022 | 0.044640933 | 0.29534326 |
| *FMO5* | -2.439071 | 1.214921 | -2.007596 | 0.044686261 | 0.29534326 |
| *SETD1B* | -1.37084 | 0.68285 | -2.007527 | 0.044693569 | 0.29534326 |
| *VPS39* | -2.629211 | 1.309796 | -2.007343 | 0.044713131 | 0.29534326 |
| *IGF1R* | -2.483717 | 1.237447 | -2.00713 | 0.044735797 | 0.29534326 |
| *ENSG00000167807* | -2.971341 | 1.480519 | -2.00696 | 0.044753955 | 0.29534326 |
| *ENSG00000256448* | -3.035725 | 1.512637 | -2.006909 | 0.044759385 | 0.29534326 |
| *GPAM* | -2.586904 | 1.289338 | -2.006382 | 0.044815535 | 0.29539158 |
| *MIR762HG* | -1.960423 | 0.977132 | -2.006303 | 0.044823954 | 0.29539158 |
| *CCDC102B* | -2.458514 | 1.225954 | -2.005387 | 0.04492165 | 0.29553707 |
| *VTI1A* | -1.129692 | 0.563493 | -2.0048 | 0.044984405 | 0.29553707 |
| *MX1* | -1.404616 | 0.700685 | -2.004634 | 0.045002187 | 0.29553707 |
| *SNX25* | -1.344006 | 0.670475 | -2.004559 | 0.045010167 | 0.29553707 |
| *BEND7* | -1.566922 | 0.781808 | -2.004228 | 0.045045692 | 0.29553707 |
| *TP53BP2* | -1.688315 | 0.84254 | -2.00384 | 0.04508721 | 0.29553707 |
| *ROM1* | -3.088515 | 1.541617 | -2.003425 | 0.045131712 | 0.29553707 |
| *CCDC157* | -2.562511 | 1.279336 | -2.003001 | 0.045177225 | 0.29553707 |
| *ENSG00000273723* | -1.589411 | 0.793682 | -2.002578 | 0.045222589 | 0.29553707 |
| *DPH6* | -2.745269 | 1.370991 | -2.002398 | 0.045241953 | 0.29553707 |
| *SENP3* | -1.385301 | 0.691926 | -2.002094 | 0.045274607 | 0.29553707 |
| *ALG10B* | -2.494353 | 1.245925 | -2.002009 | 0.04528374 | 0.29553707 |
| *NHLRC2* | -1.246961 | 0.622893 | -2.001885 | 0.04529713 | 0.29553707 |
| *DNASE1L1* | -1.642964 | 0.820718 | -2.001862 | 0.045299618 | 0.29553707 |

| | | | | | |
|---|---|---|---|---|---|
| *AXIN1* | -1.574093 | 0.786329 | -2.001826 | 0.045303472 | 0.29553707 |
| *KLK4* | -3.269269 | 1.633471 | -2.001425 | 0.045346606 | 0.29553707 |
| *B3GLCT* | -1.99768 | 0.998143 | -2.001397 | 0.045349674 | 0.29553707 |
| *ERI1* | -1.376624 | 0.687897 | -2.001207 | 0.045370094 | 0.29553707 |
| *TTN.AS1* | -1.616487 | 0.807923 | -2.000793 | 0.045414689 | 0.29553707 |
| *JAM3* | -1.945976 | 0.97286 | -2.000263 | 0.045471875 | 0.29553707 |
| *ENSG00000278238* | -2.579802 | 1.289825 | -2.000118 | 0.04548755 | 0.29553707 |
| *CRYBG1* | -1.263326 | 0.631642 | -2.000066 | 0.045493095 | 0.29553707 |
| *C5orf34* | -2.45206 | 1.226897 | -1.998587 | 0.045653059 | 0.29553707 |
| *PSD3* | -1.612943 | 0.807062 | -1.998536 | 0.045658594 | 0.29553707 |
| *FLYWCH2* | -1.119727 | 0.560323 | -1.998361 | 0.045677532 | 0.29553707 |
| *RXRA* | -2.836599 | 1.419499 | -1.998311 | 0.045682995 | 0.29553707 |
| *SIGMAR1* | -1.46666 | 0.734072 | -1.99798 | 0.045718829 | 0.29553707 |
| *ANKRD37* | -2.122283 | 1.062505 | -1.997434 | 0.04577806 | 0.29553707 |
| *ENSG00000280433* | -3.013307 | 1.508632 | -1.997377 | 0.045784273 | 0.29553707 |
| *DCAF4* | -1.663835 | 0.833065 | -1.997244 | NA | NA |
| *ZNF8* | -2.509938 | 1.256751 | -1.997164 | 0.045807334 | 0.29553707 |
| *CYP2U1* | -1.743108 | 0.872924 | -1.996863 | 0.045840115 | 0.29553707 |
| *ZNF837* | -2.283674 | 1.143785 | -1.996593 | 0.045869448 | 0.29553707 |
| *ENSG00000256591* | -3.191188 | 1.59844 | -1.996439 | 0.045886189 | NA |
| *KLHL21* | -2.099898 | 1.051868 | -1.996351 | 0.0458957 | 0.29553707 |
| *FANCD2* | -1.369955 | 0.686276 | -1.996216 | 0.0459104 | 0.29553707 |
| *TMEM87B* | -1.635191 | 0.819151 | -1.996202 | 0.045911895 | 0.29553707 |
| *SRR* | -3.497003 | 1.751929 | -1.996087 | 0.045924406 | NA |
| *ENSG00000266340* | -2.604756 | 1.305085 | -1.995852 | 0.045950049 | 0.29553707 |
| *PMS2* | -1.207678 | 0.605101 | -1.99583 | 0.045952471 | 0.29553707 |
| *RANBP9* | -1.321454 | 0.662146 | -1.995716 | 0.045964893 | 0.29553707 |
| *TESK1* | -1.867006 | 0.935689 | -1.995328 | 0.04600715 | 0.29553707 |
| *UBR7* | -1.329842 | 0.666511 | -1.99523 | 0.04601777 | 0.29553707 |
| *CHUK* | -2.103473 | 1.054344 | -1.995054 | 0.046037044 | 0.29553707 |
| *DISP1* | -2.023605 | 1.014366 | -1.994946 | 0.0460488 | 0.29553707 |
| *ENSG00000272990* | -2.567262 | 1.288008 | -1.993204 | 0.046239143 | 0.29602119 |
| *ZNF219* | -3.752319 | 1.883127 | -1.9926 | 0.04630523 | NA |
| *EDRF1* | -1.194914 | 0.599743 | -1.992376 | 0.046329866 | 0.29602119 |
| *NCEH1* | -2.966327 | 1.488849 | -1.992363 | 0.046331253 | 0.29602119 |
| *PSMG1* | -1.360691 | 0.682958 | -1.992349 | 0.046332797 | 0.29602119 |
| *TBX6* | -2.251112 | 1.129986 | -1.99216 | 0.046353518 | 0.29602119 |
| *AGAP5* | -3.285068 | 1.649034 | -1.992117 | 0.046358244 | NA |
| *ACSF2* | -1.449976 | 0.727875 | -1.992067 | 0.046363652 | 0.29602119 |
| *ECI1* | -1.377265 | 0.691469 | -1.991795 | 0.04639354 | 0.29602119 |
| *C2CD3* | -1.65586 | 0.831496 | -1.991423 | 0.046434364 | 0.29602119 |
| *LMBR1* | -1.918143 | 0.963296 | -1.991229 | 0.046455689 | 0.29602119 |
| *POLI* | -2.463948 | 1.237403 | -1.991224 | 0.046456215 | 0.29602119 |
| *LINC00562* | -2.329664 | 1.170088 | -1.991015 | 0.046479237 | 0.29602119 |
| *CCDC88A* | -2.193289 | 1.10166 | -1.990895 | 0.046492445 | 0.29602119 |
| *ENSG00000242282* | -2.712111 | 1.362286 | -1.990852 | 0.046497127 | 0.29602119 |
| *ENSG00000253891* | -3.255295 | 1.635636 | -1.990232 | 0.046565405 | NA |

| | | | | | |
|---|---|---|---|---|---|
| *CAMK2G* | -1.099166 | 0.552297 | -1.990173 | 0.046571926 | 0.2963146 |
| *TMTC3* | -2.188072 | 1.099838 | -1.989449 | 0.046651686 | 0.29663919 |
| *PPIP5K1* | -3.261628 | 1.639794 | -1.989048 | 0.04669594 | NA |
| *TIGD6* | -3.773193 | 1.897014 | -1.989017 | 0.0466993 | NA |
| *KLHDC1* | -2.622193 | 1.318473 | -1.988811 | 0.046722082 | 0.29690387 |
| *SPIN4* | -3.402602 | 1.711075 | -1.988576 | 0.046748024 | NA |
| *PSPN* | -2.763661 | 1.390079 | -1.988132 | 0.046797043 | 0.2970904 |
| *SLCO4A1* | -1.577432 | 0.793602 | -1.987686 | 0.046846415 | 0.2970904 |
| *ESRRA* | -1.535545 | 0.772569 | -1.987582 | 0.046857955 | 0.2970904 |
| *ENSG00000250073* | -3.023829 | 1.521421 | -1.987504 | 0.046866586 | 0.2970904 |
| *HEATR6* | -1.55464 | 0.782324 | -1.987207 | 0.046899482 | 0.29711642 |
| *ENSG00000237310* | -2.044942 | 1.029211 | -1.986902 | 0.046933249 | 0.29714793 |
| *TBC1D19* | -2.342134 | 1.179285 | -1.986064 | 0.047026267 | 0.29714982 |
| *ARHGEF19* | -1.582342 | 0.796778 | -1.985926 | 0.047041513 | 0.29714982 |
| *RABL3* | -1.80312 | 0.907981 | -1.985856 | 0.047049282 | 0.29714982 |
| *FHL3* | -2.64703 | 1.332969 | -1.985815 | 0.047053837 | 0.29714982 |
| *DTX3* | -2.463229 | 1.240682 | -1.985383 | 0.047101848 | 0.29714982 |
| *CATSPER2* | -1.447292 | 0.729073 | -1.985114 | 0.047131784 | 0.29714982 |
| *PPP2CB* | -2.2307 | 1.123731 | -1.985084 | 0.047135102 | 0.29714982 |
| *ZNF850* | -2.951134 | 1.4869 | -1.984757 | 0.047171568 | 0.29719816 |
| *DNAJC11* | -1.726214 | 0.870115 | -1.983891 | NA | NA |
| *MGST2* | -2.509689 | 1.265293 | -1.983484 | 0.047313432 | 0.29727795 |
| *LMNTD2* | -3.208562 | 1.618013 | -1.983026 | 0.047364518 | 0.29727795 |
| *CCDC122* | -1.544881 | 0.779248 | -1.982528 | 0.047420202 | 0.29727795 |
| *ZNF184* | -1.74587 | 0.880656 | -1.982464 | 0.047427319 | 0.29727795 |
| *ENSG00000258472* | -2.630156 | 1.326759 | -1.982391 | 0.047435496 | 0.29727795 |
| *NUBPL* | -2.082468 | 1.050611 | -1.982149 | 0.047462592 | 0.29727795 |
| *FBXO30* | -1.694318 | 0.854966 | -1.981739 | 0.047508491 | 0.29727795 |
| *ENSG00000277496* | -1.988014 | 1.003198 | -1.981677 | 0.047515425 | 0.29727795 |
| *SEPSECS* | -2.157356 | 1.088692 | -1.981603 | 0.047523696 | 0.29727795 |
| *UHRF1BP1L* | -1.676913 | 0.846299 | -1.981466 | 0.047539064 | 0.29727795 |
| *THEMIS2* | -1.476122 | 0.744972 | -1.981447 | 0.047541144 | 0.29727795 |
| *ARMC7* | -1.431147 | 0.722442 | -1.980984 | 0.047593076 | 0.29727795 |
| *TBC1D31* | -2.549962 | 1.28725 | -1.980938 | 0.047598264 | 0.29727795 |
| *SLC31A2* | -3.567965 | 1.801395 | -1.980668 | 0.047628522 | NA |
| *PLCB2* | -1.892406 | 0.955559 | -1.980418 | 0.047656623 | 0.29727795 |
| *SLC39A13* | -1.594724 | 0.805275 | -1.980346 | 0.047664645 | 0.29727795 |
| *URGCP* | -2.266372 | 1.144438 | -1.980336 | 0.047665765 | 0.29727795 |
| *ENSG00000227388* | -1.646278 | 0.831401 | -1.980125 | 0.047689484 | 0.29727795 |
| *DPY19L4* | -2.448194 | 1.236457 | -1.980007 | 0.04770274 | 0.29727795 |
| *GPHN* | -2.843423 | 1.436364 | -1.979598 | 0.047748723 | 0.29737797 |
| *ENSG00000244567* | -3.80769 | 1.923579 | -1.979481 | 0.047761842 | NA |
| *ENSG00000269246* | -3.074838 | 1.553457 | -1.979352 | 0.047776422 | 0.29737797 |
| *BAHD1* | -1.786658 | 0.902853 | -1.978903 | 0.047826949 | 0.29741172 |
| *LRRC56* | -2.376893 | 1.201184 | -1.978792 | 0.047839483 | 0.29741172 |
| *VARS* | -3.51418 | 1.776336 | -1.978331 | 0.047891351 | 0.29755493 |
| *SEC24B.AS1* | -3.452465 | 1.745881 | -1.977492 | 0.047986096 | NA |

| | | | | | |
|---|---|---|---|---|---|
| *CRY1* | -2.289243 | 1.15799 | -1.976912 | 0.048051609 | 0.29807267 |
| *ATP1B3* | -1.039214 | 0.525831 | -1.976328 | 0.048117686 | 0.29807267 |
| *NAMPT* | -1.329224 | 0.672593 | -1.976268 | 0.048124393 | 0.29807267 |
| *MAD2L1* | -3.539237 | 1.791192 | -1.975912 | 0.04816478 | 0.29807267 |
| *ZNF823* | -2.829263 | 1.432098 | -1.975607 | 0.048199306 | 0.29807267 |
| *CA5B* | -1.554503 | 0.786983 | -1.975269 | 0.048237607 | 0.29807267 |
| *ENSG00000260572* | -4.122201 | 2.087097 | -1.975088 | 0.04825814 | NA |
| *CDC37L1.DT* | -2.084225 | 1.055258 | -1.975086 | 0.048258349 | 0.29807267 |
| *ENSG00000262165* | -4.150487 | 2.101451 | -1.975058 | 0.048261539 | NA |
| *MORN1* | -3.06201 | 1.55045 | -1.974917 | 0.048277521 | 0.29807267 |
| *ENPP4* | -2.771715 | 1.403526 | -1.974823 | 0.048288232 | 0.29807267 |
| *ELL2* | -3.35586 | 1.699326 | -1.974819 | 0.048288737 | 0.29807267 |
| *KLF16* | -2.146418 | 1.086912 | -1.974786 | 0.048292394 | 0.29807267 |
| *ENSG00000271553* | -3.169659 | 1.605362 | -1.97442 | 0.048333987 | NA |
| *ENSG00000259976* | -1.387978 | 0.703041 | -1.974251 | 0.048353264 | 0.29826998 |
| *GATM* | -2.245584 | 1.137979 | -1.97331 | 0.048460284 | 0.29848057 |
| *DHX58* | -1.892204 | 0.95894 | -1.973224 | 0.048470008 | 0.29848057 |
| *MCCC1* | -1.273784 | 0.645546 | -1.973188 | 0.04847417 | 0.29848057 |
| *CORO2A* | -2.993728 | 1.517625 | -1.97264 | 0.048536558 | 0.29851309 |
| *TRMT44* | -2.196102 | 1.113284 | -1.972634 | 0.048537303 | 0.29851309 |
| *ABCB10* | -2.371427 | 1.202399 | -1.972246 | 0.048581519 | 0.29860707 |
| *MRRF* | -1.215018 | 0.61615 | -1.97195 | 0.048615346 | 0.29863713 |
| *ENSG00000240401* | -3.470054 | 1.760733 | -1.970801 | 0.048746659 | NA |
| *NCAPD3* | -2.26721 | 1.150701 | -1.970287 | 0.048805471 | 0.29915374 |
| *FAM120C* | -2.28554 | 1.160331 | -1.969732 | 0.048869151 | 0.29915374 |
| *TMEM234* | -1.459528 | 0.741017 | -1.969628 | 0.048881029 | 0.29915374 |
| *PRPF40B* | -2.471568 | 1.255094 | -1.969228 | 0.048926869 | 0.29915374 |
| *CTSC* | -1.192008 | 0.605374 | -1.969044 | 0.048948036 | 0.29915374 |
| *TEX2* | -2.615999 | 1.328932 | -1.968497 | 0.049010933 | 0.29915374 |
| *CDC37L1* | -1.253404 | 0.636735 | -1.968485 | 0.049012244 | 0.29915374 |
| *ZNF620* | -2.464361 | 1.252067 | -1.968234 | 0.049041167 | 0.29915374 |
| *ZNF284* | -2.656641 | 1.349767 | -1.968222 | 0.049042495 | 0.29915374 |
| *MATN1.AS1* | -2.675799 | 1.359921 | -1.967614 | 0.049112494 | 0.29915374 |
| *SLC37A2* | -1.785035 | 0.907257 | -1.967508 | 0.049124697 | 0.29915374 |
| *FCHSD1* | -1.732031 | 0.880424 | -1.96727 | 0.049152048 | 0.29915374 |
| *TAGLN* | -1.937697 | 0.985093 | -1.96702 | 0.049180901 | 0.29915374 |
| *SAC3D1* | -2.949279 | 1.499468 | -1.966884 | 0.049196563 | 0.29915374 |
| *CMTM3* | -1.50295 | 0.764346 | -1.966322 | 0.049261378 | 0.29915374 |
| *IGHE* | -2.136293 | 1.086851 | -1.96558 | 0.049347096 | 0.29915374 |
| *NUP214* | -1.160458 | 0.590429 | -1.965447 | 0.049362503 | 0.29915374 |
| *SEC61A1* | -1.165111 | 0.592845 | -1.965288 | 0.049380933 | 0.29915374 |
| *SRD5A3* | -1.646539 | 0.837817 | -1.965273 | 0.049382704 | 0.29915374 |
| *TMEM181* | -2.828247 | 1.439393 | -1.964888 | 0.049427178 | 0.29915374 |
| *CHKB* | -1.255349 | 0.6389 | -1.96486 | 0.049430411 | 0.29915374 |
| *TAF13* | -2.501176 | 1.272969 | -1.964836 | 0.049433191 | 0.29915374 |

| Cluster 7 | log2FC | lfcSE | stat | pvalue | padj |
|---|---|---|---|---|---|
| *VPREB3* | 2.854232 | 0.24388 | 11.703412 | < 2.22e-16 | < 2.22e-16 |
| *GRN* | 1.384602 | 0.231108 | 5.991136 | 2.08E-09 | 1.34E-05 |
| *LPCAT2* | -5.543954 | 0.964884 | -5.745721 | 9.15E-09 | 3.93E-05 |
| *CIB2* | -4.434359 | 0.792031 | -5.598717 | 2.16E-08 | 5.92E-05 |
| *LINC01800* | -4.802941 | 0.859533 | -5.587848 | 2.30E-08 | 5.92E-05 |
| *TNFSF14* | -5.086043 | 0.95761 | -5.311186 | 1.09E-07 | 0.000202244 |
| *ZNF318* | -2.077798 | 0.391348 | -5.309334 | 1.10E-07 | 0.000202244 |
| *SLC9B1* | -4.473759 | 0.847329 | -5.279838 | 1.29E-07 | 0.00020796 |
| *JCHAIN* | 2.42661 | 0.463884 | 5.23107 | 1.69E-07 | 0.000240944 |
| *LINC00618* | -4.765824 | 0.916332 | -5.200979 | 1.98E-07 | 0.000242661 |
| *TUFT1* | -4.97192 | 0.959824 | -5.180031 | 2.22E-07 | 0.000242661 |
| *CRYM* | -4.577346 | 0.885031 | -5.171961 | 2.32E-07 | 0.000242661 |
| *MS4A1* | -0.749186 | 0.145259 | -5.157569 | 2.50E-07 | 0.000242661 |
| *KLK2* | -4.17595 | 0.813331 | -5.134377 | 2.83E-07 | 0.000242661 |
| *IGLV1.51* | -4.208837 | 0.821642 | -5.122471 | 3.02E-07 | 0.000242661 |
| *NPHP1* | -3.86081 | 0.75517 | -5.112507 | 3.18E-07 | 0.000242661 |
| *LINC01991* | -3.349752 | 0.655412 | -5.110913 | 3.21E-07 | 0.000242661 |
| *MIR210HG* | -4.301304 | 0.848352 | -5.070191 | 3.97E-07 | 0.000267909 |
| *ZNF233* | -4.180884 | 0.825341 | -5.065646 | 4.07E-07 | 0.000267909 |
| *GPR161* | -4.172761 | 0.824446 | -5.06129 | 4.16E-07 | 0.000267909 |
| *ENSG00000260331* | -4.527788 | 0.899803 | -5.031979 | 4.85E-07 | 0.000297437 |
| *HIST1H2AH* | -4.482653 | 0.895069 | -5.008166 | 5.50E-07 | 0.000321389 |
| *ENSG00000274015* | -4.906912 | 0.986615 | -4.973481 | 6.58E-07 | 0.000354517 |
| *NECAB3* | -4.088924 | 0.822322 | -4.97241 | 6.61E-07 | 0.000354517 |
| *ENSG00000273165* | -4.977281 | 1.003239 | -4.961213 | 7.01E-07 | 0.000360555 |
| *EPB41L4A* | -5.219148 | 1.064983 | -4.900686 | 9.55E-07 | 0.000472628 |
| *MTRNR2L3* | -4.692298 | 0.961392 | -4.880732 | 1.06E-06 | 0.000503686 |
| *ZNF221* | -4.116133 | 0.848031 | -4.853751 | 1.21E-06 | 0.000556717 |
| *CEP72* | -4.040483 | 0.83654 | -4.829995 | 1.37E-06 | 0.000598109 |
| *P3H4* | -4.935412 | 1.022717 | -4.825787 | 1.39E-06 | 0.000598109 |
| *TEF* | -4.067284 | 0.855083 | -4.756594 | 1.97E-06 | 0.000743237 |
| *TTC30A* | -4.306932 | 0.906205 | -4.752711 | 2.01E-06 | 0.000743237 |
| *UACA* | -3.767404 | 0.792837 | -4.751802 | 2.02E-06 | 0.000743237 |
| *ENSG00000272854* | -4.764412 | 1.002702 | -4.751574 | 2.02E-06 | 0.000743237 |
| *ADAM12* | -4.339169 | 0.913464 | -4.750236 | 2.03E-06 | 0.000743237 |
| *ENSG00000224934* | -4.990534 | 1.052318 | -4.742421 | 2.11E-06 | 0.000743237 |
| *LMO2* | -4.09339 | 0.863585 | -4.739995 | 2.14E-06 | 0.000743237 |
| *ENSG00000265100* | -4.784995 | 1.013791 | -4.719902 | 2.36E-06 | 0.000785934 |
| *ENSG00000266378* | -3.936342 | 0.834331 | -4.717963 | 2.38E-06 | 0.000785934 |
| *LINC02099* | -4.588187 | 0.979534 | -4.684053 | 2.81E-06 | 0.000904734 |
| *PRR5* | -4.883462 | 1.048718 | -4.656603 | 3.21E-06 | 0.000983078 |
| *C5orf17* | -4.640488 | 0.996903 | -4.654903 | 3.24E-06 | 0.000983078 |
| *ENSG00000230747* | -4.351161 | 0.935709 | -4.65012 | 3.32E-06 | 0.000983078 |
| *ENSG00000267633* | -4.652077 | 1.001992 | -4.64283 | 3.44E-06 | 0.000983078 |
| *ARHGAP5.AS1* | -5.287809 | 1.138941 | -4.642743 | 3.44E-06 | 0.000983078 |
| *ENSG00000280011* | -3.694563 | 0.798081 | -4.629306 | 3.67E-06 | 0.001026264 |

| | | | | | |
|---|---|---|---|---|---|
| *REPS2* | -3.591461 | 0.77812 | -4.615562 | 3.92E-06 | 0.001042159 |
| *KIF18A* | -3.422656 | 0.741562 | -4.615466 | 3.92E-06 | 0.001042159 |
| *H1FX.AS1* | -4.881934 | 1.059098 | -4.609522 | 4.04E-06 | 0.001042159 |
| *FRMPD1* | -4.215548 | 0.914671 | -4.608813 | 4.05E-06 | 0.001042159 |
| *BCL9* | -4.043987 | 0.878794 | -4.601745 | 4.19E-06 | 0.001057024 |
| *ENSG00000225335* | -4.210724 | 0.91691 | -4.592296 | 4.38E-06 | 0.001084779 |
| *ENSG00000274292* | -4.220849 | 0.924366 | -4.56621 | 4.97E-06 | 0.001181146 |
| *GSTM2* | -4.329527 | 0.948581 | -4.564212 | 5.01E-06 | 0.001181146 |
| *ENSG00000272579* | -3.920037 | 0.859401 | -4.56136 | 5.08E-06 | 0.001181146 |
| *ENSG00000186019* | -4.814581 | 1.056069 | -4.558966 | 5.14E-06 | 0.001181146 |
| *TTYH2* | -4.346782 | 0.959662 | -4.529494 | 5.91E-06 | 0.001334671 |
| *KCNQ1* | -4.448622 | 0.983255 | -4.524385 | 6.06E-06 | 0.001338089 |
| *ENSG00000261079* | -3.979876 | 0.88018 | -4.521661 | 6.14E-06 | 0.001338089 |
| *ENSG00000272800* | -4.733309 | 1.048059 | -4.516264 | 6.29E-06 | 0.001349751 |
| *ATP6V0E2.AS1* | -4.354996 | 0.967784 | -4.499966 | 6.80E-06 | 0.001433599 |
| *RGS9* | -3.994482 | 0.889548 | -4.490465 | 7.11E-06 | 0.001474888 |
| *DMXL2* | -4.315991 | 0.962479 | -4.484246 | 7.32E-06 | 0.001494457 |
| *SIRPA* | -4.239828 | 0.946592 | -4.479045 | 7.50E-06 | 0.001507405 |
| *ADM2* | -3.530647 | 0.789514 | -4.471922 | 7.75E-06 | 0.001534531 |
| *ARHGAP21* | -4.252485 | 0.952805 | -4.463122 | 8.08E-06 | 0.001574728 |
| *HAVCR2* | -3.926323 | 0.88089 | -4.457221 | 8.30E-06 | 0.001594528 |
| *PPP1R26* | -4.227115 | 0.949149 | -4.453582 | 8.44E-06 | 0.001597958 |
| *ADCY4* | -4.353299 | 0.980581 | -4.43951 | 9.02E-06 | 0.001670629 |
| *SPAG5* | -4.711914 | 1.06177 | -4.437791 | 9.09E-06 | 0.001670629 |
| *ENSG00000272004* | -4.723485 | 1.066328 | -4.429674 | 9.44E-06 | 0.001673969 |
| *BCAT1* | -4.389299 | 0.991422 | -4.427274 | 9.54E-06 | 0.001673969 |
| *ENSG00000234915* | -4.65916 | 1.052498 | -4.426765 | 9.57E-06 | 0.001673969 |
| *KRTCAP3* | -3.760739 | 0.850035 | -4.424218 | 9.68E-06 | 0.001673969 |
| *VASH2* | -4.671327 | 1.056268 | -4.422482 | 9.76E-06 | 0.001673969 |
| *ENSG00000255495* | -3.807211 | 0.86172 | -4.418156 | 9.95E-06 | 0.001685353 |
| *ENSG00000271993* | -3.716114 | 0.844066 | -4.402633 | 1.07E-05 | 0.001762981 |
| *KIF15* | -4.668126 | 1.060914 | -4.400098 | 1.08E-05 | 0.001762981 |
| *LINC00937* | -3.684647 | 0.837416 | -4.400017 | 1.08E-05 | 0.001762981 |
| *APPL2* | -2.757797 | 0.627219 | -4.396864 | 1.10E-05 | 0.001766416 |
| *ENSG00000228242* | -3.975467 | 0.905726 | -4.389261 | 1.14E-05 | 0.001806725 |
| *SPA17* | -4.166914 | 0.951043 | -4.381417 | 1.18E-05 | 0.00184898 |
| *LINC00894* | -4.143467 | 0.946231 | -4.378917 | 1.19E-05 | 0.00184898 |
| *ENSG00000279714* | -4.221214 | 0.965577 | -4.371702 | 1.23E-05 | 0.001888409 |
| *IL6* | -4.076217 | 0.932966 | -4.369093 | 1.25E-05 | 0.001888624 |
| *TGFB1I1* | -4.065201 | 0.931531 | -4.364001 | 1.28E-05 | 0.001910676 |
| *ENSG00000267282* | -4.127756 | 0.946686 | -4.360217 | 1.30E-05 | 0.001921668 |
| *LINC02076* | -3.671787 | 0.845613 | -4.342158 | 1.41E-05 | 0.002047396 |
| *NR5A2* | -3.79566 | 0.874306 | -4.341339 | 1.42E-05 | 0.002047396 |
| *IFNG.AS1* | -4.316183 | 0.995578 | -4.335353 | 1.46E-05 | 0.002054564 |
| *BEX3* | -4.044332 | 0.9329 | -4.335227 | 1.46E-05 | 0.002054564 |
| *NMNAT3* | -4.192455 | 0.967501 | -4.333282 | 1.47E-05 | 0.002054564 |
| *ZNF135* | -2.822113 | 0.651795 | -4.329756 | 1.49E-05 | 0.00206529 |

| | | | | | |
|---|---|---|---|---|---|
| *NBPF3* | -3.939929 | 0.91272 | -4.31669 | 1.58E-05 | 0.002168038 |
| *SLC2A4RG* | -3.085661 | 0.715548 | -4.312307 | 1.62E-05 | 0.002188205 |
| *ENSG00000231294* | -4.53909 | 1.05368 | -4.307844 | 1.65E-05 | 0.002209552 |
| *CRYGS* | -3.709445 | 0.861707 | -4.304765 | 1.67E-05 | 0.002216095 |
| *ENSG00000266998* | -3.841419 | 0.892808 | -4.302625 | 1.69E-05 | 0.002216095 |
| *FAM230J* | -3.886918 | 0.904192 | -4.298775 | 1.72E-05 | 0.002232161 |
| *PIK3R4* | -2.842747 | 0.661827 | -4.2953 | 1.74E-05 | 0.002244742 |
| *NAGPA.AS1* | -3.959636 | 0.92269 | -4.291403 | 1.78E-05 | 0.002261889 |
| *CAPS* | -2.571078 | 0.599615 | -4.287885 | 1.80E-05 | 0.002275476 |
| *ADNP.AS1* | -3.488229 | 0.815597 | -4.276901 | 1.90E-05 | 0.00234093 |
| *ENSG00000280433* | -2.983525 | 0.69798 | -4.274512 | 1.92E-05 | 0.00234093 |
| *MMP23B* | -4.435831 | 1.038069 | -4.273157 | 1.93E-05 | 0.00234093 |
| *PDE1B* | -4.270393 | 0.999387 | -4.273013 | 1.93E-05 | 0.00234093 |
| *IGLV4.69* | -3.828319 | 0.896487 | -4.270355 | 1.95E-05 | 0.002346873 |
| *GP1BA* | -3.7887 | 0.889549 | -4.259127 | 2.05E-05 | 0.002445052 |
| *ENSG00000258940* | -3.497781 | 0.821732 | -4.256597 | 2.08E-05 | 0.002450182 |
| *CCRL2* | -4.271755 | 1.005623 | -4.247867 | 2.16E-05 | 0.00252445 |
| *ENSG00000271797* | -3.768724 | 0.888475 | -4.241791 | 2.22E-05 | 0.002570424 |
| *C7orf61* | -3.806533 | 0.899596 | -4.231379 | 2.32E-05 | 0.002668323 |
| *MOK* | -3.567918 | 0.843664 | -4.229077 | 2.35E-05 | 0.002671917 |
| *ENSG00000265778* | -3.484069 | 0.825297 | -4.221595 | 2.43E-05 | 0.002737959 |
| *ENSG00000260911* | -4.49304 | 1.0653 | -4.217628 | 2.47E-05 | 0.002752812 |
| *PCOLCE* | -2.663865 | 0.631779 | -4.216453 | 2.48E-05 | 0.002752812 |
| *LBH* | -1.190953 | 0.282945 | -4.209126 | 2.56E-05 | 0.002819302 |
| *PLK4* | -4.670154 | 1.110366 | -4.20596 | 2.60E-05 | 0.002834846 |
| *SNX21* | -4.153112 | 0.988158 | -4.202883 | 2.64E-05 | 0.002849524 |
| *ENSG00000223881* | -3.633799 | 0.866504 | -4.193634 | 2.75E-05 | 0.002943555 |
| *ENSG00000258944* | -3.89742 | 0.930255 | -4.189627 | 2.79E-05 | 0.002971244 |
| *GNRH1* | -3.226316 | 0.771254 | -4.183208 | 2.87E-05 | 0.003021525 |
| *FBLN2* | -5.529076 | 1.322084 | -4.182093 | 2.89E-05 | 0.003021525 |
| *TMEM225B* | -3.712975 | 0.889524 | -4.174116 | 2.99E-05 | 0.003104115 |
| *BIK* | -4.110607 | 0.985511 | -4.171043 | 3.03E-05 | 0.003117799 |
| *ENSG00000273314* | -3.648801 | 0.875314 | -4.168563 | 3.07E-05 | 0.003117799 |
| *ARHGAP22* | -3.708922 | 0.889928 | -4.167667 | 3.08E-05 | 0.003117799 |
| *IGKV3.20* | -3.30255 | 0.793177 | -4.163701 | 3.13E-05 | 0.003147687 |
| *PLCB3* | -3.157559 | 0.75893 | -4.160541 | 3.17E-05 | 0.003166828 |
| *SLC24A1* | -4.426526 | 1.066337 | -4.151151 | 3.31E-05 | 0.003272657 |
| *DPH1* | -3.824819 | 0.922074 | -4.148061 | 3.35E-05 | 0.003272657 |
| *ZNF286B* | -3.5125 | 0.846841 | -4.147766 | 3.36E-05 | 0.003272657 |
| *LINC02202* | -3.559638 | 0.859417 | -4.141921 | 3.44E-05 | 0.003331958 |
| *UNC79* | -4.221184 | 1.020988 | -4.134409 | 3.56E-05 | 0.003417142 |
| *CHPF* | -4.26766 | 1.033798 | -4.128136 | 3.66E-05 | 0.003485675 |
| *LMTK2* | -3.638692 | 0.88237 | -4.123769 | 3.73E-05 | 0.003526335 |
| *ENSG00000232811* | -4.929516 | 1.197754 | -4.115635 | 3.86E-05 | 0.003602824 |
| *ENSG00000261071* | -3.42605 | 0.832483 | -4.115461 | 3.86E-05 | 0.003602824 |
| *ENSG00000251330* | -4.309099 | 1.048366 | -4.1103 | 3.95E-05 | 0.0036578 |
| *ENSG00000260329* | -4.352736 | 1.059567 | -4.108035 | 3.99E-05 | 0.003667464 |

| | | | | | |
|---|---|---|---|---|---|
| *LIMA1* | -3.791788 | 0.92448 | -4.101535 | 4.10E-05 | 0.003726259 |
| *BMERB1* | -3.7337 | 0.911313 | -4.097056 | 4.18E-05 | 0.003726259 |
| *GPER1* | -3.81926 | 0.932216 | -4.096968 | 4.19E-05 | 0.003726259 |
| *RAB3B* | -4.551274 | 1.110914 | -4.096873 | 4.19E-05 | 0.003726259 |
| *ENSG00000258377* | -3.904975 | 0.953308 | -4.096239 | 4.20E-05 | 0.003726259 |
| *RGPD5* | -3.322301 | 0.811579 | -4.093626 | 4.25E-05 | 0.003742704 |
| *C17orf51* | -3.077283 | 0.752561 | -4.08908 | 4.33E-05 | 0.003780867 |
| *ENSG00000228037* | -3.797794 | 0.928983 | -4.088118 | 4.35E-05 | 0.003780867 |
| *ADD2* | -3.358689 | 0.821968 | -4.086153 | 4.39E-05 | 0.003787422 |
| *TRIM46* | -3.957146 | 0.970562 | -4.07717 | 4.56E-05 | 0.003910474 |
| *DLG2* | -3.364345 | 0.825783 | -4.074129 | 4.62E-05 | 0.003910849 |
| *PROB1* | -3.372621 | 0.828057 | -4.072933 | 4.64E-05 | 0.003910849 |
| *ACVR1B* | -3.710472 | 0.91138 | -4.071268 | 4.68E-05 | 0.003910849 |
| *SLC36A1* | -3.77719 | 0.927824 | -4.071022 | 4.68E-05 | 0.003910849 |
| *ENSG00000278817* | -3.561807 | 0.876071 | -4.065662 | 4.79E-05 | 0.003976019 |
| *LINC00987* | -3.761046 | 0.927353 | -4.055679 | 5.00E-05 | 0.004123124 |
| *ZNF391* | -3.232661 | 0.797773 | -4.052104 | 5.08E-05 | 0.004159979 |
| *ENSG00000255537* | -3.416272 | 0.843921 | -4.048096 | 5.16E-05 | 0.00420506 |
| *SPATS2L* | -3.788828 | 0.937488 | -4.041467 | 5.31E-05 | 0.004298544 |
| *MDS2* | -3.437447 | 0.851917 | -4.034956 | 5.46E-05 | 0.00439187 |
| *HIST1H1B* | -3.911168 | 0.970237 | -4.031147 | 5.55E-05 | 0.004435941 |
| *SERPINH1* | -3.786815 | 0.940696 | -4.025545 | 5.68E-05 | 0.004505825 |
| *CISH* | -5.334528 | 1.325491 | -4.024568 | 5.71E-05 | 0.004505825 |
| *GRM3* | -4.390616 | 1.091434 | -4.022797 | 5.75E-05 | 0.004512164 |
| *ENSG00000269814* | -4.01477 | 0.999378 | -4.017268 | 5.89E-05 | 0.004564297 |
| *ATXN7L2* | -3.433164 | 0.854609 | -4.017235 | 5.89E-05 | 0.004564297 |
| *LINC02021* | -3.328407 | 0.829698 | -4.011591 | 6.03E-05 | 0.00464684 |
| *CENPO* | -3.30168 | 0.823781 | -4.007959 | 6.12E-05 | 0.004690765 |
| *ENSG00000260317* | -3.555302 | 0.888885 | -3.999733 | 6.34E-05 | 0.004828088 |
| *LINC00562* | -2.752813 | 0.689172 | -3.994378 | 6.49E-05 | 0.004856361 |
| *U2AF1L5* | -4.629025 | 1.158939 | -3.994191 | 6.49E-05 | 0.004856361 |
| *IGLV2.11* | -4.955237 | 1.240613 | -3.994184 | 6.49E-05 | 0.004856361 |
| *LMNTD1* | -2.962577 | 0.745943 | -3.971587 | 7.14E-05 | 0.005310077 |
| *PTP4A3* | -2.93351 | 0.739155 | -3.968733 | 7.23E-05 | 0.005343201 |
| *IL18BP* | -3.231153 | 0.815351 | -3.962898 | 7.40E-05 | 0.005444249 |
| *CDKN2A* | -3.234637 | 0.817527 | -3.956611 | 7.60E-05 | 0.005557709 |
| *MESP1* | -3.737419 | 0.944952 | -3.955143 | 7.65E-05 | 0.005560352 |
| *ENSG00000244567* | -3.319767 | 0.839924 | -3.952463 | 7.74E-05 | 0.005582298 |
| *PBX4* | -3.420559 | 0.865633 | -3.951514 | 7.77E-05 | 0.005582298 |
| *ENSG00000262429* | -3.048579 | 0.772337 | -3.947211 | 7.91E-05 | 0.005651968 |
| *CHL1* | -3.188933 | 0.809218 | -3.940758 | 8.12E-05 | 0.005753669 |
| *ZSCAN22* | -2.800066 | 0.710625 | -3.940288 | 8.14E-05 | 0.005753669 |
| *CCDC150* | -3.608686 | 0.916472 | -3.937585 | 8.23E-05 | 0.005760706 |
| *FAM89A* | -3.672598 | 0.932754 | -3.937372 | 8.24E-05 | 0.005760706 |
| *SLC27A1* | -3.33452 | 0.847778 | -3.933246 | 8.38E-05 | 0.005804756 |
| *PLCG1* | -3.217266 | 0.81803 | -3.932946 | 8.39E-05 | 0.005804756 |
| *CCDC151* | -4.718677 | 1.201714 | -3.926622 | 8.61E-05 | 0.005927584 |

| | | | | | |
|---|---|---|---|---|---|
| *MIR34AHG* | -3.612462 | 0.920331 | -3.925175 | 8.67E-05 | 0.005931605 |
| *H3F3A* | 0.616242 | 0.157145 | 3.921491 | 8.80E-05 | 0.005962539 |
| *TIMP2* | -3.563246 | 0.908673 | -3.921374 | 8.80E-05 | 0.005962539 |
| *ENSG00000179428* | -4.54731 | 1.160547 | -3.918248 | 8.92E-05 | 0.006008755 |
| *ENSG00000273064* | -3.352114 | 0.856827 | -3.912242 | 9.14E-05 | 0.006128119 |
| *ITPRIPL1* | -3.411754 | 0.872785 | -3.909045 | 9.27E-05 | 0.006177609 |
| *CPLANE2* | -3.460296 | 0.885995 | -3.905549 | 9.40E-05 | 0.006235291 |
| *NR2F6* | -3.124952 | 0.803868 | -3.887394 | 0.000101326 | 0.006685974 |
| *LEMD2* | -2.238784 | 0.577321 | -3.877884 | 0.000105369 | 0.006882733 |
| *ZNF460.AS1* | -2.969149 | 0.765781 | -3.877284 | 0.000105629 | 0.006882733 |
| *DGAT1* | -2.931119 | 0.75624 | -3.875911 | 0.000106227 | 0.006882733 |
| *ENSG00000254162* | -3.555778 | 0.917524 | -3.875404 | 0.000106448 | 0.006882733 |
| *ABHD16A* | -3.061883 | 0.791019 | -3.870809 | 0.000108475 | 0.006923436 |
| *ENSG00000228989* | -2.886272 | 0.745693 | -3.870591 | 0.000108572 | 0.006923436 |
| *ENSG00000228436* | -2.895411 | 0.74816 | -3.870044 | 0.000108816 | 0.006923436 |
| *UCP3* | -3.758426 | 0.971391 | -3.869118 | 0.00010923 | 0.006923436 |
| *ENSG00000232874* | -3.562511 | 0.923107 | -3.859259 | 0.000113731 | 0.007173428 |
| *B4GALT1.AS1* | -3.422182 | 0.887437 | -3.856253 | 0.000115138 | 0.007177898 |
| *ANGPTL6* | -3.753424 | 0.973389 | -3.856037 | 0.00011524 | 0.007177898 |
| *PLBD2* | -2.82075 | 0.73161 | -3.855538 | 0.000115476 | 0.007177898 |
| *ENSG00000268573* | -4.08696 | 1.062423 | -3.846831 | 0.000119656 | 0.007401961 |
| *GPC2* | -3.92958 | 1.0227 | -3.842359 | 0.000121858 | 0.007497217 |
| *ENSG00000272455* | -4.284779 | 1.115519 | -3.841064 | 0.000122502 | 0.007497217 |
| *ZFP37* | -3.443268 | 0.896642 | -3.840181 | 0.000122943 | 0.007497217 |
| *LINC01001* | -3.159199 | 0.824447 | -3.8319 | 0.000127157 | 0.007717613 |
| *ASMTL.AS1* | -3.345049 | 0.875387 | -3.821225 | 0.00013279 | 0.008021659 |
| *CLIP4* | -4.212752 | 1.103885 | -3.816295 | 0.000135471 | 0.008145333 |
| *SUOX* | -3.206146 | 0.841204 | -3.811379 | 0.000138193 | 0.008270395 |
| *ZNF23* | -4.004328 | 1.052664 | -3.803994 | 0.000142382 | 0.008481593 |
| *CBX8* | -3.087937 | 0.812124 | -3.802296 | 0.000143361 | 0.008500588 |
| *HYAL3* | -2.93912 | 0.77325 | -3.800995 | 0.000144116 | 0.008506172 |
| *RAP2C.AS1* | -3.027803 | 0.796846 | -3.799732 | 0.000144853 | 0.008510581 |
| *COX16* | -1.708394 | 0.450324 | -3.793696 | 0.000148421 | 0.008680614 |
| *LINC00205* | -4.212801 | 1.112228 | -3.787712 | 0.000152041 | 0.008852075 |
| *ENSG00000250073* | -3.427297 | 0.905929 | -3.783186 | 0.000154834 | 0.008961866 |
| *SEMA3D* | -4.356997 | 1.151911 | -3.782406 | 0.00015532 | 0.008961866 |
| *ENSG00000225032* | -3.474579 | 0.919103 | -3.780401 | 0.000156576 | 0.008994044 |
| *NPIPA5* | -3.916495 | 1.038647 | -3.770765 | 0.000162748 | 0.009307017 |
| *ENSG00000240401* | -2.627216 | 0.69781 | -3.764946 | 0.000166585 | 0.009484287 |
| *ATL1* | -3.284274 | 0.872737 | -3.763189 | 0.00016776 | 0.009509126 |
| *DOK2* | -3.267438 | 0.869054 | -3.759764 | 0.000170074 | 0.009597969 |
| *IL1RAP* | -2.958248 | 0.787643 | -3.755825 | 0.000172771 | 0.009668624 |
| *LINC01816* | -4.073885 | 1.08477 | -3.755529 | 0.000172975 | 0.009668624 |
| *CCDC144A* | -3.340462 | 0.889798 | -3.754179 | 0.00017391 | 0.009668624 |
| *TRPV1* | -2.953452 | 0.786837 | -3.753574 | 0.000174331 | 0.009668624 |
| *EVI5L* | -3.297243 | 0.879326 | -3.749737 | 0.00017702 | 0.009734677 |
| *CAMKMT* | -2.209662 | 0.589463 | -3.748599 | 0.000177825 | 0.009734677 |

| | | | | | |
|---|---|---|---|---|---|
| *ENSG00000267421* | -5.173216 | 1.380277 | -3.747956 | 0.000178282 | 0.009734677 |
| *LINC00504* | -3.519666 | 0.939196 | -3.747529 | 0.000178585 | 0.009734677 |
| *ENSG00000262319* | -3.628189 | 0.968416 | -3.746519 | 0.000179305 | 0.009734677 |
| *ENSG00000261220* | -2.853401 | 0.76434 | -3.733156 | 0.000189095 | 0.010161507 |
| *PARD6B* | -2.639803 | 0.707209 | -3.732705 | 0.000189434 | 0.010161507 |
| *DEPDC4* | -4.682953 | 1.254619 | -3.73257 | 0.000189536 | 0.010161507 |
| *ELMO3* | -3.062703 | 0.820774 | -3.731483 | 0.000190356 | 0.010163101 |
| *ZNF607* | -2.945722 | 0.789841 | -3.729511 | 0.000191852 | 0.010200655 |
| *FGFRL1* | -4.041242 | 1.084162 | -3.727524 | 0.00019337 | 0.010239061 |
| *MPV17L* | -3.046469 | 0.81809 | -3.723878 | 0.000196186 | 0.010345585 |
| *ENSG00000281706* | -3.357669 | 0.902279 | -3.721322 | 0.000198183 | 0.010372764 |
| *LRP8* | -3.166717 | 0.851004 | -3.721155 | 0.000198314 | 0.010372764 |
| *ZNF133* | -2.839953 | 0.763964 | -3.717389 | 0.000201293 | 0.010485968 |
| *INSL3* | -3.850403 | 1.03711 | -3.712627 | 0.000205119 | 0.010642188 |
| *IGHV4.34* | -3.348007 | 0.902209 | -3.7109 | 0.000206524 | 0.010672062 |
| *TBC1D16* | -2.743451 | 0.741977 | -3.697489 | 0.000217743 | 0.011200504 |
| *ENSG00000278869* | -3.119107 | 0.843909 | -3.696024 | 0.000219002 | 0.011200504 |
| *ENSG00000264235* | -3.319218 | 0.898215 | -3.695349 | 0.000219585 | 0.011200504 |
| *FUT8.AS1* | -4.215064 | 1.140871 | -3.694601 | 0.000220232 | 0.011200504 |
| *ENSG00000248367* | -3.04004 | 0.82424 | -3.688296 | 0.000225761 | 0.011436501 |
| *PRELID3A* | -3.961834 | 1.074816 | -3.686056 | 0.000227756 | 0.011492296 |
| *TMEM255B* | -3.176722 | 0.863687 | -3.678095 | 0.000234983 | 0.01181064 |
| *RDH10* | -3.434936 | 0.934936 | -3.673978 | 0.000238803 | 0.011866401 |
| *AREL1* | -2.467368 | 0.671821 | -3.672658 | 0.000240041 | 0.011866401 |
| *SRRM2.AS1* | -3.395936 | 0.924758 | -3.672242 | 0.000240432 | 0.011866401 |
| *CTIF* | -3.27902 | 0.892958 | -3.672088 | 0.000240577 | 0.011866401 |
| *ENSG00000273272* | -4.170949 | 1.136189 | -3.671 | 0.000241603 | 0.011866401 |
| *TTC8* | -3.568724 | 0.972146 | -3.670976 | 0.000241626 | 0.011866401 |
| *LIX1.AS1* | -3.206987 | 0.874435 | -3.667497 | 0.000244936 | 0.011983248 |
| *ZNF713* | -2.845459 | 0.776285 | -3.665482 | 0.000246873 | 0.012007949 |
| *FAH* | -3.037122 | 0.828861 | -3.664213 | 0.000248101 | 0.012007949 |
| *CMC4* | -3.116199 | 0.850813 | -3.662614 | 0.000249655 | 0.012007949 |
| *AMIGO2* | -3.241087 | 0.884941 | -3.66249 | 0.000249775 | 0.012007949 |
| *YY2* | -3.182461 | 0.86919 | -3.661409 | 0.000250832 | 0.012007949 |
| *PHLDA3* | -4.01495 | 1.096623 | -3.661196 | 0.000251041 | 0.012007949 |
| *ENSG00000277763* | -3.739141 | 1.022285 | -3.657631 | 0.000254557 | 0.012087938 |
| *FAM161B* | -3.036298 | 0.830135 | -3.657596 | 0.000254592 | 0.012087938 |
| *PALM2* | -3.234207 | 0.884756 | -3.655477 | 0.000256704 | 0.012105386 |
| *FIRRE* | -3.802494 | 1.040257 | -3.655341 | 0.000256841 | 0.012105386 |
| *PLD6* | -2.972285 | 0.813744 | -3.652603 | 0.000259595 | 0.01219054 |
| *ASCL2* | -3.669775 | 1.005466 | -3.649824 | 0.000262421 | 0.012243123 |
| *GPX1* | 0.8225 | 0.225378 | 3.649421 | 0.000262832 | 0.012243123 |
| *RGL1* | -3.757966 | 1.029946 | -3.648701 | 0.000263569 | 0.012243123 |
| *SOX5* | -5.291275 | 1.45093 | -3.646817 | 0.000265509 | 0.012247397 |
| *ENSG00000233903* | -2.971941 | 0.814953 | -3.646763 | 0.000265565 | 0.012247397 |
| *ZNF707* | -2.327706 | 0.638945 | -3.643045 | 0.000269431 | 0.012339935 |
| *ABCD3* | -2.287726 | 0.62798 | -3.64299 | 0.00026949 | 0.012339935 |

| | | | | | |
|---|---|---|---|---|---|
| *ENSG00000237753* | -2.968918 | 0.815692 | -3.639753 | 0.0002729 | 0.012431532 |
| *NT5DC3* | -2.863669 | 0.786882 | -3.63926 | 0.000273422 | 0.012431532 |
| *ERI2* | -3.327989 | 0.915533 | -3.635027 | 0.000277951 | 0.012592953 |
| *DOCK7* | -2.84053 | 0.78197 | -3.632532 | 0.000280654 | 0.01264999 |
| *ENSG00000260778* | -3.181529 | 0.875959 | -3.632052 | 0.000281176 | 0.01264999 |
| *PKN3* | -3.980279 | 1.097693 | -3.62604 | 0.0002878 | 0.012902877 |
| *FAM83G* | -2.479763 | 0.684895 | -3.620648 | 0.000293866 | 0.013129063 |
| *CCSAP* | -2.678273 | 0.740438 | -3.617149 | 0.000297866 | 0.013261717 |
| *ENSG00000272973* | -3.169224 | 0.876514 | -3.615715 | 0.00029952 | 0.013289371 |
| *ENSG00000261879* | -2.899097 | 0.802642 | -3.611941 | 0.000303913 | 0.01343799 |
| *CCDC62* | -3.359547 | 0.932006 | -3.60464 | 0.000312586 | 0.013774129 |
| *ZNF554* | -2.932097 | 0.813782 | -3.60305 | 0.000314506 | 0.01381141 |
| *VPS33B.DT* | -3.435176 | 0.954901 | -3.597418 | 0.000321392 | 0.014065835 |
| *PAM* | -2.739065 | 0.761764 | -3.595688 | 0.000323535 | 0.014111605 |
| *WDR66* | -2.954481 | 0.822104 | -3.593803 | 0.000325886 | 0.014166147 |
| *ENSG00000266385* | -3.324618 | 0.926765 | -3.587337 | 0.000334072 | 0.014473083 |
| *FAM85B* | -3.218599 | 0.899198 | -3.57941 | 0.000344371 | 0.014869201 |
| *WNT10A* | -2.59469 | 0.725227 | -3.57776 | 0.000346551 | 0.014913285 |
| *ENSG00000272264* | -3.206755 | 0.896744 | -3.575997 | 0.000348895 | 0.014964111 |
| *ENSG00000272933* | -2.878142 | 0.80563 | -3.572537 | 0.000353539 | 0.015024343 |
| *PLEKHD1* | -4.167094 | 1.166549 | -3.572155 | 0.000354056 | 0.015024343 |
| *ANK1* | -3.000097 | 0.839933 | -3.571828 | 0.000354499 | 0.015024343 |
| *ENSG00000261420* | -3.235404 | 0.905962 | -3.571237 | 0.000355299 | 0.015024343 |
| *KDM4A.AS1* | -2.988097 | 0.836936 | -3.570283 | 0.000356596 | 0.015024343 |
| *STAT5A* | -1.663659 | 0.466042 | -3.569762 | 0.000357305 | 0.015024343 |
| *ENSG00000270127* | -3.233125 | 0.907463 | -3.562819 | 0.000366893 | 0.015377246 |
| *TSPAN4* | -3.053424 | 0.858204 | -3.557922 | 0.0003738 | 0.015615855 |
| *OPRL1* | -3.037623 | 0.854492 | -3.554887 | 0.000378141 | 0.015696767 |
| *UBAC2.AS1* | -3.06082 | 0.861023 | -3.554863 | 0.000378177 | 0.015696767 |
| *ENSG00000256950* | -3.447868 | 0.971322 | -3.549667 | 0.000385719 | 0.01595836 |
| *ENSG00000276259* | -2.720152 | 0.766871 | -3.547077 | 0.000389531 | 0.016064389 |
| *ADRB2* | -2.664915 | 0.751902 | -3.544231 | 0.000393759 | 0.016186911 |
| *CCDC146* | -3.53153 | 0.996893 | -3.542536 | 0.000396299 | 0.016239407 |
| *BOLA3.AS1* | -2.650047 | 0.74828 | -3.541516 | 0.000397834 | 0.016250583 |
| *ITGA1* | -2.938675 | 0.830118 | -3.540069 | 0.000400022 | 0.016262903 |
| *ALYREF* | -2.575437 | 0.727712 | -3.539088 | 0.000401512 | 0.016262903 |
| *ENSG00000265218* | -3.714154 | 1.049548 | -3.538815 | 0.000401928 | 0.016262903 |
| *PTPRCAP* | -3.216475 | 0.909469 | -3.53665 | 0.000405236 | 0.016345361 |
| *ENSG00000271976* | -3.5023 | 0.991105 | -3.533734 | 0.000409733 | 0.016475102 |
| *THNSL1* | -2.906977 | 0.823119 | -3.53166 | 0.00041296 | 0.016553135 |
| *HARBI1* | -2.734103 | 0.77439 | -3.530656 | 0.000414531 | 0.016564504 |
| *GTSF1* | -2.483003 | 0.704116 | -3.52641 | 0.000421234 | 0.016775329 |
| *ENSG00000228835* | -2.699685 | 0.765723 | -3.525669 | 0.000422414 | 0.016775329 |
| *ARHGEF10* | -2.812681 | 0.797978 | -3.524761 | 0.000423865 | 0.016779696 |
| *ABCB9* | -3.482918 | 0.988351 | -3.52397 | 0.000425133 | 0.016779696 |
| *CHRNE* | -4.083154 | 1.159268 | -3.522182 | 0.00042801 | 0.016790455 |
| *ENSG00000264548* | -4.289746 | 1.218168 | -3.521473 | 0.000429157 | 0.016790455 |

| | | | | | |
|---|---|---|---|---|---|
| *SFT2D3* | -2.999537 | 0.851908 | -3.520963 | 0.000429982 | 0.016790455 |
| *PUSL1* | -2.167713 | 0.615868 | -3.519766 | 0.000431928 | 0.016790455 |
| *KLHL17* | -2.990535 | 0.849781 | -3.519183 | 0.000432878 | 0.016790455 |
| *C19orf73* | -2.886415 | 0.820386 | -3.518362 | 0.00043422 | 0.016790455 |
| *ITGAV* | -3.085513 | 0.877023 | -3.518166 | 0.00043454 | 0.016790455 |
| *ASB2* | 2.581441 | 0.734278 | 3.51562 | 0.000438729 | 0.016880511 |
| *CCDC183* | -2.947207 | 0.838428 | -3.515157 | 0.000439494 | 0.016880511 |
| *RNF31* | -2.87003 | 0.816664 | -3.514334 | 0.000440858 | 0.016882491 |
| *C16orf95* | -3.952635 | 1.125101 | -3.513139 | 0.000442846 | 0.016903966 |
| *LTB4R* | -2.963365 | 0.843787 | -3.511983 | 0.000444777 | 0.016903966 |
| *ENSG00000231595* | -4.005126 | 1.14053 | -3.511635 | 0.00044536 | 0.016903966 |
| *LSM11* | -2.976659 | 0.847877 | -3.510723 | 0.000446891 | 0.016912181 |
| *ENSG00000263531* | -2.702147 | 0.770475 | -3.507118 | 0.000452988 | 0.01709264 |
| *SMIM13* | -2.89729 | 0.826436 | -3.505765 | 0.000455297 | 0.017129553 |
| *DENND6B* | 2.036788 | 0.581272 | 3.504017 | 0.000458296 | 0.017192122 |
| *MYH7B* | -3.205589 | 0.91524 | -3.502456 | 0.000460989 | 0.017242875 |
| *FRAT2* | -3.220271 | 0.919829 | -3.500944 | 0.000463613 | 0.017290732 |
| *STAT4* | -2.467322 | 0.705721 | -3.496171 | 0.000471985 | 0.017515081 |
| *C3orf33* | -2.889488 | 0.826614 | -3.49557 | 0.00047305 | 0.017515081 |
| *PRRT2* | -2.845434 | 0.814098 | -3.495197 | 0.000473712 | 0.017515081 |
| *ENSG00000182376* | -3.885772 | 1.112584 | -3.492564 | 0.000478407 | 0.017638006 |
| *ACRBP* | -3.123201 | 0.894504 | -3.491545 | 0.000480235 | 0.017654803 |
| *LINC00996* | -3.188512 | 0.913475 | -3.490531 | 0.000482062 | 0.017671487 |
| *TTLL1* | -2.811544 | 0.806753 | -3.485011 | 0.000492118 | 0.01798886 |
| *RASGEF1A* | -2.868924 | 0.823931 | -3.481995 | 0.000497693 | 0.018141136 |
| *MBOAT2* | -3.697023 | 1.062146 | -3.480712 | 0.000500083 | 0.01816471 |
| *NBPF26* | -2.680334 | 0.770181 | -3.480134 | 0.000501164 | 0.01816471 |
| *ENSG00000237357* | -2.803743 | 0.806513 | -3.476377 | 0.000508238 | 0.018369376 |
| *FRY* | -3.157204 | 0.909196 | -3.472522 | 0.000515593 | 0.018540801 |
| *SMC6* | -0.983165 | 0.283138 | -3.472381 | 0.000515863 | 0.018540801 |
| *RGMB* | -2.83056 | 0.815586 | -3.470587 | 0.000519323 | 0.018552754 |
| *NXPH4* | -4.029833 | 1.161422 | -3.469741 | 0.000520961 | 0.018552754 |
| *TCEAL9* | -3.057189 | 0.881289 | -3.468999 | 0.000522401 | 0.018552754 |
| *MEGF8* | -2.68823 | 0.775037 | -3.468518 | 0.000523337 | 0.018552754 |
| *DDIAS* | -2.639673 | 0.761045 | -3.468483 | 0.000523405 | 0.018552754 |
| *EIF2AK3.DT* | -3.23366 | 0.93385 | -3.462718 | 0.000534748 | 0.01882932 |
| *ENSG00000272374* | -3.689577 | 1.065564 | -3.462557 | 0.000535069 | 0.01882932 |
| *MAPK8IP1* | -3.23902 | 0.935572 | -3.462075 | 0.000536027 | 0.01882932 |
| *LIPE* | -1.955731 | 0.565088 | -3.460932 | 0.000538309 | 0.01882932 |
| *LRCH1* | -1.855196 | 0.536056 | -3.460824 | 0.000538524 | 0.01882932 |
| *S100A2* | -2.705099 | 0.781916 | -3.459579 | 0.00054102 | 0.018832878 |
| *NRF1* | -2.164591 | 0.625728 | -3.459314 | 0.000541553 | 0.018832878 |
| *SEPHS2* | -2.943237 | 0.851072 | -3.458271 | 0.000543654 | 0.018854971 |
| *ENSG00000277602* | -3.641887 | 1.053702 | -3.456279 | 0.000547689 | 0.018941908 |
| *SDSL* | -2.867187 | 0.829726 | -3.455583 | 0.000549105 | 0.018941908 |
| *ENSG00000240710* | -2.802249 | 0.811625 | -3.452641 | 0.000555127 | 0.019098434 |
| *GPD1L* | -3.476405 | 1.007332 | -3.4511 | 0.000558306 | 0.019156592 |

| | | | | | |
|---|---|---|---|---|---|
| *RAB23* | -2.865518 | 0.831176 | -3.447547 | 0.000565701 | 0.019293129 |
| *PDE6A* | -2.7494 | 0.79754 | -3.447352 | 0.000566111 | 0.019293129 |
| *CHKA* | -2.624369 | 0.761342 | -3.447031 | 0.000566783 | 0.019293129 |
| *JUND* | -1.016003 | 0.294961 | -3.444531 | 0.000572051 | 0.019408884 |
| *HIST1H2BF* | -3.01909 | 0.876626 | -3.443988 | 0.000573201 | 0.019408884 |
| *PRR22* | -4.013721 | 1.165682 | -3.443238 | 0.000574793 | 0.019411694 |
| *OSGEPL1.AS1* | -3.342809 | 0.971409 | -3.441198 | 0.000579145 | 0.01949918 |
| *ZNF646* | -2.524337 | 0.733706 | -3.440527 | 0.000580582 | 0.01949918 |
| *ENSG00000249604* | -4.12344 | 1.198828 | -3.43956 | 0.000582661 | 0.01949918 |
| *ERMAP* | -2.955464 | 0.859347 | -3.439196 | 0.000583445 | 0.01949918 |
| *ZNF675* | -1.889528 | 0.549688 | -3.437455 | 0.000587208 | 0.019543475 |
| *JAM2* | -2.981331 | 0.867386 | -3.437144 | 0.000587884 | 0.019543475 |
| *BTBD6* | -2.625583 | 0.764033 | -3.43648 | 0.000589327 | 0.019543475 |
| *ENSG00000269982* | -3.708108 | 1.079723 | -3.434314 | 0.000594056 | 0.019649669 |
| *ENSG00000263826* | -4.066479 | 1.18471 | -3.432467 | 0.000598117 | 0.019733261 |
| *PLAAT4* | 1.665887 | 0.485757 | 3.429463 | 0.000604778 | 0.01980767 |
| *CCDC163* | -3.063407 | 0.893343 | -3.429151 | 0.000605473 | 0.01980767 |
| *FAXDC2* | -2.864108 | 0.835357 | -3.428603 | 0.000606697 | 0.01980767 |
| *CD1D* | 2.384216 | 0.695671 | 3.427216 | 0.000609803 | 0.01980767 |
| *NHEJ1* | -3.880063 | 1.132225 | -3.426936 | 0.000610432 | 0.01980767 |
| *VASH1* | -3.969134 | 1.158366 | -3.426493 | 0.000611429 | 0.01980767 |
| *PGAP3* | -2.847163 | 0.831003 | -3.426175 | 0.000612145 | 0.01980767 |
| *MYH10* | -3.959201 | 1.155933 | -3.425112 | 0.000614545 | 0.01980767 |
| *ENSG00000183889* | -3.683788 | 1.075645 | -3.424724 | 0.000615423 | 0.01980767 |
| *CMBL* | -4.016329 | 1.172797 | -3.424573 | 0.000615767 | 0.01980767 |
| *TNNC2* | -2.946065 | 0.860653 | -3.423058 | 0.000619208 | 0.019860743 |
| *PIWIL1* | -3.20268 | 0.935775 | -3.42249 | 0.000620504 | 0.019860743 |
| *NPC2* | -0.663923 | 0.194041 | -3.421562 | 0.000622626 | 0.019879219 |
| *ZNF730* | -3.708216 | 1.084073 | -3.420634 | 0.000624753 | 0.019897757 |
| *MRVI1.AS1* | -2.995157 | 0.876215 | -3.41829 | 0.00063016 | 0.020020407 |
| *ENSG00000270681* | -3.792978 | 1.110082 | -3.416844 | 0.000633516 | 0.02007745 |
| *FAM153B* | -3.995798 | 1.169724 | -3.416016 | 0.000635445 | 0.020089111 |
| *FMNL2* | -3.491837 | 1.023469 | -3.411764 | 0.000645439 | 0.020350935 |
| *PSD3* | -2.218121 | 0.650256 | -3.411152 | 0.00064689 | 0.020350935 |
| *ENSG00000282393* | -2.821237 | 0.827508 | -3.409316 | 0.000651259 | 0.02039317 |
| *SLC15A2* | -2.446851 | 0.717708 | -3.409256 | 0.000651402 | 0.02039317 |
| *TMEM217* | -3.673709 | 1.077866 | -3.408317 | 0.000653648 | 0.02041381 |
| *FES* | -2.537902 | 0.745283 | -3.405285 | 0.000660949 | 0.020574895 |
| *ENSG00000260100* | -3.824481 | 1.123344 | -3.404551 | 0.00066273 | 0.020574895 |
| *ENSG00000264112* | -2.881966 | 0.846593 | -3.404191 | 0.000663603 | 0.020574895 |
| *MALINC1* | -3.839129 | 1.12991 | -3.397731 | 0.000679472 | 0.021016252 |
| *ENSG00000274400* | -3.820724 | 1.124962 | -3.396313 | 0.000683003 | 0.021074805 |
| *SLC35D2* | -3.25529 | 0.958945 | -3.394657 | 0.000687146 | 0.021151942 |
| *SETMAR* | -2.859525 | 0.843026 | -3.391977 | 0.000693904 | 0.021308967 |
| *ZNF20* | -2.930228 | 0.864266 | -3.390424 | 0.000697847 | 0.021365531 |
| *ENSG00000267127* | -4.091735 | 1.207021 | -3.389945 | 0.000699066 | 0.021365531 |
| *C19orf18* | -3.769226 | 1.112979 | -3.386611 | 0.000707617 | 0.021457456 |

| | | | | | |
|---|---|---|---|---|---|
| *SYNJ1* | -3.828505 | 1.130762 | -3.385773 | 0.00070978 | 0.021457456 |
| *PRDM11* | -2.839465 | 0.838705 | -3.385533 | 0.000710403 | 0.021457456 |
| *TK1* | -2.53955 | 0.750137 | -3.385449 | 0.000710619 | 0.021457456 |
| *ZNF746* | -2.501064 | 0.738787 | -3.385366 | 0.000710835 | 0.021457456 |
| *ENSG00000259363* | -2.961551 | 0.874934 | -3.384885 | 0.00071208 | 0.021457456 |
| *KCTD12* | -3.776352 | 1.11616 | -3.383343 | 0.000716091 | 0.021527909 |
| *ENSG00000159239* | -3.086558 | 0.912691 | -3.381822 | 0.000720068 | 0.021549655 |
| *WDR7* | -2.583635 | 0.764071 | -3.38141 | 0.000721149 | 0.021549655 |
| *ACTA2* | -2.842125 | 0.840603 | -3.381056 | 0.000722078 | 0.021549655 |
| *ENSG00000249476* | -2.700277 | 0.798778 | -3.38051 | 0.000723514 | 0.021549655 |
| *PAN3.AS1* | -3.636753 | 1.076217 | -3.379199 | 0.000726973 | 0.021568667 |
| *IGKV3.15* | -3.551195 | 1.050961 | -3.378998 | 0.000727505 | 0.021568667 |
| *ENSG00000271806* | -3.071796 | 0.909759 | -3.376496 | 0.000734156 | 0.021715818 |
| *EAF1.AS1* | -2.729577 | 0.808865 | -3.374575 | 0.000739297 | 0.021817748 |
| *LIMS2* | -3.526118 | 1.046932 | -3.368048 | 0.000757025 | 0.022289276 |
| *PTTG1* | -2.288439 | 0.679582 | -3.367424 | 0.00075874 | 0.022289276 |
| *ENSG00000272931* | -3.456688 | 1.027024 | -3.365731 | 0.00076341 | 0.022304903 |
| *SNTA1* | -2.819256 | 0.837734 | -3.365337 | 0.000764503 | 0.022304903 |
| *ENSG00000214265* | -2.670897 | 0.793793 | -3.364726 | 0.000766197 | 0.022304903 |
| *DOLPP1* | -2.768039 | 0.822665 | -3.364723 | 0.000766206 | 0.022304903 |
| *ENSG00000236255* | -3.550315 | 1.055869 | -3.362458 | 0.00077252 | 0.022415999 |
| *ATP2B4* | -3.06963 | 0.913008 | -3.362105 | 0.000773506 | 0.022415999 |
| *LINC01353* | -2.637751 | 0.785238 | -3.359175 | 0.000781755 | 0.022604126 |
| *CHAF1B* | -3.847435 | 1.146343 | -3.356269 | 0.000790016 | 0.022790989 |
| *COL8A1* | -3.644123 | 1.085963 | -3.35566 | 0.00079176 | 0.022790989 |
| *TIGD5* | -2.364102 | 0.705125 | -3.352741 | 0.000800157 | 0.022899112 |
| *ENSG00000271746* | -3.33608 | 0.99508 | -3.352576 | 0.000800633 | 0.022899112 |
| *PITPNM2* | -2.415833 | 0.721296 | -3.349295 | 0.000810176 | 0.022899112 |
| *LINC00924* | -3.311481 | 0.988785 | -3.349038 | 0.000810926 | 0.022899112 |
| *FAM13A.AS1* | -2.953625 | 0.882033 | -3.348658 | 0.00081204 | 0.022899112 |
| *TRAF6* | -2.15379 | 0.643217 | -3.348465 | 0.000812606 | 0.022899112 |
| *TP53I11* | -1.71587 | 0.512513 | -3.347953 | 0.000814107 | 0.022899112 |
| *ZNF775* | -2.72437 | 0.813805 | -3.347694 | 0.000814871 | 0.022899112 |
| *ENSG00000273759* | -2.750845 | 0.821719 | -3.347671 | 0.000814938 | 0.022899112 |
| *CROT* | -2.369992 | 0.707953 | -3.347668 | 0.000814944 | 0.022899112 |
| *ZNF433* | -2.673976 | 0.79877 | -3.347618 | 0.000815092 | 0.022899112 |
| *ENSG00000274561* | -2.709768 | 0.80995 | -3.3456 | 0.000821046 | 0.023016123 |
| *C1QTNF6* | -2.877584 | 0.861073 | -3.34186 | 0.000832191 | 0.023277832 |
| *SMIM2.AS1* | -3.733199 | 1.119089 | -3.335926 | 0.000850158 | 0.023728809 |
| *ZBTB20.AS2* | -2.659404 | 0.797666 | -3.333983 | 0.00085612 | 0.023843499 |
| *DOK7* | -3.276738 | 0.98313 | -3.332964 | 0.000859262 | 0.023876867 |
| *ZNF471* | -2.85143 | 0.855671 | -3.332392 | 0.000861029 | 0.023876867 |
| *ZNF57* | -2.590116 | 0.777846 | -3.329858 | 0.000868903 | 0.0240434 |
| *NACC1* | -2.002299 | 0.601525 | -3.328707 | 0.0008725 | 0.024091108 |
| *ATP6V1C2* | -3.005478 | 0.903347 | -3.327048 | 0.000877712 | 0.024106047 |
| *ENSG00000271789* | -3.33043 | 1.001095 | -3.326788 | 0.000878532 | 0.024106047 |
| *HACD1* | -3.692228 | 1.109861 | -3.326747 | 0.000878661 | 0.024106047 |

| | | | | | |
|---|---|---|---|---|---|
| *LILRA2* | -3.737501 | 1.123749 | -3.325922 | 0.000881267 | 0.024126089 |
| *BEX5* | -3.131982 | 0.942567 | -3.322821 | 0.000891122 | 0.024300639 |
| *ENSG00000259327* | -3.714848 | 1.118012 | -3.322727 | 0.00089142 | 0.024300639 |
| *AHI1* | 2.219974 | 0.668848 | 3.319099 | 0.000903085 | 0.024510479 |
| *LY6G5C* | -3.360661 | 1.012545 | -3.319024 | 0.000903327 | 0.024510479 |
| *ENSG00000236559* | -3.424284 | 1.031985 | -3.318154 | 0.000906144 | 0.024510479 |
| *ST3GAL4* | -3.457367 | 1.042012 | -3.317972 | 0.000906737 | 0.024510479 |
| *ABL1* | -2.574678 | 0.77657 | -3.315451 | 0.000914955 | 0.024680779 |
| *ENSG00000259038* | -2.722749 | 0.82168 | -3.313638 | 0.000920906 | 0.024703184 |
| *MYLK3* | -3.605337 | 1.088054 | -3.313566 | 0.000921145 | 0.024703184 |
| *SPON2* | -2.768333 | 0.835485 | -3.313444 | 0.000921546 | 0.024703184 |
| *ZNF610* | -2.879164 | 0.869843 | -3.309982 | 0.000933019 | 0.024888387 |
| *TRAPPC10* | -2.220136 | 0.670791 | -3.309727 | 0.00093387 | 0.024888387 |
| *SELENOI* | -2.558563 | 0.773071 | -3.309611 | 0.000934257 | 0.024888387 |
| *LGALS4* | -3.399472 | 1.028797 | -3.304319 | 0.000952075 | 0.025256869 |
| *ENSG00000272195* | -3.517757 | 1.064603 | -3.30429 | 0.000952173 | 0.025256869 |
| *FRY.AS1* | -2.532258 | 0.766478 | -3.303759 | 0.000953978 | 0.025256869 |
| *PIGZ* | -3.400181 | 1.029519 | -3.302688 | 0.000957629 | 0.025262942 |
| *RNFT2* | -2.692514 | 0.815377 | -3.302171 | 0.000959394 | 0.025262942 |
| *ACSS2* | -2.530886 | 0.766478 | -3.301966 | 0.000960098 | 0.025262942 |
| *FBXL14* | -3.613859 | 1.094872 | -3.300715 | 0.000964387 | 0.025324024 |
| *STX18.AS1* | -4.258193 | 1.290357 | -3.300011 | 0.000966812 | 0.025331129 |
| *ENSG00000266990* | -3.549915 | 1.075979 | -3.299242 | 0.000969463 | 0.025331129 |
| *SLC12A9.AS1* | -3.164448 | 0.959236 | -3.298923 | 0.000970564 | 0.025331129 |
| *UBAP1L* | -2.66989 | 0.810249 | -3.295148 | 0.000983699 | 0.025621982 |
| *IGHV1.24* | -3.220389 | 0.978022 | -3.292759 | 0.000992095 | 0.025669699 |
| *ARHGEF5* | -3.584559 | 1.08868 | -3.292574 | 0.000992749 | 0.025669699 |
| *CDKL5* | -2.783422 | 0.845575 | -3.291749 | 0.000995664 | 0.025669699 |
| *NR4A3* | -3.535366 | 1.074069 | -3.291562 | 0.000996328 | 0.025669699 |
| *CLEC4A* | -2.722081 | 0.827088 | -3.291163 | 0.000997742 | 0.025669699 |
| *C22orf15* | -3.366077 | 1.022903 | -3.290709 | 0.000999352 | 0.025669699 |
| *SMIM11A* | -3.736608 | 1.135731 | -3.290046 | 0.001001712 | 0.025669699 |
| *ENSG00000272990* | -2.86761 | 0.87161 | -3.290014 | 0.001001822 | 0.025669699 |
| *ENSG00000272086* | -2.711337 | 0.824329 | -3.289143 | 0.001004929 | 0.025669699 |
| *RAB33A* | -2.565657 | 0.78022 | -3.288376 | 0.001007673 | 0.025669699 |
| *IFIT1* | -3.433522 | 1.044264 | -3.287983 | 0.00100908 | 0.025669699 |
| *CCDC13* | -3.164749 | 0.962552 | -3.287874 | 0.001009471 | 0.025669699 |
| *ARHGEF40* | -3.092842 | 0.941074 | -3.286502 | 0.001014402 | 0.025744192 |
| *USP31* | -2.702913 | 0.823181 | -3.283497 | 0.001025276 | 0.025968954 |
| *SLC39A11* | -2.678219 | 0.815911 | -3.282487 | 0.001028958 | 0.025997893 |
| *ENSG00000272871* | -3.165814 | 0.964638 | -3.281868 | 0.001031218 | 0.025997893 |
| *MCU* | -2.393401 | 0.729519 | -3.280791 | 0.001035164 | 0.025997893 |
| *MEIG1* | -3.465356 | 1.056339 | -3.280535 | 0.001036105 | 0.025997893 |
| *NR6A1* | -2.923138 | 0.891086 | -3.280421 | 0.001036521 | 0.025997893 |
| *P2RX5* | 1.089591 | 0.332212 | 3.279807 | 0.00103878 | 0.026003849 |
| *GTF2IRD1* | -2.681898 | 0.818116 | -3.278141 | 0.001044931 | 0.026102123 |
| *DCUN1D3* | -3.286731 | 1.002893 | -3.277251 | 0.00104823 | 0.026102123 |

| | | | | | |
|---|---|---|---|---|---|
| *XXYLT1* | -2.878503 | 0.878383 | -3.277046 | 0.001048994 | 0.026102123 |
| *EID3* | -2.422359 | 0.7394 | -3.276114 | 0.001052459 | 0.026102123 |
| *CDKL1* | -2.850522 | 0.870183 | -3.275774 | 0.001053727 | 0.026102123 |
| *PSMA8* | -4.391236 | 1.340645 | -3.275466 | 0.001054877 | 0.026102123 |
| *BORCS8.MEF2B* | -2.859752 | 0.873441 | -3.27412 | 0.001059916 | 0.026176478 |
| *TXK* | -3.680572 | 1.124453 | -3.273211 | 0.001063331 | 0.026210501 |
| *ENSG00000261542* | -3.380166 | 1.033256 | -3.271374 | 0.001070263 | 0.026330917 |
| *SOCS6* | -3.267891 | 0.999312 | -3.270142 | 0.001074935 | 0.026395394 |
| *ZBTB42* | -3.242272 | 0.991802 | -3.269073 | 0.001079005 | 0.026419264 |
| *FAM72A* | -3.295033 | 1.008023 | -3.268809 | 0.001080013 | 0.026419264 |
| *ZNF710.AS1* | -3.222984 | 0.986362 | -3.267549 | 0.001084832 | 0.026438442 |
| *ENSG00000259146* | -4.189593 | 1.28219 | -3.267529 | 0.001084907 | 0.026438442 |
| *TMEM44.AS1* | -2.928627 | 0.896745 | -3.265842 | 0.00109139 | 0.026546157 |
| *ZNF41* | -2.731861 | 0.836789 | -3.264695 | 0.001095821 | 0.026603648 |
| *MVB12B* | -4.582504 | 1.404302 | -3.26319 | 0.001101658 | 0.026694979 |
| *MARVELD3* | -3.289954 | 1.008383 | -3.262604 | 0.001103936 | 0.026699885 |
| *CD38* | 2.363815 | 0.72464 | 3.262053 | 0.001106086 | 0.026701713 |
| *TSSK3* | -3.5626 | 1.092368 | -3.261353 | 0.001108817 | 0.026717518 |
| *PCAT1* | -3.286347 | 1.007908 | -3.260561 | 0.001111919 | 0.026742172 |
| *LINC00449* | -3.406443 | 1.045119 | -3.259382 | 0.001116553 | 0.026803516 |
| *C10orf143* | -2.803141 | 0.86017 | -3.258821 | 0.001118762 | 0.026806526 |
| *ENSG00000245522* | -2.661117 | 0.816765 | -3.258118 | 0.001121538 | 0.026823107 |
| *BACE2* | -3.014706 | 0.925471 | -3.257482 | 0.001124055 | 0.026833426 |
| *ENSG00000226853* | -3.773671 | 1.159872 | -3.253523 | 0.001139833 | 0.027058412 |
| *MORF4L2.AS1* | -2.591477 | 0.796558 | -3.253343 | 0.001140557 | 0.027058412 |
| *CNN3* | -1.489094 | 0.457722 | -3.253272 | 0.001140844 | 0.027058412 |
| *MYO1B* | -2.758694 | 0.848056 | -3.252964 | 0.001142081 | 0.027058412 |
| *SLC29A1* | -3.017917 | 0.92788 | -3.252488 | 0.001143994 | 0.027058412 |
| *ENSG00000268746* | -3.745637 | 1.152828 | -3.249086 | 0.001157763 | 0.027297985 |
| *FGD4* | -4.428305 | 1.363001 | -3.248938 | 0.001158366 | 0.027297985 |
| *KLHL25* | -4.326892 | 1.332429 | -3.247372 | 0.001164759 | 0.027398451 |
| *ENSG00000272750* | -2.599531 | 0.800938 | -3.245606 | 0.001172008 | 0.027442551 |
| *ENSG00000273058* | -4.317765 | 1.330368 | -3.245541 | 0.001172277 | 0.027442551 |
| *PART1* | -3.420974 | 1.054113 | -3.245358 | 0.001173032 | 0.027442551 |
| *KIF5C* | -2.820185 | 0.869194 | -3.244596 | 0.001176173 | 0.027466087 |
| *HIST1H2AE* | -2.692967 | 0.830205 | -3.243736 | 0.001179729 | 0.027484571 |
| *TNK1* | -3.287398 | 1.013574 | -3.243373 | 0.001181236 | 0.027484571 |
| *ENSG00000275672* | -2.369997 | 0.730939 | -3.242401 | 0.001185272 | 0.027528688 |
| *HNRNPA1* | 0.567499 | 0.175058 | 3.24177 | 0.001187899 | 0.027539982 |
| *C3orf67* | -3.541704 | 1.092821 | -3.240882 | 0.001191605 | 0.02757618 |
| *COL6A3* | -3.39679 | 1.048272 | -3.24037 | 0.001193746 | 0.02757618 |
| *ENSG00000269482* | -3.891527 | 1.201297 | -3.239439 | 0.00119765 | 0.027616771 |
| *CTF1* | -3.607046 | 1.114355 | -3.236892 | 0.001208392 | 0.027755861 |
| *PLSCR3* | -3.6174 | 1.117743 | -3.236344 | 0.001210715 | 0.027755861 |
| *STS* | -3.334793 | 1.030563 | -3.235895 | 0.001212618 | 0.027755861 |
| *ENSG00000177699* | -4.555651 | 1.407939 | -3.235688 | 0.001213498 | 0.027755861 |
| *DLGAP3* | -3.306057 | 1.02187 | -3.2353 | 0.00121515 | 0.027755861 |

| | | | | | |
|---|---|---|---|---|---|
| *CDK18* | -2.725222 | 0.84243 | -3.234954 | 0.001216624 | 0.027755861 |
| *METTL6* | -1.881308 | 0.581672 | -3.23431 | 0.00121937 | 0.027769269 |
| *ZNF850* | -2.825405 | 0.874339 | -3.231475 | 0.00123153 | 0.027996646 |
| *CYP27A1* | -4.813277 | 1.490325 | -3.229682 | 0.001239278 | 0.028077135 |
| *CENPE* | -3.723183 | 1.152814 | -3.229646 | 0.001239435 | 0.028077135 |
| *ENSG00000240291* | -2.687229 | 0.832199 | -3.229073 | 0.001241924 | 0.028084065 |
| *FRAT1* | -3.69321 | 1.144013 | -3.228294 | 0.001245309 | 0.028086187 |
| *C2CD2L* | -2.978174 | 0.922593 | -3.228047 | 0.001246383 | 0.028086187 |
| *ENSG00000272941* | -2.531636 | 0.784407 | -3.227454 | 0.001248971 | 0.0280953 |
| *ENSG00000273271* | -3.192374 | 0.989627 | -3.225837 | 0.001256048 | 0.02818437 |
| *PLCH2* | -3.593441 | 1.114055 | -3.225549 | 0.001257312 | 0.02818437 |
| *ENSG00000255455* | -3.411417 | 1.057918 | -3.224653 | 0.001261254 | 0.028195179 |
| *ENSG00000239407* | -3.713598 | 1.151719 | -3.224395 | 0.001262391 | 0.028195179 |
| *CRX* | -3.204929 | 0.994154 | -3.223775 | 0.001265129 | 0.028195179 |
| *SMIM10* | -3.299909 | 1.023723 | -3.223439 | 0.001266612 | 0.028195179 |
| *LINC02361* | -2.756593 | 0.8553 | -3.222956 | 0.00126875 | 0.028195179 |
| *SLC49A4* | -2.663981 | 0.827114 | -3.220813 | 0.001278276 | 0.028322455 |
| *SLC9A6* | -2.65395 | 0.824035 | -3.220677 | 0.00127888 | 0.028322455 |
| *ENSG00000260625* | -3.462847 | 1.076133 | -3.217861 | 0.001291504 | 0.02850521 |
| *LIX1L.AS1* | -2.668424 | 0.829257 | -3.217848 | 0.001291563 | 0.02850521 |
| *IPO13* | -2.46656 | 0.76674 | -3.216943 | 0.001295642 | 0.028510616 |
| *ALG9* | -2.496164 | 0.775975 | -3.216811 | 0.001296239 | 0.028510616 |
| *FCSK* | -2.655909 | 0.825894 | -3.215798 | 0.001300825 | 0.028562661 |
| *DNAH14* | -3.137733 | 0.975979 | -3.214959 | 0.00130463 | 0.028597394 |
| *BMPR1A* | -2.926079 | 0.910516 | -3.213649 | 0.001310596 | 0.028649867 |
| *KANK2* | -3.56335 | 1.108884 | -3.213456 | 0.001311477 | 0.028649867 |
| *ASRGL1* | -3.943349 | 1.227324 | -3.212964 | 0.001313729 | 0.028650424 |
| *ZNF577* | -2.467969 | 0.768442 | -3.211651 | 0.001319748 | 0.028732992 |
| *SLC16A13* | -2.106497 | 0.656204 | -3.210124 | 0.001326778 | 0.028837257 |
| *PLCD1* | -2.840358 | 0.884958 | -3.209596 | 0.001329217 | 0.028841553 |
| *ENSG00000273289* | -3.330307 | 1.038058 | -3.20821 | 0.00133564 | 0.028932111 |
| *ENSG00000242282* | -2.601608 | 0.811319 | -3.206641 | 0.001342945 | 0.028984954 |
| *ZBTB8A* | -2.561067 | 0.798731 | -3.206422 | 0.001343967 | 0.028984954 |
| *MBLAC2* | -2.581746 | 0.805338 | -3.205791 | 0.001346917 | 0.028984954 |
| *AMACR* | -2.772297 | 0.864788 | -3.205754 | 0.00134709 | 0.028984954 |
| *ENSG00000271971* | -2.8512 | 0.889725 | -3.204584 | 0.001352577 | 0.029017126 |
| *ARHGEF39* | -3.218791 | 1.004636 | -3.203937 | 0.001355624 | 0.029017126 |
| *SULT1A1* | -3.536972 | 1.103973 | -3.203857 | 0.001355999 | 0.029017126 |
| *ENSG00000184441* | -3.5364 | 1.104054 | -3.203104 | 0.001359549 | 0.029017126 |
| *ENSG00000261353* | -3.449319 | 1.07689 | -3.203038 | 0.001359861 | 0.029017126 |
| *MAP4K3* | -2.833783 | 0.884855 | -3.202538 | 0.001362222 | 0.029019397 |
| *TRAM2.AS1* | -2.463629 | 0.769666 | -3.200906 | 0.001369961 | 0.029091546 |
| *RBFADN* | -3.455083 | 1.079491 | -3.20066 | 0.00137113 | 0.029091546 |
| *OSBPL1A* | -2.623251 | 0.81975 | -3.200064 | 0.001373971 | 0.029091546 |
| *ENSG00000256633* | -4.158213 | 1.299713 | -3.199331 | 0.001377467 | 0.029091546 |
| *ENSG00000228395* | -3.324904 | 1.03948 | -3.198623 | 0.001380858 | 0.029091546 |
| *ENSG00000262222* | -2.550457 | 0.797443 | -3.198295 | 0.001382429 | 0.029091546 |

| | | | | | |
|---|---|---|---|---|---|
| *CCDC24* | -2.830599 | 0.885056 | -3.198216 | 0.001382805 | 0.029091546 |
| *BORCS8* | -1.598538 | 0.499851 | -3.19803 | 0.001383697 | 0.029091546 |
| *ENSG00000267698* | -3.548699 | 1.110317 | -3.196114 | 0.001392921 | 0.029186276 |
| *ENSG00000279081* | -2.67103 | 0.835816 | -3.195716 | 0.001394844 | 0.029186276 |
| *PDLIM2* | -1.717535 | 0.537455 | -3.195682 | 0.001395007 | 0.029186276 |
| *SGCB* | -2.19169 | 0.686496 | -3.192576 | 0.001410098 | 0.029454107 |
| *UPB1* | -3.188169 | 0.998839 | -3.191876 | 0.00141352 | 0.029477742 |
| *FNTB* | -2.595851 | 0.813536 | -3.190824 | 0.001418677 | 0.029526241 |
| *AMPD2* | -2.146233 | 0.672746 | -3.19026 | 0.001421448 | 0.029526241 |
| *ENSG00000235978* | -3.473085 | 1.088741 | -3.19 | 0.00142273 | 0.029526241 |
| *SAXO2* | -4.182969 | 1.312479 | -3.187076 | 0.001437192 | 0.029778347 |
| *ENSG00000259959* | -3.629273 | 1.138971 | -3.186448 | 0.001440312 | 0.029795008 |
| *CA3.AS1* | -3.729472 | 1.170773 | -3.185477 | 0.001445154 | 0.029847179 |
| *TBC1D8B* | -2.579711 | 0.810084 | -3.184497 | 0.001450057 | 0.029849431 |
| *TP53INP2* | -3.238391 | 1.016944 | -3.184435 | 0.00145037 | 0.029849431 |
| *PLP2* | -0.862914 | 0.27101 | -3.184065 | 0.001452222 | 0.029849431 |
| *PRR11* | -3.241987 | 1.018503 | -3.183089 | 0.001457127 | 0.029902472 |
| *ZNF438* | -2.485999 | 0.781195 | -3.182302 | 0.001461092 | 0.029936094 |
| *PHKB* | 2.170914 | 0.682285 | 3.18183 | 0.001463476 | 0.029937273 |
| *HLX* | -2.494917 | 0.784313 | -3.181023 | 0.00146756 | 0.029937805 |
| *CD5* | -2.690642 | 0.845975 | -3.180522 | 0.0014701 | 0.029937805 |
| *NRL* | -2.856248 | 0.898143 | -3.180171 | 0.001471879 | 0.029937805 |
| *ENSG00000261140* | -3.534467 | 1.111472 | -3.179989 | 0.001472809 | 0.029937805 |
| *ACVR2B* | -3.402902 | 1.070314 | -3.179349 | 0.001476061 | 0.029956586 |
| *ENSG00000227896* | -3.630343 | 1.142303 | -3.178092 | 0.001482478 | 0.029976508 |
| *INTS8* | -1.576284 | 0.496052 | -3.17766 | 0.00148469 | 0.029976508 |
| *ENSG00000278740* | -3.363382 | 1.05856 | -3.177317 | 0.001486443 | 0.029976508 |
| *HEATR4* | -3.406248 | 1.072195 | -3.176892 | 0.001488623 | 0.029976508 |
| *XXYLT1.AS2* | -2.690836 | 0.847006 | -3.176879 | 0.001488691 | 0.029976508 |
| *ENSG00000260572* | -3.372103 | 1.061797 | -3.175844 | 0.001494014 | 0.03003669 |
| *TMPRSS13* | -3.241814 | 1.020995 | -3.175151 | 0.001497585 | 0.030061514 |
| *WNT7B* | -3.870993 | 1.219642 | -3.173875 | 0.001504183 | 0.030070781 |
| *FOXP4* | -2.109524 | 0.664685 | -3.173719 | 0.001504992 | 0.030070781 |
| *LINC00526* | -2.152924 | 0.678363 | -3.173706 | 0.001505058 | 0.030070781 |
| *PIP5KL1* | -3.75477 | 1.183354 | -3.17299 | 0.001508775 | 0.0300983 |
| *ABHD1* | -3.190382 | 1.005634 | -3.172507 | 0.001511291 | 0.030101823 |
| *TTLL11* | -3.329322 | 1.050177 | -3.170247 | 0.001523093 | 0.030290003 |
| *ENSG00000254777* | -4.287254 | 1.352559 | -3.169735 | 0.001525783 | 0.030296677 |
| *MTHFD1L* | -2.49467 | 0.78741 | -3.168196 | 0.001533879 | 0.030335505 |
| *SGSM1* | -3.643091 | 1.15007 | -3.167714 | 0.001536428 | 0.030335505 |
| *PSAT1* | -3.667438 | 1.157793 | -3.167612 | 0.001536965 | 0.030335505 |
| *ENSG00000226822* | -4.358374 | 1.376036 | -3.167341 | 0.001538398 | 0.030335505 |
| *ENSG00000175773* | -3.843078 | 1.213427 | -3.167128 | 0.001539526 | 0.030335505 |
| *ENSG00000244733* | -4.190713 | 1.323423 | -3.166572 | 0.001542473 | 0.030344363 |
| *ENSG00000267213* | -4.227128 | 1.335099 | -3.166154 | 0.001544692 | 0.030344363 |
| *HAP1* | -4.041583 | 1.27703 | -3.16483 | 0.001551737 | 0.030436288 |
| *ITPRIPL2* | -3.367078 | 1.064071 | -3.164334 | 0.001554381 | 0.030441738 |

| | | | | | |
|---|---|---|---|---|---|
| *CAPS2* | -3.395141 | 1.073109 | -3.163836 | 0.001557045 | 0.030447568 |
| *NRAV* | -2.629968 | 0.831394 | -3.163325 | 0.001559782 | 0.030454805 |
| *ENSG00000250222* | -2.641577 | 0.83535 | -3.162239 | 0.001565609 | 0.030522259 |
| *GGT1* | -3.670505 | 1.16133 | -3.160605 | 0.001574417 | 0.030551463 |
| *C9orf40* | -2.408985 | 0.762364 | -3.159888 | 0.001578297 | 0.030551463 |
| *ENSG00000223969* | -3.399637 | 1.076025 | -3.15944 | 0.001580725 | 0.030551463 |
| *ITIH4* | -4.04646 | 1.280796 | -3.159332 | 0.001581312 | 0.030551463 |
| *ENSG00000273456* | -2.643123 | 0.836609 | -3.15933 | 0.001581321 | 0.030551463 |
| *KNTC1* | -2.323535 | 0.735453 | -3.159324 | 0.001581353 | 0.030551463 |
| *ZMIZ1* | -2.254543 | 0.713894 | -3.158091 | 0.001588062 | 0.030563108 |
| *MAFF* | -2.756849 | 0.873029 | -3.157797 | 0.001589661 | 0.030563108 |
| *SLC5A10* | -3.335722 | 1.056414 | -3.157591 | 0.001590788 | 0.030563108 |
| *UBXN6* | -1.140686 | 0.361266 | -3.157468 | 0.001591457 | 0.030563108 |
| *SYNPO* | -3.125293 | 0.989969 | -3.15696 | 0.001594231 | 0.030570742 |
| *HMMR* | -4.092758 | 1.297084 | -3.155354 | 0.001603036 | 0.030693844 |
| *ENSG00000181097* | -3.177751 | 1.007524 | -3.154022 | 0.001610371 | 0.030720727 |
| *ENSG00000272817* | -2.923508 | 0.926944 | -3.15392 | 0.001610934 | 0.030720727 |
| *FBXL16* | -3.857846 | 1.223238 | -3.153799 | 0.001611603 | 0.030720727 |
| *POLE2* | -2.776395 | 0.880717 | -3.152425 | 0.001619201 | 0.030819918 |
| *MMACHC* | -3.264064 | 1.035651 | -3.151702 | 0.001623217 | 0.030839888 |
| *ENSG00000241764* | -2.518323 | 0.799221 | -3.150972 | 0.001627279 | 0.030839888 |
| *TPH1* | -3.935287 | 1.248923 | -3.150943 | 0.001627441 | 0.030839888 |
| *LMLN* | -3.157279 | 1.002423 | -3.149649 | 0.00163467 | 0.030931318 |
| *ENSG00000248559* | -2.144734 | 0.681303 | -3.147989 | 0.001643977 | 0.03102509 |
| *PROCR* | -3.936329 | 1.25047 | -3.147879 | 0.001644597 | 0.03102509 |
| *USP27X.AS1* | -3.765597 | 1.196386 | -3.147477 | 0.001646859 | 0.03102509 |
| *CAMK4* | -4.019583 | 1.277559 | -3.146301 | 0.001653499 | 0.031104637 |
| *ENSG00000273117* | -3.295257 | 1.047709 | -3.145202 | 0.001659723 | 0.031165225 |
| *ENSG00000256116* | -4.110927 | 1.307182 | -3.144877 | 0.001661564 | 0.031165225 |
| *NKAPL* | -2.953062 | 0.939362 | -3.14369 | 0.00166832 | 0.031207698 |
| *EDN1* | -4.042754 | 1.286016 | -3.143627 | 0.001668679 | 0.031207698 |
| *ENSG00000254452* | -3.856862 | 1.22713 | -3.142994 | 0.001672295 | 0.031229921 |
| *TCAP* | -3.288011 | 1.046289 | -3.142546 | 0.001674853 | 0.031232358 |
| *ENSG00000174171* | -3.716082 | 1.182719 | -3.141982 | 0.001678081 | 0.031233876 |
| *ZNF726* | -2.472035 | 0.786976 | -3.141182 | 0.001682672 | 0.031233876 |
| *LINC01550* | -3.387971 | 1.078623 | -3.141014 | 0.001683638 | 0.031233876 |
| *POGLUT3* | -2.880948 | 0.917254 | -3.140839 | 0.001684644 | 0.031233876 |
| *BAIAP2.DT* | -3.796033 | 1.208905 | -3.140058 | 0.001689142 | 0.031253258 |
| *KIAA0391* | -3.810944 | 1.213782 | -3.139728 | 0.001691046 | 0.031253258 |
| *GSTM3* | -3.70896 | 1.181567 | -3.139017 | 0.001695154 | 0.031253258 |
| *NRSN2* | -2.410987 | 0.768301 | -3.138077 | 0.001700599 | 0.031253258 |
| *LINC01963* | -3.854619 | 1.22839 | -3.137945 | 0.00170137 | 0.031253258 |
| *WNT2B* | -2.346928 | 0.747919 | -3.137944 | 0.001701371 | 0.031253258 |
| *KIF21B* | -2.625472 | 0.836746 | -3.137717 | 0.001702692 | 0.031253258 |
| *ENSG00000229178* | -2.979573 | 0.949854 | -3.136874 | 0.001707593 | 0.031298565 |
| *MAS1* | -4.024431 | 1.283417 | -3.135716 | 0.001714352 | 0.031366383 |
| *SPDYE3* | -3.679669 | 1.174046 | -3.134177 | 0.00172337 | 0.031366383 |

| | | | | | |
|---|---|---|---|---|---|
| *STKLD1* | -3.581791 | 1.142841 | -3.134111 | 0.001723756 | 0.031366383 |
| *SLC25A10* | -3.116064 | 0.994386 | -3.133656 | 0.00172643 | 0.031366383 |
| *BHLHE40.AS1* | -3.817399 | 1.218252 | -3.133506 | 0.001727312 | 0.031366383 |
| *USP51* | -2.70076 | 0.86217 | -3.132517 | 0.001733147 | 0.031366383 |
| *SLC22A18* | -2.613754 | 0.834412 | -3.132452 | 0.001733527 | 0.031366383 |
| *ACVR1* | -2.57669 | 0.822606 | -3.13235 | 0.001734132 | 0.031366383 |
| *ENSG00000232347* | -3.062643 | 0.977759 | -3.132307 | 0.001734384 | 0.031366383 |
| *RAB11FIP5* | -2.877695 | 0.918778 | -3.132089 | 0.00173567 | 0.031366383 |
| *NECTIN1* | -3.315639 | 1.059253 | -3.130169 | 0.001747058 | 0.03144716 |
| *FBXO36* | -3.697562 | 1.181515 | -3.129509 | 0.001750986 | 0.03144716 |
| *ENSG00000215022* | -3.59168 | 1.147969 | -3.128725 | 0.001755667 | 0.03144716 |
| *AMBRA1* | -2.499262 | 0.798921 | -3.128297 | 0.001758225 | 0.03144716 |
| *LY9* | -1.434643 | 0.458694 | -3.12767 | 0.001761978 | 0.03144716 |
| *TLE2* | -3.853073 | 1.23213 | -3.127165 | 0.001765007 | 0.03144716 |
| *ANKRD50* | -3.848943 | 1.230835 | -3.127099 | 0.001765407 | 0.03144716 |
| *ENSG00000273199* | -2.446917 | 0.782594 | -3.126673 | 0.001767962 | 0.03144716 |
| *OSM* | -3.291731 | 1.052908 | -3.126325 | 0.001770058 | 0.03144716 |
| *POLN* | -3.398434 | 1.087088 | -3.126181 | 0.001770927 | 0.03144716 |
| *ENSG00000259407* | -4.041831 | 1.292935 | -3.12609 | 0.001771473 | 0.03144716 |
| *C11orf65* | -3.886708 | 1.243541 | -3.125516 | 0.001774933 | 0.03144716 |
| *CRYL1* | -2.166704 | 0.693242 | -3.125465 | 0.001775244 | 0.03144716 |
| *C2orf15* | -2.650817 | 0.848202 | -3.125218 | 0.001776734 | 0.03144716 |
| *ZNF492* | -3.117531 | 0.997604 | -3.125018 | 0.00177794 | 0.03144716 |
| *LINC01806* | -4.177491 | 1.336982 | -3.124568 | 0.001780664 | 0.03144716 |
| *MTF1* | -1.636604 | 0.523834 | -3.124278 | 0.001782419 | 0.03144716 |
| *CORO2A* | -2.516519 | 0.805545 | -3.123995 | 0.001784132 | 0.03144716 |
| *LOR* | -3.569483 | 1.142966 | -3.122999 | 0.001790181 | 0.031458384 |
| *POLR3G* | -3.812949 | 1.221059 | -3.122657 | 0.001792263 | 0.031458384 |
| *ENSG00000250155* | -3.375344 | 1.081448 | -3.121133 | 0.001801567 | 0.031458384 |
| *ZNF300* | -2.830044 | 0.906779 | -3.120987 | 0.001802463 | 0.031458384 |
| *KREMEN1* | -3.185174 | 1.02064 | -3.120762 | 0.001803836 | 0.031458384 |
| *ZGRF1* | -2.53478 | 0.812265 | -3.120631 | 0.001804641 | 0.031458384 |
| *NEURL1* | -3.873088 | 1.241478 | -3.119739 | 0.001810115 | 0.031458384 |
| *SOBP* | -4.099371 | 1.314163 | -3.119378 | 0.001812333 | 0.031458384 |
| *KCNH4* | -3.435688 | 1.101489 | -3.11913 | 0.001813858 | 0.031458384 |
| *KIF9.AS1* | -3.558305 | 1.140872 | -3.118934 | 0.001815063 | 0.031458384 |
| *LINC01521* | -3.047839 | 0.977237 | -3.118834 | 0.001815681 | 0.031458384 |
| *ENSG00000276131* | -3.7968 | 1.217445 | -3.118662 | 0.00181674 | 0.031458384 |
| *C14orf132* | -3.863501 | 1.238913 | -3.118459 | 0.001817994 | 0.031458384 |
| *CNKSR3* | -3.474014 | 1.114074 | -3.118296 | 0.001818997 | 0.031458384 |
| *CACNA1C.AS2* | -4.152247 | 1.332223 | -3.11678 | 0.001828378 | 0.031553075 |
| *TUBB4A* | -3.748202 | 1.20265 | -3.116619 | 0.001829377 | 0.031553075 |
| *ENSG00000275454* | -2.594935 | 0.833471 | -3.113407 | 0.001849405 | 0.031855823 |
| *CHRNA10* | -3.528965 | 1.134522 | -3.110531 | 0.001867515 | 0.032082555 |
| *RBPMS* | -3.234352 | 1.040267 | -3.109157 | 0.001876218 | 0.032082555 |
| *NAP1L5* | -2.996347 | 0.96373 | -3.109114 | 0.001876495 | 0.032082555 |
| *FASTKD5* | -1.962821 | 0.631447 | -3.108451 | 0.001880707 | 0.032082555 |

| | | | | | |
|---|---|---|---|---|---|
| *ADAMDEC1* | -4.010105 | 1.290275 | -3.107946 | 0.001883925 | 0.032082555 |
| *WARS2.AS1* | -2.817413 | 0.906541 | -3.107871 | 0.001884404 | 0.032082555 |
| *ENSG00000267309* | -3.936455 | 1.266633 | -3.10781 | 0.001884794 | 0.032082555 |
| *ENSG00000277283* | -2.31087 | 0.743595 | -3.107701 | 0.001885484 | 0.032082555 |
| *RAP1GAP2* | -4.194106 | 1.349956 | -3.106847 | 0.00189094 | 0.032082555 |
| *FAM174B* | -2.931478 | 0.94365 | -3.106532 | 0.001892959 | 0.032082555 |
| *KLHL11* | -3.831711 | 1.233568 | -3.106203 | 0.001895067 | 0.032082555 |
| *MCM4* | -2.429717 | 0.782215 | -3.1062 | 0.001895082 | 0.032082555 |
| *LRRC29* | -2.998494 | 0.965536 | -3.105522 | 0.001899432 | 0.032082555 |
| *DPY19L4* | -2.362026 | 0.760652 | -3.105263 | 0.001901098 | 0.032082555 |
| *ENSG00000260276* | -3.507886 | 1.129703 | -3.105139 | 0.001901896 | 0.032082555 |
| *PYGM* | -2.809763 | 0.905146 | -3.104212 | 0.001907868 | 0.032082555 |
| *ZNF674* | -2.933958 | 0.94518 | -3.104125 | 0.001908428 | 0.032082555 |
| *DRAM1* | -1.935519 | 0.623643 | -3.10357 | 0.00191201 | 0.032082555 |
| *GLS2* | -3.189181 | 1.027678 | -3.103289 | 0.001913826 | 0.032082555 |
| *ENSG00000274315* | -3.142751 | 1.012844 | -3.102896 | 0.001916367 | 0.032082555 |
| *ENSG00000269929* | -3.593978 | 1.158311 | -3.102773 | 0.001917166 | 0.032082555 |
| *PRELID2* | -2.527116 | 0.814481 | -3.102733 | 0.001917423 | 0.032082555 |
| *DISP2* | -3.139046 | 1.011842 | -3.102309 | 0.001920176 | 0.032086881 |
| *VANGL2* | -3.354456 | 1.082078 | -3.100014 | 0.001935116 | 0.032107808 |
| *RPS10P7* | -2.548739 | 0.822216 | -3.099841 | 0.001936242 | 0.032107808 |
| *ENSG00000267226* | -3.668338 | 1.1836 | -3.099304 | 0.001939758 | 0.032107808 |
| *DNM3* | -2.846744 | 0.918532 | -3.099232 | 0.00194023 | 0.032107808 |
| *ANKRD23* | -3.359365 | 1.083993 | -3.099064 | 0.001941328 | 0.032107808 |
| *ZNF440* | -2.227863 | 0.718888 | -3.09904 | 0.001941491 | 0.032107808 |
| *ENSG00000269226* | -3.34363 | 1.079087 | -3.098572 | 0.001944556 | 0.032107808 |
| *ENSG00000276216* | -3.326284 | 1.073567 | -3.098347 | 0.001946036 | 0.032107808 |
| *ENSG00000239791* | -3.577288 | 1.154598 | -3.098298 | 0.001946359 | 0.032107808 |
| *ENSG00000254837* | -2.609706 | 0.842334 | -3.098183 | 0.001947111 | 0.032107808 |
| *OR2A1.AS1* | -2.525554 | 0.81538 | -3.097395 | 0.001952296 | 0.032107808 |
| *ENSG00000268836* | -3.784681 | 1.222023 | -3.097063 | 0.001954485 | 0.032107808 |
| *ERBB2* | -2.845371 | 0.918836 | -3.096713 | 0.001956791 | 0.032107808 |
| *EFCAB7* | -2.173704 | 0.70196 | -3.09662 | 0.001957406 | 0.032107808 |
| *RMC1* | -1.646809 | 0.531846 | -3.0964 | 0.001958858 | 0.032107808 |
| *CISD1* | -1.279855 | 0.413394 | -3.095964 | 0.00196174 | 0.032114136 |
| *NT5DC4* | -3.238852 | 1.046359 | -3.095353 | 0.001965785 | 0.032139465 |
| *PAM16* | -3.206018 | 1.035878 | -3.094975 | 0.001968292 | 0.032139608 |
| *TMEM254.AS1* | -3.5232 | 1.1391 | -3.092967 | 0.00198166 | 0.032201624 |
| *ORC6* | -2.310332 | 0.746965 | -3.092959 | 0.001981713 | 0.032201624 |
| *ENSG00000276791* | -3.676214 | 1.188575 | -3.092958 | 0.001981722 | 0.032201624 |
| *CCDC136* | -2.984403 | 0.96492 | -3.092901 | 0.0019821 | 0.032201624 |
| *ZNF630* | -2.856833 | 0.923869 | -3.092248 | 0.001986466 | 0.03223185 |
| *MAP2K6* | -2.905586 | 0.93999 | -3.091083 | 0.00199428 | 0.032317886 |
| *ITPR1.DT* | -3.255172 | 1.053221 | -3.090683 | 0.001996967 | 0.032320717 |
| *GNA11* | -2.844184 | 0.920514 | -3.089779 | 0.002003056 | 0.032369376 |
| *ZSCAN20* | -3.426709 | 1.109263 | -3.089177 | 0.00200712 | 0.032369376 |
| *DOCK6* | -3.105296 | 1.005237 | -3.089118 | 0.00200752 | 0.032369376 |

| | | | | | |
|---|---|---|---|---|---|
| *RIMS3* | -3.763639 | 1.218564 | -3.088586 | 0.002011116 | 0.032382558 |
| *TGM2* | -3.343834 | 1.082795 | -3.088152 | 0.002014054 | 0.032382558 |
| *EEF1AKMT3* | -4.048325 | 1.311036 | -3.087882 | 0.002015888 | 0.032382558 |
| *FSD2* | -3.8455 | 1.245609 | -3.087246 | 0.002020206 | 0.03241146 |
| *ENSG00000237976* | -2.236498 | 0.724784 | -3.085746 | 0.002030425 | 0.032503669 |
| *PCOTH* | -3.990069 | 1.293101 | -3.085661 | 0.002031006 | 0.032503669 |
| *RPA4* | -2.852244 | 0.925416 | -3.082119 | 0.002055324 | 0.03274388 |
| *ENSG00000232698* | -3.36851 | 1.092936 | -3.082076 | 0.002055623 | 0.03274388 |
| *STK4* | -0.653587 | 0.212077 | -3.081834 | 0.002057294 | 0.03274388 |
| *KDM8* | -2.48971 | 0.807878 | -3.081791 | 0.002057595 | 0.03274388 |
| *NPIPB13* | -3.212262 | 1.042513 | -3.081268 | 0.002061208 | 0.03274388 |
| *NPR2* | -3.768583 | 1.223067 | -3.081257 | 0.002061284 | 0.03274388 |
| *SNX15* | -2.514764 | 0.81634 | -3.080533 | 0.002066301 | 0.032747506 |
| *ENSG00000272913* | -3.001542 | 0.974477 | -3.080157 | 0.002068913 | 0.032747506 |
| *MROH8* | -3.073152 | 0.997736 | -3.080124 | 0.002069148 | 0.032747506 |
| *ACSM1* | -2.673816 | 0.868197 | -3.079734 | 0.002071859 | 0.032750133 |
| *ENSG00000260285* | -3.83416 | 1.245254 | -3.079018 | 0.002076838 | 0.032788557 |
| *MIR4453HG* | -1.998438 | 0.649344 | -3.077628 | 0.002086555 | 0.032801894 |
| *TMEM104* | -3.029592 | 0.984552 | -3.077126 | 0.002090068 | 0.032801894 |
| *IGFBP4* | -2.658555 | 0.864004 | -3.077018 | 0.002090825 | 0.032801894 |
| *AP4B1.AS1* | -3.010317 | 0.978352 | -3.076925 | 0.002091476 | 0.032801894 |
| *ENSG00000259071* | -3.530101 | 1.147316 | -3.076833 | 0.002092126 | 0.032801894 |
| *ENSG00000267731* | -4.091151 | 1.329753 | -3.076624 | 0.002093589 | 0.032801894 |
| *TRIP13* | -3.031726 | 0.985495 | -3.076348 | 0.002095528 | 0.032801894 |
| *FSTL3* | -2.639375 | 0.858126 | -3.075742 | 0.002099794 | 0.032828738 |
| *ZNF572* | -2.841979 | 0.924276 | -3.074816 | 0.002106326 | 0.032890895 |
| *CHCHD4* | -1.826911 | 0.594375 | -3.073668 | 0.002114447 | 0.032977683 |
| *ENSG00000236266* | -4.84097 | 1.575424 | -3.072805 | 0.00212057 | 0.03298794 |
| *SCARB1* | -3.632663 | 1.182257 | -3.07265 | 0.002121674 | 0.03298794 |
| *LINC02132* | -3.02804 | 0.985618 | -3.072227 | 0.002124684 | 0.03298794 |
| *KHDRBS3* | -3.798935 | 1.236648 | -3.071961 | 0.002126573 | 0.03298794 |
| *FANCC* | -2.918899 | 0.950233 | -3.071772 | 0.002127923 | 0.03298794 |
| *BAG3* | -3.99269 | 1.300483 | -3.07016 | 0.002139444 | 0.033126619 |
| *ENSG00000254615* | -2.43037 | 0.792029 | -3.068534 | 0.002151115 | 0.033267307 |
| *EIF2S3B* | -3.139777 | 1.02356 | -3.067506 | 0.00215853 | 0.0333419 |
| *ENSG00000234694* | -2.853229 | 0.930286 | -3.067047 | 0.00216185 | 0.033353149 |
| *ENSG00000248544* | -2.90612 | 0.947798 | -3.066181 | 0.002168122 | 0.033371621 |
| *CLBA1* | -3.093126 | 1.008802 | -3.066138 | 0.002168432 | 0.033371621 |
| *ENSG00000270589* | -3.778253 | 1.232485 | -3.065557 | 0.002172649 | 0.033371621 |
| *DYNC2H1* | -3.014958 | 0.98381 | -3.064572 | 0.002179819 | 0.033371621 |
| *LRRC46* | -3.828617 | 1.249503 | -3.064111 | 0.002183177 | 0.033371621 |
| *C3orf14* | -2.884141 | 0.941285 | -3.064045 | 0.002183662 | 0.033371621 |
| *PPM1J* | -3.896755 | 1.271845 | -3.063861 | 0.002185003 | 0.033371621 |
| *CFAP45* | -4.123587 | 1.346085 | -3.063392 | 0.002188429 | 0.033371621 |
| *ENSG00000272323* | -2.936333 | 0.9586 | -3.063147 | 0.002190224 | 0.033371621 |
| *DACT1* | -3.872347 | 1.264179 | -3.063131 | 0.00219034 | 0.033371621 |
| *ENSG00000270091* | -3.124602 | 1.020228 | -3.062651 | 0.002193855 | 0.033371621 |

| | | | | | |
|---|---|---|---|---|---|
| *PPIAL4G* | -2.412978 | 0.787936 | -3.062402 | 0.002195687 | 0.033371621 |
| *RNLS* | -3.000897 | 0.980012 | -3.062103 | 0.00219788 | 0.033371621 |
| *ENSG00000253704* | -3.942453 | 1.287583 | -3.061902 | 0.002199358 | 0.033371621 |
| *LINC00525* | -4.041583 | 1.320233 | -3.061265 | 0.002204038 | 0.033403254 |
| *ENSG00000275418* | -3.99256 | 1.304918 | -3.059624 | 0.002216148 | 0.033459292 |
| *ASB16* | -3.98089 | 1.301131 | -3.059561 | 0.002216615 | 0.033459292 |
| *ENSG00000267281* | -3.805913 | 1.244013 | -3.059384 | 0.002217926 | 0.033459292 |
| *ENSG00000232748* | -3.44328 | 1.125492 | -3.059356 | 0.002218138 | 0.033459292 |
| *ANKS6* | -3.316957 | 1.084596 | -3.058243 | 0.002226388 | 0.033538695 |
| *SLC38A11* | -3.392451 | 1.109443 | -3.057796 | 0.002229717 | 0.033538695 |
| *ENSG00000250541* | -3.977212 | 1.300794 | -3.057526 | 0.002231723 | 0.033538695 |
| *LINC00863* | -2.994742 | 0.97956 | -3.057232 | 0.002233909 | 0.033538695 |
| *NSMCE1.DT* | -3.0141 | 0.986017 | -3.056842 | 0.002236822 | 0.033538695 |
| *ENSG00000237596* | -3.100497 | 1.014514 | -3.056139 | 0.002242071 | 0.033538695 |
| *LAT* | -2.535317 | 0.829618 | -3.056006 | 0.002243064 | 0.033538695 |
| *LRRC37A3* | -3.11424 | 1.019109 | -3.055847 | 0.002244254 | 0.033538695 |
| *TMEM9B.AS1* | -2.958333 | 0.968531 | -3.054454 | 0.002254705 | 0.033655784 |
| *FUT4* | -3.818363 | 1.25058 | -3.053275 | 0.002263586 | 0.033684844 |
| *PIF1* | -4.067583 | 1.332237 | -3.053197 | 0.002264174 | 0.033684844 |
| *USP18* | -3.092165 | 1.012778 | -3.053153 | 0.002264505 | 0.033684844 |
| *FOXRED2* | -2.500383 | 0.819112 | -3.052554 | 0.002269029 | 0.033713153 |
| *LINC01569* | -2.944353 | 0.964902 | -3.051452 | 0.002277373 | 0.033792458 |
| *ZNF490* | -3.158572 | 1.035242 | -3.051047 | 0.002280449 | 0.033792458 |
| *PET117* | -1.552173 | 0.508858 | -3.050306 | 0.002286084 | 0.033792458 |
| *ENSG00000236723* | -3.539753 | 1.160491 | -3.050219 | 0.002286746 | 0.033792458 |
| *LSINCT5* | -3.162202 | 1.036747 | -3.05012 | 0.002287498 | 0.033792458 |
| *ENSG00000269514* | -2.805407 | 0.920125 | -3.048941 | 0.002296492 | 0.033826937 |
| *ENSG00000259536* | -3.711993 | 1.217513 | -3.048832 | 0.002297333 | 0.033826937 |
| *ENSG00000278831* | -4.898547 | 1.606956 | -3.048338 | 0.002301107 | 0.033826937 |
| *TMEM237* | -2.83857 | 0.931261 | -3.048093 | 0.002302983 | 0.033826937 |
| *TMEM170B* | -3.107723 | 1.019637 | -3.047872 | 0.002304682 | 0.033826937 |
| *PEX12* | -2.92952 | 0.961328 | -3.047366 | 0.002308564 | 0.033826937 |
| *ENSG00000274828* | -2.982916 | 0.97891 | -3.04718 | 0.002309993 | 0.033826937 |
| *RABL2A* | -1.717889 | 0.563785 | -3.047067 | 0.002310863 | 0.033826937 |
| *FAM229A* | -2.250589 | 0.738792 | -3.046311 | 0.002316682 | 0.03387357 |
| *POLR2D* | -1.38385 | 0.454533 | -3.044555 | 0.002330249 | 0.034002458 |
| *TPM2* | -3.96528 | 1.30265 | -3.04401 | 0.002334473 | 0.034002458 |
| *DUSP2* | -2.86773 | 0.942118 | -3.043918 | 0.002335189 | 0.034002458 |
| *TBX6* | -2.518735 | 0.827559 | -3.043572 | 0.002337875 | 0.034002458 |
| *KIAA1324* | -3.243305 | 1.065662 | -3.043465 | 0.002338709 | 0.034002458 |
| *PANX1* | -2.698995 | 0.887107 | -3.042468 | 0.002346469 | 0.034018674 |
| *TSSK6* | -4.492856 | 1.47679 | -3.042313 | 0.002347676 | 0.034018674 |
| *HSD17B1* | -2.421345 | 0.796033 | -3.041765 | 0.002351953 | 0.034018674 |
| *ENSG00000273576* | -2.647559 | 0.870442 | -3.041627 | 0.002353029 | 0.034018674 |
| *ENSG00000275464* | -2.38512 | 0.784268 | -3.041207 | 0.002356322 | 0.034018674 |
| *RCAN1* | -4.080121 | 1.341622 | -3.041185 | 0.002356491 | 0.034018674 |
| *ENSG00000272909* | -3.95136 | 1.299384 | -3.04095 | 0.002358332 | 0.034018674 |

| | | | | | |
|---|---|---|---|---|---|
| *HESX1* | -3.345385 | 1.100552 | -3.039734 | 0.002367869 | 0.034106513 |
| *ENSG00000273619* | -3.397631 | 1.117826 | -3.039499 | 0.002369723 | 0.034106513 |
| *ENSG00000272444* | -4.357968 | 1.434115 | -3.038787 | 0.002375331 | 0.034127594 |
| *ENSG00000268129* | -3.087355 | 1.016156 | -3.038269 | 0.002379411 | 0.034127594 |
| *IGLV6.57* | -2.977575 | 0.980156 | -3.037857 | 0.002382671 | 0.034127594 |
| *TPST1* | -2.338146 | 0.769695 | -3.037757 | 0.002383461 | 0.034127594 |
| *ENSG00000267834* | -3.828061 | 1.260235 | -3.037577 | 0.002384881 | 0.034127594 |
| *WIPI1* | -2.089354 | 0.687933 | -3.037148 | 0.002388285 | 0.034127594 |
| *WFS1* | -3.05988 | 1.007546 | -3.036962 | 0.002389754 | 0.034127594 |
| *ENSG00000250132* | -2.944129 | 0.96955 | -3.036592 | 0.002392691 | 0.034131662 |
| *ENSG00000235119* | -2.488936 | 0.820046 | -3.035119 | 0.002404412 | 0.034137536 |
| *ENSG00000212939* | -2.135478 | 0.703629 | -3.03495 | 0.002405755 | 0.034137536 |
| *ENSG00000090932* | -3.295496 | 1.085884 | -3.034851 | 0.002406542 | 0.034137536 |
| *FYCO1* | -2.421996 | 0.798073 | -3.034806 | 0.002406907 | 0.034137536 |
| *DOCK9.DT* | -3.223791 | 1.062288 | -3.034763 | 0.002407247 | 0.034137536 |
| *ZHX1.C8orf76* | -2.285563 | 0.753332 | -3.033938 | 0.002413837 | 0.034137536 |
| *ATP6V1F* | -0.690871 | 0.227716 | -3.033912 | 0.002414048 | 0.034137536 |
| *KCNMB1* | -3.220598 | 1.061742 | -3.033315 | 0.002418827 | 0.034137536 |
| *ENSG00000275807* | -3.920791 | 1.292615 | -3.033224 | 0.002419559 | 0.034137536 |
| *ZNF575* | -2.870155 | 0.946242 | -3.033215 | 0.002419634 | 0.034137536 |
| *ZNF624* | -2.646696 | 0.872799 | -3.032423 | 0.002425993 | 0.034146257 |
| *CDCA8* | -3.032904 | 1.000303 | -3.031986 | 0.002429502 | 0.034146257 |
| *POPDC2* | -3.119738 | 1.028969 | -3.031907 | 0.002430142 | 0.034146257 |
| *TRIP6* | -1.847748 | 0.609452 | -3.031817 | 0.002430867 | 0.034146257 |
| *ZWINT* | -3.105726 | 1.024519 | -3.031399 | 0.00243423 | 0.034156208 |
| *FAM81A* | -3.161264 | 1.043332 | -3.029969 | 0.002445785 | 0.034213544 |
| *P2RX7* | -2.601473 | 0.858648 | -3.029732 | 0.002447708 | 0.034213544 |
| *VN1R1* | -2.55753 | 0.844215 | -3.029477 | 0.002449777 | 0.034213544 |
| *TMEM181* | -2.208907 | 0.729244 | -3.029039 | 0.002453333 | 0.034213544 |
| *ENSG00000223653* | -2.783425 | 0.919012 | -3.028714 | 0.002455973 | 0.034213544 |
| *ENSG00000272367* | -2.998019 | 0.989999 | -3.028304 | 0.002459306 | 0.034213544 |
| *SSX2IP* | -2.296127 | 0.758253 | -3.028183 | 0.002460293 | 0.034213544 |
| *ENSG00000256325* | -2.913154 | 0.962272 | -3.027369 | 0.002466925 | 0.034213544 |
| *CEACAM21* | -1.966064 | 0.649463 | -3.027214 | 0.002468189 | 0.034213544 |
| *ENSG00000229127* | -3.023658 | 0.998928 | -3.026902 | 0.002470738 | 0.034213544 |
| *ENSG00000257120* | -4.03744 | 1.333872 | -3.026857 | 0.002471111 | 0.034213544 |
| *SLC29A3* | -2.576617 | 0.851284 | -3.026741 | 0.002472059 | 0.034213544 |
| *PTOV1.AS1* | -4.126736 | 1.363494 | -3.026589 | 0.002473302 | 0.034213544 |
| *CHRNA6* | -4.060021 | 1.341573 | -3.026315 | 0.002475543 | 0.034213544 |
| *C3orf52* | -2.878639 | 0.951541 | -3.025238 | 0.002484373 | 0.034298736 |
| *ITPK1.AS1* | -3.323471 | 1.098832 | -3.024549 | 0.002490042 | 0.034321507 |
| *C15orf48* | -4.601991 | 1.521626 | -3.024389 | 0.002491357 | 0.034321507 |
| *MEIOC* | -3.823414 | 1.264516 | -3.023619 | 0.002497709 | 0.034372211 |
| *KIAA0825* | -3.638561 | 1.203607 | -3.023048 | 0.002502425 | 0.034400325 |
| *SSPN* | -3.179614 | 1.052162 | -3.021982 | 0.002511258 | 0.034482592 |
| *HEXA.AS1* | -3.823935 | 1.2655 | -3.021679 | 0.002513769 | 0.034482592 |
| *GAS8* | -2.976352 | 0.985468 | -3.020242 | 0.002525732 | 0.034609789 |

| | | | | | |
|---|---|---|---|---|---|
| *ENSG00000269910* | -4.028183 | 1.334207 | -3.019159 | 0.002534775 | 0.034685768 |
| *ENSG00000183308* | -3.872785 | 1.282832 | -3.018933 | 0.002536668 | 0.034685768 |
| *NDST1* | -4.07053 | 1.348577 | -3.01839 | 0.002541215 | 0.034686125 |
| *SLFN12L* | -3.800147 | 1.259041 | -3.018286 | 0.002542086 | 0.034686125 |
| *MYO15A* | -2.980818 | 0.987828 | -3.017548 | 0.002548284 | 0.034689643 |
| *LINC01483* | -4.678217 | 1.550416 | -3.017396 | 0.002549567 | 0.034689643 |
| *ENSG00000226644* | -3.037312 | 1.006826 | -3.016719 | 0.002555267 | 0.034689643 |
| *INTS2* | -2.10077 | 0.696446 | -3.016414 | 0.002557836 | 0.034689643 |
| *FANCE* | -2.925198 | 0.969888 | -3.016015 | 0.002561202 | 0.034689643 |
| *PFKFB3* | -1.188392 | 0.394039 | -3.015923 | 0.002561986 | 0.034689643 |
| *THRB* | -3.344608 | 1.10906 | -3.015715 | 0.002563737 | 0.034689643 |
| *PTMS* | -2.483856 | 0.823741 | -3.015337 | 0.002566941 | 0.034689643 |
| *CTRL* | -3.632205 | 1.204793 | -3.014797 | 0.002571514 | 0.034689643 |
| *IRAK3* | -2.179738 | 0.723041 | -3.014681 | 0.002572498 | 0.034689643 |
| *HSF5* | -3.318075 | 1.100664 | -3.014612 | 0.002573085 | 0.034689643 |
| *ENSG00000272991* | -3.287434 | 1.090569 | -3.014422 | 0.002574696 | 0.034689643 |
| *ENSG00000272892* | -3.312795 | 1.099272 | -3.013626 | 0.002581459 | 0.03472139 |
| *POC1A* | -3.013142 | 0.999878 | -3.01351 | 0.002582449 | 0.03472139 |
| *ENSG00000260806* | -2.843631 | 0.943972 | -3.01241 | 0.002591822 | 0.034721656 |
| *ENSG00000272156* | -2.830806 | 0.93978 | -3.0122 | 0.002593619 | 0.034721656 |
| *ENSG00000185065* | -3.49036 | 1.158848 | -3.011922 | 0.002595991 | 0.034721656 |
| *RASA4B* | -4.310749 | 1.43125 | -3.011876 | 0.002596383 | 0.034721656 |
| *C5orf22* | -2.196173 | 0.729172 | -3.011874 | 0.002596401 | 0.034721656 |
| *C1orf54* | -3.374439 | 1.120476 | -3.01161 | 0.00259866 | 0.034721656 |
| *AIF1* | -3.522893 | 1.169926 | -3.011209 | 0.002602093 | 0.03473147 |
| *FAN1* | -2.719036 | 0.903328 | -3.010021 | 0.002612296 | 0.03483152 |
| *FBXL6* | -2.945123 | 0.978654 | -3.009361 | 0.002617977 | 0.034871132 |
| *ENSG00000228192* | -4.342067 | 1.443429 | -3.008162 | 0.00262833 | 0.034919084 |
| *LINC01993* | -4.342067 | 1.443429 | -3.008162 | 0.00262833 | 0.034919084 |
| *RAPGEF5* | -2.847743 | 0.946819 | -3.007697 | 0.002632357 | 0.034919084 |
| *RNF227* | -2.525975 | 0.839881 | -3.007539 | 0.002633727 | 0.034919084 |
| *HOXB4* | -2.981695 | 0.991507 | -3.007235 | 0.002636356 | 0.034919084 |
| *TMEM177* | -2.201386 | 0.732072 | -3.007062 | 0.00263786 | 0.034919084 |
| *ENSG00000271427* | -4.237872 | 1.409577 | -3.006486 | 0.002642865 | 0.034919201 |
| *MAP3K21* | -3.289112 | 1.094027 | -3.006427 | 0.002643379 | 0.034919201 |
| *ENSG00000273080* | -2.755538 | 0.916641 | -3.006124 | 0.002646011 | 0.034919201 |
| *KCNMB4* | -3.166886 | 1.054032 | -3.004545 | 0.002659787 | 0.034974707 |
| *SIRT1* | -1.85747 | 0.618241 | -3.004443 | 0.002660677 | 0.034974707 |
| *ENSG00000267152* | -2.347241 | 0.781334 | -3.004144 | 0.002663293 | 0.034974707 |
| *IGLV1.40* | -3.283977 | 1.093174 | -3.004074 | 0.002663906 | 0.034974707 |
| *ENSG00000269051* | -3.036338 | 1.010768 | -3.003993 | 0.002664616 | 0.034974707 |
| *RAMP2.AS1* | -3.668311 | 1.221234 | -3.003775 | 0.002666526 | 0.034974707 |
| *ENSG00000253200* | -3.356187 | 1.117599 | -3.003032 | 0.00267304 | 0.034986772 |
| *IGLV3.1* | -2.885423 | 0.960853 | -3.00298 | 0.002673497 | 0.034986772 |
| *ENSG00000255337* | -3.050844 | 1.016234 | -3.002107 | 0.00268118 | 0.034986772 |
| *ENSG00000257410* | -3.975245 | 1.324179 | -3.002046 | 0.002681717 | 0.034986772 |
| *ENSG00000261335* | -3.909571 | 1.302626 | -3.001301 | 0.002688288 | 0.034986772 |

| | | | | | |
|---|---|---|---|---|---|
| *ENSG00000267751* | -3.04263 | 1.013795 | -3.001227 | 0.00268894 | 0.034986772 |
| *KCNJ14* | -3.605471 | 1.201354 | -3.001172 | 0.002689428 | 0.034986772 |
| *ATAD3C* | -3.800685 | 1.266579 | -3.000747 | 0.002693182 | 0.034986772 |
| *LINC01358* | -2.4793 | 0.826295 | -3.000503 | 0.00269534 | 0.034986772 |
| *C1QTNF12* | -3.938972 | 1.312919 | -3.000163 | 0.002698354 | 0.034986772 |
| *AURKA* | -2.850153 | 0.950185 | -2.999578 | 0.002703536 | 0.034986772 |
| *ZNF385D* | -3.483503 | 1.161359 | -2.999504 | 0.002704194 | 0.034986772 |
| *PPARGC1B* | -3.659979 | 1.220214 | -2.999456 | 0.002704624 | 0.034986772 |
| *NAV2.AS3* | -3.091292 | 1.030652 | -2.999356 | 0.002705513 | 0.034986772 |
| *ENSG00000250790* | -3.173941 | 1.058404 | -2.998799 | 0.002710457 | 0.034987603 |
| *RACGAP1* | -2.262577 | 0.75451 | -2.998737 | 0.002711016 | 0.034987603 |
| *ENSG00000272654* | -3.072691 | 1.024812 | -2.998298 | 0.002714924 | 0.035002938 |
| *CRB2* | -2.404916 | 0.802281 | -2.997598 | 0.002721161 | 0.03504292 |
| *ENSG00000267563* | -2.300195 | 0.767412 | -2.997339 | 0.002723472 | 0.03504292 |
| *ZNF449* | -2.68489 | 0.895982 | -2.996591 | 0.00273017 | 0.035094 |
| *DENND1A* | -2.300623 | 0.768028 | -2.995493 | 0.002740017 | 0.035153878 |
| *TMEM8B* | -3.15676 | 1.053847 | -2.995462 | 0.002740292 | 0.035153878 |
| *CD3D* | -4.955616 | 1.654626 | -2.995007 | 0.002744383 | 0.035171295 |
| *ENSG00000267787* | -2.646053 | 0.88382 | -2.993883 | 0.002754519 | 0.035222066 |
| *C4orf46* | -1.997228 | 0.667241 | -2.993263 | 0.002760121 | 0.035222066 |
| *PROC* | -3.036805 | 1.014743 | -2.992683 | 0.002765367 | 0.035222066 |
| *ENSG00000266677* | -3.322771 | 1.110348 | -2.99255 | 0.002766578 | 0.035222066 |
| *ENSG00000266844* | -3.987123 | 1.332356 | -2.992537 | 0.002766694 | 0.035222066 |
| *ENSG00000277368* | -3.128045 | 1.045361 | -2.99231 | 0.002768747 | 0.035222066 |
| *SIRT4* | -3.506181 | 1.171734 | -2.992301 | 0.002768828 | 0.035222066 |
| *SDC3* | -3.144964 | 1.051073 | -2.992145 | 0.002770244 | 0.035222066 |
| *CAHM* | -3.054714 | 1.021038 | -2.991772 | 0.002773629 | 0.035230294 |
| *ARMCX4* | -2.478977 | 0.828753 | -2.991212 | 0.002778728 | 0.035231687 |
| *FILIP1* | -2.911379 | 0.973418 | -2.990882 | 0.002781726 | 0.035231687 |
| *ENSG00000261845* | -2.964102 | 0.991054 | -2.990857 | 0.002781953 | 0.035231687 |
| *RHOBTB1* | -3.218499 | 1.076391 | -2.990083 | 0.002789012 | 0.035286355 |
| *DLG3* | -3.31639 | 1.109345 | -2.989503 | 0.002794317 | 0.035318735 |
| *LY6G5B* | -3.801077 | 1.271935 | -2.988421 | 0.002804226 | 0.035409202 |
| *ZNF229* | -2.695862 | 0.902282 | -2.987825 | 0.002809706 | 0.035443618 |
| *DNHD1* | -2.202949 | 0.737651 | -2.986436 | 0.002822502 | 0.035570165 |
| *WDR11.AS1* | -3.279239 | 1.098311 | -2.98571 | 0.002829207 | 0.035619768 |
| *ENSG00000232063* | -3.259212 | 1.092182 | -2.984129 | 0.002843866 | 0.035769329 |
| *TLCD2* | -2.803588 | 0.939887 | -2.982898 | 0.002855331 | 0.035834737 |
| *PLEKHG2* | -2.514286 | 0.843159 | -2.981984 | 0.002863871 | 0.035834737 |
| *CTBP1.AS* | -2.77808 | 0.931697 | -2.981741 | 0.002866143 | 0.035834737 |
| *CRYM.AS1* | -3.786915 | 1.270296 | -2.981129 | 0.00287188 | 0.035834737 |
| *LCTL* | -3.071212 | 1.030227 | -2.981102 | 0.002872128 | 0.035834737 |
| *TNRC18* | -2.222693 | 0.745611 | -2.981036 | 0.00287275 | 0.035834737 |
| *ENSG00000253891* | -2.792056 | 0.936665 | -2.980849 | 0.002874502 | 0.035834737 |
| *GK* | -2.084975 | 0.699463 | -2.980823 | 0.002874752 | 0.035834737 |
| *ENSG00000272853* | -2.799372 | 0.939249 | -2.980436 | 0.002878384 | 0.035834737 |
| *ARHGAP11A* | -2.663911 | 0.893834 | -2.980318 | 0.002879489 | 0.035834737 |

| | | | | | |
|---|---|---|---|---|---|
| *ENSG00000253106* | -3.334555 | 1.118948 | -2.98008 | 0.002881728 | 0.035834737 |
| *IGHV4.39* | -3.484764 | 1.169497 | -2.979712 | 0.002885194 | 0.035834737 |
| *SRC* | -2.964304 | 0.994832 | -2.979704 | 0.002885271 | 0.035834737 |
| *RILPL1* | -3.669064 | 1.231522 | -2.979293 | 0.002889143 | 0.035848223 |
| *HIC2* | -2.763153 | 0.927701 | -2.978494 | 0.002896683 | 0.035858308 |
| *EIF4E3* | -2.591234 | 0.869984 | -2.978483 | 0.002896788 | 0.035858308 |
| *MCOLN2* | -3.219587 | 1.081105 | -2.978051 | 0.002900875 | 0.035858308 |
| *C11orf21* | -2.931049 | 0.984242 | -2.977975 | 0.002901594 | 0.035858308 |
| *MTFR2* | -3.197322 | 1.073744 | -2.977733 | 0.00290389 | 0.035858308 |
| *EGLN3* | -3.892164 | 1.307252 | -2.977362 | 0.002907402 | 0.03586725 |
| *PRICKLE3* | -2.461964 | 0.827107 | -2.976597 | 0.002914673 | 0.03592251 |
| *C8orf89* | -3.840097 | 1.290374 | -2.975957 | 0.002920755 | 0.035949917 |
| *DPCD* | -2.202294 | 0.740074 | -2.975776 | 0.002922485 | 0.035949917 |
| *GTF2H4* | -3.452042 | 1.160189 | -2.975413 | 0.002925941 | 0.035958049 |
| *BCL2L2* | -2.312492 | 0.777281 | -2.975103 | 0.002928901 | 0.035960087 |
| *ENSG00000260526* | -3.077799 | 1.034838 | -2.974185 | 0.002937677 | 0.036021409 |
| *POLR3B* | -2.94225 | 0.989326 | -2.973995 | 0.002939495 | 0.036021409 |
| *MAMLD1* | -5.352021 | 1.800298 | -2.972853 | 0.002950457 | 0.036097567 |
| *ENSG00000271122* | -2.412908 | 0.811672 | -2.972763 | 0.00295132 | 0.036097567 |
| *ZNF75D* | -2.195433 | 0.738765 | -2.971759 | 0.002960992 | 0.036174168 |
| *EPHB6* | -1.555778 | 0.523561 | -2.971529 | 0.002963206 | 0.036174168 |
| *CPS1* | -4.485172 | 1.510104 | -2.970107 | 0.002976956 | 0.036254856 |
| *SLC26A1* | -3.508082 | 1.181156 | -2.970042 | 0.002977591 | 0.036254856 |
| *ZNF484* | -2.386594 | 0.803574 | -2.969972 | 0.002978269 | 0.036254856 |
| *LEKR1* | -2.782457 | 0.937049 | -2.969381 | 0.002984002 | 0.03629032 |
| *MYH3* | -2.339352 | 0.788042 | -2.968561 | 0.002991974 | 0.036299993 |
| *RLN2* | -2.682312 | 0.903648 | -2.968316 | 0.00299436 | 0.036299993 |
| *SPAG8* | -2.841574 | 0.957411 | -2.967976 | 0.002997681 | 0.036299993 |
| *LETM2* | -3.737732 | 1.25939 | -2.967892 | 0.002998498 | 0.036299993 |
| *MPPED2* | -3.139562 | 1.057857 | -2.96785 | 0.002998904 | 0.036299993 |
| *ENSG00000272588* | -3.05771 | 1.030503 | -2.9672 | 0.003005249 | 0.036342615 |
| *SIAE* | -2.68965 | 0.906655 | -2.966564 | 0.003011477 | 0.036352275 |
| *TRIM72* | -3.838548 | 1.293947 | -2.966541 | 0.003011699 | 0.036352275 |
| *DCBLD1* | -2.759904 | 0.930996 | -2.964465 | 0.003032096 | 0.036564177 |
| *GLCE* | -2.339167 | 0.789344 | -2.963433 | 0.003042286 | 0.036652714 |
| *TK2* | -2.649941 | 0.894365 | -2.96293 | 0.003047263 | 0.036670758 |
| *CSTF3.DT* | -3.585728 | 1.210288 | -2.962705 | 0.003049484 | 0.036670758 |
| *TIGD2* | -2.35873 | 0.796326 | -2.962018 | 0.003056303 | 0.036718447 |
| *ACVR2A* | -3.237249 | 1.093378 | -2.960777 | 0.003068643 | 0.036832309 |
| *MTCL1* | -3.714461 | 1.2551 | -2.959495 | 0.003081439 | 0.036951417 |
| *MYBL1* | -2.999162 | 1.013578 | -2.958984 | 0.003086546 | 0.036978206 |
| *FSD1L* | -1.925351 | 0.650817 | -2.958358 | 0.003092822 | 0.036997835 |
| *ENSG00000105650* | -3.279828 | 1.10879 | -2.958026 | 0.003096164 | 0.036997835 |
| *YPEL4* | -3.508084 | 1.18598 | -2.957961 | 0.003096811 | 0.036997835 |
| *ENSG00000269604* | -3.476796 | 1.175556 | -2.957576 | 0.00310068 | 0.037009699 |
| *ENSG00000225302* | -3.915325 | 1.324789 | -2.955432 | 0.003122319 | 0.03714677 |
| *LINC01176* | -3.123082 | 1.056837 | -2.955122 | 0.003125459 | 0.03714677 |

| | | | | | |
|---|---|---|---|---|---|
| *TTLL4* | -2.211565 | 0.748384 | -2.95512 | 0.003125471 | 0.03714677 |
| *PGAP1* | -2.972258 | 1.005817 | -2.955068 | 0.003125996 | 0.03714677 |
| *PLAUR* | -3.001448 | 1.015715 | -2.955009 | 0.003126599 | 0.03714677 |
| *ENSG00000282034* | -3.908283 | 1.323032 | -2.954034 | 0.003136494 | 0.037211659 |
| *TESMIN* | -2.707492 | 0.916582 | -2.953901 | 0.003137845 | 0.037211659 |
| *IFT140* | -3.467099 | 1.17394 | -2.953388 | 0.003143067 | 0.03723927 |
| *ARG2* | -3.764207 | 1.274737 | -2.952929 | 0.003147743 | 0.037260362 |
| *LINC01725* | -4.61642 | 1.564084 | -2.951516 | 0.003162181 | 0.037324174 |
| *FRG1.DT* | -4.633825 | 1.570056 | -2.951375 | 0.003163628 | 0.037324174 |
| *ZNF182* | -2.125676 | 0.720287 | -2.951152 | 0.003165911 | 0.037324174 |
| *CACNB3* | -2.752722 | 0.932763 | -2.951149 | 0.003165943 | 0.037324174 |
| *FCMR* | 1.142231 | 0.387081 | 2.950887 | 0.003168628 | 0.037324174 |
| *ENSG00000267416* | -2.92336 | 0.990734 | -2.950701 | 0.003170539 | 0.037324174 |
| *CDKN3* | -3.580399 | 1.213684 | -2.950025 | 0.003177487 | 0.037360445 |
| *ENSG00000273674* | -2.995359 | 1.015433 | -2.949836 | 0.003179427 | 0.037360445 |
| *ENSG00000251867* | -3.303905 | 1.120432 | -2.948776 | 0.003190348 | 0.037454569 |
| *DMPK* | -3.2253 | 1.094028 | -2.948096 | 0.003197374 | 0.037472806 |
| *L1CAM* | -2.343288 | 0.794857 | -2.948062 | 0.003197726 | 0.037472806 |
| *CYP4V2* | -2.205987 | 0.748529 | -2.947095 | 0.003207747 | 0.037536789 |
| *LRRC37A* | -3.688137 | 1.251634 | -2.946656 | 0.003212299 | 0.037536789 |
| *ENSG00000245213* | -3.734406 | 1.267525 | -2.946218 | 0.003216858 | 0.037536789 |
| *ENSG00000276564* | -3.338727 | 1.133229 | -2.946206 | 0.003216982 | 0.037536789 |
| *TBXAS1* | -2.681434 | 0.910155 | -2.94613 | 0.003217772 | 0.037536789 |
| *ZNF132* | -3.560202 | 1.208597 | -2.945732 | 0.003221914 | 0.037551053 |
| *AVEN* | -3.126669 | 1.061637 | -2.94514 | 0.003228083 | 0.037588907 |
| *PTPN20* | -2.946891 | 1.000716 | -2.944782 | 0.00323182 | 0.037598398 |
| *CAB39L* | -2.32987 | 0.791275 | -2.944449 | 0.003235303 | 0.037604923 |
| *DNTTIP1* | -1.405781 | 0.477512 | -2.943969 | 0.00324033 | 0.037629353 |
| *C1orf74* | -2.479207 | 0.842222 | -2.94365 | 0.003243662 | 0.037634079 |
| *AFF2* | -3.344044 | 1.13641 | -2.942638 | 0.003254287 | 0.037704614 |
| *ENSG00000277022* | -2.691259 | 0.914628 | -2.942464 | 0.003256121 | 0.037704614 |
| *CHIC1* | -2.191394 | 0.744889 | -2.941907 | 0.003261974 | 0.037704614 |
| *PRDM15* | -2.394582 | 0.814007 | -2.941723 | 0.003263917 | 0.037704614 |
| *TRABD* | 1.073856 | 0.365049 | 2.941678 | 0.003264393 | 0.037704614 |
| *ENSG00000273002* | -2.45326 | 0.834393 | -2.940172 | 0.003280299 | 0.037809039 |
| *ECT2* | -3.773141 | 1.283485 | -2.939762 | 0.003284646 | 0.037809039 |
| *ENSG00000224738* | -3.638941 | 1.237841 | -2.939748 | 0.003284789 | 0.037809039 |
| *RNF212* | -2.388044 | 0.81234 | -2.939711 | 0.003285187 | 0.037809039 |
| *RUSC1.AS1* | -3.457847 | 1.176474 | -2.939161 | 0.003291019 | 0.037842306 |
| *CLEC18A* | -3.714838 | 1.264163 | -2.938576 | 0.003297242 | 0.037876742 |
| *TNFRSF21* | -4.451572 | 1.515138 | -2.938064 | 0.003302686 | 0.037876742 |
| *MAPRE3* | -2.310943 | 0.786557 | -2.938049 | 0.003302845 | 0.037876742 |
| *EPHA4* | -4.864721 | 1.656583 | -2.936599 | 0.003318329 | 0.037984286 |
| *CDC7* | -2.740319 | 0.933174 | -2.936557 | 0.00331878 | 0.037984286 |
| *BNIPL* | -3.525742 | 1.200726 | -2.936342 | 0.003321079 | 0.037984286 |
| *GALC* | -2.366264 | 0.806064 | -2.93558 | 0.003329247 | 0.038043894 |
| *STXBP5.AS1* | -2.799702 | 0.953871 | -2.935095 | 0.003334459 | 0.038069635 |

| | | | | | |
|---|---|---|---|---|---|
| *RNF130* | -1.347366 | 0.459116 | -2.934693 | 0.003338774 | 0.038085109 |
| *CLCN4* | -2.931707 | 0.999336 | -2.933656 | 0.003349951 | 0.038178761 |
| *GSDMB* | -2.300381 | 0.784286 | -2.933091 | 0.003356058 | 0.038214512 |
| *SLC27A4* | -2.31103 | 0.788044 | -2.932617 | 0.003361187 | 0.038236497 |
| *SCN4A* | -3.729693 | 1.271912 | -2.932351 | 0.003364059 | 0.038236497 |
| *RBMS2* | -2.961725 | 1.010107 | -2.932089 | 0.003366904 | 0.038236497 |
| *ELF1* | -0.864411 | 0.295015 | -2.930064 | 0.003388921 | 0.038449683 |
| *CCDC171* | -2.750175 | 0.938686 | -2.929814 | 0.003391652 | 0.038449683 |
| *CSRNP2* | -2.350635 | 0.802622 | -2.928693 | 0.003403901 | 0.038494192 |
| *ENSG00000268093* | -3.616131 | 1.234794 | -2.928529 | 0.003405705 | 0.038494192 |
| *ALDH7A1* | -3.12629 | 1.067584 | -2.928379 | 0.00340734 | 0.038494192 |
| *RAD54B* | -4.089859 | 1.396638 | -2.928361 | 0.003407545 | 0.038494192 |
| *ENSG00000205682* | -2.777672 | 0.948709 | -2.927845 | 0.003413207 | 0.038524328 |
| *PIK3CD.AS2* | -2.418133 | 0.826379 | -2.926178 | 0.00343154 | 0.038612467 |
| *IGKV4.1* | -2.434902 | 0.832122 | -2.926136 | 0.003432009 | 0.038612467 |
| *SLC7A5* | -2.698473 | 0.92226 | -2.925936 | 0.003434213 | 0.038612467 |
| *PACRGL* | -2.419573 | 0.82697 | -2.925829 | 0.003435397 | 0.038612467 |
| *NETO2* | -2.696745 | 0.921834 | -2.925413 | 0.003439989 | 0.038612467 |
| *ADAM15* | -2.267629 | 0.775182 | -2.925286 | 0.0034414 | 0.038612467 |
| *PRKCI* | -2.096032 | 0.716578 | -2.925058 | 0.00344392 | 0.038612467 |
| *ENSG00000251143* | -3.053383 | 1.043907 | -2.924958 | 0.003445023 | 0.038612467 |
| *LINC02447* | -3.206431 | 1.096409 | -2.924485 | 0.003450269 | 0.038637603 |
| *NEXN* | -2.261227 | 0.773504 | -2.923353 | 0.003462833 | 0.038743691 |
| *CRACR2A* | -2.398496 | 0.820557 | -2.923012 | 0.003466637 | 0.038743691 |
| *SENP8* | -3.409159 | 1.166394 | -2.92282 | 0.003468775 | 0.038743691 |
| *TMEM182* | -3.111455 | 1.064826 | -2.922032 | 0.003477555 | 0.03878997 |
| *LINC01134* | -3.126584 | 1.070051 | -2.921901 | 0.003479023 | 0.03878997 |
| *HASPIN* | -3.438048 | 1.176915 | -2.921238 | 0.003486435 | 0.03878997 |
| *RGS17* | -3.5555 | 1.21714 | -2.921192 | 0.003486951 | 0.03878997 |
| *GRTP1* | -3.776923 | 1.29298 | -2.921099 | 0.003487992 | 0.03878997 |
| *ZRANB3* | -1.937428 | 0.663446 | -2.920248 | 0.003497529 | 0.038842482 |
| *ENSG00000270696* | -3.636587 | 1.245412 | -2.919988 | 0.003500452 | 0.038842482 |
| *POT1.AS1* | -3.056766 | 1.046889 | -2.919856 | 0.003501933 | 0.038842482 |
| *ASF1B* | -3.707177 | 1.269766 | -2.919576 | 0.003505083 | 0.038842482 |
| *ENSG00000273702* | -2.092369 | 0.716728 | -2.919333 | 0.003507808 | 0.038842482 |
| *ENSG00000234773* | -3.811772 | 1.305836 | -2.919028 | 0.003511244 | 0.038845766 |
| *ENSG00000272821* | -3.078969 | 1.054933 | -2.91864 | 0.003515623 | 0.038845766 |
| *ZNF443* | -2.650195 | 0.908118 | -2.918337 | 0.00351904 | 0.038845766 |
| *ENSG00000224220* | -2.922921 | 1.001606 | -2.918236 | 0.003520181 | 0.038845766 |
| *ENSG00000231105* | -3.718442 | 1.274735 | -2.917031 | 0.003533803 | 0.038933021 |
| *ENSG00000264207* | -2.956751 | 1.013645 | -2.916949 | 0.003534736 | 0.038933021 |
| *ENSG00000262621* | -2.642677 | 0.906108 | -2.916514 | 0.003539665 | 0.038933021 |
| *LSM14B* | -2.261436 | 0.775546 | -2.915928 | 0.003546324 | 0.038933021 |
| *CECR7* | -3.675091 | 1.260439 | -2.915724 | 0.003548642 | 0.038933021 |
| *ENSG00000250685* | -3.716372 | 1.274633 | -2.91564 | 0.003549603 | 0.038933021 |
| *ENSG00000231760* | -3.617099 | 1.240627 | -2.915542 | 0.003550715 | 0.038933021 |
| *LINC00106* | -3.443407 | 1.181108 | -2.915403 | 0.003552294 | 0.038933021 |

| | | | | | |
|---|---|---|---|---|---|
| *SAG* | -4.721453 | 1.619804 | -2.91483 | 0.003558822 | 0.038971369 |
| *TRIM61* | -2.269379 | 0.778722 | -2.914237 | 0.003565593 | 0.038997506 |
| *TAT.AS1* | -2.89737 | 0.994262 | -2.91409 | 0.00356727 | 0.038997506 |
| *DNAJC9.AS1* | -4.507503 | 1.546981 | -2.913741 | 0.003571258 | 0.039007959 |
| *PINLYP* | -3.089244 | 1.06059 | -2.91276 | 0.003582496 | 0.039097522 |
| *ARHGAP33* | -3.618574 | 1.242922 | -2.911343 | 0.003598785 | 0.039213622 |
| *ALDH1B1* | -2.486865 | 0.854287 | -2.911041 | 0.003602264 | 0.039213622 |
| *ST6GALNAC3* | -2.561265 | 0.879845 | -2.91104 | 0.003602277 | 0.039213622 |
| *SUV39H2* | -2.945155 | 1.01196 | -2.910346 | 0.003610286 | 0.039267582 |
| *PRKCA* | -3.595608 | 1.236048 | -2.908955 | 0.003626395 | 0.039409478 |
| *ENSG00000262580* | -2.437776 | 0.838259 | -2.908142 | 0.003635834 | 0.039468603 |
| *GNG3* | -3.599 | 1.237764 | -2.907662 | 0.003641422 | 0.039468603 |
| *LENG9* | -4.153177 | 1.428685 | -2.906992 | 0.003649227 | 0.039468603 |
| *BANK1* | -0.806032 | 0.277281 | -2.90691 | 0.003650181 | 0.039468603 |
| *ENSG00000235381* | -3.288509 | 1.131354 | -2.906703 | 0.003652598 | 0.039468603 |
| *SH2D4A* | -2.732318 | 0.940007 | -2.906699 | 0.003652649 | 0.039468603 |
| *LINC01504* | -3.639828 | 1.252245 | -2.906642 | 0.003653307 | 0.039468603 |
| *TNFRSF25* | -4.651552 | 1.600639 | -2.90606 | 0.00366011 | 0.03950892 |
| *ENSG00000272172* | -2.52508 | 0.869032 | -2.905623 | 0.003665227 | 0.039530995 |
| *ENSG00000166928* | -4.281521 | 1.474154 | -2.904392 | 0.003679673 | 0.039613671 |
| *MATN1.AS1* | -2.338528 | 0.805187 | -2.904327 | 0.003680432 | 0.039613671 |
| *FMO4* | -2.9984 | 1.032526 | -2.903945 | 0.003684925 | 0.039613671 |
| *ARHGAP10* | -4.137533 | 1.424809 | -2.903921 | 0.003685207 | 0.039613671 |
| *XKR6* | -2.37175 | 0.816992 | -2.903028 | 0.003695734 | 0.039627561 |
| *ENSG00000232098* | -3.773252 | 1.299933 | -2.902652 | 0.003700173 | 0.039627561 |
| *CRY2* | -1.915512 | 0.659924 | -2.902625 | 0.00370049 | 0.039627561 |
| *PRC1* | -3.417957 | 1.177627 | -2.90241 | 0.003703032 | 0.039627561 |
| *CENPS.CORT* | -3.884914 | 1.338519 | -2.902397 | 0.00370319 | 0.039627561 |
| *EXT1* | -4.560734 | 1.571546 | -2.902068 | 0.003707076 | 0.039627561 |
| *ENSG00000235501* | -3.223729 | 1.11087 | -2.901985 | 0.003708058 | 0.039627561 |
| *PANX2* | -2.700354 | 0.93096 | -2.900612 | 0.003724351 | 0.039768653 |
| *ENSG00000268744* | -2.855842 | 0.984877 | -2.899695 | 0.003735265 | 0.039821029 |
| *ENSG00000233757* | -3.761668 | 1.297589 | -2.898968 | 0.003743929 | 0.039821029 |
| *HSD17B14* | -2.644594 | 0.912264 | -2.898934 | 0.00374434 | 0.039821029 |
| *ACY1* | -3.28984 | 1.135033 | -2.898453 | 0.003750081 | 0.039821029 |
| *ENSG00000254428* | -3.260545 | 1.124933 | -2.898434 | 0.003750317 | 0.039821029 |
| *CDK5R1* | -4.721173 | 1.628913 | -2.898357 | 0.003751228 | 0.039821029 |
| *ARF5* | -0.832199 | 0.287143 | -2.8982 | 0.003753112 | 0.039821029 |
| *ENSG00000165521* | -4.284618 | 1.478446 | -2.898054 | 0.003754859 | 0.039821029 |
| *CDK10* | -1.505317 | 0.519529 | -2.897465 | 0.00376192 | 0.039821029 |
| *ENSG00000214797* | -2.91663 | 1.006629 | -2.897423 | 0.003762421 | 0.039821029 |
| *FKBP7* | -2.840227 | 0.980292 | -2.897328 | 0.003763559 | 0.039821029 |
| *ATPSCKMT* | -2.157937 | 0.744926 | -2.896849 | 0.003769306 | 0.039821029 |
| *SGCA* | -3.494988 | 1.206537 | -2.896709 | 0.003770988 | 0.039821029 |
| *PRDX1* | -0.754443 | 0.26046 | -2.896577 | 0.003772584 | 0.039821029 |
| *CCDC180* | -2.315914 | 0.799672 | -2.896081 | 0.003778548 | 0.039838858 |
| *RPL34.AS1* | -2.941744 | 1.015823 | -2.895922 | 0.003780465 | 0.039838858 |

| | | | | | |
|---|---|---|---|---|---|
| *ADGRE2* | -2.665403 | 0.92056 | -2.895413 | 0.003786594 | 0.039870789 |
| *HECTD4* | -1.608863 | 0.555733 | -2.895031 | 0.003791207 | 0.039886725 |
| *ZNF396* | -3.518565 | 1.215623 | -2.894454 | 0.00379819 | 0.039900767 |
| *ENSG00000204758* | -2.797126 | 0.96639 | -2.894408 | 0.003798744 | 0.039900767 |
| *CCBE1* | -3.508491 | 1.212422 | -2.893787 | 0.003806266 | 0.039928999 |
| *DNA2* | -3.06016 | 1.057534 | -2.893674 | 0.003807638 | 0.039928999 |
| *UBE2D3.AS1* | -2.802716 | 0.968893 | -2.8927 | 0.003819456 | 0.040020308 |
| *ENSG00000259408* | -3.161634 | 1.093172 | -2.892165 | 0.003825975 | 0.04002923 |
| *LINC02019* | -3.534152 | 1.221994 | -2.892119 | 0.003826529 | 0.04002923 |
| *ENSG00000262777* | -4.476827 | 1.548254 | -2.891533 | 0.003833673 | 0.040071377 |
| *JAKMIP2* | -2.62689 | 0.908663 | -2.89094 | 0.003840909 | 0.040083304 |
| *PPIL6* | -2.761713 | 0.955303 | -2.890929 | 0.003841044 | 0.040083304 |
| *TBCEL* | -2.347843 | 0.812265 | -2.890491 | 0.003846406 | 0.040106734 |
| *USP6* | -3.155952 | 1.092193 | -2.889556 | 0.003857867 | 0.040193666 |
| *ENSG00000258768* | -3.30385 | 1.143669 | -2.888817 | 0.003866943 | 0.040255626 |
| *PRSS27* | -2.580054 | 0.893297 | -2.888238 | 0.00387407 | 0.040297221 |
| *DSP* | -2.816755 | 0.97549 | -2.88753 | 0.003882802 | 0.040347315 |
| *ENSG00000276718* | -3.674272 | 1.272549 | -2.887332 | 0.003885239 | 0.040347315 |
| *LYG1* | -2.864917 | 0.99236 | -2.886975 | 0.003889657 | 0.040347315 |
| *DUSP7* | -2.920973 | 1.011845 | -2.88678 | 0.003892068 | 0.040347315 |
| *CTBP2* | -4.541451 | 1.573372 | -2.886444 | 0.003896216 | 0.040347315 |
| *MAGEE1* | -2.823726 | 0.978312 | -2.886325 | 0.003897701 | 0.040347315 |
| *JMY* | -2.204084 | 0.763863 | -2.885444 | 0.003908624 | 0.040414964 |
| *TDRKH.AS1* | -2.298328 | 0.796567 | -2.885291 | 0.003910518 | 0.040414964 |
| *CORO2B* | -3.486454 | 1.208811 | -2.884202 | 0.003924075 | 0.040470086 |
| *ARL4D* | -2.546189 | 0.882836 | -2.884103 | 0.003925305 | 0.040470086 |
| *LINC01133* | -4.407111 | 1.528254 | -2.883755 | 0.003929648 | 0.040470086 |
| *ENSG00000269886* | -4.650243 | 1.612613 | -2.883669 | 0.003930713 | 0.040470086 |
| *HIST2H4A* | -3.221615 | 1.117307 | -2.883373 | 0.003934407 | 0.040470086 |
| *ENSG00000275202* | -4.350045 | 1.508678 | -2.883348 | 0.003934723 | 0.040470086 |
| *CLDN7* | -3.154621 | 1.094633 | -2.881899 | 0.003952862 | 0.040592754 |
| *ENSG00000272927* | -2.066245 | 0.716995 | -2.881814 | 0.003953933 | 0.040592754 |
| *GOLGA6L4* | -2.657689 | 0.922283 | -2.88164 | 0.003956114 | 0.040592754 |
| *PYHIN1* | -3.666607 | 1.272805 | -2.88073 | 0.003967552 | 0.040677686 |
| *GSTO2* | -3.708607 | 1.28782 | -2.879754 | 0.00397985 | 0.040735497 |
| *ENSG00000233427* | -4.35326 | 1.511735 | -2.879645 | 0.003981234 | 0.040735497 |
| *ADAT3* | -2.696771 | 0.93658 | -2.879381 | 0.003984566 | 0.040735497 |
| *ENSG00000273270* | -1.894851 | 0.658099 | -2.879279 | 0.003985855 | 0.040735497 |
| *ENSG00000111321* | -2.447842 | 0.850344 | -2.878651 | 0.003993801 | 0.04078431 |
| *EPN2* | -2.107221 | 0.732118 | -2.878253 | 0.003998844 | 0.040803433 |
| *PON2* | -2.659489 | 0.924291 | -2.877327 | 0.004010592 | 0.04089088 |
| *AP5S1* | -1.814807 | 0.630825 | -2.876881 | 0.004016276 | 0.040916411 |
| *CSNK2A3* | -3.111957 | 1.08186 | -2.876488 | 0.004021269 | 0.040934865 |
| *KCNRG* | -3.612369 | 1.256274 | -2.875463 | 0.004034358 | 0.040997929 |
| *NFS1* | -1.980766 | 0.688884 | -2.875326 | 0.0040361 | 0.040997929 |
| *ENSG00000269194* | -2.640903 | 0.918548 | -2.875083 | 0.004039214 | 0.040997929 |
| *ZNF34* | -1.90917 | 0.664058 | -2.875005 | 0.004040209 | 0.040997929 |

| | | | | | |
|---|---|---|---|---|---|
| *ACACA* | -2.276897 | 0.792066 | -2.874631 | 0.004044994 | 0.041014139 |
| *ENSG00000274425* | -2.480902 | 0.86328 | -2.87381 | 0.004055528 | 0.041071962 |
| *ZNF354B* | -1.777216 | 0.618605 | -2.872943 | 0.004066676 | 0.041071962 |
| *ATP6V0C* | -4.122843 | 1.435072 | -2.872916 | 0.004067019 | 0.041071962 |
| *RPRML* | -4.953394 | 1.724202 | -2.872862 | 0.004067714 | 0.041071962 |
| *MADCAM1* | -2.364242 | 0.823098 | -2.87237 | 0.004074052 | 0.041071962 |
| *HSPA1L* | -4.72169 | 1.643877 | -2.872289 | 0.004075095 | 0.041071962 |
| *KCNA7* | -4.72169 | 1.643877 | -2.872289 | 0.004075095 | 0.041071962 |
| *ZFPM1* | -3.235423 | 1.126461 | -2.872201 | 0.004076233 | 0.041071962 |
| *GEM* | -2.697681 | 0.939511 | -2.871366 | 0.004087016 | 0.041140708 |
| *ENSG00000234141* | -3.723098 | 1.296715 | -2.871178 | 0.004089451 | 0.041140708 |
| *ENSG00000247121* | -2.261277 | 0.78769 | -2.870771 | 0.004094721 | 0.041152864 |
| *LYSMD4* | -2.286928 | 0.796677 | -2.870583 | 0.004097154 | 0.041152864 |
| *FGL2* | -3.076765 | 1.072084 | -2.869893 | 0.004106108 | 0.041152864 |
| *LINC00174* | -4.405093 | 1.534944 | -2.869873 | 0.004106367 | 0.041152864 |
| *TSBP1.AS1* | -3.887483 | 1.354659 | -2.869713 | 0.004108446 | 0.041152864 |
| *HYAL2* | -2.26032 | 0.787729 | -2.869413 | 0.004112349 | 0.041152864 |
| *EFNA3* | -2.752118 | 0.959178 | -2.869248 | 0.004114491 | 0.041152864 |
| *ENSG00000257605* | -4.461823 | 1.555123 | -2.869113 | 0.004116246 | 0.041152864 |
| *XRCC2* | -3.258981 | 1.136132 | -2.868488 | 0.004124385 | 0.041173865 |
| *OCRL* | -3.802276 | 1.325546 | -2.868461 | 0.004124746 | 0.041173865 |
| *SPIN2A* | -3.005655 | 1.048053 | -2.867845 | 0.004132774 | 0.041202803 |
| *ENSG00000272663* | -2.702418 | 0.942349 | -2.867748 | 0.00413405 | 0.041202803 |
| *ABCA1* | -4.613766 | 1.609347 | -2.866856 | 0.004145713 | 0.041287063 |
| *CNNM4* | -2.986292 | 1.042046 | -2.865796 | 0.00415962 | 0.041330373 |
| *DNAJC17* | -1.661677 | 0.579841 | -2.865745 | 0.00416029 | 0.041330373 |
| *TBC1D23* | -1.662882 | 0.580295 | -2.865581 | 0.004162447 | 0.041330373 |
| *CCDC134* | -2.714871 | 0.947418 | -2.865546 | 0.00416291 | 0.041330373 |
| *LAPTM5* | -0.509865 | 0.177947 | -2.865272 | 0.004166512 | 0.041334242 |
| *SHF* | -2.30998 | 0.806284 | -2.86497 | 0.004170484 | 0.041341773 |
| *C18orf54* | -2.428509 | 0.847928 | -2.864053 | 0.004182578 | 0.041429741 |
| *HCG27* | -2.576921 | 0.900043 | -2.863108 | 0.004195071 | 0.041501129 |
| *ENSG00000263080* | -3.573321 | 1.248106 | -2.862995 | 0.004196568 | 0.041501129 |
| *ARHGEF12* | -2.023397 | 0.706848 | -2.862564 | 0.004202287 | 0.041501129 |
| *FAM168A* | -2.176306 | 0.760273 | -2.862534 | 0.004202687 | 0.041501129 |
| *ENSG00000228265* | -2.651148 | 0.926441 | -2.861648 | 0.004214446 | 0.041557109 |
| *ZNF578* | -2.6187 | 0.915167 | -2.861444 | 0.004217161 | 0.041557109 |
| *LRFN4* | -3.097792 | 1.082623 | -2.861377 | 0.004218045 | 0.041557109 |
| *ZGLP1* | -3.295516 | 1.152586 | -2.859236 | 0.004246623 | 0.041806659 |
| *AGAP9* | -2.535993 | 0.887052 | -2.8589 | 0.004251133 | 0.041819057 |
| *ENSG00000273391* | -3.833492 | 1.341076 | -2.858519 | 0.004256234 | 0.041837252 |
| *ENSG00000224746* | -2.709077 | 0.947983 | -2.857728 | 0.004266858 | 0.041909662 |
| *TAS2R14* | -3.666741 | 1.283432 | -2.856981 | 0.004276918 | 0.041957883 |
| *RFC2* | -1.391225 | 0.486974 | -2.856879 | 0.004278289 | 0.041957883 |
| *LINC00630* | -4.032122 | 1.411684 | -2.85625 | 0.00428678 | 0.041997032 |
| *ENSG00000255468* | -3.454713 | 1.209612 | -2.856051 | 0.004289461 | 0.041997032 |
| *ENSG00000259715* | -2.800324 | 0.980619 | -2.855671 | 0.004294606 | 0.041997032 |

| | | | | | |
|---|---|---|---|---|---|
| *ENSG00000179979* | -3.146776 | 1.10196 | -2.855616 | 0.004295336 | 0.041997032 |
| *RCBTB2* | -4.393728 | 1.538819 | -2.855259 | 0.004300169 | 0.042004359 |
| *PCDH11Y* | -4.407703 | 1.543895 | -2.854925 | 0.004304706 | 0.042004359 |
| *KLHL42* | -2.086481 | 0.73096 | -2.85444 | 0.004311279 | 0.042004359 |
| *TNFSF13B* | -2.185985 | 0.765919 | -2.854069 | 0.004316311 | 0.042004359 |
| *ZNF282* | -2.571898 | 0.901196 | -2.853873 | 0.004318974 | 0.042004359 |
| *TMEM220.AS1* | -2.386784 | 0.83643 | -2.853536 | 0.004323556 | 0.042004359 |
| *ENSG00000258634* | -2.643618 | 0.926484 | -2.853389 | 0.004325565 | 0.042004359 |
| *NPHP4* | -3.443738 | 1.207297 | -2.852436 | 0.004338561 | 0.042004359 |
| *PPP2R5D* | -1.875303 | 0.657462 | -2.852338 | 0.004339888 | 0.042004359 |
| *LINC02453* | -2.918314 | 1.023137 | -2.852319 | 0.004340147 | 0.042004359 |
| *KPTN* | -2.164551 | 0.758881 | -2.852292 | 0.004340519 | 0.042004359 |
| *EGR2* | -3.506104 | 1.229327 | -2.852053 | 0.004343791 | 0.042004359 |
| *ENSG00000260360* | -3.822418 | 1.340268 | -2.85198 | 0.00434479 | 0.042004359 |
| *NCKIPSD* | -2.296953 | 0.805429 | -2.851837 | 0.004346737 | 0.042004359 |
| *KDM4D* | -3.577377 | 1.254413 | -2.851833 | 0.004346793 | 0.042004359 |
| *ENSG00000233654* | -3.086191 | 1.08222 | -2.851722 | 0.004348318 | 0.042004359 |
| *PCYOX1L* | -3.088096 | 1.083159 | -2.851008 | 0.004358091 | 0.042037683 |
| *ENSG00000225075* | -3.150046 | 1.104895 | -2.850992 | 0.004358302 | 0.042037683 |
| *ZDHHC23* | -2.06946 | 0.725938 | -2.850741 | 0.004361755 | 0.042039473 |
| *ENSG00000229196* | -3.678035 | 1.290615 | -2.849832 | 0.004374232 | 0.04206885 |
| *ENSG00000272092* | -3.629298 | 1.273667 | -2.849487 | 0.004378973 | 0.04206885 |
| *ENSG00000272841* | -3.352598 | 1.176804 | -2.848901 | 0.004387051 | 0.04206885 |
| *ENSG00000267198* | -3.227716 | 1.133072 | -2.848642 | 0.004390629 | 0.04206885 |
| *TRIM16L* | -2.354353 | 0.826513 | -2.848536 | 0.004392094 | 0.04206885 |
| *ENSG00000277117* | -2.341819 | 0.822158 | -2.84838 | 0.004394241 | 0.04206885 |
| *CFD* | -2.621879 | 0.920691 | -2.84773 | 0.004403231 | 0.04206885 |
| *ENSG00000258017* | -2.736207 | 0.960883 | -2.847597 | 0.004405061 | 0.04206885 |
| *ENSG00000255320* | -3.452409 | 1.212424 | -2.847526 | 0.004406043 | 0.04206885 |
| *ZNF763* | -2.60041 | 0.913237 | -2.847465 | 0.004406888 | 0.04206885 |
| *BIVM* | -2.722385 | 0.956084 | -2.847434 | 0.004407324 | 0.04206885 |
| *ENSG00000259820* | -3.027334 | 1.063288 | -2.847143 | 0.004411355 | 0.04206885 |
| *ENSG00000227374* | -4.479607 | 1.573432 | -2.847028 | 0.004412946 | 0.04206885 |
| *ENSG00000267042* | -4.589874 | 1.61224 | -2.846893 | 0.004414819 | 0.04206885 |
| *LINC02158* | -3.334326 | 1.171267 | -2.846769 | 0.004416536 | 0.04206885 |
| *FAM218A* | -2.582358 | 0.907253 | -2.846348 | 0.00442238 | 0.04206885 |
| *IL3RA* | -3.76203 | 1.321724 | -2.846304 | 0.004422994 | 0.04206885 |
| *VWCE* | -4.321357 | 1.51826 | -2.846257 | 0.004423654 | 0.04206885 |
| *B3GALT6* | -2.224017 | 0.781704 | -2.84509 | 0.004439888 | 0.042149895 |
| *USP20* | -2.100795 | 0.738421 | -2.844982 | 0.004441389 | 0.042149895 |
| *PPA1* | 1.466558 | 0.515497 | 2.844938 | 0.004442003 | 0.042149895 |
| *ENSG00000229180* | -1.849835 | 0.65033 | -2.844454 | 0.004448757 | 0.042152462 |
| *ENSG00000272030* | -3.779548 | 1.328745 | -2.844449 | 0.004448826 | 0.042152462 |
| *ENSG00000213121* | -4.355578 | 1.531622 | -2.843768 | 0.004458351 | 0.042203125 |
| *NET1* | -3.175216 | 1.116722 | -2.843335 | 0.004464407 | 0.042203125 |
| *ENSG00000279765* | -2.25811 | 0.794204 | -2.843237 | 0.004465778 | 0.042203125 |
| *DECR2* | -2.288943 | 0.80512 | -2.842983 | 0.004469343 | 0.042203125 |

| | | | | | |
|---|---|---|---|---|---|
| *SLC14A2* | -2.669229 | 0.938912 | -2.842895 | 0.004470573 | 0.042203125 |
| *CALD1* | -4.566 | 1.606597 | -2.842032 | 0.004482696 | 0.04228655 |
| *ARMCX1* | -3.129057 | 1.101093 | -2.841773 | 0.004486342 | 0.042289939 |
| *STON1* | -3.812719 | 1.342015 | -2.841041 | 0.004496657 | 0.042333328 |
| *MPZL3* | -2.778402 | 0.978279 | -2.840092 | 0.004510057 | 0.042333328 |
| *SLC25A15* | -2.840849 | 1.000276 | -2.840065 | 0.004510431 | 0.042333328 |
| *RIMS2* | -2.971005 | 1.046186 | -2.839844 | 0.004513556 | 0.042333328 |
| *HMGN3.AS1* | -3.014399 | 1.061471 | -2.839832 | 0.004513731 | 0.042333328 |
| *RNF215* | -3.02563 | 1.065428 | -2.839825 | 0.004513834 | 0.042333328 |
| *ZDHHC9* | -2.319935 | 0.816932 | -2.839815 | 0.004513976 | 0.042333328 |
| *TREML2* | -2.860669 | 1.007864 | -2.838349 | 0.004534757 | 0.042477125 |
| *BGLAP* | -3.365793 | 1.186227 | -2.837393 | 0.004548364 | 0.042477125 |
| *TMEM201* | -3.094424 | 1.090638 | -2.837261 | 0.004550237 | 0.042477125 |
| *ENSG00000268439* | -2.673507 | 0.942333 | -2.837116 | 0.004552313 | 0.042477125 |
| *ENSG00000239415* | -2.575766 | 0.907943 | -2.836925 | 0.004555036 | 0.042477125 |
| *SPINT1.AS1* | -2.135042 | 0.752616 | -2.836829 | 0.004556408 | 0.042477125 |
| *PYGB* | -2.099374 | 0.740044 | -2.836823 | 0.00455649 | 0.042477125 |
| *BIRC5* | -2.696927 | 0.950719 | -2.836725 | 0.004557891 | 0.042477125 |
| *ENSG00000239941* | -4.632587 | 1.633292 | -2.83635 | 0.00456324 | 0.042477125 |
| *TTC26* | -2.261784 | 0.797457 | -2.836245 | 0.004564748 | 0.042477125 |
| *PHLDB1* | -2.903475 | 1.023727 | -2.836182 | 0.004565637 | 0.042477125 |
| *TIGD3* | -2.904335 | 1.024113 | -2.835953 | 0.004568924 | 0.042477125 |
| *NRG4* | -3.698141 | 1.304241 | -2.835473 | 0.004575786 | 0.042510213 |
| *NPL* | -3.649688 | 1.28754 | -2.834621 | 0.004588008 | 0.042572138 |
| *ZNF711* | -2.90855 | 1.026107 | -2.834547 | 0.004589069 | 0.042572138 |
| *CA13* | -2.234107 | 0.788238 | -2.834306 | 0.004592533 | 0.042573578 |
| *IGHV1.2* | -3.419923 | 1.206718 | -2.834069 | 0.004595938 | 0.042574466 |
| *NUF2* | -3.656958 | 1.290665 | -2.83339 | 0.00460571 | 0.042634294 |
| *GNG4* | -4.396946 | 1.552366 | -2.832416 | 0.004619777 | 0.042705179 |
| *RIPOR3* | -2.227645 | 0.786487 | -2.8324 | 0.004620005 | 0.042705179 |
| *ADCY9* | -2.898782 | 1.023724 | -2.831605 | 0.004631507 | 0.042771863 |
| *PLPP3* | -2.583243 | 0.912342 | -2.831442 | 0.004633868 | 0.042771863 |
| *SNX24* | -2.955737 | 1.044072 | -2.83097 | 0.004640713 | 0.04280434 |
| *LINC00271* | -2.938595 | 1.038331 | -2.830115 | 0.004653129 | 0.042888119 |
| *LINC02413* | -3.525277 | 1.245735 | -2.829876 | 0.004656606 | 0.042889438 |
| *OXT* | -2.011744 | 0.711019 | -2.829381 | 0.004663818 | 0.042925141 |
| *STK36* | -2.814934 | 0.995115 | -2.828752 | 0.004672993 | 0.042978483 |
| *ENSG00000263004* | -2.606398 | 0.921628 | -2.828036 | 0.004683453 | 0.042978483 |
| *SLC16A1* | -2.750088 | 0.972564 | -2.827666 | 0.004688869 | 0.042978483 |
| *THSD1* | -2.77834 | 0.982706 | -2.827234 | 0.004695195 | 0.042978483 |
| *FNDC3B* | -3.055716 | 1.080837 | -2.827177 | 0.00469604 | 0.042978483 |
| *ENSG00000260030* | -3.078255 | 1.088971 | -2.826756 | 0.004702214 | 0.042978483 |
| *MAST2* | -2.30402 | 0.815245 | -2.82617 | 0.004710832 | 0.042978483 |
| *CFAP53* | -3.072916 | 1.087353 | -2.826054 | 0.004712538 | 0.042978483 |
| *ZSWIM3* | -2.676553 | 0.947115 | -2.826008 | 0.004713212 | 0.042978483 |
| *SLC26A11* | -2.335874 | 0.826595 | -2.8259 | 0.004714794 | 0.042978483 |
| *FAM238A* | -4.380313 | 1.550119 | -2.825792 | 0.004716392 | 0.042978483 |

| | | | | | |
|---|---|---|---|---|---|
| *KLHL23* | -2.333408 | 0.825845 | -2.82548 | 0.004720987 | 0.042978483 |
| *ENSG00000232934* | -3.450289 | 1.221154 | -2.825433 | 0.004721682 | 0.042978483 |
| *ZNF710* | -2.273814 | 0.804786 | -2.825365 | 0.004722677 | 0.042978483 |
| *CCNT2.AS1* | -2.72715 | 0.965256 | -2.825312 | 0.004723457 | 0.042978483 |
| *KLHL15* | -2.791693 | 0.988112 | -2.82528 | 0.004723934 | 0.042978483 |
| *ACAD11* | -3.218928 | 1.139398 | -2.825113 | 0.004726397 | 0.042978483 |
| *ENSG00000228606* | -2.77324 | 0.981772 | -2.824729 | 0.004732056 | 0.042999554 |
| *ENSG00000273055* | -3.640148 | 1.290357 | -2.821039 | 0.004786838 | 0.04345158 |
| *MST1* | -2.848161 | 1.009724 | -2.820732 | 0.004791417 | 0.04345158 |
| *RNASEH2B.AS1* | -4.064638 | 1.441042 | -2.820623 | 0.004793046 | 0.04345158 |
| *TTC24* | -2.768554 | 0.981623 | -2.820384 | 0.004796619 | 0.04345158 |
| *MAD2L1* | -2.221079 | 0.787548 | -2.820246 | 0.004798686 | 0.04345158 |
| *SGTB* | -1.773326 | 0.628904 | -2.819708 | 0.004806743 | 0.043479759 |
| *PDGFRB* | -3.11063 | 1.103222 | -2.819587 | 0.004808557 | 0.043479759 |
| *RHOBTB2* | -2.218537 | 0.786981 | -2.819047 | 0.004816642 | 0.043520908 |
| *PLPP6* | -2.053469 | 0.728482 | -2.818832 | 0.004819872 | 0.043520908 |
| *ENSG00000261242* | -4.501706 | 1.597756 | -2.817518 | 0.004839638 | 0.043648354 |
| *TMC3.AS1* | -2.497441 | 0.886421 | -2.817443 | 0.004840771 | 0.043648354 |
| *FXYD2* | -1.471203 | 0.522288 | -2.816841 | 0.004849852 | 0.043699615 |
| *NR4A2* | -2.232731 | 0.792703 | -2.816604 | 0.004853436 | 0.043701301 |
| *MIR3142HG* | -3.992278 | 1.417539 | -2.816344 | 0.004857367 | 0.043706117 |
| *PFN2* | -3.215618 | 1.141921 | -2.815973 | 0.004862978 | 0.043718386 |
| *HLCS* | -2.327809 | 0.826713 | -2.815741 | 0.004866487 | 0.043718386 |
| *ZNF410* | -3.177942 | 1.128865 | -2.815165 | 0.004875216 | 0.043718386 |
| *STK39* | -2.716101 | 0.964834 | -2.815097 | 0.004876255 | 0.043718386 |
| *ENSG00000272155* | -2.213414 | 0.786269 | -2.815084 | 0.004876452 | 0.043718386 |
| *CSGALNACT1* | -1.733002 | 0.615651 | -2.814908 | 0.004879117 | 0.043718386 |
| *ENSG00000235522* | -4.795325 | 1.704312 | -2.813642 | 0.00489838 | 0.043860441 |
| *ZNF597* | -2.936863 | 1.044093 | -2.812836 | 0.004910669 | 0.043926121 |
| *ENSG00000277767* | -3.029363 | 1.077025 | -2.812713 | 0.004912543 | 0.043926121 |
| *TLR2* | -4.290986 | 1.525732 | -2.812411 | 0.004917156 | 0.04393684 |
| *IGLV1.44* | -2.859327 | 1.016904 | -2.811797 | 0.004926551 | 0.043990236 |
| *CST3* | -2.499917 | 0.889245 | -2.81128 | 0.004934488 | 0.044018396 |
| *ZFP36L1* | -0.44311 | 0.157627 | -2.811127 | 0.004936836 | 0.044018396 |
| *RAB11FIP1* | -1.274679 | 0.453499 | -2.810763 | 0.004942422 | 0.044018396 |
| *DCLRE1A* | -3.342456 | 1.18919 | -2.8107 | 0.004943389 | 0.044018396 |
| *SLC25A25.AS1* | -2.237811 | 0.796651 | -2.809022 | 0.004969231 | 0.04417005 |
| *MYO18A* | -3.252321 | 1.157834 | -2.80897 | 0.004970033 | 0.04417005 |
| *ENSG00000116957* | -1.909536 | 0.67981 | -2.808925 | 0.004970718 | 0.04417005 |
| *HSPB1* | -1.047125 | 0.372899 | -2.808063 | 0.004984041 | 0.044257873 |
| *DCAF15* | -1.589084 | 0.565987 | -2.807636 | 0.004990654 | 0.044286032 |
| *CHEK1* | -2.695902 | 0.960336 | -2.807249 | 0.004996664 | 0.044287396 |
| *SORBS3* | -2.218958 | 0.790457 | -2.807183 | 0.004997692 | 0.044287396 |
| *HIPK1.AS1* | -2.237713 | 0.797236 | -2.806839 | 0.005003024 | 0.044304137 |
| *CDCA5* | -4.184926 | 1.491093 | -2.806617 | 0.005006471 | 0.044304166 |
| *ENSG00000228172* | -2.691437 | 0.959128 | -2.806129 | 0.005014059 | 0.044340821 |
| *RADX* | -2.499148 | 0.890742 | -2.805693 | 0.005020845 | 0.044352256 |

| | | | | | |
|---|---|---|---|---|---|
| *SSTR2* | -2.54866 | 0.90847 | -2.805442 | 0.005024759 | 0.044352256 |
| *ENSG00000263069* | -3.51794 | 1.254069 | -2.805221 | 0.005028214 | 0.044352256 |
| *ZNF547* | -2.505761 | 0.893268 | -2.805161 | 0.00502914 | 0.044352256 |
| *ENSG00000259804* | -3.551167 | 1.266149 | -2.8047 | 0.005036341 | 0.044382756 |
| *KHDC1* | -2.624925 | 0.936034 | -2.804307 | 0.005042488 | 0.044382756 |
| *IGLV3.19* | -3.351776 | 1.195237 | -2.804278 | 0.005042946 | 0.044382756 |
| *ENSG00000279571* | -4.455073 | 1.588847 | -2.803965 | 0.005047831 | 0.044387488 |
| *HSPA4L* | -2.455169 | 0.875713 | -2.803621 | 0.005053225 | 0.044387488 |
| *ZSCAN31* | -3.997185 | 1.425742 | -2.803582 | 0.005053833 | 0.044387488 |
| *ENSG00000251259* | -2.612974 | 0.932316 | -2.802669 | 0.005068173 | 0.04446552 |
| *DNAJC27.AS1* | -2.870704 | 1.024309 | -2.802576 | 0.005069629 | 0.04446552 |
| *IGSF6* | -4.445298 | 1.586451 | -2.802039 | 0.005078073 | 0.044509238 |
| *MYOM2* | -2.556652 | 0.912656 | -2.80133 | 0.005089237 | 0.044576727 |
| *INAFM2* | -4.417994 | 1.577471 | -2.800681 | 0.005099489 | 0.044608366 |
| *ANKRD34A* | -4.524763 | 1.615604 | -2.800662 | 0.005099783 | 0.044608366 |
| *ENSG00000272669* | -2.951017 | 1.053855 | -2.800213 | 0.005106892 | 0.044640207 |
| *MSS51* | -3.676186 | 1.313137 | -2.799546 | 0.005117458 | 0.044702197 |
| *ZNF426.DT* | -3.490956 | 1.247104 | -2.799249 | 0.005122161 | 0.044712918 |
| *TIGIT* | -3.222451 | 1.151394 | -2.798738 | 0.005130276 | 0.044730811 |
| *THAP9* | -2.614144 | 0.934062 | -2.798682 | 0.005131163 | 0.044730811 |
| *ZBTB33* | -1.736005 | 0.620382 | -2.798284 | 0.00513749 | 0.044755639 |
| *ENSG00000266918* | -2.412721 | 0.862311 | -2.797971 | 0.005142475 | 0.044757162 |
| *PPP1R32* | -2.892652 | 1.033889 | -2.797836 | 0.005144621 | 0.044757162 |
| *PRPF40B* | -2.176801 | 0.77814 | -2.797442 | 0.005150896 | 0.044781476 |
| *ENSG00000203279* | -2.77232 | 0.991323 | -2.796585 | 0.005164577 | 0.044801815 |
| *RNF157* | -3.675046 | 1.314234 | -2.796341 | 0.005168478 | 0.044801815 |
| *LRRC25* | -4.255215 | 1.521727 | -2.796307 | 0.005169032 | 0.044801815 |
| *TEC* | -2.176 | 0.7782 | -2.796198 | 0.005170774 | 0.044801815 |
| *ZNF66* | -2.74131 | 0.980441 | -2.795996 | 0.005173997 | 0.044801815 |
| *IL18* | -3.135429 | 1.121403 | -2.795988 | 0.005174127 | 0.044801815 |
| *ENSG00000270964* | -4.219314 | 1.509425 | -2.795312 | 0.005184967 | 0.044811117 |
| *CTSK* | -3.29173 | 1.177626 | -2.795224 | 0.005186373 | 0.044811117 |
| *CDKN2C* | -3.195585 | 1.14331 | -2.795028 | 0.005189516 | 0.044811117 |
| *NRSN2.AS1* | -2.238074 | 0.800753 | -2.794962 | 0.005190582 | 0.044811117 |
| *HPSE* | -3.53813 | 1.265953 | -2.794835 | 0.005192615 | 0.044811117 |
| *UBTD1* | -3.403823 | 1.218248 | -2.79403 | 0.005205567 | 0.044892783 |
| *ENSG00000230177* | -2.127162 | 0.761634 | -2.792894 | 0.005223883 | 0.044993694 |
| *CCNA1* | -3.519475 | 1.260164 | -2.79287 | 0.005224262 | 0.044993694 |
| *LINC00896* | -4.029944 | 1.443067 | -2.792625 | 0.005228221 | 0.044995111 |
| *ENSG00000150076* | -3.599545 | 1.289038 | -2.792427 | 0.00523142 | 0.044995111 |
| *ENSG00000203325* | -2.286497 | 0.818926 | -2.792068 | 0.005237232 | 0.04501501 |
| *ZNF236.DT* | -2.884048 | 1.033162 | -2.791476 | 0.005246824 | 0.045067344 |
| *ENSG00000262692* | -4.344501 | 1.556538 | -2.791131 | 0.005252419 | 0.045085307 |
| *TSPYL5* | -2.494422 | 0.893857 | -2.790628 | 0.005260587 | 0.045125312 |
| *MYL9* | -2.197767 | 0.787923 | -2.789315 | 0.005281961 | 0.045142747 |
| *NDST2* | -2.46223 | 0.882778 | -2.789185 | 0.005284087 | 0.045142747 |
| *AHCYL2* | -2.971302 | 1.065295 | -2.789181 | 0.005284145 | 0.045142747 |

| | | | | | |
|---|---|---|---|---|---|
| *ENSG00000254427* | -3.013906 | 1.08061 | -2.789077 | 0.005285845 | 0.045142747 |
| *ZBTB8B* | -4.315151 | 1.547241 | -2.788934 | 0.00528819 | 0.045142747 |
| *ENSG00000266086* | -4.315151 | 1.547241 | -2.788934 | 0.00528819 | 0.045142747 |
| *MIR222HG* | -2.68224 | 0.961831 | -2.78868 | 0.005292336 | 0.045142747 |
| *PTPRF* | -3.729919 | 1.337701 | -2.788305 | 0.00529846 | 0.045142747 |
| *C5* | -3.252064 | 1.166422 | -2.788069 | 0.005302328 | 0.045142747 |
| *NKX3.1* | -3.078872 | 1.104317 | -2.788033 | 0.005302922 | 0.045142747 |
| *TUBGCP5* | -2.065681 | 0.74093 | -2.787956 | 0.00530417 | 0.045142747 |
| *AFAP1L2* | -2.9749 | 1.067067 | -2.787923 | 0.00530472 | 0.045142747 |
| *ENSG00000231212* | -2.923473 | 1.048792 | -2.787468 | 0.005312176 | 0.045174297 |
| *MIXL1* | -4.374933 | 1.569691 | -2.78713 | 0.00531772 | 0.045174297 |
| *GDPD1* | -2.259145 | 0.810631 | -2.786895 | 0.005321569 | 0.045174297 |
| *SPESP1* | -3.453238 | 1.239334 | -2.786366 | 0.005330269 | 0.045174297 |
| *DCUN1D2* | -2.177143 | 0.781358 | -2.786356 | 0.005330424 | 0.045174297 |
| *KCP* | -3.266138 | 1.172231 | -2.786258 | 0.005332049 | 0.045174297 |
| *ENSG00000270024* | -4.423028 | 1.587524 | -2.786116 | 0.00533437 | 0.045174297 |
| *COMMD7* | -1.094367 | 0.392854 | -2.785679 | 0.00534157 | 0.045174297 |
| *LINC02656* | -2.552368 | 0.91626 | -2.785637 | 0.005342267 | 0.045174297 |
| *ENSG00000249141* | -3.067112 | 1.101129 | -2.785424 | 0.005345772 | 0.045174297 |
| *ENSG00000245498* | -3.593018 | 1.289971 | -2.785347 | 0.005347047 | 0.045174297 |
| *SLC27A3* | -2.418552 | 0.868864 | -2.78358 | 0.005376256 | 0.045383928 |
| *ENSG00000262420* | -3.275267 | 1.176743 | -2.783332 | 0.005380378 | 0.045383928 |
| *IZUMO4* | -1.101219 | 0.395702 | -2.782953 | 0.005386661 | 0.045383928 |
| *ENSG00000269176* | -2.875067 | 1.03312 | -2.782898 | 0.005387574 | 0.045383928 |
| *YRDC* | -2.84329 | 1.021814 | -2.782591 | 0.005392682 | 0.045383928 |
| *LARGE1* | -2.369424 | 0.851524 | -2.78257 | 0.005393023 | 0.045383928 |
| *PLK2* | -4.018963 | 1.444548 | -2.782159 | 0.00539986 | 0.045411763 |
| *ENSG00000270022* | -2.845802 | 1.023316 | -2.78096 | 0.005419836 | 0.045549984 |
| *TTYH1* | -4.090595 | 1.471742 | -2.779423 | 0.005445552 | 0.045736235 |
| *ENSG00000259520* | -4.134816 | 1.488531 | -2.777784 | 0.005473107 | 0.04592137 |
| *ING2* | -1.47853 | 0.532288 | -2.777687 | 0.005474732 | 0.04592137 |
| *EPHA1.AS1* | -2.193778 | 0.789894 | -2.777306 | 0.005481147 | 0.045941257 |
| *ENSG00000261613* | -3.605582 | 1.298316 | -2.777123 | 0.005484244 | 0.045941257 |
| *ENSG00000259153* | -2.464129 | 0.887534 | -2.776378 | 0.005496825 | 0.046016687 |
| *LY6E.DT* | -2.614447 | 0.941844 | -2.775881 | 0.005505238 | 0.046042955 |
| *GGT7* | -2.111566 | 0.760767 | -2.775574 | 0.005510439 | 0.046042955 |
| *MAPKBP1* | -2.145577 | 0.773067 | -2.775408 | 0.005513242 | 0.046042955 |
| *COQ3* | -2.759263 | 0.994204 | -2.775347 | 0.005514276 | 0.046042955 |
| *CAPN14* | -4.461281 | 1.607813 | -2.774752 | 0.005524387 | 0.046075089 |
| *LRRC27* | -1.918331 | 0.691377 | -2.774652 | 0.00552608 | 0.046075089 |
| *AHRR* | -3.418659 | 1.23228 | -2.774256 | 0.005532804 | 0.046075089 |
| *HAL* | -3.545346 | 1.278113 | -2.773891 | 0.005539025 | 0.046075089 |
| *RPL35* | -0.148872 | 0.053672 | -2.77375 | 0.00554143 | 0.046075089 |
| *CTU1* | -4.357563 | 1.571023 | -2.77371 | 0.0055421 | 0.046075089 |
| *DDX43* | -2.637649 | 0.95097 | -2.773642 | 0.005543272 | 0.046075089 |
| *ENSG00000261684* | -4.004894 | 1.444164 | -2.773158 | 0.005551523 | 0.046075089 |
| *ENSG00000255328* | -3.175841 | 1.145221 | -2.773124 | 0.005552098 | 0.046075089 |

| | | | | | |
|---|---|---|---|---|---|
| *MAST1* | -4.466241 | 1.61078 | -2.77272 | 0.005558994 | 0.046075089 |
| *ENSG00000261732* | -1.811899 | 0.653575 | -2.772289 | 0.005566364 | 0.046075089 |
| *RBKS* | -2.708021 | 0.976825 | -2.77227 | 0.005566691 | 0.046075089 |
| *DPH5* | -1.107225 | 0.399404 | -2.772196 | 0.005567947 | 0.046075089 |
| *ENSG00000280987* | -2.478818 | 0.89419 | -2.772138 | 0.005568949 | 0.046075089 |
| *DSCR9* | -3.8278 | 1.381078 | -2.771603 | 0.0055781 | 0.046075089 |
| *ENSG00000184271* | -2.45735 | 0.88662 | -2.771595 | 0.005578243 | 0.046075089 |
| *ENSG00000269737* | -3.00634 | 1.084714 | -2.771551 | 0.005579 | 0.046075089 |
| *YBEY* | -1.079212 | 0.389438 | -2.771201 | 0.005584996 | 0.046095024 |
| *VRK1* | -1.476334 | 0.532845 | -2.770663 | 0.005594223 | 0.046141579 |
| *FZD6* | -3.504192 | 1.26492 | -2.770288 | 0.00560068 | 0.046144081 |
| *ENSG00000270175* | -2.472971 | 0.892747 | -2.770068 | 0.005604465 | 0.046144081 |
| *ARL14EPL* | -2.548671 | 0.920091 | -2.77002 | 0.005605285 | 0.046144081 |
| *NEK11* | -3.737486 | 1.349386 | -2.769768 | 0.00560963 | 0.046150323 |
| *B3GNTL1* | -2.252584 | 0.81336 | -2.769479 | 0.005614604 | 0.046161734 |
| *SPATA5* | -1.950695 | 0.704458 | -2.769073 | 0.005621601 | 0.046189745 |
| *IL17RC* | -4.45461 | 1.609172 | -2.768262 | 0.005635618 | 0.046275363 |
| *GPAT2* | -4.323565 | 1.56244 | -2.767188 | 0.005654207 | 0.046333227 |
| *FITM1* | -3.221279 | 1.164163 | -2.767034 | 0.005656887 | 0.046333227 |
| *ENSG00000250075* | -2.629137 | 0.950189 | -2.766963 | 0.005658121 | 0.046333227 |
| *ITPKC* | -2.747738 | 0.993145 | -2.766704 | 0.00566262 | 0.046333227 |
| *KIAA1324L* | -2.470835 | 0.893071 | -2.766674 | 0.00566313 | 0.046333227 |
| *ENSG00000234290* | -4.347293 | 1.571435 | -2.766448 | 0.005667064 | 0.046333227 |
| *ST6GALNAC2* | -3.602314 | 1.302166 | -2.766401 | 0.005667871 | 0.046333227 |
| *ENSG00000227908* | -4.204376 | 1.520023 | -2.765995 | 0.00567494 | 0.046361555 |
| *ENSG00000258738* | -2.66284 | 0.962924 | -2.76537 | 0.005685819 | 0.046420961 |
| *SPTY2D1OS* | -3.432225 | 1.241538 | -2.764494 | 0.005701118 | 0.046479903 |
| *PIK3IP1.AS1* | -2.836221 | 1.026091 | -2.764102 | 0.005707969 | 0.046479903 |
| *RASAL2* | -2.934472 | 1.061652 | -2.764062 | 0.005708661 | 0.046479903 |
| *NINL* | -2.979253 | 1.077934 | -2.763856 | 0.005712277 | 0.046479903 |
| *ENSG00000274922* | -4.497049 | 1.627097 | -2.763848 | 0.005712417 | 0.046479903 |
| *HIST1H3H* | -3.135087 | 1.134373 | -2.763717 | 0.005714713 | 0.046479903 |
| *NEU3* | -2.561439 | 0.927064 | -2.762958 | 0.005728016 | 0.04655055 |
| *ENSG00000275180* | -2.563291 | 0.927843 | -2.762635 | 0.005733692 | 0.04655055 |
| *TTC12* | -2.499415 | 0.904732 | -2.762603 | 0.005734252 | 0.04655055 |
| *ENSG00000267011* | -3.225702 | 1.168112 | -2.761466 | 0.005754244 | 0.046661746 |
| *TIGD7* | -2.03989 | 0.738713 | -2.761412 | 0.005755203 | 0.046661746 |
| *DSCAML1* | -3.087603 | 1.118428 | -2.760664 | 0.0057684 | 0.046739298 |
| *ENSG00000259888* | -3.291486 | 1.192412 | -2.760358 | 0.005773798 | 0.046753592 |
| *ENSG00000261360* | -2.985062 | 1.081541 | -2.760009 | 0.005779974 | 0.046765401 |
| *CDYL2* | -3.756899 | 1.361262 | -2.759865 | 0.005782525 | 0.046765401 |
| *PANK1* | -2.966543 | 1.075325 | -2.75874 | 0.005802464 | 0.046896653 |
| *LINC01250* | -4.321618 | 1.566633 | -2.758539 | 0.005806044 | 0.046896653 |
| *IQCC* | -2.5641 | 0.929653 | -2.758128 | 0.005813347 | 0.046926182 |
| *C19orf33* | -2.895117 | 1.049977 | -2.757313 | 0.005827848 | 0.04696095 |
| *QPCTL* | -2.511724 | 0.910941 | -2.757284 | 0.005828376 | 0.04696095 |
| *LINC02210.CRHR1* | -3.339639 | 1.211304 | -2.757062 | 0.005832335 | 0.04696095 |

| | | | | | |
|---|---|---|---|---|---|
| *ENSG00000261441* | -3.605937 | 1.307949 | -2.75694 | 0.005834498 | 0.04696095 |
| *C16orf46* | -3.381739 | 1.226687 | -2.756807 | 0.005836875 | 0.04696095 |
| *ENSG00000272434* | -4.47823 | 1.624515 | -2.756657 | 0.005839552 | 0.04696095 |
| *PGP* | -2.027114 | 0.73543 | -2.756367 | 0.005844734 | 0.046973261 |
| *ENSG00000260060* | -3.15884 | 1.14616 | -2.75602 | 0.005850942 | 0.046993801 |
| *SLC39A14* | -3.962116 | 1.437908 | -2.755473 | 0.005860742 | 0.047020022 |
| *LINC01285* | -4.011848 | 1.456048 | -2.755299 | 0.005863858 | 0.047020022 |
| *HFE* | -2.428032 | 0.881246 | -2.755226 | 0.005865169 | 0.047020022 |
| *UTP15* | -2.044844 | 0.742342 | -2.754585 | 0.005876661 | 0.047082813 |
| *ENSG00000224905* | -3.619316 | 1.31416 | -2.754091 | 0.005885545 | 0.047115174 |
| *MSANTD3* | -2.106678 | 0.765062 | -2.753605 | 0.005894294 | 0.047115174 |
| *FEM1C* | -2.08519 | 0.75726 | -2.753597 | 0.005894436 | 0.047115174 |
| *ENSG00000257261* | -2.754088 | 1.000255 | -2.753386 | 0.005898223 | 0.047115174 |
| *LNX2* | -2.020268 | 0.733751 | -2.753343 | 0.005899009 | 0.047115174 |
| *EXOC6B* | -2.463626 | 0.894885 | -2.753008 | 0.005905049 | 0.04713416 |
| *KIZ.AS1* | -2.436049 | 0.885045 | -2.752458 | 0.005914974 | 0.047184108 |
| *PDSS1* | -2.468975 | 0.897314 | -2.751516 | 0.005932005 | 0.04729065 |
| *IER3.AS1* | -2.572568 | 0.935073 | -2.751196 | 0.005937804 | 0.047299008 |
| *SYNGAP1.AS1* | -2.227591 | 0.809723 | -2.751053 | 0.005940405 | 0.047299008 |
| *ITCH.AS1* | -3.428862 | 1.246959 | -2.74978 | 0.005963528 | 0.047440347 |
| *PHLPP1* | -2.084751 | 0.758197 | -2.749615 | 0.005966532 | 0.047440347 |
| *CDC42BPA* | -4.376074 | 1.591711 | -2.749288 | 0.005972483 | 0.047440347 |
| *SRCAP* | -2.630904 | 0.957178 | -2.748606 | 0.005984933 | 0.047440347 |
| *ENSG00000276449* | -2.270366 | 0.826118 | -2.748233 | 0.005991745 | 0.047440347 |
| *ENSG00000250616* | -2.828964 | 1.029427 | -2.748096 | 0.005994249 | 0.047440347 |
| *NUDT17* | -3.361029 | 1.223063 | -2.748043 | 0.005995213 | 0.047440347 |
| *ENSG00000260219* | -2.787931 | 1.014517 | -2.748039 | 0.005995284 | 0.047440347 |
| *SCML4* | -2.610444 | 0.949935 | -2.748023 | 0.005995581 | 0.047440347 |
| *DNAJA4* | -2.894283 | 1.05325 | -2.747956 | 0.005996804 | 0.047440347 |
| *LINC02035* | -2.771458 | 1.008591 | -2.747852 | 0.005998713 | 0.047440347 |
| *ENSG00000245975* | -4.031448 | 1.467295 | -2.747538 | 0.006004455 | 0.047456587 |
| *PPP2R3A* | -2.370426 | 0.862919 | -2.746985 | 0.006014589 | 0.047507501 |
| *ENSG00000226266* | -2.666397 | 0.971014 | -2.745992 | 0.006032827 | 0.047611697 |
| *SLC35G1* | -3.411937 | 1.242584 | -2.74584 | 0.006035626 | 0.047611697 |
| *GCNA* | -2.54785 | 0.927996 | -2.745539 | 0.006041159 | 0.047611697 |
| *TMEM121* | -3.821664 | 1.391993 | -2.745462 | 0.006042582 | 0.047611697 |
| *HIST1H2BK* | -2.158094 | 0.786122 | -2.74524 | 0.006046662 | 0.047614684 |
| *KLHL34* | -4.100939 | 1.494083 | -2.744786 | 0.006055029 | 0.04765141 |
| *WDR97* | -4.41398 | 1.608337 | -2.744438 | 0.006061468 | 0.047663704 |
| *SARM1* | -2.103814 | 0.766612 | -2.744301 | 0.006064 | 0.047663704 |
| *NATD1* | -2.497055 | 0.91001 | -2.743987 | 0.006069791 | 0.047680102 |
| *LIPT2* | -2.177118 | 0.793665 | -2.743121 | 0.006085822 | 0.047776856 |
| *ZDHHC18* | -1.945881 | 0.709549 | -2.742419 | 0.006098851 | 0.047849945 |
| *UNC5CL* | -3.248639 | 1.18478 | -2.741977 | 0.006107062 | 0.047885169 |
| *LRRC8C* | -2.141388 | 0.78125 | -2.740976 | 0.006125697 | 0.047940112 |
| *ENSG00000282386* | -2.574743 | 0.939363 | -2.740946 | 0.006126255 | 0.047940112 |
| *MAGIX* | -3.125011 | 1.140227 | -2.740691 | 0.006131009 | 0.047940112 |

| | | | | | |
|---|---|---|---|---|---|
| *ENSG00000279467* | -2.692841 | 0.982562 | -2.740633 | 0.006132103 | 0.047940112 |
| *MED27* | -2.529994 | 0.92327 | -2.740255 | 0.006139155 | 0.047940112 |
| *RFNG* | -2.057375 | 0.750804 | -2.740229 | 0.006139637 | 0.047940112 |
| *ENSG00000275413* | -3.20454 | 1.169502 | -2.740088 | 0.006142277 | 0.047940112 |
| *RIC8B* | -2.682593 | 0.979048 | -2.740002 | 0.006143875 | 0.047940112 |
| *TEX2* | -2.516946 | 0.91877 | -2.739473 | 0.006153786 | 0.047988338 |
| *UHRF1BP1* | -2.189775 | 0.799507 | -2.738906 | 0.006164403 | 0.048042016 |
| *PRSS21* | -4.292263 | 1.56729 | -2.738652 | 0.006169154 | 0.048049945 |
| *ENSG00000273373* | -2.181969 | 0.796883 | -2.738129 | 0.006178988 | 0.048085387 |
| *POMT2* | -2.536118 | 0.926263 | -2.738012 | 0.006181179 | 0.048085387 |
| *HAND2.AS1* | -3.954951 | 1.444608 | -2.737732 | 0.006186445 | 0.048097276 |
| *LLGL1* | -2.186543 | 0.798979 | -2.736672 | 0.00620642 | 0.048223436 |
| *FUT11* | -1.549017 | 0.566232 | -2.735658 | 0.006225568 | 0.048337248 |
| *KLHDC9* | -3.197609 | 1.168977 | -2.735391 | 0.006230631 | 0.048337248 |
| *ABCD1* | -2.560589 | 0.936218 | -2.735034 | 0.006237385 | 0.048337248 |
| *ZNF513* | -2.187223 | 0.799756 | -2.734862 | 0.006240649 | 0.048337248 |
| *ENSG00000245466* | -2.796497 | 1.022599 | -2.734695 | 0.006243806 | 0.048337248 |
| *CBLL1* | -1.433595 | 0.52425 | -2.734564 | 0.00624629 | 0.048337248 |
| *TTC22* | -3.268891 | 1.19548 | -2.734375 | 0.006249877 | 0.048337248 |
| *KLB* | -3.431404 | 1.255164 | -2.733828 | 0.006260267 | 0.048337248 |
| *ENSG00000255224* | -3.970289 | 1.452314 | -2.733768 | 0.006261409 | 0.048337248 |
| *ENSG00000260495* | -2.370255 | 0.867087 | -2.733584 | 0.00626491 | 0.048337248 |
| *ENTR1* | -1.789442 | 0.654618 | -2.733568 | 0.006265226 | 0.048337248 |
| *ARF4.AS1* | -2.447484 | 0.89536 | -2.733519 | 0.006266148 | 0.048337248 |
| *SYT5* | -4.19074 | 1.533384 | -2.733001 | 0.006276012 | 0.048384329 |
| *PLPPR2* | -4.334269 | 1.586493 | -2.731982 | 0.006295465 | 0.048503969 |
| *DLG4* | -4.250484 | 1.555932 | -2.731793 | 0.00629907 | 0.048503969 |
| *PIGB* | -1.793852 | 0.656813 | -2.731146 | 0.006311446 | 0.048570199 |
| *STIMATE* | -3.175454 | 1.162872 | -2.7307 | 0.006319997 | 0.048606932 |
| *DGCR6* | -2.765962 | 1.013042 | -2.730353 | 0.006326653 | 0.048623523 |
| *ENSG00000266340* | -2.488098 | 0.911327 | -2.730194 | 0.006329712 | 0.048623523 |
| *FBXO45* | -2.318563 | 0.849398 | -2.729654 | 0.006340087 | 0.048650764 |
| *NEDD4L* | -2.596456 | 0.951275 | -2.729448 | 0.006344039 | 0.048650764 |
| *ZC2HC1C* | -2.204809 | 0.807853 | -2.729221 | 0.006348406 | 0.048650764 |
| *LRATD2* | -3.186803 | 1.167716 | -2.729092 | 0.006350905 | 0.048650764 |
| *ENSG00000228005* | -3.240938 | 1.18758 | -2.729026 | 0.006352163 | 0.048650764 |
| *GSTA4* | -3.059298 | 1.121239 | -2.728498 | 0.00636235 | 0.048678369 |
| *SCRG1* | -2.67262 | 0.979539 | -2.728447 | 0.006363334 | 0.048678369 |
| *LINC01481* | -4.200504 | 1.539695 | -2.728139 | 0.006369272 | 0.048694842 |
| *DNAJB5* | -3.890297 | 1.426679 | -2.726821 | 0.006394766 | 0.048860718 |
| *ENSG00000230709* | -2.2449 | 0.823405 | -2.726363 | 0.006403659 | 0.048899631 |
| *HIST2H2BF* | -2.508832 | 0.920367 | -2.725905 | 0.006412549 | 0.048938478 |
| *ENSG00000132481* | -4.154198 | 1.524658 | -2.724676 | 0.006436461 | 0.049091847 |
| *NBEAL2* | -2.224102 | 0.81696 | -2.722412 | 0.006480731 | 0.049398934 |
| *ENSG00000276075* | -3.471097 | 1.275182 | -2.72204 | 0.006488028 | 0.049398934 |
| *LINC01004* | -3.47283 | 1.275824 | -2.722029 | 0.006488241 | 0.049398934 |
| *LRRFIP1* | -0.673056 | 0.247384 | -2.720693 | 0.00651452 | 0.049569677 |

| | | | | | |
|---|---|---|---|---|---|
| *ZNF485* | -2.044715 | 0.751631 | -2.720371 | 0.006520862 | 0.049588611 |
| *MYBPC2* | -3.140532 | 1.154951 | -2.719192 | 0.006544168 | 0.049698704 |
| *DNAJC5B* | -2.298146 | 0.845185 | -2.719106 | 0.00654587 | 0.049698704 |
| *ENSG00000229728* | -3.880276 | 1.427069 | -2.719052 | 0.006546926 | 0.049698704 |
| *ANKS1A* | -2.356635 | 0.867194 | -2.717542 | 0.006576884 | 0.049877582 |
| *TBC1D2* | -3.857391 | 1.419479 | -2.71747 | 0.006578308 | 0.049877582 |
| *NPIPB6* | -3.884665 | 1.429616 | -2.717278 | 0.00658212 | 0.049877582 |
| *SMYD3* | -1.785778 | 0.657288 | -2.716887 | 0.006589908 | 0.049892952 |
| *ENSG00000269318* | -4.107146 | 1.511844 | -2.716647 | 0.006594685 | 0.049892952 |
| *MORN4* | -4.219507 | 1.553235 | -2.716592 | 0.006595781 | 0.049892952 |
| *TTC30B* | -2.78022 | 1.023716 | -2.715812 | 0.006611341 | 0.04998127 |
| *TIGD6* | -2.38776 | 0.879294 | -2.715542 | 0.006616736 | 0.049992684 |
| *ZNF571* | -1.948713 | 0.718066 | -2.713834 | 0.006650953 | 0.05022172 |
| *ZNF831* | -4.225178 | 1.55715 | -2.713404 | 0.006659584 | 0.0502286 |
| *ENSG00000280316* | -2.409851 | 0.888193 | -2.713207 | 0.006663538 | 0.0502286 |
| *MZF1.AS1* | -2.889304 | 1.064969 | -2.713041 | 0.006666885 | 0.0502286 |
| *LINC02362* | -2.43306 | 0.896812 | -2.713012 | 0.006667479 | 0.0502286 |
| *UST* | -2.412378 | 0.889503 | -2.712051 | 0.006686837 | 0.050326495 |
| *GPRC5D.AS1* | -3.522637 | 1.298918 | -2.711978 | 0.006688296 | 0.050326495 |
| *TFR2* | -2.353497 | 0.868276 | -2.710541 | 0.00671736 | 0.050457977 |
| *PAQR4* | -4.095563 | 1.510977 | -2.71054 | 0.006717366 | 0.050457977 |
| *ENSG00000272037* | -3.237383 | 1.19439 | -2.710491 | 0.006718377 | 0.050457977 |
| *FBXL13* | -4.181355 | 1.542743 | -2.710339 | 0.006721456 | 0.050457977 |
| *ZNF74* | -2.211579 | 0.816073 | -2.710026 | 0.006727802 | 0.05047617 |
| *IL24* | -3.559364 | 1.313588 | -2.709651 | 0.0067354 | 0.050484504 |
| *HIST1H4J* | -2.172019 | 0.801606 | -2.709584 | 0.00673676 | 0.050484504 |
| *RCOR1* | -1.96949 | 0.726957 | -2.709225 | 0.006744065 | 0.050509831 |
| *ZNF613* | -2.121375 | 0.78319 | -2.708636 | 0.006756047 | 0.050557471 |
| *ENSG00000266993* | -3.323522 | 1.22706 | -2.708526 | 0.006758285 | 0.050557471 |
| *ZNF404* | -2.326762 | 0.85912 | -2.708308 | 0.006762716 | 0.050561223 |
| *TRIM2* | -4.31042 | 1.592278 | -2.707077 | 0.006787841 | 0.050693036 |
| *ENSG00000257221* | -3.16416 | 1.168966 | -2.706802 | 0.006793482 | 0.050693036 |
| *LINC00663* | -2.056705 | 0.759833 | -2.706788 | 0.006793772 | 0.050693036 |
| *ZEB2.AS1* | -2.364687 | 0.873651 | -2.706674 | 0.006796105 | 0.050693036 |
| *PRKAR2A.AS1* | -4.049832 | 1.496572 | -2.706072 | 0.006808423 | 0.050737887 |
| *ENSG00000234484* | -4.159488 | 1.537154 | -2.705966 | 0.006810592 | 0.050737887 |
| *ACCS* | -1.831461 | 0.676864 | -2.705803 | 0.006813948 | 0.050737887 |
| *ENSG00000271895* | -2.833421 | 1.047318 | -2.705406 | 0.006822095 | 0.050769167 |
| *ZNF436* | -3.429625 | 1.268135 | -2.704463 | 0.006841482 | 0.050881552 |
| *DUSP11* | 1.91346 | 0.707576 | 2.704244 | 0.006845994 | 0.050881552 |
| *ENSG00000259915* | -3.869735 | 1.431064 | -2.704096 | 0.00684906 | 0.050881552 |
| *TOLLIP.AS1* | -2.858894 | 1.057404 | -2.70369 | 0.006857429 | 0.050894781 |
| *FGF14.AS2* | -3.245785 | 1.200531 | -2.703626 | 0.006858751 | 0.050894781 |
| *ELOVL5* | -0.907018 | 0.335544 | -2.703125 | 0.006869089 | 0.050931 |
| *SMOX* | -3.484356 | 1.289067 | -2.703006 | 0.006871549 | 0.050931 |
| *SMURF1* | -2.140038 | 0.791874 | -2.702498 | 0.006882064 | 0.050955648 |
| *WDR25* | -2.304931 | 0.852959 | -2.702277 | 0.006886637 | 0.050955648 |

| | | | | | |
|---|---|---|---|---|---|
| *GNMT* | -4.049505 | 1.498661 | -2.702082 | 0.006890674 | 0.050955648 |
| *ENSG00000282246* | -2.334154 | 0.863836 | -2.70208 | 0.006890715 | 0.050955648 |
| *ENSG00000272716* | -2.508432 | 0.928468 | -2.701689 | 0.006898819 | 0.050957865 |
| *ENSG00000263809* | -2.691184 | 0.996161 | -2.701554 | 0.006901618 | 0.050957865 |
| *TEAD2* | -3.996894 | 1.479753 | -2.701054 | 0.006912009 | 0.050957865 |
| *BDH1* | -1.805174 | 0.668346 | -2.700956 | 0.006914055 | 0.050957865 |
| *ENO1.AS1* | -2.748807 | 1.017769 | -2.700816 | 0.006916963 | 0.050957865 |
| *PRDM1* | -3.150423 | 1.166542 | -2.700651 | 0.006920388 | 0.050957865 |
| *MAPK11* | -2.5291 | 0.936572 | -2.700381 | 0.006926004 | 0.050957865 |
| *TMEM52* | -3.321424 | 1.230023 | -2.700294 | 0.006927821 | 0.050957865 |
| *DIPK1B* | -2.505207 | 0.927801 | -2.700156 | 0.006930704 | 0.050957865 |
| *C6orf132* | -4.132684 | 1.530701 | -2.699864 | 0.006936781 | 0.050957865 |
| *SLC26A6* | -1.967012 | 0.728567 | -2.699837 | 0.00693735 | 0.050957865 |
| *DFFB* | -2.061024 | 0.763404 | -2.69978 | 0.006938539 | 0.050957865 |
| *ENSG00000227741* | -4.116178 | 1.524934 | -2.69925 | 0.006949597 | 0.05100996 |
| *AMY2B* | -2.612278 | 0.967854 | -2.699041 | 0.006953964 | 0.051012917 |
| *CLDN12* | -2.291188 | 0.849526 | -2.697018 | 0.006996358 | 0.051284803 |
| *CCDC102A* | -2.324917 | 0.862148 | -2.696656 | 0.007003966 | 0.051284803 |
| *ZBED2* | -1.786821 | 0.662636 | -2.696534 | 0.007006529 | 0.051284803 |
| *HCG25* | -2.446146 | 0.907161 | -2.696484 | 0.007007579 | 0.051284803 |
| *PPL* | -2.596894 | 0.963124 | -2.696323 | 0.007010956 | 0.051284803 |
| *ZFP41* | -2.374694 | 0.880792 | -2.696088 | 0.007015915 | 0.051291917 |
| *GOLGA6L10* | -2.897638 | 1.074892 | -2.695748 | 0.007023075 | 0.051301898 |
| *ENSG00000228506* | -4.237163 | 1.571855 | -2.695645 | 0.007025254 | 0.051301898 |
| *SLC16A5* | -2.747713 | 1.019431 | -2.695339 | 0.007031711 | 0.051312838 |
| *ENSG00000259939* | -4.110396 | 1.525082 | -2.695196 | 0.007034728 | 0.051312838 |
| *ARSA* | -1.742578 | 0.646753 | -2.694348 | 0.007052652 | 0.051414434 |
| *ABHD2* | -1.775244 | 0.659016 | -2.693782 | 0.007064632 | 0.051462993 |
| *YTHDC2* | -1.507478 | 0.55964 | -2.693656 | 0.007067312 | 0.051462993 |
| *ANKRD6* | -2.500483 | 0.928532 | -2.692942 | 0.007082457 | 0.051544106 |
| *PUS7* | -2.690395 | 0.999211 | -2.692518 | 0.007091466 | 0.051580492 |
| *FBXO41* | -2.567383 | 0.953727 | -2.691949 | 0.007103586 | 0.05163946 |
| *ENSG00000249209* | -3.908829 | 1.452203 | -2.691654 | 0.007109875 | 0.051655989 |
| *KCNA3* | -2.896977 | 1.076729 | -2.690536 | 0.007133743 | 0.051800154 |
| *ENSG00000244198* | -4.152675 | 1.54365 | -2.690166 | 0.007141657 | 0.051809117 |
| *PI4K2A* | -2.291575 | 0.851854 | -2.690101 | 0.007143031 | 0.051809117 |
| *ZNF436.AS1* | -2.326595 | 0.865179 | -2.689149 | 0.007163437 | 0.05191955 |
| *KLRD1* | -3.022934 | 1.124212 | -2.688937 | 0.007168 | 0.05191955 |
| *YES1* | -3.346473 | 1.244692 | -2.688596 | 0.007175322 | 0.05191955 |
| *MMAA* | -2.541706 | 0.9456 | -2.687928 | 0.007189681 | 0.05191955 |
| *C15orf65* | -2.323236 | 0.864499 | -2.687379 | 0.007201528 | 0.05191955 |
| *JMJD1C.AS1* | -2.968914 | 1.104778 | -2.687341 | 0.007202348 | 0.05191955 |
| *RWDD2A* | -2.101565 | 0.782027 | -2.687331 | 0.007202544 | 0.05191955 |
| *GINS3* | -2.848612 | 1.060042 | -2.687263 | 0.007204012 | 0.05191955 |
| *ENSG00000272696* | -3.42289 | 1.273798 | -2.687153 | 0.007206399 | 0.05191955 |
| *CCDC121* | -3.89415 | 1.44925 | -2.68701 | 0.00720948 | 0.05191955 |
| *CLHC1* | -2.180559 | 0.811635 | -2.686627 | 0.007217757 | 0.05191955 |

| | | | | | |
|---|---|---|---|---|---|
| *PTCH2* | -2.920315 | 1.087012 | -2.686553 | 0.007219341 | 0.05191955 |
| *LY86.AS1* | -4.350818 | 1.619495 | -2.686528 | 0.007219891 | 0.05191955 |
| *ARMC2* | -2.376931 | 0.884794 | -2.686423 | 0.007222164 | 0.05191955 |
| *FAM230H* | -4.283478 | 1.594602 | -2.686237 | 0.007226181 | 0.05191955 |
| *GDPD3* | -2.659691 | 0.990167 | -2.686103 | 0.007229087 | 0.05191955 |
| *PRRG4* | -2.539675 | 0.945523 | -2.686 | 0.007231301 | 0.05191955 |
| *LUC7L3* | -0.598569 | 0.22286 | -2.685849 | 0.007234569 | 0.05191955 |
| *DACT3* | -2.644594 | 0.984702 | -2.68568 | 0.007238243 | 0.05191955 |
| *BORA* | -2.062971 | 0.768147 | -2.685647 | 0.007238958 | 0.05191955 |
| *ENSG00000228302* | -2.979523 | 1.109519 | -2.685418 | 0.007243916 | 0.051926165 |
| *L2HGDH* | -2.160384 | 0.804546 | -2.685221 | 0.007248197 | 0.051927921 |
| *ENC1* | -2.525034 | 0.940648 | -2.684355 | 0.007266987 | 0.052033569 |
| *PER3* | -3.112322 | 1.159774 | -2.683558 | 0.007284325 | 0.052128705 |
| *ENSG00000273367* | -4.182381 | 1.55885 | -2.682991 | 0.00729669 | 0.052169611 |
| *SFMBT2* | -3.319919 | 1.237426 | -2.682924 | 0.00729815 | 0.052169611 |
| *PDIA3* | -0.860728 | 0.320863 | -2.682538 | 0.007306579 | 0.052176127 |
| *MIAT* | -3.090878 | 1.152233 | -2.682511 | 0.007307172 | 0.052176127 |
| *STX1A* | -4.264129 | 1.589831 | -2.682128 | 0.007315553 | 0.052180183 |
| *HS3ST3B1* | -2.42552 | 0.904453 | -2.681752 | 0.007323767 | 0.052180183 |
| *IL12A* | -2.397993 | 0.894202 | -2.681714 | 0.007324594 | 0.052180183 |
| *CRYBG3* | -2.140778 | 0.798297 | -2.681681 | 0.007325336 | 0.052180183 |
| *TRPV3* | -3.034866 | 1.131829 | -2.681381 | 0.007331893 | 0.052180183 |
| *CLDN11* | -3.436798 | 1.281731 | -2.681373 | 0.007332072 | 0.052180183 |
| *PSMD5* | -1.575579 | 0.587693 | -2.680953 | 0.007341272 | 0.052216112 |
| *ENSG00000255121* | -2.658721 | 0.991865 | -2.680527 | 0.007350631 | 0.052216112 |
| *ENSG00000272644* | -3.871655 | 1.444395 | -2.680469 | 0.007351911 | 0.052216112 |
| *PERP* | -2.524494 | 0.941834 | -2.680403 | 0.007353353 | 0.052216112 |
| *ENSG00000262413* | -3.369403 | 1.257299 | -2.679875 | 0.007364974 | 0.052269787 |
| *GADD45G* | -3.554122 | 1.326403 | -2.679519 | 0.007372793 | 0.052296434 |
| *LNX1* | -2.183629 | 0.814999 | -2.679303 | 0.007377567 | 0.052301462 |
| *KLHL26* | -2.127421 | 0.794625 | -2.677263 | 0.007422636 | 0.052591989 |
| *AGFG2* | -2.525441 | 0.943385 | -2.677 | 0.007428455 | 0.052604257 |
| *ENSG00000251034* | -4.232461 | 1.581172 | -2.676787 | 0.007433194 | 0.052608859 |
| *ENSG00000272379* | -2.651422 | 0.990676 | -2.676375 | 0.007442324 | 0.052644519 |
| *POMGNT2* | -2.382008 | 0.890235 | -2.675707 | 0.007457185 | 0.052685955 |
| *ENSG00000267317* | -2.367901 | 0.885006 | -2.675576 | 0.007460106 | 0.052685955 |
| *MAPK12* | -4.070053 | 1.521197 | -2.675559 | 0.007460466 | 0.052685955 |
| *KCNAB3* | -3.237431 | 1.210116 | -2.675306 | 0.00746611 | 0.052696894 |
| *OXNAD1* | -2.039791 | 0.762513 | -2.675088 | 0.007470966 | 0.05270226 |
| *GSEC* | -3.79852 | 1.420184 | -2.674667 | 0.007480343 | 0.052739491 |
| *SLC43A1* | -2.817811 | 1.053619 | -2.674412 | 0.007486049 | 0.05274077 |
| *ZNF805* | -3.790265 | 1.417465 | -2.673974 | 0.007495833 | 0.05274077 |
| *ENSG00000258623* | -2.59361 | 0.969955 | -2.67395 | 0.007496367 | 0.05274077 |
| *CDCA3* | -4.064456 | 1.520034 | -2.673925 | 0.00749692 | 0.05274077 |
| *LINC01209* | -4.056297 | 1.517171 | -2.673592 | 0.007504371 | 0.052756407 |
| *HAAO* | -2.482477 | 0.928564 | -2.673459 | 0.007507343 | 0.052756407 |
| *LATS2.AS1* | -2.59097 | 0.969365 | -2.672852 | 0.007520942 | 0.052789293 |

| | | | | | |
|---|---|---|---|---|---|
| *ENSG00000232412* | -3.095001 | 1.158032 | -2.672638 | 0.007525737 | 0.052789293 |
| *BCORL1* | -2.302915 | 0.861713 | -2.672485 | 0.007529166 | 0.052789293 |
| *H2BFS* | -2.493688 | 0.933242 | -2.672071 | 0.007538465 | 0.052789293 |
| *ZNF219* | -2.306529 | 0.86321 | -2.672036 | 0.007539248 | 0.052789293 |
| *C20orf144* | -3.045872 | 1.139908 | -2.672032 | 0.007539346 | 0.052789293 |
| *PWP2* | -3.303416 | 1.236322 | -2.67197 | 0.007540742 | 0.052789293 |
| *ENSG00000260641* | -3.696671 | 1.3837 | -2.671583 | 0.007549433 | 0.052818632 |
| *CCDC85C* | -2.53457 | 0.948773 | -2.671418 | 0.007553142 | 0.052818632 |
| *EPHX2* | -3.431721 | 1.284759 | -2.671101 | 0.007560286 | 0.052839869 |
| *ROM1* | -2.559892 | 0.958518 | -2.670677 | 0.007569855 | 0.052878026 |
| *FRS3* | -2.308871 | 0.864621 | -2.670385 | 0.007576437 | 0.052895291 |
| *ATG9B* | -3.381813 | 1.266928 | -2.669302 | 0.007600903 | 0.05303732 |
| *ENSG00000228863* | -3.672922 | 1.376408 | -2.668484 | 0.007619444 | 0.053084988 |
| *MYOM1* | -2.677632 | 1.003446 | -2.668437 | 0.007620499 | 0.053084988 |
| *CNNM2* | -1.999635 | 0.749383 | -2.668375 | 0.007621918 | 0.053084988 |
| *CYB561* | -2.455254 | 0.920228 | -2.668092 | 0.007628337 | 0.053084988 |
| *RAD51* | -3.787149 | 1.419423 | -2.668091 | 0.007628363 | 0.053084988 |
| *ENSG00000238287* | -2.379847 | 0.892081 | -2.667748 | 0.007636155 | 0.053091087 |
| *TSPAN32* | -2.916409 | 1.093234 | -2.667689 | 0.007637491 | 0.053091087 |
| *ZNF79* | -2.312296 | 0.867023 | -2.666937 | 0.007654603 | 0.053181306 |
| *ACACB* | -2.1892 | 0.821183 | -2.665911 | 0.007678003 | 0.053315092 |
| *FBXO9* | -1.145931 | 0.4299 | -2.665575 | 0.007685672 | 0.053316585 |
| *STEAP4* | -2.915739 | 1.093865 | -2.665539 | 0.007686505 | 0.053316585 |
| *DARS2* | -2.089831 | 0.78418 | -2.66499 | 0.007699068 | 0.053365068 |
| *DGKH* | -3.164691 | 1.187559 | -2.664871 | 0.00770179 | 0.053365068 |
| *RFESD* | -2.644511 | 0.992545 | -2.664375 | 0.007713162 | 0.053415099 |
| *ENSG00000274104* | -2.361649 | 0.886521 | -2.66395 | 0.007722897 | 0.053436731 |
| *ZNF77* | -2.086476 | 0.783299 | -2.663703 | 0.007728581 | 0.053436731 |
| *LINC01023* | -3.253782 | 1.221529 | -2.663696 | 0.007728745 | 0.053436731 |
| *ADCK1* | -2.515952 | 0.944751 | -2.663084 | 0.007742813 | 0.053505249 |
| *ASAH2B* | -2.344101 | 0.880501 | -2.662236 | 0.007762336 | 0.053585173 |
| *DDX31* | -2.130156 | 0.800143 | -2.66222 | 0.007762708 | 0.053585173 |
| *TSNARE1* | -3.454332 | 1.297724 | -2.661839 | 0.007771501 | 0.053610664 |
| *MATR3* | -2.175025 | 0.817166 | -2.661669 | 0.007775435 | 0.053610664 |
| *ZNF512B* | -2.831081 | 1.063709 | -2.661519 | 0.0077789 | 0.053610664 |
| *TNRC6B* | -0.568096 | 0.213514 | -2.660696 | 0.007797941 | 0.053702197 |
| *ZFYVE28* | -2.723595 | 1.023683 | -2.660584 | 0.007800529 | 0.053702197 |
| *CPNE2* | -3.075126 | 1.155998 | -2.660149 | 0.007810604 | 0.053725097 |
| *PODXL2* | -2.459151 | 0.924563 | -2.659797 | 0.007818786 | 0.053725097 |
| *FAM227A* | -4.07145 | 1.530779 | -2.659725 | 0.007820458 | 0.053725097 |
| *CA2* | -3.062051 | 1.151268 | -2.65972 | 0.007820557 | 0.053725097 |
| *NEB* | -2.962883 | 1.114131 | -2.659367 | 0.007828762 | 0.053752764 |
| *SH2B1* | -1.297963 | 0.488181 | -2.658774 | 0.007842548 | 0.0538187 |
| *PSMD6.AS1* | -3.339999 | 1.25661 | -2.657945 | 0.007861877 | 0.053895822 |
| *C1orf198* | -2.448144 | 0.921071 | -2.657933 | 0.007862163 | 0.053895822 |
| *KIF11* | -3.827322 | 1.440126 | -2.65763 | 0.007869217 | 0.053915448 |
| *ANXA1* | -3.673819 | 1.382717 | -2.656957 | 0.007884945 | 0.053994456 |

| | | | | | |
|---|---|---|---|---|---|
| *PPP1R3F* | -3.745095 | 1.40987 | -2.656341 | 0.007899379 | 0.054064529 |
| *ALPK1* | -1.793884 | 0.675401 | -2.656028 | 0.00790671 | 0.054072884 |
| *MBTPS2* | -2.056169 | 0.774218 | -2.655802 | 0.007911997 | 0.054072884 |
| *ENSG00000258572* | -2.425017 | 0.913153 | -2.655652 | 0.00791553 | 0.054072884 |
| *CLDN23* | -4.044959 | 1.523197 | -2.655572 | 0.00791741 | 0.054072884 |
| *GHRLOS* | -3.903784 | 1.47024 | -2.655203 | 0.007926081 | 0.054103387 |
| *ENSG00000234665* | -3.28367 | 1.236966 | -2.654617 | 0.007939855 | 0.054143853 |
| *CLUH* | -2.427324 | 0.9144 | -2.654552 | 0.007941371 | 0.054143853 |
| *CCDC51* | -1.730673 | 0.651998 | -2.654414 | 0.007944633 | 0.054143853 |
| *TSC22D1.AS1* | -4.158289 | 1.566941 | -2.653762 | 0.007960003 | 0.054219881 |
| *IGKV1.5* | -2.789788 | 1.051457 | -2.653258 | 0.007971888 | 0.054272109 |
| *PABPN1* | -0.779402 | 0.293774 | -2.65307 | 0.007976328 | 0.054273619 |
| *TMEM44* | -4.12375 | 1.554597 | -2.652618 | 0.007987026 | 0.054317685 |
| *DARS.AS1* | -2.096765 | 0.790783 | -2.651505 | 0.008013392 | 0.054448745 |
| *TNFRSF4* | -3.812563 | 1.437953 | -2.651381 | 0.008016327 | 0.054448745 |
| *EXOC8* | -2.016082 | 0.760479 | -2.651069 | 0.008023753 | 0.054448745 |
| *CTPS2* | -2.346152 | 0.885026 | -2.65094 | 0.008026812 | 0.054448745 |
| *MORN3* | -2.407462 | 0.908163 | -2.650913 | 0.008027455 | 0.054448745 |
| *ENSG00000255237* | -3.423996 | 1.291789 | -2.650585 | 0.008035252 | 0.054472911 |
| *SEPTIN8* | -4.181623 | 1.578195 | -2.649624 | 0.008058137 | 0.054598207 |
| *YIPF5* | -1.076953 | 0.406518 | -2.649213 | 0.008067958 | 0.054598207 |
| *ENSG00000262151* | -3.975427 | 1.50062 | -2.649189 | 0.008068509 | 0.054598207 |
| *KLHDC8B* | -3.344692 | 1.262578 | -2.649097 | 0.008070707 | 0.054598207 |
| *MAP3K7CL* | -2.079194 | 0.784935 | -2.648874 | 0.00807604 | 0.054605572 |
| *TANC2* | -2.804465 | 1.059075 | -2.648034 | 0.008096149 | 0.05470089 |
| *LINC02604* | -2.973483 | 1.122946 | -2.64793 | 0.00809864 | 0.05470089 |
| *VPS9D1* | -2.509257 | 0.947887 | -2.647211 | 0.008115865 | 0.054788474 |
| *C9orf139* | -3.998113 | 1.510519 | -2.646847 | 0.00812462 | 0.054801777 |
| *EGR1* | -2.756945 | 1.041677 | -2.646641 | 0.008129562 | 0.054801777 |
| *FOCAD* | -3.21855 | 1.216229 | -2.646336 | 0.008136889 | 0.054801777 |
| *HIST1H2BH* | -2.271726 | 0.858456 | -2.646291 | 0.008137968 | 0.054801777 |
| *MSH5* | -2.388134 | 0.902462 | -2.646243 | 0.008139131 | 0.054801777 |
| *SOCS7* | -2.047875 | 0.773945 | -2.646021 | 0.008144481 | 0.054809119 |
| *C17orf67* | -1.978035 | 0.747602 | -2.645838 | 0.008148885 | 0.054810093 |
| *ZNF709* | -3.784006 | 1.430556 | -2.64513 | 0.00816596 | 0.054896242 |
| *ENSG00000269246* | -2.590177 | 0.97931 | -2.6449 | 0.00817151 | 0.054904867 |
| *MORC4* | -3.636321 | 1.375159 | -2.64429 | 0.008186242 | 0.054971132 |
| *IRAK1BP1* | -2.091168 | 0.790869 | -2.644138 | 0.008189917 | 0.054971132 |
| *PRMT6* | -1.734249 | 0.655963 | -2.643823 | 0.008197559 | 0.054993738 |
| *NRG2* | -4.03364 | 1.526078 | -2.643142 | 0.008214065 | 0.055075755 |
| *PSMD6.AS2* | -2.855858 | 1.080653 | -2.642716 | 0.0082244 | 0.055091737 |
| *ZNF658* | -2.385491 | 0.902675 | -2.642691 | 0.008225012 | 0.055091737 |
| *SEC22C* | -1.285853 | 0.486637 | -2.642322 | 0.008233979 | 0.055094441 |
| *ENSG00000272693* | -4.084819 | 1.54592 | -2.642322 | 0.008233964 | 0.055094441 |
| *ENSG00000255062* | -2.538534 | 0.960971 | -2.641636 | 0.008250668 | 0.055177412 |
| *AGPAT4* | -2.218324 | 0.839812 | -2.641453 | 0.008255127 | 0.055178555 |
| *CCDC153* | -3.118634 | 1.180785 | -2.641154 | 0.008262409 | 0.055198556 |

| | | | | | |
|---|---|---|---|---|---|
| *CAMK1* | -3.523402 | 1.334561 | -2.640121 | 0.008287637 | 0.055338363 |
| *BAIAP2* | -2.837961 | 1.075069 | -2.639794 | 0.008295641 | 0.055363079 |
| *ENSG00000238260* | -2.946988 | 1.116504 | -2.639479 | 0.008303361 | 0.055385871 |
| *KCNJ5* | -4.375737 | 1.658634 | -2.638156 | 0.008335812 | 0.055555709 |
| *GALNT14* | -2.498015 | 0.946903 | -2.638089 | 0.008337458 | 0.055555709 |
| *SELPLG* | -2.466298 | 0.934997 | -2.637761 | 0.008345549 | 0.055580836 |
| *CFAP54* | -3.20165 | 1.213929 | -2.637426 | 0.008353773 | 0.055606826 |
| *FMN1* | -4.330042 | 1.641909 | -2.6372 | 0.008359353 | 0.0556152 |
| *TCFL5* | -2.358655 | 0.894628 | -2.636463 | 0.008377532 | 0.055707339 |
| *ENSG00000262312* | -2.636291 | 1.000307 | -2.635482 | 0.008401777 | 0.055839701 |
| *ADNP* | -1.113101 | 0.422392 | -2.635235 | 0.008407902 | 0.055850597 |
| *SEC24B.AS1* | -2.221291 | 0.842974 | -2.635066 | 0.008412097 | 0.055850597 |
| *RIOX2* | -1.660203 | 0.630115 | -2.634762 | 0.008419623 | 0.055853534 |
| *TMSB15B* | -2.196211 | 0.833602 | -2.634605 | 0.008423514 | 0.055853534 |
| *C3orf18* | -3.234477 | 1.227774 | -2.634424 | 0.008428005 | 0.055853534 |
| *HOOK1* | -2.682645 | 1.018334 | -2.634348 | 0.008429903 | 0.055853534 |
| *SUFU* | -2.139797 | 0.812351 | -2.634079 | 0.008436582 | 0.055858924 |
| *ENSG00000275709* | -4.032011 | 1.530812 | -2.633903 | 0.008440957 | 0.055858924 |
| *DCBLD2* | -2.462275 | 0.934884 | -2.633776 | 0.008444119 | 0.055858924 |
| *ATP8B1* | -2.256048 | 0.856635 | -2.633617 | 0.008448082 | 0.055858924 |
| *AGBL3* | -2.429923 | 0.922718 | -2.63344 | 0.008452487 | 0.055859349 |
| *ENSG00000215068* | -2.878819 | 1.093252 | -2.633262 | 0.008456905 | 0.055859853 |
| *ENSG00000248787* | -4.148543 | 1.576077 | -2.632195 | 0.00848351 | 0.056006833 |
| *HS6ST1* | -2.545478 | 0.967206 | -2.631785 | 0.008493755 | 0.056045713 |
| *CCR7* | 0.4769 | 0.181225 | 2.631533 | 0.008500054 | 0.056058532 |
| *TEDC2* | -3.261565 | 1.23989 | -2.630527 | 0.008525263 | 0.056195984 |
| *CRACR2B* | -4.124536 | 1.568195 | -2.630117 | 0.008535547 | 0.056234963 |
| *GCK* | -3.677685 | 1.398532 | -2.629676 | 0.008546636 | 0.056279205 |
| *SH3BGR* | -3.786443 | 1.440042 | -2.629398 | 0.008553613 | 0.056288646 |
| *STOML1* | -2.276719 | 0.865913 | -2.629271 | 0.008556819 | 0.056288646 |
| *OXER1* | -3.547869 | 1.349676 | -2.628682 | 0.008571659 | 0.0563305 |
| *PAQR7* | -2.281453 | 0.867912 | -2.62867 | 0.008571937 | 0.0563305 |
| *ENSG00000260805* | -2.185041 | 0.831527 | -2.627744 | 0.008595307 | 0.056430825 |
| *MLXIPL* | -4.046651 | 1.539987 | -2.627718 | 0.008595976 | 0.056430825 |
| *TRIM24* | -1.99767 | 0.760493 | -2.62681 | 0.008618935 | 0.056510223 |
| *B4GAT1* | -1.999177 | 0.76109 | -2.62673 | 0.008620971 | 0.056510223 |
| *PEAK3* | -3.973157 | 1.512615 | -2.626682 | 0.008622193 | 0.056510223 |
| *ENSG00000261177* | -4.471762 | 1.70261 | -2.626415 | 0.008628945 | 0.056510223 |
| *DHRS11* | -2.205324 | 0.839684 | -2.626373 | 0.008630029 | 0.056510223 |
| *COL4A3* | -4.041357 | 1.539151 | -2.625705 | 0.00864697 | 0.056592355 |
| *ENSG00000260879* | -2.462569 | 0.938326 | -2.624427 | 0.008679485 | 0.056776272 |
| *DHX35* | -2.357416 | 0.898373 | -2.624095 | 0.008687951 | 0.056802774 |
| *ECHDC3* | -2.090524 | 0.796845 | -2.623501 | 0.008703125 | 0.056873089 |
| *IL11RA* | -2.111923 | 0.805122 | -2.623109 | 0.008713145 | 0.056909664 |
| *ZNF81* | -2.406731 | 0.917664 | -2.622671 | 0.008724349 | 0.056929748 |
| *PRNCR1* | -2.247285 | 0.856878 | -2.622643 | 0.008725069 | 0.056929748 |
| *ENSG00000270012* | -3.608365 | 1.376223 | -2.621934 | 0.008743236 | 0.056961958 |

| | | | | | |
|---|---|---|---|---|---|
| *ENSG00000261889* | -2.49537 | 0.951749 | -2.621877 | 0.008744697 | 0.056961958 |
| *ENSG00000257831* | -2.541027 | 0.96919 | -2.621804 | 0.008746577 | 0.056961958 |
| *AGAP5* | -2.326562 | 0.887405 | -2.621759 | 0.008747732 | 0.056961958 |
| *ENSG00000272186* | -2.989987 | 1.140526 | -2.621587 | 0.00875214 | 0.056961958 |
| *FKBP8* | -0.617441 | 0.235576 | -2.62098 | 0.008767741 | 0.057034643 |
| *ENSG00000177406* | -1.945167 | 0.742238 | -2.620677 | 0.008775534 | 0.057056493 |
| *PHF2* | -2.081342 | 0.794302 | -2.620342 | 0.008784174 | 0.05708382 |
| *NDUFB2.AS1* | -2.479621 | 0.94652 | -2.619723 | 0.008800113 | 0.057131443 |
| *NIPSNAP3B* | -1.928395 | 0.736109 | -2.619713 | 0.008800382 | 0.057131443 |
| *FAS* | -1.945689 | 0.743184 | -2.618045 | 0.008843513 | 0.057368136 |
| *SHLD3* | -1.999199 | 0.763658 | -2.617926 | 0.00884659 | 0.057368136 |
| *ZNF778* | -2.038969 | 0.77889 | -2.617786 | 0.008850218 | 0.057368136 |
| *TLCD1* | -2.825362 | 1.079457 | -2.617391 | 0.008860477 | 0.057378224 |
| *TUBG2* | -2.320254 | 0.886479 | -2.617383 | 0.008860693 | 0.057378224 |
| *ZNF618* | -3.06364 | 1.170838 | -2.616622 | 0.008880455 | 0.057477271 |
| *UGDH.AS1* | -2.941209 | 1.124449 | -2.61569 | 0.008904731 | 0.057605414 |
| *VLDLR* | -3.445856 | 1.317869 | -2.614719 | 0.008930095 | 0.057725358 |
| *VASH1.AS1* | -2.451082 | 0.937494 | -2.614505 | 0.008935698 | 0.057725358 |
| *FBF1* | -2.08723 | 0.798339 | -2.614465 | 0.008936731 | 0.057725358 |
| *TTC34* | -2.273214 | 0.869541 | -2.614269 | 0.008941856 | 0.057729483 |
| *EOGT* | -3.158406 | 1.208758 | -2.612934 | 0.008976872 | 0.057926486 |
| *ENSG00000214770* | -2.036152 | 0.779417 | -2.612403 | 0.008990828 | 0.057987463 |
| *IER5L* | -3.227445 | 1.235544 | -2.612166 | 0.008997046 | 0.057998494 |
| *LINC01535* | -3.19293 | 1.222741 | -2.611289 | 0.009020168 | 0.058082381 |
| *SLC48A1* | -1.526799 | 0.584706 | -2.611224 | 0.009021866 | 0.058082381 |
| *FKBP1B* | -1.999374 | 0.765704 | -2.611159 | 0.009023601 | 0.058082381 |
| *ENSG00000271741* | -2.790994 | 1.068952 | -2.610962 | 0.009028797 | 0.058086763 |
| *NOL6* | -2.518088 | 0.964913 | -2.609654 | 0.009063383 | 0.058280135 |
| *ZNF558* | -1.91703 | 0.734853 | -2.608727 | 0.009087975 | 0.058388519 |
| *HTRA4* | -5.029205 | 1.927876 | -2.608676 | 0.00908932 | 0.058388519 |
| *DAGLB* | -1.727897 | 0.662485 | -2.608205 | 0.009101843 | 0.058388519 |
| *RTKN* | -1.919493 | 0.735975 | -2.608097 | 0.009104719 | 0.058388519 |
| *ENSG00000272335* | -2.377759 | 0.911684 | -2.608096 | 0.00910475 | 0.058388519 |
| *SMG8* | -3.507973 | 1.345085 | -2.607994 | 0.009107465 | 0.058388519 |
| *ITGA11* | -4.737151 | 1.816594 | -2.607711 | 0.009114999 | 0.058407718 |
| *SRR* | -2.166708 | 0.830954 | -2.607495 | 0.009120729 | 0.058415339 |
| *PUS10* | -2.035986 | 0.780873 | -2.60732 | 0.009125391 | 0.058416125 |
| *ZMYND10* | -3.24438 | 1.244418 | -2.607147 | 0.009130017 | 0.058416673 |
| *LINC02062* | -3.569915 | 1.369505 | -2.60672 | 0.009141415 | 0.058438542 |
| *ENSG00000277476* | -2.673821 | 1.025758 | -2.606678 | 0.009142518 | 0.058438542 |
| *FAM71F2* | -3.308587 | 1.269478 | -2.606257 | 0.009153766 | 0.058481381 |
| *MEX3A* | -3.899752 | 1.497026 | -2.604999 | 0.009187459 | 0.058667512 |
| *ENSG00000260267* | -2.213884 | 0.850168 | -2.604055 | 0.009212805 | 0.058768349 |
| *MRPS6* | 0.734447 | 0.282046 | 2.604 | 0.009214271 | 0.058768349 |
| *ENSG00000274184* | -1.998089 | 0.767362 | -2.603841 | 0.009218548 | 0.058768349 |
| *ENSG00000282951* | -3.41716 | 1.31246 | -2.603629 | 0.009224263 | 0.058768349 |
| *TUNAR* | -2.932626 | 1.126408 | -2.60352 | 0.009227193 | 0.058768349 |

| | | | | | |
|---|---|---|---|---|---|
| *ENSG00000246528* | -2.372495 | 0.91131 | -2.603391 | 0.009230655 | 0.058768349 |
| *MCPH1.AS1* | -3.818388 | 1.467124 | -2.602635 | 0.00925104 | 0.058840575 |
| *ERMN* | -2.98046 | 1.145172 | -2.602631 | 0.009251145 | 0.058840575 |
| *ENSG00000272008* | -4.015116 | 1.543115 | -2.601956 | 0.009269379 | 0.058927421 |
| *BRPF3* | -2.067033 | 0.794508 | -2.601653 | 0.009277576 | 0.058950408 |
| *LRFN1* | -2.673798 | 1.027824 | -2.601415 | 0.009283994 | 0.058962069 |
| *HOXC4* | -3.577424 | 1.375422 | -2.600965 | 0.009296187 | 0.059010377 |
| *TWSG1* | -2.285875 | 0.879361 | -2.599473 | 0.009336712 | 0.059231298 |
| *ENSG00000224610* | -3.730368 | 1.435119 | -2.599345 | 0.009340196 | 0.059231298 |
| *CYB5D1* | -3.065592 | 1.179526 | -2.599004 | 0.009349457 | 0.059260818 |
| *KCNQ5* | -2.689711 | 1.035042 | -2.59865 | 0.009359121 | 0.059292864 |
| *NEK10* | -4.01952 | 1.546887 | -2.598458 | 0.009364358 | 0.059296845 |
| *LINC02649* | -2.611137 | 1.005025 | -2.598082 | 0.009374609 | 0.059332559 |
| *WDR48* | -1.283437 | 0.494065 | -2.597709 | 0.009384792 | 0.059367807 |
| *TMEM231* | -3.33865 | 1.285631 | -2.596896 | 0.009407051 | 0.059453463 |
| *MEGF11* | -2.379374 | 0.916244 | -2.596877 | 0.009407574 | 0.059453463 |
| *TMCC1.AS1* | -2.969956 | 1.144182 | -2.595702 | 0.009439778 | 0.059549034 |
| *LRTOMT* | -1.907526 | 0.734918 | -2.595562 | 0.009443644 | 0.059549034 |
| *ZW10* | -1.939447 | 0.747239 | -2.595483 | 0.009445814 | 0.059549034 |
| *CBWD6* | -2.33762 | 0.900661 | -2.595448 | 0.009446771 | 0.059549034 |
| *EPHX1* | -2.681255 | 1.033123 | -2.595292 | 0.009451055 | 0.059549034 |
| *RERE.AS1* | -4.026487 | 1.551523 | -2.595184 | 0.009454034 | 0.059549034 |
| *TRMT5* | -2.019623 | 0.778231 | -2.595145 | 0.009455093 | 0.059549034 |
| *GGA2* | 1.215692 | 0.468571 | 2.594469 | 0.009473724 | 0.05957058 |
| *LINC02576* | -2.747224 | 1.0589 | -2.594413 | 0.00947526 | 0.05957058 |
| *THRA* | -1.868744 | 0.720303 | -2.594385 | 0.009476034 | 0.05957058 |
| *MACC1* | -2.534604 | 0.976971 | -2.594349 | 0.009477033 | 0.05957058 |
| *ENSG00000226571* | -1.899516 | 0.732434 | -2.59343 | 0.009502384 | 0.05970077 |
| *ENSG00000267136* | -3.985565 | 1.537007 | -2.593069 | 0.009512377 | 0.059734383 |
| *ENSG00000233038* | -2.514266 | 0.969868 | -2.592381 | 0.009531425 | 0.059773242 |
| *THNSL2* | -2.855029 | 1.101329 | -2.592348 | 0.009532329 | 0.059773242 |
| *CENPF* | -2.454517 | 0.946834 | -2.592342 | 0.009532501 | 0.059773242 |
| *ZMYND19* | -2.228459 | 0.859865 | -2.591638 | 0.009552018 | 0.059866449 |
| *RECK* | -1.993482 | 0.769429 | -2.590859 | 0.00957366 | 0.059972873 |
| *HMGB3* | -2.25299 | 0.86971 | -2.590506 | 0.009583488 | 0.060005226 |
| *ANKEF1* | -3.489029 | 1.347152 | -2.589929 | 0.009599576 | 0.060056012 |
| *ENSG00000251417* | -2.635333 | 1.017558 | -2.58986 | 0.009601492 | 0.060056012 |
| *GPM6A* | -2.301059 | 0.888538 | -2.589713 | 0.009605601 | 0.060056012 |
| *DDN.AS1* | -2.25899 | 0.872453 | -2.589239 | 0.009618835 | 0.060081494 |
| *SNED1* | -3.517556 | 1.358733 | -2.588851 | 0.00962968 | 0.060081494 |
| *P2RY11* | -2.285571 | 0.882934 | -2.588609 | 0.009636435 | 0.060081494 |
| *SUV39H1* | -2.084353 | 0.805223 | -2.588542 | 0.009638325 | 0.060081494 |
| *LINC01869* | -1.819084 | 0.702777 | -2.588422 | 0.00964168 | 0.060081494 |
| *PCF11.AS1* | -2.102905 | 0.812444 | -2.58837 | 0.009643146 | 0.060081494 |
| *PLEKHM3* | -3.903367 | 1.508131 | -2.588215 | 0.009647483 | 0.060081494 |
| *HECTD2* | -2.264234 | 0.87486 | -2.58811 | 0.009650425 | 0.060081494 |
| *ITIH5* | -3.859126 | 1.491209 | -2.587917 | 0.009655837 | 0.060081494 |

| | | | | | |
|---|---|---|---|---|---|
| *ITGA6.AS1* | -2.040847 | 0.788612 | -2.587897 | 0.009656371 | 0.060081494 |
| *ARHGAP44* | -2.587348 | 0.999893 | -2.587625 | 0.009664015 | 0.060099992 |
| *TM6SF1* | -3.146084 | 1.216129 | -2.586966 | 0.009682519 | 0.060185975 |
| *SYP* | -2.017419 | 0.780137 | -2.585981 | 0.009710226 | 0.060329059 |
| *FAM114A1* | -2.310389 | 0.893584 | -2.585531 | 0.009722909 | 0.060378701 |
| *KIAA0513* | -2.941513 | 1.137823 | -2.585211 | 0.009731945 | 0.060405661 |
| *PRRT3.AS1* | -4.51287 | 1.745849 | -2.584915 | 0.009740311 | 0.060428437 |
| *MBLAC1* | -2.821731 | 1.092103 | -2.583759 | 0.009773018 | 0.06049933 |
| *ENSG00000273384* | -2.654585 | 1.027422 | -2.583733 | 0.009773729 | 0.06049933 |
| *IQCN* | -2.046889 | 0.792325 | -2.583396 | 0.009783291 | 0.06049933 |
| *ENSG00000278107* | -4.008627 | 1.551696 | -2.583385 | 0.009783604 | 0.06049933 |
| *SCFD2* | -3.056396 | 1.183131 | -2.583311 | 0.00978571 | 0.06049933 |
| *BBS12* | -2.413687 | 0.93437 | -2.583225 | 0.009788157 | 0.06049933 |
| *ZNF8* | -2.056518 | 0.796105 | -2.583225 | 0.00978816 | 0.06049933 |
| *RNF32* | -2.583285 | 1.00004 | -2.583182 | 0.009789353 | 0.06049933 |
| *ENSG00000270871* | -4.137997 | 1.602018 | -2.58299 | 0.009794823 | 0.060504077 |
| *LARS2* | -2.938557 | 1.1378 | -2.582666 | 0.00980402 | 0.060531827 |
| *ENSG00000256591* | -2.357036 | 0.912732 | -2.582396 | 0.009811706 | 0.060550227 |
| *REM2* | -2.180508 | 0.844686 | -2.581443 | 0.009838826 | 0.060686885 |
| *ENSG00000254893* | -3.819535 | 1.479702 | -2.581287 | 0.009843284 | 0.060686885 |
| *PYROXD2* | -2.620448 | 1.01525 | -2.581087 | 0.009848974 | 0.060692885 |
| *ENSG00000253140* | -4.1138 | 1.594139 | -2.580577 | 0.009863534 | 0.060745347 |
| *BSCL2* | -2.778597 | 1.076784 | -2.580458 | 0.009866929 | 0.060745347 |
| *BICDL1* | -2.3431 | 0.908123 | -2.580157 | 0.009875542 | 0.060763352 |
| *ATP11A* | -1.917119 | 0.743062 | -2.580026 | 0.009879298 | 0.060763352 |
| *TVP23A* | -4.606143 | 1.785558 | -2.579666 | 0.009889585 | 0.060769829 |
| *ENSG00000257839* | -2.276017 | 0.882294 | -2.579659 | 0.009889797 | 0.060769829 |
| *ZNF596* | -2.280586 | 0.884182 | -2.579318 | 0.009899547 | 0.060800702 |
| *MUC20* | -2.563788 | 0.994127 | -2.578935 | 0.009910547 | 0.060839222 |
| *L3MBTL1* | -2.032656 | 0.788264 | -2.578649 | 0.009918749 | 0.060860536 |
| *PXDC1* | -2.247855 | 0.871858 | -2.578237 | 0.009930591 | 0.060895997 |
| *EXTL2* | -1.977308 | 0.766958 | -2.578118 | 0.009933994 | 0.060895997 |
| *CCDC32* | 1.695457 | 0.657721 | 2.577773 | 0.00994392 | 0.060903764 |
| *LINC01238* | -3.200801 | 1.241838 | -2.577471 | 0.009952633 | 0.060903764 |
| *GPSM1* | -3.927082 | 1.523642 | -2.577431 | 0.009953786 | 0.060903764 |
| *TNNT3* | -3.607384 | 1.399621 | -2.5774 | 0.009954681 | 0.060903764 |
| *SETBP1* | -1.577627 | 0.61217 | -2.577105 | 0.009963161 | 0.060903764 |
| *ENSG00000259065* | -3.922038 | 1.521887 | -2.577088 | 0.009963661 | 0.060903764 |
| *ENSG00000266490* | -2.60987 | 1.012959 | -2.576481 | 0.009981169 | 0.060981815 |
| *ENSG00000262873* | -4.279857 | 1.661259 | -2.576273 | 0.009987185 | 0.06098961 |
| *CATSPERG* | -2.44744 | 0.950165 | -2.575806 | 0.010000676 | 0.061017012 |
| *PPM1L* | -2.879284 | 1.117826 | -2.575789 | 0.010001156 | 0.061017012 |
| *FGF22* | -2.510528 | 0.974887 | -2.575199 | 0.010018257 | 0.061092377 |
| *GLI1* | -2.294114 | 0.890957 | -2.574888 | 0.01002726 | 0.061118313 |
| *ENSG00000230454* | -3.831677 | 1.488213 | -2.574683 | 0.010033192 | 0.061119068 |
| *ENSG00000277369* | -3.932848 | 1.527704 | -2.574353 | 0.010042783 | 0.061119068 |
| *FSIP2* | -2.115373 | 0.821866 | -2.573866 | 0.010056933 | 0.061119068 |

| | | | | | |
|---|---|---|---|---|---|
| *ZNF853* | -3.832563 | 1.48904 | -2.573849 | 0.01005741 | 0.061119068 |
| *PRMT2* | -0.737032 | 0.286376 | -2.573646 | 0.010063324 | 0.061119068 |
| *CDC14B* | -2.401956 | 0.933384 | -2.573385 | 0.010070914 | 0.061119068 |
| *PRH1* | -2.960371 | 1.150388 | -2.573368 | 0.010071401 | 0.061119068 |
| *ZNF839* | -2.364108 | 0.918727 | -2.573243 | 0.010075035 | 0.061119068 |
| *GPD2* | -2.084962 | 0.810251 | -2.573232 | 0.010075378 | 0.061119068 |
| *FAM66C* | -2.584995 | 1.004598 | -2.573163 | 0.010077383 | 0.061119068 |
| *ITGAX* | -2.899602 | 1.126926 | -2.57302 | 0.010081528 | 0.061119068 |
| *SPEF2* | -2.461024 | 0.956509 | -2.572922 | 0.010084385 | 0.061119068 |
| *VPS37C* | -1.861059 | 0.723437 | -2.572523 | 0.010096027 | 0.061160817 |
| *CDC37L1.DT* | -2.24952 | 0.87461 | -2.572027 | 0.010110492 | 0.061219623 |
| *VRK3* | 1.819368 | 0.707634 | 2.571057 | 0.010138871 | 0.061362586 |
| *FAM122C* | -1.95658 | 0.761337 | -2.569926 | 0.010172033 | 0.061512264 |
| *DISC1* | -3.888432 | 1.513075 | -2.569887 | 0.010173164 | 0.061512264 |
| *ENSG00000267811* | -4.071313 | 1.584354 | -2.5697 | 0.010178668 | 0.061516639 |
| *ENSG00000278238* | -2.365382 | 0.920594 | -2.56941 | 0.010187191 | 0.061539241 |
| *NQO1* | -2.734116 | 1.064756 | -2.567833 | 0.010233637 | 0.061790805 |
| *LYSMD1* | -2.199689 | 0.856713 | -2.567591 | 0.010240791 | 0.061805 |
| *ZKSCAN3* | -2.19686 | 0.8558 | -2.567025 | 0.010257515 | 0.061876907 |
| *ENSG00000237773* | -1.902298 | 0.741212 | -2.566469 | 0.010273972 | 0.06193538 |
| *CERS6* | -2.025826 | 0.7894 | -2.566285 | 0.010279423 | 0.06193538 |
| *PLXNA1* | -2.223723 | 0.866594 | -2.566047 | 0.010286474 | 0.06193538 |
| *MFAP3* | -2.417183 | 0.94205 | -2.565875 | 0.010291586 | 0.06193538 |
| *LETM1* | -1.286872 | 0.501543 | -2.565827 | 0.010293013 | 0.06193538 |
| *SVOP* | -3.623308 | 1.412222 | -2.565679 | 0.010297421 | 0.06193538 |
| *FIZ1* | -2.365406 | 0.921984 | -2.565561 | 0.010300902 | 0.06193538 |
| *C9orf163* | -3.883125 | 1.513676 | -2.565362 | 0.010306829 | 0.061942067 |
| *RAB35* | -1.42387 | 0.555149 | -2.564843 | 0.010322256 | 0.0619919 |
| *TNF* | -2.266509 | 0.883733 | -2.5647 | 0.010326512 | 0.0619919 |
| *ENSG00000277007* | -2.323186 | 0.905878 | -2.564567 | 0.010330453 | 0.0619919 |
| *THUMPD1* | -1.10305 | 0.430174 | -2.564198 | 0.010341468 | 0.0619919 |
| *TRAIP* | -2.517436 | 0.981768 | -2.564187 | 0.010341783 | 0.0619919 |
| *ENSG00000275457* | -2.028625 | 0.791283 | -2.563715 | 0.010355839 | 0.0619919 |
| *SPINDOC* | -2.168688 | 0.845938 | -2.563649 | 0.01035782 | 0.0619919 |
| *NAV1* | -2.479754 | 0.967348 | -2.563456 | 0.010363572 | 0.0619919 |
| *ALG10* | -2.246756 | 0.876473 | -2.563407 | 0.010365051 | 0.0619919 |
| *C6orf52* | -3.990712 | 1.556835 | -2.563348 | 0.010366798 | 0.0619919 |
| *ENSG00000267939* | -3.756653 | 1.465551 | -2.563304 | 0.010368118 | 0.0619919 |
| *B3GNT7* | -1.679794 | 0.655506 | -2.56259 | 0.010389452 | 0.062090611 |
| *PPP2CB* | -1.795735 | 0.7008 | -2.56241 | 0.010394867 | 0.062094126 |
| *C8orf58* | -1.827991 | 0.713495 | -2.562022 | 0.01040646 | 0.062134533 |
| *SLC25A12* | -1.973351 | 0.770412 | -2.561423 | 0.010424424 | 0.062212926 |
| *ENSG00000229729* | -2.035301 | 0.794695 | -2.561111 | 0.010433815 | 0.062240102 |
| *JUNB* | 0.872324 | 0.340712 | 2.560293 | 0.01045838 | 0.062357726 |
| *IPO11* | -1.990996 | 0.777803 | -2.55977 | 0.010474144 | 0.062404134 |
| *ENSG00000225205* | -2.255066 | 0.880984 | -2.559713 | 0.010475863 | 0.062404134 |
| *PPP1R9A* | -4.070383 | 1.590308 | -2.559495 | 0.010482451 | 0.062414485 |

| | | | | | |
|---|---|---|---|---|---|
| *SERP2* | -2.635966 | 1.029999 | -2.559193 | 0.010491534 | 0.06243967 |
| *ENSG00000268352* | -2.771673 | 1.083237 | -2.558696 | 0.010506565 | 0.06250022 |
| *PIGV* | -1.764412 | 0.689849 | -2.557679 | 0.010537325 | 0.062636907 |
| *DSEL* | -3.541186 | 1.384566 | -2.557615 | 0.010539279 | 0.062636907 |
| *B4GALT2* | -4.025378 | 1.574159 | -2.557161 | 0.010553045 | 0.062689766 |
| *ARMCX2* | -1.678805 | 0.656659 | -2.556584 | 0.010570549 | 0.062722303 |
| *ENSG00000273156* | -2.451317 | 0.958831 | -2.556568 | 0.010571055 | 0.062722303 |
| *KANK1* | -2.715596 | 1.062232 | -2.556499 | 0.010573146 | 0.062722303 |
| *CPEB2* | -2.267609 | 0.887122 | -2.556141 | 0.010584024 | 0.062757899 |
| *ENSG00000215014* | -2.000258 | 0.782969 | -2.55471 | 0.010627631 | 0.062987437 |
| *C10orf25* | -2.639064 | 1.033099 | -2.554512 | 0.010633678 | 0.062994261 |
| *ENSG00000187186* | -2.385993 | 0.934196 | -2.554059 | 0.010647508 | 0.063024642 |
| *MMP17* | -2.164478 | 0.847503 | -2.553948 | 0.010650931 | 0.063024642 |
| *IFITM3* | -2.327317 | 0.911293 | -2.553864 | 0.010653501 | 0.063024642 |
| *NR1I3* | -3.349636 | 1.311785 | -2.553494 | 0.010664801 | 0.063062498 |
| *BACH1.AS1* | -2.644511 | 1.035921 | -2.552811 | 0.010685744 | 0.06315731 |
| *AREG* | -3.806045 | 1.491217 | -2.552308 | 0.01070118 | 0.063207231 |
| *EMILIN2* | -2.171181 | 0.850704 | -2.552216 | 0.010704015 | 0.063207231 |
| *ENSG00000267260* | -3.567137 | 1.398003 | -2.551594 | 0.010723137 | 0.063274636 |
| *SMAD2* | -1.095929 | 0.429519 | -2.551525 | 0.010725265 | 0.063274636 |
| *ASPHD2* | -2.2569 | 0.884611 | -2.551292 | 0.010732449 | 0.063288001 |
| *TLR4* | -5.086459 | 1.993928 | -2.550974 | 0.010742243 | 0.063291857 |
| *OTUD7B* | -3.364107 | 1.318911 | -2.55067 | 0.010751601 | 0.063291857 |
| *ATP13A4* | -3.721872 | 1.459183 | -2.550655 | 0.010752084 | 0.063291857 |
| *ENSG00000275055* | -2.476494 | 0.970934 | -2.550632 | 0.010752778 | 0.063291857 |
| *CABP4* | -3.590612 | 1.407866 | -2.550392 | 0.010760173 | 0.063306424 |
| *ENSG00000273010* | -3.637135 | 1.426394 | -2.54988 | 0.010775988 | 0.063370492 |
| *ENSG00000258376* | -1.832873 | 0.718924 | -2.549466 | 0.010788793 | 0.063416812 |
| *ENSG00000273149* | -3.175403 | 1.245789 | -2.548908 | 0.010806073 | 0.063481032 |
| *FUT2* | -2.977597 | 1.168294 | -2.548671 | 0.010813434 | 0.063481032 |
| *ENSG00000272572* | -2.580404 | 1.012465 | -2.548636 | 0.01081452 | 0.063481032 |
| *RPS6KB2.AS1* | -2.077861 | 0.815386 | -2.548317 | 0.010824413 | 0.063482652 |
| *ENSG00000272356* | -2.504851 | 0.98296 | -2.548272 | 0.010825793 | 0.063482652 |
| *TTLL7* | -2.454219 | 0.963138 | -2.54815 | 0.010829597 | 0.063482652 |
| *NDRG2* | -2.765398 | 1.085575 | -2.547404 | 0.010852756 | 0.063589443 |
| *SLFN12* | -2.187563 | 0.858926 | -2.546859 | 0.010869723 | 0.063641631 |
| *ENSG00000226578* | -3.637253 | 1.428166 | -2.546801 | 0.010871555 | 0.063641631 |
| *CCDC154* | -1.969336 | 0.773372 | -2.546426 | 0.010883216 | 0.063651946 |
| *PDIA5* | -2.224388 | 0.873617 | -2.546181 | 0.010890863 | 0.063651946 |
| *IGHMBP2* | -1.914241 | 0.751841 | -2.546071 | 0.010894304 | 0.063651946 |
| *TXLNB* | -2.61538 | 1.027246 | -2.546011 | 0.010896188 | 0.063651946 |
| *ZNF324B* | -2.34962 | 0.922885 | -2.545951 | 0.010898052 | 0.063651946 |
| *ENSG00000259802* | -3.890316 | 1.528641 | -2.544951 | 0.010929305 | 0.063805522 |
| *MRPS24* | -1.936353 | 0.761 | -2.544485 | 0.010943906 | 0.063861789 |
| *URB2* | -2.276436 | 0.894768 | -2.544163 | 0.010953996 | 0.06389169 |
| *CDAN1* | -2.035048 | 0.800074 | -2.543574 | 0.010972493 | 0.063970578 |
| *LINC01431* | -2.174928 | 0.855259 | -2.543006 | 0.01099033 | 0.064015016 |

| | | | | | |
|---|---|---|---|---|---|
| *FAM86C1* | -2.447718 | 0.962565 | -2.542912 | 0.010993306 | 0.064015016 |
| *ZNF425* | -3.839724 | 1.510004 | -2.542856 | 0.010995041 | 0.064015016 |
| *TNFRSF9* | -4.090476 | 1.608866 | -2.542459 | 0.011007537 | 0.064048669 |
| *CPT1B* | -3.433845 | 1.350654 | -2.542357 | 0.011010776 | 0.064048669 |
| *LSR* | -1.618965 | 0.636839 | -2.54219 | 0.011016036 | 0.06405031 |
| *CHEK2* | -1.95266 | 0.768309 | -2.541504 | 0.011037667 | 0.064147089 |
| *ENSG00000269983* | -2.313012 | 0.910308 | -2.54091 | 0.01105644 | 0.064222995 |
| *ZNF517* | -3.082186 | 1.213089 | -2.540775 | 0.01106071 | 0.064222995 |
| *FBXO4* | -1.470595 | 0.578888 | -2.54038 | 0.011073219 | 0.064266628 |
| *AQP11* | -4.003223 | 1.576245 | -2.539722 | 0.011094074 | 0.064358634 |
| *HCN3* | -2.546221 | 1.002793 | -2.539129 | 0.011112881 | 0.064438682 |
| *ENSG00000229255* | -5.056356 | 1.991667 | -2.538756 | 0.011124739 | 0.064458728 |
| *CTH* | -2.869587 | 1.130388 | -2.538586 | 0.011130159 | 0.064458728 |
| *CCDC61* | -3.399895 | 1.339307 | -2.538548 | 0.011131367 | 0.064458728 |
| *AGBL2* | -2.964594 | 1.167976 | -2.538232 | 0.011141397 | 0.064487788 |
| *SYCP3* | -2.033726 | 0.801786 | -2.536493 | 0.011196896 | 0.064779886 |
| *AFDN* | -2.275893 | 0.897339 | -2.53627 | 0.011204033 | 0.064783502 |
| *ADHFE1* | -3.120857 | 1.230545 | -2.536159 | 0.011207591 | 0.064783502 |
| *ENSG00000257497* | -2.16292 | 0.852888 | -2.535995 | 0.011212847 | 0.064784777 |
| *DEGS2* | -2.880123 | 1.135845 | -2.535666 | 0.011223358 | 0.064816404 |
| *SUSD1* | -2.273611 | 0.896829 | -2.535165 | 0.011239425 | 0.064847611 |
| *FBXL19* | -2.244959 | 0.885617 | -2.534909 | 0.01124764 | 0.064847611 |
| *ENSG00000267383* | -2.277509 | 0.898463 | -2.534896 | 0.011248077 | 0.064847611 |
| *ENSG00000273486* | -2.516727 | 0.992843 | -2.53487 | 0.011248921 | 0.064847611 |
| *KCTD3* | -2.71072 | 1.069476 | -2.534625 | 0.011256776 | 0.06486383 |
| *TMEM273* | -2.012485 | 0.794098 | -2.534304 | 0.011267106 | 0.064894293 |
| *RELCH* | -1.749155 | 0.690479 | -2.533249 | 0.011301053 | 0.065060691 |
| *ENSG00000251602* | -3.479311 | 1.373697 | -2.532808 | 0.011315292 | 0.065113536 |
| *SEPTIN1* | -0.785617 | 0.310202 | -2.532599 | 0.011322028 | 0.065123169 |
| *CAMSAP1* | -1.876602 | 0.741535 | -2.530699 | 0.011383536 | 0.065356246 |
| *MYZAP* | -3.071657 | 1.213763 | -2.53069 | 0.011383841 | 0.065356246 |
| *ENSG00000260507* | -2.275933 | 0.899436 | -2.530399 | 0.011393285 | 0.065356246 |
| *LINC02550* | -5.100469 | 2.015768 | -2.530286 | 0.011396971 | 0.065356246 |
| *SLX1A* | -2.285564 | 0.903305 | -2.530225 | 0.011398946 | 0.065356246 |
| *ZFHX2* | -2.262924 | 0.894402 | -2.530099 | 0.011403039 | 0.065356246 |
| *FKBP14* | -2.14011 | 0.845895 | -2.529996 | 0.011406393 | 0.065356246 |
| *PPP1R13B* | -2.329315 | 0.920693 | -2.529959 | 0.011407583 | 0.065356246 |
| *FKBPL* | -1.726598 | 0.682467 | -2.529938 | 0.011408264 | 0.065356246 |
| *ZNF653* | -2.65873 | 1.051214 | -2.529198 | 0.011432338 | 0.065465017 |
| *ENSG00000266498* | -3.834237 | 1.516378 | -2.528549 | 0.011453511 | 0.065557085 |
| *ZNF174* | -1.855997 | 0.734308 | -2.527545 | 0.011486306 | 0.065687631 |
| *PUDP* | -1.794642 | 0.710036 | -2.527538 | 0.011486529 | 0.065687631 |
| *ENSG00000259274* | -2.653833 | 1.050843 | -2.525433 | 0.011555576 | 0.066053129 |
| *IGLV1.47* | -3.328799 | 1.318218 | -2.525227 | 0.011562358 | 0.066053168 |
| *CD68* | -1.560586 | 0.618024 | -2.525121 | 0.01156585 | 0.066053168 |
| *PDK4* | -2.393249 | 0.947872 | -2.524865 | 0.011574291 | 0.066072048 |
| *ENSG00000272256* | -3.422419 | 1.355919 | -2.524057 | 0.011600899 | 0.066154553 |

| | | | | | |
|---|---|---|---|---|---|
| *SAMD9L* | 1.77257 | 0.702332 | 2.523835 | 0.011608239 | 0.066154553 |
| *CARD16* | -0.995712 | 0.394526 | -2.523821 | 0.011608691 | 0.066154553 |
| *SMTN* | -3.291107 | 1.304027 | -2.523803 | 0.011609309 | 0.066154553 |
| *TOMM40L* | -2.194476 | 0.869588 | -2.523583 | 0.011616561 | 0.06616511 |
| *ENSG00000242539* | -2.563487 | 1.015876 | -2.523425 | 0.011621769 | 0.06616511 |
| *TENT4A* | -2.025925 | 0.802893 | -2.52328 | 0.011626589 | 0.06616511 |
| *ZNF460* | -2.289792 | 0.907533 | -2.523096 | 0.011632671 | 0.066170461 |
| *TMCC3* | -2.446538 | 0.969894 | -2.522481 | 0.011653034 | 0.066256999 |
| *PEX11A* | -2.29943 | 0.911735 | -2.522039 | 0.011667691 | 0.066281254 |
| *C15orf62* | -1.566225 | 0.62106 | -2.521856 | 0.011673741 | 0.066281254 |
| *SLC35E4* | -2.505281 | 0.993482 | -2.521716 | 0.011678396 | 0.066281254 |
| *ENSG00000265008* | -3.319298 | 1.316291 | -2.521704 | 0.011678778 | 0.066281254 |
| *NKILA* | -2.505533 | 0.993638 | -2.521576 | 0.011683056 | 0.066281254 |
| *DGKQ* | -1.999992 | 0.793247 | -2.521272 | 0.011693146 | 0.066309258 |
| *USP27X* | -2.404139 | 0.953601 | -2.521116 | 0.011698329 | 0.066309427 |
| *ENSG00000197813* | -3.182981 | 1.262733 | -2.520708 | 0.011711887 | 0.066352472 |
| *YTHDF3.AS1* | -3.858829 | 1.530938 | -2.520565 | 0.011716652 | 0.066352472 |
| *ENSG00000276651* | -2.26313 | 0.897917 | -2.520423 | 0.011721393 | 0.066352472 |
| *CACNA2D4* | -3.682236 | 1.461095 | -2.520188 | 0.011729201 | 0.066354878 |
| *RBM41* | -1.262502 | 0.500973 | -2.520101 | 0.011732132 | 0.066354878 |
| *NADK2* | -1.859695 | 0.738069 | -2.519677 | 0.01174626 | 0.06640559 |
| *RORA* | -2.274577 | 0.902815 | -2.519427 | 0.011754617 | 0.066423652 |
| *IGHV3.15* | -2.368712 | 0.940347 | -2.518976 | 0.01176966 | 0.066469859 |
| *RBM15.AS1* | -3.766077 | 1.495144 | -2.518873 | 0.011773126 | 0.066469859 |
| *SLC26A4* | -3.783976 | 1.502667 | -2.518174 | 0.01179652 | 0.066558543 |
| *MEGF9* | -1.810831 | 0.719128 | -2.518094 | 0.011799179 | 0.066558543 |
| *ENSG00000269148* | -3.047794 | 1.210626 | -2.517535 | 0.011817923 | 0.066635064 |
| *LINC01943* | -3.934978 | 1.563142 | -2.517352 | 0.011824068 | 0.066640512 |
| *RND2* | -3.502695 | 1.391603 | -2.517021 | 0.011835175 | 0.066673905 |
| *BTBD3* | -2.256055 | 0.896653 | -2.516085 | 0.011866667 | 0.066822059 |
| *SLC22A5* | -1.997706 | 0.794112 | -2.515647 | 0.011881422 | 0.06687588 |
| *SLC11A1* | -3.139352 | 1.248124 | -2.515256 | 0.011894602 | 0.066920789 |
| *ZNF816.ZNF321P* | -2.287529 | 0.909626 | -2.514803 | 0.011909889 | 0.066963896 |
| *DDX47* | -2.616202 | 1.040355 | -2.514721 | 0.011912673 | 0.066963896 |
| *GRASP* | -2.266231 | 0.901849 | -2.512872 | 0.011975278 | 0.067277052 |
| *FITM2* | -2.633176 | 1.047919 | -2.512767 | 0.011978839 | 0.067277052 |
| *IL4I1* | -4.969003 | 1.97782 | -2.512363 | 0.011992563 | 0.067310449 |
| *CLTCL1* | -3.747475 | 1.491661 | -2.512284 | 0.011995248 | 0.067310449 |
| *CRIM1* | -1.943236 | 0.773683 | -2.511671 | 0.01201611 | 0.067378845 |
| *ENSG00000279927* | -3.007928 | 1.197653 | -2.511519 | 0.012021285 | 0.067378845 |
| *ENSG00000280240* | -3.33784 | 1.329041 | -2.511464 | 0.012023147 | 0.067378845 |
| *APOBR* | -3.069865 | 1.222551 | -2.511033 | 0.012037859 | 0.067431924 |
| *NFE2L3* | -3.744309 | 1.491645 | -2.510188 | 0.012066694 | 0.067564033 |
| *OIP5* | -2.342853 | 0.933438 | -2.509919 | 0.012075886 | 0.067586091 |
| *ZNF579* | -2.366698 | 0.943062 | -2.50959 | 0.012087139 | 0.067607321 |
| *SMCO4* | -1.898444 | 0.75655 | -2.509344 | 0.012095571 | 0.067607321 |
| *MYH11* | -1.887968 | 0.752388 | -2.509301 | 0.012097028 | 0.067607321 |

| | | | | | |
|---|---|---|---|---|---|
| *ENSG00000245970* | -2.328405 | 0.927949 | -2.509194 | 0.012100696 | 0.067607321 |
| *TNKS2.AS1* | -2.733107 | 1.089374 | -2.508879 | 0.012111511 | 0.067634482 |
| *ENSG00000273145* | -3.45875 | 1.378677 | -2.508746 | 0.012116071 | 0.067634482 |
| *AIFM2* | -1.837595 | 0.732694 | -2.507998 | 0.012141741 | 0.067748389 |
| *ENSG00000269696* | -2.246151 | 0.895672 | -2.507783 | 0.012149132 | 0.067752352 |
| *HSPA1A* | -2.621526 | 1.045403 | -2.507671 | 0.012152983 | 0.067752352 |
| *PAX8* | -3.02902 | 1.208056 | -2.507351 | 0.012163978 | 0.06778428 |
| *LINGO3* | -2.033368 | 0.81112 | -2.506865 | 0.012180725 | 0.067848222 |
| *CARD6* | -2.167536 | 0.864832 | -2.506308 | 0.012199933 | 0.067925808 |
| *USP54* | -2.351434 | 0.938374 | -2.50586 | 0.012215391 | 0.067936267 |
| *ENSG00000272112* | -2.412649 | 0.962808 | -2.505847 | 0.012215836 | 0.067936267 |
| *ENSG00000224790* | -2.922289 | 1.166252 | -2.505711 | 0.012220563 | 0.067936267 |
| *SLC31A2* | -2.235121 | 0.892035 | -2.505642 | 0.012222931 | 0.067936267 |
| *GNLY* | -3.741856 | 1.493711 | -2.505074 | 0.012242583 | 0.068016113 |
| *ZCCHC14* | -2.527232 | 1.009249 | -2.504072 | 0.012277299 | 0.068151033 |
| *ENSG00000272822* | -4.778621 | 1.908344 | -2.504067 | 0.012277461 | 0.068151033 |
| *UNC13D* | -1.305464 | 0.521482 | -2.503375 | 0.012301517 | 0.068247826 |
| *MFSD14A* | -1.210028 | 0.483381 | -2.50326 | 0.012305507 | 0.068247826 |
| *C1orf112* | -2.127021 | 0.850339 | -2.501378 | 0.012371096 | 0.068582031 |
| *TMTC2* | -3.895758 | 1.557643 | -2.50106 | 0.012382228 | 0.068614181 |
| *IKZF4* | -2.283521 | 0.913096 | -2.500857 | 0.012389335 | 0.068624012 |
| *ENSG00000261248* | -2.785008 | 1.113814 | -2.500426 | 0.01240442 | 0.068678002 |
| *ZC3H15* | 1.329052 | 0.531597 | 2.500109 | 0.012415506 | 0.068709812 |
| *TMEM88* | -5.124481 | 2.050885 | -2.498668 | 0.012466106 | 0.068960183 |
| *ZNF696* | -2.180011 | 0.872601 | -2.498292 | 0.012479338 | 0.069003716 |
| *WDR11* | 1.74664 | 0.699426 | 2.497249 | 0.012516111 | 0.069177322 |
| *RUNDC3B* | -3.158855 | 1.265072 | -2.496976 | 0.012525739 | 0.069200807 |
| *FBP1* | -2.011771 | 0.805761 | -2.496735 | 0.01253425 | 0.069218107 |
| *WDHD1* | -2.146152 | 0.859722 | -2.496333 | 0.012548466 | 0.069266884 |
| *ENSG00000080031* | -2.534389 | 1.015404 | -2.495941 | 0.012562333 | 0.069301214 |
| *ATP8B2* | -2.232713 | 0.894595 | -2.495782 | 0.012567988 | 0.069301214 |
| *MYT1L* | -3.474084 | 1.392031 | -2.495694 | 0.012571085 | 0.069301214 |
| *DUSP5* | -2.425862 | 0.972075 | -2.495549 | 0.012576229 | 0.069301214 |
| *KRT8* | -2.695693 | 1.080346 | -2.495213 | 0.012588164 | 0.069304671 |
| *RABGAP1L* | 1.076536 | 0.431446 | 2.495181 | 0.012589297 | 0.069304671 |
| *TSHZ2* | -4.808351 | 1.927223 | -2.494964 | 0.012596993 | 0.069304671 |
| *C19orf66* | -0.741544 | 0.297221 | -2.494924 | 0.012598401 | 0.069304671 |
| *ENSG00000263731* | -2.608118 | 1.045579 | -2.494425 | 0.012616137 | 0.06937258 |
| *PDIK1L* | -2.0186 | 0.809621 | -2.493266 | 0.012657389 | 0.06956968 |
| *ANKRD46* | -1.433156 | 0.574921 | -2.49279 | 0.012674373 | 0.069607272 |
| *CASK* | -2.206338 | 0.885148 | -2.49262 | 0.012680463 | 0.069607272 |
| *ZNF69* | -2.263852 | 0.908254 | -2.492531 | 0.012683627 | 0.069607272 |
| *ALG11* | -1.81563 | 0.728446 | -2.492468 | 0.012685867 | 0.069607272 |
| *CXCR3* | -3.879484 | 1.556715 | -2.492096 | 0.012699165 | 0.069630496 |
| *INCENP* | -3.242538 | 1.301154 | -2.492047 | 0.012700923 | 0.069630496 |
| *ENSG00000269549* | -2.935113 | 1.178168 | -2.491251 | 0.012729402 | 0.069724801 |
| *SLC45A4* | -3.286167 | 1.319141 | -2.491141 | 0.012733359 | 0.069724801 |

| | | | | | |
|---|---|---|---|---|---|
| *FYN* | -1.858213 | 0.745937 | -2.491112 | 0.012734381 | 0.069724801 |
| *TUSC1* | -2.892223 | 1.161313 | -2.490476 | 0.012757208 | 0.069820076 |
| *ENSG00000256448* | -3.109076 | 1.248658 | -2.489935 | 0.012776654 | 0.069896771 |
| *ARSB* | -2.158408 | 0.867058 | -2.489347 | 0.0127978 | 0.06995866 |
| *ENSG00000228857* | -2.3411 | 0.940458 | -2.489318 | 0.012798841 | 0.06995866 |
| *ENSG00000273729* | -2.780754 | 1.11747 | -2.488437 | 0.012830587 | 0.070015682 |
| *KCTD21* | -2.409759 | 0.968389 | -2.48842 | 0.012831205 | 0.070015682 |
| *DCXR.DT* | -3.008566 | 1.209089 | -2.488291 | 0.012835854 | 0.070015682 |
| *ZNF48* | -1.464042 | 0.588411 | -2.488127 | 0.012841777 | 0.070015682 |
| *FKBP1C* | -4.952173 | 1.99041 | -2.488017 | 0.012845763 | 0.070015682 |
| *SETDB2* | -1.028244 | 0.41328 | -2.488007 | 0.01284611 | 0.070015682 |
| *SLC25A23* | -2.042603 | 0.820998 | -2.487953 | 0.01284808 | 0.070015682 |
| *ENSG00000250397* | -3.811635 | 1.532117 | -2.487822 | 0.012852805 | 0.070015682 |
| *LINC02285* | -2.479783 | 0.996833 | -2.487662 | 0.012858585 | 0.070017523 |
| *LEPR* | -3.53839 | 1.422835 | -2.486859 | 0.012887636 | 0.070146027 |
| *ENSG00000055483* | -1.626138 | 0.653965 | -2.486582 | 0.012897694 | 0.070171088 |
| *ENSG00000272010* | -3.62507 | 1.45852 | -2.485444 | 0.012938981 | 0.070349488 |
| *DRAXIN* | -4.723058 | 1.900405 | -2.48529 | 0.012944582 | 0.070349488 |
| *ITFG2.AS1* | -1.696706 | 0.682717 | -2.485227 | 0.012946886 | 0.070349488 |
| *SLC9A1* | -2.264698 | 0.911387 | -2.484891 | 0.012959126 | 0.070370518 |
| *HCST* | -1.628698 | 0.655459 | -2.48482 | 0.012961695 | 0.070370518 |
| *ENSG00000274987* | -3.831961 | 1.542284 | -2.484602 | 0.012969624 | 0.070383871 |
| *PACSIN3* | -3.190613 | 1.284457 | -2.484016 | 0.012991007 | 0.070470186 |
| *CYFIP1* | -3.803128 | 1.53158 | -2.48314 | 0.013022985 | 0.070613886 |
| *NRBP2* | -2.29492 | 0.924267 | -2.482962 | 0.013029484 | 0.070619365 |
| *BCKDHA* | -2.11926 | 0.853589 | -2.482763 | 0.013036785 | 0.070629182 |
| *MARS2* | -2.047123 | 0.82461 | -2.482535 | 0.013045118 | 0.070634197 |
| *ENSG00000255882* | -3.4097 | 1.373529 | -2.482438 | 0.013048689 | 0.070634197 |
| *SYAP1* | 1.772749 | 0.714169 | 2.482255 | 0.01305538 | 0.070640694 |
| *PARS2* | -1.916273 | 0.772071 | -2.481989 | 0.01306514 | 0.070663789 |
| *VWA8* | -1.953241 | 0.787076 | -2.481644 | 0.013077803 | 0.070701818 |
| *GAB2* | -2.021215 | 0.814514 | -2.481498 | 0.013083161 | 0.070701818 |
| *ENSG00000247363* | -2.227033 | 0.897598 | -2.481104 | 0.013097609 | 0.070727786 |
| *DENND2C* | -3.327856 | 1.341327 | -2.481018 | 0.01310078 | 0.070727786 |
| *ENSG00000231609* | -2.274825 | 0.916939 | -2.480889 | 0.013105519 | 0.070727786 |
| *TTLL12* | -2.10648 | 0.849124 | -2.480768 | 0.013109953 | 0.070727786 |
| *C1RL* | -1.957019 | 0.789074 | -2.480147 | 0.013132829 | 0.070821505 |
| *CACYBP* | 0.97769 | 0.394245 | 2.479902 | 0.01314186 | 0.07082248 |
| *COLCA1* | -2.247348 | 0.906258 | -2.479811 | 0.013145218 | 0.07082248 |
| *EARS2* | -2.594431 | 1.046289 | -2.479649 | 0.013151169 | 0.07082248 |
| *MOXD1* | -2.364053 | 0.953422 | -2.479545 | 0.013155027 | 0.07082248 |
| *ATXN7L1* | -1.771803 | 0.71486 | -2.47853 | 0.013192508 | 0.070983173 |
| *CHTF18* | -1.774288 | 0.71589 | -2.478438 | 0.013195908 | 0.070983173 |
| *ENSG00000277825* | -2.663435 | 1.074806 | -2.478061 | 0.01320984 | 0.07102842 |
| *MXD3* | -2.110688 | 0.851877 | -2.477691 | 0.013223553 | 0.071068697 |
| *HHAT* | -2.01108 | 0.811718 | -2.477561 | 0.013228377 | 0.071068697 |
| *ARHGAP20* | -5.10185 | 2.059503 | -2.477223 | 0.013240905 | 0.071076647 |

| | | | | | |
|---|---|---|---|---|---|
| *ZBTB32* | -5.10185 | 2.059503 | -2.477223 | 0.013240905 | 0.071076647 |
| *GTF3A* | 0.665503 | 0.268692 | 2.476821 | 0.013255829 | 0.071097515 |
| *ZNF792* | -2.629657 | 1.061707 | -2.476821 | 0.013255844 | 0.071097515 |
| *PHETA2* | -3.163399 | 1.277347 | -2.476538 | 0.013266353 | 0.071124236 |
| *ENSG00000235652* | -3.313868 | 1.338474 | -2.475855 | 0.013291743 | 0.071202618 |
| *ENSG00000254501* | -3.064306 | 1.23768 | -2.475847 | 0.013292041 | 0.071202618 |
| *GLIPR1* | -0.727409 | 0.293913 | -2.474909 | 0.013326999 | 0.071307958 |
| *LINC02018* | -2.003581 | 0.80959 | -2.474811 | 0.013330686 | 0.071307958 |
| *CKAP4* | -3.185736 | 1.287265 | -2.474809 | 0.013330743 | 0.071307958 |
| *SLC1A5* | -2.061966 | 0.83321 | -2.474725 | 0.013333873 | 0.071307958 |
| *DUSP6* | -2.602407 | 1.051709 | -2.474456 | 0.01334391 | 0.071308777 |
| *SCYL3* | -1.55934 | 0.630202 | -2.474347 | 0.013347982 | 0.071308777 |
| *INKA2* | -1.904898 | 0.769881 | -2.474276 | 0.013350652 | 0.071308777 |
| *HIST1H2AK* | -3.720245 | 1.503764 | -2.473956 | 0.013362601 | 0.071342981 |
| *BHLHE41* | -1.832426 | 0.740857 | -2.473386 | 0.013383965 | 0.071427405 |
| *NR4A1* | -2.248856 | 0.909328 | -2.473097 | 0.013394791 | 0.071455545 |
| *ENSG00000256940* | -2.373025 | 0.959657 | -2.472784 | 0.013406519 | 0.071488469 |
| *NAIF1* | -1.753104 | 0.709144 | -2.472143 | 0.013430588 | 0.071587148 |
| *ENSG00000260735* | -3.201026 | 1.294989 | -2.471856 | 0.01344135 | 0.071614847 |
| *MFSD4A* | -2.166245 | 0.876691 | -2.470934 | 0.01347608 | 0.071747005 |
| *ZBTB37* | -1.435201 | 0.580841 | -2.470901 | 0.013477307 | 0.071747005 |
| *NOCT* | -2.137561 | 0.865188 | -2.470633 | 0.013487433 | 0.07175958 |
| *DTX3* | -2.253901 | 0.91231 | -2.470543 | 0.013490823 | 0.07175958 |
| *ENSG00000242622* | -3.423054 | 1.385641 | -2.470376 | 0.013497094 | 0.071763268 |
| *PBLD* | -3.025616 | 1.225193 | -2.469502 | 0.013530141 | 0.071891299 |
| *IGHEP1* | -2.897897 | 1.173502 | -2.469443 | 0.013532348 | 0.071891299 |
| *RHOB* | -2.116531 | 0.857168 | -2.469214 | 0.013541017 | 0.071907663 |
| *LRRC8C.DT* | -2.49999 | 1.012785 | -2.468431 | 0.013570693 | 0.072035525 |
| *CDYL* | -1.70719 | 0.691653 | -2.468276 | 0.013576563 | 0.072036964 |
| *GIMAP1* | -3.143488 | 1.273728 | -2.467943 | 0.013589199 | 0.072074286 |
| *TMEM191C* | -1.63102 | 0.660964 | -2.467637 | 0.013600804 | 0.072106116 |
| *SNTB1* | -4.910057 | 1.990605 | -2.466616 | 0.013639665 | 0.072242913 |
| *ZNF718* | -2.387126 | 0.967787 | -2.46658 | 0.013641007 | 0.072242913 |
| *ERP27* | -2.641059 | 1.070765 | -2.466516 | 0.013643451 | 0.072242913 |
| *SAMD10* | -1.937079 | 0.785473 | -2.466129 | 0.013658212 | 0.072264111 |
| *ENSG00000272906* | -2.074565 | 0.841227 | -2.466117 | 0.013658686 | 0.072264111 |
| *ENSG00000253636* | -3.599656 | 1.460283 | -2.46504 | 0.013699781 | 0.072451742 |
| *TIMELESS* | -1.547423 | 0.627849 | -2.464642 | 0.013715031 | 0.072502589 |
| *C19orf54* | -2.326882 | 0.944182 | -2.464442 | 0.013722663 | 0.072513145 |
| *OTUD3* | -1.932548 | 0.78424 | -2.46423 | 0.013730791 | 0.072526307 |
| *ENSG00000227373* | -5.092837 | 2.068641 | -2.461924 | 0.013819411 | 0.072964448 |
| *GINS1* | -3.626419 | 1.473326 | -2.461383 | 0.013840257 | 0.072981394 |
| *ENSG00000246982* | -4.853658 | 1.971948 | -2.461353 | 0.01384143 | 0.072981394 |
| *ENSG00000272787* | -4.086476 | 1.660274 | -2.461326 | 0.013842439 | 0.072981394 |
| *CDKL3* | -2.083091 | 0.846354 | -2.461252 | 0.013845308 | 0.072981394 |
| *ENSG00000248932* | -3.822336 | 1.553286 | -2.460806 | 0.013862519 | 0.073031631 |
| *AGBL5* | -1.762934 | 0.716435 | -2.460705 | 0.013866449 | 0.073031631 |

| | | | | | |
|---|---|---|---|---|---|
| *VPS33B* | -1.740412 | 0.707322 | -2.460564 | 0.013871867 | 0.073031631 |
| *GRAP2* | -2.494747 | 1.014144 | -2.459953 | 0.013895509 | 0.073123346 |
| *ESAM* | -2.214069 | 0.900094 | -2.45982 | 0.013900653 | 0.073123346 |
| *TBC1D10B* | -1.746466 | 0.710149 | -2.459295 | 0.013921 | 0.073166998 |
| *EXTL3* | -2.041973 | 0.830328 | -2.459236 | 0.013923287 | 0.073166998 |
| *NUDT6* | -2.695246 | 1.096 | -2.459166 | 0.013926011 | 0.073166998 |
| *TVP23C* | -2.282099 | 0.928496 | -2.457846 | 0.013977313 | 0.073377461 |
| *IVNS1ABP* | -1.444187 | 0.587583 | -2.457842 | 0.013977474 | 0.073377461 |
| *ECE2* | -2.07266 | 0.843445 | -2.457373 | 0.013995722 | 0.073424837 |
| *COQ10A* | -1.48031 | 0.60241 | -2.457312 | 0.013998105 | 0.073424837 |
| *CTPS1* | -2.454854 | 0.999112 | -2.457037 | 0.014008843 | 0.073424837 |
| *ENSG00000262728* | -2.958721 | 1.204189 | -2.457024 | 0.014009324 | 0.073424837 |
| *PRR34* | -2.971312 | 1.20951 | -2.456624 | 0.014024925 | 0.073464112 |
| *GNGT2* | -1.747481 | 0.711363 | -2.456526 | 0.014028771 | 0.073464112 |
| *ENSG00000179094* | -2.107995 | 0.85825 | -2.456156 | 0.01404321 | 0.073464112 |
| *BTBD19* | -2.879211 | 1.172266 | -2.456107 | 0.014045138 | 0.073464112 |
| *TTC4* | -2.100202 | 0.855096 | -2.456101 | 0.014045365 | 0.073464112 |
| *ENSG00000247796* | -2.682085 | 1.09254 | -2.454907 | 0.014092101 | 0.073678611 |
| *DDR1* | -2.267503 | 0.92396 | -2.454113 | 0.014123263 | 0.073811546 |
| *ZNF283* | -1.81706 | 0.74055 | -2.453663 | 0.014140928 | 0.073873862 |
| *CUBN* | -2.125092 | 0.866146 | -2.453503 | 0.01414723 | 0.073876788 |
| *CABYR* | -2.56698 | 1.04658 | -2.452731 | 0.014177624 | 0.074005469 |
| *TNIK* | -3.436221 | 1.401421 | -2.451955 | 0.014208255 | 0.074135286 |
| *RSPH9* | -3.753701 | 1.531728 | -2.450631 | 0.014260616 | 0.07435381 |
| *ENSG00000263220* | -2.096774 | 0.855615 | -2.450604 | 0.014261693 | 0.07435381 |
| *GH1* | -3.550637 | 1.449182 | -2.450098 | 0.014281736 | 0.074378107 |
| *ENSG00000268400* | -3.077938 | 1.256261 | -2.450079 | 0.014282468 | 0.074378107 |
| *NAP1L6* | -4.49742 | 1.835645 | -2.450049 | 0.014283695 | 0.074378107 |
| *TTC13* | -1.65413 | 0.675474 | -2.448842 | 0.014331624 | 0.074587387 |
| *BACH1.IT2* | -3.500036 | 1.429318 | -2.448745 | 0.014335479 | 0.074587387 |
| *CA11* | -1.301119 | 0.531424 | -2.448364 | 0.014350642 | 0.074615491 |
| *ENSG00000273669* | -2.923223 | 1.193972 | -2.448318 | 0.014352479 | 0.074615491 |
| *ZBED4* | -1.722709 | 0.703753 | -2.447889 | 0.014369596 | 0.074651357 |
| *SVIL.AS1* | -2.492383 | 1.018191 | -2.447854 | 0.014370981 | 0.074651357 |
| *GGACT* | -2.390504 | 0.977067 | -2.446613 | 0.014420569 | 0.074873872 |
| *TRMT13* | -0.976683 | 0.399218 | -2.446491 | 0.014425455 | 0.074873872 |
| *GIHCG* | -1.13635 | 0.464638 | -2.445669 | 0.014458365 | 0.074985235 |
| *CFAP70* | -2.885025 | 1.179687 | -2.445586 | 0.014461703 | 0.074985235 |
| *IMPDH1* | -1.299031 | 0.531188 | -2.445519 | 0.014464394 | 0.074985235 |
| *GDI2* | 0.675992 | 0.276443 | 2.445319 | 0.014472414 | 0.074996597 |
| *SLC41A1* | -1.774865 | 0.725997 | -2.444728 | 0.01449613 | 0.075021886 |
| *PCLAF* | -3.044107 | 1.245172 | -2.444728 | 0.014496134 | 0.075021886 |
| *LINC01011* | -2.106746 | 0.861806 | -2.44457 | 0.014502474 | 0.075021886 |
| *LINC01551* | -2.163102 | 0.884885 | -2.444501 | 0.014505272 | 0.075021886 |
| *ERMP1* | -3.806769 | 1.557297 | -2.444472 | 0.014506447 | 0.075021886 |
| *IPPK* | -1.952846 | 0.79904 | -2.443991 | 0.014525789 | 0.075086826 |
| *GZMB* | -5.085357 | 2.080898 | -2.443828 | 0.014532346 | 0.075086826 |

| | | | | | |
|---|---|---|---|---|---|
| *ENSG00000244036* | -2.067543 | 0.846132 | -2.443524 | 0.014544593 | 0.075086826 |
| *ENSG00000261786* | -3.908967 | 1.599773 | -2.44345 | 0.014547567 | 0.075086826 |
| *CCR9* | -2.289461 | 0.936985 | -2.443435 | 0.014548182 | 0.075086826 |
| *PLGLB1* | -3.049665 | 1.248291 | -2.443071 | 0.014562872 | 0.075132507 |
| *EEF1AKMT1* | -1.72132 | 0.70469 | -2.442663 | 0.014579348 | 0.075187362 |
| *SPECC1* | -2.022294 | 0.827977 | -2.442453 | 0.014587826 | 0.075200942 |
| *TMEM140* | -1.447052 | 0.592579 | -2.441955 | 0.014607982 | 0.075231574 |
| *FRMD6.AS1* | -2.620896 | 1.073299 | -2.441906 | 0.014609968 | 0.075231574 |
| *GCM1* | -4.910006 | 2.010754 | -2.441872 | 0.014611308 | 0.075231574 |
| *SGSH* | -1.950378 | 0.798879 | -2.441395 | 0.014630653 | 0.075291765 |
| *FNBP1* | 0.705009 | 0.288785 | 2.441295 | 0.014634702 | 0.075291765 |
| *AGO2* | -1.009924 | 0.413733 | -2.441004 | 0.014646482 | 0.075322258 |
| *TMEM86B* | -3.672815 | 1.504941 | -2.440503 | 0.014666815 | 0.075396688 |
| *IGLC5* | -3.266568 | 1.338694 | -2.440115 | 0.014682596 | 0.075399214 |
| *GATD3B* | -2.211938 | 0.906532 | -2.44 | 0.014687275 | 0.075399214 |
| *ADAP1* | -2.925382 | 1.198949 | -2.439955 | 0.014689074 | 0.075399214 |
| *ZNF398* | -1.808464 | 0.7412 | -2.439914 | 0.014690746 | 0.075399214 |
| *ITSN1* | -3.076966 | 1.261208 | -2.439697 | 0.01469957 | 0.07541442 |
| *ZNF594* | -1.687939 | 0.692014 | -2.439171 | 0.014721008 | 0.075494306 |
| *RELL2* | -1.728305 | 0.708668 | -2.438809 | 0.014735755 | 0.075539827 |
| *USP46.AS1* | -4.909056 | 2.013213 | -2.438418 | 0.014751705 | 0.075561382 |
| *MMP14* | -4.909056 | 2.013213 | -2.438418 | 0.014751705 | 0.075561382 |
| *CHIT1* | -2.663528 | 1.092738 | -2.437481 | 0.014789978 | 0.075653345 |
| *TACC1* | -1.204657 | 0.494226 | -2.437462 | 0.014790787 | 0.075653345 |
| *ACOT7* | -3.555824 | 1.458853 | -2.437412 | 0.014792827 | 0.075653345 |
| *GALNT6* | -3.527535 | 1.44726 | -2.437389 | 0.014793753 | 0.075653345 |
| *ENSG00000227540* | -1.974752 | 0.810235 | -2.437259 | 0.014799057 | 0.075653345 |
| *BCYRN1* | -3.436839 | 1.410312 | -2.436935 | 0.014812352 | 0.075691237 |
| *STXBP4* | -2.121141 | 0.870659 | -2.436247 | 0.014840538 | 0.075805162 |
| *RASGRP1* | -1.8269 | 0.749937 | -2.436072 | 0.014847729 | 0.075811799 |
| *SLC4A8* | -3.109197 | 1.276616 | -2.435499 | 0.014871278 | 0.075901915 |
| *HIST1H2AG* | -1.906062 | 0.782729 | -2.435151 | 0.01488559 | 0.075944841 |
| *MOCS2* | -1.110313 | 0.456011 | -2.434836 | 0.014898522 | 0.075963459 |
| *ZNF442* | -2.171451 | 0.891864 | -2.434733 | 0.014902787 | 0.075963459 |
| *NBPF14* | -1.010559 | 0.415077 | -2.434632 | 0.014906951 | 0.075963459 |
| *LINC01754* | -2.225908 | 0.914381 | -2.434333 | 0.014919243 | 0.075986282 |
| *ZKSCAN8* | -1.597205 | 0.656142 | -2.434237 | 0.014923241 | 0.075986282 |
| *PRDM10* | -3.477991 | 1.429009 | -2.433848 | 0.014939278 | 0.076009191 |
| *ENSG00000260588* | -2.582411 | 1.061044 | -2.433841 | 0.014939554 | 0.076009191 |
| *C11orf98* | -1.842322 | 0.757098 | -2.433402 | 0.014957703 | 0.076049893 |
| *HGH1* | -2.147314 | 0.882448 | -2.433361 | 0.014959375 | 0.076049893 |
| *SIDT2* | -1.340416 | 0.550886 | -2.433199 | 0.014966085 | 0.076053955 |
| *MAP1LC3A* | -1.888425 | 0.776391 | -2.432312 | 0.015002771 | 0.076210286 |
| *NECTIN3* | -3.853994 | 1.585033 | -2.431491 | 0.015036812 | 0.076311443 |
| *DONSON* | -2.459017 | 1.011346 | -2.431428 | 0.015039418 | 0.076311443 |
| *ENSG00000270659* | -2.049729 | 0.843023 | -2.431403 | 0.015040477 | 0.076311443 |
| *ENSG00000271870* | -1.914466 | 0.787749 | -2.430299 | 0.015086386 | 0.076514201 |

| | | | | | |
|---|---|---|---|---|---|
| *PIGN* | -1.906803 | 0.784653 | -2.430122 | 0.01509376 | 0.076521438 |
| *HLA.DMA* | 0.636911 | 0.262147 | 2.429595 | 0.015115718 | 0.076602576 |
| *STARD5* | -1.722835 | 0.70926 | -2.429061 | 0.015138003 | 0.076685309 |
| *MON1A* | -2.460064 | 1.012891 | -2.428754 | 0.015150795 | 0.076719903 |
| *GFOD1* | -1.689761 | 0.696056 | -2.427622 | 0.015198168 | 0.076929516 |
| *CD302* | -4.174133 | 1.71989 | -2.426977 | 0.015225227 | 0.077036174 |
| *IFFO2* | -2.139847 | 0.881769 | -2.426767 | 0.01523403 | 0.077050416 |
| *METTL15* | -1.308599 | 0.539456 | -2.425774 | 0.015275793 | 0.07723129 |
| *SLC19A1* | -2.211076 | 0.911741 | -2.425114 | 0.015303569 | 0.077341329 |
| *SYNPO2L* | -4.573706 | 1.886301 | -2.424695 | 0.015321236 | 0.077400214 |
| *ENSG00000246731* | -3.569313 | 1.472255 | -2.424386 | 0.015334317 | 0.077435891 |
| *RTTN* | -2.208726 | 0.911161 | -2.42408 | 0.015347223 | 0.077443942 |
| *ZFP57* | -4.643641 | 1.915644 | -2.424063 | 0.015347948 | 0.077443942 |
| *PEMT* | -1.870461 | 0.771685 | -2.423866 | 0.015356283 | 0.077455623 |
| *AMER1* | -2.742278 | 1.131502 | -2.423574 | 0.015368616 | 0.077487456 |
| *ENSG00000272540* | -3.597675 | 1.484687 | -2.423188 | 0.015384981 | 0.077525607 |
| *ENSG00000249790* | -3.518808 | 1.452233 | -2.423033 | 0.01539151 | 0.077525607 |
| *BLZF1* | -1.446487 | 0.59699 | -2.422969 | 0.015394259 | 0.077525607 |
| *IGHV3.30* | -3.584698 | 1.479946 | -2.422182 | 0.01542763 | 0.077663271 |
| *ENSG00000142046* | -2.077908 | 0.858057 | -2.421644 | 0.015450468 | 0.077747821 |
| *PPIP5K1* | -2.053999 | 0.848332 | -2.421221 | 0.01546849 | 0.077795442 |
| *LMNTD2* | -2.380655 | 0.983307 | -2.42107 | 0.015474901 | 0.077795442 |
| *ARSD* | -1.844266 | 0.761811 | -2.420897 | 0.015482247 | 0.077795442 |
| *FBXL22* | -2.590863 | 1.070227 | -2.420853 | 0.015484116 | 0.077795442 |
| *LINC00886* | -1.875989 | 0.77504 | -2.420505 | 0.015498969 | 0.077839668 |
| *ENSG00000279345* | -4.768615 | 1.970547 | -2.419945 | 0.015522856 | 0.07792922 |
| *GIPC1* | -1.723491 | 0.712352 | -2.419439 | 0.015544486 | 0.077987916 |
| *SLC41A2* | -2.216096 | 0.915981 | -2.41937 | 0.015547432 | 0.077987916 |
| *ACSL6* | -3.700651 | 1.529671 | -2.419246 | 0.015552731 | 0.077987916 |
| *LGALS3BP* | -2.149327 | 0.888533 | -2.41896 | 0.015564937 | 0.078018718 |
| *TMEM169* | -2.048965 | 0.847232 | -2.418423 | 0.015587938 | 0.078103582 |
| *SDR42E1* | -2.816105 | 1.164556 | -2.418179 | 0.015598406 | 0.078117487 |
| *IGLV2.8* | -4.825999 | 1.995802 | -2.418075 | 0.015602855 | 0.078117487 |
| *BTG2* | -0.470719 | 0.194733 | -2.417257 | 0.015637977 | 0.078262873 |
| *UCK2* | -2.04206 | 0.844904 | -2.416912 | 0.015652791 | 0.07830026 |
| *RETREG1* | -1.738785 | 0.719458 | -2.4168 | 0.015657618 | 0.07830026 |
| *SH3BP5.AS1* | -1.877136 | 0.77675 | -2.416655 | 0.015663847 | 0.078300977 |
| *YEATS2* | -2.017459 | 0.83499 | -2.416148 | 0.015685673 | 0.078367102 |
| *AMDHD1* | -3.514232 | 1.454603 | -2.415938 | 0.015694715 | 0.078367102 |
| *POP1* | -1.842201 | 0.762547 | -2.415852 | 0.01569844 | 0.078367102 |
| *TNKS1BP1* | -4.984258 | 2.063206 | -2.415783 | 0.015701437 | 0.078367102 |
| *LAPTM4B* | -2.213567 | 0.916471 | -2.415316 | 0.015721578 | 0.078437201 |
| *CENPP* | -2.032386 | 0.841615 | -2.414866 | 0.015740987 | 0.078503597 |
| *ALS2* | -1.932109 | 0.80031 | -2.414202 | 0.015769707 | 0.078616357 |
| *PPP1R16A* | -1.636218 | 0.677873 | -2.413753 | 0.015789174 | 0.078682921 |
| *WNT16* | -2.997546 | 1.241949 | -2.413582 | 0.015796573 | 0.078689317 |
| *LINC00921* | -1.797654 | 0.745025 | -2.412878 | 0.015827115 | 0.07881095 |

| | | | | | |
|---|---|---|---|---|---|
| *SNAI3.AS1* | -1.869111 | 0.774743 | -2.412556 | 0.015841107 | 0.078850106 |
| *ARRDC4* | -5.155572 | 2.13742 | -2.412054 | 0.015862941 | 0.07888344 |
| *CST7* | -5.155572 | 2.13742 | -2.412054 | 0.015862941 | 0.07888344 |
| *IGKV3.11* | -2.175812 | 0.902086 | -2.411979 | 0.015866196 | 0.07888344 |
| *MPRIP.AS1* | -1.920726 | 0.796422 | -2.411694 | 0.015878612 | 0.078914678 |
| *SUMO4* | -2.339367 | 0.970219 | -2.411174 | 0.015901248 | 0.078996664 |
| *ETV2* | -2.358531 | 0.978616 | -2.410067 | 0.015949576 | 0.079206173 |
| *GPLD1* | -1.862867 | 0.773012 | -2.409882 | 0.015957681 | 0.079211566 |
| *SRFBP1* | -1.252194 | 0.51964 | -2.409736 | 0.015964061 | 0.079211566 |
| *SMKR1* | -3.285143 | 1.363345 | -2.40962 | 0.01596913 | 0.079211566 |
| *KIFC2* | -2.003677 | 0.831626 | -2.409348 | 0.015981045 | 0.079237024 |
| *XG* | -3.602242 | 1.49533 | -2.408995 | 0.015996532 | 0.079237024 |
| *MYLK* | -4.788504 | 1.987804 | -2.408942 | 0.015998852 | 0.079237024 |
| *TIRAP* | -1.723799 | 0.715584 | -2.408941 | 0.015998895 | 0.079237024 |
| *ENSG00000272936* | -4.859347 | 2.017809 | -2.408229 | 0.016030119 | 0.079353732 |
| *ENSG00000261669* | -1.978548 | 0.821614 | -2.408123 | 0.016034795 | 0.079353732 |
| *IPMK* | -1.92382 | 0.799046 | -2.407644 | 0.016055818 | 0.079422076 |
| *ENSG00000232995* | -2.24666 | 0.933182 | -2.407528 | 0.01606095 | 0.079422076 |
| *ENSG00000260018* | -2.015124 | 0.837365 | -2.406507 | 0.016105899 | 0.079613752 |
| *ENSG00000276957* | -4.811839 | 1.999632 | -2.406362 | 0.016112273 | 0.079614679 |
| *TBL1X* | -1.522197 | 0.632828 | -2.405389 | 0.016155268 | 0.079796483 |
| *CHKB.DT* | -4.142755 | 1.722479 | -2.405113 | 0.016167465 | 0.079826084 |
| *TEX9* | -3.817988 | 1.587965 | -2.404328 | 0.016202226 | 0.079931179 |
| *MKS1* | -2.228416 | 0.926845 | -2.404303 | 0.016203329 | 0.079931179 |
| *HOXB7* | -4.967767 | 2.066316 | -2.404166 | 0.016209411 | 0.079931179 |
| *MCM6* | -1.854258 | 0.771299 | -2.404072 | 0.016213599 | 0.079931179 |
| *LUCAT1* | -2.409902 | 1.002639 | -2.40356 | 0.016236283 | 0.080012355 |
| *MROH1* | -1.572104 | 0.654159 | -2.403245 | 0.016250317 | 0.080037381 |
| *METRN* | -1.960451 | 0.815779 | -2.403166 | 0.016253803 | 0.080037381 |
| *CARNS1* | -2.948709 | 1.227352 | -2.402497 | 0.016283551 | 0.080153195 |
| *LBHD1* | -3.635764 | 1.513585 | -2.402087 | 0.016301829 | 0.080206846 |
| *SH3RF1* | -5.146069 | 2.142434 | -2.401973 | 0.016306918 | 0.080206846 |
| *ENSG00000276148* | -2.239222 | 0.932428 | -2.401495 | 0.016328215 | 0.080280912 |
| *ANKRD16* | -2.169132 | 0.903296 | -2.401352 | 0.016334626 | 0.080281754 |
| *ENSG00000236140* | -2.789022 | 1.161651 | -2.400912 | 0.016354271 | 0.080318591 |
| *TBX19* | -2.079655 | 0.866197 | -2.400904 | 0.016354605 | 0.080318591 |
| *ZNF25* | -2.236468 | 0.93159 | -2.400699 | 0.016363803 | 0.0803331 |
| *ENSG00000273188* | -1.966878 | 0.819383 | -2.400438 | 0.016375477 | 0.080359749 |
| *CRNDE* | -3.537567 | 1.474038 | -2.399916 | 0.016398822 | 0.080443629 |
| *GLCCI1* | -1.397233 | 0.582329 | -2.399388 | 0.016422494 | 0.080529052 |
| *MBNL1.AS1* | -2.025414 | 0.844311 | -2.398894 | 0.016444658 | 0.080607016 |
| *ZNF487* | -1.807744 | 0.753622 | -2.398743 | 0.016451477 | 0.080609732 |
| *PARPBP* | -2.13909 | 0.892017 | -2.398038 | 0.016483178 | 0.080734315 |
| *RAB11FIP3* | -3.555315 | 1.482807 | -2.397693 | 0.016498703 | 0.080754744 |
| *ZFP69B* | -3.536159 | 1.474834 | -2.397666 | 0.016499901 | 0.080754744 |
| *LMNB2* | -1.974046 | 0.823511 | -2.39711 | 0.016524963 | 0.080805667 |
| *ENSG00000272764* | -1.894155 | 0.790205 | -2.397041 | 0.016528075 | 0.080805667 |

| | | | | | |
|---|---|---|---|---|---|
| *PLEKHA4* | -1.868272 | 0.779415 | -2.397017 | 0.016529146 | 0.080805667 |
| *ENSG00000279217* | -3.589768 | 1.497746 | -2.39678 | 0.016539861 | 0.080827344 |
| *RPH3AL* | -2.124486 | 0.886486 | -2.396524 | 0.016551402 | 0.080853032 |
| *SYCP2* | -3.255942 | 1.358849 | -2.396102 | 0.016570458 | 0.0809154 |
| *PRKRA.AS1* | -1.730929 | 0.722606 | -2.3954 | 0.016602258 | 0.081039927 |
| *EDRF1.DT* | -1.996874 | 0.833768 | -2.395 | 0.016620375 | 0.081088948 |
| *HIST3H2A* | -2.034367 | 0.849458 | -2.3949 | 0.016624904 | 0.081088948 |
| *MAP3K10* | -2.109239 | 0.880849 | -2.394552 | 0.016640666 | 0.081125709 |
| *ENSG00000271918* | -1.984588 | 0.828828 | -2.394451 | 0.016645246 | 0.081125709 |
| *ENSG00000268713* | -2.023687 | 0.845204 | -2.394317 | 0.016651356 | 0.081125709 |
| *ITGA5* | -2.118074 | 0.88473 | -2.394035 | 0.016664164 | 0.081157378 |
| *ZNF239* | -2.071953 | 0.865713 | -2.393349 | 0.01669533 | 0.081258177 |
| *EID2B* | -1.46479 | 0.612037 | -2.393302 | 0.016697491 | 0.081258177 |
| *CAPN3* | -2.142655 | 0.895655 | -2.392278 | 0.016744158 | 0.081454473 |
| *CPEB3* | -2.091542 | 0.874386 | -2.392012 | 0.016756312 | 0.081482792 |
| *ENSG00000274341* | -2.793524 | 1.168118 | -2.391474 | 0.016780854 | 0.081571309 |
| *TMSB4Y* | -2.753981 | 1.151865 | -2.39089 | 0.016807609 | 0.08166949 |
| *NAGS* | -2.080267 | 0.87013 | -2.390755 | 0.016813747 | 0.08166949 |
| *ENSG00000272870* | -2.451461 | 1.025721 | -2.389987 | 0.016848972 | 0.081772361 |
| *ENSG00000237181* | -2.197053 | 0.919334 | -2.389831 | 0.016856122 | 0.081772361 |
| *FAM185A* | -1.749297 | 0.731989 | -2.389787 | 0.016858125 | 0.081772361 |
| *ENSG00000262823* | -3.43677 | 1.438136 | -2.389739 | 0.016860346 | 0.081772361 |
| *ZNF835* | -1.998833 | 0.836521 | -2.38946 | 0.01687318 | 0.081803773 |
| *L1TD1* | -2.37722 | 0.995139 | -2.388832 | 0.016902008 | 0.08191267 |
| *FAM53B* | -1.135386 | 0.475403 | -2.388259 | 0.016928422 | 0.081979299 |
| *PARG* | -1.543896 | 0.646482 | -2.388149 | 0.01693346 | 0.081979299 |
| *LINC02166* | -3.742107 | 1.566969 | -2.388119 | 0.01693487 | 0.081979299 |
| *MFSD2A* | -4.88031 | 2.043839 | -2.387815 | 0.01694886 | 0.082016164 |
| *DPP8* | -1.138443 | 0.476854 | -2.387406 | 0.016967742 | 0.082076667 |
| *MEF2C* | 0.701973 | 0.294085 | 2.386974 | 0.016987709 | 0.082133075 |
| *CCND1* | -4.566755 | 1.913276 | -2.386877 | 0.01699217 | 0.082133075 |
| *ENSG00000266538* | -1.865624 | 0.781838 | -2.386203 | 0.017023334 | 0.082197078 |
| *TESC* | -2.508519 | 1.051266 | -2.386189 | 0.017024001 | 0.082197078 |
| *CASTOR3* | -2.058325 | 0.862604 | -2.386177 | 0.017024575 | 0.082197078 |
| *HDLBP* | 1.64107 | 0.687819 | 2.385904 | 0.017037215 | 0.082227247 |
| *LINC01410* | -3.71712 | 1.55807 | -2.38572 | 0.017045715 | 0.082237427 |
| *ENSG00000249249* | -2.100291 | 0.880468 | -2.385425 | 0.01705939 | 0.082258746 |
| *ENSG00000258302* | -3.755731 | 1.5745 | -2.385349 | 0.01706292 | 0.082258746 |
| *IGHG2* | -1.913457 | 0.802365 | -2.38477 | 0.017089799 | 0.082329193 |
| *MYO7B* | -3.607622 | 1.512819 | -2.384703 | 0.017092936 | 0.082329193 |
| *ARL15* | -1.842253 | 0.772595 | -2.384502 | 0.017102271 | 0.082329193 |
| *DAPL1* | -1.871901 | 0.785074 | -2.384363 | 0.017108695 | 0.082329193 |
| *PPARD* | -1.594674 | 0.668833 | -2.384264 | 0.0171133 | 0.082329193 |
| *H3F3C* | -2.412424 | 1.011874 | -2.384115 | 0.017120251 | 0.082329193 |
| *ENSG00000272948* | -3.146123 | 1.319668 | -2.384026 | 0.017124373 | 0.082329193 |
| *ZNF565* | -2.216228 | 0.929652 | -2.383933 | 0.017128721 | 0.082329193 |
| *ZC4H2* | -2.071913 | 0.86922 | -2.383647 | 0.01714203 | 0.082362397 |

| | | | | | |
|---|---|---|---|---|---|
| *AGPS* | -1.564523 | 0.656462 | -2.383265 | 0.01715985 | 0.082383136 |
| *GPR35* | -4.775052 | 2.003706 | -2.383111 | 0.017167029 | 0.082383136 |
| *NDC80* | -4.775052 | 2.003706 | -2.383111 | 0.017167029 | 0.082383136 |
| *ENSG00000242474* | -2.067436 | 0.867575 | -2.383005 | 0.017171957 | 0.082383136 |
| *TRG.AS1* | -2.933624 | 1.231195 | -2.382745 | 0.017184077 | 0.082410553 |
| *IGIP* | -2.244911 | 0.94246 | -2.381969 | 0.017220355 | 0.082527587 |
| *PKD2* | -2.070641 | 0.869337 | -2.381863 | 0.017225292 | 0.082527587 |
| *CNPY4* | -1.673796 | 0.702741 | -2.381811 | 0.017227722 | 0.082527587 |
| *HACE1* | -2.163205 | 0.908461 | -2.381177 | 0.017257431 | 0.082639139 |
| *ANKDD1A* | -2.62075 | 1.100762 | -2.380852 | 0.017272669 | 0.082681337 |
| *ENSG00000260274* | -2.205584 | 0.926573 | -2.380368 | 0.017295352 | 0.082759127 |
| *ZNF284* | -1.694116 | 0.711893 | -2.379735 | 0.017325099 | 0.082851964 |
| *ARHGEF19* | -1.779164 | 0.747659 | -2.379645 | 0.017329316 | 0.082851964 |
| *FBLIM1* | -1.992558 | 0.83737 | -2.379544 | 0.017334071 | 0.082851964 |
| *ENSG00000256705* | -4.849072 | 2.038072 | -2.379245 | 0.017348124 | 0.082888343 |
| *DCAF12* | -4.328142 | 1.819543 | -2.378698 | 0.017373904 | 0.082979193 |
| *ZNF212* | -1.652597 | 0.694787 | -2.378568 | 0.017380036 | 0.082979193 |
| *ZFYVE9* | -3.544798 | 1.490433 | -2.378367 | 0.017389494 | 0.082993551 |
| *RAB12* | -2.174024 | 0.914163 | -2.378158 | 0.017399373 | 0.083009914 |
| *ENSG00000272525* | -4.825765 | 2.029683 | -2.377596 | 0.017425928 | 0.083074997 |
| *C6orf47.AS1* | -4.825765 | 2.029683 | -2.377596 | 0.017425928 | 0.083074997 |
| *GRK6* | -1.559143 | 0.656113 | -2.376334 | 0.01748565 | 0.083315487 |
| *MDGA1* | -3.161911 | 1.330627 | -2.376256 | 0.017489324 | 0.083315487 |
| *ATF3* | -2.04876 | 0.862467 | -2.375465 | 0.01752684 | 0.08345452 |
| *CADM4* | -4.611853 | 1.941619 | -2.375262 | 0.017536499 | 0.08345452 |
| *ENSG00000267469* | -3.332167 | 1.402929 | -2.37515 | 0.017541829 | 0.08345452 |
| *RAB9A* | -1.076593 | 0.453284 | -2.375095 | 0.017544453 | 0.08345452 |
| *NBEA* | -2.330394 | 0.981278 | -2.374856 | 0.0175558 | 0.083477636 |
| *NKG7* | -3.422568 | 1.441386 | -2.374498 | 0.017572815 | 0.083524778 |
| *WDCP* | -1.896153 | 0.798591 | -2.374375 | 0.017578697 | 0.083524778 |
| *APLF* | -1.855426 | 0.781541 | -2.374061 | 0.017593656 | 0.083564995 |
| *COG6* | -1.787025 | 0.752817 | -2.373783 | 0.017606895 | 0.083569476 |
| *ATP1A3* | -4.623871 | 1.947903 | -2.373768 | 0.017607589 | 0.083569476 |
| *ALCAM* | -1.813648 | 0.7641 | -2.373574 | 0.017616876 | 0.083582725 |
| *ENSG00000272829* | -4.720632 | 1.989063 | -2.373295 | 0.017630196 | 0.083615087 |
| *FAM76A* | -1.124133 | 0.473912 | -2.37203 | 0.017690663 | 0.083870951 |
| *TNFSF8* | -4.698074 | 1.980846 | -2.371752 | 0.017703976 | 0.083903153 |
| *ENSG00000279210* | -3.531851 | 1.490194 | -2.370062 | 0.017785101 | 0.084230783 |
| *MPND* | -2.002378 | 0.844934 | -2.369862 | 0.017794724 | 0.084230783 |
| *NUDC* | -0.729984 | 0.308062 | -2.3696 | 0.017807335 | 0.084230783 |
| *PGAM5* | -2.200093 | 0.928497 | -2.369521 | 0.017811151 | 0.084230783 |
| *ENSG00000263264* | -2.004246 | 0.845847 | -2.369514 | 0.017811483 | 0.084230783 |
| *ENSG00000237188* | -1.715184 | 0.723876 | -2.369444 | 0.017814832 | 0.084230783 |
| *ASMTL* | -1.309801 | 0.552808 | -2.369359 | 0.017818932 | 0.084230783 |
| *NOL4L* | -3.556833 | 1.501276 | -2.369207 | 0.017826261 | 0.084234485 |
| *APOC1* | -3.169729 | 1.338319 | -2.368441 | 0.017863221 | 0.084378142 |
| *ENSG00000231113* | -1.826234 | 0.771256 | -2.36787 | 0.017890816 | 0.084477477 |

| | | | | | |
|---|---|---|---|---|---|
| *SLC25A30* | -2.091535 | 0.883402 | -2.367591 | 0.017904315 | 0.084510206 |
| *NUDT12* | -2.20003 | 0.929452 | -2.367018 | 0.017932068 | 0.084610165 |
| *TMEM135* | -2.110366 | 0.891667 | -2.366765 | 0.017944321 | 0.084636942 |
| *ENSG00000042317* | -2.04859 | 0.865838 | -2.36602 | 0.017980451 | 0.08477628 |
| *IL7R* | -2.03549 | 0.860476 | -2.365539 | 0.018003839 | 0.084855459 |
| *KNSTRN* | -1.700028 | 0.718897 | -2.364772 | 0.018041164 | 0.084897286 |
| *MFSD9* | -1.9717 | 0.833849 | -2.364577 | 0.018050651 | 0.084897286 |
| *ENSG00000229043* | -2.39947 | 1.014768 | -2.364549 | 0.018052043 | 0.084897286 |
| *ENSG00000268472* | -4.586009 | 1.939486 | -2.364549 | 0.018052026 | 0.084897286 |
| *TMEM229B* | -4.586009 | 1.939486 | -2.364549 | 0.018052026 | 0.084897286 |
| *SERAC1* | -1.816139 | 0.768092 | -2.36448 | 0.018055405 | 0.084897286 |
| *ALDH4A1* | -2.764343 | 1.169148 | -2.364408 | 0.0180589 | 0.084897286 |
| *WDR90* | -2.139448 | 0.904988 | -2.364063 | 0.018075736 | 0.084945398 |
| *SPNS3* | -2.214256 | 0.937027 | -2.363066 | 0.018124439 | 0.085143175 |
| *CKLF* | -0.967505 | 0.409463 | -2.362864 | 0.018134301 | 0.085158414 |
| *ZNF280B* | -1.963808 | 0.831225 | -2.362548 | 0.018149773 | 0.085173917 |
| *ENSG00000225187* | -3.618323 | 1.531548 | -2.362527 | 0.018150842 | 0.085173917 |
| *VSTM4* | -2.232448 | 0.945172 | -2.361949 | 0.01817913 | 0.085253086 |
| *AXIN2* | -4.509143 | 1.909154 | -2.361854 | 0.018183819 | 0.085253086 |
| *FILIP1L* | -4.733083 | 2.004035 | -2.361777 | 0.01818759 | 0.085253086 |
| *ITGA3* | -2.070146 | 0.8766 | -2.361562 | 0.018198126 | 0.08527141 |
| *SYT1* | -3.070282 | 1.300206 | -2.361382 | 0.018206961 | 0.085281752 |
| *MICOS13* | 0.930814 | 0.394288 | 2.360743 | 0.018238346 | 0.085397669 |
| *TPRN* | -2.164643 | 0.917182 | -2.360103 | 0.018269866 | 0.08551414 |
| *CELF4* | -2.934942 | 1.243851 | -2.359561 | 0.018296552 | 0.085579547 |
| *NUP210* | 1.530311 | 0.648561 | 2.359549 | 0.018297143 | 0.085579547 |
| *ENSG00000240859* | -4.517542 | 1.914799 | -2.359277 | 0.018310568 | 0.085611221 |
| *DZANK1* | -1.956918 | 0.829528 | -2.359074 | 0.018320593 | 0.085618695 |
| *SEPTIN9* | 0.577963 | 0.24502 | 2.358845 | 0.018331913 | 0.085618695 |
| *PDZD11* | -1.328445 | 0.563194 | -2.358771 | 0.018335586 | 0.085618695 |
| *ENSG00000237094* | -4.699596 | 1.992447 | -2.358706 | 0.018338783 | 0.085618695 |
| *ENSG00000264577* | -1.943637 | 0.824129 | -2.358413 | 0.01835325 | 0.085655158 |
| *DOLK* | -1.981541 | 0.840429 | -2.357774 | 0.0183849 | 0.085760121 |
| *HEATR5A* | -1.87027 | 0.793264 | -2.357689 | 0.018389071 | 0.085760121 |
| *ATG2B* | -1.680602 | 0.712908 | -2.357389 | 0.018403951 | 0.085795558 |
| *CDK2AP1* | -2.790968 | 1.184027 | -2.357183 | 0.018414179 | 0.085795558 |
| *DPF3* | -5.030491 | 2.134157 | -2.357133 | 0.018416673 | 0.085795558 |
| *LRRCC1* | -1.767832 | 0.750062 | -2.356914 | 0.018427512 | 0.085810234 |
| *SCAP* | -1.465964 | 0.622014 | -2.3568 | 0.018433161 | 0.085810234 |
| *ALG1L2* | -2.279931 | 0.967624 | -2.356215 | 0.018462224 | 0.085865073 |
| *KIF9* | -2.030877 | 0.861936 | -2.356181 | 0.01846393 | 0.085865073 |
| *CLSPN* | -2.448801 | 1.039319 | -2.35616 | 0.018464961 | 0.085865073 |
| *SLC31A1* | -1.870419 | 0.794015 | -2.355646 | 0.018490554 | 0.08594579 |
| *ENSG00000261416* | -2.308428 | 0.979998 | -2.355543 | 0.018495678 | 0.08594579 |
| *MED14OS* | -1.83898 | 0.780959 | -2.354772 | 0.018534061 | 0.086026974 |
| *LINC00638* | -2.09752 | 0.890777 | -2.354708 | 0.018537241 | 0.086026974 |
| *C5orf34* | -2.137983 | 0.907978 | -2.354665 | 0.018539412 | 0.086026974 |

| | | | | | |
|---|---|---|---|---|---|
| *STAC3* | -2.125801 | 0.902849 | -2.354547 | 0.01854529 | 0.086026974 |
| *HDAC4* | -2.078128 | 0.882639 | -2.354448 | 0.018550244 | 0.086026974 |
| *ENSG00000254281* | -2.625892 | 1.11534 | -2.354342 | 0.01855555 | 0.086026974 |
| *B3GLCT* | -2.027812 | 0.86134 | -2.354253 | 0.01855995 | 0.086026974 |
| *PHF13* | -1.885778 | 0.801064 | -2.354092 | 0.018568015 | 0.086033361 |
| *SYTL3* | -4.653954 | 1.97726 | -2.353739 | 0.01858566 | 0.086084122 |
| *ENSG00000268220* | -2.068004 | 0.87868 | -2.353533 | 0.018595939 | 0.086095551 |
| *PSPN* | -2.080602 | 0.884075 | -2.353422 | 0.01860151 | 0.086095551 |
| *PPP1R3D* | -2.254036 | 0.957873 | -2.353167 | 0.018614264 | 0.08609668 |
| *UCN* | -2.202144 | 0.935828 | -2.35315 | 0.018615137 | 0.08609668 |
| *OVCH1.AS1* | -3.317228 | 1.409947 | -2.352732 | 0.018636045 | 0.086162413 |
| *ZBTB16* | -2.478103 | 1.053499 | -2.352259 | 0.018659784 | 0.08624118 |
| *ZDHHC19* | -3.433445 | 1.460345 | -2.351118 | 0.018717091 | 0.086474977 |
| *PHOSPHO2* | -1.702735 | 0.724303 | -2.35086 | 0.018730092 | 0.086503981 |
| *CENPW* | -1.816215 | 0.772642 | -2.350656 | 0.018740339 | 0.086520253 |
| *REEP6* | -4.467298 | 1.901232 | -2.349685 | 0.018789295 | 0.086698365 |
| *JAZF1.AS1* | -4.645243 | 1.977016 | -2.349624 | 0.018792395 | 0.086698365 |
| *ZNF687.AS1* | -2.187983 | 0.931267 | -2.349468 | 0.018800276 | 0.086703639 |
| *IGLL1* | -2.748361 | 1.169872 | -2.349284 | 0.01880956 | 0.086715376 |
| *BRICD5* | -2.16333 | 0.920939 | -2.349048 | 0.018821474 | 0.086733683 |
| *CAPRIN2* | -1.614792 | 0.687456 | -2.348939 | 0.018827013 | 0.086733683 |
| *KPNA4* | -1.227898 | 0.522783 | -2.348772 | 0.018835419 | 0.086741351 |
| *ZNF571.AS1* | -2.505309 | 1.067112 | -2.347747 | 0.018887348 | 0.086949375 |
| *SLC28A2* | -2.06124 | 0.878122 | -2.347328 | 0.018908612 | 0.086978622 |
| *TMEM81* | -4.586782 | 1.954153 | -2.347197 | 0.018915231 | 0.086978622 |
| *PCNX3* | -1.801855 | 0.767677 | -2.347152 | 0.018917518 | 0.086978622 |
| *CNTLN* | -3.027293 | 1.28985 | -2.347011 | 0.018924683 | 0.086978622 |
| *ZNF469* | -4.564568 | 1.944889 | -2.346956 | 0.0189275 | 0.086978622 |
| *GDAP1* | -1.98845 | 0.847322 | -2.346746 | 0.018938144 | 0.086996463 |
| *LZTR1* | -1.602778 | 0.683143 | -2.346183 | 0.018966815 | 0.087097076 |
| *CUZD1* | -4.49517 | 1.916078 | -2.346026 | 0.018974787 | 0.087102598 |
| *OLMALINC* | -2.790734 | 1.189661 | -2.345824 | 0.018985073 | 0.087118734 |
| *ADAM9* | -4.638025 | 1.97775 | -2.345102 | 0.019021865 | 0.08723766 |
| *TSIX* | -3.920391 | 1.671773 | -2.34505 | 0.019024549 | 0.08723766 |
| *ENSG00000272181* | -4.928768 | 2.102026 | -2.344771 | 0.019038786 | 0.08727184 |
| *BBOF1* | -2.250914 | 0.960096 | -2.344468 | 0.019054231 | 0.087311534 |
| *NAB1* | -1.801628 | 0.768649 | -2.343888 | 0.019083891 | 0.087398481 |
| *TXNDC9* | -1.032301 | 0.440435 | -2.343821 | 0.019087345 | 0.087398481 |
| *SLC18B1* | -1.552922 | 0.662631 | -2.343571 | 0.019100132 | 0.087398481 |
| *AFG1L* | -1.99736 | 0.852274 | -2.343566 | 0.019100375 | 0.087398481 |
| *ENSG00000270019* | -1.583898 | 0.676014 | -2.342997 | 0.019129547 | 0.087483192 |
| *FN1* | -3.394454 | 1.448802 | -2.342939 | 0.019132486 | 0.087483192 |
| *ENSG00000255389* | -2.360484 | 1.007681 | -2.34249 | 0.019155558 | 0.08755757 |
| *ENSG00000259001* | -2.012624 | 0.859242 | -2.342324 | 0.019164058 | 0.087558506 |
| *ZXDA* | -2.220522 | 0.948041 | -2.342221 | 0.019169372 | 0.087558506 |
| *POMT1* | -1.680275 | 0.717651 | -2.341355 | 0.019213904 | 0.087700416 |
| *ENSG00000203546* | -2.225072 | 0.950336 | -2.341351 | 0.019214073 | 0.087700416 |

| | | | | | |
|---|---|---|---|---|---|
| *KCNK6* | -1.410775 | 0.602627 | -2.341043 | 0.019229949 | 0.087741754 |
| *PTK6* | -3.203917 | 1.368764 | -2.340737 | 0.019245727 | 0.087771413 |
| *ENSG00000257194* | -5.049559 | 2.15733 | -2.340652 | 0.019250092 | 0.087771413 |
| *TP53I3* | -2.216853 | 0.947165 | -2.340515 | 0.019257163 | 0.087772555 |
| *SLC41A3* | -1.226149 | 0.523947 | -2.340214 | 0.019272696 | 0.087785118 |
| *HDAC11* | -2.270499 | 0.970217 | -2.340197 | 0.019273565 | 0.087785118 |
| *PPRC1* | -1.848383 | 0.789957 | -2.339854 | 0.019291257 | 0.087804341 |
| *KIAA1328* | -1.440641 | 0.615698 | -2.339851 | 0.019291433 | 0.087804341 |
| *LRP2BP* | -2.066339 | 0.883306 | -2.339324 | 0.019318648 | 0.087872611 |
| *PIK3R2* | -3.29744 | 1.409586 | -2.339297 | 0.019320091 | 0.087872611 |
| *TST* | -2.451507 | 1.048169 | -2.338848 | 0.019343315 | 0.087947151 |
| *BOLA2.SMG1P6* | -1.551765 | 0.663572 | -2.338503 | 0.019361155 | 0.087952312 |
| *RND1* | -1.837404 | 0.785719 | -2.3385 | 0.019361343 | 0.087952312 |
| *TEX10* | -1.548782 | 0.662319 | -2.338424 | 0.019365266 | 0.087952312 |
| *DUSP16* | -4.783478 | 2.045709 | -2.338298 | 0.019371792 | 0.087952312 |
| *TEPP* | -2.141516 | 0.915897 | -2.338164 | 0.01937874 | 0.087952821 |
| *PRKAB1* | -1.520907 | 0.65089 | -2.336656 | 0.019457057 | 0.088221431 |
| *DGKZ* | -1.432457 | 0.613055 | -2.336587 | 0.019460679 | 0.088221431 |
| *YOD1* | -1.840072 | 0.787518 | -2.336547 | 0.019462752 | 0.088221431 |
| *CLCN5* | -3.345656 | 1.431911 | -2.336497 | 0.019465349 | 0.088221431 |
| *MYBL2* | -3.338895 | 1.429155 | -2.336272 | 0.019477066 | 0.088238842 |
| *SLC25A53* | -1.442218 | 0.617352 | -2.336136 | 0.019484152 | 0.088238842 |
| *TRAF3IP1* | -2.613432 | 1.11875 | -2.336028 | 0.019489764 | 0.088238842 |
| *CCNJ* | -2.136189 | 0.914534 | -2.335823 | 0.019500452 | 0.088256177 |
| *ENSG00000247373* | -2.043672 | 0.874991 | -2.335648 | 0.01950957 | 0.088266397 |
| *DLK2* | -5.134031 | 2.198795 | -2.334929 | 0.019547118 | 0.088405192 |
| *CA5A* | -4.788898 | 2.05148 | -2.334363 | 0.019576732 | 0.088508013 |
| *TRAPPC2L* | 1.472614 | 0.630893 | 2.334176 | 0.019586531 | 0.088521212 |
| *KTN1.AS1* | -3.434683 | 1.471691 | -2.333834 | 0.019604421 | 0.088546495 |
| *ABCB10* | -1.778484 | 0.762053 | -2.333806 | 0.019605888 | 0.088546495 |
| *SREK1* | -0.825971 | 0.354065 | -2.332821 | 0.019657543 | 0.088711802 |
| *ENSG00000272426* | -2.312614 | 0.991403 | -2.332669 | 0.019665538 | 0.088711802 |
| *SLC7A11* | -4.414023 | 1.892294 | -2.332631 | 0.01966753 | 0.088711802 |
| *LY6E* | -0.642266 | 0.275351 | -2.332537 | 0.019672439 | 0.088711802 |
| *TMPO.AS1* | -2.004804 | 0.859527 | -2.332451 | 0.019676963 | 0.088711802 |
| *GLB1L* | -1.812507 | 0.77719 | -2.33213 | 0.019693836 | 0.088756772 |
| *SNRPF* | -0.553277 | 0.237284 | -2.331707 | 0.01971612 | 0.088805168 |
| *ENSG00000245025* | -2.123725 | 0.91082 | -2.331664 | 0.019718378 | 0.088805168 |
| *C1QTNF3* | -2.415922 | 1.036274 | -2.331355 | 0.019734638 | 0.088818078 |
| *UBOX5* | -1.820126 | 0.780737 | -2.331293 | 0.019737911 | 0.088818078 |
| *LINC00887* | -5.029431 | 2.157428 | -2.331216 | 0.019741952 | 0.088818078 |
| *KIF13A* | -2.344575 | 1.005913 | -2.330793 | 0.019764267 | 0.088887388 |
| *DCLRE1B* | -2.244572 | 0.963184 | -2.330367 | 0.019786758 | 0.088957446 |
| *YPEL5* | 1.30863 | 0.561638 | 2.330022 | 0.01980498 | 0.089008269 |
| *ENSG00000227070* | -3.82031 | 1.639755 | -2.329806 | 0.019816429 | 0.089028628 |
| *ENSG00000275580* | -4.934653 | 2.118332 | -2.329499 | 0.019832635 | 0.089070336 |
| *GNE* | -1.738511 | 0.746467 | -2.328988 | 0.019859721 | 0.089093529 |

| | | | | | |
|---|---|---|---|---|---|
| *IGHV1.18* | -4.641533 | 1.992987 | -2.328934 | 0.019862582 | 0.089093529 |
| *AGER* | -2.106117 | 0.904373 | -2.328816 | 0.019868817 | 0.089093529 |
| *RNF208* | -4.705033 | 2.020381 | -2.328785 | 0.019870459 | 0.089093529 |
| *WDR62* | -5.095148 | 2.187935 | -2.328748 | 0.01987242 | 0.089093529 |
| *PLD2* | -1.739519 | 0.747224 | -2.327975 | 0.019913433 | 0.089246304 |
| *SEMA4F* | -1.907093 | 0.819671 | -2.326658 | 0.01998349 | 0.089529096 |
| *ENSG00000271855* | -2.672275 | 1.148635 | -2.326479 | 0.019993021 | 0.089540619 |
| *RASGRF1* | -2.224771 | 0.956434 | -2.326111 | 0.020012652 | 0.089597353 |
| *CLPB* | -1.938209 | 0.833323 | -2.325881 | 0.020024926 | 0.08962112 |
| *TRAM2* | -1.971808 | 0.847865 | -2.325616 | 0.02003903 | 0.089624042 |
| *ABCC2* | -4.845596 | 2.083583 | -2.325607 | 0.020039509 | 0.089624042 |
| *ELL3* | -2.167319 | 0.932076 | -2.32526 | 0.020058057 | 0.089675822 |
| *RPL38* | -0.152096 | 0.065441 | -2.324177 | 0.020116005 | 0.089903658 |
| *MTCH1* | 1.37291 | 0.590875 | 2.323522 | 0.020151126 | 0.090011668 |
| *ENSG00000263884* | -4.712601 | 2.028264 | -2.323465 | 0.020154163 | 0.090011668 |
| *PEX5* | -1.785935 | 0.768967 | -2.322512 | 0.020205357 | 0.090203835 |
| *MYO1D* | -4.456124 | 1.918755 | -2.322404 | 0.020211211 | 0.090203835 |
| *ADAM20* | -4.667862 | 2.010098 | -2.322206 | 0.020221856 | 0.090220048 |
| *A2M.AS1* | -5.213216 | 2.245583 | -2.321542 | 0.02025759 | 0.090348151 |
| *SH3D21* | -3.293063 | 1.418693 | -2.321196 | 0.020276286 | 0.090400198 |
| *SDCBP2* | -3.408742 | 1.469176 | -2.320172 | 0.020331551 | 0.090615194 |
| *VWA5A* | -2.5522 | 1.1003 | -2.319549 | 0.020365306 | 0.09073421 |
| *LINC01063* | -3.407132 | 1.469466 | -2.318619 | 0.020415712 | 0.090917669 |
| *ENSG00000263766* | -4.583179 | 1.976762 | -2.318529 | 0.020420616 | 0.090917669 |
| *NAA50* | -0.751787 | 0.32431 | -2.31811 | 0.020443364 | 0.090987467 |
| *IGF2BP3* | -4.734752 | 2.042763 | -2.317817 | 0.020459264 | 0.091008577 |
| *PYCR1* | -3.344672 | 1.443061 | -2.317762 | 0.020462253 | 0.091008577 |
| *CACNA1F* | -4.8672 | 2.100209 | -2.317484 | 0.020477375 | 0.091038421 |
| *WDR31* | -5.246101 | 2.26389 | -2.317295 | 0.020487654 | 0.091038421 |
| *PHGDH* | -3.856429 | 1.664228 | -2.317249 | 0.020490189 | 0.091038421 |
| *POLR1A* | -1.798026 | 0.775978 | -2.317111 | 0.020497704 | 0.091040371 |
| *THBS3* | -2.118925 | 0.914551 | -2.316902 | 0.020509087 | 0.091059495 |
| *PAQR3* | -1.968515 | 0.849685 | -2.316757 | 0.020516946 | 0.091062967 |
| *ENSG00000261584* | -2.574373 | 1.111363 | -2.31641 | 0.020535908 | 0.091115698 |
| *PLCD3* | -3.133724 | 1.352997 | -2.316136 | 0.02055086 | 0.091120182 |
| *TECPR2* | -2.103932 | 0.908382 | -2.316132 | 0.020551082 | 0.091120182 |
| *ENSG00000260228* | -4.231663 | 1.827532 | -2.315506 | 0.020585247 | 0.091240226 |
| *ENSG00000272630* | -1.989909 | 0.859513 | -2.315157 | 0.02060432 | 0.091275019 |
| *LINC00852* | -2.171719 | 0.938066 | -2.315103 | 0.020607285 | 0.091275019 |
| *ZNF510* | -1.750436 | 0.756455 | -2.314001 | 0.020667687 | 0.091511058 |
| *RAB30.DT* | -0.958114 | 0.414115 | -2.313643 | 0.020687336 | 0.091566546 |
| *DLGAP4.AS1* | -2.151914 | 0.9304 | -2.31289 | 0.020728703 | 0.091718095 |
| *PTPRJ* | -2.725799 | 1.17866 | -2.312626 | 0.020743223 | 0.091750791 |
| *PIDD1* | -1.614915 | 0.698485 | -2.312026 | 0.020776246 | 0.091865276 |
| *SPIN3* | -4.80609 | 2.079957 | -2.310668 | 0.020851209 | 0.092165064 |
| *RPS6KC1* | -1.606956 | 0.695601 | -2.310169 | 0.020878789 | 0.092255282 |
| *ENSG00000237950* | -4.580951 | 1.983532 | -2.309492 | 0.020916312 | 0.092389353 |

| | | | | | |
|---|---|---|---|---|---|
| *AZIN1.AS1* | -1.96482 | 0.85096 | -2.308945 | 0.020946615 | 0.092444325 |
| *TEPSIN* | -1.347769 | 0.583727 | -2.308903 | 0.020948955 | 0.092444325 |
| *SCN3A* | -2.60491 | 1.128314 | -2.308674 | 0.020961682 | 0.092444325 |
| *HSBP1L1* | -2.259785 | 0.97892 | -2.308447 | 0.020974266 | 0.092444325 |
| *FOSL2* | -4.561182 | 1.97588 | -2.30843 | 0.020975236 | 0.092444325 |
| *GPR171* | -3.818017 | 1.653963 | -2.308406 | 0.020976591 | 0.092444325 |
| *LRRIQ3* | -2.383589 | 1.032589 | -2.308361 | 0.020979049 | 0.092444325 |
| *C19orf81* | -4.283968 | 1.856009 | -2.308162 | 0.020990146 | 0.092450705 |
| *SLC25A35* | -2.121088 | 0.918985 | -2.308077 | 0.020994868 | 0.092450705 |
| *CUL7* | -2.022525 | 0.876451 | -2.307632 | 0.021019619 | 0.092511364 |
| *ENSG00000257354* | -1.727076 | 0.748439 | -2.307571 | 0.021023022 | 0.092511364 |
| *CSF2RB* | -2.042439 | 0.885219 | -2.30727 | 0.021039773 | 0.092548181 |
| *ENSG00000261200* | -2.93056 | 1.270201 | -2.307162 | 0.021045774 | 0.092548181 |
| *SYNGR1* | -1.70087 | 0.737272 | -2.306977 | 0.021056081 | 0.092561871 |
| *NEK3* | -2.030388 | 0.880575 | -2.305753 | 0.021124423 | 0.092830584 |
| *RFT1* | -1.653864 | 0.71766 | -2.304524 | 0.021193268 | 0.093014209 |
| *ZNF142* | -1.817874 | 0.788837 | -2.304498 | 0.021194708 | 0.093014209 |
| *IL6R* | -4.603879 | 1.997854 | -2.304412 | 0.021199511 | 0.093014209 |
| *ENSG00000260271* | -2.582812 | 1.120844 | -2.304347 | 0.021203187 | 0.093014209 |
| *PTAFR* | -2.125072 | 0.922204 | -2.30434 | 0.02120355 | 0.093014209 |
| *ENSG00000267169* | -1.830753 | 0.794517 | -2.304233 | 0.021209582 | 0.093014209 |
| *ARL5B* | -1.66545 | 0.722863 | -2.303965 | 0.021224606 | 0.093048385 |
| *TMOD1* | -2.257316 | 0.980276 | -2.302736 | 0.021293702 | 0.093319504 |
| *FBXW4* | -1.324107 | 0.575053 | -2.302583 | 0.021302328 | 0.09332552 |
| *GSTCD* | -1.823788 | 0.792308 | -2.301869 | 0.021342578 | 0.093417355 |
| *ENSG00000257433* | -1.936048 | 0.841102 | -2.301798 | 0.021346546 | 0.093417355 |
| *ATRN* | -1.893559 | 0.822648 | -2.301784 | 0.021347335 | 0.093417355 |
| *TOMM7* | -0.303749 | 0.131967 | -2.301696 | 0.021352331 | 0.093417355 |
| *TTTY14* | -2.827152 | 1.228619 | -2.30108 | 0.02138711 | 0.093537711 |
| *TMEM267* | -1.354309 | 0.588727 | -2.300402 | 0.021425438 | 0.093673498 |
| *CASZ1* | -1.916041 | 0.833216 | -2.299573 | 0.021472418 | 0.093847011 |
| *SH3D19* | -2.060476 | 0.896116 | -2.299342 | 0.021485528 | 0.093858743 |
| *C21orf58* | -2.099807 | 0.91325 | -2.299269 | 0.021489691 | 0.093858743 |
| *SMARCD3* | -4.526562 | 1.969713 | -2.298082 | 0.021557145 | 0.094117184 |
| *ANG* | -4.376693 | 1.904591 | -2.29797 | 0.021563492 | 0.094117184 |
| *JSRP1* | -3.645391 | 1.58665 | -2.297539 | 0.02158805 | 0.094165685 |
| *BATF2* | -2.68102 | 1.16692 | -2.297518 | 0.021589241 | 0.094165685 |
| *ENSG00000272606* | -1.611504 | 0.701689 | -2.296608 | 0.021641139 | 0.094360062 |
| *CD58* | 1.641577 | 0.714879 | 2.296299 | 0.021658793 | 0.094405043 |
| *CHSY1* | -1.887926 | 0.822228 | -2.296109 | 0.021669626 | 0.094420277 |
| *ZNF584* | -1.73953 | 0.757832 | -2.295402 | 0.021710111 | 0.094561454 |
| *CAMKK2* | -1.81913 | 0.79255 | -2.295286 | 0.021716725 | 0.094561454 |
| *RESF1* | -0.755257 | 0.3291 | -2.294915 | 0.021737978 | 0.094621977 |
| *ENSG00000204802* | -2.020317 | 0.880701 | -2.293988 | 0.021791209 | 0.094807787 |
| *ERCC2* | -1.605931 | 0.700083 | -2.293915 | 0.021795402 | 0.094807787 |
| *FBXW9* | -2.008471 | 0.875744 | -2.293446 | 0.02182233 | 0.094892844 |
| *RCBTB1* | -1.667349 | 0.727201 | -2.29283 | 0.021857785 | 0.095014907 |

| | | | | | |
|---|---|---|---|---|---|
| *EPOP* | -4.783894 | 2.086781 | -2.292475 | 0.021878224 | 0.095071633 |
| *NGRN* | -1.699068 | 0.741326 | -2.291931 | 0.021909643 | 0.095176023 |
| *TPRG1L* | -1.444942 | 0.630562 | -2.291516 | 0.021933609 | 0.095247975 |
| *ENSG00000277496* | -1.943738 | 0.848576 | -2.290588 | 0.021987244 | 0.095448673 |
| *TMEM38A* | -1.842634 | 0.804502 | -2.290404 | 0.021997929 | 0.095462849 |
| *PALLD* | -2.03677 | 0.889529 | -2.289719 | 0.022037619 | 0.095601702 |
| *GOLGA6L9* | -2.981743 | 1.302301 | -2.289595 | 0.022044785 | 0.095601702 |
| *TBKBP1* | -4.653318 | 2.032887 | -2.289019 | 0.022078256 | 0.095692188 |
| *ENSG00000259488* | -2.291949 | 1.001297 | -2.28898 | 0.022080524 | 0.095692188 |
| *ZNF354C* | -2.016393 | 0.880974 | -2.288822 | 0.022089724 | 0.095699823 |
| *CERCAM* | -1.89542 | 0.828178 | -2.288661 | 0.022099033 | 0.09570793 |
| *METRNL* | -2.691144 | 1.176095 | -2.288203 | 0.022125716 | 0.095791245 |
| *MYO5C* | -2.175477 | 0.950893 | -2.287826 | 0.022147656 | 0.095853984 |
| *KRT18* | -1.604252 | 0.701293 | -2.287564 | 0.022162927 | 0.095887822 |
| *RPS4Y2* | -3.523896 | 1.540752 | -2.287127 | 0.022188421 | 0.095965852 |
| *CENPU* | -2.168529 | 0.948202 | -2.28699 | 0.022196412 | 0.095968158 |
| *SGPP2* | -4.543672 | 1.987053 | -2.286638 | 0.022216955 | 0.09602471 |
| *ENSG00000225793* | -4.549416 | 1.989785 | -2.286386 | 0.022231663 | 0.096027358 |
| *SLC39A8* | -1.760512 | 0.770002 | -2.286372 | 0.022232494 | 0.096027358 |
| *RXRA* | -2.213357 | 0.96818 | -2.2861 | 0.022248422 | 0.096063908 |
| *KMO* | -1.414918 | 0.618984 | -2.285871 | 0.022261814 | 0.096089487 |
| *ENSG00000187951* | -1.657864 | 0.725342 | -2.285632 | 0.022275768 | 0.096097233 |
| *ZNF736* | -1.472298 | 0.644181 | -2.285534 | 0.02228154 | 0.096097233 |
| *TRAFD1* | -1.495594 | 0.654445 | -2.285286 | 0.022296039 | 0.096097233 |
| *HHIP.AS1* | -4.132914 | 1.808491 | -2.285284 | 0.022296203 | 0.096097233 |
| *ENSG00000268292* | -4.358353 | 1.907346 | -2.285035 | 0.022310751 | 0.096097233 |
| *GUCY2C* | -1.567313 | 0.685941 | -2.28491 | 0.022318127 | 0.096097233 |
| *THAP4* | -1.368658 | 0.599003 | -2.284893 | 0.022319079 | 0.096097233 |
| *GPAM* | -2.022175 | 0.885048 | -2.28482 | 0.022323357 | 0.096097233 |
| *EME2* | -1.937537 | 0.848221 | -2.284237 | 0.022357606 | 0.096212482 |
| *TONSL* | -1.774065 | 0.776852 | -2.283658 | 0.022391641 | 0.096326729 |
| *ERRFI1* | -4.611442 | 2.019724 | -2.283204 | 0.022418357 | 0.096409424 |
| *INPP5E* | -1.684924 | 0.738174 | -2.282556 | 0.022456533 | 0.096489023 |
| *AMT* | -1.958075 | 0.857855 | -2.282525 | 0.022458353 | 0.096489023 |
| *ANO10* | -1.96159 | 0.859401 | -2.282508 | 0.022459363 | 0.096489023 |
| *PTBP1* | -0.852974 | 0.373837 | -2.281672 | 0.022508693 | 0.096627242 |
| *TWNK* | -1.833454 | 0.803598 | -2.281556 | 0.022515585 | 0.096627242 |
| *SIK1* | -4.771191 | 2.091213 | -2.281542 | 0.022516395 | 0.096627242 |
| *SLC38A1* | -0.677387 | 0.29691 | -2.281454 | 0.022521574 | 0.096627242 |
| *EP300* | -1.010344 | 0.442914 | -2.281131 | 0.022540704 | 0.096677081 |
| *ENSG00000267397* | -4.324094 | 1.895751 | -2.28094 | 0.022552023 | 0.096693394 |
| *ENSG00000261135* | -2.462457 | 1.080323 | -2.279371 | 0.022645048 | 0.097059906 |
| *LINC00158* | -4.661272 | 2.045555 | -2.278732 | 0.022682993 | 0.09713566 |
| *ENSG00000182109* | -2.204454 | 0.967435 | -2.278658 | 0.022687384 | 0.09713566 |
| *LCA5* | -3.579109 | 1.570731 | -2.278626 | 0.022689302 | 0.09713566 |
| *ENSG00000237851* | -4.547206 | 1.995644 | -2.278565 | 0.022692919 | 0.09713566 |
| *ENSG00000231856* | -1.869723 | 0.820693 | -2.278225 | 0.022713196 | 0.097190121 |

| | | | | | |
|---|---|---|---|---|---|
| *DDTL* | -4.45289 | 1.954727 | -2.278011 | 0.022725894 | 0.097212129 |
| *SRRT* | -1.068268 | 0.469157 | -2.276994 | 0.022786571 | 0.097439283 |
| *HSPA1B* | -3.203295 | 1.407256 | -2.27627 | 0.022829858 | 0.097560062 |
| *KLRG1* | -1.970773 | 0.865874 | -2.276051 | 0.022842939 | 0.097560062 |
| *TLR7* | -1.776825 | 0.780664 | -2.276044 | 0.022843362 | 0.097560062 |
| *ITPKB.AS1* | -1.818257 | 0.79892 | -2.275893 | 0.022852409 | 0.097560062 |
| *RPL30* | -0.124959 | 0.054906 | -2.275888 | 0.022852726 | 0.097560062 |
| *SBNO2* | -1.947054 | 0.855654 | -2.275516 | 0.022874985 | 0.097576503 |
| *SAP30* | -4.709601 | 2.069699 | -2.2755 | 0.022875943 | 0.097576503 |
| *LZTS3* | -4.500383 | 1.977805 | -2.275444 | 0.022879328 | 0.097576503 |
| *ENSG00000232807* | -4.485704 | 1.972245 | -2.274415 | 0.02294103 | 0.097807236 |
| *ENSG00000254614* | -1.546634 | 0.680075 | -2.274211 | 0.022953306 | 0.097827156 |
| *CABLES2* | -3.288621 | 1.446313 | -2.273796 | 0.022978276 | 0.097871415 |
| *RAB11B.AS1* | -1.412899 | 0.621386 | -2.273785 | 0.022978903 | 0.097871415 |
| *ENSG00000269898* | -1.99541 | 0.877792 | -2.273216 | 0.023013176 | 0.097984956 |
| *TRGV5* | -2.226743 | 0.979667 | -2.272958 | 0.023028682 | 0.098018542 |
| *CHAF1A* | -1.901191 | 0.836573 | -2.272593 | 0.023050698 | 0.098079806 |
| *PLXND1* | -2.174964 | 0.957386 | -2.271774 | 0.023100152 | 0.098203413 |
| *CCDC120* | -1.998141 | 0.87956 | -2.27175 | 0.023101586 | 0.098203413 |
| *IGLV2.14* | -2.527753 | 1.112698 | -2.271733 | 0.023102645 | 0.098203413 |
| *LIN52* | -1.55626 | 0.68519 | -2.271282 | 0.023129914 | 0.098286855 |
| *ENSG00000268568* | -2.156409 | 0.949558 | -2.270962 | 0.023149284 | 0.098336692 |
| *ENSG00000130520* | -0.573624 | 0.252622 | -2.270684 | 0.02316614 | 0.098348355 |
| *ENSG00000250938* | -4.584401 | 2.018969 | -2.270664 | 0.023167317 | 0.098348355 |
| *ENSG00000260563* | -3.237993 | 1.426162 | -2.270424 | 0.02318184 | 0.098373728 |
| *MCF2L* | -4.437094 | 1.954397 | -2.270313 | 0.023188584 | 0.098373728 |
| *CYHR1* | -0.981107 | 0.432223 | -2.269908 | 0.023213153 | 0.098445499 |
| *RRP9* | -1.474512 | 0.649846 | -2.269019 | 0.023267192 | 0.09864216 |
| *GALK2* | -1.390858 | 0.613153 | -2.268368 | 0.023306775 | 0.098777427 |
| *PTCD2* | -1.541908 | 0.67989 | -2.267878 | 0.02333665 | 0.098821038 |
| *TRIM44* | 1.339828 | 0.590791 | 2.267856 | 0.023337987 | 0.098821038 |
| *EMC1.AS1* | -3.052233 | 1.345888 | -2.267821 | 0.023340105 | 0.098821038 |
| *SNX13* | -1.302517 | 0.57441 | -2.267574 | 0.023355206 | 0.098852448 |
| *ZNF287* | -1.834293 | 0.808973 | -2.267434 | 0.023363755 | 0.098856114 |
| *GZF1* | -1.653062 | 0.729247 | -2.266805 | 0.023402139 | 0.098985972 |
| *CCNA2* | -4.037019 | 1.781324 | -2.266302 | 0.023432883 | 0.09908344 |
| *CDCA7* | -4.570022 | 2.017929 | -2.264709 | 0.023530526 | 0.099463626 |
| *TRUB1* | -1.626977 | 0.718523 | -2.264333 | 0.023553624 | 0.099528566 |
| *ENSG00000229852* | -1.407436 | 0.621612 | -2.26417 | 0.023563682 | 0.099537806 |
| *ENSG00000261158* | -4.389455 | 1.938766 | -2.264046 | 0.023571283 | 0.099537806 |
| *ENSG00000238198* | -4.463469 | 1.971902 | -2.263535 | 0.023602757 | 0.09960523 |
| *MPP7* | -1.786303 | 0.789167 | -2.263529 | 0.023603112 | 0.09960523 |
| *SRPK1* | 1.633121 | 0.721532 | 2.263409 | 0.023610473 | 0.09960523 |
| *MGST2* | -2.091934 | 0.924492 | -2.262793 | 0.023648466 | 0.099732814 |
| *ZKSCAN2* | -1.960809 | 0.866877 | -2.261923 | 0.023702165 | 0.099926526 |
| *PRADC1* | -1.777825 | 0.786083 | -2.261626 | 0.023720507 | 0.099971098 |
| *LINC02242* | -3.327721 | 1.471615 | -2.261272 | 0.023742443 | 0.100030784 |

| | | | | | |
|---|---|---|---|---|---|
| *GSK3A* | -1.501026 | 0.663933 | -2.26081 | 0.023771055 | 0.10011855 |
| *TM4SF19.AS1* | -4.139753 | 1.831397 | -2.260435 | 0.023794267 | 0.100183518 |
| *CCT3* | 1.009731 | 0.446769 | 2.260071 | 0.023816826 | 0.100245697 |
| *ENSG00000259943* | -1.621058 | 0.717682 | -2.258741 | 0.023899496 | 0.100560762 |
| *MAFG.DT* | -4.319371 | 1.913262 | -2.257595 | 0.023970919 | 0.100803486 |
| *ERN1* | -3.19937 | 1.417178 | -2.257564 | 0.02397285 | 0.100803486 |
| *ZRANB1* | -1.458474 | 0.646093 | -2.257374 | 0.023984686 | 0.100820305 |
| *ENSG00000263335* | -2.199253 | 0.974404 | -2.257024 | 0.024006566 | 0.100879321 |
| *PLLP* | -4.303137 | 1.906868 | -2.256651 | 0.024029866 | 0.100944267 |
| *ENSG00000236833* | -4.491528 | 1.991469 | -2.255384 | 0.024109246 | 0.101244669 |
| *AARS2* | -1.944047 | 0.862084 | -2.255055 | 0.024129847 | 0.101271993 |
| *MSANTD2* | -1.645132 | 0.729539 | -2.255029 | 0.024131494 | 0.101271993 |
| *ACOT2* | -2.377951 | 1.054636 | -2.254759 | 0.024148449 | 0.101310104 |
| *SLC14A1* | -4.278184 | 1.898601 | -2.253335 | 0.024238058 | 0.101652899 |
| *C2orf92* | -4.244822 | 1.884224 | -2.252822 | 0.024270351 | 0.101702257 |
| *ENSG00000227518* | -4.244822 | 1.884224 | -2.252822 | 0.024270351 | 0.101702257 |
| *FSD1* | -2.506883 | 1.112813 | -2.252745 | 0.024275255 | 0.101702257 |
| *PHLDB3* | -1.401893 | 0.622393 | -2.252423 | 0.024295568 | 0.101702257 |
| *TFB1M* | -1.219823 | 0.541563 | -2.252411 | 0.024296323 | 0.101702257 |
| *TOR4A* | -1.698177 | 0.753943 | -2.252396 | 0.024297252 | 0.101702257 |
| *BAHD1* | -1.752336 | 0.77814 | -2.251956 | 0.02432506 | 0.101785543 |
| *MIR497HG* | -2.087689 | 0.927178 | -2.251659 | 0.024343821 | 0.101830932 |
| *RNPEPL1* | -1.110211 | 0.493109 | -2.251452 | 0.024356899 | 0.101852525 |
| *SLC35G5* | -2.015929 | 0.895487 | -2.25121 | 0.024372271 | 0.101883695 |
| *ACTR3B* | -1.802683 | 0.800846 | -2.250973 | 0.024387277 | 0.101913314 |
| *SLC23A3* | -2.031415 | 0.902804 | -2.250116 | 0.024441557 | 0.102106984 |
| *SPPL3* | -1.300436 | 0.578156 | -2.24928 | 0.024494683 | 0.102295712 |
| *HDX* | -2.097761 | 0.932982 | -2.248448 | 0.024547645 | 0.102445804 |
| *ENDOG* | -2.129959 | 0.947352 | -2.248328 | 0.024555263 | 0.102445804 |
| *LANCL1* | -1.553614 | 0.691079 | -2.2481 | 0.024569845 | 0.102445804 |
| *PJA1* | -1.304015 | 0.580076 | -2.248007 | 0.024575748 | 0.102445804 |
| *PRR5L* | -4.33688 | 1.929229 | -2.247987 | 0.024577037 | 0.102445804 |
| *MXRA7* | -2.23293 | 0.993311 | -2.247965 | 0.024578394 | 0.102445804 |
| *FAM111B* | -0.963139 | 0.428571 | -2.247327 | 0.024619112 | 0.102555747 |
| *PTPRK* | -1.817558 | 0.808789 | -2.24726 | 0.024623442 | 0.102555747 |
| *MFN2* | -1.567291 | 0.697466 | -2.247123 | 0.024632136 | 0.102555747 |
| *NEK6* | -1.706815 | 0.759579 | -2.247053 | 0.024636653 | 0.102555747 |
| *ENSG00000270189* | -1.541621 | 0.686127 | -2.246845 | 0.024649921 | 0.102577793 |
| *LINC01252* | -2.289869 | 1.019243 | -2.246637 | 0.024663217 | 0.10259994 |
| *NEURL4* | -1.966747 | 0.875878 | -2.245458 | 0.024738717 | 0.102880762 |
| *SLFN13* | -1.898954 | 0.845758 | -2.245269 | 0.02475088 | 0.102898086 |
| *DHCR24* | -1.924371 | 0.857295 | -2.244701 | 0.02478735 | 0.103016418 |
| *LIMD1.AS1* | -4.404831 | 1.96277 | -2.244191 | 0.024820134 | 0.103119364 |
| *ZNF599* | -2.178174 | 0.970989 | -2.243254 | 0.024880421 | 0.103336467 |
| *ENSG00000234936* | -1.879076 | 0.837862 | -2.242704 | 0.024915922 | 0.103443231 |
| *PANK3* | -1.314216 | 0.586023 | -2.242601 | 0.024922545 | 0.103443231 |
| *ADGRG5* | -2.132255 | 0.950846 | -2.242482 | 0.024930245 | 0.103443231 |

| | | | | | |
|---|---|---|---|---|---|
| *LILRB3* | -1.989644 | 0.887379 | -2.242158 | 0.024951151 | 0.1034966 |
| *IGSF22* | -2.192062 | 0.977869 | -2.241672 | 0.024982556 | 0.103593474 |
| *ATL2* | -1.560649 | 0.696334 | -2.241237 | 0.025010732 | 0.103676897 |
| *MANEA* | -1.732958 | 0.773308 | -2.240969 | 0.025028074 | 0.103715373 |
| *ENSG00000267838* | -2.350645 | 1.049111 | -2.240607 | 0.025051572 | 0.103779323 |
| *LRRC8B* | -1.706918 | 0.762081 | -2.239811 | 0.025103186 | 0.103913403 |
| *ERLIN1* | -1.716288 | 0.766276 | -2.239777 | 0.025105377 | 0.103913403 |
| *JHY* | -2.066328 | 0.922593 | -2.239696 | 0.025110641 | 0.103913403 |
| *BRIX1* | 1.624117 | 0.725178 | 2.23961 | 0.025116242 | 0.103913403 |
| *BRF1* | -1.76823 | 0.78959 | -2.239427 | 0.025128142 | 0.103929222 |
| *PREP* | -1.793708 | 0.801211 | -2.238748 | 0.025172349 | 0.104064215 |
| *ENSG00000140743* | -2.191011 | 0.978708 | -2.238677 | 0.025176957 | 0.104064215 |
| *SYT15* | -2.923976 | 1.306286 | -2.238388 | 0.02519575 | 0.104108452 |
| *ENSG00000178386* | -2.166562 | 0.968084 | -2.23799 | 0.025221678 | 0.104182129 |
| *ENSG00000272049* | -4.212955 | 1.882699 | -2.237721 | 0.025239261 | 0.104193097 |
| *CPLANE1* | -1.407949 | 0.6292 | -2.237681 | 0.025241855 | 0.104193097 |
| *DDX21* | 1.467384 | 0.655821 | 2.237477 | 0.025255174 | 0.104193097 |
| *SLC51A* | -2.546847 | 1.13828 | -2.237453 | 0.025256724 | 0.104193097 |
| *ZDHHC13* | -1.474584 | 0.65937 | -2.236353 | 0.025328634 | 0.104434316 |
| *CLNK* | -4.390754 | 1.963392 | -2.236311 | 0.025331429 | 0.104434316 |
| *COLCA2* | -1.890803 | 0.845607 | -2.236031 | 0.025349755 | 0.104476393 |
| *WDR91* | -1.933382 | 0.864863 | -2.235478 | 0.025385963 | 0.104544492 |
| *UEVLD* | -1.667394 | 0.745879 | -2.235476 | 0.025386088 | 0.104544492 |
| *CCDC15* | -2.549774 | 1.140671 | -2.235329 | 0.025395758 | 0.104544492 |
| *ZNF468* | -1.588566 | 0.710695 | -2.235227 | 0.025402445 | 0.104544492 |
| *ZNF782* | -2.158069 | 0.96551 | -2.235159 | 0.025406903 | 0.104544492 |
| *CCDC189* | -1.569822 | 0.702396 | -2.234952 | 0.025420515 | 0.104559704 |
| *HNRNPU* | 0.944997 | 0.422845 | 2.234855 | 0.025426853 | 0.104559704 |
| *UBTD2* | -1.888784 | 0.845255 | -2.234573 | 0.025445367 | 0.104602406 |
| *ENSG00000257151* | -1.863529 | 0.834016 | -2.234405 | 0.025456455 | 0.104614568 |
| *ZNF793.AS1* | -1.811657 | 0.810906 | -2.234115 | 0.025475524 | 0.104659505 |
| *ZNF486* | -1.824342 | 0.816675 | -2.233865 | 0.025491944 | 0.104693533 |
| *KLHDC1* | -2.109348 | 0.944353 | -2.233644 | 0.025506527 | 0.104720002 |
| *GLMP* | -1.638665 | 0.733808 | -2.233098 | 0.025542487 | 0.10483419 |
| *SLC19A2* | -4.290613 | 1.921982 | -2.23239 | 0.025589226 | 0.104992529 |
| *ZNF446* | -1.725333 | 0.77291 | -2.232257 | 0.025597996 | 0.10499503 |
| *ENSG00000261187* | -4.481142 | 2.007789 | -2.231879 | 0.025622992 | 0.105064065 |
| *PATZ1* | -1.819612 | 0.815367 | -2.231648 | 0.025638234 | 0.105093073 |
| *TRIM68* | -2.369077 | 1.062254 | -2.230236 | 0.025731787 | 0.105442963 |
| *MRM1* | -1.669778 | 0.748826 | -2.22986 | 0.025756766 | 0.105493635 |
| *SEPSECS* | -1.923033 | 0.862423 | -2.229803 | 0.02576055 | 0.105493635 |
| *TDRKH* | -1.971307 | 0.884408 | -2.228957 | 0.025816754 | 0.105690158 |
| *SLAIN1* | -1.947122 | 0.873663 | -2.228687 | 0.025834744 | 0.105730168 |
| *FANCB* | -1.864437 | 0.836706 | -2.228305 | 0.02586017 | 0.105800577 |
| *IVD* | -1.294609 | 0.581135 | -2.227724 | 0.025898936 | 0.105925495 |
| *RPGRIP1L* | -2.286741 | 1.026729 | -2.22721 | 0.025933212 | 0.10603198 |
| *ENSG00000271553* | -3.13323 | 1.407581 | -2.225967 | 0.026016363 | 0.106338165 |

| | | | | | |
|---|---|---|---|---|---|
| *TNFRSF12A* | -1.755952 | 0.789047 | -2.225408 | 0.026053878 | 0.106408709 |
| *ENSG00000261474* | -1.709574 | 0.768226 | -2.225354 | 0.026057501 | 0.106408709 |
| *ZNF556* | -3.53674 | 1.589304 | -2.22534 | 0.026058432 | 0.106408709 |
| *MROH6* | -2.442546 | 1.097859 | -2.224828 | 0.026092804 | 0.106515265 |
| *PELI3* | -1.834054 | 0.824526 | -2.224374 | 0.026123321 | 0.106606018 |
| *TAPT1.AS1* | -4.453344 | 2.002529 | -2.22386 | 0.026157891 | 0.106670375 |
| *CCDC34* | -1.333811 | 0.599789 | -2.2238 | 0.026161921 | 0.106670375 |
| *EXD2* | -2.115701 | 0.95145 | -2.22366 | 0.026171352 | 0.106670375 |
| *ENSG00000235475* | -4.248709 | 1.910695 | -2.223646 | 0.026172253 | 0.106670375 |
| *FAM3C* | 1.069481 | 0.48107 | 2.22313 | 0.026207041 | 0.106778338 |
| *SNX19* | -1.550756 | 0.697729 | -2.222576 | 0.026244381 | 0.106889579 |
| *VPS13B* | -1.512761 | 0.680674 | -2.222446 | 0.026253162 | 0.106889579 |
| *CEBPZOS* | -0.793911 | 0.357238 | -2.222356 | 0.026259265 | 0.106889579 |
| *RAB3A* | -2.129114 | 0.958383 | -2.221569 | 0.026312443 | 0.107072169 |
| *PINK1.AS* | -3.968137 | 1.786526 | -2.221147 | 0.026340986 | 0.107154433 |
| *ACP1* | 1.02923 | 0.463617 | 2.22 | 0.026418738 | 0.107436757 |
| *SEPT7.AS1* | -2.431608 | 1.095438 | -2.219758 | 0.026435198 | 0.107469731 |
| *SLC17A9* | -1.516842 | 0.683438 | -2.219429 | 0.026457542 | 0.107526593 |
| *HOXB3* | -2.112492 | 0.95198 | -2.219051 | 0.026483229 | 0.10758296 |
| *TMEM154* | 1.488494 | 0.670828 | 2.218891 | 0.026494168 | 0.10758296 |
| *USP49* | -2.012143 | 0.906838 | -2.218856 | 0.026496495 | 0.10758296 |
| *ENSG00000278932* | -3.211245 | 1.44761 | -2.218308 | 0.026533824 | 0.107693653 |
| *ENSG00000131845* | -2.03381 | 0.91687 | -2.21821 | 0.026540497 | 0.107693653 |
| *ANGEL1* | -2.092717 | 0.94349 | -2.218059 | 0.026550808 | 0.107701526 |
| *RGS3* | -1.656687 | 0.747108 | -2.217466 | 0.026591276 | 0.107831688 |
| *COLEC12* | -2.135529 | 0.963204 | -2.21711 | 0.026615602 | 0.107896332 |
| *LINC00926* | 0.536461 | 0.24204 | 2.216413 | 0.026663206 | 0.108055265 |
| *C5orf30* | -1.680607 | 0.758328 | -2.216201 | 0.026677715 | 0.108080026 |
| *ZAP70* | -3.456979 | 1.560325 | -2.215551 | 0.026722269 | 0.108226453 |
| *ENSG00000261770* | -2.227093 | 1.005314 | -2.215321 | 0.02673801 | 0.108256127 |
| *AK2* | -0.906774 | 0.409362 | -2.215093 | 0.026753698 | 0.108285572 |
| *KLF11* | -2.35143 | 1.061716 | -2.214744 | 0.026777614 | 0.108316092 |
| *RANGRF* | 1.540976 | 0.695783 | 2.214738 | 0.026778075 | 0.108316092 |
| *RASAL1* | -1.948043 | 0.879641 | -2.214589 | 0.026788279 | 0.108323313 |
| *IGLV2.23* | -4.182205 | 1.888593 | -2.214455 | 0.026797486 | 0.108326503 |
| *LYZ* | -3.513876 | 1.587902 | -2.212905 | 0.026904222 | 0.108696598 |
| *FGF7* | -2.199497 | 0.993983 | -2.212812 | 0.026910636 | 0.108696598 |
| *ENSG00000258056* | -1.828556 | 0.82637 | -2.212757 | 0.026914383 | 0.108696598 |
| *ENSG00000227782* | -2.859634 | 1.292427 | -2.212608 | 0.026924693 | 0.10870412 |
| *NRIP2* | -2.12046 | 0.958427 | -2.212437 | 0.026936468 | 0.108709169 |
| *PAFAH1B2* | -0.698263 | 0.315621 | -2.212345 | 0.026942841 | 0.108709169 |
| *SERTAD3* | -1.066792 | 0.482249 | -2.212119 | 0.026958473 | 0.108727228 |
| *ENSG00000204814* | -4.348872 | 1.966005 | -2.212035 | 0.026964217 | 0.108727228 |
| *ZCCHC7* | 1.067854 | 0.482897 | 2.211352 | 0.027011447 | 0.108778905 |
| *NAGPA* | -1.030511 | 0.466032 | -2.211243 | 0.027018984 | 0.108778905 |
| *ENSG00000272058* | -1.765055 | 0.798225 | -2.211226 | 0.027020178 | 0.108778905 |
| *ARMC9* | -1.739286 | 0.786572 | -2.211225 | 0.027020282 | 0.108778905 |

| | | | | | |
|---|---|---|---|---|---|
| *TPK1* | -1.541782 | 0.697282 | -2.21113 | 0.027026808 | 0.108778905 |
| *LINC00339* | -1.620374 | 0.732831 | -2.211117 | 0.027027758 | 0.108778905 |
| *PINK1* | -1.681167 | 0.760493 | -2.210629 | 0.027061511 | 0.108880694 |
| *ENSG00000176320* | -2.01653 | 0.912298 | -2.210384 | 0.027078512 | 0.108915039 |
| *AK8* | -4.550481 | 2.058853 | -2.210202 | 0.02709117 | 0.108915537 |
| *ZBTB34* | -2.550508 | 1.154004 | -2.210138 | 0.027095565 | 0.108915537 |
| *ENSG00000277959* | -2.334425 | 1.056345 | -2.209908 | 0.027111541 | 0.10892568 |
| *CNNM3* | -1.82805 | 0.827242 | -2.209813 | 0.027118173 | 0.10892568 |
| *GFI1* | -1.900146 | 0.859897 | -2.209736 | 0.027123485 | 0.10892568 |
| *MFSD1* | -1.320571 | 0.59768 | -2.209495 | 0.027140202 | 0.108958809 |
| *TRMO* | -1.559534 | 0.705988 | -2.209011 | 0.027173888 | 0.109060019 |
| *TIPARP.AS1* | -2.444347 | 1.106996 | -2.20809 | 0.027238029 | 0.109283353 |
| *ENSG00000240219* | -4.305805 | 1.950241 | -2.207832 | 0.027255979 | 0.109321285 |
| *ITGA10* | -2.241092 | 1.015187 | -2.207565 | 0.02727462 | 0.1093386 |
| *ENSG00000255139* | -4.258523 | 1.929093 | -2.207527 | 0.027277291 | 0.1093386 |
| *IQCG* | -1.44394 | 0.654426 | -2.206421 | 0.027354509 | 0.109613974 |
| *ENAH* | -4.296185 | 1.947461 | -2.206044 | 0.027380919 | 0.109685643 |
| *CCND2* | -1.897677 | 0.860309 | -2.20581 | 0.027397332 | 0.10968853 |
| *C9orf66* | -2.368924 | 1.073974 | -2.205755 | 0.027401164 | 0.10968853 |
| *ENSG00000278882* | -2.437434 | 1.105161 | -2.205502 | 0.027418899 | 0.10968853 |
| *ABCA2* | -1.840508 | 0.834515 | -2.205483 | 0.027420187 | 0.10968853 |
| *PARK7* | 0.60248 | 0.273181 | 2.205425 | 0.027424264 | 0.10968853 |
| *VPS54* | -1.632184 | 0.740285 | -2.204805 | 0.027467813 | 0.109795328 |
| *ENSG00000268218* | -1.643119 | 0.745246 | -2.204801 | 0.027468031 | 0.109795328 |
| *NOMO2* | -4.454402 | 2.0206 | -2.204495 | 0.027489544 | 0.109847195 |
| *ENSG00000273151* | -4.316768 | 1.95828 | -2.204367 | 0.027498528 | 0.109848979 |
| *F8* | -4.155911 | 1.885714 | -2.203893 | 0.027531887 | 0.109948105 |
| *INAVA* | -4.271941 | 1.939571 | -2.202518 | 0.027628707 | 0.110293249 |
| *SHPK* | -1.800416 | 0.817471 | -2.202423 | 0.027635458 | 0.110293249 |
| *ZMYM1* | -1.905913 | 0.865551 | -2.201966 | 0.027667745 | 0.11038787 |
| *HIVEP3* | -1.826717 | 0.829768 | -2.201479 | 0.027702121 | 0.110490761 |
| *ENSG00000233912* | -4.199874 | 1.908067 | -2.201114 | 0.027727941 | 0.110559471 |
| *ARHGAP18* | -1.646741 | 0.748216 | -2.200891 | 0.027743739 | 0.110588194 |
| *NOD2* | -2.758498 | 1.253468 | -2.200694 | 0.027757722 | 0.110609666 |
| *ENSG00000261644* | -3.128077 | 1.421564 | -2.200447 | 0.027775217 | 0.110634391 |
| *ETFBKMT* | -1.697711 | 0.771633 | -2.200152 | 0.02779611 | 0.110634391 |
| *F2R* | -2.107566 | 0.957927 | -2.200131 | 0.027797573 | 0.110634391 |
| *ECHS1* | 1.319472 | 0.599792 | 2.199883 | 0.027815223 | 0.110634391 |
| *TSGA13* | -4.026031 | 1.830231 | -2.199739 | 0.027825385 | 0.110634391 |
| *ZNF208* | -3.627573 | 1.649119 | -2.199704 | 0.0278279 | 0.110634391 |
| *KHDC4* | -0.7975 | 0.362549 | -2.199702 | 0.027828061 | 0.110634391 |
| *MED14* | -1.367966 | 0.621905 | -2.199636 | 0.027832713 | 0.110634391 |
| *NIPAL4* | -2.057019 | 0.935398 | -2.199085 | 0.027871897 | 0.110733394 |
| *NR1D1* | -1.872346 | 0.851436 | -2.199043 | 0.027874832 | 0.110733394 |
| *BBS10* | -2.054235 | 0.934467 | -2.198295 | 0.027928096 | 0.110877236 |
| *APOBEC3B* | -2.637592 | 1.199858 | -2.198254 | 0.027931038 | 0.110877236 |
| *SELENON* | -1.814173 | 0.82531 | -2.198171 | 0.027936893 | 0.110877236 |

| | | | | | |
|---|---|---|---|---|---|
| *DNAJC5* | -1.685698 | 0.766941 | -2.19795 | 0.027952644 | 0.110905543 |
| *NCKAP5L* | -1.788559 | 0.8143 | -2.196439 | 0.028060528 | 0.111281242 |
| *SIN3B* | -1.449934 | 0.660177 | -2.196281 | 0.028071846 | 0.111281242 |
| *ENSG00000263089* | -1.651616 | 0.752042 | -2.196174 | 0.028079485 | 0.111281242 |
| *ZNF623* | -1.526328 | 0.695005 | -2.19614 | 0.02808193 | 0.111281242 |
| *POLA1* | -1.893721 | 0.862541 | -2.195513 | 0.028126827 | 0.111424839 |
| *ZDBF2* | -1.94023 | 0.883818 | -2.195283 | 0.028143315 | 0.111455845 |
| *THEM6* | -1.640658 | 0.747453 | -2.194999 | 0.028163659 | 0.111479846 |
| *AKAP12* | -4.188918 | 1.908428 | -2.194957 | 0.028166704 | 0.111479846 |
| *MLEC* | 1.586208 | 0.722791 | 2.19456 | 0.028195158 | 0.111558149 |
| *ENSG00000228439* | -1.858938 | 0.847262 | -2.194052 | 0.028231651 | 0.111648794 |
| *NT5DC2* | -2.428167 | 1.106731 | -2.194 | 0.028235422 | 0.111648794 |
| *IKZF1* | 1.380166 | 0.629179 | 2.193601 | 0.028264142 | 0.111704042 |
| *P4HA3* | -3.696845 | 1.685314 | -2.193564 | 0.028266757 | 0.111704042 |
| *C12orf60* | -1.881445 | 0.85814 | -2.19247 | 0.028345613 | 0.11195866 |
| *ZFP30* | -1.60855 | 0.733684 | -2.192428 | 0.028348591 | 0.11195866 |
| *RFLNB* | -4.13859 | 1.888119 | -2.191912 | 0.028385857 | 0.11207144 |
| *KCTD2* | -1.829295 | 0.835095 | -2.190523 | 0.028486333 | 0.112433634 |
| *C9orf64* | -1.669963 | 0.76249 | -2.190144 | 0.028513778 | 0.112474931 |
| *ENSG00000272563* | -4.332924 | 1.97838 | -2.190137 | 0.028514279 | 0.112474931 |
| *TRAF3IP2.AS1* | -1.715951 | 0.783618 | -2.18978 | 0.028540209 | 0.112515385 |
| *TMEM128* | -1.202205 | 0.549014 | -2.189755 | 0.028542023 | 0.112515385 |
| *ENSG00000258646* | -3.263665 | 1.490957 | -2.188972 | 0.028598842 | 0.112685447 |
| *FUT10* | -1.909285 | 0.87225 | -2.18892 | 0.028602679 | 0.112685447 |
| *ARHGEF37* | -2.215587 | 1.012276 | -2.188719 | 0.02861727 | 0.112708423 |
| *PIGO* | -1.739797 | 0.795203 | -2.187865 | 0.028679467 | 0.112918818 |
| *C2orf76* | -1.464405 | 0.669494 | -2.187333 | 0.028718243 | 0.113036902 |
| *SPIN4* | -2.298168 | 1.050738 | -2.187193 | 0.02872845 | 0.113042497 |
| *RHPN1* | -1.920992 | 0.878594 | -2.18644 | 0.028783462 | 0.113224338 |
| *ZNF543* | -1.574771 | 0.72034 | -2.186149 | 0.028804718 | 0.113241087 |
| *CCDC7* | -1.931846 | 0.883679 | -2.186141 | 0.028805322 | 0.113241087 |
| *ENSG00000275636* | -4.232316 | 1.93678 | -2.185233 | 0.028871739 | 0.113371526 |
| *SLC44A5* | -3.810596 | 1.743813 | -2.185209 | 0.028873538 | 0.113371526 |
| *KLHL21* | -1.533088 | 0.70161 | -2.185098 | 0.028881632 | 0.113371526 |
| *PDE8A* | -1.516713 | 0.694119 | -2.185091 | 0.028882167 | 0.113371526 |
| *IDUA* | -4.276794 | 1.957501 | -2.184823 | 0.028901829 | 0.113371526 |
| *PAIP2B* | -1.496315 | 0.684886 | -2.184765 | 0.028906093 | 0.113371526 |
| *CPEB4* | -1.218918 | 0.557919 | -2.18476 | 0.028906414 | 0.113371526 |
| *IGHG4* | -4.428227 | 2.026903 | -2.184725 | 0.02890899 | 0.113371526 |
| *PPP1R12A* | -0.749545 | 0.343259 | -2.183612 | 0.028990761 | 0.113621533 |
| *ENSG00000242349* | -2.188449 | 1.002243 | -2.18355 | 0.028995308 | 0.113621533 |
| *CELSR1* | -2.038413 | 0.933554 | -2.183497 | 0.028999232 | 0.113621533 |
| *MYLPF* | -1.700966 | 0.779469 | -2.182213 | 0.029093848 | 0.113928832 |
| *DUSP4* | -2.46669 | 1.130373 | -2.182192 | 0.029095371 | 0.113928832 |
| *RARA* | -0.787294 | 0.360867 | -2.181672 | 0.029133773 | 0.114010185 |
| *SEMA6A.AS1* | -3.104276 | 1.422889 | -2.18167 | 0.029133869 | 0.114010185 |
| *MINPP1* | -1.526343 | 0.699693 | -2.181447 | 0.029150402 | 0.114040201 |

| | | | | | |
|---|---|---|---|---|---|
| *LRRC56* | -1.981991 | 0.909018 | -2.180364 | 0.02923048 | 0.11431872 |
| *ZNF518B* | -1.553766 | 0.712948 | -2.179353 | 0.029305475 | 0.114571769 |
| *AP3S2* | -1.652546 | 0.758309 | -2.179251 | 0.029312992 | 0.114571769 |
| *NUP107* | -0.910857 | 0.418003 | -2.179069 | 0.029326566 | 0.114590015 |
| *UCKL1.AS1* | -1.845618 | 0.847109 | -2.178725 | 0.029352073 | 0.114654865 |
| *MRFAP1L1* | -1.064344 | 0.488783 | -2.177541 | 0.029440248 | 0.114935099 |
| *NOTCH1* | -1.52755 | 0.701509 | -2.177522 | 0.029441679 | 0.114935099 |
| *LBX2.AS1* | -3.935016 | 1.807247 | -2.177354 | 0.02945418 | 0.114949025 |
| *CCDC168* | -4.39124 | 2.017035 | -2.177077 | 0.02947482 | 0.11497986 |
| *PLTP* | -2.025183 | 0.93026 | -2.177008 | 0.029479953 | 0.11497986 |
| *NSMF* | -2.002232 | 0.919817 | -2.176772 | 0.029497601 | 0.115013827 |
| *TAF1A.AS1* | -2.222845 | 1.021332 | -2.176418 | 0.029524008 | 0.115081917 |
| *MAP3K9* | -1.859707 | 0.854757 | -2.175714 | 0.029576616 | 0.115252067 |
| *MTERF2* | -1.829185 | 0.84083 | -2.175451 | 0.029596357 | 0.115261319 |
| *LINC02397* | 0.764358 | 0.351373 | 2.175349 | 0.029604007 | 0.115261319 |
| *MMP25.AS1* | -1.659697 | 0.762966 | -2.175324 | 0.029605865 | 0.115261319 |
| *ZNF319* | -2.455281 | 1.128969 | -2.174799 | 0.029645217 | 0.115379613 |
| *LINC02384* | -3.436916 | 1.580487 | -2.174592 | 0.02966067 | 0.115404849 |
| *WIZ* | -1.58032 | 0.726799 | -2.174355 | 0.029678465 | 0.115439182 |
| *ENSG00000277324* | -1.868685 | 0.859585 | -2.173939 | 0.02970968 | 0.115525673 |
| *ENSG00000223473* | -1.82022 | 0.837438 | -2.173557 | 0.029738426 | 0.115602516 |
| *SNHG31* | -4.341713 | 1.998009 | -2.173019 | 0.029778858 | 0.115706913 |
| *EIF4G2* | 0.796586 | 0.366605 | 2.172869 | 0.029790154 | 0.115706913 |
| *ZBED6CL* | -2.067621 | 0.951575 | -2.172841 | 0.029792259 | 0.115706913 |
| *EPM2A* | -2.27618 | 1.048039 | -2.171846 | 0.029867278 | 0.115963268 |
| *KATNAL2* | -4.024778 | 1.853308 | -2.171672 | 0.029880425 | 0.115979315 |
| *FAM157C* | -1.977263 | 0.911087 | -2.170223 | 0.029989952 | 0.116369333 |
| *ENSG00000233184* | -1.246637 | 0.574498 | -2.16996 | 0.03000987 | 0.116411517 |
| *SNURF* | -2.208329 | 1.017862 | -2.169576 | 0.030038975 | 0.116454591 |
| *TNFAIP1* | -1.475241 | 0.679968 | -2.169575 | 0.030039076 | 0.116454591 |
| *NDOR1* | -1.933535 | 0.891334 | -2.16926 | 0.030062951 | 0.116484182 |
| *C11orf95* | -1.878609 | 0.866069 | -2.169122 | 0.030073436 | 0.116484182 |
| *FKTN* | -1.886569 | 0.869741 | -2.169116 | 0.030073867 | 0.116484182 |
| *ZNF845* | -1.945695 | 0.897288 | -2.168417 | 0.030126981 | 0.116654789 |
| *POLQ* | -4.143826 | 1.91185 | -2.167442 | 0.030201146 | 0.11689188 |
| *ENSG00000248734* | -4.102174 | 1.892694 | -2.167374 | 0.030206381 | 0.11689188 |
| *MATK* | -2.422888 | 1.118112 | -2.166946 | 0.030238998 | 0.116923833 |
| *IDE* | -1.517785 | 0.700473 | -2.1668 | 0.030250139 | 0.116923833 |
| *DLGAP1.AS2* | -1.98556 | 0.916387 | -2.166727 | 0.030255705 | 0.116923833 |
| *CYP46A1* | -4.385872 | 2.024217 | -2.166701 | 0.030257663 | 0.116923833 |
| *ENSG00000227495* | -3.164644 | 1.460603 | -2.166669 | 0.030260073 | 0.116923833 |
| *ENSG00000275441* | -2.028395 | 0.936251 | -2.166508 | 0.030272397 | 0.116926778 |
| *ENSG00000235609* | -1.875473 | 0.865718 | -2.166379 | 0.030282239 | 0.116926778 |
| *KNL1* | -1.775507 | 0.819602 | -2.166302 | 0.030288098 | 0.116926778 |
| *ADGRB2* | -1.979739 | 0.914015 | -2.165981 | 0.03031264 | 0.116986423 |
| *HDAC7* | 1.477728 | 0.682335 | 2.165694 | 0.030334547 | 0.117035866 |
| *SWT1* | -1.681859 | 0.776766 | -2.165206 | 0.030371886 | 0.117144802 |

| | | | | | |
|---|---|---|---|---|---|
| *ENSG00000272148* | -4.406848 | 2.03545 | -2.165048 | 0.030383998 | 0.117156398 |
| *CLEC11A* | -1.715439 | 0.792462 | -2.164696 | 0.030410993 | 0.11722536 |
| *ZNF708* | -1.24732 | 0.576304 | -2.164344 | 0.030437987 | 0.117294274 |
| *FBXL18* | -1.855897 | 0.857704 | -2.163797 | 0.03047992 | 0.117394592 |
| *HIST1H2BC* | -1.737686 | 0.803084 | -2.163767 | 0.030482267 | 0.117394592 |
| *TOP2A* | -3.284977 | 1.518288 | -2.163606 | 0.030494581 | 0.117406875 |
| *ENSG00000253645* | -1.755676 | 0.811689 | -2.162993 | 0.030541739 | 0.117544707 |
| *VSIG10L* | -2.100382 | 0.9711 | -2.16289 | 0.030549675 | 0.117544707 |
| *PLPP1* | -1.880709 | 0.869578 | -2.162784 | 0.030557787 | 0.117544707 |
| *QRICH2* | -4.421802 | 2.044837 | -2.162422 | 0.030585645 | 0.117616704 |
| *ENO3* | -1.940007 | 0.897423 | -2.161753 | 0.030637191 | 0.117779723 |
| *RWDD3* | -1.931937 | 0.894063 | -2.160852 | 0.030706803 | 0.118012077 |
| *DXO* | -1.258828 | 0.582687 | -2.160385 | 0.030742879 | 0.118115445 |
| *SLC9A3R2* | -3.740323 | 1.73189 | -2.159678 | 0.030797627 | 0.118290467 |
| *DDX11* | -1.633081 | 0.756256 | -2.159429 | 0.030816871 | 0.118329059 |
| *KBTBD2* | -1.47427 | 0.682793 | -2.159174 | 0.030836633 | 0.118369619 |
| *STK17B* | -0.591631 | 0.274049 | -2.158849 | 0.030861896 | 0.118431262 |
| *ENSG00000246790* | -1.637034 | 0.758346 | -2.158689 | 0.030874266 | 0.118443407 |
| *SH2B2* | -3.977486 | 1.84351 | -2.157562 | 0.030961939 | 0.118744344 |
| *MORN1* | -1.841174 | 0.853409 | -2.157434 | 0.030971864 | 0.118747011 |
| *ZNF557* | -1.24194 | 0.575934 | -2.156393 | 0.031052956 | 0.119022457 |
| *IRS1* | -1.694484 | 0.785892 | -2.156128 | 0.031073654 | 0.11906632 |
| *ABCC4* | -4.231682 | 1.962894 | -2.155837 | 0.031096357 | 0.119117839 |
| *LACTB2.AS1* | -2.128899 | 0.987573 | -2.155688 | 0.031108029 | 0.119127087 |
| *NCEH1* | -3.035638 | 1.408622 | -2.155041 | 0.031158634 | 0.119283698 |
| *PDCD6* | -0.575435 | 0.267032 | -2.154928 | 0.031167467 | 0.119283698 |
| *FBXO30* | -1.931521 | 0.896689 | -2.15406 | 0.031235464 | 0.11950839 |
| *FCHO2* | -1.80091 | 0.836129 | -2.153866 | 0.031250709 | 0.119520392 |
| *SPIB* | 0.84196 | 0.390922 | 2.153783 | 0.031257179 | 0.119520392 |
| *IP6K1* | -1.1304 | 0.525035 | -2.153 | 0.031318642 | 0.119685315 |
| *NLRX1* | -1.789172 | 0.831014 | -2.152997 | 0.031318913 | 0.119685315 |
| *HCK* | -1.653967 | 0.768316 | -2.152717 | 0.031340964 | 0.119734022 |
| *ENSG00000272462* | -2.966315 | 1.378681 | -2.15156 | 0.031431995 | 0.120046152 |
| *NELFCD* | -0.779058 | 0.362112 | -2.15143 | 0.031442309 | 0.120049909 |
| *USP30* | -1.889252 | 0.878388 | -2.150818 | 0.031490566 | 0.120180072 |
| *ENSG00000261136* | -1.644078 | 0.764441 | -2.150693 | 0.031500439 | 0.120180072 |
| *IL6ST* | -1.660734 | 0.772204 | -2.150643 | 0.031504421 | 0.120180072 |
| *UCHL5* | -1.159812 | 0.539375 | -2.15029 | 0.03153227 | 0.120247349 |
| *ENSG00000249684* | -3.00391 | 1.397071 | -2.150148 | 0.031543482 | 0.120247349 |
| *ENSG00000258727* | -2.299992 | 1.069732 | -2.150065 | 0.031550093 | 0.120247349 |
| *MAMSTR* | -1.802607 | 0.838711 | -2.149259 | 0.031613893 | 0.120454831 |
| *CLIC3* | 1.727645 | 0.804084 | 2.148587 | 0.031667128 | 0.12060536 |
| *A4GALT* | -1.342852 | 0.625012 | -2.148524 | 0.031672147 | 0.12060536 |
| *KLRF1* | -4.148453 | 1.931036 | -2.148304 | 0.031689628 | 0.120636226 |
| *ENSG00000273240* | -4.234514 | 1.971338 | -2.148041 | 0.031710534 | 0.120680108 |
| *STRADA* | -2.238594 | 1.042347 | -2.147649 | 0.031741669 | 0.120762877 |
| *RINT1* | -1.52724 | 0.711538 | -2.146393 | 0.031841672 | 0.121080239 |

| | | | | | |
|---|---|---|---|---|---|
| *FASN* | -1.93666 | 0.902298 | -2.146365 | 0.031843905 | 0.121080239 |
| *DET1* | -1.78642 | 0.832351 | -2.146233 | 0.031854389 | 0.121080461 |
| *ZNF461* | -1.766818 | 0.823268 | -2.146103 | 0.031864727 | 0.121080461 |
| *MGAT5B* | -1.786111 | 0.832324 | -2.145932 | 0.031878399 | 0.121080461 |
| *NUSAP1* | -1.793404 | 0.835738 | -2.145892 | 0.031881604 | 0.121080461 |
| *OPN3* | -1.153191 | 0.537467 | -2.145605 | 0.031904537 | 0.121116824 |
| *SERPINF2* | -2.45958 | 1.146371 | -2.145536 | 0.031910005 | 0.121116824 |
| *ENSG00000273008* | -2.478225 | 1.155428 | -2.144854 | 0.031964485 | 0.121238853 |
| *LCMT2* | -1.849751 | 0.862441 | -2.144786 | 0.031969966 | 0.121238853 |
| *APH1B* | -1.445737 | 0.674072 | -2.14478 | 0.031970423 | 0.121238853 |
| *LAMC1* | -1.642184 | 0.765807 | -2.144385 | 0.032002029 | 0.121322956 |
| *CHCHD6* | -1.566935 | 0.730779 | -2.144199 | 0.032016967 | 0.121343835 |
| *DIP2C* | -2.769497 | 1.292036 | -2.143513 | 0.0320719 | 0.121516237 |
| *ENSG00000273723* | -1.663805 | 0.776282 | -2.143298 | 0.032089148 | 0.121545795 |
| *ENSG00000246889* | -1.894564 | 0.88414 | -2.142832 | 0.032126615 | 0.121561584 |
| *ENSG00000254741* | -3.924393 | 1.831428 | -2.142805 | 0.03212874 | 0.121561584 |
| *LINC01534* | -1.558881 | 0.727496 | -2.142803 | 0.032128894 | 0.121561584 |
| *PLD3* | -1.118366 | 0.521924 | -2.142776 | 0.032131106 | 0.121561584 |
| *KIAA1958* | -1.806633 | 0.843449 | -2.141959 | 0.032196797 | 0.121766436 |
| *MTMR3* | -1.500427 | 0.700554 | -2.141771 | 0.032211946 | 0.121766436 |
| *URGCP* | -1.656475 | 0.773421 | -2.14175 | 0.032213643 | 0.121766436 |
| *ENSG00000247765* | -3.248253 | 1.516827 | -2.141479 | 0.032235466 | 0.121813138 |
| *SYT11* | -1.723058 | 0.804888 | -2.140744 | 0.032294706 | 0.122001168 |
| *ZNF280C* | -2.911761 | 1.360783 | -2.139769 | 0.032373404 | 0.122262575 |
| *NOXA1* | -2.209803 | 1.032823 | -2.139577 | 0.032389004 | 0.122285598 |
| *NDFIP2* | -4.331754 | 2.024862 | -2.139283 | 0.032412729 | 0.122330609 |
| *EPOR* | -1.621052 | 0.757837 | -2.139051 | 0.03243153 | 0.122330609 |
| *NUDT13* | -2.602743 | 1.216823 | -2.138966 | 0.032438439 | 0.122330609 |
| *IBA57* | -1.663985 | 0.777942 | -2.138958 | 0.03243904 | 0.122330609 |
| *TMEM117* | -1.770305 | 0.827693 | -2.138842 | 0.032448463 | 0.122330609 |
| *DNAJC13* | -1.390334 | 0.650205 | -2.138301 | 0.032492355 | 0.122371969 |
| *ENSG00000273783* | -3.957124 | 1.850645 | -2.138241 | 0.032497176 | 0.122371969 |
| *ZCCHC4* | -1.70999 | 0.799745 | -2.138168 | 0.032503135 | 0.122371969 |
| *CCL4L2* | -2.578931 | 1.206147 | -2.138156 | 0.032504095 | 0.122371969 |
| *CENPV* | -1.760142 | 0.823219 | -2.13812 | 0.032506986 | 0.122371969 |
| *IQCK* | -1.852257 | 0.866632 | -2.137305 | 0.03257318 | 0.12258529 |
| *ENSG00000262165* | -3.154432 | 1.475988 | -2.137166 | 0.032584471 | 0.122591928 |
| *ZNF688* | 1.528191 | 0.715125 | 2.136957 | 0.032601488 | 0.122620096 |
| *ZNF682* | -1.553915 | 0.727252 | -2.136695 | 0.032622831 | 0.122664514 |
| *BLNK* | -0.497953 | 0.233138 | -2.135877 | 0.032689433 | 0.122861221 |
| *HEATR6* | -1.362913 | 0.638154 | -2.135711 | 0.032703015 | 0.122861221 |
| *AMN* | -2.292307 | 1.073355 | -2.135646 | 0.03270825 | 0.122861221 |
| *EDEM3* | -1.471139 | 0.688885 | -2.135536 | 0.032717244 | 0.122861221 |
| *BSPRY* | -4.14359 | 1.940367 | -2.135467 | 0.032722888 | 0.122861221 |
| *ZKSCAN7* | -2.531938 | 1.186148 | -2.134588 | 0.032794717 | 0.123072658 |
| *USP42* | -1.485096 | 0.695744 | -2.134543 | 0.032798332 | 0.123072658 |
| *IQCD* | -1.769518 | 0.82926 | -2.133853 | 0.03285483 | 0.123238727 |

| | | | | | |
|---|---|---|---|---|---|
| *SOX12* | -1.553708 | 0.728152 | -2.133768 | 0.032861745 | 0.123238727 |
| *BRCA1* | -1.680713 | 0.787744 | -2.133577 | 0.032877384 | 0.123261452 |
| *PRR12* | -1.441304 | 0.675784 | -2.132787 | 0.032942238 | 0.123436773 |
| *TSG101* | 1.452527 | 0.681051 | 2.132773 | 0.032943334 | 0.123436773 |
| *PNPLA7* | -1.766546 | 0.828364 | -2.132573 | 0.032959763 | 0.123462378 |
| *ZNF213* | -1.774035 | 0.832059 | -2.132102 | 0.032998471 | 0.123557859 |
| *GNAZ* | -2.121422 | 0.995046 | -2.131985 | 0.033008109 | 0.123557859 |
| *ENSG00000267058* | -1.504011 | 0.705475 | -2.131912 | 0.033014061 | 0.123557859 |
| *IRGQ* | -1.568875 | 0.736122 | -2.131269 | 0.033066952 | 0.123719823 |
| *FLT1* | -4.154092 | 1.949236 | -2.131139 | 0.033077714 | 0.123724111 |
| *ZFAND2B* | -0.949678 | 0.445659 | -2.130952 | 0.033093083 | 0.123739726 |
| *CCNF* | -1.850787 | 0.868566 | -2.130855 | 0.033101122 | 0.123739726 |
| *SLC38A6* | -2.202477 | 1.033805 | -2.130457 | 0.033133874 | 0.123826185 |
| *ZFAND4* | -1.865388 | 0.875973 | -2.129503 | 0.033212686 | 0.12405484 |
| *RBM11* | -3.96476 | 1.861849 | -2.129475 | 0.033215012 | 0.12405484 |
| *PRX* | -4.080301 | 1.916204 | -2.129366 | 0.033223982 | 0.12405484 |
| *ZNF555* | -1.747409 | 0.820751 | -2.129037 | 0.033251205 | 0.124120468 |
| *SSBP3* | -1.694289 | 0.795854 | -2.128895 | 0.033262915 | 0.124128169 |
| *NPIPA1* | -1.697616 | 0.797508 | -2.128651 | 0.033283185 | 0.124167798 |
| *PTPRS* | -2.210845 | 1.038989 | -2.127881 | 0.033346941 | 0.124369593 |
| *CFAP44* | -1.667648 | 0.783898 | -2.127378 | 0.033388703 | 0.124489263 |
| *IGHV3.7* | -3.43029 | 1.612553 | -2.127242 | 0.033400021 | 0.124495384 |
| *ZNF236* | -1.404357 | 0.660288 | -2.126885 | 0.033429635 | 0.124569682 |
| *SLC37A2* | -2.043116 | 0.960799 | -2.126477 | 0.033463548 | 0.124592372 |
| *TAF6L* | -1.552948 | 0.730316 | -2.126405 | 0.033469534 | 0.124592372 |
| *PPP1R12A.AS1* | -4.059071 | 1.908894 | -2.1264 | 0.033469986 | 0.124592372 |
| *SMIM11B* | -4.224969 | 1.986962 | -2.126346 | 0.033474456 | 0.124592372 |
| *VSIG10* | -1.621077 | 0.76245 | -2.126142 | 0.033491421 | 0.124619467 |
| *KDM7A.DT* | -1.752625 | 0.82441 | -2.125915 | 0.033510374 | 0.124653942 |
| *EGR3* | -4.016108 | 1.889732 | -2.125227 | 0.033567704 | 0.124831113 |
| *CD53* | -0.549657 | 0.258685 | -2.124811 | 0.033602389 | 0.124923994 |
| *TTC39B* | -1.811479 | 0.85304 | -2.123559 | 0.033707068 | 0.125276963 |
| *NHLRC4* | -2.354158 | 1.10882 | -2.123121 | 0.033743714 | 0.125349785 |
| *BCL7C* | 1.42615 | 0.671732 | 2.123092 | 0.033746145 | 0.125349785 |
| *DOP1A* | -1.669052 | 0.786253 | -2.122793 | 0.033771164 | 0.125400724 |
| *NPIPB2* | -3.177372 | 1.496857 | -2.122696 | 0.033779351 | 0.125400724 |
| *ZNF526* | -1.233591 | 0.58118 | -2.122563 | 0.033790464 | 0.125405798 |
| *MTMR10* | -1.713066 | 0.807177 | -2.122293 | 0.033813108 | 0.125453652 |
| *PIK3CD.AS1* | -1.807503 | 0.851897 | -2.121739 | 0.033859686 | 0.125590249 |
| *PCBP2* | 0.430627 | 0.203032 | 2.120984 | 0.03392314 | 0.125780228 |
| *ERCC6L2* | -1.387489 | 0.65422 | -2.12083 | 0.033936109 | 0.125780228 |
| *ENSG00000276476* | -2.109643 | 0.994774 | -2.120727 | 0.033944798 | 0.125780228 |
| *LINC01137* | -2.219362 | 1.046541 | -2.120665 | 0.033950006 | 0.125780228 |
| *ENSG00000270761* | -3.030925 | 1.42952 | -2.12024 | 0.033985839 | 0.12587674 |
| *NIP7* | -1.09694 | 0.517438 | -2.119946 | 0.034010591 | 0.125931331 |
| *PPM1F* | -1.440905 | 0.679726 | -2.119833 | 0.034020153 | 0.125931331 |
| *RIOX1* | -1.750814 | 0.826291 | -2.118883 | 0.034100352 | 0.126159481 |

| | | | | | |
|---|---|---|---|---|---|
| *FAM102A* | -1.228629 | 0.579863 | -2.118825 | 0.034105236 | 0.126159481 |
| *ENSG00000276900* | -1.235647 | 0.583209 | -2.118702 | 0.034115686 | 0.126159481 |
| *ENSG00000214248* | -3.01086 | 1.421129 | -2.118639 | 0.034121007 | 0.126159481 |
| *ZNF415* | -2.660785 | 1.256706 | -2.117269 | 0.03423701 | 0.126552027 |
| *ZNF749* | -2.626811 | 1.240878 | -2.116897 | 0.034268585 | 0.126632362 |
| *PSMG3.AS1* | -1.651505 | 0.780316 | -2.116457 | 0.034305956 | 0.126715918 |
| *ZNF85* | -1.33064 | 0.628728 | -2.116399 | 0.034310893 | 0.126715918 |
| *AVPI1* | -1.988171 | 0.93953 | -2.116134 | 0.034333439 | 0.126737987 |
| *HOMEZ* | -1.483553 | 0.701108 | -2.116011 | 0.034343895 | 0.126737987 |
| *PRCC* | -0.990821 | 0.468302 | -2.115776 | 0.034363864 | 0.126737987 |
| *FAM129A* | -2.312949 | 1.093197 | -2.115766 | 0.034364756 | 0.126737987 |
| *GNAQ* | -2.435795 | 1.151268 | -2.11575 | 0.034366118 | 0.126737987 |
| *MOV10* | -1.440506 | 0.681012 | -2.115242 | 0.034409343 | 0.126828591 |
| *MAATS1* | -4.148103 | 1.961065 | -2.115229 | 0.0344104 | 0.126828591 |
| *MAP11* | -1.514982 | 0.716478 | -2.114484 | 0.034473991 | 0.127026588 |
| *SFXN2* | -1.748227 | 0.827016 | -2.113898 | 0.034523993 | 0.127174412 |
| *ENSG00000224046* | -2.788877 | 1.319951 | -2.112864 | 0.034612369 | 0.127463468 |
| *ARRDC3.AS1* | -2.008572 | 0.950856 | -2.112382 | 0.034653663 | 0.127579023 |
| *FAM149B1* | -1.723781 | 0.816255 | -2.111817 | 0.034702114 | 0.127707198 |
| *LRRC3* | -4.561981 | 2.16033 | -2.111706 | 0.034711689 | 0.127707198 |
| *RPL12* | 0.29119 | 0.137898 | 2.111629 | 0.034718254 | 0.127707198 |
| *PLXNC1* | -1.974225 | 0.935026 | -2.111413 | 0.034736838 | 0.127739038 |
| *NMRAL1* | -1.234541 | 0.584815 | -2.110993 | 0.034772954 | 0.127827042 |
| *ENSG00000232611* | -1.567348 | 0.742501 | -2.110903 | 0.034780638 | 0.127827042 |
| *YJEFN3* | -2.040854 | 0.966988 | -2.110527 | 0.034813016 | 0.127909502 |
| *GAB1* | -1.743685 | 0.826248 | -2.110367 | 0.034826782 | 0.127923554 |
| *LINC00342* | -1.597708 | 0.757229 | -2.10994 | 0.034863485 | 0.128021821 |
| *ENSG00000231551* | -1.607372 | 0.762053 | -2.109265 | 0.034921735 | 0.128164622 |
| *PDE9A* | -2.528926 | 1.198997 | -2.109202 | 0.034927129 | 0.128164622 |
| *ABITRAM* | -1.89392 | 0.897957 | -2.109143 | 0.034932255 | 0.128164622 |
| *PIEZO1* | -1.626236 | 0.771153 | -2.108837 | 0.034958613 | 0.128224763 |
| *AOPEP* | -2.021011 | 0.958432 | -2.108664 | 0.034973617 | 0.128225374 |
| *ARNTL2* | -2.13391 | 1.012001 | -2.108605 | 0.03497871 | 0.128225374 |
| *ENSG00000262202* | -1.902697 | 0.902443 | -2.108385 | 0.034997673 | 0.128248899 |
| *ENSG00000260517* | -1.724742 | 0.818072 | -2.1083 | 0.035005062 | 0.128248899 |
| *DBR1* | -1.629402 | 0.773022 | -2.107834 | 0.035045311 | 0.128359813 |
| *PTCD1* | -1.543501 | 0.732337 | -2.107638 | 0.03506235 | 0.128385674 |
| *ENSG00000251615* | -3.132645 | 1.486467 | -2.107443 | 0.035079161 | 0.128405088 |
| *ZBP1* | -1.73934 | 0.82537 | -2.107346 | 0.035087611 | 0.128405088 |
| *MTBP* | -1.899246 | 0.901562 | -2.106618 | 0.035150689 | 0.128599351 |
| *KDM1B* | -1.725491 | 0.819134 | -2.106483 | 0.035162392 | 0.128605599 |
| *SMARCC2* | -0.867227 | 0.411897 | -2.105449 | 0.035252281 | 0.128897728 |
| *ATP6V1E1* | 1.397321 | 0.66378 | 2.105096 | 0.035282978 | 0.128973317 |
| *ATG5* | -0.967715 | 0.459738 | -2.104927 | 0.035297644 | 0.128990282 |
| *SLC16A7* | -1.12092 | 0.532608 | -2.104588 | 0.035327212 | 0.129061682 |
| *NDUFB3* | 0.930301 | 0.442171 | 2.103937 | 0.035383935 | 0.129232213 |
| *APOM* | -1.74843 | 0.83118 | -2.103553 | 0.035417498 | 0.129318089 |

| | | | | | |
|---|---|---|---|---|---|
| *VPS36* | 1.428197 | 0.679032 | 2.103284 | 0.03544093 | 0.129366936 |
| *ZNF117* | -1.679653 | 0.79871 | -2.102956 | 0.035469581 | 0.129434799 |
| *ENSG00000261351* | -1.675508 | 0.796967 | -2.102356 | 0.035522121 | 0.12955407 |
| *ZNF37A* | -0.934602 | 0.444551 | -2.102353 | 0.035522403 | 0.12955407 |
| *RNF43* | -1.443493 | 0.686666 | -2.102175 | 0.035537972 | 0.129574123 |
| *LINC01772* | -1.733743 | 0.824887 | -2.101795 | 0.035571247 | 0.129613999 |
| *PRKRA* | -0.804887 | 0.382959 | -2.101756 | 0.035574696 | 0.129613999 |
| *ENSG00000259826* | -3.282296 | 1.56173 | -2.101705 | 0.035579128 | 0.129613999 |
| *SEC24D* | -1.737454 | 0.826831 | -2.10134 | 0.035611113 | 0.129693799 |
| *EIF3J* | 1.114943 | 0.530664 | 2.101036 | 0.035637842 | 0.129754417 |
| *ENSG00000238009* | -4.038527 | 1.922485 | -2.100681 | 0.035669004 | 0.129831137 |
| *PGBD1* | -1.931264 | 0.919453 | -2.100449 | 0.035689377 | 0.129868556 |
| *KRBA2* | -1.653731 | 0.787461 | -2.10008 | 0.035721778 | 0.129949708 |
| *ERCC8* | -1.882361 | 0.896533 | -2.0996 | 0.035764025 | 0.130066622 |
| *ENSG00000271737* | -1.72356 | 0.821032 | -2.099261 | 0.035793921 | 0.130138564 |
| *ENSG00000259834* | -1.609501 | 0.766805 | -2.098969 | 0.035819603 | 0.130195151 |
| *AMZ1* | -1.808557 | 0.861982 | -2.098137 | 0.035893065 | 0.130425322 |
| *ARHGEF35* | -4.323654 | 2.06083 | -2.098016 | 0.035903705 | 0.130427152 |
| *LMBR1* | -1.49542 | 0.713016 | -2.097316 | 0.035965654 | 0.130615316 |
| *C7orf50* | -0.594603 | 0.283589 | -2.096708 | 0.036019475 | 0.130773866 |
| *CEP85* | -1.82041 | 0.868422 | -2.096228 | 0.036061937 | 0.1308911 |
| *DLAT* | -1.218277 | 0.581252 | -2.095954 | 0.036086302 | 0.130942597 |
| *EXOSC2* | -1.288456 | 0.614864 | -2.095514 | 0.036125296 | 0.131047133 |
| *PDE4A* | -4.07356 | 1.94491 | -2.094472 | 0.036217948 | 0.131338163 |
| *LHFPL4* | -1.77046 | 0.845338 | -2.094382 | 0.036225938 | 0.131338163 |
| *ENSG00000232010* | -3.985134 | 1.9029 | -2.094242 | 0.03623841 | 0.131346371 |
| *CYB5R4* | 1.384491 | 0.661333 | 2.093487 | 0.036305729 | 0.131549585 |
| *ENSG00000260257* | -1.491151 | 0.712317 | -2.09338 | 0.036315219 | 0.131549585 |
| *MASTL* | -1.789921 | 0.855084 | -2.093269 | 0.036325148 | 0.131549585 |
| *SLC30A1* | -1.904101 | 0.909736 | -2.093025 | 0.036346906 | 0.131591345 |
| *LRRC69* | -1.72665 | 0.825064 | -2.092748 | 0.036371612 | 0.131643752 |
| *C17orf49* | 1.521224 | 0.72702 | 2.092409 | 0.036401919 | 0.131716392 |
| *CNIH3* | -3.92268 | 1.87489 | -2.092219 | 0.036418907 | 0.131740816 |
| *ENSG00000271380* | -3.117346 | 1.49077 | -2.091098 | 0.036519284 | 0.132066786 |
| *MMP28* | -4.243475 | 2.029787 | -2.090601 | 0.036563844 | 0.132190777 |
| *ALKBH8* | -1.753385 | 0.838986 | -2.089887 | 0.036627974 | 0.132385434 |
| *ZNF529.AS1* | -1.498154 | 0.717168 | -2.088988 | 0.036708834 | 0.132640427 |
| *ENSG00000273183* | -3.244703 | 1.553682 | -2.088396 | 0.036762144 | 0.132788854 |
| *GPN3* | -1.043919 | 0.499901 | -2.088253 | 0.036775063 | 0.132788854 |
| *URB1* | -1.828515 | 0.875647 | -2.088188 | 0.036780872 | 0.132788854 |
| *STK17A* | 0.57731 | 0.276524 | 2.087738 | 0.036821512 | 0.132898289 |
| *KIAA0753* | -1.62358 | 0.777737 | -2.087569 | 0.036836723 | 0.132915905 |
| *ENSG00000259772* | -1.735253 | 0.831284 | -2.087437 | 0.036848684 | 0.132921788 |
| *MAGED1* | -1.503754 | 0.720684 | -2.086566 | 0.036927384 | 0.133168345 |
| *TXLNG* | -1.333144 | 0.638976 | -2.086376 | 0.03694462 | 0.133193172 |
| *EVA1B* | -1.592504 | 0.763539 | -2.085688 | 0.037006881 | 0.133380263 |
| *ZNF772* | -1.779394 | 0.853413 | -2.085032 | 0.037066419 | 0.133553837 |

| | | | | | |
|---|---|---|---|---|---|
| *DDX60* | -1.748971 | 0.838864 | -2.084928 | 0.037075799 | 0.133553837 |
| *SERHL2* | -1.752476 | 0.840638 | -2.084698 | 0.037096726 | 0.133578789 |
| *IQCE* | -1.560255 | 0.748497 | -2.084518 | 0.037113067 | 0.133578789 |
| *HEXA* | -0.678032 | 0.325272 | -2.084509 | 0.03711387 | 0.133578789 |
| *CCDC71L* | -1.520367 | 0.729428 | -2.084329 | 0.03713026 | 0.133600408 |
| *RANBP10* | -1.480733 | 0.710509 | -2.084045 | 0.037156092 | 0.133616505 |
| *ST3GAL3* | -1.991661 | 0.955682 | -2.08402 | 0.037158369 | 0.133616505 |
| *RBM26* | 1.321821 | 0.63429 | 2.083937 | 0.037165887 | 0.133616505 |
| *CMTM8* | -2.90068 | 1.392366 | -2.083274 | 0.037226225 | 0.133793043 |
| *HMGA1P4* | -1.597353 | 0.76679 | -2.083169 | 0.037235788 | 0.133793043 |
| *NDUFA2* | 0.647855 | 0.311107 | 2.08242 | 0.037304107 | 0.1340011 |
| *LINC01684* | -1.753843 | 0.842357 | -2.082067 | 0.037336349 | 0.134079487 |
| *MMP7* | -2.05915 | 0.989088 | -2.081868 | 0.037354545 | 0.134107404 |
| *SLC25A29* | -2.241887 | 1.077064 | -2.08148 | 0.037389988 | 0.134197204 |
| *FADS2* | -1.631382 | 0.783853 | -2.081234 | 0.037412493 | 0.134240533 |
| *EVI5* | -1.275888 | 0.61312 | -2.080974 | 0.037436305 | 0.134288524 |
| *BRMS1L* | -1.71733 | 0.825478 | -2.080406 | 0.037488265 | 0.134437433 |
| *LRP5L* | -1.578845 | 0.7592 | -2.079617 | 0.037560688 | 0.134659618 |
| *SOGA1* | -1.500125 | 0.721855 | -2.078152 | 0.037695393 | 0.135068595 |
| *ENSG00000273226* | -2.027268 | 0.975517 | -2.078148 | 0.037695759 | 0.135068595 |
| *HAPLN3* | -1.812229 | 0.872194 | -2.077783 | 0.037729338 | 0.135127906 |
| *ENSG00000269973* | -2.378773 | 1.144885 | -2.07774 | 0.037733315 | 0.135127906 |
| *ZNF252P.AS1* | -4.180838 | 2.012658 | -2.077272 | 0.037776492 | 0.135206596 |
| *XRCC3* | -1.669662 | 0.80378 | -2.077263 | 0.03777731 | 0.135206596 |
| *RITA1* | -0.997974 | 0.480483 | -2.077024 | 0.037799365 | 0.135206596 |
| *ENSG00000237481* | -4.206453 | 2.025235 | -2.077019 | 0.037799809 | 0.135206596 |
| *ENSG00000245317* | -1.987047 | 0.956722 | -2.076932 | 0.037807829 | 0.135206596 |
| *ZNF10* | -1.734969 | 0.835546 | -2.076451 | 0.037852288 | 0.135294965 |
| *TIAM1* | -3.683681 | 1.774039 | -2.076437 | 0.037853569 | 0.135294965 |
| *ZNF432* | -1.610691 | 0.775749 | -2.076304 | 0.037865885 | 0.1353014 |
| *CREB3* | -0.977688 | 0.471143 | -2.07514 | 0.037973589 | 0.135648576 |
| *IDNK* | -1.555503 | 0.749816 | -2.074513 | 0.038031698 | 0.135818445 |
| *PER2* | -1.726784 | 0.833133 | -2.072639 | 0.038205905 | 0.136402714 |
| *NFYC.AS1* | -1.846884 | 0.891223 | -2.072304 | 0.038237134 | 0.136476338 |
| *VAV3.AS1* | -1.712542 | 0.826515 | -2.072005 | 0.038264969 | 0.136536681 |
| *ZNF417* | -1.590828 | 0.767852 | -2.071791 | 0.03828493 | 0.136536681 |
| *NMRK1* | -1.622709 | 0.783243 | -2.071781 | 0.038285874 | 0.136536681 |
| *ADCY3* | -1.889211 | 0.911993 | -2.071518 | 0.038310395 | 0.136549783 |
| *NT5E* | -1.491174 | 0.719847 | -2.071514 | 0.038310773 | 0.136549783 |
| *DNAJC28* | -2.366276 | 1.142433 | -2.07126 | 0.038334466 | 0.136596394 |
| *ENSG00000272040* | -3.321862 | 1.60416 | -2.07078 | 0.038379318 | 0.136718351 |
| *THAP8* | -1.622955 | 0.783831 | -2.070544 | 0.038401473 | 0.136759413 |
| *ZNF470* | -1.681506 | 0.812331 | -2.069977 | 0.038454502 | 0.136863872 |
| *ST3GAL6* | -3.192942 | 1.542563 | -2.069894 | 0.038462225 | 0.136863872 |
| *CYTIP* | -0.387679 | 0.187295 | -2.069889 | 0.038462716 | 0.136863872 |
| *ENOSF1* | -1.879917 | 0.908412 | -2.069454 | 0.038503496 | 0.136971105 |
| *ZNF781* | -1.496925 | 0.723601 | -2.068715 | 0.038572826 | 0.137164049 |

| | | | | | |
|---|---|---|---|---|---|
| *SAP30L* | -1.462536 | 0.707001 | -2.068649 | 0.038579054 | 0.137164049 |
| *CLUAP1* | -1.175827 | 0.56862 | -2.067863 | 0.038652907 | 0.137388662 |
| *PSMD9* | -0.855696 | 0.413844 | -2.067678 | 0.038670339 | 0.137412663 |
| *GALNS* | -1.423138 | 0.68833 | -2.067524 | 0.038684812 | 0.13742614 |
| *SLC37A4* | -1.657122 | 0.801832 | -2.066671 | 0.038765212 | 0.137673747 |
| *TCP11L1* | -1.578923 | 0.764048 | -2.066524 | 0.03877905 | 0.137684887 |
| *HLA.G* | -2.116385 | 1.024363 | -2.06605 | 0.038823755 | 0.137805588 |
| *IGHV3.23* | -1.760601 | 0.852278 | -2.065758 | 0.038851358 | 0.137865535 |
| *TYW1B* | -1.711717 | 0.82871 | -2.06552 | 0.038873817 | 0.137907195 |
| *ZNF625* | -1.751556 | 0.848089 | -2.065296 | 0.038894971 | 0.137926488 |
| *ENSG00000124593* | -1.883593 | 0.912047 | -2.065236 | 0.038900694 | 0.137926488 |
| *MPEG1* | 1.439556 | 0.697209 | 2.064742 | 0.038947392 | 0.138054019 |
| *NAPSA* | -1.6706 | 0.809311 | -2.064226 | 0.038996312 | 0.138189354 |
| *F8A1* | -1.713755 | 0.830309 | -2.063998 | 0.039017901 | 0.138227788 |
| *QSOX1* | -1.703532 | 0.825561 | -2.063484 | 0.039066676 | 0.138362488 |
| *NAGLU* | -1.777651 | 0.861546 | -2.063326 | 0.03908162 | 0.138377328 |
| *PPP1R13L* | -1.542528 | 0.747929 | -2.062399 | 0.0391698 | 0.138651392 |
| *ESD* | 0.841589 | 0.408099 | 2.062219 | 0.039186932 | 0.138673887 |
| *DNAJC18* | -1.443627 | 0.700105 | -2.062016 | 0.039206222 | 0.138681523 |
| *DPH6* | -1.521615 | 0.737943 | -2.061969 | 0.039210646 | 0.138681523 |
| *CERNA1* | -1.789546 | 0.868808 | -2.059771 | 0.039420445 | 0.139385235 |
| *COX10* | -1.573194 | 0.76405 | -2.05902 | 0.039492314 | 0.139597059 |
| *ENSG00000260278* | -2.042167 | 0.991864 | -2.058918 | 0.039502051 | 0.139597059 |
| *KDM4A* | -1.501492 | 0.729675 | -2.057755 | 0.039613692 | 0.139938211 |
| *USP37* | -1.455218 | 0.707211 | -2.057685 | 0.039620339 | 0.139938211 |
| *LAMA5* | -1.836913 | 0.89308 | -2.056829 | 0.039702721 | 0.140172338 |
| *NIFK* | 1.383522 | 0.672698 | 2.056678 | 0.039717228 | 0.140172338 |
| *HERC3* | -1.552021 | 0.754633 | -2.056656 | 0.039719309 | 0.140172338 |
| *LMCD1* | -6.280579 | 3.054614 | -2.056096 | 0.039773266 | 0.140243916 |
| *CD109* | -6.280579 | 3.054614 | -2.056096 | 0.039773266 | 0.140243916 |
| *TRDC* | -6.280579 | 3.054614 | -2.056096 | 0.039773266 | 0.140243916 |
| *GPSM2* | -1.475885 | 0.717845 | -2.055993 | 0.039783189 | 0.140243916 |
| *ASH1L* | 0.984831 | 0.47909 | 2.055629 | 0.039818296 | 0.140329227 |
| *HLA.DQB2* | -3.91484 | 1.905017 | -2.055016 | 0.039877451 | 0.140499224 |
| *ENSG00000272894* | -1.71603 | 0.835126 | -2.054817 | 0.039896676 | 0.140528479 |
| *WSB1* | -0.607154 | 0.295518 | -2.054541 | 0.039923348 | 0.14053825 |
| *TMEM143* | -1.476292 | 0.718569 | -2.054491 | 0.039928234 | 0.14053825 |
| *SPACA9* | -1.533115 | 0.746241 | -2.054449 | 0.039932217 | 0.14053825 |
| *ZNF570* | -1.608754 | 0.783135 | -2.054247 | 0.03995175 | 0.140568546 |
| *CKB* | -1.816148 | 0.884255 | -2.053874 | 0.039987887 | 0.140633226 |
| *ENSG00000259793* | -3.82862 | 1.864135 | -2.053832 | 0.039991993 | 0.140633226 |
| *PHLDA1* | -1.965404 | 0.957448 | -2.052752 | 0.040096609 | 0.140946952 |
| *CNKSR2* | -1.742231 | 0.848757 | -2.052685 | 0.040103116 | 0.140946952 |
| *LIG4* | -1.794949 | 0.874516 | -2.052505 | 0.040120608 | 0.140969925 |
| *RAD51B* | -1.725047 | 0.840677 | -2.051974 | 0.040172142 | 0.141091341 |
| *USP10* | -0.927307 | 0.45193 | -2.051883 | 0.040181063 | 0.141091341 |
| *ENSG00000227468* | -2.58772 | 1.261188 | -2.051811 | 0.04018806 | 0.141091341 |

| | | | | | |
|---|---|---|---|---|---|
| *ENSG00000255031* | -1.709613 | 0.833319 | -2.051569 | 0.040211534 | 0.141116881 |
| *PFKL* | -0.738751 | 0.360101 | -2.05151 | 0.040217269 | 0.141116881 |
| *TMEM134* | -0.614751 | 0.299721 | -2.05108 | 0.040259191 | 0.141203419 |
| *EFNB1* | -2.830491 | 1.380033 | -2.051031 | 0.04026388 | 0.141203419 |
| *GRAMD4* | -1.905147 | 0.929032 | -2.05068 | 0.040298093 | 0.141284894 |
| *ZNF684* | -2.019168 | 0.984902 | -2.050122 | 0.040352572 | 0.141437358 |
| *PHLDB2* | -2.363567 | 1.153326 | -2.049348 | 0.040428105 | 0.141663252 |
| *ZKSCAN5* | -1.723302 | 0.840949 | -2.049236 | 0.04043904 | 0.141663252 |
| *ENSG00000241490* | -1.3572 | 0.662381 | -2.048972 | 0.040464849 | 0.141715083 |
| *ENSG00000251364* | -1.551604 | 0.757339 | -2.048756 | 0.040486 | 0.141750576 |
| *ENSG00000233483* | -2.894944 | 1.413393 | -2.048223 | 0.040538134 | 0.141894498 |
| *SAMHD1* | -1.672363 | 0.817357 | -2.046061 | 0.040750344 | 0.142598498 |
| *ERF* | -1.418898 | 0.693563 | -2.04581 | 0.040775068 | 0.14264622 |
| *DPY19L1* | -1.90152 | 0.929663 | -2.045386 | 0.040816855 | 0.142682797 |
| *PQLC2* | -1.424026 | 0.696236 | -2.045322 | 0.040823139 | 0.142682797 |
| *CEP70* | -1.879829 | 0.919129 | -2.045228 | 0.040832339 | 0.142682797 |
| *N6AMT1* | -1.490647 | 0.728862 | -2.045169 | 0.040838154 | 0.142682797 |
| *ENSG00000271133* | -2.192493 | 1.07205 | -2.045141 | 0.040840968 | 0.142682797 |
| *MYLK.AS1* | -1.578509 | 0.771989 | -2.04473 | 0.040881505 | 0.142785648 |
| *ZNF883* | -2.601767 | 1.272534 | -2.044557 | 0.040898598 | 0.142806585 |
| *KCTD21.AS1* | -2.194261 | 1.073459 | -2.044103 | 0.040943409 | 0.142924266 |
| *LILRA4* | -2.255513 | 1.103784 | -2.043438 | 0.041009144 | 0.143114906 |
| *KDM6B* | -1.020661 | 0.499684 | -2.042612 | 0.041090838 | 0.143361119 |
| *GORAB* | -1.531171 | 0.750245 | -2.040896 | 0.041261197 | 0.14391646 |
| *TAFA2* | -2.479523 | 1.215482 | -2.03995 | 0.04135526 | 0.144205456 |
| *TBXA2R* | -1.970026 | 0.966008 | -2.039348 | 0.041415275 | 0.1443756 |
| *INPP5A* | -1.545377 | 0.757855 | -2.039146 | 0.041435494 | 0.144406962 |
| *CDC25B* | -1.108935 | 0.543855 | -2.039026 | 0.041447405 | 0.144409359 |
| *IL10RB.DT* | -1.710027 | 0.83872 | -2.038853 | 0.041464667 | 0.144425212 |
| *ENSG00000261505* | -1.708787 | 0.838152 | -2.038756 | 0.041474404 | 0.144425212 |
| *ZNF862* | -3.649967 | 1.790518 | -2.038498 | 0.041500155 | 0.144475783 |
| *COL19A1* | -0.899673 | 0.441572 | -2.037434 | 0.041606549 | 0.144806997 |
| *EIF5A2* | -1.484365 | 0.728638 | -2.037178 | 0.041632195 | 0.144850958 |
| *ENSG00000102921* | -1.257665 | 0.617385 | -2.037083 | 0.041641695 | 0.144850958 |
| *RTN4IP1* | -2.065556 | 1.014176 | -2.036685 | 0.041681617 | 0.144950638 |
| *NBDY* | -0.593545 | 0.291457 | -2.036476 | 0.041702579 | 0.144956595 |
| *IL15RA* | -1.714406 | 0.841889 | -2.03638 | 0.041712185 | 0.144956595 |
| *ENSG00000261188* | -1.825501 | 0.896518 | -2.036212 | 0.041729054 | 0.144956595 |
| *TNFRSF14.AS1* | -1.391604 | 0.683462 | -2.036109 | 0.041739408 | 0.144956595 |
| *SDR42E2* | -1.889491 | 0.928001 | -2.036087 | 0.041741631 | 0.144956595 |
| *TP53BP2* | -1.472436 | 0.723203 | -2.035994 | 0.041750924 | 0.144956595 |
| *ETS2* | -3.714451 | 1.824735 | -2.035611 | 0.041789408 | 0.145051071 |
| *CHST10* | -2.786278 | 1.368882 | -2.03544 | 0.041806614 | 0.145071655 |
| *CLPTM1L* | -1.399292 | 0.687689 | -2.034773 | 0.041873706 | 0.145265293 |
| *CD79B* | 0.285921 | 0.140541 | 2.03443 | 0.041908229 | 0.145345871 |
| *BBS1* | -1.898713 | 0.934011 | -2.03286 | 0.042066659 | 0.145856023 |
| *MLLT3* | -1.726754 | 0.84954 | -2.032574 | 0.042095535 | 0.145916824 |

| | | | | | |
|---|---|---|---|---|---|
| *MNAT1* | -1.307078 | 0.643183 | -2.032201 | 0.042133297 | 0.146008386 |
| *ENSG00000251661* | -2.197128 | 1.081688 | -2.031203 | 0.042234404 | 0.146319352 |
| *SNTB2* | -1.561506 | 0.769454 | -2.02937 | 0.042420612 | 0.146924903 |
| *KRBA1* | -1.666186 | 0.82109 | -2.029237 | 0.042434135 | 0.146932188 |
| *ARL6* | -1.837231 | 0.905504 | -2.028959 | 0.042462486 | 0.1469908 |
| *ENSG00000259945* | -3.85622 | 1.900798 | -2.028738 | 0.042484997 | 0.147029172 |
| *FANCI* | -2.200113 | 1.084618 | -2.028468 | 0.042512493 | 0.147084765 |
| *NBPF15* | -1.474821 | 0.727109 | -2.028335 | 0.042526094 | 0.14709227 |
| *ENSG00000254469* | -1.564822 | 0.77164 | -2.027917 | 0.042568754 | 0.147200258 |
| *LINC02328* | -1.654117 | 0.815876 | -2.027411 | 0.042620373 | 0.147339157 |
| *ZSCAN12* | -1.732521 | 0.854919 | -2.026533 | 0.042710207 | 0.147610055 |
| *RNF114* | -0.744444 | 0.367382 | -2.026345 | 0.042729407 | 0.147636757 |
| *FGFBP3* | -2.07959 | 1.026454 | -2.025993 | 0.042765458 | 0.147718747 |
| *KIF3C* | -1.852878 | 0.9146 | -2.02589 | 0.042776098 | 0.147718747 |
| *ZNF341.AS1* | -3.68685 | 1.820013 | -2.025728 | 0.04279268 | 0.147736359 |
| *ENSG00000230555* | -1.856392 | 0.916512 | -2.025498 | 0.042816268 | 0.147778144 |
| *LLPH* | -0.749023 | 0.369912 | -2.024866 | 0.042881084 | 0.147962164 |
| *ENSG00000273329* | -1.554382 | 0.767735 | -2.024633 | 0.042905094 | 0.148005321 |
| *IGLV3.21* | -4.102846 | 2.026783 | -2.024314 | 0.04293786 | 0.148078651 |
| *PTPN14* | -1.699533 | 0.839716 | -2.023937 | 0.042976632 | 0.148159051 |
| *MPI* | -1.300564 | 0.642614 | -2.023863 | 0.042984203 | 0.148159051 |
| *JAM3* | -1.511435 | 0.747092 | -2.02309 | 0.043063828 | 0.148343017 |
| *BAIAP2L1* | -5.366324 | 2.65254 | -2.023089 | 0.043063984 | 0.148343017 |
| *ZNF582* | -1.66835 | 0.824687 | -2.02301 | 0.043072162 | 0.148343017 |
| *ZNF841* | -1.751382 | 0.865807 | -2.022831 | 0.043090538 | 0.14836659 |
| *PAXIP1* | -1.884532 | 0.931758 | -2.022556 | 0.043118975 | 0.148390841 |
| *FGF9* | -3.817908 | 1.88768 | -2.02254 | 0.043120646 | 0.148390841 |
| *CAMTA1* | 1.222003 | 0.604242 | 2.022374 | 0.043137766 | 0.148410062 |
| *ENSG00000271200* | -4.005152 | 1.981495 | -2.021278 | 0.043251018 | 0.148756916 |
| *UFSP1* | -3.816779 | 1.888514 | -2.021048 | 0.043274755 | 0.148756916 |
| *PWWP2B* | -1.42638 | 0.705789 | -2.020973 | 0.043282571 | 0.148756916 |
| *WBP1* | -2.057035 | 1.017891 | -2.02088 | 0.043292166 | 0.148756916 |
| *UBP1* | -1.362908 | 0.674427 | -2.020839 | 0.04329639 | 0.148756916 |
| *SLC29A2* | -1.467626 | 0.726329 | -2.020609 | 0.043320271 | 0.148799234 |
| *ZNF609* | -1.375346 | 0.680707 | -2.020466 | 0.043335043 | 0.148810247 |
| *PDK3* | -1.350589 | 0.668548 | -2.020182 | 0.043364546 | 0.14886678 |
| *CPNE5* | 1.389557 | 0.687871 | 2.020084 | 0.043374645 | 0.14886678 |
| *PHF21A* | -0.832018 | 0.412062 | -2.019156 | 0.043470991 | 0.149125893 |
| *LPP.AS2* | -1.598463 | 0.791658 | -2.019134 | 0.043473321 | 0.149125893 |
| *FAM111A.DT* | -1.104128 | 0.546901 | -2.018881 | 0.043499595 | 0.149173772 |
| *ARFGAP3* | -0.870018 | 0.430963 | -2.018776 | 0.043510466 | 0.149173772 |
| *GAB3* | -1.441134 | 0.714017 | -2.018346 | 0.043555279 | 0.149287632 |
| *PLEKHB1* | -1.937543 | 0.960549 | -2.01712 | 0.043683005 | 0.149685547 |
| *ETFB* | 1.201674 | 0.596287 | 2.015261 | 0.043877358 | 0.15029443 |
| *UVSSA* | -1.613316 | 0.800575 | -2.015197 | 0.043884058 | 0.15029443 |
| *PORCN* | -1.540481 | 0.764494 | -2.015033 | 0.043901163 | 0.150313004 |
| *ZNF519* | -1.454662 | 0.721968 | -2.014858 | 0.043919532 | 0.150335893 |

| | | | | | |
|---|---|---|---|---|---|
| *ZFAT* | -1.584065 | 0.786306 | -2.014566 | 0.043950173 | 0.150360693 |
| *TNFRSF10D* | -1.809024 | 0.898062 | -2.014365 | 0.043971183 | 0.150360693 |
| *C8orf82* | -1.761417 | 0.874491 | -2.01422 | 0.043986401 | 0.150360693 |
| *TRIM35* | -1.724951 | 0.85642 | -2.01414 | 0.043994799 | 0.150360693 |
| *ETV7* | -6.153896 | 3.055731 | -2.013887 | 0.044021425 | 0.150360693 |
| *BCL11B* | -6.153896 | 3.055731 | -2.013887 | 0.044021425 | 0.150360693 |
| *COL18A1* | -6.153896 | 3.055731 | -2.013887 | 0.044021425 | 0.150360693 |
| *NCK1.DT* | -1.443993 | 0.717054 | -2.013787 | 0.044031949 | 0.150360693 |
| *GNG10* | -1.897531 | 0.94227 | -2.013787 | 0.04403187 | 0.150360693 |
| *ENSG00000260077* | -1.643325 | 0.816089 | -2.013658 | 0.04404549 | 0.150367025 |
| *CDK5RAP3* | 1.190642 | 0.591365 | 2.01338 | 0.044074669 | 0.15038836 |
| *CACNB1* | -3.854067 | 1.914319 | -2.013283 | 0.044084842 | 0.15038836 |
| *NPAS1* | -3.723344 | 1.849406 | -2.013265 | 0.044086803 | 0.15038836 |
| *RPS28* | -0.109436 | 0.054369 | -2.012845 | 0.044130955 | 0.150499073 |
| *MFGE8* | -2.168808 | 1.077614 | -2.012601 | 0.044156596 | 0.150546614 |
| *ENSG00000272369* | -1.607662 | 0.798881 | -2.012391 | 0.044178727 | 0.150582166 |
| *FAM200A* | -1.419766 | 0.705648 | -2.012002 | 0.044219677 | 0.150649451 |
| *PAFAH2* | -1.771072 | 0.880262 | -2.011982 | 0.044221884 | 0.150649451 |
| *PTRHD1* | -0.651776 | 0.323989 | -2.011721 | 0.044249376 | 0.150703207 |
| *OTUD4* | -1.289203 | 0.6409 | -2.01155 | 0.044267348 | 0.150724522 |
| *LDB1* | -1.552406 | 0.77221 | -2.010343 | 0.044394947 | 0.151118992 |
| *LIMK2* | -1.560663 | 0.776636 | -2.009516 | 0.044482408 | 0.15137666 |
| *CRADD* | -1.310224 | 0.65212 | -2.009177 | 0.044518315 | 0.151458794 |
| *IGF1R* | -1.551731 | 0.772569 | -2.008535 | 0.044586517 | 0.151650732 |
| *ENSG00000218018* | -1.532475 | 0.763052 | -2.00835 | 0.04460612 | 0.151677312 |
| *NAB2* | -1.354669 | 0.674655 | -2.007942 | 0.044649412 | 0.151784408 |
| *HCRT* | -1.776586 | 0.884864 | -2.007749 | 0.044669938 | 0.151814075 |
| *C17orf99* | -1.958514 | 0.975547 | -2.007606 | 0.044685163 | 0.151825717 |
| *SENP3* | -1.513493 | 0.753973 | -2.007357 | 0.044711691 | 0.151855279 |
| *TADA2B* | -1.52872 | 0.761579 | -2.007302 | 0.044717467 | 0.151855279 |
| *ASCC3* | -1.122244 | 0.559509 | -2.005767 | 0.044881152 | 0.152370917 |
| *UBE2Q2* | -1.174291 | 0.585514 | -2.005571 | 0.044902017 | 0.152401544 |
| *C19orf44* | -1.871942 | 0.933568 | -2.005148 | 0.044947229 | 0.152458304 |
| *ENSG00000170846* | -1.608018 | 0.801987 | -2.005043 | 0.044958433 | 0.152458304 |
| *ENSG00000263470* | -5.59627 | 2.791305 | -2.004894 | 0.044974388 | 0.152458304 |
| *ZNF341* | -1.615002 | 0.805589 | -2.004746 | 0.044990258 | 0.152458304 |
| *ENTPD7* | -3.794963 | 1.893475 | -2.004231 | 0.045045287 | 0.152458304 |
| *AGAP4* | -1.539236 | 0.768044 | -2.0041 | 0.045059395 | 0.152458304 |
| *CCR1* | -6.122044 | 3.055329 | -2.003727 | 0.04509936 | 0.152458304 |
| *GIMAP7* | -6.122044 | 3.055329 | -2.003727 | 0.04509936 | 0.152458304 |
| *ANKRD20A4* | -6.122044 | 3.055329 | -2.003727 | 0.045099363 | 0.152458304 |
| *TAL2* | -6.122044 | 3.055329 | -2.003727 | 0.04509936 | 0.152458304 |
| *TNFRSF1A* | -6.122044 | 3.055329 | -2.003727 | 0.04509936 | 0.152458304 |
| *ENSG00000266088* | -6.122044 | 3.055329 | -2.003727 | 0.04509936 | 0.152458304 |
| *RASSF6* | -6.11608 | 3.052772 | -2.003451 | 0.045128902 | 0.152458304 |
| *TMX1* | -0.816628 | 0.407644 | -2.003284 | 0.045146778 | 0.152458304 |
| *EIF4ENIF1* | -1.43388 | 0.715783 | -2.003234 | 0.045152136 | 0.152458304 |

| | | | | | |
|---|---|---|---|---|---|
| *LCP2* | -6.115013 | 3.052761 | -2.003109 | 0.045165609 | 0.152458304 |
| *MSC* | -6.115013 | 3.052761 | -2.003109 | 0.045165609 | 0.152458304 |
| *KLRB1* | -6.115013 | 3.052761 | -2.003109 | 0.045165609 | 0.152458304 |
| *MGST1* | -6.115013 | 3.052761 | -2.003109 | 0.045165609 | 0.152458304 |
| *PTGER2* | -6.115013 | 3.052761 | -2.003109 | 0.045165609 | 0.152458304 |
| *IFT43* | -1.179451 | 0.588815 | -2.003091 | 0.045167565 | 0.152458304 |
| *CEP295* | -1.287061 | 0.642695 | -2.0026 | 0.045220235 | 0.152596056 |
| *BCL2* | -0.656227 | 0.327723 | -2.002383 | 0.045243577 | 0.152603382 |
| *ENSG00000261087* | -1.618697 | 0.808395 | -2.002359 | 0.045246126 | 0.152603382 |
| *TXNL4B* | -1.369054 | 0.683926 | -2.001757 | 0.045310831 | 0.152781567 |
| *MOB3B* | -1.43722 | 0.718109 | -2.001396 | 0.045349773 | 0.152872813 |
| *ENSG00000236935* | -1.60066 | 0.800578 | -1.99938 | 0.045567279 | 0.153514561 |
| *IFT46* | -1.508714 | 0.7546 | -1.999355 | 0.04556992 | 0.153514561 |
| *ZEB1.AS1* | -1.479001 | 0.739808 | -1.999169 | 0.04559008 | 0.153514561 |
| *ZNF629* | -1.686283 | 0.843527 | -1.999086 | 0.045599077 | 0.153514561 |
| *C15orf41* | -2.000199 | 1.00056 | -1.999079 | 0.045599802 | 0.153514561 |
| *ZNF628* | -1.737818 | 0.869511 | -1.998615 | 0.045650016 | 0.153643409 |
| *C19orf47* | -1.871283 | 0.936803 | -1.997519 | 0.045768821 | 0.154002986 |
| *FAM227B* | -1.74243 | 0.872917 | -1.996101 | 0.045922878 | 0.154480961 |
| *ENSG00000272760* | -1.7349 | 0.869197 | -1.99598 | 0.045936151 | 0.154485221 |
| *MIR22HG* | -1.85745 | 0.93079 | -1.995564 | 0.045981437 | 0.154597114 |
| *PLA2G15* | -1.481749 | 0.742663 | -1.995183 | 0.046022941 | 0.154696233 |
| *SPAG4* | -4.765562 | 2.389386 | -1.994472 | 0.046100529 | 0.154916559 |
| *KCNC4* | -1.540428 | 0.772399 | -1.994342 | 0.046114735 | 0.154923837 |
| *EIF4EBP3* | -1.625742 | 0.815838 | -1.992725 | 0.046291548 | 0.155477252 |
| *RC3H1* | -1.046534 | 0.525338 | -1.992114 | 0.046358542 | 0.15566163 |
| *FAM71D* | -1.633819 | 0.820475 | -1.991309 | 0.04644698 | 0.155917895 |
| *INPP4A* | -1.487993 | 0.747286 | -1.991197 | 0.046459284 | 0.155918519 |
| *STX3* | -1.182409 | 0.59408 | -1.99032 | 0.04655574 | 0.1561507 |
| *ZNF704* | -1.611516 | 0.809687 | -1.990296 | 0.046558381 | 0.1561507 |
| *ZNF852* | -1.602658 | 0.80526 | -1.990237 | 0.046564874 | 0.1561507 |
| *ZNF865* | -1.492263 | 0.750071 | -1.989495 | 0.046646629 | 0.156384099 |
| *RSRC1* | -0.801325 | 0.402801 | -1.989381 | 0.046659149 | 0.156385327 |
| *FAHD2B* | -1.577756 | 0.793519 | -1.988303 | 0.046778219 | 0.15674358 |
| *SNX30* | -1.373285 | 0.691124 | -1.987033 | 0.046918778 | 0.157173632 |
| *LINC01597* | -1.701842 | 0.856756 | -1.986378 | 0.046991356 | 0.157375788 |
| *COPE* | 0.762116 | 0.383719 | 1.986131 | 0.0470188 | 0.157411769 |
| *GPRC5C* | -1.482936 | 0.746672 | -1.986061 | 0.047026567 | 0.157411769 |
| *EIF3C* | -1.457287 | 0.733816 | -1.985901 | 0.047044311 | 0.157430209 |
| *KANSL1L* | -1.976312 | 0.995272 | -1.985701 | 0.047066485 | 0.15746346 |
| *ENSG00000262050* | -5.007887 | 2.5223 | -1.985444 | 0.047095046 | 0.157518053 |
| *KCTD19* | -2.271087 | 1.144215 | -1.984844 | 0.047161882 | 0.157700606 |
| *ENSG00000271109* | -1.51982 | 0.765989 | -1.984127 | 0.047241723 | 0.15792654 |
| *ZFP69* | -1.79245 | 0.903549 | -1.983788 | 0.047279426 | 0.158011525 |
| *CD9* | -1.3706 | 0.691023 | -1.983435 | 0.047318879 | 0.158102316 |
| *ZNF549* | -1.510405 | 0.76162 | -1.983149 | 0.047350797 | 0.158128956 |
| *APOBEC3H* | -1.659149 | 0.836632 | -1.983131 | 0.047352853 | 0.158128956 |

| | | | | | |
|---|---|---|---|---|---|
| *ELAC1* | -1.427691 | 0.719953 | -1.983033 | 0.047363721 | 0.158128956 |
| *PFAS* | -1.622361 | 0.818177 | -1.982898 | 0.047378786 | 0.158138222 |
| *VHL* | 1.022232 | 0.515669 | 1.982343 | 0.047440858 | 0.158284164 |
| *POU5F2* | -1.478229 | 0.745719 | -1.982287 | 0.047447114 | 0.158284164 |
| *ENSG00000272189* | -1.76989 | 0.892999 | -1.981962 | 0.047483545 | 0.158352507 |
| *TMCO4* | -1.434328 | 0.723719 | -1.981884 | 0.047492214 | 0.158352507 |
| *DTNB* | -1.36044 | 0.686561 | -1.981528 | 0.047532105 | 0.158444453 |
| *DGKG* | -5.462763 | 2.758099 | -1.980626 | 0.047633231 | 0.158740425 |
| *CD6* | -1.537189 | 0.776159 | -1.980508 | 0.047646498 | 0.158743525 |
| *LANCL2* | -2.160305 | 1.090934 | -1.980234 | 0.047677259 | 0.158794167 |
| *RRAGD* | -2.153531 | 1.087558 | -1.980153 | 0.047686381 | 0.158794167 |
| *ENSG00000271857* | -2.181351 | 1.101744 | -1.979907 | 0.047713931 | 0.158844798 |
| *CCDC74A* | -1.860838 | 0.939978 | -1.979662 | 0.047741548 | 0.158863888 |
| *ICA1* | -3.717185 | 1.8779 | -1.979437 | 0.047766801 | 0.158863888 |
| *STXBP5* | -1.613619 | 0.815198 | -1.97942 | 0.047768752 | 0.158863888 |
| *ERICH6.AS1* | -1.497576 | 0.756597 | -1.979357 | 0.047775794 | 0.158863888 |
| *PRR7* | -1.561626 | 0.788976 | -1.979307 | 0.047781398 | 0.158863888 |
| *AGAP2* | -1.505482 | 0.761017 | -1.978249 | 0.047900567 | 0.159208013 |
| *CHST12* | 1.347497 | 0.681184 | 1.978169 | 0.047909647 | 0.159208013 |
| *ANKLE1* | -1.614922 | 0.816587 | -1.977648 | 0.04796848 | 0.159362364 |
| *GATAD1* | -0.972865 | 0.492068 | -1.977096 | 0.048030822 | 0.159515346 |
| *TMEM185B* | -1.752975 | 0.886675 | -1.97702 | 0.048039323 | 0.159515346 |
| *ENSG00000223821* | -2.168519 | 1.097018 | -1.97674 | 0.048071015 | 0.159544813 |
| *ARMC5* | -1.616575 | 0.817824 | -1.97668 | 0.048077855 | 0.159544813 |
| *FXR2* | -1.376611 | 0.696449 | -1.976613 | 0.048085396 | 0.159544813 |
| *ENSG00000273389* | -3.034104 | 1.53533 | -1.976191 | 0.048133161 | 0.159642623 |
| *GPRASP1* | -4.340134 | 2.196349 | -1.976067 | 0.048147142 | 0.159642623 |
| *BICD2* | -1.187198 | 0.600802 | -1.976024 | 0.048152096 | 0.159642623 |
| *RNF145* | 1.326587 | 0.671429 | 1.975767 | 0.048181188 | 0.159697925 |
| *MAP3K6* | -2.157217 | 1.092156 | -1.975192 | 0.048246359 | 0.159872755 |
| *VPS37B* | 1.308666 | 0.662731 | 1.974656 | 0.048307245 | 0.160002194 |
| *RN7SL832P* | -1.915128 | 0.969913 | -1.974536 | 0.048320821 | 0.160002194 |
| *ZNF823* | -1.59917 | 0.809904 | -1.974519 | 0.048322727 | 0.160002194 |
| *ZFYVE26* | -1.484189 | 0.75174 | -1.974338 | 0.048343288 | 0.160029092 |
| *FAM53C* | -0.973102 | 0.492923 | -1.974145 | 0.048365222 | 0.160060523 |
| *WDR47* | -1.615143 | 0.818566 | -1.973138 | 0.048479874 | 0.16038976 |
| *TAF5L* | -1.562669 | 0.792006 | -1.973052 | 0.048489637 | 0.16038976 |
| *ENSG00000271576* | -1.401514 | 0.710406 | -1.972836 | 0.048514255 | 0.160412463 |
| *ENSG00000269038* | -1.81352 | 0.919327 | -1.972659 | 0.048534407 | 0.160412463 |
| *SLC2A5* | 1.474595 | 0.747533 | 1.972616 | 0.048539305 | 0.160412463 |
| *CXCL8* | -3.793505 | 1.923143 | -1.972554 | 0.048546369 | 0.160412463 |
| *ENSG00000228434* | 1.401521 | 0.710933 | 1.971382 | 0.048680226 | 0.16081347 |
| *NIPA1* | -1.579033 | 0.801107 | -1.971064 | 0.048716606 | 0.160892344 |
| *ENSG00000237513* | -1.367427 | 0.693923 | -1.970575 | 0.048772565 | 0.161035822 |
| *HOXB.AS1* | -1.895265 | 0.961881 | -1.970374 | 0.048795524 | 0.161070293 |
| *ABHD4* | -1.451709 | 0.736991 | -1.969777 | 0.048863911 | 0.161254665 |
| *FAM169A* | -1.825096 | 0.926883 | -1.96907 | 0.048945107 | 0.161481203 |

| | | | | | |
|---|---|---|---|---|---|
| *PIGBOS1* | -0.884351 | 0.449247 | -1.96852 | 0.04900823 | 0.161648011 |
| *ENSG00000281741* | -5.370472 | 2.728388 | -1.968368 | 0.049025723 | 0.16166427 |
| *RAD51C* | -1.040201 | 0.52852 | -1.968141 | 0.049051775 | 0.161708733 |
| *ENSG00000236514* | -1.602306 | 0.81449 | -1.96725 | 0.049154443 | 0.162005691 |
| *ENSG00000270006* | -3.636883 | 1.849348 | -1.966576 | 0.04923212 | 0.162184238 |
| *RASSF1.AS1* | -1.765483 | 0.897751 | -1.966561 | 0.049233826 | 0.162184238 |
| *ACTR3C* | -1.918804 | 0.975959 | -1.966071 | 0.049290413 | 0.162289437 |
| *OTUD1* | -1.498939 | 0.762405 | -1.966066 | 0.049290986 | 0.162289437 |
| *C7orf31* | -1.290706 | 0.656533 | -1.965943 | 0.049305218 | 0.162294767 |
| *ANO6* | -1.517889 | 0.772305 | -1.9654 | 0.049367964 | 0.162454679 |
| *ENSG00000272807* | -1.614009 | 0.821286 | -1.965223 | 0.0493884 | 0.162454679 |
| *SLX4* | -1.562698 | 0.795187 | -1.965195 | 0.049391677 | 0.162454679 |
| *MAP3K2.DT* | -2.27313 | 1.156892 | -1.964859 | 0.049430558 | 0.162541014 |

| Cluster 8 | log2FC | lfcSE | stat | pvalue | padj |
|---|---|---|---|---|---|
| *VPREB3* | 3.10935 | 0.122778 | 25.324971 | < 2.22e-16 | < 2.22e-16 |
| *PLAAT4* | 1.960884 | 0.210039 | 9.3358 | < 2.22e-16 | < 2.22e-16 |
| *TRABD* | 0.640191 | 0.086704 | 7.383653 | 1.54E-13 | 5.58E-10 |
| *DUS2* | 1.582962 | 0.226163 | 6.999212 | 2.57E-12 | 7.00E-09 |
| *RARA.AS1* | 2.696763 | 0.39113 | 6.894805 | 5.39E-12 | 1.17E-08 |
| *PNOC* | 1.732011 | 0.256972 | 6.740081 | 1.58E-11 | 2.87E-08 |
| *MRPS6* | 0.760093 | 0.117674 | 6.459333 | 1.05E-10 | 1.63E-07 |
| *CD1D* | 2.700403 | 0.4383 | 6.161078 | 7.23E-10 | 9.83E-07 |
| *SLC12A4* | 2.625044 | 0.429696 | 6.109078 | 1.00E-09 | 1.21E-06 |
| *AGO1* | 1.263621 | 0.220207 | 5.738339 | 9.56E-09 | 1.04E-05 |
| *MS4A1* | -0.629759 | 0.111336 | -5.656375 | 1.55E-08 | 1.53E-05 |
| *DPEP2* | 0.827363 | 0.152243 | 5.434481 | 5.50E-08 | 4.98E-05 |
| *DENND6B* | 2.202848 | 0.407365 | 5.407549 | 6.39E-08 | 5.35E-05 |
| *CD38* | 2.706547 | 0.502084 | 5.390624 | 7.02E-08 | 5.46E-05 |
| *AHI1* | 1.355534 | 0.253283 | 5.351851 | 8.71E-08 | 6.03E-05 |
| *RMDN2* | 2.072746 | 0.387529 | 5.348621 | 8.86E-08 | 6.03E-05 |
| *SLC5A3* | 1.048462 | 0.206849 | 5.068733 | 4.00E-07 | 0.000256279 |
| *HOXB3* | 3.759693 | 0.747926 | 5.026824 | 4.99E-07 | 0.000301391 |
| *SPRY1* | 1.05233 | 0.21004 | 5.010133 | 5.44E-07 | 0.000311439 |
| *GBP4* | 0.778645 | 0.156072 | 4.989007 | 6.07E-07 | 0.000330126 |
| *SEPTIN9* | 0.46807 | 0.095327 | 4.910132 | 9.10E-07 | 0.000471501 |
| *H3F3A* | 0.411428 | 0.085606 | 4.806085 | 1.54E-06 | 0.000761109 |
| *ICAM2* | 0.879563 | 0.191799 | 4.585852 | 4.52E-06 | 0.002138617 |
| *P2RX5* | 0.903187 | 0.200318 | 4.50877 | 6.52E-06 | 0.002955674 |
| *GRN* | 1.048161 | 0.233912 | 4.480996 | 7.43E-06 | 0.003233049 |
| *RRBP1* | 1.404271 | 0.316165 | 4.44158 | 8.93E-06 | 0.003736555 |
| *CD72* | 0.791993 | 0.183131 | 4.324735 | 1.53E-05 | 0.006050121 |
| *TGFBR2* | -0.693224 | 0.160452 | -4.320443 | 1.56E-05 | 0.006050121 |
| *DTX4* | 1.335757 | 0.314991 | 4.240617 | 2.23E-05 | 0.008362059 |
| *GPR183* | -0.599259 | 0.141671 | -4.229923 | 2.34E-05 | 0.008477309 |
| *SIGIRR* | 0.698336 | 0.166491 | 4.194441 | 2.74E-05 | 0.009599677 |
| *CHST15* | 0.861948 | 0.2061 | 4.182187 | 2.89E-05 | 0.009815529 |
| *TNFRSF13B* | -1.427673 | 0.343628 | -4.154704 | 3.26E-05 | 0.010737547 |
| *FCRL1* | 1.088117 | 0.264866 | 4.108172 | 3.99E-05 | 0.012760532 |
| *ENSG00000271204* | 0.499282 | 0.122284 | 4.082963 | 4.45E-05 | 0.013821053 |
| *RABGAP1L* | 0.680632 | 0.168579 | 4.03747 | 5.40E-05 | 0.01632777 |
| *MAP3K1* | 0.726835 | 0.181541 | 4.003695 | 6.24E-05 | 0.018335742 |
| *DBNL* | 0.605379 | 0.151703 | 3.990543 | 6.59E-05 | 0.018872795 |
| *CD83* | -0.463881 | 0.116666 | -3.976155 | 7.00E-05 | 0.019537152 |
| *GBP7* | 1.592185 | 0.406454 | 3.917256 | 8.96E-05 | 0.024358845 |
| *PLAAT3* | 2.993368 | 0.767086 | 3.902259 | 9.53E-05 | 0.02528678 |
| *CIB1* | 0.421383 | 0.108339 | 3.889487 | 0.00010046 | 0.026020629 |
| *GDPGP1* | 1.710475 | 0.444553 | 3.847626 | 0.00011927 | 0.030174838 |
| *CD22* | 0.77424 | 0.205237 | 3.772418 | 0.00016167 | 0.039973674 |
| *S100A4* | -0.994181 | 0.264569 | -3.757736 | 0.00017146 | 0.041450824 |
| *QPRT* | 3.380332 | 0.901268 | 3.75064 | 0.00017638 | 0.041714691 |

| | | | | | |
|---|---|---|---|---|---|
| *LINC.PINT* | 0.492244 | 0.131834 | 3.733805 | 0.00018861 | 0.043656929 |
| *MPEG1* | 0.976972 | 0.262485 | 3.722011 | 0.00019764 | 0.043976031 |
| *LCAT* | 2.008887 | 0.539811 | 3.721463 | 0.00019807 | 0.043976031 |
| *ZNF563* | 1.12665 | 0.307355 | 3.665627 | 0.00024673 | 0.05368417 |
| *RAB37* | 1.220522 | 0.337376 | 3.617688 | 0.00029725 | 0.063406763 |
| *MAP2K6* | 2.431676 | 0.675407 | 3.600314 | 0.00031783 | 0.066494345 |
| *NCF1* | 0.319515 | 0.090509 | 3.530219 | 0.00041522 | 0.085228775 |
| *ZFP36L1* | -0.418767 | 0.119695 | -3.498614 | 0.00046768 | 0.094220657 |
| *FTH1* | -0.159992 | 0.045805 | -3.492877 | 0.00047785 | 0.094517925 |
| *CD79A* | 0.418568 | 0.120266 | 3.480346 | 0.00050077 | 0.095991472 |
| *SMAD3* | 0.783538 | 0.225207 | 3.479184 | 0.00050294 | 0.095991472 |
| *IGLC2* | -1.1799 | 0.341078 | -3.459327 | 0.00054153 | 0.099765738 |
| *FAM3C* | 0.625017 | 0.180692 | 3.459009 | 0.00054217 | 0.099765738 |
| *NUP210* | 0.642716 | 0.186023 | 3.455031 | 0.00055023 | 0.099765738 |
| *LBR* | 0.490443 | 0.142812 | 3.43418 | 0.00059435 | 0.105165554 |
| *FCER2* | 1.134201 | 0.330487 | 3.431911 | 0.00059934 | 0.105165554 |
| *DPEP3* | 2.093396 | 0.61505 | 3.403619 | 0.000665 | 0.114833074 |
| *GPX1* | 0.403301 | 0.119057 | 3.387456 | 0.00070544 | 0.119913892 |
| *NDUFB9* | 0.518556 | 0.15415 | 3.363963 | 0.00076832 | 0.128593079 |
| *SERPINB6* | 0.777936 | 0.234246 | 3.321025 | 0.00089688 | 0.147835052 |
| *CD82* | -0.527695 | 0.16056 | -3.286587 | 0.00101409 | 0.164661646 |
| *DDIT4* | 1.010938 | 0.309254 | 3.26896 | 0.00107943 | 0.172693583 |
| *CD180* | 0.692766 | 0.217279 | 3.18837 | 0.00143078 | 0.22558555 |
| *NMT2* | -0.571509 | 0.179883 | -3.177122 | 0.00148745 | 0.231170454 |
| *ENSG00000278107* | 2.800645 | 0.889177 | 3.149704 | 0.00163436 | 0.247778386 |
| *ANAPC16* | 0.319636 | 0.101513 | 3.148722 | 0.00163986 | 0.247778386 |
| *CYTIP* | -0.226722 | 0.072412 | -3.131012 | 0.00174205 | 0.25961361 |
| *SESN3* | -0.506037 | 0.162255 | -3.118768 | 0.00181609 | 0.266990049 |
| *CCM2* | -0.49246 | 0.158843 | -3.100293 | 0.00193329 | 0.280430411 |
| *PUS1* | 0.89744 | 0.29066 | 3.087594 | 0.00201784 | 0.288842855 |
| *KCNN1* | 2.582871 | 0.844058 | 3.060065 | 0.00221289 | NA |
| *FAM117A* | 0.579541 | 0.189736 | 3.054455 | 0.0022547 | 0.318556972 |
| *ENSG00000233038* | 4.005717 | 1.312386 | 3.052241 | 0.0022714 | NA |
| *AGPAT5* | 0.736702 | 0.243219 | 3.028968 | 0.00245391 | 0.342257175 |
| *USF3* | -0.513871 | 0.170689 | -3.010564 | 0.00260763 | 0.359094172 |
| *SERPINF1* | -1.352308 | 0.453921 | -2.979174 | 0.00289027 | 0.393040413 |
| *ENSG00000278831* | 2.483788 | 0.838391 | 2.962567 | 0.00305086 | NA |
| *CPNE5* | 0.809985 | 0.274727 | 2.948333 | 0.00319493 | 0.429106562 |
| *PLP2* | -0.531445 | 0.180584 | -2.942924 | 0.00325128 | 0.431349972 |
| *ENSG00000233912* | 2.280094 | 0.778046 | 2.930539 | 0.00338375 | 0.432290104 |
| *ESYT1* | -0.724439 | 0.247604 | -2.925794 | 0.00343579 | 0.432290104 |
| *ENSG00000261448* | 1.725677 | 0.589825 | 2.925746 | 0.00343631 | 0.432290104 |
| *RESF1* | -0.437648 | 0.149757 | -2.922388 | 0.00347359 | 0.432290104 |
| *P2RX5.TAX1BP3* | 1.067236 | 0.365727 | 2.918119 | 0.0035215 | 0.432290104 |
| *CLIC3* | 1.001622 | 0.343256 | 2.918006 | 0.00352278 | 0.432290104 |
| *CPSF4* | -0.817719 | 0.280349 | -2.916791 | 0.00353652 | 0.432290104 |
| *CGGBP1* | -0.451469 | 0.15537 | -2.905766 | 0.00366356 | 0.439786861 |

| | | | | | |
|---|---|---|---|---|---|
| *IFI30* | 0.630148 | 0.217205 | 2.901171 | 0.00371771 | 0.439786861 |
| *USP30.AS1* | 0.809216 | 0.278939 | 2.901051 | 0.00371913 | 0.439786861 |
| *FBXO36* | 2.773995 | 0.957396 | 2.897438 | 0.00376225 | 0.440101788 |
| *IRF4* | 0.714302 | 0.250773 | 2.848407 | 0.00439387 | 0.508520526 |
| *BTBD7* | 0.614928 | 0.216149 | 2.844923 | 0.00444222 | 0.508704433 |
| *PAXX* | 0.500771 | 0.176652 | 2.834789 | 0.0045856 | 0.515158921 |
| *GSTP1* | 0.255967 | 0.090312 | 2.834253 | 0.00459329 | 0.515158921 |
| *NAA80* | -1.237574 | 0.437388 | -2.829467 | 0.00466256 | 0.517591685 |
| *RAB20* | 1.33003 | 0.474126 | 2.805225 | 0.00502815 | 0.552537214 |
| *ADA* | -0.598268 | 0.214155 | -2.793623 | 0.00521212 | 0.567026895 |
| *UBE2E2* | -0.715778 | 0.256598 | -2.789493 | 0.00527906 | 0.568622844 |
| *TUBD1* | 0.815115 | 0.29355 | 2.776747 | 0.00549059 | 0.585608564 |
| *SLC43A2* | 0.334982 | 0.121054 | 2.76722 | 0.00565366 | 0.597147458 |
| *RNF207* | 2.338817 | 0.845431 | 2.766422 | 0.00566752 | NA |
| *IRAK2* | 0.897562 | 0.325104 | 2.760846 | 0.00576518 | 0.600758575 |
| *FCMR* | 0.398427 | 0.144712 | 2.753246 | 0.00590075 | 0.600758575 |
| *SLC45A3* | 1.25908 | 0.457539 | 2.751851 | 0.00592595 | 0.600758575 |
| *LY86* | -0.399226 | 0.145182 | -2.749829 | 0.00596264 | 0.600758575 |
| *IDS* | -0.322417 | 0.117253 | -2.749756 | 0.00596396 | 0.600758575 |
| *EFHD2* | -1.16934 | 0.427635 | -2.734436 | 0.00624873 | 0.623669428 |
| *PARP14* | -0.390875 | 0.143254 | -2.728535 | 0.00636164 | 0.626643802 |
| *CLIC1* | -0.225944 | 0.082858 | -2.726874 | 0.00639374 | 0.626643802 |
| *AGPAT2* | -0.5096 | 0.187333 | -2.720291 | 0.00652246 | 0.629562954 |
| *ENSG00000089127* | 0.750298 | 0.275902 | 2.71944 | 0.00653926 | 0.629562954 |
| *ZC3H12D* | -0.985839 | 0.363633 | -2.711079 | 0.00670647 | 0.639997014 |
| *GABPB1* | 0.328487 | 0.121359 | 2.706729 | 0.00679496 | 0.642803565 |
| *RHOG* | -0.346188 | 0.128302 | -2.698234 | 0.00697083 | 0.653755989 |
| *TRMT1* | 0.501241 | 0.187081 | 2.679276 | 0.00737815 | 0.684338709 |
| *PSAT1* | 2.688362 | 1.004097 | 2.677393 | 0.00741976 | NA |
| *AP3B1* | 0.487233 | 0.18199 | 2.677258 | 0.00742274 | 0.684338709 |
| *TRANK1* | -0.558344 | 0.208927 | -2.672435 | 0.00753029 | 0.684772925 |
| *PTP4A3* | 1.927055 | 0.722055 | 2.668849 | 0.00761116 | 0.684772925 |
| *ZSWIM6* | 1.052991 | 0.395088 | 2.665203 | 0.00769419 | 0.684772925 |
| *LLGL2* | -1.682758 | 0.63179 | -2.663475 | 0.00773382 | 0.684772925 |
| *TBC1D9* | 0.737224 | 0.276828 | 2.663112 | 0.00774217 | 0.684772925 |
| *PTMA* | 0.120179 | 0.045207 | 2.658429 | 0.00785059 | 0.688762361 |
| *ENSG00000245904* | 0.267071 | 0.101055 | 2.642841 | 0.00822135 | 0.715520638 |
| *SYNGR3* | 1.515633 | 0.576124 | 2.63074 | 0.00851992 | 0.73297557 |
| *OAS3* | 1.317763 | 0.501188 | 2.629277 | 0.00855666 | 0.73297557 |
| *ENDOD1* | 0.953719 | 0.364818 | 2.614235 | 0.00894275 | 0.760064182 |
| *GPR160* | 1.010026 | 0.387964 | 2.603403 | 0.00923033 | 0.778424071 |
| *ARHGAP22* | 1.65275 | 0.636945 | 2.594807 | 0.00946439 | 0.792024029 |
| *ZNF674* | 2.009019 | 0.776988 | 2.585649 | 0.00971958 | 0.807170646 |
| *CD24* | -0.437137 | 0.169258 | -2.58267 | 0.00980391 | 0.808005309 |
| *ZNF486* | 1.203446 | 0.46674 | 2.578409 | 0.00992565 | 0.811888422 |
| *KCNC3* | 1.139557 | 0.442493 | 2.575312 | 0.01001497 | 0.813081212 |
| *MYCBP* | -1.134055 | 0.442479 | -2.562959 | 0.01037842 | 0.829405859 |

| | | | | | |
|---|---|---|---|---|---|
| *MID1IP1* | -0.39561 | 0.154373 | -2.562683 | 0.01038669 | 0.829405859 |
| *SLC2A5* | 1.004176 | 0.392275 | 2.559879 | 0.01047087 | 0.829405859 |
| *GUK1* | -0.297871 | 0.116437 | -2.558218 | 0.01052101 | 0.829405859 |
| *POLN* | 1.931495 | 0.756563 | 2.552986 | 0.01068037 | 0.831057765 |
| *CTSH* | -0.356644 | 0.139723 | -2.552518 | 0.01069474 | 0.831057765 |
| *TBK1* | -0.748704 | 0.294073 | -2.545981 | 0.01089713 | 0.836684838 |
| *ENSG00000279278* | 0.795319 | 0.312585 | 2.544328 | 0.01094881 | 0.836684838 |
| *RAB5IF* | -0.388409 | 0.152834 | -2.541371 | 0.01104185 | 0.836684838 |
| *RRS1* | 0.850014 | 0.334869 | 2.538348 | 0.01113772 | 0.836684838 |
| *FAM167A* | 1.287398 | 0.507267 | 2.537909 | 0.0111517 | 0.836684838 |
| *SCN4A* | 2.31149 | 0.912567 | 2.532953 | 0.01131062 | 0.841003302 |
| *WDR33* | 0.483167 | 0.191007 | 2.529577 | 0.01142001 | 0.841003302 |
| *WNT10A* | -1.220609 | 0.482685 | -2.528792 | 0.01144557 | 0.841003302 |
| *NUBP2* | -0.449575 | 0.177939 | -2.526563 | 0.01151848 | 0.841003302 |
| *BIN2* | 0.653349 | 0.259616 | 2.516594 | 0.01184953 | 0.859406838 |
| *SSNA1* | -0.414867 | 0.165024 | -2.513976 | 0.01193786 | 0.860079589 |
| *DIAPH1* | -0.68873 | 0.274506 | -2.508982 | 0.01210798 | 0.866596497 |
| *INTS4* | -0.761124 | 0.305559 | -2.490919 | 0.0127413 | 0.901025223 |
| *LAPTM5* | -0.205583 | 0.082545 | -2.490547 | 0.01275465 | 0.901025223 |
| *MLLT6* | -0.711983 | 0.287205 | -2.479007 | 0.01317486 | 0.919472824 |
| *SNRK* | 0.657204 | 0.265137 | 2.478738 | 0.01318483 | 0.919472824 |
| *MLXIP* | 0.520524 | 0.21039 | 2.474092 | 0.01335752 | 0.925582228 |
| *HS6ST1* | 1.244885 | 0.503678 | 2.471591 | 0.01345133 | 0.926183774 |
| *FSD1L* | -1.124781 | 0.457285 | -2.459695 | 0.01390553 | 0.94927298 |
| *OGFRL1* | -0.796514 | 0.324294 | -2.456147 | 0.01404356 | 0.94927298 |
| *ETFB* | 0.403065 | 0.164113 | 2.456022 | 0.01404844 | 0.94927298 |
| *PKN3* | 2.746113 | 1.11939 | 2.453224 | 0.01415823 | 0.950786535 |
| *PPP1R18* | 0.390186 | 0.159198 | 2.450946 | 0.01424812 | 0.950952733 |
| *GABARAPL2* | 0.296021 | 0.121019 | 2.446082 | 0.01444182 | 0.958003176 |
| *ENSG00000261335* | 1.879734 | 0.769632 | 2.442381 | 0.01459076 | 0.962017216 |
| *PARD6A* | -0.676668 | 0.278192 | -2.43238 | 0.01499995 | 0.983038575 |
| *DENND6A* | 0.884936 | 0.364209 | 2.429746 | 0.01510942 | 0.984283834 |
| *ENSG00000231856* | -1.657265 | 0.683915 | -2.423204 | 0.01538427 | 0.996223211 |
| *IKZF2* | -1.539224 | 0.636319 | -2.418948 | 0.01556545 | 0.999772489 |
| *CD79B* | 0.208247 | 0.08636 | 2.411371 | 0.01589268 | 0.999772489 |
| *P2RY8* | 0.45907 | 0.190718 | 2.407069 | 0.01608115 | 0.999772489 |
| *DMD* | -0.522894 | 0.217267 | -2.406686 | 0.01609802 | 0.999772489 |
| *ELMSAN1* | -0.445787 | 0.185243 | -2.406497 | 0.01610634 | 0.999772489 |
| *C2orf92* | 2.245006 | 0.934052 | 2.403513 | 0.01623837 | 0.999772489 |
| *MRNIP* | 0.479002 | 0.199378 | 2.402485 | 0.0162841 | 0.999772489 |
| *ENSG00000272049* | 2.118701 | 0.882078 | 2.401942 | 0.0163083 | NA |
| *FOXK1* | -0.592735 | 0.246972 | -2.400011 | 0.01639458 | 0.999772489 |
| *SLC50A1* | -0.316645 | 0.132054 | -2.397845 | 0.01649185 | 0.999772489 |
| *DIABLO* | -1.213042 | 0.505963 | -2.397493 | 0.0165077 | 0.999772489 |
| *CD27* | -1.699363 | 0.709067 | -2.396618 | 0.01654715 | 0.999772489 |
| *FOXO3* | 0.826877 | 0.345216 | 2.395249 | 0.01660906 | 0.999772489 |
| *GGA2* | 0.294302 | 0.123066 | 2.391426 | 0.01678308 | 0.999772489 |

| | | | | | |
|---|---|---|---|---|---|
| *RC3H1* | -0.680542 | 0.284782 | -2.389696 | 0.01686233 | 0.999772489 |
| *RUBCN* | -0.500626 | 0.209733 | -2.386973 | 0.01698775 | 0.999772489 |
| *EP400* | 0.567695 | 0.237968 | 2.385592 | 0.01705168 | 0.999772489 |
| *REC8* | 0.831981 | 0.349318 | 2.381731 | 0.01723148 | 0.999772489 |
| *SVOP* | 2.166892 | 0.910081 | 2.380988 | 0.01726627 | 0.999772489 |
| *JDP2* | 1.089947 | 0.457837 | 2.380644 | 0.01728241 | 0.999772489 |
| *ENSG00000230555* | 0.993364 | 0.417708 | 2.378128 | 0.0174008 | 0.999772489 |
| *TNFRSF18* | -1.374085 | 0.580252 | -2.368085 | 0.01788041 | 0.999772489 |
| *NOD2* | 3.335023 | 1.410173 | 2.364974 | 0.01803136 | NA |
| *AFTPH* | -0.493734 | 0.209094 | -2.361305 | 0.01821075 | 0.999772489 |
| *TRAF4* | -0.682176 | 0.289095 | -2.359691 | 0.01829014 | 0.999772489 |
| *SAXO2* | 1.838486 | 0.780008 | 2.357009 | 0.01842281 | NA |
| *ZNF3* | 0.659187 | 0.279883 | 2.355223 | 0.01851159 | 0.999772489 |
| *STK19* | -0.950465 | 0.403595 | -2.354999 | 0.01852277 | 0.999772489 |
| *NFE2L2* | -0.379269 | 0.161066 | -2.354743 | 0.01853551 | 0.999772489 |
| *TUSC1* | 2.005848 | 0.852638 | 2.352521 | 0.01864663 | 0.999772489 |
| *PCBP1* | -0.321614 | 0.136755 | -2.351755 | 0.01868509 | 0.999772489 |
| *PLXNB2* | 1.779051 | 0.757811 | 2.347619 | 0.01889384 | 0.999772489 |
| *NEURL1* | 2.397878 | 1.022081 | 2.346074 | 0.01897233 | 0.999772489 |
| *TRPV3* | 1.736198 | 0.740128 | 2.345807 | 0.01898593 | 0.999772489 |
| *EPC1* | 0.269933 | 0.115214 | 2.342892 | 0.01913492 | 0.999772489 |
| *PAOX* | 0.69918 | 0.298628 | 2.341308 | 0.01921629 | 0.999772489 |
| *TTLL7* | 1.840647 | 0.786695 | 2.339721 | 0.01929816 | 0.999772489 |
| *RASGEF1B* | -0.501229 | 0.214232 | -2.339656 | 0.01930153 | 0.999772489 |
| *PDCD4.AS1* | 0.710941 | 0.304179 | 2.337245 | 0.01942646 | 0.999772489 |
| *MMP11* | 0.961662 | 0.411654 | 2.336091 | 0.01948651 | 0.999772489 |
| *RERE* | -0.414363 | 0.177569 | -2.333535 | 0.01962006 | 0.999772489 |
| *DNM2* | -0.494441 | 0.211899 | -2.333378 | 0.01962831 | 0.999772489 |
| *CD47* | -0.229297 | 0.098269 | -2.333373 | 0.01962859 | 0.999772489 |
| *PKIG* | -0.424758 | 0.182161 | -2.331769 | 0.01971284 | 0.999772489 |
| *PHF20L1* | -0.391715 | 0.168115 | -2.330047 | 0.01980366 | 0.999772489 |
| *MEF2A* | 0.401038 | 0.172263 | 2.328052 | 0.01990934 | 0.999772489 |
| *SLC7A11* | 2.120331 | 0.911629 | 2.32587 | 0.02002551 | NA |
| *FZD3* | 1.398688 | 0.60208 | 2.323093 | 0.02017414 | 0.999772489 |
| *HIRA* | -0.912064 | 0.392633 | -2.322944 | 0.02018219 | 0.999772489 |
| *ZNF233* | 1.914508 | 0.827006 | 2.314986 | 0.0206137 | NA |
| *ZNF726* | 1.183755 | 0.512399 | 2.310223 | 0.0208758 | 0.999772489 |
| *ABCC10* | -1.218913 | 0.528401 | -2.306797 | 0.02106615 | 0.999772489 |
| *TIGAR* | -1.238275 | 0.536891 | -2.30638 | 0.0210894 | 0.999772489 |
| *SNX18* | 1.05339 | 0.456922 | 2.305403 | 0.02114401 | 0.999772489 |
| *ANKRD44* | 0.352077 | 0.152957 | 2.301809 | 0.02134592 | 0.999772489 |
| *MAST4* | -1.066939 | 0.463633 | -2.301257 | 0.0213771 | 0.999772489 |
| *MIB2* | -0.581068 | 0.252531 | -2.300975 | 0.02139302 | 0.999772489 |
| *CDK16* | -0.92236 | 0.401422 | -2.297728 | 0.02157726 | 0.999772489 |
| *BEST1* | -0.811995 | 0.353443 | -2.297388 | 0.02159667 | 0.999772489 |
| *CBWD2* | -0.953512 | 0.415359 | -2.295633 | 0.02169689 | 0.999772489 |
| *LSR* | -1.078956 | 0.470269 | -2.294336 | 0.02177121 | 0.999772489 |

| | | | | | |
|---|---|---|---|---|---|
| *ECHS1* | 0.443461 | 0.193354 | 2.293515 | 0.02181835 | 0.999772489 |
| *MDM4* | -0.267327 | 0.116655 | -2.291598 | 0.02192885 | 0.999772489 |
| *VPS36* | -0.39205 | 0.171256 | -2.289261 | 0.02206418 | 0.999772489 |
| *KCNH4* | 2.741203 | 1.197569 | 2.288973 | 0.02208092 | NA |
| *CKLF* | -0.476098 | 0.208067 | -2.288191 | 0.02212638 | 0.999772489 |
| *MED14* | -1.017595 | 0.444728 | -2.28813 | 0.02212996 | 0.999772489 |
| *RPS6KA4* | -0.916506 | 0.400636 | -2.287631 | 0.022159 | 0.999772489 |
| *PSD3* | -0.932144 | 0.408017 | -2.284569 | 0.02233811 | 0.999772489 |
| *MGAT2* | -0.746387 | 0.32718 | -2.281271 | 0.0225324 | 0.999772489 |
| *CYLD* | -0.366653 | 0.160726 | -2.281236 | 0.02253451 | 0.999772489 |
| *FAM76A* | -0.517618 | 0.226905 | -2.281213 | 0.02253585 | 0.999772489 |
| *PCBP4* | -1.31952 | 0.578978 | -2.279051 | 0.02266405 | 0.999772489 |
| *LINC01679* | 2.89468 | 1.271439 | 2.276696 | 0.02280441 | NA |
| *ENTPD3.AS1* | 0.612566 | 0.269889 | 2.269697 | 0.02322597 | 0.999772489 |
| *SLC25A35* | 1.14633 | 0.505288 | 2.268665 | 0.02328868 | 0.999772489 |
| *TSPYL2* | 0.540492 | 0.238311 | 2.268008 | 0.02332869 | 0.999772489 |
| *ENSG00000258539* | 1.717914 | 0.758193 | 2.265801 | 0.02346356 | 0.999772489 |
| *TKT* | -0.323238 | 0.142727 | -2.264734 | 0.023529 | 0.999772489 |
| *CYBA* | 0.223631 | 0.098769 | 2.264179 | 0.02356309 | 0.999772489 |
| *SNX6* | -0.363868 | 0.16078 | -2.263139 | 0.02362711 | 0.999772489 |
| *COQ2* | -1.170788 | 0.517384 | -2.262898 | 0.02364196 | 0.999772489 |
| *KDM6B* | -0.540082 | 0.238695 | -2.262645 | 0.02365758 | 0.999772489 |
| *PRDM1* | 2.268816 | 1.003283 | 2.261391 | 0.02373505 | NA |
| *POU2AF1* | 0.251709 | 0.111331 | 2.260906 | 0.02376509 | 0.999772489 |
| *GTF2A2* | -0.325222 | 0.144082 | -2.257203 | 0.02399539 | 0.999772489 |
| *NAMPT* | 0.56024 | 0.24834 | 2.255941 | 0.0240743 | 0.999772489 |
| *STAG2* | 0.342411 | 0.151792 | 2.255799 | 0.02408319 | 0.999772489 |
| *XYLT1* | 0.692998 | 0.307268 | 2.255353 | 0.02411117 | 0.999772489 |
| *CBX8* | 1.255189 | 0.55657 | 2.25522 | 0.02411954 | 0.999772489 |
| *TRIM38* | 0.29782 | 0.132153 | 2.253605 | 0.02422105 | 0.999772489 |
| *ZNF318* | -0.473819 | 0.210439 | -2.251579 | 0.02434888 | 0.999772489 |
| *TOMM40L* | 1.763705 | 0.783367 | 2.251442 | 0.02435758 | 0.999772489 |
| *PHC2* | -1.70886 | 0.759256 | -2.250703 | 0.02440436 | 0.999772489 |
| *LIG3* | 1.067195 | 0.474448 | 2.249339 | 0.02449091 | 0.999772489 |
| *ABR* | 0.693403 | 0.308377 | 2.248555 | 0.02454083 | 0.999772489 |
| *HSH2D* | 0.365652 | 0.162756 | 2.246632 | 0.02466354 | 0.999772489 |
| *WASF2* | 0.301539 | 0.134519 | 2.241618 | 0.02498607 | 0.999772489 |
| *DENND5A* | 0.798262 | 0.356221 | 2.240916 | 0.0250315 | 0.999772489 |
| *EHD4* | -0.649059 | 0.290012 | -2.238042 | 0.02521835 | 0.999772489 |
| *PATJ* | -0.979504 | 0.437719 | -2.237745 | 0.02523766 | 0.999772489 |
| *ZDHHC7* | -1.065171 | 0.476318 | -2.23626 | 0.02533477 | 0.999772489 |
| *ENSG00000231769* | 1.842524 | 0.824591 | 2.23447 | 0.02545216 | NA |
| *BCCIP* | 0.502405 | 0.224975 | 2.233158 | 0.02553855 | 0.999772489 |
| *FUT2* | 2.182664 | 0.977646 | 2.232572 | 0.02557722 | NA |
| *ENSG00000226380* | 0.678649 | 0.304294 | 2.230241 | 0.02573148 | 0.999772489 |
| *METAP1D* | 0.901626 | 0.404742 | 2.22766 | 0.0259032 | 0.999772489 |
| *ATP6V1A* | -0.583422 | 0.262843 | -2.21966 | 0.02644185 | 0.999772489 |

| | | | | | |
|---|---|---|---|---|---|
| *CD44* | -0.34962 | 0.157572 | -2.218793 | 0.0265008 | 0.999772489 |
| *NUDT8* | -1.359355 | 0.612924 | -2.217818 | 0.02656724 | 0.999772489 |
| *CETP* | -1.554217 | 0.701749 | -2.214776 | 0.02677542 | 0.999772489 |
| *TMCC1.AS1* | 1.873398 | 0.84677 | 2.212405 | 0.02693871 | NA |
| *ABCA2* | 1.797396 | 0.815231 | 2.204769 | 0.02747029 | 0.999772489 |
| *TOP1MT* | -0.67898 | 0.307997 | -2.204503 | 0.02748898 | 0.999772489 |
| *PXDC1* | 2.397443 | 1.087534 | 2.204476 | 0.0274909 | 0.999772489 |
| *LINC00896* | 2.041739 | 0.927007 | 2.202507 | 0.02762951 | 0.999772489 |
| *ENSG00000228201* | 2.093368 | 0.950971 | 2.201296 | 0.02771509 | NA |
| *NFATC3* | 0.445012 | 0.202266 | 2.20013 | 0.0277977 | 0.999772489 |
| *ENSG00000254859* | 2.831145 | 1.288596 | 2.197077 | 0.02801495 | NA |
| *SYTL1* | -0.362192 | 0.165045 | -2.194502 | 0.02819936 | 0.999772489 |
| *TAPT1.AS1* | 1.584252 | 0.722407 | 2.19302 | 0.02830594 | NA |
| *POU2F2* | -0.259536 | 0.118402 | -2.191985 | 0.02838062 | 0.999772489 |
| *ATP11C* | 0.664804 | 0.30334 | 2.191617 | 0.02840716 | 0.999772489 |
| *C1orf131* | -0.439392 | 0.200648 | -2.18986 | 0.02853443 | 0.999772489 |
| *ARHGAP15* | 0.290181 | 0.132558 | 2.189093 | 0.02859009 | 0.999772489 |
| *SYNE3* | -1.027197 | 0.469329 | -2.188649 | 0.02862237 | 0.999772489 |
| *TXLNG* | 0.607086 | 0.27767 | 2.186355 | 0.02878967 | 0.999772489 |
| *MRPS36* | 0.348482 | 0.159414 | 2.186018 | 0.02881431 | 0.999772489 |
| *NCF4* | -0.414935 | 0.189915 | -2.184841 | 0.02890052 | 0.999772489 |
| *CTSA* | 0.558363 | 0.255655 | 2.184045 | 0.02895891 | 0.999772489 |
| *PLEKHO1* | 0.399209 | 0.182808 | 2.183762 | 0.02897971 | 0.999772489 |
| *VPS18* | -0.961088 | 0.440283 | -2.182886 | 0.0290442 | 0.999772489 |
| *SELENOM* | -0.815375 | 0.373937 | -2.180516 | 0.02921922 | 0.999772489 |
| *TMEM242* | -0.59041 | 0.270819 | -2.180089 | 0.02925085 | 0.999772489 |
| *SIDT1* | -0.487257 | 0.223582 | -2.179323 | 0.02930769 | 0.999772489 |
| *TRIP10* | -0.84873 | 0.389471 | -2.179191 | 0.02931752 | 0.999772489 |
| *LINC002481* | 2.479693 | 1.138112 | 2.178777 | 0.02934822 | 0.999772489 |
| *FUOM* | 0.613845 | 0.281863 | 2.177812 | 0.02942007 | 0.999772489 |
| *MZB1* | -0.604746 | 0.277925 | -2.175929 | 0.02956057 | 0.999772489 |
| *ENSG00000271870* | -1.492259 | 0.686243 | -2.174535 | 0.029665 | 0.999772489 |
| *CPT2* | 0.674598 | 0.310325 | 2.17384 | 0.02971716 | 0.999772489 |
| *LRRFIP1* | -0.309855 | 0.142601 | -2.172877 | 0.02978956 | 0.999772489 |
| *UGCG* | 0.643709 | 0.296269 | 2.172714 | 0.02980185 | 0.999772489 |
| *CXCR4* | 0.127097 | 0.058624 | 2.167994 | 0.03015914 | 0.999772489 |
| *RINL* | 0.465686 | 0.214812 | 2.167876 | 0.03016811 | 0.999772489 |
| *PSMA3* | 0.300135 | 0.138453 | 2.167779 | 0.0301755 | 0.999772489 |
| *ARPC1B* | -0.192366 | 0.088795 | -2.166394 | 0.03028106 | 0.999772489 |
| *SUSD1* | 1.83975 | 0.849495 | 2.165697 | 0.03033438 | NA |
| *NICN1* | 0.801354 | 0.371225 | 2.158673 | 0.03087556 | 0.999772489 |
| *XBP1* | 0.533119 | 0.246994 | 2.158426 | 0.03089471 | 0.999772489 |
| *LARGE1* | 0.973519 | 0.452254 | 2.152593 | 0.03135064 | 0.999772489 |
| *PRR5* | 1.900181 | 0.882932 | 2.152124 | 0.03138755 | 0.999772489 |
| *VIM* | -0.577563 | 0.268378 | -2.152053 | 0.03139317 | 0.999772489 |
| *ENSG00000273384* | 1.459387 | 0.678447 | 2.151071 | 0.03147059 | 0.999772489 |
| *IRF1* | 0.257688 | 0.12 | 2.147398 | 0.03176161 | 0.999772489 |

| | | | | | |
|---|---|---|---|---|---|
| *AFG3L2* | -0.716799 | 0.33423 | -2.144629 | 0.03198253 | 0.999772489 |
| *UBE2G1* | 0.352671 | 0.1646 | 2.142594 | 0.03214569 | 0.999772489 |
| *HTR3A* | 2.143782 | 1.001448 | 2.140682 | 0.03229969 | 0.999772489 |
| *SLC45A4* | 3.213488 | 1.501714 | 2.139879 | 0.03236452 | NA |
| *ERGIC3* | -0.29267 | 0.136854 | -2.138555 | 0.03247176 | 0.999772489 |
| *HMOX1* | 0.872271 | 0.408037 | 2.137725 | 0.0325391 | 0.999772489 |
| *PRPF4B* | -0.393567 | 0.184165 | -2.13704 | 0.03259474 | 0.999772489 |
| *STX10* | -0.319896 | 0.149704 | -2.136862 | 0.0326092 | 0.999772489 |
| *NMNAT3* | 1.468994 | 0.687794 | 2.135806 | 0.03269524 | 0.999772489 |
| *CNBP* | 0.184998 | 0.086665 | 2.134643 | 0.03279022 | 0.999772489 |
| *UFSP1* | 1.805155 | 0.845871 | 2.134077 | 0.03283644 | NA |
| *PALLD* | 2.829896 | 1.326777 | 2.13291 | 0.03293209 | NA |
| *TMEM38B* | 0.603697 | 0.283714 | 2.127834 | 0.03335082 | 0.999772489 |
| *WDR19* | -0.943778 | 0.443732 | -2.126911 | 0.03342745 | 0.999772489 |
| *LINC01136* | 1.866732 | 0.879079 | 2.123508 | 0.03371128 | 0.999772489 |
| *GPT2* | 0.84609 | 0.398902 | 2.12105 | 0.03391759 | 0.999772489 |
| *LDHA* | -0.424298 | 0.200129 | -2.12012 | 0.03399596 | 0.999772489 |
| *TLR10* | -0.534576 | 0.252659 | -2.115799 | 0.03436194 | 0.999772489 |
| *TAF4B* | 0.966329 | 0.457065 | 2.114204 | 0.03449784 | 0.999772489 |
| *PDCD4* | 0.42361 | 0.200367 | 2.114166 | 0.03450112 | 0.999772489 |
| *JAK1* | 0.241805 | 0.114471 | 2.112374 | 0.03465441 | 0.999772489 |
| *ARMC2* | 1.617436 | 0.766058 | 2.111374 | 0.03474018 | NA |
| *FADS3* | -0.340359 | 0.161353 | -2.10941 | 0.03490924 | 0.999772489 |
| *TMEM120B* | -0.874515 | 0.414874 | -2.107906 | 0.03503913 | 0.999772489 |
| *RBMS1* | -0.64534 | 0.306229 | -2.10738 | 0.03508463 | 0.999772489 |
| *ENTPD5* | 1.154427 | 0.547988 | 2.106665 | 0.0351466 | 0.999772489 |
| *ENSG00000259793* | 1.894036 | 0.899835 | 2.104871 | 0.0353025 | NA |
| *KANSL2* | -0.650399 | 0.309408 | -2.102078 | 0.03554642 | 0.999772489 |
| *FYTTD1* | -0.400895 | 0.190716 | -2.102056 | 0.03554834 | 0.999772489 |
| *SEMA7A* | -1.025062 | 0.48765 | -2.102047 | 0.03554915 | 0.999772489 |
| *LINC02158* | 1.835459 | 0.873922 | 2.100254 | 0.0357065 | NA |
| *BMF* | -0.746332 | 0.355377 | -2.100115 | 0.0357187 | 0.999772489 |
| *ENSG00000253535* | 0.959535 | 0.457138 | 2.099003 | 0.03581667 | 0.999772489 |
| *NBN* | -0.506698 | 0.241478 | -2.098322 | 0.03587668 | 0.999772489 |
| *SLC41A2* | 1.306207 | 0.622604 | 2.097973 | 0.03590749 | 0.999772489 |
| *SLC33A1* | -0.742952 | 0.354308 | -2.096908 | 0.03600168 | 0.999772489 |
| *IQCC* | 1.701558 | 0.812661 | 2.09381 | 0.03627688 | 0.999772489 |
| *ZNF706* | 0.220724 | 0.105423 | 2.093697 | 0.03628697 | 0.999772489 |
| *TRADD* | -0.547967 | 0.261828 | -2.09285 | 0.03636255 | 0.999772489 |
| *BRMS1* | -0.387444 | 0.185216 | -2.091846 | 0.03645227 | 0.999772489 |
| *ENSG00000272817* | 2.03992 | 0.975483 | 2.09119 | 0.03651102 | 0.999772489 |
| *ENSG00000236723* | 1.775336 | 0.849459 | 2.089962 | 0.03662123 | NA |
| *IRAK1BP1* | 1.2773 | 0.61128 | 2.08955 | 0.03665821 | 0.999772489 |
| *CCDC144A* | 1.454103 | 0.696395 | 2.088045 | 0.03679375 | 0.999772489 |
| *MAPKAPK5* | -0.539244 | 0.258465 | -2.086333 | 0.0369485 | 0.999772489 |
| *CLCN5* | 1.711274 | 0.820875 | 2.084694 | 0.03709709 | NA |
| *CD99* | -0.447503 | 0.214857 | -2.082794 | 0.03727002 | 0.999772489 |

| | | | | | |
|---|---|---|---|---|---|
| *ZNF800* | -0.354683 | 0.170428 | -2.081125 | 0.03742242 | 0.999772489 |
| *SLC25A45* | 0.763219 | 0.366791 | 2.080802 | 0.03745199 | 0.999772489 |
| *MED28* | -0.25752 | 0.123774 | -2.080569 | 0.03747341 | 0.999772489 |
| *CIRBP* | 0.168797 | 0.081195 | 2.078898 | 0.03762673 | 0.999772489 |
| *QSOX2* | -0.690383 | 0.332313 | -2.077506 | 0.03775491 | 0.999772489 |
| *CDCA4* | -0.692 | 0.333315 | -2.076116 | 0.03788325 | 0.999772489 |
| *SEL1L3* | 0.685004 | 0.329967 | 2.075979 | 0.0378959 | 0.999772489 |
| *C8orf58* | -1.084731 | 0.522906 | -2.074429 | 0.03803946 | 0.999772489 |
| *GIGYF1* | -0.414052 | 0.19962 | -2.074199 | 0.03806084 | 0.999772489 |
| *ENSG00000268439* | 1.134628 | 0.54839 | 2.069018 | 0.03854444 | 0.999772489 |
| *SH3BGRL3* | -0.161745 | 0.078274 | -2.06639 | 0.03879167 | 0.999772489 |
| *ENSG00000263272* | -1.049571 | 0.508066 | -2.065816 | 0.03884587 | 0.999772489 |
| *KBTBD6* | -0.633443 | 0.306689 | -2.065424 | 0.03888294 | 0.999772489 |
| *TRIO* | -0.713391 | 0.345863 | -2.06264 | 0.03914681 | 0.999772489 |
| *GLIPR2* | 0.503158 | 0.244337 | 2.059284 | 0.03946705 | 0.999772489 |
| *PLEKHG1* | -0.394806 | 0.191976 | -2.05654 | 0.03973051 | 0.999772489 |
| *OSTM1* | 0.477863 | 0.23252 | 2.055148 | 0.03986475 | 0.999772489 |
| *ODR4* | 0.687911 | 0.334726 | 2.055146 | 0.03986495 | 0.999772489 |
| *APMAP* | 0.589389 | 0.286803 | 2.05503 | 0.03987611 | 0.999772489 |
| *FAM227A* | 1.624411 | 0.790491 | 2.054939 | 0.03988493 | NA |
| *ECH1* | 0.260191 | 0.126667 | 2.054133 | 0.03996281 | 0.999772489 |
| *FAM177A1* | -0.397143 | 0.193356 | -2.053949 | 0.03998063 | 0.999772489 |
| *SMAGP* | 0.44269 | 0.215564 | 2.053634 | 0.04001111 | 0.999772489 |
| *DAD1* | 0.231633 | 0.112797 | 2.053529 | 0.04002127 | 0.999772489 |
| *SH3BP1* | -0.827014 | 0.403509 | -2.049556 | 0.04040781 | 0.999772489 |
| *LINC01521* | 1.437427 | 0.70136 | 2.049485 | 0.04041467 | 0.999772489 |
| *ENSG00000272379* | 1.513524 | 0.738728 | 2.048823 | 0.04047941 | 0.999772489 |
| *ASNSD1* | 0.389146 | 0.190072 | 2.047357 | 0.04062307 | 0.999772489 |
| *RBBP7* | -0.413458 | 0.202048 | -2.046331 | 0.04072384 | 0.999772489 |
| *TMEM97* | -1.057121 | 0.516689 | -2.045954 | 0.04076091 | 0.999772489 |
| *S100A6* | -0.412724 | 0.202053 | -2.042655 | 0.04108662 | 0.999772489 |
| *DDX49* | 0.526087 | 0.257896 | 2.039914 | 0.04135886 | 0.999772489 |
| *CHP1* | -0.386487 | 0.189622 | -2.038197 | 0.04153022 | 0.999772489 |
| *CHD1* | 0.306114 | 0.150191 | 2.038174 | 0.04153256 | 0.999772489 |
| *MAN2C1* | 1.0556 | 0.518221 | 2.036969 | 0.0416531 | 0.999772489 |
| *SNRPC* | 0.268387 | 0.131824 | 2.035954 | 0.04175496 | 0.999772489 |
| *ADAMDEC1* | 1.949154 | 0.957938 | 2.03474 | 0.04187702 | NA |
| *DNAJC14* | 0.6962 | 0.342795 | 2.030954 | 0.0422597 | 0.999772489 |
| *GNAI2* | -0.334521 | 0.164735 | -2.030655 | 0.04228998 | 0.999772489 |
| *C15orf39* | 0.684695 | 0.337259 | 2.030177 | 0.04233855 | 0.999772489 |
| *UBR1* | -0.643617 | 0.317052 | -2.030004 | 0.04235614 | 0.999772489 |
| *USP33* | -0.459072 | 0.226178 | -2.029697 | 0.04238731 | 0.999772489 |
| *CD6* | -1.417473 | 0.69872 | -2.028671 | 0.04249183 | 0.999772489 |
| *DOCK10* | 0.775624 | 0.382434 | 2.028123 | 0.04254768 | 0.999772489 |
| *SNAPC3* | 0.484227 | 0.239147 | 2.024813 | 0.04288654 | 0.999772489 |
| *GYPC* | 0.257542 | 0.12723 | 2.024222 | 0.0429473 | 0.999772489 |
| *FNDC3B* | 1.691811 | 0.835812 | 2.024153 | 0.04295438 | 0.999772489 |

| | | | | | |
|---|---|---|---|---|---|
| *CLEC17A* | 0.642398 | 0.317391 | 2.023999 | 0.04297029 | 0.999772489 |
| *CARF* | 0.427179 | 0.211128 | 2.023318 | 0.04304042 | 0.999772489 |
| *FUS* | 0.164701 | 0.081489 | 2.021156 | 0.04326359 | 0.999772489 |
| *TMEM191C* | -0.877435 | 0.434286 | -2.020408 | 0.04334112 | 0.999772489 |
| *PPP4R3B* | -0.423074 | 0.209504 | -2.019411 | 0.04344449 | 0.999772489 |
| *ENSG00000259888* | 1.020969 | 0.505797 | 2.018535 | 0.04353561 | 0.999772489 |
| *LRP10* | 0.262341 | 0.129973 | 2.018421 | 0.04354748 | 0.999772489 |
| *BAG4* | -0.727055 | 0.360299 | -2.017921 | 0.04359947 | 0.999772489 |
| *PPIL2* | 0.433998 | 0.215113 | 2.017532 | 0.04364 | 0.999772489 |
| *DPH3* | -0.608129 | 0.301628 | -2.016158 | 0.04378342 | 0.999772489 |
| *AUH* | 0.798718 | 0.396273 | 2.015574 | 0.04384454 | 0.999772489 |
| *ENSG00000277654* | -0.563442 | 0.279661 | -2.014733 | 0.04393267 | 0.999772489 |
| *ENSG00000272426* | 1.032424 | 0.512627 | 2.013988 | 0.04401079 | 0.999772489 |
| *ASAH1* | 0.616475 | 0.306112 | 2.013884 | 0.0440217 | 0.999772489 |
| *FSTL3* | 1.675215 | 0.831862 | 2.013814 | 0.04402908 | NA |
| *DDHD2* | -0.719259 | 0.35726 | -2.013264 | 0.04408691 | 0.999772489 |
| *TMEM159* | -0.658787 | 0.327586 | -2.011037 | 0.04432155 | 0.999772489 |
| *NAV1* | 1.719273 | 0.855077 | 2.010664 | 0.044361 | NA |
| *TMEM183A* | -0.342784 | 0.170806 | -2.006857 | 0.04476493 | 0.999772489 |
| *CRX* | 1.490059 | 0.74305 | 2.005327 | 0.04492806 | 0.999772489 |
| *ATP1A1* | -0.491997 | 0.24552 | -2.003895 | 0.04508128 | 0.999772489 |
| *RNF168* | -0.505578 | 0.252546 | -2.00192 | 0.04529329 | 0.999772489 |
| *CNIH3* | 1.994585 | 0.996507 | 2.001577 | 0.04533027 | NA |
| *ST3GAL1* | 0.636345 | 0.317936 | 2.001488 | 0.04533984 | 0.999772489 |
| *MTPN* | 0.369384 | 0.184562 | 2.001402 | 0.04534906 | 0.999772489 |
| *EXPH5* | 1.591366 | 0.795237 | 2.001122 | 0.04537926 | NA |
| *HHEX* | -0.250948 | 0.125405 | -2.001104 | 0.04538122 | 0.999772489 |
| *MAMSTR* | 1.195692 | 0.597543 | 2.001014 | 0.04539093 | 0.999772489 |
| *NT5DC4* | 1.488717 | 0.744033 | 2.000877 | 0.04540567 | 0.999772489 |
| *ELAC1* | 0.958154 | 0.47894 | 2.000573 | 0.04543838 | 0.999772489 |
| *GPR18* | 0.908561 | 0.454397 | 1.999486 | 0.04555579 | 0.999772489 |
| *FUT7* | 1.691499 | 0.847363 | 1.996191 | 0.04591311 | 0.999772489 |
| *IQSEC1* | -0.366413 | 0.183572 | -1.996017 | 0.04593206 | 0.999772489 |
| *CTSB* | 0.426884 | 0.214029 | 1.994515 | 0.04609578 | 0.999772489 |
| *NKG7* | 2.138781 | 1.072642 | 1.993936 | 0.04615901 | NA |
| *RGS1* | -1.130538 | 0.567219 | -1.993123 | 0.04624799 | 0.999772489 |
| *IL10RA* | -0.286933 | 0.144042 | -1.992002 | 0.0463708 | 0.999772489 |
| *ZNF625* | 1.307075 | 0.656551 | 1.990822 | 0.04650051 | 0.999772489 |
| *ICA1* | 2.719174 | 1.365999 | 1.990612 | 0.04652353 | NA |
| *BBX* | 0.223993 | 0.112526 | 1.990589 | 0.0465261 | 0.999772489 |
| *RNPC3* | -0.532703 | 0.267807 | -1.989134 | 0.04668647 | 0.999772489 |
| *DICER1.AS1* | -1.024646 | 0.515221 | -1.988751 | 0.04672875 | 0.999772489 |
| *CCDC154* | 1.108468 | 0.558124 | 1.986061 | 0.04702652 | 0.999772489 |
| *FXYD5* | -0.18918 | 0.095317 | -1.984747 | 0.04717259 | 0.999772489 |
| *LPCAT4* | -0.585074 | 0.295365 | -1.980852 | 0.04760791 | 0.999772489 |
| *ENSG00000276529* | 1.870662 | 0.944649 | 1.980272 | 0.04767294 | NA |
| *TUT1* | 0.597483 | 0.301859 | 1.979342 | 0.04777749 | 0.999772489 |

| | | | | | |
|---|---|---|---|---|---|
| *PDS5B* | -0.55419 | 0.279998 | -1.979259 | 0.04778681 | 0.999772489 |
| *APOL3* | 0.535771 | 0.270764 | 1.978739 | 0.04784544 | 0.999772489 |
| *PRKCA* | 1.888652 | 0.955559 | 1.976489 | 0.04809944 | 0.999772489 |
| *C5orf17* | 1.922096 | 0.972634 | 1.976177 | 0.04813473 | NA |
| *TBCK* | 0.695505 | 0.352178 | 1.974867 | 0.0482833 | 0.999772489 |
| *CFAP410* | -0.451586 | 0.228701 | -1.974571 | 0.0483169 | 0.999772489 |
| *HOXB4* | 2.225393 | 1.127955 | 1.972945 | 0.04850184 | NA |
| *ENSG00000268746* | 1.563986 | 0.792997 | 1.972247 | 0.04858143 | NA |
| *ARL13B* | -0.981203 | 0.497527 | -1.972161 | 0.04859128 | 0.999772489 |
| *WDFY4* | -0.416112 | 0.211009 | -1.972014 | 0.04860804 | 0.999772489 |
| *DERL3* | 0.341003 | 0.173086 | 1.970138 | 0.04882253 | 0.999772489 |
| *TFE3* | -0.921791 | 0.467906 | -1.970037 | 0.04883415 | 0.999772489 |
| *ZNF239* | 1.265711 | 0.642647 | 1.969528 | 0.04889253 | 0.999772489 |
| *WDR82* | -0.51432 | 0.261256 | -1.96864 | 0.04899445 | 0.999772489 |
| *ENSG00000226853* | 1.913425 | 0.972052 | 1.968439 | 0.04901755 | 0.999772489 |
| *GFI1* | 2.728829 | 1.386615 | 1.967979 | 0.04907045 | NA |
| *PACS1* | 0.315455 | 0.160352 | 1.967269 | 0.04915226 | 0.999772489 |
| *SGMS1* | 0.599913 | 0.305164 | 1.965873 | 0.04931326 | 0.999772489 |
| *IRF2BPL* | 0.76387 | 0.388639 | 1.965499 | 0.04935651 | 0.999772489 |

| Cluster 9 | log2FC | lfcSE | stat | pvalue | padj |
|---|---|---|---|---|---|
| *VPREB3* | 4.20113 | 0.146171 | 28.741187 | < 2.22e-16 | < 2.22e-16 |
| *DENND6B* | 3.54218 | 0.202611 | 17.48262 | < 2.22e-16 | < 2.22e-16 |
| *P2RX5* | 2.231763 | 0.156485 | 14.261856 | < 2.22e-16 | < 2.22e-16 |
| *PLAAT4* | 2.687207 | 0.212616 | 12.638769 | < 2.22e-16 | < 2.22e-16 |
| *NRIP1* | 2.476226 | 0.205137 | 12.071079 | < 2.22e-16 | < 2.22e-16 |
| *SMAD3* | 1.179846 | 0.105888 | 11.142385 | < 2.22e-16 | < 2.22e-16 |
| *GRN* | 1.738068 | 0.156749 | 11.088247 | < 2.22e-16 | < 2.22e-16 |
| *SEPTIN9* | 0.950286 | 0.086125 | 11.033804 | < 2.22e-16 | < 2.22e-16 |
| *CD38* | 5.158908 | 0.489618 | 10.536598 | < 2.22e-16 | < 2.22e-16 |
| *ASB2* | 5.324524 | 0.533794 | 9.974866 | < 2.22e-16 | < 2.22e-16 |
| *CD1D* | 5.652924 | 0.580968 | 9.73018 | < 2.22e-16 | < 2.22e-16 |
| *SLC5A3* | 1.169983 | 0.127025 | 9.210632 | < 2.22e-16 | < 2.22e-16 |
| *DUS2* | 1.759822 | 0.197357 | 8.916956 | < 2.22e-16 | 4.47E-16 |
| *RABGAP1L* | 1.626291 | 0.18776 | 8.661538 | < 2.22e-16 | 4.03E-15 |
| *SLC2A5* | 3.241203 | 0.383694 | 8.44736 | < 2.22e-16 | 2.41E-14 |
| *C12orf65* | 0.630741 | 0.075564 | 8.34708 | < 2.22e-16 | 5.31E-14 |
| *RNF207* | 2.286963 | 0.308639 | 7.409839 | 1.26E-13 | 9.02E-11 |
| *DPEP2* | 1.587585 | 0.214795 | 7.39115 | 1.46E-13 | 9.81E-11 |
| *SLC12A4* | 2.092936 | 0.284384 | 7.359548 | 1.85E-13 | 1.18E-10 |
| *RARA.AS1* | 1.879972 | 0.263497 | 7.134706 | 9.70E-13 | 5.88E-10 |
| *FCRL2* | 1.518003 | 0.215055 | 7.058685 | 1.68E-12 | 9.71E-10 |
| *AHI1* | 1.056846 | 0.152232 | 6.942351 | 3.86E-12 | 2.13E-09 |
| *TRABD* | 0.663898 | 0.096121 | 6.906897 | 4.95E-12 | 2.61E-09 |
| *CD72* | 1.181273 | 0.171517 | 6.887213 | 5.69E-12 | 2.88E-09 |
| *HIC1* | 2.759468 | 0.401388 | 6.874821 | 6.21E-12 | 3.01E-09 |
| *MRPS6* | 0.801899 | 0.117638 | 6.816692 | 9.32E-12 | 4.35E-09 |
| *IRF2BPL* | 0.8984 | 0.138047 | 6.507929 | 7.62E-11 | 3.42E-08 |
| *ENDOD1* | 1.658461 | 0.259628 | 6.387848 | 1.68E-10 | 7.29E-08 |
| *LHFPL2* | 1.976063 | 0.310017 | 6.374044 | 1.84E-10 | 7.70E-08 |
| *CHST15* | 1.227661 | 0.193796 | 6.334821 | 2.38E-10 | 9.61E-08 |
| *RHOBTB3* | 2.453235 | 0.389877 | 6.292327 | 3.13E-10 | 1.22E-07 |
| *CACYBP* | 0.665342 | 0.107267 | 6.202664 | 5.55E-10 | 2.10E-07 |
| *H3F3A* | 0.462554 | 0.07744 | 5.973068 | 2.33E-09 | 8.56E-07 |
| *CD1C* | 2.263507 | 0.380394 | 5.95043 | 2.67E-09 | 9.54E-07 |
| *VPREB1* | 5.438502 | 0.926062 | 5.87272 | 4.29E-09 | 1.49E-06 |
| *LGMN* | 1.080394 | 0.185232 | 5.832655 | 5.46E-09 | 1.84E-06 |
| *MPEG1* | 1.740797 | 0.29928 | 5.816618 | 6.01E-09 | 1.97E-06 |
| *CMTM7* | 1.034133 | 0.179747 | 5.753288 | 8.75E-09 | 2.79E-06 |
| *RARA* | 0.539822 | 0.094112 | 5.735985 | 9.69E-09 | 3.02E-06 |
| *GGA2* | 0.801396 | 0.141481 | 5.664343 | 1.48E-08 | 4.48E-06 |
| *ABCB1* | 1.309048 | 0.232136 | 5.639153 | 1.71E-08 | 5.06E-06 |
| *LCAT* | 2.29004 | 0.412196 | 5.555705 | 2.76E-08 | 7.99E-06 |
| *ARL14EPL* | 2.720228 | 0.498375 | 5.458194 | 4.81E-08 | 1.36E-05 |
| *CXCR5* | 0.824101 | 0.151141 | 5.452514 | 4.97E-08 | 1.37E-05 |
| *IRAK2* | 0.78859 | 0.145333 | 5.426107 | 5.76E-08 | 1.55E-05 |
| *UBTF* | 0.721285 | 0.13304 | 5.421544 | 5.91E-08 | 1.55E-05 |

| | | | | | |
|---|---|---|---|---|---|
| *PISD* | 0.904754 | 0.166969 | 5.418697 | 6.00E-08 | 1.55E-05 |
| *PNOC* | 1.380602 | 0.256369 | 5.385214 | 7.24E-08 | 1.83E-05 |
| *BTBD7* | 0.58572 | 0.108967 | 5.375218 | 7.65E-08 | 1.89E-05 |
| *MS4A1* | -0.654199 | 0.123925 | -5.279002 | 1.30E-07 | 3.15E-05 |
| *FAM3C* | 0.486109 | 0.093176 | 5.217091 | 1.82E-07 | 4.32E-05 |
| *P2RX5.TAX1BP3* | 1.831872 | 0.357196 | 5.128484 | 2.92E-07 | 6.81E-05 |
| *XYLT1* | 1.205116 | 0.236275 | 5.100479 | 3.39E-07 | 7.76E-05 |
| *ZNF563* | 1.067412 | 0.214961 | 4.965617 | 6.85E-07 | 0.000153859 |
| *OTUD7A* | 2.807564 | 0.567778 | 4.944823 | 7.62E-07 | 0.000168112 |
| *AGPAT5* | 0.506045 | 0.102484 | 4.937787 | 7.90E-07 | 0.000171178 |
| *PTK2B* | 0.591349 | 0.12018 | 4.920522 | 8.63E-07 | 0.000183712 |
| *JCHAIN* | 2.119812 | 0.431713 | 4.910235 | 9.10E-07 | 0.000190278 |
| *SIGIRR* | 0.914882 | 0.186715 | 4.899873 | 9.59E-07 | 0.000197194 |
| *ZNF3* | 0.648306 | 0.132424 | 4.895699 | 9.80E-07 | 0.000198069 |
| *ACP5* | 0.917231 | 0.189588 | 4.838025 | 1.31E-06 | 0.000260809 |
| *CLCN3* | 0.603025 | 0.125188 | 4.816959 | 1.46E-06 | 0.000285224 |
| *AGO1* | 0.67097 | 0.139394 | 4.813474 | 1.48E-06 | 0.000285639 |
| *APOL3* | 0.904677 | 0.188653 | 4.795468 | 1.62E-06 | 0.000307651 |
| *KANSL3* | 0.895359 | 0.187264 | 4.781266 | 1.74E-06 | 0.000325128 |
| *MAP3K1* | 0.716176 | 0.152102 | 4.708522 | 2.50E-06 | 0.000458663 |
| *ZFP36L1* | -0.408682 | 0.086978 | -4.698681 | 2.62E-06 | 0.000474139 |
| *RRBP1* | 0.819423 | 0.175051 | 4.681064 | 2.85E-06 | 0.00050917 |
| *IFI30* | 0.811189 | 0.174764 | 4.64164 | 3.46E-06 | 0.000607752 |
| *CLIC3* | 2.510137 | 0.543473 | 4.618701 | 3.86E-06 | 0.000669254 |
| *PLAAT3* | 1.862902 | 0.404427 | 4.606277 | 4.10E-06 | 0.00070048 |
| *TELO2* | 0.75471 | 0.167351 | 4.509729 | 6.49E-06 | 0.001093741 |
| *RESF1* | -0.28691 | 0.064008 | -4.48238 | 7.38E-06 | 0.001226749 |
| *RMDN2* | 1.68112 | 0.375544 | 4.476498 | 7.59E-06 | 0.001243981 |
| *TSPAN18* | 3.069158 | 0.690164 | 4.446999 | 8.71E-06 | 0.001408575 |
| *C15orf39* | 0.651496 | 0.146794 | 4.438164 | 9.07E-06 | 0.001448331 |
| *NFAT5* | 0.606071 | 0.137091 | 4.420952 | 9.83E-06 | 0.001548283 |
| *CD22* | 0.782324 | 0.178897 | 4.37303 | 1.23E-05 | 0.001905876 |
| *SLC43A2* | 0.482786 | 0.111061 | 4.347033 | 1.38E-05 | 0.002119132 |
| *NCF1* | 0.624801 | 0.144051 | 4.337367 | 1.44E-05 | 0.002186785 |
| *CTSZ* | 0.512155 | 0.119335 | 4.291723 | 1.77E-05 | 0.002655435 |
| *CR2* | 1.792558 | 0.41954 | 4.272675 | 1.93E-05 | 0.002848992 |
| *FAM117A* | 0.763883 | 0.178869 | 4.270642 | 1.95E-05 | 0.002848992 |
| *SDE2* | 0.694657 | 0.165502 | 4.19726 | 2.70E-05 | 0.003901929 |
| *SPTBN1* | 0.648233 | 0.156394 | 4.144865 | 3.40E-05 | 0.004853003 |
| *ENSG00000235609* | 3.237231 | 0.785426 | 4.121623 | 3.76E-05 | 0.005307231 |
| *GBP4* | 0.750928 | 0.183967 | 4.081865 | 4.47E-05 | 0.006229945 |
| *CHCHD10* | 1.906597 | 0.468943 | 4.065737 | 4.79E-05 | 0.006601037 |
| *SRSF8* | 0.456606 | 0.112753 | 4.049621 | 5.13E-05 | 0.006993017 |
| *SEC61B* | -0.244952 | 0.06066 | -4.038096 | 5.39E-05 | 0.007263929 |
| *SMIM3* | 1.555549 | 0.385684 | 4.033223 | 5.50E-05 | 0.007334809 |
| *RAB37* | 1.577632 | 0.395995 | 3.983971 | 6.78E-05 | 0.008937229 |
| *DBNL* | 0.414232 | 0.104354 | 3.969495 | 7.20E-05 | 0.009372147 |

| | | | | | |
|---|---|---|---|---|---|
| *SLA* | 0.902515 | 0.227474 | 3.967546 | 7.26E-05 | 0.009372147 |
| *NDC1* | 0.912454 | 0.230741 | 3.954456 | 7.67E-05 | 0.009796153 |
| *CHST7* | 1.085951 | 0.277174 | 3.917936 | 8.93E-05 | 0.011286613 |
| *RDH10* | 1.568531 | 0.40321 | 3.890111 | 0.000100198 | 0.012532028 |
| *CNP* | 0.583369 | 0.150433 | 3.877946 | 0.000105342 | 0.013040904 |
| *IL1B* | 3.698407 | 0.95705 | 3.864384 | 0.00011137 | 0.013647899 |
| *CD99* | -0.355767 | 0.093117 | -3.820654 | 0.000133098 | 0.016147464 |
| *SLC25A39* | 0.359223 | 0.094684 | 3.793935 | 0.000148279 | 0.017811047 |
| *FCRL1* | 0.942339 | 0.248796 | 3.787591 | 0.000152115 | 0.018092729 |
| *ST3GAL1* | 0.65772 | 0.176646 | 3.723386 | 0.000196569 | 0.023153113 |
| *MFSD10* | -0.293517 | 0.078918 | -3.719283 | 0.000199789 | 0.023306166 |
| *JUNB* | 0.381981 | 0.103423 | 3.693401 | 0.000221275 | 0.025566738 |
| *ABI3* | 1.041609 | 0.283328 | 3.676339 | 0.000236605 | 0.027080127 |
| *CLCN7* | 0.525416 | 0.143422 | 3.663426 | 0.000248864 | 0.028216953 |
| *VHL* | 0.342618 | 0.094186 | 3.637666 | 0.00027512 | 0.030893198 |
| *NUBP1* | 0.422287 | 0.11616 | 3.63539 | 0.00027756 | 0.030893198 |
| *GPR146* | -1.158499 | 0.319722 | -3.62346 | 0.000290688 | 0.032060235 |
| *SH3BGRL3* | -0.21544 | 0.059899 | -3.59671 | 0.000322268 | 0.035223012 |
| *PALLD* | 2.841929 | 0.794161 | 3.578527 | 0.000345536 | 0.037428924 |
| *SREBF1* | 0.72351 | 0.20602 | 3.511844 | 0.00044501 | 0.047777512 |
| *RBM47* | 1.401788 | 0.400152 | 3.503136 | 0.000459815 | 0.048933977 |
| *CASP7* | 0.669455 | 0.192368 | 3.480069 | 0.000501284 | 0.052883264 |
| *PPCDC* | 0.629864 | 0.1825 | 3.451314 | 0.000557864 | 0.058344885 |
| *PXK* | 0.505045 | 0.146608 | 3.444869 | 0.000571337 | 0.059243246 |
| *ETFB* | 0.455694 | 0.13255 | 3.437914 | 0.000586214 | 0.059891008 |
| *RAB20* | 1.469759 | 0.427621 | 3.43706 | 0.000588064 | 0.059891008 |
| *HIPK2* | 0.624422 | 0.181778 | 3.435073 | 0.000592394 | 0.059891008 |
| *PUS1* | 0.516217 | 0.151158 | 3.415078 | 0.000637637 | 0.063932303 |
| *MAPKAPK2* | 0.536626 | 0.157936 | 3.397747 | 0.000679431 | 0.067564419 |
| *SPIB* | 0.469067 | 0.138279 | 3.39217 | 0.000693414 | 0.068394325 |
| *TFEB* | 0.447861 | 0.132327 | 3.384505 | 0.000713067 | 0.069765554 |
| *MCM5* | 0.386992 | 0.114863 | 3.369148 | 0.000754008 | 0.073180995 |
| *AP3S1* | 0.335713 | 0.099834 | 3.362702 | 0.000771836 | 0.074316801 |
| *FOXO3* | 0.608202 | 0.181538 | 3.350276 | 0.000807311 | 0.077120418 |
| *DENND6A* | 0.934039 | 0.279098 | 3.346629 | 0.000818005 | 0.07753151 |
| *PPFIBP2* | 0.658906 | 0.197748 | 3.332051 | 0.000862085 | 0.081076056 |
| *SLC5A5* | 2.896068 | 0.872124 | 3.320705 | 0.000897903 | 0.083795042 |
| *PHF21A* | -0.505735 | 0.153249 | -3.300084 | 0.00096656 | 0.089196777 |
| *DTX4* | 1.216386 | 0.36872 | 3.298945 | 0.000970489 | 0.089196777 |
| *CD180* | 0.976973 | 0.296647 | 3.293385 | 0.000989889 | 0.090295739 |
| *PPDPF* | -0.288643 | 0.087948 | -3.281964 | 0.001030866 | 0.093331856 |
| *DDX28* | 0.534401 | 0.163812 | 3.262271 | 0.001105236 | 0.098669751 |
| *TMEM41B* | 0.439924 | 0.134861 | 3.262052 | 0.00110609 | 0.098669751 |
| *PHKB* | 0.349303 | 0.107546 | 3.247935 | 0.001162459 | 0.102507983 |
| *CIB1* | 0.259249 | 0.079841 | 3.247065 | 0.001166016 | 0.102507983 |
| *SEPTIN2* | 0.248797 | 0.076949 | 3.233253 | 0.001223889 | 0.106821769 |
| *SLC45A3* | 0.77827 | 0.241284 | 3.225533 | 0.001257383 | 0.108961206 |

| | | | | | |
|---|---|---|---|---|---|
| *CTSA* | 0.434722 | 0.134921 | 3.222041 | 0.00127281 | 0.109515826 |
| *ENSG00000259436* | -0.661649 | 0.205597 | -3.218177 | 0.001290081 | 0.110220121 |
| *KCNH4* | 2.237763 | 0.698274 | 3.204708 | 0.001351996 | 0.114702223 |
| *SHQ1* | 0.611736 | 0.191329 | 3.197293 | 0.001387238 | 0.116874759 |
| *ST8SIA4* | -0.35755 | 0.112244 | -3.185475 | 0.001445164 | 0.120915335 |
| *SPOCK2* | 0.612804 | 0.192587 | 3.181953 | 0.001462857 | 0.121557365 |
| *PLXNB2* | 0.95683 | 0.301615 | 3.172355 | 0.001512081 | 0.124792955 |
| *GPR68* | -1.412313 | 0.44553 | -3.169966 | 0.001524568 | 0.124973346 |
| *ITGAE* | 0.367258 | 0.116549 | 3.151109 | 0.00162652 | 0.132435821 |
| *PPP1R18* | 0.427393 | 0.135836 | 3.14639 | 0.001652997 | 0.133694374 |
| *LAGE3* | -0.252754 | 0.080733 | -3.130744 | 0.00174364 | 0.140091654 |
| *MED18* | 0.673176 | 0.215723 | 3.120558 | 0.001805085 | 0.144074279 |
| *DTX1* | 1.332103 | 0.42734 | 3.117201 | 0.001825769 | 0.144772729 |
| *LZTFL1* | 0.543395 | 0.174798 | 3.108704 | 0.001879102 | 0.148034169 |
| *CLEC17A* | 0.627269 | 0.20225 | 3.101457 | 0.001925706 | 0.150726892 |
| *VRK2* | -0.460374 | 0.150921 | -3.050424 | 0.002285189 | 0.177717369 |
| *POU2AF1* | 0.362082 | 0.118944 | 3.044138 | 0.002333481 | 0.180317178 |
| *KLF3* | 0.521425 | 0.172625 | 3.020572 | 0.002522979 | 0.193726496 |
| *AKAP1* | 0.641226 | 0.214616 | 2.987789 | 0.002810038 | 0.214411197 |
| *GDPGP1* | 1.076511 | 0.360671 | 2.984744 | 0.002838155 | 0.21513566 |
| *PSAP* | 0.349393 | 0.117131 | 2.982934 | 0.002854998 | 0.21513566 |
| *FAM219B* | -0.483377 | 0.16243 | -2.975912 | 0.002921183 | 0.218764118 |
| *NFATC3* | 0.379651 | 0.127826 | 2.97006 | 0.002977414 | 0.221362851 |
| *MMP11* | 1.713663 | 0.577278 | 2.96852 | 0.002992376 | 0.221362851 |
| *BCL3* | 0.454619 | 0.154393 | 2.944546 | 0.003234286 | 0.2378082 |
| *EIF2AK4* | -0.495479 | 0.168495 | -2.940616 | 0.003275598 | 0.239394918 |
| *RASGEF1B* | -0.467752 | 0.15919 | -2.938332 | 0.003299832 | 0.23972194 |
| *AP1G2* | -0.313637 | 0.106867 | -2.934843 | 0.00333716 | 0.240990652 |
| *ZNF823* | 1.779579 | 0.608221 | 2.925875 | 0.00343489 | 0.246580419 |
| *CNR2* | 0.472462 | 0.161687 | 2.922079 | 0.003477032 | 0.248137356 |
| *ERG28* | -0.264472 | 0.09072 | -2.915248 | 0.003554061 | 0.252151266 |
| *HOXB3* | 2.533283 | 0.872236 | 2.904353 | 0.003680125 | 0.259577214 |
| *STK17A* | 0.368193 | 0.127312 | 2.892053 | 0.003827339 | 0.26840044 |
| *KCNN1* | 1.211575 | 0.420505 | 2.88124 | 0.003961136 | 0.276186785 |
| *ENSG00000268798* | 0.801229 | 0.278517 | 2.876772 | 0.004017664 | 0.278249627 |
| *EHD4* | -0.369835 | 0.128625 | -2.875288 | 0.004036592 | 0.278249627 |
| *BCAR3* | 0.667107 | 0.232202 | 2.872958 | 0.004066478 | 0.278726023 |
| *COBLL1* | -0.383112 | 0.134109 | -2.856713 | 0.004280528 | 0.291749256 |
| *LCK* | 0.90728 | 0.318297 | 2.850415 | 0.004366218 | 0.295927159 |
| *FSD1L* | -0.524079 | 0.184191 | -2.845302 | 0.004436937 | 0.299049575 |
| *BCDIN3D* | 0.401 | 0.142092 | 2.822112 | 0.004770844 | 0.319778346 |
| *FAM107B* | 0.245826 | 0.087164 | 2.820273 | 0.004798277 | 0.319849952 |
| *SLC38A1* | -0.254028 | 0.090267 | -2.814173 | 0.004890289 | 0.322752797 |
| *ZNF674.AS1* | 0.86935 | 0.308953 | 2.813862 | 0.004895031 | 0.322752797 |
| *TNK2* | 0.587259 | 0.209388 | 2.804646 | 0.005037182 | 0.330330252 |
| *GPX1* | 0.504874 | 0.18117 | 2.786748 | 0.005323984 | 0.347261137 |
| *KAT6A* | 0.295383 | 0.106191 | 2.781617 | 0.005408891 | 0.350912636 |

| | | | | | |
|---|---|---|---|---|---|
| *C16orf54* | 0.518489 | 0.187337 | 2.767687 | 0.005645569 | 0.364319365 |
| *PAOX* | 0.584958 | 0.211558 | 2.765005 | 0.005692192 | 0.36538452 |
| *DNAJC7* | 0.197254 | 0.071472 | 2.759873 | 0.005782386 | 0.369220573 |
| *CERK* | 0.787041 | 0.285649 | 2.755276 | 0.005864259 | 0.371318574 |
| *ABCD3* | -0.68901 | 0.250131 | -2.754596 | 0.005876456 | 0.371318574 |
| *DDX24* | 0.183126 | 0.066525 | 2.75273 | 0.005910069 | 0.37150754 |
| *RINL* | 0.386195 | 0.140442 | 2.749859 | 0.005962097 | 0.372846177 |
| *AHNAK* | -0.781529 | 0.284793 | -2.744198 | 0.006065889 | 0.377391619 |
| *ENSG00000254802* | 1.288778 | 0.470477 | 2.739301 | 0.006156997 | 0.381105565 |
| *RREB1* | -0.418461 | 0.153038 | -2.734363 | 0.006250105 | 0.384904972 |
| *CBX1* | -0.237011 | 0.087292 | -2.715164 | 0.006624305 | 0.405889222 |
| *DNAH1* | 1.223255 | 0.452037 | 2.706092 | 0.006808018 | 0.415049643 |
| *SIPA1L1* | -0.302267 | 0.111842 | -2.702621 | 0.006879523 | 0.417311839 |
| *ENSG00000261766* | 0.488254 | 0.18101 | 2.69738 | 0.006988752 | 0.421828559 |
| *TNIP1* | 0.261858 | 0.097422 | 2.687863 | 0.007191096 | 0.430358372 |
| *GRK3* | 0.289173 | 0.107603 | 2.687402 | 0.007201018 | 0.430358372 |
| *NUP210* | 0.413636 | 0.1544 | 2.678997 | 0.007384314 | 0.438943565 |
| *EAF2* | 0.441092 | 0.1648 | 2.676528 | 0.007438942 | 0.438943565 |
| *ZCCHC7* | 0.401552 | 0.150063 | 2.675885 | 0.007453213 | 0.438943565 |
| *LY6E* | -0.499076 | 0.187412 | -2.662992 | 0.007744936 | 0.45174774 |
| *VKORC1* | -0.235663 | 0.08853 | -2.661956 | 0.007768807 | 0.45174774 |
| *STMP1* | -0.313196 | 0.117757 | -2.659673 | 0.00782166 | 0.45174774 |
| *PXDC1* | 1.05569 | 0.397052 | 2.65882 | 0.007841496 | 0.45174774 |
| *CPSF1* | -0.54655 | 0.205635 | -2.657863 | 0.007863778 | 0.45174774 |
| *ZNF318* | -0.56865 | 0.214054 | -2.656568 | 0.007894042 | 0.45174774 |
| *RAB8B* | -0.276925 | 0.104484 | -2.650417 | 0.008039257 | 0.455456513 |
| *BMP2K* | -0.413065 | 0.15585 | -2.650394 | 0.008039806 | 0.455456513 |
| *TSPO* | -0.339414 | 0.128194 | -2.647664 | 0.008105014 | 0.455456513 |
| *IDS* | -0.199415 | 0.075322 | -2.647497 | 0.008109018 | 0.455456513 |
| *MDM4* | -0.226061 | 0.085462 | -2.645177 | 0.008164828 | 0.456477874 |
| *TMEM65* | 0.89225 | 0.338281 | 2.637602 | 0.008349449 | 0.464658324 |
| *SLC25A11* | 0.301582 | 0.114456 | 2.63492 | 0.008415705 | 0.465761572 |
| *ENSG00000225489* | 1.171301 | 0.444736 | 2.633698 | 0.008446056 | 0.465761572 |
| *STK24* | 0.335925 | 0.127945 | 2.62554 | 0.008651154 | 0.474811446 |
| *GEN1* | -0.458882 | 0.174874 | -2.624076 | 0.008688439 | 0.474811446 |
| *C7orf50* | -0.416154 | 0.158894 | -2.61906 | 0.008817248 | 0.479639548 |
| *SEL1L3* | 0.618972 | 0.236468 | 2.617569 | 0.008855857 | 0.479639548 |
| *ALOX5* | 0.308239 | 0.11789 | 2.614646 | 0.008931994 | 0.481613105 |
| *HMG20B* | -0.290299 | 0.111253 | -2.609352 | 0.009071375 | 0.486107425 |
| *FCHO1* | 0.63029 | 0.241634 | 2.608444 | 0.009095482 | 0.486107425 |
| *TEX264* | 0.303553 | 0.116581 | 2.603797 | 0.009219744 | 0.490587441 |
| *S100A6* | -0.436979 | 0.168476 | -2.59372 | 0.009494376 | 0.502994623 |
| *RHOF* | -0.24054 | 0.09284 | -2.590906 | 0.009572372 | 0.504921827 |
| *SPRY1* | 0.827068 | 0.319888 | 2.585494 | 0.009723968 | 0.510697761 |
| *GAK* | 0.337317 | 0.130728 | 2.580288 | 0.009871794 | 0.51622677 |
| *SLCO5A1* | 0.76666 | 0.297918 | 2.573393 | 0.010070681 | 0.524366971 |
| *MACROD2* | -0.689297 | 0.269408 | -2.558563 | 0.010510583 | 0.542978537 |

| | | | | | |
|---|---|---|---|---|---|
| *GABARAPL1* | 0.569782 | 0.22275 | 2.557943 | 0.010529323 | 0.542978537 |
| *SIGLEC5* | 1.173808 | 0.459083 | 2.556853 | 0.010562392 | 0.542978537 |
| *FTL* | -0.246915 | 0.09664 | -2.554998 | 0.010618849 | 0.543577528 |
| *NLRC3* | 0.822796 | 0.322748 | 2.549343 | 0.010792594 | 0.550150235 |
| *SACS* | -0.506842 | 0.199004 | -2.546895 | 0.010868622 | 0.551707635 |
| *TBC1D8* | 1.258568 | 0.494474 | 2.545269 | 0.010919366 | 0.551973944 |
| *A4GALT* | 0.997873 | 0.392906 | 2.539724 | 0.011094007 | 0.558437975 |
| *PTOV1* | -0.547501 | 0.215696 | -2.538298 | 0.0111393 | 0.558437975 |
| *OXSR1* | -0.298102 | 0.11763 | -2.534239 | 0.011269197 | 0.560376246 |
| *MRPL41* | -0.208683 | 0.082389 | -2.532913 | 0.011311893 | 0.560376246 |
| *ZBED2* | -1.377891 | 0.544025 | -2.53277 | 0.011316533 | 0.560376246 |
| *KLF6* | -0.231376 | 0.091607 | -2.525746 | 0.011545289 | 0.568264425 |
| *C16orf74* | 0.487421 | 0.193037 | 2.52501 | 0.011569511 | 0.568264425 |
| *ATP13A2* | 0.789933 | 0.31386 | 2.516836 | 0.011841398 | 0.577792002 |
| *ZNF595* | 1.052558 | 0.418293 | 2.51632 | 0.011858738 | 0.577792002 |
| *NXT1* | 0.211656 | 0.084194 | 2.513915 | 0.011939925 | 0.579420687 |
| *MTSS1* | -0.376352 | 0.149906 | -2.510598 | 0.012052693 | 0.582562844 |
| *LMNA* | -0.523587 | 0.208868 | -2.506783 | 0.012183532 | 0.583058892 |
| *ENSG00000245869* | 0.806325 | 0.321674 | 2.506654 | 0.012188006 | 0.583058892 |
| *PDCD4* | 0.45043 | 0.179733 | 2.506099 | 0.012207135 | 0.583058892 |
| *VPS13A* | -0.330554 | 0.132243 | -2.499598 | 0.012433427 | 0.58789538 |
| *SLAIN2* | -0.271698 | 0.108701 | -2.499511 | 0.012436481 | 0.58789538 |
| *TBC1D9* | 0.639017 | 0.255707 | 2.499019 | 0.012453768 | 0.58789538 |
| *ENSG00000229539* | 0.507374 | 0.203242 | 2.496407 | 0.012545855 | 0.589946969 |
| *CDC42* | -0.169791 | 0.068101 | -2.493223 | 0.012658926 | 0.592965605 |
| *TRIB1* | 1.036318 | 0.416037 | 2.490927 | 0.012741023 | 0.594515713 |
| *ATF7IP2* | 0.393143 | 0.157922 | 2.489483 | 0.012792887 | 0.594648696 |
| *TRNAU1AP* | -0.244554 | 0.098321 | -2.487307 | 0.01287143 | 0.595395116 |
| *FDFT1* | 0.261521 | 0.105195 | 2.486063 | 0.012916517 | 0.595395116 |
| *R3HDM2* | 0.406995 | 0.163783 | 2.484972 | 0.012956175 | 0.595395116 |
| *ZNF816.ZNF321P* | -1.242906 | 0.50045 | -2.483577 | 0.01300703 | 0.595476549 |
| *ING2* | -0.499854 | 0.201497 | -2.480696 | 0.01311262 | 0.598053764 |
| *BMPR1A* | -1.823765 | 0.736099 | -2.477609 | 0.013226613 | 0.600993536 |
| *MIR34AHG* | -1.18223 | 0.477687 | -2.474906 | 0.013327139 | 0.60330169 |
| *HOMER2* | -0.667714 | 0.270034 | -2.472705 | 0.013409476 | 0.604772359 |
| *BMF* | -0.293442 | 0.119355 | -2.458569 | 0.013949186 | 0.626783411 |
| *FAM234B* | -1.070308 | 0.436502 | -2.452011 | 0.014206044 | 0.635675796 |
| *TBL1X* | -0.447283 | 0.182501 | -2.450851 | 0.014251881 | 0.635675796 |
| *SGSM3* | -0.327558 | 0.134169 | -2.441385 | 0.01463104 | 0.650196988 |
| *SMARCB1* | 0.251275 | 0.103166 | 2.435626 | 0.014866048 | 0.654607402 |
| *ACO2* | -0.256851 | 0.105468 | -2.435338 | 0.014877888 | 0.654607402 |
| *ISG15* | -0.463727 | 0.190443 | -2.434991 | 0.014892157 | 0.654607402 |
| *INKA2* | -0.569365 | 0.234563 | -2.427342 | 0.015209891 | 0.663117811 |
| *NEB* | -1.096033 | 0.451543 | -2.427303 | 0.015211544 | 0.663117811 |
| *PLEKHO2* | 0.501758 | 0.206805 | 2.426231 | 0.01525654 | 0.663117811 |
| *CASZ1* | -0.550307 | 0.227039 | -2.423841 | 0.015357338 | 0.663117811 |
| *C16orf72* | 0.329925 | 0.136119 | 2.4238 | 0.015359059 | 0.663117811 |

| | | | | | |
|---|---|---|---|---|---|
| *CEP76* | 0.711725 | 0.293961 | 2.421159 | 0.015471097 | 0.665586365 |
| *CHP1* | -0.291961 | 0.120659 | -2.419721 | 0.015532423 | 0.665863444 |
| *VTI1B* | -0.219314 | 0.091011 | -2.409751 | 0.015963401 | 0.677562301 |
| *GTF2E2* | 0.330975 | 0.137352 | 2.409694 | 0.015965903 | 0.677562301 |
| *CEP350* | -0.251231 | 0.104265 | -2.409535 | 0.015972867 | 0.677562301 |
| *IGLV1.51* | -1.526973 | 0.634305 | -2.407317 | 0.016070205 | 0.679316113 |
| *GPBP1* | -0.198265 | 0.082474 | -2.403973 | 0.016217979 | 0.683182376 |
| *DDAH2* | -0.333348 | 0.138877 | -2.400314 | 0.016381004 | 0.685537486 |
| *EVI2B* | 0.226859 | 0.094559 | 2.39911 | 0.016434967 | 0.685537486 |
| *PLP2* | -0.265446 | 0.110652 | -2.398922 | 0.016443407 | 0.685537486 |
| *SERPINF1* | -1.085649 | 0.453415 | -2.39438 | 0.016648498 | 0.68963614 |
| *DNAJB11* | 0.226518 | 0.09461 | 2.394228 | 0.016655406 | 0.68963614 |
| *NRM* | 0.58774 | 0.246029 | 2.38891 | 0.016898444 | 0.695571284 |
| *PDP1* | -0.566674 | 0.237243 | -2.388585 | 0.016913413 | 0.695571284 |
| *UBXN2B* | -0.37886 | 0.15872 | -2.386977 | 0.016987552 | 0.696260056 |
| *DPM3* | -0.289425 | 0.12167 | -2.378776 | 0.017370227 | 0.70954744 |
| *MNT* | 0.447382 | 0.18847 | 2.373761 | 0.017607937 | 0.716843943 |
| *LMF2* | -0.292591 | 0.123357 | -2.371912 | 0.017696287 | 0.718031275 |
| *APOL1* | 0.648272 | 0.274187 | 2.364342 | 0.018062147 | 0.728893056 |
| *LSP1* | 0.320595 | 0.135622 | 2.363891 | 0.018084142 | 0.728893056 |
| *STK4* | -0.137928 | 0.058528 | -2.356628 | 0.018441733 | 0.740403269 |
| *CD63* | -0.261132 | 0.110909 | -2.354469 | 0.018549206 | 0.740403269 |
| *SLC9A7* | 0.354285 | 0.150478 | 2.354397 | 0.018552802 | 0.740403269 |
| *TMED2* | -0.150194 | 0.063851 | -2.352259 | 0.018659779 | 0.742230951 |
| *TMSB10* | -0.244753 | 0.104212 | -2.348616 | 0.018843349 | 0.744793898 |
| *POU2F2* | -0.209001 | 0.088992 | -2.348544 | 0.018846994 | 0.744793898 |
| *RUNX2* | 1.708259 | 0.730051 | 2.339919 | 0.01928794 | 0.75974443 |
| *MCL1* | 0.226048 | 0.096677 | 2.338177 | 0.019378065 | 0.760824214 |
| *JUN* | 0.333653 | 0.142909 | 2.33472 | 0.019558042 | 0.765309716 |
| *SNX11* | -0.22347 | 0.095877 | -2.330789 | 0.01976449 | 0.765309716 |
| *CD79B* | 0.356772 | 0.15311 | 2.330166 | 0.019797383 | 0.765309716 |
| *SEC22C* | -0.309347 | 0.132816 | -2.329143 | 0.019851511 | 0.765309716 |
| *MCM2* | 0.825798 | 0.354558 | 2.329093 | 0.019854125 | 0.765309716 |
| *GPD1L* | -1.64327 | 0.705636 | -2.328779 | 0.019870801 | 0.765309716 |
| *RGS9* | -0.826129 | 0.355601 | -2.323189 | 0.020169014 | 0.770383624 |
| *ENSG00000273188* | 1.066013 | 0.458901 | 2.322971 | 0.02018072 | 0.770383624 |
| *CHTF18* | -0.854574 | 0.367916 | -2.322742 | 0.020193043 | 0.770383624 |
| *SMARCA4* | 0.369394 | 0.159288 | 2.319038 | 0.020392995 | 0.772851463 |
| *ENSG00000272369* | -1.099261 | 0.474096 | -2.318648 | 0.020414119 | 0.772851463 |
| *FAM161A* | -0.618216 | 0.266701 | -2.318009 | 0.020448839 | 0.772851463 |
| *MIR181A2HG* | 1.400356 | 0.60478 | 2.31548 | 0.020586657 | 0.775643844 |
| *TRAPPC1* | -0.157026 | 0.067879 | -2.313308 | 0.020705702 | 0.776578264 |
| *KLF13* | 0.278118 | 0.120257 | 2.312694 | 0.020739479 | 0.776578264 |
| *ZFYVE21* | -0.422378 | 0.18273 | -2.311491 | 0.020805747 | 0.776662537 |
| *MREG* | -0.484657 | 0.210222 | -2.305456 | 0.021141065 | 0.786758888 |
| *SNX18* | 0.778032 | 0.338597 | 2.297809 | 0.021572661 | 0.798171103 |
| *ZNF785* | 0.511166 | 0.222475 | 2.297628 | 0.021582994 | 0.798171103 |

| | | | | | |
|---|---|---|---|---|---|
| *BTG2* | -0.341531 | 0.148718 | -2.296503 | 0.021647164 | 0.798171103 |
| *SYNPO* | -1.816192 | 0.791235 | -2.295388 | 0.021710886 | 0.798171103 |
| *VMP1* | -0.198915 | 0.086737 | -2.293306 | 0.021830405 | 0.800140385 |
| *DNAJB12* | 0.227664 | 0.099478 | 2.288588 | 0.022103298 | 0.807702445 |
| *UAP1L1* | -0.956752 | 0.418809 | -2.284458 | 0.022344604 | 0.812757999 |
| *ENSG00000253535* | 0.793871 | 0.34759 | 2.28393 | 0.022375632 | 0.812757999 |
| *MPHOSPH9* | 0.386918 | 0.169777 | 2.278981 | 0.022668173 | 0.82092621 |
| *AP1S1* | -0.331907 | 0.146376 | -2.267495 | 0.023360025 | 0.843463768 |
| *HDLBP* | 0.276333 | 0.121956 | 2.265849 | 0.02346064 | 0.844583041 |
| *KMT2E* | -0.192905 | 0.085197 | -2.264219 | 0.023560671 | 0.845674743 |
| *PAXX* | 0.357643 | 0.158229 | 2.260289 | 0.023803337 | 0.851864566 |
| *PKD1* | 0.495828 | 0.21971 | 2.256742 | 0.024024221 | 0.852764869 |
| *EXOC8* | 0.59023 | 0.261589 | 2.256326 | 0.024050243 | 0.852764869 |
| *PIGH* | 0.305861 | 0.135577 | 2.255992 | 0.024071162 | 0.852764869 |
| *CNR1* | 1.018237 | 0.45169 | 2.254282 | 0.024178429 | 0.852764869 |
| *MON2* | -0.419408 | 0.186052 | -2.254258 | 0.024179947 | 0.852764869 |
| *TGFB1* | 0.227257 | 0.100997 | 2.250134 | 0.024440461 | 0.859032694 |
| *TMEM14A* | -0.760144 | 0.338017 | -2.248832 | 0.024523178 | 0.859032694 |
| *FCMR* | 0.514988 | 0.229101 | 2.24787 | 0.024584511 | 0.859032694 |
| *CABLES1* | -0.886468 | 0.394514 | -2.246986 | 0.024640898 | 0.859032694 |
| *TRIM72* | 1.958977 | 0.873209 | 2.243423 | 0.024869576 | 0.860755606 |
| *PRDX4* | -0.279723 | 0.124746 | -2.242335 | 0.024939706 | 0.860755606 |
| *FXYD7* | -0.802554 | 0.358515 | -2.238548 | 0.025185317 | 0.860755606 |
| *RPA2* | 0.163575 | 0.073073 | 2.238525 | 0.025186816 | 0.860755606 |
| *RAB30* | 0.288258 | 0.128777 | 2.238432 | 0.025192908 | 0.860755606 |
| *PTGER4* | -0.553118 | 0.247163 | -2.237864 | 0.025229928 | 0.860755606 |
| *CAND1* | -0.217975 | 0.097404 | -2.237857 | 0.025230381 | 0.860755606 |
| *PTMS* | -0.720704 | 0.32215 | -2.237172 | 0.025275132 | 0.860755606 |
| *TMOD2* | -0.401922 | 0.179722 | -2.23635 | 0.025328862 | 0.860755606 |
| *ABCB4* | 0.513057 | 0.229639 | 2.234191 | 0.02547051 | 0.863151483 |
| *LILRA4* | 2.526109 | 1.131762 | 2.232015 | 0.025613999 | 0.865596217 |
| *ENSG00000260278* | -0.866375 | 0.388819 | -2.228224 | 0.025865579 | 0.869392884 |
| *DDIT4* | 0.398482 | 0.178952 | 2.226756 | 0.025963588 | 0.869392884 |
| *SLC12A6* | 0.279361 | 0.125473 | 2.226462 | 0.025983227 | 0.869392884 |
| *CBX4* | 0.801718 | 0.360158 | 2.226018 | 0.026012992 | 0.869392884 |
| *ENSG00000283013* | 0.413042 | 0.185729 | 2.223902 | 0.026155018 | 0.871379851 |
| *HIST1H2BD* | 0.361774 | 0.162742 | 2.222996 | 0.026216093 | 0.871379851 |
| *ENSG00000263394* | 0.881291 | 0.396764 | 2.221197 | 0.02633763 | 0.872813537 |
| *NDUFB6* | 0.197964 | 0.089164 | 2.220231 | 0.026403113 | 0.872813537 |
| *COPS6* | 0.167202 | 0.075369 | 2.218444 | 0.026524545 | 0.874445048 |
| *SUMO1* | -0.121373 | 0.054798 | -2.214927 | 0.026765089 | 0.879983899 |
| *PARP14* | -0.235849 | 0.106612 | -2.21221 | 0.026952177 | 0.882041009 |
| *PRKAG2* | -0.355225 | 0.160685 | -2.210696 | 0.027056907 | 0.882041009 |
| *TIPARP* | -0.275902 | 0.124804 | -2.210676 | 0.027058294 | 0.882041009 |
| *BRAT1* | 0.481258 | 0.217846 | 2.209162 | 0.0271634 | 0.882041009 |
| *EXOSC5* | 0.361271 | 0.163563 | 2.208762 | 0.027191175 | 0.882041009 |
| *CCDC154* | 0.957213 | 0.43368 | 2.207186 | 0.02730108 | 0.883244528 |

| | | | | | |
|---|---|---|---|---|---|
| *TRIM25* | 0.524036 | 0.238207 | 2.199919 | 0.027812645 | 0.89740163 |
| *CFP* | -1.937156 | 0.881814 | -2.196785 | 0.028035808 | NA |
| *ETFRF1* | 0.352526 | 0.160722 | 2.193395 | 0.028278934 | 0.910026586 |
| *ATXN10* | 0.291349 | 0.133053 | 2.189724 | 0.028544232 | 0.915294521 |
| *PSMD4* | 0.170411 | 0.077847 | 2.189046 | 0.028593523 | 0.915294521 |
| *CCAR1* | 0.201262 | 0.092084 | 2.185635 | 0.028842314 | 0.920828838 |
| *CCDC88B* | 0.257207 | 0.117841 | 2.182658 | 0.029060989 | 0.925375129 |
| *TLR10* | -0.417036 | 0.191459 | -2.178197 | 0.029391405 | 0.927971908 |
| *C16orf70* | -0.560186 | 0.257268 | -2.177441 | 0.029447706 | 0.927971908 |
| *ASH1L* | -0.195403 | 0.089805 | -2.175868 | 0.029565131 | 0.927971908 |
| *EWSR1* | 0.269711 | 0.123963 | 2.175731 | 0.029575349 | 0.927971908 |
| *IQGAP1* | -0.183179 | 0.0842 | -2.175526 | 0.029590722 | 0.927971908 |
| *PTCD3* | 0.234193 | 0.107656 | 2.175382 | 0.029601478 | 0.927971908 |
| *FLNA* | -0.274315 | 0.12631 | -2.171762 | 0.029873654 | 0.934090648 |
| *ADAM28* | 0.369721 | 0.170393 | 2.169809 | 0.030021348 | 0.9362956 |
| *DPEP3* | 2.101063 | 0.970733 | 2.164408 | 0.030433041 | NA |
| *EML2* | 0.321961 | 0.148845 | 2.163057 | 0.030536827 | 0.947957041 |
| *STIM2* | -0.406045 | 0.187735 | -2.162865 | 0.030551533 | 0.947957041 |
| *ENSG00000273472* | 0.631832 | 0.292379 | 2.161003 | 0.030695111 | 0.949011588 |
| *MYH3* | 0.757245 | 0.350727 | 2.15907 | 0.030844756 | 0.949011588 |
| *RRP36* | 0.355662 | 0.164778 | 2.158424 | 0.030894863 | 0.949011588 |
| *ENSG00000267169* | -0.841932 | 0.390078 | -2.158367 | 0.030899326 | 0.949011588 |
| *OTUD4* | 0.330881 | 0.153372 | 2.157373 | 0.030976639 | 0.949011588 |
| *PPP1R3D* | -1.286487 | 0.596742 | -2.155851 | 0.031095307 | 0.950247526 |
| *ENSG00000278376* | 0.604691 | 0.280961 | 2.152224 | 0.031379751 | 0.95646903 |
| *FRY.AS1* | -1.43633 | 0.667944 | -2.150375 | 0.031525547 | 0.95646903 |
| *MEI1* | -0.577974 | 0.269053 | -2.148179 | 0.031699504 | 0.95646903 |
| *CR1* | 1.584478 | 0.737789 | 2.147602 | 0.03174536 | 0.95646903 |
| *LINC00158* | -0.383807 | 0.178848 | -2.14599 | 0.031873783 | 0.95646903 |
| *PRKCE* | 0.356817 | 0.166317 | 2.145396 | 0.031921167 | 0.95646903 |
| *CFL1* | -0.122644 | 0.057192 | -2.144431 | 0.031998356 | 0.95646903 |
| *ARHGEF1* | 0.306842 | 0.143118 | 2.14398 | 0.032034498 | 0.95646903 |
| *ZNF33B* | -0.347691 | 0.162175 | -2.143927 | 0.032038771 | 0.95646903 |
| *C1orf50* | -0.25756 | 0.120187 | -2.142998 | 0.032113282 | 0.95646903 |
| *NSMAF* | -0.52765 | 0.24636 | -2.141781 | 0.032211075 | 0.95646903 |
| *PRMT2* | -0.21615 | 0.100941 | -2.141361 | 0.032244958 | 0.95646903 |
| *ATAD2* | 0.425762 | 0.198925 | 2.140316 | 0.032329277 | 0.956631184 |
| *ENSG00000231609* | 1.125003 | 0.526527 | 2.136648 | 0.032626599 | 0.962364926 |
| *ARL5A* | -0.176546 | 0.082654 | -2.135972 | 0.032681697 | 0.962364926 |
| *ENSG00000269482* | 1.917492 | 0.897982 | 2.135336 | 0.032733576 | NA |
| *UTP14C* | 0.513652 | 0.240882 | 2.132377 | 0.032975876 | 0.967049198 |
| *DDX10* | 0.345972 | 0.162336 | 2.131205 | 0.033072273 | 0.967049198 |
| *ASCC1* | 0.26807 | 0.125789 | 2.131112 | 0.033079906 | 0.967049198 |
| *PTTG1* | -0.395231 | 0.185947 | -2.125502 | 0.033544774 | 0.976213555 |
| *ATP8A1* | 0.417149 | 0.19627 | 2.125387 | 0.033554323 | 0.976213555 |
| *ZDHHC14* | -0.480173 | 0.226034 | -2.124338 | 0.033641896 | 0.976419821 |
| *ARID5B* | -0.3847 | 0.181216 | -2.122881 | 0.033763802 | 0.976997922 |

| | | | | | |
|---|---|---|---|---|---|
| *TPM3* | -0.198951 | 0.093749 | -2.122177 | 0.033822876 | 0.976997922 |
| *ENSG00000272843* | -1.237232 | 0.583329 | -2.120987 | 0.033922862 | 0.977025966 |
| *WDR82* | 0.225246 | 0.106235 | 2.120251 | 0.033984912 | 0.977025966 |
| *CPNE5* | 0.547758 | 0.258611 | 2.118079 | 0.034168393 | 0.977909547 |
| *TSC22D1* | -0.82205 | 0.388129 | -2.117979 | 0.034176859 | 0.977909547 |
| *GTF3A* | 0.163562 | 0.077269 | 2.116776 | 0.034278833 | 0.978519533 |
| *SMAP2* | -0.202093 | 0.095564 | -2.114727 | 0.034453263 | 0.981165578 |
| *KDM4B* | 0.269013 | 0.127266 | 2.113789 | 0.034533276 | 0.981165578 |
| *LAMTOR2* | -0.26465 | 0.125283 | -2.112421 | 0.034650341 | 0.982191437 |
| *TMEM219* | -0.220309 | 0.10439 | -2.11044 | 0.034820509 | 0.984714265 |
| *SH2B3* | 0.322157 | 0.152824 | 2.108022 | 0.035029128 | 0.988310195 |
| *POLR2M* | -0.224511 | 0.106788 | -2.102387 | 0.035519384 | 0.996406127 |
| *SLC17A9* | 0.689978 | 0.328274 | 2.101836 | 0.035567658 | 0.996406127 |
| *UNC79* | 1.692767 | 0.805582 | 2.101298 | 0.035614772 | 0.996406127 |
| *MSL3* | -0.194983 | 0.092807 | -2.100959 | 0.035644598 | 0.996406127 |
| *SUSD6* | -0.27581 | 0.131461 | -2.098041 | 0.035901521 | 0.999017259 |
| *ADO* | -0.458017 | 0.218346 | -2.097665 | 0.035934727 | 0.999017259 |
| *TANGO6* | 0.60547 | 0.288718 | 2.097096 | 0.035985043 | 0.999017259 |
| *ENSG00000259820* | 1.379818 | 0.658365 | 2.095825 | 0.036097742 | 0.999195731 |
| *ENSG00000281325* | -0.426657 | 0.203689 | -2.094648 | 0.036202284 | 0.999195731 |
| *AHCYL1* | 0.248652 | 0.118731 | 2.094241 | 0.036238553 | 0.999195731 |
| *DMD* | -0.547915 | 0.26177 | -2.093115 | 0.036338869 | 0.999689715 |
| *RHOC* | -0.371945 | 0.177825 | -2.091627 | 0.03647187 | 0.99991192 |
| *DSEL* | -1.068615 | 0.5117 | -2.088362 | 0.036765209 | 0.99991192 |
| *ZBTB20* | 0.374929 | 0.179592 | 2.087674 | 0.03682729 | 0.99991192 |
| *ZNF506* | 0.316728 | 0.151849 | 2.085802 | 0.036996537 | 0.99991192 |
| *SAMD1* | -0.486099 | 0.233231 | -2.084197 | 0.037142207 | 0.99991192 |
| *MEF2A* | 0.267289 | 0.128406 | 2.081598 | 0.037379201 | 0.99991192 |
| *ENSG00000272716* | 0.784418 | 0.377061 | 2.080349 | 0.037493533 | 0.99991192 |
| *SYVN1* | 0.313478 | 0.15076 | 2.079312 | 0.037588724 | 0.99991192 |
| *ENSG00000258623* | -0.762831 | 0.367216 | -2.077337 | 0.037770423 | 0.99991192 |
| *WDR74* | 0.396508 | 0.191552 | 2.069978 | 0.038454428 | 0.99991192 |
| *PIK3CD* | 0.273955 | 0.132577 | 2.066395 | 0.038791186 | 0.99991192 |
| *TBL1XR1* | -0.217817 | 0.10552 | -2.064221 | 0.038996749 | 0.99991192 |
| *MAD2L2* | -0.294441 | 0.142657 | -2.063974 | 0.039020134 | 0.99991192 |
| *ENSG00000089127* | 0.644411 | 0.312434 | 2.062552 | 0.039155186 | 0.99991192 |
| *DIAPH1* | -0.287393 | 0.139466 | -2.060669 | 0.03933459 | 0.99991192 |
| *MAP1S* | 0.362201 | 0.175798 | 2.060329 | 0.039367058 | 0.99991192 |
| *STX2* | -0.443047 | 0.215134 | -2.059401 | 0.039455836 | 0.99991192 |
| *TMEM191C* | 0.763947 | 0.371555 | 2.056079 | 0.03977492 | 0.99991192 |
| *COX11* | -0.280081 | 0.136224 | -2.056023 | 0.039780333 | 0.99991192 |
| *KCP* | 1.458795 | 0.710298 | 2.053779 | 0.039997092 | 0.99991192 |
| *RASGRP1* | -0.421474 | 0.205242 | -2.053549 | 0.040019393 | 0.99991192 |
| *SMC6* | -0.189885 | 0.092557 | -2.051546 | 0.040213851 | 0.99991192 |
| *SAMD4A* | -0.843929 | 0.411679 | -2.049967 | 0.040367658 | 0.99991192 |
| *SNURF* | 0.841268 | 0.410567 | 2.049041 | 0.040458143 | 0.99991192 |
| *TAF1B* | -0.469439 | 0.22925 | -2.047715 | 0.040587921 | 0.99991192 |

| | | | | | |
|---|---|---|---|---|---|
| *RBM38* | 0.249444 | 0.121892 | 2.046432 | 0.040713935 | 0.99991192 |
| *UTP3* | 0.235624 | 0.11517 | 2.045891 | 0.040767126 | 0.99991192 |
| *ZBTB39* | 0.703835 | 0.344307 | 2.044207 | 0.040933055 | 0.99991192 |
| *ATP2B1* | -0.216847 | 0.106117 | -2.043479 | 0.04100504 | 0.99991192 |
| *MIR181A1HG* | 0.468269 | 0.229174 | 2.043289 | 0.041023823 | 0.99991192 |
| *ELP4* | 0.439182 | 0.214984 | 2.04286 | 0.041066348 | 0.99991192 |
| *NEDD8* | -0.115112 | 0.056349 | -2.042825 | 0.041069732 | 0.99991192 |
| *AP3B1* | 0.432062 | 0.211645 | 2.041447 | 0.041206451 | 0.99991192 |
| *IFIT5* | -0.458412 | 0.224626 | -2.040782 | 0.04127253 | 0.99991192 |
| *TNFRSF13B* | -0.818944 | 0.401396 | -2.04024 | 0.041326425 | 0.99991192 |
| *BHLHE40* | 0.315968 | 0.154937 | 2.039329 | 0.04141718 | 0.99991192 |
| *FADS3* | -0.332959 | 0.163289 | -2.039078 | 0.041442291 | 0.99991192 |
| *MICOS10* | 0.123313 | 0.060509 | 2.037929 | 0.041557017 | 0.99991192 |
| *CD79A* | 0.319856 | 0.157203 | 2.034663 | 0.041884778 | 0.99991192 |
| *MAP3K2* | -0.276981 | 0.136294 | -2.032231 | 0.042130298 | 0.99991192 |
| *CTSB* | 0.273868 | 0.134807 | 2.031565 | 0.042197735 | 0.99991192 |
| *SAT2* | -0.211369 | 0.104099 | -2.030471 | 0.042308732 | 0.99991192 |
| *VIRMA* | 0.279086 | 0.137517 | 2.029463 | 0.04241118 | 0.99991192 |
| *CPEB4* | 0.490584 | 0.241843 | 2.02852 | 0.04250716 | 0.99991192 |
| *UBE2D4* | -0.331477 | 0.163522 | -2.027109 | 0.042651299 | 0.99991192 |
| *BNIP3* | -0.254064 | 0.125379 | -2.026368 | 0.042727134 | 0.99991192 |
| *NOP14.AS1* | -0.692084 | 0.341706 | -2.025375 | 0.042828859 | 0.99991192 |
| *TOE1* | 0.235814 | 0.116432 | 2.025337 | 0.042832799 | 0.99991192 |
| *CELF4* | -1.827377 | 0.902975 | -2.02373 | 0.042997961 | NA |
| *STRN* | 0.39695 | 0.196428 | 2.02084 | 0.043296363 | 0.99991192 |
| *TRBC2* | -0.183245 | 0.090692 | -2.020529 | 0.04332859 | 0.99991192 |
| *TGFBR2* | -0.227654 | 0.112739 | -2.019297 | 0.043456389 | 0.99991192 |
| *WARS* | 0.604007 | 0.299503 | 2.016697 | 0.043727171 | 0.99991192 |
| *CCL2* | 2.498478 | 1.239449 | 2.015797 | NA | NA |
| *RAB13* | -0.436097 | 0.216417 | -2.01508 | 0.043896276 | 0.99991192 |
| *IGFLR1* | -0.374876 | 0.18644 | -2.010701 | 0.044357056 | 0.99991192 |
| *FADS2* | 1.173248 | 0.583803 | 2.009666 | 0.044466499 | 0.99991192 |
| *IDE* | -0.573144 | 0.285269 | -2.009133 | 0.04452308 | 0.99991192 |
| *ADRB2* | -0.827001 | 0.412324 | -2.005708 | 0.044887439 | 0.99991192 |
| *MYADM* | 0.363771 | 0.181446 | 2.004837 | 0.044980466 | 0.99991192 |
| *CCT8* | 0.204432 | 0.102013 | 2.003982 | 0.045071955 | 0.99991192 |
| *LGALS1* | -0.456311 | 0.22777 | -2.003384 | 0.045136115 | 0.99991192 |
| *ATP6V1E1* | 0.242241 | 0.121023 | 2.001615 | 0.045326185 | 0.99991192 |
| *NMT2* | -0.43517 | 0.217417 | -2.001542 | 0.045334022 | 0.99991192 |
| *RPS6KB2* | -0.170357 | 0.085328 | -1.996498 | 0.045879786 | 0.99991192 |
| *SIK3* | 0.40949 | 0.205405 | 1.993577 | 0.046198304 | 0.99991192 |
| *RPP25L* | 0.255634 | 0.128242 | 1.993364 | 0.04622162 | 0.99991192 |
| *TAF9B* | -0.359349 | 0.180381 | -1.992172 | 0.046352191 | 0.99991192 |
| *EGLN2* | -0.175615 | 0.088173 | -1.991719 | 0.046401945 | 0.99991192 |
| *UTP23* | 0.254434 | 0.127748 | 1.991687 | 0.046405394 | 0.99991192 |
| *CTNND1* | -0.41456 | 0.208315 | -1.990065 | 0.046583819 | 0.99991192 |
| *SOGA1* | -0.649921 | 0.326914 | -1.988047 | 0.046806438 | 0.99991192 |

| | | | | | |
|---|---|---|---|---|---|
| *TP53I3* | -0.908808 | 0.457373 | -1.987017 | 0.046920519 | 0.99991192 |
| *GPR180* | -0.693306 | 0.349462 | -1.983924 | 0.047264299 | 0.99991192 |
| *SHISAL2A* | 0.356614 | 0.179791 | 1.983496 | 0.047312002 | 0.99991192 |
| *ENSG00000226571* | -0.640096 | 0.323103 | -1.981089 | 0.04758124 | 0.99991192 |
| *GPS2* | -0.284471 | 0.143633 | -1.980546 | 0.047642239 | 0.99991192 |
| *L1CAM* | -0.628572 | 0.317603 | -1.979112 | 0.047803394 | 0.99991192 |
| *ATG13* | 0.236143 | 0.119585 | 1.974683 | 0.048304112 | 0.99991192 |
| *CARHSP1* | 0.201049 | 0.101907 | 1.972861 | 0.048511379 | 0.99991192 |
| *SNAPIN* | -0.22874 | 0.115969 | -1.972426 | 0.048560953 | 0.99991192 |
| *CCNT2* | -0.261792 | 0.132774 | -1.971713 | 0.048642336 | 0.99991192 |
| *NRP2* | -0.400248 | 0.203137 | -1.970332 | 0.048800303 | 0.99991192 |
| *SPACA9* | -0.768868 | 0.390329 | -1.969797 | 0.048861677 | 0.99991192 |
| *EPHB6* | -0.396249 | 0.201247 | -1.96897 | 0.0489565 | 0.99991192 |
| *PFN1* | -0.158729 | 0.080667 | -1.967709 | 0.04910158 | 0.99991192 |
| *MEN1* | -0.335494 | 0.17058 | -1.96678 | 0.049208552 | 0.99991192 |
| *VPS37C* | -0.476902 | 0.24257 | -1.966036 | 0.049294426 | 0.99991192 |
| *TPRA1* | 0.375977 | 0.191241 | 1.965983 | 0.049300616 | 0.99991192 |
| *TP53I11* | -0.342961 | 0.174488 | -1.965525 | 0.049353518 | 0.99991192 |

| Cluster 10 | log2FC | lfcSE | stat | pvalue | padj |
|---|---|---|---|---|---|
| *VPREB3* | 3.983177 | 0.144048 | 27.651685 | < 2.22e-16 | < 2.22e-16 |
| *GRN* | 1.648943 | 0.091766 | 17.969007 | < 2.22e-16 | < 2.22e-16 |
| *P2RX5* | 1.833474 | 0.107016 | 17.132704 | < 2.22e-16 | < 2.22e-16 |
| *CD1D* | 5.842222 | 0.369987 | 15.790325 | < 2.22e-16 | < 2.22e-16 |
| *DENND6B* | 2.997422 | 0.198313 | 15.114605 | < 2.22e-16 | < 2.22e-16 |
| *NRIP1* | 2.153634 | 0.153913 | 13.992519 | < 2.22e-16 | < 2.22e-16 |
| *CD38* | 3.832 | 0.302384 | 12.672646 | < 2.22e-16 | < 2.22e-16 |
| *RABGAP1L* | 1.670832 | 0.135364 | 12.343296 | < 2.22e-16 | < 2.22e-16 |
| *PLAAT4* | 2.455264 | 0.216315 | 11.350397 | < 2.22e-16 | < 2.22e-16 |
| *HIC1* | 4.409995 | 0.399595 | 11.036159 | < 2.22e-16 | < 2.22e-16 |
| *MRPS6* | 0.95425 | 0.08714 | 10.950712 | < 2.22e-16 | < 2.22e-16 |
| *SLC5A3* | 1.409247 | 0.137919 | 10.217929 | < 2.22e-16 | < 2.22e-16 |
| *FCRL2* | 1.845225 | 0.182441 | 10.11411 | < 2.22e-16 | < 2.22e-16 |
| *C12orf65* | 0.947412 | 0.096472 | 9.820589 | < 2.22e-16 | < 2.22e-16 |
| *ASB2* | 5.768485 | 0.59305 | 9.726808 | < 2.22e-16 | < 2.22e-16 |
| *RNF207* | 3.276308 | 0.348802 | 9.39302 | < 2.22e-16 | < 2.22e-16 |
| *SLC2A5* | 3.370011 | 0.365318 | 9.224866 | < 2.22e-16 | < 2.22e-16 |
| *MS4A1* | -0.778299 | 0.085783 | -9.072913 | < 2.22e-16 | < 2.22e-16 |
| *CXCR5* | 0.814065 | 0.093325 | 8.72287 | < 2.22e-16 | 1.66E-15 |
| *CACYBP* | 0.908726 | 0.106092 | 8.565456 | < 2.22e-16 | 6.26E-15 |
| *SMAD3* | 1.092572 | 0.128741 | 8.486565 | < 2.22e-16 | 1.18E-14 |
| *SLC12A4* | 2.064839 | 0.246099 | 8.390282 | < 2.22e-16 | 2.56E-14 |
| *CR2* | 2.339262 | 0.284215 | 8.230598 | < 2.22e-16 | 9.42E-14 |
| *SEPTIN9* | 0.49165 | 0.060709 | 8.098435 | 5.57E-16 | 2.70E-13 |
| *CTSA* | 0.704792 | 0.08825 | 7.986324 | 1.39E-15 | 6.47E-13 |
| *SIGIRR* | 0.959555 | 0.120484 | 7.964148 | 1.66E-15 | 7.44E-13 |
| *P2RX5.TAX1BP3* | 1.823501 | 0.235258 | 7.751083 | 9.11E-15 | 3.93E-12 |
| *MAPKAPK2* | 0.969027 | 0.126805 | 7.641892 | 2.14E-14 | 8.89E-12 |
| *AHI1* | 1.15612 | 0.151991 | 7.606503 | 2.82E-14 | 1.13E-11 |
| *CMTM7* | 1.28791 | 0.169881 | 7.581229 | 3.42E-14 | 1.33E-11 |
| *IL1B* | 3.0507 | 0.408709 | 7.464232 | 8.38E-14 | 3.14E-11 |
| *ZNF563* | 1.297376 | 0.174453 | 7.436836 | 1.03E-13 | 3.75E-11 |
| *IRF2BPL* | 1.187376 | 0.160456 | 7.399994 | 1.36E-13 | 4.80E-11 |
| *XYLT1* | 1.111666 | 0.150364 | 7.393153 | 1.43E-13 | 4.91E-11 |
| *PISD* | 1.065856 | 0.144894 | 7.356117 | 1.89E-13 | 6.29E-11 |
| *RAB20* | 1.818446 | 0.248177 | 7.327207 | 2.35E-13 | 7.59E-11 |
| *CHST15* | 1.133829 | 0.156917 | 7.225672 | 4.99E-13 | 1.57E-10 |
| *RMDN2* | 1.68537 | 0.238714 | 7.060218 | 1.66E-12 | 5.09E-10 |
| *PNOC* | 1.005383 | 0.142652 | 7.047781 | 1.82E-12 | 5.42E-10 |
| *ST3GAL1* | 0.956853 | 0.135921 | 7.039798 | 1.93E-12 | 5.60E-10 |
| *ENDOD1* | 1.58254 | 0.225575 | 7.01559 | 2.29E-12 | 6.50E-10 |
| *GPX1* | 0.572024 | 0.082439 | 6.938788 | 3.95E-12 | 1.10E-09 |
| *LHFPL2* | 1.53976 | 0.228931 | 6.725862 | 1.75E-11 | 4.72E-09 |
| *PHF21A* | -0.880336 | 0.131331 | -6.703163 | 2.04E-11 | 5.39E-09 |
| *SLC5A5* | 4.873284 | 0.745746 | 6.534779 | 6.37E-11 | 1.65E-08 |
| *C1orf162* | -0.859663 | 0.133309 | -6.448634 | 1.13E-10 | 2.85E-08 |

| | | | | | |
|---|---|---|---|---|---|
| *PLAAT3* | 2.193328 | 0.345227 | 6.353284 | 2.11E-10 | 5.22E-08 |
| *RARA.AS1* | 2.683833 | 0.42503 | 6.314458 | 2.71E-10 | 6.46E-08 |
| *RHOBTB3* | 1.732839 | 0.274453 | 6.313793 | 2.72E-10 | 6.46E-08 |
| *AK2* | 0.492716 | 0.078203 | 6.300479 | 2.97E-10 | 6.90E-08 |
| *EXOSC5* | 0.628155 | 0.100586 | 6.244926 | 4.24E-10 | 9.67E-08 |
| *BTBD7* | 0.668557 | 0.107197 | 6.236723 | 4.47E-10 | 1.00E-07 |
| *ZNF3* | 0.931616 | 0.149856 | 6.216738 | 5.08E-10 | 1.11E-07 |
| *CHST7* | 1.90862 | 0.310068 | 6.155486 | 7.48E-10 | 1.61E-07 |
| *ACP5* | 0.696123 | 0.114313 | 6.089615 | 1.13E-09 | 2.38E-07 |
| *POU2F2* | -0.433043 | 0.071133 | -6.087835 | 1.14E-09 | 2.38E-07 |
| *GCHFR* | -0.572332 | 0.09471 | -6.042981 | 1.51E-09 | 3.09E-07 |
| *TMEM159* | -1.167094 | 0.194658 | -5.995626 | 2.03E-09 | 4.07E-07 |
| *MANF* | 0.647309 | 0.108233 | 5.980676 | 2.22E-09 | 4.38E-07 |
| *ZFP36L1* | -0.441244 | 0.074788 | -5.899905 | 3.64E-09 | 7.05E-07 |
| *SMC6* | -0.484552 | 0.082362 | -5.883189 | 4.02E-09 | 7.68E-07 |
| *SLC25A39* | 0.493878 | 0.084679 | 5.832327 | 5.47E-09 | 1.02E-06 |
| *CIB1* | 0.363813 | 0.06239 | 5.831268 | 5.50E-09 | 1.02E-06 |
| *CD1C* | 2.129751 | 0.366379 | 5.812975 | 6.14E-09 | 1.11E-06 |
| *MPEG1* | 0.992793 | 0.170853 | 5.810807 | 6.22E-09 | 1.11E-06 |
| *HSPA5* | 0.605095 | 0.104359 | 5.798228 | 6.70E-09 | 1.18E-06 |
| *ICAM3* | -0.441281 | 0.07625 | -5.787322 | 7.15E-09 | 1.24E-06 |
| *SRGN* | 0.65999 | 0.114093 | 5.784642 | 7.27E-09 | 1.24E-06 |
| *NME1* | 0.450916 | 0.078669 | 5.731817 | 9.94E-09 | 1.68E-06 |
| *IFI30* | 0.842136 | 0.147566 | 5.706849 | 1.15E-08 | 1.91E-06 |
| *LZTFL1* | 1.250148 | 0.219144 | 5.704698 | 1.17E-08 | 1.91E-06 |
| *VPREB1* | 4.898095 | 0.859959 | 5.695733 | 1.23E-08 | 1.98E-06 |
| *CD82* | -0.53315 | 0.093652 | -5.692909 | 1.25E-08 | 1.99E-06 |
| *JCHAIN* | 1.960433 | 0.344503 | 5.690614 | 1.27E-08 | 1.99E-06 |
| *ENSG00000254802* | 1.281403 | 0.226066 | 5.668268 | 1.44E-08 | 2.24E-06 |
| *NFAT5* | 0.657633 | 0.116603 | 5.639952 | 1.70E-08 | 2.60E-06 |
| *ACAP1* | -0.518332 | 0.091941 | -5.637669 | 1.72E-08 | 2.60E-06 |
| *ENSG00000166927* | -1.155884 | 0.205709 | -5.619012 | 1.92E-08 | 2.86E-06 |
| *DDX24* | 0.437937 | 0.078265 | 5.595578 | 2.20E-08 | 3.24E-06 |
| *TSPAN18* | 2.833456 | 0.509047 | 5.566202 | 2.60E-08 | 3.79E-06 |
| *AGPAT5* | 0.879902 | 0.158974 | 5.534896 | 3.11E-08 | 4.47E-06 |
| *CD72* | 1.180732 | 0.214156 | 5.513428 | 3.52E-08 | 4.97E-06 |
| *H3F3A* | 0.471046 | 0.085454 | 5.51226 | 3.54E-08 | 4.97E-06 |
| *WARS* | 1.094703 | 0.198862 | 5.504846 | 3.69E-08 | 5.09E-06 |
| *TSPO* | -0.528454 | 0.096017 | -5.503762 | 3.72E-08 | 5.09E-06 |
| *CR1* | 2.833472 | 0.517453 | 5.475801 | 4.36E-08 | 5.89E-06 |
| *DDX21* | 0.456191 | 0.083407 | 5.469469 | 4.51E-08 | 6.04E-06 |
| *GEN1* | -0.907255 | 0.165989 | -5.465758 | 4.61E-08 | 6.09E-06 |
| *FADS3* | -0.960922 | 0.175937 | -5.461741 | 4.71E-08 | 6.16E-06 |
| *LBH* | -0.645551 | 0.119398 | -5.406702 | 6.42E-08 | 8.30E-06 |
| *TMSB10* | -0.314885 | 0.058527 | -5.38013 | 7.44E-08 | 9.52E-06 |
| *BIN1* | -0.568549 | 0.105831 | -5.372223 | 7.78E-08 | 9.83E-06 |
| *TRABD* | 0.46774 | 0.087834 | 5.325268 | 1.01E-07 | 1.25E-05 |

| | | | | | |
|---|---|---|---|---|---|
| *RAB37* | 1.303589 | 0.244799 | 5.325148 | 1.01E-07 | 1.25E-05 |
| *ATP1B3* | 0.502045 | 0.094583 | 5.307971 | 1.11E-07 | 1.35E-05 |
| *VIM* | -1.297857 | 0.244554 | -5.307029 | 1.11E-07 | 1.35E-05 |
| *PUS1* | 0.637097 | 0.120085 | 5.305371 | 1.12E-07 | 1.35E-05 |
| *DUS2* | 0.806039 | 0.152685 | 5.279098 | 1.30E-07 | 1.54E-05 |
| *RINL* | 0.556501 | 0.106219 | 5.239205 | 1.61E-07 | 1.90E-05 |
| *PDLIM2* | -0.841925 | 0.161218 | -5.222278 | 1.77E-07 | 2.06E-05 |
| *GPSM3* | -0.363153 | 0.069651 | -5.213924 | 1.85E-07 | 2.13E-05 |
| *FXYD2* | -1.691484 | 0.326885 | -5.174553 | 2.28E-07 | 2.61E-05 |
| *SERPINF1* | -1.894627 | 0.366286 | -5.172533 | 2.31E-07 | 2.61E-05 |
| *NDUFA2* | 0.323369 | 0.062554 | 5.169432 | 2.35E-07 | 2.63E-05 |
| *LAT2* | -0.678266 | 0.131493 | -5.158177 | 2.49E-07 | 2.76E-05 |
| *LMNA* | -1.712033 | 0.334704 | -5.115066 | 3.14E-07 | 3.44E-05 |
| *MACROD2* | -0.916383 | 0.179682 | -5.100025 | 3.40E-07 | 3.69E-05 |
| *SREBF1* | 1.086003 | 0.2151 | 5.048818 | 4.45E-07 | 4.79E-05 |
| *ENSG00000237499* | -0.958226 | 0.190603 | -5.02733 | 4.97E-07 | 5.31E-05 |
| *IRF4* | 0.549635 | 0.10978 | 5.006697 | 5.54E-07 | 5.86E-05 |
| *LCAT* | 1.632951 | 0.326282 | 5.004722 | 5.59E-07 | 5.86E-05 |
| *OTUD7A* | 3.227232 | 0.648974 | 4.972821 | 6.60E-07 | 6.85E-05 |
| *NANP* | 1.233791 | 0.249027 | 4.95444 | 7.25E-07 | 7.47E-05 |
| *TLR10* | -0.571623 | 0.115486 | -4.949732 | 7.43E-07 | 7.55E-05 |
| *IQSEC1* | -0.607643 | 0.122786 | -4.948821 | 7.47E-07 | 7.55E-05 |
| *MAP3K1* | 0.629558 | 0.127734 | 4.928673 | 8.28E-07 | 8.30E-05 |
| *UBE2G1* | 0.64005 | 0.130027 | 4.922447 | 8.55E-07 | 8.50E-05 |
| *AP3S1* | 0.543852 | 0.110718 | 4.91205 | 9.01E-07 | 8.89E-05 |
| *NOLC1* | 0.534795 | 0.109214 | 4.896768 | 9.74E-07 | 9.52E-05 |
| *DNAJB11* | 0.497216 | 0.102186 | 4.86578 | 1.14E-06 | 0.000110529 |
| *DKC1* | 0.366208 | 0.075341 | 4.860702 | 1.17E-06 | 0.000112466 |
| *HNRNPLL* | 1.506956 | 0.310694 | 4.850298 | 1.23E-06 | 0.000117557 |
| *CD37* | -0.63608 | 0.131208 | -4.847864 | 1.25E-06 | 0.000118041 |
| *MRPL4* | 0.362829 | 0.075142 | 4.8286 | 1.37E-06 | 0.000128047 |
| *HIPK2* | 0.921838 | 0.190917 | 4.828481 | 1.38E-06 | 0.000128047 |
| *CD22* | 0.536267 | 0.111725 | 4.799876 | 1.59E-06 | 0.000146592 |
| *NEURL2* | 1.840873 | 0.385208 | 4.778903 | 1.76E-06 | 0.00016146 |
| *BLOC1S1* | -0.253917 | 0.053179 | -4.774727 | 1.80E-06 | 0.000163559 |
| *TP53I11* | -0.759227 | 0.159118 | -4.771485 | 1.83E-06 | 0.000164925 |
| *DDAH2* | -0.683977 | 0.143716 | -4.759229 | 1.94E-06 | 0.000173914 |
| *PLP2* | -0.729683 | 0.153397 | -4.756831 | 1.97E-06 | 0.000174648 |
| *ARL14EPL* | 4.160246 | 0.877011 | 4.743664 | 2.10E-06 | NA |
| *HCK* | -1.354058 | 0.288816 | -4.688305 | 2.75E-06 | 0.000242796 |
| *NDUFB9* | 0.252709 | 0.05415 | 4.666869 | 3.06E-06 | 0.000267515 |
| *TRBC2* | -0.348811 | 0.074884 | -4.658003 | 3.19E-06 | 0.000277212 |
| *TMEM191B* | 1.083338 | 0.23364 | 4.636774 | 3.54E-06 | 0.000303909 |
| *FKBP2* | 0.326492 | 0.070426 | 4.635971 | 3.55E-06 | 0.000303909 |
| *MYC* | 0.702075 | 0.15197 | 4.619821 | 3.84E-06 | 0.00032571 |
| *MARCHF9* | -0.675111 | 0.146172 | -4.618593 | 3.86E-06 | 0.00032571 |
| *RAB13* | -0.99161 | 0.214804 | -4.616344 | 3.91E-06 | 0.00032689 |

| | | | | | |
|---|---|---|---|---|---|
| *ENSG00000198106* | -0.919613 | 0.199361 | -4.612804 | 3.97E-06 | 0.000330134 |
| *GYPC* | 0.535855 | 0.116394 | 4.603816 | 4.15E-06 | 0.000342269 |
| *DPEP2* | 0.812522 | 0.176873 | 4.593819 | 4.35E-06 | 0.00035463 |
| *LAPTM5* | -0.420402 | 0.091521 | -4.593489 | 4.36E-06 | 0.00035463 |
| *XBP1* | 0.503329 | 0.10964 | 4.59074 | 4.42E-06 | 0.000356838 |
| *C15orf62* | -1.284964 | 0.280006 | -4.589065 | 4.45E-06 | 0.000357233 |
| *NOP16* | 0.350365 | 0.076395 | 4.586206 | 4.51E-06 | 0.000359676 |
| *PXK* | 0.693585 | 0.151298 | 4.584233 | 4.56E-06 | 0.000360619 |
| *QPRT* | 1.723283 | 0.376557 | 4.576423 | 4.73E-06 | 0.000371811 |
| *RRP9* | 0.590493 | 0.129313 | 4.566399 | 4.96E-06 | 0.000387416 |
| *RESF1* | -0.423549 | 0.093051 | -4.551793 | 5.32E-06 | 0.000412546 |
| *HNRNPC* | 0.287932 | 0.063367 | 4.54384 | 5.52E-06 | 0.000425593 |
| *GPATCH4* | 0.374344 | 0.08243 | 4.541354 | 5.59E-06 | 0.000427811 |
| *SESN3* | -0.407624 | 0.089869 | -4.53575 | 5.74E-06 | 0.000436459 |
| *RRBP1* | 0.832714 | 0.184269 | 4.519019 | 6.21E-06 | 0.00046934 |
| *TNK2* | 0.570775 | 0.126639 | 4.507083 | 6.57E-06 | 0.00049332 |
| *SDHB* | 0.343294 | 0.076259 | 4.501677 | 6.74E-06 | 0.000502794 |
| *CASP7* | 0.771851 | 0.171753 | 4.493965 | 6.99E-06 | 0.000518039 |
| *CTTNBP2NL* | -1.173194 | 0.261438 | -4.487471 | 7.21E-06 | 0.0005307 |
| *NUDC* | 0.300606 | 0.067227 | 4.471489 | 7.77E-06 | 0.000566496 |
| *PDIA6* | 0.489923 | 0.109582 | 4.470851 | 7.79E-06 | 0.000566496 |
| *PPP3CA* | -0.34058 | 0.076286 | -4.464524 | 8.02E-06 | 0.00057987 |
| *CNIH1* | 0.284805 | 0.063866 | 4.459432 | 8.22E-06 | 0.000590155 |
| *CD3EAP* | 0.666194 | 0.149499 | 4.456174 | 8.34E-06 | 0.000595513 |
| *RHOF* | -0.442465 | 0.099649 | -4.440229 | 8.99E-06 | 0.000637479 |
| *RRP15* | 0.362041 | 0.081669 | 4.433008 | 9.29E-06 | 0.000655222 |
| *ATP6V1E1* | 0.477884 | 0.107972 | 4.426 | 9.60E-06 | 0.000672786 |
| *ECE2* | 0.695423 | 0.157247 | 4.422485 | 9.76E-06 | 0.000679733 |
| *ISG20* | -0.582019 | 0.131707 | -4.419034 | 9.91E-06 | 0.000686564 |
| *CMSS1* | 0.410368 | 0.093348 | 4.396128 | 1.10E-05 | 0.000758614 |
| *ABCB1* | 1.131539 | 0.258384 | 4.379292 | 1.19E-05 | 0.000814829 |
| *PARP14* | -0.405808 | 0.092717 | -4.376848 | 1.20E-05 | 0.00081569 |
| *STK17B* | -0.440568 | 0.100666 | -4.376512 | 1.21E-05 | 0.00081569 |
| *GTPBP4* | 0.419059 | 0.095848 | 4.372133 | 1.23E-05 | 0.000827417 |
| *MRPS23* | 0.320303 | 0.073297 | 4.369925 | 1.24E-05 | 0.000831025 |
| *BCL2A1* | 0.651997 | 0.149543 | 4.359925 | 1.30E-05 | 0.000863889 |
| *GADD45GIP1* | 0.239598 | 0.054967 | 4.358947 | 1.31E-05 | 0.000863889 |
| *GBP4* | 0.711645 | 0.163866 | 4.342835 | 1.41E-05 | 0.000924512 |
| *DBNL* | 0.289311 | 0.066726 | 4.335784 | 1.45E-05 | 0.000949292 |
| *RBM47* | 1.336572 | 0.308361 | 4.334443 | 1.46E-05 | 0.000949488 |
| *C1orf216* | 0.699424 | 0.161424 | 4.33283 | 1.47E-05 | 0.000949488 |
| *PCSK7* | -0.300184 | 0.069294 | -4.332061 | 1.48E-05 | 0.000949488 |
| *GTF2E2* | 0.71023 | 0.164594 | 4.315035 | 1.60E-05 | 0.001020067 |
| *SRA1* | 0.399225 | 0.092965 | 4.294369 | 1.75E-05 | 0.001113755 |
| *SH3BGRL* | -0.283871 | 0.066439 | -4.272645 | 1.93E-05 | 0.001221364 |
| *PLXNA3* | -0.699769 | 0.164314 | -4.258735 | 2.06E-05 | 0.001292864 |
| *S100A4* | -0.880345 | 0.207318 | -4.246351 | 2.17E-05 | 0.001359051 |

| | | | | | |
|---|---|---|---|---|---|
| *LINC02201* | -1.407308 | 0.332385 | -4.23397 | 2.30E-05 | 0.001426605 |
| *MRTO4* | 0.445647 | 0.105278 | 4.23306 | 2.31E-05 | 0.001426605 |
| *KEAP1* | 0.491637 | 0.116595 | 4.216607 | 2.48E-05 | 0.001526611 |
| *NDUFAF3* | 0.291133 | 0.069148 | 4.210298 | 2.55E-05 | 0.001561617 |
| *GPR18* | 0.894958 | 0.212652 | 4.20855 | 2.57E-05 | 0.001565501 |
| *MYDGF* | 0.339681 | 0.080835 | 4.202173 | 2.64E-05 | 0.00159701 |
| *BHLHE40* | 0.535997 | 0.127567 | 4.201687 | 2.65E-05 | 0.00159701 |
| *NCL* | 0.279604 | 0.06685 | 4.182567 | 2.88E-05 | 0.001727754 |
| *SNRPD1* | 0.303879 | 0.072672 | 4.1815 | 2.90E-05 | 0.001727754 |
| *PDIA3* | 0.332568 | 0.079575 | 4.179301 | 2.92E-05 | 0.001735639 |
| *HP1BP3* | -0.29031 | 0.069524 | -4.175695 | 2.97E-05 | 0.001754415 |
| *C16orf54* | 0.549073 | 0.131577 | 4.173026 | 3.01E-05 | 0.001766142 |
| *APOL3* | 0.712473 | 0.171088 | 4.164378 | 3.12E-05 | 0.001825204 |
| *BMF* | -0.59357 | 0.142906 | -4.153569 | 3.27E-05 | 0.001904074 |
| *EBNA1BP2* | 0.431533 | 0.104057 | 4.147081 | 3.37E-05 | 0.00194908 |
| *MRPL3* | 0.383901 | 0.092758 | 4.138754 | 3.49E-05 | 0.00201117 |
| *HSPA9* | 0.341835 | 0.082633 | 4.136782 | 3.52E-05 | 0.002016798 |
| *SNRPD3* | 0.29769 | 0.071978 | 4.135851 | 3.54E-05 | 0.002016798 |
| *ATP13A2* | 0.918689 | 0.22377 | 4.105498 | 4.03E-05 | 0.002289595 |
| *SH3BGRL3* | -0.371061 | 0.09063 | -4.094221 | 4.24E-05 | 0.002392259 |
| *DCAF13* | 0.410603 | 0.100324 | 4.092778 | 4.26E-05 | 0.002395572 |
| *SYPL1* | -0.39545 | 0.096772 | -4.086396 | 4.38E-05 | 0.002450555 |
| *C15orf39* | 0.761643 | 0.186653 | 4.08053 | 4.49E-05 | 0.002501211 |
| *ADAMTS7* | -1.048672 | 0.257116 | -4.078597 | 4.53E-05 | 0.002510081 |
| *GNG11* | -1.197513 | 0.293817 | -4.075705 | 4.59E-05 | 0.002529436 |
| *ODF2L* | -0.492994 | 0.121078 | -4.071714 | 4.67E-05 | 0.002561041 |
| *ENO1* | 0.404024 | 0.099534 | 4.059154 | 4.93E-05 | 0.002690071 |
| *ACAP2* | -0.436523 | 0.107658 | -4.054713 | 5.02E-05 | 0.002728878 |
| *MCL1* | 0.350411 | 0.08653 | 4.049583 | 5.13E-05 | 0.002776408 |
| *SPOCK2* | 0.685469 | 0.169785 | 4.037271 | 5.41E-05 | 0.00291263 |
| *NDC1* | 0.681892 | 0.169431 | 4.024602 | 5.71E-05 | 0.003059789 |
| *CNP* | 0.495727 | 0.123621 | 4.010039 | 6.07E-05 | 0.003239841 |
| *FCRL1* | 0.878203 | 0.219201 | 4.006385 | 6.17E-05 | 0.003275317 |
| *WNT10A* | -1.177939 | 0.294361 | -4.001685 | 6.29E-05 | 0.003325893 |
| *EPHB6* | -0.803806 | 0.200991 | -3.999219 | 6.36E-05 | 0.003333582 |
| *GPR146* | -1.471711 | 0.36802 | -3.998997 | 6.36E-05 | 0.003333582 |
| *KDM2B* | 0.384589 | 0.096245 | 3.995947 | 6.44E-05 | 0.003361664 |
| *WDFY4* | -0.489932 | 0.122664 | -3.994084 | 6.49E-05 | 0.003373071 |
| *NHP2* | 0.269661 | 0.067855 | 3.974084 | 7.07E-05 | 0.003643345 |
| *SYNE3* | -0.996242 | 0.250711 | -3.973665 | 7.08E-05 | 0.003643345 |
| *FAM111B* | -0.87226 | 0.219604 | -3.971971 | 7.13E-05 | 0.003653195 |
| *TRAF3IP3* | -0.313002 | 0.078953 | -3.964429 | 7.36E-05 | 0.003750764 |
| *HOXB3* | 2.612695 | 0.659173 | 3.963597 | 7.38E-05 | 0.003750764 |
| *SH2D4A* | 1.931003 | 0.488103 | 3.956139 | 7.62E-05 | 0.003852913 |
| *CYTIP* | -0.462805 | 0.117142 | -3.950788 | 7.79E-05 | 0.003923035 |
| *DHRS1* | -0.547629 | 0.138747 | -3.946963 | 7.91E-05 | 0.003969046 |
| *ZNF506* | 0.643259 | 0.163414 | 3.936383 | 8.27E-05 | 0.004113909 |

| | | | | | |
|---|---|---|---|---|---|
| *MDM4* | -0.334546 | 0.08499 | -3.936307 | 8.27E-05 | 0.004113909 |
| *HPS3* | -0.653865 | 0.166681 | -3.922858 | 8.75E-05 | 0.004332043 |
| *NOC2L* | 0.347449 | 0.088666 | 3.918646 | 8.90E-05 | 0.004389755 |
| *ACSL1* | 0.635529 | 0.162426 | 3.912728 | 9.13E-05 | 0.004479786 |
| *SYTL1* | -0.470599 | 0.120309 | -3.911575 | 9.17E-05 | 0.004482331 |
| *CYB561A3* | -0.486816 | 0.12453 | -3.909231 | 9.26E-05 | 0.004507092 |
| *ENSG00000176320* | 3.56342 | 0.91365 | 3.900201 | 9.61E-05 | NA |
| *IZUMO4* | -0.559301 | 0.143534 | -3.896632 | 9.75E-05 | 0.004728225 |
| *SLC45A3* | 1.164326 | 0.299015 | 3.893872 | 9.87E-05 | 0.004762546 |
| *GPER1* | -2.971763 | 0.76434 | -3.888011 | 0.00010107 | NA |
| *PA2G4* | 0.250129 | 0.064365 | 3.886131 | 0.00010186 | 0.004896602 |
| *ARL4C* | -0.840583 | 0.216585 | -3.88108 | 0.00010399 | 0.004974851 |
| *CHST2* | 0.938527 | 0.241871 | 3.880277 | 0.00010434 | 0.004974851 |
| *CCDC124* | 0.371412 | 0.095763 | 3.878457 | 0.00010512 | 0.004991757 |
| *UBTF* | 0.449254 | 0.1161 | 3.869527 | 0.00010905 | 0.005157113 |
| *AIMP2* | 0.554656 | 0.143411 | 3.867604 | 0.00010991 | 0.005165184 |
| *PRDM2* | -0.404278 | 0.104541 | -3.867171 | 0.00011011 | 0.005165184 |
| *EIF6* | 0.262746 | 0.06802 | 3.862787 | 0.0001121 | 0.005237665 |
| *DENND3* | -0.684539 | 0.177407 | -3.85859 | 0.00011404 | 0.005307112 |
| *PEMT* | 0.666695 | 0.173369 | 3.845514 | 0.0001203 | 0.005575981 |
| *KLHL14* | -0.881591 | 0.229314 | -3.844465 | 0.00012082 | 0.005577657 |
| *TFAM* | 0.406504 | 0.105879 | 3.839341 | 0.00012337 | 0.005665483 |
| *FKBP4* | 0.372836 | 0.097133 | 3.838419 | 0.00012383 | 0.005665483 |
| *LY86* | -0.574794 | 0.149775 | -3.837726 | 0.00012418 | 0.005665483 |
| *CMIP* | -0.530022 | 0.138238 | -3.834132 | 0.00012601 | 0.00567019 |
| *COMMD6* | -0.214734 | 0.056014 | -3.833608 | 0.00012628 | 0.00567019 |
| *LPIN1* | -0.507127 | 0.132298 | -3.833219 | 0.00012648 | 0.00567019 |
| *RAD23A* | 0.186958 | 0.048782 | 3.832502 | 0.00012685 | 0.00567019 |
| *SRPRB* | 0.441734 | 0.115271 | 3.832144 | 0.00012703 | 0.00567019 |
| *MICOS10* | 0.275166 | 0.071812 | 3.831756 | 0.00012723 | 0.00567019 |
| *BRIX1* | 0.346183 | 0.090367 | 3.830865 | 0.00012769 | 0.00567019 |
| *TNRC6B* | -0.276066 | 0.072136 | -3.82704 | 0.00012969 | 0.005737074 |
| *RPN1* | 0.412912 | 0.107962 | 3.824606 | 0.00013098 | 0.005772104 |
| *LONP1* | 0.536267 | 0.140284 | 3.822738 | 0.00013198 | 0.005794079 |
| *GZMB* | 3.109875 | 0.815892 | 3.811626 | 0.00013806 | 0.006038118 |
| *ARID5B* | -0.390977 | 0.102707 | -3.806724 | 0.00014082 | 0.006125575 |
| *SIGLEC10* | 1.109806 | 0.291577 | 3.806217 | 0.00014111 | 0.006125575 |
| *KMO* | -0.686663 | 0.181087 | -3.791906 | 0.0001495 | 0.006441846 |
| *TMEM65* | 1.069985 | 0.282177 | 3.791896 | 0.0001495 | 0.006441846 |
| *NUBP1* | 0.396187 | 0.104682 | 3.784681 | 0.00015391 | 0.006607152 |
| *FBP1* | 1.835716 | 0.485759 | 3.779065 | 0.00015742 | 0.006733118 |
| *CLCF1* | -0.599467 | 0.15872 | -3.776872 | 0.00015881 | 0.006744684 |
| *BOP1* | 0.506693 | 0.134159 | 3.776812 | 0.00015885 | 0.006744684 |
| *SLCO5A1* | 1.080737 | 0.286263 | 3.775322 | 0.0001598 | 0.006760445 |
| *GPR160* | 0.781225 | 0.207596 | 3.7632 | 0.00016775 | 0.00707115 |
| *GRWD1* | 0.431138 | 0.114678 | 3.759549 | 0.00017022 | 0.007127817 |
| *CTSZ* | 0.694976 | 0.184864 | 3.759398 | 0.00017032 | 0.007127817 |

| | | | | | |
|---|---|---|---|---|---|
| *ZNF318* | -0.587752 | 0.156776 | -3.749003 | 0.00017754 | 0.007403194 |
| *RUNX2* | 1.892945 | 0.505429 | 3.745221 | 0.00018024 | 0.00748877 |
| *DMD* | -0.885594 | 0.236829 | -3.739386 | 0.00018447 | 0.007637456 |
| *CD6* | -1.357675 | 0.363418 | -3.735851 | 0.00018708 | 0.007718098 |
| *PKIG* | -0.503393 | 0.135018 | -3.728331 | 0.00019275 | 0.007923935 |
| *PRMT1* | 0.325425 | 0.087629 | 3.713692 | 0.00020426 | 0.008367357 |
| *DPEP3* | 3.160486 | 0.851123 | 3.713313 | 0.00020456 | NA |
| *IRAK2* | 1.055162 | 0.284227 | 3.712387 | 0.00020531 | 0.008381131 |
| *SLA* | 0.759604 | 0.204725 | 3.710357 | 0.00020697 | 0.008419062 |
| *ANKRD12* | -0.273394 | 0.073735 | -3.707793 | 0.00020907 | 0.008475129 |
| *PSAT1* | 0.834133 | 0.225622 | 3.697036 | 0.00021813 | 0.008811596 |
| *RAN* | 0.299508 | 0.081123 | 3.692033 | 0.00022247 | 0.008955696 |
| *HIVEP3* | 0.776831 | 0.210819 | 3.684828 | 0.00022886 | 0.009181106 |
| *CCM2* | -0.272539 | 0.074009 | -3.682529 | 0.00023093 | 0.009232517 |
| *SDF2L1* | 0.451472 | 0.122765 | 3.677534 | 0.0002355 | 0.009382888 |
| *DGAT2* | 0.660434 | 0.179712 | 3.674964 | 0.00023788 | 0.009445479 |
| *CCNH* | 0.484658 | 0.132164 | 3.667096 | 0.00024532 | 0.009707692 |
| *PSMB10* | 0.33045 | 0.090277 | 3.6604 | 0.00025182 | 0.009931169 |
| *TOP1* | 0.258836 | 0.070789 | 3.65643 | 0.00025575 | 0.010052089 |
| *ADRB2* | -1.673351 | 0.4579 | -3.654402 | 0.00025778 | 0.010097778 |
| *CCT5* | 0.282183 | 0.077355 | 3.647889 | 0.0002644 | 0.010322406 |
| *NXT1* | 0.396302 | 0.109009 | 3.63549 | 0.00027745 | 0.010795621 |
| *KIAA1109* | -0.550207 | 0.151642 | -3.628326 | 0.00028527 | 0.011062593 |
| *ACADVL* | -0.277586 | 0.07661 | -3.623369 | 0.00029079 | 0.011194402 |
| *NOP10* | 0.212292 | 0.058592 | 3.623197 | 0.00029099 | 0.011194402 |
| *TIMM17A* | 0.414431 | 0.114398 | 3.622694 | 0.00029155 | 0.011194402 |
| *DNAJC4* | -0.289542 | 0.080055 | -3.616788 | 0.00029828 | 0.011380195 |
| *HSBP1* | 0.255613 | 0.070675 | 3.616732 | 0.00029835 | 0.011380195 |
| *ENSG00000235609* | 1.963795 | 0.54366 | 3.612171 | 0.00030364 | 0.011544442 |
| *ZNF302* | -0.328726 | 0.091151 | -3.606404 | 0.00031047 | 0.011765492 |
| *PTPRE* | -0.686644 | 0.190452 | -3.605335 | 0.00031175 | 0.011775687 |
| *ENSG00000271857* | 2.024885 | 0.56234 | 3.600817 | 0.00031722 | 0.011943453 |
| *TOP2B* | 0.367658 | 0.10215 | 3.599181 | 0.00031922 | 0.011980065 |
| *SAMD9L* | -0.5931 | 0.165097 | -3.592437 | 0.0003276 | 0.012254967 |
| *DNAJB2* | -0.46749 | 0.130379 | -3.585611 | 0.00033629 | 0.012531642 |
| *EIF4A1* | 0.263482 | 0.073497 | 3.584944 | 0.00033715 | 0.012531642 |
| *COA7* | 0.50387 | 0.140599 | 3.583731 | 0.00033872 | 0.012549955 |
| *NAPSA* | 2.028539 | 0.567991 | 3.571429 | 0.00035504 | 0.013112772 |
| *LTA4H* | -0.396388 | 0.111046 | -3.569595 | 0.00035753 | 0.01311367 |
| *QARS* | -0.248389 | 0.069589 | -3.569382 | 0.00035782 | 0.01311367 |
| *PLK3* | 0.714842 | 0.200296 | 3.568928 | 0.00035845 | 0.01311367 |
| *TIMP1* | -0.885954 | 0.248719 | -3.562072 | 0.00036794 | 0.013418831 |
| *TCF3* | -0.350119 | 0.098523 | -3.553686 | 0.00037987 | 0.013810738 |
| *LDLRAD4* | 0.485942 | 0.136954 | 3.548215 | 0.00038785 | 0.014051622 |
| *LZIC* | 0.497091 | 0.140125 | 3.547495 | 0.00038891 | 0.014051622 |
| *RPF1* | 0.360472 | 0.101652 | 3.546119 | 0.00039095 | 0.014081456 |
| *DAP* | 0.551976 | 0.155792 | 3.543017 | 0.00039558 | 0.01419901 |

| | | | | | |
|---|---|---|---|---|---|
| *SNX18* | 0.760686 | 0.214743 | 3.5423 | 0.00039665 | 0.01419901 |
| *BCAR3* | 0.832837 | 0.235428 | 3.537548 | 0.00040386 | 0.014412605 |
| *MIF* | 0.353164 | 0.099895 | 3.535342 | 0.00040725 | 0.014489037 |
| *RPL10* | -0.199873 | 0.056597 | -3.531501 | 0.00041321 | 0.014656294 |
| *PPCDC* | 0.82614 | 0.234017 | 3.530254 | 0.00041516 | 0.014657224 |
| *MTSS1* | -0.574013 | 0.162616 | -3.529876 | 0.00041575 | 0.014657224 |
| *PRPF19* | 0.471438 | 0.133666 | 3.52698 | 0.00042033 | 0.01477372 |
| *EXOSC4* | 0.303182 | 0.086005 | 3.525155 | 0.00042324 | 0.014807633 |
| *ZFP36* | -0.267399 | 0.075865 | -3.524665 | 0.00042402 | 0.014807633 |
| *SMARCC1* | 0.317889 | 0.090222 | 3.523403 | 0.00042604 | 0.014807633 |
| *DIMT1* | 0.377055 | 0.107021 | 3.523191 | 0.00042638 | 0.014807633 |
| *SFXN4* | 0.48824 | 0.138773 | 3.518268 | 0.00043437 | 0.014990722 |
| *ZC3H15* | 0.218697 | 0.062163 | 3.518115 | 0.00043462 | 0.014990722 |
| *CKS2* | 0.397415 | 0.11298 | 3.517567 | 0.00043552 | 0.014990722 |
| *SNU13* | 0.216339 | 0.061545 | 3.515161 | 0.00043949 | 0.015082587 |
| *IFRD2* | 0.321526 | 0.091583 | 3.510751 | 0.00044684 | 0.015289879 |
| *APRT* | 0.192041 | 0.054763 | 3.506732 | 0.00045365 | 0.015444995 |
| *OAS3* | 1.130069 | 0.322342 | 3.505808 | 0.00045522 | 0.015444995 |
| *TENT5C* | -0.710566 | 0.202687 | -3.505729 | 0.00045536 | 0.015444995 |
| *EEF1E1* | 0.325317 | 0.093002 | 3.497956 | 0.00046884 | 0.015855982 |
| *HADHA* | -0.269795 | 0.077276 | -3.491333 | 0.00048062 | 0.016207236 |
| *GLRX3* | 0.251373 | 0.072042 | 3.489271 | 0.00048434 | 0.016228586 |
| *FCHO1* | 0.662066 | 0.189789 | 3.488437 | 0.00048585 | 0.016228586 |
| *CHCHD10* | 1.630956 | 0.467577 | 3.488103 | 0.00048646 | 0.016228586 |
| *EEF1A1* | -0.252822 | 0.072486 | -3.487853 | 0.00048692 | 0.016228586 |
| *CARD11* | 0.534835 | 0.153391 | 3.486742 | 0.00048894 | 0.016228586 |
| *GLYCTK* | 0.719994 | 0.206522 | 3.486282 | 0.00048978 | 0.016228586 |
| *ENSG00000259436* | -0.958862 | 0.275092 | -3.485611 | 0.00049101 | 0.016228586 |
| *EIF3J* | 0.248973 | 0.071498 | 3.482255 | 0.00049721 | 0.016386812 |
| *JUNB* | 0.398788 | 0.114692 | 3.477021 | 0.00050702 | 0.016662852 |
| *NIFK* | 0.25086 | 0.072195 | 3.474742 | 0.00051134 | 0.016725596 |
| *FXYD5* | -0.46507 | 0.133865 | -3.474171 | 0.00051244 | 0.016725596 |
| *IMP4* | 0.251901 | 0.072533 | 3.472933 | 0.0005148 | 0.016725596 |
| *NDUFS6* | 0.255198 | 0.073509 | 3.47164 | 0.00051729 | 0.016725596 |
| *TWISTNB* | 0.429879 | 0.12383 | 3.471518 | 0.00051752 | 0.016725596 |
| *FAM207A* | 0.367084 | 0.105742 | 3.471503 | 0.00051755 | 0.016725596 |
| *SH3TC1* | -0.620585 | 0.178892 | -3.469055 | 0.00052229 | 0.016799905 |
| *CKB* | -0.923647 | 0.266271 | -3.468825 | 0.00052274 | 0.016799905 |
| *EEF2KMT* | 0.410885 | 0.118772 | 3.459426 | 0.00054133 | 0.017339623 |
| *PWWP3A* | -0.392919 | 0.113599 | -3.458836 | 0.00054252 | 0.017339623 |
| *DAPP1* | -0.414815 | 0.12024 | -3.449908 | 0.00056078 | 0.017874193 |
| *NDUFAF4* | 0.366572 | 0.106307 | 3.448244 | 0.00056425 | 0.017935581 |
| *HSPD1* | 0.322189 | 0.09358 | 3.442921 | 0.00057547 | 0.018211773 |
| *MORC3* | -0.468611 | 0.136135 | -3.442249 | 0.0005769 | 0.018211773 |
| *POLR3GL* | -0.266113 | 0.077315 | -3.441907 | 0.00057763 | 0.018211773 |
| *TIAM1* | 1.826434 | 0.530902 | 3.440248 | 0.00058118 | 0.018236769 |
| *SNRPE* | 0.217091 | 0.063107 | 3.440073 | 0.00058156 | 0.018236769 |

| | | | | | |
|---|---|---|---|---|---|
| *C16orf72* | 0.389305 | 0.113237 | 3.437953 | 0.00058613 | 0.018330702 |
| *MFSD10* | -0.311295 | 0.090588 | -3.436379 | 0.00058955 | 0.018388105 |
| *CPNE3* | -0.342092 | 0.099688 | -3.431631 | 0.00059996 | 0.018654173 |
| *TGM2* | 1.890574 | 0.551021 | 3.431035 | 0.00060128 | 0.018654173 |
| *SLC38A5* | 0.572272 | 0.166931 | 3.428196 | 0.00060761 | 0.018800287 |
| *PAICS* | 0.329905 | 0.096454 | 3.420324 | 0.00062547 | 0.01930154 |
| *TMEM147* | 0.233822 | 0.068429 | 3.416991 | 0.00063317 | 0.019487661 |
| *TOMM5* | 0.35156 | 0.103006 | 3.412991 | 0.00064254 | 0.019723821 |
| *HSP90B1* | 0.327215 | 0.095912 | 3.411628 | 0.00064576 | 0.019753599 |
| *CALR* | 0.277494 | 0.081349 | 3.411145 | 0.00064691 | 0.019753599 |
| *PARK7* | 0.203757 | 0.059797 | 3.407487 | 0.00065564 | 0.019922647 |
| *SC5D* | -0.418216 | 0.12274 | -3.407323 | 0.00065604 | 0.019922647 |
| *C8orf58* | -0.795232 | 0.233463 | -3.406237 | 0.00065865 | 0.019922647 |
| *LARGE2* | -0.652787 | 0.191688 | -3.405467 | 0.00066051 | 0.019922647 |
| *PDHA1* | 0.374057 | 0.109847 | 3.405262 | 0.00066101 | 0.019922647 |
| *ATXN10* | 0.34376 | 0.101083 | 3.400781 | 0.00067194 | 0.020197238 |
| *CD99* | -0.520428 | 0.153072 | -3.399898 | 0.00067411 | 0.020197238 |
| *GDPGP1* | 0.886009 | 0.260637 | 3.399406 | 0.00067533 | 0.020197238 |
| *EIF4ENIF1* | 0.683243 | 0.201096 | 3.397589 | 0.00067983 | 0.020279719 |
| *MYO1C* | -0.423635 | 0.124715 | -3.396823 | 0.00068173 | 0.020284543 |
| *PTPN7* | 0.666959 | 0.196769 | 3.389561 | 0.00070005 | 0.02075045 |
| *MRPS12* | 0.261362 | 0.077132 | 3.388516 | 0.00070272 | 0.02075045 |
| *VAMP2* | -0.301367 | 0.08896 | -3.387687 | 0.00070485 | 0.02075045 |
| *KMT2E* | -0.265297 | 0.078315 | -3.38754 | 0.00070523 | 0.02075045 |
| *COA4* | 0.254222 | 0.075056 | 3.387119 | 0.00070631 | 0.02075045 |
| *TBC1D8* | 1.225927 | 0.362352 | 3.38325 | 0.00071633 | 0.020992029 |
| *NFKBIZ* | 0.434827 | 0.128638 | 3.380235 | 0.00072424 | 0.021170322 |
| *COX5A* | 0.225737 | 0.066811 | 3.37873 | 0.00072821 | 0.021233177 |
| *RSRP1* | -0.336694 | 0.099826 | -3.372793 | 0.0007441 | 0.021642088 |
| *FCER2* | 1.405165 | 0.416727 | 3.371907 | 0.0007465 | 0.021657745 |
| *FAM129C* | -0.552377 | 0.164054 | -3.367039 | 0.0007598 | 0.021945937 |
| *PFDN6* | 0.297386 | 0.088327 | 3.366892 | 0.0007602 | 0.021945937 |
| *PSMD8* | 0.239311 | 0.071122 | 3.364783 | 0.00076604 | 0.022041867 |
| *CCNG1* | -0.334817 | 0.09952 | -3.364323 | 0.00076732 | 0.022041867 |
| *ZNF44* | 0.460303 | 0.136943 | 3.361273 | 0.00077584 | 0.022231879 |
| *GLRX2* | 0.379486 | 0.112936 | 3.360176 | 0.00077893 | 0.022265463 |
| *SMIM20* | 0.463118 | 0.137901 | 3.358342 | 0.00078412 | 0.022358814 |
| *DHRS3* | 1.360827 | 0.405302 | 3.357567 | 0.00078632 | 0.022366745 |
| *SPCS2* | 0.206078 | 0.061452 | 3.353456 | 0.00079809 | 0.022619594 |
| *HIVEP2* | 0.785279 | 0.234194 | 3.353109 | 0.00079909 | 0.022619594 |
| *TIMMDC1* | 0.369367 | 0.110271 | 3.349626 | 0.00080921 | 0.02285025 |
| *RPL13A* | -0.176335 | 0.052673 | -3.347726 | 0.00081478 | 0.022951843 |
| *TRANK1* | -0.557649 | 0.166703 | -3.34517 | 0.00082232 | 0.023108401 |
| *LMF2* | -0.392408 | 0.117516 | -3.339183 | 0.00084025 | 0.023555377 |
| *ULK1* | 0.812984 | 0.243637 | 3.336865 | 0.00084729 | 0.023695626 |
| *IQGAP1* | -0.330345 | 0.099026 | -3.33593 | 0.00085015 | 0.023718487 |
| *EIF3H* | -0.201209 | 0.060332 | -3.335056 | 0.00085282 | 0.02373623 |

| | | | | | |
|---|---|---|---|---|---|
| *CERS4* | -0.373841 | 0.112242 | -3.330686 | 0.00086632 | 0.024038068 |
| *SERBP1* | 0.243561 | 0.073137 | 3.330212 | 0.0008678 | 0.024038068 |
| *CD320* | 0.355459 | 0.106796 | 3.3284 | 0.00087346 | 0.024137459 |
| *FAM30A* | -0.355616 | 0.106866 | -3.32769 | 0.00087569 | 0.024141697 |
| *ZNRD2* | 0.299934 | 0.090192 | 3.325508 | 0.00088258 | 0.024265751 |
| *BTG2* | -0.305311 | 0.091824 | -3.324944 | 0.00088436 | 0.024265751 |
| *TESK1* | 0.528849 | 0.159154 | 3.322876 | 0.00089095 | 0.024388827 |
| *SPHK1* | 0.825919 | 0.248616 | 3.322073 | 0.00089351 | 0.024401709 |
| *PRMT2* | -0.414689 | 0.12493 | -3.319365 | 0.00090222 | 0.024581883 |
| *ENSG00000259865* | -0.489144 | 0.147457 | -3.317196 | 0.00090926 | 0.024715697 |
| *SQSTM1* | 0.3016 | 0.090989 | 3.31469 | 0.00091745 | 0.024857175 |
| *CAPG* | -0.654324 | 0.197435 | -3.314121 | 0.00091932 | 0.024857175 |
| *KCNN1* | 1.59488 | 0.481379 | 3.313146 | 0.00092253 | 0.024857175 |
| *TBL3* | 0.390184 | 0.117782 | 3.312756 | 0.00092382 | 0.024857175 |
| *FKBP11* | 0.354556 | 0.107041 | 3.312353 | 0.00092515 | 0.024857175 |
| *EHD1* | 0.274836 | 0.083015 | 3.310678 | 0.0009307 | 0.024948785 |
| *UGP2* | -0.30751 | 0.093178 | -3.300232 | 0.00096605 | 0.025836817 |
| *MAP4K2* | -0.423762 | 0.128634 | -3.294333 | 0.00098655 | 0.026324695 |
| *ABCF2* | 0.33976 | 0.10333 | 3.288112 | 0.00100862 | 0.026851839 |
| *TBC1D10C* | -0.307324 | 0.093605 | -3.28319 | 0.0010264 | 0.027180749 |
| *IDS* | -0.274142 | 0.083505 | -3.282923 | 0.00102737 | 0.027180749 |
| *SLIRP* | 0.302241 | 0.092069 | 3.282755 | 0.00102798 | 0.027180749 |
| *GRPEL1* | 0.329147 | 0.100404 | 3.278214 | 0.00104466 | 0.027559141 |
| *DCTPP1* | 0.323785 | 0.098905 | 3.273683 | 0.00106156 | 0.027832687 |
| *NTHL1* | 0.426747 | 0.130366 | 3.273445 | 0.00106245 | 0.027832687 |
| *CCDC90B* | -0.254322 | 0.077696 | -3.273309 | 0.00106296 | 0.027832687 |
| *RPS27* | -0.171496 | 0.052399 | -3.272874 | 0.0010646 | 0.027832687 |
| *EIF3B* | 0.327927 | 0.100428 | 3.2653 | 0.00109348 | 0.028444197 |
| *TARS* | 0.308748 | 0.094566 | 3.26489 | 0.00109507 | 0.028444197 |
| *SLCO4A1* | 0.531094 | 0.162672 | 3.264823 | 0.00109532 | 0.028444197 |
| *TRIM25* | 0.74044 | 0.227036 | 3.261327 | 0.00110892 | 0.028718964 |
| *ZNF581* | -0.314374 | 0.096409 | -3.260836 | 0.00111084 | 0.028718964 |
| *BCL6* | -0.775876 | 0.238215 | -3.257044 | 0.00112579 | 0.029040882 |
| *FAM3C* | 0.345163 | 0.106013 | 3.25585 | 0.00113054 | 0.029098793 |
| *SRSF8* | 0.259103 | 0.079617 | 3.254374 | 0.00113643 | 0.029183217 |
| *RRS1* | 0.434724 | 0.133606 | 3.253773 | 0.00113883 | 0.029183217 |
| *CNPY2* | 0.206808 | 0.06361 | 3.251202 | 0.00114918 | 0.029383677 |
| *EEF1D* | -0.238507 | 0.073649 | -3.238447 | 0.00120182 | 0.030662308 |
| *POLR2H* | 0.301919 | 0.0933 | 3.235985 | 0.00121224 | 0.030860379 |
| *MRPL13* | 0.384859 | 0.119152 | 3.229988 | 0.00123796 | 0.031446224 |
| *ZNF524* | -0.353467 | 0.109646 | -3.223698 | 0.00126547 | 0.032075105 |
| *MMP11* | 1.022476 | 0.3173 | 3.222429 | 0.00127109 | 0.03214739 |
| *HDLBP* | 0.299192 | 0.092983 | 3.21771 | 0.00129219 | 0.03261018 |
| *SVBP* | -0.250583 | 0.077951 | -3.214617 | 0.00130619 | 0.032892163 |
| *PRR34.AS1* | -0.362624 | 0.112833 | -3.213805 | 0.00130989 | 0.032914106 |
| *PSMD11* | 0.268683 | 0.083747 | 3.208263 | 0.00133539 | 0.033482661 |
| *BTG3* | 0.445246 | 0.138893 | 3.205674 | 0.00134747 | 0.033712728 |

| | | | | | |
|---|---|---|---|---|---|
| *DPH2* | 0.346664 | 0.108204 | 3.2038 | 0.00135627 | 0.033860115 |
| *FBXL17* | -0.676075 | 0.211525 | -3.196186 | 0.00139258 | 0.034674634 |
| *LINC00996* | 1.35579 | 0.424252 | 3.195714 | 0.00139485 | 0.034674634 |
| *RFK* | 0.407971 | 0.127835 | 3.191399 | 0.00141585 | 0.035121637 |
| *GPR68* | -2.694012 | 0.846761 | -3.181551 | 0.00146489 | NA |
| *TUBB2A* | 0.898078 | 0.282362 | 3.180592 | 0.00146975 | 0.036377624 |
| *GAK* | 0.37848 | 0.119019 | 3.180002 | 0.00147274 | 0.036377624 |
| *SPINT2* | -0.327543 | 0.10305 | -3.178476 | 0.00148051 | 0.036492143 |
| *NDUFAF8* | 0.240859 | 0.075795 | 3.177779 | 0.00148408 | 0.036502694 |
| *CISD1* | 0.309074 | 0.097322 | 3.175795 | 0.00149427 | 0.036675702 |
| *RECQL* | -0.369277 | 0.116305 | -3.175085 | 0.00149793 | 0.036688148 |
| *CCDC86* | 0.415738 | 0.131166 | 3.169563 | 0.00152669 | 0.037313978 |
| *ETFB* | 0.333363 | 0.105227 | 3.168032 | 0.00153475 | 0.037432388 |
| *RHOG* | -0.252224 | 0.079661 | -3.166208 | 0.0015444 | 0.037560464 |
| *HDAC9* | 0.284674 | 0.089921 | 3.165822 | 0.00154646 | 0.037560464 |
| *ALG3* | 0.333038 | 0.10523 | 3.164866 | 0.00155154 | 0.03760555 |
| *AP1G2* | -0.305062 | 0.096533 | -3.160177 | 0.00157673 | 0.038084655 |
| *EEF2* | -0.216315 | 0.068455 | -3.159969 | 0.00157786 | 0.038084655 |
| *CCR7* | 0.310339 | 0.098243 | 3.158878 | 0.00158378 | 0.038148414 |
| *FER* | -0.697253 | 0.220926 | -3.156053 | 0.0015992 | 0.038440289 |
| *CCNT2* | -0.436783 | 0.138506 | -3.153521 | 0.00161314 | 0.038695305 |
| *WASF2* | 0.226783 | 0.07197 | 3.151052 | 0.00162684 | 0.038927928 |
| *NOP14* | 0.352428 | 0.111862 | 3.150569 | 0.00162953 | 0.038927928 |
| *MFNG* | 0.330556 | 0.104992 | 3.148391 | 0.00164172 | 0.039017206 |
| *DUSP18* | -0.602719 | 0.191468 | -3.14789 | 0.00164454 | 0.039017206 |
| *PFDN2* | 0.259425 | 0.082416 | 3.147737 | 0.0016454 | 0.039017206 |
| *VIM.AS1* | -0.630162 | 0.200237 | -3.147086 | 0.00164906 | 0.039017206 |
| *NTAN1* | -0.299439 | 0.095153 | -3.146915 | 0.00165003 | 0.039017206 |
| *TMED9* | 0.245654 | 0.078083 | 3.146049 | 0.00165492 | 0.039053435 |
| *H2AFY* | 0.244047 | 0.077714 | 3.140333 | 0.00168756 | 0.03974301 |
| *ZFP90* | -0.586822 | 0.186989 | -3.138269 | 0.00169949 | 0.039943165 |
| *RSL24D1* | 0.215323 | 0.068675 | 3.135374 | 0.00171635 | 0.040258138 |
| *ITGB7* | 0.688342 | 0.219643 | 3.13392 | 0.00172488 | 0.040376729 |
| *DDIT4* | 0.699295 | 0.223387 | 3.130418 | 0.00174558 | 0.040779267 |
| *DCTN2* | -0.283052 | 0.090451 | -3.12933 | 0.00175205 | 0.040848459 |
| *ANKRD36B* | -0.644075 | 0.205914 | -3.12788 | 0.00176072 | 0.040968499 |
| *FXYD1* | -0.894094 | 0.286114 | -3.124959 | 0.0017783 | 0.041292708 |
| *EIF3F* | -0.193173 | 0.061827 | -3.124388 | 0.00178176 | 0.041292708 |
| *CLIP4* | 1.232283 | 0.394826 | 3.121075 | 0.00180192 | 0.041676984 |
| *ENSG00000275418* | -2.655102 | 0.851238 | -3.119106 | 0.001814 | NA |
| *RUBCN* | -0.449059 | 0.143986 | -3.118762 | 0.00181613 | 0.041912786 |
| *EIF4A3* | 0.523364 | 0.167839 | 3.118244 | 0.00181932 | 0.041912786 |
| *RPS28* | -0.101494 | 0.032608 | -3.11259 | 0.00185454 | 0.042639679 |
| *MARCKSL1* | -0.441606 | 0.141989 | -3.110133 | 0.00187003 | 0.042911098 |
| *GALM* | 0.761698 | 0.245106 | 3.107634 | 0.00188591 | 0.043104877 |
| *ZNF638* | -0.321608 | 0.1035 | -3.107324 | 0.0018879 | 0.043104877 |
| *ENSG00000111540* | -0.315168 | 0.101436 | -3.107059 | 0.00188959 | 0.043104877 |

| | | | | | |
|---|---|---|---|---|---|
| *CARD8* | -0.348885 | 0.112383 | -3.104423 | 0.00190651 | 0.04340567 |
| *TMEM134* | -0.213555 | 0.068819 | -3.103136 | 0.00191482 | 0.043509746 |
| *ACACA* | 0.56745 | 0.183303 | 3.0957 | 0.00196349 | 0.044457775 |
| *POMP* | 0.22286 | 0.072003 | 3.095163 | 0.00196705 | 0.044457775 |
| *USF3* | -0.494874 | 0.159894 | -3.095019 | 0.001968 | 0.044457775 |
| *TIMM23* | 0.373916 | 0.120858 | 3.093844 | 0.00197582 | 0.044547761 |
| *FCMR* | 0.468744 | 0.151604 | 3.091889 | 0.00198887 | 0.044755406 |
| *SLC39A14* | 0.635805 | 0.205915 | 3.087702 | 0.00201711 | 0.045228313 |
| *EEF1AKNMT* | 0.502654 | 0.162797 | 3.08762 | 0.00201766 | 0.045228313 |
| *PSMD7* | 0.261507 | 0.084775 | 3.084701 | 0.00203757 | 0.045501181 |
| *STIP1* | 0.401724 | 0.130239 | 3.08451 | 0.00203887 | 0.045501181 |
| *ENSG00000089127* | 0.847796 | 0.274902 | 3.083999 | 0.00204239 | 0.045501181 |
| *PBXIP1* | -0.588174 | 0.190746 | -3.083548 | 0.00204548 | 0.045501181 |
| *TUFM* | 0.207071 | 0.067243 | 3.079464 | 0.00207374 | 0.046041709 |
| *CTSB* | 0.333205 | 0.10828 | 3.077254 | 0.00208917 | 0.046296091 |
| *RUVBL1* | 0.375638 | 0.122111 | 3.076196 | 0.0020966 | 0.04637234 |
| *ZNF652* | -0.445459 | 0.145166 | -3.068616 | 0.00215053 | 0.047408132 |
| *PINX1* | 0.397489 | 0.12954 | 3.06847 | 0.00215158 | 0.047408132 |
| *BICDL1* | 0.896475 | 0.29224 | 3.067605 | 0.00215782 | 0.047455721 |
| *NCKAP1L* | -0.269179 | 0.087826 | -3.064894 | 0.00217747 | 0.04767713 |
| *RPL41* | -0.156314 | 0.051002 | -3.064847 | 0.00217782 | 0.04767713 |
| *SEPTIN6* | -0.19496 | 0.063618 | -3.064522 | 0.00218018 | 0.04767713 |
| *RNPS1* | 0.18821 | 0.061459 | 3.062351 | 0.00219606 | 0.047934156 |
| *ARL17B* | -0.631081 | 0.206172 | -3.06095 | 0.00220636 | 0.048068903 |
| *CYBC1* | -0.250991 | 0.082038 | -3.059446 | 0.00221746 | 0.04818389 |
| *SESTD1* | -0.700181 | 0.228883 | -3.059115 | 0.00221992 | 0.04818389 |
| *SFXN3* | -0.838324 | 0.274707 | -3.051707 | 0.00227544 | 0.049297017 |
| *NNT* | -0.46397 | 0.152304 | -3.046339 | 0.00231647 | 0.050058195 |
| *ADSL* | 0.251158 | 0.082455 | 3.045987 | 0.00231918 | 0.050058195 |
| *FOXO3* | 0.491482 | 0.161425 | 3.044648 | 0.00232953 | 0.05018845 |
| *TTC3* | -0.227358 | 0.074713 | -3.043087 | 0.00234165 | 0.05035625 |
| *DNAH1* | 1.165388 | 0.383159 | 3.041525 | 0.00235383 | 0.05052478 |
| *CRTC3* | -0.349659 | 0.115102 | -3.037806 | 0.00238308 | 0.051058393 |
| *AHRR* | -1.175932 | 0.387386 | -3.035556 | 0.00240093 | 0.051287797 |
| *MYADM* | 0.640079 | 0.210895 | 3.035065 | 0.00240484 | 0.051287797 |
| *RPS27A* | -0.141766 | 0.046713 | -3.034793 | 0.00240701 | 0.051287797 |
| *SERPINE2* | 0.886072 | 0.292109 | 3.033363 | 0.00241844 | 0.051437245 |
| *TEX264* | 0.332193 | 0.109576 | 3.03162 | 0.00243246 | 0.051618633 |
| *TSFM* | 0.417804 | 0.137835 | 3.031199 | 0.00243585 | 0.051618633 |
| *HERC4* | -0.34896 | 0.115212 | -3.028843 | 0.00245492 | 0.051928246 |
| *SLC25A6* | -0.19448 | 0.06432 | -3.023627 | 0.00249764 | 0.052736015 |
| *RPL27A* | -0.130517 | 0.043184 | -3.022333 | 0.00250834 | 0.052830796 |
| *PPIL1* | 0.371309 | 0.122869 | 3.021987 | 0.00251121 | 0.052830796 |
| *SERPINB9P1* | -0.886554 | 0.293587 | -3.019733 | 0.00252998 | 0.05312948 |
| *CLIC3* | 1.82448 | 0.604696 | 3.017186 | 0.00255133 | 0.053481445 |
| *ENSG00000237188* | -0.936266 | 0.310565 | -3.014721 | 0.00257216 | 0.053821071 |
| *TIMM13* | 0.265785 | 0.088237 | 3.012158 | 0.00259397 | 0.054180046 |

| | | | | | |
|---|---|---|---|---|---|
| *TMBIM1* | -0.382994 | 0.127208 | -3.010773 | 0.00260583 | 0.054330242 |
| *SDE2* | 0.550964 | 0.183044 | 3.010008 | 0.00261241 | 0.05433937 |
| *TBC1D9* | 0.776526 | 0.258023 | 3.009525 | 0.00261656 | 0.05433937 |
| *CYCS* | 0.320198 | 0.10641 | 3.009094 | 0.00262028 | 0.05433937 |
| *CD47* | -0.30747 | 0.102208 | -3.008285 | 0.00262726 | 0.054387179 |
| *SINHCAF* | 0.241392 | 0.080318 | 3.005462 | 0.00265178 | 0.054797114 |
| *RABEPK* | 0.373907 | 0.12449 | 3.003517 | 0.00266878 | 0.055050751 |
| *PIKFYVE* | 0.425275 | 0.141703 | 3.00117 | 0.00268945 | 0.055294812 |
| *LBR* | 0.284303 | 0.094733 | 3.001093 | 0.00269012 | 0.055294812 |
| *CASP4* | -0.249975 | 0.083356 | -2.998897 | 0.00270959 | 0.055596812 |
| *SSBP2* | -0.675545 | 0.22559 | -2.99457 | 0.00274832 | 0.056276126 |
| *AP2B1* | -0.240485 | 0.080319 | -2.99412 | 0.00275237 | 0.056276126 |
| *BCCIP* | 0.286517 | 0.095747 | 2.992442 | 0.00276756 | 0.056452752 |
| *EIF5B* | 0.18811 | 0.062869 | 2.992093 | 0.00277072 | 0.056452752 |
| *EBI3* | 0.571534 | 0.191062 | 2.991356 | 0.00277742 | 0.056470812 |
| *RPS6KA2* | -0.888935 | 0.297216 | -2.990873 | 0.00278181 | 0.056470812 |
| *PIP5K1B* | 0.587975 | 0.196653 | 2.989914 | 0.00279056 | 0.056470812 |
| *LYRM9* | -0.63879 | 0.213652 | -2.989864 | 0.00279102 | 0.056470812 |
| *PNO1* | 0.397041 | 0.132888 | 2.987776 | 0.00281015 | 0.056759189 |
| *MICAL1* | -0.391573 | 0.131167 | -2.985315 | 0.00283287 | 0.057118881 |
| *LGMN* | 0.780053 | 0.261415 | 2.98396 | 0.00284544 | 0.057273057 |
| *PQLC3* | -0.529184 | 0.177409 | -2.982854 | 0.00285574 | 0.057381153 |
| *ANKRD36C* | -0.454421 | 0.152731 | -2.975295 | 0.00292707 | 0.058682841 |
| *DDX49* | 0.301944 | 0.101507 | 2.97462 | 0.00293351 | 0.058682841 |
| *ANAPC16* | 0.188552 | 0.063392 | 2.974396 | 0.00293566 | 0.058682841 |
| *HSP90AB1* | 0.154059 | 0.051813 | 2.973351 | 0.00294567 | 0.058724003 |
| *GPR155* | -0.987262 | 0.332062 | -2.973129 | 0.00294781 | 0.058724003 |
| *MRPS36* | 0.186292 | 0.062681 | 2.972051 | 0.00295817 | 0.058829693 |
| *APOBEC3C* | -0.328183 | 0.110525 | -2.969302 | 0.00298477 | 0.059257388 |
| *PAOX* | 0.581625 | 0.196169 | 2.964915 | 0.00302766 | 0.060006491 |
| *SEL1L3* | 0.655895 | 0.221374 | 2.962837 | 0.00304818 | 0.060254075 |
| *RACK1* | -0.150947 | 0.050951 | -2.962602 | 0.00305051 | 0.060254075 |
| *ZNF461* | 0.796171 | 0.268972 | 2.960054 | 0.00307585 | 0.060651552 |
| *OXT* | -0.903701 | 0.3057 | -2.956168 | 0.00311488 | 0.061317208 |
| *NOP58* | 0.199023 | 0.067348 | 2.955132 | 0.00312535 | 0.061419423 |
| *MPP6* | 0.495412 | 0.167692 | 2.954291 | 0.00313389 | 0.061472595 |
| *ENSG00000277511* | -0.875625 | 0.296438 | -2.953825 | 0.00313862 | 0.061472595 |
| *HEXD* | -0.570147 | 0.193383 | -2.948277 | 0.00319551 | 0.062481538 |
| *PSMA5* | 0.282289 | 0.095815 | 2.946181 | 0.00321724 | 0.062593568 |
| *ESYT1* | -0.35008 | 0.118837 | -2.945892 | 0.00322025 | 0.062593568 |
| *C11orf80* | -0.670199 | 0.227518 | -2.945703 | 0.00322222 | 0.062593568 |
| *SNRNP40* | 0.326758 | 0.110929 | 2.945651 | 0.00322276 | 0.062593568 |
| *RHOA* | 0.199938 | 0.067984 | 2.940967 | 0.00327189 | 0.063442032 |
| *METAP2* | 0.192238 | 0.065438 | 2.937686 | 0.00330672 | 0.064010561 |
| *ITGA4* | 0.449624 | 0.153141 | 2.936002 | 0.00332472 | 0.064231376 |
| *FAM216A* | 0.397064 | 0.135282 | 2.935089 | 0.00333453 | 0.064231376 |
| *PSMD4* | 0.227422 | 0.077484 | 2.935073 | 0.00333469 | 0.064231376 |

| | | | | | |
|---|---|---|---|---|---|
| *EMC4* | 0.265016 | 0.090341 | 2.933489 | 0.00335175 | 0.064347416 |
| *RPL7L1* | 0.229554 | 0.078253 | 2.933487 | 0.00335177 | 0.064347416 |
| *SLC44A1* | -0.596076 | 0.203388 | -2.930738 | 0.00338158 | 0.06481274 |
| *PPP1CC* | 0.210057 | 0.071735 | 2.928236 | 0.00340891 | 0.065229103 |
| *C7orf50* | -0.43475 | 0.148728 | -2.92312 | 0.00346543 | 0.066201662 |
| *CTPS1* | 0.466837 | 0.159803 | 2.921335 | 0.00348535 | 0.066385012 |
| *SNRPB* | 0.285087 | 0.097591 | 2.921237 | 0.00348644 | 0.066385012 |
| *EXOC3* | -0.463387 | 0.159072 | -2.913072 | 0.00357892 | 0.068034489 |
| *TPI1* | 0.217127 | 0.074627 | 2.909499 | 0.00362009 | 0.06870487 |
| *POLR3D* | 0.435743 | 0.150065 | 2.903688 | 0.00368796 | 0.069879023 |
| *TTC7A* | -0.529696 | 0.182456 | -2.903148 | 0.00369432 | 0.069885659 |
| *PLD2* | -0.766767 | 0.264234 | -2.901843 | 0.00370975 | 0.070063657 |
| *MTFP1* | 0.316626 | 0.109139 | 2.901132 | 0.00371818 | 0.070079357 |
| *DCUN1D5* | 0.333033 | 0.114809 | 2.900757 | 0.00372263 | 0.070079357 |
| *ATAD3B* | 0.393147 | 0.135651 | 2.898216 | 0.00375292 | 0.070436211 |
| *HINT2* | 0.258016 | 0.089031 | 2.89803 | 0.00375515 | 0.070436211 |
| *PHB* | 0.26725 | 0.092239 | 2.897362 | 0.00376315 | 0.070436211 |
| *PLEC* | -0.772642 | 0.266707 | -2.89697 | 0.00376786 | 0.070436211 |
| *CD24* | -0.261193 | 0.090171 | -2.896637 | 0.00377186 | 0.070436211 |
| *TSPAN17* | -0.360556 | 0.124519 | -2.895596 | 0.00378439 | 0.0705171 |
| *HTR3A* | 1.361698 | 0.470318 | 2.895271 | 0.00378831 | 0.0705171 |
| *PLAC8* | -0.240701 | 0.083196 | -2.893189 | 0.00381352 | 0.070872991 |
| *GNL3* | 0.318284 | 0.110234 | 2.887344 | 0.00388509 | 0.072001508 |
| *SEPTIN1* | -0.234725 | 0.081298 | -2.88722 | 0.00388662 | 0.072001508 |
| *ACP1* | 0.172766 | 0.059903 | 2.884078 | 0.00392562 | 0.072332654 |
| *NDUFAF2* | 0.326543 | 0.113224 | 2.884042 | 0.00392606 | 0.072332654 |
| *CCDC28B* | -0.515868 | 0.17887 | -2.884031 | 0.0039262 | 0.072332654 |
| *GAR1* | 0.317531 | 0.110109 | 2.883777 | 0.00392937 | 0.072332654 |
| *EIF4B* | -0.182995 | 0.063476 | -2.882883 | 0.00394054 | 0.072342104 |
| *FARSA* | 0.320236 | 0.111087 | 2.882741 | 0.00394232 | 0.072342104 |
| *OGG1* | -0.36531 | 0.126818 | -2.880585 | 0.00396938 | 0.072723963 |
| *LYAR* | 0.375672 | 0.130513 | 2.878421 | 0.00399671 | 0.072987351 |
| *OFD1* | -0.360621 | 0.125291 | -2.878271 | 0.00399862 | 0.072987351 |
| *TTC24* | -1.767619 | 0.61416 | -2.878106 | 0.00400071 | NA |
| *HSPE1* | 0.29273 | 0.101716 | 2.877922 | 0.00400304 | 0.072987351 |
| *LMAN1* | 0.318291 | 0.110615 | 2.877465 | 0.00400885 | 0.072987351 |
| *FAM111A* | -0.540288 | 0.188076 | -2.872709 | 0.00406969 | 0.073979257 |
| *L1CAM* | -1.619025 | 0.563781 | -2.871726 | 0.00408237 | 0.074026521 |
| *MDH2* | 0.213921 | 0.074497 | 2.871521 | 0.00408501 | 0.074026521 |
| *RASSF7* | -0.375728 | 0.130873 | -2.870924 | 0.00409273 | 0.07405111 |
| *RPS6* | -0.137166 | 0.04782 | -2.868362 | 0.00412603 | 0.074496691 |
| *SLC23A2* | 0.585569 | 0.20417 | 2.868045 | 0.00413017 | 0.074496691 |
| *S100A6* | -0.720512 | 0.251323 | -2.866873 | 0.00414549 | 0.074657304 |
| *ETS1* | -0.331502 | 0.115662 | -2.86612 | 0.00415537 | 0.074719593 |
| *RPF2* | 0.319184 | 0.111498 | 2.86269 | 0.00420061 | 0.075416535 |
| *GPT2* | 0.623589 | 0.218099 | 2.859203 | 0.00424707 | 0.076067797 |
| *PAXX* | 0.30484 | 0.106625 | 2.858987 | 0.00424996 | 0.076067797 |

| | | | | | |
|---|---|---|---|---|---|
| *LINC02328* | -1.143507 | 0.400542 | -2.8549 | 0.00430504 | 0.076935293 |
| *SSR2* | -0.161808 | 0.056729 | -2.852311 | 0.00434026 | 0.077445633 |
| *SMARCB1* | 0.221536 | 0.077702 | 2.851121 | 0.00435654 | 0.077585614 |
| *MCTS1* | 0.257828 | 0.090442 | 2.850763 | 0.00436144 | 0.077585614 |
| *WIPF1* | -0.26879 | 0.094378 | -2.848007 | 0.0043994 | 0.078141437 |
| *IARS* | 0.288721 | 0.101406 | 2.847173 | 0.00441094 | 0.078145465 |
| *GPATCH2L* | -0.298139 | 0.10474 | -2.846473 | 0.00442065 | 0.078145465 |
| *CCNI* | -0.175976 | 0.061826 | -2.846297 | 0.00442309 | 0.078145465 |
| *SMIM29* | -0.263765 | 0.092681 | -2.845954 | 0.00442785 | 0.078145465 |
| *DMXL1* | -0.425936 | 0.149684 | -2.845569 | 0.00443321 | 0.078145465 |
| *PRMT5* | 0.35825 | 0.126046 | 2.842219 | 0.00448008 | 0.078852044 |
| *MTHFD2* | 0.381391 | 0.134275 | 2.840379 | 0.004506 | 0.079188439 |
| *PSME2* | 0.197033 | 0.069448 | 2.837143 | 0.00455192 | 0.079874934 |
| *EPM2AIP1* | -0.259684 | 0.091573 | -2.83583 | 0.00457067 | 0.079929667 |
| *NOP56* | 0.219584 | 0.077446 | 2.835332 | 0.0045778 | 0.079929667 |
| *RPL13* | -0.137972 | 0.048667 | -2.834992 | 0.00458269 | 0.079929667 |
| *OXR1* | -0.473867 | 0.167163 | -2.834753 | 0.00458611 | 0.079929667 |
| *DISP1* | -0.978753 | 0.345297 | -2.834524 | 0.0045894 | 0.079929667 |
| *TSR1* | 0.288598 | 0.101848 | 2.833618 | 0.00460243 | 0.08003686 |
| *ZPR1* | 0.259339 | 0.09156 | 2.832456 | 0.0046192 | 0.08020858 |
| *ALYREF* | 0.50732 | 0.179171 | 2.831493 | 0.00463313 | 0.080330575 |
| *TMEM191C* | 0.686698 | 0.242644 | 2.830065 | 0.00465386 | 0.080569929 |
| *ELOF1* | 0.283015 | 0.100053 | 2.828649 | 0.00467449 | 0.080705418 |
| *BCL2L11* | -0.430633 | 0.152267 | -2.828136 | 0.004682 | 0.080705418 |
| *RAB7A* | 0.205531 | 0.072675 | 2.828101 | 0.0046825 | 0.080705418 |
| *B4GALT1* | -0.414271 | 0.146531 | -2.827186 | 0.0046959 | 0.080816706 |
| *SSBP1* | 0.273582 | 0.096879 | 2.823956 | 0.00474349 | 0.081411517 |
| *SRPK1* | 0.256901 | 0.090974 | 2.823891 | 0.00474446 | 0.081411517 |
| *CAPRIN2* | -0.720208 | 0.255117 | -2.823054 | 0.00475685 | 0.081504029 |
| *PFKFB3* | -0.635359 | 0.225195 | -2.821369 | 0.00478192 | 0.081812981 |
| *NARS* | 0.240383 | 0.085262 | 2.819333 | 0.00481236 | 0.082212838 |
| *DNPEP* | 0.223718 | 0.079513 | 2.813587 | 0.00489921 | 0.083453078 |
| *PGAM1* | 0.252408 | 0.08971 | 2.813581 | 0.0048993 | 0.083453078 |
| *IL7* | -0.55387 | 0.196968 | -2.811975 | 0.00492383 | 0.083748376 |
| *PYCR1* | 0.416762 | 0.148347 | 2.809364 | 0.00496395 | 0.084265896 |
| *FARSB* | 0.353422 | 0.125818 | 2.808987 | 0.00496976 | 0.084265896 |
| *GLOD4* | 0.300706 | 0.107067 | 2.80858 | 0.00497605 | 0.084265896 |
| *AKAP1* | 0.50145 | 0.178572 | 2.808116 | 0.00498323 | 0.084265896 |
| *YIF1A* | 0.22598 | 0.080616 | 2.803167 | 0.00506035 | 0.085228275 |
| *FXYD7* | -0.928686 | 0.331359 | -2.802654 | 0.0050684 | 0.085228275 |
| *SPIB* | 0.196651 | 0.07017 | 2.802488 | 0.00507101 | 0.085228275 |
| *SPTBN1* | 0.301197 | 0.107481 | 2.802323 | 0.0050736 | 0.085228275 |
| *ILF2* | 0.233204 | 0.083224 | 2.802122 | 0.00507678 | 0.085228275 |
| *SNRPC* | 0.2329 | 0.083348 | 2.794301 | 0.00520121 | 0.08719147 |
| *C1D* | 0.232217 | 0.083118 | 2.793835 | 0.00520871 | 0.087191562 |
| *ENSG00000244459* | -0.78642 | 0.281715 | -2.791549 | 0.00524564 | 0.087622118 |
| *ENSG00000259038* | -1.40396 | 0.502986 | -2.791249 | 0.00525051 | 0.087622118 |

| | | | | | |
|---|---|---|---|---|---|
| *RPL30* | -0.104964 | 0.03761 | -2.790847 | 0.00525703 | 0.087622118 |
| *EBF1* | -0.462589 | 0.165822 | -2.789665 | 0.00527625 | 0.087816791 |
| *GM2A* | 0.469938 | 0.168519 | 2.788639 | 0.00529301 | 0.087936495 |
| *U2AF1L4* | -0.348781 | 0.125087 | -2.788299 | 0.00529856 | 0.087936495 |
| *DDX17* | -0.185795 | 0.066678 | -2.786448 | 0.00532892 | 0.088189692 |
| *YTHDF2* | 0.270251 | 0.096988 | 2.786444 | 0.00532898 | 0.088189692 |
| *SHQ1* | 0.526961 | 0.189188 | 2.785379 | 0.00534652 | 0.088354241 |
| *SLC37A1* | -0.476667 | 0.171224 | -2.783886 | 0.00537119 | 0.088584972 |
| *EIF4E* | 0.22913 | 0.082314 | 2.783613 | 0.00537571 | 0.088584972 |
| *SSRP1* | 0.215526 | 0.077447 | 2.782889 | 0.00538772 | 0.08865737 |
| *DNAJC7* | 0.207126 | 0.074452 | 2.782002 | 0.00540247 | 0.088677861 |
| *CDK4* | 0.295512 | 0.106238 | 2.781606 | 0.00540907 | 0.088677861 |
| *TNFRSF13B* | -0.865163 | 0.311049 | -2.78144 | 0.00541183 | 0.088677861 |
| *NAA20* | 0.221484 | 0.079722 | 2.778214 | 0.00546586 | 0.089437147 |
| *LRRC59* | 0.268424 | 0.096678 | 2.776463 | 0.0054954 | 0.089779248 |
| *GNB5* | -0.312674 | 0.112632 | -2.77606 | 0.0055022 | 0.089779248 |
| *TMEM201* | 0.826319 | 0.297814 | 2.774617 | 0.00552668 | 0.090052392 |
| *NDUFB6* | 0.319485 | 0.115186 | 2.773651 | 0.00554311 | 0.090193777 |
| *CARNS1* | -1.258642 | 0.453898 | -2.772964 | 0.00555482 | 0.090258084 |
| *TPT1* | -0.164635 | 0.059426 | -2.770448 | 0.00559793 | 0.09083171 |
| *BUD23* | 0.249997 | 0.090493 | 2.762625 | 0.00573387 | 0.092907783 |
| *ECHS1* | 0.260218 | 0.094246 | 2.761038 | 0.0057618 | 0.093230609 |
| *PSMD13* | 0.226465 | 0.082059 | 2.759796 | 0.00578375 | 0.093455678 |
| *CEP135* | -0.32215 | 0.116821 | -2.757642 | 0.005822 | 0.093893201 |
| *LPP* | -0.413934 | 0.150119 | -2.757363 | 0.00582696 | 0.093893201 |
| *SPRYD4* | 0.461092 | 0.167471 | 2.753268 | 0.00590036 | 0.094944345 |
| *NICN1* | 0.670679 | 0.243749 | 2.751516 | 0.00593201 | 0.095321821 |
| *TMEM219* | -0.22795 | 0.082918 | -2.749087 | 0.00597616 | 0.095794519 |
| *AARS* | 0.352858 | 0.128359 | 2.748991 | 0.00597789 | 0.095794519 |
| *FLNA* | -0.492881 | 0.179365 | -2.747919 | 0.00599748 | 0.095976209 |
| *PFDN5* | -0.142789 | 0.05199 | -2.746485 | 0.00602377 | 0.096133547 |
| *CD5* | -1.474209 | 0.536763 | -2.746481 | 0.00602384 | 0.096133547 |
| *SLC16A3* | 0.743118 | 0.270655 | 2.745625 | 0.00603957 | 0.096252575 |
| *ATP6V0B* | 0.215607 | 0.078674 | 2.74053 | 0.00613401 | 0.097623961 |
| *PCBD2* | -0.413962 | 0.151079 | -2.740042 | 0.00614314 | 0.09763563 |
| *MYEF2* | -0.519116 | 0.189747 | -2.73583 | 0.00622231 | 0.098692352 |
| *GKAP1* | 0.535103 | 0.195607 | 2.735604 | 0.00622659 | 0.098692352 |
| *EIF2S2* | 0.224341 | 0.082064 | 2.733729 | 0.00626216 | 0.09912099 |
| *FAM210A* | 0.304515 | 0.111453 | 2.732229 | 0.00629073 | 0.099339905 |
| *MPHOSPH9* | 0.44896 | 0.164327 | 2.732107 | 0.00629306 | 0.099339905 |
| *PPFIBP2* | 0.435861 | 0.159923 | 2.725444 | 0.00642152 | 0.101230243 |
| *ZNF593* | 0.289416 | 0.106402 | 2.720027 | 0.00652765 | 0.10276418 |
| *DOP1A* | -0.887759 | 0.326786 | -2.716638 | 0.00659488 | 0.1036302 |
| *HNRNPK* | 0.144197 | 0.053085 | 2.716357 | 0.00660048 | 0.1036302 |
| *PABPC1* | -0.157667 | 0.058055 | -2.715832 | 0.00661095 | 0.103654634 |
| *PRPF31* | 0.224379 | 0.082643 | 2.715036 | 0.00662685 | 0.103764216 |
| *KCNH4* | 1.530315 | 0.564006 | 2.713296 | 0.00666175 | 0.104170429 |

| | | | | | |
|---|---|---|---|---|---|
| *GPR183* | -0.401809 | 0.148226 | -2.710782 | 0.00671247 | 0.104715208 |
| *PDCD2L* | 0.406473 | 0.149953 | 2.710677 | 0.00671459 | 0.104715208 |
| *ST6GAL1* | -0.256325 | 0.0946 | -2.709569 | 0.00673708 | 0.104840669 |
| *BANK1* | -0.454694 | 0.167832 | -2.709215 | 0.00674426 | 0.104840669 |
| *TNIP1* | 0.222058 | 0.081973 | 2.708928 | 0.0067501 | 0.104840669 |
| *PRPS1* | 0.325589 | 0.120219 | 2.708307 | 0.00676274 | 0.104840669 |
| *UBALD1* | 0.296529 | 0.109499 | 2.708064 | 0.00676769 | 0.104840669 |
| *ASF1A* | 0.343471 | 0.12689 | 2.706831 | 0.00679289 | 0.105091008 |
| *LRPAP1* | 0.261672 | 0.096744 | 2.704803 | 0.00683449 | 0.10550244 |
| *SIGLEC5* | 1.420472 | 0.525196 | 2.704651 | 0.00683762 | 0.10550244 |
| *RPP25L* | 0.286646 | 0.106052 | 2.702886 | 0.00687404 | 0.105893038 |
| *DDB2* | -0.298952 | 0.110644 | -2.701922 | 0.006894 | 0.105893038 |
| *BCL11A* | -0.261965 | 0.096971 | -2.70148 | 0.00690316 | 0.105893038 |
| *MRPL12* | 0.287653 | 0.10648 | 2.701465 | 0.00690348 | 0.105893038 |
| *PIK3CA* | -0.349272 | 0.129301 | -2.701226 | 0.00690844 | 0.105893038 |
| *PPIB* | 0.183681 | 0.068057 | 2.698939 | 0.0069561 | 0.106483168 |
| *MNT* | 0.56974 | 0.211184 | 2.697832 | 0.00697926 | 0.106697339 |
| *ISCA1* | 0.240342 | 0.089132 | 2.696466 | 0.00700795 | 0.10699546 |
| *JDP2* | 1.010952 | 0.375001 | 2.695863 | 0.00702066 | 0.106997223 |
| *CCT4* | 0.163616 | 0.060702 | 2.695382 | 0.0070308 | 0.106997223 |
| *C1QBP* | 0.297962 | 0.110555 | 2.695152 | 0.00703566 | 0.106997223 |
| *MX2* | -0.794506 | 0.294841 | -2.69469 | 0.00704541 | 0.107005675 |
| *KCTD3* | -1.427453 | 0.529995 | -2.693335 | 0.00707412 | 0.107301503 |
| *OGFRL1* | -0.481368 | 0.179189 | -2.686377 | 0.00722315 | 0.109419383 |
| *MRPS35* | 0.243689 | 0.0908 | 2.683788 | 0.00727933 | 0.110127039 |
| *NAALADL1* | -0.456754 | 0.170299 | -2.68208 | 0.0073166 | 0.110467778 |
| *C8orf33* | 0.224481 | 0.083703 | 2.681886 | 0.00732084 | 0.110467778 |
| *CAMK2G* | -0.427249 | 0.159349 | -2.681221 | 0.00733541 | 0.110544295 |
| *MAT2A* | 0.211173 | 0.078987 | 2.673524 | 0.00750588 | 0.11296689 |
| *PDE7A* | 0.301165 | 0.112698 | 2.672322 | 0.00753283 | 0.113226052 |
| *FKBP5* | 0.58127 | 0.217617 | 2.671073 | 0.00756092 | 0.113379864 |
| *ABRAXAS1* | -0.365325 | 0.136775 | -2.671 | 0.00756256 | 0.113379864 |
| *KLHL5* | -0.41492 | 0.155406 | -2.669917 | 0.00758701 | 0.113600058 |
| *SGSM3* | -0.276001 | 0.103408 | -2.669053 | 0.00760655 | 0.113746269 |
| *PDCD11* | 0.300547 | 0.112709 | 2.666572 | 0.00766292 | 0.114442183 |
| *NMT2* | -0.616984 | 0.231466 | -2.665551 | 0.00768623 | 0.114643135 |
| *PDCD5* | 0.2188 | 0.082135 | 2.66389 | 0.00772428 | 0.115063062 |
| *PDIA4* | 0.321549 | 0.120761 | 2.662698 | 0.0077517 | 0.115323874 |
| *RPL26L1* | 0.224332 | 0.084266 | 2.662185 | 0.00776351 | 0.115352065 |
| *UPP1* | 0.766389 | 0.287929 | 2.661727 | 0.00777409 | 0.115361924 |
| *SNRPG* | 0.171422 | 0.064465 | 2.659168 | 0.00783339 | 0.116093769 |
| *MYO7B* | -2.174092 | 0.817589 | -2.65915 | 0.00783381 | NA |
| *EIF3A* | 0.181735 | 0.068387 | 2.657462 | 0.00787315 | 0.116471721 |
| *SLC3A2* | 0.286677 | 0.107886 | 2.657215 | 0.00787891 | 0.116471721 |
| *IFNAR2* | 0.293397 | 0.110445 | 2.656492 | 0.00789584 | 0.116573887 |
| *CAST* | -0.296435 | 0.11162 | -2.655757 | 0.00791306 | 0.116679967 |
| *ZNF165* | 0.536538 | 0.202257 | 2.652751 | 0.00798388 | 0.117575311 |

| | | | | | |
|---|---|---|---|---|---|
| *HIGD2A* | -0.207271 | 0.078149 | -2.65226 | 0.0079955 | 0.117597483 |
| *PTGS1* | -0.720528 | 0.27196 | -2.649392 | 0.00806367 | 0.118450465 |
| *TRUB2* | 0.346772 | 0.130918 | 2.648775 | 0.00807841 | 0.118517367 |
| *SH2B3* | 0.544104 | 0.205544 | 2.647142 | 0.00811753 | 0.118941223 |
| *ETFA* | 0.272692 | 0.103137 | 2.643989 | 0.00819354 | 0.119903889 |
| *RPS23* | -0.127011 | 0.048088 | -2.641189 | 0.00826156 | 0.12074744 |
| *NPC2* | -0.329709 | 0.124864 | -2.640538 | 0.00827745 | 0.12075434 |
| *DNAJB1* | 0.363232 | 0.137588 | 2.640003 | 0.00829052 | 0.12075434 |
| *SLC38A1* | -0.243315 | 0.092168 | -2.639895 | 0.00829317 | 0.12075434 |
| *SH2D3C* | -0.50342 | 0.190825 | -2.638118 | 0.00833675 | 0.120962829 |
| *TXNRD1* | 0.294261 | 0.111548 | 2.637977 | 0.00834023 | 0.120962829 |
| *GPR137* | -0.264167 | 0.10014 | -2.637965 | 0.00834053 | 0.120962829 |
| *SRM* | 0.447198 | 0.169546 | 2.637617 | 0.00834908 | 0.120962829 |
| *GADD45A* | 0.57579 | 0.218452 | 2.635777 | 0.00839449 | 0.121469585 |
| *CHORDC1* | 0.292606 | 0.111113 | 2.633418 | 0.00845304 | 0.122164757 |
| *MATK* | 1.110841 | 0.421951 | 2.632631 | 0.00847264 | 0.122296201 |
| *GFI1* | 1.115809 | 0.423946 | 2.631962 | 0.00848934 | 0.122385278 |
| *HPRT1* | 0.226979 | 0.086256 | 2.631442 | 0.00850234 | 0.122421007 |
| *PPP2CA* | 0.237295 | 0.090199 | 2.630797 | 0.00851848 | 0.122501884 |
| *RPIA* | 0.223775 | 0.085137 | 2.628413 | 0.00857844 | 0.123211758 |
| *SAP18* | 0.143985 | 0.054821 | 2.626477 | 0.00862739 | 0.123762064 |
| *CEP126* | -0.842042 | 0.320694 | -2.625689 | 0.00864738 | 0.12382111 |
| *CCDC141* | -0.828149 | 0.315428 | -2.625476 | 0.00865279 | 0.12382111 |
| *ATG101* | 0.344421 | 0.131212 | 2.624912 | 0.00866713 | 0.123874002 |
| *CCR9* | 2.108226 | 0.80326 | 2.624587 | 0.00867542 | NA |
| *ZBTB18* | -0.516713 | 0.196929 | -2.623859 | 0.00869398 | 0.124105247 |
| *CCDC71L* | -0.609571 | 0.232386 | -2.623092 | 0.00871357 | 0.124232467 |
| *TGFBR2* | -0.235813 | 0.089924 | -2.622367 | 0.00873214 | 0.124344789 |
| *ZBTB8OS* | 0.221741 | 0.084608 | 2.620807 | 0.00877219 | 0.124762406 |
| *NASP* | 0.208072 | 0.07941 | 2.620235 | 0.00878692 | 0.124819299 |
| *WDR97* | 1.7268 | 0.65938 | 2.618822 | 0.0088234 | NA |
| *DALRD3* | -0.254569 | 0.097362 | -2.614676 | 0.0089312 | 0.126714147 |
| *MAN1B1* | -0.391738 | 0.149867 | -2.613914 | 0.00895117 | 0.126740458 |
| *TLE1* | 0.633288 | 0.242289 | 2.613773 | 0.00895484 | 0.126740458 |
| *TMEM230* | 0.283307 | 0.10844 | 2.612576 | 0.00898626 | 0.127030601 |
| *PPAN* | 0.255402 | 0.097818 | 2.610976 | 0.00902841 | 0.127471508 |
| *PDE4B* | -0.284442 | 0.108987 | -2.609861 | 0.0090579 | 0.127732862 |
| *RASSF2* | -0.340081 | 0.130393 | -2.608114 | 0.00910426 | 0.128099281 |
| *SPPL2B* | -0.402083 | 0.15417 | -2.608052 | 0.00910591 | 0.128099281 |
| *IKBKB* | -0.356026 | 0.13656 | -2.607106 | 0.00913111 | 0.128181831 |
| *MALAT1* | -0.176095 | 0.067548 | -2.606976 | 0.00913458 | 0.128181831 |
| *FAM41C* | 0.461558 | 0.177092 | 2.606319 | 0.00915211 | 0.128181831 |
| *CHCHD2* | 0.183599 | 0.070447 | 2.60618 | 0.00915585 | 0.128181831 |
| *LGALS1* | -0.817439 | 0.313725 | -2.605589 | 0.00917163 | 0.128191786 |
| *KANSL3* | 0.727623 | 0.279283 | 2.60533 | 0.00917859 | 0.128191786 |
| *PPM1K* | -0.287916 | 0.110641 | -2.602244 | 0.00926159 | 0.128962085 |
| *ALKBH2* | 0.298003 | 0.114519 | 2.60221 | 0.0092625 | 0.128962085 |

| | | | | | |
|---|---|---|---|---|---|
| *NFKBIB* | 0.262415 | 0.100859 | 2.601796 | 0.0092737 | 0.128962085 |
| *YPEL5* | -0.345547 | 0.13284 | -2.601225 | 0.00928914 | 0.128962085 |
| *MRPL47* | 0.207079 | 0.079608 | 2.601224 | 0.00928917 | 0.128962085 |
| *TCTN1* | -0.865581 | 0.332876 | -2.600313 | 0.00931388 | 0.129150925 |
| *GABARAPL1* | 0.728463 | 0.280226 | 2.599556 | 0.00933444 | 0.129213944 |
| *PSD3* | -1.038237 | 0.399462 | -2.599092 | 0.00934707 | 0.129213944 |
| *TFEB* | 0.450367 | 0.17329 | 2.598921 | 0.00935174 | 0.129213944 |
| *HM13* | 0.219106 | 0.084331 | 2.598149 | 0.00937278 | 0.12926196 |
| *ENSG00000276136* | -0.826384 | 0.318104 | -2.59784 | 0.00938121 | 0.12926196 |
| *GZF1* | 0.568627 | 0.218907 | 2.597572 | 0.00938855 | 0.12926196 |
| *CLN6* | 0.290213 | 0.111939 | 2.592599 | 0.00952538 | 0.130990847 |
| *RPL3* | -0.143583 | 0.0554 | -2.591779 | 0.00954811 | 0.131148465 |
| *SH2D2A* | 0.721915 | 0.278941 | 2.588057 | 0.00965189 | 0.132417579 |
| *MRPS7* | 0.193945 | 0.074993 | 2.586175 | 0.00970476 | 0.132986054 |
| *ICA1* | 1.945017 | 0.752661 | 2.584187 | 0.00976088 | 0.133597792 |
| *ENSG00000204758* | -1.386389 | 0.537126 | -2.581124 | 0.00984792 | 0.134622297 |
| *BARD1* | -0.482156 | 0.186829 | -2.58074 | 0.00985888 | 0.134622297 |
| *NUDT5* | 0.233279 | 0.090439 | 2.579418 | 0.0098967 | 0.134980354 |
| *ZNF277* | -0.285118 | 0.110557 | -2.578933 | 0.00991061 | 0.135003487 |
| *XRCC5* | 0.156307 | 0.060618 | 2.57855 | 0.00992161 | 0.135003487 |
| *PRELID3B* | 0.223588 | 0.086775 | 2.576655 | 0.00997615 | 0.135586992 |
| *KBTBD3* | -0.411199 | 0.159671 | -2.575281 | 0.01001588 | 0.135968223 |
| *IL10RA* | -0.394788 | 0.153368 | -2.57413 | 0.01004924 | 0.136262122 |
| *RAP2C* | -0.308446 | 0.119884 | -2.572863 | 0.01008611 | 0.136602769 |
| *NOP2* | 0.335851 | 0.130566 | 2.572265 | 0.01010354 | 0.136611048 |
| *CEBPB* | 0.756845 | 0.294267 | 2.571967 | 0.01011224 | 0.136611048 |
| *PKN1* | 0.308527 | 0.119973 | 2.571635 | 0.01012195 | 0.136611048 |
| *CEMIP2* | -0.670151 | 0.260639 | -2.571179 | 0.0101353 | 0.136632742 |
| *EML2* | 0.392483 | 0.152762 | 2.569246 | 0.01019201 | 0.137188239 |
| *DYNLT3* | -0.293847 | 0.114383 | -2.568971 | 0.01020009 | 0.137188239 |
| *PVT1* | 0.516265 | 0.201133 | 2.566792 | 0.01026442 | 0.137894063 |
| *PROC* | -1.908175 | 0.743563 | -2.56626 | 0.01028018 | NA |
| *CCDC59* | 0.176158 | 0.068704 | 2.564001 | 0.01034732 | 0.13884748 |
| *MARCHF1* | -0.523948 | 0.204438 | -2.56287 | 0.0103811 | 0.13900933 |
| *AK6* | 0.233272 | 0.091034 | 2.562479 | 0.0103928 | 0.13900933 |
| *PSMA6* | 0.267393 | 0.104369 | 2.562009 | 0.01040687 | 0.13900933 |
| *TOMM22* | 0.185726 | 0.072493 | 2.561998 | 0.01040718 | 0.13900933 |
| *EIF2S1* | 0.223707 | 0.087343 | 2.561253 | 0.01042954 | 0.139017502 |
| *PBX4* | 1.086479 | 0.424223 | 2.561102 | 0.01043406 | 0.139017502 |
| *MAFG.DT* | 0.864207 | 0.337505 | 2.560579 | 0.01044978 | 0.139017502 |
| *SF3B5* | 0.138803 | 0.054212 | 2.560386 | 0.01045559 | 0.139017502 |
| *CLIC1* | -0.277843 | 0.108536 | -2.559915 | 0.01046978 | 0.139047251 |
| *RCSD1* | -0.315054 | 0.123163 | -2.558019 | 0.01052703 | 0.139514222 |
| *UQCR11* | 0.177011 | 0.0692 | 2.557953 | 0.01052904 | 0.139514222 |
| *IER5L* | -1.392427 | 0.544435 | -2.557561 | 0.01054092 | 0.139514222 |
| *CHRNB1* | -0.422094 | 0.16517 | -2.55551 | 0.01060324 | 0.140179705 |
| *NDUFAB1* | 0.24681 | 0.096617 | 2.55451 | 0.01063375 | 0.140423444 |

| | | | | | |
|---|---|---|---|---|---|
| *CRYM* | -1.389567 | 0.544107 | -2.553847 | 0.01065401 | 0.140531476 |
| *SNX20* | -0.420019 | 0.16459 | -2.551913 | 0.01071333 | 0.14115384 |
| *ZNF33B* | -0.433343 | 0.169879 | -2.550893 | 0.01074474 | 0.141407542 |
| *M6PR* | 0.169228 | 0.066353 | 2.550429 | 0.01075905 | 0.141435952 |
| *MRPL22* | 0.189382 | 0.074276 | 2.549724 | 0.01078083 | 0.141552937 |
| *CEBPG* | 0.242785 | 0.095234 | 2.549353 | 0.01079229 | 0.141552937 |
| *USP6NL* | -0.392052 | 0.153836 | -2.548507 | 0.01081852 | 0.141737196 |
| *TMEM256* | -0.203012 | 0.079699 | -2.547222 | 0.01085843 | 0.141859106 |
| *TRMT1* | 0.216658 | 0.08506 | 2.547137 | 0.01086108 | 0.141859106 |
| *SMIM7* | -0.164317 | 0.064513 | -2.54703 | 0.0108644 | 0.141859106 |
| *FOXO4* | -1.055129 | 0.414426 | -2.546004 | 0.01089639 | 0.141936624 |
| *SLC25A15* | -1.018343 | 0.400009 | -2.545798 | 0.01090282 | 0.141936624 |
| *POLR1E* | 0.313677 | 0.12322 | 2.545666 | 0.01090694 | 0.141936624 |
| *PSMA7* | 0.193071 | 0.075869 | 2.544803 | 0.01093393 | 0.142128852 |
| *FUBP1* | 0.202377 | 0.079542 | 2.544263 | 0.01095085 | 0.142130414 |
| *NAP1L1* | -0.186165 | 0.073178 | -2.54402 | 0.01095848 | 0.142130414 |
| *LINC00847* | 0.394266 | 0.155018 | 2.543351 | 0.0109795 | 0.142244408 |
| *MCRIP2* | 0.328887 | 0.129337 | 2.542879 | 0.01099434 | 0.142278279 |
| *HDGF* | 0.246145 | 0.096845 | 2.541629 | 0.01103374 | 0.142629425 |
| *HGSNAT* | -0.48864 | 0.192396 | -2.539764 | 0.01109272 | 0.14301256 |
| *SHISAL2A* | 0.367588 | 0.144757 | 2.539342 | 0.01110611 | 0.14301256 |
| *ATP2A2* | 0.354702 | 0.139686 | 2.539276 | 0.01110823 | 0.14301256 |
| *LPAR5* | -0.468398 | 0.184471 | -2.53914 | 0.01111255 | 0.14301256 |
| *RIC8B* | -0.99791 | 0.393152 | -2.538228 | 0.01114155 | 0.143227384 |
| *PSMB3* | 0.180877 | 0.071333 | 2.535663 | 0.01122347 | 0.144067364 |
| *TWNK* | 0.441664 | 0.174198 | 2.535408 | 0.01123166 | 0.144067364 |
| *PSMD14* | 0.198312 | 0.078284 | 2.533225 | 0.01130185 | 0.144762899 |
| *CPEB4* | 0.512464 | 0.202335 | 2.532756 | 0.01131695 | 0.144762899 |
| *DRAM1* | 0.56941 | 0.224835 | 2.532563 | 0.01132321 | 0.144762899 |
| *MDM2* | 0.309435 | 0.122223 | 2.531725 | 0.01135029 | 0.144928518 |
| *NCBP2AS2* | 0.231936 | 0.091624 | 2.531392 | 0.01136108 | 0.144928518 |
| *LSM1* | 0.159296 | 0.063051 | 2.526462 | 0.01152177 | 0.146817384 |
| *MKNK2* | -0.378767 | 0.149945 | -2.526032 | 0.0115359 | 0.146836595 |
| *S1PR1* | -0.5021 | 0.198854 | -2.524967 | 0.01157091 | 0.147121294 |
| *NT5DC3* | 0.619346 | 0.245389 | 2.523942 | 0.01160472 | 0.147390076 |
| *LGALS3BP* | -1.949695 | 0.773139 | -2.521792 | 0.01167588 | NA |
| *PYCR2* | 0.250127 | 0.099241 | 2.520409 | 0.01172184 | 0.148606224 |
| *EIF3D* | -0.17809 | 0.070663 | -2.520267 | 0.01172658 | 0.148606224 |
| *CEP112* | -1.033338 | 0.410074 | -2.519882 | 0.01173941 | 0.148606224 |
| *WDR18* | 0.275584 | 0.109403 | 2.518975 | 0.0117697 | 0.148606224 |
| *MTDH* | 0.243801 | 0.096794 | 2.518764 | 0.01177674 | 0.148606224 |
| *WDFY2* | -0.442546 | 0.17573 | -2.518325 | 0.01179143 | 0.148606224 |
| *EHD4* | -0.305749 | 0.121424 | -2.518033 | 0.01180124 | 0.148606224 |
| *PPDPF* | -0.285914 | 0.113549 | -2.51799 | 0.01180266 | 0.148606224 |
| *CRIP3* | -0.702645 | 0.279112 | -2.517426 | 0.01182157 | 0.148683445 |
| *CCDC50* | -0.217938 | 0.08665 | -2.515167 | 0.0118976 | 0.149202546 |
| *COL19A1* | -0.566319 | 0.225165 | -2.515134 | 0.01189873 | 0.149202546 |

| | | | | | |
|---|---|---|---|---|---|
| *ST8SIA4* | -0.399716 | 0.158929 | -2.515057 | 0.01190132 | 0.149202546 |
| *MRPS17* | 0.288832 | 0.114864 | 2.514548 | 0.01191849 | 0.149257008 |
| *ADA* | -0.337338 | 0.13433 | -2.511254 | 0.01203031 | 0.150433099 |
| *NUDT14* | -0.569782 | 0.226913 | -2.511021 | 0.01203827 | 0.150433099 |
| *ANXA5* | -0.385043 | 0.15351 | -2.508262 | 0.01213267 | 0.151450144 |
| *WDR36* | 0.310033 | 0.123625 | 2.507861 | 0.01214644 | 0.151459014 |
| *SRSF3* | 0.174033 | 0.069405 | 2.507484 | 0.01215942 | 0.151459014 |
| *SORL1* | -0.436197 | 0.173995 | -2.506953 | 0.0121777 | 0.15152441 |
| *CD180* | 0.515492 | 0.205691 | 2.506149 | 0.01220541 | 0.151707006 |
| *SYVN1* | 0.330577 | 0.131958 | 2.505175 | 0.01223908 | 0.151903633 |
| *WDR43* | 0.248934 | 0.099388 | 2.504667 | 0.01225667 | 0.151903633 |
| *PIP4P2* | -0.618877 | 0.2471 | -2.504559 | 0.0122604 | 0.151903633 |
| *RPL34* | -0.123316 | 0.049258 | -2.503481 | 0.01229784 | 0.152205344 |
| *NUCB1* | -0.251519 | 0.100622 | -2.499636 | 0.01243209 | 0.153470659 |
| *GABBR1* | -0.53894 | 0.215616 | -2.499538 | 0.01243554 | 0.153470659 |
| *MTHFD1L* | 0.526206 | 0.210531 | 2.499421 | 0.01243965 | 0.153470659 |
| *AATF* | 0.248893 | 0.099621 | 2.498405 | 0.01247537 | 0.153655902 |
| *IFI44* | -0.937714 | 0.375349 | -2.498243 | 0.01248108 | 0.153655902 |
| *ILK* | -0.290677 | 0.116405 | -2.497107 | 0.01252113 | 0.153986017 |
| *BZW2* | 0.268247 | 0.107544 | 2.494306 | 0.01262038 | 0.154981963 |
| *FBXL20* | -0.430621 | 0.172658 | -2.49407 | 0.01262875 | 0.154981963 |
| *MBD2* | 0.29924 | 0.120048 | 2.492672 | 0.0126786 | 0.155429707 |
| *CHD7* | 0.288417 | 0.115823 | 2.490158 | 0.01276864 | 0.156368765 |
| *TMEM41B* | 0.399499 | 0.160522 | 2.488755 | 0.01281914 | 0.156791095 |
| *TBC1D2B* | -0.869015 | 0.349267 | -2.488113 | 0.01284229 | 0.156791095 |
| *ENSG00000272211* | -0.502741 | 0.20206 | -2.488078 | 0.01284356 | 0.156791095 |
| *ENSG00000225938* | -0.926219 | 0.372489 | -2.486568 | 0.0128982 | 0.157138761 |
| *DDX1* | 0.228677 | 0.091966 | 2.486544 | 0.01289905 | 0.157138761 |
| *LRRC1* | -0.84349 | 0.339525 | -2.484325 | 0.01297974 | 0.157940986 |
| *RAB11B* | -0.216475 | 0.087159 | -2.48368 | 0.01300327 | 0.157940986 |
| *PCNA* | 0.335682 | 0.135159 | 2.483615 | 0.01300563 | 0.157940986 |
| *FLOT2* | -0.271795 | 0.109489 | -2.482398 | 0.01305014 | 0.158316326 |
| *EIF3L* | -0.198101 | 0.079821 | -2.481808 | 0.01307178 | 0.158354777 |
| *ENSG00000260349* | 0.624229 | 0.251546 | 2.481569 | 0.01308054 | 0.158354777 |
| *MRPL40* | 0.216385 | 0.087326 | 2.477908 | 0.01321551 | 0.159735016 |
| *FXN* | 0.327326 | 0.132107 | 2.477733 | 0.01322201 | 0.159735016 |
| *LRRC25* | 1.679857 | 0.678879 | 2.474459 | 0.01334383 | NA |
| *NAT14* | 0.65775 | 0.265956 | 2.473155 | 0.01339259 | 0.161610067 |
| *ENSG00000267136* | 2.373139 | 0.959628 | 2.472977 | 0.01339928 | NA |
| *RAB6A* | -0.242997 | 0.098276 | -2.472601 | 0.01341339 | 0.161610067 |
| *PDCD4* | 0.397677 | 0.160857 | 2.472232 | 0.01342722 | 0.161610067 |
| *UTP11* | 0.258687 | 0.104643 | 2.472084 | 0.01343278 | 0.161610067 |
| *ATP2B1* | -0.176275 | 0.071321 | -2.471562 | 0.01345244 | 0.161679439 |
| *CCSER1* | -0.658528 | 0.266812 | -2.468134 | 0.01358195 | 0.163067545 |
| *CABLES1* | -0.863119 | 0.349824 | -2.467296 | 0.01361378 | 0.163259405 |
| *HSD17B11* | -0.281073 | 0.113961 | -2.4664 | 0.01364789 | 0.163259405 |
| *HNRNPA2B1* | 0.176156 | 0.071427 | 2.46624 | 0.01365397 | 0.163259405 |

| | | | | | |
|---|---|---|---|---|---|
| *RBM8A* | 0.13283 | 0.053867 | 2.465884 | 0.01366756 | 0.163259405 |
| *CDK19* | -0.40743 | 0.165227 | -2.46587 | 0.0136681 | 0.163259405 |
| *RPL12* | -0.147538 | 0.059855 | -2.464904 | 0.01370501 | 0.163532442 |
| *FAM167A* | 0.984911 | 0.399812 | 2.463438 | 0.01376117 | 0.164002412 |
| *SLC9B2* | 0.318432 | 0.129279 | 2.46314 | 0.01377259 | 0.164002412 |
| *RASGRP2* | -0.417422 | 0.169544 | -2.462032 | 0.01381523 | 0.164341954 |
| *HDDC2* | 0.194358 | 0.078988 | 2.460602 | 0.0138704 | 0.164712643 |
| *LIMD2* | -0.253589 | 0.103071 | -2.460332 | 0.01388085 | 0.164712643 |
| *CPNE5* | 0.619799 | 0.25194 | 2.460102 | 0.01388977 | 0.164712643 |
| *SLC29A1* | 0.605157 | 0.246023 | 2.459759 | 0.01390303 | 0.164712643 |
| *RCN1* | 0.334281 | 0.136107 | 2.456022 | 0.01404844 | 0.166133166 |
| *ITPA* | 0.29196 | 0.118879 | 2.455944 | 0.01405149 | 0.166133166 |
| *PLEKHJ1* | -0.241962 | 0.098622 | -2.453441 | 0.01414969 | 0.167124334 |
| *PRKDC* | 0.239646 | 0.097722 | 2.45234 | 0.01419304 | 0.167282076 |
| *FAM126A* | -0.380904 | 0.155331 | -2.45221 | 0.01419816 | 0.167282076 |
| *GDF11* | -0.424775 | 0.173236 | -2.452007 | 0.01420618 | 0.167282076 |
| *CHD2* | -0.247263 | 0.100904 | -2.450475 | 0.01426679 | 0.167825927 |
| *CAMLG* | -0.222047 | 0.090631 | -2.45 | 0.01428561 | 0.167877548 |
| *TRIAP1* | 0.240388 | 0.098141 | 2.449416 | 0.01430881 | 0.167980458 |
| *BTD* | -0.714298 | 0.291765 | -2.448195 | 0.01435738 | 0.168188325 |
| *CAMK2D* | -0.391557 | 0.159944 | -2.448087 | 0.0143617 | 0.168188325 |
| *CYTH4* | -0.468115 | 0.191233 | -2.447882 | 0.01436988 | 0.168188325 |
| *SNX14* | -0.333855 | 0.136407 | -2.447492 | 0.01438541 | 0.168200913 |
| *FAM214B* | -0.819856 | 0.335105 | -2.446568 | 0.01442234 | 0.168322134 |
| *TRMT10B* | -0.344883 | 0.14097 | -2.446509 | 0.01442472 | 0.168322134 |
| *GPR65* | -0.269694 | 0.11029 | -2.445312 | 0.01447269 | 0.168712657 |
| *FAM53B* | -0.362471 | 0.148323 | -2.443797 | 0.01453361 | 0.169253283 |
| *SNRNP25* | -0.202422 | 0.082904 | -2.441628 | 0.01462122 | 0.170103248 |
| *DRAP1* | -0.179718 | 0.073624 | -2.441044 | 0.01464488 | 0.170136918 |
| *SORBS3* | -0.782348 | 0.320581 | -2.440404 | 0.01467083 | 0.170136918 |
| *SLC25A19* | 0.28208 | 0.115589 | 2.440372 | 0.01467213 | 0.170136918 |
| *OTUD4* | 0.349242 | 0.143125 | 2.440114 | 0.01468261 | 0.170136918 |
| *SRSF1* | 0.182403 | 0.074773 | 2.439431 | 0.01471043 | 0.170289707 |
| *SETX* | -0.322884 | 0.132407 | -2.438571 | 0.01474547 | 0.170525639 |
| *LRRFIP1* | -0.224625 | 0.09215 | -2.437605 | 0.01478494 | 0.17057206 |
| *GRINA* | 0.280311 | 0.115004 | 2.437396 | 0.01479347 | 0.17057206 |
| *CCDC112* | -0.3996 | 0.163953 | -2.437285 | 0.01479803 | 0.17057206 |
| *FDFT1* | 0.209271 | 0.08588 | 2.43678 | 0.01481868 | 0.17057206 |
| *APOBEC3D* | -0.398101 | 0.163378 | -2.43668 | 0.01482279 | 0.17057206 |
| *RRP1* | 0.238943 | 0.098081 | 2.436175 | 0.01484349 | 0.170641448 |
| *ENSG00000225335* | -2.163301 | 0.888446 | -2.434926 | 0.01489484 | NA |
| *ZNF224* | -0.326087 | 0.133921 | -2.434917 | 0.01489521 | 0.17082895 |
| *MMP7* | 1.298591 | 0.533381 | 2.434641 | 0.01490658 | 0.17082895 |
| *PEA15* | -0.271805 | 0.111644 | -2.43456 | 0.01490989 | 0.17082895 |
| *EIF3E* | -0.155126 | 0.063724 | -2.434351 | 0.01491853 | 0.17082895 |
| *CEBPZ* | 0.239788 | 0.098548 | 2.433215 | 0.01496541 | 0.171197255 |
| *PEX2* | 0.234016 | 0.096261 | 2.431062 | 0.01505464 | 0.172003132 |

| | | | | | |
|---|---|---|---|---|---|
| *GLIPR1* | -0.364921 | 0.150124 | -2.430802 | 0.01506543 | 0.172003132 |
| *MRPL50* | 0.260567 | 0.107238 | 2.429808 | 0.01510684 | 0.172306817 |
| *SLC25A17* | 0.353095 | 0.145392 | 2.428574 | 0.01515832 | 0.172552534 |
| *PGM2L1* | -0.481016 | 0.198073 | -2.428482 | 0.01516217 | 0.172552534 |
| *MRPL18* | 0.21872 | 0.090074 | 2.428226 | 0.01517288 | 0.172552534 |
| *EHBP1* | 0.466588 | 0.192226 | 2.427284 | 0.01521235 | 0.172644842 |
| *PIK3CD* | 0.355476 | 0.146461 | 2.427106 | 0.01521982 | 0.172644842 |
| *SSB* | 0.181049 | 0.074606 | 2.426733 | 0.01523547 | 0.172644842 |
| *NAGK* | -0.274589 | 0.113157 | -2.426617 | 0.01524035 | 0.172644842 |
| *APMAP* | 0.410378 | 0.169149 | 2.426138 | 0.01526046 | 0.172704414 |
| *KLF6* | -0.301776 | 0.124414 | -2.425573 | 0.01528424 | 0.172805509 |
| *ENSG00000273247* | -0.277732 | 0.114605 | -2.423373 | 0.01537715 | 0.173687111 |
| *WDR74* | 0.26191 | 0.108109 | 2.422645 | 0.01540799 | 0.173866681 |
| *UBE2T* | 0.51974 | 0.214578 | 2.422149 | 0.01542901 | 0.173935134 |
| *FAM111A.DT* | -0.550418 | 0.227285 | -2.42171 | 0.01544769 | 0.173977144 |
| *NR6A1* | 0.585931 | 0.242005 | 2.42115 | 0.0154715 | 0.173977491 |
| *RPL11* | -0.111328 | 0.045984 | -2.421006 | 0.01547763 | 0.173977491 |
| *ZNF782* | -0.867858 | 0.3587 | -2.419453 | 0.01554388 | 0.174501155 |
| *VCP* | 0.220626 | 0.091198 | 2.419211 | 0.01555421 | 0.174501155 |
| *ZNF239* | 0.842101 | 0.348147 | 2.418805 | 0.01557158 | 0.174519814 |
| *MRPL32* | 0.234406 | 0.096923 | 2.418471 | 0.01558588 | 0.174519814 |
| *CARD19* | -0.38935 | 0.16106 | -2.417422 | 0.0156309 | 0.174855652 |
| *MECR* | 0.431486 | 0.178599 | 2.415951 | 0.01569418 | 0.175394911 |
| *ENSG00000274605* | 0.619183 | 0.256416 | 2.414755 | 0.01574581 | 0.175448185 |
| *FAM219B* | -0.277731 | 0.115021 | -2.414608 | 0.01575216 | 0.175448185 |
| *VAMP1* | -0.450289 | 0.18649 | -2.414553 | 0.01575451 | 0.175448185 |
| *USP15* | -0.212484 | 0.088005 | -2.414443 | 0.01575927 | 0.175448185 |
| *ENSG00000274265* | -0.404671 | 0.167717 | -2.412813 | 0.01582995 | 0.175820233 |
| *TBL2* | 0.418827 | 0.173587 | 2.412779 | 0.01583141 | 0.175820233 |
| *KCNK6* | -0.362884 | 0.15041 | -2.412627 | 0.01583803 | 0.175820233 |
| *PLPP3* | 1.081252 | 0.44835 | 2.411625 | 0.01588159 | 0.175996084 |
| *WIZ* | 0.42598 | 0.17664 | 2.411567 | 0.01588412 | 0.175996084 |
| *PLEKHB2* | 0.250861 | 0.104062 | 2.410678 | 0.01592292 | 0.176258098 |
| *PAFAH1B2* | -0.198739 | 0.082465 | -2.409978 | 0.01595349 | 0.176428655 |
| *IKZF3* | -0.198214 | 0.082262 | -2.409554 | 0.01597205 | 0.176466123 |
| *TMED10* | 0.200866 | 0.083492 | 2.405817 | 0.01613634 | 0.178100318 |
| *MCM3AP.AS1* | -0.53901 | 0.224074 | -2.405495 | 0.01615058 | 0.178100318 |
| *ILVBL* | 0.419047 | 0.17433 | 2.403755 | 0.01622767 | 0.178780926 |
| *UXT* | -0.09809 | 0.040833 | -2.402254 | 0.01629437 | 0.179346007 |
| *ABCA2* | 1.364388 | 0.56808 | 2.401756 | 0.01631659 | 0.17942085 |
| *RPL28* | -0.164291 | 0.068513 | -2.397943 | 0.01648745 | 0.181128358 |
| *AFF3* | -0.373938 | 0.155989 | -2.397203 | 0.01652076 | 0.181286957 |
| *LINC01800* | -1.382686 | 0.576857 | -2.396931 | 0.01653305 | 0.181286957 |
| *LINC00926* | -0.387773 | 0.161811 | -2.396454 | 0.01655457 | 0.181352019 |
| *CASD1* | 0.565885 | 0.23637 | 2.394066 | 0.01666277 | 0.182240042 |
| *IRAK3* | -1.16196 | 0.485369 | -2.393973 | 0.01666696 | 0.182240042 |
| *IGHV4.34* | -1.795103 | 0.750611 | -2.391522 | 0.0167787 | NA |

| | | | | | |
|---|---|---|---|---|---|
| *RAE1* | 0.348222 | 0.145623 | 2.391262 | 0.01679056 | 0.183303471 |
| *CD19* | -0.186597 | 0.078037 | -2.391149 | 0.01679573 | 0.183303471 |
| *RAB30* | 0.272946 | 0.11424 | 2.389234 | 0.01688356 | 0.183611251 |
| *C12orf76* | -0.350074 | 0.146538 | -2.38897 | 0.01689566 | 0.183611251 |
| *HSDL1* | -0.648605 | 0.271506 | -2.388914 | 0.01689824 | 0.183611251 |
| *PSMB7* | 0.164706 | 0.068954 | 2.388632 | 0.01691125 | 0.183611251 |
| *ATP6V0A1* | 0.470972 | 0.197205 | 2.388228 | 0.01692983 | 0.183611251 |
| *DEF6* | -0.336372 | 0.140852 | -2.388117 | 0.01693495 | 0.183611251 |
| *CD48* | -0.220906 | 0.092522 | -2.387589 | 0.0169593 | 0.183611251 |
| *S100A11* | -0.416946 | 0.174646 | -2.387374 | 0.01696923 | 0.183611251 |
| *RIPOR1* | -0.387459 | 0.162313 | -2.387109 | 0.01698147 | 0.183611251 |
| *MAGOHB* | 0.222477 | 0.093199 | 2.387103 | 0.01698175 | 0.183611251 |
| *MBP* | -0.317382 | 0.132978 | -2.386729 | 0.01699901 | 0.18362719 |
| *ZFP14* | -0.353497 | 0.148163 | -2.385864 | 0.01703907 | 0.183889195 |
| *ANAPC11* | 0.158545 | 0.06649 | 2.384504 | 0.01710215 | 0.184264055 |
| *BCLAF3* | -0.698897 | 0.293154 | -2.384063 | 0.01712267 | 0.184264055 |
| *SF3B6* | 0.149106 | 0.062546 | 2.383937 | 0.01712853 | 0.184264055 |
| *FKBP1B* | -1.030945 | 0.432488 | -2.383752 | 0.01713716 | 0.184264055 |
| *STOML2* | 0.2003 | 0.084062 | 2.382768 | 0.017183 | 0.184354872 |
| *ABLIM2* | -1.013241 | 0.425289 | -2.382475 | 0.01719671 | 0.184354872 |
| *GTF3A* | 0.210459 | 0.08834 | 2.38238 | 0.01720114 | 0.184354872 |
| *MAU2* | -0.400264 | 0.168022 | -2.382212 | 0.01720899 | 0.184354872 |
| *NEK11* | -1.727183 | 0.725323 | -2.381262 | 0.01725346 | NA |
| *TXNIP* | -0.298774 | 0.125521 | -2.380277 | 0.01729962 | 0.185155297 |
| *TMEM185A* | -0.421876 | 0.177357 | -2.378688 | 0.01737438 | 0.185635406 |
| *WNT16* | 1.038403 | 0.436552 | 2.378645 | 0.01737639 | 0.185635406 |
| *UBE2S* | 0.394132 | 0.165733 | 2.378115 | 0.0174014 | 0.185642642 |
| *NPHP3* | -0.460487 | 0.193649 | -2.377954 | 0.01740898 | 0.185642642 |
| *PSMD1* | 0.218072 | 0.091742 | 2.377001 | 0.01745404 | 0.185925784 |
| *UBA52* | -0.116282 | 0.048928 | -2.376587 | 0.01747362 | 0.185925784 |
| *SNRPF* | 0.201558 | 0.084817 | 2.376379 | 0.01748348 | 0.185925784 |
| *ST13* | -0.089024 | 0.037485 | -2.374934 | 0.01755207 | 0.186324982 |
| *SEC61B* | -0.231021 | 0.097288 | -2.374597 | 0.01756813 | 0.186324982 |
| *GSTM4* | -0.762896 | 0.321277 | -2.374577 | 0.01756907 | 0.186324982 |
| *MPG* | -0.249991 | 0.105354 | -2.372854 | 0.01765123 | 0.187025885 |
| *CCT2* | 0.172318 | 0.072631 | 2.372515 | 0.01766747 | 0.187027569 |
| *DUSP14* | 0.442962 | 0.186814 | 2.371144 | 0.01773313 | 0.18755198 |
| *C9orf72* | -0.407789 | 0.17205 | -2.370181 | 0.01777939 | 0.187870537 |
| *HDHD5* | 0.299938 | 0.126575 | 2.369646 | 0.01780511 | 0.187971513 |
| *DNAJA1* | 0.176513 | 0.074546 | 2.367828 | 0.01789285 | 0.188527372 |
| *PLEKHO1* | 0.304713 | 0.128703 | 2.367564 | 0.01790561 | 0.188527372 |
| *TCIRG1* | -0.393047 | 0.166014 | -2.367548 | 0.01790637 | 0.188527372 |
| *ZFAND2B* | -0.229896 | 0.097128 | -2.366941 | 0.01793578 | 0.188666242 |
| *HDAC6* | -0.397038 | 0.167782 | -2.366385 | 0.01796278 | 0.188779525 |
| *CPLANE1* | -0.544371 | 0.230112 | -2.365684 | 0.01799678 | 0.188901052 |
| *RPL32* | -0.09443 | 0.03992 | -2.365478 | 0.01800681 | 0.188901052 |
| *APOBR* | -1.474011 | 0.623613 | -2.363663 | 0.01809525 | NA |

| | | | | | |
|---|---|---|---|---|---|
| *ZEB1* | -0.334132 | 0.141575 | -2.360106 | 0.01826971 | 0.191367545 |
| *SYK* | -0.278706 | 0.118096 | -2.360002 | 0.01827483 | 0.191367545 |
| *UBAP2* | 0.300966 | 0.127601 | 2.358648 | 0.01834165 | 0.191894561 |
| *HHIP.AS1* | -1.354319 | 0.574416 | -2.35773 | 0.01838706 | 0.192093207 |
| *RIOK1* | 0.290508 | 0.123222 | 2.357597 | 0.01839366 | 0.192093207 |
| *TAF2* | 0.382798 | 0.162426 | 2.356757 | 0.01843531 | 0.192254499 |
| *GNAI3* | -0.21341 | 0.090561 | -2.356525 | 0.0184468 | 0.192254499 |
| *ATF4* | 0.22436 | 0.095217 | 2.356286 | 0.01845868 | 0.192254499 |
| *WNT2B* | -0.98257 | 0.417079 | -2.355837 | 0.01848101 | 0.192314952 |
| *KMT2A* | -0.232917 | 0.098929 | -2.354392 | 0.01855302 | 0.192891698 |
| *ZC3H18* | 0.260627 | 0.110736 | 2.353589 | 0.01859314 | 0.193017818 |
| *ENSG00000237773* | -0.67056 | 0.284922 | -2.353486 | 0.01859833 | 0.193017818 |
| *BAZ2A* | -0.242819 | 0.103217 | -2.352516 | 0.01864687 | 0.193349104 |
| *SERPINF2* | -1.40667 | 0.598216 | -2.35144 | 0.0187009 | 0.193736661 |
| *SNRPB2* | 0.153631 | 0.065352 | 2.350812 | 0.0187325 | 0.193891416 |
| *MCAT* | 0.342835 | 0.145925 | 2.349395 | 0.01880396 | 0.194457983 |
| *CIAPIN1* | 0.314601 | 0.133943 | 2.348773 | 0.01883537 | 0.19449766 |
| *HAX1* | 0.166378 | 0.070839 | 2.348657 | 0.01884123 | 0.19449766 |
| *UBR2* | -0.296935 | 0.126477 | -2.347744 | 0.0188875 | 0.194619775 |
| *U2AF2* | 0.248228 | 0.105744 | 2.347443 | 0.01890275 | 0.194619775 |
| *SP3* | -0.249459 | 0.106269 | -2.347434 | 0.01890324 | 0.194619775 |
| *ZCRB1* | 0.183282 | 0.078159 | 2.345005 | 0.01902685 | 0.195709301 |
| *ANKRD49* | -0.263596 | 0.112422 | -2.344694 | 0.01904271 | 0.195709301 |
| *VKORC1L1* | 0.407556 | 0.173932 | 2.343188 | 0.01911973 | 0.196327379 |
| *GBP7* | 0.808616 | 0.345465 | 2.340661 | 0.01924963 | 0.197486907 |
| *RPLP2* | -0.120131 | 0.051339 | -2.339973 | 0.01928512 | 0.197592415 |
| *ARAP2* | -0.438983 | 0.187615 | -2.339804 | 0.01929388 | 0.197592415 |
| *ABT1* | 0.219052 | 0.093663 | 2.338722 | 0.01934983 | 0.197802177 |
| *ZNF800* | -0.321501 | 0.13747 | -2.3387 | 0.01935099 | 0.197802177 |
| *SUV39H2* | 0.509764 | 0.217995 | 2.338422 | 0.01936537 | 0.197802177 |
| *CNBP* | 0.190534 | 0.081502 | 2.337788 | 0.01939827 | 0.197964448 |
| *ZNF436.AS1* | -1.05272 | 0.450537 | -2.336591 | 0.01946048 | 0.198175082 |
| *KAT2A* | 0.291251 | 0.124651 | 2.336536 | 0.0194633 | 0.198175082 |
| *PRKCI* | 0.661797 | 0.283254 | 2.336408 | 0.01947001 | 0.198175082 |
| *SLC2A1* | -0.401529 | 0.171961 | -2.335005 | 0.01954318 | 0.198745964 |
| *ZC3H7A* | -0.398127 | 0.170557 | -2.334271 | 0.01958153 | 0.198821436 |
| *PRR5* | 0.641803 | 0.275047 | 2.333433 | 0.01962544 | 0.198821436 |
| *SMURF2* | -0.480489 | 0.205916 | -2.333421 | 0.01962607 | 0.198821436 |
| *TLE3* | -0.293194 | 0.125652 | -2.333376 | 0.01962842 | 0.198821436 |
| *A1BG* | -0.300113 | 0.128625 | -2.33323 | 0.01963605 | 0.198821436 |
| *CENPP* | 0.591651 | 0.25373 | 2.33181 | 0.0197107 | 0.1994037 |
| *TAF4B* | 0.455733 | 0.19553 | 2.330757 | 0.01976615 | 0.199790991 |
| *FTSJ1* | 0.37668 | 0.161712 | 2.329325 | 0.01984185 | 0.200381973 |
| *TEC* | -0.861996 | 0.370779 | -2.324823 | 0.02008141 | 0.20262544 |
| *MFN1* | -0.339536 | 0.14608 | -2.324307 | 0.02010905 | 0.202728534 |
| *LINC02482* | -0.741507 | 0.319279 | -2.32244 | 0.02020924 | 0.20338878 |
| *STX7* | -0.194077 | 0.083566 | -2.322435 | 0.02020951 | 0.20338878 |

| | | | | | |
|---|---|---|---|---|---|
| *ATIC* | 0.208453 | 0.089776 | 2.321915 | 0.02023749 | 0.20349435 |
| *MRPS26* | 0.167751 | 0.072275 | 2.321025 | 0.02028547 | 0.203800677 |
| *PAN3* | -0.249988 | 0.107733 | -2.320443 | 0.0203169 | 0.203926389 |
| *TSTD1* | -0.236449 | 0.101911 | -2.320145 | 0.02033304 | 0.203926389 |
| *SEPTIN2* | 0.245264 | 0.105807 | 2.318031 | 0.02044762 | 0.204898891 |
| *SHPRH* | -0.279449 | 0.120658 | -2.316045 | 0.0205558 | 0.205805683 |
| *XRN1* | -0.344454 | 0.148752 | -2.31563 | 0.02057846 | 0.205855421 |
| *ENSG00000246528* | 0.906306 | 0.391519 | 2.314845 | 0.02062142 | 0.206051357 |
| *PWP1* | 0.164165 | 0.070925 | 2.314625 | 0.02063347 | 0.206051357 |
| *CCT7* | 0.202133 | 0.087376 | 2.31337 | 0.02070233 | 0.206544816 |
| *LARP4* | 0.312383 | 0.135051 | 2.313077 | 0.02071839 | 0.206544816 |
| *ENSG00000245869* | 0.518105 | 0.224154 | 2.311382 | 0.02081179 | 0.207062728 |
| *IFI27L2* | 0.291864 | 0.126301 | 2.310864 | 0.02084035 | 0.207062728 |
| *C19orf66* | -0.220862 | 0.095585 | -2.310635 | 0.02085305 | 0.207062728 |
| *RBM5* | -0.218364 | 0.094508 | -2.310536 | 0.02085848 | 0.207062728 |
| *HNRNPA3* | 0.143052 | 0.061913 | 2.310521 | 0.02085934 | 0.207062728 |
| *RPL39* | -0.097779 | 0.042327 | -2.310097 | 0.02088276 | 0.207118559 |
| *PUF60* | 0.221689 | 0.096046 | 2.308159 | 0.02099028 | 0.208007552 |
| *AMZ1* | -1.098949 | 0.47655 | -2.306053 | 0.02110769 | 0.20899311 |
| *EXOSC2* | 0.279629 | 0.121292 | 2.305427 | 0.02114265 | 0.209161224 |
| *PAK1IP1* | 0.25827 | 0.11208 | 2.304339 | 0.02120361 | 0.209586017 |
| *NEMP1* | -0.543779 | 0.236082 | -2.303344 | 0.02125947 | 0.209959837 |
| *HNRNPR* | 0.167438 | 0.072732 | 2.302117 | 0.0213286 | 0.210463861 |
| *DHTKD1* | -0.335771 | 0.14593 | -2.30091 | 0.02139673 | 0.210835445 |
| *ANKRD37* | 0.418019 | 0.181684 | 2.300808 | 0.0214025 | 0.210835445 |
| *COA1* | 0.267213 | 0.116354 | 2.296549 | 0.02164451 | 0.21303913 |
| *BATF2* | 2.117648 | 0.922151 | 2.296423 | 0.02165168 | NA |
| *ATPSCKMT* | 0.721317 | 0.314469 | 2.293765 | 0.02180398 | 0.214427297 |
| *INTS14* | 0.317167 | 0.138325 | 2.292912 | 0.02185305 | 0.214692925 |
| *UQCRFS1* | 0.130364 | 0.056862 | 2.292655 | 0.0218679 | 0.214692925 |
| *KCNC3* | 0.519308 | 0.226638 | 2.29135 | 0.02194319 | 0.215016362 |
| *RPN2* | 0.24857 | 0.108511 | 2.290742 | 0.02197837 | 0.215016362 |
| *AGO1* | 0.284126 | 0.124044 | 2.290527 | 0.02199076 | 0.215016362 |
| *S100A2* | -0.903487 | 0.394447 | -2.290513 | 0.02199158 | 0.215016362 |
| *GARS* | 0.312102 | 0.13626 | 2.290484 | 0.02199325 | 0.215016362 |
| *METTL1* | 0.345933 | 0.151096 | 2.289486 | 0.02205112 | 0.215401145 |
| *ENSG00000261766* | 0.409234 | 0.178774 | 2.289107 | 0.02207315 | 0.215435442 |
| *ATF6B* | -0.188572 | 0.082392 | -2.288728 | 0.02209518 | 0.215438242 |
| *HSPA14* | 0.281405 | 0.122967 | 2.288465 | 0.02211048 | 0.215438242 |
| *WDR83OS* | 0.146016 | 0.063846 | 2.286992 | 0.02219627 | 0.216093184 |
| *ADGRB2* | -1.747053 | 0.764221 | -2.286056 | 0.02225096 | NA |
| *RPLP1* | -0.117225 | 0.051284 | -2.285802 | 0.02226587 | 0.216528142 |
| *SYNGR3* | 0.567706 | 0.248385 | 2.285592 | 0.02227817 | 0.216528142 |
| *LINC01560* | -0.514719 | 0.225259 | -2.285005 | 0.02231254 | 0.216681212 |
| *MRPL20* | 0.201032 | 0.088007 | 2.284257 | 0.02235642 | 0.216926259 |
| *COX16* | 0.236234 | 0.103453 | 2.283492 | 0.02240139 | 0.21718151 |
| *RANBP1* | 0.260753 | 0.114351 | 2.280292 | 0.02259035 | 0.218831087 |

| | | | | | |
|---|---|---|---|---|---|
| *TFAP4* | 0.305929 | 0.134244 | 2.278904 | 0.02267276 | 0.219446641 |
| *NHLRC3* | -0.242003 | 0.106228 | -2.278152 | 0.02271751 | 0.219697015 |
| *HGH1* | 0.677401 | 0.297627 | 2.276008 | 0.02284553 | 0.220751577 |
| *ZNF569* | -0.50554 | 0.222221 | -2.274946 | 0.02290917 | 0.221088952 |
| *LMO2* | -1.037236 | 0.45597 | -2.274791 | 0.02291845 | 0.221088952 |
| *UCK2* | 0.38455 | 0.16936 | 2.270607 | 0.02317079 | 0.223337995 |
| *VPS16* | -0.397952 | 0.175363 | -2.269303 | 0.0232499 | 0.223755087 |
| *ARF1* | 0.184938 | 0.081497 | 2.26926 | 0.02325253 | 0.223755087 |
| *P4HB* | 0.156988 | 0.069191 | 2.268915 | 0.02327349 | 0.223771722 |
| *MSL3* | -0.267493 | 0.117913 | -2.268564 | 0.02329484 | 0.223782361 |
| *FCRL5* | 0.541639 | 0.238798 | 2.268192 | 0.02331753 | 0.223782361 |
| *GPN1* | 0.257013 | 0.113325 | 2.26792 | 0.02333411 | 0.223782361 |
| *SPAG1* | -0.643006 | 0.283558 | -2.267634 | 0.02335154 | 0.223782361 |
| *ATP5PF* | 0.161732 | 0.071364 | 2.266286 | 0.02343388 | 0.224386631 |
| *SETD4* | -0.451138 | 0.199197 | -2.264782 | 0.02352607 | 0.225084097 |
| *SMIM1* | -0.533648 | 0.235879 | -2.262382 | 0.02367378 | 0.22631124 |
| *ERICH6.AS1* | -0.525076 | 0.23224 | -2.260921 | 0.02376412 | 0.22697951 |
| *MAEA* | 0.27134 | 0.120029 | 2.260621 | 0.02378271 | 0.22697951 |
| *VPS51* | -0.212468 | 0.094011 | -2.260017 | 0.02382019 | 0.227025388 |
| *RBMS1* | -0.378669 | 0.167559 | -2.259915 | 0.02382654 | 0.227025388 |
| *UBXN11* | -0.363969 | 0.161094 | -2.259362 | 0.02386088 | 0.227166466 |
| *ARFGAP3* | -0.274623 | 0.121597 | -2.258471 | 0.02391632 | 0.227508171 |
| *TBL1X* | -0.355594 | 0.157524 | -2.257405 | 0.02398279 | 0.227954092 |
| *SETD5* | -0.190115 | 0.084248 | -2.256623 | 0.02403164 | 0.228231896 |
| *LATS1* | -0.423573 | 0.187836 | -2.255014 | 0.02413247 | 0.22856092 |
| *SPRY1* | 0.833803 | 0.3698 | 2.254738 | 0.02414981 | 0.22856092 |
| *TRNAU1AP* | -0.213868 | 0.094854 | -2.254699 | 0.02415225 | 0.22856092 |
| *METTL2B* | 0.293668 | 0.13025 | 2.254649 | 0.02415536 | 0.22856092 |
| *CHCHD1* | 0.159051 | 0.070548 | 2.254503 | 0.02416451 | 0.22856092 |
| *B3GNTL1* | 0.629137 | 0.279584 | 2.250265 | 0.02443214 | 0.230681377 |
| *ZBTB37* | -0.463274 | 0.205877 | -2.250245 | 0.02443341 | 0.230681377 |
| *HACD3* | 0.351435 | 0.156207 | 2.249796 | 0.02446188 | 0.230681377 |
| *PPIF* | -0.242552 | 0.107815 | -2.2497 | 0.02446801 | 0.230681377 |
| *CMC2* | 0.247278 | 0.109933 | 2.249349 | 0.02449029 | 0.230704522 |
| *TMX2* | 0.211206 | 0.093973 | 2.24751 | 0.02460746 | 0.231478926 |
| *ZC3H12D* | -0.418567 | 0.186242 | -2.247434 | 0.02461229 | 0.231478926 |
| *ANKH* | -0.506139 | 0.225449 | -2.245027 | 0.02476641 | 0.232740205 |
| *BMP2K* | -0.306161 | 0.136446 | -2.243822 | 0.02484387 | 0.233279751 |
| *DIPK1A* | 0.535025 | 0.238524 | 2.243064 | 0.0248927 | 0.233549768 |
| *TUBG1* | 0.321228 | 0.143236 | 2.242648 | 0.02491952 | 0.233612958 |
| *CCDC12* | 0.154813 | 0.06909 | 2.24076 | 0.02504165 | 0.234020448 |
| *KRCC1* | -0.194777 | 0.086937 | -2.240422 | 0.02506353 | 0.234020448 |
| *REXO2* | 0.22851 | 0.101995 | 2.240404 | 0.0250647 | 0.234020448 |
| *ALG1* | 0.383865 | 0.171338 | 2.240395 | 0.02506526 | 0.234020448 |
| *RFX1* | -0.325976 | 0.14552 | -2.240068 | 0.02508652 | 0.234020448 |
| *IL27RA* | -0.30629 | 0.136746 | -2.239841 | 0.02510128 | 0.234020448 |
| *CHRNA6* | -1.897809 | 0.84731 | -2.239805 | 0.0251036 | NA |

| | | | | | |
|---|---|---|---|---|---|
| *MBIP* | 0.323236 | 0.144314 | 2.239802 | 0.02510379 | 0.234020448 |
| *ENSG00000272817* | 1.199593 | 0.536354 | 2.236567 | 0.02531466 | 0.235797247 |
| *NUTM2B.AS1* | -0.304908 | 0.136492 | -2.23389 | 0.02549029 | 0.237243209 |
| *PDS5B* | -0.285149 | 0.127673 | -2.233421 | 0.02552123 | 0.237341269 |
| *UBB* | 0.232854 | 0.104363 | 2.231197 | 0.0256681 | 0.238516463 |
| *HOXB4* | 1.341172 | 0.601997 | 2.22787 | 0.02588917 | NA |
| *PHTF2* | -0.361024 | 0.162056 | -2.227773 | 0.02589566 | 0.240439032 |
| *ANKLE1* | 0.514927 | 0.231275 | 2.226469 | 0.02598276 | 0.240961145 |
| *NANS* | 0.287671 | 0.129214 | 2.226312 | 0.02599332 | 0.240961145 |
| *DDX10* | 0.326491 | 0.146755 | 2.224734 | 0.0260991 | 0.241537348 |
| *PSMB4* | 0.160017 | 0.07193 | 2.224607 | 0.02610762 | 0.241537348 |
| *GBP1* | -0.524909 | 0.235972 | -2.224456 | 0.02611776 | 0.241537348 |
| *NELFA* | 0.271729 | 0.122249 | 2.222743 | 0.02623315 | 0.242087287 |
| *BUD31* | 0.138244 | 0.062196 | 2.222727 | 0.02623421 | 0.242087287 |
| *MVP* | -0.389928 | 0.175434 | -2.222646 | 0.02623965 | 0.242087287 |
| *ZBED2* | -0.887632 | 0.399521 | -2.221742 | 0.02630072 | 0.242458426 |
| *PSAP* | 0.204462 | 0.092043 | 2.221364 | 0.02632631 | 0.242502224 |
| *RPL5* | -0.127448 | 0.057385 | -2.220926 | 0.02635598 | 0.242583428 |
| *TSC22D3* | -0.359062 | 0.161697 | -2.220586 | 0.02637902 | 0.242603523 |
| *AFTPH* | -0.195719 | 0.088156 | -2.220157 | 0.02640814 | 0.242679525 |
| *LINC00342* | -0.699684 | 0.315271 | -2.21931 | 0.02646567 | 0.242921159 |
| *TMEM106A* | -0.348656 | 0.157112 | -2.219155 | 0.02647619 | 0.242921159 |
| *TSEN2* | 0.374558 | 0.16882 | 2.218691 | 0.02650772 | 0.243018786 |
| *ZFP36L2* | -0.431897 | 0.194718 | -2.218057 | 0.02655091 | 0.24322308 |
| *SECISBP2L* | -0.274209 | 0.123732 | -2.216147 | 0.02668141 | 0.244226183 |
| *CNTRL* | -0.20794 | 0.093853 | -2.215597 | 0.02671915 | 0.244379346 |
| *SPCS1* | 0.160844 | 0.072639 | 2.214286 | 0.02680914 | 0.24500988 |
| *KANSL1* | -0.269785 | 0.121856 | -2.21396 | 0.0268315 | 0.245021741 |
| *CNR2* | -0.290068 | 0.131045 | -2.2135 | 0.02686317 | 0.245104953 |
| *C1orf43* | 0.159648 | 0.072134 | 2.213216 | 0.02688275 | 0.245104953 |
| *ENSG00000224046* | -1.137362 | 0.513974 | -2.212879 | 0.02690601 | 0.245124899 |
| *IKZF2* | -0.797241 | 0.360466 | -2.211693 | 0.02698787 | 0.245569216 |
| *MAPKBP1* | -0.689233 | 0.311673 | -2.211399 | 0.02700819 | 0.245569216 |
| *EMB* | -0.238708 | 0.107963 | -2.211013 | 0.02703494 | 0.245569216 |
| *SLC27A1* | -1.063551 | 0.481038 | -2.210951 | 0.02703921 | 0.245569216 |
| *CDC42* | -0.147431 | 0.066706 | -2.210161 | 0.02709401 | 0.245874944 |
| *LINC00869* | -0.273548 | 0.123802 | -2.209554 | 0.02713613 | 0.245938035 |
| *AUH* | 0.397357 | 0.179851 | 2.209364 | 0.02714937 | 0.245938035 |
| *VBP1* | 0.169854 | 0.076897 | 2.208854 | 0.02718478 | 0.245938035 |
| *PIK3CB* | -0.571714 | 0.258832 | -2.208828 | 0.02718664 | 0.245938035 |
| *PPP1R12A* | -0.227428 | 0.102976 | -2.20854 | 0.02720666 | 0.245938035 |
| *PTMA* | 0.134754 | 0.061029 | 2.208033 | 0.02724199 | 0.246013974 |
| *IMP3* | 0.180984 | 0.081974 | 2.207812 | 0.02725735 | 0.246013974 |
| *PCID2* | 0.25363 | 0.114934 | 2.206742 | 0.02733209 | 0.246497307 |
| *ITGB2* | 0.528122 | 0.239454 | 2.205524 | 0.02741733 | 0.247074565 |
| *GART* | 0.233395 | 0.105925 | 2.203394 | 0.02756699 | 0.248230956 |
| *LINC02576* | -1.205051 | 0.547029 | -2.202902 | 0.02760166 | 0.248350861 |

| | | | | | |
|---|---|---|---|---|---|
| *EVI5* | -0.35157 | 0.159631 | -2.202391 | 0.02763767 | 0.248482742 |
| *TCP11L2* | -0.488931 | 0.222034 | -2.202057 | 0.02766129 | 0.248503028 |
| *MYBBP1A* | 0.342622 | 0.155643 | 2.201335 | 0.02771234 | 0.248717335 |
| *ZNF608* | -0.532241 | 0.241805 | -2.201115 | 0.0277279 | 0.248717335 |
| *WDR46* | 0.309205 | 0.140532 | 2.200241 | 0.0277898 | 0.249080572 |
| *NUDCD2* | 0.161948 | 0.073661 | 2.19856 | 0.02790924 | 0.249958489 |
| *POC5* | -0.493838 | 0.224774 | -2.19704 | 0.02801756 | 0.25073557 |
| *HNRNPF* | 0.25299 | 0.115173 | 2.196608 | 0.02804845 | 0.250819127 |
| *CSDE1* | -0.107733 | 0.049059 | -2.19598 | 0.02809341 | 0.250877601 |
| *HDAC2* | 0.191717 | 0.087308 | 2.195862 | 0.02810186 | 0.250877601 |
| *WDTC1* | 0.299072 | 0.136213 | 2.195613 | 0.02811968 | 0.250877601 |
| *ZC3H8* | 0.269344 | 0.122697 | 2.195189 | 0.02815003 | 0.250955933 |
| *BSDC1* | -0.260045 | 0.118549 | -2.193566 | 0.02826665 | 0.251802604 |
| *ANP32A* | 0.135105 | 0.061643 | 2.191735 | 0.02839868 | 0.252765344 |
| *ZNF117* | 0.587676 | 0.268166 | 2.191465 | 0.02841818 | 0.252765344 |
| *ATP5MF* | 0.157284 | 0.071809 | 2.190319 | 0.02850112 | 0.253309432 |
| *CAMKK2* | 0.412991 | 0.188648 | 2.189214 | 0.02858126 | 0.253827757 |
| *MINDY1* | -0.807206 | 0.368896 | -2.188165 | 0.0286576 | 0.254139259 |
| *C20orf27* | 0.213952 | 0.097782 | 2.188039 | 0.02866679 | 0.254139259 |
| *SLC12A2* | -0.299952 | 0.1371 | -2.187832 | 0.02868187 | 0.254139259 |
| *IFT22* | -0.308099 | 0.140844 | -2.187514 | 0.02870504 | 0.254151024 |
| *CD2AP* | 0.290992 | 0.133077 | 2.186636 | 0.02876909 | 0.254333799 |
| *MAPK7* | -0.574061 | 0.262565 | -2.186356 | 0.02878956 | 0.254333799 |
| *RPS18* | -0.121288 | 0.055478 | -2.18622 | 0.02879954 | 0.254333799 |
| *TRIB2* | -0.745784 | 0.341158 | -2.186034 | 0.02881313 | 0.254333799 |
| *CMC1* | 0.247242 | 0.113125 | 2.185561 | 0.02884777 | 0.25442738 |
| *RABGGTA* | -0.288538 | 0.132055 | -2.184992 | 0.02888942 | 0.25442738 |
| *LSR* | -0.524627 | 0.240133 | -2.184732 | 0.0289085 | 0.25442738 |
| *ABI1* | -0.184905 | 0.084637 | -2.184694 | 0.02891131 | 0.25442738 |
| *MARS* | 0.24425 | 0.111816 | 2.184397 | 0.02893308 | 0.25442738 |
| *ENSG00000230709* | -0.761461 | 0.348838 | -2.182846 | 0.02904719 | 0.255237907 |
| *ANAPC4* | -0.393495 | 0.180298 | -2.182469 | 0.02907496 | 0.255289147 |
| *USP7* | 0.265038 | 0.121507 | 2.181252 | 0.02916476 | 0.255884438 |
| *CDC16* | -0.278111 | 0.127524 | -2.180856 | 0.02919408 | 0.255948695 |
| *HIPK3* | -0.336827 | 0.154507 | -2.180014 | 0.0292564 | 0.256226575 |
| *ATF7IP2* | 0.28269 | 0.129684 | 2.179833 | 0.02926982 | 0.256226575 |
| *NAXD* | 0.380284 | 0.174547 | 2.178694 | 0.02935438 | 0.256773537 |
| *MED27* | 0.759661 | 0.348851 | 2.177612 | 0.02943494 | 0.257071444 |
| *SUN1* | -0.227757 | 0.104598 | -2.177453 | 0.02944676 | 0.257071444 |
| *HLA.DMB* | 0.254266 | 0.116784 | 2.177231 | 0.02946336 | 0.257071444 |
| *B3GAT3* | 0.210121 | 0.096526 | 2.176843 | 0.02949231 | 0.257071444 |
| *ATP2B1.AS1* | -0.365211 | 0.167778 | -2.176754 | 0.02949892 | 0.257071444 |
| *ENSG00000226571* | -0.675431 | 0.310425 | -2.175828 | 0.02956812 | 0.257317208 |
| *TXN* | 0.249086 | 0.114481 | 2.175785 | 0.02957135 | 0.257317208 |
| *PSMB2* | 0.23429 | 0.107716 | 2.175065 | 0.02962526 | 0.257593642 |
| *AKNA* | -0.260447 | 0.119768 | -2.174595 | 0.02966049 | 0.257703328 |
| *FILIP1L* | -0.903152 | 0.415375 | -2.174306 | 0.02968218 | 0.257703328 |

| | | | | | |
|---|---|---|---|---|---|
| *LINC01869* | -0.758293 | 0.348804 | -2.173979 | 0.02970674 | 0.257724284 |
| *PRXL2C* | -0.395063 | 0.181785 | -2.17324 | 0.02976229 | 0.25801374 |
| *RPS15A* | -0.117971 | 0.054299 | -2.172609 | 0.02980978 | 0.25809232 |
| *CYB5R3* | -0.215388 | 0.099142 | -2.17253 | 0.02981572 | 0.25809232 |
| *FBXL6* | 0.688825 | 0.317171 | 2.171779 | 0.02987237 | 0.258246164 |
| *FAM107B* | 0.199096 | 0.09168 | 2.171639 | 0.02988293 | 0.258246164 |
| *ECE1* | 0.347453 | 0.160013 | 2.171411 | 0.02990008 | 0.258246164 |
| *BCYRN1* | 1.161531 | 0.535286 | 2.169926 | 0.03001244 | NA |
| *SLC25A43* | -0.39161 | 0.180492 | -2.169688 | 0.03003046 | 0.259179831 |
| *CXXC5* | -0.253755 | 0.117019 | -2.168484 | 0.0301219 | 0.259593086 |
| *DERL2* | 0.198881 | 0.091715 | 2.16847 | 0.03012297 | 0.259593086 |
| *TMCO1* | 0.155883 | 0.071941 | 2.16683 | 0.03024783 | 0.260476086 |
| *FYN* | -0.42773 | 0.197497 | -2.16575 | 0.03033029 | 0.260925989 |
| *ENSG00000233184* | -0.524109 | 0.24202 | -2.165559 | 0.03034493 | 0.260925989 |
| *ENSG00000258572* | -0.876493 | 0.404909 | -2.164669 | 0.03041305 | 0.261043504 |
| *TNFAIP8L2* | -0.411845 | 0.190261 | -2.164633 | 0.0304158 | 0.261043504 |
| *MPI* | 0.35351 | 0.163322 | 2.164501 | 0.03042591 | 0.261043504 |
| *MRPL54* | 0.188382 | 0.087099 | 2.162833 | 0.03055399 | 0.261949272 |
| *EIF2S3* | -0.214293 | 0.099096 | -2.162484 | 0.03058091 | 0.261986939 |
| *C12orf75* | -0.533373 | 0.246824 | -2.160948 | 0.03069937 | 0.262808321 |
| *NARF* | -0.199436 | 0.09234 | -2.159791 | 0.03078887 | 0.263380683 |
| *SGCB* | -0.675739 | 0.313248 | -2.157206 | 0.03098962 | 0.264582373 |
| *ATP11A* | -0.526429 | 0.244047 | -2.157085 | 0.03099903 | 0.264582373 |
| *EMP3* | -0.240663 | 0.111572 | -2.157026 | 0.03100364 | 0.264582373 |
| *MRPL36* | 0.19653 | 0.091133 | 2.156521 | 0.03104301 | 0.264582373 |
| *PHGDH* | 0.685868 | 0.318044 | 2.15652 | 0.03104306 | 0.264582373 |
| *POR* | 0.304364 | 0.141186 | 2.155769 | 0.03110169 | 0.264725603 |
| *FBXO10* | 0.513092 | 0.238014 | 2.155722 | 0.03110537 | 0.264725603 |
| *KIFC2* | -0.790816 | 0.366964 | -2.15502 | 0.03116024 | 0.264961808 |
| *CAT* | -0.354499 | 0.164535 | -2.154554 | 0.03119676 | 0.264961808 |
| *TRNT1* | 0.251344 | 0.11666 | 2.154494 | 0.03120145 | 0.264961808 |
| *IGBP1* | -0.168505 | 0.078235 | -2.153837 | 0.03125295 | 0.265061664 |
| *FCRL4* | 0.630775 | 0.292884 | 2.15367 | 0.03126604 | 0.265061664 |
| *TRADD* | -0.267085 | 0.124037 | -2.153265 | 0.03129782 | 0.265061664 |
| *CERS6* | 0.447534 | 0.207911 | 2.152526 | 0.03135595 | 0.265061664 |
| *TCEAL8* | -0.244487 | 0.113582 | -2.152518 | 0.03135657 | 0.265061664 |
| *PUM3* | 0.207416 | 0.096361 | 2.152501 | 0.03135794 | 0.265061664 |
| *GNAI2* | -0.228215 | 0.106032 | -2.152313 | 0.03137269 | 0.265061664 |
| *DAPL1* | 0.441408 | 0.20516 | 2.151535 | 0.03143403 | 0.26538714 |
| *PPA1* | 0.240366 | 0.11181 | 2.149764 | 0.0315739 | 0.266374711 |
| *NUSAP1* | -0.657608 | 0.306015 | -2.148945 | 0.03163879 | 0.266728728 |
| *RB1CC1* | -0.233639 | 0.108777 | -2.147871 | 0.03172402 | 0.267110883 |
| *AHSA1* | 0.229545 | 0.106875 | 2.147795 | 0.03173004 | 0.267110883 |
| *DESI1* | 0.359227 | 0.167411 | 2.145778 | 0.03189072 | 0.268269395 |
| *ERV3.1* | -0.176297 | 0.082201 | -2.144697 | 0.03197709 | 0.268717744 |
| *EML6* | -0.746405 | 0.348071 | -2.144402 | 0.03200072 | 0.268717744 |
| *SLC6A16* | -0.612644 | 0.285715 | -2.144244 | 0.03201331 | 0.268717744 |

| | | | | | |
|---|---|---|---|---|---|
| *CENPM* | 0.356158 | 0.166217 | 2.142734 | 0.0321345 | 0.269467595 |
| *PEX16* | -0.198417 | 0.092608 | -2.142554 | 0.03214896 | 0.269467595 |
| *TRIM23* | -0.528603 | 0.246788 | -2.141933 | 0.0321989 | 0.269691824 |
| *ISOC2* | 0.265437 | 0.12405 | 2.139762 | 0.03237404 | 0.270963698 |
| *U2SURP* | 0.141715 | 0.066269 | 2.138485 | 0.03247737 | 0.271473099 |
| *CDKL5* | -1.284388 | 0.600621 | -2.138434 | 0.03248157 | 0.271473099 |
| *ZNF490* | -1.590476 | 0.743841 | -2.138192 | 0.03250113 | NA |
| *RPL15* | -0.113445 | 0.053086 | -2.136998 | 0.03259818 | 0.272108322 |
| *MT.ND4L* | -0.401476 | 0.187894 | -2.136719 | 0.03262084 | 0.272108322 |
| *SMCHD1* | -0.184773 | 0.086485 | -2.136481 | 0.03264026 | 0.272108322 |
| *GPM6B* | -0.340289 | 0.159315 | -2.135952 | 0.03268333 | 0.272108322 |
| *TMEM107* | -0.355444 | 0.166416 | -2.135872 | 0.03268983 | 0.272108322 |
| *ATP5MD* | 0.163894 | 0.076738 | 2.135753 | 0.03269954 | 0.272108322 |
| *RELCH* | -0.558069 | 0.261331 | -2.135486 | 0.0327213 | 0.272108322 |
| *RPS25* | -0.104715 | 0.049045 | -2.135071 | 0.03275525 | 0.272196141 |
| *KARS* | 0.132082 | 0.061894 | 2.133998 | 0.03284292 | 0.272729846 |
| *PTPRC* | -0.181024 | 0.084854 | -2.133369 | 0.03289448 | 0.272963179 |
| *ITSN2* | -0.174813 | 0.081954 | -2.133055 | 0.03292022 | 0.272982079 |
| *ELOC* | 0.198345 | 0.093004 | 2.132659 | 0.03295272 | 0.273000383 |
| *ENSG00000225342* | -0.654405 | 0.306878 | -2.132456 | 0.03296936 | 0.273000383 |
| *MRPS15* | 0.174267 | 0.081734 | 2.132136 | 0.03299565 | 0.273023708 |
| *POLD2* | 0.21736 | 0.102024 | 2.130487 | 0.03313145 | 0.273862516 |
| *RAB27A* | 0.317947 | 0.149248 | 2.130333 | 0.0331441 | 0.273862516 |
| *ENSG00000245466* | -1.78772 | 0.839437 | -2.129665 | 0.03319925 | NA |
| *MGAT1* | 0.238846 | 0.112164 | 2.129445 | 0.0332175 | 0.274274232 |
| *NUS1* | 0.203884 | 0.095815 | 2.127883 | 0.03334681 | 0.275146687 |
| *MAP2K1* | -0.257083 | 0.120877 | -2.126806 | 0.03343617 | 0.275688416 |
| *ZNF596* | 0.841674 | 0.3959 | 2.125979 | 0.03350497 | 0.275922778 |
| *UBA7* | -0.358376 | 0.168577 | -2.125895 | 0.03351202 | 0.275922778 |
| *MRPS33* | 0.247464 | 0.116497 | 2.12421 | 0.03365262 | 0.276884383 |
| *ELMOD2* | -0.47408 | 0.223247 | -2.12357 | 0.03370613 | 0.277128689 |
| *MRPL37* | 0.175167 | 0.082508 | 2.123043 | 0.03375027 | 0.277135412 |
| *SEC61G* | 0.186906 | 0.088039 | 2.122991 | 0.03375459 | 0.277135412 |
| *DCLRE1C* | 0.18828 | 0.088713 | 2.122355 | 0.03380791 | 0.277278187 |
| *RGS1* | -0.514376 | 0.242377 | -2.122216 | 0.03381965 | 0.277278187 |
| *RHOC* | -0.46452 | 0.21897 | -2.121388 | 0.03388916 | 0.277652468 |
| *PSTPIP1* | -0.548097 | 0.258456 | -2.120662 | 0.03395024 | 0.277942994 |
| *RNASEK* | -0.345377 | 0.162904 | -2.120125 | 0.0339955 | 0.277942994 |
| *SNRNP48* | -0.322679 | 0.152199 | -2.120116 | 0.03399629 | 0.277942994 |
| *CD164* | 0.143144 | 0.067578 | 2.118192 | 0.03415877 | 0.279075264 |
| *SLC20A1* | 0.282293 | 0.133296 | 2.117794 | 0.03419255 | 0.279155173 |
| *RETREG1* | -0.831608 | 0.39278 | -2.117235 | 0.03423988 | 0.279164484 |
| *DLGAP1.AS1* | -0.270584 | 0.127802 | -2.117214 | 0.03424168 | 0.279164484 |
| *ENSG00000227486* | -0.355602 | 0.168024 | -2.116374 | 0.034313 | 0.279550015 |
| *ZNF677* | -0.290054 | 0.137087 | -2.115843 | 0.03435818 | 0.279722266 |
| *TEPP* | -1.300434 | 0.614754 | -2.115373 | 0.03439821 | 0.279852282 |
| *ENSG00000265206* | -0.248588 | 0.117568 | -2.114428 | 0.03447872 | 0.280311246 |

| | | | | | |
|---|---|---|---|---|---|
| *EXOSC7* | 0.258212 | 0.122159 | 2.113738 | 0.03453761 | 0.280593993 |
| *SIGLEC14* | 0.790259 | 0.373947 | 2.113295 | 0.03457553 | 0.280706001 |
| *GPATCH1* | -0.409704 | 0.193944 | -2.112489 | 0.0346445 | 0.280986371 |
| *ZNF547* | -1.015811 | 0.480937 | -2.112149 | 0.0346737 | 0.280986371 |
| *SLC35A4* | 0.219282 | 0.103824 | 2.112046 | 0.03468252 | 0.280986371 |
| *KLRD1* | 1.427827 | 0.676049 | 2.112018 | 0.03468487 | NA |
| *CEP70* | 0.709156 | 0.335854 | 2.111501 | 0.03472927 | 0.2811693 |
| *FEM1A* | 0.248211 | 0.117612 | 2.110427 | 0.03482157 | 0.281720577 |
| *SNX12* | 0.229204 | 0.108635 | 2.109859 | 0.03487049 | 0.281920242 |
| *TNPO3* | 0.266106 | 0.126156 | 2.109352 | 0.03491421 | 0.282077726 |
| *ENSG00000187186* | -0.831156 | 0.394153 | -2.108711 | 0.03496952 | 0.282328524 |
| *SLC2A3* | -0.449679 | 0.213413 | -2.107085 | 0.03511024 | 0.283268054 |
| *CYSLTR1* | -0.477315 | 0.226735 | -2.10517 | 0.0352765 | 0.284287142 |
| *DLGAP4* | -0.239325 | 0.11369 | -2.105068 | 0.03528543 | 0.284287142 |
| *MAATS1* | -1.78493 | 0.848006 | -2.104857 | 0.03530378 | NA |
| *FAM227A* | 1.636851 | 0.77766 | 2.104841 | 0.03530512 | NA |
| *FLII* | -0.252362 | 0.119951 | -2.103883 | 0.03538867 | 0.284747111 |
| *HEATR6* | -0.444721 | 0.211387 | -2.103817 | 0.03539443 | 0.284747111 |
| *MRPS34* | 0.165696 | 0.078769 | 2.10357 | 0.03541594 | 0.284747111 |
| *PIK3CG* | 0.433686 | 0.206287 | 2.102347 | 0.0355229 | 0.28540985 |
| *ZNF736* | -0.459192 | 0.218452 | -2.102025 | 0.03555104 | 0.285438816 |
| *EIF1* | 0.158352 | 0.07535 | 2.101542 | 0.03559345 | 0.285582208 |
| *CYTH1* | 0.186672 | 0.088911 | 2.099549 | 0.03576855 | 0.286789327 |
| *EIF3J.DT* | -0.245593 | 0.117009 | -2.098925 | 0.03582349 | 0.287032012 |
| *CYC1* | 0.199775 | 0.095194 | 2.098616 | 0.03585074 | 0.287052637 |
| *LIMS2* | -1.034836 | 0.493178 | -2.098303 | 0.0358784 | 0.287076517 |
| *ENSG00000253636* | 1.762343 | 0.840008 | 2.098008 | 0.03590444 | NA |
| *ENSG00000235501* | -1.536639 | 0.732476 | -2.09787 | 0.03591661 | NA |
| *ESR2* | -0.484351 | 0.230907 | -2.097602 | 0.03594028 | 0.287372596 |
| *OAS2* | 0.346788 | 0.165348 | 2.097325 | 0.0359648 | 0.287372596 |
| *ADRM1* | 0.20277 | 0.096776 | 2.095261 | 0.03614779 | 0.288391581 |
| *CCS* | -0.242608 | 0.115797 | -2.095116 | 0.03616071 | 0.288391581 |
| *NETO2* | 0.641932 | 0.306407 | 2.095029 | 0.03616845 | 0.288391581 |
| *MRPS10* | 0.184697 | 0.088188 | 2.094344 | 0.03622931 | 0.288391581 |
| *MTAP* | 0.298639 | 0.142605 | 2.094168 | 0.03624504 | 0.288391581 |
| *TTYH3* | 0.477712 | 0.228118 | 2.094146 | 0.03624698 | 0.288391581 |
| *ENSG00000267519* | -0.329438 | 0.15733 | -2.093934 | 0.03626585 | 0.288391581 |
| *CCDC146* | -1.327642 | 0.634125 | -2.093658 | 0.03629047 | NA |
| *TIMM21* | 0.302323 | 0.144406 | 2.093565 | 0.03629871 | 0.288455745 |
| *FCHSD2* | -0.315206 | 0.150582 | -2.093253 | 0.03632662 | 0.288480468 |
| *PSMB6* | 0.164245 | 0.078537 | 2.09132 | 0.03649939 | 0.289654812 |
| *CD52* | -0.244582 | 0.11703 | -2.089911 | 0.03662577 | 0.290073152 |
| *IGLC2* | -0.300523 | 0.143816 | -2.08964 | 0.03665019 | 0.290073152 |
| *TPRG1L* | 0.376761 | 0.180299 | 2.089639 | 0.03665023 | 0.290073152 |
| *MAP3K20* | -0.436602 | 0.208938 | -2.089621 | 0.03665184 | 0.290073152 |
| *SDCBP* | 0.304631 | 0.145831 | 2.088934 | 0.03671364 | 0.290313244 |
| *C8orf37* | -0.833882 | 0.399229 | -2.088729 | 0.03673209 | 0.290313244 |

| | | | | | |
|---|---|---|---|---|---|
| *SMYD2* | -0.311964 | 0.149446 | -2.087468 | 0.03684586 | 0.29101475 |
| *ABCF1* | 0.177861 | 0.085232 | 2.086775 | 0.03690849 | 0.291311619 |
| *CPEB3* | 0.783095 | 0.375423 | 2.0859 | 0.03698769 | 0.29156306 |
| *POLR1C* | 0.230076 | 0.110302 | 2.085869 | 0.03699047 | 0.29156306 |
| *ENSG00000257275* | -0.769386 | 0.369032 | -2.084877 | 0.03708049 | 0.291989052 |
| *VPS9D1.AS1* | 0.51586 | 0.247453 | 2.084678 | 0.03709856 | 0.291989052 |
| *NDUFC2* | 0.208123 | 0.099846 | 2.084444 | 0.03711981 | 0.291989052 |
| *RPS4X* | -0.217917 | 0.104567 | -2.084 | 0.03716019 | 0.291996676 |
| *ZCCHC18* | -0.655902 | 0.314782 | -2.083669 | 0.03719027 | 0.291996676 |
| *RRP8* | 0.244277 | 0.117237 | 2.083605 | 0.03719607 | 0.291996676 |
| *ADD3* | -0.233034 | 0.111863 | -2.083203 | 0.0372327 | 0.292033373 |
| *RREB1* | -0.299261 | 0.143668 | -2.083003 | 0.03725095 | 0.292033373 |
| *ITFG1* | 0.254654 | 0.122308 | 2.082069 | 0.03733613 | 0.292488623 |
| *FAM136A* | 0.174119 | 0.083642 | 2.081722 | 0.03736788 | 0.292488623 |
| *KLF13* | 0.242432 | 0.116468 | 2.081541 | 0.03738444 | 0.292488623 |
| *DRAM2* | -0.254997 | 0.122585 | -2.080163 | 0.03751055 | 0.29327803 |
| *XPNPEP1* | -0.334403 | 0.160822 | -2.079338 | 0.0375863 | 0.293672906 |
| *RHBDD3* | 0.397229 | 0.191095 | 2.078703 | 0.03764464 | 0.293739889 |
| *BTBD19* | 0.728173 | 0.350303 | 2.078695 | 0.03764537 | 0.293739889 |
| *STAG3* | -0.402985 | 0.193912 | -2.078181 | 0.03769272 | 0.293912256 |
| *FOXO1* | -0.317185 | 0.152656 | -2.077771 | 0.03773042 | 0.294009193 |
| *PSEN1* | -0.324425 | 0.156365 | -2.074796 | 0.0380054 | 0.295953724 |
| *PRICKLE1* | 0.597339 | 0.288048 | 2.073746 | 0.03810287 | 0.296514257 |
| *CNFN* | -0.518492 | 0.250135 | -2.072845 | 0.03818672 | 0.296968079 |
| *PSMC2* | 0.231299 | 0.111604 | 2.072499 | 0.03821892 | 0.297019982 |
| *AMN1* | -0.378217 | 0.182644 | -2.070792 | 0.03837824 | 0.297911624 |
| *ST6GALNAC4* | -0.216503 | 0.104555 | -2.070721 | 0.03838487 | 0.297911624 |
| *RILPL2* | 0.324118 | 0.156586 | 2.069906 | 0.03846113 | 0.298155314 |
| *CDK5RAP3* | -0.225902 | 0.109143 | -2.069776 | 0.03847337 | 0.298155314 |
| *MFSD2A* | 0.739553 | 0.357347 | 2.069564 | 0.03849315 | 0.298155314 |
| *AAK1* | 0.574539 | 0.277759 | 2.068478 | 0.0385951 | 0.298746118 |
| *ATN1* | -0.347129 | 0.167858 | -2.067992 | 0.03864081 | 0.298901033 |
| *PLEKHF2* | -0.230776 | 0.11161 | -2.067701 | 0.03866818 | 0.298913993 |
| *NPLOC4* | 0.286458 | 0.138567 | 2.067282 | 0.03870756 | 0.299019776 |
| *ENSG00000255240* | -0.805549 | 0.389852 | -2.066293 | 0.03880078 | 0.299540963 |
| *AP2A1* | -0.259881 | 0.125809 | -2.065675 | 0.03885922 | 0.299793175 |
| *HDAC10* | -0.235521 | 0.114147 | -2.06331 | 0.03908318 | 0.301321203 |
| *SERTAD2* | -0.302841 | 0.146853 | -2.06221 | 0.03918772 | 0.301890097 |
| *HCCS* | 0.234386 | 0.11367 | 2.061988 | 0.03920887 | 0.301890097 |
| *STKLD1* | -2.523992 | 1.224232 | -2.061695 | 0.0392368 | NA |
| *C14orf28* | -0.458231 | 0.222379 | -2.060592 | 0.03934202 | 0.302714975 |
| *CDCA8* | -0.997516 | 0.484189 | -2.060177 | 0.0393816 | 0.302760037 |
| *CRIPT* | 0.15995 | 0.077646 | 2.059986 | 0.03939992 | 0.302760037 |
| *AKT1S1* | 0.291547 | 0.141553 | 2.059634 | 0.03943359 | 0.302818724 |
| *KHDC4* | -0.187486 | 0.091105 | -2.057925 | 0.03959732 | 0.303603185 |
| *LPP.AS2* | -0.7828 | 0.380398 | -2.057846 | 0.03960493 | 0.303603185 |
| *ENSG00000228463* | 0.367942 | 0.178831 | 2.05748 | 0.03964005 | 0.303603185 |

| | | | | | |
|---|---|---|---|---|---|
| *NCBP2* | 0.148302 | 0.07208 | 2.05745 | 0.03964296 | 0.303603185 |
| *FAM120AOS* | -0.234529 | 0.114004 | -2.057208 | 0.03966622 | 0.303603185 |
| *TINCR* | -1.549777 | 0.753878 | -2.055741 | 0.03980746 | NA |
| *DUS1L* | 0.26395 | 0.128404 | 2.055619 | 0.03981925 | 0.304574029 |
| *QTRT1* | 0.195839 | 0.095287 | 2.055257 | 0.03985416 | 0.304640802 |
| *ANAPC5* | 0.137019 | 0.066703 | 2.054181 | 0.0399582 | 0.305235525 |
| *DDX18* | 0.117058 | 0.057028 | 2.052644 | 0.04010712 | 0.306172042 |
| *CXorf21* | -0.302714 | 0.147497 | -2.052333 | 0.04013736 | 0.30620197 |
| *GVQW3* | -0.735838 | 0.358634 | -2.051779 | 0.04019115 | 0.306213996 |
| *RPUSD2* | 0.381338 | 0.185858 | 2.051775 | 0.04019157 | 0.306213996 |
| *ASAH1* | 0.204231 | 0.09965 | 2.049498 | 0.04041346 | 0.307702995 |
| *CTSC* | 0.374477 | 0.182798 | 2.048584 | 0.04050282 | 0.308132033 |
| *HIST3H2A* | 0.739747 | 0.361137 | 2.04838 | 0.04052278 | 0.308132033 |
| *RABEP2* | -0.243528 | 0.118944 | -2.047421 | 0.04061679 | 0.308645157 |
| *ZNF688* | -0.305086 | 0.149051 | -2.04686 | 0.0406718 | 0.308798946 |
| *PRDM8* | -0.654865 | 0.320012 | -2.046375 | 0.04071947 | 0.308798946 |
| *NBPF19* | -0.48839 | 0.23868 | -2.046209 | 0.04073579 | 0.308798946 |
| *STX17* | -0.282446 | 0.138069 | -2.045684 | 0.04078751 | 0.308798946 |
| *ICAM2* | 0.407455 | 0.199196 | 2.045501 | 0.04080546 | 0.308798946 |
| *EXD3* | -0.524482 | 0.256408 | -2.0455 | 0.04080562 | 0.308798946 |
| *RFX5* | 0.217665 | 0.106421 | 2.045325 | 0.04082283 | 0.308798946 |
| *RGL1* | 1.035414 | 0.506389 | 2.044702 | 0.04088425 | 0.309062635 |
| *DEGS2* | -1.25745 | 0.6152 | -2.043969 | 0.04095663 | 0.309408698 |
| *ATP6V1D* | 0.233916 | 0.114462 | 2.043612 | 0.04099194 | 0.30947426 |
| *SIGMAR1* | 0.286911 | 0.140428 | 2.043123 | 0.04104028 | 0.30947426 |
| *EARS2* | 0.442844 | 0.216754 | 2.043074 | 0.04104511 | 0.30947426 |
| *STRAP* | 0.150408 | 0.073636 | 2.042581 | 0.04109397 | 0.30964202 |
| *CSNK1E* | -0.307375 | 0.15054 | -2.041811 | 0.04117031 | 0.309936876 |
| *RPS12* | -0.122589 | 0.060044 | -2.041649 | 0.04118639 | 0.309936876 |
| *MLXIPL* | 1.741798 | 0.853216 | 2.041451 | 0.04120599 | NA |
| *SMG9* | -0.31441 | 0.154054 | -2.04091 | 0.0412598 | 0.310042726 |
| *DOK1* | -0.343103 | 0.168117 | -2.040861 | 0.04126467 | 0.310042726 |
| *ENSG00000272182* | 0.803545 | 0.393759 | 2.040703 | 0.0412804 | 0.310042726 |
| *FGL2* | -1.384502 | 0.678645 | -2.040099 | 0.04134042 | NA |
| *TMF1* | -0.221687 | 0.108688 | -2.039655 | 0.04138475 | 0.310625914 |
| *SCAI* | -0.31883 | 0.156345 | -2.03928 | 0.04142209 | 0.310705727 |
| *NT5C3A* | -0.222859 | 0.109381 | -2.037468 | 0.04160321 | 0.31186327 |
| *DAAM1* | -0.402914 | 0.197834 | -2.036621 | 0.04168807 | 0.312282963 |
| *MMACHC* | 0.689199 | 0.338444 | 2.036373 | 0.04171289 | 0.312282963 |
| *HMGN2* | -0.113368 | 0.055683 | -2.03595 | 0.04175538 | 0.312335172 |
| *KIF5C* | -1.13762 | 0.558816 | -2.035769 | 0.04177355 | 0.312335172 |
| *PSMB1* | 0.161986 | 0.07959 | 2.035272 | 0.04182351 | 0.312507843 |
| *CDK2AP1* | 0.670507 | 0.329497 | 2.034941 | 0.04185686 | 0.312556281 |
| *TMED3* | 0.208739 | 0.10261 | 2.034297 | 0.04192163 | 0.312747634 |
| *MANBA* | -0.255948 | 0.125825 | -2.034152 | 0.04193625 | 0.312747634 |
| *TIMM44* | 0.187279 | 0.092111 | 2.03318 | 0.04203436 | 0.313278499 |
| *DHX33* | 0.28938 | 0.142366 | 2.032651 | 0.04208784 | 0.313296177 |

| | | | | | |
|---|---|---|---|---|---|
| *ZNF263* | 0.27875 | 0.137138 | 2.032623 | 0.04209059 | 0.313296177 |
| *KCNQ5* | 0.543538 | 0.267586 | 2.031265 | 0.04222811 | 0.313972279 |
| *JAK2* | -0.561369 | 0.276374 | -2.031193 | 0.0422354 | 0.313972279 |
| *EPS8L2* | -0.519076 | 0.25564 | -2.030494 | 0.04230633 | 0.314298713 |
| *FAM135A* | -0.424096 | 0.208963 | -2.029521 | 0.04240525 | 0.314571959 |
| *MYO1G* | -0.326637 | 0.16097 | -2.02918 | 0.04243992 | 0.314571959 |
| *RTCB* | 0.233379 | 0.115016 | 2.029098 | 0.04244833 | 0.314571959 |
| *ATM* | -0.165363 | 0.081507 | -2.028826 | 0.04247601 | 0.314571959 |
| *ENSG00000273669* | -1.154871 | 0.56932 | -2.028512 | 0.04250806 | 0.314571959 |
| *MRFAP1L1* | 0.246039 | 0.121303 | 2.028311 | 0.04252856 | 0.314571959 |
| *HMBS* | 0.234123 | 0.11543 | 2.028273 | 0.04253238 | 0.314571959 |
| *HK2* | 0.597318 | 0.294558 | 2.027843 | 0.0425763 | 0.314610495 |
| *PPM1L* | 0.90217 | 0.444925 | 2.027692 | 0.04259167 | 0.314610495 |
| *OPTN* | -0.437817 | 0.216001 | -2.026921 | 0.04267054 | 0.314993066 |
| *ACOT7* | 0.489751 | 0.241721 | 2.0261 | 0.04275454 | 0.315136813 |
| *ANXA2R* | -0.367086 | 0.181189 | -2.025983 | 0.04276654 | 0.315136813 |
| *COQ7* | 0.303943 | 0.150026 | 2.025937 | 0.04277128 | 0.315136813 |
| *QPCT* | 0.763099 | 0.376757 | 2.02544 | 0.04282219 | 0.315312282 |
| *DDT* | 0.147782 | 0.07302 | 2.02386 | 0.04298459 | 0.316307877 |
| *LINC01138* | -0.43373 | 0.214387 | -2.023119 | 0.0430609 | 0.316669092 |
| *TCF4* | -0.302304 | 0.149463 | -2.022596 | 0.04311486 | 0.316766403 |
| *KIF2A* | -0.230317 | 0.113879 | -2.022463 | 0.04312859 | 0.316766403 |
| *MRPS28* | 0.274733 | 0.135883 | 2.021841 | 0.04319283 | 0.316860542 |
| *PIAS2* | -0.241256 | 0.119327 | -2.021811 | 0.04319588 | 0.316860542 |
| *RPP25* | 0.419647 | 0.207758 | 2.019878 | 0.04339601 | 0.318128015 |
| *C4orf36* | -0.695668 | 0.344948 | -2.016737 | 0.043723 | 0.32032329 |
| *IRAK1* | 0.200749 | 0.099612 | 2.015319 | 0.04387128 | 0.321207378 |
| *AP4B1* | -0.440645 | 0.218711 | -2.014743 | 0.04393157 | 0.321446447 |
| *NUP210* | 0.220127 | 0.109305 | 2.013884 | 0.04402168 | 0.321746574 |
| *CCDC144A* | 0.844036 | 0.419121 | 2.013825 | 0.0440279 | 0.321746574 |
| *ANKRD54* | 0.267023 | 0.132711 | 2.012063 | 0.04421329 | 0.32223855 |
| *ZER1* | 0.5831 | 0.289821 | 2.011931 | 0.04422717 | 0.32223855 |
| *N4BP3* | -0.330362 | 0.164218 | -2.011727 | 0.04424877 | 0.32223855 |
| *PSMC5* | 0.143784 | 0.07148 | 2.011521 | 0.04427044 | 0.32223855 |
| *MDK* | 0.822298 | 0.408822 | 2.011386 | 0.04428468 | 0.32223855 |
| *CCT6A* | 0.15886 | 0.078986 | 2.011252 | 0.04429882 | 0.32223855 |
| *KIAA1841* | -0.742391 | 0.369137 | -2.011155 | 0.04430904 | 0.32223855 |
| *RPSA* | -0.135636 | 0.067449 | -2.010951 | 0.0443306 | 0.32223855 |
| *ENSG00000197180* | -0.428138 | 0.212917 | -2.01082 | 0.0443445 | 0.32223855 |
| *UQCRQ* | 0.182743 | 0.090918 | 2.009966 | 0.04443481 | 0.322693207 |
| *LTB* | -0.337357 | 0.167879 | -2.00952 | 0.04448199 | 0.322834366 |
| *CEPT1* | -0.20778 | 0.103416 | -2.009168 | 0.04451936 | 0.322904136 |
| *RPL7A* | -0.125297 | 0.062395 | -2.008126 | 0.04462995 | 0.323504559 |
| *AK1* | 0.496099 | 0.247087 | 2.007788 | 0.04466583 | 0.32356302 |
| *DMTF1* | -0.240869 | 0.12003 | -2.006751 | 0.04477621 | 0.32416082 |
| *PPARA* | -0.439956 | 0.219466 | -2.004669 | 0.04499847 | 0.325567255 |
| *RGS19* | -0.209592 | 0.104571 | -2.004311 | 0.0450368 | 0.325642118 |

| | | | | | |
|---|---|---|---|---|---|
| *PCP2* | -1.099172 | 0.548702 | -2.003223 | 0.0451534 | 0.326282385 |
| *NRSN2.AS1* | -0.748102 | 0.37354 | -2.002734 | 0.0452058 | 0.326458288 |
| *NSUN6* | -0.30726 | 0.153535 | -2.001233 | 0.04536725 | 0.327257709 |
| *ERMARD* | 0.481285 | 0.240502 | 2.001167 | 0.0453744 | 0.327257709 |
| *ENSG00000260979* | 0.735893 | 0.367777 | 2.000921 | 0.04540089 | 0.327257709 |
| *DCBLD2* | -0.927033 | 0.463393 | -2.000532 | 0.04544288 | 0.327357567 |
| *ENSG00000272843* | -1.711458 | 0.855544 | -2.000432 | 0.04545364 | NA |
| *FOXRED2* | 0.676423 | 0.338183 | 2.000171 | 0.0454818 | 0.327435177 |
| *VSIG10L* | 1.366172 | 0.68319 | 1.999697 | 0.04553295 | NA |
| *TMEM208* | 0.149648 | 0.074848 | 1.99936 | 0.04556937 | 0.327862701 |
| *PSMA3* | 0.186423 | 0.09328 | 1.998536 | 0.04565863 | 0.328301941 |
| *KIF20B* | -0.235018 | 0.117645 | -1.997683 | 0.04575107 | 0.328499232 |
| *JMJD8* | 0.293168 | 0.146763 | 1.997567 | 0.04576361 | 0.328499232 |
| *PFDN4* | 0.170172 | 0.085193 | 1.997501 | 0.04577078 | 0.328499232 |
| *UBE3D* | 0.336388 | 0.168464 | 1.996798 | 0.04584714 | 0.32884441 |
| *STRBP* | -0.171188 | 0.085758 | -1.996168 | 0.04591559 | 0.328927788 |
| *CNR1* | 0.724619 | 0.363045 | 1.99595 | 0.04593933 | 0.328927788 |
| *YARS* | 0.210092 | 0.105261 | 1.995911 | 0.04594358 | 0.328927788 |
| *SF3A3* | 0.188622 | 0.09453 | 1.995358 | 0.04600382 | 0.329156454 |
| *MSTO1* | 0.360871 | 0.18091 | 1.994759 | 0.0460692 | 0.329421666 |
| *UPF3B* | -0.182692 | 0.091616 | -1.994115 | 0.0461395 | 0.329721733 |
| *HSPH1* | 0.220146 | 0.110502 | 1.992233 | 0.04634552 | 0.330774724 |
| *KLF3* | 0.339185 | 0.170279 | 1.991929 | 0.04637884 | 0.330774724 |
| *TFB2M* | 0.320477 | 0.160896 | 1.991831 | 0.04638966 | 0.330774724 |
| *CISD2* | 0.172477 | 0.086597 | 1.991731 | 0.04640058 | 0.330774724 |
| *RPGR* | -0.450263 | 0.226102 | -1.991412 | 0.04643566 | 0.330822113 |
| *NFE2L1* | 0.251859 | 0.12656 | 1.990029 | 0.04658774 | 0.331702419 |
| *TOB2* | -0.199162 | 0.100122 | -1.989195 | 0.04667971 | 0.332153972 |
| *APPBP2* | -0.31515 | 0.158481 | -1.988561 | 0.04674968 | 0.332257442 |
| *AGPAT2* | -0.238397 | 0.119904 | -1.988231 | 0.04678614 | 0.332257442 |
| *PELO* | 0.360678 | 0.181429 | 1.987984 | 0.04681343 | 0.332257442 |
| *SASH3* | -0.241936 | 0.121702 | -1.987929 | 0.04681955 | 0.332257442 |
| *CARS2* | 0.320978 | 0.161542 | 1.986968 | 0.04692594 | 0.332257442 |
| *UNC45A* | -0.236137 | 0.118853 | -1.98679 | 0.04694568 | 0.332257442 |
| *TRAP1* | 0.27768 | 0.139765 | 1.986763 | 0.04694871 | 0.332257442 |
| *PHF11* | -0.20197 | 0.101662 | -1.986688 | 0.04695702 | 0.332257442 |
| *INKA1* | -0.510893 | 0.2572 | -1.986363 | 0.04699304 | 0.332257442 |
| *DTD2* | 0.303642 | 0.15287 | 1.98628 | 0.04700223 | 0.332257442 |
| *PGK1* | 0.260144 | 0.130974 | 1.986224 | 0.0470084 | 0.332257442 |
| *RAB11FIP1* | -0.23847 | 0.120112 | -1.985399 | 0.04710011 | 0.332703539 |
| *METRN* | 0.492511 | 0.24821 | 1.984253 | 0.0472276 | 0.333165886 |
| *BRMS1* | 0.139578 | 0.070349 | 1.984088 | 0.04724602 | 0.333165886 |
| *POLR2K* | 0.167935 | 0.084643 | 1.984039 | 0.04725148 | 0.333165886 |
| *ENSG00000225489* | 0.613169 | 0.309171 | 1.983268 | 0.04733749 | 0.333502345 |
| *PDK3* | -0.421913 | 0.212755 | -1.983098 | 0.04735653 | 0.333502345 |
| *CCDC57* | -0.23867 | 0.120435 | -1.981734 | 0.04750901 | 0.334045277 |
| *SYNC* | -0.321062 | 0.162015 | -1.981676 | 0.04751548 | 0.334045277 |

| | | | | | |
|---|---|---|---|---|---|
| *RBM28* | 0.254226 | 0.128291 | 1.981638 | 0.04751976 | 0.334045277 |
| *HERC1* | -0.348193 | 0.175878 | -1.979741 | 0.04773269 | 0.335339465 |
| *TBXAS1* | 0.859633 | 0.434726 | 1.977413 | 0.04799497 | 0.33691414 |
| *TSSC4* | 0.185185 | 0.093658 | 1.977238 | 0.04801475 | 0.33691414 |
| *ENSG00000274536* | -0.46241 | 0.233969 | -1.976374 | 0.04811248 | 0.337287041 |
| *ACSM3* | -0.499009 | 0.252573 | -1.975705 | 0.04818823 | 0.337287041 |
| *NMRK1* | -0.731328 | 0.370196 | -1.975517 | 0.04820955 | 0.337287041 |
| *BRWD3* | -0.460698 | 0.233215 | -1.975423 | 0.04822013 | 0.337287041 |
| *PDPK1* | -0.24661 | 0.12484 | -1.975416 | 0.04822101 | 0.337287041 |
| *SPN* | 0.490584 | 0.248391 | 1.975043 | 0.04826324 | 0.337287041 |
| *PHF20L1* | -0.222503 | 0.112674 | -1.974751 | 0.04829636 | 0.337287041 |
| *FAH* | 0.586778 | 0.297157 | 1.974639 | 0.04830908 | 0.337287041 |
| *BRCA2* | -0.40212 | 0.203668 | -1.974386 | 0.04833789 | 0.337287041 |
| *PLXNB2* | 0.762668 | 0.386316 | 1.974211 | 0.04835781 | 0.337287041 |
| *OGA* | -0.148608 | 0.075296 | -1.97364 | 0.04842271 | 0.337537306 |
| *USP3* | -0.185763 | 0.09415 | -1.973065 | 0.04848812 | 0.337537649 |
| *ZNF431* | -0.202435 | 0.102599 | -1.973062 | 0.04848854 | 0.337537649 |
| *CRLF3* | -0.199035 | 0.100939 | -1.971834 | 0.04862856 | 0.337537649 |
| *CLK4* | -0.301907 | 0.15314 | -1.971447 | 0.04867282 | 0.337537649 |
| *BEND5* | -0.434497 | 0.220414 | -1.971279 | 0.04869195 | 0.337537649 |
| *PLEKHG2* | -0.714852 | 0.362665 | -1.97111 | 0.04871133 | 0.337537649 |
| *CRYM.AS1* | -1.3473 | 0.683535 | -1.971076 | 0.04871513 | 0.337537649 |
| *LAPTM4B* | 0.713477 | 0.361978 | 1.97105 | 0.04871817 | 0.337537649 |
| *NUDCD3* | -0.23053 | 0.116963 | -1.970963 | 0.04872811 | 0.337537649 |
| *ZNF322* | -0.322267 | 0.163529 | -1.970706 | 0.04875749 | 0.337537649 |
| *INTS6* | -0.181002 | 0.09185 | -1.970613 | 0.04876814 | 0.337537649 |
| *LINC01572* | 0.693094 | 0.351741 | 1.970466 | 0.04878504 | 0.337537649 |
| *FAM118B* | 0.419666 | 0.212992 | 1.970336 | 0.04879993 | 0.337537649 |
| *MBD4* | -0.172939 | 0.087809 | -1.969498 | 0.0488959 | 0.338000505 |
| *ENSG00000274184* | -0.901064 | 0.457606 | -1.969082 | 0.04894368 | 0.338129929 |
| *SEMA4A* | 0.887266 | 0.450734 | 1.968492 | 0.04901141 | 0.338185077 |
| *ENSG00000160602* | -0.248527 | 0.126254 | -1.96847 | 0.04901395 | 0.338185077 |
| *ERP29* | -0.118496 | 0.060204 | -1.968254 | 0.04903887 | 0.338185077 |
| *TEP1* | -0.476538 | 0.242145 | -1.967989 | 0.04906925 | 0.338194099 |
| *TTC39C* | -0.491645 | 0.250037 | -1.96629 | 0.04926507 | 0.339048645 |
| *LTA* | -0.370686 | 0.188526 | -1.966228 | 0.04927228 | 0.339048645 |
| *VPS13C* | -0.201181 | 0.102323 | -1.966131 | 0.0492835 | 0.339048645 |
| *CCND3* | 0.159973 | 0.081374 | 1.965903 | 0.04930981 | 0.339048645 |
| *NDUFV1* | 0.170792 | 0.0869 | 1.96539 | 0.04936914 | 0.339256104 |
| *DOT1L* | 0.30988 | 0.157709 | 1.964884 | 0.04942769 | 0.339457948 |
