## Supplemental Methods for "Single-cell Landscape Analysis Unravels Molecular Programming of the Human B Cell Compartment in Chronic GVHD"

*Additional Information on Patient and HD blood samples*

Single-cell RNA-Seq was performed on peripheral B cells from allo-HCT patients chosen at random, with the desire to gain new knowledge about the molecular mechanisms that emerge in the allo-HCT setting to drive B cell hyper-responsiveness and cGVHD development in some patients (loss of tolerance). Anonymous HD samples were buffy coats purchased from Gulf Coast Regional Blood Center (Houston, TX). Viably frozen PBMCs from allo-HCT patients or HDs were prepared by Ficoll-Paque® PLUS (GE Healthcare) gradient separation prior to storage in a liquid nitrogen freezer. For some experiments, purified B cells were additionally isolated immediately following PBMC gradient separation using a Human B Cell Isolation Kit II (Miltenyi Biotech), washed in cold Dulbecco's phosphate-buffered saline, and stored as B cell pellets at -80°C. The characteristics of allo-HCT patients who provided the samples used for single-cell RNA-Seq analysis and the samples used for additional experiments are described in **Supplemental Tables 1** and **2,** respectively.

*B cell culture, single-cell RNA-Seq library preparation, and library sequencing*

B cells were purified from viably cryopreserved allo-HCT patient PBMCs using the Human B Cell Isolation Kit II (Miltenyi Biotech). B cells were then cultured for 18 h at 5% $CO_2$ in RPMI-1640 medium containing 10% fetal bovine serum (FBS) and 55 μM 2-Mercaptoethanol (Gibco). In some wells, cells were treated with 100 nM ATRA (MP Biomedicals, cat# 190269) for the entire culture period. The cells were then harvested, and dead cells were removed using an EasySep™ Dead Cell Removal (Annexin V) Kit (STEMCELL Technologies, Inc.). Live B cells (in HBSS containing

2% FBS) were immediately transferred on ice to the Duke Molecular Genomics Core for 10X Genomics Chromium™ Single-Cell 3' library generation according to the manufacturer's instructions and the Core Facility's standard methods, targeting 10,000 B cells per sample. 10X Genomics libraries were then transferred directly to the Duke Center for Genomic and Computational Biology, Sequencing and Genomic Technologies Shared Resource, for sequencing. All individual 10X Genomics libraries (16 total) were pooled and then applied evenly across all 4 lanes of a NovaSeq S4 Flow Cell (Illumina) with 100-bp paired-end sequencing at a depth of ~700 million reads per sample (~70,000 reads per cell).

*Single-cell RNA-Seq data analysis on purified B cells from allo-HCT patients*

The raw data in BCL format were de-multiplexed into FASTQs by Illumina's bcl2fastq (https://support.illumina.com). The extraction of cellular barcodes and unique molecular identifiers (UMI sequences) and genome alignment of the biological sequences were performed using the Cell Ranger pipeline (version 2.1.1, https://support.10xgenomics.com) to generate a filtered sparse matrix of gene read counts per cell. The 16 individual matrices were integrated using Stuart et al.'s method to remove batch effects (*1*). Any cells in which fewer than 500 genes were detected and those with greater than 5% mitochondrial gene counts were considered poor quality and excluded from the analysis. Any genes detected in fewer than 150 cells were likewise excluded. Visualizations of single-cell-level transcription profiles in UMAP space using color was performed using R package Seurat v3.1.4 (*1*). Trajectory analysis was performed to infer the relative relationship among cell clusters over 'pseudotime' using the package Slingshot (*2*).

*Comparison of allo-HCT patient single-cell RNA-Seq data to a publicly available dataset on purified B cells from HDs, HIV patients, and malaria-exposed individuals*

To compare scRNA data from this study to data published by Holla *et al.* (*3*), both datasets were filtered for cell quality using the same thresholds (cells with more than 5% mitochondrial gene counts, counts from fewer than 200 genes, or counts from more than 2,500 genes were excluded). High-quality cells were then filtered to retain only B cells, with cell type inferred based on signature genes from CellAssign (*4*) and expression of the union of those signature genes and the top 2000 most variable genes (following variance stabilizing transformation (*5*)) using the R package scSorter (*6*). To normalize the datasets, gene counts per cell were transformed to counts per million reads mapped (CPM) using the Seurat R package. To compare gene expression, data from the present study and *Holla et al.*(*3*) were integrated following steps recommended by the developers of Seurat. Five cohorts were integrated to account for batch effects, as follows: Holla *et al*. data (*3*) were split by sample type (HD vs. HIV vs. Malaria), and allo-HCT data was split by the two sequencing runs. To investigate biologically-relevant genes across studies and sample types, we generated a heatmap with the following attributes: We identified each subset based on a detection threshold of 50 CPM for *ITGAX* and *CD27* expression. To illustrate distribution of gene expression, values were plotted lowest to highest for each gene of interest and within each B cell subset ($CD27^{+}ITGAX^{-}$, $CD27^{-}ITGAX^{+}$, and $CD27^{+}ITGAX^{+}$) and sample type. Populations with more than 100 B cells were randomly down-sampled to 100 cells. Data analyses and plot generation were carried out in the R statistical environment with packages from the comprehensive R archive network (CRAN; https://cran.r-project.org/) and the Bioconductor project (*7*). Code used to replicate the analyses and figures followed principles of reproducible analysis, leveraging knitr dynamic reports and git source code management (*8*).

*Dermal skin cell isolation and single-cell RNA-Seq analysis*

The skin biopsies were washed with cold DPBS and digested with 1X dispase (ThermoFisher Scientific) overnight at 4ºC. The dermis was mechanically separated from the epidermis and then digested at 37ºC and 5% CO2 with 0.25% trypsin/EDTA for 1.5 hr. Dermal cell suspensions were then passed through 70 μm and 40 μm cell strainers, then pelleted by centrifugation and resuspended in Keratinocyte-SFM (ThermoFisher Scientific) containing 0.4% BSA. Single-cell suspensions were then subjected to 10X Genomics Chromium library generation and Illumina sequencing as described above. Signature gene analysis was performed using the R package Seurat to identify B cells within dermal cell clusters. DEG analysis was performed using the R package DESeq2, as described for blood B cells above. The skin cell scRNA-Seq library data will be made available through the Gene Expression Omnibus (GEO) database (https://www.ncbi.nlm.nih.gov/geo/) following peer review.

*Quantitative PCR (qPCR)*

Frozen (-80°C) B cell pellets were thawed on ice, and total RNA was isolated using the RNeasy Plus Mini Kit (Qiagen) and quantified with a Qubit® 3.0 Fluorometer (ThermoFisher). RNA (500 ng) was reverse-transcribed with an iScript cDNA Synthesis Kit (BioRad). *CKS2* gene expression was assessed by qPCR using *ACTB* (*β-ACTIN*) as the housekeeping gene. Primers were designed with the NCBI primer BLAST program: *CKS2*, forward 5'-ACGAGTACCGGCATGTTATGT-3' and reverse 5'-GCCTAGACTCTGTTGGACACC-3'; *ACTB*: forward, 5'-

GCTGTGCTACGTCGCCCT-3’ and reverse, 5’-AAGGTAGTTTCGTGGATGCC-3’. Amplification was performed using the iTaq Universal SYBR® Green Supermix (Bio-Rad) with an annealing temperature of 60°C on an ABI StepOne Plus™ Real-time PCR System (ThermoFisher Scientific). Each 20-µl reaction contained 2.5 ng of cDNA, with each primer used at a concentration of 250 nM. Data analysis was performed using Step One Software v2.3 software, and relative quantitation of gene expression was achieved using the standard-curve method. Fold change in expression in each sample was calculated using the average gene expression in patients with No cGVHD.

*Flow cytometry and PhenoGraph analysis*

Viably frozen PBMCs from allo-HCT patients were surface stained using antibodies against CD19, CD11c (ITGAX), CD21, CD24, CD27, EBI2 (GPR183), and TACI, along with their corresponding isotype control antibodies. 7-AAD (BioLegend) was used to assess cell viability. PhenoGraph cluster analysis was then performed on the remaining 6 markers after first pre-gating on viable B cells ($CD19^+$ $7AAD^-$), using this available function in Flowjo in combination with R. A detailed summary of all antibodies used is as follows: CD19 Pacific blue (clone J3-119, Beckman Coulter, cat#A86355); CD24 BV510 (clone ML5, BioLegend, cat#311126); CD21 FITC (Bu32, BioLegend, cat#354910); TACI PE (clone 1A1, BioLegend, cat#311906), EBI2 (GPR183) PE (clone SA313E4, BioLegend, cat#368912); CD11c APC (clone 3.9, BioLegend, cat#301614); CD27 PE-Cy7 (clone O323, eBioscience, cat#25-0279-42); IgD APC-H7 (clone IA6-2, BD Biosciences, cat#561305). Isotype control antibodies used for flow cytometry were: Mouse IgG2a BV510 (clone MOPC-173, BioLegend, cat#400268); Mouse IgG1 FITC (clone MOPC-21, BioLegend, cat#400110); Rat IgG2a PE (clone R35-95, BD Biosciences, cat#553930); Mouse

IgG2a PE (clone MOPC-173, BioLegend, cat#400214); Mouse IgG1 APC (clone MOPC-21, BioLegend, cat#400120); Mouse IgG1 PE-Cy7 (clone MOPC-21, BioLegend, cat#400126); Mouse IgG2a APC-H7 (clone G155-178, BD Biosciences, cat#560897).

*Phosphoarray analysis*

Proteome Profiler™ Human Phospho-Kinase Array Kits (R&D Systems) were utilized according to the manufacturer's instructions to analyze whole cell lysates of purified, untreated B cell samples from patients with Active cGVHD or No cGVHD (*n*=3 each). Spot densities on the arrays were quantified using ImageJ software (https://imagej.nih.gov/ij/).

*Western blot analysis*

Frozen (-80°C) pellets of purified B cells from allo-HCT patients with Active cGVHD (*n*=4) or No cGVHD (*n*=4) were lysed using a commercially available whole cell lysis buffer (Roche Complete Lysis-M) with freshly added protease inhibitors (Roche). Protein lysates were then denatured and loaded into wells of a 4–12% Bis-Tris gel (ThermoFisher) for electrophoresis under reducing conditions. Proteins were transferred from the gels to a nitrocellulose membrane using dry electroblotting (20 V for 9 min). Membranes were blocked in 2% fish gelatin buffer for 75 min and then incubated with rabbit anti-human $P27^{KIP1}$ polyclonal antibody (C-19, Santa Cruz Biotechnology, cat#sc-528, lot#KO413) overnight at 4°C. The membrane was thoroughly washed in Tris-buffered saline containing Tween 20 (TBST) before incubation for 1 h with donkey anti-rabbit polyclonal secondary antibody labeled with a near-infrared fluorescent dye (IRDye® 680,

LI-COR, cat#926-68073, lot#C50821-05). The membrane was then washed thoroughly in TBST, and fluorescently labeled proteins were detected using an LI-COR Odyssey CLX Imaging System.
